# E proteins orchestrate dynamic transcriptional cascades to suppress ILC2 differentiation

**DOI:** 10.1101/2020.03.26.008037

**Authors:** Vincent Peng, Constantin Georgescu, Anna Bakowska, Liangyue Qian, Jonathan D Wren, Xiao-Hong Sun

**Affiliations:** Oklahoma Medical Research Foundation, Program in Arthritis and Clinical Immunology, Oklahoma City, OK; Department of Pathology and Immunology, Washington University School of Medicine, St Louis, MO; Oklahoma Medical Research Foundation, Program in Genes and Human Diseases, Oklahoma City, OK; Department of Cell Biology, University of Oklahoma Health Sciences Center, Oklahoma City, OK; Department of Microbiology and Immunology, University of Oklahoma Health Sciences Center, Oklahoma City, OK

## Abstract

The basic helix-loop-helix transcription factors collectively called E proteins powerfully suppress the differentiation of group2 innate lymphoid cells from bone marrow and thymic progenitors. Here we investigated the underlying molecular mechanisms using inducible gain and loss of function approaches in ILC2s and their precursors, respectively. Cross-examination of RNA sequencing and ATAC sequencing data obtained at different time points reveals a set of genes which are likely direct targets of E proteins. Consequently, a widespread down-regulation of chromatin accessibility occurs at a later time point, possibly due to the activation of transcriptional repressor genes such as *Cbfa2t3* and *Jdp2*. The large number of genes repressed by gain of E protein function leads to the down-regulation of a transcriptional network important for ILC2 differentiation.

**Summary:** Differentiation of group 2 innate lymphoid cells is forcefully repressed by E protein transcription factors. This report elucidates how E proteins repress a transcriptional network important for ILC2 differentiation by up-regulating the expression of transcriptional repressors.

---

Group 2 innate lymphoid cells (ILC2s) are a subset of innate lymphoid cells (ILCs) which play important roles in initiating type 2 immunity (1). They are enriched in barrier locations such as the lung, intestine and skin (2). They respond to signals released by tissue damages, namely IL-25, IL-33 and TSLP, and produce type 2 cytokines primarily consisting of IL-5, IL-9 and IL-13 (3). However, ILC2s exist as heterogeneous populations with diverse characteristics and functionality due to the influences of the tissues in which they reside and the cell origins from which they arise (4;5). A complete knowledge of the ontogeny of ILC2s will help understand ILC2-mediated immunity.

ILC2s are generated during fetal, neonatal and adult stages, among which the neonatal phase contributes to the majority of the tissue-resident ILC2s (2). ILC2s differentiate from progenitors such as common lymphoid progenitors (CLP) or lymphoid-multipotent progenitors (LMPP), which are also responsible for making B and T lymphoid cells as well as NK cells (6). ILC2 differentiation has been thought to take place in the fetal liver and bone marrow (3). However, we have established that this also occurs in the thymus, where ILC2s not only differentiate from multipotent early T cell progenitors (ETP or DN1 cells) but also from committed T cell precursors, namely CD4^-^CD8^-^ CD25^+^c-kit^-^ (DN3) cells (5;7;8). In fact, the thymus is a fertile niche for ILC2 differentiation when a critical checkpoint that ensures proper T cell development is disabled.

The basic helix-loop-helix family to transcription factors, collectively called E proteins, serve as a critical checkpoint to promote B and T cell development and suppress ILC2 differentiation (5;7-10). In the thymus, the *Tcf3* and *Tcf12* genes are expressed to produce their respective E proteins, E12 and E47 from *Tcf3* and HEB from *Tcf12*.

These proteins dimerize among themselves and activate transcription by binding to the DNA sequences called E boxes. When both genes are deleted using a thymus-specifically expressed Cre, *plck*-Cre, T cell development is blocked at the DN3 stage when the Cre transgene is first expressed (7;11). Consequently, an abundance of ILC2s are generated in *Tcf3* and *Tcf12* double knock-out mice (dKO). E protein function can be abolished by their naturally occurring inhibitors, Id proteins, which dimerize with E proteins and prevent DNA binding. When Id1 is ectopically expressed in the thymus, T cell development is arrested and ILC2 development is drastically enhanced, leading to accumulation of 10 to 100-fold more ILC2s in different tissues (7). The promotion of ILC2 differentiation in the absence of E protein activities are not simply the result of T cell developmental blockade because numerous other animal models with T cell defects (e.g., *Rag1*^-/-^ mice) do not exhibit such robust enhancement of ILC2 production (Bajana and Sun, unpublished). Thus, it is likely that E proteins play a specific role in suppressing ILC2 differentiation.

To understand the underlying mechanisms, we sought to elucidate the transcriptional programs controlled by E proteins in the context of ILC2 differentiation. We measured transcriptomes and chromatin accessibilities in T cell precursors and in ILC2s upon inducible loss or gain of E protein function. We observed dynamic changes in gene regulation, which ultimately leads to a suppression of ILC2 transcriptional programs by E proteins.

## Results and Discussion

### Altered gene expression immediately following inducible E protein ablation in T cell precursors

In addition to our previous report of the substantial increase in ILC2 numbers in *Tcf3* and *Tcf12* double knockout mice, we have also shown that deletion of these genes using tamoxifen-inducible Cre enhances ILC2 differentiation from CLP, DN1 and DN3 cells by 20-40 fold when cultured on OP9-DL1 stromal cells for 11-14 days (5;7). To detect changes in gene expression immediately following E protein ablation, we isolated sufficient DN1 and DN3 cells for RNA sequencing after brief cultures on OP9-DL1 stromal cells. We seeded stromal cells with DN1 or DN3 cells isolated from ROSA26-CreERT2;ROSA26-Stop-tdTomato;*Tcf3*^f/f^;*Tcf12*^f/f^ (dKO) mice or control mice without the floxed alleles of the E protein genes. Four days later, tamoxifen was added and tdTomato^+^ (indicative of successful Cre-mediated deletion) cells were sorted 24 and 72 hours later for RNA sequencing (Fig. 1a). At these time points, few ILC2s were detected (5), thus allowing the evaluation of early changes in gene expression in ILC2 precursors.

**Figure 1.**
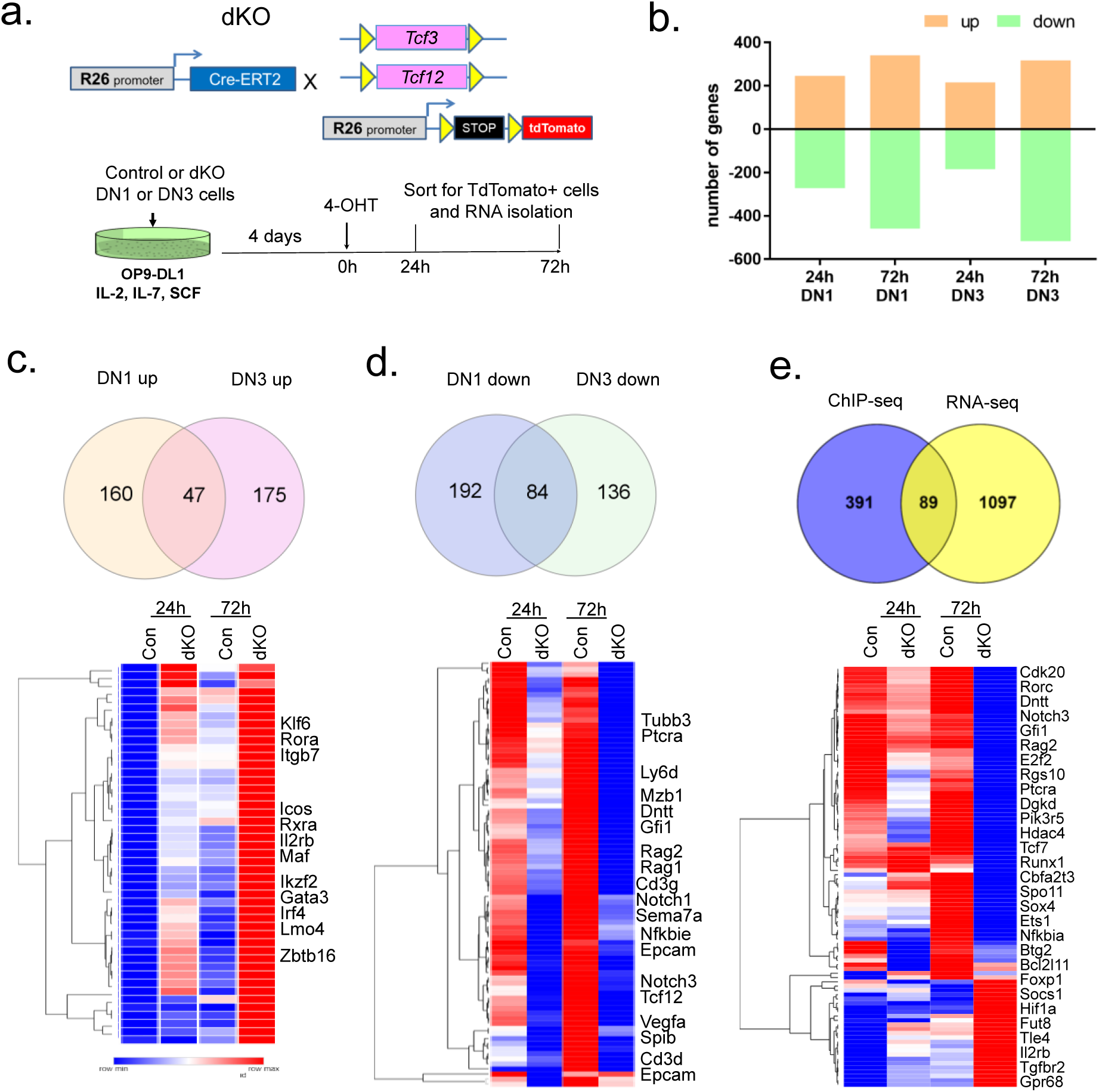
Transcriptome changes immediately after E protein ablation. **(a)** Experimental design. DN1 and DN3 thymocytes from ROSA26-CreERT2;ROSA26-stop-tdTomato;E2A^f/f^;HEB^f/f^ (dKO) and ROSA26-CreERT2;ROSA26-stop-tdTomato (control) mice were co-cultured on OP9-DL1 stromal cells. Biological duplicates were collected 24 and 72 hours (h) later and sorted for tdTomato expression before RNA isolation and sequencing. (b) Numbers of differentially expressed genes comparing dKO and control at indicated time points and cell preparations. Genes with a log2 fold change (LFC) > 1 and false discovery rate (FDR) < 0.05 were counted. **(c and d)** Venn diagrams show the overlaps of genes up and down regulated in DN1 and DN3-derived cells. Expression levels of genes commonly found in both DN1 and DN3-derived cells are shown in heatmaps below each Venn diagram with representative genes listed. (**e**) Venn diagram compares protein-coding genes differentially expressed in DN3-derived cells to genes assigned to anti-E protein (E47) ChIP-seq peaks detected in ex vivo Rag2^-/-^ thymocytes. Fisher test was used to determine the statistical significance of the overlap (p<0.00001). Heatmap shows the relative levels of expression in DN3-derived cells with representatives listed on the side.

The transcriptomes of dKO and controls at 24h and 72h were compared separately (Fig.1b). Although there were substantial overlaps between the differential genes at 24h and 72h in DN1 and DN3 cells, differences were also found in terms of the magnitude of changes and genes differentially expressed. DN3 cells harbor a higher level of endogenous E proteins than DN1 cells, mostly due to the up-regulation of *Tcf12* (http://rstats.immgen.org/Skyline/skyline.html), and thus the peak time of the changes for some of the genes were delayed compared to DN1 cells. Overall, we obtained lists of 1140 and 1376 differentially regulated protein coding genes at either time point in DN1 and DN3 cells, respectively (supplemental table 1).

DN1 and DN3 cells, being at different developmental stages, have distinct transcriptomes. Since E protein ablation promoted ILC2 differentiation from both cell types, a common set of differential genes will be extremely informative for understanding how E proteins suppress ILC2 differentiation. Remarkably, a profound number of genes up-regulated due to E protein ablation are known to be associated with ILC differentiation. This list consists of *Zbtb16, Rora, Maf, Lmo4, Klf6, Icos, Itgb7, Ikzf2, Irf4*, and *Gata3* (Fig. 1c). Conversely, among the genes down-regulated by the loss of E proteins are genes previously known to be activated by E proteins and important for T cell development (Fig. 1d) (12). These include *Rag1, Rag2, Notch1, Notch3, Cd3g, Cd3d, Ptcra* and *Spib*. These data indicate an instrumental role for E-protein activity in controlling the switch between ILC2 and T cell development.

Not all of the genes found in our analyses are directly regulated by E proteins. In order to identify direct targets, we compared the list of genes altered in DN3 cells to E2A-ChIP-seq data generated by Murre and colleagues (Fig. 1e) (12). Eighty-nine of differentially regulated protein-coding genes in cultured DN3 cells were found to be bound by E2A proteins in *Rag2*^-/-^ thymocytes, which are mostly cells arrested at the DN3 stage, suggesting that these genes are likely direct targets of E proteins. Although the overlap between our RNA-seq results and the ChIP-seq data is highly significant in a Fisher test, the small number of genes overlapped in the two datasets may be due to changes in gene expression which already occurred after a 4-day culture of DN3 cells on OP9-DL1 stroma before E protein deletion such that their transcriptional profile is intrinsically different from that of ex-vivo Rag2^-/-^ DN3 cells, which lack pre-TCR signaling.

### Transcript levels altered by inducible E protein activity in ILC2s

We next took a complementary approach by examining gene expression in ILC2s after overexpression of an inducible E protein, E47-ER, in which E47 was fused to a modified hormone-binding domain of estrogen receptor (13). E47 activity can then be quickly induced by releasing the fusion protein from the cytoplasm to the nucleus upon binding to tamoxifen. We made use of our Id1 transgenic mice, in which a large amount of functional ILC2s are produced in the thymus (7). Thymic ILC2s were harvested from these mice by sorting for Lin^-^Thy1^+^ICOS^+^ST2^+^ cells and stimulated to proliferate with IL-2, IL-7, IL25 and IL-33. These cells were then transduced with retroviruses expressing E47-ER and EGFP or EGFP alone at a relatively high efficiency (about 30%).

Transduced cells were sorted for EGFP expression and expanded before treatments with tamoxifen for 4 and 16 hours, respectively and RNA sequencing was performed (Fig. 2a). These short time points were chosen to detect immediate effects of E47 and to avoid any toxicity of E protein known to exist in B and T cells. Interestingly, ILC2s appeared to have tolerated E47 remarkably well within 16 hours (data not shown).

**Figure 2.**
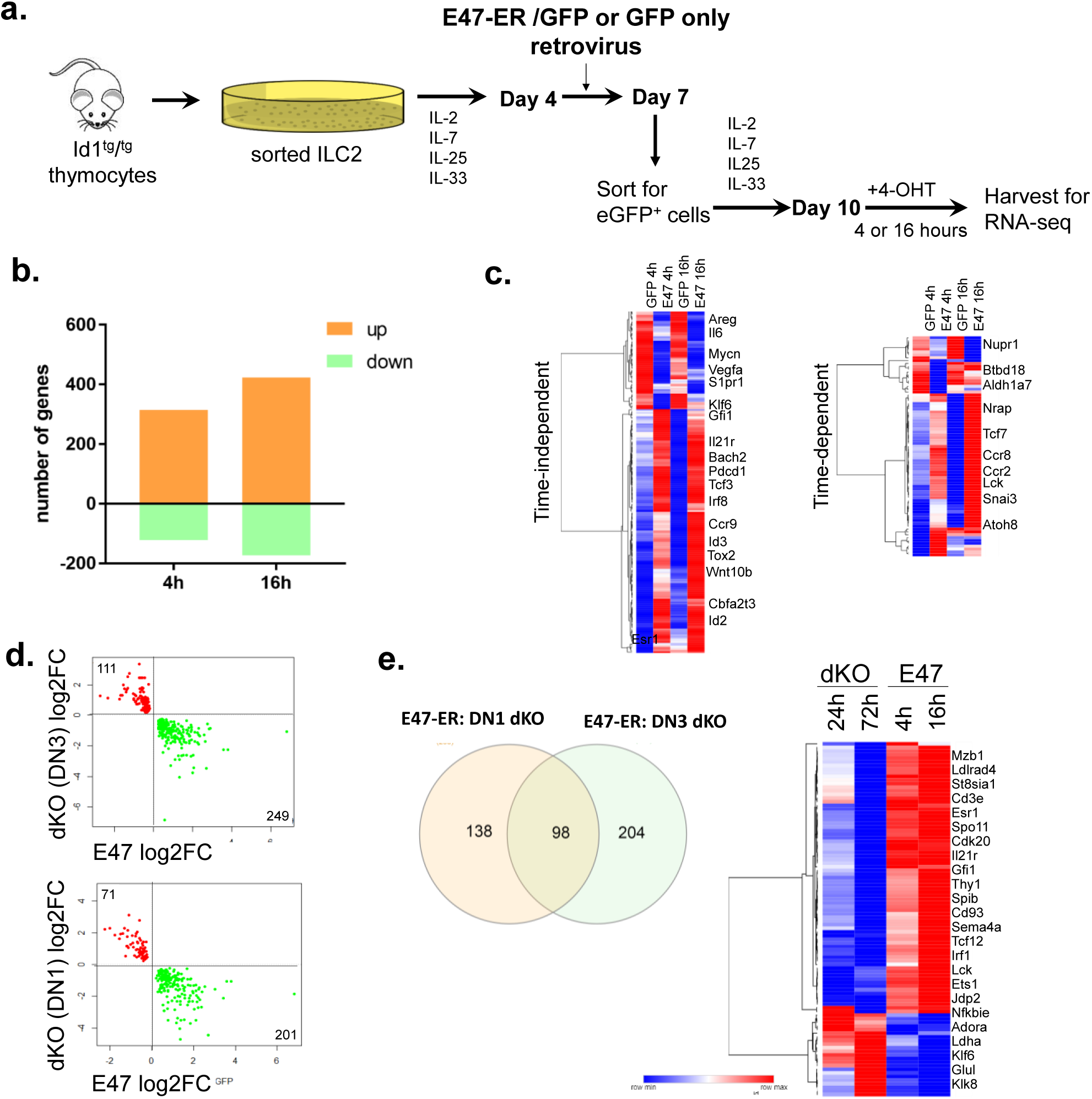
Transcriptome changes upon inducible expression of E47-ER. **(a)** Experimental scheme. **(b)** Numbers of differentially expressed genes comparing E47-ER/EGFP to EGFP-transduced cells at 4h and 16h. Genes with LFC > 0.5 and FDR < 0.05 were counted. **(c)** Expression levels of genes altered similarly (time-independent) and differently (time-dependent) at 4h and 16h are separately shown in heatmaps with genes of interest listed. **(d)** Gene expression changed in opposite direction in E protein deficient and E47-ER expressing cells are plotted. FDR<0.08. **(e)** Venn diagram shows the number of shared protein-coding genes in (d) between DN1 and DN3-derived cells. Relative levels of the 98 common genes are displayed in the heatmap with representatives listed.

At 4 and 16 hours post induction, 436 and 596 genes were found to be differentially regulated, respectively (Fig. 2b; supplemental table 1). In both cases, many more genes were found to be up-regulated by E47 than down-regulated. The differential genes can be divided into two sets: time-independent and time dependent (Fig. 2c). The former includes genes whose expression changed statistically similarly at 24h and 72h, which may be indicative of direct effects of E47. The latter consist of genes which exhibit slower kinetics in their activation or are indirectly affected by E47.

To narrow down the list of genes controlled by E proteins, we compared genes altered by loss of E proteins in either DN1 or DN3 cells (Fig. 1) to those affected by E47 overexpression in ILC2s, aiming to find genes altered in opposite directions. Genes that passed the threshold of combined log2 fold change (LFC) > 1 and false discovery rate (FDR) < 0.05 were obtained (Fig. 2d; supplemental table 1). Between these two sets, we found 98 protein coding genes in common, which might be more relevant to the suppression of ILC2 differentiation (Fig. 2e). It is noted that the majority of the genes in this group are known to be direct targets of E proteins such as *Tcf12, Cdk20, Gfi1, and Ets1* (Fig. 1e). We reasoned that shortage of down-regulated genes found in this assay was because the 16-hour window is too narrow to detect decreases in mRNA levels, which heavily depend on the repression of transcription by newly made repressors and turnover of existing transcripts.

### Dynamic changes in chromatin accessibility upon inducible E47 expression

To better understand how E proteins inhibit ILC2 differentiation or function, we need to further examine the genes directly or indirectly down-regulated by overexpression of E47 in ILC2s. We thus determined the chromatin accessibility in ILC2s using the technique called Assay for Transposase-Accessible Chromatin using sequencing (ATAC-seq) (14). We used retroviral transduced cells carrying E47-ER or a control vector treated as described in Fig. 2a for ATAC-seq.

Using a threshold of over 2-fold change with a FDR < 0.01, we found about 1300 peaks increased by E47 at 4h but less than 100 peaks decreased (Fig. 3b; supplemental table 1). However, by 16h, 815 peaks were elevated by E47 but 4735 peaks were reduced, which suggest major changes in chromatin accessibility, resulting in transcriptional repression (Fig. 3b; supplemental table 1). These changes correspond to less than 2% of all peaks detected, suggesting that the assays were specific to E47 activity rather than nonspecific effects. Approximately 80% of differential peaks are associated with protein coding genes. Regarding genomic distribution, most peaks reside in the intronic and intergenic regions (Fig. 3c), consistent with the primary roles of E proteins and other transcription factors in regulating transcription through enhancers or super-enhancers.

**Figure 3.**
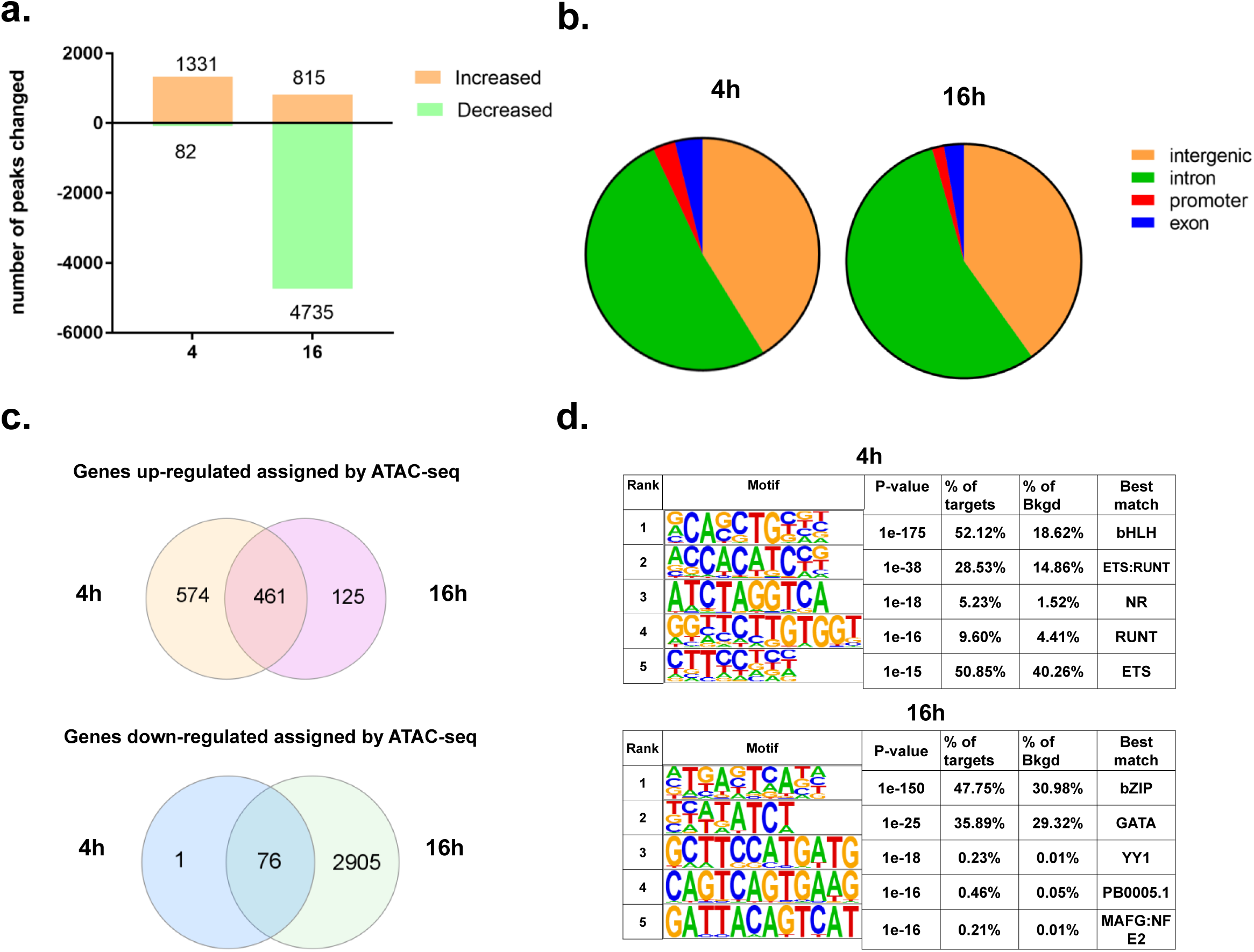
ATAC-seq analyses in E47-ER expressing cells. (a) Cells used in ATAC-seq analyses were treated as described in Fig. 2a. Numbers of peaks whose levels were increases or decreased comparing E47-ER expressing and control cells at 4h and 16h are as indicated. The cutoffs for peak calling are LFC > 1 and FDR<0.01. (b) Distribution of differential peaks across the genome at each time point. (c) Numbers of genes assigned to the peaks and those that overlap between the two time points. (d) Tables showing the top 5 most enriched transcription factor binding motifs in differentially accessible peaks at 4h and 16h. Binding sites that best match the transcription factor families are shown in the right column. NR stands for nuclear receptors.

The differential peaks obtained between E47-ER and control cells were then assigned to the nearest annotated gene. The numbers of up-regulated genes are 973 and 666 at 4h and 16h post tamoxifen addition, respectively. Among them, 483 genes are common at both time points. The smaller number of up-regulated genes at 16h might be attributed to negative feedback mechanisms. In contrast, the numbers of down-regulated genes were drastically different: 82 versus 2635 at 4h and 16h respectively. Most of the genes found to be repressed at 4h remained down-regulated at 16h. Since E47 is best known to be a transcriptional activator, the profound increase in genes inhibited by E47 induction at 16h suggests the induction of transcriptional repressors by E proteins.

Analyses of transcription factor binding sites within differential peaks reveals that at 4h, E boxes recognized by basic helix-loop-helix transcription factors such as E proteins were most abundant, representing over 50% of the binding sites found compared to a background of 18% in all open chromatin regions (Fig.3d). This suggests the direct involvement of E47 in the increased chromatin accessibility observed (Fig. 3b). Interestingly, by 16h, the predominant binding sites found in the differential peaks have shifted to the sites bound by bZip and GATA transcription factors. The known motif analysis revealed binding sites for different subgroups of bZip proteins including ATF3, BATF, JUN, FOS, BACH2 and MAF (supplemental Fig. 1a). The large increase in the number of down-regulated regions (∼4,735 peaks) may result from the secondary effects of the gene products initially elevated by E47 induction. These repressed motifs predominated over the E boxes directly associated with E47 binding. Indeed, when only elevated peaks were analyzed for their binding sites, E boxes remained the most prevalent (data not shown). Together, these analyses illustrated a dynamic change in chromatin accessibility within the 16 hours after induction of E47 activity.

### Genes initially activated by E47 may lead to repression of ILC2 genes later

We next correlated the ATAC-seq data with the RNA-seq results in EGFP or E47-ER/EGFP retrovirus transduced ILC2s after the same treatments with tamoxifen. We looked for genes whose expression patterns changed in the same manner upon E47 induction. The thresholds of ATAC-seq data were set at a LFC > 1 and FDR < 0.01. Since we reasoned that changes in mRNA levels, particularly the reduction of transcripts, may be slower than alterations in chromatin accessibility, the threshold for the RNA-seq data was set at FDR < 0.05 without the LFC cutoff to maximize the detection of any trend of gene expression (Fig. 4a). Four hours after E47 induction, the majority of the common differential genes are up-regulated. In contrast, by 16h, the majority of genes were downregulated. Additionally, we observed a significant fraction of genes that were positively increased by RNA-seq but negatively regulated by ATAC-seq, which may illustrate the enhanced sensitivity of ATAC-seq in detection of dynamic gene repression at 16h. The delayed appearance of down-regulated genes supports our hypothesis that E47 indirectly causes gene repression.

**Figure 4.**
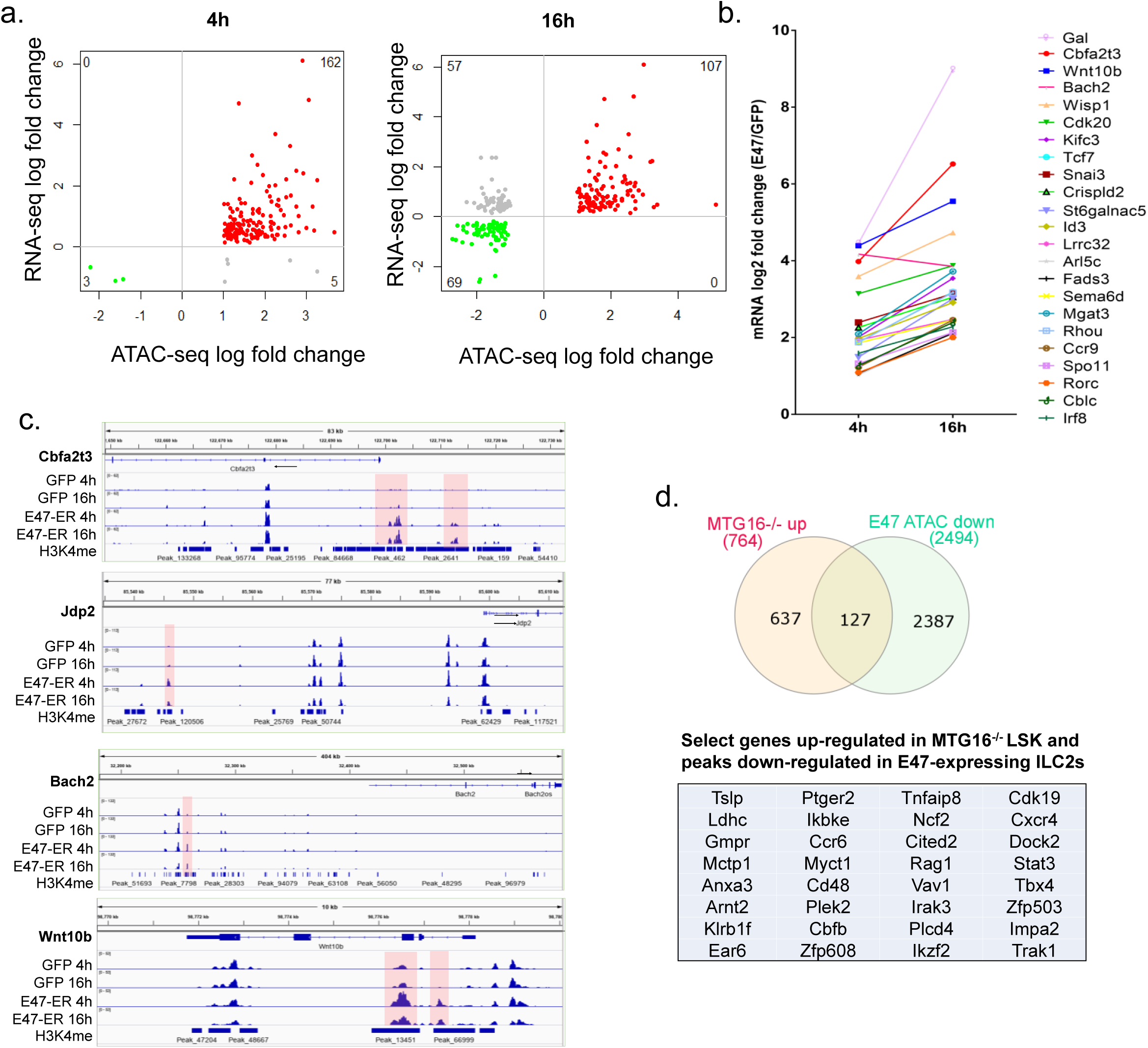
Initial transcriptional activation followed up widespread gene repression. (a) Correlation of mRNA levels and chromatin accessibility. LFC of ATAC peaks and mRNA levels in E47-ER expressing cells at each time point are plotted on x and y axis, respectively. (b) Top up-regulated genes at 4h and 16h are plotted. (c) ATAC peak tracks at indicated time points and cells are shown for representative genes as indicated. H3K4me ChIP-seq data were generated using thymocytes by the ENCODE consortium. (d) Comparison of genes repressed by MTG16 (encoded by *Cbfa2t3*) in hematopoietic stem cells (LSK) and genes repressed in E47-ER expressing ILC2s at 16h. Fisher test shows the statistical significance of the overlap (p<0.00001). A select list of the genes are shown below the Venn diagram.

We then set out to identify putative candidates that may mediate the profound chromatin remodeling observed at 16h. Fig. 4b shows a list of more dramatically up-regulated genes at both 4h and 16h. Among them, several transcriptional repressors stood out. The signal tracks of a few select genes are displayed in Fig. 4c. For example, expression of *Cbfa2t3*, which encodes MTG16 (also called ETO2), was significantly enhanced at 4h and continued to increase by 16h. The differential peaks were detected approximately 1kb, 3kb and 13kb upstream of the transcription start site. These peaks overlap with the region observed in chromatin immunoprecipitation against methylated K4 of histone 3 (H3K4me) in thymocytes, suggesting that these regions may contain enhancers. MTG16 is a known transcriptional repressor capable of recruiting histone deacetylases (15;16). It not only interacts with E proteins but also forms super complexes with GATA1, E proteins, Tal1, Lmo2 and Ldb in erythroid cells (17;18). MTG16 is also important for T cell development, as well as the differentiation of plasmacytoid dendritic cells (19;20). It is thus conceivable that E proteins recruit MTG16 and cooperate with GATA3 to suppress the ILC2 fate in T cell precursors where these proteins all exist. Interestingly, *Cbfa2t3* has been shown to be expressed at very low levels in ILC2Ps compared to their precursors in the bone marrow (supplemental Fig. 1b) (21), suggesting that down-regulating this gene is necessary for ILC2 differentiation. ILC2s in small intestine also express *Cbfa2t3* at a low level.

Furthermore, we found that 127 genes up-regulated in *Cbf2at3*^*-/-*^ hematopoietic stem cells (LSK) overlap with the genes assigned to diminished peaks at 16h in ILC2s (*p* < 2.2e-16) (Fig. 4d). Considering the difference of the cell types in which the two sets of data were generated, the overlap is remarkable and supports the repressive role of *Cbfa2t3* products in ILC2s.

However, there are likely to be other repressors which mediate the suppressive function of E proteins in addition to MTG16. Jdp2 is a transcriptional repressor dimerizing with the Jun family of leucine-zipper transcription factors (22), many of which are expressed in ILC2s. Jdp2 can recruit histone deacetylase 3, thus regulating histone modification and chromatin assembly (23;24). This is particularly relevant in our study because we observed widespread decreases in ATAC peaks at 16h, and these peaks are predominantly populated with bZip transcription factor binding sites (Fig. 3e; supplemental Fig. 1a). Thus, Jdp2 is a probable candidate for mediating gene repression at 16h. Another up-regulated bZip transcription repressor is Bach2 (25). Its binding sites are readily detectable in ATAC peaks at 16h (Supplemental Fig. 1a).

Considering the inhibition of Th2 differentiation by Bach2 (26;27), it could also contribute to the suppression of the ILC2 fate.

Finally, several genes encoding proteins related to the Wnt signaling pathway, *Wnt10b, Wisp* and *Snai*, are on top of the list of up-regulated genes shown in Fig. 4b. Given the crucial roles of Wnt signaling in regulating various developmental programs (28;29), it may play an important role in ILC2 suppression.

### Down-regulation of a network of transcriptional programs for ILC2 differentiation by E47

The large amount of ATAC peaks found to be reduced by E47 at 16h may provide plausible hints as to how E proteins suppress ILC2 differentiation. We thus performed pathway analysis on the genes regulated in similar manners at 16h as detected using RNA-seq and ATAC-seq (Fig. 4a). Consistent with our expectations, the top canonical pathway found to be down-regulated by E47 is related to T helper 2 cells. The top-ranking network of transcriptional programs includes key transcription factors and signaling molecules known to be ILC signatures such as IL-7R, TOX, IRF4, MAF, KLF6 and RBPJ (Fig. 5a) (4;5;21;30). The reduction in chromatin accessibility of the key ILC2 signature genes is striking. For example, 15 peaks assigned to *Maf* were found to be diminished by at least 2 fold (supplemental Fig. 2). Likewise, 3 peaks each were reduced for *Tox* and *Irf4*, respectively.

**Figure 5.**
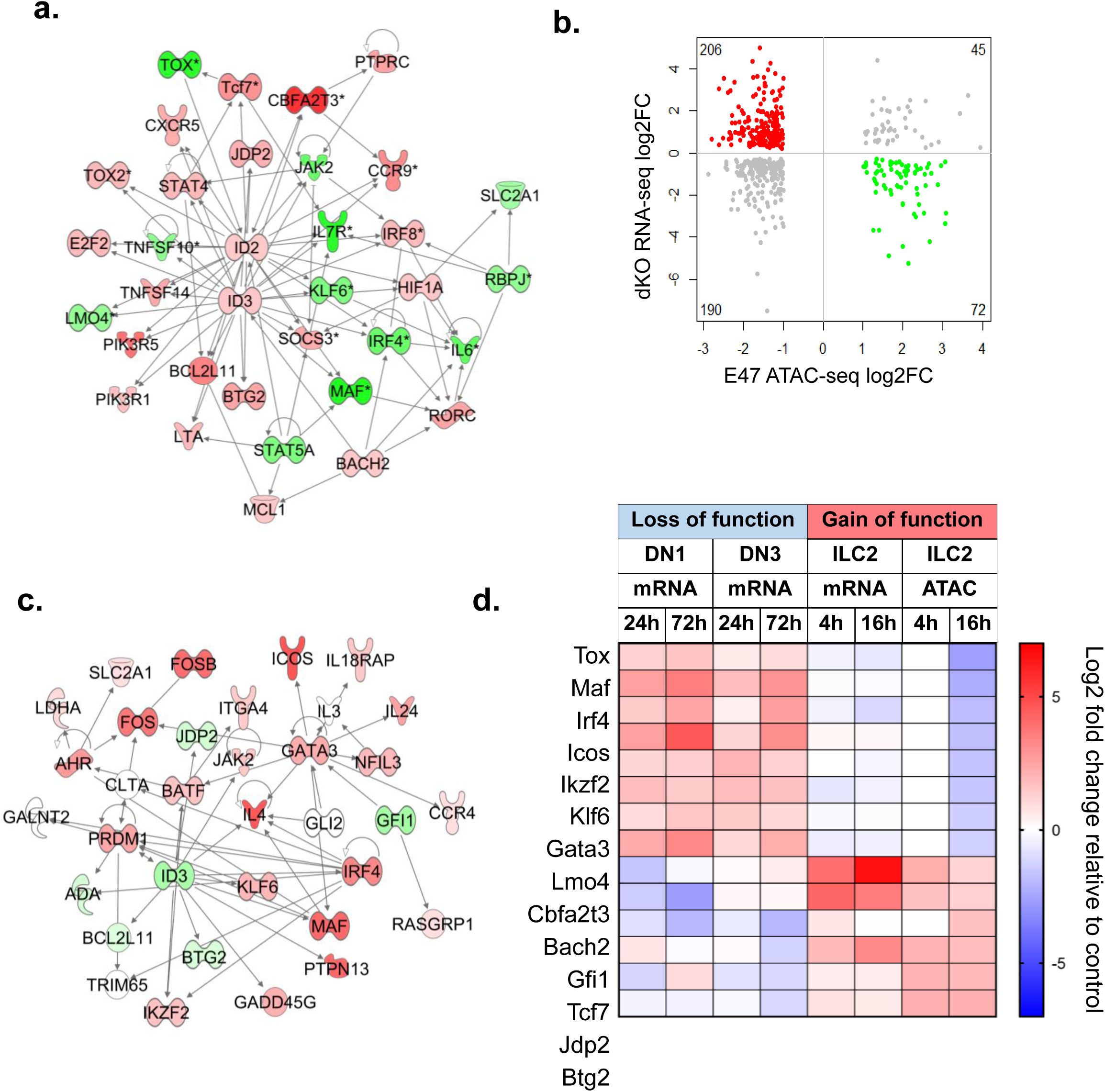
Loss and gain of E protein function impact a similar transcription program regulating ILC2 differentiation. (a) Ingenuity pathway analysis of differential peaks detected at 16h after E47-ER induction. Top transcription network (Score=45) is shown. Up and down-regulated genes are depicted in red and green. Multiple peaks assigned to the same gene is indicated with an asterisk. Arrows show connections between two genes, mostly by positive or negative influence in expression. (b) Inverse correlation between the effects of loss and gain of E protein function. LFC of differentially expressed genes in both DN1 and DN3-derived cells is plotted against the changes in E47 expressing cells at 16h. Genes increased and decreased in E protein deficient cells are highlighted in red and green, respectively. (c) Top network identified using the ingenuity pathway analysis of inversely correlated genes as shown in (b) (Score=38). (d) Summary of the expression profiles of key genes regulated by E proteins in relation to ILC2 differentiation.

*Id2* and *Id3* were up-regulated by E47 in the treated cells because these genes are known to be activated by E proteins in a feedback loop that helps maintain the net activity of E proteins (31). Since E47-ER overexpressed in the cells would likely outcompete the endogenous Id proteins, the small increases in Id2 and Id3 mRNA levels may not be consequential. However, the major nodes formed around Id2 and Id3 highlight the importance of the helix-loop-helix family of transcription factors such as E47 in regulating ILC2 differentiation and their interplay with other transcription factors (Fig. 5a).

To correlate the ATAC-seq data with gene expression changes caused by E protein ablation in both DN1 and DN3-initiated ILC2 cultures (Fig. 1), we plotted fold changes in mRNA levels and ATAC-seq peaks (Fig 5b). We focused on the inverse relationship between the two datasets. There are 206 and 72 genes up- and down-regulated upon E protein gene deletion, which are assigned to peaks decreased and increased in ATAC-seq, respectively. Ingenuity pathway analysis of these commonly regulated genes revealed the top regulatory network consisting of key regulatory proteins in ILC2 differentiation. The same molecules found to be down-regulated by E47 expression in ILC2s, IRF4, MAF and KLF6, are shown to be up-regulated during ILC2 differentiation of E protein deficient DN1 and DN3 thymocytes (Fig. 5a and 5c). Conversely, genes activated by E47 are diminished with E protein ablation, which include *Id3, Jdp2, Jak2, Bcl2l11* and *Btg2*. In addition, several more genes encoding transcription factors potentially important for ILC2 differentiation, such as *Gata3, Batf, Nfil3, Ahr, Ikzf2, Prdm1, Fos* and *Fosb* were also deemed to be up-regulated in E protein deficient DN1 and DN3 cells and down-regulated by E47 as detected by ATAC-seq.

Collectively, this study uses inducible loss and gain of E protein function models to cross-examine transcription programs in the context of ILC2 differentiating cells or ILC2s and the consensus is summarized in Fig. 5d. By following changes in mRNA levels or chromatin accessibility shortly after alterations in E protein activity, we have a unique opportunity to postulate a cascade of regulatory events that lead to E protein-mediated suppression of ILC2 differentiation. Induction of E47 in ILC2s quickly led to the up-regulation of genes like *Cbfa2t3* and *Jdp2*, which in turn cause coordinated transcriptional repression of a network of transcriptional programs important for the ILC2 cell fate.

E proteins are known to promote B and T lymphoid cell commitment, partly through their ability to facilitate the production of proteins involved in antigen receptor signaling. However, simply pushing the B and T cell-specific function may not be sufficient and suppression of other cell fates such as the myeloid and innate lymphoid paths could also be necessary (32;33). Conversely, the differentiation into ILC2 or other ILCs requires the inhibition of E protein function by their inhibitors, the Id proteins. Id2 is well known to be essential of ILC differentiation (34). Id2 and sometimes Id3 may act by antagonizing the E protein activities we have elucidated in this study to promote ILC differentiation from in common lymphoid progenitors in the bone marrow or T cell precursors in the thymus.

## Methods

### Mice and OP9 stromal culture

ROSA26-CreERT2;ROSA26-Stop-tdTomato;*Tcf3*^f/f^;*Tcf12*^f/f^ (dKO) mice and controls lacking to floxed *Tcf3* and *Tcf12* are as described (5;7). CLP from the bone marrow and DN1 or DN3 cells from the thymus were sorted and cultured as described (5). Briefly, cells were seeded at 100 or 1000 cells/well in 48-well plates containing a layer of OP9-DL1 stromal cells plated 24 hours before. The culture medium contains 20% bovine fetal calf serum (FCS), 30ng/ml of each IL-2, IL-7 and stem cell factor (all from R&D systems) in α-MEM (Gibco, ThermoFisher). On day 4, 4-hydroxy-tamoxifen (4-OHT) was added to a concentration of 1 µM. Cells were harvested 24 and 72 hours later and CD45^+^tdTomato^+^ were sorted and used for RNA sequencing.

### Retroviral transduction in ILC2s

To purify ILC2s from the thymus of plck-Id1^tg/tg^ mice, thymocytes were stained with anti-CD90.2 (53-2.1), anti-ST2 (DIH9), anti-ICOS (C398.4A) and a cocktail of lineage (Lin) specific antibodies: anti-FcεR (MAR-1), anti-B220 (RA3-6B2), anti-CD19 (ID3), anti-Mac-1 (M1/70), anti-Gr-1 (R86-8C5), anti-CD11c (N418), anti-NK1.1 (PK136), anti-Ter-119 (Ter-119), anti-CD3ε (145-2C11), anti-CD5 (53-7.3), anti-CD8α (53-6.7), anti-TCRβ (H57-597), and anti-γδTCR (GL-3; ebioscience). All antibodies are from Biolegend unless specified otherwise. Lin-CD90.2+ICOS+ST2+ cells were sorted on a FACSAria II (BD Biosciences). The cells were plated on 96-well U-bottom plates at 80,000 cells/well in α-MEM medium containing 10% FCS, 10ng/ml each IL-2, IL-7, IL-25 and IL-33. Cells were split 1:3 on day 3 and placed onto 96-well flat-bottom plates and spin-infected with retrovirus the next day. Spin-infection was performed by mixing retroviral stocks and cultured cells at a 1:1 ratio plus the addition of polybrene to a concentration of 8 µg/ml. The retroviral stocks are as described (13). The cells were centrifuged for 1.5 hours at 1,300 rpm before the plates were returned to the 37°C incubator with 5% CO_2_. After 6 hours, the culture was replaced with fresh medium and incubated for 2 days. Transduced cells were sorted for the expression of EGFP and cultured for 3 additional days in U-bottom plates with the same medium. These cells with or without E47-ER were induced with 1 µM 4-OHT for 4 or 16 hours and harvested.

### RNA and ATAC sequencing

RNA was isolated using the Trizol reagent per vender’s instruction (Life Technology). RNA sequencing (RNA-seq) was performed in duplicate by the Clinical Genomics Center at the Oklahoma Medical Research Foundation (OMRF) using the Ovation RNA-Seq System (NuGEN Technologies Inc., San Carlos, CA) followed by KAPA Hyper library preparation kits (KAPA Biosystems, Wilmington, MA) on an Illumina NextSeq 500 sequencing platform with 76-bp, paired-end reads (Illumina, San Diego, CA).

ATAC-sequencing was also performed in duplicate for each time point and cell type. ATAC transpose reactions were carried out using the NX#-TDE1, TAGMENT DNA enzyme and buffer from Illumina. The reaction was purified using Qiagen miniElute columns, which is followed by PCR amplification with matching primers. The PCR products were purified using magnetic beads and eluted in EB buffer (Qiagen). The products were sequenced using an Illumina NextSeq 500 sequencing platform with 76-bp, paired-end reads.

### Analyses of RNA sequencing data

Raw sequencing reads (in a FASTQ format) were trimmed using Trimmomatic to remove any low-quality bases at the beginning or the end of sequencing reads as well as any adapter sequences (Bolger et al., 2014). Trimmed sequencing reads were aligned to the Mus musculus genome reference (GRCm38/mm10) using STAR v.2.4.0h (35). Gene-level read counts were determined using HTSeq v.0.5.3p9 (36) with the GENCODE Release M10 (GRCm38) annotation. Only autosomal genes coding for lncRNAs, miRNAs, and protein-coding mRNAs were selected for analyses.

Read-count normalization and differentially expressed analyses was performed using the edgeR package from Bioconductor. Expression values normalized with the voom function were analyzed for differential expression using the standard functions of the limma program. Each transcript expression variation was tested with the moderated t-statistics and the corresponding p values were adjusted for multiple testing using false discovery rate (FDR). Unless specified, sets of differentially expressed transcripts were filtered requiring at least two fold change in expression and a FDR below 0.05. Overrepresented functional gene sets (GO, KEGG pathways) were identified using specialized R Bioconductor packages. Ingenuity Pathway Analysis (IPA, QIAGEN, Redwood City CA) was used to explore significant gene networks and pathways interactively.

### ATAC Data Processing and Analysis

Alignment and peak-calling of ATAC sequencing data were performed using the ENCODE-DCC ATAC-Seq pipeline (https://github.com/ENCODE-DCC/atac-seq-pipeline (37). Briefly, data were aligned to mm10 genome using Bowtie2 (38). Aligned data were filtered for PCR duplicates, blacklisted regions, and shifted +4bp for the + strand and - 5bp for the – strand. Peaks were called using MACS2 (39) and filtered by Irreproducible Discovery Rate (IDR < 0.1) (https://github.com/nboley/idr). Differential peak calling was performed using differential binding analysis of ChIP-Seq peak data (DiffBind, http://bioconductor.org/packages/release/bioc/vignettes/DiffBind/inst/doc/DiffBind.pdf.) (40), where a consensus peakset of 60,543 peaks was derived from all samples, and a window of 150 bp was centered on peak summits for further downstream analysis. DESeq2 (41) was used to perform differential peak analysis with DE peaks identified by q < 0.05 and absolute log fold-change > 1. Motif enrichment was performed using Hypergeometric Optimization of Motif EnRichment program (HOMER) version 4.1 (42), where the background was set as the union of all ATAC peaks.

## Author Contribution

V.P. and C.G. carried out bioinformatics analyses; A.B., L.Q. and X.-H. S. generated the data. V.P., C.G., J.D.W. and X.-H. S. analyzed the data and wrote the manuscript.

## Acknowledgements

We are grateful to Dr. Graham Wiley for advice in the ATAC-seq experiments and to the Clinical Genomics Center and the Flow cytometry facility at the Oklahoma Medical Research Foundation for outstanding technical assistance. We thank Dr. Marco Colonna for advice and support. This work was supported by grants from the NIH (1R01AI126851), and the Presbyterian Health Foundation to X.-H.S. X.-H.S holds the Lew and Mira Ward Chair in Biomedical Research at the Oklahoma Medical Research Foundation. V.P. is supported by NIH T32DK077653-28.

## Figure legends

**Supplemental Figure 1.**
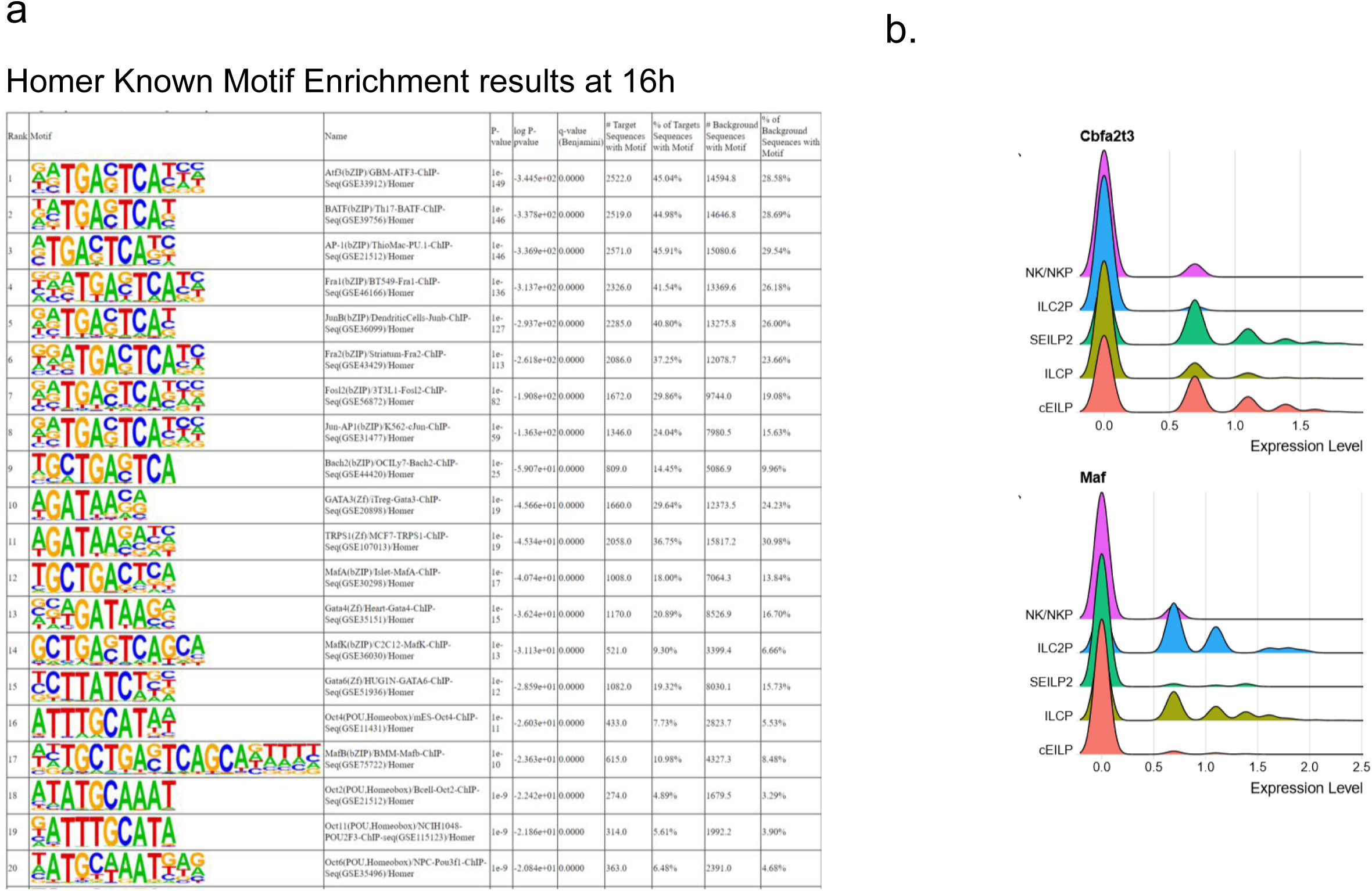
Additional analyses of ATAC-seq data. **(a)** Known motif analysis of the ATAC peaks at 16h. Top 20 motifs are shown. **(b)** Analyses of single cell sequencing data of bone marrow ILC progenitors for indicated genes. SEILP2: specified early innate lymphoid progenitors cluster 2; cEILP committed EILP, ILCP: innate lymphoid cell progenitor; ALP: all lymphoid progenitors.

**Supplemental Figure 2.**
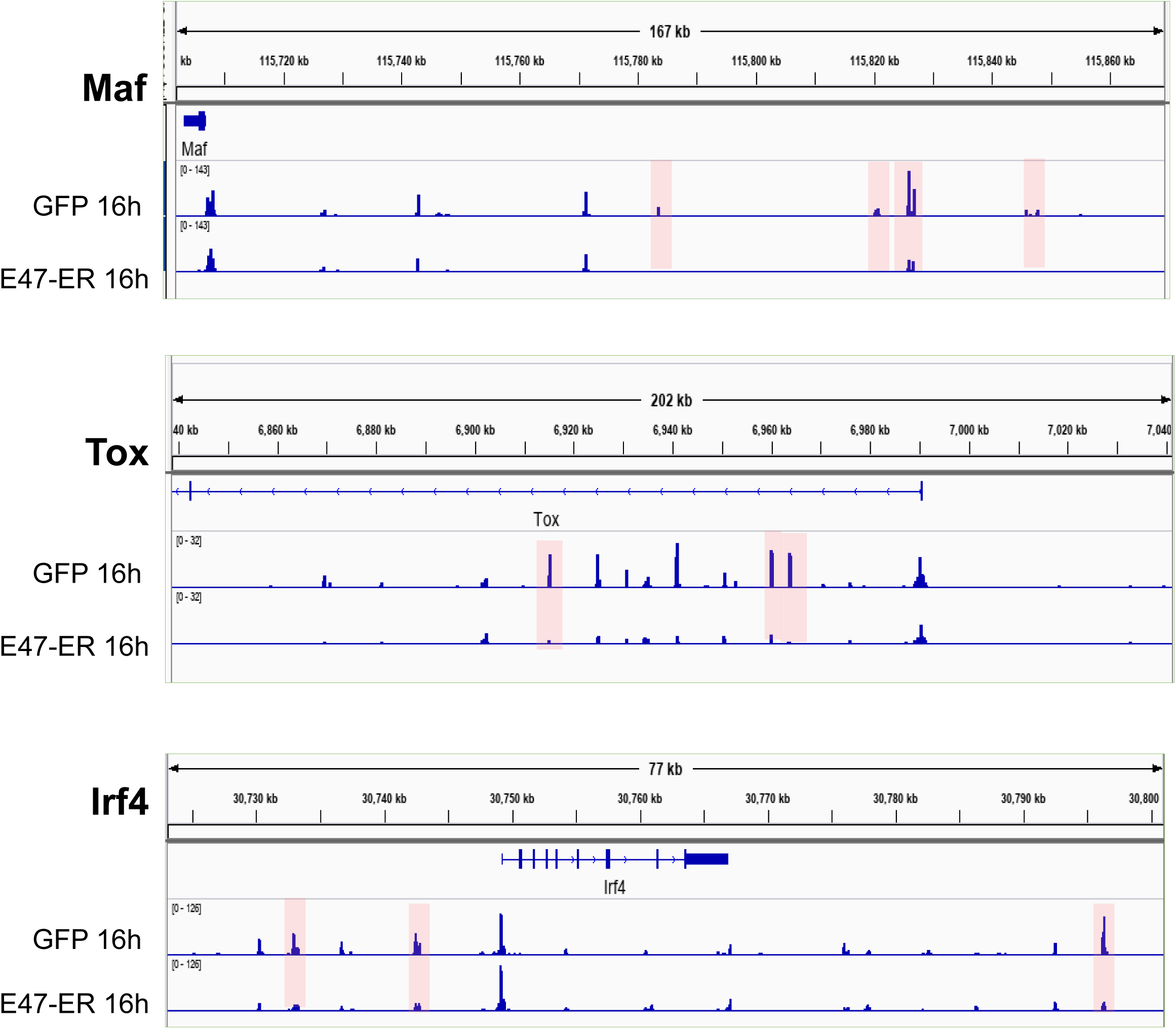
ATAC-seq peak tracks detected at 16h. Genes of interest are listed on the left. Representative differential peaks for *Maf* are shown.

