## Supplemental table 1 for "E proteins orchestrate dynamic transcriptional cascades to suppress ILC2 differentiation"

| dKO 24h | Con 24h | dKO 72h | con 72h | sample_1 | sample_2 | logFC | test_stat | p_value | q_value | pvlltr | Geneid | biotype | SYMBOL | GENENAME | ENTREZID | Chr | Start | End | Strand | UniGene |
|---|---|---|---|---|---|---|---|---|---|---|---|---|---|---|---|---|---|---|---|---|
| 6 | 7.92 | 5.95 | 7.88 | DN1ab | DN1cd | 1.92 | 39.86 | 1.50E-09 | 2.50E-05 | 0.91 | ENSMUSG00000048758 | protein_coding | Rpl29 | ribosomal protein L29 | 19944 | 9 | 106429454 | 106431568 | 1 | Mm.288696 Mm.359377 Mm.431965 |
| 7.02 | 1.65 | 7.18 | 1.4 | DN1ab | DN1cd | -5.57 | -37.69 | 2.20E-09 | 2.50E-05 | 0.18 | ENSMUSG00000082399 | processed_pseudogene | Gm14036 | predicted gene 14036 | NA | 2 | 134803949 | 134804428 | 1 | |
| 5.56 | 6.88 | 5.56 | 6.88 | DN1ab | DN1cd | 1.32 | 27.2 | 2.10E-08 | 0.00016 | 0.98 | ENSMUSG00000097906 | processed_pseudogene | Gm9625 | predicted gene 9625 | NA | 13 | 67810246 | 67811200 | 1 | |
| 10.34 | 3.34 | 10.57 | 3.28 | DN1ab | DN1cd | -7.16 | -26.01 | 2.90E-08 | 0.00017 | 0.64 | ENSMUSG00000086503 | lincRNA | Xist | inactive X specific transcripts | NA | X | 103460366 | 103483254 | -1 | |
| 4.76 | 6.32 | 5.43 | 7.04 | DN1ab | DN1cd | 1.58 | 21.4 | 1.10E-07 | 0.00049 | 0.82 | ENSMUSG00000040345 | protein_coding | Arhgap9 | Rho GTPase activating protein 9 | 216445 | 10 | 127321964 | 127329943 | 1 | Mm.227198 |
| 2.67 | 5.99 | 2.29 | 5.83 | DN1ab | DN1cd | 3.38 | 21.01 | 1.30E-07 | 0.00049 | 0.57 | ENSMUSG00000074516 | processed_pseudogene | Gm10709 | predicted gene 10709 | NA | 9 | 7751725 | 7752204 | 1 | |
| 6.99 | 8.67 | 6.93 | 8.9 | DN1ab | DN1cd | 1.78 | 19.11 | 2.50E-07 | 0.00081 | 0.15 | ENSMUSG00000059751 | processed_pseudogene | Rps3a3 | ribosomal protein S3A3 | NA | 13 | 108670603 | 108671397 | 1 | |
| 4.91 | 6.32 | 4.71 | 6.13 | DN1ab | DN1cd | 1.41 | 18.6 | 3.00E-07 | 0.00083 | 0.98 | ENSMUSG00000078154 | protein_coding | Gm12184 | predicted gene 12184 | NA | 11 | 48825346 | 48826655 | -1 | |
| 6.16 | 3.12 | 6.85 | 4.19 | DN1ab | DN1cd | -2.84 | -18.35 | 3.30E-07 | 0.00083 | 0.24 | ENSMUSG00000052336 | protein_coding | Cx3cr1 | chemokine (C-X3-C motif) receptor 1 | 13051 | 9 | 119901616 | 120069879 | -1 | Mm.44065 |
| -3.82 | 3.44 | -2.81 | 3.8 | DN1ab | DN1cd | 7.07 | 17.93 | 3.80E-07 | 0.00088 | 0.5 | ENSMUSG00000056673 | protein_coding | Kdm5d | lysine (K)-specific demethylase 5D | 20592 | Y | 897788 | 956786 | 1 | Mm.262676 |
| 9.32 | 8.06 | 9.45 | 7.92 | DN1ab | DN1cd | -1.36 | -17.69 | 4.20E-07 | 0.00088 | 0.079 | ENSMUSG00000066362 | processed_pseudogene | Rps13-ps1 | ribosomal protein S13, pseudogene 1 | NA | 8 | 87047689 | 87048226 | 1 | |
| 6.45 | 5.42 | 6.43 | 5.38 | DN1ab | DN1cd | -1.04 | -16.87 | 5.90E-07 | 0.001 | 0.86 | ENSMUSG00000036046 | protein_coding | 5031439G07Rik | RIKEN cDNA 5031439G07 gene | 223739 | 15 | 84943936 | 84988551 | -1 | Mm.323925 |
| -3.82 | 0.45 | -3.94 | 0.92 | DN1ab | DN1cd | 4.47 | 16.69 | 6.30E-07 | 0.001 | 0.33 | ENSMUSG00000101059 | processed_pseudogene | Gm4017 | predicted gene 4017 | NA | Y | 1268548 | 1268885 | -1 | |
| 3.08 | 4.79 | 3.6 | 5.44 | DN1ab | DN1cd | 1.76 | 15.63 | 9.90E-07 | 0.0015 | 0.61 | ENSMUSG00000027994 | protein_coding | Mcub | mitochondrial calcium uniporter dominant negative beta subunit | 66815 | 3 | 129914960 | 129970206 | -1 | Mm.31056 |
| 2.21 | 5.67 | 3.44 | 6.23 | DN1ab | DN1cd | 3.25 | 15.15 | 1.20E-06 | 0.0018 | 0.16 | ENSMUSG00000005125 | protein_coding | Ndrg1 | N-myc downstream regulated gene 1 | 17988 | 15 | 66929318 | 66969640 | -1 | Mm.30837 |
| 8.7 | 7.71 | 8.97 | 8.2 | DN1ab | DN1cd | -0.9 | -14.56 | 1.60E-06 | 0.0019 | 0.069 | ENSMUSG00000040699 | protein_coding | Limd2 | LIM domain containing 2 | 67803 | 11 | 106156256 | 106160860 | -1 | Mm.21687 |
| 3.65 | 5.3 | 3.84 | 5.58 | DN1ab | DN1cd | 1.68 | 14.53 | 1.60E-06 | 0.0019 | 0.75 | ENSMUSG00000022307 | protein_coding | Oxr1 | oxidation resistance 1 | 170719 | 15 | 41447482 | 41861048 | 1 | Mm.254267 |
| 0.04 | 6.51 | -0.47 | 6.53 | DN1ab | DN1cd | 6.62 | 14.49 | 1.70E-06 | 0.0019 | 0.63 | ENSMUSG00000069045 | protein_coding | Ddx3y | DEAD (Asp-Glu-Ala-Asp) box polypeptide 3, Y-linked | 26900 | Y | 1260771 | 1286629 | -1 | Mm.302938 |
| -0.26 | -3.86 | -0.11 | -4.02 | DN1ab | DN1cd | -3.72 | -14.49 | 1.70E-06 | 0.0019 | 0.59 | ENSMUSG00000083611 | processed_pseudogene | Gm8832 | predicted gene 8832 | NA | X | 168366464 | 168366858 | 1 | |
| 6.37 | 7.68 | 6.58 | 8.07 | DN1ab | DN1cd | 1.37 | 14.15 | 2.00E-06 | 0.0021 | 0.4 | ENSMUSG00000020484 | protein_coding | Xbp1 | X-box binding protein 1 | 22433 | 11 | 5520659 | 5525893 | 1 | Mm.469937 |
| -2.83 | 4.97 | -1.56 | 5.36 | DN1ab | DN1cd | 7.55 | 14.12 | 2.00E-06 | 0.0021 | 0.5 | ENSMUSG00000068457 | protein_coding | Uty | ubiquitously transcribed tetratricopeptide repeat gene, Y chromosome | 22290 | Y | 1096861 | 1245759 | -1 | Mm.20477 |
| 6.76 | 5.64 | 7.35 | 6.17 | DN1ab | DN1cd | -1.15 | -13.9 | 2.20E-06 | 0.0022 | 0.71 | ENSMUSG00000006519 | protein_coding | Cyba | cytochrome b-245, alpha polypeptide | 13057 | 8 | 122424776 | 122432930 | -1 | Mm.271671 |
| -3.14 | 3.12 | -1.14 | 3.72 | DN1ab | DN1cd | 5.86 | 13.81 | 2.30E-06 | 0.0022 | 0.14 | ENSMUSG00000042757 | protein_coding | Tmem108 | transmembrane protein 108 | 81907 | 9 | 103482947 | 103761837 | -1 | Mm.384704 |
| 6.71 | 7.38 | 6.79 | 7.49 | DN1ab | DN1cd | 0.68 | 13.39 | 2.90E-06 | 0.0026 | 0.75 | ENSMUSG00000018899 | protein_coding | Irf1 | interferon regulatory factor 1 | 16362 | 11 | 53770014 | 53778374 | 1 | Mm.105218 |
| 5.69 | 4.84 | 6.7 | 5.78 | DN1ab | DN1cd | -0.88 | -13.35 | 2.90E-06 | 0.0026 | 0.63 | ENSMUSG00000051735 | protein_coding | Rinl | Ras and Rab interactor-like | 320435 | 7 | 28788969 | 28798963 | 1 | Mm.74632 |
| 8.15 | 9.2 | 7.69 | 8.95 | DN1ab | DN1cd | 1.12 | 13.11 | 3.30E-06 | 0.0028 | 0.27 | ENSMUSG00000026558 | protein_coding | Uck2 | uridine-cytidine kinase 2 | 80914 | 1 | 167222883 | 167285320 | -1 | Mm.280895 Mm.440156 |
| 4 | 0.04 | 5.33 | 0.61 | DN1ab | DN1cd | -4.39 | -13.04 | 3.40E-06 | 0.0028 | 0.29 | ENSMUSG00000030703 | protein_coding | Gdpd3 | glycerophosphodiester phosphodiesterase domain containing 3 | 68616 | 7 | 126766334 | 126775649 | 1 | Mm.246881 |
| 6.45 | 7.46 | 7.09 | 7.82 | DN1ab | DN1cd | 0.91 | 12.71 | 4.10E-06 | 0.0032 | 0.034 | ENSMUSG00000020027 | protein_coding | Socs2 | suppressor of cytokine signaling 2 | 216233 | 10 | 95385362 | 95417180 | -1 | Mm.474283 Mm.490329 |
| 4.86 | 6.39 | 4.37 | 6.18 | DN1ab | DN1cd | 1.62 | 12.49 | 4.60E-06 | 0.0033 | 0.35 | ENSMUSG00000038604 | protein_coding | Ripor1 | RHO family interacting cell polarization regulator 1 | 75687 | 8 | 105605255 | 105622219 | 1 | Mm.41261 |
| 2.26 | 4.59 | 3.24 | 4.9 | DN1ab | DN1cd | 2.11 | 12.44 | 4.70E-06 | 0.0033 | 0.046 | ENSMUSG00000045394 | protein_coding | Epcam | epithelial cell adhesion molecule | 17075 | 17 | 87635979 | 87651129 | 1 | Mm.4259 |
| 5.07 | 6.18 | 5.4 | 6.77 | DN1ab | DN1cd | 1.2 | 12.43 | 4.70E-06 | 0.0033 | 0.23 | ENSMUSG00000026721 | protein_coding | Rabgap1l | RAB GTPase activating protein 1-like | 29809 | 1 | 160219174 | 160793211 | -1 | Mm.25833 |
| 5.52 | 6.57 | 4.82 | 5.85 | DN1ab | DN1cd | 1.04 | 12.05 | 5.80E-06 | 0.0039 | 0.93 | ENSMUSG00000027714 | protein_coding | Exosc9 | exosome component 9 | 50911 | 3 | 36552606 | 36565727 | 1 | Mm.116711 |
| 5.74 | 3.71 | 6.85 | 4.82 | DN1ab | DN1cd | -2.03 | -11.87 | 6.50E-06 | 0.0041 | 0.98 | ENSMUSG00000024190 | protein_coding | Dusp1 | dual specificity phosphatase 1 | 19252 | 17 | 26505590 | 26508519 | -1 | Mm.239041 |
| 9.31 | 7.56 | 11.75 | 9.61 | DN1ab | DN1cd | -1.92 | -11.86 | 6.50E-06 | 0.0041 | 0.26 | ENSMUSG00000001020 | protein_coding | S100a4 | S100 calcium binding protein A4 | 20198 | 3 | 90603771 | 90606045 | 1 | Mm.3925 |
| 0.44 | 3.02 | 2.06 | 4.45 | DN1ab | DN1cd | 2.52 | 11.82 | 6.60E-06 | 0.0041 | 0.71 | ENSMUSG00000042156 | protein_coding | Dzip1 | DAZ interacting protein 1 | 66573 | 14 | 118875520 | 118925460 | -1 | Mm.87456 |
| -3.82 | -0.44 | -3.94 | -1.03 | DN1ab | DN1cd | 3.23 | 11.76 | 6.90E-06 | 0.0041 | 0.45 | ENSMUSG00000097699 | bidirectional_promoter_lncRNA | 5430400D12Rik | RIKEN cDNA 5430400D12 gene | NA | 7 | 79920723 | 79922521 | 1 | |
| 5.52 | 6.44 | 5.69 | 6.75 | DN1ab | DN1cd | 0.97 | 11.74 | 7.00E-06 | 0.0041 | 0.45 | ENSMUSG00000021108 | protein_coding | Prkch | protein kinase C, eta | 18755 | 12 | 73584796 | 73778185 | 1 | Mm.341677 |
| 3.61 | 1.66 | 5.25 | 3.42 | DN1ab | DN1cd | -1.88 | -11.64 | 7.40E-06 | 0.0041 | 0.73 | ENSMUSG00000038642 | protein_coding | Ctss | cathepsin S | 13040 | 3 | 95526786 | 95556403 | 1 | Mm.3619 |
| -0.42 | -3.86 | 0.26 | -4.02 | DN1ab | DN1cd | -3.77 | -11.64 | 7.40E-06 | 0.0041 | 0.22 | ENSMUSG00000102286 | TEC | Gm37004 | predicted gene, 37004 | NA | 2 | 78972672 | 79098851 | 1 | |
| 3.6 | 1.49 | 3.61 | 1.14 | DN1ab | DN1cd | -2.27 | -11.59 | 7.60E-06 | 0.0041 | 0.4 | ENSMUSG00000096712 | processed_pseudogene | Gm15454 | predicted gene 15454 | NA | 1 | 134518081 | 134518563 | 1 | |
| 7.82 | 6.9 | 8.56 | 7.4 | DN1ab | DN1cd | -1.01 | -11.54 | 7.80E-06 | 0.0041 | 0.2 | ENSMUSG00000031328 | protein_coding | Flna | filamin, alpha | 192176 | X | 74223461 | 74249820 | -1 | Mm.295533 |
| 3.76 | -1.28 | 3.89 | -0.44 | DN1ab | DN1cd | -4.63 | -11.43 | 8.30E-06 | 0.0043 | 0.43 | ENSMUSG00000048078 | protein_coding | Tenm4 | teneurin transmembrane protein 4 | 23966 | 7 | 96171246 | 96911093 | 1 | Mm.254610 Mm.391678 |
| 7.59 | 6.97 | 7.93 | 7.29 | DN1ab | DN1cd | -0.63 | -11.34 | 8.80E-06 | 0.0044 | 0.86 | ENSMUSG00000038366 | protein_coding | Lasp1 | LIM and SH3 protein 1 | 16796 | 11 | 97799000 | 97838764 | 1 | Mm.271967 Mm.470698 |
| 9.94 | 9.25 | 9.96 | 9.26 | DN1ab | DN1cd | -0.69 | -11.28 | 9.10E-06 | 0.0044 | 0.92 | ENSMUSG00000062611 | processed_pseudogene | Rps3a2 | ribosomal protein S3A2 | NA | 14 | 88123017 | 88123808 | -1 | |
| -3.14 | 2.3 | -2.36 | 2.65 | DN1ab | DN1cd | 5.32 | 11.21 | 9.50E-06 | 0.0044 | 0.71 | ENSMUSG00000099876 | lincRNA | Gm29650 | predicted gene 29650 | NA | Y | 1048393 | 1049134 | 1 | |
| 7.32 | 7.83 | 7.45 | 7.97 | DN1ab | DN1cd | 0.51 | 11.2 | 9.60E-06 | 0.0044 | 0.95 | ENSMUSG00000063696 | processed_pseudogene | Gm8730 | predicted pseudogene 8730 | NA | 8 | 102864845 | 102865798 | -1 | |
| 10.5 | 10 | 10.88 | 10.47 | DN1ab | DN1cd | -0.46 | -11.2 | 9.60E-06 | 0.0044 | 0.3 | ENSMUSG00000008348 | protein_coding | Ubc | ubiquitin C | 22190 | 5 | 125385965 | 125390202 | -1 | Mm.331 Mm.419997 |
| 5.17 | 6.16 | 4.73 | 6.06 | DN1ab | DN1cd | 1.11 | 11.19 | 9.60E-06 | 0.0044 | 0.087 | ENSMUSG00000034663 | protein_coding | Bmp2k | BMP2 inducible kinase | 140780 | 5 | 96997689 | 97091867 | 1 | Mm.281490 Mm.392853 |
| 6.06 | 6.9 | 5.87 | 7.03 | DN1ab | DN1cd | 0.96 | 11.11 | 1.00E-05 | 0.0044 | 0.063 | ENSMUSG00000022472 | protein_coding | Desi1 | desumoylating isopeptidase 1 | 28075 | 15 | 81992523 | 82033862 | -1 | Mm.474453 |
| 5.75 | 6.57 | 5.72 | 6.8 | DN1ab | DN1cd | 0.92 | 11.1 | 1.00E-05 | 0.0044 | 0.14 | ENSMUSG00000021892 | protein_coding | Sh3bp5 | SH3-domain binding protein 5 (BTK-associated) | 24056 | 14 | 31359880 | 31436078 | -1 | Mm.383198 Mm.438866 |
| 6.38 | 5.13 | 6.69 | 5.1 | DN1ab | DN1cd | -1.39 | -11.1 | 1.00E-05 | 0.0044 | 0.2 | ENSMUSG00000004633 | protein_coding | Chn2 | chimerin 2 | 69993 | 6 | 54039554 | 54301810 | 1 | Mm.321147 Mm.448157 |
| 6.14 | 6.84 | 5.97 | 6.64 | DN1ab | DN1cd | 0.69 | 11.05 | 1.00E-05 | 0.0044 | 0.88 | ENSMUSG00000032254 | protein_coding | Kif23 | kinesin family member 23 | 71819 | 9 | 61915905 | 61946774 | -1 | Mm.259374 |
| 6.33 | 5.8 | 7.35 | 6.83 | DN1ab | DN1cd | -0.52 | -11 | 1.10E-05 | 0.0044 | 0.88 | ENSMUSG00000056888 | protein_coding | Glipr1 | GLI pathogenesis-related 1 (glioma) | 73690 | 10 | 111985448 | 112002631 | -1 | Mm.173790 |
| 6.33 | 3.09 | 8.32 | 4.51 | DN1ab | DN1cd | -3.56 | -10.98 | 1.10E-05 | 0.0044 | 0.42 | ENSMUSG00000049103 | protein_coding | Ccr2 | chemokine (C-C motif) receptor 2 | 12772 | 9 | 124101950 | 124113557 | 1 | Mm.6272 |
| 6.61 | 5.03 | 6.84 | 5.77 | DN1ab | DN1cd | -1.36 | -10.93 | 1.10E-05 | 0.0044 | 0.03 | ENSMUSG00000086236 | antisense | 5830418P13Rik | RIKEN cDNA 5830418P13 gene | NA | 9 | 103423578 | 103461260 | 1 | |
| 0.69 | 3.15 | 0.14 | 3.25 | DN1ab | DN1cd | 2.67 | 10.91 | 1.10E-05 | 0.0044 | 0.25 | ENSMUSG00000025227 | protein_coding | Mfsd13a | major facilitator superfamily domain containing 13a | 75146 | 19 | 46341121 | 46375252 | 1 | Mm.159290 |
| 2.81 | -2.28 | 2.66 | -1.01 | DN1ab | DN1cd | -4.35 | -10.91 | 1.10E-05 | 0.0044 | 0.067 | ENSMUSG00000084403 | processed_pseudogene | Rps15a-ps8 | ribosomal protein S15A, pseudogene 8 | NA | 4 | 53791070 | 53791456 | -1 | |
| 2.44 | 0.12 | 3.04 | 1.06 | DN1ab | DN1cd | -2.14 | -10.84 | 1.20E-05 | 0.0045 | 0.41 | ENSMUSG00000020474 | protein_coding | Polm | polymerase (DNA directed), mu | 54125 | 11 | 5827860 | 5838016 | -1 | Mm.260194 |
| 5.53 | 2.62 | 6.58 | 4.04 | DN1ab | DN1cd | -2.7 | -10.76 | 1.30E-05 | 0.0045 | 0.5 | ENSMUSG00000038128 | protein_coding | Camk4 | calcium/calmodulin-dependent protein kinase IV | 12326 | 18 | 32934931 | 33191228 | 1 | Mm.222329 Mm.410854 |
| 6.34 | 5.71 | 7.24 | 6.47 | DN1ab | DN1cd | -0.68 | -10.74 | 1.30E-05 | 0.0045 | 0.29 | ENSMUSG00000030605 | protein_coding | Mfge8 | milk fat globule-EGF factor 8 protein | 17304 | 7 | 79133768 | 79149060 | -1 | Mm.1451 |
| 3.89 | 6.44 | 3.82 | 6.81 | DN1ab | DN1cd | 2.7 | 10.74 | 1.30E-05 | 0.0045 | 0.45 | ENSMUSG00000034361 | protein_coding | Cpne2 | copine II | 234577 | 8 | 94532990 | 94570531 | 1 | Mm.291815 |
| 5.68 | 4.93 | 6.57 | 5.69 | DN1ab | DN1cd | -0.81 | -10.74 | 1.30E-05 | 0.0045 | 0.42 | ENSMUSG00000055805 | protein_coding | Fmnl1 | formin-like 1 | 57778 | 11 | 103171107 | 103198901 | 1 | Mm.138913 |
| 7.29 | 5.99 | 9.06 | 7.44 | DN1ab | DN1cd | -1.44 | -10.71 | 1.30E-05 | 0.0045 | 0.25 | ENSMUSG00000001025 | protein_coding | S100a6 | S100 calcium binding protein A6 (calcyclin) | 20200 | 3 | 90612882 | 90624181 | 1 | Mm.100144 |
| 3.4 | 4.76 | 3.93 | 5.34 | DN1ab | DN1cd | 1.38 | 10.67 | 1.30E-05 | 0.0046 | 0.85 | ENSMUSG00000022995 | protein_coding | Enah | ENAH actin regulator | 13800 | 1 | 181896384 | 182019990 | -1 | Mm.87759 Mm.389224 |
| 6.27 | 3.34 | 7.82 | 5.02 | DN1ab | DN1cd | -2.86 | -10.62 | 1.40E-05 | 0.0046 | 0.84 | ENSMUSG00000026483 | protein_coding | Fam129a | family with sequence similarity 129, member A | 63913 | 1 | 151571186 | 151721939 | 1 | Mm.157700 Mm.482559 |
| 7.61 | 4.97 | 8.73 | 6.17 | DN1ab | DN1cd | -2.6 | -10.62 | 1.40E-05 | 0.0046 | 0.88 | ENSMUSG00000074607 | protein_coding | Tox2 | TOX high mobility group box family member 2 | 269389 | 2 | 163203125 | 163324170 | 1 | Mm.207709 |
| 7.7 | 8.38 | 7.72 | 8.25 | DN1ab | DN1cd | 0.63 | 10.59 | 1.40E-05 | 0.0046 | 0.27 | ENSMUSG00000025544 | protein_coding | Tm9sf2 | transmembrane 9 superfamily member 2 | 68059 | 14 | 122107038 | 122159604 | 1 | Mm.275191 |
| 8.98 | 9.84 | 8.43 | 9.62 | DN1ab | DN1cd | 0.98 | 10.56 | 1.40E-05 | 0.0046 | 0.087 | ENSMUSG00000021213 | protein_coding | Akr1c13 | aldo-keto reductase family 1, member C13 | 27384 | 13 | 4191150 | 4205596 | 1 | Mm.27447 |
| 5.49 | 2.65 | 6.16 | 2.86 | DN1ab | DN1cd | -3.07 | -10.55 | 1.40E-05 | 0.0046 | 0.47 | ENSMUSG00000066687 | protein_coding | Zbtb16 | zinc finger and BTB domain containing 16 | 235320 | 9 | 48654297 | 48836222 | -1 | Mm.457803 |
| 7.04 | 6.45 | 7.33 | 6.55 | DN1ab | DN1cd | -0.66 | -10.5 | 1.50E-05 | 0.0046 | 0.17 | ENSMUSG00000032014 | protein_coding | Oaf | out at first homolog | 102644 | 9 | 43221235 | 43239911 | -1 | Mm.246479 |
| 2.31 | 3.96 | 1.92 | 4.35 | DN1ab | DN1cd | 1.92 | 10.5 | 1.50E-05 | 0.0046 | 0.027 | ENSMUSG00000096847 | protein_coding | Tmem151b | transmembrane protein 151B | 210573 | 17 | 45541952 | 45549677 | -1 | Mm.329064 Mm.491193 |
| 7.22 | 6.12 | 8.27 | 7.21 | DN1ab | DN1cd | -1.08 | -10.44 | 1.50E-05 | 0.0047 | 0.89 | ENSMUSG00000001281 | protein_coding | Itgb7 | integrin beta 7 | 16421 | 15 | 102215995 | 102231944 | -1 | Mm.58 |
| -2.24 | 1.42 | -1.83 | 2.15 | DN1ab | DN1cd | 3.76 | 10.37 | 1.60E-05 | 0.0049 | 0.71 | ENSMUSG00000074657 | protein_coding | Kif5a | kinesin family member 5A | 16572 | 10 | 127225696 | 127263348 | -1 | Mm.30355 |
| 6.22 | 5.46 | 6.73 | 6 | DN1ab | DN1cd | -0.75 | -10.31 | 1.70E-05 | 0.005 | 0.89 | ENSMUSG00000003644 | protein_coding | Rps6ka1 | ribosomal protein S6 kinase polypeptide 1 | 20111 | 4 | 133847290 | 133887797 | -1 | Mm.301827 |
| 2.39 | -1.54 | 2.47 | -0.25 | DN1ab | DN1cd | -3.27 | -10.29 | 1.70E-05 | 0.005 | 0.044 | ENSMUSG00000091785 | processed_pseudogene | Rpl29-ps5 | ribosomal protein L29, pseudogene 5 | NA | 18 | 30459082 | 30459544 | -1 | |
| 8.87 | 9.37 | 8.99 | 9.5 | DN1ab | DN1cd | 0.5 | 10.27 | 1.70E-05 | 0.005 | 0.97 | ENSMUSG00000090862 | protein_coding | Rps13 | ribosomal protein S13 | 68052 | 7 | 116331505 | 116334195 | -1 | Mm.14798 Mm.345443 Mm.425541 |
| -3.82 | -0.33 | -2.81 | 0.06 | DN1ab | DN1cd | 3.27 | 10.07 | 1.90E-05 | 0.0056 | 0.41 | ENSMUSG00000046410 | protein_coding | Kcnk6 | potassium inwardly-rectifying channel, subfamily K, member 6 | 52150 | 7 | 29221926 | 29232515 | -1 | Mm.24877 |
| 6.75 | 6.11 | 7.47 | 6.66 | DN1ab | DN1cd | -0.7 | -10.05 | 2.00E-05 | 0.0056 | 0.25 | ENSMUSG00000037012 | protein_coding | Hk1 | hexokinase 1 | 15275 | 10 | 62268855 | 62379908 | -1 | Mm.196605 |
| 3.31 | 1.25 | 3.84 | 2.28 | DN1ab | DN1cd | -1.8 | -10.03 | 2.00E-05 | 0.0056 | 0.17 | ENSMUSG00000079227 | protein_coding | Ccr5 | chemokine (C-C motif) receptor 5 | 12774 | 9 | 124121543 | 124147699 | 1 | Mm.14302 |
| 2.59 | 1.41 | 2.62 | 1.11 | DN1ab | DN1cd | -1.32 | -10.03 | 2.00E-05 | 0.0056 | 0.22 | ENSMUSG00000094446 | processed_pseudogene | Gm6344 | predicted gene 6344 | NA | 13 | 55085777 | 55086259 | 1 | |
| 4.2 | 2.76 | 4.94 | 3.51 | DN1ab | DN1cd | -1.44 | -9.98 | 2.10E-05 | 0.0056 | 0.99 | ENSMUSG00000029924 | protein_coding | Slc37a3 | solute carrier family 37 (glycerol-3-phosphate transporter), member 3 | 72144 | 6 | 39334773 | 39377675 | -1 | Mm.482167 |
| 1.13 | 3.53 | 0.7 | 3.71 | DN1ab | DN1cd | 2.59 | 9.97 | 2.10E-05 | 0.0056 | 0.3 | ENSMUSG00000079481 | protein_coding | Nhsl2 | NHS-like 2 | 100042480 | X | 101849385 | 102092055 | 1 | Mm.232 |
| 3.41 | 4.92 | 2.51 | 4.11 | DN1ab | DN1cd | 1.54 | 9.93 | 2.10E-05 | 0.0056 | 0.81 | ENSMUSG00000033752 | protein_coding | Mnd1 | meiotic nuclear divisions 1 | 76915 | 3 | 84087933 | 84155786 | -1 | Mm.274829 |
| 7.97 | 7.47 | 8.48 | 7.99 | DN1ab | DN1cd | -0.49 | -9.93 | 2.10E-05 | 0.0056 | 0.93 | ENSMUSG00000030536 | protein_coding | Iqgap1 | IQ motif containing GTPase activating protein 1 | 29875 | 7 | 80711583 | 80825974 | -1 | Mm.207619 |
| 7.06 | 6.06 | 7.25 | 6.35 | DN1ab | DN1cd | -0.96 | -9.88 | 2.20E-05 | 0.0057 | 0.66 | ENSMUSG00000031309 | protein_coding | Rps6ka3 | ribosomal protein S6 kinase polypeptide 3 | 110651 | X | 159210307 | 159368244 | 1 | Mm.328476 Mm.392829 |
| 5.79 | 4.2 | 5.99 | 4.8 | DN1ab | DN1cd | -1.42 | -9.8 | 2.30E-05 | 0.006 | 0.18 | ENSMUSG00000000244 | protein_coding | Tspan32 | tetraspanin 32 | 27027 | 7 | 143005046 | 143019644 | 1 | Mm.28172 |
| 11.18 | 10.67 | 11.22 | 10.5 | DN1ab | DN1cd | -0.59 | -9.73 | 2.40E-05 | 0.0062 | 0.092 | ENSMUSG00000094344 | processed_pseudogene | Gm11942 | predicted gene 11942 | NA | 4 | 32964888 | 32965232 | 1 | |
| 5.86 | 5.16 | 6.8 | 5.77 | DN1ab | DN1cd | -0.84 | -9.7 | 2.50E-05 | 0.0063 | 0.05 | ENSMUSG00000040111 | protein_coding | Gramd1b | GRAM domain containing 1B | 235283 | 9 | 40293233 | 40531383 | -1 | Mm.138434 |
| 6.24 | 5.27 | 6.52 | 5.27 | DN1ab | DN1cd | -1.08 | -9.68 | 2.50E-05 | 0.0063 | 0.24 | ENSMUSG00000030137 | protein_coding | Tuba8 | tubulin, alpha 8 | 53857 | 6 | 121210696 | 121226854 | 1 | Mm.32884 |
| 7.14 | 6.36 | 8.09 | 7.16 | DN1ab | DN1cd | -0.84 | -9.64 | 2.60E-05 | 0.0064 | 0.44 | ENSMUSG00000026223 | protein_coding | Itm2c | integral membrane protein 2C | 64294 | 1 | 85894281 | 85908675 | 1 | Mm.29870 |
| 3.22 | 1.38 | 3.41 | 1.39 | DN1ab | DN1cd | -1.92 | -9.63 | 2.60E-05 | 0.0064 | 0.7 | ENSMUSG00000094497 | processed_pseudogene | Gm8210 | predicted pseudogene 8210 | NA | 1 | 43189156 | 43189638 | 1 | |
| 9.62 | 7.32 | 11.04 | 7.79 | DN1ab | DN1cd | -2.7 | -9.61 | 2.60E-05 | 0.0064 | 0.089 | ENSMUSG00000028266 | protein_coding | Lmo4 | LIM domain only 4 | 16911 | 3 | 144188530 | 144205220 | -1 | Mm.29187 |
| 6.99 | 5.61 | 7.44 | 6.32 | DN1ab | DN1cd | -1.27 | -9.58 | 2.70E-05 | 0.0064 | 0.37 | ENSMUSG00000040212 | protein_coding | Emp3 | epithelial membrane protein 3 | 13732 | 7 | 45918023 | 45921404 | -1 | Mm.20829 |
| 4.98 | 3.85 | 5.96 | 4.9 | DN1ab | DN1cd | -1.1 | -9.56 | 2.70E-05 | 0.0065 | 0.77 | ENSMUSG00000084399 | transcribed_processed_pseudogene | Gm15666 | predicted gene 15666 | NA | 10 | 105505804 | 105508599 | -1 | |
| 4.71 | 2.9 | 6.1 | 4.27 | DN1ab | DN1cd | -1.82 | -9.55 | 2.80E-05 | 0.0065 | 0.95 | ENSMUSG00000026285 | protein_coding | Pdcd1 | programmed cell death 1 | 18566 | 1 | 94038305 | 94052553 | -1 | Mm.5024 |
| 3.12 | 4.41 | 2.86 | 4 | DN1ab | DN1cd | 1.24 | 9.46 | 2.90E-05 | 0.0067 | 0.63 | ENSMUSG00000053716 | protein_coding | Dusp7 | dual specificity phosphatase 7 | 235584 | 9 | 106368632 | 106375724 | 1 | Mm.275584 |
| -2.24 | 0.53 | -3.49 | -0.03 | DN1ab | DN1cd | 2.99 | 9.45 | 3.00E-05 | 0.0067 | 0.33 | ENSMUSG00000049551 | protein_coding | Fzd9 | frizzled class receptor 9 | 14371 | 5 | 135248938 | 135251230 | -1 | Mm.6256 |
| 5.2 | 4.5 | 5.18 | 4.55 | DN1ab | DN1cd | -0.67 | -9.43 | 3.00E-05 | 0.0067 | 0.67 | ENSMUSG00000030423 | protein_coding | Pop4 | processing of precursor 4, ribonuclease P/MRP family, (S. cerevisiae) | 66161 | 7 | 38261996 | 38271423 | -1 | Mm.22284 |
| 7.96 | 6.71 | 8.94 | 7.33 | DN1ab | DN1cd | -1.39 | -9.42 | 3.00E-05 | 0.0067 | 0.25 | ENSMUSG00000041272 | protein_coding | Tox | thymocyte selection-associated high mobility group box | 252838 | 4 | 6686353 | 6991557 | -1 | Mm.435828 Mm.488241 |
| 8.85 | 9.26 | 8.34 | 8.79 | DN1ab | DN1cd | 0.42 | 9.42 | 3.00E-05 | 0.0067 | 0.65 | ENSMUSG00000078249 | protein_coding | Hmga1b | high mobility group AT-hook 1B | 111241 | 11 | 120762794 | 120764419 | 1 | Mm.426212 |
| 6.77 | 7.7 | 6.75 | 8.13 | DN1ab | DN1cd | 1.09 | 9.4 | 3.10E-05 | 0.0067 | 0.063 | ENSMUSG00000032228 | protein_coding | Tcf12 | transcription factor 12 | 21406 | 9 | 71842688 | 72111871 | -1 | Mm.171615 Mm.480117 |
| 5.24 | 5.79 | 5.05 | 5.74 | DN1ab | DN1cd | 0.6 | 9.32 | 3.20E-05 | 0.007 | 0.34 | ENSMUSG00000081434 | processed_pseudogene | Gm14165 | predicted gene 14165 | NA | 2 | 152353615 | 152354566 | 1 | |
| 7.97 | 8.5 | 8.09 | 8.56 | DN1ab | DN1cd | 0.5 | 9.25 | 3.40E-05 | 0.0072 | 0.63 | ENSMUSG00000069972 | processed_pseudogene | Rps13-ps2 | ribosomal protein S13, pseudogene 2 | NA | 7 | 88530446 | 88530898 | -1 | |
| 1.72 | -1.7 | 3.42 | -0.18 | DN1ab | DN1cd | -3.51 | -9.25 | 3.40E-05 | 0.0072 | 0.84 | ENSMUSG00000106570 | TEC | C230071H17Rik | RIKEN cDNA C230071H17 gene | NA | 5 | 131882966 | 131885938 | -1 | |
| 1.7 | 3.57 | 0.34 | 2.85 | DN1ab | DN1cd | 2.07 | 9.24 | 3.40E-05 | 0.0072 | 0.21 | ENSMUSG00000063455 | protein_coding | D630045J12Rik | RIKEN cDNA D630045J12 gene | 330286 | 6 | 38123174 | 38254009 | -1 | Mm.277163 |
| 7.09 | 6.32 | 7.2 | 6.61 | DN1ab | DN1cd | -0.7 | -9.21 | 3.50E-05 | 0.0072 | 0.31 | ENSMUSG00000029455 | protein_coding | Aldh2 | aldehyde dehydrogenase 2, mitochondrial | 11669 | 5 | 121566027 | 121593824 | -1 | Mm.284446 |
| 2.2 | 3.83 | 1.95 | 4.52 | DN1ab | DN1cd | 1.95 | 9.21 | 3.50E-05 | 0.0072 | 0.02 | ENSMUSG00000079037 | protein_coding | Prnp | prion protein | 19122 | 2 | 131909928 | 131938429 | 1 | Mm.648 |
| 6.76 | 6.13 | 7.22 | 6.53 | DN1ab | DN1cd | -0.65 | -9.19 | 3.50E-05 | 0.0072 | 0.7 | ENSMUSG00000039156 | protein_coding | Stim2 | stromal interaction molecule 2 | 116873 | 5 | 53998499 | 54121057 | 1 | Mm.265985 |
| 2.48 | 3.96 | 2.34 | 4.31 | DN1ab | DN1cd | 1.64 | 9.16 | 3.60E-05 | 0.0073 | 0.22 | ENSMUSG00000023951 | protein_coding | Vegfa | vascular endothelial growth factor A | 22339 | 17 | 46016993 | 46032369 | -1 | Mm.282184 |
| 2.97 | 4.14 | 3.11 | 4.52 | DN1ab | DN1cd | 1.25 | 9.14 | 3.70E-05 | 0.0074 | 0.44 | ENSMUSG00000046841 | protein_coding | Ckap4 | cytoskeleton-associated protein 4 | 216197 | 10 | 84526305 | 84534062 | -1 | Mm.334999 |
| 4.83 | 6.2 | 4.36 | 6.36 | DN1ab | DN1cd | 1.58 | 9.12 | 3.70E-05 | 0.0074 | 0.075 | ENSMUSG00000052751 | protein_coding | Repin1 | replication initiator 1 | 58887 | 6 | 48593883 | 48599082 | 1 | Mm.219183 Mm.410678 |
| 4.2 | 5.05 | 4.67 | 5.65 | DN1ab | DN1cd | 0.9 | 9.1 | 3.80E-05 | 0.0074 | 0.58 | ENSMUSG00000030103 | protein_coding | Bhlhe40 | basic helix-loop-helix family, member e40 | 20893 | 6 | 108660629 | 108666925 | 1 | Mm.2436 |
| 2.37 | -0.68 | 5.17 | 1.47 | DN1ab | DN1cd | -3.47 | -9.09 | 3.80E-05 | 0.0074 | 0.45 | ENSMUSG00000022747 | protein_coding | St3gal6 | ST3 beta-galactoside alpha-2,3-sialyltransferase 6 | 54613 | 16 | 58468125 | 58524243 | -1 | Mm.212742 |
| 8.26 | 7.48 | 9.1 | 8.25 | DN1ab | DN1cd | -0.81 | -9.08 | 3.90E-05 | 0.0074 | 0.7 | ENSMUSG00000004207 | protein_coding | Psap | prosaposin | 19156 | 10 | 60277627 | 60302597 | 1 | Mm.29978 Mm.277498 |
| -1.56 | 1.84 | -2.01 | 1.3 | DN1ab | DN1cd | 3.38 | 9.07 | 3.90E-05 | 0.0074 | 0.92 | ENSMUSG00000035539 | protein_coding | Ccdc180 | coiled-coil domain containing 180 | 381522 | 4 | 45890303 | 45950774 | 1 | Mm.54220 |
| 5.85 | 7.34 | 5.4 | 7.11 | DN1ab | DN1cd | 1.57 | 9.06 | 3.90E-05 | 0.0074 | 0.57 | ENSMUSG00000035561 | protein_coding | Aldh1b1 | aldehyde dehydrogenase 1 family, member B1 | 72535 | 4 | 45799022 | 45804604 | 1 | Mm.331583 |
| 3.89 | 4.64 | 3.56 | 4.29 | DN1ab | DN1cd | 0.74 | 9.04 | 4.00E-05 | 0.0075 | 0.88 | ENSMUSG00000020818 | protein_coding | Mfsd11 | major facilitator superfamily domain containing 11 | 69900 | 11 | 116852440 | 116875635 | 1 | Mm.348004 Mm.484423 |
| 8.97 | 9.34 | 8.36 | 8.87 | DN1ab | DN1cd | 0.42 | 9.03 | 4.00E-05 | 0.0075 | 0.18 | ENSMUSG00000046711 | protein_coding | Hmga1 | high mobility group AT-hook 1 | 15361 | 17 | 27556497 | 27563674 | 1 | Mm.4438 Mm.426212 Mm.440289 |
| 0.28 | -3.12 | 2.31 | -1.83 | DN1ab | DN1cd | -3.72 | -9.01 | 4.10E-05 | 0.0075 | 0.42 | ENSMUSG00000105726 | TEC | Gm42443 | predicted gene 42443 | NA | 5 | 131694721 | 131695875 | -1 | |
| -1.54 | 2.26 | 0.1 | 3.04 | DN1ab | DN1cd | 3.53 | 9 | 4.10E-05 | 0.0075 | 0.34 | ENSMUSG00000097604 | lincRNA | Gm17322 | predicted gene, 17322 | NA | 9 | 57998041 | 58007451 | 1 | |
| 3.58 | 6.46 | 1.85 | 6.38 | DN1ab | DN1cd | 3.4 | 8.91 | 4.40E-05 | 0.008 | 0.028 | ENSMUSG00000067750 | protein_coding | Khdc1a | KH domain containing 1A | 368204 | 1 | 21349641 | 21352200 | 1 | Mm.26006 |
| 6.38 | 5.81 | 6.59 | 5.91 | DN1ab | DN1cd | -0.61 | -8.89 | 4.40E-05 | 0.008 | 0.43 | ENSMUSG00000021076 | protein_coding | Actr10 | ARP10 actin-related protein 10 | 56444 | 12 | 70937857 | 70964718 | 1 | Mm.29317 |
| 3.46 | 5.56 | 2 | 5.19 | DN1ab | DN1cd | 2.45 | 8.88 | 4.50E-05 | 0.008 | 0.054 | ENSMUSG00000062380 | protein_coding | Tubb3 | tubulin, beta 3 class III | 22152 | 8 | 123411424 | 123422015 | 1 | Mm.40068 |
| 0.19 | -3.12 | 3.3 | 0.36 | DN1ab | DN1cd | -3.12 | -8.86 | 4.50E-05 | 0.0081 | 0.62 | ENSMUSG00000073010 | protein_coding | Gm5127 | predicted gene 5127 | 331493 | X | 106583186 | 106710557 | 1 | Mm.249319 |
| 5.11 | 4.13 | 5.6 | 4.74 | DN1ab | DN1cd | -0.93 | -8.82 | 4.70E-05 | 0.0082 | 0.57 | ENSMUSG00000029050 | protein_coding | Ski | ski sarcoma viral oncogene homolog (avian) | 20481 | 4 | 155154075 | 155222592 | -1 | Mm.28520 |
| 2.91 | 1.8 | 3.6 | 1.95 | DN1ab | DN1cd | -1.34 | -8.81 | 4.70E-05 | 0.0082 | 0.077 | ENSMUSG00000043251 | protein_coding | Exoc3l | exocyst complex component 3-like | 277978 | 8 | 105289924 | 105296101 | -1 | Mm.40228 |
| 1.8 | 3.42 | 1.85 | 3.46 | DN1ab | DN1cd | 1.62 | 8.79 | 4.80E-05 | 0.0082 | 0.98 | ENSMUSG00000031788 | protein_coding | Kifc3 | kinesin family member C3 | 16582 | 8 | 95099828 | 95202812 | -1 | Mm.378951 |
| 6.89 | 6.15 | 7.66 | 6.97 | DN1ab | DN1cd | -0.72 | -8.79 | 4.80E-05 | 0.0082 | 0.78 | ENSMUSG00000024063 | protein_coding | Lbh | limb-bud and heart | 77889 | 17 | 72918305 | 72941947 | 1 | Mm.154109 |
| 4.91 | 3.06 | 6.5 | 4.85 | DN1ab | DN1cd | -1.74 | -8.79 | 4.80E-05 | 0.0082 | 0.63 | ENSMUSG00000024696 | protein_coding | Lpxn | leupaxin | 107321 | 19 | 12798606 | 12833807 | 1 | Mm.313136 |
| 1.95 | 4.62 | 2.24 | 3.79 | DN1ab | DN1cd | 2.32 | 8.78 | 4.80E-05 | 0.0082 | 0.033 | ENSMUSG00000031762 | protein_coding | Mt2 | metallothionein 2 | 17750 | 8 | 94172664 | 94173568 | 1 | Mm.147226 |
| -3.82 | 1.17 | -2.81 | 0.28 | DN1ab | DN1cd | 4.42 | 8.77 | 4.80E-05 | 0.0082 | 0.076 | ENSMUSG00000025754 | protein_coding | Agbl1 | ATP/GTP binding protein-like 1 | 244071 | 7 | 76229887 | 77124698 | 1 | Mm.322829 |
| 8.6 | 8.05 | 8.73 | 8.02 | DN1ab | DN1cd | -0.61 | -8.76 | 4.90E-05 | 0.0082 | 0.32 | ENSMUSG00000031827 | protein_coding | Cotl1 | coactosin-like 1 (Dictyostelium) | 72042 | 8 | 119809222 | 119840544 | -1 | Mm.393405 |
| 7.91 | 6.12 | 8.72 | 7.31 | DN1ab | DN1cd | -1.63 | -8.74 | 4.90E-05 | 0.0082 | 0.36 | ENSMUSG00000000078 | protein_coding | Klf6 | Kruppel-like factor 6 | 23849 | 13 | 5861482 | 5870394 | 1 | Mm.275036 |
| 4.86 | 5.52 | 4.99 | 5.51 | DN1ab | DN1cd | 0.61 | 8.74 | 5.00E-05 | 0.0082 | 0.4 | ENSMUSG00000033713 | protein_coding | Foxn3 | forkhead box N3 | 71375 | 12 | 99190078 | 99563582 | -1 | Mm.341972 Mm.448530 Mm.489662 |
| 10.69 | 11.66 | 9.83 | 10.86 | DN1ab | DN1cd | 0.99 | 8.73 | 5.00E-05 | 0.0082 | 0.84 | ENSMUSG00000030795 | protein_coding | Fus | fused in sarcoma | 233908 | 7 | 127967457 | 127985701 | 1 | Mm.277680 |
| 7.18 | 6.23 | 8.35 | 7.48 | DN1ab | DN1cd | -0.92 | -8.72 | 5.00E-05 | 0.0082 | 0.75 | ENSMUSG00000041959 | protein_coding | S100a10 | S100 calcium binding protein A10 (calpactin) | 20194 | 3 | 93555080 | 93564643 | 1 | Mm.1 |
| -1.01 | -3.86 | -0.59 | -4.02 | DN1ab | DN1cd | -3.07 | -8.71 | 5.10E-05 | 0.0082 | 0.46 | ENSMUSG00000070424 | protein_coding | Art5 | ADP-ribosyltransferase 5 | 11875 | 7 | 102096879 | 102109489 | -1 | Mm.2816 |
| 4.48 | 5.37 | 4.36 | 5.44 | DN1ab | DN1cd | 0.95 | 8.7 | 5.10E-05 | 0.0082 | 0.42 | ENSMUSG00000000276 | protein_coding | Dgke | diacylglycerol kinase, epsilon | 56077 | 11 | 89035179 | 89066850 | -1 | Mm.153695 |
| 5.45 | 4.79 | 5.53 | 4.81 | DN1ab | DN1cd | -0.68 | -8.68 | 5.20E-05 | 0.0083 | 0.74 | ENSMUSG00000054702 | protein_coding | Ap1s3 | adaptor-related protein complex AP-1, sigma 3 | 252903 | 1 | 79606876 | 79671972 | -1 | Mm.32409 |
| 6.01 | 4.83 | 7.34 | 6.36 | DN1ab | DN1cd | -1.09 | -8.62 | 5.40E-05 | 0.0084 | 0.49 | ENSMUSG00000031129 | protein_coding | Slc9a9 | solute carrier family 9 (sodium/hydrogen exchanger), member 9 | 331004 | 9 | 94669909 | 95230445 | 1 | Mm.326125 |
| 1.84 | 3.12 | 1.77 | 3.48 | DN1ab | DN1cd | 1.42 | 8.61 | 5.50E-05 | 0.0084 | 0.25 | ENSMUSG00000087380 | processed_transcript | 2210408F21Rik | RIKEN cDNA 2210408F21 gene | NA | 6 | 31220351 | 31337404 | 1 | |
| 6.69 | 8.08 | 6.74 | 8.04 | DN1ab | DN1cd | 1.36 | 8.6 | 5.50E-05 | 0.0084 | 0.82 | ENSMUSG00000027368 | protein_coding | Dusp2 | dual specificity phosphatase 2 | 13537 | 2 | 127336159 | 127338376 | 1 | Mm.4729 |
| 6.98 | 8.15 | 6.97 | 8.57 | DN1ab | DN1cd | 1.32 | 8.59 | 5.50E-05 | 0.0084 | 0.2 | ENSMUSG00000002204 | protein_coding | Napsa | napsin A aspartic peptidase | 16541 | 7 | 44572380 | 44586862 | 1 | Mm.383181 |
| 7.42 | 7.87 | 7.56 | 8.07 | DN1ab | DN1cd | 0.47 | 8.59 | 5.50E-05 | 0.0084 | 0.65 | ENSMUSG00000019188 | protein_coding | H13 | histocompatibility 13 | 14950 | 2 | 152669461 | 152708670 | 1 | Mm.277327 |
| 5.9 | 5.04 | 6.35 | 5.36 | DN1ab | DN1cd | -0.91 | -8.59 | 5.50E-05 | 0.0084 | 0.6 | ENSMUSG00000032402 | protein_coding | Smad3 | SMAD family member 3 | 17127 | 9 | 63646767 | 63757994 | -1 | Mm.7320 |
| 6.52 | 7 | 6.1 | 6.57 | DN1ab | DN1cd | 0.48 | 8.56 | 5.70E-05 | 0.0086 | 0.98 | ENSMUSG00000032640 | protein_coding | Chsy1 | chondroitin sulfate synthase 1 | 269941 | 7 | 66109515 | 66173798 | 1 | Mm.333916 |
| 6.07 | 5.63 | 6.57 | 6.06 | DN1ab | DN1cd | -0.47 | -8.54 | 5.70E-05 | 0.0086 | 0.61 | ENSMUSG00000020198 | protein_coding | Ap3d1 | adaptor-related protein complex 3, delta 1 subunit | 11776 | 10 | 80706956 | 80742264 | -1 | Mm.27082 Mm.209294 |
| 5.32 | 5.84 | 5.15 | 5.92 | DN1ab | DN1cd | 0.61 | 8.54 | 5.80E-05 | 0.0086 | 0.097 | ENSMUSG00000050075 | protein_coding | Gpr171 | G protein-coupled receptor 171 | 229323 | 3 | 59096448 | 59101821 | -1 | Mm.123648 |
| 0.48 | -3.86 | 1.45 | -1.82 | DN1ab | DN1cd | -3.89 | -8.52 | 5.90E-05 | 0.0087 | 0.27 | ENSMUSG00000042793 | protein_coding | Lgr6 | leucine-rich repeat-containing G protein-coupled receptor 6 | 329252 | 1 | 134983301 | 135105276 | -1 | Mm.279594 |
| 4.04 | 3.04 | 4.49 | 3.4 | DN1ab | DN1cd | -1.04 | -8.49 | 6.00E-05 | 0.0088 | 0.76 | ENSMUSG00000028988 | protein_coding | Ctnnbip1 | catenin beta interacting protein 1 | 67087 | 4 | 149518236 | 149566437 | 1 | Mm.299735 Mm.474667 |
| 3.56 | 2.29 | 4.84 | 3.75 | DN1ab | DN1cd | -1.18 | -8.47 | 6.10E-05 | 0.0089 | 0.54 | ENSMUSG00000002668 | protein_coding | Dennd1c | DENN/MADD domain containing 1C | 70785 | 17 | 57065905 | 57078514 | -1 | Mm.284447 |
| -0.28 | -3.86 | 1.56 | -4.02 | DN1ab | DN1cd | -4.4 | -8.44 | 6.20E-05 | 0.009 | 0.045 | ENSMUSG00000025952 | protein_coding | Crygc | crystallin, gamma C | 12966 | 1 | 65071525 | 65073689 | -1 | Mm.491115 |
| 2.99 | -1.39 | 3.12 | -0.1 | DN1ab | DN1cd | -3.7 | -8.44 | 6.20E-05 | 0.009 | 0.21 | ENSMUSG00000052560 | protein_coding | Cpne8 | copine VIII | 66871 | 15 | 90487482 | 90679432 | -1 | Mm.290991 |
| 3.78 | 5.47 | 2.36 | 4.85 | DN1ab | DN1cd | 1.95 | 8.43 | 6.20E-05 | 0.009 | 0.1 | ENSMUSG00000028332 | protein_coding | Hemgn | hemogen | 93966 | 4 | 46393989 | 46413506 | -1 | Mm.25793 |
| -3.82 | 0.45 | -2.73 | 1.43 | DN1ab | DN1cd | 4.24 | 8.42 | 6.30E-05 | 0.009 | 0.93 | ENSMUSG00000047907 | protein_coding | Tshz2 | teashirt zinc finger family member 2 | 228911 | 2 | 169633013 | 170071816 | 1 | Mm.315789 Mm.468546 Mm.482636 |
| 3.48 | 0.24 | 4.49 | 0.7 | DN1ab | DN1cd | -3.54 | -8.42 | 6.30E-05 | 0.009 | 0.55 | ENSMUSG00000056758 | protein_coding | Hmga2 | high mobility group AT-hook 2 | 15364 | 10 | 120361275 | 120476469 | -1 | Mm.157190 |
| 7.16 | 3.01 | 9.19 | 5.63 | DN1ab | DN1cd | -3.79 | -8.39 | 6.40E-05 | 0.0091 | 0.57 | ENSMUSG00000032238 | protein_coding | Rora | RAR-related orphan receptor alpha | 19883 | 9 | 68653786 | 69388246 | 1 | Mm.378450 Mm.391890 Mm.427266 Mm.445802 Mm.491288 |
| 7.68 | 6.4 | 7.37 | 6.56 | DN1ab | DN1cd | -1.1 | -8.37 | 6.50E-05 | 0.0091 | 0.066 | ENSMUSG00000023828 | protein_coding | Slc22a3 | solute carrier family 22 (organic cation transporter), member 3 | 20519 | 17 | 12418960 | 12507704 | -1 | Mm.99252 |
| -3.82 | -1.28 | -3.94 | -1.7 | DN1ab | DN1cd | 2.43 | 8.37 | 6.60E-05 | 0.0091 | 0.66 | ENSMUSG00000040935 | protein_coding | Padi6 | peptidyl arginine deiminase, type VI | 242726 | 4 | 140727355 | 140742643 | -1 | Mm.271661 |
| 2.69 | 1.33 | 3.27 | 2.33 | DN1ab | DN1cd | -1.16 | -8.37 | 6.60E-05 | 0.0091 | 0.13 | ENSMUSG00000022324 | protein_coding | Matn2 | matrilin 2 | 17181 | 15 | 34306677 | 34436273 | 1 | Mm.396856 |
| -1.92 | 0.65 | -0.1 | 2.18 | DN1ab | DN1cd | 2.47 | 8.29 | 7.00E-05 | 0.0096 | 0.67 | ENSMUSG00000089718 | lincRNA | 2310075C17Rik | RIKEN cDNA 2310075C17 gene | NA | 9 | 112227579 | 112228242 | 1 | |
| 5.07 | 3.94 | 5.25 | 4.58 | DN1ab | DN1cd | -0.93 | -8.27 | 7.10E-05 | 0.0097 | 0.03 | ENSMUSG00000031534 | protein_coding | Smim19 | small integral membrane protein 19 | 102032 | 8 | 22462614 | 22476882 | -1 | Mm.378231 |
| 4.65 | 3.24 | 5.37 | 3.14 | DN1ab | DN1cd | -1.76 | -8.26 | 7.20E-05 | 0.0097 | 0.044 | ENSMUSG00000038807 | protein_coding | Rap1gap2 | RAP1 GTPase activating protein 2 | 380711 | 11 | 74383356 | 74610915 | -1 | Mm.425690 |
| 4.7 | 5.44 | 4.94 | 6.03 | DN1ab | DN1cd | 0.87 | 8.25 | 7.20E-05 | 0.0097 | 0.11 | ENSMUSG00000050921 | protein_coding | P2ry10 | purinergic receptor P2Y, G-protein coupled 10 | 78826 | X | 107088492 | 107104974 | 1 | Mm.74639 |
| -3.82 | -1.28 | -3.94 | -1.46 | DN1ab | DN1cd | 2.52 | 8.24 | 7.20E-05 | 0.0097 | 0.94 | ENSMUSG00000032735 | protein_coding | Ablim3 | actin binding LIM protein family, member 3 | 319713 | 18 | 61799395 | 61911852 | -1 | Mm.329478 |

| | | | | | | | | | | | | | | | | | | | | |
|---|---|---|---|---|---|---|---|---|---|---|---|---|---|---|---|---|---|---|---|---|
| 5.64 | 4.73 | 6.7 | 6.04 | DN1ab | DN1cd | -0.8 | -8.21 | 7.40E-05 | 0.0099 | 0.23 | ENSMUSG00000037706 | protein_coding | Cd81 | CD81 antigen | 12520 | 7 | 143052739 | 143067934 | 1 | Mm.806 |
| 2 | 4.64 | 1.31 | 3.59 | DN1ab | DN1cd | 2.53 | 8.18 | 7.60E-05 | 0.01 | 0.62 | ENSMUSG00000100967 | protein_coding | Gm29666 | predicted gene 29666 | NA | 15 | 84910438 | 84920038 | -1 | |
| 3.32 | 4.34 | 3.35 | 4.29 | DN1ab | DN1cd | 0.99 | 8.17 | 7.70E-05 | 0.01 | 0.78 | ENSMUSG00000074682 | protein_coding | Zcchc3 | zinc finger, CCHC domain containing 3 | 67917 | 2 | 152411955 | 152415027 | -1 | Mm.391810 |
| 5.66 | 4.33 | 6.22 | 4.95 | DN1ab | DN1cd | -1.31 | -8.17 | 7.70E-05 | 0.01 | 0.87 | ENSMUSG00000040322 | protein_coding | Slc25a24 | solute carrier family 25 (mitochondrial carrier, phosphate carrier), member 24 | 229731 | 3 | 109123149 | 109168457 | 1 | Mm.33574 |
| 1.37 | 2.52 | 1.4 | 2.85 | DN1ab | DN1cd | 1.25 | 8.17 | 7.70E-05 | 0.01 | 0.39 | ENSMUSG00000043157 | protein_coding | Arl11 | ADP-ribosylation factor-like 11 | 219144 | 14 | 61309753 | 61311936 | 1 | Mm.206566 |
| 6.27 | 6.75 | 5.95 | 6.71 | DN1ab | DN1cd | 0.58 | 8.17 | 7.70E-05 | 0.01 | 0.038 | ENSMUSG00000033767 | protein_coding | Tmem131l | transmembrane 131 like | 229473 | 3 | 83897655 | 84040175 | -1 | Mm.28838 |
| 3.86 | 2.63 | 4.99 | 3.83 | DN1ab | DN1cd | -1.2 | -8.07 | 8.30E-05 | 0.011 | 0.81 | ENSMUSG00000029923 | protein_coding | Rab19 | RAB19, member RAS oncogene family | 19331 | 6 | 39381175 | 39390380 | 1 | Mm.123866 |
| 1.92 | 0.35 | 3.53 | 2.17 | DN1ab | DN1cd | -1.46 | -8.04 | 8.50E-05 | 0.011 | 0.6 | ENSMUSG00000000562 | protein_coding | Adora3 | adenosine A3 receptor | 11542 | 3 | 105870858 | 105908926 | 1 | Mm.235024 |
| 0.88 | -1.54 | 2.28 | 0.23 | DN1ab | DN1cd | -2.23 | -8.02 | 8.60E-05 | 0.011 | 0.55 | ENSMUSG00000002324 | protein_coding | Rec8 | REC8 meiotic recombination protein | 56739 | 14 | 55618037 | 55625395 | 1 | Mm.23149 |
| 4.5 | 3.88 | 4.91 | 4.18 | DN1ab | DN1cd | -0.66 | -8.02 | 8.60E-05 | 0.011 | 0.56 | ENSMUSG00000043061 | protein_coding | Tmem18 | transmembrane protein 18 | 211986 | 12 | 30584426 | 30591215 | 1 | Mm.34035 |
| 7.84 | 8.51 | 8.4 | 8.87 | DN1ab | DN1cd | 0.59 | 7.98 | 8.90E-05 | 0.011 | 0.23 | ENSMUSG00000076752 | TR_C_gene | Tcrg-C2 | T-cell receptor gamma, constant 2 | NA | 13 | 19304649 | 19307551 | -1 | |
| 1.31 | -0.74 | 2.16 | -0.44 | DN1ab | DN1cd | -2.31 | -7.97 | 9.00E-05 | 0.011 | 0.37 | ENSMUSG00000012123 | protein_coding | Crybg2 | crystallin beta-gamma domain containing 2 | 230806 | 4 | 134060815 | 134092504 | 1 | Mm.240080 |
| -1.81 | 1.53 | -1.01 | 2.67 | DN1ab | DN1cd | 3.45 | 7.96 | 9.10E-05 | 0.011 | 0.74 | ENSMUSG00000022652 | protein_coding | Morc1 | microrchidia 1 | 17450 | 16 | 48431237 | 48630900 | 1 | Mm.250060 |
| 5.79 | 6.33 | 5.34 | 6.01 | DN1ab | DN1cd | 0.59 | 7.96 | 9.10E-05 | 0.011 | 0.45 | ENSMUSG00000043421 | protein_coding | Hilpda | hypoxia inducible lipid droplet associated | 69573 | 6 | 29272488 | 29275446 | 1 | Mm.45161 |
| 4.56 | 3.34 | 5.77 | 4.47 | DN1ab | DN1cd | -1.26 | -7.95 | 9.10E-05 | 0.011 | 0.81 | ENSMUSG00000052684 | protein_coding | Jun | jun proto-oncogene | 16476 | 4 | 95049034 | 95052222 | -1 | Mm.275071 |
| 7.17 | 6.19 | 7.4 | 6.42 | DN1ab | DN1cd | -0.98 | -7.92 | 9.40E-05 | 0.011 | 0.99 | ENSMUSG00000062232 | protein_coding | Rapgef2 | Rap guanine nucleotide exchange factor (GEF) 2 | 76089 | 3 | 79062516 | 79286517 | -1 | Mm.31220 Mm.462396 |
| 4.35 | 5.1 | 4.41 | 5.26 | DN1ab | DN1cd | 0.79 | 7.88 | 9.60E-05 | 0.012 | 0.66 | ENSMUSG00000083757 | transcribed_processed_pseudogene | Rps15a-ps4 | ribosomal protein S15A, pseudogene 4 | NA | 4 | 132219988 | 132220489 | -1 | |
| -3.82 | -1.54 | -3.94 | -0.59 | DN1ab | DN1cd | 2.69 | 7.88 | 9.70E-05 | 0.012 | 0.13 | ENSMUSG00000058626 | protein_coding | Capn11 | calpain 11 | 268958 | 17 | 45630204 | 45659325 | -1 | Mm.426026 |
| 6.93 | 6.51 | 7.48 | 7.08 | DN1ab | DN1cd | -0.41 | -7.86 | 9.80E-05 | 0.012 | 0.85 | ENSMUSG00000028649 | protein_coding | Macf1 | microtubule-actin crosslinking factor 1 | 11426 | 4 | 123349633 | 123684360 | -1 | Mm.402299 Mm.485351 |
| -0.86 | 1.07 | -1.83 | 0.67 | DN1ab | DN1cd | 2.11 | 7.86 | 9.80E-05 | 0.012 | 0.36 | ENSMUSG00000041075 | protein_coding | Fzd7 | frizzled class receptor 7 | 14369 | 1 | 59482424 | 59486955 | 1 | Mm.297906 |
| 0.89 | 3.64 | 1.05 | 3.31 | DN1ab | DN1cd | 2.6 | 7.86 | 9.80E-05 | 0.012 | 0.53 | ENSMUSG00000091695 | protein_coding | Gm17669 | predicted gene, 17669 | NA | 18 | 67562387 | 67562851 | 1 | |
| 0.63 | -3.01 | 0.85 | -3.24 | DN1ab | DN1cd | -3.83 | -7.86 | 9.80E-05 | 0.012 | 0.68 | ENSMUSG00000081860 | processed_pseudogene | Gm11533 | predicted gene 11533 | NA | 11 | 97037007 | 97037483 | 1 | |
| 0.09 | -1.93 | 0.13 | -1.7 | DN1ab | DN1cd | -1.94 | -7.85 | 9.90E-05 | 0.012 | 0.75 | ENSMUSG00000026809 | protein_coding | Spaca9 | sperm acrosome associated 9 | 69987 | 2 | 28692080 | 28699727 | -1 | Mm.239376 |
| 2.4 | 0.45 | 3.4 | 1.39 | DN1ab | DN1cd | -1.98 | -7.82 | 1.00E-04 | 0.012 | 0.91 | ENSMUSG00000112734 | sense_intronic | Gm47644 | predicted gene, 47644 | NA | 10 | 53337753 | 53345085 | -1 | |
| 4.16 | 4.74 | 4.28 | 4.73 | DN1ab | DN1cd | 0.53 | 7.81 | 1.00E-04 | 0.012 | 0.44 | ENSMUSG00000072694 | protein_coding | 1500011B03Rik | RIKEN cDNA 1500011B03 gene | 330173 | 5 | 114808196 | 114823468 | -1 | Mm.482205 |
| 6.66 | 6.2 | 7.15 | 6.5 | DN1ab | DN1cd | -0.53 | -7.79 | 1.00E-04 | 0.012 | 0.17 | ENSMUSG00000005054 | protein_coding | Cstb | cystatin B | 13014 | 10 | 78425669 | 78427622 | 1 | Mm.6095 |
| -0.5 | -3.86 | 2.28 | -1.64 | DN1ab | DN1cd | -3.59 | -7.79 | 1.00E-04 | 0.012 | 0.59 | ENSMUSG00000106557 | TEC | Gm42442 | predicted gene 42442 | NA | 5 | 131697221 | 131698353 | -1 | |
| 8.08 | 8.42 | 7.81 | 8.09 | DN1ab | DN1cd | 0.32 | 7.79 | 1.00E-04 | 0.012 | 0.52 | ENSMUSG00000021427 | protein_coding | Ssr1 | signal sequence receptor, alpha | 107513 | 13 | 37966605 | 37994217 | -1 | Mm.426670 Mm.490298 |
| 2.32 | 3.85 | 2.24 | 4.08 | DN1ab | DN1cd | 1.63 | 7.77 | 0.00011 | 0.012 | 0.52 | ENSMUSG00000043587 | protein_coding | Pxylp1 | 2-phosphoxylose phosphatase 1 | 235534 | 9 | 96823336 | 96892669 | -1 | Mm.233031 |
| 2.33 | 3.78 | 1.93 | 3.46 | DN1ab | DN1cd | 1.48 | 7.74 | 0.00011 | 0.012 | 0.88 | ENSMUSG00000032478 | protein_coding | Nme6 | NME/NM23 nucleoside diphosphate kinase 6 | 54369 | 9 | 109832749 | 109843116 | 1 | Mm.12951 |
| 4.31 | 1.4 | 6.14 | 2.52 | DN1ab | DN1cd | -3.33 | -7.71 | 0.00011 | 0.012 | 0.46 | ENSMUSG00000029673 | protein_coding | Auts2 | autism susceptibility candidate 2 | NA | 5 | 131437333 | 132543344 | -1 | |
| 5.33 | 4.29 | 5.42 | 4.69 | DN1ab | DN1cd | -0.91 | -7.71 | 0.00011 | 0.012 | 0.22 | ENSMUSG00000038871 | protein_coding | Bpgm | 2,3-bisphosphoglycerate mutase | 12183 | 6 | 34476207 | 34505613 | 1 | Mm.282863 |
| 5.38 | 4.65 | 6.04 | 5.31 | DN1ab | DN1cd | -0.73 | -7.7 | 0.00011 | 0.012 | 0.98 | ENSMUSG00000005447 | protein_coding | Pafah1b3 | platelet-activating factor acetylhydrolase, isoform 1b, subunit 3 | 18476 | 7 | 25295049 | 25297986 | -1 | Mm.597 |
| 7.76 | 7.22 | 7.6 | 7.2 | DN1ab | DN1cd | -0.48 | -7.69 | 0.00011 | 0.012 | 0.29 | ENSMUSG00000000751 | protein_coding | Rpa1 | replication protein A1 | 68275 | 11 | 75298166 | 75348324 | -1 | Mm.102574 Mm.180734 Mm.254027 Mm.472099 |
| 0.42 | -3.86 | 1.82 | -1.51 | DN1ab | DN1cd | -3.87 | -7.69 | 0.00011 | 0.012 | 0.39 | ENSMUSG00000000869 | protein_coding | Il4 | interleukin 4 | 16189 | 11 | 53602982 | 53618669 | -1 | Mm.276360 |
| 2.83 | 1.64 | 2.28 | 1.33 | DN1ab | DN1cd | -1.09 | -7.68 | 0.00011 | 0.012 | 0.43 | ENSMUSG00000020160 | protein_coding | Meis1 | Meis homeobox 1 | 17268 | 11 | 18879817 | 19018985 | -1 | Mm.356578 Mm.445192 |
| 4.47 | 3.22 | 5.96 | 4.39 | DN1ab | DN1cd | -1.4 | -7.67 | 0.00011 | 0.012 | 0.4 | ENSMUSG00000024300 | protein_coding | Myo1f | myosin IF | 17916 | 17 | 33555707 | 33607764 | 1 | Mm.42019 |
| 5.36 | 5.97 | 5.46 | 6.28 | DN1ab | DN1cd | 0.69 | 7.66 | 0.00012 | 0.013 | 0.29 | ENSMUSG00000040433 | protein_coding | Zbtb38 | zinc finger and BTB domain containing 38 | 245007 | 9 | 96682770 | 96752831 | -1 | Mm.485954 |
| 1.69 | 3.5 | 2.31 | 3.96 | DN1ab | DN1cd | 1.75 | 7.66 | 0.00012 | 0.013 | 0.77 | ENSMUSG00000023947 | protein_coding | Nfkbie | nuclear factor of kappa light polypeptide gene enhancer in B cells inhibitor, epsilon | 18037 | 17 | 45555703 | 45563169 | 1 | Mm.57043 |
| 5.25 | 4.51 | 5.2 | 4.66 | DN1ab | DN1cd | -0.66 | -7.65 | 0.00012 | 0.013 | 0.29 | ENSMUSG00000097230 | lincRNA | Gm26853 | predicted gene, 26853 | NA | 8 | 19789798 | 19798548 | 1 | |
| 5.72 | 4.94 | 6.4 | 5.2 | DN1ab | DN1cd | -0.95 | -7.64 | 0.00012 | 0.013 | 0.11 | ENSMUSG00000038304 | protein_coding | Cd160 | CD160 antigen | 54215 | 3 | 96798763 | 96829351 | -1 | Mm.34693 |
| 6.05 | 6.43 | 5.62 | 6.07 | DN1ab | DN1cd | 0.41 | 7.62 | 0.00012 | 0.013 | 0.6 | ENSMUSG00000022351 | protein_coding | Sqle | squalene epoxidase | 20775 | 15 | 59315077 | 59331192 | 1 | Mm.296169 |
| -1.5 | -3.86 | -2.36 | -4.02 | DN1ab | DN1cd | -2.11 | -7.6 | 0.00012 | 0.013 | 0.25 | ENSMUSG00000094257 | processed_pseudogene | Ap3s1-ps2 | adaptor-related protein complex 3, sigma 1 subunit, pseudogene 2 | NA | 8 | 94405211 | 94405789 | 1 | |
| 3.3 | 1.88 | 5.51 | 3.11 | DN1ab | DN1cd | -1.92 | -7.59 | 0.00012 | 0.013 | 0.037 | ENSMUSG00000036526 | protein_coding | Card11 | caspase recruitment domain family, member 11 | 108723 | 5 | 140872990 | 141000582 | -1 | Mm.46187 |
| 5.33 | 5.88 | 5.59 | 6.3 | DN1ab | DN1cd | 0.61 | 7.58 | 0.00012 | 0.013 | 0.37 | ENSMUSG00000027506 | protein_coding | Tpd52 | tumor protein D52 | 21985 | 3 | 8925593 | 9004723 | -1 | Mm.371590 |
| 3.7 | 4.72 | 3.93 | 4.93 | DN1ab | DN1cd | 1.01 | 7.57 | 0.00013 | 0.013 | 0.95 | ENSMUSG00000025035 | protein_coding | Arl3 | ADP-ribosylation factor-like 3 | 56350 | 19 | 46531109 | 46573085 | -1 | Mm.22085 |
| 6.58 | 6.94 | 6.68 | 7.16 | DN1ab | DN1cd | 0.4 | 7.56 | 0.00013 | 0.013 | 0.3 | ENSMUSG00000027508 | protein_coding | Pag1 | phosphoprotein associated with glycosphingolipid microdomains 1 | 94212 | 3 | 9687479 | 9833679 | -1 | Mm.23897 Mm.486503 |
| 3.29 | 0.94 | 4.78 | 0.83 | DN1ab | DN1cd | -3.11 | -7.56 | 0.00013 | 0.013 | 0.038 | ENSMUSG00000032420 | protein_coding | Nt5e | 5' nucleotidase, ecto | 23959 | 9 | 88327197 | 88372092 | 1 | Mm.244235 |
| 0.19 | 1.83 | 0.97 | 2.16 | DN1ab | DN1cd | 1.49 | 7.55 | 0.00013 | 0.013 | 0.32 | ENSMUSG00000104704 | TEC | Gm43133 | predicted gene 43133 | NA | 5 | 123561525 | 123561987 | -1 | |
| 2.15 | 3.27 | 3.08 | 4.28 | DN1ab | DN1cd | 1.15 | 7.55 | 0.00013 | 0.013 | 0.83 | ENSMUSG00000056144 | protein_coding | Trim34a | tripartite motif-containing 34A | 94094 | 7 | 104244457 | 104262234 | 1 | Mm.326945 |
| 4.54 | 5.26 | 5.17 | 5.61 | DN1ab | DN1cd | 0.62 | 7.54 | 0.00013 | 0.013 | 0.11 | ENSMUSG00000041831 | protein_coding | Sytl3 | synaptotagmin-like 3 | 83672 | 17 | 6659093 | 6738044 | 1 | Mm.442201 |
| -0.81 | 2.32 | -2.28 | 2.09 | DN1ab | DN1cd | 3.5 | 7.53 | 0.00013 | 0.013 | 0.25 | ENSMUSG00000021263 | protein_coding | Degs2 | delta(4)-desaturase, sphingolipid 2 | 70059 | 12 | 108678711 | 108702315 | -1 | Mm.207605 |
| 2.02 | 3.52 | 3.62 | 4.7 | DN1ab | DN1cd | 1.35 | 7.51 | 0.00013 | 0.013 | 0.3 | ENSMUSG00000066258 | protein_coding | Trim12a | tripartite motif-containing 12A | 76681 | 7 | 104299894 | 104315466 | -1 | Mm.458309 Mm.486388 |
| 6.67 | 7.27 | 6.45 | 7.08 | DN1ab | DN1cd | 0.62 | 7.5 | 0.00013 | 0.013 | 0.86 | ENSMUSG00000031604 | protein_coding | Msmo1 | methylsterol monoxygenase 1 | 66234 | 8 | 64718139 | 64733792 | -1 | Mm.30119 |
| 5.01 | 6.25 | 4.88 | 6.16 | DN1ab | DN1cd | 1.25 | 7.49 | 0.00013 | 0.013 | 0.92 | ENSMUSG00000032018 | protein_coding | Sc5d | sterol-C5-desaturase | 235293 | 9 | 42251595 | 42264256 | -1 | Mm.32700 Mm.392107 Mm.470142 |
| 2.4 | 0.3 | 4.1 | 1.78 | DN1ab | DN1cd | -2.22 | -7.48 | 0.00013 | 0.013 | 0.74 | ENSMUSG00000020424 | protein_coding | Castor1 | cytosolic arginine sensor for mTORC1 subunit 1 | 71962 | 11 | 4218225 | 4222409 | 1 | Mm.7852 |
| 4.58 | 5.07 | 4.09 | 4.73 | DN1ab | DN1cd | 0.55 | 7.48 | 0.00013 | 0.013 | 0.37 | ENSMUSG00000018428 | protein_coding | Akap1 | A kinase (PRKA) anchor protein 1 | 11640 | 11 | 88830792 | 88864586 | -1 | Mm.2969 |
| 2.69 | 1.26 | 3.67 | 2.17 | DN1ab | DN1cd | -1.47 | -7.48 | 0.00014 | 0.013 | 0.88 | ENSMUSG00000028977 | protein_coding | Casz1 | castor zinc finger 1 | 69743 | 4 | 148804429 | 148954889 | 1 | Mm.233879 |
| 6.51 | 7.13 | 5.83 | 6.68 | DN1ab | DN1cd | 0.7 | 7.47 | 0.00014 | 0.013 | 0.3 | ENSMUSG00000044573 | protein_coding | Acp1 | acid phosphatase 1, soluble | 11431 | 12 | 30893326 | 30911589 | -1 | Mm.359831 |
| 5.57 | 4.38 | 6.6 | 4.92 | DN1ab | DN1cd | -1.4 | -7.47 | 0.00014 | 0.013 | 0.22 | ENSMUSG00000030142 | protein_coding | Clec4e | C-type lectin domain family 4, member e | 56619 | 6 | 123281789 | 123289870 | -1 | Mm.248327 |
| 8.76 | 9.14 | 8.54 | 8.88 | DN1ab | DN1cd | 0.37 | 7.45 | 0.00014 | 0.013 | 0.71 | ENSMUSG00000053317 | protein_coding | Sec61b | Sec61 beta subunit | 66212 | 4 | 47474658 | 47483242 | 1 | Mm.20188 |
| 5.19 | 4.54 | 5.29 | 4.28 | DN1ab | DN1cd | -0.79 | -7.45 | 0.00014 | 0.013 | 0.1 | ENSMUSG00000031749 | protein_coding | St3gal2 | ST3 beta-galactoside alpha-2,3-sialyltransferase 2 | 20444 | 8 | 110919922 | 110972480 | 1 | Mm.200388 |
| 5.66 | 6.54 | 5.88 | 6.89 | DN1ab | DN1cd | 0.93 | 7.45 | 0.00014 | 0.013 | 0.67 | ENSMUSG00000015355 | protein_coding | Cd48 | CD48 antigen | 12506 | 1 | 171682009 | 171705258 | 1 | Mm.1738 |
| 3.87 | 2.7 | 4.4 | 3.69 | DN1ab | DN1cd | -0.96 | -7.45 | 0.00014 | 0.013 | 0.062 | ENSMUSG00000002602 | protein_coding | Axl | AXL receptor tyrosine kinase | 26362 | 7 | 25757273 | 25788705 | -1 | Mm.4128 |
| 6.58 | 5.66 | 6.6 | 5.91 | DN1ab | DN1cd | -0.83 | -7.43 | 0.00014 | 0.013 | 0.34 | ENSMUSG00000032221 | protein_coding | Mns1 | meiosis-specific nuclear structural protein 1 | 17427 | 9 | 72438011 | 72462025 | 1 | Mm.387671 |
| 4.68 | 2.23 | 6.81 | 3.79 | DN1ab | DN1cd | -2.77 | -7.43 | 0.00014 | 0.013 | 0.48 | ENSMUSG00000069833 | protein_coding | Ahnak | AHNAK nucleoprotein (desmoyokin) | 66395 | 19 | 8989284 | 9076919 | 1 | Mm.203866 Mm.392302 |
| 2.71 | -0.88 | 4 | 1.62 | DN1ab | DN1cd | -2.8 | -7.42 | 0.00014 | 0.013 | 0.13 | ENSMUSG00000022868 | protein_coding | Ahsg | alpha-2-HS-glycoprotein | 11625 | 16 | 22891277 | 22899449 | 1 | Mm.197554 |
| 6.26 | 5.45 | 7.31 | 6.26 | DN1ab | DN1cd | -0.91 | -7.42 | 0.00014 | 0.013 | 0.37 | ENSMUSG00000038155 | protein_coding | Gstp2 | glutathione S-transferase, pi 2 | 14869 | 19 | 4040285 | 4042236 | -1 | Mm.441542 |
| 2.65 | -1.18 | 2.98 | -2.85 | DN1ab | DN1cd | -4.8 | -7.41 | 0.00014 | 0.013 | 0.13 | ENSMUSG00000006154 | protein_coding | Eps8l1 | EPS8-like 1 | 67425 | 7 | 4460674 | 4480487 | 1 | Mm.360392 |
| 0.57 | -1.93 | 2.84 | -0.1 | DN1ab | DN1cd | -2.72 | -7.4 | 0.00014 | 0.013 | 0.59 | ENSMUSG00000032220 | protein_coding | Myo1e | myosin IE | 71602 | 9 | 70207350 | 70399766 | 1 | Mm.249311 Mm.444670 |
| 8.71 | 8.09 | 8.48 | 8.15 | DN1ab | DN1cd | -0.51 | -7.39 | 0.00015 | 0.013 | 0.031 | ENSMUSG00000025066 | protein_coding | Sfr1 | SWI5 dependent recombination repair 1 | 67788 | 19 | 47731756 | 47735588 | 1 | Mm.181610 |
| -3.82 | -1.1 | -3.94 | -0.89 | DN1ab | DN1cd | 2.84 | 7.38 | 0.00015 | 0.014 | 0.71 | ENSMUSG00000071234 | protein_coding | Syndig1l | synapse differentiation inducing 1 like | 627191 | 12 | 84677277 | 84698831 | -1 | Mm.152484 |
| 2.76 | -0.28 | 3.47 | 1.56 | DN1ab | DN1cd | -2.36 | -7.38 | 0.00015 | 0.014 | 0.075 | ENSMUSG00000034177 | protein_coding | Rnf43 | ring finger protein 43 | 207742 | 11 | 87662722 | 87735539 | 1 | Mm.440230 |
| -0.5 | -3.12 | 2.58 | -0.56 | DN1ab | DN1cd | -2.85 | -7.37 | 0.00015 | 0.014 | 0.54 | ENSMUSG00000021411 | protein_coding | Pxdc1 | PX domain containing 1 | 66895 | 13 | 34627834 | 34653486 | -1 | Mm.28943 |
| 3.95 | 5.81 | 2.85 | 5.42 | DN1ab | DN1cd | 2.09 | 7.36 | 0.00015 | 0.014 | 0.27 | ENSMUSG00000063320 | protein_coding | 1190007I07Rik | RIKEN cDNA 1190007I07 gene | 544717 | 10 | 82619851 | 82623228 | -1 | Mm.334867 |
| 2.48 | 1.22 | 3.68 | 2.26 | DN1ab | DN1cd | -1.34 | -7.35 | 0.00015 | 0.014 | 0.68 | ENSMUSG00000063889 | protein_coding | Crem | cAMP responsive element modulator | 12916 | 18 | 3266048 | 3337748 | -1 | Mm.5244 |
| 4.83 | 3.25 | 6.87 | 4.35 | DN1ab | DN1cd | -2.04 | -7.34 | 0.00015 | 0.014 | 0.084 | ENSMUSG00000068227 | protein_coding | Il2rb | interleukin 2 receptor, beta chain | 16185 | 15 | 78479256 | 78495271 | -1 | Mm.35287 Mm.384038 |
| 6.41 | 5.77 | 7.34 | 6.39 | DN1ab | DN1cd | -0.77 | -7.33 | 0.00015 | 0.014 | 0.15 | ENSMUSG00000022565 | protein_coding | Plec | plectin | 18810 | 15 | 76170974 | 76232574 | -1 | Mm.234912 |
| 1.91 | 0.03 | 2.63 | 0.5 | DN1ab | DN1cd | -2 | -7.33 | 0.00015 | 0.014 | 0.68 | ENSMUSG00000032737 | protein_coding | Inppl1 | inositol polyphosphate phosphatase-like 1 | 16332 | 7 | 101822632 | 101838229 | -1 | Mm.476000 |
| 6.69 | 6.26 | 7.13 | 6.41 | DN1ab | DN1cd | -0.54 | -7.33 | 0.00015 | 0.014 | 0.04 | ENSMUSG00000009090 | protein_coding | Ap1b1 | adaptor protein complex AP-1, beta 1 subunit | 11764 | 11 | 4986824 | 5042791 | 1 | Mm.274816 |
| 5.64 | 6.49 | 5.48 | 6.53 | DN1ab | DN1cd | 0.93 | 7.32 | 0.00016 | 0.014 | 0.5 | ENSMUSG00000057531 | protein_coding | Dtnbp1 | dystrobrevin binding protein 1 | 94245 | 13 | 44922075 | 45002147 | -1 | Mm.352311 |
| 5.94 | 5.48 | 6.26 | 5.92 | DN1ab | DN1cd | -0.41 | -7.31 | 0.00016 | 0.014 | 0.35 | ENSMUSG00000021549 | protein_coding | Rasa1 | RAS p21 protein activator 1 | 218397 | 13 | 85214780 | 85289130 | -1 | Mm.259653 |
| 3.43 | -0.68 | 4.56 | 2.04 | DN1ab | DN1cd | -2.99 | -7.29 | 0.00016 | 0.014 | 0.066 | ENSMUSG00000031397 | protein_coding | Tktl1 | transketolase-like 1 | 83553 | X | 74177259 | 74208500 | 1 | Mm.25057 |
| 8.54 | 7.91 | 8.88 | 7.92 | DN1ab | DN1cd | -0.76 | -7.29 | 0.00016 | 0.014 | 0.12 | ENSMUSG00000027009 | protein_coding | Itga4 | integrin alpha 4 | 16401 | 2 | 79255426 | 79333123 | 1 | Mm.31903 |
| 6.78 | 5.97 | 7.72 | 6.77 | DN1ab | DN1cd | -0.86 | -7.29 | 0.00016 | 0.014 | 0.6 | ENSMUSG00000031304 | protein_coding | Il2rg | interleukin 2 receptor, gamma chain | 16186 | X | 101264378 | 101268255 | -1 | Mm.2923 |
| 5.29 | 5.74 | 5.59 | 6.21 | DN1ab | DN1cd | 0.52 | 7.28 | 0.00016 | 0.014 | 0.28 | ENSMUSG00000001741 | protein_coding | Il16 | interleukin 16 | 16170 | 7 | 83642825 | 83745726 | -1 | Mm.10137 |
| 9.99 | 9.39 | 10.61 | 9.55 | DN1ab | DN1cd | -0.77 | -7.27 | 0.00016 | 0.014 | 0.016 | ENSMUSG00000068220 | protein_coding | Lgals1 | lectin, galactose binding, soluble 1 | 16852 | 15 | 78926725 | 78930465 | 1 | Mm.43831 |
| 5.4 | 4.7 | 5.78 | 5.25 | DN1ab | DN1cd | -0.63 | -7.26 | 0.00016 | 0.014 | 0.37 | ENSMUSG00000003199 | protein_coding | Mpnd | MPN domain containing | 68047 | 17 | 56005711 | 56016790 | 1 | Mm.358710 |
| 7.43 | 6.73 | 8.7 | 7.63 | DN1ab | DN1cd | -0.85 | -7.26 | 0.00016 | 0.014 | 0.11 | ENSMUSG00000023048 | protein_coding | Prr13 | proline rich 13 | 66151 | 15 | 102459028 | 102462806 | 1 | Mm.393955 |
| 2.56 | 3.86 | 2.74 | 5.03 | DN1ab | DN1cd | 1.65 | 7.26 | 0.00016 | 0.014 | 0.021 | ENSMUSG00000038146 | protein_coding | Notch3 | notch 3 | 18131 | 17 | 32120820 | 32166880 | -1 | Mm.439741 |
| 1.35 | -0.52 | 1.93 | 0.5 | DN1ab | DN1cd | -1.66 | -7.25 | 0.00016 | 0.014 | 0.37 | ENSMUSG00000106427 | TEC | Gm42820 | predicted gene 42820 | NA | 3 | 98121588 | 98124650 | 1 | |
| 4.45 | 5.65 | 4.24 | 5.92 | DN1ab | DN1cd | 1.36 | 7.21 | 0.00017 | 0.015 | 0.23 | ENSMUSG00000036452 | protein_coding | Arhgap26 | Rho GTPase activating protein 26 | 71302 | 18 | 38601534 | 39376284 | 1 | Mm.329396 |
| 7.1 | 8.05 | 6.24 | 7.43 | DN1ab | DN1cd | 1.03 | 7.2 | 0.00017 | 0.015 | 0.44 | ENSMUSG00000027478 | protein_coding | Dnmt3b | DNA methyltransferase 3B | 13436 | 2 | 153649450 | 153687730 | 1 | Mm.89772 Mm.491804 |
| 5.1 | 5.52 | 4.81 | 5.44 | DN1ab | DN1cd | 0.5 | 7.19 | 0.00017 | 0.015 | 0.17 | ENSMUSG00000025986 | protein_coding | Slc39a10 | solute carrier family 39 (zinc transporter), member 10 | 227059 | 1 | 46807544 | 46892852 | -1 | Mm.233889 Mm.475398 |
| 0.04 | 1.74 | 0.81 | 2.72 | DN1ab | DN1cd | 1.77 | 7.19 | 0.00017 | 0.015 | 0.72 | ENSMUSG00000042401 | protein_coding | Crtac1 | cartilage acidic protein 1 | 72832 | 19 | 42283037 | 42431783 | -1 | Mm.313558 |
| 1.67 | 3.22 | 0.92 | 2.82 | DN1ab | DN1cd | 1.66 | 7.19 | 0.00017 | 0.015 | 0.51 | ENSMUSG00000036858 | protein_coding | Ptcra | pre T cell antigen receptor alpha | 19208 | 17 | 46755663 | 46763712 | -1 | Mm.215173 |
| -3.82 | -0.24 | -2.81 | -0.71 | DN1ab | DN1cd | 3.08 | 7.19 | 0.00017 | 0.015 | 0.1 | ENSMUSG00000112294 | bidirectional_promoter_lncRNA | Gm4129 | predicted gene 4129 | NA | 10 | 123196695 | 123199418 | 1 | |
| 4.28 | 3.52 | 4.99 | 4.11 | DN1ab | DN1cd | -0.81 | -7.18 | 0.00017 | 0.015 | 0.64 | ENSMUSG00000024426 | protein_coding | Atat1 | alpha tubulin acetyltransferase 1 | 73242 | 17 | 35897595 | 35910075 | -1 | Mm.273155 |
| 5.37 | 4.51 | 6.43 | 5.38 | DN1ab | DN1cd | -0.95 | -7.18 | 0.00018 | 0.015 | 0.52 | ENSMUSG00000103653 | processed_pseudogene | Gstp-ps | glutathione S-transferase, pi, pseudogene | NA | 1 | 192073820 | 192074450 | -1 | |
| 7.18 | 6.26 | 7.33 | 6.24 | DN1ab | DN1cd | -0.99 | -7.17 | 0.00018 | 0.015 | 0.59 | ENSMUSG00000090460 | processed_pseudogene | Gm17511 | predicted gene, 17511 | NA | 7 | 127286044 | 127286385 | -1 | |
| -0.4 | -3.86 | 0.74 | -4.02 | DN1ab | DN1cd | -3.98 | -7.16 | 0.00018 | 0.015 | 0.28 | ENSMUSG00000106450 | TEC | Gm43018 | predicted gene 43018 | NA | 5 | 132439255 | 132440529 | -1 | |
| 6.02 | 5.57 | 5.95 | 5.56 | DN1ab | DN1cd | -0.43 | -7.16 | 0.00018 | 0.015 | 0.67 | ENSMUSG00000038126 | protein_coding | Mphosph9 | M-phase phosphoprotein 9 | 269702 | 5 | 124250959 | 124327972 | -1 | Mm.60224 |
| 7.15 | 4.58 | 9.59 | 6.1 | DN1ab | DN1cd | -3.02 | -7.16 | 0.00018 | 0.015 | 0.3 | ENSMUSG00000055435 | protein_coding | Maf | avian musculoaponeurotic fibrosarcoma oncogene homolog | 17132 | 8 | 115682942 | 115707794 | -1 | Mm.275549 Mm.439772 Mm.488377 |
| 5.35 | 4.95 | 5.82 | 5.36 | DN1ab | DN1cd | -0.42 | -7.15 | 0.00018 | 0.015 | 0.62 | ENSMUSG00000000605 | protein_coding | Clcn4 | chloride channel, voltage-sensitive 4 | 12727 | 7 | 7282309 | 7300851 | -1 | Mm.297883 Mm.488576 |
| 4.27 | 3.18 | 5.46 | 4.16 | DN1ab | DN1cd | -1.19 | -7.14 | 0.00018 | 0.015 | 0.58 | ENSMUSG00000029822 | protein_coding | Osbpl3 | oxysterol binding protein-like 3 | 71720 | 6 | 50293330 | 50456201 | -1 | Mm.44153 |
| 5.52 | 4.97 | 5.95 | 5.34 | DN1ab | DN1cd | -0.57 | -7.14 | 0.00018 | 0.015 | 0.72 | ENSMUSG00000074825 | protein_coding | Itpripl1 | inositol 1,4,5-triphosphate receptor interacting protein-like 1 | 73338 | 2 | 127138772 | 127143442 | -1 | Mm.57559 |
| 9.23 | 9.64 | 9 | 9.59 | DN1ab | DN1cd | 0.48 | 7.13 | 0.00018 | 0.015 | 0.26 | ENSMUSG00000019982 | protein_coding | Myb | myeloblastosis oncogene | 17863 | 10 | 21124935 | 21160984 | -1 | Mm.52109 Mm.473872 |
| 8.39 | 9.1 | 8.88 | 9.61 | DN1ab | DN1cd | 0.72 | 7.12 | 0.00018 | 0.015 | 0.94 | ENSMUSG00000024014 | protein_coding | Pim1 | proviral integration site 1 | 18712 | 17 | 29490753 | 29496112 | 1 | Mm.405293 Mm.485038 |
| 5.89 | 5.18 | 6.31 | 5.67 | DN1ab | DN1cd | -0.68 | -7.12 | 0.00018 | 0.015 | 0.73 | ENSMUSG00000034786 | protein_coding | Gpsm3 | G-protein signalling modulator 3 (AGS3-like, C. elegans) | 106512 | 17 | 34589806 | 34591754 | 1 | Mm.26584 |
| 5.63 | 4.95 | 6 | 5.54 | DN1ab | DN1cd | -0.59 | -7.11 | 0.00019 | 0.015 | 0.2 | ENSMUSG00000020732 | protein_coding | Rab37 | RAB37, member RAS oncogene family | 58222 | 11 | 115091431 | 115162236 | 1 | Mm.143789 |
| 3.64 | 2.14 | 5.55 | 3.58 | DN1ab | DN1cd | -1.74 | -7.09 | 0.00019 | 0.015 | 0.37 | ENSMUSG00000046532 | protein_coding | Ar | androgen receptor | 11835 | X | 98148769 | 98323215 | 1 | Mm.39005 Mm.394224 Mm.439657 |
| 3.93 | 3.43 | 4.67 | 3.99 | DN1ab | DN1cd | -0.57 | -7.08 | 0.00019 | 0.015 | 0.32 | ENSMUSG00000008384 | protein_coding | Sertad1 | SERTA domain containing 1 | 55942 | 7 | 27486910 | 27490316 | 1 | Mm.153684 |
| 4.06 | 4.84 | 4.04 | 4.91 | DN1ab | DN1cd | 0.81 | 7.08 | 0.00019 | 0.015 | 0.72 | ENSMUSG00000076441 | protein_coding | Ass1 | argininosuccinate synthetase 1 | 11898 | 2 | 31470207 | 31520672 | 1 | Mm.3217 |
| 6.23 | 5.81 | 6.22 | 5.81 | DN1ab | DN1cd | -0.42 | -7.08 | 0.00019 | 0.015 | 0.93 | ENSMUSG00000071796 | transcribed_unprocessed_pseudogene | 6820431F20Rik | RIKEN cDNA 6820431F20 gene | NA | 8 | 20262613 | 20297432 | -1 | |
| 4.02 | 5.13 | 4.2 | 5.54 | DN1ab | DN1cd | 1.19 | 7.07 | 0.00019 | 0.015 | 0.56 | ENSMUSG00000049717 | protein_coding | Lig4 | ligase IV, DNA, ATP-dependent | 319583 | 8 | 9969049 | 9977686 | -1 | Mm.80584 |
| 4.36 | 3.13 | 5.84 | 4.52 | DN1ab | DN1cd | -1.27 | -7.06 | 0.00019 | 0.015 | 0.82 | ENSMUSG00000003363 | protein_coding | Pld3 | phospholipase D family, member 3 | 18807 | 7 | 27532000 | 27553218 | -1 | Mm.6483 |
| 1.42 | 0.13 | 2.3 | 0.97 | DN1ab | DN1cd | -1.31 | -7.06 | 0.00019 | 0.015 | 0.92 | ENSMUSG00000112239 | processed_pseudogene | Gm17823 | predicted gene, 17823 | NA | 10 | 53299137 | 53300142 | -1 | |
| 5.19 | 6.07 | 4.19 | 5.56 | DN1ab | DN1cd | 1.05 | 7.03 | 2.00E-04 | 0.015 | 0.13 | ENSMUSG00000078202 | protein_coding | Nrarp | Notch-regulated ankyrin repeat protein | 67122 | 2 | 25180758 | 25183339 | 1 | Mm.46539 |
| 2.16 | 0.8 | 2.64 | 1.56 | DN1ab | DN1cd | -1.23 | -7.02 | 2.00E-04 | 0.015 | 0.46 | ENSMUSG00000033102 | protein_coding | Cdc14b | CDC14 cell division cycle 14B | 218294 | 13 | 64189268 | 64275290 | -1 | Mm.25335 |
| 3.37 | 0.93 | 4 | 2.3 | DN1ab | DN1cd | -2.03 | -7.01 | 2.00E-04 | 0.015 | 0.23 | ENSMUSG00000070056 | protein_coding | Mfhas1 | malignant fibrous histiocytoma amplified sequence 1 | 52065 | 8 | 35587798 | 35679449 | 1 | Mm.103246 |
| 1.4 | -1.93 | 2.1 | -1.39 | DN1ab | DN1cd | -3.4 | -7 | 2.00E-04 | 0.015 | 0.89 | ENSMUSG00000046442 | protein_coding | Ppm1e | protein phosphatase 1E (PP2C domain containing) | 320472 | 11 | 87226906 | 87359023 | -1 | Mm.341988 |
| 7.37 | 6.88 | 8.05 | 7.23 | DN1ab | DN1cd | -0.61 | -7 | 2.00E-04 | 0.015 | 0.046 | ENSMUSG00000031799 | protein_coding | Tpm4 | tropomyosin 4 | 326618 | 8 | 72134975 | 72153142 | 1 | Mm.295124 |
| 7 | 6.48 | 7.33 | 6.5 | DN1ab | DN1cd | -0.64 | -7 | 0.00021 | 0.015 | 0.092 | ENSMUSG00000071662 | protein_coding | Polr2g | polymerase (RNA) II (DNA directed) polypeptide G | 67710 | 19 | 8793129 | 8798557 | -1 | Mm.22117 |
| 2.71 | 4.49 | 3.55 | 4.99 | DN1ab | DN1cd | 1.66 | 6.97 | 0.00021 | 0.016 | 0.52 | ENSMUSG00000020297 | protein_coding | Nsg2 | neuron specific gene family member 2 | 18197 | 11 | 32000463 | 32059202 | 1 | Mm.3304 |
| 2.42 | 0.8 | 4.92 | 2.05 | DN1ab | DN1cd | -2.28 | -6.97 | 0.00021 | 0.016 | 0.042 | ENSMUSG00000025094 | protein_coding | Slc18a2 | solute carrier family 18 (vesicular monoamine), member 2 | 214084 | 19 | 59260878 | 59296012 | 1 | Mm.268797 Mm.361919 |
| 5.97 | 6.48 | 5.89 | 6.3 | DN1ab | DN1cd | 0.48 | 6.96 | 0.00021 | 0.016 | 0.49 | ENSMUSG00000017286 | protein_coding | Glod4 | glyoxalase domain containing 4 | 67201 | 11 | 76120987 | 76243725 | -1 | Mm.482113 |
| 6.22 | 6.88 | 6.67 | 7.08 | DN1ab | DN1cd | 0.56 | 6.95 | 0.00021 | 0.016 | 0.14 | ENSMUSG00000022305 | protein_coding | Lrp12 | low density lipoprotein-related protein 12 | 239393 | 15 | 39870589 | 39943994 | -1 | Mm.39874 |
| 5.73 | 6.36 | 5.45 | 6.08 | DN1ab | DN1cd | 0.63 | 6.94 | 0.00022 | 0.016 | 0.97 | ENSMUSG00000094870 | protein_coding | Zfp131 | zinc finger protein 131 | 72465 | 13 | 119765187 | 119790889 | -1 | Mm.139115 Mm.446549 |
| -3.82 | -1.54 | -3.94 | -0.48 | DN1ab | DN1cd | 2.73 | 6.93 | 0.00022 | 0.016 | 0.16 | ENSMUSG00000039269 | protein_coding | 2300002M23Rik | RIKEN cDNA 2300002M23 gene | 69542 | 17 | 35567450 | 35568952 | 1 | Mm.265291 |
| 5.98 | 6.46 | 5.85 | 6.46 | DN1ab | DN1cd | 0.52 | 6.92 | 0.00022 | 0.016 | 0.41 | ENSMUSG00000036097 | protein_coding | Slf2 | SMC5-SMC6 complex localization factor 2 | 226151 | 19 | 44931119 | 44983787 | 1 | Mm.212609 |
| -0.71 | -3.86 | 1.23 | -4.02 | DN1ab | DN1cd | -3.98 | -6.92 | 0.00022 | 0.016 | 0.063 | ENSMUSG00000105624 | TEC | Gm43187 | predicted gene 43187 | NA | 5 | 132490343 | 132492739 | -1 | |
| -2.62 | 1.13 | -3.94 | 1.26 | DN1ab | DN1cd | 4.21 | 6.92 | 0.00022 | 0.016 | 0.3 | ENSMUSG00000051682 | protein_coding | Treml4 | triggering receptor expressed on myeloid cells-like 4 | 224840 | 17 | 48264295 | 48275360 | 1 | Mm.131234 |
| 4.86 | 4.13 | 4.88 | 4.35 | DN1ab | DN1cd | -0.66 | -6.92 | 0.00022 | 0.016 | 0.33 | ENSMUSG00000037148 | protein_coding | Arhgap10 | Rho GTPase activating protein 10 | 78514 | 8 | 77250366 | 77517953 | -1 | Mm.165974 |
| 7.58 | 7.95 | 7.41 | 7.87 | DN1ab | DN1cd | 0.4 | 6.91 | 0.00022 | 0.016 | 0.46 | ENSMUSG00000031807 | protein_coding | Pgls | 6-phosphogluconolactonase | 66171 | 8 | 71592176 | 71601092 | 1 | Mm.282284 Mm.402679 |
| 5.59 | 5.16 | 5.8 | 5.45 | DN1ab | DN1cd | -0.4 | -6.89 | 0.00023 | 0.016 | 0.48 | ENSMUSG00000026848 | protein_coding | Tor1b | torsin family 1, member B | 30934 | 2 | 30952959 | 30959015 | 1 | Mm.249164 |
| 5.07 | 3.08 | 5.43 | 4.11 | DN1ab | DN1cd | -1.66 | -6.89 | 0.00023 | 0.016 | 0.18 | ENSMUSG00000008136 | protein_coding | Fhl2 | four and a half LIM domains 2 | 14200 | 1 | 43123074 | 43196984 | -1 | Mm.6799 |
| 7.51 | 8.06 | 7.3 | 7.68 | DN1ab | DN1cd | 0.49 | 6.89 | 0.00023 | 0.016 | 0.32 | ENSMUSG00000016528 | protein_coding | Mapkapk2 | MAP kinase-activated protein kinase 2 | 17164 | 1 | 131053700 | 131097826 | -1 | Mm.221235 Mm.490983 |
| 5.45 | 4.83 | 6.16 | 5.56 | DN1ab | DN1cd | -0.61 | -6.88 | 0.00023 | 0.016 | 0.93 | ENSMUSG00000022105 | protein_coding | Rb1 | RB transcriptional corepressor 1 | 19645 | 14 | 73183673 | 73325822 | -1 | Mm.273862 |
| -3.82 | -0.96 | -3.94 | -0.89 | DN1ab | DN1cd | 2.93 | 6.88 | 0.00023 | 0.016 | 0.85 | ENSMUSG00000105933 | sense_intronic | Gm43260 | predicted gene 43260 | NA | 3 | 104180612 | 104183041 | -1 | |
| 4.55 | 3.44 | 5.02 | 4.22 | DN1ab | DN1cd | -0.97 | -6.88 | 0.00023 | 0.016 | 0.3 | ENSMUSG00000034863 | protein_coding | Ano8 | anoctamin 8 | 382014 | 8 | 71476019 | 71485963 | -1 | Mm.260360 Mm.475713 |
| 8.24 | 8.58 | 7.73 | 8.35 | DN1ab | DN1cd | 0.44 | 6.87 | 0.00023 | 0.016 | 0.016 | ENSMUSG00000022797 | protein_coding | Tfrc | transferrin receptor | 22042 | 16 | 32608920 | 32632794 | 1 | Mm.28683 |
| 4.54 | 3.45 | 6.03 | 4.04 | DN1ab | DN1cd | -1.49 | -6.87 | 0.00023 | 0.016 | 0.021 | ENSMUSG00000026509 | protein_coding | Capn2 | calpain 2 | 12334 | 1 | 182467260 | 182517608 | -1 | Mm.19306 Mm.474138 |
| 4.96 | 3.72 | 5.41 | 3.97 | DN1ab | DN1cd | -1.33 | -6.86 | 0.00023 | 0.016 | 0.65 | ENSMUSG00000026435 | protein_coding | Slc45a3 | solute carrier family 45, member 3 | 212980 | 1 | 131962967 | 131982969 | 1 | Mm.200307 |
| 2.09 | 3.69 | 2.8 | 4.18 | DN1ab | DN1cd | 1.52 | 6.85 | 0.00023 | 0.016 | 0.67 | ENSMUSG00000020178 | protein_coding | Adora2a | adenosine A2a receptor | 11540 | 10 | 75316877 | 75334784 | 1 | Mm.333734 |
| 3.85 | 3.05 | 4.51 | 3.67 | DN1ab | DN1cd | -0.82 | -6.85 | 0.00023 | 0.016 | 0.91 | ENSMUSG00000071649 | protein_coding | B3gat3 | beta-1,3-glucuronyltransferase 3 (glucuronosyltransferase I) | 72727 | 19 | 8920374 | 8927234 | 1 | Mm.259 |
| -1.81 | -3.86 | -0.77 | -4.02 | DN1ab | DN1cd | -2.48 | -6.85 | 0.00023 | 0.016 | 0.11 | ENSMUSG00000024397 | protein_coding | Aif1 | allograft inflammatory factor 1 | 11629 | 17 | 35170991 | 35176068 | -1 | Mm.10747 |
| 6.21 | 5.88 | 6.24 | 5.78 | DN1ab | DN1cd | -0.38 | -6.85 | 0.00023 | 0.016 | 0.33 | ENSMUSG00000074746 | protein_coding | Pdzd8 | PDZ domain containing 8 | 107368 | 19 | 59296084 | 59345780 | -1 | Mm.27250 Mm.268797 Mm.361919 |
| 7.75 | 6.63 | 8.38 | 7.08 | DN1ab | DN1cd | -1.19 | -6.85 | 0.00023 | 0.016 | 0.65 | ENSMUSG00000025997 | protein_coding | Ikzf2 | IKAROS family zinc finger 2 | 22779 | 1 | 69531214 | 69687245 | -1 | Mm.106343 |
| 6.65 | 6.1 | 7.34 | 6.49 | DN1ab | DN1cd | -0.66 | -6.83 | 0.00024 | 0.016 | 0.12 | ENSMUSG00000022102 | protein_coding | Dok2 | docking protein 2 | 13449 | 14 | 70766036 | 70778495 | 1 | Mm.243323 |
| 3.88 | 3.2 | 4.39 | 3.44 | DN1ab | DN1cd | -0.79 | -6.82 | 0.00024 | 0.016 | 0.29 | ENSMUSG00000031728 | protein_coding | Zfp821 | zinc finger protein 821 | 75871 | 8 | 109705546 | 109724932 | 1 | Mm.180410 Mm.286964 |
| 7.04 | 6.67 | 7.78 | 7.23 | DN1ab | DN1cd | -0.44 | -6.82 | 0.00024 | 0.016 | 0.2 | ENSMUSG00000061288 | protein_coding | Taok3 | TAO kinase 3 | 330177 | 5 | 117120129 | 117275219 | 1 | Mm.24343 Mm.487518 |
| 3.86 | 3.22 | 4.36 | 3.76 | DN1ab | DN1cd | -0.62 | -6.8 | 0.00024 | 0.017 | 0.85 | ENSMUSG00000104724 | TEC | Gm43162 | predicted gene 43162 | NA | 3 | 138976312 | 138978081 | -1 | |
| 4.65 | 3.75 | 6.29 | 5.52 | DN1ab | DN1cd | -0.84 | -6.8 | 0.00025 | 0.017 | 0.64 | ENSMUSG00000026222 | protein_coding | Sp100 | nuclear antigen Sp100 | 20684 | 1 | 85649988 | 85709998 | 1 | Mm.290906 Mm.486514 |
| 0.95 | 2.72 | 0.51 | 2.74 | DN1ab | DN1cd | 1.92 | 6.79 | 0.00025 | 0.017 | 0.49 | ENSMUSG00000078240 | protein_coding | Gm3550 | predicted gene 3550 | NA | 18 | 34736359 | 34738014 | 1 | |
| 6.49 | 5.86 | 7.33 | 6.26 | DN1ab | DN1cd | -0.81 | -6.79 | 0.00025 | 0.017 | 0.06 | ENSMUSG00000024789 | protein_coding | Jak2 | Janus kinase 2 | 16452 | 19 | 29251828 | 29313080 | 1 | Mm.275839 |
| 0.65 | -2.78 | 1.55 | -3.24 | DN1ab | DN1cd | -4.02 | -6.78 | 0.00025 | 0.017 | 0.28 | ENSMUSG00000104994 | TEC | Gm42989 | predicted gene 42989 | NA | 5 | 132370376 | 132372681 | -1 | |
| 3.07 | 3.78 | 2.33 | 3.43 | DN1ab | DN1cd | 0.85 | 6.77 | 0.00025 | 0.017 | 0.14 | ENSMUSG00000031586 | protein_coding | Rbpms | RNA binding protein gene with multiple splicing | 19663 | 8 | 33782643 | 33929863 | -1 | Mm.323997 |
| 8.26 | 7.67 | 8.98 | 8.3 | DN1ab | DN1cd | -0.62 | -6.76 | 0.00026 | 0.017 | 0.63 | ENSMUSG00000026728 | protein_coding | Vim | vimentin | 22352 | 2 | 13573927 | 13582826 | 1 | Mm.268000 |
| 1.51 | -2.24 | 2.72 | -1.82 | DN1ab | DN1cd | -4.13 | -6.75 | 0.00026 | 0.017 | 0.56 | ENSMUSG00000104936 | TEC | 2610011E03Rik | RIKEN cDNA 2610011E03 gene | NA | 5 | 131996179 | 131998413 | -1 | |
| 3.6 | 1.93 | 4.2 | 2.47 | DN1ab | DN1cd | -1.7 | -6.75 | 0.00026 | 0.017 | 0.92 | ENSMUSG00000097296 | lincRNA | Gm26532 | predicted gene, 26532 | NA | 8 | 84208920 | 84209609 | 1 | |
| -0.74 | -3.86 | -2.36 | -4.02 | DN1ab | DN1cd | -2.59 | -6.75 | 0.00026 | 0.017 | 0.055 | ENSMUSG00000091618 | protein_coding | H60c | histocompatibility 60c | 670558 | 10 | 3256208 | 3267771 | -1 | Mm.398582 |
| 4.18 | 2.9 | 5.79 | 3.62 | DN1ab | DN1cd | -1.7 | -6.74 | 0.00026 | 0.017 | 0.07 | ENSMUSG00000015846 | protein_coding | Rxra | retinoid X receptor alpha | 20181 | 2 | 27676440 | 27762957 | 1 | Mm.24624 |
| 3.14 | 2.18 | 4.18 | 2.83 | DN1ab | DN1cd | -1.13 | -6.74 | 0.00026 | 0.017 | 0.28 | ENSMUSG00000112571 | TEC | Gm48207 | predicted gene, 48207 | NA | 10 | 105532984 | 105534493 | -1 | |
| -3.82 | -0.87 | -2.36 | -0.71 | DN1ab | DN1cd | 2.48 | 6.73 | 0.00026 | 0.017 | 0.086 | ENSMUSG00000103835 | sense_intronic | Gm37612 | predicted gene, 37612 | NA | 9 | 106870674 | 106871501 | 1 | |
| 4.79 | 5.71 | 4.13 | 5.54 | DN1ab | DN1cd | 1.09 | 6.73 | 0.00026 | 0.017 | 0.16 | ENSMUSG00000030161 | protein_coding | Gabarapl1 | gamma-aminobutyric acid (GABA) A receptor-associated protein-like 1 | 57436 | 6 | 129533160 | 129542346 | 1 | Mm.14638 |
| 7.05 | 6.7 | 7.5 | 6.99 | DN1ab | DN1cd | -0.41 | -6.73 | 0.00026 | 0.017 | 0.23 | ENSMUSG00000025135 | protein_coding | Anapc11 | anaphase promoting complex subunit 11 | 66156 | 11 | 120598421 | 120608198 | 1 | Mm.21645 |
| 5.33 | 5.93 | 5.3 | 5.67 | DN1ab | DN1cd | 0.52 | 6.72 | 0.00026 | 0.017 | 0.16 | ENSMUSG00000033257 | protein_coding | Ttll4 | tubulin tyrosine ligase-like family, member 4 | 67534 | 1 | 74661745 | 74703730 | 1 | Mm.260714 |

| | | | | | | | | | | | | | | | | | | | | |
|---|---|---|---|---|---|---|---|---|---|---|---|---|---|---|---|---|---|---|---|---|
| -3.82 | -1.93 | -3.94 | -1.46 | DN1ab | DN1cd | 2.11 | 6.72 | 0.00026 | 0.017 | 0.4 | ENSMUSG00000111514 | lincRNA | E230014E18Rik | RIKEN cDNA E230014E18 gene | NA | 10 | 84518045 | 84523270 | 1 | |
| 3.78 | 4.6 | 4.17 | 5.04 | DN1ab | DN1cd | 0.84 | 6.71 | 0.00027 | 0.017 | 0.88 | ENSMUSG00000021298 | protein_coding | Gpr132 | G protein-coupled receptor 132 | 56696 | 12 | 112850873 | 112868228 | -1 | Mm.20455 |
| 4.83 | 3.9 | 5.16 | 4.05 | DN1ab | DN1cd | -1 | -6.71 | 0.00027 | 0.017 | 0.58 | ENSMUSG00000026035 | protein_coding | Ppil3 | peptidylprolyl isomerase (cyclophilin)-like 3 | 70225 | 1 | 58430994 | 58445486 | -1 | Mm.340195 |
| -3.82 | -1.93 | -3.94 | -1.03 | DN1ab | DN1cd | 2.27 | 6.7 | 0.00027 | 0.017 | 0.15 | ENSMUSG00000103670 | antisense | Gm37622 | predicted gene, 37622 | NA | 1 | 155231304 | 155233159 | 1 | |
| 4.25 | 3.07 | 4.6 | 3.37 | DN1ab | DN1cd | -1.2 | -6.69 | 0.00027 | 0.017 | 0.91 | ENSMUSG00000017897 | protein_coding | Eya2 | EYA transcriptional coactivator and phosphatase 2 | 14049 | 2 | 165595032 | 165771727 | 1 | Mm.282719 |
| 0.58 | -1.54 | 0.83 | -0.89 | DN1ab | DN1cd | -1.95 | -6.68 | 0.00027 | 0.017 | 0.54 | ENSMUSG00000073000 | lincRNA | Gm10451 | predicted gene 10451 | NA | 12 | 76444497 | 76451336 | 1 | |
| 8.6 | 8.29 | 8.58 | 8.21 | DN1ab | DN1cd | -0.34 | -6.68 | 0.00027 | 0.017 | 0.61 | ENSMUSG00000026341 | protein_coding | Actr3 | ARP3 actin-related protein 3 | 74117 | 1 | 125392905 | 125435727 | -1 | Mm.183102 Mm.471990 |
| 3.73 | 1.91 | 5.58 | 4.09 | DN1ab | DN1cd | -1.63 | -6.67 | 0.00028 | 0.018 | 0.56 | ENSMUSG00000026604 | protein_coding | Ptpn14 | protein tyrosine phosphatase, non-receptor type 14 | 19250 | 1 | 189728268 | 189876695 | 1 | Mm.4498 |
| 0.57 | -1.54 | 1.34 | 0.06 | DN1ab | DN1cd | -1.72 | -6.65 | 0.00028 | 0.018 | 0.11 | ENSMUSG00000007867 | protein_coding | Ift43 | intraflagellar transport 43 | 76411 | 12 | 86082541 | 86162459 | 1 | Mm.292345 |
| 5.4 | 5.94 | 5.44 | 6.02 | DN1ab | DN1cd | 0.56 | 6.65 | 0.00028 | 0.018 | 0.83 | ENSMUSG00000024513 | protein_coding | Mbd2 | methyl-CpG binding domain protein 2 | 17191 | 18 | 70568189 | 70626131 | 1 | Mm.322 |
| 3.47 | 2.88 | 4.36 | 3.59 | DN1ab | DN1cd | -0.67 | -6.64 | 0.00028 | 0.018 | 0.43 | ENSMUSG00000037946 | protein_coding | Fgd3 | FYVE, RhoGEF and PH domain containing 3 | 30938 | 13 | 49261554 | 49320311 | -1 | Mm.291089 |
| 3.85 | 2.45 | 4.65 | 3.85 | DN1ab | DN1cd | -1.1 | -6.64 | 0.00029 | 0.018 | 0.057 | ENSMUSG00000019947 | protein_coding | Arid5b | AT rich interactive domain 5B (MRF1-like) | 71371 | 10 | 68092520 | 68278740 | -1 | Mm.440357 |
| 6.61 | 5.89 | 6.67 | 6.04 | DN1ab | DN1cd | -0.68 | -6.62 | 0.00029 | 0.018 | 0.68 | ENSMUSG00000020642 | protein_coding | Rnf144a | ring finger protein 144A | 108089 | 12 | 26300964 | 26415254 | -1 | Mm.214932 Mm.407984 |
| 0.84 | 2.94 | 0.67 | 2.44 | DN1ab | DN1cd | 1.99 | 6.61 | 0.00029 | 0.018 | 0.64 | ENSMUSG00000108095 | lincRNA | Gm44067 | predicted gene, 44067 | NA | 6 | 128843786 | 128846959 | -1 | |
| 5.15 | 5.95 | 5.38 | 5.87 | DN1ab | DN1cd | 0.69 | 6.6 | 0.00029 | 0.018 | 0.16 | ENSMUSG00000027999 | protein_coding | Pla2g12a | phospholipase A2, group XIIA | 66350 | 3 | 129878606 | 129895825 | 1 | Mm.151951 |
| 5.88 | 4.93 | 5.97 | 5.48 | DN1ab | DN1cd | -0.76 | -6.6 | 3.00E-04 | 0.018 | 0.035 | ENSMUSG00000037369 | protein_coding | Kdm6a | lysine (K)-specific demethylase 6A | 22289 | X | 18162575 | 18279936 | 1 | Mm.257498 Mm.417709 |
| 3.17 | 0.28 | 3.05 | 1.25 | DN1ab | DN1cd | -2.3 | -6.6 | 3.00E-04 | 0.018 | 0.13 | ENSMUSG00000002104 | protein_coding | Rapsn | receptor-associated protein of the synapse | 19400 | 2 | 91035620 | 91045729 | 1 | Mm.1272 |
| 4.46 | 4.99 | 4.39 | 5.19 | DN1ab | DN1cd | 0.63 | 6.59 | 3.00E-04 | 0.018 | 0.19 | ENSMUSG00000039753 | protein_coding | Fbxl5 | F-box and leucine-rich repeat protein 5 | 242960 | 5 | 43744615 | 43821638 | -1 | Mm.25794 |
| 6.77 | 5.92 | 7.91 | 6.66 | DN1ab | DN1cd | -1.01 | -6.59 | 3.00E-04 | 0.018 | 0.21 | ENSMUSG00000018340 | protein_coding | Anxa6 | annexin A6 | 11749 | 11 | 54979108 | 55033445 | -1 | Mm.265347 |
| -2.24 | -3.86 | -2.36 | -4.02 | DN1ab | DN1cd | -1.64 | -6.59 | 3.00E-04 | 0.018 | 0.94 | ENSMUSG00000105868 | TEC | Gm43766 | predicted gene 43766 | NA | 14 | 52258811 | 52264227 | 1 | |
| -2.24 | -3.86 | -2.36 | -4.02 | DN1ab | DN1cd | -1.64 | -6.59 | 3.00E-04 | 0.018 | 0.94 | ENSMUSG00000049526 | protein_coding | Tmem202 | transmembrane protein 202 | 73893 | 9 | 59518685 | 59539847 | -1 | Mm.216135 |
| 5.9 | 4.22 | 7.35 | 4.88 | DN1ab | DN1cd | -2.04 | -6.58 | 3.00E-04 | 0.018 | 0.23 | ENSMUSG00000039308 | protein_coding | Ndst2 | N-deacetylase/N-sulfotransferase (heparan glucosaminyl) 2 | 17423 | 14 | 20723730 | 20734562 | -1 | Mm.4084 |
| 3.4 | 4.64 | 2.59 | 4.27 | DN1ab | DN1cd | 1.39 | 6.58 | 3.00E-04 | 0.018 | 0.36 | ENSMUSG00000036186 | protein_coding | Fam69b | family with sequence similarity 69, member B | 56279 | 2 | 26628457 | 26636497 | 1 | Mm.279863 |
| 5.01 | 4.19 | 6.06 | 5.23 | DN1ab | DN1cd | -0.82 | -6.56 | 0.00031 | 0.019 | 0.96 | ENSMUSG00000064043 | processed_transcript | Trerf1 | transcriptional regulating factor 1 | 224829 | 17 | 47140875 | 47361958 | 1 | Mm.260989 |
| 1.28 | -1.14 | 1.45 | -0.59 | DN1ab | DN1cd | -2.23 | -6.55 | 0.00031 | 0.019 | 0.61 | ENSMUSG00000019935 | protein_coding | Slc17a8 | solute carrier family 17 (sodium-dependent inorganic phosphate cotransporter), member 8 | 216227 | 10 | 89574020 | 89621253 | -1 | Mm.233921 |
| 0.25 | -3.12 | 2.61 | -1.03 | DN1ab | DN1cd | -3.5 | -6.55 | 0.00031 | 0.019 | 0.82 | ENSMUSG00000085133 | sense_intronic | B930095G15Rik | RIKEN cDNA B930095G15 gene | NA | 14 | 121248706 | 121253971 | 1 | |
| 7.57 | 7.86 | 7.45 | 7.82 | DN1ab | DN1cd | 0.32 | 6.55 | 0.00031 | 0.019 | 0.53 | ENSMUSG00000021670 | protein_coding | Hmgcr | 3-hydroxy-3-methylglutaryl-Coenzyme A reductase | 15357 | 13 | 96648967 | 96670936 | -1 | Mm.316652 |
| 5.03 | 5.65 | 3.95 | 4.66 | DN1ab | DN1cd | 0.65 | 6.55 | 0.00031 | 0.019 | 0.7 | ENSMUSG00000057359 | processed_transcript | Gm17494 | predicted gene, 17494 | NA | 3 | 122255958 | 122293720 | -1 | |
| 3.63 | 2.95 | 4.34 | 3.77 | DN1ab | DN1cd | -0.63 | -6.54 | 0.00031 | 0.019 | 0.62 | ENSMUSG00000061731 | protein_coding | Ext1 | exostoses (multiple) 1 | 14042 | 15 | 53064038 | 53346159 | -1 | Mm.309395 |
| -2.24 | -3.86 | -1.83 | -4.02 | DN1ab | DN1cd | -1.83 | -6.53 | 0.00031 | 0.019 | 0.36 | ENSMUSG00000107577 | TEC | Gm44103 | predicted gene, 44103 | NA | 6 | 116260434 | 116264095 | -1 | |
| 6.96 | 6.41 | 7.87 | 7.11 | DN1ab | DN1cd | -0.64 | -6.52 | 0.00032 | 0.019 | 0.34 | ENSMUSG00000000594 | protein_coding | Gm2a | GM2 ganglioside activator protein | 14667 | 11 | 55098115 | 55113029 | 1 | Mm.287807 |
| 2.46 | -0.6 | 2.52 | -0.12 | DN1ab | DN1cd | -2.84 | -6.52 | 0.00032 | 0.019 | 0.66 | ENSMUSG00000026836 | protein_coding | Acvr1 | activin A receptor, type 1 | 11477 | 2 | 58388644 | 58567157 | -1 | Mm.689 |
| 4.61 | 2.41 | 5.97 | 4.42 | DN1ab | DN1cd | -1.84 | -6.51 | 0.00032 | 0.019 | 0.27 | ENSMUSG00000026012 | protein_coding | Cd28 | CD28 antigen | 12487 | 1 | 60716800 | 60773359 | 1 | Mm.255003 |
| 6.32 | 6.98 | 6.35 | 6.96 | DN1ab | DN1cd | 0.65 | 6.51 | 0.00032 | 0.019 | 0.81 | ENSMUSG00000055723 | protein_coding | Rras2 | related RAS viral (r-ras) oncogene 2 | 66922 | 7 | 114046782 | 114117781 | -1 | Mm.276572 |
| 2.92 | 4.39 | 4.16 | 5.08 | DN1ab | DN1cd | 1.28 | 6.51 | 0.00032 | 0.019 | 0.19 | ENSMUSG00000027333 | protein_coding | Smox | spermine oxidase | 228608 | 2 | 131491496 | 131525922 | 1 | Mm.136586 |
| -3.82 | -2.28 | -3.94 | -1.7 | DN1ab | DN1cd | 1.8 | 6.5 | 0.00032 | 0.019 | 0.25 | ENSMUSG00000040329 | protein_coding | Il7 | interleukin 7 | 16196 | 3 | 7569994 | 7613760 | -1 | Mm.3825 |
| -3.82 | -2.28 | -3.94 | -1.7 | DN1ab | DN1cd | 1.8 | 6.5 | 0.00032 | 0.019 | 0.25 | ENSMUSG00000038296 | protein_coding | Galnt18 | polypeptide N-acetylgalactosaminyltransferase 18 | 233733 | 7 | 111471661 | 111779977 | -1 | Mm.323322 |
| -3.82 | -2.28 | -3.94 | -1.7 | DN1ab | DN1cd | 1.8 | 6.5 | 0.00032 | 0.019 | 0.25 | ENSMUSG00000096629 | protein_coding | Gm3383 | predicted gene 3383 | 100041515 | 14 | 5607620 | 5807982 | -1 | Mm.443041 Mm.484345 |
| 7.2 | 7.99 | 6.95 | 8.19 | DN1ab | DN1cd | 0.95 | 6.5 | 0.00032 | 0.019 | 0.15 | ENSMUSG00000025492 | protein_coding | Ifitm3 | interferon induced transmembrane protein 3 | 66141 | 7 | 141009586 | 141010770 | -1 | Mm.141021 |
| 6.09 | 5.47 | 6.47 | 6.12 | DN1ab | DN1cd | -0.52 | -6.5 | 0.00033 | 0.019 | 0.086 | ENSMUSG00000071379 | protein_coding | Hpcal1 | hippocalcin-like 1 | 53602 | 12 | 17690856 | 17791933 | 1 | Mm.313619 Mm.489835 |
| 9.17 | 9.44 | 8.57 | 8.87 | DN1ab | DN1cd | 0.29 | 6.49 | 0.00033 | 0.019 | 0.72 | ENSMUSG00000058799 | protein_coding | Nap1l1 | nucleosome assembly protein 1-like 1 | 53605 | 10 | 111473223 | 111498150 | 1 | Mm.290407 |
| 6.85 | 7.27 | 6.64 | 7.06 | DN1ab | DN1cd | 0.42 | 6.48 | 0.00033 | 0.019 | 0.93 | ENSMUSG00000027708 | protein_coding | Dcun1d1 | DCN1, defective in cullin neddylation 1, domain containing 1 (S. cerevisiae) | 114893 | 3 | 35892105 | 35937445 | -1 | Mm.374810 Mm.379305 Mm.474495 |
| 4.06 | 5.16 | 3.06 | 5.13 | DN1ab | DN1cd | 1.43 | 6.48 | 0.00033 | 0.019 | 0.02 | ENSMUSG00000055541 | protein_coding | Lair1 | leukocyte-associated Ig-like receptor 1 | 52855 | 7 | 4003402 | 4063204 | -1 | Mm.290880 |
| 4.78 | 5.28 | 4.54 | 5.26 | DN1ab | DN1cd | 0.58 | 6.47 | 0.00033 | 0.019 | 0.27 | ENSMUSG00000063439 | protein_coding | B9d2 | B9 protein domain 2 | 232987 | 7 | 25680780 | 25686558 | 1 | Mm.22849 |
| -0.86 | 1.2 | 0.33 | 2.58 | DN1ab | DN1cd | 2.12 | 6.47 | 0.00033 | 0.019 | 0.81 | ENSMUSG00000112023 | protein_coding | Lilr4b | leukocyte immunoglobulin-like receptor, subfamily B, member 4B | 14727 | 10 | 51480632 | 51486703 | 1 | Mm.439661 |
| 7.76 | 7.39 | 7.85 | 7.5 | DN1ab | DN1cd | -0.36 | -6.46 | 0.00034 | 0.019 | 0.86 | ENSMUSG00000009013 | protein_coding | Dynll1 | dynein light chain LC8-type 1 | 56455 | 5 | 115297110 | 115300999 | -1 | Mm.256858 |
| 2.43 | 3.83 | 0.7 | 3.04 | DN1ab | DN1cd | 1.7 | 6.45 | 0.00034 | 0.019 | 0.092 | ENSMUSG00000022246 | protein_coding | Rai14 | retinoic acid induced 14 | 75646 | 15 | 10568969 | 10714624 | -1 | Mm.212395 |
| 7.68 | 8.03 | 7.5 | 7.96 | DN1ab | DN1cd | 0.38 | 6.45 | 0.00034 | 0.019 | 0.42 | ENSMUSG00000019471 | protein_coding | Cdc37 | cell division cycle 37 | 12539 | 9 | 21133222 | 21149982 | -1 | Mm.32331 |
| 4.43 | 5.03 | 3.75 | 4.31 | DN1ab | DN1cd | 0.59 | 6.45 | 0.00034 | 0.019 | 0.81 | ENSMUSG00000020413 | protein_coding | Hus1 | HUS1 checkpoint clamp component | 15574 | 11 | 8993137 | 9011191 | -1 | Mm.42201 |
| 6.46 | 6.99 | 6.2 | 7.01 | DN1ab | DN1cd | 0.63 | 6.45 | 0.00034 | 0.019 | 0.21 | ENSMUSG00000040850 | protein_coding | Psme4 | proteasome (prosome, macropain) activator subunit 4 | 103554 | 11 | 30771726 | 30880361 | 1 | Mm.240066 |
| 6.33 | 5.38 | 6.94 | 5.88 | DN1ab | DN1cd | -0.99 | -6.44 | 0.00034 | 0.019 | 0.74 | ENSMUSG00000013033 | protein_coding | Adgrl1 | adhesion G protein-coupled receptor L1 | 330814 | 8 | 83900105 | 83941954 | 1 | Mm.260733 |
| 5.87 | 6.42 | 5.61 | 6.09 | DN1ab | DN1cd | 0.53 | 6.43 | 0.00034 | 0.019 | 0.73 | ENSMUSG00000000743 | protein_coding | Chmp1a | charged multivesicular body protein 1A | 234852 | 8 | 123204264 | 123212763 | -1 | Mm.41596 |
| 4.66 | 5.29 | 5.04 | 5.59 | DN1ab | DN1cd | 0.59 | 6.43 | 0.00035 | 0.019 | 0.72 | ENSMUSG00000034453 | protein_coding | Polr3b | polymerase (RNA) III (DNA directed) polypeptide B | 70428 | 10 | 84622292 | 84727178 | 1 | Mm.335007 Mm.472000 |
| 5.55 | 5.19 | 5.89 | 5.53 | DN1ab | DN1cd | -0.36 | -6.43 | 0.00035 | 0.019 | 0.95 | ENSMUSG00000020778 | protein_coding | Ten1 | TEN1 telomerase capping complex subunit | 69535 | 11 | 116198855 | 116215318 | 1 | Mm.390347 |
| 5.2 | 6.14 | 4.89 | 6.3 | DN1ab | DN1cd | 1.11 | 6.42 | 0.00035 | 0.019 | 0.21 | ENSMUSG00000029408 | protein_coding | Abcb9 | ATP-binding cassette, sub-family B (MDR/TAP), member 9 | 56325 | 5 | 124061530 | 124095798 | -1 | Mm.254839 |
| 3.63 | 2.95 | 4.46 | 3.75 | DN1ab | DN1cd | -0.7 | -6.42 | 0.00035 | 0.019 | 0.92 | ENSMUSG00000022159 | protein_coding | Rab2b | RAB2B, member RAS oncogene family | 76338 | 14 | 52261759 | 52279545 | -1 | Mm.32870 |
| 1.22 | -0.43 | 2.07 | 0.68 | DN1ab | DN1cd | -1.53 | -6.42 | 0.00035 | 0.019 | 0.62 | ENSMUSG00000078546 | protein_coding | Zfp995 | zinc finger protein 995 | 70081 | 17 | 21879570 | 21909926 | -1 | Mm.245648 |
| 5 | 3.29 | 5.71 | 3.78 | DN1ab | DN1cd | -1.82 | -6.41 | 0.00035 | 0.019 | 0.73 | ENSMUSG00000041895 | protein_coding | Wipi1 | WD repeat domain, phosphoinositide interacting 1 | 52639 | 11 | 109573331 | 109611967 | -1 | Mm.35817 |
| 7.28 | 6.37 | 7.49 | 6.9 | DN1ab | DN1cd | -0.78 | -6.41 | 0.00035 | 0.019 | 0.22 | ENSMUSG00000037138 | protein_coding | Aff3 | AF4/FMR2 family, member 3 | 16764 | 1 | 38177326 | 38664955 | -1 | Mm.336679 |
| -0.53 | 1.74 | 1.03 | 2.37 | DN1ab | DN1cd | 1.96 | 6.41 | 0.00035 | 0.019 | 0.16 | ENSMUSG00000036053 | protein_coding | Fmnl2 | formin-like 2 | 71409 | 2 | 52857860 | 53133804 | 1 | Mm.394427 |
| 6.32 | 5.58 | 7.12 | 6.42 | DN1ab | DN1cd | -0.72 | -6.4 | 0.00036 | 0.019 | 0.87 | ENSMUSG00000004665 | protein_coding | Cnn2 | calponin 2 | 12798 | 10 | 79988584 | 79996062 | 1 | Mm.157770 |
| 4.86 | 6.8 | 2.74 | 6.28 | DN1ab | DN1cd | 2.46 | 6.4 | 0.00036 | 0.019 | 0.037 | ENSMUSG00000034041 | protein_coding | Lyl1 | lymphoblastomic leukemia 1 | 17095 | 8 | 84701457 | 84704715 | 1 | Mm.4925 |
| 4.09 | 2.03 | 4.37 | 3.16 | DN1ab | DN1cd | -1.63 | -6.4 | 0.00036 | 0.019 | 0.092 | ENSMUSG00000026883 | protein_coding | Dab2ip | disabled 2 interacting protein | 69601 | 2 | 35558266 | 35730994 | 1 | Mm.29629 |
| 4.72 | 4.03 | 5.24 | 4.55 | DN1ab | DN1cd | -0.69 | -6.4 | 0.00036 | 0.019 | 1 | ENSMUSG00000050410 | protein_coding | Tcf19 | transcription factor 19 | 106795 | 17 | 35512734 | 35516824 | -1 | Mm.11434 Mm.475434 |
| -1.54 | -3.86 | -0.25 | -4.02 | DN1ab | DN1cd | -2.86 | -6.39 | 0.00036 | 0.019 | 0.12 | ENSMUSG00000107799 | antisense | Gm38825 | predicted gene, 38825 | NA | 6 | 67179799 | 67239710 | -1 | |
| -2.24 | -3.86 | -2.15 | -4.02 | DN1ab | DN1cd | -1.71 | -6.39 | 0.00036 | 0.019 | 0.69 | ENSMUSG00000107450 | processed_pseudogene | Gm2522 | predicted gene 2522 | NA | 6 | 4390920 | 4391241 | 1 | |
| 6.87 | 7.52 | 6.58 | 7.36 | DN1ab | DN1cd | 0.7 | 6.38 | 0.00036 | 0.019 | 0.61 | ENSMUSG00000027510 | protein_coding | Rbm38 | RNA binding motif protein 38 | 56190 | 2 | 173020498 | 173034734 | 1 | Mm.485782 |
| -1.81 | -3.86 | -2.36 | -4.02 | DN1ab | DN1cd | -1.91 | -6.38 | 0.00036 | 0.019 | 0.58 | ENSMUSG00000104598 | antisense | Gm30214 | predicted gene, 30214 | NA | 14 | 53032417 | 53039466 | -1 | |
| -1.81 | -3.86 | -2.36 | -4.02 | DN1ab | DN1cd | -1.91 | -6.38 | 0.00036 | 0.019 | 0.58 | ENSMUSG00000051065 | protein_coding | Mb21d2 | Mab-21 domain containing 2 | 239796 | 16 | 28826176 | 28929673 | -1 | Mm.155887 |
| 0.09 | 2.44 | 0.36 | 2.35 | DN1ab | DN1cd | 2.23 | 6.38 | 0.00036 | 0.019 | 0.67 | ENSMUSG00000074037 | protein_coding | Mc1r | melanocortin 1 receptor | 17199 | 8 | 123407107 | 123410744 | 1 | Mm.324942 |
| 4.38 | 5.03 | 4.86 | 5.32 | DN1ab | DN1cd | 0.58 | 6.38 | 0.00037 | 0.019 | 0.35 | ENSMUSG00000025372 | protein_coding | Baiap2 | brain-specific angiogenesis inhibitor 1-associated protein 2 | 108100 | 11 | 119942763 | 120006782 | 1 | Mm.197534 |
| 8.08 | 8.98 | 8.3 | 8.8 | DN1ab | DN1cd | 0.76 | 6.37 | 0.00037 | 0.02 | 0.11 | ENSMUSG00000060019 | protein_coding | Gm10073 | predicted pseudogene 10073 | NA | 8 | 106572966 | 106573461 | -1 | |
| 7.9 | 8.54 | 7.28 | 8.3 | DN1ab | DN1cd | 0.77 | 6.37 | 0.00037 | 0.02 | 0.14 | ENSMUSG00000024613 | protein_coding | Tcof1 | treacle ribosome biogenesis factor 1 | 21453 | 18 | 60813755 | 60848971 | -1 | Mm.486251 |
| 3.6 | 2.99 | 4.16 | 3.48 | DN1ab | DN1cd | -0.64 | -6.36 | 0.00037 | 0.02 | 0.75 | ENSMUSG00000078771 | protein_coding | Evi2a | ecotropic viral integration site 2a | 14017 | 11 | 79526560 | 79530609 | -1 | Mm.439665 |
| -3.82 | -1.88 | -3.94 | -1.7 | DN1ab | DN1cd | 2.05 | 6.35 | 0.00037 | 0.02 | 0.68 | ENSMUSG00000045589 | protein_coding | Frrs1l | ferric-chelate reductase 1 like | 230235 | 4 | 56957173 | 56990391 | -1 | Mm.207654 |
| 8.59 | 9.3 | 8.92 | 9.28 | DN1ab | DN1cd | 0.58 | 6.34 | 0.00038 | 0.02 | 0.053 | ENSMUSG00000034165 | protein_coding | Ccnd3 | cyclin D3 | 12445 | 17 | 47505051 | 47599691 | 1 | Mm.27291 |
| -1.5 | -3.86 | -1.14 | -2.44 | DN1ab | DN1cd | -1.98 | -6.34 | 0.00038 | 0.02 | 0.1 | ENSMUSG00000095150 | processed_pseudogene | Rps19-ps13 | ribosomal protein S19, pseudogene 13 | NA | 18 | 40726269 | 40726709 | 1 | |
| 2.97 | 4.55 | 1.85 | 4.72 | DN1ab | DN1cd | 2.01 | 6.34 | 0.00038 | 0.02 | 0.042 | ENSMUSG00000022500 | protein_coding | Litaf | LPS-induced TN factor | 56722 | 16 | 10959275 | 11066157 | -1 | Mm.294753 |
| 7.8 | 7.49 | 8.06 | 7.82 | DN1ab | DN1cd | -0.28 | -6.32 | 0.00039 | 0.02 | 0.52 | ENSMUSG00000029004 | protein_coding | Kmt2e | lysine (K)-specific methyltransferase 2E | 69188 | 5 | 23434441 | 23504235 | 1 | Mm.205190 Mm.403814 |
| 4.64 | 3.73 | 5.13 | 4.41 | DN1ab | DN1cd | -0.83 | -6.31 | 0.00039 | 0.02 | 0.54 | ENSMUSG00000071647 | protein_coding | Eml3 | echinoderm microtubule associated protein like 3 | 225898 | 19 | 8929552 | 8941582 | 1 | Mm.31008 |
| 6.06 | 5.39 | 6.97 | 6.03 | DN1ab | DN1cd | -0.78 | -6.3 | 0.00039 | 0.02 | 0.34 | ENSMUSG00000022372 | protein_coding | Sla | src-like adaptor | 20491 | 15 | 66780819 | 66831829 | -1 | Mm.7601 |
| -1.54 | -3.86 | 0.38 | -1.7 | DN1ab | DN1cd | -2.23 | -6.29 | 4.00E-04 | 0.02 | 0.77 | ENSMUSG00000092563 | processed_pseudogene | Gm3617 | predicted gene 3617 | NA | 1 | 39314899 | 39315453 | 1 | |
| 0.36 | -3.86 | 2.98 | -3.24 | DN1ab | DN1cd | -5.09 | -6.29 | 4.00E-04 | 0.02 | 0.24 | ENSMUSG00000021219 | protein_coding | Rgs6 | regulator of G-protein signaling 6 | 50779 | 12 | 82588292 | 83162056 | 1 | Mm.126176 Mm.153013 |
| 2.87 | 1.52 | 3.4 | 1.7 | DN1ab | DN1cd | -1.5 | -6.29 | 4.00E-04 | 0.02 | 0.5 | ENSMUSG00000028680 | protein_coding | Plk3 | polo like kinase 3 | 12795 | 4 | 117128655 | 117133963 | -1 | Mm.259022 |
| 4.44 | 5.13 | 4.78 | 5.53 | DN1ab | DN1cd | 0.71 | 6.28 | 4.00E-04 | 0.02 | 0.79 | ENSMUSG00000027636 | protein_coding | Sla2 | Src-like-adaptor 2 | 77799 | 2 | 156872457 | 156887192 | -1 | Mm.29264 Mm.387161 |
| 0.98 | 2.58 | 1.57 | 3.21 | DN1ab | DN1cd | 1.61 | 6.27 | 4.00E-04 | 0.021 | 0.94 | ENSMUSG00000020399 | protein_coding | Havcr2 | hepatitis A virus cellular receptor 2 | 171285 | 11 | 46454935 | 46481255 | 1 | Mm.72168 |
| 5.74 | 6.39 | 5.36 | 6.31 | DN1ab | DN1cd | 0.75 | 6.26 | 0.00041 | 0.021 | 0.27 | ENSMUSG00000001123 | protein_coding | Lgals9 | lectin, galactose binding, soluble 9 | 16859 | 11 | 78962974 | 78984946 | -1 | Mm.341434 |
| 4.8 | 6.31 | 4.01 | 5.87 | DN1ab | DN1cd | 1.63 | 6.26 | 0.00041 | 0.021 | 0.55 | ENSMUSG00000031765 | protein_coding | Mt1 | metallothionein 1 | 17748 | 8 | 94179082 | 94180327 | 1 | Mm.192991 Mm.388249 |
| 5.86 | 6.49 | 5.98 | 7.02 | DN1ab | DN1cd | 0.78 | 6.26 | 0.00041 | 0.021 | 0.11 | ENSMUSG00000066892 | protein_coding | Fbxl12 | F-box and leucine-rich repeat protein 12 | 30843 | 9 | 20618118 | 20644767 | -1 | Mm.24608 Mm.390165 Mm.489685 |
| 5.11 | 5.7 | 5.61 | 6.56 | DN1ab | DN1cd | 0.72 | 6.26 | 0.00041 | 0.021 | 0.15 | ENSMUSG00000063286 | transcribed_unprocessed_pseudogene | Gm8995 | predicted gene 8995 | NA | 7 | 106504239 | 106511671 | -1 | |
| 2.38 | 4.33 | 2.2 | 4.32 | DN1ab | DN1cd | 2 | 6.26 | 0.00041 | 0.021 | 0.82 | ENSMUSG00000004709 | protein_coding | Cd244a | CD244 molecule A | 18106 | 1 | 171559193 | 171609746 | 1 | Mm.2299 |
| 6.08 | 6.56 | 5.83 | 6.12 | DN1ab | DN1cd | 0.41 | 6.25 | 0.00041 | 0.021 | 0.21 | ENSMUSG00000038943 | protein_coding | Prc1 | protein regulator of cytokinesis 1 | 233406 | 7 | 80294450 | 80316259 | 1 | Mm.227274 |
| 1.46 | 0.3 | 2.29 | 0.37 | DN1ab | DN1cd | -1.49 | -6.24 | 0.00042 | 0.021 | 0.12 | ENSMUSG00000081564 | processed_pseudogene | Gm13717 | predicted gene 13717 | NA | 6 | 29750761 | 29751125 | 1 | |
| 4.77 | 5.92 | 5.12 | 5.98 | DN1ab | DN1cd | 1.05 | 6.23 | 0.00042 | 0.021 | 0.45 | ENSMUSG00000026463 | protein_coding | Atp2b4 | ATPase, Ca++ transporting, plasma membrane 4 | 381290 | 1 | 133699457 | 133801041 | -1 | Mm.188617 |
| 3.77 | 3.03 | 4.39 | 3.44 | DN1ab | DN1cd | -0.83 | -6.22 | 0.00043 | 0.021 | 0.48 | ENSMUSG00000073139 | protein_coding | Tmem185a | transmembrane protein 185A | 236848 | X | 70459753 | 70477180 | -1 | Mm.24569 |
| 6.34 | 5.83 | 6.63 | 6.01 | DN1ab | DN1cd | -0.55 | -6.21 | 0.00043 | 0.021 | 0.6 | ENSMUSG00000047824 | protein_coding | Pygo2 | pygopus 2 | 68911 | 3 | 89430214 | 89435128 | 1 | Mm.22521 Mm.487671 |
| 6.44 | 6.77 | 6.56 | 7.19 | DN1ab | DN1cd | 0.44 | 6.2 | 0.00043 | 0.021 | 0.02 | ENSMUSG00000020134 | protein_coding | Peli1 | pellino 1 | 67245 | 11 | 21091291 | 21150323 | 1 | Mm.28957 |
| -3.82 | -1.31 | -2.28 | -0.13 | DN1ab | DN1cd | 2.38 | 6.2 | 0.00043 | 0.021 | 0.68 | ENSMUSG00000103662 | TEC | Gm34294 | predicted gene, 34294 | NA | 2 | 169624056 | 169628318 | -1 | |
| 4.91 | 4.12 | 5.18 | 4.45 | DN1ab | DN1cd | -0.77 | -6.2 | 0.00043 | 0.021 | 0.84 | ENSMUSG00000028221 | protein_coding | Pip4p2 | phosphatidylinositol-4,5-bisphosphate 4-phosphatase 2 | 72519 | 4 | 14864076 | 14915176 | 1 | Mm.279158 |
| 0.73 | 2.45 | -0.48 | 2.48 | DN1ab | DN1cd | 2.13 | 6.19 | 0.00044 | 0.022 | 0.085 | ENSMUSG00000033032 | protein_coding | Afap1l1 | actin filament associated protein 1-like 1 | 106877 | 18 | 61730261 | 61786702 | -1 | Mm.26632 |
| 4.64 | 3.87 | 5.16 | 4.46 | DN1ab | DN1cd | -0.74 | -6.18 | 0.00044 | 0.022 | 0.78 | ENSMUSG00000022517 | protein_coding | Mgrn1 | mahogunin, ring finger 1 | 17237 | 16 | 4886249 | 4938296 | 1 | Mm.291326 Mm.391848 |
| 6.25 | 5.75 | 6.3 | 5.75 | DN1ab | DN1cd | -0.52 | -6.18 | 0.00044 | 0.022 | 0.75 | ENSMUSG00000035329 | protein_coding | Fbxo33 | F-box protein 33 | 70611 | 12 | 59200655 | 59219725 | -1 | Mm.311026 |
| 4.16 | 2.45 | 4.56 | 2.76 | DN1ab | DN1cd | -1.75 | -6.17 | 0.00044 | 0.022 | 0.88 | ENSMUSG00000020279 | protein_coding | Il9r | interleukin 9 receptor | 16199 | 11 | 32187541 | 32200279 | -1 | Mm.384 |
| 6.7 | 5.83 | 7.81 | 6.4 | DN1ab | DN1cd | -1.09 | -6.17 | 0.00045 | 0.022 | 0.14 | ENSMUSG00000030987 | protein_coding | Stim1 | stromal interaction molecule 1 | 20866 | 7 | 102267806 | 102437319 | 1 | Mm.645 |
| -2.24 | 0.62 | -0.81 | 2.29 | DN1ab | DN1cd | 2.94 | 6.17 | 0.00045 | 0.022 | 0.83 | ENSMUSG00000029082 | protein_coding | Bst1 | bone marrow stromal cell antigen 1 | 12182 | 5 | 43818885 | 43843986 | 1 | Mm.246332 |
| -3.82 | -0.96 | -3.49 | -1.03 | DN1ab | DN1cd | 2.72 | 6.16 | 0.00045 | 0.022 | 0.69 | ENSMUSG00000022240 | protein_coding | Ctnnd2 | catenin (cadherin associated protein), delta 2 | 18163 | 15 | 30172593 | 31029341 | 1 | Mm.321648 |
| 6.52 | 7.04 | 6.69 | 7.58 | DN1ab | DN1cd | 0.66 | 6.16 | 0.00045 | 0.022 | 0.11 | ENSMUSG00000022148 | protein_coding | Fyb | FYN binding protein | 23880 | 15 | 6522853 | 6663313 | 1 | Mm.170905 |
| 6.34 | 5.5 | 7.35 | 6.27 | DN1ab | DN1cd | -0.93 | -6.16 | 0.00045 | 0.022 | 0.46 | ENSMUSG00000060803 | protein_coding | Gstp1 | glutathione S-transferase, pi 1 | 14870 | 19 | 4035407 | 4037921 | -1 | Mm.299292 |
| 3.95 | 2.04 | 6.65 | 3.82 | DN1ab | DN1cd | -2.43 | -6.16 | 0.00045 | 0.022 | 0.28 | ENSMUSG00000041235 | protein_coding | Chd7 | chromodomain helicase DNA binding protein 7 | 320790 | 4 | 8690406 | 8867659 | 1 | Mm.138792 |
| 1.44 | 3.58 | -0.35 | 3.58 | DN1ab | DN1cd | 2.71 | 6.16 | 0.00045 | 0.022 | 0.047 | ENSMUSG00000030559 | protein_coding | Rab38 | RAB38, member RAS oncogene family | 72433 | 7 | 88430273 | 88491572 | 1 | Mm.276669 |
| 6.55 | 5.61 | 6.63 | 5.67 | DN1ab | DN1cd | -0.95 | -6.15 | 0.00045 | 0.022 | 0.93 | ENSMUSG00000025026 | protein_coding | Add3 | adducin 3 (gamma) | 27360 | 19 | 53140445 | 53247399 | 1 | Mm.426080 Mm.491114 |
| 4.54 | 4.12 | 5.21 | 4.65 | DN1ab | DN1cd | -0.48 | -6.15 | 0.00046 | 0.022 | 0.41 | ENSMUSG00000015149 | protein_coding | Sirt2 | sirtuin 2 | 64383 | 7 | 28766735 | 28788661 | 1 | Mm.272443 |
| 4.83 | 4.21 | 5.01 | 4.31 | DN1ab | DN1cd | -0.65 | -6.14 | 0.00046 | 0.022 | 0.75 | ENSMUSG00000029434 | protein_coding | Vps33a | VPS33A CORVET/HOPS core subunit | 77573 | 5 | 123528659 | 123573038 | -1 | Mm.41372 |
| -2.15 | 2.17 | -2.81 | 3.2 | DN1ab | DN1cd | 4.83 | 6.14 | 0.00046 | 0.022 | 0.36 | ENSMUSG00000032278 | protein_coding | Paqr5 | progestin and adipoQ receptor family member V | 74090 | 9 | 61953481 | 62026856 | -1 | Mm.273267 |
| 1.72 | 0.29 | 2.04 | 0.93 | DN1ab | DN1cd | -1.29 | -6.14 | 0.00046 | 0.022 | 0.48 | ENSMUSG00000062300 | protein_coding | Nectin2 | nectin cell adhesion molecule 2 | 19294 | 7 | 19716644 | 19750483 | -1 | Mm.4341 |
| 6.79 | 7.22 | 6.3 | 6.79 | DN1ab | DN1cd | 0.45 | 6.14 | 0.00046 | 0.022 | 0.71 | ENSMUSG00000093314 | miRNA | Mir5136 | microRNA 5136 | NA | 19 | 8888699 | 8888774 | -1 | |
| 9.93 | 10.22 | 9.46 | 9.87 | DN1ab | DN1cd | 0.34 | 6.13 | 0.00046 | 0.022 | 0.33 | ENSMUSG00000026202 | protein_coding | Tuba4a | tubulin, alpha 4A | 22145 | 1 | 75214228 | 75219865 | -1 | Mm.1155 |
| 8.39 | 7.93 | 9.08 | 8.67 | DN1ab | DN1cd | -0.44 | -6.12 | 0.00047 | 0.022 | 0.76 | ENSMUSG00000091955 | protein_coding | Gm9844 | predicted pseudogene 9844 | NA | 7 | 24862213 | 24862697 | 1 | |
| 4.36 | 6.23 | 3.08 | 6.39 | DN1ab | DN1cd | 2.35 | 6.12 | 0.00047 | 0.022 | 0.069 | ENSMUSG00000045659 | protein_coding | Plekha7 | pleckstrin homology domain containing, family A member 7 | 233765 | 7 | 116123485 | 116308376 | -1 | Mm.3741 Mm.393600 |
| 4 | 3.15 | 4.85 | 3.75 | DN1ab | DN1cd | -0.96 | -6.12 | 0.00047 | 0.022 | 0.45 | ENSMUSG00000000791 | protein_coding | Il12rb1 | interleukin 12 receptor, beta 1 | 16161 | 8 | 70808449 | 70821424 | 1 | Mm.731 |
| 4.89 | 5.43 | 4.85 | 5.74 | DN1ab | DN1cd | 0.67 | 6.11 | 0.00047 | 0.022 | 0.11 | ENSMUSG00000052949 | protein_coding | Rnf157 | ring finger protein 157 | 217340 | 11 | 116336353 | 116413032 | -1 | Mm.426344 |
| 1.23 | 2.62 | 0.75 | 3.29 | DN1ab | DN1cd | 1.78 | 6.11 | 0.00047 | 0.022 | 0.046 | ENSMUSG00000082292 | processed_pseudogene | Gm12250 | predicted gene 12250 | NA | 11 | 58187739 | 58189012 | 1 | |
| 6.74 | 7.16 | 6.8 | 7.57 | DN1ab | DN1cd | 0.54 | 6.11 | 0.00047 | 0.022 | 0.046 | ENSMUSG00000016933 | protein_coding | Plcg1 | phospholipase C, gamma 1 | 18803 | 2 | 160731300 | 160775760 | 1 | Mm.44463 |
| 4.28 | 3.7 | 5.04 | 4.03 | DN1ab | DN1cd | -0.75 | -6.1 | 0.00048 | 0.022 | 0.068 | ENSMUSG00000049804 | protein_coding | Armcx4 | armadillo repeat containing, X-linked 4 | 100503043 | X | 134686519 | 134696757 | 1 | Mm.31961 |
| 6.03 | 5.69 | 6.4 | 5.97 | DN1ab | DN1cd | -0.37 | -6.09 | 0.00048 | 0.022 | 0.53 | ENSMUSG00000002332 | protein_coding | Dhrs1 | dehydrogenase/reductase (SDR family) member 1 | 52585 | 14 | 55739020 | 55745690 | -1 | Mm.21623 |
| 3.12 | 2.14 | 3.46 | 2.36 | DN1ab | DN1cd | -1.03 | -6.07 | 0.00049 | 0.023 | 0.77 | ENSMUSG00000109873 | TEC | Gm45407 | predicted gene 45407 | NA | 8 | 77368841 | 77372516 | -1 | |
| 5.19 | 5.68 | 5.25 | 5.98 | DN1ab | DN1cd | 0.58 | 6.07 | 0.00049 | 0.023 | 0.24 | ENSMUSG00000038893 | protein_coding | Fam117a | family with sequence similarity 117, member A | 215512 | 11 | 95337018 | 95381872 | 1 | Mm.262113 |
| 3.31 | 4.37 | 1.97 | 3.11 | DN1ab | DN1cd | 1.09 | 6.06 | 5.00E-04 | 0.023 | 0.85 | ENSMUSG00000096356 | protein_coding | Olfr889 | olfactory receptor 889 | 258475 | 9 | 38115198 | 38121245 | 1 | Mm.377522 |
| 6.25 | 5.95 | 6.22 | 5.77 | DN1ab | DN1cd | -0.35 | -6.06 | 5.00E-04 | 0.023 | 0.26 | ENSMUSG00000048371 | protein_coding | Pdp2 | pyruvate dehyrogenase phosphatase catalytic subunit 2 | 382051 | 8 | 104591451 | 104599026 | 1 | Mm.290750 |
| 4.6 | 4.01 | 5.43 | 4.96 | DN1ab | DN1cd | -0.54 | -6.05 | 5.00E-04 | 0.023 | 0.54 | ENSMUSG00000059714 | protein_coding | Flot1 | flotillin 1 | 14251 | 17 | 35823230 | 35832791 | 1 | Mm.2931 |
| 0.75 | -0.88 | 0.67 | -0.32 | DN1ab | DN1cd | -1.35 | -6.05 | 5.00E-04 | 0.023 | 0.16 | ENSMUSG00000074284 | antisense | Gm10658 | predicted gene 10658 | NA | 9 | 57056630 | 57071966 | -1 | |
| 7.04 | 6.6 | 7.62 | 6.99 | DN1ab | DN1cd | -0.52 | -6.04 | 0.00051 | 0.023 | 0.31 | ENSMUSG00000052331 | protein_coding | Ankrd44 | ankyrin repeat domain 44 | 329154 | 1 | 54645340 | 54926387 | -1 | Mm.397910 |
| 6.02 | 6.43 | 4.97 | 5.73 | DN1ab | DN1cd | 0.54 | 6.04 | 0.00051 | 0.023 | 0.05 | ENSMUSG00000025758 | protein_coding | Plk4 | polo like kinase 4 | 20873 | 3 | 40800019 | 40816883 | 1 | Mm.3794 |
| 7.19 | 6.86 | 7.29 | 6.83 | DN1ab | DN1cd | -0.38 | -6.03 | 0.00051 | 0.023 | 0.37 | ENSMUSG00000036504 | protein_coding | Phpt1 | phosphohistidine phosphatase 1 | 75454 | 2 | 25573430 | 25575224 | -1 | Mm.458079 |
| 8.23 | 7.94 | 8.49 | 8.21 | DN1ab | DN1cd | -0.29 | -6.03 | 0.00051 | 0.023 | 0.94 | ENSMUSG00000036241 | protein_coding | Ube2r2 | ubiquitin-conjugating enzyme E2R 2 | 67615 | 4 | 41135743 | 41193380 | 1 | Mm.389540 |
| 6.55 | 6.2 | 6.75 | 6.41 | DN1ab | DN1cd | -0.34 | -6.02 | 0.00052 | 0.023 | 0.95 | ENSMUSG00000005656 | protein_coding | Snx6 | sorting nexin 6 | 72183 | 12 | 54746349 | 54795703 | -1 | Mm.28240 Mm.424023 |
| 0.84 | 4.51 | -2.72 | 4.28 | DN1ab | DN1cd | 4.65 | 6.02 | 0.00052 | 0.023 | 0.033 | ENSMUSG00000069805 | protein_coding | Fbp1 | fructose bisphosphatase 1 | 14121 | 13 | 62864753 | 62888282 | -1 | Mm.423078 |
| 4.12 | 2.47 | 4.6 | 3.53 | DN1ab | DN1cd | -1.37 | -6.02 | 0.00052 | 0.023 | 0.22 | ENSMUSG00000057191 | protein_coding | AB124611 | cDNA sequence AB124611 | 382062 | 9 | 21526176 | 21545333 | 1 | Mm.325205 |
| 6.83 | 7.42 | 6.07 | 6.55 | DN1ab | DN1cd | 0.55 | 6.02 | 0.00052 | 0.023 | 0.62 | ENSMUSG00000097195 | processed_transcript | Snhg5 | small nucleolar RNA host gene 5 | NA | 9 | 88521050 | 88522929 | -1 | |
| 5.37 | 4.9 | 6.49 | 6.04 | DN1ab | DN1cd | -0.46 | -6.02 | 0.00052 | 0.023 | 0.91 | ENSMUSG00000036587 | protein_coding | Fut7 | fucosyltransferase 7 | 14347 | 2 | 25423267 | 25426374 | 1 | Mm.1203 |
| 4.52 | 5.05 | 3.89 | 4.94 | DN1ab | DN1cd | 0.72 | 6.01 | 0.00052 | 0.023 | 0.022 | ENSMUSG00000043872 | protein_coding | Zmym1 | zinc finger, MYM domain containing 1 | 68310 | 4 | 127047094 | 127061152 | -1 | Mm.273806 |
| 1.19 | 3.21 | 2.22 | 3.9 | DN1ab | DN1cd | 1.9 | 6 | 0.00053 | 0.023 | 0.64 | ENSMUSG00000105703 | processed_transcript | Gm43305 | predicted gene 43305 | NA | 14 | 54292443 | 54297456 | -1 | |
| 3.56 | 2.74 | 3.83 | 3.31 | DN1ab | DN1cd | -0.69 | -6 | 0.00053 | 0.023 | 0.21 | ENSMUSG00000102243 | TEC | Gm37718 | predicted gene, 37718 | NA | 1 | 139155152 | 139157782 | 1 | |
| 5.46 | 6.01 | 5.09 | 6.08 | DN1ab | DN1cd | 0.71 | 6 | 0.00053 | 0.023 | 0.061 | ENSMUSG00000021236 | protein_coding | Entpd5 | ectonucleoside triphosphate diphosphohydrolase 5 | 12499 | 12 | 84373857 | 84409029 | -1 | Mm.10211 |
| 7.23 | 6.48 | 8.34 | 7.84 | DN1ab | DN1cd | -0.65 | -6 | 0.00053 | 0.023 | 0.3 | ENSMUSG00000025647 | protein_coding | Shisa5 | shisa family member 5 | 66940 | 9 | 109038565 | 109057777 | 1 | Mm.196533 Mm.475600 |
| 7.42 | 6.99 | 7.53 | 6.72 | DN1ab | DN1cd | -0.57 | -5.99 | 0.00053 | 0.023 | 0.052 | ENSMUSG00000026827 | protein_coding | Gpd2 | glycerol phosphate dehydrogenase 2, mitochondrial | 14571 | 2 | 57237635 | 57370719 | 1 | Mm.3711 |
| -3.82 | -1.93 | -3.94 | -2.07 | DN1ab | DN1cd | 1.88 | 5.99 | 0.00053 | 0.023 | 0.98 | ENSMUSG00000081217 | processed_pseudogene | Gm15268 | predicted gene 15268 | NA | X | 141627959 | 141628468 | 1 | |
| -3.82 | -1.88 | -3.94 | -1.46 | DN1ab | DN1cd | 2.14 | 5.99 | 0.00054 | 0.024 | 0.5 | ENSMUSG00000042064 | protein_coding | Myo3b | myosin IIIB | 329421 | 2 | 70039126 | 70429198 | 1 | Mm.99648 |
| 5.8 | 5.11 | 6.44 | 5.23 | DN1ab | DN1cd | -0.9 | -5.98 | 0.00054 | 0.024 | 0.078 | ENSMUSG00000025417 | protein_coding | Pip4k2c | phosphatidylinositol-5-phosphate 4-kinase, type II, gamma | 117150 | 10 | 127197067 | 127211585 | -1 | Mm.22682 |
| 5.24 | 5.71 | 4.89 | 5.5 | DN1ab | DN1cd | 0.52 | 5.97 | 0.00054 | 0.024 | 0.46 | ENSMUSG00000038213 | protein_coding | Tapbpl | TAP binding protein-like | 213233 | 6 | 125223933 | 125231860 | -1 | Mm.296902 |
| 4.98 | 4.57 | 4.86 | 4.15 | DN1ab | DN1cd | -0.52 | -5.97 | 0.00055 | 0.024 | 0.08 | ENSMUSG00000030615 | protein_coding | Tmem126a | transmembrane protein 126A | 66271 | 7 | 90450700 | 90457229 | -1 | Mm.28963 |
| 1.58 | -0.08 | 3.14 | 0.7 | DN1ab | DN1cd | -2.03 | -5.96 | 0.00055 | 0.024 | 0.27 | ENSMUSG00000034675 | protein_coding | Dbn1 | drebrin 1 | 56320 | 13 | 55473429 | 55488111 | -1 | Mm.19016 |
| 2.46 | 1.59 | 3.7 | 2.43 | DN1ab | DN1cd | -1.05 | -5.94 | 0.00056 | 0.024 | 0.29 | ENSMUSG00000045827 | protein_coding | Serpinb9 | serine (or cysteine) peptidase inhibitor, clade B, member 9 | 20723 | 13 | 33003250 | 33017957 | 1 | Mm.245041 Mm.272569 |
| 5.83 | 5.28 | 5.81 | 5.14 | DN1ab | DN1cd | -0.6 | -5.94 | 0.00056 | 0.024 | 0.62 | ENSMUSG00000019878 | protein_coding | Hsf2 | heat shock factor 2 | 15500 | 10 | 57486385 | 57513135 | 1 | Mm.261395 Mm.447362 |
| 5.88 | 5.42 | 6.38 | 5.82 | DN1ab | DN1cd | -0.5 | -5.93 | 0.00056 | 0.024 | 0.62 | ENSMUSG00000029106 | protein_coding | Add1 | adducin 1 (alpha) | 11518 | 5 | 34573664 | 34632308 | 1 | Mm.289106 |
| 7.23 | 7.54 | 7.42 | 7.75 | DN1ab | DN1cd | 0.32 | 5.93 | 0.00057 | 0.024 | 0.86 | ENSMUSG00000004535 | protein_coding | Tax1bp1 | Tax1 (human T cell leukemia virus type I) binding protein 1 | 52440 | 6 | 52713729 | 52766780 | 1 | Mm.389757 |
| 8.94 | 8.5 | 9.95 | 9.49 | DN1ab | DN1cd | -0.45 | -5.93 | 0.00057 | 0.024 | 0.88 | ENSMUSG00000028581 | protein_coding | Laptm5 | lysosomal-associated protein transmembrane 5 | 16792 | 4 | 130913125 | 130936141 | 1 | Mm.271868 |
| 3.91 | 4.77 | 3.43 | 4.4 | DN1ab | DN1cd | 0.9 | 5.93 | 0.00057 | 0.024 | 0.76 | ENSMUSG00000051335 | protein_coding | Gfod1 | glucose-fructose oxidoreductase domain containing 1 | 328232 | 13 | 43195245 | 43304172 | -1 | Mm.441399 |
| 6.28 | 6.72 | 5.94 | 6.42 | DN1ab | DN1cd | 0.46 | 5.91 | 0.00058 | 0.025 | 0.83 | ENSMUSG00000091905 | processed_pseudogene | Gm6395 | predicted gene 6395 | NA | 12 | 64786311 | 64787001 | 1 | |
| 7.68 | 6.84 | 8.49 | 7.35 | DN1ab | DN1cd | -0.96 | -5.91 | 0.00058 | 0.025 | 0.39 | ENSMUSG00000020077 | protein_coding | Srgn | serglycin | 19073 | 10 | 62493833 | 62527451 | -1 | Mm.338790 |
| 4.86 | 4.27 | 5.27 | 4.81 | DN1ab | DN1cd | -0.53 | -5.91 | 0.00058 | 0.025 | 0.55 | ENSMUSG00000047810 | protein_coding | Ccdc88b | coiled-coil domain containing 88B | 78317 | 19 | 6844623 | 6858211 | -1 | Mm.329596 |
| 1.61 | 3.25 | 1.55 | 3.23 | DN1ab | DN1cd | 1.66 | 5.9 | 0.00058 | 0.025 | 0.95 | ENSMUSG00000073274 | processed_transcript | Gm14636 | predicted gene 14636 | NA | X | 19156981 | 19167241 | -1 | |
| -2.91 | 0.13 | -1.02 | 1.1 | DN1ab | DN1cd | 2.72 | 5.9 | 0.00058 | 0.025 | 0.37 | ENSMUSG00000037325 | protein_coding | Bbs7 | Bardet-Biedl syndrome 7 (human) | 71492 | 3 | 36573142 | 36613477 | -1 | Mm.286187 |
| 8.06 | 8.42 | 7.73 | 8.18 | DN1ab | DN1cd | 0.39 | 5.9 | 0.00058 | 0.025 | 0.52 | ENSMUSG00000022710 | protein_coding | Usp7 | ubiquitin specific peptidase 7 | 252870 | 16 | 8689595 | 8792308 | -1 | Mm.295330 |
| 4.16 | 4.8 | 4.28 | 4.9 | DN1ab | DN1cd | 0.63 | 5.9 | 0.00058 | 0.025 | 0.94 | ENSMUSG00000022515 | protein_coding | Anks3 | ankyrin repeat and sterile alpha motif domain containing 3 | 72615 | 16 | 4941436 | 4964205 | -1 | Mm.290628 |
| 5.93 | 6.42 | 6.4 | 7.35 | DN1ab | DN1cd | 0.66 | 5.9 | 0.00059 | 0.025 | 0.033 | ENSMUSG00000049502 | protein_coding | Dtx3l | deltex 3-like, E3 ubiquitin ligase | 209200 | 16 | 35926511 | 35939151 | -1 | Mm.390852 |
| 4.75 | 2.17 | 8.18 | 3.62 | DN1ab | DN1cd | -3.74 | -5.88 | 0.00059 | 0.025 | 0.13 | ENSMUSG00000026009 | protein_coding | Icos | inducible T cell co-stimulator | 54167 | 1 | 60977927 | 61000320 | 1 | Mm.42044 |
| 0.65 | -2.78 | 1.21 | -1.03 | DN1ab | DN1cd | -2.9 | -5.88 | 6.00E-04 | 0.025 | 0.26 | ENSMUSG00000024059 | protein_coding | Clip4 | CAP-GLY domain containing linker protein family, member 4 | 78785 | 17 | 71768473 | 71864273 | 1 | Mm.196382 |

| | | | | | | | | | | | | | | | | | | | | |
|---|---|---|---|---|---|---|---|---|---|---|---|---|---|---|---|---|---|---|---|---|
| 2 | 2.9 | 2.5 | 3.75 | DN1ab | DN1cd | 1.03 | 5.87 | 6.00E-04 | 0.025 | 0.36 | ENSMUSG00000021477 | protein_coding | Ctsl | cathepsin L | 13039 | 13 | 64359337 | 64370890 | -1 | Mm.930 Mm.471755 |
| 7.84 | 7.56 | 7.86 | 7.34 | DN1ab | DN1cd | -0.37 | -5.87 | 6.00E-04 | 0.025 | 0.074 | ENSMUSG00000022043 | protein_coding | Trim35 | tripartite motif-containing 35 | 66854 | 14 | 66297031 | 66311424 | 1 | Mm.43871 |
| 5.05 | 5.61 | 4.64 | 5.24 | DN1ab | DN1cd | 0.57 | 5.86 | 0.00061 | 0.025 | 0.86 | ENSMUSG00000005718 | protein_coding | Tfap4 | transcription factor AP4 | 83383 | 16 | 4544661 | 4559854 | -1 | Mm.273210 |
| 3.99 | 4.55 | 3.57 | 4.25 | DN1ab | DN1cd | 0.6 | 5.85 | 0.00061 | 0.025 | 0.59 | ENSMUSG00000017697 | protein_coding | Ada | adenosine deaminase | 11486 | 2 | 163726584 | 163750239 | -1 | Mm.388 |
| 5.17 | 4.18 | 6.12 | 5.4 | DN1ab | DN1cd | -0.87 | -5.85 | 0.00061 | 0.025 | 0.41 | ENSMUSG00000000290 | protein_coding | Itgb2 | integrin beta 2 | 16414 | 10 | 77530252 | 77565708 | 1 | Mm.1137 |
| -3.82 | -1.06 | -0.69 | 0.78 | DN1ab | DN1cd | 2.27 | 5.85 | 0.00061 | 0.025 | 0.1 | ENSMUSG00000041460 | protein_coding | Cacna2d4 | calcium channel, voltage-dependent, alpha 2/delta subunit 4 | 319734 | 6 | 119236526 | 119352407 | 1 | Mm.440773 |
| -0.34 | 2.15 | -2.27 | 1.91 | DN1ab | DN1cd | 3.01 | 5.85 | 0.00061 | 0.025 | 0.13 | ENSMUSG00000048442 | protein_coding | Smim5 | small integral membrane protein 5 | 66528 | 11 | 115899966 | 115906269 | 1 | Mm.275740 |
| -2.91 | -0.44 | -3.49 | -0.71 | DN1ab | DN1cd | 2.58 | 5.85 | 0.00061 | 0.025 | 0.77 | ENSMUSG00000101959 | protein_coding | Ldhal6b | lactate dehydrogenase A-like 6B | 106557 | 17 | 5417323 | 5418767 | -1 | Mm.432764 |
| 6.39 | 6.03 | 6.47 | 6.21 | DN1ab | DN1cd | -0.33 | -5.85 | 0.00062 | 0.025 | 0.41 | ENSMUSG00000005609 | protein_coding | Ctr9 | CTR9 homolog, Paf1/RNA polymerase II complex component | 22083 | 7 | 111028951 | 111056377 | 1 | Mm.255858 |
| 2.58 | 1.79 | 3.61 | 2.42 | DN1ab | DN1cd | -0.97 | -5.85 | 0.00062 | 0.025 | 0.26 | ENSMUSG00000056498 | protein_coding | Tmem154 | transmembrane protein 154 | 320782 | 3 | 84666192 | 84704575 | 1 | Mm.107370 |
| -3.82 | -1.54 | -3.94 | -1.83 | DN1ab | DN1cd | 2.22 | 5.85 | 0.00062 | 0.025 | 0.85 | ENSMUSG00000071691 | protein_coding | Gm960 | predicted gene 960 | 381196 | 19 | 4625734 | 4698668 | -1 | Mm.297789 |
| 10.55 | 9.6 | 10.51 | 9.87 | DN1ab | DN1cd | -0.83 | -5.85 | 0.00062 | 0.025 | 0.32 | ENSMUSG00000017493 | protein_coding | Igfbp4 | insulin-like growth factor binding protein 4 | 16010 | 11 | 99041244 | 99054392 | 1 | Mm.233799 |
| 2.85 | 1.91 | 3.59 | 2.04 | DN1ab | DN1cd | -1.2 | -5.84 | 0.00062 | 0.025 | 0.14 | ENSMUSG00000030108 | protein_coding | Slc6a13 | solute carrier family 6 (neurotransmitter transporter, GABA), member 13 | 14412 | 6 | 121300227 | 121337733 | 1 | Mm.258596 |
| 5.68 | 6.3 | 5.89 | 6.51 | DN1ab | DN1cd | 0.63 | 5.82 | 0.00064 | 0.026 | 0.98 | ENSMUSG00000034765 | protein_coding | Dusp5 | dual specificity phosphatase 5 | 240672 | 19 | 53529109 | 53542431 | 1 | Mm.482424 |
| 2.02 | 1.22 | 2.47 | 1.56 | DN1ab | DN1cd | -0.85 | -5.82 | 0.00064 | 0.026 | 0.72 | ENSMUSG00000042271 | protein_coding | Nxt2 | nuclear transport factor 2-like export factor 2 | 237082 | X | 142226770 | 142239692 | 1 | Mm.41899 |
| 6.29 | 6.71 | 6.3 | 6.69 | DN1ab | DN1cd | 0.41 | 5.81 | 0.00064 | 0.026 | 0.89 | ENSMUSG00000030168 | protein_coding | Adipor2 | adiponectin receptor 2 | 68465 | 6 | 119353150 | 119417704 | -1 | Mm.291826 |
| 2.17 | 1.38 | 2.51 | 1.14 | DN1ab | DN1cd | -1.02 | -5.81 | 0.00064 | 0.026 | 0.11 | ENSMUSG00000018001 | protein_coding | Cyth3 | cytohesin 3 | 19159 | 5 | 143622447 | 143710250 | 1 | Mm.281003 |
| 7.93 | 7.64 | 7.84 | 7.56 | DN1ab | DN1cd | -0.29 | -5.81 | 0.00064 | 0.026 | 0.95 | ENSMUSG00000027304 | protein_coding | Rtf1 | RTF1, Paf1/RNA polymerase II complex component | 76246 | 2 | 119675068 | 119735407 | 1 | Mm.33437 |
| 2.21 | 1.24 | 2.94 | 1.52 | DN1ab | DN1cd | -1.16 | -5.81 | 0.00064 | 0.026 | 0.29 | ENSMUSG00000071648 | protein_coding | Rom1 | rod outer segment membrane protein 1 | 19881 | 19 | 8927382 | 8929356 | -1 | Mm.426094 |
| 7.07 | 6.31 | 7.05 | 6.29 | DN1ab | DN1cd | -0.76 | -5.8 | 0.00065 | 0.026 | 1 | ENSMUSG00000013089 | protein_coding | Etv5 | ets variant 5 | 104156 | 16 | 22381309 | 22439719 | -1 | Mm.155708 |
| -0.82 | 3.07 | -1 | 1.76 | DN1ab | DN1cd | 3.56 | 5.8 | 0.00065 | 0.026 | 0.44 | ENSMUSG00000042873 | protein_coding | Lhfpl4 | lipoma HMGIC fusion partner-like protein 4 | 269788 | 6 | 113168090 | 113195384 | -1 | Mm.37943 |
| 1.89 | 1.09 | 3.42 | 2.22 | DN1ab | DN1cd | -0.99 | -5.8 | 0.00065 | 0.026 | 0.28 | ENSMUSG00000027309 | protein_coding | 4930402H24Rik | RIKEN cDNA 4930402H24 gene | 228602 | 2 | 130706200 | 130906406 | -1 | Mm.271990 Mm.392671 |
| 3.76 | 2.62 | 4.56 | 3.02 | DN1ab | DN1cd | -1.32 | -5.79 | 0.00065 | 0.026 | 0.42 | ENSMUSG00000041308 | protein_coding | Sntb2 | syntrophin, basic 2 | 20650 | 8 | 106935750 | 107019714 | 1 | Mm.30228 |
| -3.82 | -2.28 | -3.94 | -1.46 | DN1ab | DN1cd | 1.89 | 5.79 | 0.00065 | 0.026 | 0.18 | ENSMUSG00000090219 | protein_coding | 4930516K23Rik | RIKEN cDNA 4930516K23 gene | 259081 | 7 | 104053600 | 104061445 | -1 | Mm.347669 Mm.466608 |
| 6.27 | 5.61 | 7.26 | 6.08 | DN1ab | DN1cd | -0.86 | -5.78 | 0.00066 | 0.026 | 0.088 | ENSMUSG00000017466 | protein_coding | Timp2 | tissue inhibitor of metalloproteinase 2 | 21858 | 11 | 118301069 | 118355740 | -1 | Mm.206505 |
| 5.55 | 5.01 | 5.56 | 5.1 | DN1ab | DN1cd | -0.51 | -5.77 | 0.00066 | 0.026 | 0.7 | ENSMUSG00000025332 | protein_coding | Kdm5c | lysine (K)-specific demethylase 5C | 20591 | X | 152233020 | 152274535 | 1 | Mm.142655 Mm.482277 |
| 5.46 | 6.26 | 4.78 | 6.28 | DN1ab | DN1cd | 1.04 | 5.76 | 0.00067 | 0.027 | 0.051 | ENSMUSG00000027010 | protein_coding | Slc25a12 | solute carrier family 25 (mitochondrial carrier, Aralar), member 12 | 78830 | 2 | 71271063 | 71367749 | -1 | Mm.30928 Mm.146696 |
| 2.69 | 3.82 | 2.4 | 3.85 | DN1ab | DN1cd | 1.24 | 5.76 | 0.00068 | 0.027 | 0.51 | ENSMUSG00000005802 | protein_coding | Slc30a4 | solute carrier family 30 (zinc transporter), member 4 | 22785 | 2 | 122681233 | 122702663 | -1 | Mm.27801 |
| 6.63 | 6.09 | 7.09 | 6.65 | DN1ab | DN1cd | -0.5 | -5.75 | 0.00068 | 0.027 | 0.63 | ENSMUSG00000045165 | protein_coding | AI467606 | expressed sequence AI467606 | 101602 | 7 | 127091359 | 127093986 | 1 | Mm.490357 |
| 6.46 | 5.93 | 7.2 | 6.29 | DN1ab | DN1cd | -0.68 | -5.75 | 0.00068 | 0.027 | 0.1 | ENSMUSG00000024480 | protein_coding | Ap3s1 | adaptor-related protein complex 3, sigma 1 subunit | 11777 | 18 | 46741876 | 46790826 | 1 | Mm.312536 |
| 2.27 | 1.45 | 1.78 | 0.56 | DN1ab | DN1cd | -0.97 | -5.75 | 0.00068 | 0.027 | 0.29 | ENSMUSG00000088990 | scaRNA | Gm22767 | predicted gene, 22767 | NA | 4 | 132838383 | 132838547 | -1 | |
| 3.04 | 3.9 | 2.49 | 3.42 | DN1ab | DN1cd | 0.88 | 5.74 | 0.00069 | 0.027 | 0.84 | ENSMUSG00000001521 | protein_coding | Tulp3 | tubby-like protein 3 | 22158 | 6 | 128321161 | 128355851 | -1 | Mm.291783 Mm.439490 |
| 9.26 | 9.58 | 9.31 | 9.58 | DN1ab | DN1cd | 0.3 | 5.74 | 0.00069 | 0.027 | 0.65 | ENSMUSG00000032097 | protein_coding | Ddx6 | DEAD (Asp-Glu-Ala-Asp) box polypeptide 6 | 13209 | 9 | 44604892 | 44640731 | 1 | Mm.267061 |
| 6.24 | 5.84 | 6.58 | 6.15 | DN1ab | DN1cd | -0.41 | -5.73 | 0.00069 | 0.027 | 0.84 | ENSMUSG00000028447 | protein_coding | Dctn3 | dynactin 3 | 53598 | 4 | 41714798 | 41723170 | -1 | Mm.29025 |
| 6.82 | 7.37 | 6.35 | 6.92 | DN1ab | DN1cd | 0.56 | 5.73 | 0.00069 | 0.027 | 0.92 | ENSMUSG00000046598 | protein_coding | Bdh1 | 3-hydroxybutyrate dehydrogenase, type 1 | 71911 | 16 | 31422280 | 31458901 | 1 | Mm.293470 |
| 7.36 | 7.71 | 6.71 | 7.37 | DN1ab | DN1cd | 0.46 | 5.73 | 7.00E-04 | 0.027 | 0.047 | ENSMUSG00000053398 | protein_coding | Phgdh | 3-phosphoglycerate dehydrogenase | 236539 | 3 | 98313170 | 98339990 | -1 | Mm.16898 Mm.371997 |
| 6.12 | 5.63 | 6.88 | 6.06 | DN1ab | DN1cd | -0.62 | -5.72 | 7.00E-04 | 0.027 | 0.14 | ENSMUSG00000026094 | protein_coding | Stk17b | serine/threonine kinase 17b (apoptosis-inducing) | 98267 | 1 | 53755506 | 53785224 | -1 | Mm.25559 |
| 6.28 | 6.69 | 6.16 | 6.95 | DN1ab | DN1cd | 0.54 | 5.72 | 7.00E-04 | 0.027 | 0.035 | ENSMUSG00000046668 | protein_coding | Cxxc5 | CXXC finger 5 | 67393 | 18 | 35829818 | 35861687 | 1 | Mm.94560 |
| -0.34 | 1.3 | 0.71 | 2.64 | DN1ab | DN1cd | 1.74 | 5.71 | 0.00071 | 0.027 | 0.68 | ENSMUSG00000073902 | unprocessed_pseudogene | Gm1966 | predicted gene 1966 | NA | 7 | 106596743 | 106604035 | -1 | |
| -3.82 | -1.1 | -3.94 | -2.44 | DN1ab | DN1cd | 2.29 | 5.7 | 0.00072 | 0.027 | 0.16 | ENSMUSG00000025017 | protein_coding | Pik3ap1 | phosphoinositide-3-kinase adaptor protein 1 | 83490 | 19 | 41274218 | 41385070 | -1 | Mm.222266 |
| 8.21 | 7.23 | 8.97 | 7.43 | DN1ab | DN1cd | -1.2 | -5.7 | 0.00072 | 0.028 | 0.21 | ENSMUSG00000015619 | protein_coding | Gata3 | GATA binding protein 3 | 14462 | 2 | 9857078 | 9890034 | -1 | Mm.313866 |
| 6.27 | 5.84 | 6.42 | 5.93 | DN1ab | DN1cd | -0.46 | -5.69 | 0.00072 | 0.028 | 0.76 | ENSMUSG00000053768 | protein_coding | Chchd3 | coiled-coil-helix-coiled-coil-helix domain containing 3 | 66075 | 6 | 32790976 | 33060260 | -1 | Mm.21501 |
| 5.99 | 6.4 | 5.75 | 6.31 | DN1ab | DN1cd | 0.46 | 5.69 | 0.00072 | 0.028 | 0.43 | ENSMUSG00000020516 | protein_coding | Rps6kb1 | ribosomal protein S6 kinase, polypeptide 1 | 72508 | 11 | 86498871 | 86544805 | -1 | Mm.394280 Mm.446624 Mm.479484 |
| 6.48 | 7.03 | 6.31 | 6.55 | DN1ab | DN1cd | 0.44 | 5.68 | 0.00073 | 0.028 | 0.046 | ENSMUSG00000020572 | protein_coding | Nampt | nicotinamide phosphoribosyltransferase | 59027 | 12 | 32819545 | 32853349 | 1 | Mm.202727 |
| 4.42 | 4.78 | 4.87 | 5.47 | DN1ab | DN1cd | 0.45 | 5.68 | 0.00073 | 0.028 | 0.16 | ENSMUSG00000017756 | protein_coding | Slc12a7 | solute carrier family 12, member 7 | 20499 | 13 | 73733094 | 73816754 | 1 | Mm.275800 |
| -1.29 | -3.86 | -1.93 | -4.02 | DN1ab | DN1cd | -2.39 | -5.67 | 0.00074 | 0.028 | 0.62 | ENSMUSG00000108068 | processed_pseudogene | Gm44075 | predicted gene, 44075 | NA | 6 | 63783852 | 63784067 | -1 | |
| -2.62 | 1.35 | -3.94 | 0.8 | DN1ab | DN1cd | 4.21 | 5.67 | 0.00074 | 0.028 | 0.67 | ENSMUSG00000099655 | protein_coding | 2310034G01Rik | RIKEN cDNA 2310034G01 gene | NA | 19 | 46348201 | 46350302 | -1 | |
| 7.08 | 7.75 | 6.61 | 7.77 | DN1ab | DN1cd | 0.84 | 5.67 | 0.00074 | 0.028 | 0.12 | ENSMUSG00000032946 | protein_coding | Rasgrp2 | RAS, guanyl releasing protein 2 | 19395 | 19 | 6399340 | 6415216 | 1 | Mm.77017 |
| 6.29 | 5.79 | 7.23 | 6.33 | DN1ab | DN1cd | -0.66 | -5.66 | 0.00075 | 0.028 | 0.092 | ENSMUSG00000041736 | protein_coding | Tspo | translocator protein | 12257 | 15 | 83563592 | 83574203 | 1 | Mm.1508 |
| 7.41 | 8.08 | 7.17 | 7.93 | DN1ab | DN1cd | 0.71 | 5.66 | 0.00075 | 0.028 | 0.77 | ENSMUSG00000105879 | processed_pseudogene | Gm6204 | predicted gene 6204 | NA | 3 | 53692422 | 53692878 | -1 | |
| 2.83 | 2.04 | 3.63 | 2.56 | DN1ab | DN1cd | -0.91 | -5.64 | 0.00076 | 0.029 | 0.44 | ENSMUSG00000104835 | lincRNA | Gm5547 | predicted gene 5547 | NA | 3 | 105815540 | 105817357 | 1 | |
| 8.67 | 8.47 | 8.69 | 8.37 | DN1ab | DN1cd | -0.25 | -5.64 | 0.00076 | 0.029 | 0.16 | ENSMUSG00000024758 | protein_coding | Rtn3 | reticulon 3 | 20168 | 19 | 7425901 | 7483289 | -1 | Mm.246990 |
| -3.14 | -0.43 | -2.36 | 0.4 | DN1ab | DN1cd | 2.73 | 5.63 | 0.00077 | 0.029 | 0.97 | ENSMUSG00000026399 | protein_coding | Cd55 | CD55 molecule, decay accelerating factor for complement | 13136 | 1 | 130439027 | 130462744 | -1 | Mm.101591 |
| 2.3 | 3.74 | 2.48 | 4.65 | DN1ab | DN1cd | 1.69 | 5.63 | 0.00077 | 0.029 | 0.28 | ENSMUSG00000026576 | protein_coding | Atp1b1 | ATPase, Na+/K+ transporting, beta 1 polypeptide | 11931 | 1 | 164437109 | 164458355 | -1 | Mm.4550 |
| 10.84 | 11.15 | 10.17 | 10.42 | DN1ab | DN1cd | 0.29 | 5.62 | 0.00078 | 0.029 | 0.61 | ENSMUSG00000057113 | protein_coding | Npm1 | nucleophosmin 1 | 18148 | 11 | 33152287 | 33163206 | -1 | Mm.485384 |
| 6.03 | 5.56 | 6.66 | 5.83 | DN1ab | DN1cd | -0.61 | -5.62 | 0.00078 | 0.029 | 0.1 | ENSMUSG00000014444 | protein_coding | Piezo1 | piezo-type mechanosensitive ion channel component 1 | 234839 | 8 | 122481698 | 122551329 | -1 | Mm.37324 Mm.482288 |
| 2.04 | 3.12 | 2.14 | 3.17 | DN1ab | DN1cd | 1.06 | 5.61 | 0.00078 | 0.029 | 0.92 | ENSMUSG00000030123 | protein_coding | Plxnd1 | plexin D1 | 67784 | 6 | 115954811 | 115995005 | -1 | Mm.3085 |
| 5.1 | 5.68 | 4.86 | 5.33 | DN1ab | DN1cd | 0.55 | 5.61 | 0.00078 | 0.029 | 0.6 | ENSMUSG00000066640 | protein_coding | Fbxl18 | F-box and leucine-rich repeat protein 18 | 231863 | 5 | 142866946 | 142895421 | -1 | Mm.89753 Mm.361645 Mm.454456 |
| -1.14 | 1 | 0 | 3.37 | DN1ab | DN1cd | 2.57 | 5.61 | 0.00079 | 0.029 | 0.22 | ENSMUSG00000008193 | protein_coding | Spib | Spi-B transcription factor (Spi-1/PU.1 related) | 272382 | 7 | 44525993 | 44532071 | -1 | Mm.8012 |
| 5.58 | 5.18 | 5.76 | 5.28 | DN1ab | DN1cd | -0.43 | -5.6 | 0.00079 | 0.029 | 0.63 | ENSMUSG00000031988 | protein_coding | Vps26b | VPS26 retromer complex component B | 69091 | 9 | 27007771 | 27030065 | -1 | Mm.218717 Mm.440390 |
| -1.54 | -3.86 | -2.36 | -4.02 | DN1ab | DN1cd | -2.08 | -5.6 | 8.00E-04 | 0.029 | 0.43 | ENSMUSG00000026452 | protein_coding | Syt2 | synaptotagmin II | 20980 | 1 | 134646677 | 134762593 | 1 | Mm.5102 |
| 0.19 | -1.53 | 2.38 | -0.13 | DN1ab | DN1cd | -2.09 | -5.6 | 8.00E-04 | 0.029 | 0.32 | ENSMUSG00000022817 | protein_coding | Itgb5 | integrin beta 5 | 16419 | 16 | 33829665 | 33949338 | 1 | Mm.6424 |
| 2.58 | 1.54 | 2.7 | 2.15 | DN1ab | DN1cd | -0.82 | -5.58 | 0.00081 | 0.029 | 0.11 | ENSMUSG00000034118 | protein_coding | Tpst1 | protein-tyrosine sulfotransferase 1 | 22021 | 5 | 130073326 | 130135729 | 1 | Mm.332720 |
| 4.64 | 5.07 | 4.19 | 4.82 | DN1ab | DN1cd | 0.5 | 5.58 | 0.00082 | 0.03 | 0.32 | ENSMUSG00000035310 | protein_coding | Lin54 | lin-54 homolog (C. elegans) | 231506 | 5 | 100441918 | 100500639 | -1 | Mm.212568 |
| 8.42 | 8.78 | 7.89 | 8.17 | DN1ab | DN1cd | 0.33 | 5.57 | 0.00082 | 0.03 | 0.57 | ENSMUSG00000022752 | protein_coding | Tomm70a | translocase of outer mitochondrial membrane 70A | 28185 | 16 | 57121703 | 57156705 | 1 | Mm.196933 Mm.213292 |
| 0.8 | 2.31 | 1.27 | 2.3 | DN1ab | DN1cd | 1.35 | 5.57 | 0.00082 | 0.03 | 0.39 | ENSMUSG00000036718 | protein_coding | Micall2 | MICAL-like 2 | 231830 | 5 | 139706696 | 139736336 | -1 | Mm.240510 |
| -0.53 | -3.12 | 0.23 | -1.46 | DN1ab | DN1cd | -2.24 | -5.57 | 0.00082 | 0.03 | 0.3 | ENSMUSG00000019876 | protein_coding | Pkib | protein kinase inhibitor beta, cAMP dependent, testis specific | 18768 | 10 | 57631981 | 57741112 | 1 | Mm.262135 |
| 6.97 | 7.78 | 6.26 | 6.93 | DN1ab | DN1cd | 0.76 | 5.56 | 0.00083 | 0.03 | 0.65 | ENSMUSG00000055296 | protein_coding | Tmem245 | transmembrane protein 245 | 242474 | 4 | 56866923 | 56947437 | -1 | Mm.296632 Mm.486370 |
| 0.19 | -1.88 | 2.32 | -1.01 | DN1ab | DN1cd | -2.63 | -5.56 | 0.00083 | 0.03 | 0.21 | ENSMUSG00000026018 | protein_coding | Ica1l | islet cell autoantigen 1-like | 70375 | 1 | 59982490 | 60043184 | -1 | Mm.17834 |
| -1.81 | 0.52 | -1.47 | 1.26 | DN1ab | DN1cd | 2.46 | 5.56 | 0.00083 | 0.03 | 0.71 | ENSMUSG00000086527 | antisense | Gm15856 | predicted gene 15856 | NA | 6 | 121223692 | 121226153 | -1 | |
| 2.87 | 3.75 | 1.48 | 2.46 | DN1ab | DN1cd | 0.91 | 5.55 | 0.00084 | 0.03 | 0.82 | ENSMUSG00000097467 | antisense | Gm26737 | predicted gene, 26737 | NA | 9 | 44112923 | 44121960 | 1 | |
| -3.82 | -1.54 | -3.94 | 0.62 | DN1ab | DN1cd | 3.18 | 5.55 | 0.00084 | 0.03 | 0.036 | ENSMUSG00000100277 | processed_transcript | 1810053B23Rik | RIKEN cDNA 1810053B23 gene | NA | 16 | 93342504 | 93366668 | -1 | |
| 1.13 | -0.77 | 0.85 | -2.07 | DN1ab | DN1cd | -2.32 | -5.55 | 0.00084 | 0.03 | 0.25 | ENSMUSG00000059031 | protein_coding | Olfr482 | olfactory receptor 482 | 258728 | 7 | 108092999 | 108097586 | -1 | Mm.349555 |
| 6.92 | 6.52 | 6.66 | 6.38 | DN1ab | DN1cd | -0.36 | -5.55 | 0.00084 | 0.03 | 0.42 | ENSMUSG00000032381 | protein_coding | Ciao2a | cytosolic iron-sulfur assembly component 2A | 68250 | 9 | 66126611 | 66138955 | 1 | Mm.27107 |
| 6.5 | 5.94 | 7.13 | 6.1 | DN1ab | DN1cd | -0.74 | -5.54 | 0.00084 | 0.03 | 0.092 | ENSMUSG00000049470 | protein_coding | Aff4 | AF4/FMR2 family, member 4 | 93736 | 11 | 53350833 | 53421830 | 1 | Mm.395281 |
| 7.91 | 8.2 | 7.86 | 8.48 | DN1ab | DN1cd | 0.41 | 5.54 | 0.00085 | 0.03 | 0.017 | ENSMUSG00000052397 | protein_coding | Ezr | ezrin | 22350 | 17 | 6738041 | 6782784 | -1 | Mm.277812 |
| -1.44 | 0.72 | -3.94 | -0.85 | DN1ab | DN1cd | 2.44 | 5.54 | 0.00085 | 0.03 | 0.36 | ENSMUSG00000104606 | lincRNA | Gm43409 | predicted gene 43409 | NA | 5 | 123188701 | 123191588 | -1 | |
| -3.82 | -2.28 | -2.81 | -0.56 | DN1ab | DN1cd | 1.81 | 5.54 | 0.00085 | 0.03 | 0.33 | ENSMUSG00000078490 | protein_coding | Cfap74 | cilia and flagella associated protein 74 | 544678 | 4 | 155409190 | 155466823 | 1 | Mm.472798 |
| -1.71 | 0.5 | -2.36 | 0.86 | DN1ab | DN1cd | 2.55 | 5.53 | 0.00085 | 0.03 | 0.34 | ENSMUSG00000043953 | protein_coding | Ccrl2 | chemokine (C-C motif) receptor-like 2 | 54199 | 9 | 111054486 | 111057519 | -1 | Mm.7336 |
| -2.24 | -3.86 | -1.22 | -4.02 | DN1ab | DN1cd | -2.05 | -5.52 | 0.00086 | 0.03 | 0.13 | ENSMUSG00000065299 | snRNA | Gm23138 | predicted gene, 23138 | NA | 4 | 108498044 | 108498146 | 1 | |
| 6.22 | 6.74 | 5.94 | 6.18 | DN1ab | DN1cd | 0.42 | 5.52 | 0.00087 | 0.03 | 0.072 | ENSMUSG00000105987 | processed_transcript | AI506816 | expressed sequence AI506816 | NA | 5 | 23698296 | 23712667 | -1 | |
| 2.76 | 3.84 | 0.83 | 2.83 | DN1ab | DN1cd | 1.38 | 5.51 | 0.00088 | 0.03 | 0.081 | ENSMUSG00000097839 | lincRNA | Gm26909 | predicted gene, 26909 | NA | 8 | 22644861 | 22646286 | 1 | |
| 10.28 | 9.54 | 11.09 | 10.67 | DN1ab | DN1cd | -0.61 | -5.51 | 0.00088 | 0.03 | 0.17 | ENSMUSG00000092341 | lincRNA | Malat1 | metastasis associated lung adenocarcinoma transcript 1 (non-coding RNA) | NA | 19 | 5795690 | 5802672 | -1 | |
| 2.57 | 3.42 | 2.67 | 4.49 | DN1ab | DN1cd | 1.21 | 5.5 | 0.00088 | 0.03 | 0.017 | ENSMUSG00000045441 | protein_coding | Gprin3 | GPRIN family member 3 | 243385 | 6 | 59347226 | 59426294 | -1 | Mm.138080 Mm.485751 |
| 0.27 | 1.91 | -0.13 | 2.39 | DN1ab | DN1cd | 1.93 | 5.49 | 0.00089 | 0.03 | 0.27 | ENSMUSG00000068417 | protein_coding | Pnp2 | purine-nucleoside phosphorylase 2 | 667034 | 14 | 50955992 | 50964749 | 1 | Mm.482171 |
| 6.63 | 7.5 | 6.87 | 7.36 | DN1ab | DN1cd | 0.74 | 5.49 | 9.00E-04 | 0.03 | 0.19 | ENSMUSG00000068129 | protein_coding | Cst7 | cystatin F (leukocystatin) | 13011 | 2 | 150570415 | 150578944 | 1 | Mm.12965 |
| 3.72 | 2.67 | 5.04 | 4.41 | DN1ab | DN1cd | -0.85 | -5.49 | 9.00E-04 | 0.03 | 0.19 | ENSMUSG00000030246 | protein_coding | Ldhb | lactate dehydrogenase B | 16832 | 6 | 142490249 | 142507957 | -1 | Mm.9745 |
| 7.61 | 7.84 | 7.19 | 7.51 | DN1ab | DN1cd | 0.27 | 5.49 | 9.00E-04 | 0.03 | 0.41 | ENSMUSG00000052738 | protein_coding | Suclg1 | succinate-CoA ligase, GDP-forming, alpha subunit | 56451 | 6 | 73248382 | 73276911 | 1 | Mm.29845 |
| 0.36 | -3.86 | 3.69 | -2.61 | DN1ab | DN1cd | -5.13 | -5.48 | 9.00E-04 | 0.03 | 0.3 | ENSMUSG00000063142 | protein_coding | Kcnma1 | potassium large conductance calcium-activated channel, subfamily M, alpha member 1 | 16531 | 14 | 23289431 | 24014491 | -1 | Mm.343607 Mm.486347 |
| 3.14 | 4.11 | 3.91 | 4.42 | DN1ab | DN1cd | 0.81 | 5.48 | 9.00E-04 | 0.03 | 0.14 | ENSMUSG00000069874 | protein_coding | Irgm2 | immunity-related GTPase family M member 2 | 54396 | 11 | 58199618 | 58222782 | 1 | Mm.458517 |
| 5.72 | 6.96 | 4.25 | 6.46 | DN1ab | DN1cd | 1.57 | 5.48 | 9.00E-04 | 0.03 | 0.11 | ENSMUSG00000060404 | protein_coding | Olfr1369-ps1 | olfactory receptor 1369, pseudogene 1 | NA | 13 | 21111483 | 21119095 | 1 | |
| 2.72 | -0.06 | 5.08 | 0.31 | DN1ab | DN1cd | -3.83 | -5.48 | 0.00091 | 0.031 | 0.17 | ENSMUSG00000104969 | sense_intronic | Gm43445 | predicted gene 43445 | NA | 3 | 144197172 | 144199262 | -1 | |
| 1.91 | 2.76 | 1.28 | 2.72 | DN1ab | DN1cd | 1.05 | 5.48 | 0.00091 | 0.031 | 0.15 | ENSMUSG00000061186 | protein_coding | Sfmbt2 | Scm-like with four mbt domains 2 | 353282 | 2 | 10370510 | 10595253 | 1 | Mm.329991 |
| 5.57 | 4.81 | 5.26 | 4.86 | DN1ab | DN1cd | -0.62 | -5.45 | 0.00094 | 0.031 | 0.11 | ENSMUSG00000081992 | processed_pseudogene | Gm13408 | predicted gene 13408 | NA | 2 | 34564999 | 34565796 | 1 | |
| 4.24 | 3.42 | 5.04 | 3.5 | DN1ab | DN1cd | -1.12 | -5.44 | 0.00094 | 0.031 | 0.077 | ENSMUSG00000039176 | protein_coding | Polg | polymerase (DNA directed), gamma | 18975 | 7 | 79446231 | 79466362 | -1 | Mm.3616 |
| 5.99 | 6.89 | 6.19 | 6.62 | DN1ab | DN1cd | 0.73 | 5.43 | 0.00095 | 0.032 | 0.089 | ENSMUSG00000055817 | protein_coding | Mta3 | metastasis associated 3 | 116871 | 17 | 83706163 | 83821516 | 1 | Mm.277668 Mm.443827 |
| 1.09 | 2.3 | 1.27 | 3.78 | DN1ab | DN1cd | 1.67 | 5.43 | 0.00096 | 0.032 | 0.024 | ENSMUSG00000021779 | protein_coding | Thrb | thyroid hormone receptor beta | 21834 | 14 | 17660261 | 18038090 | 1 | Mm.32563 Mm.491616 |
| 6.29 | 6.63 | 6.25 | 6.69 | DN1ab | DN1cd | 0.37 | 5.42 | 0.00096 | 0.032 | 0.5 | ENSMUSG00000081431 | processed_pseudogene | Gm15483 | predicted gene 15483 | NA | 7 | 133738479 | 133738934 | -1 | |
| 3.12 | 1.12 | 4.61 | 3.53 | DN1ab | DN1cd | -1.47 | -5.42 | 0.00096 | 0.032 | 0.086 | ENSMUSG00000076755 | TR_V_gene | Tcrg-V1 | T cell receptor gamma, variable 1 | NA | 13 | 19339989 | 19340454 | 1 | |
| 3.12 | -0.36 | 4.27 | 2.38 | DN1ab | DN1cd | -2.42 | -5.42 | 0.00097 | 0.032 | 0.088 | ENSMUSG00000020183 | protein_coding | Cpm | carboxypeptidase M | 70574 | 10 | 117629500 | 117687352 | 1 | Mm.339332 |
| 5.73 | 5.33 | 5.6 | 5.34 | DN1ab | DN1cd | -0.34 | -5.42 | 0.00097 | 0.032 | 0.35 | ENSMUSG00000026187 | protein_coding | Xrcc5 | X-ray repair complementing defective repair in Chinese hamster cells 5 | 22596 | 1 | 72307421 | 72394952 | 1 | Mm.246952 |
| 7.05 | 7.41 | 7.22 | 7.77 | DN1ab | DN1cd | 0.43 | 5.41 | 0.00098 | 0.032 | 0.29 | ENSMUSG00000027472 | protein_coding | Pdrg1 | p53 and DNA damage regulated 1 | 68559 | 2 | 153008890 | 153015427 | -1 | Mm.12746 |
| 6.4 | 6.87 | 5.85 | 6.07 | DN1ab | DN1cd | 0.39 | 5.39 | 0.00099 | 0.032 | 0.084 | ENSMUSG00000034203 | protein_coding | Chchd4 | coiled-coil-helix-coiled-coil-helix domain containing 4 | 72170 | 6 | 91462172 | 91473546 | -1 | Mm.41622 |
| 10.51 | 10.78 | 10.42 | 10.99 | DN1ab | DN1cd | 0.38 | 5.39 | 0.001 | 0.032 | 0.02 | ENSMUSG00000064363 | protein_coding | mt-Nd4 | mitochondrially encoded NADH dehydrogenase 4 | 17719 | MT | 10167 | 11544 | 1 | |
| 5.41 | 5.88 | 5.07 | 5.92 | DN1ab | DN1cd | 0.61 | 5.38 | 0.001 | 0.033 | 0.1 | ENSMUSG00000022971 | protein_coding | Ifnar2 | interferon (alpha and beta) receptor 2 | 15976 | 16 | 91372783 | 91405589 | 1 | Mm.6834 |
| 2.89 | 0.85 | 4.35 | 3.32 | DN1ab | DN1cd | -1.45 | -5.37 | 0.001 | 0.033 | 0.053 | ENSMUSG00000024334 | protein_coding | H2-Oa | histocompatibility 2, O region alpha locus | 15001 | 17 | 34092340 | 34095234 | 1 | Mm.116 |
| 9.41 | 9.84 | 9.02 | 9.57 | DN1ab | DN1cd | 0.47 | 5.37 | 0.001 | 0.033 | 0.55 | ENSMUSG00000052833 | protein_coding | Sae1 | SUMO1 activating enzyme subunit 1 | 56459 | 7 | 16320234 | 16387806 | -1 | Mm.258530 |
| -0.85 | 0.83 | -3.94 | -1.03 | DN1ab | DN1cd | 2.04 | 5.36 | 0.001 | 0.033 | 0.15 | ENSMUSG00000106536 | TEC | Gm42508 | predicted gene 42508 | NA | 3 | 97262355 | 97265646 | 1 | |
| 7.13 | 6.75 | 6.95 | 6.7 | DN1ab | DN1cd | -0.33 | -5.35 | 0.001 | 0.033 | 0.32 | ENSMUSG00000030091 | protein_coding | Nup210 | nucleoporin 210 | 54563 | 6 | 91013068 | 91116829 | -1 | Mm.28162 |
| 2.71 | 1.32 | 3.92 | 1.5 | DN1ab | DN1cd | -1.85 | -5.35 | 0.001 | 0.033 | 0.14 | ENSMUSG00000021356 | protein_coding | Irf4 | interferon regulatory factor 4 | 16364 | 13 | 30749226 | 30766976 | 1 | Mm.4677 |
| 0.45 | 1.99 | 0.57 | 2.84 | DN1ab | DN1cd | 1.78 | 5.35 | 0.001 | 0.033 | 0.33 | ENSMUSG00000034265 | protein_coding | Zdhhc14 | zinc finger, DHHC domain containing 14 | 224454 | 17 | 5492557 | 5753811 | 1 | Mm.487663 Mm.491143 |
| 6.11 | 8.07 | 5 | 7.82 | DN1ab | DN1cd | 2.25 | 5.35 | 0.001 | 0.033 | 0.37 | ENSMUSG00000073940 | protein_coding | Hbb-bt | hemoglobin, beta adult t chain | 101488143 | 7 | 103812524 | 103813996 | -1 | Mm.467412 |
| 3.76 | 4.2 | 3.31 | 3.98 | DN1ab | DN1cd | 0.52 | 5.34 | 0.001 | 0.033 | 0.29 | ENSMUSG00000007216 | protein_coding | Zfp775 | zinc finger protein 775 | 243372 | 6 | 48602063 | 48623227 | 1 | Mm.435190 |
| 4.86 | 4.32 | 5.28 | 4.3 | DN1ab | DN1cd | -0.71 | -5.34 | 0.0011 | 0.033 | 0.11 | ENSMUSG00000052609 | protein_coding | Plekhg3 | pleckstrin homology domain containing, family G (with RhoGef domain) member 3 | 263406 | 12 | 76530891 | 76580488 | 1 | Mm.291854 |
| 0.53 | 2.24 | -0.63 | 2.52 | DN1ab | DN1cd | 2.18 | 5.34 | 0.0011 | 0.033 | 0.094 | ENSMUSG00000046546 | protein_coding | Fam43a | family with sequence similarity 43, member A | 224093 | 16 | 30599723 | 30602797 | 1 | Mm.28244 |
| 5.66 | 6.03 | 5.41 | 5.95 | DN1ab | DN1cd | 0.43 | 5.33 | 0.0011 | 0.033 | 0.34 | ENSMUSG00000021891 | protein_coding | Mettl6 | methyltransferase like 6 | 67011 | 14 | 31473578 | 31495040 | -1 | Mm.291731 |
| 5.07 | 5.42 | 4.96 | 5.41 | DN1ab | DN1cd | 0.38 | 5.33 | 0.0011 | 0.033 | 0.54 | ENSMUSG00000021371 | protein_coding | Mcur1 | mitochondrial calcium uniporter regulator 1 | 76137 | 13 | 43538393 | 43560191 | -1 | Mm.426290 |
| 7.97 | 7.21 | 8.3 | 7.85 | DN1ab | DN1cd | -0.64 | -5.33 | 0.0011 | 0.033 | 0.22 | ENSMUSG00000060131 | protein_coding | Atp8b4 | ATPase, class I, type 8B, member 4 | 241633 | 2 | 126320973 | 126500674 | -1 | Mm.222235 |
| 7.8 | 8.31 | 7.51 | 8.52 | DN1ab | DN1cd | 0.69 | 5.33 | 0.0011 | 0.033 | 0.047 | ENSMUSG00000061887 | protein_coding | Ssbp3 | single-stranded DNA binding protein 3 | 72475 | 4 | 106910701 | 107049694 | 1 | Mm.195635 Mm.445960 Mm.474861 |
| 6.54 | 6.83 | 5.99 | 6.4 | DN1ab | DN1cd | 0.33 | 5.33 | 0.0011 | 0.033 | 0.35 | ENSMUSG00000040331 | protein_coding | Nsmce4a | NSE4 homolog A, SMC5-SMC6 complex component | 67872 | 7 | 130532526 | 130573118 | -1 | Mm.21714 Mm.411879 |
| 4.66 | 3.92 | 4.88 | 3.36 | DN1ab | DN1cd | -1.04 | -5.33 | 0.0011 | 0.033 | 0.037 | ENSMUSG00000025289 | protein_coding | Prdx4 | peroxiredoxin 4 | 53381 | X | 155323918 | 155340754 | -1 | Mm.247542 |
| 7.23 | 6.74 | 7.55 | 7.33 | DN1ab | DN1cd | -0.39 | -5.32 | 0.0011 | 0.033 | 0.044 | ENSMUSG00000040430 | protein_coding | Pitpnc1 | phosphatidylinositol transfer protein, cytoplasmic 1 | 71795 | 11 | 107207892 | 107470699 | -1 | Mm.397413 Mm.476186 |
| 1.92 | 3.36 | 3.08 | 5.28 | DN1ab | DN1cd | 1.72 | 5.32 | 0.0011 | 0.033 | 0.28 | ENSMUSG00000078606 | protein_coding | Gm4070 | predicted gene 4070 | 100042856 | 7 | 105895139 | 105953967 | -1 | Mm.425473 Mm.458429 |
| -2.91 | -0.16 | -3.94 | 0.68 | DN1ab | DN1cd | 3.41 | 5.32 | 0.0011 | 0.033 | 0.18 | ENSMUSG00000005883 | protein_coding | Spo11 | SPO11 meiotic protein covalently bound to DSB | 26972 | 2 | 172977700 | 172993576 | 1 | Mm.23495 |
| 4.78 | 4.34 | 5.56 | 4.85 | DN1ab | DN1cd | -0.55 | -5.32 | 0.0011 | 0.033 | 0.22 | ENSMUSG00000020272 | protein_coding | Stk10 | serine/threonine kinase 10 | 20868 | 11 | 32533305 | 32624587 | 1 | Mm.8235 |
| 5.87 | 6.35 | 5.2 | 6.06 | DN1ab | DN1cd | 0.61 | 5.31 | 0.0011 | 0.033 | 0.11 | ENSMUSG00000019790 | protein_coding | Stxbp5 | syntaxin binding protein 5 (tomosyn) | 78808 | 10 | 9755547 | 9901079 | -1 | Mm.331751 |
| 0.31 | -3.12 | 4.35 | -3.22 | DN1ab | DN1cd | -5.31 | -5.31 | 0.0011 | 0.033 | 0.022 | ENSMUSG00000015843 | protein_coding | Rxrg | retinoid X receptor gamma | 20183 | 1 | 167598384 | 167639623 | 1 | Mm.3475 |
| 6.03 | 5.69 | 6.69 | 6.04 | DN1ab | DN1cd | -0.45 | -5.31 | 0.0011 | 0.033 | 0.061 | ENSMUSG00000001794 | protein_coding | Capns1 | calpain, small subunit 1 | 12336 | 7 | 30186936 | 30198811 | -1 | Mm.439782 |
| -1.5 | -3.86 | -1.22 | -2.44 | DN1ab | DN1cd | -1.95 | -5.3 | 0.0011 | 0.034 | 0.14 | ENSMUSG00000082576 | processed_pseudogene | Gm12058 | predicted gene 12058 | NA | 11 | 22786811 | 22787247 | -1 | |
| -3.14 | 0.38 | -2.96 | -0.71 | DN1ab | DN1cd | 3.12 | 5.3 | 0.0011 | 0.034 | 0.35 | ENSMUSG00000108365 | TEC | Gm44951 | predicted gene 44951 | NA | 7 | 80036705 | 80038728 | 1 | |
| 3.62 | 2.95 | 4.92 | 3.54 | DN1ab | DN1cd | -0.97 | -5.29 | 0.0011 | 0.034 | 0.041 | ENSMUSG00000036769 | protein_coding | Wdr44 | WD repeat domain 44 | 72404 | X | 23693051 | 23806025 | 1 | Mm.423 |
| -3.14 | 0.38 | -1.94 | -0.03 | DN1ab | DN1cd | 3 | 5.29 | 0.0011 | 0.034 | 0.2 | ENSMUSG00000042265 | protein_coding | Trem1 | triggering receptor expressed on myeloid cells 1 | 58217 | 17 | 48232768 | 48246924 | 1 | Mm.248352 |
| 6.08 | 5.7 | 6.15 | 5.91 | DN1ab | DN1cd | -0.32 | -5.28 | 0.0011 | 0.034 | 0.3 | ENSMUSG00000068921 | protein_coding | Dap3 | death associated protein 3 | 65111 | 3 | 88920803 | 88951181 | -1 | Mm.29028 Mm.400397 |
| 12.62 | 12.89 | 12.27 | 12.4 | DN1ab | DN1cd | 0.22 | 5.26 | 0.0011 | 0.034 | 0.15 | ENSMUSG00000023944 | protein_coding | Hsp90ab1 | heat shock protein 90 alpha (cytosolic), class B member 1 | 15516 | 17 | 45567775 | 45573271 | -1 | Mm.2180 |
| 4.03 | 4.49 | 4.12 | 4.77 | DN1ab | DN1cd | 0.53 | 5.26 | 0.0011 | 0.034 | 0.42 | ENSMUSG00000032571 | protein_coding | Pik3r4 | phosphoinositide-3-kinase regulatory subunit 4 | 75669 | 9 | 105642978 | 105687657 | 1 | Mm.274830 |
| 4.29 | 5.5 | 2.28 | 4.52 | DN1ab | DN1cd | 1.54 | 5.26 | 0.0012 | 0.034 | 0.1 | ENSMUSG00000000861 | protein_coding | Bcl11a | B cell CLL/lymphoma 11A (zinc finger protein) | 14025 | 11 | 24078056 | 24174123 | 1 | Mm.53687 |
| 3.79 | 4.27 | 3.11 | 4.03 | DN1ab | DN1cd | 0.64 | 5.25 | 0.0012 | 0.034 | 0.075 | ENSMUSG00000027654 | protein_coding | Fam83d | family with sequence similarity 83, member D | 71878 | 2 | 158768093 | 158786637 | 1 | Mm.85162 Mm.396400 |
| 7.48 | 6.58 | 7.67 | 7.28 | DN1ab | DN1cd | -0.69 | -5.25 | 0.0012 | 0.034 | 0.044 | ENSMUSG00000026034 | protein_coding | Clk1 | CDC-like kinase 1 | 12747 | 1 | 58410189 | 58424066 | -1 | Mm.1761 |
| 5.43 | 6.21 | 5.63 | 6.97 | DN1ab | DN1cd | 0.98 | 5.24 | 0.0012 | 0.034 | 0.17 | ENSMUSG00000024617 | protein_coding | Camk2a | calcium/calmodulin-dependent protein kinase II alpha | 12322 | 18 | 60925618 | 60988152 | 1 | Mm.131530 Mm.446530 |
| 3.75 | 5.83 | 1.81 | 5.28 | DN1ab | DN1cd | 2.53 | 5.23 | 0.0012 | 0.035 | 0.2 | ENSMUSG00000031074 | protein_coding | Fgf3 | fibroblast growth factor 3 | 14174 | 7 | 144838083 | 144844436 | 1 | Mm.4947 |
| 4.58 | 4.27 | 4.98 | 4.51 | DN1ab | DN1cd | -0.38 | -5.22 | 0.0012 | 0.035 | 0.33 | ENSMUSG00000024782 | protein_coding | Ak3 | adenylate kinase 3 | 56248 | 19 | 29020833 | 29047961 | -1 | Mm.196067 |
| 1.15 | -0.44 | 1.64 | 0.84 | DN1ab | DN1cd | -1.21 | -5.22 | 0.0012 | 0.035 | 0.088 | ENSMUSG00000033644 | protein_coding | Piwil2 | piwi-like RNA-mediated gene silencing 2 | 57746 | 14 | 70372477 | 70429383 | -1 | Mm.85253 |
| -0.42 | 1.11 | -1.83 | 0.36 | DN1ab | DN1cd | 1.74 | 5.22 | 0.0012 | 0.035 | 0.39 | ENSMUSG00000052125 | bidirectional_promoter_lncRNA | F730043M19Rik | RIKEN cDNA F730043M19 gene | NA | 12 | 33111711 | 33147597 | -1 | |
| -0.49 | -3.12 | 1.17 | -4.02 | DN1ab | DN1cd | -3.67 | -5.22 | 0.0012 | 0.035 | 0.065 | ENSMUSG00000043969 | protein_coding | Emx2 | empty spiracles homeobox 2 | 13797 | 19 | 59458372 | 59465357 | 1 | Mm.245394 |
| 3.56 | 4.06 | 3.05 | 3.77 | DN1ab | DN1cd | 0.58 | 5.21 | 0.0012 | 0.035 | 0.38 | ENSMUSG00000061119 | protein_coding | Prcp | prolylcarboxypeptidase (angiotensinase C) | 72461 | 7 | 92874470 | 92934583 | 1 | Mm.389969 |
| 2.49 | 3.33 | 2.01 | 3.52 | DN1ab | DN1cd | 1.08 | 5.2 | 0.0012 | 0.035 | 0.13 | ENSMUSG00000000552 | protein_coding | Zfp385a | zinc finger protein 385A | 29813 | 15 | 103313895 | 103340093 | -1 | Mm.14099 |
| 9.55 | 9.3 | 9.7 | 9.15 | DN1ab | DN1cd | -0.37 | -5.2 | 0.0012 | 0.035 | 0.032 | ENSMUSG00000027940 | protein_coding | Tpm3 | tropomyosin 3, gamma | 59069 | 3 | 90072649 | 90100902 | 1 | Mm.240839 Mm.421791 Mm.442249 Mm.490309 |
| 4.63 | 2.92 | 4.52 | 3.58 | DN1ab | DN1cd | -1.35 | -5.2 | 0.0012 | 0.035 | 0.15 | ENSMUSG00000026556 | protein_coding | Vangl2 | VANGL planar cell polarity 2 | 93840 | 1 | 172000960 | 172028444 | -1 | Mm.36148 Mm.392110 |
| 5.37 | 4.87 | 6.14 | 5.19 | DN1ab | DN1cd | -0.68 | -5.19 | 0.0012 | 0.035 | 0.08 | ENSMUSG00000004270 | protein_coding | Lpcat3 | lysophosphatidylcholine acyltransferase 3 | 14792 | 6 | 124663027 | 124704418 | 1 | Mm.273915 |
| 8 | 8.47 | 7.66 | 7.95 | DN1ab | DN1cd | 0.41 | 5.19 | 0.0012 | 0.035 | 0.3 | ENSMUSG00000028884 | protein_coding | Rpa2 | replication protein A2 | 19891 | 4 | 132768332 | 132778752 | 1 | Mm.2870 |
| 7.42 | 7.99 | 6.63 | 6.83 | DN1ab | DN1cd | 0.44 | 5.18 | 0.0013 | 0.036 | 0.028 | ENSMUSG00000085241 | processed_transcript | Snhg3 | small nucleolar RNA host gene 3 | NA | 4 | 132348014 | 132353686 | -1 | |
| 7.82 | 7.47 | 8.3 | 7.57 | DN1ab | DN1cd | -0.49 | -5.17 | 0.0013 | 0.036 | 0.041 | ENSMUSG00000046722 | protein_coding | Cdc42se1 | CDC42 small effector 1 | 57912 | 3 | 95228732 | 95236409 | 1 | Mm.28189 Mm.331208 |
| 2.61 | 3.34 | 2.48 | 3.75 | DN1ab | DN1cd | 0.92 | 5.17 | 0.0013 | 0.036 | 0.16 | ENSMUSG00000039033 | protein_coding | Tasp1 | taspase, threonine aspartase 1 | 75812 | 2 | 139833480 | 140066805 | -1 | Mm.293449 |
| 0.6 | 1.75 | -0.39 | 2.12 | DN1ab | DN1cd | 1.62 | 5.17 | 0.0013 | 0.036 | 0.021 | ENSMUSG00000071235 | protein_coding | Vrtn | vertebrae development associated | 432677 | 12 | 84641019 | 84651455 | 1 | Mm.311913 |
| -1.22 | -3.86 | -2.81 | -4.02 | DN1ab | DN1cd | -2.12 | -5.17 | 0.0013 | 0.036 | 0.091 | ENSMUSG00000022582 | protein_coding | Ly6g | lymphocyte antigen 6 complex, locus G | 546644 | 15 | 75155240 | 75159126 | 1 | |
| 5.91 | 6.51 | 5.47 | 6.56 | DN1ab | DN1cd | 0.77 | 5.17 | 0.0013 | 0.036 | 0.11 | ENSMUSG00000037486 | protein_coding | Asxl2 | additional sex combs like 2, transcriptional regulator | 75302 | 12 | 3426857 | 3506852 | 1 | Mm.120475 |
| -3.82 | -1.93 | -3.94 | -0.67 | DN1ab | DN1cd | 2.42 | 5.16 | 0.0013 | 0.036 | 0.16 | ENSMUSG00000086526 | lincRNA | Gm12762 | predicted gene 12762 | NA | 7 | 34230420 | 34234102 | -1 | |
| 3.07 | 2.09 | 2.92 | 2.38 | DN1ab | DN1cd | -0.8 | -5.15 | 0.0013 | 0.036 | 0.18 | ENSMUSG00000026826 | protein_coding | Nr4a2 | nuclear receptor subfamily 4, group A, member 2 | 18227 | 2 | 57106830 | 57124003 | -1 | Mm.3507 |
| 4.14 | 4.84 | 4.13 | 4.5 | DN1ab | DN1cd | 0.59 | 5.15 | 0.0013 | 0.036 | 0.18 | ENSMUSG00000055491 | protein_coding | Pprc1 | peroxisome proliferative activated receptor, gamma, coactivator-related 1 | 226169 | 19 | 46044886 | 46072915 | 1 | Mm.2415 |
| 1.98 | 3.46 | 1.1 | 4.03 | DN1ab | DN1cd | 1.96 | 5.14 | 0.0013 | 0.036 | 0.062 | ENSMUSG00000020900 | protein_coding | Myh10 | myosin, heavy polypeptide 10, non-muscle | 77579 | 11 | 68691559 | 68816632 | 1 | Mm.218233 |
| 3.83 | 3.29 | 5.63 | 4.39 | DN1ab | DN1cd | -0.85 | -5.13 | 0.0013 | 0.036 | 0.019 | ENSMUSG00000018819 | protein_coding | Lsp1 | lymphocyte specific 1 | 16985 | 7 | 142460809 | 142494867 | 1 | Mm.234003 |
| 1.63 | 2.42 | 3.17 | 4.41 | DN1ab | DN1cd | 0.97 | 5.13 | 0.0013 | 0.036 | 0.28 | ENSMUSG00000028339 | protein_coding | Col15a1 | collagen, type XV, alpha 1 | 12819 | 4 | 47208161 | 47313167 | 1 | Mm.233547 |
| 7.64 | 6.91 | 8.15 | 6.9 | DN1ab | DN1cd | -0.92 | -5.13 | 0.0013 | 0.036 | 0.17 | ENSMUSG00000026159 | protein_coding | Agfg1 | ArfGAP with FG repeats 1 | 15463 | 1 | 82839483 | 82901182 | 1 | Mm.392569 Mm.433409 |
| 5.21 | 4.82 | 5.51 | 4.77 | DN1ab | DN1cd | -0.52 | -5.12 | 0.0013 | 0.036 | 0.098 | ENSMUSG00000021978 | protein_coding | Extl3 | exostoses (multiple)-like 3 | 54616 | 14 | 65052060 | 65149855 | -1 | Mm.103748 |
| -2.91 | 0.13 | -3.94 | 1.07 | DN1ab | DN1cd | 3.72 | 5.11 | 0.0013 | 0.037 | 0.22 | ENSMUSG00000041361 | protein_coding | Myzap | myocardial zonula adherens protein | 102371 | 9 | 71504347 | 71592360 | -1 | Mm.27585 |
| 0.44 | -3.86 | 0.07 | -2.07 | DN1ab | DN1cd | -3.41 | -5.1 | 0.0014 | 0.037 | 0.11 | ENSMUSG00000026235 | protein_coding | Epha4 | Eph receptor A4 | 13838 | 1 | 77367185 | 77515088 | -1 | Mm.400747 |
| 3.42 | 2.06 | 3.17 | 2.42 | DN1ab | DN1cd | -1.09 | -5.1 | 0.0014 | 0.037 | 0.16 | ENSMUSG00000041231 | protein_coding | Ublcp1 | ubiquitin-like domain containing CTD phosphatase 1 | 79560 | 11 | 44454571 | 44470498 | -1 | Mm.338784 |

| | | | | | | | | | | | | | | | | | | | | |
|---|---|---|---|---|---|---|---|---|---|---|---|---|---|---|---|---|---|---|---|---|
| 8.42 | 8.26 | 9.14 | 8.86 | DN1ab | DN1cd | -0.21 | -5.09 | 0.0014 | 0.037 | 0.17 | ENSMUSG00000045962 | protein_coding | Wnk1 | WNK lysine deficient protein kinase 1 | 232341 | 6 | 119923969 | 120038672 | -1 | Mm.333349 |
| 4.91 | 5.49 | 4.66 | 5.62 | DN1ab | DN1cd | 0.71 | 5.07 | 0.0014 | 0.038 | 0.2 | ENSMUSG00000028528 | protein_coding | Dnajc6 | DnaJ heat shock protein family (Hsp40) member C6 | 72685 | 4 | 101496648 | 101642799 | 1 | Mm.76494 |
| 5.31 | 5.81 | 5.24 | 5.51 | DN1ab | DN1cd | 0.42 | 5.06 | 0.0014 | 0.038 | 0.18 | ENSMUSG00000032508 | protein_coding | Myd88 | myeloid differentiation primary response gene 88 | 17874 | 9 | 119335934 | 119341411 | -1 | Mm.213003 |
| 7.81 | 7.55 | 8.3 | 7.87 | DN1ab | DN1cd | -0.33 | -5.06 | 0.0014 | 0.038 | 0.2 | ENSMUSG00000026219 | protein_coding | Trip12 | thyroid hormone receptor interactor 12 | 14897 | 1 | 84721189 | 84840516 | -1 | Mm.209265 |
| 7.12 | 6.85 | 7.45 | 6.98 | DN1ab | DN1cd | -0.34 | -5.06 | 0.0014 | 0.038 | 0.17 | ENSMUSG00000020476 | protein_coding | Dbnl | drebrin-like | 13169 | 11 | 5788488 | 5800962 | 1 | Mm.256925 |
| 1.48 | -0.44 | 2.42 | -0.85 | DN1ab | DN1cd | -2.5 | -5.06 | 0.0014 | 0.038 | 0.19 | ENSMUSG00000021696 | protein_coding | Elovl7 | ELOVL family member 7, elongation of long chain fatty acids (yeast) | 74559 | 13 | 108214404 | 108285683 | 1 | Mm.286127 |
| 4.17 | 3.01 | 4.55 | 3.97 | DN1ab | DN1cd | -0.9 | -5.06 | 0.0014 | 0.038 | 0.1 | ENSMUSG00000025408 | protein_coding | Ddit3 | DNA-damage inducible transcript 3 | 13198 | 10 | 127290774 | 127296288 | 1 | Mm.110220 |
| 6.46 | 7.37 | 5.33 | 7.04 | DN1ab | DN1cd | 1.19 | 5.06 | 0.0014 | 0.038 | 0.11 | ENSMUSG00000053310 | protein_coding | Nrgn | neurogranin | 64011 | 9 | 37544492 | 37552904 | -1 | Mm.335065 |
| 2 | 3.13 | 2.98 | 3.42 | DN1ab | DN1cd | 0.88 | 5.05 | 0.0014 | 0.038 | 0.045 | ENSMUSG00000037405 | protein_coding | Icam1 | intercellular adhesion molecule 1 | 15894 | 9 | 21015985 | 21028817 | 1 | Mm.435508 |
| 3.85 | 4.59 | 3.14 | 4.47 | DN1ab | DN1cd | 0.95 | 5.05 | 0.0014 | 0.038 | 0.14 | ENSMUSG00000068036 | protein_coding | Afdn | afadin, adherens junction formation factor | 17356 | 17 | 13760539 | 13906150 | 1 | Mm.59167 |
| 7.36 | 9.01 | 5.29 | 8.8 | DN1ab | DN1cd | 2.27 | 5.05 | 0.0015 | 0.038 | 0.035 | ENSMUSG00000031785 | protein_coding | Adgrg1 | adhesion G protein-coupled receptor G1 | 14766 | 8 | 94974751 | 95014217 | 1 | Mm.290834 |
| 5.34 | 5.62 | 5.27 | 5.78 | DN1ab | DN1cd | 0.36 | 5.04 | 0.0015 | 0.038 | 0.14 | ENSMUSG00000035770 | protein_coding | Dync1li2 | dynein, cytoplasmic 1 light intermediate chain 2 | 234663 | 8 | 104417680 | 104443047 | -1 | Mm.289583 |
| 0.42 | -3.86 | 1.29 | -1.01 | DN1ab | DN1cd | -3.42 | -5.03 | 0.0015 | 0.039 | 0.16 | ENSMUSG00000020690 | protein_coding | Efcab3 | EF-hand calcium binding domain 3 | 70894 | 11 | 105063592 | 105117537 | 1 | Mm.87320 |
| 5.56 | 4.77 | 6.83 | 5.23 | DN1ab | DN1cd | -1.12 | -5.03 | 0.0015 | 0.039 | 0.063 | ENSMUSG00000038437 | protein_coding | Mllt6 | myeloid/lymphoid or mixed-lineage leukemia; translocated to, 6 | 246198 | 11 | 97663414 | 97685463 | 1 | Mm.23685 |
| 4.4 | 5.05 | 4.8 | 5.09 | DN1ab | DN1cd | 0.52 | 5.02 | 0.0015 | 0.039 | 0.089 | ENSMUSG00000038976 | protein_coding | Ppp1r9b | protein phosphatase 1, regulatory subunit 9B | 217124 | 11 | 94991035 | 95006899 | 1 | Mm.229087 |
| 2.53 | 0.5 | 1.95 | 0.93 | DN1ab | DN1cd | -1.56 | -5.01 | 0.0015 | 0.039 | 0.11 | ENSMUSG00000011463 | protein_coding | Cpb1 | carboxypeptidase B1 (tissue) | 76703 | 3 | 20248264 | 20275733 | -1 | Mm.34692 |
| -3.14 | -0.68 | -3.94 | 0.05 | DN1ab | DN1cd | 3.03 | 5.01 | 0.0015 | 0.039 | 0.25 | ENSMUSG00000086194 | unprocessed_pseudogene | Gm16329 | predicted gene 16329 | NA | 1 | 21354005 | 21355116 | 1 | |
| 9.54 | 9.83 | 8.88 | 9.51 | DN1ab | DN1cd | 0.41 | 5 | 0.0015 | 0.039 | 0.028 | ENSMUSG00000066551 | protein_coding | Hmgb1 | high mobility group box 1 | 15289 | 5 | 149046702 | 149184489 | -1 | Mm.207047 Mm.313345 |
| 3.19 | 3.93 | 2.87 | 4.44 | DN1ab | DN1cd | 1.04 | 5 | 0.0015 | 0.039 | 0.042 | ENSMUSG00000046718 | protein_coding | Bst2 | bone marrow stromal cell antigen 2 | 69550 | 8 | 71534255 | 71537456 | -1 | Mm.260325 |
| 5.29 | 4.85 | 6.01 | 5.03 | DN1ab | DN1cd | -0.65 | -4.99 | 0.0015 | 0.039 | 0.021 | ENSMUSG00000042608 | protein_coding | Stk40 | serine/threonine kinase 40 | 74178 | 4 | 126103957 | 126141029 | 1 | Mm.440269 |
| 5.73 | 5.41 | 6.09 | 5.58 | DN1ab | DN1cd | -0.39 | -4.99 | 0.0016 | 0.04 | 0.27 | ENSMUSG00000057738 | protein_coding | Sptan1 | spectrin alpha, non-erythrocytic 1 | 20740 | 2 | 29965560 | 30031451 | 1 | Mm.204969 |
| 1.08 | 3.31 | -1.39 | 2.19 | DN1ab | DN1cd | 2.63 | 4.97 | 0.0016 | 0.04 | 0.27 | ENSMUSG00000048001 | protein_coding | Hes5 | hes family bHLH transcription factor 5 | 15208 | 4 | 154960923 | 154962371 | 1 | Mm.137268 |
| 1.22 | 3.57 | -0.78 | 3.58 | DN1ab | DN1cd | 2.98 | 4.97 | 0.0016 | 0.04 | 0.12 | ENSMUSG00000041722 | protein_coding | Khdc1c | KH domain containing 1C | 433278 | 1 | 21368331 | 21370845 | 1 | Mm.26006 Mm.484599 |
| -1.22 | 0.21 | -2.81 | -0.03 | DN1ab | DN1cd | 1.88 | 4.96 | 0.0016 | 0.04 | 0.088 | ENSMUSG00000035964 | protein_coding | Tmem59l | transmembrane protein 59-like | 67937 | 8 | 70483867 | 70487358 | -1 | Mm.23002 |
| 8.94 | 8.56 | 9.5 | 8.8 | DN1ab | DN1cd | -0.51 | -4.96 | 0.0016 | 0.04 | 0.12 | ENSMUSG00000090841 | protein_coding | Myl6 | myosin, light polypeptide 6, alkali, smooth muscle and non-muscle | 17904 | 10 | 128490860 | 128494145 | -1 | Mm.337074 |
| 7.63 | 8.07 | 7.13 | 7.81 | DN1ab | DN1cd | 0.53 | 4.95 | 0.0016 | 0.04 | 0.29 | ENSMUSG00000026511 | protein_coding | Srp9 | signal recognition particle 9 | 27058 | 1 | 182124737 | 182132407 | 1 | Mm.273196 Mm.303071 |
| 6.4 | 6.72 | 6.12 | 6.74 | DN1ab | DN1cd | 0.43 | 4.95 | 0.0016 | 0.041 | 0.091 | ENSMUSG00000025609 | protein_coding | Mkln1 | muskelin 1, intracellular mediator containing kelch motifs | 27418 | 6 | 31398735 | 31516811 | 1 | Mm.238038 Mm.392838 Mm.440531 |
| 7.84 | 8.3 | 6.5 | 7.45 | DN1ab | DN1cd | 0.62 | 4.94 | 0.0016 | 0.041 | 0.049 | ENSMUSG00000030189 | protein_coding | Ybx3 | Y box protein 3 | 56449 | 6 | 131364855 | 131388476 | -1 | Mm.458000 |
| -1.29 | -3.12 | -0.88 | -4.02 | DN1ab | DN1cd | -2.31 | -4.94 | 0.0016 | 0.041 | 0.19 | ENSMUSG00000107251 | TEC | Gm43102 | predicted gene 43102 | NA | 5 | 75737882 | 75740205 | 1 | |
| 4.69 | 3.96 | 6.29 | 4.89 | DN1ab | DN1cd | -1.02 | -4.94 | 0.0016 | 0.041 | 0.1 | ENSMUSG00000035891 | protein_coding | Cerk | ceramide kinase | 223753 | 15 | 86139128 | 86186141 | -1 | Mm.222685 |
| 4.38 | 4.96 | 3.58 | 4.58 | DN1ab | DN1cd | 0.73 | 4.93 | 0.0017 | 0.041 | 0.19 | ENSMUSG00000029186 | protein_coding | Pi4k2b | phosphatidylinositol 4-kinase type 2 beta | 67073 | 5 | 52741574 | 52769340 | 1 | Mm.248647 |
| -0.74 | 1.06 | 0.44 | 1.37 | DN1ab | DN1cd | 1.5 | 4.93 | 0.0017 | 0.041 | 0.19 | ENSMUSG00000026688 | protein_coding | Mgst3 | microsomal glutathione S-transferase 3 | 66447 | 1 | 167371966 | 167393841 | -1 | Mm.218286 |
| 6.82 | 7.19 | 7.18 | 7.37 | DN1ab | DN1cd | 0.3 | 4.93 | 0.0017 | 0.041 | 0.19 | ENSMUSG00000001419 | protein_coding | Mef2d | myocyte enhancer factor 2D | 17261 | 3 | 88142372 | 88172086 | 1 | Mm.28184 Mm.485397 |
| -1.01 | -3.86 | -2.81 | -4.02 | DN1ab | DN1cd | -2.25 | -4.92 | 0.0017 | 0.041 | 0.08 | ENSMUSG00000102588 | TEC | Gm37444 | predicted gene, 37444 | NA | 1 | 14255552 | 14255990 | -1 | |
| 2.33 | 3.5 | 1.2 | 3.61 | DN1ab | DN1cd | 1.59 | 4.92 | 0.0017 | 0.041 | 0.062 | ENSMUSG00000032717 | protein_coding | Mdfi | MyoD family inhibitor | 17240 | 17 | 47815328 | 47834691 | -1 | Mm.329100 |
| 3.37 | 4.55 | 2.13 | 4.64 | DN1ab | DN1cd | 1.63 | 4.9 | 0.0017 | 0.042 | 0.044 | ENSMUSG00000022272 | protein_coding | Myo10 | myosin X | 17909 | 15 | 25622525 | 25813673 | 1 | Mm.60590 |
| 5.81 | 6.13 | 5.88 | 6.44 | DN1ab | DN1cd | 0.41 | 4.89 | 0.0017 | 0.042 | 0.19 | ENSMUSG00000049878 | protein_coding | Rlf | rearranged L-myc fusion sequence | 109263 | 4 | 121145373 | 121215084 | -1 | Mm.215745 |
| -0.59 | -3.12 | 4.16 | -1.46 | DN1ab | DN1cd | -3.86 | -4.89 | 0.0017 | 0.042 | 0.038 | ENSMUSG00000020383 | protein_coding | Il13 | interleukin 13 | 16163 | 11 | 53631324 | 53634702 | -1 | Mm.1284 |
| 3.18 | 4.04 | 2.4 | 4.13 | DN1ab | DN1cd | 1.16 | 4.88 | 0.0018 | 0.042 | 0.07 | ENSMUSG00000035342 | protein_coding | Lzts2 | leucine zipper, putative tumor suppressor 2 | 226154 | 19 | 45015176 | 45027104 | 1 | Mm.28204 Mm.329877 |
| -0.02 | 1.17 | 0.41 | 2.28 | DN1ab | DN1cd | 1.43 | 4.88 | 0.0018 | 0.042 | 0.3 | ENSMUSG00000026639 | protein_coding | Lamb3 | laminin, beta 3 | 16780 | 1 | 193207699 | 193343878 | 1 | Mm.435441 |
| 6.25 | 7 | 5.83 | 7.46 | DN1ab | DN1cd | 1.07 | 4.87 | 0.0018 | 0.042 | 0.044 | ENSMUSG00000030263 | protein_coding | Lrmp | lymphoid-restricted membrane protein | 16970 | 6 | 145115653 | 145174934 | 1 | Mm.843 |
| 5.78 | 5.42 | 5.91 | 5.29 | DN1ab | DN1cd | -0.46 | -4.86 | 0.0018 | 0.043 | 0.19 | ENSMUSG00000032872 | protein_coding | Cyb5r4 | cytochrome b5 reductase 4 | 266690 | 9 | 87022014 | 87077774 | 1 | Mm.30166 Mm.317460 |
| 3.3 | 3.93 | 2.72 | 4 | DN1ab | DN1cd | 0.86 | 4.86 | 0.0018 | 0.043 | 0.069 | ENSMUSG00000001053 | protein_coding | N4bp3 | NEDD4 binding protein 3 | 212706 | 11 | 51643063 | 51650842 | -1 | Mm.41760 |
| -3.82 | -0.44 | -2.01 | -0.58 | DN1ab | DN1cd | 2.71 | 4.85 | 0.0018 | 0.043 | 0.094 | ENSMUSG00000084964 | processed_transcript | Gm15503 | predicted gene 15503 | NA | 7 | 128408968 | 128412341 | -1 | |
| 8.61 | 9.37 | 7.83 | 9.07 | DN1ab | DN1cd | 0.93 | 4.85 | 0.0018 | 0.043 | 0.25 | ENSMUSG00000004099 | protein_coding | Dnmt1 | DNA methyltransferase (cytosine-5) 1 | 13433 | 9 | 20907209 | 20959888 | -1 | Mm.128580 Mm.485562 |
| 7.86 | 8.08 | 7.95 | 8.46 | DN1ab | DN1cd | 0.33 | 4.84 | 0.0018 | 0.043 | 0.019 | ENSMUSG00000000409 | protein_coding | Lck | lymphocyte protein tyrosine kinase | 16818 | 4 | 129548344 | 129573641 | -1 | Mm.293753 |
| 4.79 | 5.29 | 3.9 | 4.84 | DN1ab | DN1cd | 0.66 | 4.83 | 0.0019 | 0.043 | 0.13 | ENSMUSG00000035268 | protein_coding | Pkig | protein kinase inhibitor, gamma | 18769 | 2 | 163658386 | 163726158 | 1 | Mm.10091 |
| 5.38 | 5.88 | 5.43 | 6.46 | DN1ab | DN1cd | 0.69 | 4.83 | 0.0019 | 0.043 | 0.067 | ENSMUSG00000034480 | protein_coding | Diaph2 | diaphanous related formin 2 | 54004 | X | 129749742 | 130465834 | 1 | Mm.10763 |
| 2.9 | 1.37 | 3.22 | 2.49 | DN1ab | DN1cd | -1.14 | -4.82 | 0.0019 | 0.044 | 0.08 | ENSMUSG00000000881 | protein_coding | Dlg3 | discs large MAGUK scaffold protein 3 | 53310 | X | 100767722 | 100818410 | 1 | Mm.4615 |
| 0.98 | -1.14 | 5.1 | -0.24 | DN1ab | DN1cd | -3.81 | -4.81 | 0.0019 | 0.044 | 0.025 | ENSMUSG00000025813 | protein_coding | Homer2 | homer scaffolding protein 2 | 26557 | 7 | 81600481 | 81707527 | -1 | Mm.228 |
| 2.98 | 1.68 | 4.41 | 1.87 | DN1ab | DN1cd | -1.86 | -4.8 | 0.0019 | 0.044 | 0.11 | ENSMUSG00000028063 | protein_coding | Lmna | lamin A | 16905 | 3 | 88480147 | 88509956 | -1 | Mm.243014 Mm.471227 |
| 1.82 | 0.3 | 2.25 | -0.77 | DN1ab | DN1cd | -2.1 | -4.79 | 0.002 | 0.045 | 0.093 | ENSMUSG00000109602 | antisense | Gm31597 | predicted gene, 31597 | NA | 7 | 45742049 | 45750257 | 1 | |
| 8.18 | 7.95 | 8.33 | 7.8 | DN1ab | DN1cd | -0.34 | -4.78 | 0.002 | 0.045 | 0.028 | ENSMUSG00000058126 | protein_coding | Tpm3-rs7 | tropomyosin 3, related sequence 7 | NA | 14 | 113314608 | 113316754 | 1 | |
| 6.14 | 6.66 | 5.92 | 6.14 | DN1ab | DN1cd | 0.41 | 4.78 | 0.002 | 0.045 | 0.098 | ENSMUSG00000031781 | protein_coding | Ciapin1 | cytokine induced apoptosis inhibitor 1 | 109006 | 8 | 94819804 | 94838358 | -1 | Mm.27582 Mm.472546 |
| 5 | 3.58 | 7.56 | 4.32 | DN1ab | DN1cd | -2.27 | -4.76 | 0.002 | 0.046 | 0.043 | ENSMUSG00000037940 | protein_coding | Inpp4b | inositol polyphosphate-4-phosphatase, type II | 234515 | 8 | 81342556 | 82127914 | 1 | Mm.324468 |
| 6.44 | 7.1 | 6.05 | 7.26 | DN1ab | DN1cd | 0.86 | 4.76 | 0.002 | 0.046 | 0.14 | ENSMUSG00000022528 | protein_coding | Hes1 | hes family bHLH transcription factor 1 | 15205 | 16 | 30064384 | 30067796 | 1 | Mm.390859 |
| 0.1 | 1.73 | 0.06 | 3.19 | DN1ab | DN1cd | 2.15 | 4.75 | 0.002 | 0.046 | 0.11 | ENSMUSG00000022197 | protein_coding | Pdzd2 | PDZ domain containing 2 | 68070 | 15 | 12359711 | 12739924 | -1 | Mm.423205 |
| 7.2 | 6.99 | 6.98 | 6.58 | DN1ab | DN1cd | -0.28 | -4.75 | 0.002 | 0.046 | 0.15 | ENSMUSG00000020250 | protein_coding | Txnrd1 | thioredoxin reductase 1 | 50493 | 10 | 82833951 | 82897712 | 1 | Mm.210155 Mm.471159 |
| 6.18 | 6.96 | 7.53 | 7.8 | DN1ab | DN1cd | 0.59 | 4.75 | 0.002 | 0.046 | 0.029 | ENSMUSG00000023067 | protein_coding | Cdkn1a | cyclin-dependent kinase inhibitor 1A (P21) | 12575 | 17 | 29090976 | 29100727 | 1 | Mm.195663 |
| 2.92 | 2.25 | 4.14 | 2.72 | DN1ab | DN1cd | -0.99 | -4.75 | 0.002 | 0.046 | 0.069 | ENSMUSG00000020439 | protein_coding | Smtn | smoothelin | 29856 | 11 | 3517523 | 3540612 | -1 | Mm.188516 |
| 4.81 | 4.15 | 5.12 | 4.8 | DN1ab | DN1cd | -0.52 | -4.75 | 0.002 | 0.046 | 0.13 | ENSMUSG00000002763 | protein_coding | Pex6 | peroxisomal biogenesis factor 6 | 224824 | 17 | 46711463 | 46725541 | 1 | Mm.299399 |
| 7.49 | 7.16 | 8.19 | 7.5 | DN1ab | DN1cd | -0.47 | -4.75 | 0.0021 | 0.046 | 0.064 | ENSMUSG00000008475 | protein_coding | Arpc5 | actin related protein 2/3 complex, subunit 5 | 67771 | 1 | 152766542 | 152775597 | 1 | Mm.288974 |
| -1.29 | 0.36 | -2.81 | 0.3 | DN1ab | DN1cd | 2.13 | 4.75 | 0.0021 | 0.046 | 0.13 | ENSMUSG00000024034 | protein_coding | Tmprss3 | transmembrane protease, serine 3 | 140765 | 17 | 31179272 | 31198975 | -1 | Mm.214638 |
| 6.62 | 7.06 | 6.22 | 7.22 | DN1ab | DN1cd | 0.64 | 4.74 | 0.0021 | 0.046 | 0.03 | ENSMUSG00000023942 | protein_coding | Slc29a1 | solute carrier family 29 (nucleoside transporters), member 1 | 63959 | 17 | 45585200 | 45599606 | -1 | Mm.29744 Mm.486335 |
| 7.03 | 6.77 | 7.57 | 7.04 | DN1ab | DN1cd | -0.36 | -4.74 | 0.0021 | 0.046 | 0.064 | ENSMUSG00000015575 | protein_coding | Atp6v0e | ATPase, H+ transporting, lysosomal V0 subunit E | 11974 | 17 | 26676396 | 26699644 | 1 | Mm.22602 |
| 4.84 | 5.37 | 3.56 | 4.63 | DN1ab | DN1cd | 0.72 | 4.73 | 0.0021 | 0.046 | 0.087 | ENSMUSG00000047221 | protein_coding | Fam185a | family with sequence similarity 185, member A | 330050 | 5 | 21424958 | 21482124 | 1 | Mm.40655 |
| 6.01 | 6.35 | 5.41 | 5.99 | DN1ab | DN1cd | 0.42 | 4.72 | 0.0021 | 0.046 | 0.22 | ENSMUSG00000048327 | protein_coding | Ckap2l | cytoskeleton associated protein 2-like | 70466 | 2 | 129268210 | 129297212 | -1 | Mm.45785 Mm.491209 |
| 6.94 | 7.56 | 6.36 | 7.81 | DN1ab | DN1cd | 0.92 | 4.7 | 0.0022 | 0.047 | 0.025 | ENSMUSG00000045382 | protein_coding | Cxcr4 | chemokine (C-X-C motif) receptor 4 | 12767 | 1 | 128588199 | 128592293 | -1 | Mm.1401 |
| 7.51 | 7.29 | 7.67 | 7.24 | DN1ab | DN1cd | -0.3 | -4.69 | 0.0022 | 0.047 | 0.14 | ENSMUSG00000041926 | protein_coding | Rnpep | arginyl aminopeptidase (aminopeptidase B) | 215615 | 1 | 135262712 | 135284084 | -1 | Mm.291048 |
| -2.24 | -0.17 | -1.14 | -0.44 | DN1ab | DN1cd | 1.61 | 4.68 | 0.0022 | 0.048 | 0.044 | ENSMUSG00000084048 | processed_pseudogene | Gm12042 | predicted gene 12042 | NA | 11 | 21295560 | 21295860 | 1 | |
| 3.62 | 4.46 | 4.6 | 4.96 | DN1ab | DN1cd | 0.66 | 4.67 | 0.0022 | 0.048 | 0.1 | ENSMUSG00000032089 | protein_coding | Il10ra | interleukin 10 receptor, alpha | 16154 | 9 | 45253837 | 45269149 | -1 | Mm.379327 |
| 3.58 | 4.53 | 3.15 | 4.86 | DN1ab | DN1cd | 1.22 | 4.67 | 0.0022 | 0.048 | 0.18 | ENSMUSG00000085399 | processed_transcript | Foxd2os | forkhead box D2, opposite strand | NA | 4 | 114909257 | 114921967 | 1 | |
| -0.11 | -1.7 | 3.48 | -0.39 | DN1ab | DN1cd | -2.66 | -4.67 | 0.0022 | 0.048 | 0.03 | ENSMUSG00000024793 | protein_coding | Tnfrsf25 | tumor necrosis factor receptor superfamily, member 25 | 85030 | 4 | 152115934 | 152120119 | 1 | Mm.101198 |
| 2.09 | 0.82 | 2.72 | 2.15 | DN1ab | DN1cd | -0.93 | -4.67 | 0.0023 | 0.048 | 0.073 | ENSMUSG00000073427 | protein_coding | Gm4924 | predicted gene 4924 | 237412 | 10 | 82354255 | 82422882 | 1 | Mm.189519 |
| 4.79 | 5.59 | 3.91 | 5.46 | DN1ab | DN1cd | 1.06 | 4.66 | 0.0023 | 0.048 | 0.11 | ENSMUSG00000021457 | protein_coding | Syk | spleen tyrosine kinase | 20963 | 13 | 52583173 | 52648792 | 1 | Mm.375031 |
| 4.68 | 4.29 | 5.59 | 4.77 | DN1ab | DN1cd | -0.56 | -4.66 | 0.0023 | 0.048 | 0.064 | ENSMUSG00000074342 | protein_coding | I830077J02Rik | RIKEN cDNA I830077J02 gene | 433638 | 3 | 105924358 | 105932664 | -1 | Mm.239081 |
| 4.34 | 4.65 | 4.03 | 4.6 | DN1ab | DN1cd | 0.4 | 4.65 | 0.0023 | 0.049 | 0.16 | ENSMUSG00000027088 | protein_coding | Phospho2 | phosphatase, orphan 2 | 73373 | 2 | 69789623 | 69800005 | 1 | Mm.9621 |
| -1.92 | 0.6 | -1.01 | -0.05 | DN1ab | DN1cd | 2.03 | 4.64 | 0.0023 | 0.049 | 0.094 | ENSMUSG00000092212 | polymorphic_pseudogene | Slc22a13b-ps | solute carrier family 22 (organic cation transporter), member 13b, pseudogene | NA | 9 | 119220488 | 119231692 | -1 | |
| 5.67 | 5.97 | 5.16 | 5.86 | DN1ab | DN1cd | 0.44 | 4.64 | 0.0023 | 0.049 | 0.028 | ENSMUSG00000025451 | protein_coding | Paip1 | polyadenylate binding protein-interacting protein 1 | 218693 | 13 | 119428601 | 119458218 | 1 | Mm.490243 |
| 1.24 | -0.41 | 2.63 | 1.9 | DN1ab | DN1cd | -1.16 | -4.63 | 0.0023 | 0.049 | 0.053 | ENSMUSG00000097729 | processed_transcript | 2310015A10Rik | RIKEN cDNA 2310015A10 gene | NA | 12 | 80120546 | 80132844 | -1 | |
| 7.01 | 7.32 | 6.73 | 7.29 | DN1ab | DN1cd | 0.4 | 4.62 | 0.0024 | 0.049 | 0.19 | ENSMUSG00000045973 | protein_coding | Slc25a51 | solute carrier family 25, member 51 | 230125 | 4 | 45395923 | 45408766 | -1 | Mm.260210 |
| 4.04 | 4.44 | 4.06 | 5.01 | DN1ab | DN1cd | 0.6 | 4.62 | 0.0024 | 0.049 | 0.022 | ENSMUSG00000054676 | protein_coding | 1600014C10Rik | RIKEN cDNA 1600014C10 gene | 72244 | 7 | 38183217 | 38197568 | 1 | Mm.187554 |
| 5.89 | 6.47 | 5.81 | 6.05 | DN1ab | DN1cd | 0.46 | 4.62 | 0.0024 | 0.049 | 0.1 | ENSMUSG00000038459 | protein_coding | Abhd17c | abhydrolase domain containing 17C | 70178 | 7 | 84109356 | 84151893 | -1 | Mm.23914 |
| -0.03 | 2.38 | -1.13 | 4.52 | DN1ab | DN1cd | 3.45 | 4.61 | 0.0024 | 0.049 | 0.027 | ENSMUSG00000024353 | protein_coding | Mzb1 | marginal zone B and B1 cell-specific protein 1 | 69816 | 18 | 35647266 | 35649367 | -1 | Mm.27252 |
| 4.1 | 3.68 | 4.84 | 3.9 | DN1ab | DN1cd | -0.63 | -4.61 | 0.0024 | 0.049 | 0.048 | ENSMUSG00000036661 | protein_coding | Dennd3 | DENN/MADD domain containing 3 | 105841 | 15 | 73512560 | 73572242 | 1 | Mm.322557 |
| 1.13 | 2.71 | -0.24 | 2.84 | DN1ab | DN1cd | 2.07 | 4.61 | 0.0024 | 0.049 | 0.12 | ENSMUSG00000067586 | protein_coding | S1pr3 | sphingosine-1-phosphate receptor 3 | 13610 | 13 | 51408639 | 51422797 | 1 | Mm.136736 |
| 5.59 | 4.93 | 6.52 | 5.27 | DN1ab | DN1cd | -0.89 | -4.6 | 0.0024 | 0.049 | 0.14 | ENSMUSG00000020437 | protein_coding | Myo1g | myosin IG | 246177 | 11 | 6506548 | 6520965 | -1 | Mm.239554 |
| 2.04 | 1.22 | 3.38 | 1.76 | DN1ab | DN1cd | -1.18 | -4.6 | 0.0024 | 0.05 | 0.12 | ENSMUSG00000032997 | protein_coding | Chpf | chondroitin polymerizing factor | 74241 | 1 | 75474569 | 75479307 | -1 | Mm.260577 |
| 8.2 | 7.86 | 8.64 | 8.03 | DN1ab | DN1cd | -0.44 | -4.6 | 0.0024 | 0.05 | 0.16 | ENSMUSG00000024048 | protein_coding | Myl12a | myosin, light chain 12A, regulatory, non-sarcomeric | 67268 | 17 | 70993656 | 71002878 | -1 | Mm.489605 |
| 3.71 | 3.25 | 4.38 | 3.49 | DN1ab | DN1cd | -0.63 | -4.6 | 0.0025 | 0.05 | 0.11 | ENSMUSG00000033436 | protein_coding | Armcx2 | armadillo repeat containing, X-linked 2 | 67416 | X | 134804145 | 134809221 | -1 | Mm.285969 |
| 5.32 | 5.66 | 4.73 | 5.34 | DN1ab | DN1cd | 0.44 | 4.59 | 0.0025 | 0.05 | 0.2 | ENSMUSG00000072582 | protein_coding | Ptrh2 | peptidyl-tRNA hydrolase 2 | 217057 | 11 | 86683985 | 86692457 | 1 | Mm.24868 |
| 5.56 | 5.25 | 6.11 | 5.49 | DN1ab | DN1cd | -0.43 | -4.59 | 0.0025 | 0.05 | 0.12 | ENSMUSG00000020736 | protein_coding | Nt5c | 5',3'-nucleotidase, cytosolic | 50773 | 11 | 115490420 | 115491862 | -1 | Mm.390379 |
| -0.09 | 1.89 | -1.13 | 3.01 | DN1ab | DN1cd | 2.69 | 4.59 | 0.0025 | 0.05 | 0.076 | ENSMUSG00000020251 | protein_coding | Glt8d2 | glycosyltransferase 8 domain containing 2 | 74782 | 10 | 82650433 | 82690650 | -1 | Mm.191281 |
| 6.54 | 6.92 | 5.73 | 6.48 | DN1ab | DN1cd | 0.51 | 4.59 | 0.0025 | 0.05 | 0.11 | ENSMUSG00000017861 | protein_coding | Mybl2 | myeloblastosis oncogene-like 2 | 17865 | 2 | 163054687 | 163084688 | 1 | Mm.4594 |
| 1.1 | 2.67 | 0.13 | 3.77 | DN1ab | DN1cd | 2.26 | 4.58 | 0.0025 | 0.05 | 0.033 | ENSMUSG00000027864 | protein_coding | Ptgfrn | prostaglandin F2 receptor negative regulator | 19221 | 3 | 101040232 | 101110278 | -1 | Mm.24807 |
| 4.79 | 4.34 | 5.34 | 4.51 | DN1ab | DN1cd | -0.6 | -4.58 | 0.0025 | 0.05 | 0.17 | ENSMUSG00000000056 | protein_coding | Narf | nuclear prelamin A recognition factor | 67608 | 11 | 121237253 | 121255856 | 1 | Mm.291832 |
| 6.13 | 6.47 | 5.72 | 6.41 | DN1ab | DN1cd | 0.47 | 4.58 | 0.0025 | 0.05 | 0.09 | ENSMUSG00000021054 | protein_coding | Sgpp1 | sphingosine-1-phosphate phosphatase 1 | 81535 | 12 | 75714249 | 75735729 | -1 | Mm.280199 |
| 7.66 | 7.42 | 7.91 | 7.4 | DN1ab | DN1cd | -0.34 | -4.57 | 0.0025 | 0.05 | 0.058 | ENSMUSG00000018326 | protein_coding | Ywhab | tyrosine 3-monooxygenase/tryptophan 5-monooxygenase activation protein, beta polypeptide | 54401 | 2 | 163994960 | 164018588 | 1 | Mm.34319 Mm.491117 |
| -2.24 | -3.86 | -0.57 | -4.02 | DN1ab | DN1cd | -2.29 | -4.57 | 0.0025 | 0.05 | 0.069 | ENSMUSG00000018387 | protein_coding | Shroom1 | shroom family member 1 | 71774 | 11 | 53457205 | 53467766 | 1 | Mm.274370 |
| 5.59 | 5.3 | 6.22 | 5.59 | DN1ab | DN1cd | -0.43 | -4.57 | 0.0025 | 0.05 | 0.082 | ENSMUSG00000008859 | protein_coding | Rala | v-ral simian leukemia viral oncogene A (ras related) | 56044 | 13 | 17880571 | 17944239 | -1 | Mm.27348 |
| 3.47 | 4.23 | 2.82 | 4.48 | DN1ab | DN1cd | 1.08 | 4.57 | 0.0025 | 0.051 | 0.057 | ENSMUSG00000041684 | protein_coding | Bivm | basic, immunoglobulin-like variable motif containing | 246229 | 1 | 44118957 | 44144770 | 1 | Mm.386883 |
| 4.3 | 5.01 | 3.77 | 5.07 | DN1ab | DN1cd | 0.92 | 4.55 | 0.0026 | 0.051 | 0.17 | ENSMUSG00000073002 | protein_coding | Vamp5 | vesicle-associated membrane protein 5 | 53620 | 6 | 72368794 | 72380468 | -1 | Mm.440321 |
| 2.17 | 3.21 | 1.31 | 3.47 | DN1ab | DN1cd | 1.43 | 4.55 | 0.0026 | 0.051 | 0.084 | ENSMUSG00000039629 | protein_coding | Strip2 | striatin interacting protein 2 | 320609 | 6 | 29917012 | 29959681 | 1 | Mm.56097 |
| -1.81 | -0.17 | -2.81 | 0.09 | DN1ab | DN1cd | 2.08 | 4.53 | 0.0026 | 0.051 | 0.21 | ENSMUSG00000087575 | lincRNA | Gm12976 | predicted gene 12976 | NA | 4 | 129261965 | 129265010 | 1 | |
| 7.81 | 8.07 | 7.51 | 8.14 | DN1ab | DN1cd | 0.39 | 4.53 | 0.0026 | 0.051 | 0.02 | ENSMUSG00000008730 | protein_coding | Hipk1 | homeodomain interacting protein kinase 1 | 15257 | 3 | 103739815 | 103791563 | -1 | Mm.20827 |
| 2.82 | 3.43 | 2.3 | 3.67 | DN1ab | DN1cd | 0.88 | 4.53 | 0.0027 | 0.052 | 0.042 | ENSMUSG00000032501 | protein_coding | Trib1 | tribbles pseudokinase 1 | 211770 | 15 | 59648350 | 59657099 | 1 | Mm.40298 |
| 5.42 | 5.92 | 5.25 | 6.25 | DN1ab | DN1cd | 0.69 | 4.53 | 0.0027 | 0.052 | 0.11 | ENSMUSG00000029776 | protein_coding | Hibadh | 3-hydroxyisobutyrate dehydrogenase | 58875 | 6 | 52546228 | 52640389 | -1 | Mm.286458 |
| 8.16 | 7.84 | 7.69 | 7.56 | DN1ab | DN1cd | -0.25 | -4.52 | 0.0027 | 0.052 | 0.091 | ENSMUSG00000020570 | protein_coding | Sypl | synaptophysin-like protein | 19027 | 12 | 32953891 | 32979860 | 1 | Mm.246304 |
| -0.62 | 1.59 | -2.81 | 2.01 | DN1ab | DN1cd | 3.02 | 4.52 | 0.0027 | 0.052 | 0.059 | ENSMUSG00000036111 | protein_coding | Lmo1 | LIM domain only 1 | 109594 | 7 | 109138572 | 109175207 | -1 | Mm.360145 |
| 5.96 | 5.57 | 6.73 | 5.93 | DN1ab | DN1cd | -0.55 | -4.52 | 0.0027 | 0.052 | 0.1 | ENSMUSG00000022175 | protein_coding | Lrp10 | low-density lipoprotein receptor-related protein 10 | 65107 | 14 | 54464137 | 54471497 | 1 | Mm.472719 |
| 9.68 | 9.38 | 10.13 | 9.4 | DN1ab | DN1cd | -0.46 | -4.52 | 0.0027 | 0.052 | 0.026 | ENSMUSG00000001175 | protein_coding | Calm1 | calmodulin 1 | 12313 | 12 | 100199435 | 100209814 | 1 | Mm.285993 |
| 4.31 | 3.89 | 4.81 | 4.01 | DN1ab | DN1cd | -0.57 | -4.52 | 0.0027 | 0.052 | 0.15 | ENSMUSG00000025232 | protein_coding | Hexa | hexosaminidase A | 15211 | 9 | 59539540 | 59565109 | 1 | Mm.2284 |
| 6.96 | 6.48 | 8.04 | 6.95 | DN1ab | DN1cd | -0.71 | -4.51 | 0.0027 | 0.052 | 0.041 | ENSMUSG00000021728 | protein_coding | Emb | embigin | 13723 | 13 | 117208536 | 117274415 | 1 | Mm.274926 |
| 4.48 | 3.96 | 6.02 | 4.77 | DN1ab | DN1cd | -0.83 | -4.51 | 0.0027 | 0.052 | 0.039 | ENSMUSG00000025959 | protein_coding | Klf7 | Kruppel-like factor 7 (ubiquitous) | 93691 | 1 | 64029447 | 64122282 | -1 | Mm.29466 Mm.458200 |
| 3.07 | 4.28 | 1.41 | 3.86 | DN1ab | DN1cd | 1.62 | 4.51 | 0.0027 | 0.052 | 0.1 | ENSMUSG00000040447 | protein_coding | Spns2 | spinster homolog 2 | 216892 | 11 | 72451638 | 72489904 | -1 | Mm.212812 |
| -0.81 | 0.99 | 0.33 | 0.97 | DN1ab | DN1cd | 1.41 | 4.51 | 0.0027 | 0.052 | 0.072 | ENSMUSG00000050394 | protein_coding | Armcx6 | armadillo repeat containing, X-linked 6 | 278097 | X | 134748461 | 134751417 | -1 | Mm.113272 |
| 6.91 | 7.15 | 6.84 | 7.33 | DN1ab | DN1cd | 0.33 | 4.5 | 0.0027 | 0.052 | 0.091 | ENSMUSG00000024750 | protein_coding | Zfand5 | zinc finger, AN1-type domain 5 | 22682 | 19 | 21272278 | 21282289 | 1 | Mm.292405 Mm.379919 |
| -2.91 | -1.06 | -3.94 | -0.48 | DN1ab | DN1cd | 2.45 | 4.49 | 0.0028 | 0.053 | 0.17 | ENSMUSG00000111690 | TEC | Gm48720 | predicted gene, 48720 | NA | 9 | 45008142 | 45009347 | -1 | |
| 3.12 | 3.67 | 3.33 | 4.48 | DN1ab | DN1cd | 0.78 | 4.49 | 0.0028 | 0.053 | 0.09 | ENSMUSG00000062939 | protein_coding | Stat4 | signal transducer and activator of transcription 4 | 20849 | 1 | 51987148 | 52107189 | 1 | Mm.1550 |
| 6.15 | 5.6 | 6.19 | 5.96 | DN1ab | DN1cd | -0.43 | -4.47 | 0.0029 | 0.053 | 0.095 | ENSMUSG00000005514 | protein_coding | Por | P450 (cytochrome) oxidoreductase | 18984 | 5 | 135670033 | 135735326 | 1 | Mm.3863 |
| 6.18 | 6.47 | 5.76 | 6.44 | DN1ab | DN1cd | 0.43 | 4.46 | 0.0029 | 0.054 | 0.036 | ENSMUSG00000025742 | protein_coding | Prps2 | phosphoribosyl pyrophosphate synthetase 2 | 110639 | X | 167346322 | 167382749 | -1 | Mm.272955 |
| 0.91 | 0.13 | 2.81 | 0.92 | DN1ab | DN1cd | -1.28 | -4.45 | 0.0029 | 0.054 | 0.04 | ENSMUSG00000074170 | protein_coding | Plekhf1 | pleckstrin homology domain containing, family F (with FYVE domain) member 1 | 72287 | 7 | 38216972 | 38228016 | -1 | Mm.333798 |
| 5 | 4.36 | 5.18 | 4.94 | DN1ab | DN1cd | -0.48 | -4.45 | 0.0029 | 0.054 | 0.055 | ENSMUSG00000031367 | protein_coding | Ap1s2 | adaptor-related protein complex 1, sigma 2 subunit | 108012 | X | 163909017 | 163933666 | 1 | Mm.426783 |
| 5.38 | 5.64 | 4.94 | 5.55 | DN1ab | DN1cd | 0.39 | 4.42 | 0.003 | 0.055 | 0.044 | ENSMUSG00000030254 | protein_coding | Rad18 | RAD18 E3 ubiquitin protein ligase | 58186 | 6 | 112619850 | 112696686 | -1 | Mm.103812 |
| 11.29 | 10.97 | 11.52 | 10.86 | DN1ab | DN1cd | -0.45 | -4.42 | 0.003 | 0.055 | 0.12 | ENSMUSG00000062825 | protein_coding | Actg1 | actin, gamma, cytoplasmic 1 | 11465 | 11 | 120345690 | 120348542 | -1 | Mm.196173 Mm.426706 Mm.470166 |
| -1.81 | -3.86 | 0.65 | -3.22 | DN1ab | DN1cd | -2.71 | -4.42 | 0.003 | 0.055 | 0.16 | ENSMUSG00000030114 | protein_coding | Klrg1 | killer cell lectin-like receptor subfamily G, member 1 | 50928 | 6 | 122270596 | 122282882 | -1 | Mm.20434 |
| 5.34 | 6.65 | 3.83 | 6.69 | DN1ab | DN1cd | 1.84 | 4.41 | 0.0031 | 0.056 | 0.07 | ENSMUSG00000042379 | protein_coding | Esm1 | endothelial cell-specific molecule 1 | 71690 | 13 | 113209659 | 113218098 | 1 | Mm.38929 |
| 7.13 | 6.62 | 7.85 | 7.66 | DN1ab | DN1cd | -0.38 | -4.4 | 0.0031 | 0.056 | 0.058 | ENSMUSG00000004612 | protein_coding | Nkg7 | natural killer cell group 7 sequence | 72310 | 7 | 43437073 | 43438249 | 1 | Mm.34613 |
| 8.73 | 9.07 | 8.28 | 9.06 | DN1ab | DN1cd | 0.5 | 4.4 | 0.0031 | 0.056 | 0.052 | ENSMUSG00000024222 | protein_coding | Fkbp5 | FK506 binding protein 5 | 14229 | 17 | 28399094 | 28517527 | -1 | Mm.276405 |
| 5.97 | 5.51 | 5.76 | 5.58 | DN1ab | DN1cd | -0.36 | -4.4 | 0.0031 | 0.056 | 0.092 | ENSMUSG00000037339 | protein_coding | Fam53a | family with sequence similarity 53, member A | 74504 | 5 | 33600347 | 33629635 | -1 | Mm.294695 |
| 1.85 | 2.73 | 0.59 | 2.6 | DN1ab | DN1cd | 1.26 | 4.39 | 0.0031 | 0.056 | 0.051 | ENSMUSG00000062866 | protein_coding | Phactr2 | phosphatase and actin regulator 2 | 215789 | 10 | 13207717 | 13474412 | -1 | Mm.283359 Mm.487722 |
| 3.27 | 2.62 | 6.61 | 4.92 | DN1ab | DN1cd | -1.18 | -4.36 | 0.0032 | 0.057 | 0.039 | ENSMUSG00000026879 | protein_coding | Gsn | gelsolin | 227753 | 2 | 35256380 | 35307892 | 1 | Mm.443 Mm.21109 |
| 4.2 | 4.69 | 4.17 | 5.21 | DN1ab | DN1cd | 0.69 | 4.36 | 0.0033 | 0.058 | 0.083 | ENSMUSG00000043505 | protein_coding | Gimap5 | GTPase, IMAP family member 5 | 317757 | 6 | 48746197 | 48754210 | 1 | Mm.455917 |
| 5.57 | 5.14 | 6.05 | 5.88 | DN1ab | DN1cd | -0.33 | -4.33 | 0.0034 | 0.059 | 0.069 | ENSMUSG00000032905 | protein_coding | Atg12 | autophagy related 12 | 67526 | 18 | 46730144 | 46741580 | -1 | Mm.9852 |
| -2.49 | 0.38 | -3.94 | 2.12 | DN1ab | DN1cd | 3.97 | 4.33 | 0.0034 | 0.059 | 0.094 | ENSMUSG00000021872 | protein_coding | Rnase10 | ribonuclease, RNase A family, 10 (non-active) | 75019 | 14 | 51007751 | 51010758 | 1 | Mm.318775 |
| 3.38 | 4.09 | 2.88 | 4.48 | DN1ab | DN1cd | 1.03 | 4.31 | 0.0035 | 0.06 | 0.066 | ENSMUSG00000024544 | protein_coding | Ldlrad4 | low density lipoprotein receptor class A domain containing 4 | 52662 | 18 | 67933257 | 68255549 | 1 | Mm.440183 Mm.470001 |
| 0.64 | 3.17 | -1.29 | 3.85 | DN1ab | DN1cd | 3.34 | 4.31 | 0.0035 | 0.06 | 0.12 | ENSMUSG00000101586 | antisense | Gm29017 | predicted gene 29017 | NA | 1 | 59151047 | 59169320 | 1 | |
| 6.25 | 6.65 | 5.62 | 6.66 | DN1ab | DN1cd | 0.63 | 4.31 | 0.0035 | 0.06 | 0.018 | ENSMUSG00000029366 | protein_coding | Dck | deoxycytidine kinase | 13178 | 5 | 88764996 | 88783281 | 1 | Mm.298892 |
| 0.78 | 2.54 | -0.63 | 3.12 | DN1ab | DN1cd | 2.41 | 4.3 | 0.0035 | 0.06 | 0.092 | ENSMUSG00000032864 | protein_coding | Rag2 | recombination activating gene 2 | 19374 | 2 | 101624718 | 101632529 | 1 | Mm.4988 |
| 8.13 | 7.75 | 8.11 | 8 | DN1ab | DN1cd | -0.27 | -4.3 | 0.0035 | 0.06 | 0.02 | ENSMUSG00000049932 | protein_coding | H2afx | H2A histone family, member X | 15270 | 9 | 44334694 | 44336077 | 1 | Mm.245931 |
| 0.73 | 2 | 0.23 | 2.73 | DN1ab | DN1cd | 1.7 | 4.29 | 0.0036 | 0.061 | 0.15 | ENSMUSG00000056367 | protein_coding | Actr3b | ARP3 actin-related protein 3B | 242894 | 5 | 25759997 | 25850688 | 1 | Mm.150319 |
| 4.07 | 4.47 | 3.83 | 4.82 | DN1ab | DN1cd | 0.62 | 4.29 | 0.0036 | 0.061 | 0.033 | ENSMUSG00000030745 | protein_coding | Il21r | interleukin 21 receptor | 60504 | 7 | 125603429 | 125633570 | 1 | Mm.155643 |
| 3.08 | 2.37 | 5.39 | 3.43 | DN1ab | DN1cd | -1.29 | -4.27 | 0.0036 | 0.061 | 0.021 | ENSMUSG00000031155 | protein_coding | Pim2 | proviral integration site 2 | 18715 | X | 7878262 | 7883432 | 1 | Mm.347478 |
| 2.44 | 1.77 | 3.8 | 2.2 | DN1ab | DN1cd | -1.07 | -4.26 | 0.0037 | 0.062 | 0.056 | ENSMUSG00000021550 | protein_coding | 2210016F16Rik | RIKEN cDNA 2210016F16 gene | 70153 | 13 | 58379817 | 58385225 | -1 | Mm.296610 |
| 1.78 | 1.08 | 3.54 | 1.68 | DN1ab | DN1cd | -1.22 | -4.25 | 0.0038 | 0.062 | 0.027 | ENSMUSG00000013698 | protein_coding | Pea15a | phosphoprotein enriched in astrocytes 15A | 18611 | 1 | 172196728 | 172206804 | -1 | Mm.544 |
| 0.04 | 1.31 | -0.83 | 2.49 | DN1ab | DN1cd | 1.98 | 4.25 | 0.0038 | 0.062 | 0.018 | ENSMUSG00000031026 | protein_coding | Trim66 | tripartite motif-containing 66 | 330627 | 7 | 109449006 | 109508134 | -1 | Mm.183528 |
| 6.02 | 5.71 | 6.54 | 5.9 | DN1ab | DN1cd | -0.44 | -4.24 | 0.0038 | 0.063 | 0.11 | ENSMUSG00000026979 | protein_coding | Psd4 | pleckstrin and Sec7 domain containing 4 | 215632 | 2 | 24367580 | 24414954 | 1 | Mm.490434 |
| 2.7 | 2.04 | 3.83 | 2.14 | DN1ab | DN1cd | -1.09 | -4.23 | 0.0038 | 0.063 | 0.033 | ENSMUSG00000047123 | protein_coding | Ticam1 | toll-like receptor adaptor molecule 1 | 106759 | 17 | 56269319 | 56276786 | -1 | Mm.203952 |
| 2.6 | 1.54 | 4.43 | 1.83 | DN1ab | DN1cd | -1.75 | -4.23 | 0.0038 | 0.063 | 0.05 | ENSMUSG00000018398 | protein_coding | 8-Sep | septin 8 | 20362 | 11 | 53519257 | 53549565 | 1 | Mm.274399 Mm.359575 |
| 0.6 | 1.75 | 0.02 | 3.04 | DN1ab | DN1cd | 1.81 | 4.22 | 0.0039 | 0.064 | 0.02 | ENSMUSG00000019768 | protein_coding | Esr1 | estrogen receptor 1 (alpha) | 13982 | 10 | 4611593 | 5005614 | 1 | Mm.9213 Mm.463262 Mm.489063 |
| 3.54 | 4.14 | 2.78 | 4.01 | DN1ab | DN1cd | 0.82 | 4.21 | 0.0039 | 0.064 | 0.13 | ENSMUSG00000054520 | protein_coding | Sh3bp2 | SH3-domain binding protein 2 | 24055 | 5 | 34525838 | 34563641 | 1 | Mm.5012 |
| 2.24 | 3.28 | 2.75 | 5.32 | DN1ab | DN1cd | 1.6 | 4.21 | 0.0039 | 0.064 | 0.037 | ENSMUSG00000045868 | protein_coding | Gvin1 | GTPase, very large interferon inducible 1 | 74558 | 7 | 106156556 | 106215326 | -1 | Mm.458429 |
| 1.32 | -0.32 | 4.8 | 0.45 | DN1ab | DN1cd | -3.01 | -4.2 | 0.004 | 0.065 | 0.046 | ENSMUSG00000027276 | protein_coding | Jag1 | jagged 1 | 16449 | 2 | 137081456 | 137116644 | -1 | Mm.22398 |
| 6.89 | 5.99 | 6.83 | 6.54 | DN1ab | DN1cd | -0.66 | -4.19 | 0.004 | 0.065 | 0.05 | ENSMUSG00000045414 | protein_coding | 1190002N15Rik | RIKEN cDNA 1190002N15 gene | 68861 | 9 | 94517864 | 94538081 | -1 | Mm.258746 |
| 3.88 | 2.42 | 3.41 | 2.91 | DN1ab | DN1cd | -1.03 | -4.18 | 0.0041 | 0.065 | 0.036 | ENSMUSG00000042129 | protein_coding | Rassf4 | Ras association (RalGDS/AF-6) domain family member 4 | 213391 | 6 | 116633008 | 116673952 | -1 | Mm.257931 |
| 4.96 | 5.3 | 4.85 | 5.6 | DN1ab | DN1cd | 0.49 | 4.17 | 0.0041 | 0.066 | 0.085 | ENSMUSG00000039879 | protein_coding | Heca | hdc homolog, cell cycle regulator | 380629 | 10 | 17868612 | 17948067 | -1 | Mm.276430 Mm.473073 |
| 4.89 | 4.47 | 5.18 | 4.22 | DN1ab | DN1cd | -0.63 | -4.17 | 0.0041 | 0.066 | 0.074 | ENSMUSG00000031861 | protein_coding | Lpar2 | lysophosphatidic acid receptor 2 | 53978 | 8 | 69822429 | 69831102 | 1 | Mm.23253 |
| 4 | 4.86 | 3.83 | 5.61 | DN1ab | DN1cd | 1.19 | 4.16 | 0.0042 | 0.066 | 0.13 | ENSMUSG00000045349 | protein_coding | Sh2d5 | SH2 domain containing 5 | 230863 | 4 | 138250403 | 138261332 | 1 | Mm.482408 |
| 0.1 | -0.99 | 1.91 | -1.03 | DN1ab | DN1cd | -1.87 | -4.14 | 0.0043 | 0.067 | 0.026 | ENSMUSG00000020419 | protein_coding | Hormad2 | HORMA domain containing 2 | 75828 | 11 | 4345814 | 4441105 | -1 | Mm.159592 |
| 1.96 | 2.5 | 1.52 | 2.82 | DN1ab | DN1cd | 0.82 | 4.13 | 0.0043 | 0.068 | 0.055 | ENSMUSG00000032567 | protein_coding | Aste1 | asteroid homolog 1 | 66595 | 9 | 105395392 | 105412229 | 1 | Mm.84941 |
| 7.59 | 8.05 | 6.81 | 7.83 | DN1ab | DN1cd | 0.66 | 4.13 | 0.0043 | 0.068 | 0.09 | ENSMUSG00000030199 | protein_coding | Etv6 | ets variant 6 | 14011 | 6 | 134035700 | 134270158 | 1 | Mm.269995 |
| -0.19 | -1.93 | 3.96 | -0.59 | DN1ab | DN1cd | -3.05 | -4.13 | 0.0044 | 0.068 | 0.044 | ENSMUSG00000021200 | protein_coding | Asb2 | ankyrin repeat and SOCS box-containing 2 | 65256 | 12 | 103321142 | 103356001 | -1 | Mm.27159 Mm.321278 |
| 2.62 | 3.94 | 0.59 | 4.12 | DN1ab | DN1cd | 2.05 | 4.12 | 0.0044 | 0.068 | 0.019 | ENSMUSG00000025780 | protein_coding | Itih5 | inter-alpha (globulin) inhibitor H5 | 209378 | 2 | 10153571 | 10256529 | 1 | Mm.313876 |
| 2.38 | 3.51 | 1.06 | 4.05 | DN1ab | DN1cd | 1.76 | 4.12 | 0.0044 | 0.068 | 0.023 | ENSMUSG00000098557 | protein_coding | Kctd12 | potassium channel tetramerisation domain containing 12 | 239217 | 14 | 102976581 | 102982637 | -1 | Mm.246466 |
| 1.99 | 2.89 | 0.97 | 3.03 | DN1ab | DN1cd | 1.3 | 4.1 | 0.0045 | 0.069 | 0.074 | ENSMUSG00000057265 | protein_coding | Bbof1 | basal body orientation factor 1 | 72873 | 12 | 84409071 | 84442887 | 1 | Mm.440428 |
| 4.32 | 4.62 | 4.15 | 4.87 | DN1ab | DN1cd | 0.46 | 4.09 | 0.0045 | 0.07 | 0.052 | ENSMUSG00000031349 | protein_coding | Nsdhl | NAD(P) dependent steroid dehydrogenase-like | 18194 | X | 72918557 | 72958514 | 1 | Mm.38792 |
| 6.59 | 6.4 | 6.87 | 6.43 | DN1ab | DN1cd | -0.29 | -4.06 | 0.0047 | 0.072 | 0.074 | ENSMUSG00000062014 | protein_coding | Gmfb | glia maturation factor, beta | 63985 | 14 | 46808149 | 46822242 | -1 | Mm.87312 |
| 5.88 | 5.19 | 7.3 | 5.63 | DN1ab | DN1cd | -1.08 | -4.03 | 0.0049 | 0.074 | 0.06 | ENSMUSG00000032009 | protein_coding | Sesn3 | sestrin 3 | 75747 | 9 | 14275067 | 14333101 | 1 | Mm.325126 Mm.439776 |

| | | | | | | | | | | | | | | | | | | | | |
|---|---|---|---|---|---|---|---|---|---|---|---|---|---|---|---|---|---|---|---|---|
| 6.18 | 6.7 | 6.58 | 6.73 | DN1ab | DN1cd | 0.39 | 4.01 | 0.005 | 0.075 | 0.047 | ENSMUSG00000022136 | protein_coding | Dnajc3 | DnaJ heat shock protein family (Hsp40) member C3 | 100037258 | 14 | 118937976 | 118981697 | 1 | Mm.12616 |
| 2.95 | 3.61 | 2.59 | 4.09 | DN1ab | DN1cd | 0.96 | 3.99 | 0.0052 | 0.076 | 0.086 | ENSMUSG00000101013 | antisense | A630072M18Rik | RIKEN cDNA A630072M18 gene | NA | 5 | 20950989 | 20956398 | 1 | |
| 5.84 | 6.09 | 5.19 | 5.78 | DN1ab | DN1cd | 0.37 | 3.98 | 0.0052 | 0.076 | 0.064 | ENSMUSG00000038047 | protein_coding | Haus6 | HAUS augmin-like complex, subunit 6 | 230376 | 4 | 86578855 | 86612055 | -1 | Mm.74730 |
| 3.79 | 3.41 | 4.69 | 3.78 | DN1ab | DN1cd | -0.6 | -3.98 | 0.0052 | 0.076 | 0.076 | ENSMUSG00000078695 | protein_coding | Cisd3 | CDGSH iron sulfur domain 3 | 217149 | 11 | 97685826 | 97688621 | 1 | Mm.29659 |
| 4.13 | 4.56 | 3.64 | 4.62 | DN1ab | DN1cd | 0.63 | 3.98 | 0.0053 | 0.077 | 0.085 | ENSMUSG00000059743 | protein_coding | Fdps | farnesyl diphosphate synthetase | 110196 | 3 | 89093588 | 89101959 | -1 | Mm.39472 Mm.335450 Mm.490094 |
| 0.26 | 2.47 | -2.8 | 3 | DN1ab | DN1cd | 3.33 | 3.98 | 0.0053 | 0.077 | 0.031 | ENSMUSG00000031786 | protein_coding | Drc7 | dynein regulatory complex subunit 7 | 330830 | 8 | 95055103 | 95078141 | 1 | Mm.277112 |
| 7.94 | 7.6 | 8.78 | 7.9 | DN1ab | DN1cd | -0.55 | -3.96 | 0.0054 | 0.078 | 0.045 | ENSMUSG00000019189 | protein_coding | Rnf145 | ring finger protein 145 | 74315 | 11 | 44518964 | 44565520 | 1 | Mm.156771 |
| 6.25 | 5.87 | 6.86 | 5.94 | DN1ab | DN1cd | -0.59 | -3.93 | 0.0056 | 0.08 | 0.07 | ENSMUSG00000040710 | protein_coding | St8sia4 | ST8 alpha-N-acetyl-neuraminide alpha-2,8-sialyltransferase 4 | 20452 | 1 | 95587682 | 95667571 | -1 | Mm.306228 |
| 6.04 | 5.62 | 7.06 | 5.88 | DN1ab | DN1cd | -0.72 | -3.9 | 0.0059 | 0.082 | 0.028 | ENSMUSG00000061577 | protein_coding | Adgrg5 | adhesion G protein-coupled receptor G5 | 382045 | 8 | 94923694 | 94943280 | 1 | Mm.334726 |
| 7.22 | 7.05 | 7.17 | 6.74 | DN1ab | DN1cd | -0.27 | -3.89 | 0.0059 | 0.082 | 0.063 | ENSMUSG00000027397 | protein_coding | Slc20a1 | solute carrier family 20, member 1 | 20515 | 2 | 129198764 | 129211616 | 1 | Mm.272675 |
| 1.99 | 3.02 | -0.88 | 2.06 | DN1ab | DN1cd | 1.63 | 3.88 | 0.0059 | 0.083 | 0.018 | ENSMUSG00000040016 | protein_coding | Ptger3 | prostaglandin E receptor 3 (subtype EP3) | 19218 | 3 | 157566892 | 157645888 | 1 | Mm.30424 |
| 3.33 | 4.2 | 3.25 | 3.43 | DN1ab | DN1cd | 0.64 | 3.88 | 0.006 | 0.083 | 0.03 | ENSMUSG00000046994 | protein_coding | Mars2 | methionine-tRNA synthetase 2 (mitochondrial) | 212679 | 1 | 55237177 | 55248470 | 1 | Mm.19223 |
| 1.99 | 2.78 | 0.11 | 2.06 | DN1ab | DN1cd | 1.17 | 3.88 | 0.006 | 0.083 | 0.058 | ENSMUSG00000020186 | protein_coding | Csrp2 | cysteine and glycine-rich protein 2 | 13008 | 10 | 110919789 | 110939622 | 1 | Mm.2020 |
| 10.05 | 9.89 | 10.08 | 9.69 | DN1ab | DN1cd | -0.24 | -3.87 | 0.0061 | 0.083 | 0.075 | ENSMUSG00000028691 | protein_coding | Prdx1 | peroxiredoxin 1 | 18477 | 4 | 116685544 | 116700822 | 1 | Mm.30929 |
| 7.72 | 7.97 | 7.62 | 8.23 | DN1ab | DN1cd | 0.38 | 3.85 | 0.0062 | 0.085 | 0.061 | ENSMUSG00000032745 | protein_coding | Gpbp1 | GC-rich promoter binding protein 1 | 73274 | 13 | 111425680 | 111490111 | -1 | Mm.57286 Mm.441830 |
| 5.8 | 5.15 | 5.99 | 5.84 | DN1ab | DN1cd | -0.45 | -3.85 | 0.0062 | 0.085 | 0.016 | ENSMUSG00000031066 | protein_coding | Usp11 | ubiquitin specific peptidase 11 | 236733 | X | 20703906 | 20720539 | 1 | Mm.34489 |
| 6.85 | 6.71 | 7.34 | 6.94 | DN1ab | DN1cd | -0.24 | -3.82 | 0.0064 | 0.086 | 0.028 | ENSMUSG00000029552 | protein_coding | Tes | testis derived transcript | 21753 | 6 | 17065149 | 17105828 | 1 | Mm.436548 |
| 6.21 | 6 | 6.58 | 5.95 | DN1ab | DN1cd | -0.37 | -3.82 | 0.0065 | 0.087 | 0.016 | ENSMUSG00000031516 | protein_coding | Dctn6 | dynactin 6 | 22428 | 8 | 34090420 | 34108796 | -1 | Mm.150373 |
| -0.28 | 1.25 | -3.5 | 0.06 | DN1ab | DN1cd | 2.15 | 3.81 | 0.0065 | 0.087 | 0.095 | ENSMUSG00000102540 | antisense | Gm38134 | predicted gene, 38134 | NA | 9 | 108010976 | 108016608 | -1 | |
| 4.58 | 4.92 | 4.24 | 5.1 | DN1ab | DN1cd | 0.53 | 3.8 | 0.0067 | 0.088 | 0.058 | ENSMUSG00000038312 | protein_coding | Edem2 | ER degradation enhancer, mannosidase alpha-like 2 | 108687 | 2 | 155701677 | 155729475 | -1 | Mm.273706 |
| 4.4 | 4.97 | 3.62 | 5.22 | DN1ab | DN1cd | 0.94 | 3.79 | 0.0067 | 0.088 | 0.029 | ENSMUSG00000041444 | protein_coding | Arhgap32 | Rho GTPase activating protein 32 | 330914 | 9 | 32116136 | 32268446 | 1 | Mm.46683 |
| 4.13 | 3.83 | 4.84 | 4.02 | DN1ab | DN1cd | -0.51 | -3.77 | 0.0069 | 0.09 | 0.044 | ENSMUSG00000028466 | protein_coding | Creb3 | cAMP responsive element binding protein 3 | 12913 | 4 | 43562332 | 43567060 | 1 | Mm.12407 |
| 4.62 | 5.18 | 3.73 | 5.13 | DN1ab | DN1cd | 0.85 | 3.76 | 0.007 | 0.091 | 0.064 | ENSMUSG00000059182 | protein_coding | Skap2 | src family associated phosphoprotein 2 | 54353 | 6 | 51857422 | 52012549 | -1 | Mm.221479 Mm.392558 |
| -1.81 | -0.55 | -3.94 | -0.48 | DN1ab | DN1cd | 2.03 | 3.75 | 0.007 | 0.091 | 0.039 | ENSMUSG00000105707 | TEC | Gm43753 | predicted gene 43753 | NA | 5 | 62656799 | 62660107 | -1 | |
| 6.78 | 7.12 | 6.19 | 7.11 | DN1ab | DN1cd | 0.55 | 3.73 | 0.0072 | 0.093 | 0.045 | ENSMUSG00000024219 | protein_coding | Anks1 | ankyrin repeat and SAM domain containing 1 | 224650 | 17 | 27909306 | 28062775 | 1 | Mm.32556 |
| 3.5 | 3.1 | 4.49 | 3.42 | DN1ab | DN1cd | -0.67 | -3.71 | 0.0074 | 0.094 | 0.052 | ENSMUSG00000059708 | protein_coding | Akap17b | A kinase (PRKA) anchor protein 17B | 338351 | X | 36608314 | 36645395 | -1 | Mm.116704 |
| 5.04 | 4.7 | 5.88 | 4.89 | DN1ab | DN1cd | -0.6 | -3.68 | 0.0078 | 0.096 | 0.038 | ENSMUSG00000049871 | protein_coding | Nlrc3 | NLR family, CARD domain containing 3 | 268857 | 16 | 3945007 | 3976632 | -1 | Mm.215994 |
| 0.23 | -3.01 | 0.75 | -0.13 | DN1ab | DN1cd | -2.17 | -3.65 | 0.0081 | 0.098 | 0.031 | ENSMUSG00000083107 | processed_pseudogene | Gm5303 | predicted gene 5303 | NA | 6 | 28361914 | 28363165 | -1 | |
| -1.92 | -3.86 | 2.39 | -2.85 | DN1ab | DN1cd | -3.19 | -3.65 | 0.0081 | 0.099 | 0.055 | ENSMUSG00000015879 | protein_coding | Fam184b | family with sequence similarity 184, member B | 58227 | 5 | 45529705 | 45639614 | -1 | Mm.266466 Mm.402857 |
| 0.99 | 2.69 | -0.6 | 3.75 | DN1ab | DN1cd | 2.57 | 3.62 | 0.0084 | 0.1 | 0.072 | ENSMUSG00000030283 | protein_coding | St8sia1 | ST8 alpha-N-acetyl-neuraminide alpha-2,8-sialyltransferase 1 | 20449 | 6 | 142821545 | 142964452 | -1 | Mm.260838 |
| 3.62 | 3.1 | 5.76 | 4.03 | DN1ab | DN1cd | -1.06 | -3.61 | 0.0085 | 0.1 | 0.025 | ENSMUSG00000041390 | protein_coding | Mdfic | MyoD family inhibitor domain containing | 16543 | 6 | 15720661 | 15802169 | 1 | Mm.1314 |
| 0.6 | 1.74 | -0.24 | 2.96 | DN1ab | DN1cd | 1.87 | 3.58 | 0.0089 | 0.1 | 0.046 | ENSMUSG00000032679 | protein_coding | Cd59a | CD59a antigen | 12509 | 2 | 104095801 | 104115354 | 1 | Mm.247265 |
| 7.29 | 7.52 | 7.23 | 7.88 | DN1ab | DN1cd | 0.38 | 3.56 | 0.0091 | 0.1 | 0.044 | ENSMUSG00000002028 | protein_coding | Kmt2a | lysine (K)-specific methyltransferase 2A | 214162 | 9 | 44803355 | 44881296 | -1 | Mm.2389 |
| -1.01 | 0.51 | -2.81 | 1.88 | DN1ab | DN1cd | 2.61 | 3.56 | 0.0091 | 0.1 | 0.023 | ENSMUSG00000097429 | lincRNA | Gm26520 | predicted gene, 26520 | NA | 12 | 3343816 | 3357153 | -1 | |
| 6.12 | 5.88 | 6.72 | 5.94 | DN1ab | DN1cd | -0.45 | -3.54 | 0.0093 | 0.11 | 0.021 | ENSMUSG00000024038 | protein_coding | Ndufv3 | NADH:ubiquinone oxidoreductase core subunit V3 | 78330 | 17 | 31520115 | 31531326 | 1 | Mm.28349 |
| 4.98 | 5.23 | 4.73 | 5.51 | DN1ab | DN1cd | 0.44 | 3.53 | 0.0095 | 0.11 | 0.02 | ENSMUSG00000018425 | protein_coding | Dhx40 | DEAH (Asp-Glu-Ala-His) box polypeptide 40 | 67487 | 11 | 86768846 | 86807746 | -1 | Mm.260627 |
| 5 | 4.83 | 5.68 | 5.11 | DN1ab | DN1cd | -0.33 | -3.52 | 0.0096 | 0.11 | 0.018 | ENSMUSG00000020522 | protein_coding | Mfap3 | microfibrillar-associated protein 3 | 216760 | 11 | 57518664 | 57533815 | 1 | Mm.30501 |
| 1.54 | 2.15 | 0.54 | 2.45 | DN1ab | DN1cd | 1.07 | 3.52 | 0.0096 | 0.11 | 0.025 | ENSMUSG00000034987 | protein_coding | Hrh2 | histamine receptor H2 | 15466 | 13 | 54192129 | 54236180 | 1 | Mm.57243 |
| 3.07 | 4.16 | 1.55 | 4.61 | DN1ab | DN1cd | 1.76 | 3.51 | 0.0098 | 0.11 | 0.05 | ENSMUSG00000033542 | protein_coding | Arhgef5 | Rho guanine nucleotide exchange factor (GEF) 5 | 54324 | 6 | 43265582 | 43289320 | 1 | Mm.320978 |
| 1.97 | 2.76 | 0.41 | 2.66 | DN1ab | DN1cd | 1.29 | 3.51 | 0.0098 | 0.11 | 0.047 | ENSMUSG00000022793 | protein_coding | B4galt4 | UDP-Gal:betaGlcNAc beta 1,4-galactosyltransferase, polypeptide 4 | 56375 | 16 | 38742264 | 38769049 | 1 | Mm.182377 |
| 12.6 | 12.4 | 13.14 | 12.55 | DN1ab | DN1cd | -0.35 | -3.49 | 0.01 | 0.11 | 0.043 | ENSMUSG00000029580 | protein_coding | Actb | actin, beta | 11461 | 5 | 142903115 | 142906754 | -1 | Mm.328431 Mm.391967 Mm.469717 |
| 5.4 | 5.63 | 5.04 | 5.81 | DN1ab | DN1cd | 0.43 | 3.48 | 0.01 | 0.11 | 0.018 | ENSMUSG00000004105 | protein_coding | Angptl2 | angiopoietin-like 2 | 26360 | 2 | 33216069 | 33247717 | 1 | Mm.208919 |
| 0.92 | 1.69 | -0.36 | 2.02 | DN1ab | DN1cd | 1.33 | 3.47 | 0.01 | 0.11 | 0.027 | ENSMUSG00000037759 | protein_coding | Ptger2 | prostaglandin E receptor 2 (subtype EP2) | 19217 | 14 | 44988195 | 45003820 | 1 | Mm.4630 |
| 3.82 | 4.08 | 3.86 | 4.64 | DN1ab | DN1cd | 0.46 | 3.46 | 0.01 | 0.11 | 0.04 | ENSMUSG00000028093 | protein_coding | Acp6 | acid phosphatase 6, lysophosphatidic | 66659 | 3 | 97158777 | 97177299 | 1 | Mm.101368 |
| 3 | 3.33 | 2.48 | 3.45 | DN1ab | DN1cd | 0.56 | 3.45 | 0.011 | 0.11 | 0.038 | ENSMUSG00000050147 | protein_coding | F2rl3 | coagulation factor II (thrombin) receptor-like 3 | 14065 | 8 | 72761880 | 72763874 | 1 | Mm.12948 |
| 5.18 | 5.95 | 3.6 | 6.17 | DN1ab | DN1cd | 1.4 | 3.44 | 0.011 | 0.11 | 0.017 | ENSMUSG00000032086 | protein_coding | Bace1 | beta-site APP cleaving enzyme 1 | 23821 | 9 | 45838580 | 45864399 | 1 | Mm.24044 |
| 4.06 | 4.45 | 3.34 | 4.49 | DN1ab | DN1cd | 0.67 | 3.44 | 0.011 | 0.11 | 0.042 | ENSMUSG00000052446 | protein_coding | Zfp961 | zinc finger protein 961 | 234413 | 8 | 71951038 | 71993792 | 1 | Mm.334607 |
| 3.86 | 4.34 | 2.95 | 4.37 | DN1ab | DN1cd | 0.82 | 3.41 | 0.011 | 0.12 | 0.048 | ENSMUSG00000014075 | protein_coding | Tctex1d2 | Tctex1 domain containing 2 | 66061 | 16 | 32419702 | 32429099 | 1 | Mm.302693 |
| 4.42 | 4.16 | 4.74 | 3.96 | DN1ab | DN1cd | -0.45 | -3.4 | 0.011 | 0.12 | 0.04 | ENSMUSG00000047777 | protein_coding | Phf13 | PHD finger protein 13 | 230936 | 4 | 151989633 | 151996258 | -1 | Mm.25582 |
| 2.35 | 4 | -0.31 | 4.5 | DN1ab | DN1cd | 2.67 | 3.4 | 0.011 | 0.12 | 0.047 | ENSMUSG00000050108 | protein_coding | Bpifc | BPI fold containing family C | 270757 | 10 | 85959340 | 86011895 | -1 | Mm.107214 |
| 1.19 | 0.78 | 3.02 | 1.5 | DN1ab | DN1cd | -0.9 | -3.4 | 0.011 | 0.12 | 0.019 | ENSMUSG00000001270 | protein_coding | Ckb | creatine kinase, brain | 12709 | 12 | 111669361 | 111672338 | -1 | Mm.16831 |
| -3.14 | -1.93 | -3.94 | -0.05 | DN1ab | DN1cd | 2.25 | 3.39 | 0.011 | 0.12 | 0.033 | ENSMUSG00000086745 | antisense | Tspan2os | tetraspanin 2, opposite strand | NA | 3 | 102720231 | 102735417 | -1 | |
| 9.61 | 9.8 | 8.87 | 9.5 | DN1ab | DN1cd | 0.35 | 3.37 | 0.012 | 0.12 | 0.032 | ENSMUSG00000062461 | processed_pseudogene | Gm5453 | predicted gene 5453 | NA | 13 | 98904224 | 98904670 | 1 | |
| 2.8 | 1.54 | 3.1 | 2.82 | DN1ab | DN1cd | -0.79 | -3.37 | 0.012 | 0.12 | 0.021 | ENSMUSG00000026544 | protein_coding | Dusp23 | dual specificity phosphatase 23 | 68440 | 1 | 172630764 | 172632962 | -1 | Mm.33024 |
| 5.5 | 5.81 | 5.03 | 6.09 | DN1ab | DN1cd | 0.58 | 3.35 | 0.012 | 0.12 | 0.016 | ENSMUSG00000026116 | protein_coding | Tmem131 | transmembrane protein 131 | 56030 | 1 | 36792191 | 36943666 | -1 | Mm.186943 |
| 9.5 | 9.41 | 9.66 | 9.35 | DN1ab | DN1cd | -0.17 | -3.32 | 0.013 | 0.13 | 0.017 | ENSMUSG00000028788 | protein_coding | Ptp4a2 | protein tyrosine phosphatase 4a2 | 19244 | 4 | 129811219 | 129850003 | 1 | Mm.193688 |
| -1.14 | 0.07 | -3.94 | 0.23 | DN1ab | DN1cd | 2.2 | 3.3 | 0.013 | 0.13 | 0.018 | ENSMUSG00000104034 | TEC | 2900092N22Rik | RIKEN cDNA 2900092N22 gene | NA | 2 | 14673404 | 14674812 | 1 | |
| 4.85 | 5.86 | 3.16 | 6.59 | DN1ab | DN1cd | 1.85 | 3.26 | 0.014 | 0.13 | 0.025 | ENSMUSG00000060063 | protein_coding | Alox5ap | arachidonate 5-lipoxygenase activating protein | 11690 | 5 | 149264767 | 149288153 | 1 | Mm.19844 |
| -3.82 | -1.14 | -1.14 | -0.89 | DN1ab | DN1cd | 1.8 | 3.25 | 0.014 | 0.13 | 0.016 | ENSMUSG00000111090 | TEC | Gm30698 | predicted gene, 30698 | NA | 10 | 29362148 | 29365345 | 1 | |
| 10.02 | 9.66 | 9.96 | 9.91 | DN1ab | DN1cd | -0.24 | -3.25 | 0.014 | 0.13 | 0.02 | ENSMUSG00000025203 | protein_coding | Scd2 | stearoyl-Coenzyme A desaturase 2 | 20250 | 19 | 44293676 | 44306864 | 1 | Mm.487021 |
| 4.61 | 4.8 | 4.49 | 5.22 | DN1ab | DN1cd | 0.4 | 3.23 | 0.014 | 0.13 | 0.018 | ENSMUSG00000027452 | protein_coding | Acss1 | acyl-CoA synthetase short-chain family member 1 | 68738 | 2 | 150618105 | 150668500 | -1 | Mm.7044 |
| 2.28 | 3.78 | -0.44 | 4.2 | DN1ab | DN1cd | 2.51 | 3.21 | 0.015 | 0.14 | 0.046 | ENSMUSG00000064147 | protein_coding | Rab44 | RAB44, member RAS oncogene family | 442827 | 17 | 29114145 | 29148980 | 1 | Mm.440341 |
| 3.6 | 4.17 | 2.67 | 4.61 | DN1ab | DN1cd | 1.06 | 3.18 | 0.015 | 0.14 | 0.034 | ENSMUSG00000044252 | protein_coding | Osbpl1a | oxysterol binding protein-like 1A | 64291 | 18 | 12755314 | 12941841 | -1 | Mm.259470 |
| 1.85 | 1.2 | 2.7 | 0.41 | DN1ab | DN1cd | -1.27 | -3.17 | 0.016 | 0.14 | 0.03 | ENSMUSG00000069266 | protein_coding | Hist1h4b | histone cluster 1, H4b | 326620 | 13 | 23757013 | 23757409 | 1 | Mm.261642 |
| -1.01 | -0.1 | -2.28 | 0.88 | DN1ab | DN1cd | 1.71 | 3.16 | 0.016 | 0.14 | 0.032 | ENSMUSG00000026204 | protein_coding | Ptprn | protein tyrosine phosphatase, receptor type, N | 19275 | 1 | 75247027 | 75264502 | -1 | Mm.2902 |
| 6.97 | 6.77 | 8.47 | 7.69 | DN1ab | DN1cd | -0.43 | -3.15 | 0.016 | 0.14 | 0.016 | ENSMUSG00000020614 | protein_coding | Fam20a | family with sequence similarity 20, member A | 208659 | 11 | 109669749 | 109722279 | -1 | Mm.208662 |
| 5.23 | 5.37 | 4.61 | 5.1 | DN1ab | DN1cd | 0.26 | 3.15 | 0.016 | 0.14 | 0.035 | ENSMUSG00000005899 | protein_coding | Smpd4 | sphingomyelin phosphodiesterase 4 | 77626 | 16 | 17619354 | 17644828 | 1 | Mm.34659 |
| 7.73 | 7.08 | 7.68 | 7.62 | DN1ab | DN1cd | -0.42 | -3.14 | 0.016 | 0.14 | 0.016 | ENSMUSG00000048490 | protein_coding | Nrip1 | nuclear receptor interacting protein 1 | 268903 | 16 | 76287400 | 76373827 | -1 | Mm.74711 Mm.447018 Mm.455873 |
| 1.73 | 3.05 | -1.66 | 2.95 | DN1ab | DN1cd | 2.37 | 3.13 | 0.017 | 0.15 | 0.026 | ENSMUSG00000057914 | protein_coding | Cacnb2 | calcium channel, voltage-dependent, beta 2 subunit | 12296 | 2 | 14603088 | 14987908 | 1 | Mm.313930 |
| 4.75 | 5.18 | 4.59 | 4.61 | DN1ab | DN1cd | 0.29 | 3.11 | 0.017 | 0.15 | 0.016 | ENSMUSG00000010914 | protein_coding | Pdhx | pyruvate dehydrogenase complex, component X | 27402 | 2 | 103021075 | 103073513 | -1 | Mm.315011 |
| 4.65 | 4.98 | 4.42 | 5.7 | DN1ab | DN1cd | 0.68 | 3.08 | 0.018 | 0.15 | 0.019 | ENSMUSG00000042745 | protein_coding | Id1 | inhibitor of DNA binding 1 | 15901 | 2 | 152736251 | 152737410 | 1 | Mm.444 |
| 1.42 | 2.46 | -0.29 | 3.88 | DN1ab | DN1cd | 2.12 | 3.02 | 0.019 | 0.16 | 0.016 | ENSMUSG00000048612 | protein_coding | Myof | myoferlin | 226101 | 19 | 37899036 | 38043577 | -1 | Mm.34674 |
| 2.5 | 2.79 | 2.19 | 3.34 | DN1ab | DN1cd | 0.61 | 3.02 | 0.019 | 0.16 | 0.021 | ENSMUSG00000052539 | protein_coding | Magi3 | membrane associated guanylate kinase, WW and PDZ domain containing 3 | 99470 | 3 | 104013259 | 104220374 | -1 | Mm.264849 |
| 4.39 | 4.65 | 4.02 | 5.08 | DN1ab | DN1cd | 0.55 | 3.01 | 0.019 | 0.16 | 0.017 | ENSMUSG00000039153 | protein_coding | Runx2 | runt related transcription factor 2 | 12393 | 17 | 44495987 | 44814797 | -1 | Mm.391013 Mm.391017 Mm.491666 |
| -2.91 | -3.86 | -0.39 | -4.02 | DN1ab | DN1cd | -1.91 | -3 | 0.02 | 0.16 | 0.026 | ENSMUSG00000096037 | rRNA | n-R5s136 | nuclear encoded rRNA 5S 136 | NA | 8 | 123599861 | 123599979 | -1 | |
| -2.91 | -3.86 | -0.19 | -4.02 | DN1ab | DN1cd | -1.99 | -2.99 | 0.02 | 0.16 | 0.019 | ENSMUSG00000108090 | processed_pseudogene | Gm6637 | predicted gene 6637 | NA | 6 | 122295664 | 122296045 | 1 | |
| 5.52 | 5.85 | 5 | 6.39 | DN1ab | DN1cd | 0.72 | 2.89 | 0.023 | 0.18 | 0.024 | ENSMUSG00000051124 | protein_coding | Gimap9 | GTPase, IMAP family member 9 | 317758 | 6 | 48676129 | 48679114 | 1 | Mm.116971 |
| 6.75 | 6.88 | 6.8 | 7.38 | DN1ab | DN1cd | 0.3 | 2.85 | 0.024 | 0.18 | 0.021 | ENSMUSG00000050965 | protein_coding | Prkca | protein kinase C, alpha | 18750 | 11 | 107933387 | 108343928 | -1 | Mm.222178 |
| 6.58 | 6.39 | 7.46 | 6.56 | DN1ab | DN1cd | -0.46 | -2.76 | 0.028 | 0.19 | 0.018 | ENSMUSG00000004319 | protein_coding | Clcn3 | chloride channel, voltage-sensitive 3 | 12725 | 8 | 60910389 | 60983300 | -1 | Mm.25263 Mm.259751 |
| -3.14 | -3.86 | -0.25 | -4.02 | DN1ab | DN1cd | -1.82 | -2.63 | 0.034 | 0.21 | 0.016 | ENSMUSG00000107676 | TEC | Gm44178 | predicted gene, 44178 | NA | 6 | 88919708 | 88922637 | 1 | |
| 4.73 | 4.88 | 3.87 | 4.72 | DN1ab | DN1cd | 0.41 | 2.58 | 0.037 | 0.22 | 0.016 | ENSMUSG00000030200 | protein_coding | Bcl2l14 | BCL2-like 14 (apoptosis facilitator) | 66813 | 6 | 134396318 | 134438736 | 1 | Mm.474472 |
| 4.91 | 6.34 | 3.74 | 8.15 | DN1ab_ltr | DN1cd | 2.48 | 4.54 | 0.0026 | 0.051 | 1.10E-05 | ENSMUSG00000025014 | protein_coding | Dntt | deoxynucleotidyltransferase, terminal | 21673 | 19 | 41029275 | 41059523 | 1 | Mm.25620 |
| 6.75 | 4.41 | 7.08 | 6.25 | DN1ab_ltr | DN1cd | -1.62 | -5.61 | 0.00078 | 0.029 | 1.40E-05 | ENSMUSG00000000682 | protein_coding | Cd52 | CD52 antigen | 23833 | 4 | 134082448 | 134095082 | -1 | Mm.24130 |
| 5.27 | 5.56 | 4.41 | 6.52 | DN1ab_ltr | DN1cd | 0.96 | 2.84 | 0.025 | 0.18 | 2.90E-05 | ENSMUSG00000022010 | protein_coding | Tsc22d1 | TSC22 domain family, member 1 | 21807 | 14 | 76414961 | 76507765 | 1 | Mm.153272 |
| 5.27 | 6.44 | 4.58 | 8.1 | DN1ab_ltr | DN1cd | 2.02 | 4.62 | 0.0024 | 0.049 | 2.90E-05 | ENSMUSG00000000031 | lincRNA | H19 | H19, imprinted maternally expressed transcript | NA | 7 | 142575529 | 142578143 | -1 | |
| 6.02 | 4.95 | 5.62 | 6.03 | DN1ab_ltr | DN1cd | -0.46 | -1.62 | 0.15 | 0.47 | 3.10E-05 | ENSMUSG00000064373 | protein_coding | Selenop | selenoprotein P | 20363 | 15 | 3268547 | 3280508 | 1 | Mm.392203 |
| -3.82 | -3.86 | 1.82 | -4.02 | DN1ab_ltr | DN1cd | -2.13 | -1.97 | 0.089 | 0.36 | 3.80E-05 | ENSMUSG00000037463 | protein_coding | Fbxo27 | F-box protein 27 | 233040 | 7 | 28692849 | 28699338 | 1 | Mm.116116 |
| 3.19 | 4.01 | 2.77 | 5.49 | DN1ab_ltr | DN1cd | 1.51 | 4.27 | 0.0036 | 0.062 | 4.20E-05 | ENSMUSG00000032093 | protein_coding | Cd3e | CD3 antigen, epsilon polypeptide | 12501 | 9 | 44998740 | 45009627 | -1 | Mm.210361 |
| 2.68 | 3.96 | 0.75 | 5.43 | DN1ab_ltr | DN1cd | 2.43 | 3.9 | 0.0058 | 0.082 | 4.60E-05 | ENSMUSG00000034634 | protein_coding | Ly6d | lymphocyte antigen 6 complex, locus D | 17068 | 15 | 74762056 | 74763620 | -1 | Mm.878 |
| 7.16 | 6.05 | 6.53 | 8.06 | DN1ab_ltr | DN1cd | -0.02 | -0.04 | 0.97 | 0.99 | 4.60E-05 | ENSMUSG00000007872 | protein_coding | Id3 | inhibitor of DNA binding 3 | 15903 | 4 | 136143497 | 136145755 | 1 | Mm.110 |
| 6.85 | 6.53 | 6.56 | 7.25 | DN1ab_ltr | DN1cd | 0.06 | 0.33 | 0.75 | 0.93 | 5.20E-05 | ENSMUSG00000037685 | protein_coding | Atp8a1 | ATPase, aminophospholipid transporter (APLT), class I, type 8A, member 1 | 11980 | 5 | 67618140 | 67847434 | -1 | Mm.153230 |
| -3.82 | -3.86 | 1.54 | -4.02 | DN1ab_ltr | DN1cd | -2.02 | -1.96 | 0.09 | 0.36 | 7.50E-05 | ENSMUSG00000022286 | protein_coding | Grhl2 | grainyhead like transcription factor 2 | 252973 | 15 | 37233036 | 37363569 | 1 | Mm.244612 |
| 3.68 | 2.88 | 8.34 | 3.33 | DN1ab_ltr | DN1cd | -2.79 | -3.39 | 0.011 | 0.12 | 9.90E-05 | ENSMUSG00000033577 | protein_coding | Myo6 | myosin VI | 17920 | 9 | 80165031 | 80311729 | 1 | Mm.4040 |
| -3.82 | -3.86 | 1.13 | -4.02 | DN1ab_ltr | DN1cd | -1.88 | -1.95 | 0.092 | 0.37 | 0.00012 | ENSMUSG00000021662 | protein_coding | Arhgef28 | Rho guanine nucleotide exchange factor (GEF) 28 | 110596 | 13 | 97899469 | 98206439 | -1 | Mm.252718 |
| 5.36 | 5.68 | 7.87 | 6.01 | DN1ab_ltr | DN1cd | -0.52 | -1.25 | 0.25 | 0.59 | 0.00012 | ENSMUSG00000030530 | protein_coding | Furin | furin (paired basic amino acid cleaving enzyme) | 18550 | 7 | 80388585 | 80405436 | -1 | Mm.5241 |
| -1.5 | -3.86 | -3.94 | -1.7 | DN1ab_ltr | DN1cd | -0.71 | -0.81 | 0.44 | 0.77 | 0.00015 | ENSMUSG00000025934 | protein_coding | Gsta3 | glutathione S-transferase, alpha 3 | 14859 | 1 | 21240589 | 21265661 | 1 | Mm.394593 |
| 2.25 | 1.34 | 1.9 | 3.96 | DN1ab_ltr | DN1cd | 0.48 | 0.82 | 0.44 | 0.77 | 0.00015 | ENSMUSG00000016194 | protein_coding | Hsd11b1 | hydroxysteroid 11-beta dehydrogenase 1 | 15483 | 1 | 193221634 | 193264075 | -1 | Mm.28328 |
| 4.99 | 5.42 | 5.57 | 4.88 | DN1ab_ltr | DN1cd | 0.03 | 0.13 | 0.9 | 0.96 | 0.00016 | ENSMUSG00000036499 | protein_coding | Eea1 | early endosome antigen 1 | 216238 | 10 | 95940650 | 96045518 | 1 | Mm.210035 Mm.490373 |
| -1.29 | -3.86 | -3.94 | -1.7 | DN1ab_ltr | DN1cd | -0.84 | -0.92 | 0.39 | 0.72 | 0.00017 | ENSMUSG00000024245 | protein_coding | Tmem178 | transmembrane protein 178 | 68027 | 17 | 80944632 | 81001816 | 1 | Mm.357108 Mm.474997 |
| -3.82 | -1.28 | 1.41 | -2.44 | DN1ab_ltr | DN1cd | 0.26 | 0.22 | 0.84 | 0.93 | 0.00018 | ENSMUSG00000036172 | protein_coding | Cd200r3 | CD200 receptor 3 | 74603 | 16 | 44943678 | 44981380 | 1 | Mm.297337 |
| -1.81 | -3.86 | -3.28 | 1.47 | DN1ab_ltr | DN1cd | 0.66 | 0.5 | 0.63 | 0.89 | 0.00018 | ENSMUSG00000086291 | sense_intronic | Gm15513 | predicted gene 15513 | NA | 5 | 34211810 | 34213802 | -1 | |
| 6.84 | 6.65 | 7.87 | 6.61 | DN1ab_ltr | DN1cd | -0.59 | -2.88 | 0.023 | 0.18 | 0.00019 | ENSMUSG00000079184 | protein_coding | Mphosph8 | M-phase phosphoprotein 8 | 75339 | 14 | 56668248 | 56697430 | 1 | Mm.152466 |
| 1.48 | 1.82 | 6.64 | 2.85 | DN1ab_ltr | DN1cd | -1.54 | -1.9 | 0.1 | 0.39 | 0.00019 | ENSMUSG00000028965 | protein_coding | Tnfrsf9 | tumor necrosis factor receptor superfamily, member 9 | 21942 | 4 | 150914562 | 150946102 | 1 | Mm.244187 |
| 7.26 | 7.19 | 7.04 | 7.84 | DN1ab_ltr | DN1cd | 0.26 | 1.58 | 0.16 | 0.49 | 0.00019 | ENSMUSG00000039168 | protein_coding | Dap | death-associated protein | 223453 | 15 | 31224314 | 31274341 | 1 | Mm.222867 |
| 5.86 | 4.75 | 3.96 | 5.81 | DN1ab_ltr | DN1cd | 0.09 | 0.16 | 0.87 | 0.95 | 0.00021 | ENSMUSG00000016494 | protein_coding | Cd34 | CD34 antigen | 12490 | 1 | 194938819 | 194961279 | 1 | Mm.29798 |
| 1.75 | 1.71 | 5.26 | 1.84 | DN1ab_ltr | DN1cd | -1.46 | -2.21 | 0.063 | 0.3 | 0.00021 | ENSMUSG00000025161 | protein_coding | Slc16a3 | solute carrier family 16 (monocarboxylic acid transporters), member 3 | 80879 | 11 | 120948480 | 120960868 | 1 | Mm.28632 |
| -3.82 | -3.86 | 0.49 | -4.02 | DN1ab_ltr | DN1cd | -1.65 | -1.94 | 0.094 | 0.37 | 0.00022 | ENSMUSG00000040035 | protein_coding | Disp2 | dispatched RND tramsporter family member 2 | 214240 | 2 | 118779719 | 118811293 | 1 | Mm.221499 |
| 4.07 | 4.86 | 5.65 | 3.9 | DN1ab_ltr | DN1cd | -0.09 | -0.19 | 0.86 | 0.94 | 0.00023 | ENSMUSG00000022180 | protein_coding | Slc7a8 | solute carrier family 7 (cationic amino acid transporter, y+ system), member 8 | 50934 | 14 | 54722209 | 54781946 | -1 | Mm.276831 |
| 4.93 | 5.24 | 6.08 | 5.07 | DN1ab_ltr | DN1cd | -0.17 | -0.7 | 0.51 | 0.82 | 0.00023 | ENSMUSG00000032290 | protein_coding | Ptpn9 | protein tyrosine phosphatase, non-receptor type 9 | 56294 | 9 | 56994923 | 57062807 | 1 | Mm.325643 |
| 8.89 | 9 | 9.6 | 8.75 | DN1ab_ltr | DN1cd | -0.24 | -1.31 | 0.23 | 0.57 | 0.00025 | ENSMUSG00000063229 | protein_coding | Ldha | lactate dehydrogenase A | 16828 | 7 | 46841475 | 46855627 | 1 | Mm.29324 |
| 6.65 | 6.19 | 8.99 | 6.86 | DN1ab_ltr | DN1cd | -1.13 | -3.5 | 0.0099 | 0.11 | 0.00026 | ENSMUSG00000045136 | protein_coding | Tubb2b | tubulin, beta 2B class IIB | 73710 | 13 | 34126748 | 34130466 | -1 | Mm.379227 Mm.472121 |
| 5.45 | 5.43 | 6.92 | 5.41 | DN1ab_ltr | DN1cd | -0.59 | -2.05 | 0.079 | 0.34 | 0.00027 | ENSMUSG00000027639 | protein_coding | Samhd1 | SAM domain and HD domain, 1 | 56045 | 2 | 157097533 | 157135265 | -1 | Mm.248478 Mm.468781 |
| 7.32 | 7.35 | 8.42 | 7.47 | DN1ab_ltr | DN1cd | -0.34 | -1.76 | 0.12 | 0.43 | 0.00027 | ENSMUSG00000030695 | protein_coding | Aldoa | aldolase A, fructose-bisphosphate | 11674 | 7 | 126795234 | 126800751 | -1 | Mm.275831 |
| 8.53 | 8.61 | 10 | 9.2 | DN1ab_ltr | DN1cd | -0.26 | -1.52 | 0.17 | 0.5 | 3.00E-04 | ENSMUSG00000076749 | TR_C_gene | Tcrg-C1 | T cell receptor gamma, constant 1 | NA | 13 | 19214103 | 19217024 | 1 | |
| 6 | 5.78 | 6.59 | 5.52 | DN1ab_ltr | DN1cd | -0.53 | -3.26 | 0.014 | 0.13 | 0.00032 | ENSMUSG00000033629 | protein_coding | Hacd3 | 3-hydroxyacyl-CoA dehydratase 3 | 57874 | 9 | 64986983 | 65021693 | -1 | Mm.308180 |
| 6.44 | 5.93 | 6.83 | 7.1 | DN1ab_ltr | DN1cd | -0.2 | -1.29 | 0.24 | 0.58 | 0.00032 | ENSMUSG00000019820 | protein_coding | Utrn | utrophin | 22288 | 10 | 12382188 | 12869365 | -1 | Mm.331784 Mm.418026 |
| -1.29 | -3.86 | -3.49 | -0.25 | DN1ab_ltr | DN1cd | -0.37 | -0.33 | 0.75 | 0.93 | 0.00034 | ENSMUSG00000087336 | antisense | Gm15860 | predicted gene 15860 | NA | 5 | 123332726 | 123344437 | -1 | |
| -0.86 | -3.86 | -2.15 | -0.71 | DN1ab_ltr | DN1cd | -1.33 | -1.55 | 0.17 | 0.5 | 0.00035 | ENSMUSG00000049709 | protein_coding | Nlrp10 | NLR family, pyrin domain containing 10 | 244202 | 7 | 108921852 | 108930178 | -1 | Mm.37991 |
| 5 | 3.85 | 7.02 | 4.42 | DN1ab_ltr | DN1cd | -1.79 | -6.24 | 0.00042 | 0.021 | 0.00039 | ENSMUSG00000019256 | protein_coding | Ahr | aryl-hydrocarbon receptor | 11622 | 12 | 35497974 | 35535038 | -1 | Mm.341377 |
| -3.82 | -3.86 | 0.72 | -4.02 | DN1ab_ltr | DN1cd | -1.73 | -1.92 | 0.097 | 0.38 | 4.00E-04 | ENSMUSG00000025468 | protein_coding | Caly | calcyon neuron-specific vesicular protein | 68566 | 7 | 140069880 | 140082548 | -1 | Mm.44241 |
| -1.22 | -3.86 | -3.94 | -2.44 | DN1ab_ltr | DN1cd | -1.15 | -1.45 | 0.19 | 0.53 | 4.00E-04 | ENSMUSG00000106813 | processed_pseudogene | Gm7181 | predicted gene 7181 | NA | 5 | 42499981 | 42500414 | -1 | |
| 3.56 | 0.68 | 3.14 | 2.11 | DN1ab_ltr | DN1cd | -1.83 | -5.01 | 0.0015 | 0.039 | 0.00041 | ENSMUSG00000030532 | protein_coding | Hddc3 | HD domain containing 3 | 68695 | 7 | 80343104 | 80350547 | 1 | Mm.21171 |
| 7.87 | 8.11 | 7.2 | 8.41 | DN1ab_ltr | DN1cd | 0.6 | 3.17 | 0.015 | 0.14 | 0.00043 | ENSMUSG00000047139 | protein_coding | Cd24a | CD24a antigen | 12484 | 10 | 43578284 | 43584265 | 1 | Mm.29742 |
| 8.72 | 8.83 | 8.36 | 9.11 | DN1ab_ltr | DN1cd | 0.35 | 2.81 | 0.026 | 0.18 | 0.00044 | ENSMUSG00000015937 | protein_coding | H2afy | H2A histone family, member Y | 26914 | 13 | 56073619 | 56136361 | -1 | Mm.283802 Mm.478369 |
| 3.44 | 3.41 | 5.41 | 3.87 | DN1ab_ltr | DN1cd | -0.64 | -2.17 | 0.066 | 0.31 | 0.00044 | ENSMUSG00000055116 | protein_coding | Arntl | aryl hydrocarbon receptor nuclear translocator-like | 11865 | 7 | 113207465 | 113314126 | 1 | Mm.33970 Mm.440371 |
| 3.69 | 5.04 | 5.79 | 5.41 | DN1ab_ltr | DN1cd | 0.73 | 2.19 | 0.064 | 0.3 | 0.00046 | ENSMUSG00000003882 | protein_coding | Il7r | interleukin 7 receptor | 16197 | 15 | 9505788 | 9530176 | -1 | Mm.389 |
| 6.06 | 5.74 | 7.19 | 5.92 | DN1ab_ltr | DN1cd | -0.68 | -3.72 | 0.0074 | 0.093 | 0.00046 | ENSMUSG00000019951 | protein_coding | Uhrf1bp1l | UHRF1 (ICBP90) binding protein 1-like | 75089 | 10 | 89744991 | 89819871 | 1 | Mm.268922 Mm.446134 |
| -3.82 | -3.86 | -0.24 | -4.02 | DN1ab_ltr | DN1cd | -1.38 | -1.92 | 0.096 | 0.38 | 0.00046 | ENSMUSG00000047562 | protein_coding | Mmp10 | matrix metallopeptidase 10 | 17384 | 9 | 7502352 | 7510240 | 1 | Mm.14126 |
| -3.82 | -1.88 | -0.88 | -4.02 | DN1ab_ltr | DN1cd | 0.09 | 0.09 | 0.93 | 0.97 | 0.00048 | ENSMUSG00000106106 | rRNA | | 18s RNA, related sequence 5 | NA | 17 | 39846353 | 39848827 | -1 | |
| 6.82 | 6.95 | 6.45 | 7.27 | DN1ab_ltr | DN1cd | 0.39 | 2.94 | 0.021 | 0.17 | 0.00049 | ENSMUSG00000053477 | protein_coding | Tcf4 | transcription factor 4 | 21413 | 18 | 69343356 | 69689079 | 1 | Mm.4269 |
| 1.84 | 1.22 | 0.17 | 2.4 | DN1ab_ltr | DN1cd | 0.55 | 0.97 | 0.36 | 0.7 | 0.00049 | ENSMUSG00000000440 | protein_coding | Pparg | peroxisome proliferator activated receptor gamma | 19016 | 6 | 115360951 | 115490399 | 1 | Mm.3020 |
| 4.31 | 3.42 | 3.85 | 4.78 | DN1ab_ltr | DN1cd | -0.1 | -0.28 | 0.79 | 0.93 | 5.00E-04 | ENSMUSG00000042797 | protein_coding | Aqp11 | aquaporin 11 | 66333 | 7 | 97724006 | 97738289 | -1 | Mm.29756 |
| 7 | 6.95 | 7.86 | 7.02 | DN1ab_ltr | DN1cd | -0.35 | -2.26 | 0.058 | 0.28 | 0.00052 | ENSMUSG00000023456 | protein_coding | Tpi1 | triosephosphate isomerase 1 | 21991 | 6 | 124810586 | 124814296 | -1 | Mm.4222 Mm.439915 |
| -3.82 | -1.93 | -1.83 | -4.02 | DN1ab_ltr | DN1cd | 0.41 | 0.51 | 0.62 | 0.89 | 0.00053 | ENSMUSG00000034768 | protein_coding | Asb16 | ankyrin repeat and SOCS box-containing 16 | 217217 | 11 | 102268743 | 102279462 | 1 | Mm.216432 |
| -3.82 | -1.28 | -2.36 | -4.02 | DN1ab_ltr | DN1cd | 1.03 | 1.27 | 0.25 | 0.59 | 0.00055 | ENSMUSG00000110386 | lincRNA | Gm42031 | predicted gene, 42031 | NA | 8 | 85125872 | 85132289 | -1 | |
| 1.09 | -0.88 | -2.36 | 0.15 | DN1ab_ltr | DN1cd | -0.04 | -0.05 | 0.96 | 0.98 | 0.00055 | ENSMUSG00000007888 | protein_coding | Crlf1 | cytokine receptor-like factor 1 | 12931 | 8 | 70493158 | 70504081 | 1 | Mm.335584 |
| -3.82 | -2.28 | -1.23 | -4.02 | DN1ab_ltr | DN1cd | -0.04 | -0.04 | 0.97 | 0.99 | 0.00056 | ENSMUSG00000010492 | antisense | Uckl1os | uridine-cytidine kinase 1-like 1, opposite strand | NA | 2 | 181578480 | 181585170 | 1 | |
| -2.24 | -1.06 | -3.94 | 0.78 | DN1ab_ltr | DN1cd | 2.55 | 3.66 | 0.0079 | 0.097 | 0.00056 | ENSMUSG00000112148 | protein_coding | Lilrb4a | leukocyte immunoglobulin-like receptor, subfamily B, member 4A | 14728 | 10 | 51490956 | 51496611 | 1 | Mm.34408 |
| 2.22 | 1.24 | 3.75 | 0.5 | DN1ab_ltr | DN1cd | -1.88 | -4.17 | 0.0041 | 0.066 | 0.00057 | ENSMUSG00000037922 | protein_coding | Bank1 | B cell scaffold protein with ankyrin repeats 1 | 242248 | 3 | 136053363 | 136326066 | -1 | Mm.30832 |
| 3.63 | 2.3 | 2.56 | 2.99 | DN1ab_ltr | DN1cd | -0.55 | -1.57 | 0.16 | 0.49 | 6.00E-04 | ENSMUSG00000044017 | protein_coding | Adgrd1 | adhesion G protein-coupled receptor D1 | 243277 | 5 | 129096750 | 129204599 | 1 | Mm.440501 |
| -3.82 | -3.86 | -3.94 | -0.44 | DN1ab_ltr | DN1cd | 1.3 | 1.87 | 0.1 | 0.39 | 6.00E-04 | ENSMUSG00000060275 | protein_coding | Nrg2 | neuregulin 2 | 100042150 | 18 | 36017707 | 36197380 | -1 | Mm.380390 |
| -0.35 | -3.86 | -3.94 | -1.83 | DN1ab_ltr | DN1cd | -1.47 | -1.35 | 0.22 | 0.56 | 0.00062 | ENSMUSG00000020282 | protein_coding | Rhbdf1 | rhomboid 5 homolog 1 | 13650 | 11 | 32209585 | 32222300 | -1 | Mm.11545 |
| -3.82 | -3.86 | -0.36 | -4.02 | DN1ab_ltr | DN1cd | -1.34 | -1.91 | 0.098 | 0.38 | 0.00063 | ENSMUSG00000111933 | lincRNA | Gm36660 | predicted gene, 36660 | NA | 12 | 80018415 | 80081434 | -1 | |
| 7.26 | 7.15 | 8.2 | 6.43 | DN1ab_ltr | DN1cd | -0.72 | -2.24 | 0.059 | 0.29 | 0.00063 | ENSMUSG00000027533 | protein_coding | Fabp5 | fatty acid binding protein 5, epidermal | 16592 | 3 | 10012548 | 10016607 | 1 | Mm.741 |
| 0.19 | 0.38 | 3.29 | 0.88 | DN1ab_ltr | DN1cd | -0.88 | -1.7 | 0.13 | 0.44 | 0.00064 | ENSMUSG00000039783 | protein_coding | Kmo | kynurenine 3-monooxygenase (kynurenine 3-hydroxylase) | 98256 | 1 | 175620381 | 175662116 | 1 | Mm.27217 |
| 4.85 | 4.61 | 4.07 | 5.87 | DN1ab_ltr | DN1cd | 0.57 | 1.4 | 0.2 | 0.54 | 0.00064 | ENSMUSG00000027778 | protein_coding | Ift80 | intraflagellar transport 80 | 68259 | 3 | 68892499 | 69004570 | -1 | Mm.389451 |
| -1.22 | -3.86 | -3.94 | -2.07 | DN1ab_ltr | DN1cd | -1.02 | -1.17 | 0.28 | 0.62 | 0.00064 | ENSMUSG00000097805 | antisense | Gm17473 | predicted gene, 17473 | NA | 15 | 42676241 | 42704616 | 1 | |
| 5.43 | 4.68 | 8.53 | 5.17 | DN1ab_ltr | DN1cd | -1.86 | -3.57 | 0.0089 | 0.1 | 0.00065 | ENSMUSG00000015437 | protein_coding | Gzmb | granzyme B | 14939 | 14 | 56258838 | 56262260 | -1 | Mm.14874 |
| 1.97 | -0.52 | -0.4 | 1.36 | DN1ab_ltr | DN1cd | -0.35 | -0.4 | 0.7 | 0.92 | 0.00066 | ENSMUSG00000027797 | protein_coding | Dclk1 | doublecortin-like kinase 1 | 13175 | 3 | 55242364 | 55539068 | 1 | Mm.393242 Mm.472264 |
| 2.25 | 3.69 | 0.28 | 3.94 | DN1ab_ltr | DN1cd | 2.16 | 5.15 | 0.0013 | 0.036 | 0.00068 | ENSMUSG00000047904 | protein_coding | Sstr2 | somatostatin receptor 2 | 20606 | 11 | 113619342 | 113626028 | 1 | |
| 8.34 | 7.13 | 8.19 | 7.89 | DN1ab_ltr | DN1cd | -0.85 | -4.71 | 0.0021 | 0.047 | 0.00068 | ENSMUSG00000001865 | protein_coding | Cpa3 | carboxypeptidase A3, mast cell | 12873 | 3 | 20215620 | 20242181 | -1 | Mm.1135 |
| 0.92 | 1.24 | -0.31 | 2.95 | DN1ab_ltr | DN1cd | 1.44 | 2.48 | 0.042 | 0.24 | 7.00E-04 | ENSMUSG00000052914 | protein_coding | Cyp2j6 | cytochrome P450, family 2, subfamily j, polypeptide 6 | 13110 | 4 | 96516138 | 96553661 | -1 | Mm.98200 |
| -3.82 | -1.54 | -1.02 | -2.44 | DN1ab_ltr | DN1cd | 0.95 | 1.32 | 0.23 | 0.57 | 7.00E-04 | ENSMUSG00000097462 | antisense | 9530026P05Rik | RIKEN cDNA 9530026P05 gene | NA | 6 | 92940329 | 93272979 | 1 | |
| 4.32 | 3.93 | 2.93 | 5.25 | DN1ab_ltr | DN1cd | 0.69 | 1.28 | 0.24 | 0.58 | 0.00072 | ENSMUSG00000044265 | protein_coding | Olfm5 | olfactomedin 5 | 244180 | 7 | 104153013 | 104164831 | -1 | Mm.17149 |
| -2.24 | -3.86 | -3.49 | -0.85 | DN1ab_ltr | DN1cd | -0.06 | -0.07 | 0.95 | 0.98 | 0.00073 | ENSMUSG00000062743 | protein_coding | Zfp677 | zinc finger protein 677 | 210503 | 17 | 21383748 | 21399265 | 1 | Mm.33443 |
| -1.92 | -3.86 | -3.94 | -0.77 | DN1ab_ltr | DN1cd | -0.05 | -0.05 | 0.96 | 0.98 | 0.00074 | ENSMUSG00000111617 | TEC | Gm47148 | predicted gene, 47148 | NA | 9 | 122993127 | 122993704 | 1 | |
| 4.23 | 4.78 | 3.32 | 5.52 | DN1ab_ltr | DN1cd | 1.15 | 3.57 | 0.009 | 0.1 | 0.00075 | ENSMUSG00000036908 | protein_coding | Unc93b1 | unc-93 homolog B1, TLR signaling regulator | 54445 | 19 | 3935186 | 3949340 | 1 | Mm.262094 |
| 2.75 | 3.32 | 0.04 | 2.81 | DN1ab_ltr | DN1cd | 1.3 | 3.09 | 0.018 | 0.15 | 0.00076 | ENSMUSG00000049881 | transcribed_processed_pseudogene | 2810025M15Rik | RIKEN cDNA 2810025M15 gene | NA | 1 | 157412352 | 157420236 | 1 | |
| -0.49 | -2.28 | -3.94 | -1.28 | DN1ab_ltr | DN1cd | -0.14 | -0.16 | 0.88 | 0.95 | 0.00076 | ENSMUSG00000015766 | protein_coding | Eps8 | epidermal growth factor receptor pathway substrate 8 | 13860 | 6 | 137477245 | 137654876 | -1 | Mm.235346 Mm.412332 |
| 1.23 | 0.21 | 6.09 | -0.01 | DN1ab_ltr | DN1cd | -3.32 | -3.24 | 0.014 | 0.13 | 0.00076 | ENSMUSG00000018168 | protein_coding | Ikzf3 | IKAROS family zinc finger 3 | 22780 | 11 | 98464902 | 98546031 | -1 | Mm.133367 Mm.459157 |
| 2.23 | 2.86 | 5.12 | 2.82 | DN1ab_ltr | DN1cd | -0.49 | -0.85 | 0.42 | 0.75 | 0.00077 | ENSMUSG00000040183 | protein_coding | Ankrd6 | ankyrin repeat domain 6 | 140577 | 4 | 32804035 | 32950841 | -1 | Mm.214766 |
| 6.01 | 6.35 | 4.96 | 6.04 | DN1ab_ltr | DN1cd | 0.6 | 4.12 | 0.0044 | 0.068 | 0.00078 | ENSMUSG00000037242 | protein_coding | Clic4 | chloride intracellular channel 4 (mitochondrial) | 29876 | 4 | 135213969 | 135272814 | -1 | Mm.257765 |
| 0.16 | -1.54 | -1.14 | 0.86 | DN1ab_ltr | DN1cd | -0.05 | -0.07 | 0.95 | 0.98 | 0.00078 | ENSMUSG00000074896 | protein_coding | Ifit3 | interferon-induced protein with tetratricopeptide repeats 3 | 15959 | 19 | 34583531 | 34588731 | 1 | Mm.426079 |
| -0.65 | 0.13 | 2.62 | 0.88 | DN1ab_ltr | DN1cd | -0.21 | -0.43 | 0.68 | 0.91 | 0.00079 | ENSMUSG00000021675 | protein_coding | F2rl2 | coagulation factor II (thrombin) receptor-like 2 | 14064 | 13 | 95696853 | 95702739 | 1 | Mm.440952 |
| 7.29 | 7.75 | 8.23 | 7.96 | DN1ab_ltr | DN1cd | 0.19 | 1.31 | 0.23 | 0.57 | 0.00079 | ENSMUSG00000032294 | protein_coding | Pkm | pyruvate kinase, muscle | 18746 | 9 | 59656368 | 59679375 | 1 | Mm.326167 Mm.488724 |
| 8.65 | 8.76 | 9.03 | 8.57 | DN1ab_ltr | DN1cd | -0.1 | -0.86 | 0.42 | 0.75 | 8.00E-04 | ENSMUSG00000000420 | protein_coding | Galnt1 | polypeptide N-acetylgalactosaminyltransferase 1 | 14423 | 18 | 24205344 | 24286818 | 1 | Mm.30249 |
| -2.24 | -3.86 | -3.94 | -1.28 | DN1ab_ltr | DN1cd | -0.08 | -0.09 | 0.93 | 0.97 | 0.00089 | ENSMUSG00000030513 | protein_coding | Pcsk6 | proprotein convertase subtilisin/kexin type 6 | 18553 | 7 | 65861734 | 66050386 | 1 | Mm.294007 |
| 0.16 | 0.22 | 0.15 | 2.72 | DN1ab_ltr | DN1cd | 1.09 | 2.17 | 0.067 | 0.31 | 0.00091 | ENSMUSG00000032128 | protein_coding | Robo3 | roundabout guidance receptor 3 | 19649 | 9 | 37415669 | 37433246 | -1 | Mm.212826 |

| | | | | | | | | | | | | | | | | | | | | |
|---|---|---|---|---|---|---|---|---|---|---|---|---|---|---|---|---|---|---|---|---|
| 2.56 | 2.32 | 2.11 | 3.63 | DN1ab_Itr | DN1cd | 0.48 | 1.35 | 0.22 | 0.55 | 0.00092 | ENSMUSG00000026004 | protein_coding | Kansl1l | KAT8 regulatory NSL complex subunit 1-like | 68691 | 1 | 66719248 | 66817562 | -1 | Mm.240912 Mm.383276 Mm.461139 Mm.482481 |
| 7.39 | 7.42 | 7.65 | 6.84 | DN1ab_Itr | DN1cd | -0.27 | -1.65 | 0.14 | 0.46 | 0.00093 | ENSMUSG00000005667 | protein_coding | Mthfd2 | methylenetetrahydrofolate dehydrogenase (NAD+ dependent), methenyltetrahydrofolate cyclohydrolase | 17768 | 6 | 83305691 | 83325908 | -1 | Mm.443 Mm.475021 |
| 3.15 | 3.65 | 2.47 | 4.61 | DN1ab_Itr | DN1cd | 1.11 | 3.43 | 0.011 | 0.12 | 0.00093 | ENSMUSG00000047821 | protein_coding | Trim16 | tripartite motif-containing 16 | 94092 | 11 | 62820231 | 62850808 | 1 | Mm.117087 |
| -3.82 | -3.86 | -0.12 | -4.02 | DN1ab_Itr | DN1cd | -1.43 | -1.89 | 0.1 | 0.39 | 0.00094 | ENSMUSG00000070323 | protein_coding | Mmp27 | matrix metallopeptidase 27 | 234911 | 9 | 7571396 | 7581885 | 1 | Mm.334940 |
| -1.14 | -3.86 | -3.94 | -2.07 | DN1ab_Itr | DN1cd | -1.07 | -1.19 | 0.27 | 0.61 | 0.00094 | ENSMUSG00000105858 | lincRNA | 2310074N15Rik | RIKEN cDNA 2310074N15 gene | NA | 5 | 24447908 | 24448489 | 1 | |
| 3.15 | 2.31 | 2.88 | 4.03 | DN1ab_Itr | DN1cd | 0.06 | 0.15 | 0.89 | 0.95 | 0.00095 | ENSMUSG00000075010 | lincRNA | AW112010 | expressed sequence AW112010 | NA | 19 | 11047617 | 11050566 | -1 | |
| 2.91 | 3.41 | 3.15 | 2.4 | DN1ab_Itr | DN1cd | 0.08 | 0.32 | 0.76 | 0.93 | 0.00095 | ENSMUSG00000097994 | antisense | Gm26982 | predicted gene, 26982 | NA | 6 | 113714971 | 113716121 | 1 | |
| 5.34 | 5.14 | 5.56 | 4.38 | DN1ab_Itr | DN1cd | -0.56 | -2.89 | 0.023 | 0.17 | 0.00098 | ENSMUSG00000042195 | protein_coding | Slc35f2 | solute carrier family 35, member F2 | 72022 | 9 | 53771538 | 53818154 | 1 | Mm.26159 |
| -1.29 | -3.86 | -2.81 | -1.22 | DN1ab_Itr | DN1cd | -1.05 | -1.29 | 0.24 | 0.58 | 0.00099 | ENSMUSG00000028794 | protein_coding | A3galt2 | alpha 1,3-galactosyltransferase 2 (isoglobotriaosylceramide synthase) | 215493 | 4 | 128755364 | 128769298 | 1 | Mm.300900 |
| 4.38 | 4.26 | 5.55 | 4.55 | DN1ab_Itr | DN1cd | -0.46 | -2.63 | 0.034 | 0.21 | 0.001 | ENSMUSG00000024639 | protein_coding | Gnaq | guanine nucleotide binding protein, alpha q polypeptide | 14682 | 19 | 16132831 | 16387463 | 1 | Mm.439701 Mm.441601 |
| 8.71 | 7.97 | 8.38 | 8.87 | DN1ab_Itr | DN1cd | -0.25 | -1.03 | 0.34 | 0.67 | 0.001 | ENSMUSG00000026581 | protein_coding | Sell | selectin, lymphocyte | 20343 | 1 | 164061982 | 164084181 | 1 | Mm.1461 |
| 6.08 | 6.4 | 5.76 | 6.67 | DN1ab_Itr | DN1cd | 0.54 | 4.63 | 0.0023 | 0.049 | 0.001 | ENSMUSG00000024378 | protein_coding | Stard4 | StAR-related lipid transfer (START) domain containing 4 | 170459 | 18 | 33199355 | 33213862 | -1 | Mm.127058 Mm.393277 |
| 5.63 | 5.35 | 6.9 | 5.79 | DN1ab_Itr | DN1cd | -0.6 | -3.61 | 0.0085 | 0.1 | 0.001 | ENSMUSG00000018507 | protein_coding | Trpv2 | transient receptor potential cation channel, subfamily V, member 2 | 22368 | 11 | 62574486 | 62600515 | 1 | Mm.288064 |
| 6.1 | 6.01 | 9.2 | 7.18 | DN1ab_Itr | DN1cd | -0.86 | -2.22 | 0.062 | 0.29 | 0.001 | ENSMUSG00000020644 | protein_coding | Id2 | inhibitor of DNA binding 2 | 15902 | 12 | 25093799 | 25097140 | -1 | Mm.34871 |
| -3.82 | -3.86 | -0.56 | -4.02 | DN1ab_Itr | DN1cd | -1.27 | -1.89 | 0.1 | 0.39 | 0.001 | ENSMUSG00000103170 | lincRNA | Gm37170 | predicted gene, 37170 | NA | 3 | 52198248 | 52200489 | 1 | |
| -3.82 | -3.86 | -0.56 | -4.02 | DN1ab_Itr | DN1cd | -1.27 | -1.89 | 0.1 | 0.39 | 0.001 | ENSMUSG00000028435 | protein_coding | Aqp3 | aquaporin 3 | 11828 | 4 | 41092722 | 41098183 | -1 | Mm.34043 |
| -3.82 | -3.86 | -3.94 | -0.85 | DN1ab_Itr | DN1cd | 1.12 | 1.81 | 0.11 | 0.41 | 0.0011 | ENSMUSG00000054256 | protein_coding | Msi1 | musashi RNA-binding protein 1 | 17690 | 5 | 115429599 | 115455698 | 1 | Mm.5077 Mm.391995 |
| 6.26 | 6.04 | 5.66 | 6.12 | DN1ab_Itr | DN1cd | 0.04 | 0.26 | 0.8 | 0.93 | 0.0011 | ENSMUSG00000015880 | protein_coding | Ncapg | non-SMC condensin I complex, subunit G | 54392 | 5 | 45669919 | 45700546 | 1 | Mm.32012 |
| 6.54 | 6.6 | 7.11 | 6.49 | DN1ab_Itr | DN1cd | -0.19 | -1.45 | 0.19 | 0.53 | 0.0011 | ENSMUSG00000030042 | protein_coding | Pole4 | polymerase (DNA-directed), epsilon 4 (p12 subunit) | 66979 | 6 | 82618992 | 82705365 | -1 | Mm.195753 |
| -3.82 | -3.86 | -0.06 | -4.02 | DN1ab_Itr | DN1cd | -1.45 | -1.88 | 0.1 | 0.39 | 0.0011 | ENSMUSG00000099025 | lincRNA | Gm27162 | predicted gene 27162 | NA | 9 | 32904937 | 32928966 | -1 | |
| 1.42 | 3.61 | -0.57 | 4.72 | DN1ab_Itr | DN1cd | 3.17 | 5.37 | 0.001 | 0.033 | 0.0011 | ENSMUSG00000052730 | processed_transcript | Gm5111 | predicted gene 5111 | 330305 | 6 | 48589445 | 48590584 | 1 | Mm.123868 |
| 1.89 | 2.95 | -0.19 | 2.95 | DN1ab_Itr | DN1cd | 1.75 | 4.36 | 0.0033 | 0.058 | 0.0011 | ENSMUSG00000049807 | protein_coding | Arhgap23 | Rho GTPase activating protein 23 | 58996 | 11 | 97415533 | 97502402 | 1 | Mm.394118 |
| 4.47 | 5.17 | 5.67 | 5.15 | DN1ab_Itr | DN1cd | 0.26 | 1.07 | 0.32 | 0.66 | 0.0011 | ENSMUSG00000024143 | protein_coding | Rhoq | ras homolog family member Q | 104215 | 17 | 86963082 | 87000069 | 1 | Mm.826 |
| -0.09 | -3.12 | -1.27 | 0.23 | DN1ab_Itr | DN1cd | -1.1 | -1.2 | 0.27 | 0.6 | 0.0011 | ENSMUSG00000031150 | protein_coding | Ccdc120 | coiled-coil domain containing 120 | 54648 | X | 7731714 | 7750905 | -1 | Mm.63342 |
| 5.73 | 5.51 | 7.45 | 6.13 | DN1ab_Itr | DN1cd | -0.65 | -2.93 | 0.022 | 0.17 | 0.0012 | ENSMUSG00000059040 | protein_coding | Eno1b | enolase 1B, retrotransposed | 433182 | 18 | 48045335 | 48048378 | 1 | Mm.70666 Mm.372357 |
| 1.44 | 0.66 | 0.08 | 1.9 | DN1ab_Itr | DN1cd | 0.31 | 0.59 | 0.58 | 0.86 | 0.0012 | ENSMUSG00000047669 | protein_coding | Msl3l2 | MSL3 like 2 | 73390 | 10 | 56106917 | 56116880 | 1 | Mm.181475 |
| 0.44 | -1.93 | -0.03 | 0.93 | DN1ab_Itr | DN1cd | -0.78 | -1.14 | 0.29 | 0.63 | 0.0012 | ENSMUSG00000040734 | protein_coding | Ppp1r13l | protein phosphatase 1, regulatory subunit 13 like | 333654 | 7 | 19359749 | 19378533 | 1 | Mm.333640 |
| -1.22 | -3.01 | 2.97 | -4.02 | DN1ab_Itr | DN1cd | -3.89 | -3.71 | 0.0075 | 0.094 | 0.0012 | ENSMUSG00000050010 | protein_coding | Shisa3 | shisa family member 3 | 330096 | 5 | 67607882 | 67623552 | 1 | Mm.442747 |
| 4.65 | 4.47 | 4.65 | 5.48 | DN1ab_Itr | DN1cd | 0.22 | 1.06 | 0.33 | 0.67 | 0.0012 | ENSMUSG00000104339 | TEC | C130089K02Rik | RIKEN cDNA C130089K02 gene | NA | 3 | 52057285 | 52064244 | -1 | |
| 5.29 | 5.22 | 6.67 | 4.81 | DN1ab_Itr | DN1cd | -0.73 | -2.07 | 0.077 | 0.33 | 0.0012 | ENSMUSG00000058173 | protein_coding | Smco4 | single-pass membrane protein with coiled-coil domains 4 | 170748 | 9 | 15493432 | 15545260 | 1 | Mm.86572 |
| 4.77 | 4.51 | 5.33 | 6.23 | DN1ab_Itr | DN1cd | 0.22 | 0.92 | 0.39 | 0.72 | 0.0012 | ENSMUSG00000068744 | protein_coding | Psrc1 | proline/serine-rich coiled-coil 1 | 56742 | 3 | 108383839 | 108388231 | 1 | Mm.389499 |
| -1.01 | 0.38 | -3.94 | 0.37 | DN1ab_Itr | DN1cd | 2.36 | 4.18 | 0.0041 | 0.066 | 0.0012 | ENSMUSG00000094525 | TR_V_gene | Trbv12-2 | T cell receptor beta, variable 12-2 | NA | 6 | 41118851 | 41119364 | 1 | |
| -1.01 | -1.88 | -3.94 | -0.71 | DN1ab_Itr | DN1cd | 0.67 | 0.82 | 0.44 | 0.77 | 0.0013 | ENSMUSG00000080941 | processed_pseudogene | Gm16056 | predicted gene 16056 | NA | 5 | 26859967 | 26860101 | -1 | |
| 5.59 | 5.35 | 6.83 | 5.13 | DN1ab_Itr | DN1cd | -0.79 | -2.72 | 0.029 | 0.2 | 0.0013 | ENSMUSG00000021831 | protein_coding | Ero1l | ERO1-like (S. cerevisiae) | 50527 | 14 | 45283087 | 45318771 | -1 | Mm.387108 |
| 4.1 | 3.91 | 3.58 | 4.44 | DN1ab_Itr | DN1cd | 0.22 | 1.03 | 0.34 | 0.67 | 0.0013 | ENSMUSG00000041975 | protein_coding | Mettl8 | methyltransferase like 8 | 228019 | 2 | 70964561 | 71055583 | -1 | Mm.490343 |
| 4.35 | 4.44 | 5.6 | 4.72 | DN1ab_Itr | DN1cd | -0.29 | -1.47 | 0.18 | 0.52 | 0.0013 | ENSMUSG00000032515 | protein_coding | Csrnp1 | cysteine-serine-rich nuclear protein 1 | 215418 | 9 | 119971163 | 119984658 | -1 | Mm.491094 |
| -0.81 | 0.92 | 0.88 | -0.44 | DN1ab_Itr | DN1cd | 0.75 | 1.29 | 0.24 | 0.58 | 0.0014 | ENSMUSG00000111341 | TEC | Gm47111 | predicted gene, 47111 | NA | 9 | 121843822 | 121846937 | -1 | |
| 6.15 | 6.7 | 5.36 | 6.84 | DN1ab_Itr | DN1cd | 0.88 | 4.76 | 0.002 | 0.045 | 0.0014 | ENSMUSG00000038866 | protein_coding | Zcchc2 | zinc finger, CCHC domain containing 2 | 227449 | 1 | 105990406 | 106034074 | 1 | Mm.210188 |
| 0.69 | 0.38 | -1.48 | 2.03 | DN1ab_Itr | DN1cd | 1.19 | 1.55 | 0.17 | 0.5 | 0.0014 | ENSMUSG00000051618 | protein_coding | Ubqln3 | ubiquilin 3 | 244178 | 7 | 104140623 | 104143279 | -1 | Mm.334092 |
| 7.16 | 6.72 | 8.4 | 6.73 | DN1ab_Itr | DN1cd | -0.91 | -3.7 | 0.0076 | 0.095 | 0.0014 | ENSMUSG00000019943 | protein_coding | Atp2b1 | ATPase, Ca++ transporting, plasma membrane 1 | 67972 | 10 | 98914406 | 99026143 | 1 | Mm.166944 Mm.471912 |
| 8.46 | 7.93 | 9.09 | 7.72 | DN1ab_Itr | DN1cd | -0.85 | -5.04 | 0.0015 | 0.038 | 0.0015 | ENSMUSG00000044894 | protein_coding | Uqcrq | ubiquinol-cytochrome c reductase, complex III subunit VII | 22272 | 11 | 53427922 | 53430831 | -1 | Mm.251621 |
| 4.04 | 3.21 | 3.83 | 4.05 | DN1ab_Itr | DN1cd | -0.38 | -1.77 | 0.12 | 0.42 | 0.0015 | ENSMUSG00000041263 | protein_coding | Rusc1 | RUN and SH3 domain containing 1 | 72296 | 3 | 89083981 | 89093311 | -1 | Mm.27687 |
| 3.61 | 4.2 | 2.44 | 4.93 | DN1ab_Itr | DN1cd | 1.27 | 3.36 | 0.012 | 0.12 | 0.0015 | ENSMUSG00000028634 | protein_coding | Hivep3 | human immunodeficiency virus type I enhancer binding protein 3 | 16656 | 4 | 119733784 | 120138045 | 1 | Mm.302758 Mm.394479 Mm.422538 Mm.487430 Mm.488325 |
| 2.24 | 4.79 | -0.42 | 4.58 | DN1ab_Itr | DN1cd | 3.3 | 7.04 | 2.00E-04 | 0.015 | 0.0015 | ENSMUSG00000032092 | protein_coding | Mpzl2 | myelin protein zero-like 2 | 14012 | 9 | 45039799 | 45054015 | 1 | Mm.33240 |
| 2.09 | 2.12 | 3.49 | 1.82 | DN1ab_Itr | DN1cd | -0.61 | -1.79 | 0.12 | 0.42 | 0.0015 | ENSMUSG00000086742 | processed_transcript | Gm16201 | predicted gene 16201 | NA | 7 | 139950396 | 139978977 | 1 | |
| 4.46 | 5.27 | 3.99 | 6.1 | DN1ab_Itr | DN1cd | 1.28 | 4.96 | 0.0016 | 0.04 | 0.0015 | ENSMUSG00000031402 | protein_coding | Mpp1 | membrane protein, palmitoylated | 17524 | X | 75109733 | 75131016 | -1 | Mm.391267 |
| 4.18 | 4.44 | 3.24 | 4.73 | DN1ab_Itr | DN1cd | 0.71 | 2.9 | 0.023 | 0.17 | 0.0015 | ENSMUSG00000036591 | protein_coding | Arhgap21 | Rho GTPase activating protein 21 | 71435 | 2 | 20847919 | 20968881 | -1 | Mm.28507 Mm.465492 |
| 4.07 | 4.05 | 2.9 | 4.96 | DN1ab_Itr | DN1cd | 0.77 | 1.85 | 0.11 | 0.4 | 0.0015 | ENSMUSG00000056531 | protein_coding | Ccdc18 | coiled-coil domain containing 18 | 73254 | 5 | 108132875 | 108233628 | 1 | Mm.441231 |
| 5.31 | 5.22 | 4.31 | 5.86 | DN1ab_Itr | DN1cd | 0.53 | 1.61 | 0.15 | 0.47 | 0.0015 | ENSMUSG00000033083 | protein_coding | Tbc1d4 | TBC1 domain family, member 4 | 210789 | 14 | 101442360 | 101609191 | -1 | Mm.320639 |
| 6.07 | 5.75 | 7.63 | 6.22 | DN1ab_Itr | DN1cd | -0.74 | -3.34 | 0.012 | 0.12 | 0.0016 | ENSMUSG00000063524 | protein_coding | Eno1 | enolase 1, alpha non-neuron | 13806 | 4 | 150236721 | 150248879 | 1 | Mm.70666 Mm.372357 Mm.372389 |
| -1.01 | -0.33 | -3.94 | 0.06 | DN1ab_Itr | DN1cd | 1.85 | 2.81 | 0.026 | 0.18 | 0.0016 | ENSMUSG00000036912 | protein_coding | Piwil4 | piwi-like RNA-mediated gene silencing 4 | 330890 | 9 | 14696230 | 14740733 | -1 | Mm.334969 |
| 6.55 | 6.54 | 7.12 | 6.15 | DN1ab_Itr | DN1cd | -0.36 | -1.86 | 0.11 | 0.4 | 0.0016 | ENSMUSG00000032624 | protein_coding | Eml4 | echinoderm microtubule associated protein like 4 | 78798 | 17 | 83350931 | 83480361 | 1 | Mm.295565 |
| -1.92 | -2.28 | -3.94 | -0.71 | DN1ab_Itr | DN1cd | 0.99 | 1.37 | 0.21 | 0.55 | 0.0016 | ENSMUSG00000018126 | protein_coding | Baiap2l2 | BAI1-associated protein 2-like 2 | 207495 | 15 | 79258195 | 79285537 | -1 | Mm.309647 |
| -3.82 | -1.14 | -2.81 | -4.02 | DN1ab_Itr | DN1cd | 1.29 | 1.67 | 0.14 | 0.46 | 0.0016 | ENSMUSG00000081660 | unprocessed_pseudogene | Gm11787 | predicted gene 11787 | NA | 4 | 3496978 | 3516515 | -1 | |
| 1.72 | 2.84 | 1.8 | 4.82 | DN1ab_Itr | DN1cd | 1.81 | 4.75 | 0.002 | 0.046 | 0.0017 | ENSMUSG00000052160 | protein_coding | Pld4 | phospholipase D family, member 4 | 104759 | 12 | 112760655 | 112768990 | 1 | Mm.203915 |
| 3.72 | 3.95 | 6.39 | 5.11 | DN1ab_Itr | DN1cd | -0.4 | -1.27 | 0.24 | 0.58 | 0.0017 | ENSMUSG00000024235 | protein_coding | Map3k8 | mitogen-activated protein kinase kinase kinase 8 | 26410 | 18 | 4331327 | 4353015 | -1 | Mm.3275 |
| 10.15 | 10.42 | 9.11 | 10.46 | DN1ab_Itr | DN1cd | 0.66 | 3.03 | 0.019 | 0.16 | 0.0017 | ENSMUSG00000023132 | protein_coding | Gzma | granzyme A | 14938 | 13 | 113093825 | 113100981 | -1 | Mm.15510 |
| 2.85 | 5.17 | 1.45 | 6.08 | DN1ab_Itr | DN1cd | 3.06 | 6.79 | 0.00025 | 0.017 | 0.0018 | ENSMUSG00000061311 | protein_coding | Rag1 | recombination activating gene 1 | 19373 | 2 | 101638282 | 101649501 | -1 | Mm.828 |
| 6.34 | 7.09 | 6.42 | 8.15 | DN1ab_Itr | DN1cd | 1.11 | 5.67 | 0.00074 | 0.028 | 0.0018 | ENSMUSG00000051285 | protein_coding | Pcmtd1 | protein-L-isoaspartate (D-aspartate) O-methyltransferase domain containing 1 | 319263 | 1 | 7088920 | 7173628 | 1 | Mm.21539 |
| -1.5 | -2.28 | -3.94 | -0.77 | DN1ab_Itr | DN1cd | 0.7 | 0.88 | 0.41 | 0.74 | 0.0018 | ENSMUSG00000040434 | protein_coding | Large2 | LARGE xylosyl- and glucuronyltransferase 2 | 228366 | 2 | 92365046 | 92374869 | -1 | Mm.31158 |
| 0.36 | -3.86 | -0.57 | -1.13 | DN1ab_Itr | DN1cd | -2.72 | -3.64 | 0.0082 | 0.099 | 0.0019 | ENSMUSG00000035969 | protein_coding | Rusc2 | RUN and SH3 domain containing 2 | 100213 | 4 | 43381979 | 43427088 | 1 | Mm.235730 Mm.401159 |
| -3.82 | -0.7 | -1.12 | -1.7 | DN1ab_Itr | DN1cd | 1.86 | 2.54 | 0.039 | 0.23 | 0.0019 | ENSMUSG00000070304 | protein_coding | Scn2b | sodium channel, voltage-gated, type II, beta | 72821 | 9 | 45117782 | 45130070 | 1 | Mm.229373 Mm.491371 |
| 5.29 | 5.21 | 4.36 | 5.36 | DN1ab_Itr | DN1cd | 0.32 | 1.47 | 0.19 | 0.53 | 0.0019 | ENSMUSG00000025950 | protein_coding | Idh1 | isocitrate dehydrogenase 1 (NADP+), soluble | 15926 | 1 | 65158616 | 65186500 | -1 | Mm.9925 |
| 6.99 | 7.19 | 6.81 | 7.65 | DN1ab_Itr | DN1cd | 0.44 | 3.36 | 0.012 | 0.12 | 0.0019 | ENSMUSG00000009585 | protein_coding | Apobec3 | apolipoprotein B mRNA editing enzyme, catalytic polypeptide 3 | 80287 | 15 | 79891659 | 79915906 | 1 | Mm.284059 Mm.432615 |
| -0.23 | -0.1 | 1.22 | -1.7 | DN1ab_Itr | DN1cd | -0.94 | -1.54 | 0.17 | 0.5 | 0.0019 | ENSMUSG00000025779 | protein_coding | Ly96 | lymphocyte antigen 96 | 17087 | 1 | 16688051 | 16709611 | 1 | Mm.116844 |
| -3.82 | -3.86 | -3.94 | -1.03 | DN1ab_Itr | DN1cd | 1.04 | 1.76 | 0.12 | 0.43 | 0.002 | ENSMUSG00000112481 | lincRNA | Gm29684 | predicted gene, 29684 | NA | 10 | 95120582 | 95145825 | -1 | |
| 5.26 | 5.33 | 4.28 | 5.54 | DN1ab_Itr | DN1cd | 0.5 | 2.08 | 0.076 | 0.33 | 0.002 | ENSMUSG00000002997 | protein_coding | Prkar2b | protein kinase, cAMP dependent regulatory, type II beta | 19088 | 12 | 31958476 | 32061296 | -1 | Mm.25594 |
| 6.15 | 5.03 | 6.9 | 6.53 | DN1ab_Itr | DN1cd | -0.8 | -5.2 | 0.0012 | 0.035 | 0.002 | ENSMUSG00000095041 | protein_coding | | NA | NA | JH584304.1 | 52190 | 59690 | -1 | |
| -3.82 | -3.86 | -0.88 | -4.02 | DN1ab_Itr | DN1cd | -1.15 | -1.85 | 0.11 | 0.4 | 0.002 | ENSMUSG00000003309 | protein_coding | Ap1m2 | adaptor protein complex AP-1, mu 2 subunit | 11768 | 9 | 21294275 | 21312337 | -1 | Mm.22239 |
| 3.68 | 4.47 | 4.84 | 4.59 | DN1ab_Itr | DN1cd | 0.41 | 1.99 | 0.087 | 0.36 | 0.002 | ENSMUSG00000027030 | protein_coding | Stk39 | serine/threonine kinase 39 | 53416 | 2 | 68210445 | 68472268 | -1 | Mm.198414 |
| 8.64 | 8.1 | 9.31 | 8.17 | DN1ab_Itr | DN1cd | -0.77 | -6.25 | 0.00041 | 0.021 | 0.002 | ENSMUSG00000025366 | protein_coding | Esyt1 | extended synaptotagmin-like protein 1 | 23943 | 10 | 128509965 | 128525871 | -1 | Mm.66056 |
| -3.82 | -3.86 | -3.94 | -1.22 | DN1ab_Itr | DN1cd | 0.97 | 1.74 | 0.13 | 0.43 | 0.0021 | ENSMUSG00000097934 | lincRNA | 6720483E21Rik | RIKEN cDNA 6720483E21 gene | NA | 1 | 20888650 | 20890473 | -1 | |
| -3.82 | -3.86 | -3.94 | -1.22 | DN1ab_Itr | DN1cd | 0.97 | 1.74 | 0.13 | 0.43 | 0.0021 | ENSMUSG00000104784 | antisense | Gm42984 | predicted gene 42984 | NA | 5 | 44226587 | 44243548 | 1 | |
| -3.82 | -3.86 | -3.94 | -1.22 | DN1ab_Itr | DN1cd | 0.97 | 1.74 | 0.13 | 0.43 | 0.0021 | ENSMUSG00000066607 | protein_coding | Insyn1 | inhibitory synaptic factor 1 | 319477 | 9 | 58488603 | 58499780 | 1 | Mm.153674 |
| -3.82 | -3.86 | -3.94 | -1.22 | DN1ab_Itr | DN1cd | 0.97 | 1.74 | 0.13 | 0.43 | 0.0021 | ENSMUSG00000091007 | protein_coding | D630036H23Rik | RIKEN cDNA D630036H23 gene | NA | 12 | 36381242 | 36382181 | -1 | |
| 7.29 | 7.33 | 6.92 | 7.59 | DN1ab_Itr | DN1cd | 0.27 | 2.2 | 0.064 | 0.3 | 0.0021 | ENSMUSG00000013663 | protein_coding | Pten | phosphatase and tensin homolog | 19211 | 19 | 32757497 | 32826160 | 1 | Mm.245395 |
| 1.4 | 1.99 | 1.35 | 3.95 | DN1ab_Itr | DN1cd | 1.36 | 3.33 | 0.013 | 0.13 | 0.0021 | ENSMUSG00000034586 | protein_coding | Hid1 | HID1 domain containing | 217310 | 11 | 115347707 | 115367756 | -1 | Mm.205707 |
| 4.9 | 5.76 | 4.21 | 6.14 | DN1ab_Itr | DN1cd | 1.24 | 5.73 | 0.00069 | 0.027 | 0.0021 | ENSMUSG00000029275 | protein_coding | Gfi1 | growth factor independent 1 transcription repressor | 14581 | 5 | 107716657 | 107726036 | -1 | Mm.2078 Mm.453139 |
| 5.21 | 5.07 | 4.78 | 5.62 | DN1ab_Itr | DN1cd | 0.24 | 1.2 | 0.27 | 0.6 | 0.0022 | ENSMUSG00000057367 | protein_coding | Birc2 | baculoviral IAP repeat-containing 2 | 11797 | 9 | 7818227 | 7837064 | -1 | Mm.335659 Mm.400826 |
| -3.82 | -3.86 | -0.82 | -4.02 | DN1ab_Itr | DN1cd | -1.17 | -1.85 | 0.11 | 0.4 | 0.0022 | ENSMUSG00000029228 | protein_coding | Lnx1 | ligand of numb-protein X 1 | 16924 | 5 | 74592447 | 74702912 | -1 | Mm.440403 |
| 6.43 | 5.47 | 8.15 | 6.12 | DN1ab_Itr | DN1cd | -1.4 | -6.36 | 0.00037 | 0.02 | 0.0022 | ENSMUSG00000002489 | protein_coding | Tiam1 | T cell lymphoma invasion and metastasis 1 | 21844 | 16 | 89787111 | 90143769 | -1 | Mm.310902 |
| 8.63 | 8.55 | 7.89 | 8.32 | DN1ab_Itr | DN1cd | 0.11 | 1.05 | 0.33 | 0.67 | 0.0022 | ENSMUSG00000038252 | protein_coding | Ncapd2 | non-SMC condensin I complex, subunit D2 | 68298 | 6 | 125168007 | 125191701 | -1 | Mm.257590 |
| 4.41 | 5.51 | 3.98 | 7.37 | DN1ab_Itr | DN1cd | 1.92 | 4.17 | 0.0041 | 0.066 | 0.0022 | ENSMUSG00000032094 | protein_coding | Cd3d | CD3 antigen, delta polypeptide | 12500 | 9 | 44981786 | 44987339 | 1 | Mm.4527 |
| 5.88 | 6.26 | 6.18 | 7.19 | DN1ab_Itr | DN1cd | 0.62 | 4.85 | 0.0018 | 0.043 | 0.0022 | ENSMUSG00000026313 | protein_coding | Hdac4 | histone deacetylase 4 | 208727 | 1 | 91928779 | 92195699 | -1 | Mm.318567 |
| 3.98 | 5.29 | 2.7 | 5.43 | DN1ab_Itr | DN1cd | 1.79 | 6.33 | 0.00038 | 0.02 | 0.0022 | ENSMUSG00000035547 | protein_coding | Capn5 | calpain 5 | 12337 | 7 | 98121559 | 98178274 | -1 | Mm.326847 |
| 7.11 | 7.1 | 6.39 | 6.87 | DN1ab_Itr | DN1cd | 0.17 | 1.63 | 0.15 | 0.47 | 0.0023 | ENSMUSG00000040084 | protein_coding | Bub1b | BUB1B, mitotic checkpoint serine/threonine kinase | 12236 | 2 | 118598211 | 118641591 | 1 | Mm.29133 |
| 6.51 | 6.47 | 7.16 | 6.36 | DN1ab_Itr | DN1cd | -0.32 | -2.03 | 0.082 | 0.34 | 0.0023 | ENSMUSG00000026786 | protein_coding | Apbb1ip | amyloid beta (A4) precursor protein-binding, family B, member 1 interacting protein | 54519 | 2 | 22774094 | 22875653 | 1 | Mm.14255 |
| 1 | -1.54 | 0.92 | 1.05 | DN1ab_Itr | DN1cd | -1.16 | -2.07 | 0.077 | 0.33 | 0.0023 | ENSMUSG00000018500 | protein_coding | Adora2b | adenosine A2b receptor | 11541 | 11 | 62248984 | 62266453 | 1 | Mm.40740 |
| 2.56 | 2.22 | 5.37 | 3.05 | DN1ab_Itr | DN1cd | -1.23 | -3 | 0.02 | 0.16 | 0.0023 | ENSMUSG00000044345 | protein_coding | Marveld1 | MARVEL (membrane-associating) domain containing 1 | 277010 | 19 | 42147400 | 42151703 | 1 | Mm.206250 |
| 7.54 | 7.93 | 7.2 | 8.43 | DN1ab_Itr | DN1cd | 0.7 | 4.08 | 0.0046 | 0.07 | 0.0024 | ENSMUSG00000110275 | processed_pseudogene | Gm5905 | predicted gene 5905 | NA | 7 | 141177885 | 141178464 | -1 | |
| 2.99 | 4.56 | 2.43 | 6.58 | DN1ab_Itr | DN1cd | 2.45 | 4.77 | 0.002 | 0.045 | 0.0024 | ENSMUSG00000026011 | protein_coding | Ctla4 | cytotoxic T-lymphocyte-associated protein 4 | 12477 | 1 | 60887000 | 60915832 | 1 | Mm.390 |
| 3.11 | 3.34 | 1.28 | 3.4 | DN1ab_Itr | DN1cd | 0.9 | 2.37 | 0.05 | 0.26 | 0.0024 | ENSMUSG00000007682 | protein_coding | Dio2 | deiodinase, iodothyronine, type II | 13371 | 12 | 90724552 | 90738438 | -1 | Mm.21389 Mm.408815 |
| 4.54 | 2.89 | 4.99 | 4.3 | DN1ab_Itr | DN1cd | -1.18 | -5.8 | 0.00065 | 0.026 | 0.0024 | ENSMUSG00000043391 | protein_coding | 2510009E07Rik | RIKEN cDNA 2510009E07 gene | 72190 | 16 | 21649045 | 21694665 | -1 | Mm.131606 |
| 7.01 | 6.92 | 7.76 | 7.13 | DN1ab_Itr | DN1cd | -0.29 | -2.61 | 0.035 | 0.22 | 0.0025 | ENSMUSG00000030403 | protein_coding | Vasp | vasodilator-stimulated phosphoprotein | 22323 | 7 | 19256929 | 19271817 | -1 | Mm.9684 |
| 4.63 | 6 | 3.46 | 6.11 | DN1ab_Itr | DN1cd | 1.81 | 7.11 | 0.00019 | 0.015 | 0.0025 | ENSMUSG00000048285 | protein_coding | Frmd6 | FERM domain containing 6 | 319710 | 12 | 70825514 | 70902234 | 1 | Mm.2962 |
| 3.34 | 3.35 | 2.86 | 4.25 | DN1ab_Itr | DN1cd | 0.55 | 1.92 | 0.096 | 0.38 | 0.0025 | ENSMUSG00000036158 | protein_coding | Prickle1 | prickle planar cell polarity protein 1 | 106042 | 15 | 93499114 | 93595891 | -1 | Mm.150314 |
| 5.97 | 5.09 | 6.95 | 5.06 | DN1ab_Itr | DN1cd | -1.28 | -6.19 | 0.00044 | 0.022 | 0.0025 | ENSMUSG00000032554 | protein_coding | Trf | transferrin | 22041 | 9 | 103204001 | 103230444 | -1 | Mm.37214 |
| 6.67 | 6.63 | 6.82 | 6.01 | DN1ab_Itr | DN1cd | -0.32 | -2.01 | 0.084 | 0.35 | 0.0025 | ENSMUSG00000000628 | protein_coding | Hk2 | hexokinase 2 | 15277 | 6 | 82725025 | 82774454 | -1 | Mm.255848 Mm.469167 |
| 9.65 | 9.41 | 9.91 | 9.14 | DN1ab_Itr | DN1cd | -0.43 | -3.99 | 0.0052 | 0.076 | 0.0025 | ENSMUSG00000026547 | protein_coding | Tagln2 | transgelin 2 | 21346 | 1 | 172500047 | 172507380 | 1 | Mm.271711 |
| 4.16 | 4.76 | 3.25 | 5.07 | DN1ab_Itr | DN1cd | 1.04 | 4.18 | 0.0041 | 0.066 | 0.0026 | ENSMUSG00000033721 | protein_coding | Vav3 | vav 3 oncogene | 57257 | 3 | 109340653 | 109685698 | 1 | Mm.282257 |
| 1.23 | 1.04 | 0.74 | 2.28 | DN1ab_Itr | DN1cd | 0.51 | 1.41 | 0.2 | 0.54 | 0.0026 | ENSMUSG00000054203 | protein_coding | Ifi205 | interferon activated gene 205 | 226695 | 1 | 174011998 | 174031810 | -1 | Mm.425061 |
| 4.56 | 5.14 | 5.01 | 4.88 | DN1ab_Itr | DN1cd | 0.32 | 2.25 | 0.059 | 0.29 | 0.0026 | ENSMUSG00000040548 | protein_coding | Tex2 | testis expressed gene 2 | 21763 | 11 | 106502147 | 106613423 | -1 | Mm.102407 |
| -3.82 | -3.86 | -0.73 | -4.02 | DN1ab_Itr | DN1cd | -1.21 | -1.83 | 0.11 | 0.4 | 0.0027 | ENSMUSG00000086127 | processed_transcript | Gm11934 | predicted gene 11934 | NA | 4 | 33303569 | 33310137 | -1 | |
| -0.4 | 1.24 | -3.94 | 1.47 | DN1ab_Itr | DN1cd | 2.83 | 3.82 | 0.0065 | 0.087 | 0.0027 | ENSMUSG00000002100 | protein_coding | Mybpc3 | myosin binding protein C, cardiac | 17868 | 2 | 91118144 | 91136516 | 1 | Mm.10728 |
| 4.23 | 2.81 | 2.91 | 3.13 | DN1ab_Itr | DN1cd | -0.72 | -2.1 | 0.074 | 0.33 | 0.0027 | ENSMUSG00000036030 | protein_coding | Prtg | protogenin | 235472 | 9 | 72806874 | 72917291 | 1 | Mm.31696 Mm.469160 |
| 4.1 | 5.69 | 2.77 | 5.97 | DN1ab_Itr | DN1cd | 2.13 | 6.59 | 3.00E-04 | 0.018 | 0.0027 | ENSMUSG00000038963 | protein_coding | Slco4a1 | solute carrier organic anion transporter family, member 4a1 | 108115 | 2 | 180456245 | 180474867 | 1 | Mm.133687 |
| 4.69 | 4.4 | 5.87 | 4.64 | DN1ab_Itr | DN1cd | -0.67 | -3.41 | 0.011 | 0.12 | 0.0027 | ENSMUSG00000024937 | protein_coding | Ehbp1l1 | EH domain binding protein 1-like 1 | 114601 | 19 | 5707376 | 5726317 | -1 | Mm.210447 |
| 6.22 | 6.69 | 6.08 | 7.17 | DN1ab_Itr | DN1cd | 0.7 | 5.48 | 9.00E-04 | 0.03 | 0.0027 | ENSMUSG00000046985 | protein_coding | Tapt1 | transmembrane anterior posterior transformation 1 | 231225 | 5 | 44175154 | 44226626 | -1 | Mm.30688 Mm.369319 |
| 0.28 | 2.8 | -3.94 | 3.93 | DN1ab_Itr | DN1cd | 4.15 | 3.97 | 0.0053 | 0.077 | 0.0027 | ENSMUSG00000046844 | protein_coding | Vat1l | vesicle amine transport protein 1 like | 270097 | 8 | 114205612 | 114374071 | 1 | Mm.334825 |
| 1.1 | -1.14 | -0.49 | -0.12 | DN1ab_Itr | DN1cd | -1.04 | -1.88 | 0.1 | 0.39 | 0.0028 | ENSMUSG00000027848 | protein_coding | Olfml3 | olfactomedin-like 3 | 99543 | 3 | 103722222 | 103738001 | -1 | Mm.211535 |
| 1.76 | 3.17 | 4.42 | 4.11 | DN1ab_Itr | DN1cd | 0.78 | 2.21 | 0.063 | 0.3 | 0.0028 | ENSMUSG00000057897 | protein_coding | Camk2b | calcium/calmodulin-dependent protein kinase II, beta | 12323 | 11 | 5969644 | 6066362 | -1 | Mm.439733 |
| -3.82 | -3.86 | -1.27 | -4.02 | DN1ab_Itr | DN1cd | -1.01 | -1.84 | 0.11 | 0.4 | 0.0028 | ENSMUSG00000051219 | protein_coding | 3100002H09Rik | RIKEN cDNA 3100002H09 gene | NA | 4 | 124609931 | 124610803 | -1 | |
| -3.82 | -3.86 | -1.27 | -4.02 | DN1ab_Itr | DN1cd | -1.01 | -1.84 | 0.11 | 0.4 | 0.0028 | ENSMUSG00000047129 | protein_coding | 1700113H08Rik | RIKEN cDNA 1700113H08 gene | 76640 | 10 | 87058045 | 87230589 | 1 | Mm.78423 |
| 0.43 | 0.78 | -3.94 | -0.1 | DN1ab_Itr | DN1cd | 1.48 | 2.14 | 0.069 | 0.31 | 0.0028 | ENSMUSG00000093765 | antisense | Gm20658 | predicted gene 20658 | NA | 12 | 116102769 | 116138648 | -1 | |
| 2.11 | 3.09 | 0.7 | 3.85 | DN1ab_Itr | DN1cd | 1.73 | 3.94 | 0.0055 | 0.079 | 0.0028 | ENSMUSG00000034881 | protein_coding | Tbxa2r | thromboxane A2 receptor | 21390 | 10 | 81328731 | 81335172 | 1 | Mm.4545 |
| 6.32 | 6.09 | 6.87 | 6.05 | DN1ab_Itr | DN1cd | -0.45 | -3.72 | 0.0073 | 0.093 | 0.0028 | ENSMUSG00000032350 | protein_coding | Gclc | glutamate-cysteine ligase, catalytic subunit | 14629 | 9 | 77754535 | 77794485 | 1 | Mm.89888 |
| 9.43 | 8.72 | 8.9 | 9.06 | DN1ab_Itr | DN1cd | -0.37 | -2.07 | 0.077 | 0.33 | 0.0029 | ENSMUSG00000000782 | protein_coding | Tcf7 | transcription factor 7, T cell specific | 21414 | 11 | 52252371 | 52283014 | -1 | Mm.31630 |
| 6.13 | 5.52 | 7.22 | 5.67 | DN1ab_Itr | DN1cd | -0.97 | -4.98 | 0.0016 | 0.04 | 0.0029 | ENSMUSG00000039316 | protein_coding | Rftn1 | raftlin lipid raft linker 1 | 76438 | 17 | 49992257 | 50190674 | -1 | Mm.41854 |
| 5.32 | 5.29 | 5.07 | 5.86 | DN1ab_Itr | DN1cd | 0.28 | 1.7 | 0.13 | 0.45 | 0.0029 | ENSMUSG00000022779 | protein_coding | Top3b | topoisomerase (DNA) III beta | 21976 | 16 | 16870736 | 16892990 | 1 | Mm.326089 |
| 4.05 | 4.17 | 4.77 | 3.81 | DN1ab_Itr | DN1cd | -0.28 | -1.26 | 0.25 | 0.59 | 0.0029 | ENSMUSG00000004317 | protein_coding | Clcn5 | chloride channel, voltage-sensitive 5 | 12728 | X | 7153810 | 7319358 | -1 | Mm.486257 |
| 0.65 | 1.03 | -1.04 | 2.06 | DN1ab_Itr | DN1cd | 1.38 | 2.48 | 0.042 | 0.24 | 0.003 | ENSMUSG00000029279 | protein_coding | Brdt | bromodomain, testis-specific | 114642 | 5 | 107331159 | 107387058 | 1 | Mm.182836 |
| 4.24 | 3.86 | 6.52 | 3.89 | DN1ab_Itr | DN1cd | -1.3 | -2.75 | 0.028 | 0.19 | 0.003 | ENSMUSG00000018821 | protein_coding | Avpi1 | arginine vasopressin-induced 1 | 69534 | 19 | 42123273 | 42129059 | -1 | Mm.30060 |
| 5.16 | 4.87 | 5.61 | 4.57 | DN1ab_Itr | DN1cd | -0.57 | -3.69 | 0.0076 | 0.095 | 0.003 | ENSMUSG00000003948 | protein_coding | Mmd | monocyte to macrophage differentiation-associated | 67468 | 11 | 90249456 | 90278589 | 1 | Mm.277518 Mm.488129 |
| -3.82 | -1.31 | -3.94 | -4.02 | DN1ab_Itr | DN1cd | 1.57 | 2.97 | 0.021 | 0.16 | 0.003 | ENSMUSG00000064489 | snRNA | Gm23296 | predicted gene, 23296 | NA | 12 | 52604485 | 52604591 | -1 | |
| -3.82 | -1.31 | -3.94 | -4.02 | DN1ab_Itr | DN1cd | 1.57 | 2.97 | 0.021 | 0.16 | 0.003 | ENSMUSG00000072919 | protein_coding | Noxred1 | NADP+ dependent oxidoreductase domain containing 1 | 71275 | 12 | 87221040 | 87238732 | -1 | Mm.79198 |
| 7.26 | 7.4 | 7.18 | 8.06 | DN1ab_Itr | DN1cd | 0.41 | 2.7 | 0.031 | 0.2 | 0.0032 | ENSMUSG00000039852 | protein_coding | Rere | arginine glutamic acid dipeptide (RE) repeats | 68703 | 4 | 150281646 | 150621966 | 1 | Mm.291274 |
| -1.54 | 1.34 | -3.94 | 2.56 | DN1ab_Itr | DN1cd | 4.03 | 5.59 | 0.00081 | 0.029 | 0.0032 | ENSMUSG00000075330 | lincRNA | A930003A15Rik | RIKEN cDNA A930003A15 gene | NA | 16 | 19876567 | 19884274 | -1 | |
| 7.42 | 7.05 | 7.24 | 7.33 | DN1ab_Itr | DN1cd | -0.2 | -2.03 | 0.082 | 0.35 | 0.0033 | ENSMUSG00000060261 | protein_coding | Gtf2i | general transcription factor II I | 14886 | 5 | 134237834 | 134314760 | -1 | Mm.261570 Mm.412191 Mm.466495 |
| 7.51 | 7.48 | 7.13 | 7.57 | DN1ab_Itr | DN1cd | 0.15 | 1.52 | 0.17 | 0.5 | 0.0033 | ENSMUSG00000001151 | protein_coding | Pcnt | pericentrin (kendrin) | 18541 | 10 | 76351263 | 76442786 | -1 | Mm.251794 |
| -0.09 | -0.06 | -3.94 | 0.15 | DN1ab_Itr | DN1cd | 1.44 | 1.74 | 0.12 | 0.43 | 0.0034 | ENSMUSG00000030650 | protein_coding | Tmc5 | transmembrane channel-like gene family 5 | 74424 | 7 | 118597297 | 118675086 | 1 | Mm.11068 |
| 1.68 | -0.44 | 1.53 | 1.55 | DN1ab_Itr | DN1cd | -0.99 | -2.16 | 0.067 | 0.31 | 0.0035 | ENSMUSG00000032595 | protein_coding | Cdhr4 | cadherin-related family member 4 | 69398 | 9 | 107992497 | 107999684 | 1 | Mm.476887 |
| 4.4 | 4.64 | 3.1 | 4.28 | DN1ab_Itr | DN1cd | 0.57 | 2.95 | 0.021 | 0.17 | 0.0035 | ENSMUSG00000040618 | protein_coding | Pck2 | phosphoenolpyruvate carboxykinase 2 (mitochondrial) | 74551 | 14 | 55540266 | 55551242 | 1 | Mm.29856 Mm.491140 |
| 5.58 | 5.57 | 4.72 | 5.36 | DN1ab_Itr | DN1cd | 0.23 | 1.7 | 0.13 | 0.44 | 0.0035 | ENSMUSG00000001911 | protein_coding | Nfix | nuclear factor I/X | 18032 | 8 | 84699876 | 84800344 | -1 | Mm.9394 Mm.487058 |
| 5.06 | 5.67 | 5.33 | 6.71 | DN1ab_Itr | DN1cd | 0.89 | 5.56 | 0.00083 | 0.03 | 0.0036 | ENSMUSG00000028657 | protein_coding | Ppt1 | palmitoyl-protein thioesterase 1 | 19063 | 4 | 122836242 | 122859175 | 1 | Mm.277719 |
| 2.38 | 2.54 | 0.98 | 3.8 | DN1ab_Itr | DN1cd | 1.17 | 2.11 | 0.073 | 0.32 | 0.0036 | ENSMUSG00000025810 | protein_coding | Nrp1 | neuropilin 1 | 18186 | 8 | 128358604 | 128503363 | 1 | Mm.271745 |
| 8.12 | 7.76 | 7.76 | 7.83 | DN1ab_Itr | DN1cd | -0.2 | -2.2 | 0.063 | 0.3 | 0.0036 | ENSMUSG00000031004 | protein_coding | Mki67 | antigen identified by monoclonal antibody Ki 67 | 17345 | 7 | 135689784 | 135716361 | -1 | Mm.4078 |
| -3.82 | -3.86 | -3.94 | -0.89 | DN1ab_Itr | DN1cd | 1.11 | 1.73 | 0.13 | 0.44 | 0.0036 | ENSMUSG00000025020 | protein_coding | Slit1 | slit guidance ligand 1 | 20562 | 19 | 41600257 | 41743665 | -1 | Mm.39155 Mm.40322 |
| 4.39 | 4.94 | 4.5 | 4.34 | DN1ab_Itr | DN1cd | 0.3 | 2.07 | 0.077 | 0.33 | 0.0036 | ENSMUSG00000019861 | protein_coding | Gopc | golgi associated PDZ and coiled-coil motif containing | 94221 | 10 | 52335850 | 52382124 | -1 | Mm.390258 |
| 4.56 | 5.19 | 4.05 | 5.87 | DN1ab_Itr | DN1cd | 1.07 | 4.36 | 0.0032 | 0.057 | 0.0036 | ENSMUSG00000022471 | protein_coding | Xrcc6 | X-ray repair complementing defective repair in Chinese hamster cells 6 | 14375 | 15 | 81987835 | 82040085 | 1 | Mm.288809 |
| 6.51 | 6.32 | 7.46 | 6.5 | DN1ab_Itr | DN1cd | -0.48 | -3 | 0.02 | 0.16 | 0.0036 | ENSMUSG00000021886 | protein_coding | Gpr65 | G-protein coupled receptor 65 | 14744 | 12 | 98268635 | 98276644 | 1 | Mm.378924 |
| -3.82 | -3.86 | -0.05 | -4.02 | DN1ab_Itr | DN1cd | -1.45 | -1.79 | 0.12 | 0.42 | 0.0037 | ENSMUSG00000071567 | NA | NA | NA | NA | NA | NA | NA | NA | |
| 3.93 | 4.07 | 2.98 | 4.12 | DN1ab_Itr | DN1cd | 0.5 | 2.43 | 0.046 | 0.25 | 0.0037 | ENSMUSG00000062190 | protein_coding | Lancl2 | LanC (bacterial lantibiotic synthetase component C)-like 2 | 71835 | 6 | 57702540 | 57739439 | 1 | Mm.274904 |
| 8.62 | 9.42 | 8.41 | 9.68 | DN1ab_Itr | DN1cd | 0.97 | 9.94 | 2.10E-05 | 0.0056 | 0.0039 | ENSMUSG00000069662 | protein_coding | Marcks | myristoylated alanine rich protein kinase C substrate | 17118 | 10 | 37133375 | 37138920 | -1 | Mm.30059 |
| -1.01 | -1.14 | 3.21 | -1.49 | DN1ab_Itr | DN1cd | -1.97 | -2.03 | 0.081 | 0.34 | 0.0039 | ENSMUSG00000053199 | protein_coding | Arhgap20 | Rho GTPase activating protein 20 | 244867 | 9 | 51765337 | 51853856 | 1 | Mm.400352 Mm.443529 |
| 9.47 | 9.37 | 10.75 | 9.69 | DN1ab_Itr | DN1cd | -0.47 | -2.34 | 0.051 | 0.26 | 0.0039 | ENSMUSG00000027763 | protein_coding | Mbnl1 | muscleblind like splicing factor 1 | 56758 | 3 | 60472830 | 60629750 | 1 | Mm.255723 |
| 6.25 | 6.39 | 6.51 | 7.38 | DN1ab_Itr | DN1cd | 0.42 | 2.74 | 0.029 | 0.19 | 0.004 | ENSMUSG00000104876 | TR_C_gene | Trdc | T cell receptor delta, constant region | NA | 14 | 54142851 | 54148868 | 1 | |
| 6.8 | 6.81 | 6.27 | 6.77 | DN1ab_Itr | DN1cd | 0.19 | 1.86 | 0.11 | 0.4 | 0.004 | ENSMUSG00000037466 | protein_coding | Tedc1 | tubulin epsilon and delta complex 1 | 104732 | 12 | 113156421 | 113166048 | 1 | Mm.87616 |
| 2.88 | 2.68 | 4.5 | 2.87 | DN1ab_Itr | DN1cd | -0.78 | -2.57 | 0.037 | 0.22 | 0.0041 | ENSMUSG00000005824 | protein_coding | Tnfsf14 | tumor necrosis factor (ligand) superfamily, member 14 | 50930 | 17 | 57189492 | 57194177 | -1 | Mm.483369 |
| -1.01 | 1.37 | 0.58 | 0.3 | DN1ab_Itr | DN1cd | 1.55 | 2.91 | 0.023 | 0.17 | 0.0041 | ENSMUSG00000042087 | TEC | 4933440N22Rik | RIKEN cDNA 4933440N22 gene | NA | 6 | 117905808 | 117907760 | -1 | |
| 2.76 | 2.94 | 2.38 | 3.75 | DN1ab_Itr | DN1cd | 0.64 | 2.57 | 0.037 | 0.22 | 0.0041 | ENSMUSG00000023367 | protein_coding | Tmem176a | transmembrane protein 176A | 66058 | 6 | 48840919 | 48847071 | 1 | Mm.27061 |
| 8.64 | 8.61 | 7.91 | 8.88 | DN1ab_Itr | DN1cd | 0.34 | 1.66 | 0.14 | 0.46 | 0.0042 | ENSMUSG00000034349 | protein_coding | Smc4 | structural maintenance of chromosomes 4 | 70099 | 3 | 69004738 | 69034623 | 1 | Mm.206841 |
| 6.01 | 7.48 | 5.34 | 8.04 | DN1ab_Itr | DN1cd | 1.9 | 7.56 | 0.00013 | 0.013 | 0.0042 | ENSMUSG00000026923 | protein_coding | Notch1 | notch 1 | 18128 | 2 | 26457903 | 26516663 | -1 | Mm.290610 |
| -1.81 | -1.53 | -3.94 | 0.36 | DN1ab_Itr | DN1cd | 1.84 | 2.18 | 0.065 | 0.3 | 0.0042 | ENSMUSG00000034845 | protein_coding | Plvap | plasmalemma vesicle associated protein | 84094 | 8 | 71497765 | 71511752 | -1 | Mm.260768 |
| 5.45 | 5.86 | 4.87 | 6.23 | DN1ab_Itr | DN1cd | 0.76 | 3.86 | 0.0061 | 0.084 | 0.0042 | ENSMUSG00000090019 | protein_coding | Gimap1 | GTPase, IMAP family member 1 | 16205 | 6 | 48739054 | 48743795 | 1 | Mm.252599 |
| 3.69 | 3.93 | 1.86 | 4.42 | DN1ab_Itr | DN1cd | 1.07 | 2.23 | 0.061 | 0.29 | 0.0043 | ENSMUSG00000050377 | protein_coding | Il31ra | interleukin 31 receptor A | 218624 | 13 | 112519898 | 112594360 | -1 | Mm.380801 |
| 3.01 | 4.42 | 2.38 | 5.07 | DN1ab_Itr | DN1cd | 1.84 | 7.04 | 2.00E-04 | 0.015 | 0.0043 | ENSMUSG00000040270 | protein_coding | Bach2 | BTB and CNC homology, basic leucine zipper transcription factor 2 | 12014 | 4 | 32238804 | 32586108 | 1 | Mm.491223 |
| -3.82 | -3.86 | -1.48 | -4.02 | DN1ab_Itr | DN1cd | -0.94 | -1.81 | 0.11 | 0.41 | 0.0043 | ENSMUSG00000105816 | processed_transcript | D030025E07Rik | RIKEN cDNA D030025E07 gene | NA | 3 | 128117015 | 128231606 | -1 | |
| -3.82 | -3.86 | -1.48 | -4.02 | DN1ab_Itr | DN1cd | -0.94 | -1.81 | 0.11 | 0.41 | 0.0043 | ENSMUSG00000057286 | protein_coding | St6galnac2 | ST6 (alpha-N-acetyl-neuraminyl-2,3-beta-galactosyl-1,3)-N-acetylgalactosaminide alpha-2,6-sialyltransferase 2 | 20446 | 11 | 116675303 | 116694868 | -1 | Mm.3947 |
| -3.82 | -3.86 | 0.78 | -4.02 | DN1ab_Itr | DN1cd | -1.75 | -1.77 | 0.12 | 0.42 | 0.0043 | ENSMUSG00000038132 | protein_coding | Rbm24 | RNA binding motif protein 24 | 666794 | 13 | 46418434 | 46431095 | 1 | Mm.38269 |
| 5.28 | 5.27 | 4.86 | 5.91 | DN1ab_Itr | DN1cd | 0.39 | 1.78 | 0.12 | 0.42 | 0.0043 | ENSMUSG00000054619 | protein_coding | Mettl7a1 | methyltransferase like 7A1 | 70152 | 15 | 100304140 | 100328662 | 1 | Mm.488086 Mm.490271 |
| 4.71 | 4.83 | 4.56 | 5.46 | DN1ab_Itr | DN1cd | 0.41 | 2.52 | 0.04 | 0.23 | 0.0043 | ENSMUSG00000040528 | protein_coding | Milr1 | mast cell immunoglobulin like receptor 1 | 380732 | 11 | 106751226 | 106768794 | 1 | Mm.343952 |
| -1.01 | -1.06 | -3.49 | -0.56 | DN1ab_Itr | DN1cd | 1.06 | 1.7 | 0.13 | 0.44 | 0.0044 | ENSMUSG00000101800 | processed_pseudogene | Rnf170-ps | ring finger protein 170, pseudogene | NA | 7 | 26072073 | 26072931 | 1 | |
| 6.49 | 7.12 | 6.22 | 7.37 | DN1ab_Itr | DN1cd | 0.81 | 7.42 | 0.00014 | 0.013 | 0.0044 | ENSMUSG00000027111 | protein_coding | Itga6 | integrin alpha 6 | 16403 | 2 | 71745616 | 71858416 | 1 | Mm.225096 |
| 3.24 | 3.31 | 3.14 | 4.24 | DN1ab_Itr | DN1cd | -0.48 | 2.18 | 0.065 | 0.3 | 0.0044 | ENSMUSG00000021140 | protein_coding | Pcnx | pecanex homolog | 54604 | 12 | 81860023 | 82000924 | 1 | Mm.86584 |
| 4.8 | 4.91 | 3.64 | 4.46 | DN1ab_Itr | DN1cd | 0.36 | 2.44 | 0.045 | 0.25 | 0.0044 | ENSMUSG00000021391 | protein_coding | Cenpp | centromere protein P | 66336 | 13 | 49464023 | 49652785 | 1 | Mm.440323 |
| 4.41 | 4.93 | 2.88 | 5.48 | DN1ab_Itr | DN1cd | 1.26 | 2.93 | 0.022 | 0.17 | 0.0044 | ENSMUSG00000021915 | protein_coding | Sgk3 | serum/glucocorticoid regulated kinase 3 | 170755 | 1 | 9798107 | 9900845 | 1 | Mm.336410 |

| | | | | | | | | | | | | | | | | | | | | |
|---|---|---|---|---|---|---|---|---|---|---|---|---|---|---|---|---|---|---|---|---|
| 4.29 | 4.88 | 4.08 | 5.58 | DN1ab_ltr | DN1cd | 0.93 | 4.9 | 0.0017 | 0.042 | 0.0045 | ENSMUSG00000105519 | TEC | E430021H15Rik | RIKEN cDNA E430021H15 gene | NA | 5 | 44213801 | 44217259 | -1 | |
| -3.82 | -3.86 | -1.38 | -4.02 | DN1ab_ltr | DN1cd | -0.97 | -1.8 | 0.11 | 0.41 | 0.0045 | ENSMUSG00000110551 | processed_pseudogene | Gm45750 | predicted gene 45750 | NA | 8 | 104223747 | 104224100 | 1 | |
| -3.82 | -3.86 | -1.38 | -4.02 | DN1ab_ltr | DN1cd | -0.97 | -1.8 | 0.11 | 0.41 | 0.0045 | ENSMUSG00000110887 | processed_pseudogene | Cbx3-ps8 | chromobox 3, pseudogene 8 | NA | 9 | 7723147 | 7723694 | 1 | |
| 2.79 | 4.08 | 1.74 | 3.93 | DN1ab_ltr | DN1cd | 1.59 | 8.62 | 5.40E-05 | 0.0084 | 0.0045 | ENSMUSG00000072568 | protein_coding | Fam84b | family with sequence similarity 84, member B | 399603 | 15 | 60818994 | 60853778 | -1 | Mm.411968 |
| 5.95 | 5.34 | 6.94 | 5.23 | DN1ab_ltr | DN1cd | -1.04 | -4.44 | 0.003 | 0.054 | 0.0046 | ENSMUSG00000022791 | protein_coding | Tnk2 | tyrosine kinase, non-receptor, 2 | 51789 | 16 | 32643874 | 32683493 | 1 | Mm.251115 |
| -3.82 | -1.54 | -3.94 | -4.02 | DN1ab_ltr | DN1cd | 1.42 | 2.88 | 0.023 | 0.18 | 0.0047 | ENSMUSG00000080748 | processed_pseudogene | Gm15661 | predicted gene 15661 | NA | 7 | 88448675 | 88449230 | -1 | |
| 5.34 | 5.3 | 5.78 | 5.22 | DN1ab_ltr | DN1cd | -0.24 | -2.22 | 0.062 | 0.29 | 0.0048 | ENSMUSG00000028668 | protein_coding | Eloa | elongin A | 27224 | 4 | 136003368 | 136021763 | -1 | Mm.27663 |
| -3.14 | -1.88 | -3.94 | 1.5 | DN1ab_ltr | DN1cd | 2.95 | 3.3 | 0.013 | 0.13 | 0.0048 | ENSMUSG00000028544 | protein_coding | Slc5a9 | solute carrier family 5 (sodium/glucose cotransporter), member 9 | 230612 | 4 | 111875375 | 111902918 | -1 | Mm.26630 |
| -3.82 | -3.86 | -1.62 | -4.02 | DN1ab_ltr | DN1cd | -0.89 | -1.8 | 0.11 | 0.41 | 0.0049 | ENSMUSG00000097040 | processed_transcript | 2610316D01Rik | RIKEN cDNA 2610316D01 gene | NA | 3 | 45280437 | 45378260 | -1 | |
| -3.82 | -3.86 | -1.62 | -4.02 | DN1ab_ltr | DN1cd | -0.89 | -1.8 | 0.11 | 0.41 | 0.0049 | ENSMUSG00000107363 | TEC | Gm43373 | predicted gene 43373 | NA | 5 | 69217421 | 69220290 | -1 | |
| -3.82 | -3.86 | -1.62 | -4.02 | DN1ab_ltr | DN1cd | -0.89 | -1.8 | 0.11 | 0.41 | 0.0049 | ENSMUSG00000076483 | TR_J_gene | Trbj1-1 | T cell receptor beta joining 1-1 | NA | 6 | 41533864 | 41533911 | 1 | |
| -3.82 | -3.86 | -1.62 | -4.02 | DN1ab_ltr | DN1cd | -0.89 | -1.8 | 0.11 | 0.41 | 0.0049 | ENSMUSG00000109754 | lincRNA | Gm39214 | predicted gene, 39214 | NA | 8 | 85730816 | 85765095 | 1 | |
| -3.82 | -3.86 | -1.62 | -4.02 | DN1ab_ltr | DN1cd | -0.89 | -1.8 | 0.11 | 0.41 | 0.0049 | ENSMUSG00000088184 | snRNA | Gm23137 | predicted gene, 23137 | NA | 11 | 87444474 | 87444580 | -1 | |
| -3.82 | -3.86 | -1.62 | -4.02 | DN1ab_ltr | DN1cd | -0.89 | -1.8 | 0.11 | 0.41 | 0.0049 | ENSMUSG00000055782 | protein_coding | Abcd2 | ATP-binding cassette, sub-family D (ALD), member 2 | 26874 | 15 | 91145871 | 91191799 | -1 | Mm.295456 |
| -1.5 | -3.86 | -0.47 | -0.22 | DN1ab_ltr | DN1cd | -1.36 | -2.45 | 0.044 | 0.24 | 0.0049 | ENSMUSG00000036196 | protein_coding | Slc26a8 | solute carrier family 26, member 8 | 224661 | 17 | 28637783 | 28690233 | -1 | |
| 5.69 | 6.37 | 4.42 | 6.9 | DN1ab_ltr | DN1cd | 1.32 | 3.51 | 0.0098 | 0.11 | 0.005 | ENSMUSG00000020323 | protein_coding | Prss57 | protease, serine 57 | 73106 | 10 | 79781474 | 79790961 | -1 | Mm.491308 |
| -3.82 | -3.86 | -1.3 | -4.02 | DN1ab_ltr | DN1cd | -1 | -1.79 | 0.12 | 0.42 | 0.005 | ENSMUSG00000040009 | protein_coding | Gnaz | guanine nucleotide binding protein, alpha z subunit | 14687 | 10 | 74967177 | 75016907 | 1 | Mm.32595 Mm.393508 |
| -3.82 | -3.86 | -1.3 | -4.02 | DN1ab_ltr | DN1cd | -1 | -1.79 | 0.12 | 0.42 | 0.005 | ENSMUSG00000040828 | protein_coding | Catsperd | cation channel sperm associated auxiliary subunit delta | 106757 | 17 | 56628143 | 56664456 | 1 | Mm.23401 |
| -2.24 | -2.28 | -1.62 | -4.02 | DN1ab_ltr | DN1cd | -0.9 | -1.82 | 0.11 | 0.41 | 0.0051 | ENSMUSG00000098428 | unprocessed_pseudogene | Gm2260 | predicted gene 2260 | 100039484 | 14 | 26016431 | 26025837 | 1 | |
| 4.12 | 4.74 | 3.24 | 4.95 | DN1ab_ltr | DN1cd | 1 | 4.38 | 0.0032 | 0.057 | 0.0051 | ENSMUSG00000013846 | protein_coding | St3gal1 | ST3 beta-galactoside alpha-2,3-sialyltransferase 1 | 20442 | 15 | 67102875 | 67176830 | -1 | Mm.248334 Mm.442417 Mm.461082 |
| 7.86 | 7.48 | 8.76 | 7.78 | DN1ab_ltr | DN1cd | -0.61 | -4.82 | 0.0019 | 0.044 | 0.0052 | ENSMUSG00000030830 | protein_coding | Itgal | integrin alpha L | 16408 | 7 | 127296260 | 127335138 | 1 | Mm.1618 |
| 5.02 | 6.24 | 3.91 | 6.72 | DN1ab_ltr | DN1cd | 1.77 | 5.34 | 0.001 | 0.033 | 0.0052 | ENSMUSG00000021211 | protein_coding | Akr1c12 | aldo-keto reductase family 1, member C12 | 622402 | 13 | 4268176 | 4279433 | -1 | Mm.27447 Mm.380356 |
| -2.91 | -3.86 | 0.95 | -4.02 | DN1ab_ltr | DN1cd | -2.4 | -2.84 | 0.025 | 0.18 | 0.0053 | ENSMUSG00000021587 | protein_coding | Pcsk1 | proprotein convertase subtilisin/kexin type 1 | 18548 | 13 | 75089826 | 75134861 | 1 | Mm.1333 Mm.394672 |
| 4.04 | 4.59 | 3.95 | 5.42 | DN1ab_ltr | DN1cd | 0.9 | 4.66 | 0.0023 | 0.048 | 0.0053 | ENSMUSG00000001143 | protein_coding | Lman2l | lectin, mannose-binding 2-like | 214895 | 1 | 36419871 | 36445271 | -1 | Mm.287854 |
| 11.28 | 11.4 | 11.1 | 11.65 | DN1ab_ltr | DN1cd | 0.28 | 3.06 | 0.018 | 0.15 | 0.0053 | ENSMUSG00000064370 | protein_coding | mt-Cytb | mitochondrially encoded cytochrome b | 17711 | MT | 14145 | 15288 | 1 | |
| -0.36 | -0.18 | -3.94 | 0.63 | DN1ab_ltr | DN1cd | 1.74 | 1.9 | 0.099 | 0.39 | 0.0054 | ENSMUSG00000090214 | antisense | Gm15657 | predicted gene 15657 | NA | 16 | 33770430 | 33796874 | -1 | |
| 3.61 | 3.8 | 3.43 | 5.03 | DN1ab_ltr | DN1cd | 0.74 | 2.46 | 0.043 | 0.24 | 0.0054 | ENSMUSG00000003992 | protein_coding | Ssbp2 | single-stranded DNA binding protein 2 | 66970 | 13 | 91460283 | 91703429 | 1 | Mm.343095 |
| 7.05 | 8.12 | 6.38 | 8.24 | DN1ab_ltr | DN1cd | 1.35 | 8.07 | 8.30E-05 | 0.011 | 0.0054 | ENSMUSG00000015143 | protein_coding | Actn1 | actinin, alpha 1 | 109711 | 12 | 80167547 | 80260371 | -1 | Mm.253564 Mm.403477 |
| 3.22 | 3.86 | 2.8 | 4.74 | DN1ab_ltr | DN1cd | 1.12 | 4.06 | 0.0047 | 0.072 | 0.0054 | ENSMUSG00000052889 | protein_coding | Prkcb | protein kinase C, beta | 18751 | 7 | 122288751 | 122634402 | 1 | Mm.207496 Mm.446371 |
| -1.29 | -3.86 | -3.94 | -4.02 | DN1ab_ltr | DN1cd | -1.67 | -3.19 | 0.015 | 0.14 | 0.0055 | ENSMUSG00000048215 | lincRNA | A630023P12Rik | RIKEN cDNA A630023P12 gene | NA | 5 | 110514923 | 110525980 | 1 | |
| -1.29 | -3.86 | -3.94 | -4.02 | DN1ab_ltr | DN1cd | -1.67 | -3.19 | 0.015 | 0.14 | 0.0055 | ENSMUSG00000105345 | lincRNA | BC030343 | cDNA sequence BC030343 | NA | 5 | 143270075 | 143272513 | 1 | |
| -1.29 | -3.86 | -3.94 | -4.02 | DN1ab_ltr | DN1cd | -1.67 | -3.19 | 0.015 | 0.14 | 0.0055 | ENSMUSG00000089663 | antisense | Gm15651 | predicted gene 15651 | NA | 16 | 22548553 | 22551149 | 1 | |
| 5.19 | 5.32 | 4.75 | 5.47 | DN1ab_ltr | DN1cd | 0.35 | 2.83 | 0.025 | 0.18 | 0.0055 | ENSMUSG00000038503 | protein_coding | Mesd | mesoderm development LRP chaperone | 67943 | 7 | 83884466 | 83901532 | 1 | Mm.117365 |
| 2.02 | 2.4 | 0.72 | 3.15 | DN1ab_ltr | DN1cd | 1.13 | 2.61 | 0.035 | 0.22 | 0.0055 | ENSMUSG00000024008 | protein_coding | Cpne5 | copine V | 240058 | 17 | 29155556 | 29238167 | -1 | Mm.39905 |
| 3.44 | 3.49 | 2.95 | 3.88 | DN1ab_ltr | DN1cd | 0.38 | 2.04 | 0.081 | 0.34 | 0.0055 | ENSMUSG00000028035 | protein_coding | Dnajb4 | DnaJ heat shock protein family (Hsp40) member B4 | 67035 | 3 | 152178511 | 152210302 | -1 | Mm.46746 Mm.458538 |
| 1.42 | 3.22 | 2.78 | 2.6 | DN1ab_ltr | DN1cd | 1.15 | 2.83 | 0.025 | 0.18 | 0.0055 | ENSMUSG00000000531 | protein_coding | Grasp | GRP1 (general receptor for phosphoinositides 1)-associated scaffold protein | 56149 | 15 | 101224188 | 101232756 | 1 | Mm.276573 |
| -3.82 | -3.86 | 1.62 | -3.24 | DN1ab_ltr | DN1cd | -1.76 | -1.74 | 0.12 | 0.43 | 0.0057 | ENSMUSG00000035459 | protein_coding | Stab2 | stabilin 2 | 192188 | 10 | 86841198 | 87008025 | -1 | Mm.279611 |
| 1.65 | 2.08 | 3.13 | 4.79 | DN1ab_ltr | DN1cd | 0.94 | 3.57 | 0.0089 | 0.1 | 0.0057 | ENSMUSG00000104713 | protein_coding | Gbp6 | guanylate binding protein 6 | 100702 | 5 | 105270702 | 105293698 | -1 | Mm.491210 |
| -1.22 | -1.14 | -3.94 | -1.03 | DN1ab_ltr | DN1cd | 1.11 | 1.86 | 0.1 | 0.4 | 0.0058 | ENSMUSG00000105945 | antisense | Gm43570 | predicted gene 43570 | NA | 3 | 59002471 | 59006158 | -1 | |
| 5.5 | 5.55 | 5.32 | 6.15 | DN1ab_ltr | DN1cd | 0.34 | 2.06 | 0.078 | 0.34 | 0.0058 | ENSMUSG00000029313 | protein_coding | Aff1 | AF4/FMR2 family, member 1 | 17355 | 5 | 103692374 | 103855322 | 1 | Mm.6949 |
| 5.36 | 5.7 | 4.81 | 5.84 | DN1ab_ltr | DN1cd | 0.59 | 4.02 | 0.005 | 0.074 | 0.0059 | ENSMUSG00000028385 | protein_coding | Snx30 | sorting nexin family member 30 | 209131 | 4 | 59805840 | 59904737 | 1 | Mm.259705 Mm.490979 |
| 5.5 | 5.23 | 6.47 | 5.42 | DN1ab_ltr | DN1cd | -0.57 | -3.44 | 0.011 | 0.12 | 0.0059 | ENSMUSG00000012609 | protein_coding | Ttll5 | tubulin tyrosine ligase-like family, member 5 | 320244 | 12 | 85824659 | 86061893 | 1 | Mm.132172 |
| -3.82 | -1.28 | -3.94 | -4.02 | DN1ab_ltr | DN1cd | 1.59 | 2.88 | 0.024 | 0.18 | 0.006 | ENSMUSG00000025666 | protein_coding | Tmem47 | transmembrane protein 47 | 192216 | X | 81070698 | 81097872 | 1 | Mm.490421 |
| 3.71 | 2.65 | 6.47 | 3.84 | DN1ab_ltr | DN1cd | -1.83 | -5.29 | 0.0011 | 0.034 | 0.006 | ENSMUSG00000032323 | protein_coding | Cyp11a1 | cytochrome P450, family 11, subfamily a, polypeptide 1 | 13070 | 9 | 58006411 | 58027023 | 1 | Mm.302865 |
| 3.85 | 3.99 | 2.95 | 4.23 | DN1ab_ltr | DN1cd | 0.56 | 2.32 | 0.053 | 0.27 | 0.006 | ENSMUSG00000089774 | protein_coding | Slc5a3 | solute carrier family 5 (inositol transporters), member 3 | 53881 | 16 | 92058322 | 92087473 | 1 | Mm.217354 Mm.451146 |
| 8.27 | 8.51 | 8.14 | 8.92 | DN1ab_ltr | DN1cd | 0.44 | 3.82 | 0.0065 | 0.087 | 0.0061 | ENSMUSG00000030067 | protein_coding | Foxp1 | forkhead box P1 | 108655 | 6 | 98925338 | 99522721 | -1 | Mm.234965 Mm.392884 Mm.485266 Mm.487716 |
| -1.01 | -3.86 | -3.94 | -4.02 | DN1ab_ltr | DN1cd | -1.85 | -3.15 | 0.016 | 0.14 | 0.0061 | ENSMUSG00000008489 | protein_coding | Elavl2 | ELAV like RNA binding protein 1 | 15569 | 4 | 91250763 | 91400785 | -1 | Mm.318042 |
| -1.01 | -3.86 | -3.94 | -4.02 | DN1ab_ltr | DN1cd | -1.85 | -3.15 | 0.016 | 0.14 | 0.0061 | ENSMUSG00000053528 | lincRNA | A530021J07Rik | Riken cDNA A530021J07 gene | NA | 7 | 83061783 | 83179429 | 1 | |
| 6.85 | 7.63 | 6.29 | 7.99 | DN1ab_ltr | DN1cd | 1.11 | 5.67 | 0.00074 | 0.028 | 0.0061 | ENSMUSG00000020577 | protein_coding | Tspan13 | tetraspanin 13 | 66109 | 12 | 36014557 | 36042500 | -1 | Mm.254663 |
| 2.93 | 4.61 | 1.22 | 4.43 | DN1ab_ltr | DN1cd | 2.17 | 6.86 | 0.00023 | 0.016 | 0.0061 | ENSMUSG00000066235 | protein_coding | Pomgnt2 | protein O-linked mannose beta 1,4-N-acetylglucosaminyltransferase 2 | 215494 | 9 | 121981606 | 121997110 | -1 | Mm.402151 |
| 4.27 | 4.02 | 5.54 | 4.33 | DN1ab_ltr | DN1cd | -0.64 | -3.05 | 0.019 | 0.16 | 0.0061 | ENSMUSG00000021196 | protein_coding | Pfkp | phosphofructokinase, platelet | 56421 | 13 | 6579768 | 6648777 | -1 | Mm.273874 |
| -1.81 | 0.21 | -3.94 | 0.88 | DN1ab_ltr | DN1cd | 2.98 | 5.09 | 0.0014 | 0.037 | 0.0064 | ENSMUSG00000050138 | protein_coding | Kcnk12 | potassium channel, subfamily K, member 12 | 210741 | 17 | 87745801 | 87797994 | -1 | Mm.312046 |
| 6.29 | 6.38 | 6 | 6.71 | DN1ab_ltr | DN1cd | 0.32 | 2.47 | 0.043 | 0.24 | 0.0064 | ENSMUSG00000037111 | protein_coding | Setd7 | SET domain containing (lysine methyltransferase) 7 | 73251 | 3 | 51515319 | 51560879 | -1 | Mm.192111 |
| 4.14 | 4.27 | 3.42 | 4.36 | DN1ab_ltr | DN1cd | 0.42 | 2.47 | 0.043 | 0.24 | 0.0065 | ENSMUSG00000034601 | protein_coding | 2700049A03Rik | RIKEN cDNA 2700049A03 gene | 76967 | 12 | 71136848 | 71243303 | 1 | Mm.87290 |
| 6.9 | 6.37 | 6.69 | 6.66 | DN1ab_ltr | DN1cd | -0.34 | -3.23 | 0.014 | 0.13 | 0.0065 | ENSMUSG00000024095 | protein_coding | Hnrnpll | heterogeneous nuclear ribonucleoprotein L-like | 72692 | 17 | 80029487 | 80062334 | -1 | Mm.64579 |
| 4.18 | 3.75 | 5.33 | 4.16 | DN1ab_ltr | DN1cd | -0.74 | -4.66 | 0.0023 | 0.048 | 0.0066 | ENSMUSG00000020097 | protein_coding | Sgpl1 | sphingosine phosphate lyase 1 | 20397 | 10 | 61098642 | 61147703 | -1 | Mm.412319 Mm.431862 |
| 7.09 | 7.23 | 6.98 | 7.75 | DN1ab_ltr | DN1cd | 0.38 | 2.77 | 0.028 | 0.19 | 0.0066 | ENSMUSG00000062604 | protein_coding | Srpk2 | serine/arginine-rich protein specific kinase 2 | 20817 | 5 | 23503264 | 23684617 | -1 | Mm.288728 |
| 5.44 | 6.31 | 5.46 | 6.98 | DN1ab_ltr | DN1cd | 1.1 | 7.91 | 9.40E-05 | 0.011 | 0.0067 | ENSMUSG00000040253 | protein_coding | Gbp7 | guanylate binding protein 7 | 229900 | 3 | 142530342 | 142550149 | 1 | Mm.45740 |
| 6.33 | 6.54 | 6.25 | 6.92 | DN1ab_ltr | DN1cd | 0.38 | 3.8 | 0.0066 | 0.088 | 0.0068 | ENSMUSG00000039968 | protein_coding | Rsbn1l | round spermatid basic protein 1-like | 242860 | 5 | 20893028 | 20956398 | -1 | Mm.254286 |
| 5.43 | 6.21 | 4.61 | 6.5 | DN1ab_ltr | DN1cd | 1.18 | 4.99 | 0.0016 | 0.04 | 0.0069 | ENSMUSG00000000303 | protein_coding | Cdh1 | cadherin 1 | 12550 | 8 | 106603351 | 106670246 | 1 | Mm.35605 |
| 4.4 | 5.03 | 4.14 | 5.82 | DN1ab_ltr | DN1cd | 1.02 | 4.52 | 0.0027 | 0.052 | 0.0069 | ENSMUSG00000037972 | protein_coding | Snn | stannin | 20621 | 16 | 11060945 | 11074985 | 1 | Mm.325800 |
| 5.98 | 5.36 | 6.82 | 5.74 | DN1ab_ltr | DN1cd | -0.8 | -7.94 | 9.20E-05 | 0.011 | 0.007 | ENSMUSG00000015966 | protein_coding | Il17rb | interleukin 17 receptor B | 50905 | 14 | 29996135 | 30008896 | -1 | Mm.269363 |
| -2.91 | -1.31 | -3.94 | 1.26 | DN1ab_ltr | DN1cd | 3.03 | 3.88 | 0.006 | 0.083 | 0.007 | ENSMUSG00000020811 | protein_coding | Wscd1 | WSC domain containing 1 | 216881 | 11 | 71749920 | 71789647 | 1 | Mm.48648 |
| 9.12 | 9.02 | 10.04 | 9.37 | DN1ab_ltr | DN1cd | -0.32 | -2.58 | 0.036 | 0.22 | 0.0071 | ENSMUSG00000039936 | protein_coding | Pik3cd | phosphatidylinositol-4,5-bisphosphate 3-kinase catalytic subunit delta | 18707 | 4 | 149649168 | 149702571 | -1 | Mm.229108 |
| 8.78 | 9.23 | 8.03 | 9.21 | DN1ab_ltr | DN1cd | 0.71 | 4.57 | 0.0025 | 0.05 | 0.0071 | ENSMUSG00000045817 | protein_coding | Zfp36l2 | zinc finger protein 36, C3H type-like 2 | 12193 | 17 | 84183931 | 84187947 | -1 | Mm.259321 |
| 0.44 | 0.72 | -1.83 | 1.47 | DN1ab_ltr | DN1cd | 1.37 | 2.13 | 0.071 | 0.32 | 0.0071 | ENSMUSG00000046031 | protein_coding | Calhm6 | calcium homeostasis modulator family member 6 | 215900 | 10 | 34126067 | 34127984 | -1 | Mm.34479 |
| 0.93 | 2.3 | -0.16 | 3.94 | DN1ab_ltr | DN1cd | 2.31 | 4 | 0.0051 | 0.075 | 0.0072 | ENSMUSG00000059305 | protein_coding | Vpreb1 | pre-B lymphocyte gene 1 | 22362 | 16 | 16868403 | 16870843 | -1 | Mm.57574 |
| 6.33 | 5.98 | 7.27 | 6.2 | DN1ab_ltr | DN1cd | -0.63 | -4.09 | 0.0046 | 0.07 | 0.0073 | ENSMUSG00000021109 | protein_coding | Hif1a | hypoxia inducible factor 1, alpha subunit | 15251 | 12 | 73901375 | 73947530 | 1 | Mm.3879 Mm.446610 |
| 4.2 | 4.91 | 3.28 | 5.11 | DN1ab_ltr | DN1cd | 1.11 | 4.66 | 0.0023 | 0.048 | 0.0073 | ENSMUSG00000025921 | protein_coding | Rdh10 | retinol dehydrogenase 10 (all-trans) | 98711 | 1 | 16105774 | 16133734 | 1 | Mm.274376 |
| -1.14 | -3.86 | -3.94 | -4.02 | DN1ab_ltr | DN1cd | -1.77 | -3.13 | 0.017 | 0.15 | 0.0074 | ENSMUSG00000110018 | transcribed_processed_pseudogene | 5430437J10Rik | RIKEN cDNA 5430437J10 gene | NA | 15 | 5496266 | 5594984 | 1 | |
| 4.71 | 4.62 | 6.25 | 5.35 | DN1ab_ltr | DN1cd | -0.41 | -2.35 | 0.051 | 0.26 | 0.0075 | ENSMUSG00000045763 | protein_coding | Basp1 | brain abundant, membrane attached signal protein 1 | 70350 | 15 | 25363277 | 25413764 | -1 | Mm.29586 |
| 5.01 | 5.39 | 3.96 | 5.41 | DN1ab_ltr | DN1cd | 0.76 | 3.34 | 0.012 | 0.12 | 0.0076 | ENSMUSG00000085795 | protein_coding | Zfp703 | zinc finger protein 703 | 353310 | 8 | 26977325 | 26981461 | 1 | Mm.212469 |
| -1.54 | -0.32 | -3.49 | 1.96 | DN1ab_ltr | DN1cd | 2.77 | 3.04 | 0.019 | 0.16 | 0.0078 | ENSMUSG00000034009 | protein_coding | Rxfp1 | relaxin/insulin-like family peptide receptor 1 | 381489 | 3 | 79641611 | 79737880 | -1 | Mm.331086 |
| -0.65 | -3.86 | 0.09 | -0.39 | DN1ab_ltr | DN1cd | -2.1 | -3.51 | 0.0098 | 0.11 | 0.0079 | ENSMUSG00000097261 | lincRNA | 4930532G15Rik | RIKEN cDNA 4930532G15 gene | NA | 1 | 185359461 | 185362865 | -1 | |
| 4.23 | 4.14 | 4.62 | 3.64 | DN1ab_ltr | DN1cd | -0.42 | -2.18 | 0.065 | 0.3 | 0.0079 | ENSMUSG00000044447 | protein_coding | Dock5 | dedicator of cytokinesis 5 | 68813 | 14 | 67752135 | 67933442 | -1 | Mm.258155 |
| 5.69 | 5.83 | 5.6 | 6.36 | DN1ab_ltr | DN1cd | 0.37 | 2.76 | 0.028 | 0.19 | 0.0079 | ENSMUSG00000041777 | protein_coding | Cir1 | corepressor interacting with RBPJ, 1 | 66935 | 2 | 73283105 | 73312701 | -1 | Mm.292140 Mm.361271 |
| 5.73 | 5.19 | 8.45 | 6.3 | DN1ab_ltr | DN1cd | -1.22 | -3.41 | 0.011 | 0.12 | 0.008 | ENSMUSG00000041954 | protein_coding | Tnfrsf18 | tumor necrosis factor receptor superfamily, member 18 | 21936 | 4 | 156026164 | 156028895 | 1 | Mm.3180 |
| 5.22 | 4.54 | 5.66 | 5.43 | DN1ab_ltr | DN1cd | -0.49 | -4.92 | 0.0017 | 0.041 | 0.008 | ENSMUSG00000079685 | protein_coding | Ulbp1 | UL16 binding protein 1 | 77777 | 10 | 7440362 | 7473638 | -1 | Mm.442442 |
| 5.22 | 5.44 | 4.54 | 5.59 | DN1ab_ltr | DN1cd | 0.53 | 2.98 | 0.02 | 0.16 | 0.0081 | ENSMUSG00000028378 | protein_coding | Ptgr1 | prostaglandin reductase 1 | 67103 | 4 | 58965439 | 58987119 | -1 | Mm.34497 |
| 2.08 | 1.71 | 4.16 | 2.33 | DN1ab_ltr | DN1cd | -1 | -3.07 | 0.018 | 0.15 | 0.0082 | ENSMUSG00000002983 | protein_coding | Relb | avian reticuloendotheliosis viral (v-rel) oncogene related B | 19698 | 7 | 19606217 | 19629438 | -1 | Mm.1741 |
| 4.21 | 3.78 | 5.15 | 3.58 | DN1ab_ltr | DN1cd | -0.88 | -3.56 | 0.0091 | 0.1 | 0.0082 | ENSMUSG00000038843 | protein_coding | Gcnt1 | glucosaminyl (N-acetyl) transferase 1, core 2 | 14537 | 19 | 17326141 | 17356667 | -1 | Mm.244825 |
| -0.17 | 0.52 | -3.5 | 0.99 | DN1ab_ltr | DN1cd | 1.98 | 2.45 | 0.044 | 0.24 | 0.0082 | ENSMUSG00000035578 | protein_coding | Iqcg | IQ motif containing G | 69707 | 16 | 33012683 | 33056218 | -1 | Mm.227918 |
| 5.67 | 6.81 | 4.16 | 7.04 | DN1ab_ltr | DN1cd | 1.74 | 4.69 | 0.0022 | 0.047 | 0.0084 | ENSMUSG00000022824 | protein_coding | Muc13 | mucin 13, epithelial transmembrane | 17063 | 16 | 33794037 | 33819934 | 1 | Mm.3177 |
| 6.05 | 5.71 | 6.81 | 5.93 | DN1ab_ltr | DN1cd | -0.54 | -4.56 | 0.0026 | 0.051 | 0.0084 | ENSMUSG00000020743 | protein_coding | Mif4gd | MIF4G domain containing | 69674 | 11 | 115607918 | 115612969 | -1 | Mm.390387 Mm.489360 |
| 5.33 | 5.82 | 4.75 | 5.75 | DN1ab_ltr | DN1cd | 0.67 | 6.05 | 5.00E-04 | 0.023 | 0.0084 | ENSMUSG00000027459 | protein_coding | Fam110a | family with sequence similarity 110, member A | 73847 | 2 | 151969398 | 151980219 | -1 | Mm.272444 Mm.404253 |
| 4.11 | 6.67 | 1.88 | 6.88 | DN1ab_ltr | DN1cd | 3.34 | 6.54 | 0.00031 | 0.019 | 0.0085 | ENSMUSG00000027858 | protein_coding | Tspan2 | tetraspanin 2 | 70747 | 3 | 102734529 | 102801513 | 1 | Mm.27469 |
| -0.5 | -3.86 | -1.24 | -1.28 | DN1ab_ltr | DN1cd | -2.09 | -2.89 | 0.023 | 0.17 | 0.0085 | ENSMUSG00000019178 | protein_coding | Styxl1 | serine/threonine/tyrosine interacting-like 1 | 76571 | 5 | 135747220 | 135778385 | -1 | Mm.247666 |
| 5.96 | 6.4 | 4.94 | 6.06 | DN1ab_ltr | DN1cd | 0.68 | 4.77 | 0.002 | 0.045 | 0.0086 | ENSMUSG00000024538 | protein_coding | Ppic | peptidylprolyl isomerase C | 19038 | 18 | 53406341 | 53418007 | -1 | Mm.4587 |
| 5.21 | 5.48 | 5.11 | 6.34 | DN1ab_ltr | DN1cd | 0.63 | 3.01 | 0.02 | 0.16 | 0.0086 | ENSMUSG00000024975 | protein_coding | Pdcd4 | programmed cell death 4 | 18569 | 19 | 53892231 | 53929861 | 1 | Mm.1605 |
| 6.37 | 5.79 | 7.7 | 6.35 | DN1ab_ltr | DN1cd | -0.89 | -5.24 | 0.0012 | 0.034 | 0.0086 | ENSMUSG00000032479 | protein_coding | Map4 | microtubule-associated protein 4 | 17758 | 9 | 109931460 | 110083955 | 1 | Mm.217318 Mm.443428 Mm.484451 |
| 2.89 | 3.59 | 2.92 | 4.52 | DN1ab_ltr | DN1cd | 1.03 | 5.32 | 0.0011 | 0.033 | 0.0086 | ENSMUSG00000026919 | protein_coding | Lcn4 | lipocalin 4 | 16821 | 2 | 26667674 | 26671282 | -1 | Mm.123925 |
| 3.11 | 3.48 | 2.56 | 4.22 | DN1ab_ltr | DN1cd | 0.85 | 3.04 | 0.019 | 0.16 | 0.0086 | ENSMUSG00000059336 | protein_coding | Slc14a1 | solute carrier family 14 (urea transporter), member 1 | 108052 | 18 | 78100091 | 78142119 | -1 | Mm.33832 |
| 1.66 | 2.01 | -1.27 | 1.51 | DN1ab_ltr | DN1cd | 1.16 | 2.26 | 0.058 | 0.28 | 0.0087 | ENSMUSG00000026180 | protein_coding | Cxcr2 | chemokine (C-X-C motif) receptor 2 | 12765 | 1 | 74153989 | 74161246 | 1 | Mm.234466 |
| 3.93 | 4.85 | 2.78 | 4.89 | DN1ab_ltr | DN1cd | 1.33 | 5.14 | 0.0013 | 0.036 | 0.0087 | ENSMUSG00000058794 | protein_coding | Nfe2 | nuclear factor, erythroid derived 2 | 18022 | 15 | 103248212 | 103258403 | -1 | Mm.457989 Mm.489030 |
| 2.39 | 2.18 | 3.05 | 1.73 | DN1ab_ltr | DN1cd | -0.63 | -2.6 | 0.035 | 0.22 | 0.0088 | ENSMUSG00000038618 | protein_coding | Rassf7 | Ras association (RalGDS/AF-6) domain family (N-terminal) member 7 | 66985 | 7 | 141215860 | 141218658 | 1 | Mm.21202 |
| 1.54 | 2.11 | 0.94 | 3.07 | DN1ab_ltr | DN1cd | 1.15 | 3.37 | 0.012 | 0.12 | 0.009 | ENSMUSG00000003206 | protein_coding | Ebi3 | Epstein-Barr virus induced gene 3 | 50498 | 17 | 55952640 | 55957022 | 1 | Mm.256798 |
| 7.07 | 7.33 | 6.68 | 7.6 | DN1ab_ltr | DN1cd | 0.5 | 3.51 | 0.0097 | 0.11 | 0.009 | ENSMUSG00000030275 | protein_coding | Etnk1 | ethanolamine kinase 1 | 75320 | 6 | 143167221 | 143232307 | 1 | Mm.272548 |
| 5.73 | 5.4 | 6.4 | 5.62 | DN1ab_ltr | DN1cd | -0.5 | -5.07 | 0.0014 | 0.038 | 0.009 | ENSMUSG00000039841 | protein_coding | Zfp800 | zinc finger protein 800 | 627049 | 6 | 28239927 | 28398005 | -1 | Mm.84441 Mm.393511 |
| 4.17 | 3.65 | 5.59 | 4.28 | DN1ab_ltr | DN1cd | -0.85 | -4.81 | 0.0019 | 0.044 | 0.0091 | ENSMUSG00000110206 | protein_coding | Flt3l | FMS-like tyrosine kinase 3 ligand | 14256 | 7 | 45129788 | 45136432 | -1 | Mm.1658 |
| 4.15 | 5.78 | 2.93 | 6.28 | DN1ab_ltr | DN1cd | 2.21 | 6.04 | 0.00051 | 0.023 | 0.0091 | ENSMUSG00000028434 | protein_coding | Epb41l4b | erythrocyte membrane protein band 4.1 like 4b | 54357 | 4 | 56991972 | 57143437 | -1 | Mm.28217 Mm.381403 |
| 7.17 | 8.31 | 6.07 | 9.2 | DN1ab_ltr | DN1cd | 1.85 | 4.32 | 0.0034 | 0.06 | 0.0095 | ENSMUSG00000005474 | protein_coding | Myl10 | myosin, light chain 10, regulatory | 59310 | 5 | 136693146 | 136701094 | 1 | Mm.5178 |
| 0.74 | 1.9 | -1.14 | 2.3 | DN1ab_ltr | DN1cd | 1.93 | 3.96 | 0.0054 | 0.078 | 0.0096 | ENSMUSG00000056130 | protein_coding | Ticam2 | toll-like receptor adaptor molecule 2 | 225471 | 18 | 46559155 | 46574533 | -1 | Mm.149280 |
| 7.88 | 7.39 | 9.26 | 7.92 | DN1ab_ltr | DN1cd | -0.82 | -4.36 | 0.0033 | 0.058 | 0.0098 | ENSMUSG00000049866 | protein_coding | Arl4c | ADP-ribosylation factor-like 4C | 320982 | 1 | 88673125 | 88702221 | -1 | Mm.27968 Mm.477970 |
| 4.7 | 4.43 | 5.94 | 5.05 | DN1ab_ltr | DN1cd | -0.53 | -3.84 | 0.0063 | 0.085 | 0.0098 | ENSMUSG00000054065 | protein_coding | Pkp3 | plakophilin 3 | 56460 | 7 | 141078218 | 141090510 | 1 | Mm.350037 |
| 5.72 | 6.08 | 5.45 | 6.37 | DN1ab_ltr | DN1cd | 0.57 | 4.61 | 0.0024 | 0.049 | 0.0099 | ENSMUSG00000009681 | protein_coding | Bcr | breakpoint cluster region | 110279 | 10 | 75060592 | 75184921 | 1 | Mm.333722 |
| -1.92 | -0.55 | -3.94 | -0.12 | DN1ab_ltr | DN1cd | 2.24 | 4.21 | 0.0039 | 0.064 | 0.01 | ENSMUSG00000001023 | protein_coding | S100a5 | S100 calcium binding protein A5 | 20199 | 3 | 90608523 | 90611780 | 1 | Mm.19946 |
| 4.79 | 5.43 | 3.73 | 5.4 | DN1ab_ltr | DN1cd | 1 | 4.49 | 0.0028 | 0.053 | 0.01 | ENSMUSG00000034330 | protein_coding | Plcg2 | phospholipase C, gamma 2 | 234779 | 8 | 117498291 | 117635142 | 1 | Mm.192699 |
| 5.66 | 6.14 | 5.05 | 6.59 | DN1ab_ltr | DN1cd | 0.87 | 3.79 | 0.0067 | 0.088 | 0.01 | ENSMUSG00000021871 | protein_coding | Gm49342 | predicted gene, 49342 | NA | 14 | 50944517 | 50963136 | 1 | |
| 7.9 | 6.85 | 8.23 | 7.72 | DN1ab_ltr | DN1cd | -0.84 | -6.9 | 0.00022 | 0.016 | 0.01 | ENSMUSG00000018474 | protein_coding | Chd3 | chromodomain helicase DNA binding protein 3 | 216848 | 11 | 69343273 | 69369406 | -1 | Mm.178246 |
| -1.5 | -1.06 | -3.94 | -0.77 | DN1ab_ltr | DN1cd | 1.44 | 2.4 | 0.047 | 0.25 | 0.01 | ENSMUSG00000086331 | sense_intronic | Gm16310 | predicted gene 16310 | NA | 16 | 92094568 | 92096490 | 1 | |
| -1.14 | -0.7 | -3.29 | 0.6 | DN1ab_ltr | DN1cd | 1.73 | 2.27 | 0.057 | 0.28 | 0.011 | ENSMUSG00000016150 | protein_coding | Tenm1 | teneurin transmembrane protein 1 | 23963 | X | 42527866 | 43429126 | -1 | Mm.327698 |
| 5.97 | 5.43 | 5.78 | 5.7 | DN1ab_ltr | DN1cd | -0.36 | -3.52 | 0.0096 | 0.11 | 0.011 | ENSMUSG00000057506 | protein_coding | Bloc1s2 | biogenesis of lysosomal organelles complex-1, subunit 2 | 73689 | 19 | 44139266 | 44146446 | -1 | Mm.124685 |
| 2.45 | 3.22 | 0.78 | 2.89 | DN1ab_ltr | DN1cd | 1.23 | 4.26 | 0.0037 | 0.062 | 0.011 | ENSMUSG00000042190 | protein_coding | Cmklr1 | chemokine-like receptor 1 | 14747 | 5 | 113612354 | 113650426 | -1 | Mm.5196 |
| 3.66 | 2.36 | 3.1 | 2.93 | DN1ab_ltr | DN1cd | -0.8 | -3.11 | 0.017 | 0.15 | 0.011 | ENSMUSG00000022114 | protein_coding | Spry2 | sprouty RTK signaling antagonist 2 | 24064 | 14 | 105891947 | 105896819 | -1 | Mm.89982 Mm.489870 |
| 5.07 | 4.25 | 4.71 | 4.55 | DN1ab_ltr | DN1cd | -0.55 | -3.67 | 0.0079 | 0.097 | 0.011 | ENSMUSG00000016552 | protein_coding | Foxred2 | FAD-dependent oxidoreductase domain containing 2 | 239554 | 15 | 77940522 | 77956722 | -1 | Mm.295605 |
| 0.14 | 0.72 | -2.02 | 1.15 | DN1ab_ltr | DN1cd | 1.48 | 2.62 | 0.034 | 0.22 | 0.011 | ENSMUSG00000036295 | protein_coding | Lrrn3 | leucine rich repeat protein 3, neuronal | 16981 | 12 | 41451668 | 41486431 | -1 | Mm.404786 Mm.491383 |
| -0.86 | -0.18 | -3.94 | -0.59 | DN1ab_ltr | DN1cd | 1.58 | 2.73 | 0.029 | 0.2 | 0.011 | ENSMUSG00000082079 | protein_coding | Dnmt3c | DNA methyltransferase 3C | NA | 2 | 153696652 | 153729907 | 1 | |
| 3.85 | 4.36 | 3.66 | 4.79 | DN1ab_ltr | DN1cd | 0.73 | 5.34 | 0.0011 | 0.033 | 0.011 | ENSMUSG00000029291 | protein_coding | Rufy3 | RUN and FYVE domain containing 3 | 52822 | 5 | 88565040 | 88651392 | 1 | Mm.270469 |
| 5.11 | 3.52 | 4.9 | 4.42 | DN1ab_ltr | DN1cd | -1.09 | -4.25 | 0.0037 | 0.062 | 0.011 | ENSMUSG00000009894 | protein_coding | Snap47 | synaptosomal-associated protein, 47 | 67826 | 11 | 59407134 | 59451186 | -1 | Mm.26680 |
| 4.29 | 4.5 | 4.35 | 5.3 | DN1ab_ltr | DN1cd | 0.49 | 2.98 | 0.02 | 0.16 | 0.011 | ENSMUSG00000020564 | protein_coding | Atxn7l1 | ataxin 7-like 1 | 380753 | 12 | 33147693 | 33373185 | 1 | Mm.41540 Mm.352037 |
| 8.77 | 8.49 | 9.04 | 8.48 | DN1ab_ltr | DN1cd | -0.39 | -6.06 | 0.00049 | 0.023 | 0.012 | ENSMUSG00000000563 | protein_coding | Atp5f1 | ATP synthase, H+ transporting, mitochondrial F0 complex, subunit B1 | 11950 | 3 | 105942698 | 105960099 | -1 | Mm.251152 |
| 2.4 | 1.56 | 4.48 | 1.97 | DN1ab_ltr | DN1cd | -1.59 | -4.2 | 0.004 | 0.065 | 0.012 | ENSMUSG00000052512 | protein_coding | Nav2 | neuron navigator 2 | 78286 | 7 | 48908716 | 49610090 | 1 | Mm.333881 Mm.471488 |
| 1.62 | 2.17 | 0.5 | 2.19 | DN1ab_ltr | DN1cd | 0.95 | 3.82 | 0.0065 | 0.087 | 0.012 | ENSMUSG00000033697 | protein_coding | Arhgap39 | Rho GTPase activating protein 39 | 223666 | 15 | 76723985 | 76818170 | -1 | Mm.322931 |
| 0.29 | 1.07 | -0.88 | 2.14 | DN1ab_ltr | DN1cd | 1.58 | 3.21 | 0.015 | 0.14 | 0.012 | ENSMUSG00000032109 | protein_coding | Nlrx1 | NLR family member X1 | 270151 | 9 | 44252717 | 44268599 | -1 | Mm.3957 |
| -0.03 | 0.44 | -2.8 | 0.15 | DN1ab_ltr | DN1cd | 1.31 | 2.41 | 0.046 | 0.25 | 0.012 | ENSMUSG00000017950 | protein_coding | Hnf4a | hepatic nuclear factor 4, alpha | 15378 | 2 | 163506808 | 163572910 | 1 | Mm.202383 |
| 5.17 | 4.86 | 6.09 | 5.27 | DN1ab_ltr | DN1cd | -0.51 | -4.39 | 0.0031 | 0.056 | 0.012 | ENSMUSG00000022973 | protein_coding | Synj1 | synaptojanin 1 | 104015 | 16 | 90936092 | 91011308 | -1 | Mm.187079 |
| 6.24 | 6.69 | 6.47 | 6.47 | DN1ab_ltr | DN1cd | 0.29 | 2.97 | 0.021 | 0.16 | 0.012 | ENSMUSG00000020307 | protein_coding | Cdc34 | cell division cycle 34 | 216150 | 10 | 79682195 | 79688398 | 1 | Mm.21981 |
| 5.59 | 5.35 | 6.12 | 5.29 | DN1ab_ltr | DN1cd | -0.47 | -3.59 | 0.0087 | 0.1 | 0.012 | ENSMUSG00000021936 | protein_coding | Mapk8 | mitogen-activated protein kinase 8 | 26419 | 14 | 33377898 | 33447158 | -1 | Mm.21495 |
| 7.06 | 7.19 | 7.02 | 7.5 | DN1ab_ltr | DN1cd | 0.26 | 3.28 | 0.013 | 0.13 | 0.012 | ENSMUSG00000024816 | protein_coding | Frmd8 | FERM domain containing 8 | 67457 | 19 | 5849702 | 5875274 | -1 | Mm.202092 |
| 2.82 | 3.19 | 2.46 | 3.87 | DN1ab_ltr | DN1cd | 0.76 | 3.24 | 0.014 | 0.13 | 0.012 | ENSMUSG00000024669 | protein_coding | Cd5 | CD5 antigen | 12507 | 19 | 10718143 | 10738974 | -1 | Mm.779 |
| 4.13 | 4.46 | 3.73 | 4.84 | DN1ab_ltr | DN1cd | 0.62 | 3.6 | 0.0087 | 0.1 | 0.012 | ENSMUSG00000026727 | protein_coding | Rsu1 | Ras suppressor protein 1 | 20163 | 2 | 13076821 | 13271415 | -1 | Mm.905 |
| 4 | 3.68 | 5.3 | 3.72 | DN1ab_ltr | DN1cd | -0.82 | -2.9 | 0.023 | 0.17 | 0.012 | ENSMUSG00000058672 | protein_coding | Tubb2a | tubulin, beta 2A class IIA | 22151 | 13 | 34074274 | 34078007 | -1 | Mm.422827 |
| 6.32 | 7.06 | 5.97 | 7.28 | DN1ab_ltr | DN1cd | 0.94 | 7.43 | 0.00014 | 0.013 | 0.012 | ENSMUSG00000020346 | protein_coding | Mgat1 | mannoside acetylglucosaminyltransferase 1 | 17308 | 11 | 49244191 | 49263030 | 1 | Mm.196933 |
| 3.42 | 4.46 | 2.61 | 4.71 | DN1ab_ltr | DN1cd | 1.41 | 6.05 | 5.00E-04 | 0.023 | 0.012 | ENSMUSG00000011008 | protein_coding | Mcoln2 | mucolipin 2 | 68279 | 3 | 146149833 | 146195513 | 1 | Mm.116862 |
| 8.14 | 7.44 | 9.36 | 8.08 | DN1ab_ltr | DN1cd | -0.93 | -7.07 | 0.00019 | 0.015 | 0.012 | ENSMUSG00000032231 | protein_coding | Anxa2 | annexin A2 | 12306 | 9 | 69453620 | 69491795 | 1 | Mm.238343 |
| 5.7 | 7.55 | 5.48 | 8.11 | DN1ab_ltr | DN1cd | 2.12 | 12.43 | 4.70E-06 | 0.0033 | 0.012 | ENSMUSG00000033295 | protein_coding | Ptprf | protein tyrosine phosphatase, receptor type, F | 19268 | 4 | 118208213 | 118291405 | -1 | Mm.29855 |
| 4.37 | 4.9 | 4.24 | 5.29 | DN1ab_ltr | DN1cd | 0.72 | 6.34 | 0.00038 | 0.02 | 0.013 | ENSMUSG00000030852 | protein_coding | Tacc2 | transforming, acidic coiled-coil containing protein 2 | 57752 | 7 | 130577438 | 130764785 | 1 | Mm.86322 |
| 6.78 | 6.52 | 7.35 | 6.59 | DN1ab_ltr | DN1cd | -0.45 | -3.93 | 0.0056 | 0.08 | 0.013 | ENSMUSG00000063410 | protein_coding | Stk24 | serine/threonine kinase 24 | 223255 | 14 | 121286343 | 121380011 | -1 | Mm.390756 Mm.442081 Mm.472866 |
| 3.69 | 4.51 | 1.74 | 4.34 | DN1ab_ltr | DN1cd | 1.42 | 3.65 | 0.008 | 0.098 | 0.013 | ENSMUSG00000034664 | protein_coding | Itga2b | integrin alpha 2b | 16399 | 11 | 102453297 | 102470122 | -1 | Mm.26646 |
| 4.11 | 3.83 | 5.08 | 3.99 | DN1ab_ltr | DN1cd | -0.6 | -3.27 | 0.013 | 0.13 | 0.013 | ENSMUSG00000041439 | protein_coding | Mfsd6 | major facilitator superfamily domain containing 6 | 98682 | 1 | 52656286 | 52727462 | -1 | Mm.491173 |
| 0.84 | 3.62 | -0.11 | 4.32 | DN1ab_ltr | DN1cd | 3.29 | 9.2 | 3.50E-05 | 0.0072 | 0.013 | ENSMUSG00000038264 | protein_coding | Sema7a | sema domain, immunoglobulin domain (Ig), and GPI membrane anchor, (semaphorin) 7A | 20361 | 9 | 57940112 | 57962865 | 1 | Mm.335187 |
| 2.55 | 5.22 | 1.48 | 6.37 | DN1ab_ltr | DN1cd | 3.38 | 7.04 | 2.00E-04 | 0.015 | 0.013 | ENSMUSG00000002033 | protein_coding | Cd3g | CD3 antigen, gamma polypeptide | 12502 | 9 | 44969572 | 44980431 | -1 | Mm.335106 |
| 7.45 | 7.63 | 6.62 | 7.2 | DN1ab_ltr | DN1cd | 0.32 | 3.56 | 0.0091 | 0.1 | 0.013 | ENSMUSG00000045273 | protein_coding | Cenph | centromere protein H | 26886 | 13 | 100759674 | 100775899 | -1 | Mm.273502 |
| 2.91 | 2.72 | 3.84 | 2.92 | DN1ab_ltr | DN1cd | -0.48 | -2.85 | 0.024 | 0.18 | 0.013 | ENSMUSG00000038936 | protein_coding | Sccpdh | saccharopine dehydrogenase (putative) | 109232 | 1 | 179668210 | 179687189 | 1 | Mm.136458 Mm.491204 |
| 4.59 | 5.18 | 3.24 | 5.29 | DN1ab_ltr | DN1cd | 1.11 | 3.41 | 0.011 | 0.12 | 0.014 | ENSMUSG00000008305 | protein_coding | Tle1 | transducin-like enhancer of split 1 | 21885 | 4 | 72117142 | 72200919 | -1 | Mm.278444 Mm.388819 Mm.399716 |
| 4.82 | 5.42 | 3.84 | 5.72 | DN1ab_ltr | DN1cd | 1.06 | 3.69 | 0.0077 | 0.095 | 0.014 | ENSMUSG00000037999 | protein_coding | Arap2 | ArfGAP with RhoGAP domain, ankyrin repeat and PH domain 2 | 212285 | 5 | 62602445 | 62766159 | -1 | Mm.244403 |
| 0.79 | 2.06 | -2.8 | 1.88 | DN1ab_ltr | DN1cd | 2.36 | 3.19 | 0.015 | 0.14 | 0.014 | ENSMUSG00000026815 | protein_coding | Gfi1b | growth factor independent 1B | 14582 | 2 | 28609450 | 28621982 | -1 | Mm.373385 |
| -3.82 | -1.54 | -3.94 | 0.6 | DN1ab_ltr | DN1cd | 3.18 | 6.17 | 0.00045 | 0.022 | 0.014 | ENSMUSG00000105501 | lincRNA | 5330426L24Rik | RIKEN cDNA 5330426L24 gene | NA | 14 | 54349462 | 54351636 | -1 | |
| 2.1 | 1.06 | 3.2 | 0.62 | DN1ab_ltr | DN1cd | -1.67 | -4.73 | 0.0021 | 0.046 | 0.014 | ENSMUSG00000033949 | protein_coding | Trim36 | tripartite motif-containing 36 | 28105 | 18 | 46165302 | 46212607 | -1 | Mm.277382 |
| 6.22 | 6.38 | 5.37 | 6.11 | DN1ab_ltr | DN1cd | 0.37 | 2.78 | 0.027 | 0.19 | 0.014 | ENSMUSG00000089726 | lincRNA | Mir17hg | Mir17 host gene (non-protein coding) | NA | 14 | 115042879 | 115046727 | 1 | |
| -2.91 | -3.86 | 0.2 | -4.02 | DN1ab_ltr | DN1cd | -2.12 | -2.89 | 0.023 | 0.17 | 0.015 | ENSMUSG00000105289 | TEC | Gm43483 | predicted gene 43483 | NA | 5 | 131639948 | 131642807 | -1 | |
| 2.48 | 3.24 | 0.77 | 3.06 | DN1ab_ltr | DN1cd | 1.28 | 3.77 | 0.0069 | 0.09 | 0.015 | ENSMUSG00000032715 | protein_coding | Trib3 | tribbles pseudokinase 3 | 228775 | 2 | 152337422 | 152344032 | -1 | Mm.276018 |
| 8.95 | 9.54 | 8.7 | 9.79 | DN1ab_ltr | DN1cd | 0.77 | 6.9 | 0.00022 | 0.016 | 0.015 | ENSMUSG00000017707 | protein_coding | Serinc3 | serine incorporator 3 | 26943 | 2 | 163623272 | 163645131 | -1 | Mm.218473 Mm.281691 |
| -2.24 | 4.52 | -3.94 | 4.9 | DN1ab_ltr | DN1cd | 7.34 | 16.7 | 6.30E-07 | 0.001 | 0.015 | ENSMUSG00000069049 | protein_coding | Eif2s3y | eukaryotic translation initiation factor 2, subunit 3, structural gene Y-linked | 26908 | Y | 1010543 | 1028847 | 1 | Mm.250909 |
| -1.29 | -3.86 | -0.43 | -0.71 | DN1ab_ltr | DN1cd | -1.69 | -3.24 | 0.014 | 0.13 | 0.015 | ENSMUSG00000112437 | lincRNA | Gm35188 | predicted gene, 35188 | NA | 12 | 52496326 | 52581409 | -1 | |
| 2.36 | 1.89 | 3.54 | 1.12 | DN1ab_ltr | DN1cd | -1.21 | -2.72 | 0.029 | 0.2 | 0.015 | ENSMUSG00000069310 | protein_coding | Hist1h3c | histone cluster 1, H3c | 319148 | 13 | 23744973 | 23745602 | -1 | Mm.262517 |
| 5.32 | 6.3 | 6.56 | 6.66 | DN1ab_ltr | DN1cd | 0.65 | 3.31 | 0.013 | 0.13 | 0.015 | ENSMUSG00000032578 | protein_coding | Cish | cytokine inducible SH2-containing protein | 12700 | 9 | 107296026 | 107302784 | 1 | Mm.4592 |
| 6.78 | 7.2 | 6.37 | 7.48 | DN1ab_ltr | DN1cd | 0.67 | 4.3 | 0.0035 | 0.06 | 0.015 | ENSMUSG00000027435 | protein_coding | Cd93 | CD93 antigen | 17064 | 2 | 148436640 | 148443563 | -1 | Mm.681 |
| 10.84 | 3.34 | 10.8 | 2.55 | DN3ab | DN3cd | -7.78 | -38.98 | 3.10E-11 | 7.20E-07 | 0.056 | ENSMUSG00000086503 | lincRNA | Xist | inactive X specific transcripts | NA | X | 103460366 | 103483254 | -1 | |
| 7.22 | 1.5 | 6.9 | 1.39 | DN3ab | DN3cd | -5.62 | -28.37 | 5.20E-10 | 5.90E-06 | 0.61 | ENSMUSG00000082399 | processed_pseudogene | Gm14036 | predicted gene 14036 | NA | 2 | 134803949 | 134804428 | 1 | |
| 6.04 | 8.03 | 5.92 | 7.64 | DN3ab | DN3cd | 1.87 | 24.36 | 2.00E-09 | 1.50E-05 | 0.047 | ENSMUSG00000048758 | protein_coding | Rpl29 | ribosomal protein L29 | 19944 | 9 | 106429454 | 106431568 | 1 | Mm.288696 Mm.359377 Mm.431965 |
| 5.38 | 3.13 | 6.32 | 4.07 | DN3ab | DN3cd | -2.25 | -23.13 | 3.10E-09 | 1.80E-05 | 0.99 | ENSMUSG00000030352 | protein_coding | Tspan9 | tetraspanin 9 | 109246 | 6 | 127961396 | 128143594 | -1 | Mm.21814 Mm.29608 |
| 2.69 | 5.88 | 1.95 | 5.62 | DN3ab | DN3cd | 3.35 | 21.42 | 6.00E-09 | 2.80E-05 | 0.13 | ENSMUSG00000074516 | processed_pseudogene | Gm10709 | predicted gene 10709 | NA | 9 | 7751725 | 7752204 | 1 | |
| 5.97 | 7.18 | 5.6 | 6.99 | DN3ab | DN3cd | 1.28 | 18.35 | 2.30E-08 | 7.30E-05 | 0.14 | ENSMUSG00000029722 | protein_coding | Agfg2 | ArfGAP with FG repeats 2 | 231801 | 5 | 137650463 | 137684726 | -1 | Mm.260869 |
| 4.7 | 6.5 | 4.06 | 5.7 | DN3ab | DN3cd | 1.74 | 18.15 | 2.50E-08 | 7.30E-05 | 0.4 | ENSMUSG00000078154 | protein_coding | Gm12184 | predicted gene 12184 | NA | 11 | 48825346 | 48826655 | -1 | |
| 7.83 | 6.01 | 9.21 | 7.53 | DN3ab | DN3cd | -1.74 | -16.44 | 6.00E-08 | 0.00012 | 0.55 | ENSMUSG00000000078 | protein_coding | Klf6 | Kruppel-like factor 6 | 23849 | 13 | 5861482 | 5870394 | 1 | Mm.275036 |
| 9.33 | 10.47 | 8.61 | 9.81 | DN3ab | DN3cd | 1.16 | 16.2 | 6.80E-08 | 0.00012 | 0.67 | ENSMUSG00000094248 | protein_coding | Hist1h2ao | histone cluster 1, H2ao | 665433 | 13 | 21810465 | 21810944 | 1 | Mm.250564 Mm.422826 |

| | | | | | | | | | | | | | | | | | | | | |
|---|---|---|---|---|---|---|---|---|---|---|---|---|---|---|---|---|---|---|---|---|
| 6.74 | 5.09 | 8.04 | 6.62 | DN3ab | DN3cd | -1.51 | -16.2 | 6.80E-08 | 0.00012 | 0.2 | ENSMUSG00000054892 | protein_coding | Txk | TXK tyrosine kinase | 22165 | 5 | 72695978 | 72752777 | -1 | Mm.3264 |
| 4.04 | 1.26 | 6.33 | 3.59 | DN3ab | DN3cd | -2.75 | -16.11 | 7.10E-08 | 0.00012 | 0.92 | ENSMUSG00000008136 | protein_coding | Fhl2 | four and a half LIM domains 2 | 14200 | 1 | 43123074 | 43196984 | -1 | Mm.6799 |
| 5.75 | 0.83 | 5.83 | 1.35 | DN3ab | DN3cd | -4.67 | -16.02 | 7.50E-08 | 0.00012 | 0.46 | ENSMUSG00000030703 | protein_coding | Gdpd3 | glycerophosphodiester phosphodiesterase domain containing 3 | 68616 | 7 | 126766334 | 126775649 | 1 | Mm.246881 |
| 5.61 | 4.19 | 6.95 | 5.74 | DN3ab | DN3cd | -1.29 | -15.95 | 7.80E-08 | 0.00012 | 0.19 | ENSMUSG00000072620 | protein_coding | Slfn2 | schlafen 2 | 20556 | 11 | 83065112 | 83070678 | 1 | Mm.278689 |
| 1.41 | 4.44 | 2.84 | 6.25 | DN3ab | DN3cd | 3.27 | 15.94 | 7.80E-08 | 0.00012 | 0.38 | ENSMUSG00000002204 | protein_coding | Napsa | napsin A aspartic peptidase | 16541 | 7 | 44572380 | 44586862 | 1 | Mm.383181 |
| 5.74 | 6.97 | 5.31 | 6.64 | DN3ab | DN3cd | 1.27 | 15.72 | 8.80E-08 | 0.00013 | 0.53 | ENSMUSG00000097906 | processed_pseudogene | Gm9625 | predicted gene 9625 | NA | 13 | 67810246 | 67811200 | 1 | |
| 6.91 | 5.57 | 7.78 | 6.48 | DN3ab | DN3cd | -1.32 | -15.32 | 1.10E-07 | 0.00015 | 0.84 | ENSMUSG00000040430 | protein_coding | Pitpnc1 | phosphatidylinositol transfer protein, cytoplasmic 1 | 71795 | 11 | 107207892 | 107470699 | -1 | Mm.397413 Mm.476186 |
| -2.43 | 3.59 | -3.72 | 3.98 | DN3ab | DN3cd | 6.84 | 14.44 | 1.80E-07 | 2.00E-04 | 0.054 | ENSMUSG00000056673 | protein_coding | Kdm5d | lysine (K)-specific demethylase 5D | 20592 | Y | 897788 | 956786 | 1 | Mm.262676 |
| -0.47 | 6.2 | -0.69 | 6.57 | DN3ab | DN3cd | 6.92 | 14.37 | 1.90E-07 | 2.00E-04 | 0.56 | ENSMUSG00000069045 | protein_coding | Ddx3y | DEAD (Asp-Glu-Ala-Asp) box polypeptide 3, Y-linked | 26900 | Y | 1260771 | 1286629 | -1 | Mm.302938 |
| 6.15 | 7.28 | 5.65 | 6.88 | DN3ab | DN3cd | 1.18 | 14.34 | 1.90E-07 | 2.00E-04 | 0.55 | ENSMUSG00000032254 | protein_coding | Kif23 | kinesin family member 23 | 71819 | 9 | 61915905 | 61946774 | -1 | Mm.259374 |
| 8.65 | 7.47 | 8.8 | 7.59 | DN3ab | DN3cd | -1.2 | -14.03 | 2.30E-07 | 0.00023 | 0.87 | ENSMUSG00000026159 | protein_coding | Agfg1 | ArfGAP with FG repeats 1 | 15463 | 1 | 82839483 | 82901182 | 1 | Mm.392569 Mm.433409 |
| 4.2 | 5.82 | 3.81 | 5.75 | DN3ab | DN3cd | 1.75 | 13.94 | 2.50E-07 | 0.00023 | 0.2 | ENSMUSG00000029406 | protein_coding | Pitpnm2 | phosphatidylinositol transfer protein, membrane-associated 2 | 19679 | 5 | 124118690 | 124249760 | -1 | Mm.44261 |
| 4.47 | 2.45 | 4.95 | 3.06 | DN3ab | DN3cd | -1.95 | -13.76 | 2.80E-07 | 0.00024 | 0.67 | ENSMUSG00000067219 | protein_coding | Nipal1 | NIPA-like domain containing 1 | 70701 | 5 | 72647795 | 72671078 | 1 | Mm.38884 |
| 7.74 | 5.83 | 10.36 | 8.14 | DN3ab | DN3cd | -2.11 | -13.68 | 2.90E-07 | 0.00024 | 0.38 | ENSMUSG00000017493 | protein_coding | Igfbp4 | insulin-like growth factor binding protein 4 | 16010 | 11 | 99041244 | 99054392 | 1 | Mm.233799 |
| 0.79 | -1.35 | 5.35 | 2.65 | DN3ab | DN3cd | -2.64 | -13.58 | 3.10E-07 | 0.00024 | 0.37 | ENSMUSG00000021069 | protein_coding | Pygl | liver glycogen phosphorylase | 110095 | 12 | 70190811 | 70231488 | -1 | Mm.256926 Mm.447796 |
| 6.9 | 5.74 | 7.93 | 6.65 | DN3ab | DN3cd | -1.23 | -13.49 | 3.30E-07 | 0.00024 | 0.52 | ENSMUSG00000036006 | protein_coding | Ripor2 | RHO family interacting cell polarization regulator 2 | 193385 | 13 | 24582189 | 24733816 | 1 | Mm.217319 |
| 7.08 | 5.97 | 8.26 | 6.79 | DN3ab | DN3cd | -1.31 | -13.47 | 3.30E-07 | 0.00024 | 0.041 | ENSMUSG00000028064 | protein_coding | Sema4a | sema domain, immunoglobulin domain (Ig), transmembrane domain (TM) and short cytoplasmic domain, (semaphorin) 4A | 20351 | 3 | 88435959 | 88461182 | -1 | Mm.439752 |
| 9.29 | 10.65 | 9.59 | 10.59 | DN3ab | DN3cd | 1.18 | 13.22 | 3.90E-07 | 0.00028 | 0.032 | ENSMUSG00000034165 | protein_coding | Ccnd3 | cyclin D3 | 12445 | 17 | 47505051 | 47599691 | 1 | Mm.27291 |
| 4.15 | 2.44 | 4.06 | 2.31 | DN3ab | DN3cd | -1.73 | -13.15 | 4.00E-07 | 0.00028 | 0.88 | ENSMUSG00000083833 | processed_pseudogene | Gm13841 | predicted gene 13841 | NA | 5 | 115801677 | 115802159 | -1 | |
| 2.77 | -0.35 | 4.21 | 1.12 | DN3ab | DN3cd | -3.1 | -13.03 | 4.40E-07 | 0.00029 | 0.95 | ENSMUSG00000026435 | protein_coding | Slc45a3 | solute carrier family 45, member 3 | 212980 | 1 | 131962967 | 131982969 | 1 | Mm.200307 |
| -2.06 | 3.02 | -2.95 | 3.03 | DN3ab | DN3cd | 5.51 | 12.97 | 4.50E-07 | 3.00E-04 | 0.28 | ENSMUSG00000099876 | lincRNA | Gm29650 | predicted gene 29650 | NA | Y | 1048393 | 1049134 | 1 | |
| -4.01 | 0.47 | -3.72 | 2.2 | DN3ab | DN3cd | 5.24 | 12.87 | 4.90E-07 | 3.00E-04 | 0.05 | ENSMUSG00000100277 | processed_transcript | 1810053B23Rik | RIKEN cDNA 1810053B23 gene | NA | 16 | 93342504 | 93366668 | -1 | |
| 5.01 | 3.52 | 7.9 | 5.99 | DN3ab | DN3cd | -1.8 | -12.86 | 4.90E-07 | 3.00E-04 | 0.18 | ENSMUSG00000004612 | protein_coding | Nkg7 | natural killer cell group 7 sequence | 72310 | 7 | 43437073 | 43438249 | 1 | Mm.34613 |
| -1.37 | 4.41 | -2.13 | 5.14 | DN3ab | DN3cd | 6.47 | 12.77 | 5.20E-07 | 3.00E-04 | 0.13 | ENSMUSG00000068457 | protein_coding | Uty | ubiquitously transcribed tetratricopeptide repeat gene, Y chromosome | 22290 | Y | 1096861 | 1245759 | -1 | Mm.20477 |
| 5.09 | 3.15 | 6.32 | 4.55 | DN3ab | DN3cd | -1.84 | -12.75 | 5.20E-07 | 3.00E-04 | 0.59 | ENSMUSG00000020732 | protein_coding | Rab37 | RAB37, member RAS oncogene family | 58222 | 11 | 115091431 | 115162236 | 1 | Mm.143789 |
| 6.26 | 7.27 | 5.62 | 6.82 | DN3ab | DN3cd | 1.08 | 12.71 | 5.40E-07 | 3.00E-04 | 0.27 | ENSMUSG00000069272 | protein_coding | Hist1h2ae | histone cluster 1, H2ae | 319166 | 13 | 23570517 | 23571220 | -1 | Mm.261665 |
| 3.77 | 2.28 | 5.61 | 3.55 | DN3ab | DN3cd | -1.87 | -12.66 | 5.60E-07 | 3.00E-04 | 0.043 | ENSMUSG00000027907 | protein_coding | S100a11 | S100 calcium binding protein A11 | 20195 | 3 | 93520488 | 93526287 | 1 | Mm.280038 |
| 7.48 | 8.53 | 6.75 | 8.09 | DN3ab | DN3cd | 1.17 | 12.66 | 5.60E-07 | 3.00E-04 | 0.12 | ENSMUSG00000061615 | protein_coding | Hist1h2ab | histone cluster 1, H2ab | 319172 | 13 | 23751126 | 23751598 | 1 | Mm.423402 |
| 6.65 | 7.79 | 5.67 | 7.19 | DN3ab | DN3cd | 1.28 | 12.62 | 5.70E-07 | 3.00E-04 | 0.047 | ENSMUSG00000018983 | protein_coding | E2f2 | E2F transcription factor 2 | 242705 | 4 | 136172394 | 136196057 | 1 | Mm.307932 |
| -3.22 | 1.75 | -3.72 | 2.06 | DN3ab | DN3cd | 5.37 | 12.57 | 5.90E-07 | 3.00E-04 | 0.34 | ENSMUSG00000071234 | protein_coding | Syndig1l | synapse differentiation inducing 1 like | 627191 | 12 | 84677277 | 84698831 | -1 | Mm.152484 |
| 6.46 | 5.63 | 7.37 | 6.24 | DN3ab | DN3cd | -0.99 | -12.45 | 6.40E-07 | 0.00031 | 0.042 | ENSMUSG00000019843 | protein_coding | Fyn | Fyn proto-oncogene | 14360 | 10 | 39368855 | 39565381 | 1 | Mm.4848 |
| -4.01 | 0.32 | -3.72 | 0.58 | DN3ab | DN3cd | 4.32 | 12.35 | 6.90E-07 | 0.00033 | 0.96 | ENSMUSG00000047517 | protein_coding | Dmbt1 | deleted in malignant brain tumors 1 | 12945 | 7 | 131032053 | 131121630 | 1 | Mm.4138 |
| 5.09 | 6.53 | 4.62 | 6.31 | DN3ab | DN3cd | 1.54 | 12.23 | 7.40E-07 | 0.00033 | 0.32 | ENSMUSG00000022895 | protein_coding | Ets2 | E26 avian leukemia oncogene 2, 3' domain | 23872 | 16 | 95702075 | 95721051 | 1 | Mm.290207 |
| 4.83 | 2.11 | 7.61 | 4.33 | DN3ab | DN3cd | -3.14 | -12.13 | 8.00E-07 | 0.00034 | 0.37 | ENSMUSG00000032238 | protein_coding | Rora | RAR-related orphan receptor alpha | 19883 | 9 | 68653786 | 69388246 | 1 | Mm.378450 Mm.391890 Mm.427266 Mm.445802 Mm.491288 |
| 6.02 | 5.06 | 6.88 | 5.86 | DN3ab | DN3cd | -0.99 | -12.1 | 8.10E-07 | 0.00034 | 0.68 | ENSMUSG00000028159 | protein_coding | Dapp1 | dual adaptor for phosphotyrosine and 3-phosphoinositides 1 | 26377 | 3 | 137931007 | 137981545 | -1 | Mm.254835 Mm.470427 |
| 3.24 | 0.42 | 6.43 | 3.12 | DN3ab | DN3cd | -3.22 | -12.04 | 8.50E-07 | 0.00035 | 0.5 | ENSMUSG00000024511 | protein_coding | Rab27b | RAB27B, member RAS oncogene family | 80718 | 18 | 69979131 | 70141605 | -1 | Mm.246753 |
| 3.01 | 4.91 | 2.76 | 4.52 | DN3ab | DN3cd | 1.84 | 11.92 | 9.30E-07 | 0.00036 | 0.7 | ENSMUSG00000020205 | protein_coding | Phlda1 | pleckstrin homology like domain, family A, member 1 | 21664 | 10 | 111506286 | 111508645 | 1 | Mm.3117 |
| 6.95 | 5.65 | 7.56 | 6.45 | DN3ab | DN3cd | -1.2 | -11.89 | 9.40E-07 | 0.00036 | 0.38 | ENSMUSG00000021892 | protein_coding | Sh3bp5 | SH3-domain binding protein 5 (BTK-associated) | 24056 | 14 | 31359880 | 31436078 | -1 | Mm.383198 Mm.438866 |
| 6.85 | 5.93 | 7.41 | 6.17 | DN3ab | DN3cd | -1.08 | -11.89 | 9.50E-07 | 0.00036 | 0.063 | ENSMUSG00000036943 | protein_coding | Rab8b | RAB8B, member RAS oncogene family | 235442 | 9 | 66843664 | 66919687 | -1 | Mm.260376 |
| 2.19 | 4.6 | 2.24 | 5.44 | DN3ab | DN3cd | 2.78 | 11.85 | 9.70E-07 | 0.00036 | 0.078 | ENSMUSG00000034361 | protein_coding | Cpne2 | copine II | 234577 | 8 | 94532990 | 94570531 | 1 | Mm.291815 |
| 6.99 | 6.04 | 7.86 | 7 | DN3ab | DN3cd | -0.91 | -11.84 | 9.80E-07 | 0.00036 | 0.54 | ENSMUSG00000026223 | protein_coding | Itm2c | integral membrane protein 2C | 64294 | 1 | 85894281 | 85908675 | 1 | Mm.29870 |
| 8.89 | 7.95 | 10.43 | 9.32 | DN3ab | DN3cd | -1.04 | -11.79 | 1.00E-06 | 0.00037 | 0.35 | ENSMUSG00000079523 | protein_coding | Tmsb10 | thymosin, beta 10 | 19240 | 6 | 72957347 | 72958748 | -1 | Mm.3532 Mm.391824 Mm.436717 |
| 5.16 | 3.2 | 6.05 | 3.96 | DN3ab | DN3cd | -2.03 | -11.76 | 1.00E-06 | 0.00037 | 0.71 | ENSMUSG00000019970 | protein_coding | Sgk1 | serum/glucocorticoid regulated kinase 1 | 20393 | 10 | 21882184 | 21999903 | 1 | Mm.28405 |
| 4.82 | 3.44 | 6.06 | 4.79 | DN3ab | DN3cd | -1.31 | -11.74 | 1.10E-06 | 0.00037 | 0.66 | ENSMUSG00000040811 | protein_coding | Eml2 | echinoderm microtubule associated protein like 2 | 72205 | 7 | 19176421 | 19206482 | 1 | Mm.154121 |
| 2.63 | -3.92 | 2.74 | -2.8 | DN3ab | DN3cd | -6.1 | -11.73 | 1.10E-06 | 0.00037 | 0.34 | ENSMUSG00000084403 | processed_pseudogene | Rps15a-ps8 | ribosomal protein S15A, pseudogene 8 | NA | 4 | 53791070 | 53791456 | -1 | |
| 6.48 | 7.64 | 5.93 | 7.09 | DN3ab | DN3cd | 1.16 | 11.62 | 1.10E-06 | 0.00038 | 0.98 | ENSMUSG00000080999 | unprocessed_pseudogene | Gm11336 | predicted gene 11336 | NA | 13 | 23746166 | 23746499 | -1 | |
| 2.67 | 0.47 | 6.31 | 3.36 | DN3ab | DN3cd | -2.83 | -11.6 | 1.20E-06 | 0.00038 | 0.25 | ENSMUSG00000026483 | protein_coding | Fam129a | family with sequence similarity 129, member A | 63913 | 1 | 151571186 | 151721939 | 1 | Mm.157700 Mm.482559 |
| 9.41 | 10.46 | 8.79 | 9.77 | DN3ab | DN3cd | 1.03 | 11.48 | 1.30E-06 | 0.00039 | 0.72 | ENSMUSG00000094777 | protein_coding | Hist1h2ap | histone cluster 1, H2ap | 319171 | 13 | 21833026 | 21833507 | -1 | Mm.250564 Mm.422826 |
| 6.94 | 8.09 | 6.44 | 7.49 | DN3ab | DN3cd | 1.11 | 11.48 | 1.30E-06 | 0.00039 | 0.63 | ENSMUSG00000069270 | protein_coding | Hist1h2ac | histone cluster 1, H2ac | 319164 | 13 | 23681467 | 23683948 | -1 | Mm.261973 |
| -1.49 | 3.18 | -0.59 | 3.94 | DN3ab | DN3cd | 4.59 | 11.48 | 1.30E-06 | 0.00039 | 0.87 | ENSMUSG00000042757 | protein_coding | Tmem108 | transmembrane protein 108 | 81907 | 9 | 103482947 | 103761837 | -1 | Mm.384704 |
| 8.37 | 8.87 | 8.07 | 8.59 | DN3ab | DN3cd | 0.51 | 11.43 | 1.30E-06 | 0.00039 | 0.87 | ENSMUSG00000052397 | protein_coding | Ezr | ezrin | 22350 | 17 | 6738041 | 6782784 | -1 | Mm.277812 |
| 3.23 | 4.61 | 2.81 | 4.24 | DN3ab | DN3cd | 1.4 | 11.43 | 1.30E-06 | 0.00039 | 0.81 | ENSMUSG00000047409 | protein_coding | Ctdspl | CTD (carboxy-terminal domain, RNA polymerase II, polypeptide A) small phosphatase-like | 69274 | 9 | 118926453 | 119043998 | 1 | Mm.439928 |
| 2.15 | -1.6 | 1.86 | -2.43 | DN3ab | DN3cd | -3.99 | -11.43 | 1.30E-06 | 0.00039 | 0.43 | ENSMUSG00000091785 | processed_pseudogene | Rpl29-ps5 | ribosomal protein L29, pseudogene 5 | NA | 18 | 30459082 | 30459544 | -1 | |
| 7.37 | 5.96 | 7.46 | 6.33 | DN3ab | DN3cd | -1.27 | -11.41 | 1.30E-06 | 0.00039 | 0.21 | ENSMUSG00000045414 | protein_coding | 1190002N15Rik | RIKEN cDNA 1190002N15 gene | 68861 | 9 | 94517864 | 94538081 | -1 | Mm.258746 |
| 7.96 | 8.78 | 7.48 | 8.48 | DN3ab | DN3cd | 0.89 | 11.3 | 1.40E-06 | 0.00042 | 0.25 | ENSMUSG00000026558 | protein_coding | Uck2 | uridine-cytidine kinase 2 | 80914 | 1 | 167222883 | 167285320 | -1 | Mm.280895 Mm.440156 |
| 4.03 | 2.76 | 6.32 | 4.6 | DN3ab | DN3cd | -1.58 | -11.2 | 1.60E-06 | 0.00044 | 0.14 | ENSMUSG00000026509 | protein_coding | Capn2 | calpain 2 | 12334 | 1 | 182467260 | 182517608 | -1 | Mm.19306 Mm.474138 |
| 7.49 | 8.5 | 6.74 | 8.13 | DN3ab | DN3cd | 1.17 | 11.17 | 1.60E-06 | 0.00044 | 0.058 | ENSMUSG00000071516 | protein_coding | Hist1h2ai | histone cluster 1, H2ai | 319191 | 13 | 21716422 | 21716814 | 1 | Mm.14767 |
| 4.89 | 3.94 | 5.86 | 4.78 | DN3ab | DN3cd | -1.02 | -11.15 | 1.60E-06 | 0.00044 | 0.5 | ENSMUSG00000024642 | protein_coding | Tle4 | transducin-like enhancer of split 4 | 21888 | 19 | 14448072 | 14598051 | -1 | Mm.103638 |
| 7.85 | 6.9 | 8.82 | 7.54 | DN3ab | DN3cd | -1.13 | -11.15 | 1.60E-06 | 0.00044 | 0.1 | ENSMUSG00000046722 | protein_coding | Cdc42se1 | CDC42 small effector 1 | 57912 | 3 | 95228732 | 95236409 | 1 | Mm.28189 Mm.331208 |
| 6.61 | 4.18 | 9.18 | 7.21 | DN3ab | DN3cd | -2.09 | -11.14 | 1.60E-06 | 0.00044 | 0.29 | ENSMUSG00000001281 | protein_coding | Itgb7 | integrin beta 7 | 16421 | 15 | 102215995 | 102231944 | -1 | Mm.58 |
| 7.6 | 6.61 | 8.17 | 7.16 | DN3ab | DN3cd | -1 | -11.1 | 1.70E-06 | 0.00045 | 0.88 | ENSMUSG00000026655 | protein_coding | Fam107b | family with sequence similarity 107, member B | 66540 | 2 | 3570488 | 3782142 | 1 | Mm.277864 |
| 4.74 | 1.09 | 6.92 | 3.3 | DN3ab | DN3cd | -3.62 | -11.07 | 1.70E-06 | 0.00045 | 0.96 | ENSMUSG00000021250 | protein_coding | Fos | FBJ osteosarcoma oncogene | 14281 | 12 | 85473890 | 85477273 | 1 | Mm.246513 |
| 9.08 | 10.01 | 8.06 | 9.38 | DN3ab | DN3cd | 1.08 | 10.98 | 1.80E-06 | 0.00048 | 0.038 | ENSMUSG00000004099 | protein_coding | Dnmt1 | DNA methyltransferase (cytosine-5) 1 | 13433 | 9 | 20907209 | 20959888 | -1 | Mm.128580 Mm.485562 |
| 6.73 | 7.71 | 6.23 | 7.42 | DN3ab | DN3cd | 1.07 | 10.95 | 1.90E-06 | 0.00048 | 0.3 | ENSMUSG00000076617 | IG_C_gene | Ighm | immunoglobulin heavy constant mu | NA | 12 | 113418558 | 113422730 | -1 | |
| 6.84 | 5.97 | 7.41 | 6.45 | DN3ab | DN3cd | -0.91 | -10.91 | 1.90E-06 | 0.00048 | 0.57 | ENSMUSG00000060216 | protein_coding | Arrb2 | arrestin, beta 2 | 216869 | 11 | 70432635 | 70440828 | 1 | Mm.203747 |
| 10.93 | 10.29 | 11.02 | 10.13 | DN3ab | DN3cd | -0.75 | -10.89 | 2.00E-06 | 0.00049 | 0.072 | ENSMUSG00000094344 | processed_pseudogene | Gm11942 | predicted gene 11942 | NA | 4 | 32964888 | 32965232 | 1 | |
| 4.66 | 3.47 | 5.75 | 4.34 | DN3ab | DN3cd | -1.32 | -10.88 | 2.00E-06 | 0.00049 | 0.39 | ENSMUSG00000024696 | protein_coding | Lpxn | leupaxin | 107321 | 19 | 12798606 | 12833807 | 1 | Mm.313136 |
| 7.87 | 8.65 | 7.11 | 7.84 | DN3ab | DN3cd | 0.76 | 10.84 | 2.00E-06 | 5.00E-04 | 0.72 | ENSMUSG00000020649 | protein_coding | Rrm2 | ribonucleotide reductase M2 | 20135 | 12 | 24708241 | 24714146 | 1 | Mm.99 Mm.471826 |
| 10.93 | 9.38 | 12.16 | 10.99 | DN3ab | DN3cd | -1.32 | -10.81 | 2.10E-06 | 5.00E-04 | 0.12 | ENSMUSG00000023132 | protein_coding | Gzma | granzyme A | 14938 | 13 | 113093825 | 113100981 | -1 | Mm.15510 |
| 5.69 | 4.3 | 7.39 | 5.91 | DN3ab | DN3cd | -1.45 | -10.79 | 2.10E-06 | 0.00051 | 0.75 | ENSMUSG00000029552 | protein_coding | Tes | testis derived transcript | 21753 | 6 | 17065149 | 17105828 | 1 | Mm.436548 |
| 8.36 | 9.16 | 7.46 | 8.27 | DN3ab | DN3cd | 0.8 | 10.77 | 2.20E-06 | 0.00051 | 0.97 | ENSMUSG00000001228 | protein_coding | Uhrf1 | ubiquitin-like, containing PHD and RING finger domains, 1 | 18140 | 17 | 56303321 | 56323486 | 1 | Mm.42196 |
| 7.03 | 5.34 | 8.7 | 6.75 | DN3ab | DN3cd | -1.84 | -10.75 | 2.20E-06 | 0.00051 | 0.47 | ENSMUSG00000023048 | protein_coding | Prr13 | proline rich 13 | 66151 | 15 | 102459028 | 102462806 | 1 | Mm.393955 |
| 5.55 | 6.41 | 4.74 | 5.87 | DN3ab | DN3cd | 0.97 | 10.67 | 2.30E-06 | 0.00053 | 0.13 | ENSMUSG00000030346 | protein_coding | Rad51ap1 | RAD51 associated protein 1 | 19362 | 6 | 126923050 | 126939587 | -1 | Mm.204634 |
| 3.46 | 2.28 | 4.9 | 3.3 | DN3ab | DN3cd | -1.44 | -10.67 | 2.30E-06 | 0.00053 | 0.1 | ENSMUSG00000061086 | protein_coding | Myl4 | myosin, light polypeptide 4 | 17896 | 11 | 104550663 | 104595753 | 1 | Mm.390355 |
| 5.56 | 6.58 | 4.82 | 6.08 | DN3ab | DN3cd | 1.11 | 10.64 | 2.40E-06 | 0.00053 | 0.27 | ENSMUSG00000080076 | unprocessed_pseudogene | Hist1h2aj | histone cluster 1, H2aj | NA | 13 | 21721457 | 21721808 | -1 | |
| 8.3 | 8.93 | 7.58 | 8.46 | DN3ab | DN3cd | 0.73 | 10.59 | 2.50E-06 | 0.00055 | 0.062 | ENSMUSG00000021377 | protein_coding | Dek | DEK oncogene (DNA binding) | 110052 | 13 | 47084775 | 47106201 | -1 | Mm.131150 |
| 3.63 | 1.66 | 3.22 | 1.12 | DN3ab | DN3cd | -2.02 | -10.56 | 2.50E-06 | 0.00056 | 0.75 | ENSMUSG00000094497 | processed_pseudogene | Gm8210 | predicted pseudogene 8210 | NA | 1 | 43189156 | 43189638 | 1 | |
| 7.48 | 8.05 | 7.17 | 7.77 | DN3ab | DN3cd | 0.59 | 10.54 | 2.60E-06 | 0.00056 | 0.81 | ENSMUSG00000063696 | processed_pseudogene | Gm8730 | predicted pseudogene 8730 | NA | 8 | 102864845 | 102865798 | -1 | |
| 7.18 | 8.24 | 6.63 | 7.62 | DN3ab | DN3cd | 1.03 | 10.53 | 2.60E-06 | 0.00056 | 0.74 | ENSMUSG00000069301 | protein_coding | Hist1h2ag | histone cluster 1, H2ag | 319167 | 13 | 22042460 | 22042944 | -1 | Mm.263165 |
| 6.07 | 3.74 | 7.88 | 5.05 | DN3ab | DN3cd | -2.64 | -10.46 | 2.70E-06 | 0.00059 | 0.36 | ENSMUSG00000055148 | protein_coding | Klf2 | Kruppel-like factor 2 (lung) | 16598 | 8 | 72319033 | 72321656 | 1 | Mm.26938 |
| 1.84 | -3.92 | 1.97 | -3.11 | DN3ab | DN3cd | -5.45 | -10.42 | 2.80E-06 | 6.00E-04 | 0.54 | ENSMUSG00000066687 | protein_coding | Zbtb16 | zinc finger and BTB domain containing 16 | 235320 | 9 | 48654297 | 48836222 | -1 | Mm.457803 |
| 7.53 | 8.52 | 6.74 | 8.1 | DN3ab | DN3cd | 1.14 | 10.41 | 2.90E-06 | 6.00E-04 | 0.076 | ENSMUSG00000071478 | protein_coding | Hist1h2ad | histone cluster 1, H2ad | 319165 | 13 | 23574381 | 23574952 | 1 | Mm.390553 |
| 7.76 | 7.29 | 8.13 | 7.54 | DN3ab | DN3cd | -0.53 | -10.39 | 2.90E-06 | 6.00E-04 | 0.24 | ENSMUSG00000048865 | protein_coding | Arhgap30 | Rho GTPase activating protein 30 | 226652 | 1 | 171388954 | 171410298 | 1 | Mm.251048 |
| 0.07 | -3.92 | -0.18 | -4.02 | DN3ab | DN3cd | -3.92 | -10.38 | 2.90E-06 | 6.00E-04 | 0.84 | ENSMUSG00000083611 | processed_pseudogene | Gm8832 | predicted gene 8832 | NA | X | 168366464 | 168366858 | 1 | |
| 6.45 | 4.96 | 8.84 | 6.74 | DN3ab | DN3cd | -1.89 | -10.37 | 3.00E-06 | 6.00E-04 | 0.1 | ENSMUSG00000041959 | protein_coding | S100a10 | S100 calcium binding protein A10 (calpactin) | 20194 | 3 | 93555080 | 93564643 | 1 | Mm.1 |
| 7.58 | 6.5 | 9.59 | 7.96 | DN3ab | DN3cd | -1.42 | -10.3 | 3.10E-06 | 0.00063 | 0.039 | ENSMUSG00000049866 | protein_coding | Arl4c | ADP-ribosylation factor-like 4C | 320982 | 1 | 88673125 | 88702221 | -1 | Mm.27968 Mm.477970 |
| 6.1 | 7.11 | 5.33 | 6.47 | DN3ab | DN3cd | 1.07 | 10.28 | 3.20E-06 | 0.00063 | 0.57 | ENSMUSG00000069302 | protein_coding | Hist1h2ah | histone cluster 1, H2ah | 319168 | 13 | 22035164 | 22035568 | -1 | Mm.250564 Mm.262527 |
| 3.15 | 5.57 | 2.8 | 5.66 | DN3ab | DN3cd | 2.6 | 10.27 | 3.20E-06 | 0.00063 | 0.43 | ENSMUSG00000105703 | processed_transcript | Gm43305 | predicted gene 43305 | NA | 14 | 54292443 | 54297456 | -1 | |
| 1.82 | -1.12 | 6.2 | 1.59 | DN3ab | DN3cd | -4.28 | -10.25 | 3.20E-06 | 0.00063 | 0.11 | ENSMUSG00000002565 | protein_coding | Scin | scinderin | 20259 | 12 | 40059769 | 40134228 | -1 | Mm.2416 |
| 8.87 | 9.53 | 7.92 | 8.52 | DN3ab | DN3cd | 0.64 | 10.21 | 3.40E-06 | 0.00064 | 0.66 | ENSMUSG00000029730 | protein_coding | Mcm7 | minichromosome maintenance complex component 7 | 17220 | 5 | 138164583 | 138172422 | -1 | Mm.378965 |
| 2.51 | 1 | 4.98 | 2.93 | DN3ab | DN3cd | -1.92 | -10.17 | 3.50E-06 | 0.00065 | 0.2 | ENSMUSG00000024300 | protein_coding | Myo1f | myosin IF | 17916 | 17 | 33555707 | 33607764 | 1 | Mm.42019 |
| 4.27 | 2.12 | 4.62 | 2.85 | DN3ab | DN3cd | -1.94 | -10.16 | 3.50E-06 | 0.00065 | 0.34 | ENSMUSG00000041895 | protein_coding | Wipi1 | WD repeat domain, phosphoinositide interacting 1 | 52639 | 11 | 109573331 | 109611967 | -1 | Mm.35817 |
| -4.01 | -0.12 | -3.72 | 0.65 | DN3ab | DN3cd | 4.13 | 10.16 | 3.50E-06 | 0.00065 | 0.56 | ENSMUSG00000020811 | protein_coding | Wscd1 | WSC domain containing 1 | 216881 | 11 | 71749920 | 71789647 | 1 | Mm.48648 |
| 3.01 | 0.4 | 5.1 | 3.1 | DN3ab | DN3cd | -2.13 | -10.13 | 3.60E-06 | 0.00066 | 0.24 | ENSMUSG00000063605 | protein_coding | Ccdc102a | coiled-coil domain containing 102A | 234582 | 8 | 94902869 | 94918098 | -1 | Mm.277871 |
| 6.18 | 5.03 | 7.54 | 6 | DN3ab | DN3cd | -1.37 | -10.11 | 3.60E-06 | 0.00066 | 0.16 | ENSMUSG00000032009 | protein_coding | Sesn3 | sestrin 3 | 75747 | 9 | 14275067 | 14333101 | 1 | Mm.325126 Mm.439776 |
| 2.53 | 3.91 | 2.69 | 4.28 | DN3ab | DN3cd | 1.48 | 10.1 | 3.70E-06 | 0.00066 | 0.47 | ENSMUSG00000023947 | protein_coding | Nfkbie | nuclear factor of kappa light polypeptide gene enhancer in B cells inhibitor, epsilon | 18037 | 17 | 45555703 | 45563169 | 1 | Mm.57043 |
| 7.4 | 7.97 | 7.16 | 7.87 | DN3ab | DN3cd | 0.63 | 10.03 | 3.90E-06 | 7.00E-04 | 0.27 | ENSMUSG00000026421 | protein_coding | Csrp1 | cysteine and glycine-rich protein 1 | 13007 | 1 | 135720061 | 135752232 | 1 | Mm.196484 |
| 4.28 | 5.3 | 3.72 | 4.91 | DN3ab | DN3cd | 1.09 | 10.03 | 3.90E-06 | 7.00E-04 | 0.46 | ENSMUSG00000017697 | protein_coding | Ada | adenosine deaminase | 11486 | 2 | 163726584 | 163750239 | -1 | Mm.388 |
| 6.22 | 4.99 | 7.97 | 6.24 | DN3ab | DN3cd | -1.53 | -9.98 | 4.00E-06 | 0.00072 | 0.11 | ENSMUSG00000004665 | protein_coding | Cnn2 | calponin 2 | 12798 | 10 | 79988584 | 79996062 | 1 | Mm.157770 |
| 7.21 | 6.13 | 7.57 | 6.47 | DN3ab | DN3cd | -1.09 | -9.96 | 4.10E-06 | 0.00072 | 0.91 | ENSMUSG00000030671 | protein_coding | Pde3b | phosphodiesterase 3B, cGMP-inhibited | 18576 | 7 | 114415281 | 114539251 | 1 | Mm.430730 |
| 1.51 | -1.11 | 3.48 | 1.07 | DN3ab | DN3cd | -2.47 | -9.89 | 4.40E-06 | 0.00075 | 0.7 | ENSMUSG00000034028 | protein_coding | Cd226 | CD226 antigen | 225825 | 18 | 89197431 | 89270201 | 1 | Mm.139293 |
| 4.93 | 3.23 | 6.07 | 4.34 | DN3ab | DN3cd | -1.72 | -9.88 | 4.40E-06 | 0.00075 | 0.94 | ENSMUSG00000029468 | protein_coding | P2rx7 | purinergic receptor P2X, ligand-gated ion channel, 7 | 18439 | 5 | 122643911 | 122691432 | 1 | Mm.42026 |
| 6.86 | 5.99 | 7.12 | 6.43 | DN3ab | DN3cd | -0.78 | -9.87 | 4.40E-06 | 0.00075 | 0.24 | ENSMUSG00000021076 | protein_coding | Actr10 | ARP10 actin-related protein 10 | 56444 | 12 | 70937857 | 70964718 | 1 | Mm.29317 |
| 4.76 | 5.81 | 4.23 | 5.44 | DN3ab | DN3cd | 1.11 | 9.84 | 4.50E-06 | 0.00076 | 0.51 | ENSMUSG00000063021 | protein_coding | Hist1h2ak | histone cluster 1, H2ak | 319169 | 13 | 21753435 | 21753827 | -1 | Mm.370338 |
| 8.83 | 9.32 | 8.73 | 9.34 | DN3ab | DN3cd | 0.54 | 9.82 | 4.60E-06 | 0.00077 | 0.23 | ENSMUSG00000030742 | protein_coding | Lat | linker for activation of T cells | 16797 | 7 | 126363824 | 126369570 | -1 | Mm.10280 |
| 3.82 | 5.27 | 3.3 | 5.09 | DN3ab | DN3cd | 1.58 | 9.8 | 4.70E-06 | 0.00077 | 0.3 | ENSMUSG00000038963 | protein_coding | Slco4a1 | solute carrier organic anion transporter family, member 4a1 | 108115 | 2 | 180456245 | 180474867 | 1 | Mm.133687 |
| 6.19 | 5.16 | 6.26 | 5.53 | DN3ab | DN3cd | -0.88 | -9.8 | 4.70E-06 | 0.00077 | 0.065 | ENSMUSG00000108633 | TEC | Gm44694 | predicted gene 44694 | NA | 7 | 75707885 | 75709182 | 1 | |
| 0.15 | 3.93 | 0.52 | 3.84 | DN3ab | DN3cd | 3.55 | 9.78 | 4.80E-06 | 0.00077 | 0.53 | ENSMUSG00000019235 | protein_coding | Rps6kl1 | ribosomal protein S6 kinase-like 1 | 238323 | 12 | 85135249 | 85151264 | -1 | Mm.44829 |
| 6.66 | 6.02 | 7.09 | 6.29 | DN3ab | DN3cd | -0.72 | -9.78 | 4.80E-06 | 0.00077 | 0.26 | ENSMUSG00000022808 | protein_coding | Snx4 | sorting nexin 4 | 69150 | 16 | 33251442 | 33300269 | 1 | Mm.28196 Mm.396285 |
| 3.81 | 1 | 3.35 | 1.12 | DN3ab | DN3cd | -2.53 | -9.77 | 4.80E-06 | 0.00077 | 0.26 | ENSMUSG00000096712 | processed_pseudogene | Gm15454 | predicted gene 15454 | NA | 1 | 134518081 | 134518563 | 1 | |
| 6.45 | 7.26 | 5.42 | 6.36 | DN3ab | DN3cd | 0.86 | 9.77 | 4.80E-06 | 0.00077 | 0.45 | ENSMUSG00000017861 | protein_coding | Mybl2 | myeloblastosis oncogene-like 2 | 17865 | 2 | 163054687 | 163084688 | 1 | Mm.4594 |
| 6.72 | 5.51 | 7.62 | 6.66 | DN3ab | DN3cd | -1.07 | -9.74 | 4.90E-06 | 0.00078 | 0.27 | ENSMUSG00000026222 | protein_coding | Sp100 | nuclear antigen Sp100 | 20684 | 1 | 85649988 | 85709998 | 1 | Mm.290906 Mm.486514 |
| 5.33 | 6.28 | 5.59 | 6.64 | DN3ab | DN3cd | 1 | 9.7 | 5.10E-06 | 8.00E-04 | 0.62 | ENSMUSG00000030745 | protein_coding | Il21r | interleukin 21 receptor | 60504 | 7 | 125603429 | 125633570 | 1 | Mm.155643 |
| 7.29 | 8.27 | 6.31 | 7.66 | DN3ab | DN3cd | 1.12 | 9.68 | 5.20E-06 | 8.00E-04 | 0.11 | ENSMUSG00000023942 | protein_coding | Slc29a1 | solute carrier family 29 (nucleoside transporters), member 1 | 63959 | 17 | 45585200 | 45599606 | -1 | Mm.29744 Mm.486335 |
| 6.42 | 7.35 | 5.58 | 6.96 | DN3ab | DN3cd | 1.11 | 9.68 | 5.20E-06 | 8.00E-04 | 0.042 | ENSMUSG00000061991 | protein_coding | Hist1h2af | histone cluster 1, H2af | 319173 | 13 | 23533906 | 23534304 | 1 | Mm.377878 |
| 1.54 | 3.56 | 4.57 | 6 | DN3ab | DN3cd | 1.55 | 9.67 | 5.20E-06 | 8.00E-04 | 0.12 | ENSMUSG00000099413 | lincRNA | Gm17767 | predicted gene, 17767 | NA | 1 | 51507086 | 51596718 | -1 | |
| 4.35 | 3.14 | 5.1 | 4.09 | DN3ab | DN3cd | -1.09 | -9.67 | 5.30E-06 | 8.00E-04 | 0.38 | ENSMUSG00000024851 | protein_coding | Pitpnm1 | phosphatidylinositol transfer protein, membrane-associated 1 | 18739 | 19 | 4099998 | 4113965 | 1 | Mm.1860 |
| 7.93 | 8.5 | 7.16 | 7.78 | DN3ab | DN3cd | 0.59 | 9.66 | 5.30E-06 | 8.00E-04 | 0.7 | ENSMUSG00000005410 | protein_coding | Mcm5 | minichromosome maintenance complex component 5 | 17218 | 8 | 75109569 | 75128439 | 1 | Mm.5048 |
| 7.35 | 7.95 | 7.31 | 7.78 | DN3ab | DN3cd | 0.54 | 9.65 | 5.30E-06 | 8.00E-04 | 0.24 | ENSMUSG00000036202 | protein_coding | Rif1 | replication timing regulatory factor 1 | 51869 | 2 | 52072832 | 52122383 | 1 | Mm.254530 Mm.389247 |
| 1.19 | 3.82 | 1.07 | 3.08 | DN3ab | DN3cd | 2.37 | 9.63 | 5.40E-06 | 8.00E-04 | 0.2 | ENSMUSG00000091695 | protein_coding | Gm17669 | predicted gene, 17669 | NA | 18 | 67562387 | 67562851 | 1 | |
| 8.19 | 7.69 | 8.89 | 8.5 | DN3ab | DN3cd | -0.45 | -9.63 | 5.40E-06 | 8.00E-04 | 0.21 | ENSMUSG00000019189 | protein_coding | Rnf145 | ring finger protein 145 | 74315 | 11 | 44518964 | 44565520 | 1 | Mm.156771 |
| 6.2 | 5.08 | 7.86 | 6.19 | DN3ab | DN3cd | -1.44 | -9.62 | 5.50E-06 | 8.00E-04 | 0.061 | ENSMUSG00000000594 | protein_coding | Gm2a | GM2 ganglioside activator protein | 14667 | 11 | 55098115 | 55113029 | 1 | Mm.287807 |
| 3.59 | 4.55 | 4.01 | 5.14 | DN3ab | DN3cd | 1.05 | 9.61 | 5.50E-06 | 0.00081 | 0.39 | ENSMUSG00000041515 | protein_coding | Irf8 | interferon regulatory factor 8 | 15900 | 8 | 120736358 | 120756694 | 1 | Mm.334861 |
| 8.42 | 7.78 | 8.36 | 7.78 | DN3ab | DN3cd | -0.62 | -9.58 | 5.70E-06 | 0.00082 | 0.61 | ENSMUSG00000025066 | protein_coding | Sfr1 | SWI5 dependent recombination repair 1 | 67788 | 19 | 47731756 | 47735588 | 1 | Mm.181610 |
| 5.76 | 4.84 | 6.98 | 5.76 | DN3ab | DN3cd | -1.09 | -9.57 | 5.70E-06 | 0.00082 | 0.19 | ENSMUSG00000003644 | protein_coding | Rps6ka1 | ribosomal protein S6 kinase polypeptide 1 | 20111 | 4 | 133847290 | 133887797 | -1 | Mm.301827 |
| 10.1 | 8.92 | 12.82 | 11.07 | DN3ab | DN3cd | -1.57 | -9.57 | 5.70E-06 | 0.00082 | 0.11 | ENSMUSG00000049775 | protein_coding | Tmsb4x | thymosin, beta 4, X chromosome | 19241 | X | 167207093 | 167209315 | -1 | Mm.142729 Mm.490263 |
| 6.02 | 5.26 | 6.8 | 5.98 | DN3ab | DN3cd | -0.79 | -9.52 | 6.00E-06 | 0.00085 | 0.74 | ENSMUSG00000049470 | protein_coding | Aff4 | AF4/FMR2 family, member 4 | 93736 | 11 | 53350833 | 53421830 | 1 | Mm.395281 |
| 9.1 | 9.86 | 8.95 | 9.45 | DN3ab | DN3cd | 0.64 | 9.5 | 6.10E-06 | 0.00086 | 0.035 | ENSMUSG00000090862 | protein_coding | Rps13 | ribosomal protein S13 | 68052 | 7 | 116331505 | 116334195 | -1 | Mm.14798 Mm.345443 Mm.425541 |
| 4.92 | 2 | 6.78 | 4.73 | DN3ab | DN3cd | -2.25 | -9.47 | 6.20E-06 | 0.00087 | 0.12 | ENSMUSG00000061577 | protein_coding | Adgrg5 | adhesion G protein-coupled receptor G5 | 382045 | 8 | 94923694 | 94943280 | 1 | Mm.334726 |
| 5.91 | 5.19 | 6.6 | 5.62 | DN3ab | DN3cd | -0.85 | -9.46 | 6.30E-06 | 0.00087 | 0.13 | ENSMUSG00000042700 | protein_coding | Sipa1l1 | signal-induced proliferation-associated 1 like 1 | 217692 | 12 | 82169320 | 82451786 | 1 | Mm.261333 |
| 2.03 | 3.74 | 1.5 | 3.7 | DN3ab | DN3cd | 1.9 | 9.45 | 6.30E-06 | 0.00087 | 0.23 | ENSMUSG00000025780 | protein_coding | Itih5 | inter-alpha (globulin) inhibitor H5 | 209378 | 2 | 10153571 | 10256529 | 1 | Mm.313876 |
| 3.56 | 4.7 | 3.4 | 4.61 | DN3ab | DN3cd | 1.17 | 9.44 | 6.40E-06 | 0.00087 | 0.79 | ENSMUSG00000020785 | protein_coding | Camkk1 | calcium/calmodulin-dependent protein kinase kinase 1, alpha | 55984 | 11 | 73019008 | 73042073 | 1 | Mm.9998 |
| 1.09 | 3.12 | 1.75 | 3.76 | DN3ab | DN3cd | 2.02 | 9.44 | 6.40E-06 | 0.00087 | 0.96 | ENSMUSG00000004085 | protein_coding | Map3k20 | mitogen-activated protein kinase kinase kinase 20 | 65964 | 2 | 72285637 | 72442610 | 1 | Mm.304143 Mm.314618 |
| -0.76 | -3.92 | 3.25 | 0.6 | DN3ab | DN3cd | -2.77 | -9.43 | 6.40E-06 | 0.00087 | 0.45 | ENSMUSG00000073010 | protein_coding | Gm5127 | predicted gene 5127 | 331493 | X | 106583186 | 106710557 | 1 | Mm.249319 |
| 6.46 | 5.52 | 7.37 | 6.01 | DN3ab | DN3cd | -1.16 | -9.41 | 6.50E-06 | 0.00088 | 0.072 | ENSMUSG00000022102 | protein_coding | Dok2 | docking protein 2 | 13449 | 14 | 70766036 | 70778495 | 1 | Mm.243323 |
| 5.51 | 4.68 | 5.81 | 5.11 | DN3ab | DN3cd | -0.76 | -9.36 | 6.90E-06 | 0.00092 | 0.42 | ENSMUSG00000035133 | protein_coding | Arhgap5 | Rho GTPase activating protein 5 | 11855 | 12 | 52503972 | 52571975 | 1 | Mm.35059 Mm.466962 Mm.474991 |
| 3.79 | 0.26 | 3.3 | 0.12 | DN3ab | DN3cd | -3.38 | -9.3 | 7.20E-06 | 0.00095 | 0.64 | ENSMUSG00000052560 | protein_coding | Cpne8 | copine VIII | 66871 | 15 | 90487482 | 90679432 | -1 | Mm.290991 |
| 3.68 | 4.93 | 3.32 | 4.4 | DN3ab | DN3cd | 1.18 | 9.28 | 7.30E-06 | 0.00096 | 0.5 | ENSMUSG00000069309 | protein_coding | Hist1h2an | histone cluster 1, H2an | 319170 | 13 | 21786826 | 21787218 | -1 | |
| 7.57 | 8.24 | 7.04 | 7.73 | DN3ab | DN3cd | 0.68 | 9.27 | 7.40E-06 | 0.00096 | 0.96 | ENSMUSG00000028884 | protein_coding | Rpa2 | replication protein A2 | 19891 | 4 | 132768332 | 132778752 | 1 | Mm.2870 |
| 7.51 | 8.05 | 6.74 | 7.56 | DN3ab | DN3cd | 0.65 | 9.27 | 7.40E-06 | 0.00096 | 0.034 | ENSMUSG00000019773 | protein_coding | Fbxo5 | F-box protein 5 | 67141 | 10 | 5799160 | 5805600 | -1 | Mm.197520 |
| 5.3 | 4.48 | 6.86 | 5.72 | DN3ab | DN3cd | -1.02 | -9.26 | 7.50E-06 | 0.00096 | 0.13 | ENSMUSG00000056888 | protein_coding | Glipr1 | GLI pathogenesis-related 1 (glioma) | 73690 | 10 | 111985448 | 112002631 | -1 | Mm.173790 |
| 7.02 | 6.04 | 7.6 | 6.14 | DN3ab | DN3cd | -1.21 | -9.26 | 7.50E-06 | 0.00096 | 0.053 | ENSMUSG00000020601 | protein_coding | Trib2 | tribbles pseudokinase 2 | 217410 | 12 | 15791727 | 15816922 | -1 | Mm.491136 |
| 1.09 | -0.93 | 3.91 | 1.3 | DN3ab | DN3cd | -2.46 | -9.24 | 7.60E-06 | 0.00097 | 0.34 | ENSMUSG00000036019 | protein_coding | Tmtc2 | transmembrane and tetratricopeptide repeat containing 2 | 278279 | 10 | 105187663 | 105574451 | -1 | Mm.25225 |
| 3.8 | 1.41 | 7.78 | 3.46 | DN3ab | DN3cd | -3.87 | -9.24 | 7.60E-06 | 0.00097 | 0.04 | ENSMUSG00000045092 | protein_coding | S1pr1 | sphingosine-1-phosphate receptor 1 | 13609 | 3 | 115710433 | 115715072 | -1 | Mm.982 |
| 5.02 | 3.37 | 6.1 | 4.89 | DN3ab | DN3cd | -1.37 | -9.23 | 7.70E-06 | 0.00097 | 0.15 | ENSMUSG00000025278 | protein_coding | Flnb | filamin, beta | 286940 | 14 | 7817957 | 7951588 | 1 | Mm.489652 |
| 5.63 | 4.67 | 7.38 | 6.38 | DN3ab | DN3cd | -0.99 | -9.21 | 7.80E-06 | 0.00098 | 0.87 | ENSMUSG00000056671 | protein_coding | Prelid2 | PRELI domain containing 2 | 77619 | 18 | 41875696 | 41951194 | -1 | Mm.30649 |
| 5.88 | 5.13 | 6.9 | 5.82 | DN3ab | DN3cd | -0.93 | -9.18 | 8.00E-06 | 0.00099 | 0.079 | ENSMUSG00000021676 | protein_coding | Iqgap2 | IQ motif containing GTPase activating protein 2 | 544963 | 13 | 95627177 | 95891922 | -1 | Mm.38878 |
| 2.18 | 4.21 | 1.76 | 4.35 | DN3ab | DN3cd | 2.25 | 9.17 | 8.00E-06 | 0.00099 | 0.27 | ENSMUSG00000034265 | protein_coding | Zdhhc14 | zinc finger, DHHC domain containing 14 | 224454 | 17 | 5492557 | 5753811 | 1 | Mm.487663 Mm.491143 |
| 2.21 | 0.62 | 5.23 | 2.67 | DN3ab | DN3cd | -2.33 | -9.17 | 8.10E-06 | 0.00099 | 0.094 | ENSMUSG00000046280 | protein_coding | She | src homology 2 domain-containing transforming protein E | 214547 | 3 | 89831370 | 89858834 | 1 | Mm.225 |
| 4.98 | 2.22 | 5.72 | 2.61 | DN3ab | DN3cd | -2.96 | -9.17 | 8.10E-06 | 0.00099 | 0.61 | ENSMUSG00000003545 | protein_coding | Fosb | FBJ osteosarcoma oncogene B | 14282 | 7 | 19302696 | 19310051 | -1 | Mm.248335 |
| 3.25 | 4.44 | 3.39 | 4.81 | DN3ab | DN3cd | 1.3 | 9.16 | 8.20E-06 | 0.001 | 0.41 | ENSMUSG00000030122 | protein_coding | Ptms | parathymosin | 69202 | 6 | 124913681 | 124920103 | -1 | Mm.142187 |
| 8.36 | 8.82 | 7.75 | 8.22 | DN3ab | DN3cd | 0.46 | 9.12 | 8.40E-06 | 0.001 | 0.88 | ENSMUSG00000029472 | protein_coding | Anapc5 | anaphase-promoting complex subunit 5 | 59008 | 5 | 122787459 | 122821339 | -1 | Mm.332667 |
| 6.36 | 4.47 | 7.69 | 6.45 | DN3ab | DN3cd | -1.48 | -9.02 | 9.20E-06 | 0.0011 | 0.043 | ENSMUSG00000060131 | protein_coding | Atp8b4 | ATPase, class I, type 8B, member 4 | 241633 | 2 | 126320973 | 126500674 | -1 | Mm.222235 |
| 6.75 | 6.22 | 7.36 | 6.71 | DN3ab | DN3cd | -0.6 | -8.99 | 9.50E-06 | 0.0011 | 0.37 | ENSMUSG00000026827 | protein_coding | Gpd2 | glycerol phosphate dehydrogenase 2, mitochondrial | 14571 | 2 | 57237635 | 57370719 | 1 | Mm.3711 |
| 1.2 | -2.34 | 4.34 | 2.11 | DN3ab | DN3cd | -2.44 | -8.98 | 9.60E-06 | 0.0011 | 0.061 | ENSMUSG00000040229 | protein_coding | Gpr34 | G protein-coupled receptor 34 | 23890 | X | 13632089 | 13640858 | 1 | Mm.487326 |
| -4.01 | 0.59 | -3.72 | 1.6 | DN3ab | DN3cd | 4.96 | 8.95 | 9.90E-06 | 0.0012 | 0.53 | ENSMUSG00000031750 | protein_coding | Il34 | interleukin 34 | 76527 | 8 | 110741829 | 110805924 | -1 | Mm.77697 |
| 6.17 | 5.57 | 6.71 | 5.88 | DN3ab | DN3cd | -0.72 | -8.9 | 1.00E-05 | 0.0012 | 0.14 | ENSMUSG00000028228 | protein_coding | Cpne3 | copine III | 70568 | 4 | 19519254 | 19570108 | -1 | Mm.38390 |
| 6.07 | 5.08 | 6.05 | 5.36 | DN3ab | DN3cd | -0.85 | -8.87 | 1.10E-05 | 0.0012 | 0.1 | ENSMUSG00000037369 | protein_coding | Kdm6a | lysine (K)-specific demethylase 6A | 22289 | X | 18162575 | 18279936 | 1 | Mm.257498 Mm.417709 |
| -2.43 | 3.32 | -2.6 | 3.08 | DN3ab | DN3cd | 5.71 | 8.84 | 1.10E-05 | 0.0012 | 0.96 | ENSMUSG00000041361 | protein_coding | Myzap | myocardial zonula adherens protein | 102371 | 9 | 71504347 | 71592360 | -1 | Mm.27585 |
| -1.12 | 4.58 | -1.92 | 4.55 | DN3ab | DN3cd | 6.04 | 8.8 | 1.10E-05 | 0.0013 | 0.6 | ENSMUSG00000069049 | protein_coding | Eif2s3y | eukaryotic translation initiation factor 2, subunit 3, structural gene Y-linked | 26908 | Y | 1010543 | 1028847 | 1 | Mm.250909 |
| 0.69 | -3.92 | 0.27 | -3.34 | DN3ab | DN3cd | -4.17 | -8.79 | 1.10E-05 | 0.0013 | 0.29 | ENSMUSG00000081860 | processed_pseudogene | Gm11533 | predicted gene 11533 | NA | 11 | 97037007 | 97037483 | 1 | |
| 5.83 | 3.93 | 10.12 | 6.52 | DN3ab | DN3cd | -3.18 | -8.76 | 1.20E-05 | 0.0013 | 0.031 | ENSMUSG00000033577 | protein_coding | Myo6 | myosin VI | 17920 | 9 | 80165031 | 80311729 | 1 | Mm.4040 |
| 3.61 | 4.62 | 3.6 | 4.9 | DN3ab | DN3cd | 1.14 | 8.74 | 1.20E-05 | 0.0013 | 0.26 | ENSMUSG00000022307 | protein_coding | Oxr1 | oxidation resistance 1 | 170719 | 15 | 41447482 | 41861048 | 1 | Mm.254267 |
| 4.86 | 3.65 | 6.66 | 5.63 | DN3ab | DN3cd | -1.09 | -8.73 | 1.20E-05 | 0.0013 | 0.51 | ENSMUSG00000020101 | protein_coding | Vsir | V-set immunoregulatory receptor | 74048 | 10 | 60346851 | 60372684 | 1 | Mm.273584 |
| 2.65 | 1.57 | 5.53 | 3.78 | DN3ab | DN3cd | -1.6 | -8.72 | 1.20E-05 | 0.0013 | 0.12 | ENSMUSG00000038642 | protein_coding | Ctss | cathepsin S | 13040 | 3 | 95526786 | 95556403 | 1 | Mm.3619 |
| 4.37 | 3.32 | 5.96 | 4.56 | DN3ab | DN3cd | -1.28 | -8.68 | 1.30E-05 | 0.0014 | 0.26 | ENSMUSG00000036526 | protein_coding | Card11 | caspase recruitment domain family, member 11 | 108723 | 5 | 140872990 | 141000582 | -1 | Mm.46187 |
| 3.99 | 3.06 | 6.46 | 4.9 | DN3ab | DN3cd | -1.37 | -8.67 | 1.30E-05 | 0.0014 | 0.049 | ENSMUSG00000024399 | protein_coding | Ltb | lymphotoxin B | 16994 | 17 | 35194439 | 35196320 | 1 | Mm.1715 |

| | | | | | | | | | | | | | | | | | | | | |
|---|---|---|---|---|---|---|---|---|---|---|---|---|---|---|---|---|---|---|---|---|
| 7.87 | 7.38 | 8.46 | 7.81 | DN3ab | DN3cd | -0.57 | -8.66 | 1.30E-05 | 0.0014 | 0.2 | ENSMUSG00000028530 | protein_coding | Jak1 | Janus kinase 1 | 16451 | 4 | 101152367 | 101265282 | -1 | Mm.289657 |
| 8.69 | 7.94 | 9.73 | 8.68 | DN3ab | DN3cd | -0.91 | -8.65 | 1.30E-05 | 0.0014 | 0.17 | ENSMUSG00000031207 | protein_coding | Msn | moesin | 17698 | X | 96096042 | 96168552 | 1 | Mm.138876 |
| 5.47 | 6.27 | 5.06 | 5.66 | DN3ab | DN3cd | 0.72 | 8.62 | 1.30E-05 | 0.0014 | 0.19 | ENSMUSG00000105987 | processed_transcript | AI506816 | expressed sequence AI506816 | NA | 5 | 23698296 | 23712667 | -1 | |
| 5.07 | 3.93 | 6.2 | 4.73 | DN3ab | DN3cd | -1.32 | -8.59 | 1.40E-05 | 0.0014 | 0.3 | ENSMUSG00000038732 | protein_coding | Mboat1 | membrane bound O-acyltransferase domain containing 1 | 218121 | 13 | 30136489 | 30246717 | 1 | Mm.89682 |
| -2.43 | 0.81 | -3.72 | -0.46 | DN3ab | DN3cd | 3.24 | 8.59 | 1.40E-05 | 0.0014 | 0.98 | ENSMUSG00000039269 | protein_coding | 2300002M23Rik | RIKEN cDNA 2300002M23 gene | 69542 | 17 | 35567450 | 35568952 | 1 | Mm.265291 |
| 5.8 | 4.89 | 7.08 | 5.71 | DN3ab | DN3cd | -1.17 | -8.57 | 1.40E-05 | 0.0014 | 0.089 | ENSMUSG00000024789 | protein_coding | Jak2 | Janus kinase 2 | 16452 | 19 | 29251828 | 29313080 | 1 | Mm.275839 |
| 8.21 | 8.77 | 8.03 | 8.48 | DN3ab | DN3cd | 0.5 | 8.56 | 1.40E-05 | 0.0015 | 0.36 | ENSMUSG00000069972 | processed_pseudogene | Rps13-ps2 | ribosomal protein S13, pseudogene 2 | NA | 7 | 88530446 | 88530898 | -1 | |
| 6.94 | 6.11 | 6.87 | 6.27 | DN3ab | DN3cd | -0.72 | -8.55 | 1.40E-05 | 0.0015 | 0.17 | ENSMUSG00000017466 | protein_coding | Timp2 | tissue inhibitor of metalloproteinase 2 | 21858 | 11 | 118301069 | 118355740 | -1 | Mm.206505 |
| 7.21 | 6 | 9.23 | 7.4 | DN3ab | DN3cd | -1.59 | -8.48 | 1.50E-05 | 0.0016 | 0.099 | ENSMUSG00000025647 | protein_coding | Shisa5 | shisa family member 5 | 66940 | 9 | 109038565 | 109057777 | 1 | Mm.196533 Mm.475600 |
| 5.04 | 6.26 | 4.76 | 5.94 | DN3ab | DN3cd | 1.2 | 8.47 | 1.50E-05 | 0.0016 | 0.89 | ENSMUSG00000027714 | protein_coding | Exosc9 | exosome component 9 | 50911 | 3 | 36552606 | 36565727 | 1 | Mm.116711 |
| 2.52 | 5.61 | -0.25 | 4.75 | DN3ab | DN3cd | 3.41 | 8.46 | 1.50E-05 | 0.0016 | 0.068 | ENSMUSG00000036858 | protein_coding | Ptcra | pre T cell antigen receptor alpha | 19208 | 17 | 46755663 | 46763712 | -1 | Mm.215173 |
| 6.66 | 6.15 | 7.12 | 6.56 | DN3ab | DN3cd | -0.54 | -8.44 | 1.60E-05 | 0.0016 | 0.68 | ENSMUSG00000021969 | protein_coding | Zdhhc20 | zinc finger, DHHC domain containing 20 | 75965 | 14 | 57832703 | 57890276 | -1 | Mm.29044 |
| -0.7 | -3.92 | 1.56 | -1.48 | DN3ab | DN3cd | -3.12 | -8.41 | 1.60E-05 | 0.0016 | 0.8 | ENSMUSG00000024675 | protein_coding | Ms4a4c | membrane-spanning 4-domains, subfamily A, member 4C | 64380 | 19 | 11404770 | 11427246 | 1 | Mm.440715 |
| 3.5 | 2.42 | 4.31 | 2.75 | DN3ab | DN3cd | -1.34 | -8.4 | 1.60E-05 | 0.0016 | 0.11 | ENSMUSG00000051890 | protein_coding | Klhdc1 | kelch domain containing 1 | 271005 | 12 | 69241176 | 69284632 | 1 | Mm.180969 |
| 5.74 | 6.48 | 4.74 | 5.77 | DN3ab | DN3cd | 0.84 | 8.4 | 1.60E-05 | 0.0016 | 0.15 | ENSMUSG00000026669 | protein_coding | Mcm10 | minichromosome maintenance 10 replication initiation factor | 70024 | 2 | 4989714 | 5012791 | -1 | Mm.23828 Mm.447935 |
| 9.14 | 8.42 | 10.23 | 9.46 | DN3ab | DN3cd | -0.75 | -8.39 | 1.70E-05 | 0.0016 | 0.74 | ENSMUSG00000039936 | protein_coding | Pik3cd | phosphatidylinositol-4,5-bisphosphate 3-kinase catalytic subunit delta | 18707 | 4 | 149649168 | 149702571 | -1 | Mm.229108 |
| 3.65 | 6.02 | 2.26 | 5.86 | DN3ab | DN3cd | 2.72 | 8.38 | 1.70E-05 | 0.0016 | 0.079 | ENSMUSG00000030844 | protein_coding | Rgs10 | regulator of G-protein signalling 10 | 67865 | 7 | 128373621 | 128418758 | -1 | Mm.18635 |
| 6.78 | 7.42 | 6.23 | 6.9 | DN3ab | DN3cd | 0.65 | 8.37 | 1.70E-05 | 0.0016 | 0.82 | ENSMUSG00000022360 | protein_coding | Atad2 | ATPase family, AAA domain containing 2 | 70472 | 15 | 58094044 | 58135082 | -1 | Mm.221758 |
| 7.92 | 8.41 | 7.06 | 7.7 | DN3ab | DN3cd | 0.55 | 8.32 | 1.80E-05 | 0.0017 | 0.26 | ENSMUSG00000030978 | protein_coding | Rrm1 | ribonucleotide reductase M1 | 20133 | 7 | 102441695 | 102469771 | 1 | Mm.197486 Mm.415177 |
| -2.61 | 1.99 | -3.72 | 2.06 | DN3ab | DN3cd | 5.18 | 8.31 | 1.80E-05 | 0.0017 | 0.36 | ENSMUSG00000065191 | snRNA | Gm23458 | predicted gene, 23458 | NA | 5 | 123360832 | 123361000 | 1 | |
| 4.57 | 5.42 | 3.94 | 4.58 | DN3ab | DN3cd | 0.77 | 8.31 | 1.80E-05 | 0.0017 | 0.22 | ENSMUSG00000023908 | protein_coding | Pkmyt1 | protein kinase, membrane associated tyrosine/threonine 1 | 268930 | 17 | 23726336 | 23736735 | 1 | Mm.182193 |
| 2.56 | -0.94 | 1.69 | -0.92 | DN3ab | DN3cd | -3.1 | -8.3 | 1.80E-05 | 0.0017 | 0.23 | ENSMUSG00000008734 | protein_coding | Gprc5b | G protein-coupled receptor, family C, group 5, member B | 64297 | 7 | 118972047 | 118995211 | -1 | Mm.103439 |
| 5.93 | 6.64 | 5.28 | 6.14 | DN3ab | DN3cd | 0.77 | 8.3 | 1.80E-05 | 0.0017 | 0.46 | ENSMUSG00000033970 | protein_coding | Rfc3 | replication factor C (activator 1) 3 | 69263 | 5 | 151642756 | 151651242 | -1 | Mm.12553 |
| 5.17 | 4.5 | 5.31 | 4.66 | DN3ab | DN3cd | -0.66 | -8.29 | 1.80E-05 | 0.0017 | 0.95 | ENSMUSG00000041040 | protein_coding | Fam117b | family with sequence similarity 117, member B | 72750 | 1 | 59913006 | 59985346 | 1 | Mm.258250 |
| 8.28 | 7.46 | 9.88 | 8.84 | DN3ab | DN3cd | -0.95 | -8.28 | 1.80E-05 | 0.0017 | 0.4 | ENSMUSG00000091955 | protein_coding | Gm9844 | predicted pseudogene 9844 | NA | 7 | 24862213 | 24862697 | 1 | |
| 6.12 | 5.53 | 6.63 | 5.77 | DN3ab | DN3cd | -0.73 | -8.25 | 1.90E-05 | 0.0017 | 0.098 | ENSMUSG00000015837 | protein_coding | Sqstm1 | sequestosome 1 | 18412 | 11 | 50199366 | 50210827 | -1 | Mm.40828 |
| 6.66 | 5.89 | 7.36 | 6.61 | DN3ab | DN3cd | -0.76 | -8.24 | 1.90E-05 | 0.0017 | 0.91 | ENSMUSG00000041736 | protein_coding | Tspo | translocator protein | 12257 | 15 | 83563592 | 83574203 | 1 | Mm.1508 |
| 4.37 | 5.2 | 3.5 | 4.28 | DN3ab | DN3cd | 0.81 | 8.23 | 1.90E-05 | 0.0018 | 0.8 | ENSMUSG00000007080 | protein_coding | Pole | polymerase (DNA directed), epsilon | 18973 | 5 | 110286306 | 110337474 | 1 | Mm.35061 |
| 6.61 | 5.9 | 7.23 | 6.69 | DN3ab | DN3cd | -0.62 | -8.21 | 2.00E-05 | 0.0018 | 0.24 | ENSMUSG00000028496 | protein_coding | Mllt3 | myeloid/lymphoid or mixed-lineage leukemia; translocated to, 3 | 70122 | 4 | 87769925 | 88033364 | -1 | Mm.288898 Mm.472285 |
| 6.08 | 6.58 | 5.87 | 6.45 | DN3ab | DN3cd | 0.53 | 8.2 | 2.00E-05 | 0.0018 | 0.57 | ENSMUSG00000052825 | processed_pseudogene | Gm9892 | predicted gene 9892 | NA | 8 | 52196065 | 52197056 | -1 | |
| 5.21 | 4.34 | 5.87 | 4.59 | DN3ab | DN3cd | -1.08 | -8.19 | 2.00E-05 | 0.0018 | 0.11 | ENSMUSG00000020520 | protein_coding | Galnt10 | polypeptide N-acetylgalactosaminyltransferase 10 | 171212 | 11 | 57645442 | 57787514 | 1 | Mm.271670 |
| 2.94 | 3.98 | 2.91 | 4.04 | DN3ab | DN3cd | 1.08 | 8.16 | 2.10E-05 | 0.0018 | 0.72 | ENSMUSG00000024669 | protein_coding | Cd5 | CD5 antigen | 12507 | 19 | 10718143 | 10738974 | -1 | Mm.779 |
| 4.41 | 3.5 | 5.49 | 4.47 | DN3ab | DN3cd | -0.98 | -8.15 | 2.10E-05 | 0.0019 | 0.66 | ENSMUSG00000032089 | protein_coding | Il10ra | interleukin 10 receptor, alpha | 16154 | 9 | 45253837 | 45269149 | -1 | Mm.379327 |
| 5.13 | 4.39 | 5.88 | 4.94 | DN3ab | DN3cd | -0.85 | -8.14 | 2.10E-05 | 0.0019 | 0.34 | ENSMUSG00000032175 | protein_coding | Tyk2 | tyrosine kinase 2 | 54721 | 9 | 21104068 | 21131243 | -1 | Mm.20249 Mm.450004 |
| 5.09 | 5.73 | 4.71 | 5.62 | DN3ab | DN3cd | 0.76 | 8.12 | 2.10E-05 | 0.0019 | 0.13 | ENSMUSG00000024056 | protein_coding | Ndc80 | NDC80 kinetochore complex component | 67052 | 17 | 71496100 | 71526857 | -1 | Mm.225956 |
| 7.56 | 8.17 | 7.36 | 7.9 | DN3ab | DN3cd | 0.58 | 8.12 | 2.10E-05 | 0.0019 | 0.62 | ENSMUSG00000097415 | lincRNA | AU020206 | expressed sequence AU020206 | NA | 7 | 75769038 | 75782099 | -1 | |
| 0.73 | 4.63 | -0.97 | 5.01 | DN3ab | DN3cd | 4.48 | 8.12 | 2.10E-05 | 0.0019 | 0.081 | ENSMUSG00000048612 | protein_coding | Myof | myoferlin | 226101 | 19 | 37899036 | 38043577 | -1 | Mm.34674 |
| 3.19 | 4.07 | 3.17 | 4.22 | DN3ab | DN3cd | 0.96 | 8.12 | 2.10E-05 | 0.0019 | 0.47 | ENSMUSG00000018796 | protein_coding | Acsl1 | acyl-CoA synthetase long-chain family member 1 | 14081 | 8 | 46471037 | 46536051 | 1 | Mm.210323 |
| 5.33 | 6.03 | 5.05 | 5.61 | DN3ab | DN3cd | 0.64 | 8.1 | 2.20E-05 | 0.0019 | 0.38 | ENSMUSG00000020029 | protein_coding | Nudt4 | nudix (nucleoside diphosphate linked moiety X)-type motif 4 | 71207 | 10 | 95547007 | 95564146 | -1 | Mm.24397 |
| 6.24 | 5.69 | 6.84 | 6.26 | DN3ab | DN3cd | -0.56 | -8.09 | 2.20E-05 | 0.0019 | 0.79 | ENSMUSG00000049744 | protein_coding | Arhgap15 | Rho GTPase activating protein 15 | 76117 | 2 | 43748824 | 44395953 | 1 | Mm.217350 |
| 5.03 | 5.94 | 4.33 | 5.77 | DN3ab | DN3cd | 1.12 | 8.08 | 2.20E-05 | 0.0019 | 0.042 | ENSMUSG00000031505 | protein_coding | Naxd | NAD(P)HX dehydratase | 69225 | 8 | 11497506 | 11514960 | 1 | Mm.64911 Mm.267998 |
| 8.52 | 9.02 | 7.66 | 8.32 | DN3ab | DN3cd | 0.56 | 8.08 | 2.20E-05 | 0.0019 | 0.25 | ENSMUSG00000028693 | protein_coding | Nasp | nuclear autoantigenic sperm protein (histone-binding) | 50927 | 4 | 116601052 | 116627941 | -1 | Mm.257181 |
| 2.6 | 1.25 | 3.71 | 2.42 | DN3ab | DN3cd | -1.32 | -8.06 | 2.30E-05 | 0.0019 | 0.86 | ENSMUSG00000047123 | protein_coding | Ticam1 | toll-like receptor adaptor molecule 1 | 106759 | 17 | 56269319 | 56276786 | -1 | Mm.203952 |
| 3.34 | 0.33 | 3.82 | 1.68 | DN3ab | DN3cd | -2.46 | -8.02 | 2.40E-05 | 0.002 | 0.17 | ENSMUSG00000020183 | protein_coding | Cpm | carboxypeptidase M | 70574 | 10 | 117629500 | 117687352 | 1 | Mm.339332 |
| 3.15 | 4.49 | 2.96 | 4.37 | DN3ab | DN3cd | 1.37 | 7.98 | 2.50E-05 | 0.002 | 0.83 | ENSMUSG00000024670 | protein_coding | Cd6 | CD6 antigen | 12511 | 19 | 10789341 | 10830058 | -1 | Mm.290897 |
| 4.5 | 3.57 | 5.21 | 4.04 | DN3ab | DN3cd | -1.06 | -7.95 | 2.50E-05 | 0.0021 | 0.37 | ENSMUSG00000023088 | protein_coding | Abcc1 | ATP-binding cassette, sub-family C (CFTR/MRP), member 1 | 17250 | 16 | 14361558 | 14475737 | 1 | Mm.196634 |
| 11.3 | 11.67 | 11.2 | 11.5 | DN3ab | DN3cd | 0.34 | 7.92 | 2.60E-05 | 0.0021 | 0.43 | ENSMUSG00000101111 | unprocessed_pseudogene | Gm28437 | predicted gene 28437 | NA | 1 | 24613189 | 24613971 | -1 | |
| 6.92 | 6.34 | 7.75 | 6.8 | DN3ab | DN3cd | -0.77 | -7.91 | 2.60E-05 | 0.0021 | 0.033 | ENSMUSG00000041272 | protein_coding | Tox | thymocyte selection-associated high mobility group box | 252838 | 4 | 6686353 | 6991557 | -1 | Mm.435828 Mm.488241 |
| 6.13 | 5.63 | 6.84 | 6.08 | DN3ab | DN3cd | -0.63 | -7.9 | 2.70E-05 | 0.0021 | 0.073 | ENSMUSG00000051510 | protein_coding | Mafg | v-maf musculoaponeurotic fibrosarcoma oncogene family, protein G (avian) | 17134 | 11 | 120625117 | 120633600 | -1 | Mm.268010 Mm.292957 Mm.393664 |
| 5.19 | 4.59 | 6.05 | 5.32 | DN3ab | DN3cd | -0.67 | -7.9 | 2.70E-05 | 0.0021 | 0.44 | ENSMUSG00000043008 | protein_coding | Klhl6 | kelch-like 6 | 239743 | 16 | 19946496 | 19983037 | -1 | Mm.86699 |
| 11.24 | 11.63 | 11.16 | 11.47 | DN3ab | DN3cd | 0.35 | 7.89 | 2.70E-05 | 0.0021 | 0.47 | ENSMUSG00000064358 | protein_coding | mt-Co3 | mitochondrially encoded cytochrome c oxidase III | 17710 | MT | 8607 | 9390 | 1 | |
| 7.7 | 6.9 | 8.05 | 7.45 | DN3ab | DN3cd | -0.7 | -7.88 | 2.70E-05 | 0.0022 | 0.25 | ENSMUSG00000038366 | protein_coding | Lasp1 | LIM and SH3 protein 1 | 16796 | 11 | 97799000 | 97838764 | 1 | Mm.271967 Mm.470698 |
| 8.76 | 9.39 | 7.94 | 8.9 | DN3ab | DN3cd | 0.76 | 7.87 | 2.70E-05 | 0.0022 | 0.098 | ENSMUSG00000046711 | protein_coding | Hmga1 | high mobility group AT-hook 1 | 15361 | 17 | 27556497 | 27563674 | 1 | Mm.4438 Mm.426212 Mm.440289 |
| 5.66 | 4.74 | 6.6 | 5.86 | DN3ab | DN3cd | -0.81 | -7.87 | 2.70E-05 | 0.0022 | 0.37 | ENSMUSG00000051735 | protein_coding | Rinl | Ras and Rab interactor-like | 320435 | 7 | 28788969 | 28798963 | 1 | Mm.74632 |
| 7.2 | 7.93 | 6.71 | 7.24 | DN3ab | DN3cd | 0.65 | 7.86 | 2.80E-05 | 0.0022 | 0.24 | ENSMUSG00000047832 | protein_coding | Cdca4 | cell division cycle associated 4 | 71963 | 12 | 112820229 | 112829423 | -1 | Mm.313185 |
| 1.84 | -0.93 | 1.55 | -2.11 | DN3ab | DN3cd | -3.14 | -7.86 | 2.80E-05 | 0.0022 | 0.26 | ENSMUSG00000082328 | processed_pseudogene | Gm13921 | predicted gene 13921 | NA | 2 | 102174290 | 102174752 | 1 | |
| 8.01 | 8.59 | 8.03 | 8.62 | DN3ab | DN3cd | 0.59 | 7.84 | 2.80E-05 | 0.0022 | 0.94 | ENSMUSG00000060019 | protein_coding | Gm10073 | predicted pseudogene 10073 | NA | 8 | 106572966 | 106573461 | -1 | |
| 6.39 | 5.01 | 7.43 | 6.55 | DN3ab | DN3cd | -1.09 | -7.84 | 2.80E-05 | 0.0022 | 0.061 | ENSMUSG00000031129 | protein_coding | Slc9a9 | solute carrier family 9 (sodium/hydrogen exchanger), member 9 | 331004 | 9 | 94669909 | 95230445 | 1 | Mm.326125 |
| 9.08 | 8.57 | 9.12 | 8.43 | DN3ab | DN3cd | -0.6 | -7.83 | 2.90E-05 | 0.0022 | 0.24 | ENSMUSG00000040699 | protein_coding | Limd2 | LIM domain containing 2 | 67803 | 11 | 106156256 | 106160860 | -1 | Mm.21687 |
| 6.67 | 5.98 | 7.19 | 6.46 | DN3ab | DN3cd | -0.71 | -7.81 | 2.90E-05 | 0.0022 | 0.82 | ENSMUSG00000075284 | protein_coding | Wipf1 | WAS/WASL interacting protein family, member 1 | 215280 | 2 | 73429610 | 73529734 | -1 | Mm.223504 |
| 2.86 | 3.91 | 2.27 | 3.68 | DN3ab | DN3cd | 1.19 | 7.81 | 2.90E-05 | 0.0022 | 0.23 | ENSMUSG00000025507 | protein_coding | Pidd1 | p53 induced death domain protein 1 | 57913 | 7 | 141438113 | 141444025 | -1 | Mm.334321 |
| 4.67 | 3.96 | 5.75 | 4.77 | DN3ab | DN3cd | -0.86 | -7.81 | 2.90E-05 | 0.0022 | 0.23 | ENSMUSG00000047446 | protein_coding | Arl4a | ADP-ribosylation factor-like 4A | 11861 | 12 | 40005447 | 40038025 | -1 | Mm.12723 |
| 8.04 | 8.87 | 7.84 | 9.14 | DN3ab | DN3cd | 1.05 | 7.81 | 2.90E-05 | 0.0022 | 0.074 | ENSMUSG00000032094 | protein_coding | Cd3d | CD3 antigen, delta polypeptide | 12500 | 9 | 44981786 | 44987339 | 1 | Mm.4527 |
| 7.16 | 6.59 | 7.54 | 7.03 | DN3ab | DN3cd | -0.54 | -7.75 | 3.10E-05 | 0.0023 | 0.65 | ENSMUSG00000037926 | protein_coding | Ssh2 | slingshot protein phosphatase 2 | 237860 | 11 | 77216287 | 77460220 | 1 | Mm.440381 |
| 5.74 | 7.16 | 3.81 | 6.14 | DN3ab | DN3cd | 1.68 | 7.73 | 3.20E-05 | 0.0024 | 0.049 | ENSMUSG00000062380 | protein_coding | Tubb3 | tubulin, beta 3 class III | 22152 | 8 | 123411424 | 123422015 | 1 | Mm.40068 |
| 5.17 | 4.26 | 5.49 | 4.6 | DN3ab | DN3cd | -0.9 | -7.71 | 3.20E-05 | 0.0024 | 0.95 | ENSMUSG00000031488 | protein_coding | Rab11fip1 | RAB11 family interacting protein 1 (class I) | 75767 | 8 | 27138773 | 27174646 | -1 | Mm.205335 |
| 5.92 | 5.03 | 6.55 | 5.73 | DN3ab | DN3cd | -0.86 | -7.7 | 3.20E-05 | 0.0024 | 0.77 | ENSMUSG00000024197 | protein_coding | Plin3 | perilipin 3 | 66905 | 17 | 56277476 | 56292873 | -1 | Mm.311696 |
| 5.09 | 4.45 | 6.03 | 5.15 | DN3ab | DN3cd | -0.77 | -7.7 | 3.30E-05 | 0.0024 | 0.22 | ENSMUSG00000040433 | protein_coding | Zbtb38 | zinc finger and BTB domain containing 38 | 245007 | 9 | 96682770 | 96752831 | -1 | Mm.485954 |
| -1.02 | -3.92 | -0.58 | -4.02 | DN3ab | DN3cd | -3.15 | -7.69 | 3.30E-05 | 0.0024 | 0.5 | ENSMUSG00000112411 | TEC | Gm19337 | predicted gene, 19337 | NA | 10 | 63250932 | 63252515 | 1 | |
| 3.54 | 2.51 | 4.55 | 3.32 | DN3ab | DN3cd | -1.15 | -7.67 | 3.40E-05 | 0.0025 | 0.52 | ENSMUSG00000040813 | protein_coding | Tex264 | testis expressed gene 264 | 21767 | 9 | 106658746 | 106685927 | -1 | Mm.22266 |
| 6.2 | 6.83 | 5.27 | 6.23 | DN3ab | DN3cd | 0.76 | 7.65 | 3.40E-05 | 0.0025 | 0.082 | ENSMUSG00000062075 | protein_coding | Lmnb2 | lamin B2 | 16907 | 10 | 80901203 | 80918245 | -1 | Mm.7362 |
| 3.87 | 2.4 | 5.22 | 4.18 | DN3ab | DN3cd | -1.18 | -7.65 | 3.40E-05 | 0.0025 | 0.19 | ENSMUSG00000050212 | protein_coding | Eva1b | eva-1 homolog B (C. elegans) | 230752 | 4 | 126147744 | 126149875 | 1 | Mm.41569 |
| 4.97 | 4.03 | 7.57 | 6.05 | DN3ab | DN3cd | -1.34 | -7.65 | 3.40E-05 | 0.0025 | 0.12 | ENSMUSG00000026879 | protein_coding | Gsn | gelsolin | 227753 | 2 | 35256380 | 35307892 | 1 | Mm.443 Mm.21109 |
| 5.79 | 5.27 | 6.47 | 5.68 | DN3ab | DN3cd | -0.66 | -7.65 | 3.40E-05 | 0.0025 | 0.097 | ENSMUSG00000023277 | protein_coding | Twf2 | twinfilin actin binding protein 2 | 23999 | 9 | 106203108 | 106215389 | 1 | Mm.274346 |
| 5.12 | 5.94 | 4.23 | 5.39 | DN3ab | DN3cd | 0.95 | 7.63 | 3.50E-05 | 0.0025 | 0.19 | ENSMUSG00000058773 | protein_coding | Hist1h1b | histone cluster 1, H1b | 56702 | 13 | 21779883 | 21780625 | -1 | Mm.221314 |
| 4.65 | 3.52 | 5.99 | 4.41 | DN3ab | DN3cd | -1.4 | -7.62 | 3.50E-05 | 0.0025 | 0.24 | ENSMUSG00000046876 | protein_coding | Atxn1 | ataxin 1 | 20238 | 13 | 45549755 | 45965008 | -1 | Mm.342683 Mm.342686 Mm.486641 Mm.487047 |
| 5.74 | 6.34 | 4.95 | 5.76 | DN3ab | DN3cd | 0.68 | 7.62 | 3.50E-05 | 0.0025 | 0.21 | ENSMUSG00000017550 | protein_coding | Atad5 | ATPase family, AAA domain containing 5 | 237877 | 11 | 80089400 | 80135794 | 1 | Mm.89551 |
| 0.79 | 3.31 | -0.03 | 2.69 | DN3ab | DN3cd | 2.59 | 7.6 | 3.60E-05 | 0.0026 | 0.79 | ENSMUSG00000034485 | protein_coding | Uaca | uveal autoantigen with coiled-coil domains and ankyrin repeats | 72565 | 9 | 60794542 | 60880370 | 1 | Mm.68819 |
| 5.48 | 4.6 | 6.4 | 4.9 | DN3ab | DN3cd | -1.2 | -7.6 | 3.60E-05 | 0.0026 | 0.034 | ENSMUSG00000026426 | protein_coding | Arl8a | ADP-ribosylation factor-like 8A | 68724 | 1 | 135146824 | 135156269 | 1 | Mm.174998 |
| 5.3 | 5.93 | 5.52 | 6.4 | DN3ab | DN3cd | 0.75 | 7.6 | 3.60E-05 | 0.0026 | 0.18 | ENSMUSG00000027381 | protein_coding | Bcl2l11 | BCL2-like 11 (apoptosis facilitator) | 12125 | 2 | 128126038 | 128162547 | 1 | Mm.141083 Mm.453214 |
| 8.52 | 8.22 | 8.54 | 8.1 | DN3ab | DN3cd | -0.36 | -7.59 | 3.60E-05 | 0.0026 | 0.11 | ENSMUSG00000000346 | protein_coding | Dazap2 | DAZ associated protein 2 | 23994 | 15 | 100615349 | 100620761 | 1 | Mm.486175 |
| 5.61 | 4.62 | 6.62 | 5.62 | DN3ab | DN3cd | -0.99 | -7.57 | 3.70E-05 | 0.0026 | 0.98 | ENSMUSG00000040111 | protein_coding | Gramd1b | GRAM domain containing 1B | 235283 | 9 | 40293233 | 40531383 | -1 | Mm.138434 |
| 4.38 | 3.49 | 4.99 | 4.2 | DN3ab | DN3cd | -0.83 | -7.57 | 3.70E-05 | 0.0026 | 0.65 | ENSMUSG00000019842 | protein_coding | Traf3ip2 | TRAF3 interacting protein 2 | 103213 | 10 | 39612934 | 39655307 | 1 | Mm.436686 |
| 5.85 | 5.23 | 6.55 | 5.85 | DN3ab | DN3cd | -0.66 | -7.56 | 3.70E-05 | 0.0026 | 0.65 | ENSMUSG00000024772 | protein_coding | Ehd1 | EH-domain containing 1 | 13660 | 19 | 6276725 | 6300096 | 1 | Mm.30169 |
| 2.38 | 4.5 | 1.42 | 4.27 | DN3ab | DN3cd | 2.36 | 7.56 | 3.80E-05 | 0.0026 | 0.28 | ENSMUSG00000021779 | protein_coding | Thrb | thyroid hormone receptor beta | 21834 | 14 | 17660261 | 18038090 | 1 | Mm.32563 Mm.491616 |
| 5.99 | 5.49 | 6.85 | 6 | DN3ab | DN3cd | -0.69 | -7.56 | 3.80E-05 | 0.0026 | 0.03 | ENSMUSG00000022528 | protein_coding | Hes1 | hes family bHLH transcription factor 1 | 15205 | 16 | 30064384 | 30067796 | 1 | Mm.390859 |
| 6.98 | 6.47 | 7.59 | 6.86 | DN3ab | DN3cd | -0.62 | -7.55 | 3.80E-05 | 0.0026 | 0.16 | ENSMUSG00000031950 | protein_coding | Gabarapl2 | gamma-aminobutyric acid (GABA) A receptor-associated protein-like 2 | 93739 | 8 | 111940703 | 111953612 | 1 | Mm.371666 |
| 4.6 | 3.21 | 5.67 | 4.7 | DN3ab | DN3cd | -1.13 | -7.53 | 3.90E-05 | 0.0026 | 0.17 | ENSMUSG00000022014 | protein_coding | Epsti1 | epithelial stromal interaction 1 (breast) | 108670 | 14 | 77904239 | 78002657 | 1 | Mm.68134 |
| 8.7 | 9.34 | 7.92 | 8.84 | DN3ab | DN3cd | 0.75 | 7.53 | 3.90E-05 | 0.0026 | 0.18 | ENSMUSG00000078249 | protein_coding | Hmga1b | high mobility group AT-hook 1B | 111241 | 11 | 120762794 | 120764419 | 1 | Mm.426212 |
| 4.9 | 5.88 | 4.4 | 5.87 | DN3ab | DN3cd | 1.18 | 7.52 | 3.90E-05 | 0.0026 | 0.12 | ENSMUSG00000020393 | protein_coding | Kremen1 | kringle containing transmembrane protein 1 | 84035 | 11 | 5191552 | 5261558 | -1 | Mm.209989 |
| 5.67 | 4.91 | 6.14 | 5.29 | DN3ab | DN3cd | -0.8 | -7.51 | 3.90E-05 | 0.0027 | 0.7 | ENSMUSG00000041035 | protein_coding | Gm17018 | predicted gene 17018 | 226162 | 19 | 45560615 | 45579763 | 1 | Mm.21978 |
| 6.16 | 5.65 | 7.34 | 6.46 | DN3ab | DN3cd | -0.71 | -7.5 | 4.00E-05 | 0.0027 | 0.035 | ENSMUSG00000053819 | protein_coding | Camk2d | calcium/calmodulin-dependent protein kinase II, delta | 108058 | 3 | 126596302 | 126846326 | 1 | Mm.255822 Mm.442379 |
| 7.1 | 7.57 | 6.31 | 6.96 | DN3ab | DN3cd | 0.54 | 7.5 | 4.00E-05 | 0.0027 | 0.21 | ENSMUSG00000006715 | protein_coding | Gmnn | geminin | 57441 | 13 | 24751845 | 24761923 | -1 | Mm.12239 |
| 3.72 | 2.72 | 3.71 | 2.48 | DN3ab | DN3cd | -1.1 | -7.49 | 4.10E-05 | 0.0027 | 0.44 | ENSMUSG00000045410 | protein_coding | Akr1e1 | aldo-keto reductase family 1, member E1 | 56043 | 13 | 4590750 | 4609174 | -1 | Mm.251908 |
| 5.02 | 6.2 | 4.55 | 5.73 | DN3ab | DN3cd | 1.18 | 7.48 | 4.10E-05 | 0.0027 | 0.99 | ENSMUSG00000020492 | protein_coding | Ska2 | spindle and kinetochore associated complex subunit 2 | 66140 | 11 | 87109234 | 87124635 | 1 | Mm.45008 |
| 2.15 | 3.62 | 2.1 | 3.48 | DN3ab | DN3cd | 1.43 | 7.48 | 4.10E-05 | 0.0027 | 0.82 | ENSMUSG00000074682 | protein_coding | Zcchc3 | zinc finger, CCHC domain containing 3 | 67917 | 2 | 152411955 | 152415027 | -1 | Mm.391810 |
| 6.64 | 7.3 | 6.12 | 6.61 | DN3ab | DN3cd | 0.59 | 7.48 | 4.10E-05 | 0.0027 | 0.28 | ENSMUSG00000017499 | protein_coding | Cdc6 | cell division cycle 6 | 23834 | 11 | 98907801 | 98923940 | 1 | Mm.20912 |
| -4.01 | -0.2 | -3.72 | -0.61 | DN3ab | DN3cd | 3.49 | 7.48 | 4.10E-05 | 0.0027 | 0.45 | ENSMUSG00000073402 | protein_coding | Gm8909 | predicted gene 8909 | 667977 | 17 | 36164443 | 36168537 | -1 | Mm.441651 Mm.489198 |
| 2.53 | 1.14 | 3.2 | 1.56 | DN3ab | DN3cd | -1.52 | -7.47 | 4.10E-05 | 0.0027 | 0.53 | ENSMUSG00000040451 | protein_coding | Sgms1 | sphingomyelin synthase 1 | 208449 | 19 | 32122727 | 32389714 | -1 | Mm.329810 |
| 3.32 | 2.12 | 4.92 | 3.11 | DN3ab | DN3cd | -1.59 | -7.46 | 4.20E-05 | 0.0027 | 0.16 | ENSMUSG00000054752 | protein_coding | Fsd1l | fibronectin type III and SPRY domain containing 1-like | 319636 | 4 | 53631471 | 53707009 | 1 | Mm.103464 Mm.487999 |
| 3.15 | 4.64 | 2.76 | 3.97 | DN3ab | DN3cd | 1.39 | 7.45 | 4.20E-05 | 0.0027 | 0.47 | ENSMUSG00000020185 | protein_coding | E2f7 | E2F transcription factor 7 | 52679 | 10 | 110745439 | 110787384 | 1 | Mm.11747 |
| 5.85 | 6.59 | 5.52 | 6.09 | DN3ab | DN3cd | 0.67 | 7.44 | 4.20E-05 | 0.0027 | 0.36 | ENSMUSG00000020471 | protein_coding | Pold2 | polymerase (DNA directed), delta 2, regulatory subunit | 18972 | 11 | 5872180 | 5878292 | -1 | Mm.35788 |
| 3.99 | 3.1 | 5.23 | 3.92 | DN3ab | DN3cd | -1.14 | -7.44 | 4.30E-05 | 0.0027 | 0.18 | ENSMUSG00000024457 | protein_coding | Trim26 | tripartite motif-containing 26 | 22670 | 17 | 36837134 | 36859398 | 1 | Mm.34587 |
| 7.39 | 7.03 | 8.08 | 7.72 | DN3ab | DN3cd | -0.36 | -7.43 | 4.30E-05 | 0.0027 | 1 | ENSMUSG00000032504 | protein_coding | Pdcd6ip | programmed cell death 6 interacting protein | 18571 | 9 | 113651744 | 113708259 | -1 | Mm.29816 |
| 6.19 | 5.33 | 7.22 | 6.43 | DN3ab | DN3cd | -0.82 | -7.42 | 4.30E-05 | 0.0028 | 0.75 | ENSMUSG00000066621 | protein_coding | Tecpr1 | tectonin beta-propeller repeat containing 1 | 70381 | 5 | 144194442 | 144223615 | -1 | Mm.239412 Mm.458463 |
| 6.68 | 6.21 | 6.87 | 6.19 | DN3ab | DN3cd | -0.57 | -7.39 | 4.50E-05 | 0.0028 | 0.16 | ENSMUSG00000029106 | protein_coding | Add1 | adducin 1 (alpha) | 11518 | 5 | 34573664 | 34632308 | 1 | Mm.289106 |
| 6 | 5.22 | 5.94 | 5.38 | DN3ab | DN3cd | -0.67 | -7.38 | 4.50E-05 | 0.0028 | 0.22 | ENSMUSG00000028995 | protein_coding | Fam126a | family with sequence similarity 126, member A | 84652 | 5 | 23915276 | 24030690 | -1 | Mm.304976 Mm.486278 |
| 3.07 | 0.06 | 2.43 | -0.34 | DN3ab | DN3cd | -2.91 | -7.35 | 4.70E-05 | 0.0029 | 0.78 | ENSMUSG00000106219 | lincRNA | 5830416I19Rik | RIKEN cDNA 5830416I19 gene | NA | 5 | 64044978 | 64050228 | 1 | |
| 7.66 | 8.05 | 7.19 | 7.62 | DN3ab | DN3cd | 0.41 | 7.34 | 4.70E-05 | 0.0029 | 0.74 | ENSMUSG00000004642 | protein_coding | Slbp | stem-loop binding protein | 20492 | 5 | 33634952 | 33652574 | -1 | Mm.4172 |
| 7.34 | 6.83 | 8.42 | 7.92 | DN3ab | DN3cd | -0.5 | -7.34 | 4.70E-05 | 0.0029 | 0.99 | ENSMUSG00000007891 | protein_coding | Ctsd | cathepsin D | 13033 | 7 | 142375911 | 142388038 | -1 | Mm.231395 |
| 4.21 | 2.96 | 5.08 | 3.87 | DN3ab | DN3cd | -1.22 | -7.34 | 4.70E-05 | 0.0029 | 0.89 | ENSMUSG00000045362 | protein_coding | Tnfrsf26 | tumor necrosis factor receptor superfamily, member 26 | 244237 | 7 | 143607659 | 143628722 | -1 | Mm.247498 |
| 6.85 | 5.92 | 7.95 | 6.97 | DN3ab | DN3cd | -0.96 | -7.33 | 4.80E-05 | 0.0029 | 0.85 | ENSMUSG00000021065 | protein_coding | Fut8 | fucosyltransferase 8 | 53618 | 12 | 77238125 | 77476338 | 1 | Mm.35628 Mm.489686 |
| 7.11 | 7.71 | 6.15 | 6.88 | DN3ab | DN3cd | 0.64 | 7.31 | 4.80E-05 | 0.003 | 0.48 | ENSMUSG00000037313 | protein_coding | Tacc3 | transforming, acidic coiled-coil containing protein 3 | 21335 | 5 | 33658128 | 33678995 | 1 | Mm.379024 |
| 2.02 | 4.09 | 1.76 | 3.34 | DN3ab | DN3cd | 1.87 | 7.3 | 4.90E-05 | 0.003 | 0.37 | ENSMUSG00000100967 | protein_coding | Gm29666 | predicted gene 29666 | NA | 15 | 84910438 | 84920038 | -1 | |
| 5.82 | 5.18 | 6.73 | 5.76 | DN3ab | DN3cd | -0.81 | -7.29 | 4.90E-05 | 0.003 | 0.12 | ENSMUSG00000040710 | protein_coding | St8sia4 | ST8 alpha-N-acetyl-neuraminide alpha-2,8-sialyltransferase 4 | 20452 | 1 | 95587682 | 95667571 | -1 | Mm.306228 |
| 5.32 | 4.4 | 6.15 | 4.63 | DN3ab | DN3cd | -1.23 | -7.29 | 5.00E-05 | 0.003 | 0.062 | ENSMUSG00000038679 | protein_coding | Trps1 | transcriptional repressor GATA binding 1 | 83925 | 15 | 50654752 | 50890463 | -1 | Mm.30466 |
| 6.5 | 7.06 | 5.53 | 6.08 | DN3ab | DN3cd | 0.56 | 7.29 | 5.00E-05 | 0.003 | 0.98 | ENSMUSG00000028066 | protein_coding | Pmf1 | polyamine-modulated factor 1 | 67037 | 3 | 88394143 | 88410331 | -1 | Mm.36715 |
| 6.17 | 6.7 | 5.51 | 6.15 | DN3ab | DN3cd | 0.57 | 7.29 | 5.00E-05 | 0.003 | 0.5 | ENSMUSG00000002835 | protein_coding | Chaf1a | chromatin assembly factor 1, subunit A (p150) | 27221 | 17 | 56040439 | 56072289 | 1 | Mm.391010 |
| 5.34 | 6.61 | 5.34 | 6.13 | DN3ab | DN3cd | 1.06 | 7.28 | 5.00E-05 | 0.003 | 0.087 | ENSMUSG00000027994 | protein_coding | Mcub | mitochondrial calcium uniporter dominant negative beta subunit | 66815 | 3 | 129914960 | 129970206 | -1 | Mm.31056 |
| 6.85 | 7.28 | 6.02 | 6.56 | DN3ab | DN3cd | 0.47 | 7.27 | 5.10E-05 | 0.003 | 0.37 | ENSMUSG00000024735 | protein_coding | Prpf19 | pre-mRNA processing factor 19 | 28000 | 19 | 10895231 | 10909559 | 1 | Mm.491104 |
| 5.32 | 5.96 | 5.31 | 6.06 | DN3ab | DN3cd | 0.69 | 7.26 | 5.10E-05 | 0.0031 | 0.6 | ENSMUSG00000046668 | protein_coding | Cxxc5 | CXXC finger 5 | 67393 | 18 | 35829818 | 35861687 | 1 | Mm.94560 |
| -4.01 | -0.7 | -3.72 | 0.9 | DN3ab | DN3cd | 4.03 | 7.25 | 5.20E-05 | 0.0031 | 0.24 | ENSMUSG00000111169 | lincRNA | Gm34004 | predicted gene, 34004 | NA | 9 | 61081951 | 61117839 | 1 | |
| 6.04 | 6.63 | 5.43 | 5.97 | DN3ab | DN3cd | 0.57 | 7.21 | 5.40E-05 | 0.0032 | 0.74 | ENSMUSG00000000028 | protein_coding | Cdc45 | cell division cycle 45 | 12544 | 16 | 18780447 | 18811987 | -1 | Mm.1248 |
| 5.77 | 5.19 | 5.89 | 5.3 | DN3ab | DN3cd | -0.59 | -7.2 | 5.50E-05 | 0.0032 | 0.95 | ENSMUSG00000021819 | protein_coding | Zswim8 | zinc finger SWIM-type containing 8 | 268721 | 14 | 20707552 | 20723619 | 1 | Mm.275082 |
| 5.84 | 5.08 | 7.04 | 5.76 | DN3ab | DN3cd | -1.05 | -7.2 | 5.50E-05 | 0.0032 | 0.059 | ENSMUSG00000025557 | protein_coding | Slc15a1 | solute carrier family 15 (oligopeptide transporter), member 1 | 56643 | 14 | 121459621 | 121505252 | -1 | Mm.155618 |
| 8.46 | 7.96 | 8.95 | 8.59 | DN3ab | DN3cd | -0.43 | -7.19 | 5.50E-05 | 0.0032 | 0.26 | ENSMUSG00000022108 | protein_coding | Itm2b | integral membrane protein 2B | 16432 | 14 | 73362226 | 73385289 | -1 | Mm.4266 Mm.392973 |
| 7.31 | 6.73 | 7.69 | 7.16 | DN3ab | DN3cd | -0.55 | -7.18 | 5.60E-05 | 0.0032 | 0.77 | ENSMUSG00000037822 | protein_coding | Smim14 | small integral membrane protein 14 | 68552 | 5 | 65446844 | 65537184 | -1 | Mm.10709 Mm.393232 |
| 3.99 | 3.19 | 5.09 | 4.07 | DN3ab | DN3cd | -0.93 | -7.15 | 5.80E-05 | 0.0033 | 0.41 | ENSMUSG00000040435 | protein_coding | Ppp1r15a | protein phosphatase 1, regulatory subunit 15A | 17872 | 7 | 45522916 | 45526268 | -1 | Mm.4048 |
| 6.49 | 5.75 | 7.67 | 6.54 | DN3ab | DN3cd | -0.96 | -7.14 | 5.80E-05 | 0.0033 | 0.14 | ENSMUSG00000032712 | protein_coding | Resf1 | retroelement silencing factor 1 | 67246 | 6 | 149309414 | 149335663 | 1 | Mm.333515 |
| 6.24 | 5.76 | 6.88 | 6.21 | DN3ab | DN3cd | -0.58 | -7.13 | 5.90E-05 | 0.0034 | 0.27 | ENSMUSG00000032328 | protein_coding | Tmem30a | transmembrane protein 30A | 69981 | 9 | 79768943 | 79793507 | -1 | Mm.486324 |
| 8.34 | 7.94 | 8.94 | 8.64 | DN3ab | DN3cd | -0.35 | -7.12 | 5.90E-05 | 0.0034 | 0.33 | ENSMUSG00000028822 | protein_coding | Tmem50a | transmembrane protein 50A | 71817 | 4 | 134897849 | 134915024 | -1 | Mm.88349 |
| 7.1 | 7.61 | 6.46 | 7.1 | DN3ab | DN3cd | 0.56 | 7.12 | 5.90E-05 | 0.0034 | 0.42 | ENSMUSG00000032477 | protein_coding | Cdc25a | cell division cycle 25A | 12530 | 9 | 109875579 | 109893895 | 1 | Mm.307103 |
| 4.68 | 5.33 | 4.04 | 4.9 | DN3ab | DN3cd | 0.73 | 7.12 | 6.00E-05 | 0.0034 | 0.3 | ENSMUSG00000021714 | protein_coding | Cenpk | centromere protein K | 60411 | 13 | 104228611 | 104252392 | 1 | Mm.281498 |
| 3.44 | 2.18 | 4.18 | 3.21 | DN3ab | DN3cd | -1.09 | -7.1 | 6.10E-05 | 0.0034 | 0.36 | ENSMUSG00000059708 | protein_coding | Akap17b | A kinase (PRKA) anchor protein 17B | 338351 | X | 36608314 | 36645395 | -1 | Mm.116704 |
| 5.28 | 4.66 | 6.31 | 5.66 | DN3ab | DN3cd | -0.63 | -7.1 | 6.10E-05 | 0.0034 | 0.86 | ENSMUSG00000001248 | protein_coding | Gramd1a | GRAM domain containing 1A | 52857 | 7 | 31130127 | 31155896 | -1 | Mm.27366 |
| 3.69 | 7.5 | 0.69 | 7.16 | DN3ab | DN3cd | 4.15 | 7.09 | 6.10E-05 | 0.0034 | 0.13 | ENSMUSG00000032092 | protein_coding | Mpzl2 | myelin protein zero-like 2 | 14012 | 9 | 45039799 | 45054015 | 1 | Mm.33240 |
| 7.1 | 7.52 | 6.33 | 6.88 | DN3ab | DN3cd | 0.47 | 7.09 | 6.10E-05 | 0.0034 | 0.33 | ENSMUSG00000023015 | protein_coding | Racgap1 | Rac GTPase-activating protein 1 | 26934 | 15 | 99620496 | 99651656 | -1 | Mm.273804 |
| 3.49 | 2.42 | 3.84 | 2.87 | DN3ab | DN3cd | -1.02 | -7.09 | 6.10E-05 | 0.0034 | 0.71 | ENSMUSG00000021867 | protein_coding | Tmem254b | transmembrane protein 254b | 100039257 | 14 | 26203369 | 26207041 | -1 | Mm.33344 |
| 6.15 | 6.69 | 5.67 | 6.26 | DN3ab | DN3cd | 0.56 | 7.09 | 6.20E-05 | 0.0034 | 0.8 | ENSMUSG00000022718 | protein_coding | Dgcr8 | DGCR8, microprocessor complex subunit | 94223 | 16 | 18253948 | 18289246 | -1 | Mm.447352 Mm.482156 |
| 9.58 | 10.26 | 8.76 | 9.74 | DN3ab | DN3cd | 0.8 | 7.08 | 6.20E-05 | 0.0034 | 0.2 | ENSMUSG00000040204 | protein_coding | Pclaf | PCNA clamp associated factor | 68026 | 9 | 65890237 | 65903266 | 1 | Mm.351273 Mm.460895 |
| 3.93 | 2.76 | 4.36 | 3.58 | DN3ab | DN3cd | -0.95 | -7.08 | 6.20E-05 | 0.0034 | 0.13 | ENSMUSG00000027605 | protein_coding | Acss2 | acyl-CoA synthetase short-chain family member 2 | 60525 | 2 | 155517948 | 155585724 | 1 | Mm.255026 |
| 5.19 | 4.45 | 5.93 | 5.4 | DN3ab | DN3cd | -0.63 | -7.07 | 6.30E-05 | 0.0034 | 0.23 | ENSMUSG00000030560 | protein_coding | Ctsc | cathepsin C | 13032 | 7 | 88278085 | 88310888 | 1 | Mm.89295 Mm.322945 Mm.483300 |
| -4.01 | -0.93 | -3.72 | -0.04 | DN3ab | DN3cd | 3.39 | 7.07 | 6.30E-05 | 0.0034 | 0.54 | ENSMUSG00000032511 | protein_coding | Scn5a | sodium channel, voltage-gated, type V, alpha | 20271 | 9 | 119483408 | 119579016 | -1 | Mm.103584 |
| 7.19 | 7.77 | 6.65 | 7.15 | DN3ab | DN3cd | 0.54 | 7.07 | 6.30E-05 | 0.0034 | 0.61 | ENSMUSG00000015176 | protein_coding | Nolc1 | nucleolar and coiled-body phosphoprotein 1 | 70769 | 19 | 46075863 | 46085530 | -1 | Mm.402190 Mm.486316 |
| 4.87 | 5.64 | 4.21 | 5.08 | DN3ab | DN3cd | 0.81 | 7.07 | 6.30E-05 | 0.0034 | 0.66 | ENSMUSG00000049577 | protein_coding | Zfpm1 | zinc finger protein, multitype 1 | 22761 | 8 | 122282141 | 122337251 | 1 | Mm.3105 Mm.390068 |
| 3.44 | 4.91 | 3.99 | 5.08 | DN3ab | DN3cd | 1.27 | 7.05 | 6.50E-05 | 0.0035 | 0.31 | ENSMUSG00000027333 | protein_coding | Smox | spermine oxidase | 228608 | 2 | 131491496 | 131525922 | 1 | Mm.136586 |
| 7.74 | 7.4 | 8.13 | 7.54 | DN3ab | DN3cd | -0.46 | -7.02 | 6.70E-05 | 0.0035 | 0.031 | ENSMUSG00000059923 | protein_coding | Grb2 | growth factor receptor bound protein 2 | 14784 | 11 | 115644045 | 115708597 | -1 | Mm.439649 Mm.490413 |
| 6.6 | 6.11 | 7.51 | 6.66 | DN3ab | DN3cd | -0.68 | -7 | 6.80E-05 | 0.0036 | 0.06 | ENSMUSG00000037012 | protein_coding | Hk1 | hexokinase 1 | 15275 | 10 | 62268855 | 62379908 | -1 | Mm.196605 |
| 3.83 | 2.87 | 4.08 | 3.3 | DN3ab | DN3cd | -0.87 | -7 | 6.80E-05 | 0.0036 | 0.49 | ENSMUSG00000020654 | protein_coding | Adcy3 | adenylate cyclase 3 | 104111 | 12 | 4133103 | 4213525 | 1 | Mm.489807 |
| 6.69 | 5.94 | 7.51 | 6.41 | DN3ab | DN3cd | -0.93 | -6.98 | 7.00E-05 | 0.0036 | 0.2 | ENSMUSG00000040212 | protein_coding | Emp3 | epithelial membrane protein 3 | 13732 | 7 | 45918023 | 45921404 | -1 | Mm.20829 |
| 9.61 | 10.14 | 8.66 | 9.12 | DN3ab | DN3cd | 0.5 | 6.97 | 7.00E-05 | 0.0037 | 0.65 | ENSMUSG00000027203 | protein_coding | Dut | deoxyuridine triphosphatase | 110074 | 2 | 125247190 | 125258608 | 1 | Mm.282499 Mm.471988 |
| 2.2 | 3.53 | 1.49 | 3.45 | DN3ab | DN3cd | 1.56 | 6.95 | 7.20E-05 | 0.0037 | 0.17 | ENSMUSG00000049539 | protein_coding | Hist1h1a | histone cluster 1, H1a | 80838 | 13 | 23763666 | 23764406 | 1 | Mm.480242 |
| 4.86 | 5.46 | 4.17 | 4.97 | DN3ab | DN3cd | 0.68 | 6.95 | 7.20E-05 | 0.0037 | 0.28 | ENSMUSG00000024791 | protein_coding | Cdca5 | cell division cycle associated 5 | 67849 | 19 | 6084983 | 6091777 | 1 | Mm.23526 |
| 1.34 | -0.24 | 3.07 | 0.19 | DN3ab | DN3cd | -2.32 | -6.95 | 7.20E-05 | 0.0037 | 0.031 | ENSMUSG00000000627 | protein_coding | Sema4f | sema domain, immunoglobulin domain (Ig), TM domain, and short cytoplasmic domain | 20355 | 6 | 82911885 | 82939769 | -1 | Mm.270543 |
| 9.52 | 9.19 | 10.34 | 9.79 | DN3ab | DN3cd | -0.44 | -6.94 | 7.20E-05 | 0.0037 | 0.077 | ENSMUSG00000029622 | protein_coding | Arpc1b | actin related protein 2/3 complex, subunit 1B | 11867 | 5 | 145114215 | 145130705 | 1 | Mm.30010 |
| 5.06 | 4.23 | 5.63 | 4.72 | DN3ab | DN3cd | -0.87 | -6.94 | 7.30E-05 | 0.0037 | 0.75 | ENSMUSG00000028559 | protein_coding | Osbpl9 | oxysterol binding protein-like 9 | 100273 | 4 | 109061145 | 109202272 | -1 | Mm.366315 |
| 6.24 | 6.74 | 5.51 | 6.03 | DN3ab | DN3cd | 0.51 | 6.93 | 7.30E-05 | 0.0037 | 0.85 | ENSMUSG00000027018 | protein_coding | Hat1 | histone aminotransferase 1 | 107435 | 2 | 71388958 | 71441622 | 1 | Mm.272472 |
| 5.75 | 5.24 | 5.86 | 5.14 | DN3ab | DN3cd | -0.61 | -6.93 | 7.30E-05 | 0.0037 | 0.23 | ENSMUSG00000032872 | protein_coding | Cyb5r4 | cytochrome b5 reductase 4 | 266690 | 9 | 87022014 | 87077774 | 1 | Mm.30166 Mm.317460 |
| 5.94 | 6.45 | 5.44 | 6.01 | DN3ab | DN3cd | 0.54 | 6.93 | 7.30E-05 | 0.0037 | 0.73 | ENSMUSG00000036097 | protein_coding | Slf2 | SMC5-SMC6 complex localization factor 2 | 226151 | 19 | 44931119 | 44983787 | 1 | Mm.212609 |
| -1.45 | 1.21 | -2.9 | 1.08 | DN3ab | DN3cd | 3.27 | 6.89 | 7.70E-05 | 0.0038 | 0.15 | ENSMUSG00000025227 | protein_coding | Mfsd13a | major facilitator superfamily domain containing 13a | 75146 | 19 | 46341121 | 46375252 | 1 | Mm.159290 |
| 2.91 | 4 | 1.98 | 3.77 | DN3ab | DN3cd | 1.32 | 6.88 | 7.70E-05 | 0.0038 | 0.058 | ENSMUSG00000023938 | protein_coding | Aars2 | alanyl-tRNA synthetase 2, mitochondrial | 224805 | 17 | 45506841 | 45520843 | 1 | Mm.329063 |
| 4.51 | 5.34 | 3.53 | 4.81 | DN3ab | DN3cd | 1 | 6.88 | 7.70E-05 | 0.0038 | 0.13 | ENSMUSG00000029554 | protein_coding | Mad1l1 | MAD1 mitotic arrest deficient 1-like 1 | 17120 | 5 | 140008689 | 140321552 | -1 | Mm.27250 |
| 2.65 | -3.14 | 3.73 | 0.19 | DN3ab | DN3cd | -4.31 | -6.87 | 7.80E-05 | 0.0038 | 0.079 | ENSMUSG00000031397 | protein_coding | Tktl1 | transketolase-like 1 | 83553 | X | 74177259 | 74208500 | 1 | Mm.25057 |
| 2.73 | 3.48 | 2.92 | 4.1 | DN3ab | DN3cd | 0.96 | 6.87 | 7.80E-05 | 0.0038 | 0.099 | ENSMUSG00000064372 | Mt_tRNA | mt-Tp | mitochondrially encoded tRNA proline | NA | MT | 15356 | 15422 | -1 | |
| 4.91 | 5.63 | 3.73 | 4.64 | DN3ab | DN3cd | 0.78 | 6.87 | 7.80E-05 | 0.0038 | 0.46 | ENSMUSG00000032783 | protein_coding | Troap | trophinin associated protein | 78733 | 15 | 99074575 | 99083409 | 1 | Mm.124905 |
| 7.03 | 6.53 | 7.53 | 7.13 | DN3ab | DN3cd | -0.45 | -6.85 | 8.00E-05 | 0.0039 | 0.49 | ENSMUSG00000038608 | protein_coding | Dock10 | dedicator of cytokinesis 10 | 210293 | 1 | 80501073 | 80758527 | -1 | Mm.133473 |
| -0.12 | 2.06 | -1.76 | 1.28 | DN3ab | DN3cd | 2.47 | 6.85 | 8.00E-05 | 0.0039 | 0.24 | ENSMUSG00000111617 | TEC | Gm47148 | predicted gene, 47148 | NA | 9 | 122993127 | 122993704 | 1 | |

| | | | | | | | | | | | | | | | | | | | | |
|---|---|---|---|---|---|---|---|---|---|---|---|---|---|---|---|---|---|---|---|---|
| 6.7 | 6.29 | 6.89 | 6.41 | DN3ab | DN3cd | -0.44 | -6.84 | 8.10E-05 | 0.0039 | 0.54 | ENSMUSG00000033159 | protein_coding | Cnppd1 | cyclin Pas1/PHO80 domain containing 1 | 69171 | 1 | 75134554 | 75142711 | -1 | Mm.28193 |
| 6.4 | 6.81 | 5.62 | 6.19 | DN3ab | DN3cd | 0.48 | 6.84 | 8.10E-05 | 0.0039 | 0.25 | ENSMUSG00000027323 | protein_coding | Rad51 | RAD51 recombinase | 19361 | 2 | 119112793 | 119147445 | 1 | Mm.330492 |
| -0.41 | 2.05 | 0.76 | 2.76 | DN3ab | DN3cd | 2.19 | 6.84 | 8.10E-05 | 0.0039 | 0.48 | ENSMUSG00000097465 | NA | NA | NA | NA | NA | NA | NA | NA | |
| 5.49 | 6.09 | 4.68 | 5.39 | DN3ab | DN3cd | 0.65 | 6.83 | 8.20E-05 | 0.0039 | 0.57 | ENSMUSG00000026683 | protein_coding | Nuf2 | NUF2, NDC80 kinetochore complex component | 66977 | 1 | 169497934 | 169531464 | -1 | Mm.151315 |
| -1.27 | -3.92 | 1.4 | -1.73 | DN3ab | DN3cd | -2.93 | -6.83 | 8.20E-05 | 0.0039 | 0.58 | ENSMUSG00000024209 | protein_coding | 1700061G19Rik | RIKEN cDNA 1700061G19 gene | 78625 | 17 | 56875477 | 56888904 | 1 | Mm.160106 |
| -1.45 | 1.71 | -2.36 | 2.16 | DN3ab | DN3cd | 3.8 | 6.83 | 8.20E-05 | 0.0039 | 0.22 | ENSMUSG00000062785 | protein_coding | Kcnc3 | potassium voltage gated channel, Shaw-related subfamily, member 3 | 16504 | 7 | 44590664 | 44604754 | 1 | Mm.40312 |
| 3.99 | 2.96 | 5.82 | 4.07 | DN3ab | DN3cd | -1.49 | -6.82 | 8.30E-05 | 0.004 | 0.11 | ENSMUSG00000044199 | protein_coding | S1pr4 | sphingosine-1-phosphate receptor 4 | 13611 | 10 | 81497747 | 81500132 | -1 | Mm.33065 |
| 5.34 | 4.7 | 6.59 | 5.71 | DN3ab | DN3cd | -0.78 | -6.81 | 8.30E-05 | 0.004 | 0.34 | ENSMUSG00000018507 | protein_coding | Trpv2 | transient receptor potential cation channel, subfamily V, member 2 | 22368 | 11 | 62574486 | 62600515 | 1 | Mm.288064 |
| 5.53 | 6.08 | 5.11 | 5.6 | DN3ab | DN3cd | 0.53 | 6.8 | 8.50E-05 | 0.004 | 0.73 | ENSMUSG00000030528 | protein_coding | Blm | Bloom syndrome, RecQ like helicase | 12144 | 7 | 80454733 | 80535119 | -1 | Mm.12932 |
| 4.83 | 4.33 | 5.7 | 4.89 | DN3ab | DN3cd | -0.67 | -6.8 | 8.50E-05 | 0.004 | 0.1 | ENSMUSG00000006920 | protein_coding | Ezh1 | enhancer of zeste 1 polycomb repressive complex 2 subunit | 14055 | 11 | 101191115 | 101226463 | -1 | Mm.5027 |
| 6.35 | 6.82 | 5.71 | 6.53 | DN3ab | DN3cd | 0.62 | 6.79 | 8.60E-05 | 0.0041 | 0.038 | ENSMUSG00000040331 | protein_coding | Nsmce4a | NSE4 homolog A, SMC5-SMC6 complex component | 67872 | 7 | 130532526 | 130573118 | -1 | Mm.21714 Mm.411879 |
| 3.85 | 2.43 | 3.77 | 2.9 | DN3ab | DN3cd | -1.14 | -6.78 | 8.70E-05 | 0.0041 | 0.08 | ENSMUSG00000031565 | protein_coding | Fgfr1 | fibroblast growth factor receptor 1 | 14182 | 8 | 25513654 | 25575718 | 1 | Mm.265716 |
| 7.53 | 7.13 | 7.62 | 7.09 | DN3ab | DN3cd | -0.46 | -6.78 | 8.70E-05 | 0.0041 | 0.34 | ENSMUSG00000022048 | protein_coding | Dpysl2 | dihydropyrimidinase-like 2 | 12934 | 14 | 66802864 | 66868688 | -1 | Mm.352648 Mm.475100 |
| 5.25 | 5.79 | 4.96 | 5.55 | DN3ab | DN3cd | 0.56 | 6.77 | 8.70E-05 | 0.0041 | 0.75 | ENSMUSG00000081434 | processed_pseudogene | Gm14165 | predicted gene 14165 | NA | 2 | 152353615 | 152354566 | 1 | |
| 3.71 | 3 | 4.44 | 3.49 | DN3ab | DN3cd | -0.84 | -6.77 | 8.70E-05 | 0.0041 | 0.33 | ENSMUSG00000039886 | protein_coding | Tmem120a | transmembrane protein 120A | 215210 | 5 | 135735485 | 135744448 | -1 | Mm.11688 |
| -1.45 | -3.92 | -0.16 | -4.02 | DN3ab | DN3cd | -3.19 | -6.75 | 9.00E-05 | 0.0042 | 0.12 | ENSMUSG00000054383 | protein_coding | Pnma1 | paraneoplastic antigen MA1 | 70481 | 12 | 84146131 | 84148489 | -1 | Mm.444348 |
| 0.05 | 2.78 | 0.78 | 2.86 | DN3ab | DN3cd | 2.39 | 6.74 | 9.00E-05 | 0.0042 | 0.37 | ENSMUSG00000024008 | protein_coding | Cpne5 | copine V | 240058 | 17 | 29155556 | 29238167 | -1 | Mm.39905 |
| 7.57 | 8.04 | 6.71 | 7.48 | DN3ab | DN3cd | 0.59 | 6.74 | 9.10E-05 | 0.0042 | 0.096 | ENSMUSG00000006585 | protein_coding | Cdt1 | chromatin licensing and DNA replication factor 1 | 67177 | 8 | 122568015 | 122573554 | 1 | Mm.21873 |
| 7.66 | 7.97 | 7.44 | 7.89 | DN3ab | DN3cd | 0.37 | 6.73 | 9.10E-05 | 0.0043 | 0.18 | ENSMUSG00000020608 | protein_coding | Smc6 | structural maintenance of chromosomes 6 | 67241 | 12 | 11265886 | 11319785 | 1 | Mm.173953 |
| 6.34 | 5.89 | 6.74 | 6.2 | DN3ab | DN3cd | -0.5 | -6.73 | 9.20E-05 | 0.0043 | 0.55 | ENSMUSG00000018567 | protein_coding | Gabarap | gamma-aminobutyric acid receptor associated protein | 56486 | 11 | 69991143 | 69994951 | 1 | Mm.272460 |
| 7.06 | 6.59 | 7.41 | 6.75 | DN3ab | DN3cd | -0.56 | -6.73 | 9.20E-05 | 0.0043 | 0.3 | ENSMUSG00000020570 | protein_coding | Sypl | synaptophysin-like protein | 19027 | 12 | 32953891 | 32979860 | 1 | Mm.246304 |
| 5.39 | 5.92 | 4.84 | 5.71 | DN3ab | DN3cd | 0.67 | 6.72 | 9.30E-05 | 0.0043 | 0.074 | ENSMUSG00000034329 | protein_coding | Brip1 | BRCA1 interacting protein C-terminal helicase 1 | 237911 | 11 | 86058138 | 86201193 | -1 | Mm.186143 |
| 5.57 | 5.16 | 6.13 | 5.61 | DN3ab | DN3cd | -0.46 | -6.72 | 9.30E-05 | 0.0043 | 0.35 | ENSMUSG00000028173 | protein_coding | Wls | wntless WNT ligand secretion mediator | 68151 | 3 | 159839672 | 159938664 | 1 | Mm.6766 |
| 5.47 | 6 | 5.22 | 6.09 | DN3ab | DN3cd | 0.68 | 6.71 | 9.30E-05 | 0.0043 | 0.073 | ENSMUSG00000027010 | protein_coding | Slc25a12 | solute carrier family 25 (mitochondrial carrier, Aralar), member 12 | 78830 | 2 | 71271063 | 71367749 | -1 | Mm.30928 Mm.146696 |
| 5.62 | 4.92 | 7.22 | 6.03 | DN3ab | DN3cd | -0.99 | -6.7 | 9.40E-05 | 0.0043 | 0.092 | ENSMUSG00000022565 | protein_coding | Plec | plectin | 18810 | 15 | 76170974 | 76232574 | -1 | Mm.234912 |
| 6.27 | 6.77 | 5.87 | 6.34 | DN3ab | DN3cd | 0.49 | 6.7 | 9.50E-05 | 0.0043 | 0.79 | ENSMUSG00000008690 | protein_coding | Ncaph2 | non-SMC condensin II complex, subunit H2 | 52683 | 15 | 89355719 | 89372826 | 1 | Mm.143167 |
| 4.92 | 5.55 | 4.36 | 5.07 | DN3ab | DN3cd | 0.66 | 6.69 | 9.60E-05 | 0.0044 | 0.71 | ENSMUSG00000025395 | protein_coding | Prim1 | DNA primase, p49 subunit | 19075 | 10 | 128015168 | 128030037 | 1 | Mm.2903 |
| 2.54 | 0.66 | 2.96 | 0.51 | DN3ab | DN3cd | -2.14 | -6.68 | 9.60E-05 | 0.0044 | 0.39 | ENSMUSG00000032737 | protein_coding | Inppl1 | inositol polyphosphate phosphatase-like 1 | 16332 | 7 | 101822632 | 101838229 | -1 | Mm.476000 |
| 7.09 | 6.61 | 7.22 | 6.8 | DN3ab | DN3cd | -0.45 | -6.68 | 9.70E-05 | 0.0044 | 0.67 | ENSMUSG00000022016 | protein_coding | Akap11 | A kinase (PRKA) anchor protein 11 | 219181 | 14 | 78492246 | 78536808 | -1 | Mm.89413 |
| 5.91 | 6.44 | 5.66 | 6.05 | DN3ab | DN3cd | 0.47 | 6.68 | 9.70E-05 | 0.0044 | 0.33 | ENSMUSG00000035478 | protein_coding | Mbd3 | methyl-CpG binding domain protein 3 | 17192 | 10 | 80392539 | 80399550 | -1 | Mm.7142 |
| 4.7 | 3.83 | 4.81 | 3.93 | DN3ab | DN3cd | -0.88 | -6.67 | 9.80E-05 | 0.0044 | 0.95 | ENSMUSG00000022517 | protein_coding | Mgrn1 | mahogunin, ring finger 1 | 17237 | 16 | 4886249 | 4938296 | 1 | Mm.291326 Mm.391848 |
| 4.3 | 3.71 | 5.24 | 4.53 | DN3ab | DN3cd | -0.66 | -6.66 | 9.90E-05 | 0.0044 | 0.57 | ENSMUSG00000038894 | protein_coding | Irs2 | insulin receptor substrate 2 | 384783 | 8 | 10984681 | 11008458 | -1 | Mm.407207 |
| 5.63 | 6.25 | 5.29 | 5.82 | DN3ab | DN3cd | 0.58 | 6.66 | 9.90E-05 | 0.0044 | 0.6 | ENSMUSG00000027242 | protein_coding | Wdr76 | WD repeat domain 76 | 241627 | 2 | 121506723 | 121544860 | 1 | Mm.293284 |
| 8.56 | 8.86 | 8 | 8.37 | DN3ab | DN3cd | 0.33 | 6.65 | 1.00E-04 | 0.0044 | 0.53 | ENSMUSG00000061360 | protein_coding | Phf5a | PHD finger protein 5A | 68479 | 15 | 81864520 | 81871911 | -1 | Mm.271715 |
| 5.73 | 5.18 | 6 | 5.59 | DN3ab | DN3cd | -0.48 | -6.65 | 1.00E-04 | 0.0044 | 0.31 | ENSMUSG00000019877 | protein_coding | Serinc1 | serine incorporator 1 | 56442 | 10 | 57515774 | 57532530 | -1 | Mm.29344 |
| 4.24 | 3.7 | 5.36 | 4.64 | DN3ab | DN3cd | -0.65 | -6.63 | 1.00E-04 | 0.0045 | 0.33 | ENSMUSG00000032846 | processed_transcript | Zswim6 | zinc finger SWIM-type containing 6 | 67263 | 13 | 107724617 | 107890064 | -1 | Mm.490418 |
| -1.12 | 1.5 | -2.6 | 1.32 | DN3ab | DN3cd | 3.2 | 6.63 | 1.00E-04 | 0.0045 | 0.17 | ENSMUSG00000030607 | protein_coding | Acan | aggrecan | 11595 | 7 | 79053483 | 79115099 | 1 | Mm.358571 |
| 6.51 | 5.99 | 6.84 | 6.1 | DN3ab | DN3cd | -0.63 | -6.62 | 1.00E-04 | 0.0045 | 0.28 | ENSMUSG00000030805 | protein_coding | Stx4a | syntaxin 4A (placental) | 20909 | 7 | 127824294 | 127849019 | 1 | Mm.24867 |
| 7.95 | 7.49 | 8.07 | 7.68 | DN3ab | DN3cd | -0.43 | -6.62 | 1.00E-04 | 0.0045 | 0.54 | ENSMUSG00000020315 | protein_coding | Sptbn1 | spectrin beta, non-erythrocytic 1 | 20742 | 11 | 30099395 | 30268175 | -1 | Mm.123110 Mm.466085 |
| 5.88 | 6.49 | 5.3 | 5.79 | DN3ab | DN3cd | 0.56 | 6.61 | 1.00E-04 | 0.0046 | 0.5 | ENSMUSG00000035198 | protein_coding | Tubg1 | tubulin, gamma 1 | 103733 | 11 | 101119938 | 101126419 | 1 | Mm.479145 |
| 4.19 | 5.14 | 3.55 | 5.17 | DN3ab | DN3cd | 1.21 | 6.61 | 1.00E-04 | 0.0046 | 0.061 | ENSMUSG00000039840 | protein_coding | Epg5 | ectopic P-granules autophagy protein 5 homolog (C. elegans) | 100502841 | 18 | 77938467 | 78035027 | 1 | Mm.297782 Mm.491179 |
| 7.92 | 8.39 | 7.95 | 8.42 | DN3ab | DN3cd | 0.47 | 6.61 | 0.00011 | 0.0046 | 0.99 | ENSMUSG00000001472 | protein_coding | Tcf25 | transcription factor 25 (basic helix-loop-helix) | 66855 | 8 | 123373753 | 123403835 | 1 | Mm.178818 Mm.295438 |
| 7.19 | 6.44 | 7.53 | 6.93 | DN3ab | DN3cd | -0.67 | -6.57 | 0.00011 | 0.0047 | 0.48 | ENSMUSG00000045165 | protein_coding | AI467606 | expressed sequence AI467606 | 101602 | 7 | 127091359 | 127093986 | 1 | Mm.490357 |
| 5.5 | 4.97 | 5.97 | 5.15 | DN3ab | DN3cd | -0.67 | -6.57 | 0.00011 | 0.0047 | 0.13 | ENSMUSG00000018999 | protein_coding | Slc35b4 | solute carrier family 35, member B4 | 58246 | 6 | 34153380 | 34177111 | -1 | Mm.245527 |
| 6.55 | 6.97 | 6.01 | 6.43 | DN3ab | DN3cd | 0.42 | 6.55 | 0.00011 | 0.0048 | 0.99 | ENSMUSG00000028953 | protein_coding | Abcf2 | ATP-binding cassette, sub-family F (GCN20), member 2 | 27407 | 5 | 24565345 | 24577467 | -1 | Mm.21629 |
| 4.71 | 5.84 | 4.2 | 5.02 | DN3ab | DN3cd | 1.01 | 6.54 | 0.00011 | 0.0048 | 0.34 | ENSMUSG00000028212 | protein_coding | Ccne2 | cyclin E2 | 12448 | 4 | 11191351 | 11204779 | 1 | Mm.35867 |
| 3.59 | 2.16 | 4.87 | 3.96 | DN3ab | DN3cd | -1.07 | -6.54 | 0.00011 | 0.0049 | 0.12 | ENSMUSG00000018750 | protein_coding | Zbtb4 | zinc finger and BTB domain containing 4 | 75580 | 11 | 69765912 | 69784023 | 1 | Mm.426361 |
| 4.91 | 5.55 | 3.89 | 5.04 | DN3ab | DN3cd | 0.82 | 6.51 | 0.00012 | 0.005 | 0.032 | ENSMUSG00000021485 | protein_coding | Mxd3 | Max dimerization protein 3 | 17121 | 13 | 55325168 | 55329823 | -1 | Mm.20350 |
| 5.68 | 4.24 | 6.12 | 5.26 | DN3ab | DN3cd | -1.12 | -6.5 | 0.00012 | 0.005 | 0.084 | ENSMUSG00000024620 | protein_coding | Pdgfrb | platelet derived growth factor receptor, beta polypeptide | 18596 | 18 | 61045150 | 61085061 | 1 | Mm.4146 |
| 2.11 | 3.33 | 0.79 | 2.93 | DN3ab | DN3cd | 1.5 | 6.5 | 0.00012 | 0.005 | 0.044 | ENSMUSG00000097839 | lincRNA | Gm26909 | predicted gene, 26909 | NA | 8 | 22644861 | 22646286 | 1 | |
| 5.64 | 4.84 | 5.83 | 5.26 | DN3ab | DN3cd | -0.69 | -6.49 | 0.00012 | 0.0051 | 0.28 | ENSMUSG00000032657 | protein_coding | Fam189b | family with sequence similarity 189, member B | 68521 | 3 | 89183143 | 89189295 | 1 | Mm.274708 |
| 7.26 | 7.75 | 6.79 | 7.09 | DN3ab | DN3cd | 0.41 | 6.46 | 0.00012 | 0.0052 | 0.12 | ENSMUSG00000028851 | protein_coding | Nudc | nudC nuclear distribution protein | 18221 | 4 | 133532542 | 133545996 | -1 | Mm.69 Mm.393592 |
| 7.6 | 7.11 | 7.9 | 7.57 | DN3ab | DN3cd | -0.42 | -6.46 | 0.00012 | 0.0052 | 0.2 | ENSMUSG00000024456 | protein_coding | Diaph1 | diaphanous related formin 1 | 13367 | 18 | 37843601 | 37935476 | -1 | Mm.195916 |
| 5.28 | 4.7 | 6.26 | 5.34 | DN3ab | DN3cd | -0.76 | -6.45 | 0.00013 | 0.0052 | 0.15 | ENSMUSG00000020736 | protein_coding | Nt5c | 5',3'-nucleotidase, cytosolic | 50773 | 11 | 115490420 | 115491862 | -1 | Mm.390379 |
| 3.42 | 4.45 | 3.71 | 4.37 | DN3ab | DN3cd | 0.85 | 6.43 | 0.00013 | 0.0053 | 0.15 | ENSMUSG00000008999 | protein_coding | Bmp7 | bone morphogenetic protein 7 | 12162 | 2 | 172868012 | 172940321 | -1 | Mm.595 Mm.392150 |
| 10.05 | 10.37 | 9.37 | 9.85 | DN3ab | DN3cd | 0.38 | 6.43 | 0.00013 | 0.0053 | 0.2 | ENSMUSG00000028333 | protein_coding | Anp32b | acidic (leucine-rich) nuclear phosphoprotein 32 family, member B | 67628 | 4 | 46450902 | 46472657 | 1 | Mm.263913 |
| 6.95 | 6.39 | 7.01 | 6.63 | DN3ab | DN3cd | -0.47 | -6.43 | 0.00013 | 0.0053 | 0.19 | ENSMUSG00000020458 | protein_coding | Rtn4 | reticulon 4 | 68585 | 11 | 29692947 | 29744331 | 1 | Mm.192580 Mm.440639 Mm.488364 |
| 3.79 | 2.96 | 3.69 | 2.53 | DN3ab | DN3cd | -0.97 | -6.42 | 0.00013 | 0.0054 | 0.28 | ENSMUSG00000003190 | protein_coding | Bcl2l12 | BCL2-like 12 (proline rich) | 75736 | 7 | 44991222 | 44998712 | -1 | Mm.261006 |
| -2.06 | 0.25 | -2.13 | 0.47 | DN3ab | DN3cd | 2.45 | 6.42 | 0.00013 | 0.0054 | 0.69 | ENSMUSG00000102956 | TEC | Gm37017 | predicted gene, 37017 | NA | 1 | 86988270 | 86989016 | 1 | |
| 5.22 | 4.35 | 5.66 | 5.13 | DN3ab | DN3cd | -0.69 | -6.41 | 0.00013 | 0.0054 | 0.1 | ENSMUSG00000009905 | protein_coding | Kdsr | 3-ketodihydrosphingosine reductase | 70750 | 1 | 106720459 | 106759727 | -1 | Mm.458911 |
| 7.35 | 6.83 | 7.94 | 7.21 | DN3ab | DN3cd | -0.63 | -6.4 | 0.00013 | 0.0054 | 0.31 | ENSMUSG00000052331 | protein_coding | Ankrd44 | ankyrin repeat domain 44 | 329154 | 1 | 54645340 | 54926387 | -1 | Mm.397910 |
| 7.94 | 8.23 | 7.74 | 8.2 | DN3ab | DN3cd | 0.37 | 6.4 | 0.00013 | 0.0054 | 0.13 | ENSMUSG00000070697 | protein_coding | Utp3 | UTP3 small subunit processome component | 65961 | 5 | 88554462 | 88556090 | 1 | Mm.490364 |
| 3.51 | 2.68 | 3.55 | 2.33 | DN3ab | DN3cd | -1 | -6.38 | 0.00014 | 0.0055 | 0.2 | ENSMUSG00000072680 | protein_coding | Tmem254c | transmembrane protein 254c | 100039192 | 14 | 26063919 | 26067355 | -1 | Mm.33344 |
| 6.24 | 5.57 | 6.49 | 6.1 | DN3ab | DN3cd | -0.53 | -6.38 | 0.00014 | 0.0055 | 0.062 | ENSMUSG00000017376 | protein_coding | Nlk | nemo like kinase | 18099 | 11 | 78567168 | 78697373 | -1 | Mm.9001 |
| 0.66 | 2.3 | 0.81 | 2.42 | DN3ab | DN3cd | 1.62 | 6.37 | 0.00014 | 0.0056 | 0.96 | ENSMUSG00000074580 | transcribed_processed_pseudogene | 4931440P22Rik | RIKEN cDNA 4931440P22 gene | NA | 3 | 65507257 | 65529379 | -1 | |
| 3.53 | 2.65 | 4.29 | 3.06 | DN3ab | DN3cd | -1.07 | -6.37 | 0.00014 | 0.0056 | 0.3 | ENSMUSG00000033416 | protein_coding | Gucd1 | guanylyl cyclase domain containing 1 | 68778 | 10 | 75351111 | 75517972 | -1 | Mm.29382 |
| 7.46 | 7.86 | 6.91 | 7.29 | DN3ab | DN3cd | 0.39 | 6.36 | 0.00014 | 0.0056 | 0.92 | ENSMUSG00000025134 | protein_coding | Alyref | Aly/REF export factor | 21681 | 11 | 120592121 | 120598365 | -1 | Mm.1886 |
| 8.39 | 8.65 | 8.29 | 8.57 | DN3ab | DN3cd | 0.27 | 6.34 | 0.00014 | 0.0057 | 0.89 | ENSMUSG00000031902 | protein_coding | Nfatc3 | nuclear factor of activated T cells, cytoplasmic, calcineurin dependent 3 | 18021 | 8 | 106058840 | 106130537 | 1 | Mm.383185 |
| 5.09 | 3.74 | 6.37 | 5.55 | DN3ab | DN3cd | -1.01 | -6.34 | 0.00014 | 0.0057 | 0.11 | ENSMUSG00000019960 | protein_coding | Dusp6 | dual specificity phosphatase 6 | 67603 | 10 | 99263231 | 99267489 | 1 | Mm.1791 |
| 3.77 | 4.69 | 3.07 | 4.75 | DN3ab | DN3cd | 1.22 | 6.34 | 0.00014 | 0.0057 | 0.04 | ENSMUSG00000027435 | protein_coding | Cd93 | CD93 antigen | 17064 | 2 | 148436640 | 148443563 | -1 | Mm.681 |
| 4.73 | 5.38 | 3.93 | 4.6 | DN3ab | DN3cd | 0.65 | 6.33 | 0.00014 | 0.0057 | 0.9 | ENSMUSG00000039748 | protein_coding | Exo1 | exonuclease 1 | 26909 | 1 | 175880581 | 175913489 | 1 | Mm.283046 |
| 3.44 | 4.24 | 2.87 | 4.26 | DN3ab | DN3cd | 1.03 | 6.33 | 0.00014 | 0.0057 | 0.05 | ENSMUSG00000061533 | protein_coding | Cep128 | centrosomal protein 128 | 75216 | 12 | 90998492 | 91384409 | -1 | Mm.312204 |
| 4.22 | 3.05 | 4.3 | 3.49 | DN3ab | DN3cd | -0.99 | -6.33 | 0.00014 | 0.0057 | 0.24 | ENSMUSG00000097353 | lincRNA | A430046D13Rik | Riken cDNA A430046D13 gene | NA | 10 | 128499364 | 128504242 | 1 | |
| 4.37 | 3.72 | 5.11 | 4.01 | DN3ab | DN3cd | -0.88 | -6.33 | 0.00015 | 0.0057 | 0.089 | ENSMUSG00000037960 | protein_coding | Card19 | caspase recruitment domain family, member 19 | 68480 | 13 | 49202950 | 49216037 | -1 | Mm.3336 |
| 5.4 | 4.87 | 5.69 | 5.22 | DN3ab | DN3cd | -0.5 | -6.33 | 0.00015 | 0.0057 | 0.72 | ENSMUSG00000034560 | protein_coding | Washc4 | WASH complex subunit 4 | 319277 | 10 | 83543752 | 83596473 | 1 | Mm.491194 |
| 6.23 | 6.73 | 5.19 | 5.81 | DN3ab | DN3cd | 0.55 | 6.31 | 0.00015 | 0.0057 | 0.53 | ENSMUSG00000026196 | protein_coding | Bard1 | BRCA1 associated RING domain 1 | 12021 | 1 | 71027498 | 71103146 | -1 | Mm.10764 Mm.490738 |
| 5.78 | 5.16 | 6.36 | 5.24 | DN3ab | DN3cd | -0.87 | -6.31 | 0.00015 | 0.0057 | 0.057 | ENSMUSG00000025417 | protein_coding | Pip4k2c | phosphatidylinositol-5-phosphate 4-kinase, type II, gamma | 117150 | 10 | 127197067 | 127211585 | -1 | Mm.22682 |
| 5.82 | 6.27 | 5.69 | 6.49 | DN3ab | DN3cd | 0.61 | 6.31 | 0.00015 | 0.0058 | 0.046 | ENSMUSG00000020901 | protein_coding | Pik3r5 | phosphoinositide-3-kinase regulatory subunit 5 | 320207 | 11 | 68432121 | 68497849 | 1 | Mm.244960 |
| 7.8 | 8.13 | 7.4 | 7.87 | DN3ab | DN3cd | 0.39 | 6.3 | 0.00015 | 0.0058 | 0.24 | ENSMUSG00000026020 | protein_coding | Nop58 | NOP58 ribonucleoprotein | 55989 | 1 | 59684971 | 59719044 | 1 | Mm.220367 |
| 6.95 | 6.3 | 6.81 | 6.42 | DN3ab | DN3cd | -0.52 | -6.3 | 0.00015 | 0.0058 | 0.1 | ENSMUSG00000027775 | protein_coding | Mfsd1 | major facilitator superfamily domain containing 1 | 66868 | 3 | 67582741 | 67604237 | 1 | Mm.491153 |
| 3.57 | 4.82 | 3.17 | 5.31 | DN3ab | DN3cd | 1.63 | 6.29 | 0.00015 | 0.0058 | 0.078 | ENSMUSG00000052160 | protein_coding | Pld4 | phospholipase D family, member 4 | 104759 | 12 | 112760655 | 112768990 | 1 | Mm.203915 |
| 5.11 | 5.76 | 4.2 | 5.23 | DN3ab | DN3cd | 0.79 | 6.29 | 0.00015 | 0.0058 | 0.13 | ENSMUSG00000084279 | processed_pseudogene | Gm13232 | predicted gene 13232 | NA | 4 | 145729581 | 145730212 | -1 | |
| 9.5 | 9.89 | 8.95 | 9.63 | DN3ab | DN3cd | 0.51 | 6.27 | 0.00016 | 0.0059 | 0.064 | ENSMUSG00000025364 | protein_coding | Pa2g4 | proliferation-associated 2G4 | 18813 | 10 | 128557766 | 128565987 | -1 | Mm.4742 |
| 4.38 | 5.3 | 4.52 | 5.8 | DN3ab | DN3cd | 1.1 | 6.26 | 0.00016 | 0.0059 | 0.32 | ENSMUSG00000023951 | protein_coding | Vegfa | vascular endothelial growth factor A | 22339 | 17 | 46016993 | 46032369 | -1 | Mm.282184 |
| 7.7 | 8.23 | 6.19 | 7.14 | DN3ab | DN3cd | 0.68 | 6.25 | 0.00016 | 0.006 | 0.046 | ENSMUSG00000040084 | protein_coding | Bub1b | BUB1B, mitotic checkpoint serine/threonine kinase | 12236 | 2 | 118598211 | 118641591 | 1 | Mm.29133 |
| 7.81 | 7.24 | 8.11 | 7.71 | DN3ab | DN3cd | -0.48 | -6.25 | 0.00016 | 0.006 | 0.31 | ENSMUSG00000038612 | protein_coding | Mcl1 | myeloid cell leukemia sequence 1 | 17210 | 3 | 95658788 | 95663176 | 1 | Mm.1639 |
| 4.31 | 2.81 | 5.14 | 4.3 | DN3ab | DN3cd | -1.09 | -6.24 | 0.00016 | 0.006 | 0.052 | ENSMUSG00000042349 | protein_coding | Ikbke | inhibitor of kappaB kinase epsilon | 56489 | 1 | 131254343 | 131279606 | -1 | Mm.386783 |
| 3.7 | 2.57 | 3.84 | 2.96 | DN3ab | DN3cd | -1 | -6.23 | 0.00016 | 0.0061 | 0.44 | ENSMUSG00000035235 | protein_coding | Trim13 | tripartite motif-containing 13 | 66597 | 14 | 61598247 | 61605946 | 1 | Mm.491258 |
| 7.08 | 7.58 | 6.37 | 7.03 | DN3ab | DN3cd | 0.57 | 6.22 | 0.00016 | 0.0061 | 0.42 | ENSMUSG00000006678 | protein_coding | Pola1 | polymerase (DNA directed), alpha 1 | 18968 | X | 93304767 | 93632155 | -1 | Mm.1923 |
| 5.49 | 5.95 | 5.2 | 5.74 | DN3ab | DN3cd | 0.5 | 6.22 | 0.00017 | 0.0061 | 0.64 | ENSMUSG00000038388 | protein_coding | Mpp6 | membrane protein, palmitoylated 6 (MAGUK p55 subfamily member 6) | 56524 | 6 | 50110241 | 50198939 | 1 | Mm.41288 Mm.248361 Mm.463873 |
| 9.21 | 8.88 | 9.59 | 9.05 | DN3ab | DN3cd | -0.43 | -6.21 | 0.00017 | 0.0062 | 0.1 | ENSMUSG00000022378 | protein_coding | Fam49b | family with sequence similarity 49, member B | 223601 | 15 | 63929097 | 64060478 | -1 | Mm.246108 |
| 8.12 | 8.5 | 8.04 | 8.34 | DN3ab | DN3cd | 0.34 | 6.2 | 0.00017 | 0.0062 | 0.45 | ENSMUSG00000089715 | protein_coding | Cbx6 | chromobox 6 | 494448 | 15 | 79823896 | 79834688 | -1 | Mm.38438 |
| 4.97 | 5.52 | 4.42 | 5.11 | DN3ab | DN3cd | 0.61 | 6.2 | 0.00017 | 0.0062 | 0.5 | ENSMUSG00000015748 | protein_coding | Prpf3 | pre-mRNA processing factor 3 | 70767 | 3 | 95830124 | 95855885 | -1 | Mm.279872 |
| 6.68 | 6.26 | 6.87 | 6.32 | DN3ab | DN3cd | -0.48 | -6.2 | 0.00017 | 0.0062 | 0.44 | ENSMUSG00000067851 | protein_coding | Arfgef1 | ADP-ribosylation factor guanine nucleotide-exchange factor 1(brefeldin A-inhibited) | 211673 | 1 | 10137571 | 10232670 | -1 | Mm.229141 |
| 3.89 | 4.61 | 3.31 | 4.4 | DN3ab | DN3cd | 0.87 | 6.18 | 0.00017 | 0.0063 | 0.19 | ENSMUSG00000051235 | protein_coding | Gen1 | GEN1, Holliday junction 5' flap endonuclease | 209334 | 12 | 11238920 | 11265801 | -1 | Mm.215295 |
| 8.75 | 9.14 | 7.83 | 8.28 | DN3ab | DN3cd | 0.41 | 6.18 | 0.00017 | 0.0063 | 0.7 | ENSMUSG00000063480 | protein_coding | Snu13 | SNU13 homolog, small nuclear ribonucleoprotein (U4/U6.U5) | 20826 | 15 | 82040525 | 82047598 | -1 | Mm.299312 |
| 6.11 | 6.56 | 5.51 | 6.28 | DN3ab | DN3cd | 0.57 | 6.17 | 0.00017 | 0.0063 | 0.07 | ENSMUSG00000040354 | protein_coding | Mars | methionine-tRNA synthetase | 216443 | 10 | 127296221 | 127311786 | -1 | Mm.28173 |
| 3.83 | 4.64 | 3.22 | 3.92 | DN3ab | DN3cd | 0.77 | 6.17 | 0.00018 | 0.0064 | 0.67 | ENSMUSG00000072980 | protein_coding | Oip5 | Opa interacting protein 5 | 70645 | 2 | 119609512 | 119618469 | -1 | Mm.246412 |
| 4.68 | 3.56 | 5 | 4.32 | DN3ab | DN3cd | -0.88 | -6.16 | 0.00018 | 0.0064 | 0.11 | ENSMUSG00000026657 | protein_coding | Frmd4a | FERM domain containing 4A | 209630 | 2 | 4017717 | 4614043 | 1 | Mm.37932 Mm.396281 Mm.487653 |
| 6.19 | 5.78 | 6.31 | 5.62 | DN3ab | DN3cd | -0.54 | -6.16 | 0.00018 | 0.0064 | 0.084 | ENSMUSG00000021987 | protein_coding | Mtmr6 | myotubularin related protein 6 | 219135 | 14 | 60265228 | 60302370 | 1 | Mm.247007 |
| 5.98 | 6.52 | 5.08 | 5.61 | DN3ab | DN3cd | 0.53 | 6.15 | 0.00018 | 0.0064 | 0.98 | ENSMUSG00000002274 | protein_coding | Metrn | meteorin, glial cell differentiation regulator | 70083 | 17 | 25794571 | 25797136 | -1 | Mm.41925 |
| 8.69 | 9.12 | 7.92 | 8.5 | DN3ab | DN3cd | 0.49 | 6.15 | 0.00018 | 0.0065 | 0.38 | ENSMUSG00000028312 | protein_coding | Smc2 | structural maintenance of chromosomes 2 | 14211 | 4 | 52439243 | 52488260 | 1 | Mm.2999 |
| 3.46 | 4.31 | 3.18 | 4.41 | DN3ab | DN3cd | 1.01 | 6.14 | 0.00018 | 0.0065 | 0.25 | ENSMUSG00000098090 | transcribed_unprocessed_pseudogene | 2700099C18Rik | RIKEN cDNA 2700099C18 gene | NA | 17 | 94750100 | 94775132 | 1 | |
| 3.92 | 4.67 | 2.88 | 3.87 | DN3ab | DN3cd | 0.83 | 6.14 | 0.00018 | 0.0065 | 0.41 | ENSMUSG00000078521 | protein_coding | Aunip | aurora kinase A and ninein interacting protein | 69885 | 4 | 134510999 | 134523927 | 1 | Mm.141592 |
| 4.82 | 5.41 | 3.87 | 4.56 | DN3ab | DN3cd | 0.62 | 6.13 | 0.00018 | 0.0065 | 0.65 | ENSMUSG00000022881 | protein_coding | Rfc4 | replication factor C (activator 1) 4 | 106344 | 16 | 23113943 | 23127737 | -1 | Mm.386835 |
| 6.34 | 6.82 | 5.55 | 6.42 | DN3ab | DN3cd | 0.64 | 6.13 | 0.00018 | 0.0065 | 0.05 | ENSMUSG00000015880 | protein_coding | Ncapg | non-SMC condensin I complex, subunit G | 54392 | 5 | 45669919 | 45700546 | 1 | Mm.32012 |
| 10.11 | 10.53 | 9.89 | 10.18 | DN3ab | DN3cd | 0.37 | 6.11 | 0.00019 | 0.0066 | 0.32 | ENSMUSG00000074656 | protein_coding | Eif2s2 | eukaryotic translation initiation factor 2, subunit 2 (beta) | 67204 | 2 | 154871410 | 154892935 | -1 | Mm.377134 Mm.470088 |
| 7.27 | 7.62 | 6.92 | 7.28 | DN3ab | DN3cd | 0.35 | 6.11 | 0.00019 | 0.0066 | 0.96 | ENSMUSG00000007670 | protein_coding | Khsrp | KH-type splicing regulatory protein | 16549 | 17 | 57021051 | 57031522 | -1 | Mm.34296 |
| 6.5 | 6.12 | 6.56 | 6.13 | DN3ab | DN3cd | -0.4 | -6.11 | 0.00019 | 0.0066 | 0.7 | ENSMUSG00000016534 | protein_coding | Lamp2 | lysosomal-associated membrane protein 2 | 16784 | X | 38401357 | 38456454 | -1 | Mm.486 Mm.414016 |
| 8.04 | 8.49 | 7.37 | 7.87 | DN3ab | DN3cd | 0.47 | 6.09 | 0.00019 | 0.0068 | 0.79 | ENSMUSG00000025001 | protein_coding | Hells | helicase, lymphoid specific | 15201 | 19 | 38930915 | 38971051 | 1 | Mm.57223 Mm.392920 Mm.486446 |
| -1.82 | 0.71 | -2.95 | 0.12 | DN3ab | DN3cd | 2.77 | 6.08 | 0.00019 | 0.0068 | 0.55 | ENSMUSG00000009216 | protein_coding | Fam163b | family with sequence similarity 163, member B | 109349 | 2 | 27110380 | 27142491 | -1 | Mm.314080 |
| 6.42 | 5.88 | 6.91 | 6.55 | DN3ab | DN3cd | -0.45 | -6.08 | 0.00019 | 0.0068 | 0.21 | ENSMUSG00000032384 | protein_coding | Csnk1g1 | casein kinase 1, gamma 1 | 214897 | 9 | 65908924 | 66045015 | 1 | Mm.325816 |
| 5.07 | 4.32 | 6.12 | 4.81 | DN3ab | DN3cd | -1.05 | -6.08 | 0.00019 | 0.0068 | 0.093 | ENSMUSG00000020272 | protein_coding | Stk10 | serine/threonine kinase 10 | 20868 | 11 | 32533305 | 32624587 | 1 | Mm.8235 |
| 7.52 | 7.87 | 7.03 | 7.67 | DN3ab | DN3cd | 0.48 | 6.07 | 2.00E-04 | 0.0068 | 0.038 | ENSMUSG00000020994 | protein_coding | Pnn | pinin | 18949 | 12 | 59066884 | 59073998 | 1 | Mm.22347 |
| 6.23 | 5.62 | 6.51 | 6.13 | DN3ab | DN3cd | -0.49 | -6.06 | 2.00E-04 | 0.0069 | 0.16 | ENSMUSG00000029407 | protein_coding | Uso1 | USO1 vesicle docking factor | 56041 | 5 | 92137938 | 92202798 | 1 | Mm.15868 |
| 5.5 | 5.1 | 5.59 | 5.03 | DN3ab | DN3cd | -0.47 | -6.06 | 2.00E-04 | 0.0069 | 0.27 | ENSMUSG00000017831 | protein_coding | Rab5a | RAB5A, member RAS oncogene family | 271457 | 17 | 53479234 | 53507680 | 1 | Mm.329123 |
| -4.01 | -1.96 | -3.72 | 0 | DN3ab | DN3cd | 2.95 | 6.05 | 2.00E-04 | 0.0069 | 0.061 | ENSMUSG00000105041 | TEC | Gm42676 | predicted gene 42676 | NA | 5 | 13525723 | 13528408 | 1 | |
| 3.69 | 3.12 | 4.63 | 3.53 | DN3ab | DN3cd | -0.86 | -6.05 | 2.00E-04 | 0.0069 | 0.041 | ENSMUSG00000053192 | protein_coding | Mllt11 | myeloid/lymphoid or mixed-lineage leukemia; translocated to, 11 | 56772 | 3 | 95219135 | 95232171 | -1 | Mm.331208 |
| 5.69 | 5.27 | 6.42 | 5.66 | DN3ab | DN3cd | -0.6 | -6.03 | 0.00021 | 0.007 | 0.062 | ENSMUSG00000103653 | processed_pseudogene | Gstp-ps | glutathione S-transferase, pi, pseudogene | NA | 1 | 192073820 | 192074450 | -1 | |
| 3.99 | 3.04 | 3.79 | 2.69 | DN3ab | DN3cd | -1.01 | -6.03 | 0.00021 | 0.007 | 0.65 | ENSMUSG00000048280 | protein_coding | Zfp738 | zinc finger protein 738 | 408068 | 13 | 67658685 | 67687071 | -1 | Mm.435551 |
| -1.69 | -3.92 | 0.34 | -3.34 | DN3ab | DN3cd | -3.01 | -6.03 | 0.00021 | 0.007 | 0.13 | ENSMUSG00000103632 | TEC | Gm38329 | predicted gene, 38329 | NA | 1 | 160727734 | 160728160 | -1 | |
| 10.36 | 10.55 | 9.74 | 10.04 | DN3ab | DN3cd | 0.23 | 6.01 | 0.00021 | 0.0071 | 0.13 | ENSMUSG00000039630 | protein_coding | Hnrnpu | heterogeneous nuclear ribonucleoprotein U | 51810 | 1 | 178321108 | 178337797 | -1 | Mm.86589 Mm.453372 Mm.482674 |
| 3.46 | 1.9 | 6.06 | 2.66 | DN3ab | DN3cd | -2.71 | -6.01 | 0.00021 | 0.0071 | 0.034 | ENSMUSG00000022419 | protein_coding | Deptor | DEP domain containing MTOR-interacting protein | 97998 | 15 | 55112317 | 55259271 | 1 | Mm.295397 Mm.393497 |
| 6.82 | 6.45 | 7.04 | 6.75 | DN3ab | DN3cd | -0.33 | -6 | 0.00021 | 0.0072 | 0.51 | ENSMUSG00000031812 | protein_coding | Map1lc3b | microtubule-associated protein 1 light chain 3 beta | 67443 | 8 | 121590361 | 121598760 | 1 | Mm.28357 |
| 4.94 | 5.4 | 4.22 | 4.97 | DN3ab | DN3cd | 0.57 | 6 | 0.00021 | 0.0072 | 0.13 | ENSMUSG00000097589 | processed_transcript | Dleu2 | deleted in lymphocytic leukemia, 2 | NA | 14 | 61602839 | 61682373 | -1 | |
| 4.34 | 5.41 | 3.31 | 5.36 | DN3ab | DN3cd | 1.42 | 6 | 0.00022 | 0.0072 | 0.032 | ENSMUSG00000027864 | protein_coding | Ptgfrn | prostaglandin F2 receptor negative regulator | 19221 | 3 | 101040232 | 101110278 | -1 | Mm.24807 |
| 0.83 | 2.06 | 1.22 | 3.23 | DN3ab | DN3cd | 1.65 | 5.99 | 0.00022 | 0.0072 | 0.14 | ENSMUSG00000040714 | protein_coding | Klc3 | kinesin light chain 3 | 232943 | 7 | 19394437 | 19404104 | -1 | Mm.213403 |
| 9.18 | 9.63 | 8.22 | 8.82 | DN3ab | DN3cd | 0.5 | 5.98 | 0.00022 | 0.0073 | 0.4 | ENSMUSG00000019942 | protein_coding | Cdk1 | cyclin-dependent kinase 1 | 12534 | 10 | 69335146 | 69352938 | -1 | Mm.281367 |
| 5.98 | 3.42 | 6.69 | 5.35 | DN3ab | DN3cd | -1.77 | -5.98 | 0.00022 | 0.0073 | 0.037 | ENSMUSG00000054672 | protein_coding | 5830411N06Rik | RIKEN cDNA 5830411N06 gene | 244234 | 7 | 140247284 | 140300736 | 1 | Mm.85125 |
| 6.13 | 6.62 | 5.78 | 6.11 | DN3ab | DN3cd | 0.42 | 5.95 | 0.00023 | 0.0074 | 0.21 | ENSMUSG00000034889 | protein_coding | Cactin | cactin, spliceosome C complex subunit | 70312 | 10 | 81321103 | 81326242 | 1 | Mm.87628 |
| 5.31 | 5.89 | 4.75 | 5.78 | DN3ab | DN3cd | 0.77 | 5.95 | 0.00023 | 0.0075 | 0.065 | ENSMUSG00000063659 | protein_coding | Zbtb18 | zinc finger and BTB domain containing 18 | 30928 | 1 | 177442351 | 177450764 | 1 | Mm.330700 |
| 5.28 | 3.77 | 6.23 | 5.44 | DN3ab | DN3cd | -1.06 | -5.94 | 0.00023 | 0.0075 | 0.034 | ENSMUSG00000042207 | protein_coding | Kdm5b | lysine (K)-specific demethylase 5B | 75605 | 1 | 134560171 | 134635285 | 1 | Mm.28995 Mm.391994 |
| 7.27 | 9.38 | 6.08 | 8.64 | DN3ab | DN3cd | 2.27 | 5.92 | 0.00024 | 0.0076 | 0.59 | ENSMUSG00000030867 | protein_coding | Plk1 | polo like kinase 1 | 18817 | 7 | 122159439 | 122169873 | 1 | Mm.16525 |
| 7.48 | 7.94 | 7.04 | 7.34 | DN3ab | DN3cd | 0.39 | 5.92 | 0.00024 | 0.0076 | 0.25 | ENSMUSG00000056209 | protein_coding | Npm3 | nucleoplasmin 3 | 18150 | 19 | 45747734 | 45749591 | -1 | Mm.1406 |
| 6.63 | 5.8 | 6.45 | 5.89 | DN3ab | DN3cd | -0.71 | -5.91 | 0.00024 | 0.0077 | 0.28 | ENSMUSG00000044408 | protein_coding | Sptssa | serine palmitoyltransferase, small subunit A | 104725 | 12 | 54645392 | 54656573 | -1 | Mm.485026 |
| 2.39 | 1.34 | 3.69 | 1.61 | DN3ab | DN3cd | -1.62 | -5.9 | 0.00024 | 0.0077 | 0.039 | ENSMUSG00000029994 | protein_coding | Anxa4 | annexin A4 | 11746 | 6 | 86736840 | 86793584 | -1 | Mm.259702 |
| 5.82 | 5.25 | 6.1 | 5.75 | DN3ab | DN3cd | -0.46 | -5.9 | 0.00024 | 0.0078 | 0.14 | ENSMUSG00000031659 | protein_coding | Adcy7 | adenylate cyclase 7 | 11513 | 8 | 88272403 | 88329962 | 1 | Mm.288206 |
| 3.42 | 4.21 | 3.09 | 4.36 | DN3ab | DN3cd | 1 | 5.88 | 0.00025 | 0.0079 | 0.15 | ENSMUSG00000052565 | protein_coding | Hist1h1d | histone cluster 1, H1d | 14957 | 13 | 23553262 | 23560686 | 1 | Mm.247046 |
| 4.62 | 5.14 | 3.6 | 4.57 | DN3ab | DN3cd | 0.68 | 5.88 | 0.00025 | 0.0079 | 0.04 | ENSMUSG00000039187 | protein_coding | Fanci | Fanconi anemia, complementation group I | 208836 | 7 | 79391929 | 79450264 | 1 | Mm.349493 |
| 5.87 | 6.51 | 5.14 | 5.66 | DN3ab | DN3cd | 0.59 | 5.87 | 0.00025 | 0.0079 | 0.57 | ENSMUSG00000036875 | protein_coding | Dna2 | DNA replication helicase/nuclease 2 | 327762 | 10 | 62947026 | 62974185 | 1 | Mm.21492 |
| 4.77 | 3.54 | 5.12 | 4.47 | DN3ab | DN3cd | -0.91 | -5.86 | 0.00025 | 0.0079 | 0.04 | ENSMUSG00000036833 | protein_coding | Pnpla7 | patatin-like phospholipase domain containing 7 | 241274 | 2 | 24976033 | 25054057 | 1 | Mm.389243 |
| 5.1 | 4.52 | 6.09 | 5.08 | DN3ab | DN3cd | -0.82 | -5.85 | 0.00026 | 0.008 | 0.11 | ENSMUSG00000074825 | protein_coding | Itpripl1 | inositol 1,4,5-triphosphate receptor interacting protein-like 1 | 73338 | 2 | 127138772 | 127143442 | -1 | Mm.57559 |
| 5.08 | 5.74 | 4.84 | 5.81 | DN3ab | DN3cd | 0.8 | 5.84 | 0.00026 | 0.008 | 0.26 | ENSMUSG00000016181 | protein_coding | Diexf | digestive organ expansion factor homolog (zebrafish) | 215193 | 1 | 193091104 | 193130272 | -1 | Mm.173758 Mm.491390 |
| 5.45 | 5.87 | 4.8 | 5.45 | DN3ab | DN3cd | 0.52 | 5.84 | 0.00026 | 0.0081 | 0.18 | ENSMUSG00000038416 | protein_coding | Cdc16 | CDC16 cell division cycle 16 | 69957 | 8 | 13757676 | 13781938 | 1 | Mm.182412 |
| 5.06 | 5.59 | 4.11 | 4.94 | DN3ab | DN3cd | 0.64 | 5.84 | 0.00026 | 0.0081 | 0.17 | ENSMUSG00000022466 | protein_coding | Rpap3 | RNA polymerase II associated protein 3 | 71919 | 15 | 97675097 | 97705825 | -1 | Mm.12255 |
| 3.3 | 2.54 | 4.21 | 2.86 | DN3ab | DN3cd | -1.08 | -5.84 | 0.00026 | 0.0081 | 0.098 | ENSMUSG00000018378 | protein_coding | Cuedc1 | CUE domain containing 1 | 103841 | 11 | 88098058 | 88194140 | 1 | Mm.374846 |
| 5.64 | 5.01 | 6.51 | 6.11 | DN3ab | DN3cd | -0.51 | -5.83 | 0.00026 | 0.0081 | 0.17 | ENSMUSG00000030605 | protein_coding | Mfge8 | milk fat globule-EGF factor 8 protein | 17304 | 7 | 79133768 | 79149060 | -1 | Mm.1451 |
| 1.31 | 2.96 | 1.12 | 2.19 | DN3ab | DN3cd | 1.42 | 5.83 | 0.00026 | 0.0081 | 0.23 | ENSMUSG00000078240 | protein_coding | Gm3550 | predicted gene 3550 | NA | 18 | 34736359 | 34738014 | 1 | |
| 5.13 | 5.57 | 4.56 | 5.23 | DN3ab | DN3cd | 0.53 | 5.8 | 0.00027 | 0.0083 | 0.22 | ENSMUSG00000029547 | protein_coding | Ints1 | integrator complex subunit 1 | 68510 | 5 | 139751282 | 139775674 | -1 | Mm.292942 |
| 7.91 | 6.96 | 8.87 | 6.94 | DN3ab | DN3cd | -1.44 | -5.8 | 0.00028 | 0.0083 | 0.035 | ENSMUSG00000029603 | protein_coding | Dtx1 | deltex 1, E3 ubiquitin ligase | 14357 | 5 | 120680202 | 120711927 | -1 | Mm.1645 |
| 5.29 | 4.33 | 5.09 | 4.62 | DN3ab | DN3cd | -0.72 | -5.79 | 0.00028 | 0.0084 | 0.034 | ENSMUSG00000097230 | lincRNA | Gm26853 | predicted gene, 26853 | NA | 8 | 19789798 | 19798548 | 1 | |
| 0.66 | 3.13 | 1.19 | 2.64 | DN3ab | DN3cd | 1.97 | 5.79 | 0.00028 | 0.0084 | 0.12 | ENSMUSG00000032010 | protein_coding | Usp2 | ubiquitin specific peptidase 2 | 53376 | 9 | 44067021 | 44095627 | 1 | Mm.272770 |
| 3.67 | 3.03 | 4.35 | 3.33 | DN3ab | DN3cd | -0.84 | -5.79 | 0.00028 | 0.0084 | 0.16 | ENSMUSG00000037722 | protein_coding | Gnpnat1 | glucosamine-phosphate N-acetyltransferase 1 | 54342 | 14 | 45376334 | 45404913 | -1 | Mm.312945 |
| 4.13 | 4.77 | 3.33 | 4.42 | DN3ab | DN3cd | 0.8 | 5.79 | 0.00028 | 0.0084 | 0.091 | ENSMUSG00000017146 | protein_coding | Brca1 | breast cancer 1, early onset | 12189 | 11 | 101488764 | 101551955 | -1 | Mm.244975 |
| 5.52 | 5.12 | 6.49 | 5.76 | DN3ab | DN3cd | -0.58 | -5.78 | 0.00028 | 0.0084 | 0.083 | ENSMUSG00000021365 | protein_coding | Nedd9 | neural precursor cell expressed, developmentally down-regulated gene 9 | 18003 | 13 | 41309581 | 41487362 | -1 | Mm.288980 |
| 4.14 | 2.82 | 4.57 | 3.8 | DN3ab | DN3cd | -1.01 | -5.76 | 0.00029 | 0.0086 | 0.11 | ENSMUSG00000029822 | protein_coding | Osbpl3 | oxysterol binding protein-like 3 | 71720 | 6 | 50293330 | 50456201 | -1 | Mm.44153 |
| 5.35 | 5.87 | 4.61 | 5.14 | DN3ab | DN3cd | 0.52 | 5.75 | 0.00029 | 0.0086 | 0.96 | ENSMUSG00000022177 | protein_coding | Haus4 | HAUS augmin-like complex, subunit 4 | 219072 | 14 | 54541785 | 54554616 | -1 | Mm.202152 |
| 8.59 | 9.02 | 8.37 | 8.6 | DN3ab | DN3cd | 0.34 | 5.75 | 0.00029 | 0.0086 | 0.085 | ENSMUSG00000041859 | protein_coding | Mcm3 | minichromosome maintenance complex component 3 | 17215 | 1 | 20802968 | 20820312 | -1 | Mm.4502 |
| 4 | 4.62 | 3.52 | 4.19 | DN3ab | DN3cd | 0.64 | 5.72 | 3.00E-04 | 0.0088 | 0.83 | ENSMUSG00000031669 | protein_coding | Gins3 | GINS complex subunit 3 (Psf3 homolog) | 78833 | 8 | 95633453 | 95645060 | 1 | Mm.35546 |
| 4.11 | 4.76 | 3.1 | 3.99 | DN3ab | DN3cd | 0.73 | 5.72 | 3.00E-04 | 0.0088 | 0.38 | ENSMUSG00000027454 | protein_coding | Gins1 | GINS complex subunit 1 (Psf1 homolog) | 69270 | 2 | 150905400 | 150931280 | 1 | Mm.271603 |
| 0.37 | 2.29 | -0.97 | 2.75 | DN3ab | DN3cd | 2.52 | 5.71 | 3.00E-04 | 0.0088 | 0.034 | ENSMUSG00000032278 | protein_coding | Paqr5 | progestin and adipoQ receptor family member V | 74090 | 9 | 61953481 | 62026856 | -1 | Mm.273267 |
| 5.36 | 4.91 | 5.94 | 5.21 | DN3ab | DN3cd | -0.59 | -5.7 | 0.00031 | 0.0089 | 0.15 | ENSMUSG00000027678 | protein_coding | Ncoa3 | nuclear receptor coactivator 3 | 17979 | 2 | 165992636 | 166073242 | 1 | Mm.469663 |
| 7.96 | 8.43 | 7.02 | 7.95 | DN3ab | DN3cd | 0.66 | 5.69 | 0.00031 | 0.009 | 0.037 | ENSMUSG00000034333 | protein_coding | Zbed4 | zinc finger, BED type containing 4 | 223773 | 15 | 88751660 | 88784516 | 1 | Mm.396622 |
| 7.57 | 8.01 | 6.69 | 7.54 | DN3ab | DN3cd | 0.61 | 5.68 | 0.00032 | 0.009 | 0.044 | ENSMUSG00000036768 | protein_coding | Kif15 | kinesin family member 15 | 209737 | 9 | 122951046 | 123018733 | 1 | Mm.278920 |
| 7.87 | 8.2 | 7.63 | 8.11 | DN3ab | DN3cd | 0.4 | 5.68 | 0.00032 | 0.0091 | 0.29 | ENSMUSG00000029623 | protein_coding | Pdap1 | PDGFA associated protein 1 | 231887 | 5 | 145128769 | 145140238 | -1 | Mm.188851 |
| 6.48 | 7.07 | 5.26 | 6.19 | DN3ab | DN3cd | 0.72 | 5.67 | 0.00032 | 0.0091 | 0.2 | ENSMUSG00000022033 | protein_coding | Pbk | PDZ binding kinase | 52033 | 14 | 65805837 | 65817822 | 1 | Mm.24337 |
| 2.25 | 3.58 | 1.79 | 2.56 | DN3ab | DN3cd | 1.13 | 5.66 | 0.00033 | 0.0092 | 0.17 | ENSMUSG00000032478 | protein_coding | Nme6 | NME/NM23 nucleoside diphosphate kinase 6 | 54369 | 9 | 109832749 | 109843116 | 1 | Mm.12951 |
| 6.92 | 7.39 | 6.05 | 6.88 | DN3ab | DN3cd | 0.61 | 5.65 | 0.00033 | 0.0093 | 0.092 | ENSMUSG00000037474 | protein_coding | Dtl | denticleless E3 ubiquitin protein ligase | 76843 | 1 | 191537356 | 191575544 | -1 | Mm.189102 |

| | | | | | | | | | | | | | | | | | | | | |
|---|---|---|---|---|---|---|---|---|---|---|---|---|---|---|---|---|---|---|---|---|
| 3.25 | 4.04 | 2.55 | 3.82 | DN3ab | DN3cd | 0.97 | 5.64 | 0.00033 | 0.0093 | 0.16 | ENSMUSG00000014786 | protein_coding | Slc9a5 | solute carrier family 9 (sodium/hydrogen exchanger), member 5 | 277973 | 8 | 105348843 | 105369881 | 1 | Mm.334766 |
| 2.44 | 3.51 | 1.63 | 2.76 | DN3ab | DN3cd | 1.09 | 5.63 | 0.00034 | 0.0093 | 0.87 | ENSMUSG00000034773 | protein_coding | BC030867 | cDNA sequence BC030867 | 217216 | 11 | 102248882 | 102265187 | 1 | Mm.247335 |
| -1.82 | -3.92 | -0.41 | -4.02 | DN3ab | DN3cd | -2.87 | -5.62 | 0.00034 | 0.0094 | 0.12 | ENSMUSG00000071573 | protein_coding | Rnls | renalase, FAD-dependent amine oxidase | 67795 | 19 | 33137747 | 33392295 | -1 | Mm.204591 |
| 0.27 | 2.16 | -0.86 | 2.72 | DN3ab | DN3cd | 2.51 | 5.62 | 0.00034 | 0.0094 | 0.05 | ENSMUSG00000047907 | protein_coding | Tshz2 | teashirt zinc finger family member 2 | 228911 | 2 | 169633013 | 170071816 | 1 | Mm.315789 Mm.468546 Mm.482636 |
| -0.43 | -3.92 | 1.26 | -0.51 | DN3ab | DN3cd | -2.46 | -5.61 | 0.00035 | 0.0095 | 0.031 | ENSMUSG00000020155 | protein_coding | Kcnmb1 | potassium large conductance calcium-activated channel, subfamily M, beta member 1 | 16533 | 11 | 33963013 | 33973641 | 1 | Mm.6206 |
| 7.74 | 7.46 | 8.26 | 7.67 | DN3ab | DN3cd | -0.44 | -5.58 | 0.00036 | 0.0097 | 0.032 | ENSMUSG00000034724 | protein_coding | Cnot6l | CCR4-NOT transcription complex, subunit 6-like | 231464 | 5 | 96070333 | 96164171 | -1 | Mm.28374 Mm.384746 |
| 6.68 | 7.1 | 5.85 | 6.59 | DN3ab | DN3cd | 0.55 | 5.57 | 0.00036 | 0.0098 | 0.11 | ENSMUSG00000007050 | protein_coding | Lsm2 | LSM2 homolog, U6 small nuclear RNA and mRNA degradation associated | 27756 | 17 | 34981862 | 34985891 | 1 | Mm.165735 |
| 5.27 | 5.69 | 4.94 | 5.56 | DN3ab | DN3cd | 0.51 | 5.57 | 0.00037 | 0.0098 | 0.27 | ENSMUSG00000046111 | protein_coding | Cep295 | centrosomal protein 295 | 319675 | 9 | 15316915 | 15357788 | -1 | Mm.218317 |
| 10.3 | 11.21 | 9.5 | 11.19 | DN3ab | DN3cd | 1.23 | 5.57 | 0.00037 | 0.0098 | 0.077 | ENSMUSG00000030795 | protein_coding | Fus | fused in sarcoma | 233908 | 7 | 127967457 | 127985701 | 1 | Mm.277680 |
| 5.78 | 4.99 | 7.19 | 5.56 | DN3ab | DN3cd | -1.26 | -5.55 | 0.00037 | 0.0099 | 0.052 | ENSMUSG00000015355 | protein_coding | Cd48 | CD48 antigen | 12506 | 1 | 171682009 | 171705258 | 1 | Mm.1738 |
| 0.1 | 2.53 | -2.36 | 1.71 | DN3ab | DN3cd | 2.94 | 5.54 | 0.00038 | 0.01 | 0.15 | ENSMUSG00000104669 | antisense | Gm38411 | predicted gene, 38411 | NA | 3 | 94368612 | 94377245 | -1 | |
| 6.42 | 6.99 | 5.6 | 6.57 | DN3ab | DN3cd | 0.73 | 5.53 | 0.00038 | 0.01 | 0.14 | ENSMUSG00000025742 | protein_coding | Prps2 | phosphoribosyl pyrophosphate synthetase 2 | 110639 | X | 167346322 | 167382749 | -1 | Mm.272955 |
| 2.84 | -0.7 | -0.74 | -2.43 | DN3ab | DN3cd | -3.01 | -5.53 | 0.00039 | 0.01 | 0.11 | ENSMUSG00000046997 | protein_coding | Spsb4 | splA/ryanodine receptor domain and SOCS box containing 4 | 211949 | 9 | 96943482 | 97018843 | -1 | Mm.491271 |
| 3.53 | 4.24 | 3.42 | 4.81 | DN3ab | DN3cd | 1.03 | 5.51 | 0.00039 | 0.01 | 0.054 | ENSMUSG00000092837 | ribozyme | Rpph1 | ribonuclease P RNA component H1 | NA | 14 | 50807449 | 50807767 | -1 | |
| 6.88 | 6.59 | 7.11 | 6.56 | DN3ab | DN3cd | -0.41 | -5.51 | 0.00039 | 0.01 | 0.05 | ENSMUSG00000015023 | protein_coding | Ddx19a | DEAD (Asp-Glu-Ala-Asp) box polypeptide 19a | 13680 | 8 | 110974996 | 110997800 | -1 | Mm.287901 |
| 4.17 | 3.54 | 5.19 | 3.91 | DN3ab | DN3cd | -0.98 | -5.51 | 0.00039 | 0.01 | 0.054 | ENSMUSG00000035199 | protein_coding | Arl6ip5 | ADP-ribosylation factor-like 6 interacting protein 5 | 65106 | 6 | 97210689 | 97233315 | 1 | Mm.291014 |
| 5.68 | 5.03 | 5.43 | 5.02 | DN3ab | DN3cd | -0.54 | -5.49 | 4.00E-04 | 0.01 | 0.2 | ENSMUSG00000019461 | protein_coding | Plscr3 | phospholipid scramblase 3 | 70310 | 11 | 69846376 | 69852058 | 1 | Mm.28874 |
| 6.48 | 6.88 | 5.87 | 6.46 | DN3ab | DN3cd | 0.48 | 5.49 | 0.00041 | 0.01 | 0.28 | ENSMUSG00000097195 | processed_transcript | Snhg5 | small nucleolar RNA host gene 5 | NA | 9 | 88521050 | 88522929 | -1 | |
| 6.51 | 5.98 | 7.65 | 6.48 | DN3ab | DN3cd | -0.88 | -5.47 | 0.00042 | 0.011 | 0.035 | ENSMUSG00000069094 | protein_coding | Pde7a | phosphodiesterase 7A | 18583 | 3 | 19223108 | 19311322 | -1 | Mm.355614 |
| 4.05 | 3.14 | 3.51 | 2.98 | DN3ab | DN3cd | -0.75 | -5.45 | 0.00042 | 0.011 | 0.14 | ENSMUSG00000025436 | protein_coding | Atp23 | ATP23 metallopeptidase and ATP synthase assembly factor homolog | 68876 | 10 | 126868427 | 126901355 | -1 | Mm.276769 |
| 6.76 | 6.49 | 6.84 | 6.37 | DN3ab | DN3cd | -0.36 | -5.45 | 0.00043 | 0.011 | 0.14 | ENSMUSG00000062014 | protein_coding | Gmfb | glia maturation factor, beta | 63985 | 14 | 46808149 | 46822242 | -1 | Mm.87312 |
| 1.27 | 2.62 | 1.39 | 2.28 | DN3ab | DN3cd | 1.14 | 5.45 | 0.00043 | 0.011 | 0.26 | ENSMUSG00000108095 | lincRNA | Gm44067 | predicted gene, 44067 | NA | 6 | 128843786 | 128846959 | -1 | |
| 7.47 | 7.16 | 7.93 | 7.31 | DN3ab | DN3cd | -0.46 | -5.44 | 0.00043 | 0.011 | 0.05 | ENSMUSG00000022781 | protein_coding | Pak2 | p21 (RAC1) activated kinase 2 | 224105 | 16 | 32016290 | 32079342 | -1 | Mm.234204 |
| 6.54 | 6.97 | 5.63 | 6.47 | DN3ab | DN3cd | 0.59 | 5.43 | 0.00044 | 0.011 | 0.053 | ENSMUSG00000029363 | protein_coding | Rfc5 | replication factor C (activator 1) 5 | 72151 | 5 | 117378103 | 117389047 | -1 | Mm.27997 |
| 6.66 | 6.28 | 7.05 | 6.25 | DN3ab | DN3cd | -0.59 | -5.43 | 0.00044 | 0.011 | 0.033 | ENSMUSG00000024856 | protein_coding | Cdk2ap2 | CDK2-associated protein 2 | 52004 | 19 | 4097182 | 4099019 | 1 | Mm.29609 |
| 5.88 | 5.23 | 7.05 | 5.65 | DN3ab | DN3cd | -1.05 | -5.42 | 0.00044 | 0.011 | 0.039 | ENSMUSG00000020437 | protein_coding | Myo1g | myosin IG | 246177 | 11 | 6506548 | 6520965 | -1 | Mm.239554 |
| 3.85 | 4.44 | 3.14 | 4.27 | DN3ab | DN3cd | 0.8 | 5.41 | 0.00045 | 0.011 | 0.051 | ENSMUSG00000091264 | protein_coding | Smim13 | small integral membrane protein 13 | 108934 | 13 | 41249844 | 41276577 | 1 | Mm.38578 |
| 4.77 | 5.29 | 4.58 | 5.62 | DN3ab | DN3cd | 0.76 | 5.41 | 0.00045 | 0.011 | 0.047 | ENSMUSG00000038623 | protein_coding | Tm6sf1 | transmembrane 6 superfamily member 1 | 107769 | 7 | 81859001 | 81884434 | 1 | Mm.221412 Mm.392934 |
| 3.59 | 1.68 | 2.72 | 1.85 | DN3ab | DN3cd | -1.45 | -5.41 | 0.00045 | 0.011 | 0.034 | ENSMUSG00000030532 | protein_coding | Hddc3 | HD domain containing 3 | 68695 | 7 | 80343104 | 80350547 | 1 | Mm.21171 |
| 8.66 | 8.85 | 8.47 | 8.78 | DN3ab | DN3cd | 0.24 | 5.37 | 0.00047 | 0.012 | 0.15 | ENSMUSG00000027479 | protein_coding | Mapre1 | microtubule-associated protein, RP/EB family, member 1 | 13589 | 2 | 153741274 | 153773310 | 1 | Mm.143877 |
| 9.71 | 9.89 | 9.75 | 10.13 | DN3ab | DN3cd | 0.28 | 5.37 | 0.00047 | 0.012 | 0.031 | ENSMUSG00000101249 | unprocessed_pseudogene | Gm29216 | predicted gene 29216 | NA | 1 | 24615706 | 24616197 | -1 | |
| 5.7 | 6.08 | 5.28 | 5.86 | DN3ab | DN3cd | 0.47 | 5.36 | 0.00048 | 0.012 | 0.26 | ENSMUSG00000050890 | protein_coding | Pdik1l | PDLIM1 interacting kinase 1 like | 230809 | 4 | 134275002 | 134287895 | -1 | Mm.22778 |
| 5.35 | 5.99 | 4.01 | 5.14 | DN3ab | DN3cd | 0.8 | 5.36 | 0.00048 | 0.012 | 0.12 | ENSMUSG00000054115 | protein_coding | Skp2 | S-phase kinase-associated protein 2 (p45) | 27401 | 15 | 9111985 | 9155425 | -1 | Mm.35584 |
| 8.59 | 8.93 | 7.6 | 8.25 | DN3ab | DN3cd | 0.46 | 5.35 | 0.00048 | 0.012 | 0.059 | ENSMUSG00000028873 | protein_coding | Cdca8 | cell division cycle associated 8 | 52276 | 4 | 124918465 | 124939311 | -1 | Mm.28038 |
| 5.4 | 5.01 | 5.4 | 4.78 | DN3ab | DN3cd | -0.5 | -5.35 | 0.00048 | 0.012 | 0.19 | ENSMUSG00000022337 | protein_coding | Emc2 | ER membrane protein complex subunit 2 | 66736 | 15 | 43477229 | 43527763 | 1 | Mm.244512 |
| 5.14 | 4.39 | 4.93 | 3.62 | DN3ab | DN3cd | -0.97 | -5.32 | 5.00E-04 | 0.012 | 0.12 | ENSMUSG00000026035 | protein_coding | Ppil3 | peptidylprolyl isomerase (cyclophilin)-like 3 | 70225 | 1 | 58430994 | 58445486 | -1 | Mm.340195 |
| 5.23 | 5.89 | 4.28 | 5.29 | DN3ab | DN3cd | 0.79 | 5.3 | 0.00052 | 0.012 | 0.24 | ENSMUSG00000095865 | processed_pseudogene | Gm13237 | predicted gene 13237 | NA | 4 | 146564453 | 146565087 | -1 | |
| 3.12 | 3.77 | 2.77 | 3.93 | DN3ab | DN3cd | 0.86 | 5.28 | 0.00053 | 0.013 | 0.1 | ENSMUSG00000111030 | TEC | Gm47234 | predicted gene, 47234 | NA | 9 | 44634195 | 44635103 | 1 | |
| 1.07 | 2.29 | 0.15 | 2.36 | DN3ab | DN3cd | 1.58 | 5.26 | 0.00054 | 0.013 | 0.09 | ENSMUSG00000074892 | protein_coding | B3galt5 | UDP-Gal:betaGlcNAc beta 1,3-galactosyltransferase, polypeptide 5 | 93961 | 16 | 96235801 | 96319859 | 1 | Mm.154783 |
| 5.37 | 4.89 | 6.65 | 5.56 | DN3ab | DN3cd | -0.82 | -5.25 | 0.00055 | 0.013 | 0.034 | ENSMUSG00000021585 | protein_coding | Cast | calpastatin | 12380 | 13 | 74692368 | 74808810 | -1 | Mm.29163 Mm.441995 |
| 5.48 | 5.84 | 5.34 | 6.08 | DN3ab | DN3cd | 0.54 | 5.25 | 0.00055 | 0.013 | 0.037 | ENSMUSG00000032536 | protein_coding | Trak1 | trafficking protein, kinesin binding 1 | 67095 | 9 | 121297502 | 121474918 | 1 | Mm.491112 |
| 4.55 | 4.14 | 5.48 | 4.62 | DN3ab | DN3cd | -0.66 | -5.25 | 0.00056 | 0.013 | 0.049 | ENSMUSG00000036402 | protein_coding | Gng12 | guanine nucleotide binding protein (G protein), gamma 12 | 14701 | 6 | 66896397 | 67021350 | 1 | Mm.234342 |
| 5.78 | 6.15 | 4.66 | 5.29 | DN3ab | DN3cd | 0.46 | 5.22 | 0.00058 | 0.013 | 0.14 | ENSMUSG00000058290 | protein_coding | Espl1 | extra spindle pole bodies 1, separase | 105988 | 15 | 102296266 | 102324357 | 1 | Mm.288324 |
| 6.48 | 6.1 | 6.82 | 6.01 | DN3ab | DN3cd | -0.58 | -5.18 | 6.00E-04 | 0.014 | 0.045 | ENSMUSG00000050708 | protein_coding | Ftl1 | ferritin light polypeptide 1 | 14325 | 7 | 45457944 | 45459884 | -1 | Mm.30357 Mm.348374 Mm.431913 Mm.491102 |
| 3.62 | 2.85 | 3.74 | 2.34 | DN3ab | DN3cd | -1.03 | -5.18 | 0.00061 | 0.014 | 0.096 | ENSMUSG00000072676 | protein_coding | Tmem254a | transmembrane protein 254a | 66039 | 14 | 25923300 | 25927672 | -1 | Mm.33344 |
| 4.38 | 5.3 | 3.58 | 5.18 | DN3ab | DN3cd | 1.18 | 5.18 | 0.00061 | 0.014 | 0.15 | ENSMUSG00000003882 | protein_coding | Il7r | interleukin 7 receptor | 16197 | 15 | 9505788 | 9530176 | -1 | Mm.389 |
| 6.16 | 6.56 | 5.3 | 5.94 | DN3ab | DN3cd | 0.49 | 5.15 | 0.00063 | 0.014 | 0.23 | ENSMUSG00000029910 | protein_coding | Mad2l1 | MAD2 mitotic arrest deficient-like 1 | 56150 | 6 | 66535390 | 66547220 | 1 | Mm.489653 |
| 2.03 | 3.37 | 2.29 | 2.98 | DN3ab | DN3cd | 1.04 | 5.14 | 0.00064 | 0.014 | 0.096 | ENSMUSG00000091383 | processed_pseudogene | Hist1h2al | histone cluster 1, H2al | NA | 13 | 51202727 | 51203065 | -1 | |
| 5.4 | 5.89 | 4.23 | 5.27 | DN3ab | DN3cd | 0.68 | 5.14 | 0.00064 | 0.014 | 0.032 | ENSMUSG00000041147 | protein_coding | Brca2 | breast cancer 2, early onset | 12190 | 5 | 150522630 | 150570329 | 1 | Mm.236256 |
| -0.65 | 1.28 | -2.9 | 1.21 | DN3ab | DN3cd | 2.82 | 5.11 | 0.00067 | 0.014 | 0.035 | ENSMUSG00000032936 | protein_coding | Camkv | CaM kinase-like vesicle-associated | 235604 | 9 | 107935674 | 107949691 | 1 | Mm.274540 |
| 5.32 | 5.77 | 5.18 | 6.12 | DN3ab | DN3cd | 0.68 | 5.09 | 0.00068 | 0.014 | 0.049 | ENSMUSG00000036181 | protein_coding | Hist1h1c | histone cluster 1, H1c | 50708 | 13 | 23738808 | 23740367 | 1 | Mm.193539 |
| 2.46 | 3.34 | 1.83 | 3.32 | DN3ab | DN3cd | 1.11 | 5.07 | 7.00E-04 | 0.015 | 0.17 | ENSMUSG00000032352 | protein_coding | Lrrc1 | leucine rich repeat containing 1 | 214345 | 9 | 77430823 | 77544870 | -1 | Mm.28534 Mm.482664 |
| 1.83 | 3.05 | 1.42 | 3.59 | DN3ab | DN3cd | 1.61 | 5.07 | 7.00E-04 | 0.015 | 0.13 | ENSMUSG00000030123 | protein_coding | Plxnd1 | plexin D1 | 67784 | 6 | 115954811 | 115995005 | -1 | Mm.3085 |
| 5.19 | 5.7 | 4.82 | 5.1 | DN3ab | DN3cd | 0.42 | 5.06 | 0.00071 | 0.015 | 0.14 | ENSMUSG00000034875 | protein_coding | Nudt19 | nudix (nucleoside diphosphate linked moiety X)-type motif 19 | 110959 | 7 | 35547185 | 35556304 | -1 | Mm.410206 Mm.418631 Mm.474965 |
| 6.32 | 6.05 | 6.77 | 6.23 | DN3ab | DN3cd | -0.41 | -5 | 0.00078 | 0.016 | 0.068 | ENSMUSG00000018171 | protein_coding | Vmp1 | vacuole membrane protein 1 | 75909 | 11 | 86583865 | 86683836 | -1 | Mm.390398 Mm.491180 |
| 3.43 | 4.02 | 3.07 | 4.23 | DN3ab | DN3cd | 0.84 | 4.98 | 0.00079 | 0.016 | 0.075 | ENSMUSG00000104814 | TEC | Gm42979 | predicted gene 42979 | NA | 5 | 129556147 | 129558374 | 1 | |
| 2.43 | 3.17 | 1.35 | 2.56 | DN3ab | DN3cd | 0.89 | 4.98 | 8.00E-04 | 0.016 | 0.19 | ENSMUSG00000047757 | protein_coding | Fancb | Fanconi anemia, complementation group B | 237211 | X | 164980592 | 164997272 | 1 | Mm.336317 |
| 4.58 | 5.32 | 4.19 | 4.52 | DN3ab | DN3cd | 0.58 | 4.96 | 0.00082 | 0.016 | 0.054 | ENSMUSG00000034321 | protein_coding | Exosc1 | exosome component 1 | 66583 | 19 | 41922980 | 41933314 | -1 | Mm.289086 |
| 2.61 | 3.43 | 1.65 | 3.15 | DN3ab | DN3cd | 1.06 | 4.96 | 0.00082 | 0.016 | 0.12 | ENSMUSG00000104197 | TEC | Gm37632 | predicted gene, 37632 | NA | 1 | 137971174 | 137972315 | 1 | |
| 8.46 | 8.68 | 7.93 | 8.33 | DN3ab | DN3cd | 0.3 | 4.94 | 0.00083 | 0.016 | 0.13 | ENSMUSG00000022797 | protein_coding | Tfrc | transferrin receptor | 22042 | 16 | 32608920 | 32632794 | 1 | Mm.28683 |
| 6.03 | 6.32 | 5.78 | 6.39 | DN3ab | DN3cd | 0.43 | 4.94 | 0.00084 | 0.016 | 0.043 | ENSMUSG00000091905 | processed_pseudogene | Gm6395 | predicted gene 6395 | NA | 12 | 64786311 | 64787001 | 1 | |
| 8.72 | 8.39 | 9.3 | 8.56 | DN3ab | DN3cd | -0.54 | -4.92 | 0.00086 | 0.017 | 0.051 | ENSMUSG00000076749 | TR_C_gene | Tcrg-C1 | T cell receptor gamma, constant 1 | NA | 13 | 19214103 | 19217024 | 1 | |
| 8.74 | 9.03 | 8.58 | 9.21 | DN3ab | DN3cd | 0.44 | 4.91 | 0.00087 | 0.017 | 0.047 | ENSMUSG00000043091 | protein_coding | Tuba1c | tubulin, alpha 1C | 22146 | 15 | 99029891 | 99038110 | 1 | Mm.88212 Mm.485827 |
| 4.31 | 4.69 | 4.09 | 4.91 | DN3ab | DN3cd | 0.58 | 4.91 | 0.00087 | 0.017 | 0.041 | ENSMUSG00000027634 | protein_coding | Ndrg3 | N-myc downstream regulated gene 3 | 29812 | 2 | 156927345 | 156992056 | -1 | Mm.279256 |
| 6.74 | 7.01 | 6 | 6.48 | DN3ab | DN3cd | 0.36 | 4.9 | 0.00089 | 0.017 | 0.14 | ENSMUSG00000000838 | protein_coding | Fmr1 | fragile X mental retardation 1 | 14265 | X | 68678541 | 68717963 | 1 | Mm.3451 |
| 7.8 | 8.01 | 7.89 | 8.37 | DN3ab | DN3cd | 0.34 | 4.88 | 0.00091 | 0.017 | 0.036 | ENSMUSG00000022255 | protein_coding | Mtdh | metadherin | 67154 | 15 | 34082694 | 34145624 | 1 | Mm.130883 |
| 2.15 | 2.96 | 1.79 | 3.62 | DN3ab | DN3cd | 1.26 | 4.86 | 0.00093 | 0.017 | 0.032 | ENSMUSG00000032549 | protein_coding | Rab6b | RAB6B, member RAS oncogene family | 270192 | 9 | 103111787 | 103185276 | 1 | Mm.490252 |
| 5.89 | 6.3 | 5.63 | 6.56 | DN3ab | DN3cd | 0.65 | 4.86 | 0.00094 | 0.018 | 0.04 | ENSMUSG00000030557 | protein_coding | Mef2a | myocyte enhancer factor 2A | 17258 | 7 | 67231163 | 67372858 | -1 | Mm.132788 Mm.426509 Mm.491250 |
| 4.87 | 5.21 | 4.58 | 5.3 | DN3ab | DN3cd | 0.51 | 4.83 | 0.00097 | 0.018 | 0.049 | ENSMUSG00000027459 | protein_coding | Fam110a | family with sequence similarity 110, member A | 73847 | 2 | 151969398 | 151980219 | -1 | Mm.272444 Mm.404253 |
| 10.32 | 10.56 | 10.06 | 10.58 | DN3ab | DN3cd | 0.36 | 4.83 | 0.00097 | 0.018 | 0.052 | ENSMUSG00000022205 | protein_coding | Sub1 | SUB1 homolog (S. cerevisiae) | 20024 | 15 | 11981339 | 11996983 | -1 | Mm.41746 |
| 10.73 | 11.03 | 10.36 | 10.48 | DN3ab | DN3cd | 0.22 | 4.83 | 0.00098 | 0.018 | 0.031 | ENSMUSG00000026234 | protein_coding | Ncl | nucleolin | 17975 | 1 | 86344719 | 86359400 | -1 | Mm.154378 Mm.474153 |
| 4.61 | 3.61 | 5.16 | 4.71 | DN3ab | DN3cd | -0.68 | -4.81 | 0.00099 | 0.018 | 0.035 | ENSMUSG00000030047 | protein_coding | Arhgap25 | Rho GTPase activating protein 25 | 232201 | 6 | 87458545 | 87533259 | -1 | Mm.119564 |
| 4.33 | 4.88 | 3.41 | 4.63 | DN3ab | DN3cd | 0.79 | 4.79 | 0.001 | 0.019 | 0.033 | ENSMUSG00000087177 | antisense | E130307A14Rik | RIKEN cDNA E130307A14 gene | NA | 10 | 39621412 | 39732007 | -1 | |
| 3.04 | 3.81 | 1.36 | 2.66 | DN3ab | DN3cd | 0.9 | 4.78 | 0.001 | 0.019 | 0.21 | ENSMUSG00000047003 | protein_coding | Zfp41 | zinc finger protein 41 | 22701 | 15 | 75616679 | 75629827 | 1 | Mm.87210 |
| 4.01 | 4.74 | 2.89 | 4.19 | DN3ab | DN3cd | 0.92 | 4.74 | 0.0011 | 0.02 | 0.16 | ENSMUSG00000056665 | protein_coding | Them6 | thioesterase superfamily member 6 | 223626 | 15 | 74721204 | 74724639 | 1 | Mm.268603 |
| 7.76 | 7.57 | 8.09 | 7.67 | DN3ab | DN3cd | -0.3 | -4.74 | 0.0011 | 0.02 | 0.044 | ENSMUSG00000057058 | protein_coding | Skap1 | src family associated phosphoprotein 1 | 78473 | 11 | 96464593 | 96759130 | 1 | Mm.395840 Mm.446811 |
| 6.26 | 6.63 | 5.57 | 6.34 | DN3ab | DN3cd | 0.54 | 4.73 | 0.0011 | 0.02 | 0.078 | ENSMUSG00000032264 | protein_coding | Zw10 | zw10 kinetochore protein | 26951 | 9 | 49055613 | 49078775 | 1 | Mm.24791 |
| 2.43 | 3.32 | 1.73 | 3.66 | DN3ab | DN3cd | 1.3 | 4.7 | 0.0012 | 0.02 | 0.049 | ENSMUSG00000038319 | protein_coding | Kcnh2 | potassium voltage-gated channel, subfamily H (eag-related), member 2 | 16511 | 5 | 24319589 | 24351604 | -1 | Mm.6539 |
| 4.22 | 3.88 | 4.79 | 4 | DN3ab | DN3cd | -0.57 | -4.68 | 0.0012 | 0.021 | 0.037 | ENSMUSG00000037750 | protein_coding | Fam222b | family with sequence similarity 222, member B | 216971 | 11 | 78094660 | 78156700 | 1 | Mm.119387 |
| 5.56 | 6.1 | 5.04 | 6.2 | DN3ab | DN3cd | 0.8 | 4.65 | 0.0013 | 0.021 | 0.062 | ENSMUSG00000024795 | protein_coding | Kif20b | kinesin family member 20B | 240641 | 19 | 34922361 | 34975745 | 1 | Mm.278471 |
| 5.01 | 5.43 | 4.04 | 4.93 | DN3ab | DN3cd | 0.59 | 4.63 | 0.0013 | 0.022 | 0.064 | ENSMUSG00000034311 | protein_coding | Kif4 | kinesin family member 4 | 16571 | X | 100625737 | 100727214 | 1 | Mm.383091 |
| 5.38 | 5.82 | 4.2 | 5.2 | DN3ab | DN3cd | 0.64 | 4.61 | 0.0013 | 0.022 | 0.036 | ENSMUSG00000002055 | protein_coding | Spag5 | sperm associated antigen 5 | 54141 | 11 | 78301529 | 78322457 | 1 | Mm.24250 |
| 5.49 | 5.93 | 4.52 | 5.46 | DN3ab | DN3cd | 0.63 | 4.59 | 0.0014 | 0.022 | 0.065 | ENSMUSG00000073705 | protein_coding | Cenps | centromere protein S | 69928 | 4 | 149127121 | 149137629 | -1 | Mm.260893 |
| 7.65 | 8.09 | 6.94 | 7.12 | DN3ab | DN3cd | 0.34 | 4.59 | 0.0014 | 0.022 | 0.056 | ENSMUSG00000030357 | protein_coding | Fkbp4 | FK506 binding protein 4 | 14228 | 6 | 128429735 | 128438677 | -1 | Mm.12758 |
| 0.35 | 1.79 | -1.76 | 1.42 | DN3ab | DN3cd | 1.98 | 4.58 | 0.0014 | 0.023 | 0.042 | ENSMUSG00000104639 | sense_intronic | Gm42939 | predicted gene 42939 | NA | 3 | 101407793 | 101409997 | 1 | |
| 6.41 | 6.66 | 5.77 | 6.29 | DN3ab | DN3cd | 0.36 | 4.55 | 0.0014 | 0.023 | 0.074 | ENSMUSG00000058440 | protein_coding | Nrf1 | nuclear respiratory factor 1 | 18181 | 6 | 30047988 | 30153458 | 1 | Mm.259258 |
| 7.43 | 7.65 | 6.6 | 7.11 | DN3ab | DN3cd | 0.34 | 4.54 | 0.0015 | 0.024 | 0.046 | ENSMUSG00000024151 | protein_coding | Msh2 | mutS homolog 2 | 17685 | 17 | 87672330 | 87723713 | 1 | Mm.4619 |
| 3.11 | 2.58 | 3.76 | 2.46 | DN3ab | DN3cd | -0.9 | -4.45 | 0.0017 | 0.025 | 0.036 | ENSMUSG00000062373 | protein_coding | Tmem65 | transmembrane protein 65 | 74868 | 15 | 58782269 | 58823638 | -1 | Mm.293754 |
| 4.51 | 5.12 | 3.12 | 4.32 | DN3ab | DN3cd | 0.79 | 4.44 | 0.0017 | 0.026 | 0.12 | ENSMUSG00000034023 | protein_coding | Fancd2 | Fanconi anemia, complementation group D2 | 211651 | 6 | 113531682 | 113597017 | 1 | Mm.160061 |
| 1.37 | 2.76 | -0.61 | 2.59 | DN3ab | DN3cd | 1.87 | 4.31 | 0.002 | 0.029 | 0.05 | ENSMUSG00000064080 | protein_coding | Fbln2 | fibulin 2 | 14115 | 6 | 91212455 | 91272540 | 1 | Mm.249146 Mm.410235 |
| 2.21 | -0.08 | -0.56 | -1.06 | DN3ab | DN3cd | -1.79 | -4.3 | 0.002 | 0.029 | 0.031 | ENSMUSG00000077391 | snoRNA | Gm24336 | predicted gene, 24336 | NA | 19 | 44113979 | 44114124 | -1 | |
| 1.18 | 2.57 | -0.73 | 2.09 | DN3ab | DN3cd | 1.77 | 4.27 | 0.0021 | 0.03 | 0.12 | ENSMUSG00000074037 | protein_coding | Mc1r | melanocortin 1 receptor | 17199 | 8 | 123407107 | 123410744 | 1 | Mm.324942 |
| 0.43 | 2.14 | -1.51 | 2.73 | DN3ab | DN3cd | 2.53 | 4.23 | 0.0023 | 0.031 | 0.032 | ENSMUSG00000021263 | protein_coding | Degs2 | delta(4)-desaturase, sphingolipid 2 | 70059 | 12 | 108678711 | 108702315 | -1 | Mm.207605 |
| 2.02 | 2.72 | 0.85 | 2.51 | DN3ab | DN3cd | 1.02 | 4.18 | 0.0025 | 0.032 | 0.04 | ENSMUSG00000104164 | TEC | Gm38248 | predicted gene, 38248 | NA | 1 | 127978905 | 127984687 | -1 | |
| 6.14 | 6.37 | 5.4 | 5.98 | DN3ab | DN3cd | 0.37 | 3.99 | 0.0033 | 0.038 | 0.038 | ENSMUSG00000019738 | protein_coding | Polr2i | polymerase (RNA) II (DNA directed) polypeptide I | 69920 | 7 | 30231948 | 30233390 | 1 | Mm.440028 |
| 3.53 | 4.06 | 2.2 | 3.54 | DN3ab | DN3cd | 0.78 | 3.86 | 0.0039 | 0.043 | 0.042 | ENSMUSG00000078773 | protein_coding | Rad54b | RAD54 homolog B (S. cerevisiae) | 623474 | 4 | 11558922 | 11615805 | 1 | Mm.491200 |
| 4.31 | 6.48 | 7.6 | 7.45 | DN3ab_ltr | DN3cd | 0.58 | 1.59 | 0.15 | 0.4 | 5.40E-08 | ENSMUSG00000023067 | protein_coding | Cdkn1a | cyclin-dependent kinase inhibitor 1A (P21) | 12575 | 17 | 29090976 | 29100727 | 1 | Mm.195663 |
| 9.73 | 9.52 | 11.11 | 9.66 | DN3ab_ltr | DN3cd | -0.87 | -4.09 | 0.0028 | 0.035 | 8.80E-08 | ENSMUSG00000027763 | protein_coding | Mbnl1 | muscleblind like splicing factor 1 | 56758 | 3 | 60472830 | 60629750 | 1 | Mm.255723 |
| 7.96 | 7.78 | 6.45 | 8.23 | DN3ab_ltr | DN3cd | 0.61 | 1.85 | 0.098 | 0.33 | 1.80E-07 | ENSMUSG00000066892 | protein_coding | Fbxl12 | F-box and leucine-rich repeat protein 12 | 30843 | 9 | 20618118 | 20644767 | -1 | Mm.24608 Mm.390165 Mm.489685 |
| 9.51 | 8.96 | 8.71 | 9.48 | DN3ab_ltr | DN3cd | 0.04 | 0.17 | 0.87 | 0.94 | 8.50E-07 | ENSMUSG00000022475 | protein_coding | Hdac7 | histone deacetylase 7 | 56233 | 15 | 97792664 | 97844502 | -1 | Mm.384027 |
| 7.79 | 8.04 | 6.43 | 7.93 | DN3ab_ltr | DN3cd | 0.74 | 3.55 | 0.0064 | 0.059 | 9.60E-07 | ENSMUSG00000026605 | protein_coding | Cenpf | centromere protein F | 108000 | 1 | 189640606 | 189688086 | -1 | Mm.129746 |
| 6.2 | 5.27 | 4.83 | 7.02 | DN3ab_ltr | DN3cd | 0.46 | 0.86 | 0.41 | 0.7 | 9.60E-07 | ENSMUSG00000029530 | protein_coding | Ccr9 | chemokine (C-C motif) receptor 9 | 12769 | 9 | 123678439 | 123784330 | 1 | Mm.440604 Mm.442383 |
| 10.74 | 10.45 | 11.75 | 10.18 | DN3ab_ltr | DN3cd | -0.92 | -4.18 | 0.0025 | 0.032 | 1.00E-06 | ENSMUSG00000062825 | protein_coding | Actg1 | actin, gamma, cytoplasmic 1 | 11465 | 11 | 120345690 | 120348542 | -1 | Mm.196173 Mm.426706 Mm.470166 |
| 10.05 | 10.08 | 9.22 | 10.17 | DN3ab_ltr | DN3cd | 0.42 | 2.67 | 0.026 | 0.15 | 1.10E-06 | ENSMUSG00000041126 | protein_coding | H2afv | H2A histone family, member V | 77605 | 11 | 6427229 | 6444443 | -1 | Mm.27624 |
| 10.06 | 9.54 | 9.39 | 9.85 | DN3ab_ltr | DN3cd | -0.08 | -0.49 | 0.64 | 0.85 | 1.40E-06 | ENSMUSG00000069662 | protein_coding | Marcks | myristoylated alanine rich protein kinase C substrate | 17118 | 10 | 37133375 | 37138920 | -1 | Mm.30059 |
| 6.57 | 6.38 | 5.02 | 6.89 | DN3ab_ltr | DN3cd | 0.61 | 1.75 | 0.11 | 0.36 | 1.60E-06 | ENSMUSG00000015314 | protein_coding | Slamf6 | SLAM family member 6 | 30925 | 1 | 171917515 | 171953170 | 1 | Mm.486404 |
| 5.27 | 6.7 | 8.29 | 7.67 | DN3ab_ltr | DN3cd | 0.1 | 0.31 | 0.76 | 0.92 | 2.40E-06 | ENSMUSG00000020644 | protein_coding | Id2 | inhibitor of DNA binding 2 | 15902 | 12 | 25093799 | 25097140 | -1 | Mm.34871 |
| 6.85 | 5.91 | 5.44 | 6.36 | DN3ab_ltr | DN3cd | -0.17 | -0.55 | 0.6 | 0.83 | 2.60E-06 | ENSMUSG00000027778 | protein_coding | Ift80 | intraflagellar transport 80 | 68259 | 3 | 68892499 | 69004570 | -1 | Mm.389451 |
| 4.97 | 5.7 | 1.88 | 6.49 | DN3ab_ltr | DN3cd | 1.44 | 2.78 | 0.022 | 0.13 | 2.60E-06 | ENSMUSG00000053977 | protein_coding | Cd8a | CD8 antigen, alpha chain | 12525 | 6 | 71373427 | 71379173 | 1 | Mm.1858 |
| 8.01 | 7.04 | 7.45 | 8.28 | DN3ab_ltr | DN3cd | -0.09 | -0.29 | 0.78 | 0.92 | 3.20E-06 | ENSMUSG00000037235 | protein_coding | Mxd4 | Max dimerization protein 4 | 17122 | 5 | 34173883 | 34187720 | -1 | Mm.259260 Mm.391777 |
| 5.24 | 4.94 | 3.42 | 5.47 | DN3ab_ltr | DN3cd | 0.5 | 1.29 | 0.23 | 0.52 | 3.50E-06 | ENSMUSG00000018427 | protein_coding | Ypel2 | yippee like 2 | 77864 | 11 | 86936425 | 86993707 | -1 | Mm.191894 |
| 6.71 | 6.12 | 6.46 | 7.27 | DN3ab_ltr | DN3cd | 0.11 | 0.45 | 0.66 | 0.87 | 3.50E-06 | ENSMUSG00000050921 | protein_coding | P2ry10 | purinergic receptor P2Y, G-protein coupled 10 | 78826 | X | 107088492 | 107104974 | 1 | Mm.74639 |
| 10.4 | 10.36 | 9.7 | 10.68 | DN3ab_ltr | DN3cd | 0.41 | 2.31 | 0.047 | 0.21 | 3.90E-06 | ENSMUSG00000048251 | protein_coding | Bcl11b | B cell leukemia/lymphoma 11B | 58208 | 12 | 107910403 | 108003602 | -1 | Mm.392694 |
| 7.75 | 7.93 | 8.4 | 7.57 | DN3ab_ltr | DN3cd | -0.3 | -1.72 | 0.12 | 0.37 | 4.50E-06 | ENSMUSG00000032966 | protein_coding | Fkbp1a | FK506 binding protein 1a | 14225 | 2 | 151542483 | 151561692 | 1 | Mm.278458 |
| 8.29 | 7.88 | 9.51 | 8.13 | DN3ab_ltr | DN3cd | -0.92 | -5.41 | 0.00045 | 0.011 | 4.60E-06 | ENSMUSG00000032231 | protein_coding | Anxa2 | annexin A2 | 12306 | 9 | 69453620 | 69491795 | 1 | Mm.238343 |
| 9.23 | 8.3 | 9.14 | 9.35 | DN3ab_ltr | DN3cd | -0.35 | -1.76 | 0.11 | 0.35 | 4.90E-06 | ENSMUSG00000037266 | protein_coding | Rsrp1 | arginine/serine rich protein 1 | 27981 | 4 | 134923592 | 134927671 | 1 | Mm.331964 |
| 5.01 | 4.86 | 7.3 | 5.25 | DN3ab_ltr | DN3cd | -1.26 | -3.85 | 0.004 | 0.044 | 5.00E-06 | ENSMUSG00000032290 | protein_coding | Ptpn9 | protein tyrosine phosphatase, non-receptor type 9 | 56294 | 9 | 56994923 | 57062807 | 1 | Mm.325643 |
| 10.57 | 10.69 | 9.44 | 10.54 | DN3ab_ltr | DN3cd | 0.51 | 3.07 | 0.014 | 0.098 | 6.20E-06 | ENSMUSG00000028832 | protein_coding | Stmn1 | stathmin 1 | 16765 | 4 | 134468320 | 134473843 | 1 | Mm.378957 |
| 9.03 | 9.03 | 8.49 | 9.54 | DN3ab_ltr | DN3cd | 0.47 | 2.58 | 0.03 | 0.16 | 6.40E-06 | ENSMUSG00000030287 | protein_coding | Itpr2 | inositol 1,4,5-triphosphate receptor 2 | 16439 | 6 | 146108299 | 146502223 | -1 | Mm.7800 |
| 8.32 | 8.94 | 7.28 | 9.01 | DN3ab_ltr | DN3cd | 1.07 | 5.62 | 0.00035 | 0.0094 | 7.00E-06 | ENSMUSG00000032228 | protein_coding | Tcf12 | transcription factor 12 | 21406 | 9 | 71842688 | 72111871 | -1 | Mm.171615 Mm.480117 |
| 6.63 | 7 | 4.74 | 7.48 | DN3ab_ltr | DN3cd | 1.2 | 3.04 | 0.014 | 0.1 | 7.10E-06 | ENSMUSG00000020388 | protein_coding | Pdlim4 | PDZ and LIM domain 4 | 30794 | 11 | 54054928 | 54069014 | -1 | Mm.21830 |
| 4.22 | 5.38 | 1.49 | 5.66 | DN3ab_ltr | DN3cd | 1.73 | 4.17 | 0.0025 | 0.032 | 7.80E-06 | ENSMUSG00000015476 | protein_coding | Prrt1 | proline-rich transmembrane protein 1 | 260297 | 17 | 34629533 | 34633126 | 1 | Mm.390965 |
| 7.69 | 6.65 | 7.15 | 7.34 | DN3ab_ltr | DN3cd | -0.46 | -2.14 | 0.061 | 0.25 | 7.80E-06 | ENSMUSG00000038128 | protein_coding | Camk4 | calcium/calmodulin-dependent protein kinase IV | 12326 | 18 | 32934931 | 33191228 | 1 | Mm.222329 Mm.410854 |
| 6.39 | 6.79 | 6.86 | 6.01 | DN3ab_ltr | DN3cd | -0.17 | -0.76 | 0.47 | 0.74 | 8.20E-06 | ENSMUSG00000046229 | protein_coding | Scand1 | SCAN domain-containing 1 | 19018 | 2 | 156311847 | 156312704 | -1 | Mm.389346 |
| 1.22 | 0.82 | -3.72 | 1.78 | DN3ab_ltr | DN3cd | 1.47 | 1.53 | 0.16 | 0.43 | 8.80E-06 | ENSMUSG00000100552 | lincRNA | Gm29019 | predicted gene 29019 | NA | 15 | 82982725 | 82987830 | -1 | |
| 8.83 | 8.37 | 7.98 | 8.9 | DN3ab_ltr | DN3cd | 0.15 | 0.64 | 0.54 | 0.79 | 9.20E-06 | ENSMUSG00000076431 | protein_coding | Sox4 | SRY (sex determining region Y)-box 4 | 20677 | 13 | 28948919 | 28953713 | -1 | Mm.240627 Mm.455819 |
| 6.76 | 6.43 | 5.51 | 6.78 | DN3ab_ltr | DN3cd | 0.33 | 1.19 | 0.27 | 0.56 | 1.00E-05 | ENSMUSG00000026594 | protein_coding | Ralgps2 | Ral GEF with PH domain and SH3 binding motif 2 | 78255 | 1 | 156804166 | 156939626 | -1 | Mm.28376 Mm.396243 |
| 1.44 | 2.65 | -3.72 | 2.53 | DN3ab_ltr | DN3cd | 2.34 | 3.16 | 0.012 | 0.089 | 1.10E-05 | ENSMUSG00000044505 | protein_coding | Lingo4 | leucine rich repeat and Ig domain containing 4 | 320747 | 3 | 94398517 | 94404501 | 1 | Mm.213001 |
| 7.12 | 7.11 | 6.23 | 7.2 | DN3ab_ltr | DN3cd | 0.4 | 2.38 | 0.042 | 0.2 | 1.10E-05 | ENSMUSG00000003778 | protein_coding | Brd8 | bromodomain containing 8 | 78656 | 18 | 34598615 | 34624601 | -1 | Mm.45602 Mm.411740 |
| 6.09 | 5.94 | 3.75 | 6.38 | DN3ab_ltr | DN3cd | 0.7 | 1.56 | 0.15 | 0.42 | 1.10E-05 | ENSMUSG00000056531 | protein_coding | Ccdc18 | coiled-coil domain containing 18 | 73254 | 5 | 108132875 | 108233628 | 1 | Mm.441231 |
| 7.14 | 7.16 | 8.52 | 7.23 | DN3ab_ltr | DN3cd | -0.66 | -2.87 | 0.019 | 0.12 | 1.10E-05 | ENSMUSG00000029076 | protein_coding | Sdf4 | stromal cell derived factor 4 | 20318 | 4 | 155992872 | 156013610 | 1 | Mm.293517 |
| 3.24 | 1.54 | 1.45 | 4.24 | DN3ab_ltr | DN3cd | 0.33 | 0.42 | 0.68 | 0.88 | 1.10E-05 | ENSMUSG00000037224 | protein_coding | Zfyve28 | zinc finger, FYVE domain containing 28 | 231125 | 5 | 34194893 | 34288449 | -1 | Mm.113388 |
| 8.76 | 8.59 | 9.68 | 8.5 | DN3ab_ltr | DN3cd | -0.68 | -3.83 | 0.0042 | 0.045 | 1.20E-05 | ENSMUSG00000090841 | protein_coding | Myl6 | myosin, light polypeptide 6, alkali, smooth muscle and non-muscle | 17904 | 10 | 128490860 | 128494145 | -1 | Mm.337074 |
| 9.47 | 9.6 | 8.54 | 9.57 | DN3ab_ltr | DN3cd | 0.5 | 3.26 | 0.01 | 0.081 | 1.20E-05 | ENSMUSG00000000409 | protein_coding | Lck | lymphocyte protein tyrosine kinase | 16818 | 4 | 129548344 | 129573641 | -1 | Mm.293753 |
| 4.65 | 3.67 | 3.52 | 4.69 | DN3ab_ltr | DN3cd | -0.03 | -0.08 | 0.94 | 0.97 | 1.20E-05 | ENSMUSG00000104467 | TEC | Gm37660 | predicted gene, 37660 | NA | 9 | 71840005 | 71841736 | -1 | |
| 0.69 | 2 | -3.72 | 2.77 | DN3ab_ltr | DN3cd | 2.9 | 3.44 | 0.0075 | 0.067 | 1.40E-05 | ENSMUSG00000039109 | protein_coding | F13a1 | coagulation factor XIII, A1 subunit | 74145 | 13 | 36867178 | 37050244 | -1 | Mm.235105 |
| 0.16 | -0.37 | -3.72 | 1.74 | DN3ab_ltr | DN3cd | 2.21 | 2.09 | 0.067 | 0.26 | 1.40E-05 | ENSMUSG00000108985 | TEC | A930030B08Rik | RIKEN cDNA A930030B08 gene | NA | 7 | 99900192 | 99904156 | -1 | |
| 4.09 | 3.14 | 2.35 | 3.73 | DN3ab_ltr | DN3cd | -0.12 | -0.3 | 0.77 | 0.92 | 1.50E-05 | ENSMUSG00000079110 | protein_coding | Capn3 | calpain 3 | 12335 | 2 | 120456019 | 120504913 | 1 | Mm.458021 Mm.485295 |
| 7.69 | 7.33 | 7.5 | 8.21 | DN3ab_ltr | DN3cd | 0.16 | 0.87 | 0.41 | 0.7 | 1.60E-05 | ENSMUSG00000028248 | protein_coding | Pnisr | PNN interacting serine/arginine-rich | 66625 | 4 | 21847583 | 21876475 | 1 | Mm.100117 |
| 8.55 | 8.9 | 7.27 | 8.7 | DN3ab_ltr | DN3cd | 0.77 | 4.11 | 0.0027 | 0.034 | 1.60E-05 | ENSMUSG00000027306 | protein_coding | Nusap1 | nucleolar and spindle associated protein 1 | 108907 | 2 | 119618298 | 119651244 | 1 | Mm.290015 Mm.439488 |
| 4.88 | 4.76 | 7.71 | 5.59 | DN3ab_ltr | DN3cd | -1.37 | -4.03 | 0.003 | 0.037 | 1.60E-05 | ENSMUSG00000021831 | protein_coding | Ero1l | ERO1-like (S. cerevisiae) | 50527 | 14 | 45283087 | 45318771 | -1 | Mm.387108 |
| 5.38 | 5.77 | 7.1 | 6.09 | DN3ab_ltr | DN3cd | -0.39 | -1.58 | 0.15 | 0.41 | 1.60E-05 | ENSMUSG00000021196 | protein_coding | Pfkp | phosphofructokinase, platelet | 56421 | 13 | 6579768 | 6648777 | -1 | Mm.273874 |
| 5.7 | 5.91 | 3.6 | 5.93 | DN3ab_ltr | DN3cd | 0.84 | 2.46 | 0.036 | 0.18 | 1.60E-05 | ENSMUSG00000027220 | protein_coding | Syt13 | synaptotagmin XIII | 80976 | 2 | 92915098 | 92956058 | 1 | Mm.33490 |
| 9.63 | 9.29 | 8.53 | 9.49 | DN3ab_ltr | DN3cd | 0.2 | 0.88 | 0.4 | 0.69 | 1.60E-05 | ENSMUSG00000034349 | protein_coding | Smc4 | structural maintenance of chromosomes 4 | 70099 | 3 | 69004738 | 69034623 | 1 | Mm.206841 |
| 4.59 | 3.25 | 4.11 | 4.98 | DN3ab_ltr | DN3cd | -0.14 | -0.35 | 0.73 | 0.91 | 1.70E-05 | ENSMUSG00000001986 | protein_coding | Gria3 | glutamate receptor, ionotropic, AMPA3 (alpha 3) | 53623 | X | 41400854 | 41678601 | 1 | Mm.327681 |
| 6.07 | 5.22 | 8.46 | 6.02 | DN3ab_ltr | DN3cd | -1.79 | -6.53 | 0.00012 | 0.0049 | 1.70E-05 | ENSMUSG00000020077 | protein_coding | Srgn | serglycin | 19073 | 10 | 62493833 | 62527451 | -1 | Mm.338790 |
| 6.16 | 5.26 | 6.09 | 6.71 | DN3ab_ltr | DN3cd | -0.1 | -0.36 | 0.73 | 0.91 | 1.70E-05 | ENSMUSG00000090019 | protein_coding | Gimap1 | GTPase, IMAP family member 1 | 16205 | 6 | 48739054 | 48743795 | 1 | Mm.252599 |
| 6.46 | 6.28 | 5.87 | 6.87 | DN3ab_ltr | DN3cd | 0.35 | 1.68 | 0.13 | 0.38 | 1.80E-05 | ENSMUSG00000037111 | protein_coding | Setd7 | SET domain containing (lysine methyltransferase) 7 | 73251 | 3 | 51515319 | 51560879 | -1 | Mm.192111 |
| 3.65 | 2.79 | 2.9 | 5.39 | DN3ab_ltr | DN3cd | 0.87 | 1.47 | 0.18 | 0.45 | 1.80E-05 | ENSMUSG00000079071 | protein_coding | Gm14085 | predicted gene 14085 | 381417 | 2 | 122484941 | 122528040 | 1 | Mm.482295 |
| 9.89 | 9.75 | 8.45 | 9.7 | DN3ab_ltr | DN3cd | 0.4 | 1.69 | 0.13 | 0.38 | 2.10E-05 | ENSMUSG00000000782 | protein_coding | Tcf7 | transcription factor 7, T cell specific | 21414 | 11 | 52252371 | 52283014 | -1 | Mm.31630 |
| 9.57 | 9.53 | 8.99 | 9.74 | DN3ab_ltr | DN3cd | 0.31 | 2.23 | 0.053 | 0.23 | 2.10E-05 | ENSMUSG00000018379 | protein_coding | Srsf1 | serine/arginine-rich splicing factor 1 | 110809 | 11 | 88047373 | 88053755 | 1 | Mm.391719 |
| 6.4 | 6.97 | 4.45 | 6.7 | DN3ab_ltr | DN3cd | 1.11 | 4 | 0.0032 | 0.038 | 2.20E-05 | ENSMUSG00000022471 | protein_coding | Xrcc6 | X-ray repair complementing defective repair in Chinese hamster cells 6 | 14375 | 15 | 81987835 | 82040085 | 1 | Mm.288809 |
| 7.36 | 7.53 | 7.88 | 7.16 | DN3ab_ltr | DN3cd | -0.24 | -1.54 | 0.16 | 0.42 | 2.20E-05 | ENSMUSG00000025132 | protein_coding | Arhgdia | Rho GDP dissociation inhibitor (GDI) alpha | 192662 | 11 | 120578104 | 120581624 | -1 | Mm.30016 Mm.456388 Mm.474783 |
| 5.87 | 5.58 | 4.22 | 6.18 | DN3ab_ltr | DN3cd | 0.54 | 1.41 | 0.19 | 0.47 | 2.30E-05 | ENSMUSG00000003992 | protein_coding | Ssbp2 | single-stranded DNA binding protein 2 | 66970 | 13 | 91460283 | 91703429 | 1 | Mm.343095 |
| 10.32 | 10.53 | 9.38 | 10.27 | DN3ab_ltr | DN3cd | 0.48 | 4.12 | 0.0027 | 0.034 | 2.30E-05 | ENSMUSG00000026434 | protein_coding | Nucks1 | nuclear casein kinase and cyclin-dependent kinase substrate 1 | 98415 | 1 | 131910534 | 131936321 | 1 | Mm.246869 Mm.392848 |
| 6.33 | 6.55 | 3.73 | 6.74 | DN3ab_ltr | DN3cd | 1 | 2.27 | 0.05 | 0.22 | 2.50E-05 | ENSMUSG00000042797 | protein_coding | Aqp11 | aquaporin 11 | 66333 | 7 | 97724006 | 97738289 | -1 | Mm.29756 |
| 9.09 | 8.67 | 7.96 | 8.87 | DN3ab_ltr | DN3cd | 0.13 | 0.57 | 0.58 | 0.82 | 2.50E-05 | ENSMUSG00000029705 | protein_coding | Cux1 | cut-like homeobox 1 | 13047 | 5 | 136248135 | 136567490 | -1 | Mm.320317 Mm.436792 |
| 3.03 | 3.27 | 1 | 4.19 | DN3ab_ltr | DN3cd | 1.06 | 2.25 | 0.052 | 0.22 | 2.70E-05 | ENSMUSG00000050179 | lincRNA | A930002I21Rik | RIKEN cDNA A930002I21 gene | NA | 3 | 107090880 | 107098401 | 1 | |
| 9.41 | 9.69 | 8.21 | 9.28 | DN3ab_ltr | DN3cd | 0.59 | 4.23 | 0.0023 | 0.031 | 2.80E-05 | ENSMUSG00000017716 | protein_coding | Birc5 | baculoviral IAP repeat-containing 5 | 11799 | 11 | 117849251 | 117855743 | 1 | Mm.8552 |
| 5.79 | 4.47 | 4.71 | 5.36 | DN3ab_ltr | DN3cd | -0.43 | -1.21 | 0.26 | 0.55 | 3.00E-05 | ENSMUSG00000025245 | protein_coding | Lztfl1 | leucine zipper transcription factor-like 1 | 93730 | 9 | 123694416 | 123717697 | -1 | Mm.378263 Mm.475504 |
| 4.93 | 6.1 | 2.04 | 6.26 | DN3ab_ltr | DN3cd | 1.74 | 4.1 | 0.0027 | 0.034 | 3.10E-05 | ENSMUSG00000038587 | protein_coding | Akap12 | A kinase (PRKA) anchor protein (gravin) 12 | 83397 | 10 | 4266380 | 4359470 | 1 | Mm.27481 |
| 5.26 | 4.59 | 4.81 | 5.7 | DN3ab_ltr | DN3cd | 0.09 | 0.33 | 0.75 | 0.92 | 3.10E-05 | ENSMUSG00000035126 | protein_coding | Wdr78 | WD repeat domain 78 | 242584 | 4 | 103038065 | 103114555 | -1 | Mm.374877 |
| 5.25 | 4.14 | 5.09 | 5.51 | DN3ab_ltr | DN3cd | -0.29 | -1.07 | 0.31 | 0.61 | 3.20E-05 | ENSMUSG00000031166 | protein_coding | Wdr13 | WD repeat domain 13 | 73447 | X | 8123301 | 8132892 | -1 | Mm.284162 |
| 6.31 | 5.97 | 5.46 | 6.38 | DN3ab_ltr | DN3cd | 0.2 | 0.92 | 0.38 | 0.68 | 3.20E-05 | ENSMUSG00000022191 | protein_coding | Drosha | drosha, ribonuclease type III | 14000 | 15 | 12824815 | 12935291 | 1 | Mm.293142 |
| 7.11 | 7.24 | 5.76 | 7.07 | DN3ab_ltr | DN3cd | 0.59 | 2.89 | 0.018 | 0.12 | 3.40E-05 | ENSMUSG00000027160 | protein_coding | Ccdc34 | coiled-coil domain containing 34 | 68201 | 2 | 110017817 | 110173360 | 1 | Mm.181767 |
| 8.75 | 8.69 | 9.54 | 8.68 | DN3ab_ltr | DN3cd | -0.46 | -3.26 | 0.01 | 0.081 | 3.40E-05 | ENSMUSG00000006304 | protein_coding | Arpc2 | actin related protein 2/3 complex, subunit 2 | 76709 | 1 | 74236084 | 74268209 | 1 | Mm.337038 |
| 6.57 | 6.74 | 5.57 | 7.26 | DN3ab_ltr | DN3cd | 0.81 | 3.01 | 0.015 | 0.1 | 3.50E-05 | ENSMUSG00000022469 | protein_coding | Rapgef3 | Rap guanine nucleotide exchange factor (GEF) 3 | 223864 | 15 | 97744770 | 97767972 | -1 | Mm.24028 |

| | | | | | | | | | | | | | | | | | | | | |
|---|---|---|---|---|---|---|---|---|---|---|---|---|---|---|---|---|---|---|---|---|
| 6.75 | 5.64 | 6.49 | 7.09 | DN3ab_ltr | DN3cd | -0.22 | -0.73 | 0.48 | 0.75 | 3.60E-05 | ENSMUSG00000034614 | protein_coding | Pik3ip1 | phosphoinositide-3-kinase interacting protein 1 | 216505 | 11 | 3330401 | 3342971 | 1 | Mm.390323 |
| 5.18 | 5.81 | 6.7 | 5.96 | DN3ab_ltr | DN3cd | -0.12 | -0.48 | 0.64 | 0.85 | 3.60E-05 | ENSMUSG00000025791 | protein_coding | Pgm1 | phosphoglucomutase 1 | 72157 | 4 | 99929414 | 99987294 | 1 | Mm.217764 |
| 1.54 | 2.85 | -3.72 | 3 | DN3ab_ltr | DN3cd | 2.47 | 3.11 | 0.013 | 0.094 | 3.60E-05 | ENSMUSG00000038390 | protein_coding | Gpr162 | G protein-coupled receptor 162 | 14788 | 6 | 124858444 | 124863983 | -1 | Mm.2514 |
| 2 | 2.68 | -1.4 | 3.7 | DN3ab_ltr | DN3cd | 1.55 | 2.46 | 0.037 | 0.18 | 3.70E-05 | ENSMUSG00000048148 | protein_coding | Nwd1 | NACHT and WD repeat domain containing 1 | 319555 | 8 | 72646711 | 72717876 | 1 | Mm.390076 |
| 5.03 | 4.78 | 6.49 | 5.01 | DN3ab_ltr | DN3cd | -0.92 | -4.16 | 0.0025 | 0.032 | 3.70E-05 | ENSMUSG00000026288 | protein_coding | Inpp5d | inositol polyphosphate-5-phosphatase D | 16331 | 1 | 87620312 | 87720507 | 1 | Mm.15105 |
| 4.92 | 3.68 | 5.59 | 5.97 | DN3ab_ltr | DN3cd | -0.21 | -0.76 | 0.47 | 0.74 | 3.90E-05 | ENSMUSG00000093661 | protein_coding | Eif4e3 | eukaryotic translation initiation factor 4E member 3 | 66892 | 6 | 99625135 | 99666771 | -1 | Mm.32889 |
| 6.12 | 5.45 | 5.6 | 6.33 | DN3ab_ltr | DN3cd | -0.01 | -0.02 | 0.98 | 0.99 | 3.90E-05 | ENSMUSG00000040746 | protein_coding | Rnf167 | ring finger protein 167 | 70510 | 11 | 70647235 | 70651421 | 1 | Mm.261818 |
| 6.27 | 6.1 | 5.71 | 6.83 | DN3ab_ltr | DN3cd | 0.42 | 1.82 | 0.1 | 0.34 | 4.10E-05 | ENSMUSG00000029313 | protein_coding | Aff1 | AF4/FMR2 family, member 1 | 17355 | 5 | 103692374 | 103855322 | 1 | Mm.6949 |
| 6.7 | 6.6 | 5.78 | 6.83 | DN3ab_ltr | DN3cd | 0.39 | 1.89 | 0.092 | 0.31 | 4.20E-05 | ENSMUSG00000053477 | protein_coding | Tcf4 | transcription factor 4 | 21413 | 18 | 69343356 | 69689079 | 1 | Mm.4269 |
| 9.21 | 9.5 | 10.31 | 9.65 | DN3ab_ltr | DN3cd | -0.21 | -1.24 | 0.25 | 0.54 | 4.20E-05 | ENSMUSG00000063229 | protein_coding | Ldha | lactate dehydrogenase A | 16828 | 7 | 46841475 | 46855627 | 1 | Mm.29324 |
| 7.68 | 7.41 | 7.13 | 7.63 | DN3ab_ltr | DN3cd | 0.07 | 0.52 | 0.62 | 0.84 | 4.20E-05 | ENSMUSG00000019978 | protein_coding | Epb41l2 | erythrocyte membrane protein band 4.1 like 2 | 13822 | 10 | 25359798 | 25523519 | 1 | Mm.306026 |
| 4.84 | 4.97 | 2.55 | 5.26 | DN3ab_ltr | DN3cd | 0.82 | 2.01 | 0.076 | 0.28 | 4.30E-05 | ENSMUSG00000036158 | protein_coding | Prickle1 | prickle planar cell polarity protein 1 | 106042 | 15 | 93499114 | 93595891 | -1 | Mm.150314 |
| 3.75 | 3.67 | 1.88 | 4.12 | DN3ab_ltr | DN3cd | 0.61 | 1.6 | 0.15 | 0.4 | 4.30E-05 | ENSMUSG00000076882 | TR_J_gene | Traj49 | T cell receptor alpha joining 49 | NA | 14 | 54168686 | 54168747 | 1 | |
| 6.67 | 6.8 | 5.29 | 6.7 | DN3ab_ltr | DN3cd | 0.62 | 2.78 | 0.022 | 0.13 | 4.40E-05 | ENSMUSG00000033952 | protein_coding | Aspm | abnormal spindle microtubule assembly | 12316 | 1 | 139454772 | 139494091 | 1 | Mm.168523 |
| 7.01 | 7.44 | 5.45 | 7.03 | DN3ab_ltr | DN3cd | 0.84 | 4.26 | 0.0022 | 0.03 | 4.40E-05 | ENSMUSG00000036223 | protein_coding | Ska1 | spindle and kinetochore associated complex subunit 1 | 66468 | 18 | 74195299 | 74207818 | -1 | Mm.283406 |
| 5.95 | 5.9 | 4.9 | 6.44 | DN3ab_ltr | DN3cd | 0.61 | 2.18 | 0.058 | 0.24 | 4.40E-05 | ENSMUSG00000105519 | TEC | E430021H15Rik | RIKEN cDNA E430021H15 gene | NA | 5 | 44213801 | 44217259 | -1 | |
| 3.93 | 4.74 | 4.31 | 6.98 | DN3ab_ltr | DN3cd | 1.84 | 5.54 | 0.00038 | 0.01 | 4.50E-05 | ENSMUSG00000065911 | misc_RNA | Gm24447 | predicted gene, 24447 | NA | 15 | 93234363 | 93234700 | 1 | |
| 8.59 | 8.18 | 8.64 | 8.9 | DN3ab_ltr | DN3cd | -0.08 | -0.64 | 0.54 | 0.8 | 4.70E-05 | ENSMUSG00000030067 | protein_coding | Foxp1 | forkhead box P1 | 108655 | 6 | 98925338 | 99522721 | -1 | Mm.234965 Mm.392884 Mm.485266 Mm.487716 |
| 5.13 | 4.07 | 4.51 | 5.2 | DN3ab_ltr | DN3cd | -0.19 | -0.62 | 0.55 | 0.8 | 4.70E-05 | ENSMUSG00000027589 | protein_coding | Pcmtd2 | protein-L-isoaspartate (D-aspartate) O-methyltransferase domain containing 2 | 245867 | 2 | 181837854 | 181857461 | 1 | Mm.491085 |
| 5.2 | 3.21 | 3.19 | 3.34 | DN3ab_ltr | DN3cd | -1.2 | -3.24 | 0.01 | 0.082 | 5.00E-05 | ENSMUSG00000040536 | protein_coding | Necab1 | N-terminal EF-hand calcium binding protein 1 | 69352 | 4 | 14952245 | 15149794 | -1 | Mm.389031 |
| 7.22 | 6.82 | 6.57 | 7.61 | DN3ab_ltr | DN3cd | 0.26 | 1.01 | 0.34 | 0.63 | 5.10E-05 | ENSMUSG00000102744 | TEC | 5830444F18Rik | RIKEN cDNA 5830444F18 gene | NA | 17 | 51791470 | 51793461 | -1 | |
| 8.99 | 8.72 | 7.66 | 8.92 | DN3ab_ltr | DN3cd | 0.35 | 1.3 | 0.23 | 0.51 | 5.20E-05 | ENSMUSG00000029267 | protein_coding | Mtf2 | metal response element binding transcription factor 2 | 17765 | 5 | 108065674 | 108109004 | 1 | Mm.257149 |
| 2.5 | 2.19 | 8.07 | 5.24 | DN3ab_ltr | DN3cd | -2.48 | -7.92 | 2.60E-05 | 0.0021 | 5.30E-05 | ENSMUSG00000028965 | protein_coding | Tnfrsf9 | tumor necrosis factor receptor superfamily, member 9 | 21942 | 4 | 150914562 | 150946102 | 1 | Mm.244187 |
| 3.42 | 3.79 | 2.31 | 4.69 | DN3ab_ltr | DN3cd | 1.12 | 3.17 | 0.012 | 0.088 | 5.30E-05 | ENSMUSG00000076928 | TR_C_gene | Trac | T cell receptor alpha constant | NA | 14 | 54220521 | 54224206 | 1 | |
| 7.59 | 7.79 | 8.11 | 7.37 | DN3ab_ltr | DN3cd | -0.24 | -1.44 | 0.18 | 0.46 | 5.30E-05 | ENSMUSG00000005667 | protein_coding | Mthfd2 | methylenetetrahydrofolate dehydrogenase (NAD+ dependent), methenyltetrahydrofolate cyclohydrolase | 17768 | 6 | 83305691 | 83325908 | -1 | Mm.443 Mm.475021 |
| 10.24 | 9.09 | 9.38 | 9.64 | DN3ab_ltr | DN3cd | -0.51 | -2.03 | 0.073 | 0.27 | 5.40E-05 | ENSMUSG00000045136 | protein_coding | Tubb2b | tubulin, beta 2B class IIB | 73710 | 13 | 34126748 | 34130466 | -1 | Mm.379227 Mm.472121 |
| 6.64 | 6.25 | 8.29 | 6.45 | DN3ab_ltr | DN3cd | -1.17 | -4.49 | 0.0016 | 0.025 | 5.50E-05 | ENSMUSG00000036093 | protein_coding | Arl5a | ADP-ribosylation factor-like 5A | 75423 | 2 | 52397951 | 52424901 | -1 | Mm.314378 |
| 6.07 | 6.34 | 4.99 | 6.29 | DN3ab_ltr | DN3cd | 0.67 | 3.73 | 0.0048 | 0.049 | 5.60E-05 | ENSMUSG00000038379 | protein_coding | Ttk | Ttk protein kinase | 22137 | 9 | 83834689 | 83872389 | 1 | Mm.1904 |
| 5.07 | 4.46 | 4.26 | 5.1 | DN3ab_ltr | DN3cd | 0.03 | 0.1 | 0.92 | 0.96 | 5.70E-05 | ENSMUSG00000021514 | protein_coding | Zfp369 | zinc finger protein 369 | 170936 | 13 | 65278814 | 65304221 | 1 | Mm.151546 |
| 7.25 | 6.95 | 6.36 | 7.18 | DN3ab_ltr | DN3cd | 0.19 | 0.93 | 0.38 | 0.67 | 5.90E-05 | ENSMUSG00000029426 | protein_coding | Scarb2 | scavenger receptor class B, member 2 | 12492 | 5 | 92441314 | 92506833 | -1 | Mm.297964 |
| 1.57 | 1.36 | 0.98 | 4 | DN3ab_ltr | DN3cd | 1.41 | 2.42 | 0.039 | 0.19 | 6.00E-05 | ENSMUSG00000021565 | protein_coding | Slc6a19 | solute carrier family 6 (neurotransmitter transporter), member 19 | 74338 | 13 | 73679745 | 73704865 | -1 | Mm.271635 |
| 9.01 | 8.75 | 7.53 | 8.64 | DN3ab_ltr | DN3cd | 0.27 | 1.12 | 0.29 | 0.59 | 6.00E-05 | ENSMUSG00000015217 | protein_coding | Hmgb3 | high mobility group box 3 | 15354 | X | 71555918 | 71560676 | 1 | Mm.336087 |
| 5.56 | 4.98 | 5.39 | 6.14 | DN3ab_ltr | DN3cd | 0.09 | 0.39 | 0.71 | 0.89 | 6.00E-05 | ENSMUSG00000027219 | protein_coding | Slc28a2 | solute carrier family 28 (sodium-coupled nucleoside transporter), member 2 | 269346 | 2 | 122426477 | 122461137 | 1 | Mm.29510 |
| 6.73 | 6.72 | 6.44 | 7.52 | DN3ab_ltr | DN3cd | 0.51 | 2.58 | 0.03 | 0.16 | 6.00E-05 | ENSMUSG00000039852 | protein_coding | Rere | arginine glutamic acid dipeptide (RE) repeats | 68703 | 4 | 150281646 | 150621966 | 1 | Mm.291274 |
| 5.79 | 5.15 | 4.36 | 5.35 | DN3ab_ltr | DN3cd | -0.01 | -0.04 | 0.97 | 0.99 | 6.10E-05 | ENSMUSG00000027035 | protein_coding | Cers6 | ceramide synthase 6 | 241447 | 2 | 68861441 | 69114282 | 1 | Mm.222222 Mm.393211 |
| 3.44 | 3.67 | 0.43 | 3.42 | DN3ab_ltr | DN3cd | 0.74 | 1.91 | 0.09 | 0.31 | 6.20E-05 | ENSMUSG00000027075 | protein_coding | Slc43a1 | solute carrier family 43, member 1 | 72401 | 2 | 84838850 | 84863594 | 1 | Mm.22716 |
| 6.38 | 6.72 | 5.45 | 6.71 | DN3ab_ltr | DN3cd | 0.71 | 4.31 | 0.002 | 0.029 | 6.20E-05 | ENSMUSG00000026708 | protein_coding | Cenpl | centromere protein L | 70454 | 1 | 161070713 | 161086724 | 1 | Mm.248212 |
| 6.54 | 5.81 | 5.13 | 6.13 | DN3ab_ltr | DN3cd | -0.03 | -0.1 | 0.92 | 0.96 | 6.20E-05 | ENSMUSG00000022822 | protein_coding | Abcc5 | ATP-binding cassette, sub-family C (CFTR/MRP), member 5 | 27416 | 16 | 20331303 | 20426394 | -1 | Mm.20845 |
| 6.17 | 6.17 | 7.99 | 6.8 | DN3ab_ltr | DN3cd | -0.67 | -3.21 | 0.011 | 0.084 | 6.20E-05 | ENSMUSG00000056708 | protein_coding | Ier5 | immediate early response 5 | 15939 | 1 | 155096361 | 155099636 | -1 | Mm.12246 |
| 8.56 | 8.69 | 7.54 | 8.68 | DN3ab_ltr | DN3cd | 0.54 | 3.03 | 0.015 | 0.1 | 6.30E-05 | ENSMUSG00000021213 | protein_coding | Akr1c13 | aldo-keto reductase family 1, member C13 | 27384 | 13 | 4191150 | 4205596 | 1 | Mm.27447 |
| 7.56 | 7.57 | 6.83 | 7.62 | DN3ab_ltr | DN3cd | 0.35 | 2.48 | 0.036 | 0.18 | 6.30E-05 | ENSMUSG00000001151 | protein_coding | Pcnt | pericentrin (kendrin) | 18541 | 10 | 76351263 | 76442786 | -1 | Mm.251794 |
| 6.72 | 6.37 | 6.37 | 6.92 | DN3ab_ltr | DN3cd | 0.07 | 0.43 | 0.68 | 0.88 | 6.30E-05 | ENSMUSG00000046897 | protein_coding | Zfp740 | zinc finger protein 740 | 68744 | 15 | 102203249 | 102215606 | 1 | Mm.352945 |
| 9.67 | 9.36 | 9.16 | 9.63 | DN3ab_ltr | DN3cd | 0.04 | 0.29 | 0.78 | 0.92 | 6.40E-05 | ENSMUSG00000028759 | protein_coding | Hp1bp3 | heterochromatin protein 1, binding protein 3 | 15441 | 4 | 138216296 | 138244683 | 1 | Mm.348429 Mm.399941 |
| 6.8 | 6.21 | 6.81 | 7.18 | DN3ab_ltr | DN3cd | -0.1 | -0.6 | 0.56 | 0.8 | 6.40E-05 | ENSMUSG00000078671 | protein_coding | Chd2 | chromodomain helicase DNA binding protein 2 | 244059 | 7 | 73426638 | 73541830 | -1 | Mm.34539 Mm.34955 |
| 2.64 | 2.33 | -1.09 | 3.72 | DN3ab_ltr | DN3cd | 0.66 | 0.91 | 0.39 | 0.68 | 6.40E-05 | ENSMUSG00000060985 | protein_coding | Tdrd5 | tudor domain containing 5 | 214575 | 1 | 156255296 | 156303664 | -1 | Mm.252800 |
| 3.19 | 1.71 | 2.58 | 3.79 | DN3ab_ltr | DN3cd | 0.03 | 0.06 | 0.95 | 0.98 | 6.40E-05 | ENSMUSG00000109787 | TEC | Gm45286 | predicted gene 45286 | NA | 8 | 77574219 | 77575568 | 1 | |
| 6.94 | 6.43 | 5.75 | 6.34 | DN3ab_ltr | DN3cd | -0.07 | -0.35 | 0.74 | 0.91 | 6.60E-05 | ENSMUSG00000026012 | protein_coding | Cd28 | CD28 antigen | 12487 | 1 | 60716800 | 60773359 | 1 | Mm.255003 |
| 4.1 | 5 | 4.52 | 7.15 | DN3ab_ltr | DN3cd | 1.85 | 5.96 | 0.00023 | 0.0074 | 6.90E-05 | ENSMUSG00000065037 | misc_RNA | Rn7sk | RNA, 7SK, nuclear | NA | 9 | 78175303 | 78175633 | -1 | |
| 5.8 | 6.27 | 6.48 | 5.94 | DN3ab_ltr | DN3cd | -0.02 | -0.09 | 0.93 | 0.97 | 7.00E-05 | ENSMUSG00000014444 | protein_coding | Piezo1 | piezo-type mechanosensitive ion channel component 1 | 234839 | 8 | 122481698 | 122551329 | -1 | Mm.37324 Mm.482288 |
| 6.67 | 6.44 | 5.48 | 6.33 | DN3ab_ltr | DN3cd | 0.19 | 1.01 | 0.34 | 0.63 | 7.20E-05 | ENSMUSG00000058672 | protein_coding | Tubb2a | tubulin, beta 2A class IIA | 22151 | 13 | 34074274 | 34078007 | -1 | Mm.422827 |
| 6.5 | 6.12 | 5.71 | 6.66 | DN3ab_ltr | DN3cd | 0.21 | 0.86 | 0.41 | 0.7 | 7.50E-05 | ENSMUSG00000051124 | protein_coding | Gimap9 | GTPase, IMAP family member 9 | 317758 | 6 | 48676129 | 48679114 | 1 | Mm.116971 |
| 7.63 | 8.24 | 2.46 | 9.07 | DN3ab_ltr | DN3cd | 1.32 | 1.89 | 0.092 | 0.31 | 7.80E-05 | ENSMUSG00000061311 | protein_coding | Rag1 | recombination activating gene 1 | 19373 | 2 | 101638282 | 101649501 | -1 | Mm.828 |
| 4.46 | 4.14 | 3.89 | 5.29 | DN3ab_ltr | DN3cd | 0.49 | 1.58 | 0.15 | 0.41 | 7.80E-05 | ENSMUSG00000030337 | protein_coding | Vamp1 | vesicle-associated membrane protein 1 | 22317 | 6 | 125215551 | 125245964 | 1 | Mm.32321 |
| 5.7 | 6.44 | 4.28 | 6.46 | DN3ab_ltr | DN3cd | 1.25 | 5.02 | 0.00075 | 0.016 | 8.40E-05 | ENSMUSG00000001143 | protein_coding | Lman2l | lectin, mannose-binding 2-like | 214895 | 1 | 36419871 | 36445271 | -1 | Mm.287854 |
| 7.36 | 7.22 | 6.73 | 7.6 | DN3ab_ltr | DN3cd | 0.31 | 1.72 | 0.12 | 0.37 | 8.40E-05 | ENSMUSG00000031007 | protein_coding | Atp6ap2 | ATPase, H+ transporting, lysosomal accessory protein 2 | 70495 | X | 12587801 | 12617049 | 1 | Mm.25148 |
| 2.13 | 2.21 | 0.79 | 3.62 | DN3ab_ltr | DN3cd | 1.11 | 2.29 | 0.048 | 0.21 | 8.40E-05 | ENSMUSG00000041750 | polymorphic_pseudogene | Cd1d2 | CD1d2 antigen | 12480 | 3 | 86986551 | 86989780 | 1 | Mm.389465 |
| 7.46 | 7.1 | 7.11 | 7.47 | DN3ab_ltr | DN3cd | -0.03 | -0.2 | 0.84 | 0.93 | 8.50E-05 | ENSMUSG00000027522 | protein_coding | Stx16 | syntaxin 16 | 228960 | 2 | 174076308 | 174099771 | 1 | Mm.277977 |
| 7.2 | 6.56 | 6.79 | 6.97 | DN3ab_ltr | DN3cd | -0.26 | -1.76 | 0.11 | 0.35 | 8.50E-05 | ENSMUSG00000033326 | protein_coding | Kdm4a | lysine (K)-specific demethylase 4A | 230674 | 4 | 118136957 | 118180043 | -1 | Mm.234234 |
| 6.58 | 5.87 | 3.99 | 5.43 | DN3ab_ltr | DN3cd | -0.07 | -0.21 | 0.84 | 0.92 | 8.80E-05 | ENSMUSG00000040209 | protein_coding | Zfp704 | zinc finger protein 704 | 170753 | 3 | 9427020 | 9610085 | -1 | Mm.212908 |
| 5.09 | 5.57 | 6.09 | 5.48 | DN3ab_ltr | DN3cd | -0.08 | -0.38 | 0.71 | 0.9 | 8.90E-05 | ENSMUSG00000002881 | protein_coding | Nab1 | Ngfi-A binding protein 1 | 17936 | 1 | 52457294 | 52500679 | -1 | Mm.25903 |
| 6.77 | 6.31 | 6.05 | 6.45 | DN3ab_ltr | DN3cd | -0.09 | -0.55 | 0.59 | 0.83 | 9.10E-05 | ENSMUSG00000045193 | protein_coding | Cirbp | cold inducible RNA binding protein | 12696 | 10 | 80165985 | 80172786 | 1 | Mm.17898 |
| 5.32 | 5.11 | 6.81 | 5.02 | DN3ab_ltr | DN3cd | -1.02 | -3.59 | 0.006 | 0.057 | 9.20E-05 | ENSMUSG00000050471 | protein_coding | Fam118b | family with sequence similarity 118, member B | 109229 | 9 | 35216965 | 35267805 | -1 | Mm.24197 |
| 4.6 | 4.17 | 3.48 | 4.48 | DN3ab_ltr | DN3cd | 0.13 | 0.51 | 0.62 | 0.84 | 9.30E-05 | ENSMUSG00000028795 | protein_coding | Ccdc28b | coiled coil domain containing 28B | 66264 | 4 | 129619274 | 129623947 | -1 | Mm.383215 |
| 7.87 | 8.14 | 9.38 | 8.59 | DN3ab_ltr | DN3cd | -0.31 | -1.65 | 0.13 | 0.38 | 9.30E-05 | ENSMUSG00000030695 | protein_coding | Aldoa | aldolase A, fructose-bisphosphate | 11674 | 7 | 126795234 | 126800751 | -1 | Mm.275831 |
| -0.47 | -0.08 | -3.72 | 2.39 | DN3ab_ltr | DN3cd | 3.19 | 3.07 | 0.014 | 0.098 | 9.40E-05 | ENSMUSG00000027737 | protein_coding | Slc7a11 | solute carrier family 7 (cationic amino acid transporter, y+ system), member 11 | 26570 | 3 | 49892526 | 50443614 | -1 | Mm.260988 |
| 4.52 | 3.64 | 3.45 | 4.63 | DN3ab_ltr | DN3cd | 0.03 | 0.08 | 0.94 | 0.97 | 9.40E-05 | ENSMUSG00000029311 | protein_coding | Hsd17b11 | hydroxysteroid (17-beta) dehydrogenase 11 | 114664 | 5 | 103989762 | 104021919 | -1 | Mm.46019 Mm.258051 |
| 5.62 | 5.33 | 5.06 | 6.22 | DN3ab_ltr | DN3cd | 0.38 | 1.44 | 0.18 | 0.46 | 9.50E-05 | ENSMUSG00000035891 | protein_coding | Cerk | ceramide kinase | 223753 | 15 | 86139128 | 86186141 | -1 | Mm.222685 |
| 8.83 | 8.47 | 9.83 | 8.7 | DN3ab_ltr | DN3cd | -0.75 | -5.43 | 0.00043 | 0.011 | 9.50E-05 | ENSMUSG00000026547 | protein_coding | Tagln2 | transgelin 2 | 21346 | 1 | 172500047 | 172507380 | 1 | Mm.271711 |
| 7.65 | 8.39 | 6.48 | 8.78 | DN3ab_ltr | DN3cd | 1.38 | 5.01 | 0.00076 | 0.016 | 9.60E-05 | ENSMUSG00000032503 | protein_coding | Arpp21 | cyclic AMP-regulated phosphoprotein, 21 | 74100 | 9 | 112065091 | 112235938 | -1 | Mm.297444 Mm.487515 |
| 4.89 | 4.76 | 8.03 | 6.62 | DN3ab_ltr | DN3cd | -1.01 | -4.71 | 0.0012 | 0.02 | 9.60E-05 | ENSMUSG00000025492 | protein_coding | Ifitm3 | interferon induced transmembrane protein 3 | 66141 | 7 | 141009586 | 141010770 | -1 | Mm.141021 |
| 0.68 | 1.69 | -3.72 | 1.77 | DN3ab_ltr | DN3cd | 2.37 | 3.16 | 0.012 | 0.089 | 9.80E-05 | ENSMUSG00000105708 | processed_transcript | Gm36070 | predicted gene, 36070 | NA | 3 | 94356507 | 94360842 | -1 | |
| -0.11 | -0.93 | -2.95 | 1.41 | DN3ab_ltr | DN3cd | 1.72 | 1.82 | 0.1 | 0.34 | 1.00E-04 | ENSMUSG00000105149 | TEC | Gm42983 | predicted gene 42983 | NA | 5 | 44224510 | 44225324 | 1 | |
| 6.97 | 7.27 | 4.17 | 8 | DN3ab_ltr | DN3cd | 1.33 | 2.29 | 0.048 | 0.21 | 1.00E-04 | ENSMUSG00000034634 | protein_coding | Ly6d | lymphocyte antigen 6 complex, locus D | 17068 | 15 | 74762056 | 74763620 | -1 | Mm.878 |
| 3.98 | 4.03 | 2.15 | 3.9 | DN3ab_ltr | DN3cd | 0.54 | 1.91 | 0.089 | 0.31 | 0.00011 | ENSMUSG00000059027 | processed_transcript | 9630013D21Rik | RIKEN cDNA 9630013D21 gene | NA | 4 | 109651529 | 109657853 | -1 | |
| 7 | 6.32 | 6.61 | 7.31 | DN3ab_ltr | DN3cd | -0.01 | -0.03 | 0.98 | 0.99 | 0.00011 | ENSMUSG00000047888 | protein_coding | Tnrc6b | trinucleotide repeat containing 6b | 213988 | 15 | 80711313 | 80941085 | 1 | Mm.131328 Mm.412092 |
| -1.69 | -3.92 | -3.72 | -0.73 | DN3ab_ltr | DN3cd | 0.41 | 0.43 | 0.68 | 0.88 | 0.00011 | ENSMUSG00000004151 | protein_coding | Etv1 | ets variant 1 | 14009 | 12 | 38779380 | 38870484 | 1 | Mm.4866 |
| 8.47 | 8.65 | 8.77 | 8.38 | DN3ab_ltr | DN3cd | -0.08 | -0.8 | 0.44 | 0.72 | 0.00011 | ENSMUSG00000074884 | protein_coding | Serf2 | small EDRK-rich factor 2 | 378702 | 2 | 121449195 | 121458313 | 1 | Mm.262252 |
| 7.18 | 6.72 | 7.25 | 8.03 | DN3ab_ltr | DN3cd | 0.18 | 0.81 | 0.44 | 0.71 | 0.00011 | ENSMUSG00000028211 | protein_coding | Trp53inp1 | transformation related protein 53 inducible nuclear protein 1 | 60599 | 4 | 11156431 | 11174379 | 1 | Mm.393018 Mm.468195 |
| 4.89 | 4.61 | 6.58 | 4.75 | DN3ab_ltr | DN3cd | -1.13 | -4.03 | 0.0031 | 0.037 | 0.00012 | ENSMUSG00000052837 | protein_coding | Junb | jun B proto-oncogene | 16477 | 8 | 84974484 | 84978718 | -1 | Mm.1167 |
| 6.36 | 6.89 | 5.35 | 7.53 | DN3ab_ltr | DN3cd | 1.22 | 4.13 | 0.0026 | 0.034 | 0.00012 | ENSMUSG00000028657 | protein_coding | Ppt1 | palmitoyl-protein thioesterase 1 | 19063 | 4 | 122836242 | 122859175 | 1 | Mm.277719 |
| 12.7 | 12.77 | 13.57 | 12.69 | DN3ab_ltr | DN3cd | -0.41 | -2.35 | 0.044 | 0.2 | 0.00012 | ENSMUSG00000029580 | protein_coding | Actb | actin, beta | 11461 | 5 | 142903115 | 142906754 | -1 | Mm.328431 Mm.391967 Mm.469717 |
| 6.09 | 6.19 | 6.91 | 6.15 | DN3ab_ltr | DN3cd | -0.33 | -2.03 | 0.073 | 0.27 | 0.00012 | ENSMUSG00000031770 | protein_coding | Herpud1 | homocysteine-inducible, endoplasmic reticulum stress-inducible, ubiquitin-like domain member 1 | 64209 | 8 | 94386438 | 94395377 | 1 | Mm.29151 |
| -0.88 | -3.92 | -3.72 | -1.23 | DN3ab_ltr | DN3cd | -0.51 | -0.51 | 0.63 | 0.84 | 0.00012 | ENSMUSG00000062127 | protein_coding | Cttnbp2nl | CTTNBP2 N-terminal like | 80281 | 3 | 105001915 | 105053146 | -1 | Mm.200327 |
| 7.08 | 7.34 | 5.83 | 7.07 | DN3ab_ltr | DN3cd | 0.63 | 3.61 | 0.0058 | 0.056 | 0.00012 | ENSMUSG00000031245 | protein_coding | Hmgn5 | high-mobility group nucleosome binding domain 5 | 50887 | X | 109004534 | 109013404 | -1 | Mm.298443 |
| 5.08 | 4.44 | 4.54 | 5.47 | DN3ab_ltr | DN3cd | 0.11 | 0.38 | 0.71 | 0.9 | 0.00012 | ENSMUSG00000024781 | protein_coding | Lipa | lysosomal acid lipase A | 16889 | 19 | 34492318 | 34527474 | -1 | Mm.157545 |
| 7.25 | 7.37 | 7.64 | 7.05 | DN3ab_ltr | DN3cd | -0.21 | -1.65 | 0.13 | 0.39 | 0.00012 | ENSMUSG00000004865 | protein_coding | Srpk1 | serine/arginine-rich protein specific kinase 1 | 20815 | 17 | 28587648 | 28622709 | -1 | Mm.15252 Mm.479188 |
| 6.66 | 6.05 | 5.85 | 6.58 | DN3ab_ltr | DN3cd | -0.02 | -0.07 | 0.95 | 0.98 | 0.00012 | ENSMUSG00000022604 | protein_coding | Cep97 | centrosomal protein 97 | 74201 | 16 | 55899888 | 55934855 | -1 | Mm.178427 |
| 6.11 | 5.56 | 5.74 | 6.34 | DN3ab_ltr | DN3cd | 0 | 0 | 1 | 1 | 0.00012 | ENSMUSG00000041777 | protein_coding | Cir1 | corepressor interacting with RBPJ, 1 | 66935 | 2 | 73283105 | 73312701 | -1 | Mm.292140 Mm.361271 |
| 3.57 | 2.03 | 3.54 | 3.99 | DN3ab_ltr | DN3cd | -0.31 | -0.88 | 0.4 | 0.69 | 0.00013 | ENSMUSG00000037363 | protein_coding | Letm2 | leucine zipper-EF-hand containing transmembrane protein 2 | 270035 | 8 | 25578490 | 25597582 | -1 | Mm.268349 |
| 8.86 | 8.71 | 8.57 | 8.96 | DN3ab_ltr | DN3cd | 0.1 | 1 | 0.34 | 0.64 | 0.00013 | ENSMUSG00000022185 | protein_coding | Acin1 | apoptotic chromatin condensation inducer 1 | 56215 | 14 | 54642161 | 54686931 | -1 | Mm.297078 |
| 6.61 | 6.67 | 5.88 | 6.7 | DN3ab_ltr | DN3cd | 0.38 | 2.78 | 0.022 | 0.13 | 0.00013 | ENSMUSG00000036572 | protein_coding | Upf3b | UPF3 regulator of nonsense transcripts homolog B (yeast) | 68134 | X | 37091678 | 37110322 | -1 | Mm.271160 |
| 3.2 | 2.2 | 3.36 | 4.27 | DN3ab_ltr | DN3cd | 0.16 | 0.47 | 0.65 | 0.85 | 0.00013 | ENSMUSG00000081769 | protein_coding | Gm12216 | predicted gene 12216 | NA | 11 | 53783418 | 53859256 | -1 | |
| 6.74 | 6.71 | 5.9 | 6.72 | DN3ab_ltr | DN3cd | 0.33 | 2.14 | 0.062 | 0.25 | 0.00013 | ENSMUSG00000020069 | protein_coding | Hnrnph3 | heterogeneous nuclear ribonucleoprotein H3 | 432467 | 10 | 63014664 | 63024217 | -1 | Mm.274784 |
| 0.89 | 4.16 | -3.72 | 4.49 | DN3ab_ltr | DN3cd | 4.5 | 5.77 | 0.00028 | 0.0085 | 0.00013 | ENSMUSG00000069805 | protein_coding | Fbp1 | fructose bisphosphatase 1 | 14121 | 13 | 62864753 | 62888282 | -1 | Mm.423078 |
| 5.73 | 5.79 | 2.33 | 6.88 | DN3ab_ltr | DN3cd | 0.92 | 1.43 | 0.19 | 0.47 | 0.00013 | ENSMUSG00000053044 | protein_coding | Cd8b1 | CD8 antigen, beta chain 1 | 12526 | 6 | 71322788 | 71337494 | 1 | Mm.333148 |
| 6.64 | 6.05 | 6.3 | 6.73 | DN3ab_ltr | DN3cd | -0.1 | -0.53 | 0.61 | 0.84 | 0.00013 | ENSMUSG00000058318 | protein_coding | Phf21a | PHD finger protein 21A | 192285 | 2 | 92093117 | 92364666 | 1 | Mm.330408 |
| 6.5 | 6.56 | 6.12 | 7.03 | DN3ab_ltr | DN3cd | 0.45 | 2.85 | 0.019 | 0.12 | 0.00013 | ENSMUSG00000020346 | protein_coding | Mgat1 | mannoside acetylglucosaminyltransferase 1 | 17308 | 11 | 49244191 | 49263030 | 1 | Mm.196933 |
| 4.63 | 4.4 | 4.03 | 5.08 | DN3ab_ltr | DN3cd | 0.34 | 1.46 | 0.18 | 0.45 | 0.00013 | ENSMUSG00000041346 | protein_coding | Wrap53 | WD repeat containing, antisense to Trp53 | 216853 | 11 | 69561758 | 69580255 | -1 | Mm.31586 |
| 5.31 | 4.79 | 4.28 | 5.27 | DN3ab_ltr | DN3cd | 0.1 | 0.38 | 0.71 | 0.9 | 0.00013 | ENSMUSG00000035967 | protein_coding | Ints6l | integrator complex subunit 6 like | 236790 | X | 56454857 | 56507843 | 1 | Mm.474772 |
| 0.77 | 0.53 | 4.58 | 1.04 | DN3ab_ltr | DN3cd | -2.43 | -4.26 | 0.0022 | 0.03 | 0.00013 | ENSMUSG00000043154 | protein_coding | Ppp2r3a | protein phosphatase 2, regulatory subunit B'', alpha | 235542 | 9 | 101105084 | 101251795 | -1 | Mm.335386 |
| 7.28 | 6.57 | 6.4 | 6.72 | DN3ab_ltr | DN3cd | -0.26 | -1.4 | 0.2 | 0.47 | 0.00014 | ENSMUSG00000034248 | protein_coding | Slc25a37 | solute carrier family 25, member 37 | 67712 | 14 | 69241848 | 69285112 | -1 | Mm.293635 |
| 6.1 | 6.12 | 6.97 | 6.09 | DN3ab_ltr | DN3cd | -0.43 | -2.61 | 0.029 | 0.16 | 0.00014 | ENSMUSG00000063410 | protein_coding | Stk24 | serine/threonine kinase 24 | 223255 | 14 | 121286343 | 121380011 | -1 | Mm.390756 Mm.442081 Mm.472866 |
| 9.76 | 9.79 | 10.41 | 9.57 | DN3ab_ltr | DN3cd | -0.39 | -2.44 | 0.038 | 0.18 | 0.00014 | ENSMUSG00000001175 | protein_coding | Calm1 | calmodulin 1 | 12313 | 12 | 100199435 | 100209814 | 1 | Mm.285993 |
| 5.9 | 4.8 | 4.34 | 4.51 | DN3ab_ltr | DN3cd | -0.64 | -2.82 | 0.02 | 0.13 | 0.00014 | ENSMUSG00000105199 | sense_intronic | Gm43581 | predicted gene 43581 | NA | 3 | 104642994 | 104645857 | 1 | |
| 8.65 | 8.38 | 9.1 | 8.26 | DN3ab_ltr | DN3cd | -0.54 | -5.1 | 0.00068 | 0.014 | 0.00014 | ENSMUSG00000000563 | protein_coding | Atp5f1 | ATP synthase, H+ transporting, mitochondrial F0 complex, subunit B1 | 11950 | 3 | 105942698 | 105960099 | -1 | Mm.251152 |
| 9.64 | 9.42 | 9.14 | 9.58 | DN3ab_ltr | DN3cd | 0.08 | 0.65 | 0.53 | 0.79 | 0.00014 | ENSMUSG00000063511 | protein_coding | Snrnp70 | small nuclear ribonucleoprotein 70 (U1) | 20637 | 7 | 45376453 | 45395742 | -1 | Mm.216386 |
| 5.95 | 5.77 | 5.22 | 6.21 | DN3ab_ltr | DN3cd | 0.33 | 1.56 | 0.15 | 0.42 | 0.00015 | ENSMUSG00000062761 | protein_coding | Zfp512 | zinc finger protein 512 | 269639 | 5 | 31452431 | 31481754 | 1 | Mm.259127 |
| 7.3 | 6.98 | 6.67 | 7.18 | DN3ab_ltr | DN3cd | 0.05 | 0.31 | 0.77 | 0.92 | 0.00015 | ENSMUSG00000045973 | protein_coding | Slc25a51 | solute carrier family 25, member 51 | 230125 | 4 | 45395923 | 45408766 | -1 | Mm.260210 |
| 3.89 | 3.75 | 2.98 | 4.75 | DN3ab_ltr | DN3cd | 0.66 | 1.91 | 0.089 | 0.31 | 0.00015 | ENSMUSG00000082585 | processed_pseudogene | Gm15387 | predicted gene 15387 | NA | 15 | 74093686 | 74094333 | -1 | |
| 6.08 | 6.31 | 5.25 | 6.41 | DN3ab_ltr | DN3cd | 0.61 | 3.64 | 0.0055 | 0.054 | 0.00015 | ENSMUSG00000032763 | protein_coding | Ilvbl | ilvB (bacterial acetolactate synthase)-like | 216136 | 10 | 78574346 | 78584502 | 1 | Mm.2644 |
| 5.4 | 5.37 | 5.55 | 6.42 | DN3ab_ltr | DN3cd | 0.43 | 2.62 | 0.028 | 0.15 | 0.00015 | ENSMUSG00000058761 | protein_coding | Rnf169 | ring finger protein 169 | 108937 | 7 | 99920254 | 99980448 | -1 | Mm.323087 |
| 6.17 | 5.53 | 5.8 | 6.01 | DN3ab_ltr | DN3cd | -0.24 | -1.53 | 0.16 | 0.42 | 0.00015 | ENSMUSG00000078429 | protein_coding | Ctdsp2 | CTD (carboxy-terminal domain, RNA polymerase II, polypeptide A) small phosphatase 2 | 52468 | 10 | 126978717 | 126999975 | 1 | Mm.29490 |
| 6.51 | 6.07 | 6.07 | 6.67 | DN3ab_ltr | DN3cd | 0.05 | 0.24 | 0.82 | 0.92 | 0.00015 | ENSMUSG00000047989 | protein_coding | Ino80c | INO80 complex subunit C | 225280 | 18 | 24104757 | 24121953 | -1 | Mm.258973 |
| 0.29 | 0.2 | -3.72 | 1.04 | DN3ab_ltr | DN3cd | 1.82 | 2.08 | 0.067 | 0.26 | 0.00016 | ENSMUSG00000102687 | TEC | Gm37219 | predicted gene, 37219 | NA | 1 | 119630744 | 119631669 | -1 | |
| 5.62 | 5.63 | 5.07 | 6.1 | DN3ab_ltr | DN3cd | 0.46 | 2.48 | 0.035 | 0.18 | 0.00016 | ENSMUSG00000079555 | protein_coding | Haus3 | HAUS augmin-like complex, subunit 3 | 231123 | 5 | 34153880 | 34169527 | -1 | Mm.319286 |
| 2.83 | 2.03 | 0.29 | 3.77 | DN3ab_ltr | DN3cd | 0.49 | 0.69 | 0.51 | 0.77 | 0.00016 | ENSMUSG00000005883 | protein_coding | Spo11 | SPO11 meiotic protein covalently bound to DSB | 26972 | 2 | 172977700 | 172993576 | 1 | Mm.23495 |
| 6.25 | 6.49 | 3.6 | 6.89 | DN3ab_ltr | DN3cd | 1.09 | 2.18 | 0.058 | 0.24 | 0.00016 | ENSMUSG00000020323 | protein_coding | Prss57 | protease, serine 57 | 73106 | 10 | 79781474 | 79790961 | -1 | Mm.491308 |
| -1.69 | -3.92 | -3.72 | -0.87 | DN3ab_ltr | DN3cd | 0.31 | 0.33 | 0.75 | 0.92 | 0.00016 | ENSMUSG00000087272 | lincRNA | Gm15409 | predicted gene 15409 | NA | 5 | 148959278 | 148961712 | 1 | |
| 4.9 | 4.24 | 4.05 | 5.16 | DN3ab_ltr | DN3cd | 0.13 | 0.41 | 0.69 | 0.88 | 0.00016 | ENSMUSG00000028988 | protein_coding | Ctnnbip1 | catenin beta interacting protein 1 | 67087 | 4 | 149518236 | 149566437 | 1 | Mm.299735 Mm.474667 |
| 5.41 | 3.74 | 5.58 | 4.93 | DN3ab_ltr | DN3cd | -1.09 | -5.86 | 0.00025 | 0.0079 | 0.00017 | ENSMUSG00000031431 | protein_coding | Tsc22d3 | TSC22 domain family, member 3 | 14605 | X | 140539528 | 140600659 | -1 | Mm.22216 |
| 4.88 | 4.73 | 6.25 | 5 | DN3ab_ltr | DN3cd | -0.75 | -3.7 | 0.005 | 0.051 | 0.00017 | ENSMUSG00000003382 | protein_coding | Etv3 | ets variant 3 | 27049 | 3 | 87525407 | 87540156 | 1 | Mm.219460 |
| 2.04 | 3.19 | -0.86 | 4.05 | DN3ab_ltr | DN3cd | 1.89 | 3.41 | 0.0079 | 0.069 | 0.00017 | ENSMUSG00000022197 | protein_coding | Pdzd2 | PDZ domain containing 2 | 68070 | 15 | 12359711 | 12739924 | -1 | Mm.423205 |
| 7.49 | 7.32 | 6.88 | 7.37 | DN3ab_ltr | DN3cd | 0.12 | 0.97 | 0.36 | 0.65 | 0.00017 | ENSMUSG00000060261 | protein_coding | Gtf2i | general transcription factor II I | 14886 | 5 | 134237834 | 134314760 | -1 | Mm.261570 Mm.412191 Mm.466495 |
| 2.57 | 3.11 | 0.07 | 3.36 | DN3ab_ltr | DN3cd | 1.16 | 2.72 | 0.024 | 0.14 | 0.00018 | ENSMUSG00000020251 | protein_coding | Glt8d2 | glycosyltransferase 8 domain containing 2 | 74782 | 10 | 82650433 | 82690650 | -1 | Mm.191281 |
| 3.31 | 2.9 | 5.66 | 3.06 | DN3ab_ltr | DN3cd | -1.69 | -4.23 | 0.0023 | 0.031 | 0.00018 | ENSMUSG00000021356 | protein_coding | Irf4 | interferon regulatory factor 4 | 16364 | 13 | 30749226 | 30766976 | 1 | Mm.4677 |
| 7.3 | 6.94 | 6.97 | 7.42 | DN3ab_ltr | DN3cd | 0.02 | 0.13 | 0.9 | 0.95 | 0.00018 | ENSMUSG00000030336 | protein_coding | Cd27 | CD27 antigen | 21940 | 6 | 125232622 | 125237010 | -1 | Mm.367714 |
| 9.7 | 9.94 | 8.6 | 9.66 | DN3ab_ltr | DN3cd | 0.56 | 3.82 | 0.0042 | 0.045 | 0.00018 | ENSMUSG00000066551 | protein_coding | Hmgb1 | high mobility group box 1 | 15289 | 5 | 149046702 | 149184489 | -1 | Mm.207047 Mm.313345 |
| 4.82 | 5.25 | 5.65 | 4.27 | DN3ab_ltr | DN3cd | -0.38 | -1.15 | 0.28 | 0.58 | 0.00018 | ENSMUSG00000058173 | protein_coding | Smco4 | single-pass membrane protein with coiled-coil domains 4 | 170748 | 9 | 15493432 | 15545260 | 1 | Mm.86572 |
| 3.83 | 4.48 | 2.79 | 5.54 | DN3ab_ltr | DN3cd | 1.44 | 3.84 | 0.0041 | 0.044 | 0.00018 | ENSMUSG00000104145 | TEC | D130019J16Rik | RIKEN cDNA D130019J16 gene | NA | 1 | 137935189 | 137937346 | 1 | |
| 7.31 | 7 | 6.96 | 7.48 | DN3ab_ltr | DN3cd | 0.08 | 0.51 | 0.62 | 0.84 | 0.00018 | ENSMUSG00000070733 | protein_coding | Fryl | FRY like transcription coactivator | 72313 | 5 | 73019987 | 73256619 | -1 | Mm.260594 |
| 8.27 | 8.12 | 8.8 | 7.88 | DN3ab_ltr | DN3cd | -0.51 | -3.61 | 0.0058 | 0.056 | 0.00018 | ENSMUSG00000006699 | protein_coding | Cdc42 | cell division cycle 42 | 12540 | 4 | 137319696 | 137357720 | -1 | Mm.1022 Mm.447553 |
| 8.85 | 8.45 | 9.74 | 8.71 | DN3ab_ltr | DN3cd | -0.73 | -6.2 | 0.00017 | 0.0062 | 0.00018 | ENSMUSG00000021998 | protein_coding | Lcp1 | lymphocyte cytosolic protein 1 | 18826 | 14 | 75131101 | 75230842 | 1 | Mm.153911 Mm.484415 |
| 4.92 | 3.58 | 3.72 | 4.14 | DN3ab_ltr | DN3cd | -0.58 | -1.81 | 0.1 | 0.34 | 0.00019 | ENSMUSG00000044770 | protein_coding | Scml4 | Scm polycomb group protein like 4 | 268297 | 10 | 42860370 | 42960780 | 1 | Mm.98731 |
| -0.88 | -3.92 | -3.72 | -0.77 | DN3ab_ltr | DN3cd | -0.17 | -0.15 | 0.88 | 0.95 | 0.00019 | ENSMUSG00000071042 | protein_coding | Rasgrp3 | RAS, guanyl releasing protein 3 | 240168 | 17 | 75435896 | 75529054 | 1 | Mm.329203 |
| 6.27 | 6.16 | 5.67 | 6.55 | DN3ab_ltr | DN3cd | 0.33 | 1.8 | 0.11 | 0.34 | 0.00019 | ENSMUSG00000047412 | protein_coding | Zbtb44 | zinc finger and BTB domain containing 44 | 235132 | 9 | 31030644 | 31075885 | 1 | Mm.239355 Mm.393131 |
| 3.08 | 2.56 | 2.33 | 3.79 | DN3ab_ltr | DN3cd | 0.4 | 1.09 | 0.3 | 0.6 | 0.00019 | ENSMUSG00000050957 | protein_coding | Insl6 | insulin-like 6 | 27356 | 19 | 29321344 | 29325356 | -1 | Mm.306511 |
| 7.84 | 8.09 | 8.23 | 7.89 | DN3ab_ltr | DN3cd | -0.03 | -0.27 | 0.79 | 0.92 | 0.00019 | ENSMUSG00000001627 | protein_coding | Ifrd1 | interferon-related developmental regulator 1 | 15982 | 12 | 40201567 | 40248504 | -1 | Mm.168 |
| -0.43 | -1.73 | -3.72 | 0.23 | DN3ab_ltr | DN3cd | 1.31 | 1.34 | 0.21 | 0.49 | 2.00E-04 | ENSMUSG00000097211 | antisense | BC065403 | cDNA sequence BC065403 | NA | 9 | 72361735 | 72409945 | -1 | |
| 6.55 | 5.94 | 6.31 | 6.59 | DN3ab_ltr | DN3cd | -0.18 | -1.11 | 0.29 | 0.59 | 2.00E-04 | ENSMUSG00000034297 | protein_coding | Med13 | mediator complex subunit 13 | 327987 | 11 | 86267033 | 86357602 | -1 | Mm.341886 |
| 8.26 | 8.46 | 7.66 | 8.65 | DN3ab_ltr | DN3cd | 0.55 | 3.8 | 0.0043 | 0.046 | 0.00021 | ENSMUSG00000032187 | protein_coding | Smarca4 | SWI/SNF related, matrix associated, actin dependent regulator of chromatin, subfamily a, member 4 | 20586 | 9 | 21616169 | 21704230 | 1 | Mm.286593 |
| 7.81 | 7.9 | 8.62 | 7.92 | DN3ab_ltr | DN3cd | -0.3 | -2.06 | 0.069 | 0.27 | 0.00021 | ENSMUSG00000001467 | protein_coding | Cyp51 | cytochrome P450, family 51 | 13121 | 5 | 4081145 | 4104746 | -1 | Mm.140158 |
| 6.76 | 6.81 | 8.35 | 7.38 | DN3ab_ltr | DN3cd | -0.52 | -2.77 | 0.022 | 0.13 | 0.00021 | ENSMUSG00000059040 | protein_coding | Eno1b | enolase 1B, retrotransposed | 433182 | 18 | 48045335 | 48048378 | 1 | Mm.70666 Mm.372357 |
| 6.65 | 6.53 | 5.87 | 6.68 | DN3ab_ltr | DN3cd | 0.28 | 1.62 | 0.14 | 0.4 | 0.00021 | ENSMUSG00000036893 | protein_coding | Ehmt1 | euchromatic histone methyltransferase 1 | 77683 | 2 | 24789928 | 24919614 | -1 | Mm.24176 |
| 6.37 | 5.54 | 5.76 | 5.74 | DN3ab_ltr | DN3cd | -0.47 | -3.15 | 0.012 | 0.09 | 0.00021 | ENSMUSG00000001761 | protein_coding | Smo | smoothened, frizzled class receptor | 319757 | 6 | 29735503 | 29761365 | 1 | Mm.29279 |
| 7.45 | 7.12 | 8.38 | 7.18 | DN3ab_ltr | DN3cd | -0.77 | -4.8 | 0.001 | 0.018 | 0.00021 | ENSMUSG00000034595 | protein_coding | Ppp1r18 | protein phosphatase 1, regulatory subunit 18 | 76448 | 17 | 35865593 | 35875596 | 1 | Mm.359982 Mm.421125 |
| 8.21 | 8.54 | 7.46 | 8.67 | DN3ab_ltr | DN3cd | 0.71 | 4.45 | 0.0017 | 0.026 | 0.00021 | ENSMUSG00000024613 | protein_coding | Tcof1 | treacle ribosome biogenesis factor 1 | 21453 | 18 | 60813755 | 60848971 | -1 | Mm.486251 |
| 6.88 | 6.94 | 8.52 | 7.54 | DN3ab_ltr | DN3cd | -0.52 | -2.77 | 0.022 | 0.13 | 0.00021 | ENSMUSG00000063524 | protein_coding | Eno1 | enolase 1, alpha non-neuron | 13806 | 4 | 150236721 | 150248879 | 1 | Mm.70666 Mm.372357 Mm.372389 |
| 5.68 | 4.66 | 11.47 | 7.36 | DN3ab_ltr | DN3cd | -3.41 | -7.21 | 5.40E-05 | 0.0032 | 0.00022 | ENSMUSG00000015437 | protein_coding | Gzmb | granzyme B | 14939 | 14 | 56258838 | 56262260 | -1 | Mm.14874 |
| 5.63 | 5.6 | 6.16 | 5.21 | DN3ab_ltr | DN3cd | -0.47 | -2.72 | 0.024 | 0.14 | 0.00022 | ENSMUSG00000033629 | protein_coding | Hacd3 | 3-hydroxyacyl-CoA dehydratase 3 | 57874 | 9 | 64986983 | 65021693 | -1 | Mm.308180 |
| 4.17 | 3.74 | 3.23 | 4.8 | DN3ab_ltr | DN3cd | 0.43 | 1.18 | 0.27 | 0.56 | 0.00022 | ENSMUSG00000104432 | TEC | A430027C01Rik | RIKEN cDNA A430027C01 gene | NA | 9 | 72011376 | 72013019 | -1 | |
| 7.87 | 7.93 | 9.52 | 8.69 | DN3ab_ltr | DN3cd | -0.45 | -2.76 | 0.022 | 0.13 | 0.00022 | ENSMUSG00000038037 | protein_coding | Socs1 | suppressor of cytokine signaling 1 | 12703 | 16 | 10782240 | 10785536 | -1 | Mm.130 |
| 9.23 | 9.33 | 8.93 | 10 | DN3ab_ltr | DN3cd | 0.56 | 3.11 | 0.013 | 0.093 | 0.00023 | ENSMUSG00000020009 | protein_coding | Ifngr1 | interferon gamma receptor 1 | 15979 | 10 | 19591949 | 19610229 | 1 | Mm.549 |
| 3.61 | 3.87 | 5.74 | 4.74 | DN3ab_ltr | DN3cd | -0.55 | -2.42 | 0.039 | 0.19 | 0.00023 | ENSMUSG00000027544 | protein_coding | Nfatc2 | nuclear factor of activated T cells, cytoplasmic, calcineurin dependent 2 | 18019 | 2 | 168476410 | 168601657 | -1 | Mm.116802 |
| 3.29 | 2.25 | 1.02 | 3.48 | DN3ab_ltr | DN3cd | 0.12 | 0.2 | 0.84 | 0.93 | 0.00023 | ENSMUSG00000007682 | protein_coding | Dio2 | deiodinase, iodothyronine, type II | 13371 | 12 | 90724552 | 90738438 | -1 | Mm.21389 Mm.408815 |
| 3.26 | 5.16 | 0.02 | 5.8 | DN3ab_ltr | DN3cd | 2.45 | 4.88 | 9.00E-04 | 0.017 | 0.00023 | ENSMUSG00000063109 | antisense | Dgkeos | diacylglycerol kinase, epsilon, opposite strand | NA | 11 | 89064729 | 89069098 | 1 | |
| 4.79 | 4.58 | 6.39 | 4.99 | DN3ab_ltr | DN3cd | -0.88 | -3.99 | 0.0033 | 0.038 | 0.00023 | ENSMUSG00000053560 | protein_coding | Ier2 | immediate early response 2 | 15936 | 8 | 84661331 | 84662852 | -1 | Mm.399 |
| 4.37 | 6.01 | 5.83 | 6.39 | DN3ab_ltr | DN3cd | 1.03 | 5.18 | 0.00061 | 0.014 | 0.00024 | ENSMUSG00000059895 | protein_coding | Ptp4a3 | protein tyrosine phosphatase 4a3 | 19245 | 15 | 73723145 | 73758766 | 1 | Mm.390807 |
| 7.3 | 7.06 | 7.05 | 7.64 | DN3ab_ltr | DN3cd | 0.15 | 0.99 | 0.35 | 0.64 | 0.00024 | ENSMUSG00000062604 | protein_coding | Srpk2 | serine/arginine-rich protein specific kinase 2 | 20817 | 5 | 23503264 | 23684617 | -1 | Mm.288728 |
| 7.62 | 7.66 | 8.19 | 7.59 | DN3ab_ltr | DN3cd | -0.27 | -2.21 | 0.055 | 0.23 | 0.00024 | ENSMUSG00000022831 | protein_coding | Hcls1 | hematopoietic cell specific Lyn substrate 1 | 15163 | 16 | 36934983 | 36963212 | 1 | Mm.4091 |
| 0.77 | -1.17 | -1.76 | 1.65 | DN3ab_ltr | DN3cd | 0.6 | 0.61 | 0.56 | 0.8 | 0.00024 | ENSMUSG00000003410 | protein_coding | Elavl3 | ELAV like RNA binding protein 3 | 15571 | 9 | 22015005 | 22052023 | -1 | Mm.390167 |
| 7.22 | 6.39 | 7.2 | 7.37 | DN3ab_ltr | DN3cd | -0.32 | -1.73 | 0.12 | 0.36 | 0.00024 | ENSMUSG00000002897 | protein_coding | Il17ra | interleukin 17 receptor A | 16172 | 6 | 120463247 | 120487559 | 1 | Mm.4481 |
| 7.61 | 7.8 | 7.95 | 7.49 | DN3ab_ltr | DN3cd | -0.11 | -0.94 | 0.37 | 0.66 | 0.00025 | ENSMUSG00000044147 | protein_coding | Arf6 | ADP-ribosylation factor 6 | 11845 | 12 | 69372150 | 69373783 | 1 | Mm.27308 |
| 10 | 10.09 | 9.86 | 9.35 | DN3ab_ltr | DN3cd | -0.17 | -1.51 | 0.17 | 0.43 | 0.00025 | ENSMUSG00000028691 | protein_coding | Prdx1 | peroxiredoxin 1 | 18477 | 4 | 116685544 | 116700822 | 1 | Mm.30929 |
| 6.85 | 7.01 | 5.64 | 6.68 | DN3ab_ltr | DN3cd | 0.5 | 3.14 | 0.012 | 0.091 | 0.00025 | ENSMUSG00000032218 | protein_coding | Ccnb2 | cyclin B2 | 12442 | 9 | 70407692 | 70421547 | -1 | Mm.22592 |
| 8.01 | 8.28 | 7.19 | 8.29 | DN3ab_ltr | DN3cd | 0.62 | 4.02 | 0.0031 | 0.037 | 0.00026 | ENSMUSG00000005698 | protein_coding | Ctcf | CCCTC-binding factor | 13018 | 8 | 105636568 | 105682922 | 1 | Mm.269474 |
| 4.05 | 4.65 | 3.95 | 5.94 | DN3ab_ltr | DN3cd | 1.28 | 4.92 | 0.00085 | 0.017 | 0.00026 | ENSMUSG00000016756 | protein_coding | Cmah | cytidine monophospho-N-acetylneuraminic acid hydroxylase | 12763 | 13 | 24327404 | 24477377 | 1 | Mm.8396 |
| 6.73 | 6.53 | 5.59 | 6.32 | DN3ab_ltr | DN3cd | 0.17 | 1.03 | 0.33 | 0.63 | 0.00026 | ENSMUSG00000032902 | protein_coding | Slc16a1 | solute carrier family 16 (monocarboxylic acid transporters), member 1 | 20501 | 3 | 104638668 | 104658462 | 1 | Mm.9086 |

| | | | | | | | | | | | | | | | | | | | | |
|---|---|---|---|---|---|---|---|---|---|---|---|---|---|---|---|---|---|---|---|---|
| 8.04 | 8.08 | 8.51 | 7.98 | DN3ab_ltr | DN3cd | -0.24 | -2.25 | 0.051 | 0.22 | 0.00026 | ENSMUSG00000000420 | protein_coding | Galnt1 | polypeptide N-acetylgalactosaminyltransferase 1 | 14423 | 18 | 24205344 | 24286818 | 1 | Mm.30249 |
| 1.24 | 1.28 | -2.9 | 2.25 | DN3ab_ltr | DN3cd | 1.53 | 1.78 | 0.11 | 0.35 | 0.00026 | ENSMUSG00000079076 | transcribed_processed_pseudogene | Gm3086 | predicted gene 3086 | NA | 12 | 69963409 | 69969736 | 1 | |
| 3.66 | 3.61 | 1.92 | 5.44 | DN3ab_ltr | DN3cd | 1.16 | 1.86 | 0.097 | 0.32 | 0.00026 | ENSMUSG00000076911 | TR_J_gene | Traj18 | T cell receptor alpha joining 18 | NA | 14 | 54200777 | 54200842 | 1 | |
| 4.84 | 5.19 | 5.45 | 4.67 | DN3ab_ltr | DN3cd | -0.18 | -0.84 | 0.42 | 0.71 | 0.00026 | ENSMUSG00000050188 | protein_coding | Lsm10 | U7 snRNP-specific Sm-like protein LSM10 | 116748 | 4 | 126096623 | 126098584 | 1 | Mm.41865 |
| 10.37 | 10.36 | 9.76 | 10.23 | DN3ab_ltr | DN3cd | 0.19 | 2.09 | 0.067 | 0.26 | 0.00026 | ENSMUSG00000059005 | protein_coding | Hnrnpa3 | heterogeneous nuclear ribonucleoprotein A3 | 229279 | 2 | 75659261 | 75669407 | 1 | Mm.316306 Mm.331491 Mm.379375 |
| 5.19 | 5.63 | 3.76 | 6.04 | DN3ab_ltr | DN3cd | 1.1 | 3.37 | 0.0085 | 0.072 | 0.00027 | ENSMUSG00000045775 | protein_coding | Slc16a5 | solute carrier family 16 (monocarboxylic acid transporters), member 5 | 217316 | 11 | 115462474 | 115474398 | 1 | Mm.25773 |
| 2.6 | 1.96 | 1.31 | 2.76 | DN3ab_ltr | DN3cd | 0.19 | 0.48 | 0.64 | 0.85 | 0.00027 | ENSMUSG00000047342 | protein_coding | Zfp286 | zinc finger protein 286 | 192651 | 11 | 62752577 | 62789462 | -1 | Mm.168530 |
| 4.2 | 5.56 | 2.74 | 5.65 | DN3ab_ltr | DN3cd | 1.84 | 6.89 | 7.60E-05 | 0.0038 | 0.00027 | ENSMUSG00000042035 | protein_coding | Igsf3 | immunoglobulin superfamily, member 3 | 78908 | 3 | 101377083 | 101463059 | 1 | Mm.257997 Mm.396180 Mm.489667 |
| 5.74 | 5.9 | 4.53 | 5.59 | DN3ab_ltr | DN3cd | 0.5 | 3.05 | 0.014 | 0.099 | 0.00027 | ENSMUSG00000014633 | protein_coding | Cmc2 | COX assembly mitochondrial protein 2 | 66531 | 8 | 116888685 | 116921455 | -1 | Mm.4011 |
| 5.94 | 5.95 | 4.66 | 5.59 | DN3ab_ltr | DN3cd | 0.35 | 2.1 | 0.066 | 0.26 | 0.00027 | ENSMUSG00000028678 | protein_coding | Kif2c | kinesin family member 2C | 73804 | 4 | 117159639 | 117182639 | -1 | Mm.247651 |
| 3.63 | 3.55 | 2.37 | 4.17 | DN3ab_ltr | DN3cd | 0.61 | 1.81 | 0.1 | 0.34 | 0.00027 | ENSMUSG00000017639 | protein_coding | Rab11fip4 | RAB11 family interacting protein 4 (class II) | 268451 | 11 | 79591212 | 79698023 | 1 | Mm.130656 Mm.491175 |
| 6.27 | 6.29 | 5.87 | 6.69 | DN3ab_ltr | DN3cd | 0.39 | 2.6 | 0.029 | 0.16 | 0.00027 | ENSMUSG00000021054 | protein_coding | Sgpp1 | sphingosine-1-phosphate phosphatase 1 | 81535 | 12 | 75714249 | 75735729 | -1 | Mm.280199 |
| 6.4 | 6.26 | 4.61 | 6.25 | DN3ab_ltr | DN3cd | 0.49 | 1.55 | 0.15 | 0.42 | 0.00027 | ENSMUSG00000029254 | protein_coding | Stap1 | signal transducing adaptor family member 1 | 56792 | 5 | 86071746 | 86106125 | 1 | Mm.131237 |
| 2.78 | 3.62 | -2.09 | 4.49 | DN3ab_ltr | DN3cd | 1.57 | 2.22 | 0.054 | 0.23 | 0.00028 | ENSMUSG00000028150 | protein_coding | Rorc | RAR-related orphan receptor gamma | 19885 | 3 | 94372794 | 94398276 | 1 | Mm.4372 |
| 6.75 | 6.63 | 6.31 | 7.06 | DN3ab_ltr | DN3cd | 0.28 | 1.73 | 0.12 | 0.36 | 0.00028 | ENSMUSG00000050965 | protein_coding | Prkca | protein kinase C, alpha | 18750 | 11 | 107933387 | 108343928 | -1 | Mm.222178 |
| 7.57 | 7.8 | 6.64 | 7.87 | DN3ab_ltr | DN3cd | 0.65 | 3.59 | 0.006 | 0.057 | 0.00028 | ENSMUSG00000045328 | protein_coding | Cenpe | centromere protein E | 229841 | 3 | 135212537 | 135273611 | 1 | Mm.161470 |
| 6.6 | 6.87 | 8.64 | 7.23 | DN3ab_ltr | DN3cd | -0.68 | -2.2 | 0.056 | 0.23 | 0.00028 | ENSMUSG00000019943 | protein_coding | Atp2b1 | ATPase, Ca++ transporting, plasma membrane 1 | 67972 | 10 | 98914406 | 99026143 | 1 | Mm.166944 Mm.471912 |
| 6.43 | 5.73 | 8.2 | 6.11 | DN3ab_ltr | DN3cd | -1.47 | -5.73 | 3.00E-04 | 0.0087 | 0.00028 | ENSMUSG00000028163 | protein_coding | Nfkb1 | nuclear factor of kappa light polypeptide gene enhancer in B cells 1, p105 | 18033 | 3 | 135584655 | 135691547 | -1 | Mm.256765 |
| 1.87 | 2.41 | 4.78 | 3.38 | DN3ab_ltr | DN3cd | -0.85 | -2.58 | 0.03 | 0.16 | 0.00028 | ENSMUSG00000059479 | protein_coding | B3gnt8 | UDP-GlcNAc:betaGal beta-1,3-N-acetylglucosaminyltransferase 8 | 232984 | 7 | 25626654 | 25635100 | 1 | Mm.192369 |
| 4.86 | 3.63 | 3.68 | 3.73 | DN3ab_ltr | DN3cd | -0.71 | -3 | 0.015 | 0.1 | 0.00028 | ENSMUSG00000042272 | protein_coding | Sestd1 | SEC14 and spectrin domains 1 | 228071 | 2 | 77180340 | 77280592 | -1 | Mm.27120 |
| 7.8 | 7.74 | 7.03 | 7.71 | DN3ab_ltr | DN3cd | 0.26 | 1.89 | 0.092 | 0.31 | 0.00028 | ENSMUSG00000002297 | protein_coding | Dbf4 | DBF4 zinc finger | 27214 | 5 | 8396973 | 8422716 | -1 | Mm.292470 |
| 9.52 | 8.68 | 10.76 | 9.38 | DN3ab_ltr | DN3cd | -1.14 | -11.27 | 1.50E-06 | 0.00042 | 0.00029 | ENSMUSG00000026770 | protein_coding | Il2ra | interleukin 2 receptor, alpha chain | 16184 | 2 | 11642807 | 11693193 | 1 | Mm.915 |
| 8.08 | 8.65 | 8.54 | 8.37 | DN3ab_ltr | DN3cd | 0.22 | 1.62 | 0.14 | 0.4 | 0.00029 | ENSMUSG00000025153 | protein_coding | Fasn | fatty acid synthase | 14104 | 11 | 120805846 | 120824547 | -1 | Mm.236443 |
| 5.16 | 3.61 | 5.58 | 5.66 | DN3ab_ltr | DN3cd | -0.54 | -1.83 | 0.1 | 0.33 | 0.00029 | ENSMUSG00000042675 | protein_coding | Ypel3 | yippee like 3 | 66090 | 7 | 126776955 | 126780514 | 1 | Mm.27114 |
| 5.92 | 5.67 | 5.38 | 5.96 | DN3ab_ltr | DN3cd | 0.12 | 0.75 | 0.47 | 0.74 | 0.00029 | ENSMUSG00000027452 | protein_coding | Acss1 | acyl-CoA synthetase short-chain family member 1 | 68738 | 2 | 150618105 | 150668500 | -1 | Mm.7044 |
| 3.99 | 5.22 | 1.02 | 5.58 | DN3ab_ltr | DN3cd | 1.78 | 3.86 | 0.004 | 0.044 | 0.00029 | ENSMUSG00000027858 | protein_coding | Tspan2 | tetraspanin 2 | 70747 | 3 | 102734529 | 102801513 | 1 | Mm.27469 |
| -3.22 | -2.34 | 1.76 | -3.34 | DN3ab_ltr | DN3cd | -2.67 | -2.43 | 0.038 | 0.18 | 0.00029 | ENSMUSG00000024793 | protein_coding | Tnfrsf25 | tumor necrosis factor receptor superfamily, member 25 | 85030 | 4 | 152115934 | 152120119 | 1 | Mm.101198 |
| 6.79 | 6.58 | 6.5 | 7.16 | DN3ab_ltr | DN3cd | 0.2 | 1.22 | 0.25 | 0.55 | 3.00E-04 | ENSMUSG00000057110 | protein_coding | Cntrl | centriolin | 26920 | 2 | 35109492 | 35178822 | 1 | Mm.231332 |
| 6.82 | 6.73 | 6.65 | 7.3 | DN3ab_ltr | DN3cd | 0.26 | 1.9 | 0.091 | 0.31 | 3.00E-04 | ENSMUSG00000026750 | protein_coding | Psmb7 | proteasome (prosome, macropain) subunit, beta type 7 | 19177 | 2 | 38588036 | 38644087 | -1 | Mm.389251 |
| 6.74 | 6.56 | 6.74 | 7.53 | DN3ab_ltr | DN3cd | 0.31 | 1.72 | 0.12 | 0.37 | 3.00E-04 | ENSMUSG00000094974 | processed_pseudogene | Rps19-ps2 | ribosomal protein S19, pseudogene 2 | NA | 14 | 52902793 | 52903230 | 1 | |
| 4.65 | 3.55 | 4.3 | 4.5 | DN3ab_ltr | DN3cd | -0.44 | -1.81 | 0.1 | 0.34 | 3.00E-04 | ENSMUSG00000042063 | protein_coding | Zfp386 | zinc finger protein 386 (Kruppel-like) | 56220 | 12 | 116047724 | 116063360 | 1 | Mm.254997 |
| 6.88 | 7.29 | 5.98 | 7.32 | DN3ab_ltr | DN3cd | 0.8 | 4.64 | 0.0013 | 0.022 | 3.00E-04 | ENSMUSG00000022372 | protein_coding | Sla | src-like adaptor | 20491 | 15 | 66780819 | 66831829 | -1 | Mm.7601 |
| 1.21 | 3.36 | -2.9 | 3.47 | DN3ab_ltr | DN3cd | 3.1 | 4.71 | 0.0011 | 0.02 | 3.00E-04 | ENSMUSG00000020787 | protein_coding | P2rx1 | purinergic receptor P2X, ligand-gated ion channel, 1 | 18436 | 11 | 72999103 | 73015200 | 1 | Mm.25722 |
| 4.26 | 2.91 | 2.64 | 3.07 | DN3ab_ltr | DN3cd | -0.69 | -2.15 | 0.061 | 0.24 | 3.00E-04 | ENSMUSG00000045005 | protein_coding | Fzd5 | frizzled class receptor 5 | 14367 | 1 | 64730558 | 64737751 | -1 | Mm.150813 Mm.470210 |
| 5.93 | 5.97 | 4.65 | 6.68 | DN3ab_ltr | DN3cd | 0.84 | 2.32 | 0.046 | 0.21 | 3.00E-04 | ENSMUSG00000049717 | protein_coding | Lig4 | ligase IV, DNA, ATP-dependent | 319583 | 8 | 9969049 | 9977686 | -1 | Mm.80584 |
| 0.16 | -0.02 | 5.47 | 1.75 | DN3ab_ltr | DN3cd | -3.02 | -5.75 | 0.00029 | 0.0086 | 3.00E-04 | ENSMUSG00000022667 | protein_coding | Cd200r1 | CD200 receptor 1 | 57781 | 16 | 44765736 | 44794978 | 1 | Mm.416152 |
| 4.94 | 3.89 | 4.13 | 4.57 | DN3ab_ltr | DN3cd | -0.36 | -1.3 | 0.23 | 0.51 | 3.00E-04 | ENSMUSG00000032392 | protein_coding | Parp16 | poly (ADP-ribose) polymerase family, member 16 | 214424 | 9 | 65195353 | 65239224 | 1 | Mm.31129 |
| 3.9 | 4.01 | 2.82 | 4.67 | DN3ab_ltr | DN3cd | 0.78 | 2.46 | 0.036 | 0.18 | 0.00031 | ENSMUSG00000090626 | protein_coding | Tex9 | testis expressed gene 9 | 21778 | 9 | 72450394 | 72492212 | -1 | Mm.14485 |
| 7.72 | 7.69 | 8.13 | 7.41 | DN3ab_ltr | DN3cd | -0.35 | -2.69 | 0.025 | 0.14 | 0.00031 | ENSMUSG00000026430 | protein_coding | Rassf5 | Ras association (RalGDS/AF-6) domain family member 5 | 54354 | 1 | 131176410 | 131245258 | -1 | Mm.248291 |
| 8.7 | 7.57 | 8.81 | 8.38 | DN3ab_ltr | DN3cd | -0.78 | -5.96 | 0.00023 | 0.0074 | 0.00032 | ENSMUSG00000066406 | protein_coding | Akap13 | A kinase (PRKA) anchor protein 13 | 75547 | 7 | 75455534 | 75754609 | 1 | Mm.216107 |
| 5.3 | 5.43 | 4.31 | 5.49 | DN3ab_ltr | DN3cd | 0.55 | 2.83 | 0.02 | 0.13 | 0.00032 | ENSMUSG00000048285 | protein_coding | Frmd6 | FERM domain containing 6 | 319710 | 12 | 70825514 | 70902234 | 1 | Mm.2962 |
| 5.22 | 3.8 | 5.67 | 5.51 | DN3ab_ltr | DN3cd | -0.66 | -2.84 | 0.02 | 0.12 | 0.00032 | ENSMUSG00000057143 | protein_coding | Trim12c | tripartite motif-containing 12C | 319236 | 7 | 104338754 | 104353362 | -1 | Mm.425502 |
| 4.52 | 4.23 | 3.53 | 4.56 | DN3ab_ltr | DN3cd | 0.24 | 0.99 | 0.35 | 0.64 | 0.00032 | ENSMUSG00000050332 | protein_coding | Amer1 | APC membrane recruitment 1 | 72345 | X | 95420318 | 95444872 | -1 | Mm.182867 |
| 6.3 | 6.93 | 4.83 | 6.82 | DN3ab_ltr | DN3cd | 1.12 | 4.6 | 0.0013 | 0.022 | 0.00032 | ENSMUSG00000029275 | protein_coding | Gfi1 | growth factor independent 1 transcription repressor | 14581 | 5 | 107716657 | 107726036 | -1 | Mm.2078 Mm.453139 |
| 6.25 | 6.27 | 5.35 | 6.12 | DN3ab_ltr | DN3cd | 0.33 | 2.33 | 0.045 | 0.2 | 0.00033 | ENSMUSG00000021932 | protein_coding | Rnaseh2b | ribonuclease H2, subunit B | 67153 | 14 | 62292589 | 62372992 | 1 | Mm.26040 |
| 7.83 | 6.67 | 6.55 | 6.98 | DN3ab_ltr | DN3cd | -0.49 | -1.66 | 0.13 | 0.38 | 0.00033 | ENSMUSG00000047604 | protein_coding | Frat2 | frequently rearranged in advanced T cell lymphomas 2 | 212398 | 19 | 41845972 | 41848132 | -1 | Mm.222723 |
| 7.3 | 7.17 | 6.6 | 7.39 | DN3ab_ltr | DN3cd | 0.27 | 1.57 | 0.15 | 0.41 | 0.00033 | ENSMUSG00000004043 | protein_coding | Stat5a | signal transducer and activator of transcription 5A | 20850 | 11 | 100859351 | 100885169 | 1 | Mm.277403 |
| 3.56 | 3.45 | 2.22 | 4.21 | DN3ab_ltr | DN3cd | 0.65 | 1.72 | 0.12 | 0.37 | 0.00034 | ENSMUSG00000098120 | lincRNA | Gm5914 | predicted gene 5914 | NA | 8 | 105624336 | 105637485 | -1 | |
| 6.28 | 6.64 | 5.45 | 7.16 | DN3ab_ltr | DN3cd | 0.94 | 3.74 | 0.0047 | 0.049 | 0.00035 | ENSMUSG00000084129 | processed_pseudogene | Hmgb1-ps1 | high mobility group box 1, pseudogene 1 | NA | 11 | 41141763 | 41142396 | 1 | |
| 6.7 | 6.49 | 6.69 | 7.42 | DN3ab_ltr | DN3cd | 0.26 | 1.46 | 0.18 | 0.46 | 0.00035 | ENSMUSG00000096681 | processed_pseudogene | Rps19-ps1 | ribosomal protein S19, pseudogene 1 | NA | 14 | 53213730 | 53214167 | 1 | |
| 8.84 | 10 | 5.62 | 9.78 | DN3ab_ltr | DN3cd | 1.95 | 3.98 | 0.0033 | 0.039 | 0.00036 | ENSMUSG00000025014 | protein_coding | Dntt | deoxynucleotidyltransferase, terminal | 21673 | 19 | 41029275 | 41059523 | 1 | Mm.25620 |
| 4.67 | 4.93 | 3.21 | 4.69 | DN3ab_ltr | DN3cd | 0.66 | 3.08 | 0.013 | 0.097 | 0.00036 | ENSMUSG00000021391 | protein_coding | Cenpp | centromere protein P | 66336 | 13 | 49464023 | 49652785 | -1 | Mm.440323 |
| 3.39 | 3.45 | 0.07 | 3.4 | DN3ab_ltr | DN3cd | 0.61 | 1.33 | 0.22 | 0.5 | 0.00036 | ENSMUSG00000043460 | protein_coding | Elfn2 | leucine rich repeat and fibronectin type III, extracellular 2 | 207393 | 15 | 78667331 | 78718771 | -1 | Mm.323188 |
| 7.59 | 7.47 | 6.54 | 7.43 | DN3ab_ltr | DN3cd | 0.3 | 1.6 | 0.14 | 0.4 | 0.00037 | ENSMUSG00000022952 | protein_coding | Runx1 | runt related transcription factor 1 | 12394 | 16 | 92601466 | 92826149 | -1 | Mm.4081 Mm.470227 |
| 3.43 | 4.99 | -1.4 | 5.12 | DN3ab_ltr | DN3cd | 1.99 | 3.81 | 0.0043 | 0.045 | 0.00037 | ENSMUSG00000075330 | lincRNA | A930003A15Rik | RIKEN cDNA A930003A15 gene | NA | 16 | 19876567 | 19884274 | -1 | |
| 0.16 | 0.76 | 3.83 | 1.61 | DN3ab_ltr | DN3cd | -1.46 | -3.06 | 0.014 | 0.099 | 0.00037 | ENSMUSG00000039783 | protein_coding | Kmo | kynurenine 3-monooxygenase (kynurenine 3-hydroxylase) | 98256 | 1 | 175620381 | 175662116 | 1 | Mm.27217 |
| 7.21 | 7.27 | 9.91 | 8.65 | DN3ab_ltr | DN3cd | -0.78 | -3.32 | 0.0091 | 0.075 | 0.00037 | ENSMUSG00000000184 | protein_coding | Ccnd2 | cyclin D2 | 12444 | 6 | 127125162 | 127152193 | -1 | Mm.333406 |
| 6.59 | 6.19 | 7.61 | 6.26 | DN3ab_ltr | DN3cd | -0.88 | -4.96 | 0.00081 | 0.016 | 0.00038 | ENSMUSG00000032026 | protein_coding | Rexo2 | RNA exonuclease 2 | 104444 | 9 | 48468512 | 48480623 | -1 | Mm.21911 |
| 5.69 | 5.9 | 5.49 | 6.61 | DN3ab_ltr | DN3cd | 0.65 | 3.73 | 0.0048 | 0.049 | 0.00038 | ENSMUSG00000020423 | protein_coding | Btg2 | B cell translocation gene 2, anti-proliferative | 12227 | 1 | 134075170 | 134079120 | -1 | Mm.392646 |
| 4.58 | 4.38 | 3.71 | 4.59 | DN3ab_ltr | DN3cd | 0.24 | 1.17 | 0.27 | 0.57 | 0.00038 | ENSMUSG00000021756 | protein_coding | Il6st | interleukin 6 signal transducer | 16195 | 13 | 112464070 | 112510086 | 1 | Mm.4364 |
| 6.68 | 6.53 | 7.9 | 6.75 | DN3ab_ltr | DN3cd | -0.67 | -3.58 | 0.0061 | 0.057 | 0.00039 | ENSMUSG00000021109 | protein_coding | Hif1a | hypoxia inducible factor 1, alpha subunit | 15251 | 12 | 73901375 | 73947530 | 1 | Mm.3879 Mm.446610 |
| 8.22 | 8.37 | 7.15 | 7.85 | DN3ab_ltr | DN3cd | 0.36 | 3.53 | 0.0066 | 0.06 | 0.00039 | ENSMUSG00000029177 | protein_coding | Cenpa | centromere protein A | 12615 | 5 | 30666777 | 30674830 | 1 | Mm.290563 |
| 4.62 | 3.62 | 4.27 | 4.52 | DN3ab_ltr | DN3cd | -0.37 | -1.59 | 0.15 | 0.41 | 0.00039 | ENSMUSG00000106205 | TEC | C230096K16Rik | RIKEN cDNA C230096K16 gene | NA | 5 | 64804014 | 64805735 | 1 | |
| 7.97 | 8.41 | 6.72 | 8.02 | DN3ab_ltr | DN3cd | 0.77 | 4.89 | 0.00089 | 0.017 | 0.00039 | ENSMUSG00000052889 | protein_coding | Prkcb | protein kinase C, beta | 18751 | 7 | 122288751 | 122634402 | 1 | Mm.207496 Mm.446371 |
| 1.5 | 2.26 | -1.16 | 3.87 | DN3ab_ltr | DN3cd | 1.84 | 2.63 | 0.028 | 0.15 | 0.00039 | ENSMUSG00000108132 | lincRNA | Gm44175 | predicted gene, 44175 | NA | 6 | 71354351 | 71360829 | -1 | |
| 2.8 | 3.06 | 0.59 | 4.32 | DN3ab_ltr | DN3cd | 1.2 | 2.07 | 0.068 | 0.26 | 0.00039 | ENSMUSG00000022508 | protein_coding | Bcl6 | B cell leukemia/lymphoma 6 | 12053 | 16 | 23965052 | 23988852 | -1 | Mm.347398 |
| 4.6 | 3.17 | 4.11 | 4.2 | DN3ab_ltr | DN3cd | -0.65 | -2.27 | 0.05 | 0.22 | 0.00039 | ENSMUSG00000034640 | protein_coding | Tiparp | TCDD-inducible poly(ADP-ribose) polymerase | 99929 | 3 | 65528410 | 65555518 | 1 | Mm.246398 |
| 4.82 | 5.36 | 3.26 | 5.44 | DN3ab_ltr | DN3cd | 1.08 | 3.72 | 0.0049 | 0.05 | 4.00E-04 | ENSMUSG00000022353 | protein_coding | Mtss1 | metastasis suppressor 1 | 211401 | 15 | 58941234 | 59082005 | -1 | Mm.215481 Mm.394414 |
| 3.53 | 3.05 | 7.93 | 5.21 | DN3ab_ltr | DN3cd | -2.22 | -6.37 | 0.00014 | 0.0056 | 4.00E-04 | ENSMUSG00000037940 | protein_coding | Inpp4b | inositol polyphosphate-4-phosphatase, type II | 234515 | 8 | 81342556 | 82127914 | 1 | Mm.324468 |
| 6.13 | 6.15 | 6.96 | 6.21 | DN3ab_ltr | DN3cd | -0.37 | -2.5 | 0.034 | 0.17 | 0.00041 | ENSMUSG00000024038 | protein_coding | Ndufv3 | NADH:ubiquinone oxidoreductase core subunit V3 | 78330 | 17 | 31520115 | 31531326 | 1 | Mm.28349 |
| 5.3 | 4.53 | 5.17 | 5.29 | DN3ab_ltr | DN3cd | -0.32 | -1.93 | 0.086 | 0.3 | 0.00041 | ENSMUSG00000027130 | protein_coding | Slc12a6 | solute carrier family 12, member 6 | 107723 | 2 | 112265825 | 112363163 | 1 | Mm.491155 |
| 7.26 | 7.35 | 6.08 | 6.8 | DN3ab_ltr | DN3cd | 0.33 | 2.84 | 0.02 | 0.12 | 0.00041 | ENSMUSG00000027011 | protein_coding | Ube2e3 | ubiquitin-conjugating enzyme E2E 3 | 22193 | 2 | 78868124 | 78921293 | 1 | Mm.1485 Mm.393087 |
| 7.07 | 7.03 | 7.7 | 6.99 | DN3ab_ltr | DN3cd | -0.37 | -2.84 | 0.02 | 0.12 | 0.00042 | ENSMUSG00000015575 | protein_coding | Atp6v0e | ATPase, H+ transporting, lysosomal V0 subunit E | 11974 | 17 | 26676396 | 26699644 | 1 | Mm.22602 |
| 0.51 | 0.6 | 3.42 | 0.86 | DN3ab_ltr | DN3cd | -1.6 | -3.3 | 0.0094 | 0.077 | 0.00042 | ENSMUSG00000039713 | protein_coding | Plekhg5 | pleckstrin homology domain containing, family G (with RhoGef domain) member 5 | 269608 | 4 | 152072498 | 152115400 | 1 | Mm.332102 Mm.486442 |
| 4.84 | 4.69 | 3.95 | 4.95 | DN3ab_ltr | DN3cd | 0.31 | 1.47 | 0.18 | 0.45 | 0.00042 | ENSMUSG00000022210 | protein_coding | Dhrs4 | dehydrogenase/reductase (SDR family) member 4 | 28200 | 14 | 55478758 | 55490340 | 1 | Mm.27427 |
| 1.84 | 2.03 | 3.99 | 2.08 | DN3ab_ltr | DN3cd | -1.07 | -2.73 | 0.023 | 0.14 | 0.00042 | ENSMUSG00000028680 | protein_coding | Plk3 | polo like kinase 3 | 12795 | 4 | 117128655 | 117133963 | -1 | Mm.259022 |
| -0.84 | 1.88 | -3.72 | 2.52 | DN3ab_ltr | DN3cd | 4.27 | 6.42 | 0.00013 | 0.0054 | 0.00042 | ENSMUSG00000016262 | protein_coding | Sertad4 | SERTA domain containing 4 | 214791 | 1 | 192844488 | 192855756 | -1 | Mm.22806 |
| 8.37 | 8.27 | 7.56 | 8.53 | DN3ab_ltr | DN3cd | 0.36 | 1.81 | 0.1 | 0.34 | 0.00042 | ENSMUSG00000061887 | protein_coding | Ssbp3 | single-stranded DNA binding protein 3 | 72475 | 4 | 106910701 | 107049694 | 1 | Mm.195635 Mm.445960 Mm.474861 |
| 6.15 | 5.5 | 6.3 | 6.44 | DN3ab_ltr | DN3cd | -0.25 | -1.66 | 0.13 | 0.38 | 0.00042 | ENSMUSG00000036333 | protein_coding | Kidins220 | kinase D-interacting substrate 220 | 77480 | 12 | 24974925 | 25063152 | 1 | Mm.250641 Mm.475357 |
| 6.98 | 6.87 | 7.56 | 6.55 | DN3ab_ltr | DN3cd | -0.54 | -3.13 | 0.012 | 0.091 | 0.00043 | ENSMUSG00000032575 | protein_coding | Manf | mesencephalic astrocyte-derived neurotrophic factor | 74840 | 9 | 106838312 | 106891979 | -1 | Mm.29778 |
| 3.94 | 4.23 | 2.05 | 4.7 | DN3ab_ltr | DN3cd | 0.96 | 2.41 | 0.04 | 0.19 | 0.00043 | ENSMUSG00000021703 | protein_coding | Serinc5 | serine incorporator 5 | 218442 | 13 | 92611091 | 92711947 | 1 | Mm.297970 |
| 4.95 | 4.91 | 6.8 | 5.95 | DN3ab_ltr | DN3cd | -0.53 | -3.49 | 0.007 | 0.063 | 0.00044 | ENSMUSG00000009687 | protein_coding | Fxyd5 | FXYD domain-containing ion transport regulator 5 | 18301 | 7 | 31032722 | 31042481 | -1 | Mm.1870 |
| 6.09 | 6.31 | 4.96 | 7.22 | DN3ab_ltr | DN3cd | 1.06 | 2.83 | 0.02 | 0.13 | 0.00044 | ENSMUSG00000098243 | processed_transcript | Gm4258 | predicted gene 4258 | NA | 1 | 137898595 | 138000384 | 1 | |
| 7.46 | 6.6 | 7.4 | 7.24 | DN3ab_ltr | DN3cd | -0.52 | -3.97 | 0.0033 | 0.039 | 0.00044 | ENSMUSG00000026034 | protein_coding | Clk1 | CDC-like kinase 1 | 12747 | 1 | 58410189 | 58424066 | -1 | Mm.1761 |
| 8.2 | 8.44 | 7.48 | 8.41 | DN3ab_ltr | DN3cd | 0.53 | 4.05 | 0.003 | 0.036 | 0.00045 | ENSMUSG00000029687 | protein_coding | Ezh2 | enhancer of zeste 2 polycomb repressive complex 2 subunit | 14056 | 6 | 47530139 | 47595341 | -1 | Mm.246688 |
| 6.55 | 6.59 | 5.74 | 6.92 | DN3ab_ltr | DN3cd | 0.53 | 2.51 | 0.034 | 0.17 | 0.00045 | ENSMUSG00000020297 | protein_coding | Nsg2 | neuron specific gene family member 2 | 18197 | 11 | 32000463 | 32059202 | 1 | Mm.3304 |
| 6.51 | 6.56 | 5.84 | 6.73 | DN3ab_ltr | DN3cd | 0.41 | 2.65 | 0.027 | 0.15 | 0.00045 | ENSMUSG00000026019 | protein_coding | Wdr12 | WD repeat domain 12 | 57750 | 1 | 60069785 | 60098645 | -1 | Mm.281079 |
| 5.58 | 5.28 | 9.54 | 7.47 | DN3ab_ltr | DN3cd | -1.57 | -5.24 | 0.00056 | 0.013 | 0.00045 | ENSMUSG00000041954 | protein_coding | Tnfrsf18 | tumor necrosis factor receptor superfamily, member 18 | 21936 | 4 | 156026164 | 156028895 | 1 | Mm.3180 |
| 4.77 | 3.87 | 5.16 | 5.24 | DN3ab_ltr | DN3cd | -0.34 | -1.85 | 0.098 | 0.33 | 0.00046 | ENSMUSG00000057554 | protein_coding | Lgals8 | lectin, galactose binding, soluble 8 | 56048 | 13 | 12439415 | 12464944 | -1 | Mm.474138 Mm.486765 |
| 4.52 | 4.8 | 2.07 | 4.44 | DN3ab_ltr | DN3cd | 0.76 | 2.27 | 0.05 | 0.22 | 0.00046 | ENSMUSG00000036412 | protein_coding | Arsi | arylsulfatase i | 545260 | 18 | 60911780 | 60918561 | 1 | Mm.20147 |
| 3.11 | 3.96 | 3.74 | 6.19 | DN3ab_ltr | DN3cd | 1.78 | 5.97 | 0.00022 | 0.0073 | 0.00046 | ENSMUSG00000065824 | misc_RNA | Gm26315 | predicted gene, 26315 | NA | 2 | 7076094 | 7076414 | -1 | |
| 3.26 | 3 | 2.53 | 4.32 | DN3ab_ltr | DN3cd | 0.67 | 1.73 | 0.12 | 0.36 | 0.00047 | ENSMUSG00000059146 | protein_coding | Ntrk3 | neurotrophic tyrosine kinase, receptor, type 3 | 18213 | 7 | 78175959 | 78738012 | -1 | Mm.119373 Mm.33496 Mm.421361 |
| 10.92 | 10.92 | 9.54 | 10.46 | DN3ab_ltr | DN3cd | 0.34 | 2.06 | 0.07 | 0.27 | 0.00048 | ENSMUSG00000015749 | protein_coding | Anp32e | acidic (leucine-rich) nuclear phosphoprotein 32 family, member E | 66471 | 3 | 95929246 | 95947390 | 1 | Mm.218657 |
| 4.79 | 5.12 | 3.57 | 5.1 | DN3ab_ltr | DN3cd | 0.76 | 3.46 | 0.0073 | 0.065 | 0.00048 | ENSMUSG00000062510 | protein_coding | Nsl1 | NSL1, MIS12 kinetochore complex component | 381318 | 1 | 191063012 | 191086474 | 1 | Mm.32504 |
| 7.09 | 6.94 | 6.01 | 6.96 | DN3ab_ltr | DN3cd | 0.3 | 1.49 | 0.17 | 0.44 | 0.00048 | ENSMUSG00000098318 | lincRNA | Lockd | lncRNA downstream of Cdkn1b | NA | 6 | 134929092 | 134956798 | 1 | |
| 8.17 | 8.04 | 8.96 | 7.89 | DN3ab_ltr | DN3cd | -0.59 | -3.3 | 0.0095 | 0.077 | 0.00048 | ENSMUSG00000024048 | protein_coding | Myl12a | myosin, light chain 12A, regulatory, non-sarcomeric | 67268 | 17 | 70993656 | 71002878 | -1 | Mm.489605 |
| 7.38 | 7.68 | 6.3 | 7.24 | DN3ab_ltr | DN3cd | 0.56 | 4.64 | 0.0013 | 0.021 | 0.00048 | ENSMUSG00000027469 | protein_coding | Tpx2 | TPX2, microtubule-associated | 72119 | 2 | 152847964 | 152895321 | 1 | Mm.407737 |
| 9.11 | 9.17 | 9.22 | 8.88 | DN3ab_ltr | DN3cd | -0.13 | -1.67 | 0.13 | 0.38 | 0.00049 | ENSMUSG00000057322 | protein_coding | Rpl38 | ribosomal protein L38 | 67671 | 11 | 114668524 | 114672331 | 1 | Mm.238817 Mm.371629 |
| 4.92 | 4.84 | 6.26 | 5.19 | DN3ab_ltr | DN3cd | -0.62 | -3.33 | 0.0089 | 0.074 | 0.00049 | ENSMUSG00000039942 | protein_coding | Ptger4 | prostaglandin E receptor 4 (subtype EP4) | 19219 | 15 | 5206661 | 5244187 | -1 | Mm.18509 |
| 2.8 | 2.54 | 5.73 | 3.91 | DN3ab_ltr | DN3cd | -1.38 | -5.14 | 0.00064 | 0.014 | 0.00049 | ENSMUSG00000064023 | protein_coding | Klk8 | kallikrein related-peptidase 8 | 259277 | 7 | 43797577 | 43803826 | 1 | Mm.458023 |
| 7.77 | 8.14 | 6.88 | 8.09 | DN3ab_ltr | DN3cd | 0.72 | 4.63 | 0.0013 | 0.022 | 5.00E-04 | ENSMUSG00000026918 | protein_coding | Brd3 | bromodomain containing 3 | 67382 | 2 | 27445579 | 27507662 | -1 | Mm.28721 |
| 6.35 | 6.15 | 5.3 | 6.24 | DN3ab_ltr | DN3cd | 0.27 | 1.27 | 0.24 | 0.52 | 5.00E-04 | ENSMUSG00000024219 | protein_coding | Anks1 | ankyrin repeat and SAM domain containing 1 | 224650 | 17 | 27909306 | 28062775 | 1 | Mm.32556 |
| 7.43 | 7.44 | 7.13 | 7.83 | DN3ab_ltr | DN3cd | 0.33 | 2.58 | 0.03 | 0.16 | 5.00E-04 | ENSMUSG00000021495 | protein_coding | Fam193b | family with sequence similarity 193, member B | 212483 | 13 | 55539316 | 55571120 | -1 | Mm.35661 |
| 4.56 | 4.62 | 2.01 | 4.29 | DN3ab_ltr | DN3cd | 0.57 | 1.63 | 0.14 | 0.39 | 5.00E-04 | ENSMUSG00000057606 | protein_coding | Colq | collagen-like tail subunit (single strand of homotrimer) of asymmetric acetylcholinesterase | 382864 | 14 | 31523082 | 31591343 | -1 | Mm.239821 |
| 3.29 | 3.46 | 0.91 | 4.07 | DN3ab_ltr | DN3cd | 0.89 | 1.83 | 0.1 | 0.33 | 0.00051 | ENSMUSG00000029279 | protein_coding | Brdt | bromodomain, testis-specific | 114642 | 5 | 107331159 | 107387058 | 1 | Mm.182836 |
| 13.19 | 13.21 | 13.04 | 13.74 | DN3ab_ltr | DN3cd | 0.34 | 2.72 | 0.024 | 0.14 | 0.00051 | ENSMUSG00000064339 | Mt_rRNA | mt-Rnr2 | mitochondrially encoded 16S rRNA | NA | MT | 1094 | 2675 | 1 | |
| 5.48 | 5.85 | 4.08 | 5.5 | DN3ab_ltr | DN3cd | 0.73 | 3.85 | 0.004 | 0.044 | 0.00051 | ENSMUSG00000039396 | protein_coding | Neil3 | nei like 3 (E. coli) | 234258 | 8 | 53586867 | 53639065 | -1 | Mm.281749 |
| 6.36 | 7.57 | 6.06 | 8.31 | DN3ab_ltr | DN3cd | 1.7 | 8.73 | 1.20E-05 | 0.0013 | 0.00052 | ENSMUSG00000007872 | protein_coding | Id3 | inhibitor of DNA binding 3 | 15903 | 4 | 136143497 | 136145755 | 1 | Mm.110 |
| 9.11 | 9.13 | 8.69 | 9.19 | DN3ab_ltr | DN3cd | 0.23 | 2.51 | 0.034 | 0.17 | 0.00052 | ENSMUSG00000022858 | protein_coding | Tra2b | transformer 2 beta | 20462 | 16 | 22244549 | 22266005 | -1 | Mm.210352 Mm.389440 Mm.392922 Mm.442193 |
| 7.18 | 7.37 | 6.47 | 7.44 | DN3ab_ltr | DN3cd | 0.52 | 3.63 | 0.0056 | 0.054 | 0.00052 | ENSMUSG00000020166 | protein_coding | Cnot2 | CCR4-NOT transcription complex, subunit 2 | 72068 | 10 | 116485161 | 116581511 | -1 | Mm.351553 |
| 5.59 | 5.65 | 4.66 | 5.6 | DN3ab_ltr | DN3cd | 0.41 | 2.51 | 0.034 | 0.17 | 0.00052 | ENSMUSG00000022070 | protein_coding | Bora | bora, aurora kinase A activator | 77744 | 14 | 99046222 | 99074540 | 1 | Mm.23503 |
| 6.46 | 6.57 | 6.79 | 6.26 | DN3ab_ltr | DN3cd | -0.19 | -1.53 | 0.16 | 0.42 | 0.00053 | ENSMUSG00000022174 | protein_coding | Dad1 | defender against cell death 1 | 13135 | 14 | 54235479 | 54254104 | -1 | Mm.319038 |
| 4.07 | 3.93 | 8.02 | 4.93 | DN3ab_ltr | DN3cd | -2.15 | -4.16 | 0.0025 | 0.033 | 0.00053 | ENSMUSG00000027639 | protein_coding | Samhd1 | SAM domain and HD domain, 1 | 56045 | 2 | 157097533 | 157135265 | -1 | Mm.248478 Mm.468781 |
| 2.6 | 2.99 | 0.4 | 3.19 | DN3ab_ltr | DN3cd | 0.99 | 2.49 | 0.035 | 0.17 | 0.00053 | ENSMUSG00000057265 | protein_coding | Bbof1 | basal body orientation factor 1 | 72873 | 12 | 84409071 | 84442887 | 1 | Mm.440428 |
| 6.75 | 6.6 | 7.41 | 6.46 | DN3ab_ltr | DN3cd | -0.54 | -3.5 | 0.0068 | 0.062 | 0.00054 | ENSMUSG00000005103 | protein_coding | Wdr1 | WD repeat domain 1 | 22388 | 5 | 38526813 | 38563221 | -1 | Mm.2654 Mm.393965 Mm.406120 |
| 3.1 | 1.28 | 3.89 | 3.72 | DN3ab_ltr | DN3cd | -0.64 | -2.27 | 0.05 | 0.22 | 0.00054 | ENSMUSG00000099757 | processed_transcript | BE692007 | expressed sequence BE692007 | NA | 19 | 11470166 | 11484578 | -1 | |
| 6.78 | 6.58 | 5.81 | 7.04 | DN3ab_ltr | DN3cd | 0.42 | 1.56 | 0.15 | 0.42 | 0.00054 | ENSMUSG00000049295 | protein_coding | Zfp219 | zinc finger protein 219 | 69890 | 14 | 52006077 | 52020733 | -1 | Mm.475652 |
| 4.45 | 3.34 | 4.46 | 4.63 | DN3ab_ltr | DN3cd | -0.4 | -1.67 | 0.13 | 0.38 | 0.00054 | ENSMUSG00000083041 | processed_pseudogene | Gmfg-ps | glia maturation factor, gamma, pseudogene | NA | 6 | 4893352 | 4893778 | -1 | |
| 7.85 | 8.14 | 6.35 | 7.36 | DN3ab_ltr | DN3cd | 0.55 | 4.14 | 0.0026 | 0.033 | 0.00054 | ENSMUSG00000041431 | protein_coding | Ccnb1 | cyclin B1 | 268697 | 13 | 100778650 | 100786570 | -1 | Mm.260114 Mm.380027 Mm.482545 |
| 8.71 | 8.77 | 8.19 | 9.02 | DN3ab_ltr | DN3cd | 0.4 | 2.81 | 0.021 | 0.13 | 0.00054 | ENSMUSG00000020863 | protein_coding | Luc7l3 | LUC7-like 3 (S. cerevisiae) | 67684 | 11 | 94287890 | 94321988 | -1 | Mm.30927 |
| 6.61 | 6.65 | 6.19 | 6.87 | DN3ab_ltr | DN3cd | 0.33 | 2.69 | 0.025 | 0.14 | 0.00055 | ENSMUSG00000038324 | protein_coding | Trpc4ap | transient receptor potential cation channel, subfamily C, member 4 associated protein | 56407 | 2 | 155634271 | 155692384 | -1 | Mm.234944 Mm.268304 Mm.469980 |
| 1.16 | 2.22 | -1.01 | 4.64 | DN3ab_ltr | DN3cd | 2.41 | 3.04 | 0.014 | 0.1 | 0.00056 | ENSMUSG00000086291 | sense_intronic | Gm15513 | predicted gene 15513 | NA | 5 | 34211810 | 34213802 | -1 | |
| 6.45 | 6.77 | 5.22 | 6.44 | DN3ab_ltr | DN3cd | 0.66 | 3.99 | 0.0033 | 0.038 | 0.00056 | ENSMUSG00000048327 | protein_coding | Ckap2l | cytoskeleton associated protein 2-like | 70466 | 2 | 129268210 | 129297212 | -1 | Mm.45785 Mm.491209 |
| 7.43 | 7.28 | 6.29 | 7.52 | DN3ab_ltr | DN3cd | 0.42 | 1.65 | 0.13 | 0.39 | 0.00058 | ENSMUSG00000030541 | protein_coding | Idh2 | isocitrate dehydrogenase 2 (NADP+), mitochondrial | 269951 | 7 | 80094846 | 80115392 | -1 | Mm.246432 Mm.290925 |
| 6.07 | 6.12 | 4.99 | 5.91 | DN3ab_ltr | DN3cd | 0.39 | 2.39 | 0.041 | 0.19 | 0.00058 | ENSMUSG00000027331 | protein_coding | Knstrn | kinetochore-localized astrin/SPAG5 binding | 51944 | 2 | 118814003 | 118853957 | 1 | Mm.252695 |
| 6.43 | 6.66 | 8.01 | 6.85 | DN3ab_ltr | DN3cd | -0.52 | -1.97 | 0.08 | 0.29 | 0.00059 | ENSMUSG00000020572 | protein_coding | Nampt | nicotinamide phosphoribosyltransferase | 59027 | 12 | 32819545 | 32853349 | 1 | Mm.202727 |
| 5.78 | 5.81 | 3.5 | 5.55 | DN3ab_ltr | DN3cd | 0.61 | 1.77 | 0.11 | 0.35 | 0.00059 | ENSMUSG00000004105 | protein_coding | Angptl2 | angiopoietin-like 2 | 26360 | 2 | 33216069 | 33247717 | 1 | Mm.208919 |
| 2.39 | 2.33 | 0.9 | 2.93 | DN3ab_ltr | DN3cd | 0.66 | 1.71 | 0.12 | 0.37 | 6.00E-04 | ENSMUSG00000030446 | protein_coding | Zfp273 | zinc finger protein 273 | 212569 | 13 | 67813740 | 67827002 | 1 | Mm.358842 |
| 4.41 | 4.29 | 3.39 | 4.7 | DN3ab_ltr | DN3cd | 0.44 | 1.65 | 0.13 | 0.39 | 0.00062 | ENSMUSG00000039741 | protein_coding | Bahcc1 | BAH domain and coiled-coil containing 1 | 268515 | 11 | 120232947 | 120292296 | 1 | Mm.309498 |
| 3.59 | 3.92 | 1.37 | 4.03 | DN3ab_ltr | DN3cd | 0.9 | 2.35 | 0.044 | 0.2 | 0.00062 | ENSMUSG00000009739 | protein_coding | Pou6f1 | POU domain, class 6, transcription factor 1 | 19009 | 15 | 100575318 | 100599984 | -1 | Mm.28825 |
| 5.08 | 5.02 | 3.96 | 4.83 | DN3ab_ltr | DN3cd | 0.29 | 1.69 | 0.13 | 0.37 | 0.00063 | ENSMUSG00000032235 | protein_coding | Ice2 | interactor of little elongation complex ELL subunit 2 | 93697 | 9 | 69397906 | 69433122 | 1 | Mm.35911 |
| 3.53 | 2.62 | 9.67 | 5.66 | DN3ab_ltr | DN3cd | -3.53 | -8.3 | 1.80E-05 | 0.0017 | 0.00063 | ENSMUSG00000050335 | protein_coding | Lgals3 | lectin, galactose binding, soluble 3 | 16854 | 14 | 47367751 | 47386160 | 1 | Mm.248615 |
| -4.01 | 0.53 | -1.4 | -0.8 | DN3ab_ltr | DN3cd | 2.87 | 3.81 | 0.0043 | 0.046 | 0.00064 | ENSMUSG00000104606 | lincRNA | Gm43409 | predicted gene 43409 | NA | 5 | 123188701 | 123191588 | -1 | |
| 6.97 | 6.46 | 8.58 | 6.94 | DN3ab_ltr | DN3cd | -1.13 | -5.25 | 0.00056 | 0.013 | 0.00065 | ENSMUSG00000032434 | protein_coding | Cmtm6 | CKLF-like MARVEL transmembrane domain containing 6 | 67213 | 9 | 114731116 | 114749344 | 1 | Mm.28858 |
| 4.32 | 4.29 | 3.27 | 5 | DN3ab_ltr | DN3cd | 0.67 | 2.04 | 0.072 | 0.27 | 0.00065 | ENSMUSG00000030341 | protein_coding | Tnfrsf1a | tumor necrosis factor receptor superfamily, member 1a | 21937 | 6 | 125349362 | 125362484 | 1 | Mm.1258 Mm.474976 |
| 7.46 | 6.49 | 7.13 | 6.89 | DN3ab_ltr | DN3cd | -0.63 | -4.44 | 0.0017 | 0.026 | 0.00065 | ENSMUSG00000035000 | protein_coding | Dpp4 | dipeptidylpeptidase 4 | 13482 | 2 | 62330073 | 62412231 | -1 | Mm.1151 |
| 6.07 | 5.9 | 6.79 | 5.8 | DN3ab_ltr | DN3cd | -0.58 | -3.69 | 0.0051 | 0.051 | 0.00066 | ENSMUSG00000023791 | protein_coding | Pigx | phosphatidylinositol glycan anchor biosynthesis, class X | 72084 | 16 | 32084416 | 32099740 | -1 | Mm.37613 |
| 3.59 | 4.04 | 2.46 | 4.42 | DN3ab_ltr | DN3cd | 0.98 | 3.52 | 0.0067 | 0.061 | 0.00066 | ENSMUSG00000026004 | protein_coding | Kansl1l | KAT8 regulatory NSL complex subunit 1-like | 68691 | 1 | 66719248 | 66817562 | -1 | Mm.240912 Mm.383276 Mm.461139 Mm.482481 |
| 6.09 | 5.97 | 5.81 | 6.47 | DN3ab_ltr | DN3cd | 0.25 | 1.66 | 0.13 | 0.38 | 0.00066 | ENSMUSG00000043241 | protein_coding | Upf2 | UPF2 regulator of nonsense transcripts homolog (yeast) | 326622 | 2 | 5951469 | 6056703 | 1 | Mm.31102 |
| 0.89 | 3.15 | -2.9 | 3.43 | DN3ab_ltr | DN3cd | 3.3 | 4.87 | 0.00093 | 0.017 | 0.00067 | ENSMUSG00000084993 | sense_intronic | Gm12305 | predicted gene 12305 | NA | 11 | 68269558 | 68270879 | -1 | |
| 0.23 | -3.92 | -1.19 | -0.2 | DN3ab_ltr | DN3cd | -1.8 | -1.83 | 0.1 | 0.33 | 0.00067 | ENSMUSG00000095530 | snoRNA | Snora19 | small nucleolar RNA, H/ACA box 19 | NA | 19 | 60774330 | 60774461 | -1 | |
| 7.65 | 7.57 | 8.16 | 7.32 | DN3ab_ltr | DN3cd | -0.43 | -2.97 | 0.016 | 0.11 | 0.00067 | ENSMUSG00000028670 | protein_coding | Lypla2 | lysophospholipase 2 | 26394 | 4 | 135968224 | 135972626 | -1 | Mm.34302 |
| 3.83 | 4.54 | 2.36 | 5.3 | DN3ab_ltr | DN3cd | 1.42 | 3.61 | 0.0058 | 0.056 | 0.00067 | ENSMUSG00000027165 | protein_coding | B230118H07Rik | RIKEN cDNA B230118H07 gene | 68170 | 2 | 101560781 | 101649532 | -1 | Mm.31263 |
| 8.95 | 9.1 | 8.35 | 9.18 | DN3ab_ltr | DN3cd | 0.44 | 3.42 | 0.0078 | 0.068 | 0.00068 | ENSMUSG00000018666 | protein_coding | Cbx1 | chromobox 1 | 12412 | 11 | 96789127 | 96808640 | 1 | Mm.29055 |
| 2.69 | 2.92 | 1.38 | 4.62 | DN3ab_ltr | DN3cd | 1.36 | 2.46 | 0.036 | 0.18 | 0.00068 | ENSMUSG00000029205 | protein_coding | Chrna9 | cholinergic receptor, nicotinic, alpha polypeptide 9 | 231252 | 5 | 65934921 | 65977326 | 1 | Mm.140015 Mm.486671 |
| 7.85 | 7.83 | 7.65 | 8.25 | DN3ab_ltr | DN3cd | 0.28 | 2.35 | 0.044 | 0.2 | 0.00069 | ENSMUSG00000021025 | protein_coding | Nfkbia | nuclear factor of kappa light polypeptide gene enhancer in B cells inhibitor, alpha | 18035 | 12 | 55489410 | 55492647 | -1 | Mm.170515 |
| 8.96 | 8.57 | 8.63 | 8.69 | DN3ab_ltr | DN3cd | -0.18 | -2.15 | 0.061 | 0.24 | 0.00069 | ENSMUSG00000028410 | protein_coding | Dnaja1 | DnaJ heat shock protein family (Hsp40) member A1 | 15502 | 4 | 40722150 | 40737149 | 1 | Mm.27897 |
| 5.84 | 6.25 | 5.38 | 6.53 | DN3ab_ltr | DN3cd | 0.74 | 5.26 | 0.00055 | 0.013 | 0.00069 | ENSMUSG00000068735 | protein_coding | Trp53i11 | transformation related protein 53 inducible protein 11 | 277414 | 2 | 93187548 | 93201759 | 1 | Mm.41033 |
| 7.98 | 8.02 | 8.36 | 7.68 | DN3ab_ltr | DN3cd | -0.3 | -2.17 | 0.059 | 0.24 | 0.00069 | ENSMUSG00000021670 | protein_coding | Hmgcr | 3-hydroxy-3-methylglutaryl-Coenzyme A reductase | 15357 | 13 | 96648967 | 96670936 | -1 | Mm.316652 |
| 8.25 | 7.68 | 9.46 | 7.82 | DN3ab_ltr | DN3cd | -1.12 | -5.53 | 0.00038 | 0.01 | 7.00E-04 | ENSMUSG00000025366 | protein_coding | Esyt1 | extended synaptotagmin-like protein 1 | 23943 | 10 | 128509965 | 128525871 | -1 | Mm.66056 |
| 7.07 | 7.56 | 5.95 | 7.14 | DN3ab_ltr | DN3cd | 0.76 | 5.81 | 0.00027 | 0.0083 | 0.00072 | ENSMUSG00000030677 | protein_coding | Kif22 | kinesin family member 22 | 110033 | 7 | 127027729 | 127042471 | -1 | Mm.286488 |
| 4.17 | 4.13 | 5.84 | 4.47 | DN3ab_ltr | DN3cd | -0.79 | -3.14 | 0.012 | 0.091 | 0.00072 | ENSMUSG00000057762 | protein_coding | Gm6169 | predicted gene 6169 | NA | 13 | 97098208 | 97099237 | -1 | |
| 9.54 | 9.77 | 9.13 | 9.77 | DN3ab_ltr | DN3cd | 0.41 | 5.21 | 0.00058 | 0.013 | 0.00072 | ENSMUSG00000004880 | protein_coding | Lbr | lamin B receptor | 98386 | 1 | 181815335 | 181843046 | -1 | Mm.4538 |
| 6.32 | 5.87 | 7.23 | 6.08 | DN3ab_ltr | DN3cd | -0.81 | -5.94 | 0.00023 | 0.0075 | 0.00073 | ENSMUSG00000016087 | protein_coding | Fli1 | Friend leukemia integration 1 | 14247 | 9 | 32422204 | 32542861 | -1 | Mm.258908 |
| 4.54 | 4.12 | 5.52 | 3.99 | DN3ab_ltr | DN3cd | -0.97 | -4.53 | 0.0015 | 0.024 | 0.00073 | ENSMUSG00000059714 | protein_coding | Flot1 | flotillin 1 | 14251 | 17 | 35823230 | 35832791 | 1 | Mm.2931 |
| 5.03 | 4.59 | 9.85 | 7.31 | DN3ab_ltr | DN3cd | -2.05 | -6.1 | 0.00019 | 0.0067 | 0.00074 | ENSMUSG00000001025 | protein_coding | S100a6 | S100 calcium binding protein A6 (calcyclin) | 20200 | 3 | 90612882 | 90624181 | 1 | Mm.100144 |
| 6.72 | 6.49 | 7.78 | 6.5 | DN3ab_ltr | DN3cd | -0.77 | -3.81 | 0.0042 | 0.045 | 0.00074 | ENSMUSG00000028843 | protein_coding | Sh3bgrl3 | SH3 domain binding glutamic acid-rich protein-like 3 | 73723 | 4 | 134127406 | 134128789 | -1 | Mm.22240 |
| 6.33 | 5.91 | 6.86 | 5.79 | DN3ab_ltr | DN3cd | -0.73 | -5.86 | 0.00025 | 0.0079 | 0.00074 | ENSMUSG00000024965 | protein_coding | Fermt3 | fermitin family member 3 | 108101 | 19 | 6998958 | 7019469 | -1 | Mm.157591 |
| 6.52 | 6.48 | 5.97 | 6.67 | DN3ab_ltr | DN3cd | 0.28 | 2.01 | 0.076 | 0.28 | 0.00076 | ENSMUSG00000056763 | protein_coding | Cspp1 | centrosome and spindle pole associated protein 1 | 211660 | 1 | 10037987 | 10136768 | 1 | Mm.244046 |
| 7.6 | 6.62 | 7.04 | 7.02 | DN3ab_ltr | DN3cd | -0.54 | -2.93 | 0.017 | 0.11 | 0.00077 | ENSMUSG00000035150 | protein_coding | Eif2s3x | eukaryotic translation initiation factor 2, subunit 3, structural gene X-linked | 26905 | X | 94188707 | 94212862 | -1 | Mm.218851 |
| 4.7 | 6.13 | 2.61 | 5.83 | DN3ab_ltr | DN3cd | 1.87 | 6.28 | 0.00015 | 0.0059 | 0.00077 | ENSMUSG00000031785 | protein_coding | Adgrg1 | adhesion G protein-coupled receptor G1 | 14766 | 8 | 94974751 | 95014217 | 1 | Mm.290834 |
| 3.14 | 3 | 1.67 | 4.16 | DN3ab_ltr | DN3cd | 0.82 | 1.69 | 0.13 | 0.38 | 0.00077 | ENSMUSG00000037579 | protein_coding | Kcnh3 | potassium voltage-gated channel, subfamily H (eag-related), member 3 | 16512 | 15 | 99224861 | 99242817 | 1 | Mm.374793 |
| 2.64 | 2.84 | 1.96 | 4.03 | DN3ab_ltr | DN3cd | 1.01 | 2.8 | 0.021 | 0.13 | 0.00078 | ENSMUSG00000040093 | protein_coding | Bmf | BCL2 modifying factor | 171543 | 2 | 118528757 | 118549687 | -1 | Mm.210125 |
| 1.38 | 1.08 | 2.85 | 0.17 | DN3ab_ltr | DN3cd | -1.38 | -2.99 | 0.015 | 0.11 | 0.00078 | ENSMUSG00000010751 | protein_coding | Tnfrsf22 | tumor necrosis factor receptor superfamily, member 22 | 79202 | 7 | 143634806 | 143649661 | -1 | Mm.261384 |
| 6.4 | 6.32 | 5.73 | 6.37 | DN3ab_ltr | DN3cd | 0.23 | 1.66 | 0.13 | 0.38 | 0.00079 | ENSMUSG00000041740 | protein_coding | Rnf10 | ring finger protein 10 | 50849 | 5 | 115241412 | 115272898 | -1 | Mm.30051 |
| 6.71 | 6.79 | 6.16 | 6.89 | DN3ab_ltr | DN3cd | 0.37 | 2.99 | 0.015 | 0.11 | 0.00082 | ENSMUSG00000059409 | protein_coding | Ppp2r5d | protein phosphatase 2, regulatory subunit B', delta | 21770 | 17 | 46682991 | 46705111 | -1 | Mm.295009 Mm.329083 Mm.443214 |

| | | | | | | | | | | | | | | | | | | | | | |
|---|---|---|---|---|---|---|---|---|---|---|---|---|---|---|---|---|---|---|---|---|---|
| 6.36 | 6.64 | 5.18 | 6.95 | DN3ab_Itr | DN3cd | 0.88 | 3.16 | 0.012 | 0.089 | 0.00082 | ENSMUSG00000051934 | protein_coding | Spats2 | spermatogenesis associated, serine-rich 2 | 72572 | | 15 | 99125916 | 99213215 | 1 | Mm.276650 |
| 0.55 | 3.44 | -2.9 | 3.91 | DN3ab_Itr | DN3cd | 4 | 5.86 | 0.00025 | 0.0079 | 0.00082 | ENSMUSG00000028544 | protein_coding | Slc5a9 | solute carrier family 5 (sodium/glucose cotransporter), member 9 | 230612 | | 4 | 111875375 | 111902918 | -1 | Mm.26630 |
| 3.36 | 3.24 | 1.98 | 3.64 | DN3ab_Itr | DN3cd | 0.51 | 1.54 | 0.16 | 0.42 | 0.00082 | ENSMUSG00000106951 | lincRNA | 5930430L01Rik | RIKEN cDNA 5930430L01 gene | NA | | 5 | 148990056 | 148995223 | -1 | |
| 9.62 | 10.14 | 8.57 | 9.62 | DN3ab_Itr | DN3cd | 0.73 | 7.28 | 5.00E-05 | 0.003 | 0.00083 | ENSMUSG00000056394 | protein_coding | Lig1 | ligase I, DNA, ATP-dependent | 16881 | | 7 | 13277283 | 13311433 | 1 | Mm.288179 Mm.421129 |
| 5.72 | 5.45 | 6.83 | 5.7 | DN3ab_Itr | DN3cd | -0.72 | -4.31 | 0.002 | 0.029 | 0.00083 | ENSMUSG00000034765 | protein_coding | Dusp5 | dual specificity phosphatase 5 | 240672 | | 19 | 53529109 | 53542431 | 1 | Mm.482424 |
| 4.03 | 4.61 | 2.05 | 4.94 | DN3ab_Itr | DN3cd | 1.21 | 3.06 | 0.014 | 0.099 | 0.00085 | ENSMUSG00000030283 | protein_coding | St8sia1 | ST8 alpha-N-acetyl-neuraminide alpha-2,8-sialyltransferase 1 | 20449 | | 6 | 142821545 | 142964452 | -1 | Mm.260838 |
| 5.3 | 4.39 | 5.14 | 5.22 | DN3ab_Itr | DN3cd | -0.41 | -2.12 | 0.063 | 0.25 | 0.00085 | ENSMUSG00000040652 | protein_coding | Oaz2 | ornithine decarboxylase antizyme 2 | 18247 | | 9 | 65668001 | 65690300 | 1 | Mm.116749 |
| -4.01 | -3.92 | -0.18 | -4.02 | DN3ab_Itr | DN3cd | -1.95 | -2.52 | 0.033 | 0.17 | 0.00086 | ENSMUSG00000028435 | protein_coding | Aqp3 | aquaporin 3 | 11828 | | 4 | 41092722 | 41098183 | -1 | Mm.34043 |
| 7.86 | 8.07 | 7.13 | 7.85 | DN3ab_Itr | DN3cd | 0.43 | 4.35 | 0.0019 | 0.028 | 0.00088 | ENSMUSG00000020496 | protein_coding | Rnf187 | ring finger protein 187 | 108660 | | 11 | 58932288 | 58938916 | -1 | Mm.249986 |
| 2.98 | 3.1 | 1.13 | 2.97 | DN3ab_Itr | DN3cd | 0.6 | 1.99 | 0.078 | 0.29 | 0.00088 | ENSMUSG00000044807 | protein_coding | Zfp354c | zinc finger protein 354C | 30944 | | 11 | 50811086 | 50827724 | -1 | Mm.103674 |
| 3.9 | 4.33 | 1.52 | 3.53 | DN3ab_Itr | DN3cd | 0.77 | 3.05 | 0.014 | 0.1 | 0.00089 | ENSMUSG00000042155 | protein_coding | Klhl23 | kelch-like 23 | 277396 | | 2 | 69821944 | 69836651 | 1 | Mm.138073 |
| 6.76 | 6.85 | 5.87 | 6.97 | DN3ab_Itr | DN3cd | 0.51 | 2.67 | 0.026 | 0.15 | 0.00089 | ENSMUSG00000052031 | protein_coding | Tagap1 | T cell activation GTPase activating protein 1 | 380608 | | 17 | 6955011 | 6961156 | -1 | Mm.371734 |
| 5.84 | 5.84 | 5.22 | 6.21 | DN3ab_Itr | DN3cd | 0.43 | 2.26 | 0.051 | 0.22 | 0.00089 | ENSMUSG00000021891 | protein_coding | Mettl6 | methyltransferase like 6 | 67011 | | 14 | 31473578 | 31495040 | -1 | Mm.291731 |
| 3.39 | 4.81 | 4.25 | 4.16 | DN3ab_Itr | DN3cd | 0.68 | 2.31 | 0.047 | 0.21 | 0.00089 | ENSMUSG00000032344 | protein_coding | Cgas | cyclic GMP-AMP synthase | 214763 | | 9 | 78430526 | 78443237 | -1 | Mm.101559 |
| 6.04 | 5.97 | 5.78 | 6.41 | DN3ab_Itr | DN3cd | 0.26 | 1.91 | 0.089 | 0.31 | 0.00089 | ENSMUSG00000025898 | protein_coding | Cwf19l2 | CWF19-like 2, cell cycle control (S. pombe) | 244672 | | 9 | 3403592 | 3479236 | 1 | Mm.245482 |
| 4.5 | 4.75 | 3.37 | 4.73 | DN3ab_Itr | DN3cd | 0.65 | 3.17 | 0.012 | 0.088 | 0.00089 | ENSMUSG00000003531 | protein_coding | Dgcr6 | DiGeorge syndrome critical region gene 6 | 13353 | | 16 | 18052860 | 18071632 | 1 | Mm.27155 |
| 1.44 | 1.86 | -1.36 | 2.05 | DN3ab_Itr | DN3cd | 1.17 | 2.3 | 0.047 | 0.21 | 9.00E-04 | ENSMUSG00000063522 | protein_coding | 2010109I03Rik | RIKEN cDNA 2010109I03 gene | 67038 | | 15 | 74876987 | 74897021 | -1 | Mm.154566 |
| 4.4 | 4.48 | 2.81 | 4.2 | DN3ab_Itr | DN3cd | 0.49 | 2.05 | 0.071 | 0.27 | 9.00E-04 | ENSMUSG00000073434 | protein_coding | Wdr90 | WD repeat domain 90 | 106618 | | 17 | 25844771 | 25861501 | -1 | Mm.35470 |
| 6.09 | 6.16 | 4.99 | 6.52 | DN3ab_Itr | DN3cd | 0.66 | 2.4 | 0.04 | 0.19 | 9.00E-04 | ENSMUSG00000103038 | TEC | Gm37124 | predicted gene, 37124 | NA | | 1 | 137924388 | 137926379 | 1 | |
| 1.47 | 1.58 | 5.31 | 3.44 | DN3ab_Itr | DN3cd | -1.47 | -4.8 | 0.001 | 0.018 | 9.00E-04 | ENSMUSG00000025473 | protein_coding | Adam8 | a disintegrin and metallopeptidase domain 8 | 11501 | | 7 | 139978932 | 139992562 | -1 | Mm.15969 |
| 7.48 | 8.46 | 4.52 | 7.76 | DN3ab_Itr | DN3cd | 1.57 | 4.16 | 0.0025 | 0.033 | 0.00091 | ENSMUSG00000091705 | protein_coding | H2-Q2 | histocompatibility 2, Q region locus 2 | 15013 | | 17 | 35342242 | 35346762 | 1 | Mm.427969 |
| 7.29 | 7.32 | 6.9 | 7.57 | DN3ab_Itr | DN3cd | 0.32 | 2.59 | 0.03 | 0.16 | 0.00091 | ENSMUSG00000020721 | protein_coding | Helz | helicase with zinc finger domain | 78455 | | 11 | 107547930 | 107693826 | 1 | Mm.272047 Mm.394603 Mm.490425 |
| 4.4 | 4.85 | 3.41 | 5.48 | DN3ab_Itr | DN3cd | 1.08 | 3.52 | 0.0067 | 0.061 | 0.00091 | ENSMUSG00000102142 | sense_intronic | Gm26930 | predicted gene, 26930 | NA | | 1 | 137932041 | 137933284 | 1 | |
| 0.23 | 2.29 | -0.27 | 4.57 | DN3ab_Itr | DN3cd | 3.27 | 6.07 | 2.00E-04 | 0.0068 | 0.00092 | ENSMUSG00000008193 | protein_coding | Spib | Spi-B transcription factor (Spi-1/PU.1 related) | 272382 | | 7 | 44525993 | 44532071 | -1 | Mm.8012 |
| 4.08 | 1.3 | 5.15 | 4.05 | DN3ab_Itr | DN3cd | -1.49 | -5.38 | 0.00047 | 0.012 | 0.00093 | ENSMUSG00000064262 | protein_coding | Gimap8 | GTPase, IMAP family member 8 | 243374 | | 6 | 48647234 | 48660875 | 1 | Mm.86514 |
| 3.18 | 3.12 | 1.79 | 3.57 | DN3ab_Itr | DN3cd | 0.57 | 1.68 | 0.13 | 0.38 | 0.00094 | ENSMUSG00000066643 | protein_coding | Wdr35 | WD repeat domain 35 | 74682 | | 12 | 8973892 | 9028847 | 1 | Mm.87389 |
| 8.87 | 8.95 | 8.38 | 8.92 | DN3ab_Itr | DN3cd | 0.29 | 3.22 | 0.011 | 0.083 | 0.00094 | ENSMUSG00000045799 | processed_pseudogene | Gm9800 | predicted gene 9800 | NA | | 14 | 8800000 | 8800335 | -1 | |
| -0.56 | -0.49 | -3.72 | 0.64 | DN3ab_Itr | DN3cd | 2.09 | 2.5 | 0.034 | 0.17 | 0.00094 | ENSMUSG00000050066 | processed_transcript | A130023I24Rik | RIKEN cDNA A130023I24 gene | NA | | 7 | 134227474 | 134230705 | -1 | |
| 3.27 | 3.24 | 2.35 | 3.74 | DN3ab_Itr | DN3cd | 0.53 | 1.97 | 0.081 | 0.29 | 0.00096 | ENSMUSG00000047996 | protein_coding | Prrg1 | proline rich Gla (G-carboxyglutamic acid) 1 | 546336 | X | | 78449613 | 78583896 | -1 | Mm.211712 |
| 8.8 | 8.85 | 8.78 | 9.22 | DN3ab_Itr | DN3cd | 0.23 | 3.14 | 0.012 | 0.091 | 0.00098 | ENSMUSG00000004207 | protein_coding | Psap | prosaposin | 19156 | | 10 | 60277627 | 60302597 | 1 | Mm.29978 Mm.277498 |
| 5.15 | 5.38 | 3.67 | 5.99 | DN3ab_Itr | DN3cd | 0.99 | 2.54 | 0.032 | 0.17 | 0.00099 | ENSMUSG00000030246 | protein_coding | Ldhb | lactate dehydrogenase B | 16832 | | 6 | 142490249 | 142507957 | -1 | Mm.9745 |
| 4.43 | 4.41 | 3.88 | 4.85 | DN3ab_Itr | DN3cd | 0.41 | 2.15 | 0.06 | 0.24 | 0.001 | ENSMUSG00000032743 | protein_coding | D430042O09Rik | RIKEN cDNA D430042O09 gene | 233865 | | 7 | 125707888 | 125874793 | 1 | Mm.274389 |
| 6.75 | 6.92 | 6.49 | 7.34 | DN3ab_Itr | DN3cd | 0.48 | 3.65 | 0.0055 | 0.053 | 0.001 | ENSMUSG00000029249 | protein_coding | Rest | RE1-silencing transcription factor | 19712 | | 5 | 77265491 | 77286432 | 1 | Mm.28840 Mm.482359 |
| 7.3 | 7.3 | 7.63 | 6.95 | DN3ab_Itr | DN3cd | -0.32 | -2.41 | 0.04 | 0.19 | 0.001 | ENSMUSG00000079426 | protein_coding | Arpc4 | actin related protein 2/3 complex, subunit 4 | 68089 | | 6 | 113378115 | 113390448 | 1 | Mm.289306 Mm.472172 |
| 4.11 | 5.61 | 0.89 | 5.52 | DN3ab_Itr | DN3cd | 1.97 | 4.56 | 0.0014 | 0.023 | 0.001 | ENSMUSG00000025810 | protein_coding | Nrp1 | neuropilin 1 | 18186 | | 8 | 128358604 | 128503363 | 1 | Mm.271745 |
| 4.79 | 5.33 | 3.68 | 5.23 | DN3ab_Itr | DN3cd | 0.9 | 4.72 | 0.0011 | 0.02 | 0.001 | ENSMUSG00000019845 | protein_coding | Tube1 | epsilon-tubulin 1 | 71924 | | 10 | 39133976 | 39152542 | 1 | Mm.297956 |
| 2.69 | 2.65 | 1.94 | 4.02 | DN3ab_Itr | DN3cd | 0.91 | 2.21 | 0.055 | 0.23 | 0.001 | ENSMUSG00000028523 | protein_coding | Tctex1d1 | Tctex1 domain containing 1 | 67344 | | 4 | 102978606 | 103005594 | 1 | Mm.111346 |
| 5.39 | 6.1 | 5.14 | 6.51 | DN3ab_Itr | DN3cd | 1.01 | 7.72 | 3.20E-05 | 0.0024 | 0.001 | ENSMUSG00000027636 | protein_coding | Sla2 | Src-like-adaptor 2 | 77799 | | 2 | 156872457 | 156887192 | -1 | Mm.29264 Mm.387161 |
| 1.6 | 2.48 | 0.03 | 3.29 | DN3ab_Itr | DN3cd | 1.65 | 3.78 | 0.0045 | 0.047 | 0.001 | ENSMUSG00000022652 | protein_coding | Morc1 | microrchidia 1 | 17450 | | 16 | 48431237 | 48630900 | 1 | Mm.250060 |
| 6.59 | 6.75 | 5.76 | 6.67 | DN3ab_Itr | DN3cd | 0.47 | 3.31 | 0.0093 | 0.076 | 0.001 | ENSMUSG00000023940 | protein_coding | Sgo1 | shugoshin 1 | 72415 | | 17 | 53674786 | 53689333 | -1 | Mm.153202 |
| 6.44 | 6.68 | 5.34 | 6.43 | DN3ab_Itr | DN3cd | 0.57 | 3.57 | 0.0062 | 0.058 | 0.001 | ENSMUSG00000041498 | protein_coding | Kif14 | kinesin family member 14 | 381293 | | 1 | 136466343 | 136531511 | 1 | Mm.458468 |
| 4.52 | 3.12 | 4.42 | 4.37 | DN3ab_Itr | DN3cd | -0.65 | -2.47 | 0.036 | 0.18 | 0.001 | ENSMUSG00000043391 | protein_coding | 2510009E07Rik | RIKEN cDNA 2510009E07 gene | 72190 | | 16 | 21649045 | 21694665 | -1 | Mm.131606 |
| 8.62 | 8.35 | 9 | 8.18 | DN3ab_Itr | DN3cd | -0.53 | -4.99 | 0.00078 | 0.016 | 0.001 | ENSMUSG00000020547 | protein_coding | Bzw2 | basic leucine zipper and W2 domains 2 | 66912 | | 12 | 36091835 | 36158080 | -1 | Mm.288669 |
| 5.55 | 5.88 | 4.51 | 5.98 | DN3ab_Itr | DN3cd | 0.77 | 3.54 | 0.0065 | 0.06 | 0.0011 | ENSMUSG00000029438 | protein_coding | Bcl7a | B cell CLL/lymphoma 7A | 77045 | | 5 | 123343834 | 123374992 | 1 | Mm.151960 |
| 6.07 | 6.51 | 5.05 | 6.29 | DN3ab_Itr | DN3cd | 0.75 | 4.84 | 0.00095 | 0.018 | 0.0011 | ENSMUSG00000047534 | protein_coding | Mis18bp1 | MIS18 binding protein 1 | 217653 | | 12 | 65132734 | 65172604 | -1 | Mm.21144 |
| 8.55 | 8.71 | 7.75 | 8.43 | DN3ab_Itr | DN3cd | 0.37 | 3.66 | 0.0054 | 0.053 | 0.0011 | ENSMUSG00000009575 | protein_coding | Cbx5 | chromobox 5 | 12419 | | 15 | 103191544 | 103239816 | -1 | Mm.262059 |
| 6.34 | 6.32 | 6.27 | 6.89 | DN3ab_Itr | DN3cd | 0.29 | 2.31 | 0.047 | 0.21 | 0.0011 | ENSMUSG00000026743 | protein_coding | Mllt10 | myeloid/lymphoid or mixed-lineage leukemia; translocated to, 10 | 17354 | | 2 | 18055237 | 18212388 | 1 | Mm.209175 |
| 1.47 | 1.59 | 3.22 | 1.45 | DN3ab_Itr | DN3cd | -0.94 | -2.52 | 0.033 | 0.17 | 0.0011 | ENSMUSG00000072568 | protein_coding | Fam84b | family with sequence similarity 84, member B | 399603 | | 15 | 60818994 | 60853778 | -1 | Mm.411968 |
| 7.69 | 7.64 | 8.18 | 7.59 | DN3ab_Itr | DN3cd | -0.31 | -2.91 | 0.017 | 0.11 | 0.0011 | ENSMUSG00000033220 | protein_coding | Rac2 | Rac family small GTPase 2 | 19354 | | 15 | 78559167 | 78572783 | -1 | Mm.1972 |
| 5.44 | 5.09 | 7.8 | 6.5 | DN3ab_Itr | DN3cd | -0.94 | -5.29 | 0.00053 | 0.012 | 0.0011 | ENSMUSG00000030530 | protein_coding | Furin | furin (paired basic amino acid cleaving enzyme) | 18550 | | 7 | 80388585 | 80405436 | -1 | Mm.5241 |
| 5.53 | 6.17 | 3.87 | 6.25 | DN3ab_Itr | DN3cd | 1.21 | 3.83 | 0.0042 | 0.045 | 0.0012 | ENSMUSG00000000276 | protein_coding | Dgke | diacylglycerol kinase, epsilon | 56077 | | 11 | 89035179 | 89066850 | -1 | Mm.153695 |
| -0.04 | -0.68 | 3.24 | -1.06 | DN3ab_Itr | DN3cd | -2.65 | -3.7 | 0.0051 | 0.051 | 0.0012 | ENSMUSG00000038007 | protein_coding | Acer2 | alkaline ceramidase 2 | 230379 | | 4 | 86874396 | 86934822 | 1 | Mm.45019 Mm.458684 |
| 5.93 | 6.33 | 4.7 | 5.92 | DN3ab_Itr | DN3cd | 0.7 | 4.44 | 0.0017 | 0.026 | 0.0012 | ENSMUSG00000020808 | protein_coding | Pimreg | PICALM interacting mitotic regulator | 109212 | | 11 | 72042032 | 72047370 | 1 | Mm.33118 |
| 8.43 | 8.03 | 8.86 | 8.02 | DN3ab_Itr | DN3cd | -0.61 | -7.04 | 6.50E-05 | 0.0035 | 0.0012 | ENSMUSG00000026341 | protein_coding | Actr3 | ARP3 actin-related protein 3 | 74117 | | 1 | 125392905 | 125435727 | -1 | Mm.183102 Mm.471990 |
| 4.08 | 3.87 | 5.52 | 4.11 | DN3ab_Itr | DN3cd | -0.87 | -3.73 | 0.0048 | 0.049 | 0.0012 | ENSMUSG00000027878 | protein_coding | Notch2 | notch 2 | 18129 | | 3 | 98013527 | 98150361 | 1 | Mm.485843 |
| 4.14 | 4.08 | 3.77 | 5.16 | DN3ab_Itr | DN3cd | 0.63 | 2.22 | 0.054 | 0.23 | 0.0012 | ENSMUSG00000021211 | protein_coding | Akr1c12 | aldo-keto reductase family 1, member C12 | 622402 | | 13 | 4268176 | 4279433 | -1 | Mm.27447 Mm.380356 |
| 8.49 | 8.75 | 8.14 | 8.77 | DN3ab_Itr | DN3cd | 0.43 | 5.77 | 0.00028 | 0.0085 | 0.0012 | ENSMUSG00000032011 | protein_coding | Thy1 | thymus cell antigen 1, theta | 21838 | | 9 | 44043384 | 44048579 | 1 | Mm.3951 |
| 5.46 | 5.68 | 4.66 | 5.67 | DN3ab_Itr | DN3cd | 0.55 | 3.58 | 0.0061 | 0.057 | 0.0012 | ENSMUSG00000021871 | protein_coding | Gm49342 | predicted gene, 49342 | NA | | 14 | 50944517 | 50963136 | 1 | |
| 5.49 | 6.83 | 3.21 | 7.07 | DN3ab_Itr | DN3cd | 2.01 | 4.62 | 0.0013 | 0.022 | 0.0012 | ENSMUSG00000096847 | protein_coding | Tmem151b | transmembrane protein 151B | 210573 | | 17 | 45541952 | 45549677 | -1 | Mm.329064 Mm.491193 |
| -0.13 | 1.16 | -3.72 | 2.51 | DN3ab_Itr | DN3cd | 3.29 | 3.47 | 0.0072 | 0.064 | 0.0012 | ENSMUSG00000034009 | protein_coding | Rxfp1 | relaxin/insulin-like family peptide receptor 1 | 381489 | | 3 | 79641611 | 79737880 | -1 | Mm.331086 |
| 4.09 | 5.54 | 0.88 | 5 | DN3ab_Itr | DN3cd | 1.84 | 5 | 0.00077 | 0.016 | 0.0012 | ENSMUSG00000079507 | protein_coding | H2-Q1 | histocompatibility 2, Q region locus 1 | 15006 | | 17 | 35320405 | 35325099 | 1 | Mm.466882 |
| 7.63 | 8.01 | 6.78 | 7.88 | DN3ab_Itr | DN3cd | 0.68 | 4.98 | 0.00079 | 0.016 | 0.0012 | ENSMUSG00000064326 | protein_coding | Siva1 | SIVA1, apoptosis-inducing factor | 30954 | | 12 | 112644679 | 112649149 | 1 | Mm.289812 |
| 0.79 | 0.76 | -2.9 | 0.86 | DN3ab_Itr | DN3cd | 1.18 | 1.69 | 0.13 | 0.38 | 0.0012 | ENSMUSG00000041605 | protein_coding | Inava | innate immunity activator | 67313 | | 1 | 136213531 | 136234264 | -1 | Mm.36763 |
| 6.52 | 6.87 | 5.53 | 6.58 | DN3ab_Itr | DN3cd | 0.63 | 4.59 | 0.0014 | 0.022 | 0.0012 | ENSMUSG00000001517 | protein_coding | Foxm1 | forkhead box M1 | 14235 | | 6 | 128362967 | 128376146 | 1 | Mm.42148 |
| 8.36 | 8.54 | 7.49 | 8.91 | DN3ab_Itr | DN3cd | 0.72 | 2.99 | 0.016 | 0.11 | 0.0012 | ENSMUSG00000022468 | protein_coding | Endou | endonuclease, polyU-specific | 19011 | | 15 | 97711015 | 97731339 | -1 | Mm.1001 |
| 1.27 | 1.47 | -2.9 | 0.81 | DN3ab_Itr | DN3cd | 1.07 | 1.79 | 0.11 | 0.35 | 0.0013 | ENSMUSG00000001039 | protein_coding | B9d1 | B9 protein domain 1 | 27078 | | 11 | 61505144 | 61512931 | 1 | Mm.261226 |
| 8.06 | 6.68 | 8.33 | 7.67 | DN3ab_Itr | DN3cd | -1 | -7.1 | 6.10E-05 | 0.0034 | 0.0013 | ENSMUSG00000026581 | protein_coding | Sell | selectin, lymphocyte | 20343 | | 1 | 164061982 | 164084181 | 1 | Mm.1461 |
| 7.72 | 8.29 | 6 | 7.74 | DN3ab_Itr | DN3cd | 0.99 | 4.53 | 0.0015 | 0.024 | 0.0013 | ENSMUSG00000022472 | protein_coding | Desi1 | desumoylating isopeptidase 1 | 28075 | | 15 | 81992523 | 82033862 | -1 | Mm.474453 |
| 3.54 | 1.21 | 2.52 | 2.2 | DN3ab_Itr | DN3cd | -1.3 | -3.28 | 0.0097 | 0.079 | 0.0013 | ENSMUSG00000106859 | TEC | Gm42737 | predicted gene 42737 | NA | | 5 | 67757576 | 67759330 | -1 | |
| 4.29 | 4.28 | 3.27 | 4.53 | DN3ab_Itr | DN3cd | 0.48 | 1.95 | 0.083 | 0.3 | 0.0013 | ENSMUSG00000019971 | protein_coding | Cep290 | centrosomal protein 290 | 216274 | | 10 | 100487558 | 100574840 | 1 | Mm.229114 |
| 4.91 | 5.65 | 5.13 | 5.07 | DN3ab_Itr | DN3cd | 0.38 | 2.43 | 0.038 | 0.18 | 0.0013 | ENSMUSG00000031960 | protein_coding | Aars | alanyl-tRNA synthetase | 234734 | | 8 | 111033144 | 111057664 | 1 | Mm.24174 Mm.491269 |
| 7.49 | 7.54 | 6.53 | 7.33 | DN3ab_Itr | DN3cd | 0.35 | 2.47 | 0.036 | 0.18 | 0.0013 | ENSMUSG00000026565 | protein_coding | Pou2f1 | POU domain, class 2, transcription factor 1 | 18986 | | 1 | 165865154 | 166002678 | -1 | Mm.245261 |
| 5.13 | 5.92 | 3.94 | 6.31 | DN3ab_Itr | DN3cd | 1.38 | 4.64 | 0.0013 | 0.022 | 0.0013 | ENSMUSG00000053702 | protein_coding | Nebl | nebulette | 74103 | | 2 | 17343909 | 17731464 | -1 | Mm.120298 Mm.256298 Mm.404039 Mm.440498 |
| 0.56 | 0.94 | -0.08 | 2.76 | DN3ab_Itr | DN3cd | 1.52 | 3.1 | 0.013 | 0.094 | 0.0013 | ENSMUSG00000026601 | protein_coding | Axdnd1 | axonemal dynein light chain domain containing 1 | 77352 | | 1 | 156323509 | 156421159 | -1 | Mm.428709 |
| 3.45 | 2.86 | 6.8 | 4.73 | DN3ab_Itr | DN3cd | -1.69 | -6.66 | 9.90E-05 | 0.0044 | 0.0013 | ENSMUSG00000022885 | protein_coding | St6gal1 | beta galactoside alpha 2,6 sialyltransferase 1 | 20440 | | 16 | 23224740 | 23360350 | 1 | Mm.149029 Mm.489708 |
| 5.58 | 3.88 | 5.17 | 4.73 | DN3ab_Itr | DN3cd | -1.05 | -4.22 | 0.0023 | 0.031 | 0.0013 | ENSMUSG00000021559 | protein_coding | Dapk1 | death associated protein kinase 1 | 69635 | | 13 | 60601947 | 60763191 | 1 | Mm.24103 |
| 7.01 | 7.08 | 6.74 | 7.29 | DN3ab_Itr | DN3cd | 0.29 | 2.97 | 0.016 | 0.11 | 0.0013 | ENSMUSG00000018707 | protein_coding | Dync1h1 | dynein cytoplasmic 1 heavy chain 1 | 13424 | | 12 | 110601452 | 110666945 | 1 | Mm.181430 |
| 7.12 | 7.07 | 8.37 | 7.58 | DN3ab_Itr | DN3cd | -0.45 | -3.14 | 0.012 | 0.091 | 0.0014 | ENSMUSG00000004319 | protein_coding | Clcn3 | chloride channel, voltage-sensitive 3 | 12725 | | 8 | 60910389 | 60983300 | -1 | Mm.25263 Mm.259751 |
| 3.05 | 4.57 | 0.43 | 5.12 | DN3ab_Itr | DN3cd | 2.11 | 4.36 | 0.0019 | 0.027 | 0.0014 | ENSMUSG00000023274 | protein_coding | Cd4 | CD4 antigen | 12504 | | 6 | 124864692 | 124888221 | -1 | Mm.2209 |
| 8.33 | 8.35 | 7.92 | 8.48 | DN3ab_Itr | DN3cd | 0.27 | 2.47 | 0.036 | 0.18 | 0.0014 | ENSMUSG00000037608 | protein_coding | Bclaf1 | BCL2-associated transcription factor 1 | 72567 | | 10 | 20312469 | 20344613 | 1 | Mm.294783 |
| 6.19 | 5.9 | 7.42 | 6.19 | DN3ab_Itr | DN3cd | -0.79 | -4.27 | 0.0022 | 0.03 | 0.0014 | ENSMUSG00000032413 | protein_coding | Rasa2 | RAS p21 protein activator 2 | 114713 | | 9 | 96539300 | 96631617 | -1 | Mm.124502 |
| 4.56 | 3.81 | 7.49 | 5.27 | DN3ab_Itr | DN3cd | -1.74 | -6.44 | 0.00013 | 0.0053 | 0.0014 | ENSMUSG00000060591 | protein_coding | Ifitm2 | interferon induced transmembrane protein 2 | 80876 | | 7 | 140954837 | 140955987 | -1 | Mm.379266 |
| 5.69 | 5.98 | 5.38 | 6.4 | DN3ab_Itr | DN3cd | 0.62 | 4.29 | 0.0021 | 0.029 | 0.0014 | ENSMUSG00000016940 | protein_coding | Kctd2 | potassium channel tetramerisation domain containing 2 | 70382 | | 11 | 115420128 | 115431274 | 1 | Mm.276299 |
| 7.02 | 4.65 | 6.06 | 4.91 | DN3ab_Itr | DN3cd | -1.81 | -7.58 | 3.70E-05 | 0.0026 | 0.0014 | ENSMUSG00000029455 | protein_coding | Aldh2 | aldehyde dehydrogenase 2, mitochondrial | 11669 | | 5 | 121566027 | 121593824 | -1 | Mm.284446 |
| 5.55 | 5.81 | 4.91 | 5.91 | DN3ab_Itr | DN3cd | 0.57 | 3.92 | 0.0036 | 0.041 | 0.0014 | ENSMUSG00000079470 | protein_coding | Utp14b | UTP14B small subunit processome component | 195434 | | 1 | 78658038 | 78671512 | 1 | Mm.276016 |
| 4.81 | 4.93 | 3.71 | 4.84 | DN3ab_Itr | DN3cd | 0.5 | 2.58 | 0.03 | 0.16 | 0.0014 | ENSMUSG00000030177 | protein_coding | Ccdc77 | coiled-coil domain containing 77 | 67200 | | 6 | 120324323 | 120364374 | -1 | Mm.45767 |
| 5.45 | 2.85 | 4.4 | 3.47 | DN3ab_Itr | DN3cd | -1.78 | -5.41 | 0.00045 | 0.011 | 0.0014 | ENSMUSG00000033083 | protein_coding | Tbc1d4 | TBC1 domain family, member 4 | 210789 | | 14 | 101442360 | 101609191 | -1 | Mm.320639 |
| 7.56 | 7.01 | 7.29 | 7.32 | DN3ab_Itr | DN3cd | -0.27 | -2.39 | 0.041 | 0.19 | 0.0014 | ENSMUSG00000050379 | protein_coding | 6-Sep | septin 6 | 56526 | X | | 36911326 | 36991794 | -1 | Mm.260036 Mm.470184 |
| 6.18 | 6.14 | 7.14 | 6.31 | DN3ab_Itr | DN3cd | -0.45 | -2.84 | 0.02 | 0.12 | 0.0014 | ENSMUSG00000028149 | protein_coding | Rap1gds1 | RAP1, GTP-GDP dissociation stimulator 1 | 229877 | | 3 | 138925902 | 139075201 | -1 | Mm.28893 Mm.400828 |
| 10.81 | 10.99 | 10.82 | 11.56 | DN3ab_Itr | DN3cd | 0.45 | 4.05 | 0.003 | 0.036 | 0.0015 | ENSMUSG00000064337 | Mt_rRNA | mt-Rnr1 | mitochondrially encoded 12S rRNA | NA | MT | | 70 | 1024 | 1 | |
| 3.35 | 5.19 | 1.33 | 5.72 | DN3ab_Itr | DN3cd | 2.45 | 5.75 | 0.00029 | 0.0086 | 0.0015 | ENSMUSG00000026576 | protein_coding | Atp1b1 | ATPase, Na+/K+ transporting, beta 1 polypeptide | 11931 | | 1 | 164437109 | 164458355 | -1 | Mm.4550 |
| 9.83 | 8.96 | 9.63 | 9.38 | DN3ab_Itr | DN3cd | -0.57 | -4.64 | 0.0013 | 0.021 | 0.0015 | ENSMUSG00000002107 | protein_coding | Celf2 | CUGBP, Elav-like family member 2 | 14007 | | 2 | 6539694 | 7509563 | -1 | Mm.398543 |
| 3.32 | 3.18 | 5.67 | 3.93 | DN3ab_Itr | DN3cd | -1.16 | -3.84 | 0.0041 | 0.044 | 0.0015 | ENSMUSG00000015846 | protein_coding | Rxra | retinoid X receptor alpha | 20181 | | 2 | 27676440 | 27762957 | 1 | Mm.24624 |
| 4.56 | 3.59 | 3.94 | 3.93 | DN3ab_Itr | DN3cd | -0.55 | -2.84 | 0.02 | 0.12 | 0.0015 | ENSMUSG00000041180 | protein_coding | Hectd2 | HECT domain E3 ubiquitin protein ligase 2 | 226098 | | 19 | 36554639 | 36621135 | 1 | Mm.61107 |
| 9.59 | 9.57 | 9.85 | 9.47 | DN3ab_Itr | DN3cd | -0.19 | -2.61 | 0.028 | 0.15 | 0.0015 | ENSMUSG00000003429 | protein_coding | Rps11 | ribosomal protein S11 | 27207 | | 7 | 45122379 | 45124439 | -1 | Mm.196538 Mm.389435 |
| 6.63 | 6.91 | 6.14 | 7.05 | DN3ab_Itr | DN3cd | 0.56 | 4.48 | 0.0016 | 0.025 | 0.0015 | ENSMUSG00000005417 | protein_coding | Mprip | myosin phosphatase Rho interacting protein | 26936 | | 11 | 59661305 | 59780860 | 1 | Mm.2402 |
| 7.17 | 7.65 | 5.77 | 7.62 | DN3ab_Itr | DN3cd | 1.01 | 3.87 | 0.0039 | 0.043 | 0.0015 | ENSMUSG00000030263 | protein_coding | Lrmp | lymphoid-restricted membrane protein | 16970 | | 6 | 145115653 | 145174934 | 1 | Mm.843 |
| 7.47 | 7.1 | 8.73 | 7.4 | DN3ab_Itr | DN3cd | -0.88 | -4.62 | 0.0013 | 0.022 | 0.0015 | ENSMUSG00000031827 | protein_coding | Cotl1 | coactosin-like 1 (Dictyostelium) | 72042 | | 8 | 119809222 | 119840544 | -1 | Mm.393405 |
| 9.4 | 9.63 | 8.04 | 8.99 | DN3ab_Itr | DN3cd | 0.49 | 3.6 | 0.0059 | 0.056 | 0.0015 | ENSMUSG00000003038 | protein_coding | Hmgn2 | high mobility group nucleosomal binding domain 2 | 15331 | | 4 | 133964738 | 133968650 | -1 | Mm.319660 |
| 6.64 | 6.69 | 5.92 | 6.5 | DN3ab_Itr | DN3cd | 0.27 | 2.57 | 0.031 | 0.16 | 0.0015 | ENSMUSG00000031575 | protein_coding | Ash2l | ASH2 like histone lysine methyltransferase complex subunit | 23808 | | 8 | 25815996 | 25847694 | -1 | Mm.27706 |
| 3.83 | 3.34 | 5.14 | 3.74 | DN3ab_Itr | DN3cd | -1 | -5.43 | 0.00044 | 0.011 | 0.0015 | ENSMUSG00000032750 | protein_coding | Gab3 | growth factor receptor bound protein 2-associated protein 3 | 210710 | X | | 74966843 | 75085458 | -1 | Mm.267056 |
| 4.54 | 4.3 | 6.1 | 4.96 | DN3ab_Itr | DN3cd | -0.76 | -4.28 | 0.0021 | 0.029 | 0.0016 | ENSMUSG00000030103 | protein_coding | Bhlhe40 | basic helix-loop-helix family, member e40 | 20893 | | 6 | 108660629 | 108666925 | 1 | Mm.2436 |
| 5.81 | 5.01 | 5.52 | 5.49 | DN3ab_Itr | DN3cd | -0.43 | -2.86 | 0.019 | 0.12 | 0.0016 | ENSMUSG00000032870 | protein_coding | Smap2 | small ArfGAP 2 | 69780 | | 4 | 120968317 | 121017247 | -1 | Mm.271819 |
| 6.74 | 7.15 | 5.56 | 6.83 | DN3ab_Itr | DN3cd | 0.74 | 4.46 | 0.0016 | 0.025 | 0.0016 | ENSMUSG00000012443 | protein_coding | Kif11 | kinesin family member 11 | 16551 | | 19 | 37376403 | 37421859 | 1 | Mm.42203 |
| 7.81 | 7.43 | 8.8 | 7.89 | DN3ab_Itr | DN3cd | -0.66 | -6.11 | 0.00019 | 0.0066 | 0.0016 | ENSMUSG00000026970 | protein_coding | Rbms1 | RNA binding motif, single stranded interacting protein 1 | 56878 | | 2 | 60750193 | 60963192 | -1 | Mm.259667 Mm.482338 |
| 5.22 | 5.23 | 4.01 | 5.12 | DN3ab_Itr | DN3cd | 0.41 | 1.95 | 0.083 | 0.3 | 0.0016 | ENSMUSG00000027379 | protein_coding | Bub1 | BUB1, mitotic checkpoint serine/threonine kinase | 12235 | | 2 | 127801122 | 127831865 | -1 | Mm.2185 |
| 4.7 | 5.59 | 4.74 | 4.84 | DN3ab_Itr | DN3cd | 0.55 | 3.56 | 0.0063 | 0.058 | 0.0016 | ENSMUSG00000063954 | protein_coding | Hist2h2aa2 | histone cluster 2, H2aa2 | 319192 | | 3 | 96239749 | 96240276 | -1 | Mm.5220 Mm.478832 Mm.485412 |
| 5.26 | 5.52 | 4.46 | 5.8 | DN3ab_Itr | DN3cd | 0.71 | 3.36 | 0.0086 | 0.072 | 0.0016 | ENSMUSG00000062646 | protein_coding | Ganc | glucosidase, alpha; neutral C | 76051 | | 2 | 120403896 | 120461700 | 1 | Mm.38851 |
| 6.52 | 6.24 | 7.27 | 6.44 | DN3ab_Itr | DN3cd | -0.56 | -5.11 | 0.00067 | 0.014 | 0.0016 | ENSMUSG00000038155 | protein_coding | Gstp2 | glutathione S-transferase, pi 2 | 14869 | | 19 | 4040285 | 4042236 | -1 | Mm.441542 |
| 4.24 | 4.31 | 3.19 | 4.28 | DN3ab_Itr | DN3cd | 0.45 | 2.27 | 0.05 | 0.22 | 0.0016 | ENSMUSG00000027635 | protein_coding | Dsn1 | DSN1 homolog, MIS12 kinetochore complex component | 66934 | | 2 | 156995265 | 157007154 | -1 | Mm.25410 |
| 7.94 | 8.69 | 8.15 | 9.61 | DN3ab_Itr | DN3cd | 1.11 | 7.82 | 2.90E-05 | 0.0022 | 0.0017 | ENSMUSG00000097971 | lincRNA | Gm26917 | predicted gene, 26917 | NA | | 17 | 39843013 | 39846341 | 1 | |
| 7 | 7 | 6.6 | 7.24 | DN3ab_Itr | DN3cd | 0.29 | 2.29 | 0.048 | 0.21 | 0.0017 | ENSMUSG00000032621 | protein_coding | Srek1 | splicing regulatory glutamine/lysine-rich protein 1 | 218543 | | 13 | 103739348 | 103774608 | -1 | Mm.458176 Mm.459433 |
| 5.24 | 5.66 | 4.79 | 6.14 | DN3ab_Itr | DN3cd | 0.83 | 4.57 | 0.0014 | 0.023 | 0.0017 | ENSMUSG00000028460 | protein_coding | Sit1 | suppression inducing transmembrane adaptor 1 | 54390 | | 4 | 43482081 | 43483734 | -1 | Mm.31775 |
| 5.37 | 5.62 | 4.45 | 5.6 | DN3ab_Itr | DN3cd | 0.61 | 3.45 | 0.0074 | 0.066 | 0.0017 | ENSMUSG00000022034 | protein_coding | Esco2 | establishment of sister chromatid cohesion N-acetyltransferase 2 | 71988 | | 14 | 65819038 | 65833994 | -1 | Mm.249280 |
| 3.99 | 3.66 | 5.24 | 3.83 | DN3ab_Itr | DN3cd | -0.91 | -4.27 | 0.0022 | 0.03 | 0.0017 | ENSMUSG00000028599 | protein_coding | Tnfrsf1b | tumor necrosis factor receptor superfamily, member 1b | 21938 | | 4 | 145213463 | 145246870 | -1 | Mm.235328 |
| 1.31 | 1.57 | 0.85 | 3.62 | DN3ab_Itr | DN3cd | 1.48 | 2.97 | 0.016 | 0.11 | 0.0017 | ENSMUSG00000065259 | snoRNA | Snora30 | small nucleolar RNA, H/ACA box 30 | NA | | 7 | 127527874 | 127528003 | 1 | |
| 8.64 | 8.91 | 8.26 | 9.36 | DN3ab_Itr | DN3cd | 0.65 | 3.92 | 0.0036 | 0.041 | 0.0017 | ENSMUSG00000032035 | protein_coding | Ets1 | E26 avian leukemia oncogene 1, 5' domain | 23871 | | 9 | 32636221 | 32757820 | 1 | Mm.292415 |
| 5.89 | 5.76 | 6.39 | 5.68 | DN3ab_Itr | DN3cd | -0.41 | -3.45 | 0.0074 | 0.066 | 0.0017 | ENSMUSG00000006589 | protein_coding | Aprt | adenine phosphoribosyl transferase | 11821 | | 8 | 122574635 | 122576909 | -1 | Mm.1786 |
| -1.02 | -0.48 | -2.95 | 1.31 | DN3ab_Itr | DN3cd | 2.47 | 3.3 | 0.0094 | 0.077 | 0.0017 | ENSMUSG00000052730 | processed_transcript | Gm5111 | predicted gene 5111 | 330305 | | 6 | 48589445 | 48590584 | 1 | Mm.123868 |
| 5.09 | 6.41 | 6.23 | 6.74 | DN3ab_Itr | DN3cd | 0.88 | 5.42 | 0.00044 | 0.011 | 0.0017 | ENSMUSG00000027863 | protein_coding | Cd2 | CD2 antigen | 12481 | | 3 | 101275899 | 101287939 | -1 | Mm.22842 |
| 7.23 | 6.7 | 8.33 | 7.09 | DN3ab_Itr | DN3cd | -0.91 | -6.41 | 0.00013 | 0.0054 | 0.0017 | ENSMUSG00000031799 | protein_coding | Tpm4 | tropomyosin 4 | 326618 | | 8 | 72134975 | 72153142 | 1 | Mm.295124 |
| 2.67 | 3.46 | 0.1 | 3.8 | DN3ab_Itr | DN3cd | 1.42 | 2.97 | 0.016 | 0.11 | 0.0018 | ENSMUSG00000021483 | protein_coding | Cdk20 | cyclin-dependent kinase 20 | 105278 | | 13 | 64432314 | 64441773 | 1 | Mm.74982 |
| 3.42 | 3.53 | 2.65 | 4.08 | DN3ab_Itr | DN3cd | 0.65 | 2.49 | 0.035 | 0.17 | 0.0018 | ENSMUSG00000052539 | protein_coding | Magi3 | membrane associated guanylate kinase, WW and PDZ domain containing 3 | 99470 | | 3 | 104013259 | 104220374 | -1 | Mm.264849 |
| 6.45 | 5.4 | 5.89 | 5.62 | DN3ab_Itr | DN3cd | -0.7 | -4.47 | 0.0016 | 0.025 | 0.0018 | ENSMUSG00000005696 | protein_coding | Sh2d1a | SH2 domain containing 1A | 20400 | X | | 42502453 | 42529330 | 1 | Mm.441197 |
| 6.95 | 5.91 | 8.3 | 6.64 | DN3ab_Itr | DN3cd | -1.38 | -11.26 | 1.50E-06 | 0.00042 | 0.0018 | ENSMUSG00000018340 | protein_coding | Anxa6 | annexin A6 | 11749 | | 11 | 54979108 | 55033445 | -1 | Mm.265347 |
| 7.15 | 6.36 | 7.16 | 7.14 | DN3ab_Itr | DN3cd | -0.41 | -2.65 | 0.027 | 0.15 | 0.0018 | ENSMUSG00000034160 | protein_coding | Ogt | O-linked N-acetylglucosamine (GlcNAc) transferase (UDP-N-acetylglucosamine:polypeptide-N-acetylglucosaminyl transferase) | 108155 | X | | 101640060 | 101684351 | 1 | Mm.487332 Mm.491168 |
| -1.63 | -1.6 | -3.72 | 0.54 | DN3ab_Itr | DN3cd | 2.4 | 2.84 | 0.02 | 0.12 | 0.0018 | ENSMUSG00000032128 | protein_coding | Robo3 | roundabout guidance receptor 3 | 19649 | | 9 | 37415669 | 37433246 | -1 | Mm.212826 |
| 2.21 | 0.72 | 5.3 | 1.86 | DN3ab_Itr | DN3cd | -2.83 | -7.77 | 3.00E-05 | 0.0023 | 0.0018 | ENSMUSG00000074813 | lincRNA | Gm14005 | predicted gene 14005 | NA | | 2 | 128178319 | 128502765 | -1 | |
| 5.09 | 4.91 | 5.98 | 4.92 | DN3ab_Itr | DN3cd | -0.62 | -3.51 | 0.0068 | 0.062 | 0.0019 | ENSMUSG00000026393 | protein_coding | Nek7 | NIMA (never in mitosis gene a)-related expressed kinase 7 | 59125 | | 1 | 138482875 | 138620141 | -1 | Mm.143817 Mm.486373 |
| 9.77 | 9.7 | 10.36 | 9.82 | DN3ab_Itr | DN3cd | -0.3 | -3.23 | 0.011 | 0.083 | 0.0019 | ENSMUSG00000026864 | protein_coding | Hspa5 | heat shock protein 5 | 14828 | | 2 | 34771970 | 34777547 | 1 | Mm.330160 Mm.470180 Mm.474909 |
| 5.95 | 5.32 | 8.59 | 6.81 | DN3ab_Itr | DN3cd | -1.37 | -6.22 | 0.00016 | 0.0061 | 0.0019 | ENSMUSG00000032578 | protein_coding | Cish | cytokine inducible SH2-containing protein | 12700 | | 9 | 107296026 | 107302784 | 1 | Mm.4592 |
| 6.5 | 6.45 | 7.1 | 6.36 | DN3ab_Itr | DN3cd | -0.39 | -2.78 | 0.022 | 0.13 | 0.0019 | ENSMUSG00000032112 | protein_coding | Trappc4 | trafficking protein particle complex 4 | 60409 | | 9 | 44403698 | 44407600 | -1 | Mm.29814 |
| 7.66 | 7.33 | 8.42 | 7.37 | DN3ab_Itr | DN3cd | -0.69 | -4.83 | 0.00098 | 0.018 | 0.0019 | ENSMUSG00000030830 | protein_coding | Itgal | integrin alpha L | 16408 | | 7 | 127296260 | 127335138 | 1 | Mm.1618 |
| 4.27 | 4.24 | 5.3 | 4.25 | DN3ab_Itr | DN3cd | -0.56 | -2.76 | 0.022 | 0.13 | 0.0019 | ENSMUSG00000044768 | protein_coding | D1Ertd622e | DNA segment, Chr 1, ERATO Doi 622, expressed | 52392 | | 1 | 97606318 | 97662074 | -1 | Mm.12309 |
| 4.33 | 2.26 | 6.22 | 2.99 | DN3ab_Itr | DN3cd | -2.76 | -11.95 | 9.10E-07 | 0.00036 | 0.0019 | ENSMUSG00000034573 | protein_coding | Ptpn13 | protein tyrosine phosphatase, non-receptor type 13 | 19249 | | 5 | 103425192 | 103598303 | 1 | Mm.3414 |
| 1.8 | 2.83 | -0.28 | 3.07 | DN3ab_Itr | DN3cd | 1.63 | 3.98 | 0.0033 | 0.038 | 0.0019 | ENSMUSG00000029848 | protein_coding | Stra8 | stimulated by retinoic acid gene 8 | 20899 | | 6 | 34920163 | 34939344 | 1 | Mm.5171 |
| 5.51 | 5.47 | 6.58 | 5.65 | DN3ab_Itr | DN3cd | -0.5 | -2.83 | 0.02 | 0.13 | 0.0019 | ENSMUSG00000037553 | protein_coding | Zdhhc18 | zinc finger, DHHC domain containing 18 | 503610 | | 4 | 133605299 | 133650154 | -1 | Mm.331948 |
| 8.86 | 8.5 | 10.51 | 9.15 | DN3ab_Itr | DN3cd | -0.93 | -4.7 | 0.0012 | 0.02 | 0.0019 | ENSMUSG00000068220 | protein_coding | Lgals1 | lectin, galactose binding, soluble 1 | 16852 | | 15 | 78926725 | 78930465 | 1 | Mm.43831 |
| 5.93 | 5.51 | 7.02 | 5.76 | DN3ab_Itr | DN3cd | -0.86 | -5.06 | 0.00072 | 0.015 | 0.002 | ENSMUSG00000038437 | protein_coding | Mllt6 | myeloid/lymphoid or mixed-lineage leukemia; translocated to, 6 | 246198 | | 11 | 97663414 | 97685463 | 1 | Mm.23685 |
| 4.41 | 3.46 | 6.72 | 4.17 | DN3ab_Itr | DN3cd | -1.92 | -6.19 | 0.00017 | 0.0062 | 0.002 | ENSMUSG00000028525 | protein_coding | Pde4b | phosphodiesterase 4B, cAMP specific | 18578 | | 4 | 102087543 | 102607259 | 1 | Mm.20181 |
| 2.41 | 4.75 | -0.64 | 4.57 | DN3ab_Itr | DN3cd | 2.79 | 6.66 | 9.90E-05 | 0.0044 | 0.002 | ENSMUSG00000016150 | protein_coding | Tenm1 | teneurin transmembrane protein 1 | 23963 | X | | 42527866 | 43429126 | -1 | Mm.327698 |
| 6.42 | 6.35 | 7.67 | 6.78 | DN3ab_Itr | DN3cd | -0.51 | -3.14 | 0.012 | 0.091 | 0.002 | ENSMUSG00000006519 | protein_coding | Cyba | cytochrome b-245, alpha polypeptide | 13057 | | 8 | 122424776 | 122432930 | -1 | Mm.271671 |
| 9.11 | 9.14 | 8.49 | 9.33 | DN3ab_Itr | DN3cd | 0.39 | 2.41 | 0.04 | 0.19 | 0.002 | ENSMUSG00000019982 | protein_coding | Myb | myeloblastosis oncogene | 17863 | | 10 | 21124935 | 21160984 | -1 | Mm.52109 Mm.473872 |
| 7.55 | 8.27 | 6.88 | 8.43 | DN3ab_Itr | DN3cd | 1.09 | 6.63 | 1.00E-04 | 0.0045 | 0.002 | ENSMUSG00000026923 | protein_coding | Notch1 | notch 1 | 18128 | | 2 | 26457903 | 26516663 | -1 | Mm.290610 |
| 4.14 | 4.6 | 2.12 | 3.86 | DN3ab_Itr | DN3cd | 0.78 | 3.43 | 0.0076 | 0.067 | 0.002 | ENSMUSG00000030086 | protein_coding | Chchd6 | coiled-coil-helix-coiled-coil-helix domain containing 6 | 66098 | | 6 | 89383146 | 89595652 | -1 | Mm.20313 |
| 4.59 | 4.63 | 2.94 | 4.8 | DN3ab_Itr | DN3cd | 0.62 | 1.84 | 0.1 | 0.33 | 0.002 | ENSMUSG00000056895 | protein_coding | Hist3h2ba | histone cluster 3, H2ba | 78303 | | 11 | 58948920 | 58949533 | 1 | Mm.28022 |
| 6.99 | 6.92 | 7.23 | 6.69 | DN3ab_Itr | DN3cd | -0.29 | -2.96 | 0.016 | 0.11 | 0.0021 | ENSMUSG00000028792 | protein_coding | Ak2 | adenylate kinase 2 | 11637 | | 4 | 128991958 | 129011529 | 1 | Mm.29460 |
| 5.53 | 5.83 | 5.41 | 6.31 | DN3ab_Itr | DN3cd | 0.59 | 4.79 | 0.001 | 0.019 | 0.0021 | ENSMUSG00000094870 | protein_coding | Zfp131 | zinc finger protein 131 | 72465 | | 13 | 119765187 | 119790889 | -1 | Mm.139115 Mm.446549 |
| 4.81 | 4.5 | 6.26 | 5.16 | DN3ab_Itr | DN3cd | -0.76 | -4.77 | 0.0011 | 0.019 | 0.0021 | ENSMUSG00000031146 | protein_coding | Plp2 | proteolipid protein 2 | 18824 | X | | 7667941 | 7671390 | -1 | Mm.18565 Mm.432064 Mm.463231 |
| -0.65 | 0.13 | -2.13 | 2.26 | DN3ab_Itr | DN3cd | 2.53 | 3.47 | 0.0072 | 0.065 | 0.0022 | ENSMUSG00000035211 | protein_coding | Xrra1 | X-ray radiation resistance associated 1 | 446101 | | 7 | 99859116 | 99917822 | 1 | Mm.440757 |
| 7.78 | 7.97 | 6.98 | 7.97 | DN3ab_Itr | DN3cd | 0.53 | 3.26 | 0.01 | 0.081 | 0.0022 | ENSMUSG00000026915 | protein_coding | Strbp | spermatid perinuclear RNA binding protein | 20744 | | 2 | 37483228 | 37703859 | -1 | Mm.237095 Mm.477856 |
| 4.35 | 4.44 | 3.28 | 5.24 | DN3ab_Itr | DN3cd | 0.84 | 2.3 | 0.047 | 0.21 | 0.0022 | ENSMUSG00000037972 | protein_coding | Snn | stannin | 20621 | | 16 | 11060945 | 11074985 | 1 | Mm.325800 |
| 6.52 | 4.4 | 6.62 | 5.57 | DN3ab_Itr | DN3cd | -1.53 | -7.13 | 5.90E-05 | 0.0034 | 0.0022 | ENSMUSG00000032014 | protein_coding | Oaf | out at first homolog | 102644 | | 9 | 43221235 | 43239911 | -1 | Mm.246479 |
| 5.31 | 5.51 | 2.91 | 6.06 | DN3ab_Itr | DN3cd | 1.02 | 1.94 | 0.085 | 0.3 | 0.0022 | ENSMUSG00000108207 | processed_transcript | 1810059H22Rik | RIKEN cDNA 1810059H22 gene | NA | | 5 | 148953748 | 148970122 | -1 | |
| 6.01 | 5.58 | 11.14 | 8.67 | DN3ab_Itr | DN3cd | -2.02 | -5.99 | 0.00022 | 0.0073 | 0.0022 | ENSMUSG00000001020 | protein_coding | S100a4 | S100 calcium binding protein A4 | 20198 | | 3 | 90603771 | 90606045 | 1 | Mm.3925 |
| 7.56 | 7.43 | 8.11 | 7.41 | DN3ab_Itr | DN3cd | -0.41 | -3.6 | 0.0059 | 0.056 | 0.0022 | ENSMUSG00000025135 | protein_coding | Anapc11 | anaphase promoting complex subunit 11 | 66156 | | 11 | 120598421 | 120608198 | 1 | Mm.21645 |
| 6.07 | 6.41 | 5.19 | 6.25 | DN3ab_Itr | DN3cd | 0.63 | 4.46 | 0.0016 | 0.025 | 0.0022 | ENSMUSG00000020330 | protein_coding | Hmmr | hyaluronan mediated motility receptor (RHAMM) | 15366 | | 11 | 40701395 | 40733422 | -1 | Mm.116997 |
| 6.53 | 6.21 | 7.32 | 6.5 | DN3ab_Itr | DN3cd | -0.58 | -5.64 | 0.00034 | 0.0093 | 0.0023 | ENSMUSG00000060803 | protein_coding | Gstp1 | glutathione S-transferase, pi 1 | 14870 | | 19 | 4035407 | 4037921 | -1 | Mm.299292 |
| 2.55 | 2.78 | -0.61 | 2.4 | DN3ab_Itr | DN3cd | 0.74 | 1.69 | 0.13 | 0.38 | 0.0023 | ENSMUSG00000027378 | protein_coding | Nphp1 | nephronophthisis 1 (juvenile) homolog (human) | 53885 | | 2 | 127740732 | 127788897 | -1 | Mm.210766 |
| 6.27 | 6.13 | 7.11 | 5.97 | DN3ab_Itr | DN3cd | -0.62 | -3.13 | 0.012 | 0.092 | 0.0023 | ENSMUSG00000025823 | protein_coding | Pdia4 | protein disulfide isomerase associated 4 | 12304 | | 6 | 47796141 | 47813430 | -1 | Mm.2442 |
| 4.14 | 4.06 | 6.64 | 5.27 | DN3ab_Itr | DN3cd | -0.92 | -3.77 | 0.0046 | 0.048 | 0.0023 | ENSMUSG00000056737 | protein_coding | Capg | capping protein (actin filament), gelsolin-like | 12332 | | 6 | 72544391 | 72562983 | 1 | Mm.18626 |
| 4.44 | 2.16 | 3.76 | 2.82 | DN3ab_Itr | DN3cd | -1.6 | -5.88 | 0.00025 | 0.0079 | 0.0023 | ENSMUSG00000095105 | protein_coding | Edaradd | EDAR (ectodysplasin-A receptor)-associated death domain | 171211 | | 13 | 12472632 | 12520438 | -1 | Mm.159671 |
| 5.12 | 5.18 | 4.5 | 5.22 | DN3ab_Itr | DN3cd | 0.34 | 2.53 | 0.033 | 0.17 | 0.0023 | ENSMUSG00000022020 | protein_coding | Naa16 | N(alpha)-acetyltransferase 16, NatA auxiliary subunit | 66897 | | 14 | 79325269 | 79390778 | -1 | Mm.24425 |
| 5.9 | 5.78 | 6.5 | 5.72 | DN3ab_Itr | DN3cd | -0.45 | -3.29 | 0.0095 | 0.078 | 0.0024 | ENSMUSG00000029148 | protein_coding | Nrbp1 | nuclear receptor binding protein 1 | 192292 | | 5 | 31240864 | 31251566 | 1 | Mm.292040 |
| 6.6 | 6.87 | 5.53 | 7.02 | DN3ab_Itr | DN3cd | 0.77 | 3.2 | 0.011 | 0.085 | 0.0024 | ENSMUSG00000034353 | protein_coding | Ramp1 | receptor (calcitonin) activity modifying protein 1 | 51801 | | 1 | 91179822 | 91225196 | 1 | Mm.3272 |
| 6.23 | 6.8 | 5.12 | 6.76 | DN3ab_Itr | DN3cd | 0.99 | 4.7 | 0.0012 | 0.02 | 0.0024 | ENSMUSG00000032221 | protein_coding | Mns1 | meiosis-specific nuclear structural protein 1 | 17427 | | 9 | 72438011 | 72462025 | 1 | Mm.387671 |
| 7.85 | 6.87 | 8.65 | 7.05 | DN3ab_Itr | DN3cd | -1.29 | -10.23 | 3.30E-06 | 0.00063 | 0.0024 | ENSMUSG00000031328 | protein_coding | Flna | filamin, alpha | 192176 | X | | 74223461 | 74249820 | -1 | Mm.295533 |

| | | | | | | | | | | | | | | | | | | | | |
|---|---|---|---|---|---|---|---|---|---|---|---|---|---|---|---|---|---|---|---|---|
| 7.1 | 7.2 | 6.22 | 7.06 | DN3ab_Itr | DN3cd | 0.41 | 2.73 | 0.023 | 0.14 | 0.0024 | ENSMUSG00000038025 | protein_coding | Phf2 | PHD finger protein 2 | 18676 | 13 | 48801750 | 48871119 | -1 | Mm.486213 |
| 5.15 | 5.44 | 4.15 | 5.59 | DN3ab_Itr | DN3cd | 0.74 | 3.27 | 0.0098 | 0.079 | 0.0024 | ENSMUSG00000001524 | protein_coding | Gtf2h4 | general transcription factor II H, polypeptide 4 | 14885 | 17 | 35667730 | 35673739 | -1 | Mm.10182 |
| 2.76 | 2.29 | 4.9 | 3 | DN3ab_Itr | DN3cd | -1.38 | -4.92 | 0.00086 | 0.017 | 0.0025 | ENSMUSG00000023571 | protein_coding | C1qtnf12 | C1q and tumor necrosis factor related 12 | 67389 | 4 | 155962318 | 155966629 | 1 | Mm.29140 |
| 3.06 | 3.33 | 1.68 | 3.9 | DN3ab_Itr | DN3cd | 0.94 | 2.49 | 0.035 | 0.18 | 0.0025 | ENSMUSG00000044033 | protein_coding | Ccdc141 | coiled-coil domain containing 141 | 545428 | 2 | 77009902 | 77170636 | -1 | Mm.26579 |
| 5.41 | 6.03 | 4.85 | 6.15 | DN3ab_Itr | DN3cd | 0.92 | 6.65 | 1.00E-04 | 0.0044 | 0.0026 | ENSMUSG00000070738 | protein_coding | Dgkd | diacylglycerol kinase, delta | 227333 | 1 | 87853287 | 87945180 | 1 | Mm.277217 |
| 5.14 | 4.59 | 6.07 | 4.36 | DN3ab_Itr | DN3cd | -1.11 | -4.71 | 0.0012 | 0.02 | 0.0026 | ENSMUSG00000024190 | protein_coding | Dusp1 | dual specificity phosphatase 1 | 19252 | 17 | 26505590 | 26508519 | -1 | Mm.239041 |
| 7.71 | 7.85 | 7.65 | 8.25 | DN3ab_Itr | DN3cd | 0.35 | 3.79 | 0.0044 | 0.046 | 0.0026 | ENSMUSG00000094483 | protein_coding | Purb | purine rich element binding protein B | 19291 | 11 | 6467599 | 6475917 | -1 | Mm.296150 Mm.390332 Mm.474938 |
| 4.33 | 4.75 | 1.64 | 3.72 | DN3ab_Itr | DN3cd | 0.75 | 2.79 | 0.021 | 0.13 | 0.0026 | ENSMUSG00000040658 | protein_coding | Dnph1 | 2'-deoxynucleoside 5'-phosphate N-hydrolase 1 | 381101 | 17 | 46496711 | 46499624 | 1 | Mm.35801 |
| 5.82 | 5.66 | 6.64 | 5.76 | DN3ab_Itr | DN3cd | -0.52 | -3.5 | 0.0069 | 0.062 | 0.0027 | ENSMUSG00000001794 | protein_coding | Capns1 | calpain, small subunit 1 | 12336 | 7 | 30186936 | 30198811 | -1 | Mm.439782 |
| 7.75 | 8.23 | 6.88 | 8.05 | DN3ab_Itr | DN3cd | 0.77 | 5.54 | 0.00038 | 0.01 | 0.0027 | ENSMUSG00000057406 | protein_coding | Nsd2 | nuclear receptor binding SET domain protein 2 | 107823 | 5 | 33820725 | 33897975 | 1 | Mm.19892 Mm.332320 Mm.491382 |
| 4.83 | 4.92 | 4.08 | 4.92 | DN3ab_Itr | DN3cd | 0.4 | 2.6 | 0.029 | 0.16 | 0.0027 | ENSMUSG00000031262 | protein_coding | Cenpi | centromere protein I | 102920 | X | 134308084 | 134362639 | 1 | Mm.34903 |
| 3.34 | 4.03 | 2.43 | 4.37 | DN3ab_Itr | DN3cd | 1.15 | 4.66 | 0.0012 | 0.021 | 0.0028 | ENSMUSG00000001053 | protein_coding | N4bp3 | NEDD4 binding protein 3 | 212706 | 11 | 51643063 | 51650842 | -1 | Mm.41760 |
| 7.98 | 7.81 | 8.59 | 7.87 | DN3ab_Itr | DN3cd | -0.44 | -3.97 | 0.0034 | 0.039 | 0.0028 | ENSMUSG00000052681 | protein_coding | Rap1b | RAS related protein 1b | 215449 | 10 | 117813871 | 117846035 | -1 | Mm.233009 Mm.491159 |
| 3.62 | 4.19 | 3.05 | 4.67 | DN3ab_Itr | DN3cd | 1.01 | 4.7 | 0.0012 | 0.02 | 0.0028 | ENSMUSG00000027329 | protein_coding | Spef1 | sperm flagellar 1 | 70997 | 2 | 131170261 | 131187282 | -1 | Mm.252722 |
| 4.29 | 5.11 | 4.4 | 4.45 | DN3ab_Itr | DN3cd | 0.49 | 3.06 | 0.014 | 0.099 | 0.0028 | ENSMUSG00000013846 | protein_coding | St3gal1 | ST3 beta-galactoside alpha-2,3-sialyltransferase 1 | 20442 | 15 | 67102875 | 67176830 | -1 | Mm.248334 Mm.442417 Mm.461082 |
| 1.51 | 2.12 | -0.15 | 2.54 | DN3ab_Itr | DN3cd | 1.27 | 3.17 | 0.012 | 0.088 | 0.0029 | ENSMUSG00000077506 | snoRNA | Scarna9 | small Cajal body-specific RNA 9 | NA | 9 | 15326264 | 15326333 | -1 | |
| 2.53 | 4.51 | -1.16 | 4.73 | DN3ab_Itr | DN3cd | 2.5 | 4.66 | 0.0012 | 0.021 | 0.0029 | ENSMUSG00000072553 | protein_coding | Gm525 | predicted gene 525 | 217071 | 11 | 89073841 | 89093040 | 1 | Mm.342127 |
| 3.88 | 4.28 | 2.84 | 4.31 | DN3ab_Itr | DN3cd | 0.79 | 3.7 | 0.005 | 0.051 | 0.0029 | ENSMUSG00000081622 | processed_pseudogene | Gm5937 | predicted gene 5937 | NA | X | 77456586 | 77457228 | 1 | |
| 6.44 | 5.81 | 6.15 | 6.08 | DN3ab_Itr | DN3cd | -0.37 | -3.17 | 0.012 | 0.088 | 0.0029 | ENSMUSG00000028630 | protein_coding | Dyrk2 | dual-specificity tyrosine-(Y)-phosphorylation regulated kinase 2 | 69181 | 10 | 118855603 | 118870209 | -1 | Mm.45565 Mm.458174 |
| 5.53 | 5.77 | 5.44 | 6.45 | DN3ab_Itr | DN3cd | 0.61 | 3.83 | 0.0041 | 0.045 | 0.0029 | ENSMUSG00000022106 | protein_coding | Rcbtb2 | regulator of chromosome condensation (RCC1) and BTB (POZ) domain containing protein 2 | 105670 | 14 | 73123037 | 73207843 | 1 | Mm.280068 |
| -0.24 | 0.85 | -3.72 | 1.33 | DN3ab_Itr | DN3cd | 2.67 | 3.36 | 0.0085 | 0.072 | 0.0029 | ENSMUSG00000108232 | antisense | Gm43916 | predicted gene, 43916 | NA | 6 | 120951901 | 120952867 | 1 | |
| 4.6 | 4.87 | 3.66 | 5 | DN3ab_Itr | DN3cd | 0.68 | 3.16 | 0.012 | 0.089 | 0.0029 | ENSMUSG00000025036 | protein_coding | Sfxn2 | sideroflexin 2 | 94279 | 19 | 46573365 | 46596898 | 1 | Mm.296837 |
| 4.98 | 4.5 | 7.89 | 6.15 | DN3ab_Itr | DN3cd | -1.33 | -5.54 | 0.00038 | 0.01 | 0.0029 | ENSMUSG00000028480 | protein_coding | Glipr2 | GLI pathogenesis-related 2 | 384009 | 4 | 43957401 | 43979118 | 1 | Mm.22213 |
| 12.72 | 12.85 | 11.99 | 12.52 | DN3ab_Itr | DN3cd | 0.28 | 3.56 | 0.0063 | 0.059 | 0.0029 | ENSMUSG00000026238 | protein_coding | Ptma | prothymosin alpha | 19231 | 1 | 86526726 | 86530712 | 1 | Mm.19187 |
| 7.7 | 7.91 | 6.81 | 7.64 | DN3ab_Itr | DN3cd | 0.47 | 3.81 | 0.0043 | 0.046 | 0.003 | ENSMUSG00000038384 | protein_coding | Setd1b | SET domain containing 1B | 208043 | 5 | 123142193 | 123168629 | 1 | Mm.250391 |
| 5.3 | 5.62 | 4 | 5.13 | DN3ab_Itr | DN3cd | 0.6 | 3.75 | 0.0047 | 0.048 | 0.003 | ENSMUSG00000041064 | protein_coding | Pif1 | PIF1 5'-to-3' DNA helicase | 208084 | 9 | 65587160 | 65595967 | 1 | Mm.211032 |
| 5.63 | 5.57 | 6.02 | 5.31 | DN3ab_Itr | DN3cd | -0.36 | -2.67 | 0.026 | 0.15 | 0.003 | ENSMUSG00000036835 | protein_coding | Psenen | presenilin enhancer gamma secretase subunit | 66340 | 7 | 30561863 | 30563627 | -1 | Mm.86557 |
| 13.08 | 13.16 | 12.55 | 13.24 | DN3ab_Itr | DN3cd | 0.34 | 2.88 | 0.018 | 0.12 | 0.0031 | ENSMUSG00000024608 | protein_coding | Rps14 | ribosomal protein S14 | 20044 | 18 | 60774510 | 60778546 | 1 | Mm.43778 |
| 6.13 | 5.93 | 6.81 | 5.93 | DN3ab_Itr | DN3cd | -0.54 | -3.86 | 0.004 | 0.044 | 0.0031 | ENSMUSG00000039316 | protein_coding | Rftn1 | raftlin lipid raft linker 1 | 76438 | 17 | 49992257 | 50190674 | -1 | Mm.41854 |
| 1.41 | 2.19 | -0.23 | 2.99 | DN3ab_Itr | DN3cd | 1.57 | 3.31 | 0.0092 | 0.076 | 0.0031 | ENSMUSG00000086060 | lincRNA | Gm16084 | predicted gene 16084 | NA | 12 | 107147164 | 107160207 | -1 | |
| 0.37 | 1.18 | -1.76 | 2.41 | DN3ab_Itr | DN3cd | 2.02 | 3.03 | 0.014 | 0.1 | 0.0031 | ENSMUSG00000027249 | protein_coding | F2 | coagulation factor II | 14061 | 2 | 91625320 | 91636414 | -1 | Mm.89048 |
| -1.45 | -0.12 | -3.72 | 0.6 | DN3ab_Itr | DN3cd | 2.83 | 4.51 | 0.0015 | 0.024 | 0.0032 | ENSMUSG00000036091 | protein_coding | Hyal3 | hyaluronoglucosaminidase 3 | 109685 | 9 | 107580818 | 107587359 | 1 | Mm.447916 |
| 4.03 | 3.04 | 5.82 | 3.7 | DN3ab_Itr | DN3cd | -1.67 | -7.34 | 4.70E-05 | 0.0029 | 0.0032 | ENSMUSG00000037992 | protein_coding | Rara | retinoic acid receptor, alpha | 19401 | 11 | 98927818 | 98974942 | 1 | Mm.439744 |
| 8.77 | 5.92 | 9.44 | 7.91 | DN3ab_Itr | DN3cd | -2.06 | -7.82 | 2.90E-05 | 0.0022 | 0.0032 | ENSMUSG00000024677 | protein_coding | Ms4a6b | membrane-spanning 4-domains, subfamily A, member 6B | 69774 | 19 | 11516512 | 11531256 | 1 | Mm.436639 |
| 7.95 | 8.08 | 7.31 | 8.02 | DN3ab_Itr | DN3cd | 0.38 | 3.23 | 0.011 | 0.083 | 0.0032 | ENSMUSG00000019818 | protein_coding | Cd164 | CD164 antigen | 53599 | 10 | 41519414 | 41531048 | 1 | Mm.269815 |
| 7.82 | 8.08 | 6.93 | 7.71 | DN3ab_Itr | DN3cd | 0.47 | 4.47 | 0.0016 | 0.025 | 0.0033 | ENSMUSG00000011179 | protein_coding | Odc1 | ornithine decarboxylase, structural 1 | 18263 | 12 | 17544794 | 17551505 | 1 | Mm.34102 Mm.472891 |
| 3.09 | 3.47 | 3.73 | 5.22 | DN3ab_Itr | DN3cd | 1.03 | 4.52 | 0.0015 | 0.024 | 0.0033 | ENSMUSG00000066258 | protein_coding | Trim12a | tripartite motif-containing 12A | 76681 | 7 | 104299894 | 104315466 | -1 | Mm.458309 Mm.486388 |
| 8.71 | 8.31 | 10.27 | 9.12 | DN3ab_Itr | DN3cd | -0.82 | -5.43 | 0.00044 | 0.011 | 0.0033 | ENSMUSG00000024014 | protein_coding | Pim1 | proviral integration site 1 | 18712 | 17 | 29490753 | 29496112 | 1 | Mm.405293 Mm.485038 |
| 4.1 | 4.32 | 2.83 | 4.27 | DN3ab_Itr | DN3cd | 0.64 | 2.66 | 0.026 | 0.15 | 0.0034 | ENSMUSG00000028035 | protein_coding | Dnajb4 | DnaJ heat shock protein family (Hsp40) member B4 | 67035 | 3 | 152178511 | 152210302 | -1 | Mm.46746 Mm.458538 |
| 3.34 | 2.8 | 4.63 | 3 | DN3ab_Itr | DN3cd | -1.13 | -5 | 0.00077 | 0.016 | 0.0034 | ENSMUSG00000078695 | protein_coding | Cisd3 | CDGSH iron sulfur domain 3 | 217149 | 11 | 97685826 | 97688621 | 1 | Mm.29659 |
| -2.06 | 1.28 | -3.72 | 2.4 | DN3ab_Itr | DN3cd | 4.75 | 8.12 | 2.10E-05 | 0.0019 | 0.0035 | ENSMUSG00000047501 | protein_coding | Cldn4 | claudin 4 | 12740 | 5 | 134945119 | 134946934 | -1 | Mm.7339 |
| 7.32 | 7.93 | 6.5 | 7.55 | DN3ab_Itr | DN3cd | 0.79 | 8.58 | 1.40E-05 | 0.0014 | 0.0035 | ENSMUSG00000042489 | protein_coding | Clspn | claspin | 269582 | 4 | 126556935 | 126593903 | 1 | Mm.243962 |
| 3.81 | 0.52 | 5.03 | 3.71 | DN3ab_Itr | DN3cd | -1.7 | -5.29 | 0.00052 | 0.012 | 0.0035 | ENSMUSG00000020034 | protein_coding | Tcp11l2 | t-complex 11 (mouse) like 2 | 216198 | 10 | 84576626 | 84614359 | 1 | Mm.296142 |
| 5.8 | 5.6 | 6.43 | 5.43 | DN3ab_Itr | DN3cd | -0.59 | -3.59 | 0.006 | 0.057 | 0.0037 | ENSMUSG00000059248 | protein_coding | 9-Sep | septin 9 | 53860 | 11 | 117199661 | 117362325 | 1 | Mm.38450 Mm.451420 |
| 7.64 | 7.82 | 6.71 | 7.86 | DN3ab_Itr | DN3cd | 0.59 | 2.99 | 0.016 | 0.11 | 0.0037 | ENSMUSG00000046985 | protein_coding | Tapt1 | transmembrane anterior posterior transformation 1 | 231225 | 5 | 44175154 | 44226626 | -1 | Mm.30688 Mm.369319 |
| 8.18 | 7.84 | 8.82 | 8.13 | DN3ab_Itr | DN3cd | -0.52 | -6.98 | 7.00E-05 | 0.0036 | 0.0037 | ENSMUSG00000030220 | protein_coding | Arhgdib | Rho, GDP dissociation inhibitor (GDI) beta | 11857 | 6 | 136923655 | 136941899 | -1 | Mm.2241 |
| 5.95 | 5.56 | 6.45 | 5.54 | DN3ab_Itr | DN3cd | -0.64 | -5.88 | 0.00025 | 0.0079 | 0.0037 | ENSMUSG00000032402 | protein_coding | Smad3 | SMAD family member 3 | 17127 | 9 | 63646767 | 63757994 | -1 | Mm.7320 |
| 9.34 | 9.59 | 8.03 | 8.94 | DN3ab_Itr | DN3cd | 0.49 | 3.76 | 0.0046 | 0.048 | 0.0037 | ENSMUSG00000070713 | protein_coding | Gm10282 | predicted pseudogene 10282 | NA | 8 | 72304110 | 72305260 | -1 | |
| 5.86 | 4.81 | 5.9 | 5.5 | DN3ab_Itr | DN3cd | -0.72 | -5.24 | 0.00056 | 0.013 | 0.0038 | ENSMUSG00000032812 | protein_coding | Arap1 | ArfGAP with RhoGAP domain, ankyrin repeat and PH domain 1 | 69710 | 7 | 101348067 | 101412586 | 1 | Mm.277687 |
| 4.1 | 4.99 | 2.54 | 4.83 | DN3ab_Itr | DN3cd | 1.31 | 4.93 | 0.00085 | 0.017 | 0.0038 | ENSMUSG00000032501 | protein_coding | Trib1 | tribbles pseudokinase 1 | 211770 | 15 | 59648350 | 59657099 | 1 | Mm.40298 |
| 6.46 | 6.19 | 7.36 | 6.63 | DN3ab_Itr | DN3cd | -0.52 | -5.33 | 5.00E-04 | 0.012 | 0.0038 | ENSMUSG00000021728 | protein_coding | Emb | embigin | 13723 | 13 | 117208536 | 117274415 | 1 | Mm.274926 |
| 4.19 | 3.71 | 4.69 | 2.94 | DN3ab_Itr | DN3cd | -1.01 | -3.85 | 0.004 | 0.044 | 0.0038 | ENSMUSG00000040570 | protein_coding | Rundc3b | RUN domain containing 3B | 242819 | 5 | 8490334 | 8622952 | -1 | Mm.332192 |
| 4.24 | 3.33 | 5.77 | 3.89 | DN3ab_Itr | DN3cd | -1.47 | -7.37 | 4.60E-05 | 0.0029 | 0.0038 | ENSMUSG00000032666 | protein_coding | 1700025G04Rik | RIKEN cDNA 1700025G04 gene | 69399 | 1 | 151852403 | 152090125 | -1 | Mm.278031 |
| 8.34 | 8.52 | 7.68 | 8.36 | DN3ab_Itr | DN3cd | 0.39 | 3.81 | 0.0043 | 0.046 | 0.0038 | ENSMUSG00000032481 | protein_coding | Smarcc1 | SWI/SNF related, matrix associated, actin dependent regulator of chromatin, subfamily c, member 1 | 20588 | 9 | 110117708 | 110240178 | 1 | Mm.85410 |
| 3.53 | 3.21 | 4.9 | 3.68 | DN3ab_Itr | DN3cd | -0.84 | -4.48 | 0.0016 | 0.025 | 0.0039 | ENSMUSG00000021959 | protein_coding | Lats2 | large tumor suppressor 2 | 50523 | 14 | 57689662 | 57758388 | -1 | Mm.347899 |
| 7.05 | 7.29 | 5.94 | 6.63 | DN3ab_Itr | DN3cd | 0.42 | 4.48 | 0.0016 | 0.025 | 0.0039 | ENSMUSG00000003779 | protein_coding | Kif20a | kinesin family member 20A | 19348 | 18 | 34624624 | 34633265 | 1 | Mm.258846 |
| 4.28 | 5.06 | 2.4 | 5.04 | DN3ab_Itr | DN3cd | 1.29 | 3.75 | 0.0047 | 0.048 | 0.0039 | ENSMUSG00000067750 | protein_coding | Khdc1a | KH domain containing 1A | 368204 | 1 | 21349641 | 21352200 | 1 | Mm.26006 |
| 1.71 | 2.28 | 0.56 | 3 | DN3ab_Itr | DN3cd | 1.25 | 3.29 | 0.0096 | 0.078 | 0.0039 | ENSMUSG00000021614 | protein_coding | Vcan | versican | 13003 | 13 | 89655312 | 89742509 | -1 | Mm.158700 Mm.410783 |
| 5.19 | 6.25 | 4.68 | 6.7 | DN3ab_Itr | DN3cd | 1.47 | 7.48 | 4.10E-05 | 0.0027 | 0.004 | ENSMUSG00000002033 | protein_coding | Cd3g | CD3 antigen, gamma polypeptide | 12502 | 9 | 44969572 | 44980431 | -1 | Mm.335106 |
| 6.17 | 6.56 | 5.38 | 6.28 | DN3ab_Itr | DN3cd | 0.6 | 5.61 | 0.00035 | 0.0095 | 0.004 | ENSMUSG00000039994 | protein_coding | Timeless | timeless circadian clock 1 | 21853 | 10 | 128232065 | 128252941 | 1 | Mm.6458 |
| 3.89 | 4.32 | 2.49 | 4.79 | DN3ab_Itr | DN3cd | 1.06 | 2.88 | 0.018 | 0.12 | 0.0041 | ENSMUSG00000035547 | protein_coding | Capn5 | calpain 5 | 12337 | 7 | 98121559 | 98178274 | -1 | Mm.326847 |
| 7.73 | 8.33 | 7.57 | 7.61 | DN3ab_Itr | DN3cd | 0.37 | 3.2 | 0.011 | 0.085 | 0.0041 | ENSMUSG00000020923 | protein_coding | Ubtf | upstream binding transcription factor, RNA polymerase I | 21429 | 11 | 102304560 | 102319742 | -1 | Mm.2845 |
| 6.43 | 6.75 | 5.66 | 6.65 | DN3ab_Itr | DN3cd | 0.59 | 4.28 | 0.0021 | 0.029 | 0.0041 | ENSMUSG00000031821 | protein_coding | Gins2 | GINS complex subunit 2 (Psf2 homolog) | 272551 | 8 | 120578633 | 120589304 | -1 | Mm.275910 Mm.334856 |
| 6.96 | 8.52 | 6.75 | 8.89 | DN3ab_Itr | DN3cd | 1.83 | 14.96 | 1.30E-07 | 0.00017 | 0.0042 | ENSMUSG00000059751 | processed_pseudogene | Rps3a3 | ribosomal protein S3A3 | NA | 13 | 108670603 | 108671397 | 1 | |
| 7.83 | 7.92 | 7.49 | 8.08 | DN3ab_Itr | DN3cd | 0.31 | 2.97 | 0.016 | 0.11 | 0.0042 | ENSMUSG00000002102 | protein_coding | Psmc3 | proteasome (prosome, macropain) 26S subunit, ATPase 3 | 19182 | 2 | 91054009 | 91066369 | 1 | Mm.289832 |
| 4.09 | 3.86 | 5.19 | 4.07 | DN3ab_Itr | DN3cd | -0.71 | -3.77 | 0.0046 | 0.048 | 0.0042 | ENSMUSG00000038650 | protein_coding | Rnh1 | ribonuclease/angiogenin inhibitor 1 | 107702 | 7 | 141160326 | 141172857 | -1 | Mm.197493 Mm.279485 |
| 5.29 | 4.31 | 8.36 | 5.77 | DN3ab_Itr | DN3cd | -2.05 | -6.51 | 0.00012 | 0.005 | 0.0043 | ENSMUSG00000058755 | protein_coding | Osm | oncostatin M | 18413 | 11 | 4236420 | 4241026 | 1 | Mm.131422 |
| 7.6 | 7.34 | 8.34 | 7.42 | DN3ab_Itr | DN3cd | -0.59 | -4.31 | 0.002 | 0.028 | 0.0043 | ENSMUSG00000008475 | protein_coding | Arpc5 | actin related protein 2/3 complex, subunit 5 | 67771 | 1 | 152766542 | 152775597 | 1 | Mm.288974 |
| 5.15 | 5.38 | 4.8 | 5.68 | DN3ab_Itr | DN3cd | 0.52 | 3.84 | 0.0041 | 0.044 | 0.0044 | ENSMUSG00000022474 | protein_coding | Pmm1 | phosphomannomutase 1 | 29858 | 15 | 81951108 | 81960930 | -1 | Mm.18939 |
| 4.16 | 4.65 | 2.96 | 4.3 | DN3ab_Itr | DN3cd | 0.77 | 4.48 | 0.0016 | 0.025 | 0.0044 | ENSMUSG00000019214 | protein_coding | Chtf18 | CTF18, chromosome transmission fidelity factor 18 | 214901 | 17 | 25718926 | 25727419 | -1 | Mm.102540 |
| 4.43 | 6.82 | 6.09 | 6.91 | DN3ab_Itr | DN3cd | 1.48 | 4.57 | 0.0014 | 0.023 | 0.0044 | ENSMUSG00000005125 | protein_coding | Ndrg1 | N-myc downstream regulated gene 1 | 17988 | 15 | 66929318 | 66969640 | -1 | Mm.30837 |
| 5.52 | 5.87 | 4.63 | 5.88 | DN3ab_Itr | DN3cd | 0.71 | 3.81 | 0.0043 | 0.046 | 0.0044 | ENSMUSG00000092281 | processed_pseudogene | Hmgb1-ps7 | high-mobility group high mobility group box 1, pseudogene 7 | NA | 7 | 42020836 | 42021483 | 1 | |
| 7.21 | 7.67 | 6.09 | 6.99 | DN3ab_Itr | DN3cd | 0.63 | 6.78 | 8.70E-05 | 0.0041 | 0.0045 | ENSMUSG00000023505 | protein_coding | Cdca3 | cell division cycle associated 3 | 14793 | 6 | 124829547 | 124833701 | 1 | Mm.285723 |
| 4 | 4.89 | 3.01 | 4.97 | DN3ab_Itr | DN3cd | 1.28 | 5.95 | 0.00023 | 0.0074 | 0.0045 | ENSMUSG00000038264 | protein_coding | Sema7a | sema domain, immunoglobulin domain (Ig), and GPI membrane anchor, (semaphorin) 7A | 20361 | 9 | 57940112 | 57962865 | 1 | Mm.335187 |
| 3.74 | 4.33 | 0.23 | 4.18 | DN3ab_Itr | DN3cd | 1.07 | 2.19 | 0.057 | 0.23 | 0.0045 | ENSMUSG00000032864 | protein_coding | Rag2 | recombination activating gene 2 | 19374 | 2 | 101624718 | 101632529 | 1 | Mm.4988 |
| 1.43 | -3.12 | 2.91 | 1.11 | DN3ab_Itr | DN3cd | -2.53 | -4.93 | 0.00085 | 0.016 | 0.0045 | ENSMUSG00000024679 | protein_coding | Ms4a6d | membrane-spanning 4-domains, subfamily A, member 6D | 68774 | 19 | 11586604 | 11604849 | -1 | Mm.290390 |
| -0.58 | 0.78 | -3.72 | 0.17 | DN3ab_Itr | DN3cd | 2.35 | 4.42 | 0.0017 | 0.026 | 0.0045 | ENSMUSG00000056032 | antisense | BC018473 | cDNA sequence BC018473 | NA | 11 | 116752167 | 116759373 | 1 | |
| 6.37 | 5.75 | 6.17 | 6.01 | DN3ab_Itr | DN3cd | -0.41 | -4.2 | 0.0024 | 0.031 | 0.0045 | ENSMUSG00000010142 | protein_coding | Tnfrsf13b | tumor necrosis factor receptor superfamily, member 13b | 57916 | 11 | 61126755 | 61149372 | 1 | Mm.265915 |
| 4.84 | 4.37 | 5.42 | 4.03 | DN3ab_Itr | DN3cd | -0.9 | -4.63 | 0.0013 | 0.022 | 0.0046 | ENSMUSG00000020978 | protein_coding | Klhdc2 | kelch domain containing 2 | 69554 | 12 | 69296681 | 69310687 | 1 | Mm.234368 |
| -0.7 | -0.03 | -1.78 | 1.73 | DN3ab_Itr | DN3cd | 2.05 | 3.37 | 0.0084 | 0.071 | 0.0047 | ENSMUSG00000111006 | TEC | Gm10826 | predicted gene 10826 | NA | 10 | 21030490 | 21033793 | -1 | |
| 1.84 | 2.24 | 0.62 | 2.84 | DN3ab_Itr | DN3cd | 1.05 | 2.83 | 0.02 | 0.13 | 0.0047 | ENSMUSG00000054934 | protein_coding | Kcnmb4 | potassium large conductance calcium-activated channel, subfamily M, beta member 4 | 58802 | 10 | 116417861 | 116473878 | -1 | Mm.440652 |
| 4.65 | 6.1 | 3.21 | 6.26 | DN3ab_Itr | DN3cd | 1.96 | 6.27 | 0.00015 | 0.0059 | 0.0048 | ENSMUSG00000021477 | protein_coding | Ctsl | cathepsin L | 13039 | 13 | 64359337 | 64370890 | -1 | Mm.930 Mm.471755 |
| 6.32 | 6.1 | 6.98 | 6.18 | DN3ab_Itr | DN3cd | -0.51 | -4.1 | 0.0027 | 0.034 | 0.005 | ENSMUSG00000031103 | protein_coding | Elf4 | E74-like factor 4 (ets domain transcription factor) | 56501 | X | 48411046 | 48463132 | -1 | Mm.36808 |
| 6.22 | 6.43 | 5.58 | 6.31 | DN3ab_Itr | DN3cd | 0.43 | 3.89 | 0.0038 | 0.042 | 0.005 | ENSMUSG00000035293 | protein_coding | G2e3 | G2/M-phase specific E3 ubiquitin ligase | 217558 | 12 | 51348061 | 51376986 | 1 | Mm.253264 |
| 11.29 | 11.75 | 10.37 | 11.22 | DN3ab_Itr | DN3cd | 0.61 | 7.64 | 3.50E-05 | 0.0025 | 0.0051 | ENSMUSG00000023004 | protein_coding | Tuba1b | tubulin, alpha 1B | 22143 | 15 | 98931425 | 98934565 | -1 | Mm.392113 Mm.491454 |
| 8.19 | 6.18 | 8.56 | 7.29 | DN3ab_Itr | DN3cd | -1.61 | -10.26 | 3.20E-06 | 0.00063 | 0.0051 | ENSMUSG00000025997 | protein_coding | Ikzf2 | IKAROS family zinc finger 2 | 22779 | 1 | 69531214 | 69687245 | -1 | Mm.106343 |
| 3.65 | 5.75 | 3.65 | 5.08 | DN3ab_Itr | DN3cd | 1.81 | 12.52 | 6.10E-07 | 3.00E-04 | 0.0051 | ENSMUSG00000063320 | protein_coding | 1190007I07Rik | RIKEN cDNA 1190007I07 gene | 544717 | 10 | 82619851 | 82623228 | -1 | Mm.334867 |
| 1.89 | 3.5 | -1.82 | 3.93 | DN3ab_Itr | DN3cd | 2.34 | 3.51 | 0.0068 | 0.062 | 0.0053 | ENSMUSG00000048058 | protein_coding | Ldlrad3 | low density lipoprotein receptor class A domain containing 3 | 241576 | 2 | 101950203 | 102186385 | -1 | Mm.489740 |
| 6.58 | 6.67 | 5.99 | 6.53 | DN3ab_Itr | DN3cd | 0.28 | 2.95 | 0.017 | 0.11 | 0.0053 | ENSMUSG00000019715 | protein_coding | Gle1 | GLE1 RNA export mediator (yeast) | 74412 | 2 | 29935426 | 29960371 | 1 | Mm.275121 |
| 6.17 | 6.32 | 5.92 | 6.86 | DN3ab_Itr | DN3cd | 0.52 | 3.11 | 0.013 | 0.094 | 0.0053 | ENSMUSG00000064368 | protein_coding | mt-Nd6 | mitochondrially encoded NADH dehydrogenase 6 | 17722 | MT | 13552 | 14070 | -1 | |
| 7.09 | 6 | 7.38 | 7.04 | DN3ab_Itr | DN3cd | -0.69 | -4.39 | 0.0018 | 0.027 | 0.0054 | ENSMUSG00000035621 | protein_coding | Midn | midnolin | 59090 | 10 | 80148272 | 80158368 | 1 | Mm.143813 |
| 5.66 | 6.03 | 4.8 | 5.81 | DN3ab_Itr | DN3cd | 0.62 | 4.56 | 0.0014 | 0.023 | 0.0054 | ENSMUSG00000018858 | protein_coding | Mrpl58 | mitochondrial ribosomal protein L58 | 68572 | 11 | 115403752 | 115412674 | 1 | Mm.271062 |
| 6.75 | 6.42 | 7.28 | 6.53 | DN3ab_Itr | DN3cd | -0.53 | -5.71 | 0.00031 | 0.0088 | 0.0054 | ENSMUSG00000005054 | protein_coding | Cstb | cystatin B | 13014 | 10 | 78425669 | 78427622 | 1 | Mm.6095 |
| 6.91 | 7.04 | 5.81 | 6.45 | DN3ab_Itr | DN3cd | 0.33 | 3.07 | 0.014 | 0.098 | 0.0055 | ENSMUSG00000036639 | protein_coding | Nudt1 | nudix (nucleoside diphosphate linked moiety X)-type motif 1 | 17766 | 5 | 140321656 | 140338137 | 1 | Mm.118846 |
| 3.28 | 4.8 | 3.93 | 4.5 | DN3ab_Itr | DN3cd | 1.05 | 5.14 | 0.00064 | 0.014 | 0.0055 | ENSMUSG00000079491 | polymorphic_pseudogene | H2-T10 | histocompatibility 2, T region locus 10 | NA | 17 | 36115876 | 36121465 | -1 | |
| 6.9 | 7.22 | 5.92 | 7.01 | DN3ab_Itr | DN3cd | 0.63 | 3.98 | 0.0033 | 0.039 | 0.0056 | ENSMUSG00000064210 | protein_coding | Ano6 | anoctamin 6 | 105722 | 15 | 95790843 | 95974751 | 1 | Mm.38087 |
| 8.46 | 8.02 | 8.36 | 8.25 | DN3ab_Itr | DN3cd | -0.28 | -3.97 | 0.0033 | 0.039 | 0.0056 | ENSMUSG00000029131 | protein_coding | Dnajb6 | DnaJ heat shock protein family (Hsp40) member B6 | 23950 | 5 | 29735688 | 29818134 | 1 | Mm.290110 |
| 3.93 | 4.2 | 2.66 | 4.39 | DN3ab_Itr | DN3cd | 0.78 | 2.64 | 0.027 | 0.15 | 0.0056 | ENSMUSG00000030852 | protein_coding | Tacc2 | transforming, acidic coiled-coil containing protein 2 | 57752 | 7 | 130577438 | 130764785 | 1 | Mm.86322 |
| 6.23 | 6.12 | 6.53 | 5.88 | DN3ab_Itr | DN3cd | -0.36 | -3.12 | 0.012 | 0.092 | 0.0057 | ENSMUSG00000049922 | protein_coding | Slc35c1 | solute carrier family 35, member C1 | 228368 | 2 | 92452764 | 92460538 | -1 | Mm.273990 |
| 4.17 | 3.95 | 5.94 | 4.94 | DN3ab_Itr | DN3cd | -0.7 | -4.25 | 0.0022 | 0.03 | 0.0057 | ENSMUSG00000028793 | protein_coding | Rnf19b | ring finger protein 19B | 75234 | 4 | 129058271 | 129085886 | 1 | Mm.259672 Mm.486170 |
| 5.56 | 5.99 | 4.74 | 5.93 | DN3ab_Itr | DN3cd | 0.74 | 4.56 | 0.0014 | 0.023 | 0.0057 | ENSMUSG00000068466 | processed_pseudogene | Gm5518 | predicted gene 5518 | NA | 19 | 29160118 | 29160765 | 1 | |
| 4.11 | 2.95 | 4.01 | 3.77 | DN3ab_Itr | DN3cd | -0.68 | -3.4 | 0.008 | 0.069 | 0.0058 | ENSMUSG00000020883 | protein_coding | Fbxl20 | F-box and leucine-rich repeat protein 20 | 72194 | 11 | 98082556 | 98150403 | -1 | Mm.218350 |
| 9.34 | 9.65 | 9.07 | 9.63 | DN3ab_Itr | DN3cd | 0.42 | 7.52 | 3.90E-05 | 0.0026 | 0.0058 | ENSMUSG00000030654 | protein_coding | Arl6ip1 | ADP-ribosylation factor-like 6 interacting protein 1 | 54208 | 7 | 118118891 | 118129662 | -1 | Mm.29924 |
| 5.11 | 5.88 | 4.96 | 5.07 | DN3ab_Itr | DN3cd | 0.5 | 3.55 | 0.0063 | 0.059 | 0.0058 | ENSMUSG00000002068 | protein_coding | Ccne1 | cyclin E1 | 12447 | 7 | 38097984 | 38107534 | -1 | Mm.16110 |
| 9.88 | 8.96 | 9.66 | 9.3 | DN3ab_Itr | DN3cd | -0.66 | -5.54 | 0.00038 | 0.01 | 0.0059 | ENSMUSG00000062611 | processed_pseudogene | Rps3a2 | ribosomal protein S3A2 | NA | 14 | 88123017 | 88123808 | -1 | |
| 5.2 | 5.58 | 4.5 | 6.11 | DN3ab_Itr | DN3cd | 0.91 | 3.5 | 0.0069 | 0.062 | 0.006 | ENSMUSG00000079037 | protein_coding | Prnp | prion protein | 19122 | 2 | 131909928 | 131938429 | 1 | Mm.648 |
| 0.37 | -0.22 | 3.04 | 0.69 | DN3ab_Itr | DN3cd | -1.76 | -4.83 | 0.00097 | 0.018 | 0.006 | ENSMUSG00000026796 | protein_coding | Fam129b | family with sequence similarity 129, member B | 227737 | 2 | 32876114 | 32925254 | 1 | Mm.301646 |
| 8.01 | 6.35 | 10.75 | 8.37 | DN3ab_Itr | DN3cd | -2.14 | -14.81 | 1.50E-07 | 0.00018 | 0.006 | ENSMUSG00000026728 | protein_coding | Vim | vimentin | 22352 | 2 | 13573927 | 13582826 | 1 | Mm.268000 |
| 3.14 | 3.45 | 1.56 | 3.33 | DN3ab_Itr | DN3cd | 0.76 | 2.62 | 0.028 | 0.15 | 0.0061 | ENSMUSG00000108455 | TEC | Gm44745 | predicted gene 44745 | NA | 7 | 122463438 | 122466160 | 1 | |
| 8.35 | 9.07 | 8.17 | 8.4 | DN3ab_Itr | DN3cd | 0.51 | 4.93 | 0.00084 | 0.016 | 0.0061 | ENSMUSG00000056612 | protein_coding | Ppp1r14b | protein phosphatase 1, regulatory inhibitor subunit 14B | 18938 | 19 | 6975048 | 6977324 | 1 | Mm.140 |
| 10.47 | 10.65 | 9.85 | 10.42 | DN3ab_Itr | DN3cd | 0.34 | 4.19 | 0.0024 | 0.032 | 0.0062 | ENSMUSG00000029836 | protein_coding | Cbx3 | chromobox 3 | 12417 | 6 | 51470360 | 51483704 | 1 | Mm.280968 Mm.393602 Mm.458234 Mm.490361 |
| 7.02 | 7.45 | 5.83 | 6.92 | DN3ab_Itr | DN3cd | 0.68 | 4.92 | 0.00086 | 0.017 | 0.0063 | ENSMUSG00000045273 | protein_coding | Cenph | centromere protein H | 26886 | 13 | 100759674 | 100775899 | -1 | Mm.273502 |
| 5.32 | 5.49 | 4.18 | 5.06 | DN3ab_Itr | DN3cd | 0.42 | 2.85 | 0.019 | 0.12 | 0.0063 | ENSMUSG00000022322 | protein_coding | Shcbp1 | Shc SH2-domain binding protein 1 | 20419 | 8 | 4735976 | 4779567 | -1 | Mm.37801 |
| 7.56 | 7.22 | 8.16 | 7.23 | DN3ab_Itr | DN3cd | -0.63 | -4.85 | 0.00095 | 0.018 | 0.0065 | ENSMUSG00000068798 | protein_coding | Rap1a | RAS-related protein 1a | 109905 | 3 | 105727267 | 105801336 | -1 | Mm.101922 Mm.365961 Mm.458171 |
| 3.3 | 1.6 | 2.99 | 2.48 | DN3ab_Itr | DN3cd | -1.07 | -4.06 | 0.0029 | 0.036 | 0.0065 | ENSMUSG00000008307 | protein_coding | 1700109H08Rik | RIKEN cDNA 1700109H08 gene | 77036 | 5 | 3571566 | 3584356 | 1 | Mm.389069 |
| -0.33 | 2.23 | -0.18 | 4.21 | DN3ab_Itr | DN3cd | 3.47 | 8.55 | 1.40E-05 | 0.0015 | 0.0066 | ENSMUSG00000024353 | protein_coding | Mzb1 | marginal zone B and B1 cell-specific protein 1 | 69816 | 18 | 35647266 | 35649367 | -1 | Mm.27252 |
| 6.16 | 5.18 | 7.64 | 5.92 | DN3ab_Itr | DN3cd | -1.4 | -8.92 | 1.00E-05 | 0.0012 | 0.0066 | ENSMUSG00000026094 | protein_coding | Stk17b | serine/threonine kinase 17b (apoptosis-inducing) | 98267 | 1 | 53755506 | 53785224 | -1 | Mm.25559 |
| 4.94 | 4.48 | 6.43 | 4.88 | DN3ab_Itr | DN3cd | -1.06 | -4.55 | 0.0014 | 0.023 | 0.0066 | ENSMUSG00000040548 | protein_coding | Tex2 | testis expressed gene 2 | 21763 | 11 | 106502147 | 106613423 | -1 | Mm.102407 |
| 5.5 | 5.7 | 5.12 | 5.89 | DN3ab_Itr | DN3cd | 0.45 | 3.62 | 0.0057 | 0.055 | 0.0067 | ENSMUSG00000009647 | protein_coding | Mcu | mitochondrial calcium uniporter | 215999 | 10 | 59446984 | 59616692 | -1 | Mm.296470 |
| 0.27 | 2.42 | -1.51 | 3.29 | DN3ab_Itr | DN3cd | 3.04 | 5.58 | 0.00036 | 0.0097 | 0.0067 | ENSMUSG00000070304 | protein_coding | Scn2b | sodium channel, voltage-gated, type II, beta | 72821 | 9 | 45117782 | 45130070 | 1 | Mm.229373 Mm.491371 |
| 6.65 | 6.8 | 6.14 | 6.8 | DN3ab_Itr | DN3cd | 0.37 | 3.28 | 0.0098 | 0.079 | 0.0067 | ENSMUSG00000023089 | protein_coding | Ndufa5 | NADH:ubiquinone oxidoreductase subunit A5 | 68202 | 6 | 24518666 | 24528013 | -1 | Mm.275780 |
| 6.77 | 6.24 | 8.37 | 7.28 | DN3ab_Itr | DN3cd | -0.85 | -7.09 | 6.10E-05 | 0.0034 | 0.0067 | ENSMUSG00000030403 | protein_coding | Vasp | vasodilator-stimulated phosphoprotein | 22323 | 7 | 19256929 | 19271817 | -1 | Mm.9684 |
| 6.96 | 6.74 | 6.81 | 6.12 | DN3ab_Itr | DN3cd | -0.42 | -4.14 | 0.0026 | 0.033 | 0.0069 | ENSMUSG00000020250 | protein_coding | Txnrd1 | thioredoxin reductase 1 | 50493 | 10 | 82833951 | 82897712 | 1 | Mm.210155 Mm.471159 |
| 6.41 | 6.07 | 7.63 | 6.64 | DN3ab_Itr | DN3cd | -0.69 | -5 | 0.00077 | 0.016 | 0.0069 | ENSMUSG00000024308 | protein_coding | Tapbp | TAP binding protein | 21356 | 17 | 33915899 | 33929292 | 1 | Mm.154457 |
| 4.64 | 4.33 | 5.44 | 3.85 | DN3ab_Itr | DN3cd | -0.9 | -3.29 | 0.0096 | 0.078 | 0.0069 | ENSMUSG00000039048 | protein_coding | Foxred1 | FAD-dependent oxidoreductase domain containing 1 | 235169 | 9 | 35204206 | 35211055 | -1 | Mm.138512 |
| -1.02 | 0.66 | -2.9 | 2.35 | DN3ab_Itr | DN3cd | 3.42 | 4.43 | 0.0017 | 0.026 | 0.007 | ENSMUSG00000022650 | protein_coding | Retnlb | resistin like beta | 57263 | 16 | 48816856 | 48818892 | 1 | Mm.21123 |
| 3.28 | 3.68 | 1.06 | 2.9 | DN3ab_Itr | DN3cd | 0.72 | 2.78 | 0.022 | 0.13 | 0.007 | ENSMUSG00000041912 | protein_coding | Tdrkh | tudor and KH domain containing protein | 72634 | 3 | 94413273 | 94434668 | 1 | Mm.331198 Mm.470795 |
| 4.1 | 4.36 | 3.21 | 4.21 | DN3ab_Itr | DN3cd | 0.54 | 3.36 | 0.0086 | 0.072 | 0.007 | ENSMUSG00000026779 | protein_coding | Mastl | microtubule associated serine/threonine kinase-like | 67121 | 2 | 23115606 | 23156024 | -1 | Mm.253721 |
| 4.78 | 5.33 | 3.34 | 4.83 | DN3ab_Itr | DN3cd | 0.85 | 4.43 | 0.0017 | 0.026 | 0.007 | ENSMUSG00000068101 | protein_coding | Cenpm | centromere protein M | 66570 | 15 | 82233779 | 82244748 | -1 | Mm.23596 |
| 3.68 | 4.3 | 2.63 | 4.49 | DN3ab_Itr | DN3cd | 1.07 | 4.17 | 0.0025 | 0.032 | 0.007 | ENSMUSG00000034881 | protein_coding | Tbxa2r | thromboxane A2 receptor | 21390 | 10 | 81328731 | 81335172 | 1 | Mm.4545 |
| 1.99 | 0.72 | 4.64 | 1.88 | DN3ab_Itr | DN3cd | -2.29 | -7.51 | 4.00E-05 | 0.0027 | 0.0071 | ENSMUSG00000047415 | protein_coding | Gpr68 | G protein-coupled receptor 68 | 238377 | 12 | 100876682 | 100908198 | -1 | Mm.32160 |
| 2.78 | 3.59 | 0.52 | 3.79 | DN3ab_Itr | DN3cd | 1.39 | 3.09 | 0.013 | 0.096 | 0.0071 | ENSMUSG00000028434 | protein_coding | Epb41l4b | erythrocyte membrane protein band 4.1 like 4b | 54357 | 4 | 56991972 | 57143437 | -1 | Mm.28217 Mm.381403 |
| 7.08 | 6.08 | 7.33 | 6.99 | DN3ab_Itr | DN3cd | -0.66 | -4.61 | 0.0013 | 0.022 | 0.0072 | ENSMUSG00000039159 | protein_coding | Ube2h | ubiquitin-conjugating enzyme E2H | 22214 | 6 | 30211289 | 30304539 | -1 | Mm.332967 Mm.391754 |
| 0.23 | 1.44 | -1.16 | 2.62 | DN3ab_Itr | DN3cd | 2.17 | 3.99 | 0.0033 | 0.038 | 0.0072 | ENSMUSG00000066026 | protein_coding | Dhrs3 | dehydrogenase/reductase (SDR family) member 3 | 20148 | 4 | 144892827 | 144928209 | 1 | Mm.14063 Mm.181278 |
| -2.06 | -1.11 | -3.72 | -0.07 | DN3ab_Itr | DN3cd | 2.37 | 3.94 | 0.0035 | 0.04 | 0.0073 | ENSMUSG00000107049 | TEC | Gm43771 | predicted gene 43771 | NA | 5 | 65963851 | 65965428 | 1 | |
| 4.79 | 5.04 | 3.78 | 4.69 | DN3ab_Itr | DN3cd | 0.49 | 3.5 | 0.0068 | 0.062 | 0.0073 | ENSMUSG00000041840 | protein_coding | Haus1 | HAUS augmin-like complex, subunit 1 | 225745 | 18 | 77757567 | 77767780 | -1 | Mm.39293 |
| 8.4 | 8.94 | 7.49 | 8.5 | DN3ab_Itr | DN3cd | 0.72 | 7.21 | 5.40E-05 | 0.0032 | 0.0073 | ENSMUSG00000031004 | protein_coding | Mki67 | antigen identified by monoclonal antibody Ki 67 | 17345 | 7 | 135689784 | 135716361 | -1 | Mm.4078 |
| 4.3 | 4.76 | 3.47 | 4.74 | DN3ab_Itr | DN3cd | 0.77 | 4.43 | 0.0017 | 0.026 | 0.0073 | ENSMUSG00000083822 | processed_pseudogene | Hmgb1-ps5 | high mobility group box 1, pseudogene 5 | NA | 3 | 95530418 | 95530974 | 1 | |
| 7.61 | 7.46 | 8.12 | 7.59 | DN3ab_Itr | DN3cd | -0.34 | -4.07 | 0.0029 | 0.036 | 0.0074 | ENSMUSG00000014867 | protein_coding | Surf4 | surfeit gene 4 | 20932 | 2 | 26920040 | 26933928 | -1 | Mm.300594 |
| 6.61 | 5.3 | 6.83 | 6.45 | DN3ab_Itr | DN3cd | -0.81 | -4.02 | 0.0031 | 0.037 | 0.0074 | ENSMUSG00000095041 | protein_coding | | NA | NA | JH584304.1 | 52190 | 59690 | -1 | |
| 7.16 | 6.96 | 7.54 | 6.85 | DN3ab_Itr | DN3cd | -0.43 | -4.06 | 0.0029 | 0.036 | 0.0074 | ENSMUSG00000025724 | protein_coding | Sec11a | SEC11 homolog A, signal peptidase complex subunit | 56529 | 7 | 80904889 | 80947780 | -1 | Mm.43760 |
| 4.89 | 5.62 | 4.66 | 6.33 | DN3ab_Itr | DN3cd | 1.16 | 5.73 | 3.00E-04 | 0.0087 | 0.0074 | ENSMUSG00000034586 | protein_coding | Hid1 | HID1 domain containing | 217310 | 11 | 115347707 | 115367756 | -1 | Mm.205707 |
| 3.14 | 2.03 | 5.28 | 2.86 | DN3ab_Itr | DN3cd | -1.95 | -7.11 | 6.00E-05 | 0.0034 | 0.0075 | ENSMUSG00000037606 | protein_coding | Osbpl5 | oxysterol binding protein-like 5 | 79196 | 7 | 143688762 | 143756985 | -1 | Mm.21199 |
| 5.79 | 6.23 | 4.63 | 5.78 | DN3ab_Itr | DN3cd | 0.7 | 4.68 | 0.0012 | 0.021 | 0.0075 | ENSMUSG00000027326 | protein_coding | Knl1 | kinetochore scaffold 1 | 76464 | 2 | 119047119 | 119105501 | 1 | Mm.202518 Mm.392877 |
| -2.06 | -0.49 | -3.72 | 0.55 | DN3ab_Itr | DN3cd | 2.98 | 4.96 | 0.00082 | 0.016 | 0.0075 | ENSMUSG00000107271 | lincRNA | BC028471 | cDNA sequence BC028471 | NA | 5 | 149367636 | 149393596 | 1 | |
| 3.9 | 4.28 | 1.69 | 4.01 | DN3ab_Itr | DN3cd | 0.84 | 2.33 | 0.045 | 0.2 | 0.0077 | ENSMUSG00000104413 | antisense | Gm37065 | predicted gene, 37065 | NA | 1 | 171909957 | 171919356 | -1 | |
| 7.62 | 7.76 | 7.29 | 7.81 | DN3ab_Itr | DN3cd | 0.31 | 3.69 | 0.0051 | 0.051 | 0.0077 | ENSMUSG00000034566 | protein_coding | Atp5h | ATP synthase, H+ transporting, mitochondrial F0 complex, subunit D | 71679 | 11 | 115415689 | 115419962 | -1 | Mm.371641 |
| 3.99 | 4.22 | 3.35 | 4.42 | DN3ab_Itr | DN3cd | 0.58 | 3.15 | 0.012 | 0.09 | 0.0077 | ENSMUSG00000044469 | protein_coding | Tnfaip8l1 | tumor necrosis factor, alpha-induced protein 8-like 1 | 66443 | 17 | 56162477 | 56173955 | 1 | Mm.2312 |
| 4.02 | 3.33 | 5.78 | 4.23 | DN3ab_Itr | DN3cd | -1.22 | -6.58 | 0.00011 | 0.0047 | 0.0078 | ENSMUSG00000038497 | protein_coding | Tmco3 | transmembrane and coiled-coil domains 3 | 234076 | 8 | 13288190 | 13322924 | 1 | Mm.219813 |
| 4.31 | 4.63 | 2.99 | 4.26 | DN3ab_Itr | DN3cd | 0.63 | 3.21 | 0.011 | 0.084 | 0.0078 | ENSMUSG00000093726 | sense_overlapping | Gm20667 | predicted gene 20667 | NA | 14 | 67673177 | 67675106 | -1 | |
| 5.65 | 6.02 | 5.04 | 6.04 | DN3ab_Itr | DN3cd | 0.64 | 4.61 | 0.0013 | 0.022 | 0.0078 | ENSMUSG00000081738 | processed_pseudogene | Hmgb1-ps2 | high mobility group box 1, pseudogene 2 | NA | X | 146175942 | 146176577 | 1 | |
| 2.89 | 3.8 | 1.4 | 3.64 | DN3ab_Itr | DN3cd | 1.3 | 4.9 | 0.00088 | 0.017 | 0.0079 | ENSMUSG00000029847 | protein_coding | Slc23a4 | solute carrier family 23 member 4 | 243753 | 6 | 34945278 | 34988404 | -1 | Mm.440221 |
| 5.26 | 5.94 | 4.33 | 5.62 | DN3ab_Itr | DN3cd | 0.91 | 7.04 | 6.50E-05 | 0.0035 | 0.0079 | ENSMUSG00000048922 | protein_coding | Cdca2 | cell division cycle associated 2 | 108912 | 14 | 67676331 | 67715841 | -1 | Mm.33831 |
| 8.31 | 8.76 | 7.43 | 8.3 | DN3ab_Itr | DN3cd | 0.62 | 6.89 | 7.70E-05 | 0.0038 | 0.0079 | ENSMUSG00000024660 | protein_coding | Incenp | inner centromere protein | 16319 | 19 | 9872297 | 9899533 | -1 | Mm.29755 |
| 5.39 | 6.19 | 3.51 | 6.08 | DN3ab_Itr | DN3cd | 1.34 | 3.79 | 0.0044 | 0.046 | 0.008 | ENSMUSG00000038146 | protein_coding | Notch3 | notch 3 | 18131 | 17 | 32120820 | 32166880 | -1 | Mm.439741 |
| 5.32 | 5.52 | 4.4 | 5.44 | DN3ab_Itr | DN3cd | 0.54 | 2.98 | 0.016 | 0.11 | 0.008 | ENSMUSG00000082809 | processed_pseudogene | Gm14150 | predicted gene 14150 | NA | 2 | 151458956 | 151460309 | 1 | |
| 4.82 | 3.79 | 6.42 | 4.45 | DN3ab_Itr | DN3cd | -1.57 | -7.8 | 2.90E-05 | 0.0022 | 0.0081 | ENSMUSG00000043252 | protein_coding | Tmem64 | transmembrane protein 64 | 100201 | 4 | 15265831 | 15286753 | 1 | Mm.394111 |
| 4.22 | 4.55 | 2.91 | 4.13 | DN3ab_Itr | DN3cd | 0.61 | 3.27 | 0.0098 | 0.079 | 0.0081 | ENSMUSG00000032965 | protein_coding | Ift57 | intraflagellar transport 57 | 73916 | 16 | 49699233 | 49765126 | 1 | Mm.241276 Mm.404371 |
| 7.59 | 7.22 | 8 | 7.16 | DN3ab_Itr | DN3cd | -0.59 | -5.8 | 0.00027 | 0.0083 | 0.0082 | ENSMUSG00000082319 | processed_pseudogene | Gm8822 | predicted gene 8822 | NA | X | 81820384 | 81822730 | -1 | |
| 6.22 | 6.75 | 5.37 | 6.36 | DN3ab_Itr | DN3cd | 0.71 | 7.11 | 6.00E-05 | 0.0034 | 0.0082 | ENSMUSG00000024601 | protein_coding | Isoc1 | isochorismatase domain containing 1 | 66307 | 18 | 58659482 | 58679570 | 1 | Mm.182574 |
| 3.92 | 2.78 | 8.18 | 5.09 | DN3ab_Itr | DN3cd | -2.66 | -7.55 | 3.80E-05 | 0.0026 | 0.0083 | ENSMUSG00000026009 | protein_coding | Icos | inducible T cell co-stimulator | 54167 | 1 | 60977927 | 61000320 | 1 | Mm.42044 |
| 5.4 | 5.74 | 5.19 | 6.21 | DN3ab_Itr | DN3cd | 0.66 | 4.37 | 0.0019 | 0.027 | 0.0084 | ENSMUSG00000039828 | protein_coding | Wdr70 | WD repeat domain 70 | 545085 | 15 | 7873055 | 8099209 | -1 | Mm.321312 |
| 8.76 | 9.34 | 6.93 | 8.5 | DN3ab_Itr | DN3cd | 0.92 | 4.48 | 0.0016 | 0.025 | 0.0084 | ENSMUSG00000055612 | protein_coding | Cdca7 | cell division cycle associated 7 | 66953 | 2 | 72476159 | 72486893 | 1 | Mm.270676 |
| 10.61 | 11.07 | 9.8 | 10.62 | DN3ab_Itr | DN3cd | 0.61 | 8.07 | 2.20E-05 | 0.0019 | 0.0085 | ENSMUSG00000020914 | protein_coding | Top2a | topoisomerase (DNA) II alpha | 21973 | 11 | 98992943 | 99024189 | -1 | Mm.4237 |
| 4.83 | 3.79 | 5.23 | 4.87 | DN3ab_Itr | DN3cd | -0.66 | -4.36 | 0.0019 | 0.027 | 0.0086 | ENSMUSG00000043505 | protein_coding | Gimap5 | GTPase, IMAP family member 5 | 317757 | 6 | 48746197 | 48754210 | 1 | Mm.455917 |
| -2.06 | -3.92 | 1.45 | -4.02 | DN3ab_Itr | DN3cd | -3.98 | -5.06 | 0.00071 | 0.015 | 0.0086 | ENSMUSG00000021640 | protein_coding | Naip1 | NLR family, apoptosis inhibitory protein 1 | 17940 | 13 | 100407764 | 100452869 | -1 | Mm.6898 |
| 6.9 | 6.33 | 7.18 | 5.92 | DN3ab_Itr | DN3cd | -0.88 | -5.84 | 0.00026 | 0.0081 | 0.0087 | ENSMUSG00000090460 | processed_pseudogene | Gm17511 | predicted gene, 17511 | NA | 7 | 127286044 | 127286385 | -1 | |

| | | | | | | | | | | | | | | | | | | | | |
|---|---|---|---|---|---|---|---|---|---|---|---|---|---|---|---|---|---|---|---|---|
| 4.69 | 4.45 | 6.03 | 5.05 | DN3ab_Itr | DN3cd | -0.65 | -4.05 | 0.003 | 0.036 | 0.0087 | ENSMUSG00000005873 | protein_coding | Reep5 | receptor accessory protein 5 | 13476 | 18 | 34344889 | 34373415 | -1 | Mm.280075 |
| 4.04 | 3.85 | 4.87 | 4.04 | DN3ab_Itr | DN3cd | -0.53 | -3.66 | 0.0053 | 0.053 | 0.0087 | ENSMUSG00000024955 | protein_coding | Esrra | estrogen related receptor, alpha | 26379 | 19 | 6910977 | 6921813 | -1 | Mm.386776 |
| 6.02 | 6.87 | 5.49 | 5.72 | DN3ab_Itr | DN3cd | 0.61 | 4.53 | 0.0015 | 0.024 | 0.0087 | ENSMUSG00000028633 | protein_coding | Ctps | cytidine 5'-triphosphate synthase | 51797 | 4 | 120539868 | 120570276 | -1 | Mm.1815 |
| 8.04 | 8.36 | 7.8 | 8.42 | DN3ab_Itr | DN3cd | 0.46 | 6.88 | 7.70E-05 | 0.0038 | 0.0087 | ENSMUSG00000045969 | protein_coding | Ing1 | inhibitor of growth family, member 1 | 26356 | 8 | 11555571 | 11563251 | 1 | Mm.25709 |
| 4.53 | 4.26 | 5.07 | 4.08 | DN3ab_Itr | DN3cd | -0.62 | -3.85 | 0.004 | 0.044 | 0.0088 | ENSMUSG00000071647 | protein_coding | Eml3 | echinoderm microtubule associated protein like 3 | 225898 | 19 | 8929552 | 8941582 | 1 | Mm.31008 |
| 8.82 | 9.1 | 8.31 | 8.96 | DN3ab_Itr | DN3cd | 0.44 | 5.3 | 0.00052 | 0.012 | 0.0089 | ENSMUSG00000041133 | protein_coding | Smc1a | structural maintenance of chromosomes 1A | 24061 | X | 152016428 | 152062694 | 1 | Mm.482095 |
| 1.62 | 2.17 | 0.27 | 2.55 | DN3ab_Itr | DN3cd | 1.14 | 3.1 | 0.013 | 0.095 | 0.0089 | ENSMUSG00000102374 | TEC | Gm38387 | predicted gene, 38387 | NA | 1 | 66769290 | 66771866 | -1 | |
| 3.83 | 3.34 | 4.87 | 3.43 | DN3ab_Itr | DN3cd | -0.99 | -4.71 | 0.0011 | 0.02 | 0.0089 | ENSMUSG00000034403 | protein_coding | Pja1 | praja ring finger ubiquitin ligase 1 | 18744 | X | 99465734 | 99471273 | -1 | Mm.8211 |
| -0.56 | 0.98 | -3.72 | 1.24 | DN3ab_Itr | DN3cd | 2.97 | 3.99 | 0.0033 | 0.038 | 0.009 | ENSMUSG00000016179 | protein_coding | Camk1g | calcium/calmodulin-dependent protein kinase I gamma | 215303 | 1 | 193346346 | 193370298 | -1 | Mm.40329 |
| 5.19 | 6.07 | 4.11 | 6.23 | DN3ab_Itr | DN3cd | 1.34 | 5.11 | 0.00067 | 0.014 | 0.009 | ENSMUSG00000020900 | protein_coding | Myh10 | myosin, heavy polypeptide 10, non-muscle | 77579 | 11 | 68691559 | 68816632 | 1 | Mm.218233 |
| 2.23 | 2.59 | 0.65 | 2.42 | DN3ab_Itr | DN3cd | 0.79 | 2.68 | 0.025 | 0.14 | 0.0091 | ENSMUSG00000056832 | protein_coding | Ttc26 | tetratricopeptide repeat domain 26 | 264134 | 6 | 38381469 | 38427647 | 1 | Mm.253446 |
| 7.24 | 7.44 | 6.54 | 7.37 | DN3ab_Itr | DN3cd | 0.47 | 3.41 | 0.008 | 0.069 | 0.0093 | ENSMUSG00000005370 | protein_coding | Msh6 | mutS homolog 6 | 17688 | 17 | 87975062 | 87990892 | 1 | Mm.18210 |
| 7.28 | 7.41 | 6.71 | 7.28 | DN3ab_Itr | DN3cd | 0.32 | 3.28 | 0.0097 | 0.079 | 0.0093 | ENSMUSG00000075266 | protein_coding | Cenpw | centromere protein W | 66311 | 10 | 30194569 | 30200560 | -1 | Mm.251779 |
| 4.83 | 5.39 | 3.76 | 5.35 | DN3ab_Itr | DN3cd | 0.94 | 4.27 | 0.0022 | 0.03 | 0.0093 | ENSMUSG00000026134 | protein_coding | Prim2 | DNA primase, p58 subunit | 19076 | 1 | 33453810 | 33669795 | -1 | Mm.27705 |
| 6.87 | 6.29 | 7.97 | 6.81 | DN3ab_Itr | DN3cd | -0.89 | -6.92 | 7.40E-05 | 0.0038 | 0.0094 | ENSMUSG00000031304 | protein_coding | Il2rg | interleukin 2 receptor, gamma chain | 16186 | X | 101264378 | 101268255 | -1 | Mm.2923 |
| 4.74 | 4.52 | 5.13 | 4.34 | DN3ab_Itr | DN3cd | -0.49 | -3.82 | 0.0042 | 0.045 | 0.0094 | ENSMUSG00000017615 | protein_coding | Tnfaip1 | tumor necrosis factor, alpha-induced protein 1 (endothelial) | 21927 | 11 | 78522850 | 78536332 | -1 | Mm.386774 |
| 5.29 | 2.66 | 5.17 | 3.61 | DN3ab_Itr | DN3cd | -2.02 | -8.53 | 1.50E-05 | 0.0015 | 0.0096 | ENSMUSG00000038807 | protein_coding | Rap1gap2 | RAP1 GTPase activating protein 2 | 380711 | 11 | 74383356 | 74610915 | -1 | Mm.425690 |
| 7.86 | 8.27 | 7.9 | 8.01 | DN3ab_Itr | DN3cd | 0.27 | 3.92 | 0.0036 | 0.041 | 0.0096 | ENSMUSG00000022391 | protein_coding | Rangap1 | RAN GTPase activating protein 1 | 19387 | 15 | 81704248 | 81745530 | -1 | Mm.270975 Mm.475271 |
| 3.15 | 2.51 | 5.26 | 3.56 | DN3ab_Itr | DN3cd | -1.34 | -5.95 | 0.00023 | 0.0075 | 0.0096 | ENSMUSG00000030156 | protein_coding | Cd69 | CD69 antigen | 12515 | 6 | 129267325 | 129275436 | -1 | Mm.74745 |
| 11.06 | 11.31 | 11.2 | 11.63 | DN3ab_Itr | DN3cd | 0.33 | 8.01 | 2.40E-05 | 0.002 | 0.0098 | ENSMUSG00000064351 | protein_coding | mt-Co1 | mitochondrially encoded cytochrome c oxidase I | 17708 | MT | 5328 | 6872 | 1 | |
| 8.85 | 9.27 | 7.47 | 8.51 | DN3ab_Itr | DN3cd | 0.65 | 4.85 | 0.00094 | 0.018 | 0.0098 | ENSMUSG00000038252 | protein_coding | Ncapd2 | non-SMC condensin I complex, subunit D2 | 68298 | 6 | 125168007 | 125191701 | -1 | Mm.257590 |
| 2.2 | 3.49 | -0.11 | 3.72 | DN3ab_Itr | DN3cd | 1.89 | 3.97 | 0.0034 | 0.039 | 0.0099 | ENSMUSG00000020723 | protein_coding | Cacng4 | calcium channel, voltage-dependent, gamma subunit 4 | 54377 | 11 | 107732357 | 107794557 | -1 | Mm.103724 |
| 5.35 | 3.69 | 5.94 | 5.05 | DN3ab_Itr | DN3cd | -1.2 | -6.98 | 6.90E-05 | 0.0036 | 0.01 | ENSMUSG00000037295 | protein_coding | Ldlrap1 | low density lipoprotein receptor adaptor protein 1 | 100017 | 4 | 134741554 | 134768024 | -1 | Mm.27486 Mm.482148 |
| 8.41 | 7.25 | 8.37 | 7.88 | DN3ab_Itr | DN3cd | -0.83 | -5.61 | 0.00035 | 0.0095 | 0.01 | ENSMUSG00000018474 | protein_coding | Chd3 | chromodomain helicase DNA binding protein 3 | 216848 | 11 | 69343273 | 69369406 | -1 | Mm.178246 |
| 5.32 | 4.43 | 5.17 | 4.93 | DN3ab_Itr | DN3cd | -0.58 | -3.95 | 0.0034 | 0.039 | 0.01 | ENSMUSG00000006127 | protein_coding | Inpp5k | inositol polyphosphate 5-phosphatase K | 19062 | 11 | 75630988 | 75648871 | 1 | Mm.1458 |
| 5.47 | 4.55 | 6.15 | 5.87 | DN3ab_Itr | DN3cd | -0.56 | -3.89 | 0.0037 | 0.042 | 0.01 | ENSMUSG00000033446 | protein_coding | Lpar6 | lysophosphatidic acid receptor 6 | 67168 | 14 | 73237895 | 73243294 | 1 | Mm.390681 |
| 6.6 | 5.93 | 8.11 | 6.87 | DN3ab_Itr | DN3cd | -0.99 | -7.93 | 2.60E-05 | 0.0021 | 0.01 | ENSMUSG00000028161 | protein_coding | Ppp3ca | protein phosphatase 3, catalytic subunit, alpha isoform | 19055 | 3 | 136670124 | 136937727 | 1 | Mm.331389 Mm.486983 |
| 5.11 | 5.83 | 4.25 | 5.8 | DN3ab_Itr | DN3cd | 1.04 | 5.81 | 0.00027 | 0.0083 | 0.01 | ENSMUSG00000025035 | protein_coding | Arl3 | ADP-ribosylation factor-like 3 | 56350 | 19 | 46531109 | 46573085 | -1 | Mm.22085 |
| 5.18 | 5.36 | 4.45 | 5.22 | DN3ab_Itr | DN3cd | 0.42 | 3.22 | 0.011 | 0.084 | 0.01 | ENSMUSG00000031402 | protein_coding | Mpp1 | membrane protein, palmitoylated | 17524 | X | 75109733 | 75131016 | -1 | Mm.391267 |
| 6.66 | 6.23 | 6.98 | 6.02 | DN3ab_Itr | DN3cd | -0.68 | -5.66 | 0.00033 | 0.0092 | 0.01 | ENSMUSG00000062382 | protein_coding | Ftl1-ps1 | ferritin light polypeptide 1, pseudogene 1 | 100862446 | 13 | 74399694 | 74407346 | 1 | |
| -2.43 | -0.48 | -3.72 | 0.59 | DN3ab_Itr | DN3cd | 3.2 | 5.88 | 0.00025 | 0.0079 | 0.01 | ENSMUSG00000082700 | processed_pseudogene | Gm13792 | predicted gene 13792 | NA | 2 | 92872887 | 92873309 | -1 | |
| 6.18 | 5.94 | 6.97 | 6.29 | DN3ab_Itr | DN3cd | -0.47 | -4.65 | 0.0012 | 0.021 | 0.01 | ENSMUSG00000041920 | protein_coding | Slc16a6 | solute carrier family 16 (monocarboxylic acid transporters), member 6 | 104681 | 11 | 109450855 | 109473598 | -1 | Mm.265874 |
| 5.78 | 5.01 | 6.75 | 5.41 | DN3ab_Itr | DN3cd | -1.07 | -8.23 | 1.90E-05 | 0.0018 | 0.01 | ENSMUSG00000044026 | protein_coding | Slc35g1 | solute carrier family 35, member G1 | 240660 | 19 | 38395980 | 38405607 | 1 | Mm.262085 |
| 1.66 | 3.08 | -0.97 | 2.95 | DN3ab_Itr | DN3cd | 1.95 | 4.25 | 0.0022 | 0.03 | 0.011 | ENSMUSG00000079550 | protein_coding | Mpp4 | membrane protein, palmitoylated 4 (MAGUK p55 subfamily member 4) | 227157 | 1 | 59120935 | 59163389 | -1 | Mm.474049 |
| 8.63 | 8.74 | 8.32 | 8.69 | DN3ab_Itr | DN3cd | 0.22 | 3.62 | 0.0057 | 0.055 | 0.011 | ENSMUSG00000017221 | protein_coding | Psmd3 | proteasome (prosome, macropain) 26S subunit, non-ATPase, 3 | 22123 | 11 | 98682554 | 98695979 | 1 | Mm.12194 |
| 4.16 | 3.68 | 4.78 | 3.48 | DN3ab_Itr | DN3cd | -0.87 | -4.62 | 0.0013 | 0.022 | 0.011 | ENSMUSG00000041650 | protein_coding | Pcca | propionyl-Coenzyme A carboxylase, alpha polypeptide | 110821 | 14 | 122534324 | 122891100 | 1 | Mm.23876 |
| 9.55 | 9.82 | 8.57 | 9.34 | DN3ab_Itr | DN3cd | 0.46 | 4.24 | 0.0023 | 0.031 | 0.011 | ENSMUSG00000019961 | protein_coding | Tmpo | thymopoietin | 21917 | 10 | 91147571 | 91181315 | -1 | Mm.159684 Mm.439157 |
| 3.16 | 3.52 | 1.62 | 3.13 | DN3ab_Itr | DN3cd | 0.71 | 2.96 | 0.016 | 0.11 | 0.011 | ENSMUSG00000031629 | protein_coding | Cenpu | centromere protein U | 71876 | 8 | 46552028 | 46580007 | 1 | Mm.22108 Mm.217385 |
| 6.65 | 5.87 | 8.79 | 7.14 | DN3ab_Itr | DN3cd | -1.31 | -6.95 | 7.20E-05 | 0.0037 | 0.011 | ENSMUSG00000068227 | protein_coding | Il2rb | interleukin 2 receptor, beta chain | 16185 | 15 | 78479256 | 78495271 | -1 | Mm.35287 Mm.384038 |
| 5.95 | 6.77 | 4.4 | 6.62 | DN3ab_Itr | DN3cd | 1.31 | 4.43 | 0.0017 | 0.026 | 0.012 | ENSMUSG00000020167 | protein_coding | Tcf3 | transcription factor 3 | 21423 | 10 | 80409514 | 80433647 | -1 | Mm.3406 |
| 3.4 | 2.51 | 5.23 | 3.21 | DN3ab_Itr | DN3cd | -1.58 | -6.39 | 0.00014 | 0.0055 | 0.012 | ENSMUSG00000032579 | protein_coding | Hemk1 | HemK methyltransferase family member 1 | 69536 | 9 | 107327084 | 107338350 | -1 | Mm.259467 |
| 6.87 | 7.29 | 7 | 7.88 | DN3ab_Itr | DN3cd | 0.65 | 6.24 | 0.00016 | 0.006 | 0.012 | ENSMUSG00000032316 | protein_coding | Clk3 | CDC-like kinase 3 | 102414 | 9 | 57750624 | 57765860 | -1 | Mm.25720 |
| 5.92 | 6.3 | 5.05 | 6.22 | DN3ab_Itr | DN3cd | 0.7 | 4.03 | 0.0031 | 0.037 | 0.012 | ENSMUSG00000037242 | protein_coding | Clic4 | chloride intracellular channel 4 (mitochondrial) | 29876 | 4 | 135213969 | 135272814 | -1 | Mm.257765 |
| 5.48 | 5.12 | 6.13 | 5.28 | DN3ab_Itr | DN3cd | -0.61 | -5.36 | 0.00048 | 0.012 | 0.012 | ENSMUSG00000036948 | protein_coding | BC037034 | cDNA sequence BC037034 | 231807 | 5 | 138259656 | 138264033 | -1 | Mm.164493 |
| 11.12 | 11.38 | 9.93 | 10.71 | DN3ab_Itr | DN3cd | 0.45 | 3.99 | 0.0032 | 0.038 | 0.012 | ENSMUSG00000037894 | protein_coding | H2afz | H2A histone family, member Z | 51788 | 3 | 137864487 | 137866922 | 1 | Mm.117541 Mm.372513 Mm.465508 |
| 0.14 | 1.59 | -1.76 | 2.59 | DN3ab_Itr | DN3cd | 2.5 | 3.97 | 0.0034 | 0.039 | 0.012 | ENSMUSG00000047904 | protein_coding | Sstr2 | somatostatin receptor 2 | 20606 | 11 | 113619342 | 113626028 | 1 | |
| 8.06 | 8.35 | 6.95 | 7.94 | DN3ab_Itr | DN3cd | 0.56 | 3.71 | 0.0049 | 0.05 | 0.012 | ENSMUSG00000096006 | processed_pseudogene | Gm21596 | predicted gene, 21596 | NA | 2 | 113919247 | 113920562 | -1 | |
| 7.02 | 7.43 | 6.58 | 7.35 | DN3ab_Itr | DN3cd | 0.57 | 6.86 | 8.00E-05 | 0.0039 | 0.012 | ENSMUSG00000029191 | protein_coding | Rfc1 | replication factor C (activator 1) 1 | 19687 | 5 | 65261850 | 65335670 | -1 | Mm.148877 |
| 4.56 | 4.93 | 3.67 | 4.69 | DN3ab_Itr | DN3cd | 0.61 | 4.21 | 0.0023 | 0.031 | 0.012 | ENSMUSG00000037243 | protein_coding | Zfp692 | zinc finger protein 692 | 103836 | 11 | 58307069 | 58314627 | 1 | Mm.41440 |
| 3.51 | 2.96 | 4.64 | 3.34 | DN3ab_Itr | DN3cd | -0.97 | -5.6 | 0.00035 | 0.0096 | 0.012 | ENSMUSG00000028221 | protein_coding | Pip4p2 | phosphatidylinositol-4,5-bisphosphate 4-phosphatase 2 | 72519 | 4 | 14864076 | 14915176 | 1 | Mm.279158 |
| 5.6 | 5.95 | 5.7 | 6.54 | DN3ab_Itr | DN3cd | 0.59 | 5.33 | 5.00E-04 | 0.012 | 0.012 | ENSMUSG00000018381 | protein_coding | Abi3 | ABI gene family, member 3 | 66610 | 11 | 95830074 | 95842476 | -1 | Mm.485545 |
| -1.63 | -0.46 | -3.72 | 0.23 | DN3ab_Itr | DN3cd | 2.57 | 4.02 | 0.0031 | 0.037 | 0.012 | ENSMUSG00000036356 | protein_coding | Csgalnact1 | chondroitin sulfate N-acetylgalactosaminyltransferase 1 | 234356 | 8 | 68356781 | 68735146 | -1 | Mm.334569 Mm.489712 |
| -1.02 | -3.12 | 4.71 | -0.18 | DN3ab_Itr | DN3cd | -4.16 | -7.41 | 4.40E-05 | 0.0028 | 0.012 | ENSMUSG00000032420 | protein_coding | Nt5e | 5' nucleotidase, ecto | 23959 | 9 | 88327197 | 88372092 | 1 | Mm.244235 |
| 5.65 | 4.88 | 6.05 | 5.79 | DN3ab_Itr | DN3cd | -0.49 | -4.24 | 0.0023 | 0.031 | 0.012 | ENSMUSG00000109901 | protein_coding | Chmp1b | charged multivesicular body protein 1B | 67064 | 18 | 67203531 | 67207880 | 1 | Mm.73777 |
| 6.52 | 7.58 | 6.31 | 6.69 | DN3ab_Itr | DN3cd | 0.78 | 5.06 | 0.00071 | 0.015 | 0.013 | ENSMUSG00000021000 | protein_coding | Mia2 | melanoma inhibitory activity 2 | 338320 | 12 | 59095799 | 59191583 | 1 | Mm.244118 Mm.213215 Mm.438038 |
| 6.79 | 6.98 | 6.16 | 6.84 | DN3ab_Itr | DN3cd | 0.4 | 3.54 | 0.0065 | 0.06 | 0.013 | ENSMUSG00000028394 | protein_coding | Pole3 | polymerase (DNA directed), epsilon 3 (p17 subunit) | 59001 | 4 | 62522649 | 62525068 | -1 | Mm.27444 |
| 3.67 | 4.35 | 2.5 | 4.02 | DN3ab_Itr | DN3cd | 0.96 | 5.25 | 0.00055 | 0.013 | 0.013 | ENSMUSG00000034303 | protein_coding | Ccdc15 | coiled-coil domain containing 15 | 245902 | 9 | 37275835 | 37348432 | -1 | Mm.390150 |
| 3.77 | 2.81 | 5.84 | 4.19 | DN3ab_Itr | DN3cd | -1.42 | -9.46 | 6.30E-06 | 0.00087 | 0.013 | ENSMUSG00000042842 | protein_coding | Serpinb6b | serine (or cysteine) peptidase inhibitor, clade B, member 6b | 20708 | 13 | 32965209 | 32979067 | 1 | Mm.36526 |
| 8.37 | 7.3 | 9.22 | 6.99 | DN3ab_Itr | DN3cd | -1.63 | -6.28 | 0.00015 | 0.0059 | 0.013 | ENSMUSG00000028266 | protein_coding | Lmo4 | LIM domain only 4 | 16911 | 3 | 144188530 | 144205220 | -1 | Mm.29187 |
| 3.17 | 2.32 | 5.72 | 3.53 | DN3ab_Itr | DN3cd | -1.77 | -6.25 | 0.00016 | 0.006 | 0.013 | ENSMUSG00000038304 | protein_coding | Cd160 | CD160 antigen | 54215 | 3 | 96798763 | 96829351 | -1 | Mm.34693 |
| 4.03 | 3.02 | 3.75 | 3.43 | DN3ab_Itr | DN3cd | -0.68 | -4.2 | 0.0024 | 0.031 | 0.013 | ENSMUSG00000040929 | protein_coding | Rfx3 | regulatory factor X, 3 (influences HLA class II expression) | 19726 | 19 | 27761721 | 28011166 | -1 | Mm.313321 |
| 8.27 | 7.26 | 8.47 | 7.82 | DN3ab_Itr | DN3cd | -0.83 | -9.9 | 4.30E-06 | 0.00075 | 0.013 | ENSMUSG00000057230 | protein_coding | Aak1 | AP2 associated kinase 1 | 269774 | 6 | 86849517 | 87003223 | 1 | Mm.221038 |
| 7.56 | 7.36 | 7.74 | 7.2 | DN3ab_Itr | DN3cd | -0.36 | -4.45 | 0.0017 | 0.025 | 0.013 | ENSMUSG00000031628 | protein_coding | Casp3 | caspase 3 | 12367 | 8 | 46617291 | 46639689 | 1 | Mm.34405 |
| 8.47 | 9.53 | 7.63 | 9.45 | DN3ab_Itr | DN3cd | 1.37 | 8.14 | 2.10E-05 | 0.0019 | 0.013 | ENSMUSG00000033295 | protein_coding | Ptprf | protein tyrosine phosphatase, receptor type, F | 19268 | 4 | 118208213 | 118291405 | -1 | Mm.29855 |
| 5.29 | 4.74 | 6.03 | 4.86 | DN3ab_Itr | DN3cd | -0.86 | -6.03 | 0.00021 | 0.007 | 0.013 | ENSMUSG00000035329 | protein_coding | Fbxo33 | F-box protein 33 | 70611 | 12 | 59200655 | 59219725 | -1 | Mm.311026 |
| 0.37 | 2.71 | 2.27 | 3.16 | DN3ab_Itr | DN3cd | 1.38 | 4.37 | 0.0019 | 0.027 | 0.013 | ENSMUSG00000045394 | protein_coding | Epcam | epithelial cell adhesion molecule | 17075 | 17 | 87635979 | 87651129 | 1 | Mm.4259 |
| 6.33 | 6.66 | 5.94 | 6.87 | DN3ab_Itr | DN3cd | 0.6 | 4.48 | 0.0016 | 0.025 | 0.013 | ENSMUSG00000038459 | protein_coding | Abhd17c | abhydrolase domain containing 17C | 70178 | 7 | 84109356 | 84151893 | -1 | Mm.23914 |
| 0.63 | 2.8 | -1.51 | 2.91 | DN3ab_Itr | DN3cd | 2.82 | 5.99 | 0.00022 | 0.0073 | 0.014 | ENSMUSG00000064147 | protein_coding | Rab44 | RAB44, member RAS oncogene family | 442827 | 17 | 29114145 | 29148980 | 1 | Mm.440341 |
| 9.21 | 9.48 | 8.56 | 9.14 | DN3ab_Itr | DN3cd | 0.4 | 5.65 | 0.00033 | 0.0092 | 0.014 | ENSMUSG00000002477 | protein_coding | Snrpd1 | small nuclear ribonucleoprotein D1 | 20641 | 18 | 10617775 | 10642079 | 1 | Mm.603 |
| 3.14 | 3.9 | 1.78 | 4.48 | DN3ab_Itr | DN3cd | 1.4 | 3.38 | 0.0083 | 0.071 | 0.014 | ENSMUSG00000022861 | protein_coding | Dgkg | diacylglycerol kinase, gamma | 110197 | 16 | 22468461 | 22657221 | -1 | Mm.194986 Mm.448158 |
| 7.06 | 6.35 | 8.04 | 6.84 | DN3ab_Itr | DN3cd | -0.97 | -8.62 | 1.30E-05 | 0.0014 | 0.014 | ENSMUSG00000039168 | protein_coding | Dap | death-associated protein | 223453 | 15 | 31224314 | 31274341 | 1 | Mm.222867 |
| 7.86 | 8.19 | 6.43 | 7.32 | DN3ab_Itr | DN3cd | 0.54 | 4.32 | 0.002 | 0.028 | 0.014 | ENSMUSG00000027715 | protein_coding | Ccna2 | cyclin A2 | 12428 | 3 | 36564865 | 36572150 | -1 | Mm.4189 |
| 4.4 | 3.99 | 5.78 | 4.66 | DN3ab_Itr | DN3cd | -0.82 | -5.05 | 0.00072 | 0.015 | 0.014 | ENSMUSG00000059456 | protein_coding | Ptk2b | PTK2 protein tyrosine kinase 2 beta | 19229 | 14 | 66153257 | 66281052 | -1 | Mm.21613 |
| 6.27 | 6.6 | 5.57 | 6.32 | DN3ab_Itr | DN3cd | 0.5 | 5.22 | 0.00057 | 0.013 | 0.014 | ENSMUSG00000030980 | protein_coding | Knop1 | lysine rich nucleolar protein 1 | 66356 | 7 | 118842222 | 118856254 | -1 | Mm.482278 |
| 4.15 | 4.5 | 3.53 | 4.53 | DN3ab_Itr | DN3cd | 0.61 | 4.04 | 0.003 | 0.037 | 0.014 | ENSMUSG00000043872 | protein_coding | Zmym1 | zinc finger, MYM domain containing 1 | 68310 | 4 | 127047094 | 127061152 | -1 | Mm.273806 |
| 6.16 | 4.67 | 6.37 | 5.32 | DN3ab_Itr | DN3cd | -1.26 | -12.26 | 7.30E-07 | 0.00033 | 0.014 | ENSMUSG00000086236 | antisense | 5830418P13Rik | RIKEN cDNA 5830418P13 gene | NA | 9 | 103423578 | 103461260 | 1 | |
| 6.52 | 5.78 | 6.6 | 6.33 | DN3ab_Itr | DN3cd | -0.51 | -4.62 | 0.0013 | 0.022 | 0.014 | ENSMUSG00000045098 | protein_coding | Kmt5b | lysine methyltransferase 5B | 225888 | 19 | 3767421 | 3818303 | 1 | Mm.278578 |
| 2.37 | 2.81 | 0.98 | 2.65 | DN3ab_Itr | DN3cd | 0.82 | 3.04 | 0.014 | 0.1 | 0.014 | ENSMUSG00000081943 | processed_pseudogene | Gm9078 | predicted gene 9078 | NA | X | 101062260 | 101063058 | 1 | |
| 9.46 | 9.2 | 9.9 | 9.34 | DN3ab_Itr | DN3cd | -0.41 | -5.75 | 0.00029 | 0.0086 | 0.014 | ENSMUSG00000028745 | protein_coding | Capzb | capping protein (actin filament) muscle Z-line, beta | 12345 | 4 | 139192899 | 139291818 | 1 | Mm.2945 |
| 7.16 | 6.92 | 7.53 | 6.97 | DN3ab_Itr | DN3cd | -0.39 | -5.19 | 6.00E-04 | 0.013 | 0.014 | ENSMUSG00000015714 | protein_coding | Cers2 | ceramide synthase 2 | 76893 | 3 | 95314791 | 95323599 | 1 | Mm.181009 |
| 5.43 | 5.21 | 5.9 | 5.21 | DN3ab_Itr | DN3cd | -0.44 | -4.05 | 0.003 | 0.036 | 0.014 | ENSMUSG00000033352 | protein_coding | Map2k4 | mitogen-activated protein kinase kinase 4 | 26398 | 11 | 65688243 | 65788297 | -1 | Mm.412922 |
| 6.85 | 6.54 | 7.92 | 7.15 | DN3ab_Itr | DN3cd | -0.56 | -5.21 | 0.00058 | 0.013 | 0.015 | ENSMUSG00000024516 | protein_coding | Sec11c | SEC11 homolog C, signal peptidase complex subunit | 66286 | 18 | 65800549 | 65817665 | 1 | Mm.27800 |
| 6.65 | 7.12 | 5.78 | 6.68 | DN3ab_Itr | DN3cd | 0.65 | 6.61 | 1.00E-04 | 0.0046 | 0.015 | ENSMUSG00000028702 | protein_coding | Rad54l | RAD54 like (S. cerevisiae) | 19366 | 4 | 116094264 | 116123690 | -1 | Mm.3655 |
| 7.34 | 6.97 | 8.19 | 7.27 | DN3ab_Itr | DN3cd | -0.65 | -5.1 | 0.00067 | 0.014 | 0.015 | ENSMUSG00000020134 | protein_coding | Peli1 | pellino 1 | 67245 | 11 | 21091291 | 21150323 | 1 | Mm.28957 |
| 5.54 | 5.81 | 4.83 | 5.75 | DN3ab_Itr | DN3cd | 0.54 | 3.63 | 0.0056 | 0.054 | 0.015 | ENSMUSG00000069135 | protein_coding | Fgfr1op | Fgfr1 oncogene partner | 75296 | 17 | 8165501 | 8196804 | 1 | Mm.227250 |
| 2.13 | 2.95 | 0.46 | 3.25 | DN3ab_Itr | DN3cd | 1.38 | 3.35 | 0.0087 | 0.073 | 0.015 | ENSMUSG00000039137 | protein_coding | Whrn | whirlin | 73750 | 4 | 63414910 | 63495991 | -1 | Mm.300397 |
| 7.74 | 8.11 | 7.16 | 7.95 | DN3ab_Itr | DN3cd | 0.55 | 5.86 | 0.00025 | 0.0079 | 0.015 | ENSMUSG00000031807 | protein_coding | Pgls | 6-phosphogluconolactonase | 66171 | 8 | 71592176 | 71601092 | 1 | Mm.282284 Mm.402679 |
| 5.84 | 6.31 | 4.95 | 5.97 | DN3ab_Itr | DN3cd | 0.69 | 5.41 | 0.00045 | 0.011 | 0.015 | ENSMUSG00000021707 | protein_coding | Dhfr | dihydrofolate reductase | 13361 | 13 | 92354726 | 92389053 | 1 | Mm.23695 |
| 3.12 | 1.6 | 4.04 | 3.42 | DN3ab_Itr | DN3cd | -0.91 | -4.54 | 0.0015 | 0.024 | 0.016 | ENSMUSG00000031539 | protein_coding | Ap3m2 | adaptor-related protein complex 3, mu 2 subunit | 64933 | 8 | 22787354 | 22805622 | -1 | Mm.390023 |
| -0.84 | 1.36 | -2.95 | 1.44 | DN3ab_Itr | DN3cd | 3.14 | 6.09 | 0.00019 | 0.0067 | 0.016 | ENSMUSG00000050910 | protein_coding | Cdr2l | cerebellar degeneration-related protein 2-like | 237988 | 11 | 115381916 | 115396132 | 1 | Mm.102872 |
| 4.11 | 2.36 | 8.01 | 5.15 | DN3ab_Itr | DN3cd | -2.64 | -12.82 | 5.00E-07 | 3.00E-04 | 0.016 | ENSMUSG00000048376 | protein_coding | F2r | coagulation factor II (thrombin) receptor | 14062 | 13 | 95601803 | 95618487 | -1 | Mm.24816 |
| 4.4 | 4.76 | 2.74 | 4.05 | DN3ab_Itr | DN3cd | 0.63 | 3.15 | 0.012 | 0.09 | 0.016 | ENSMUSG00000036086 | protein_coding | Zranb3 | zinc finger, RAN-binding domain containing 3 | 226409 | 1 | 127954184 | 128103047 | -1 | Mm.425048 |
| 6.75 | 7.28 | 6.6 | 7.68 | DN3ab_Itr | DN3cd | 0.79 | 6.21 | 0.00017 | 0.0062 | 0.016 | ENSMUSG00000026313 | protein_coding | Hdac4 | histone deacetylase 4 | 208727 | 1 | 91928779 | 92195699 | -1 | Mm.318567 |
| 6.56 | 6.9 | 6.44 | 7.32 | DN3ab_Itr | DN3cd | 0.6 | 4.78 | 0.001 | 0.019 | 0.016 | ENSMUSG00000032883 | protein_coding | Acsl3 | acyl-CoA synthetase long-chain family member 3 | 74205 | 1 | 78657825 | 78707743 | 1 | Mm.276016 |
| 6.4 | 5.98 | 6.97 | 5.84 | DN3ab_Itr | DN3cd | -0.77 | -4.68 | 0.0012 | 0.021 | 0.016 | ENSMUSG00000028931 | protein_coding | Kcnab2 | potassium voltage-gated channel, shaker-related subfamily, beta member 2 | 16498 | 4 | 152390742 | 152477871 | -1 | Mm.388924 |
| 4.06 | 3.67 | 4.52 | 3.23 | DN3ab_Itr | DN3cd | -0.8 | -3.8 | 0.0043 | 0.046 | 0.016 | ENSMUSG00000041594 | protein_coding | Tmtc4 | transmembrane and tetratricopeptide repeat containing 4 | 70551 | 14 | 122918971 | 122984035 | -1 | Mm.271988 |
| -0.01 | -1.73 | 0.35 | -4.02 | DN3ab_Itr | DN3cd | -2.97 | -4.74 | 0.0011 | 0.02 | 0.016 | ENSMUSG00000047205 | protein_coding | Dusp18 | dual specificity phosphatase 18 | 75219 | 11 | 3895240 | 3901296 | 1 | Mm.32588 |
| 5.26 | 4.32 | 6.1 | 4.52 | DN3ab_Itr | DN3cd | -1.27 | -8.42 | 1.60E-05 | 0.0016 | 0.016 | ENSMUSG00000015966 | protein_coding | Il17rb | interleukin 17 receptor B | 50905 | 14 | 29996135 | 30008896 | -1 | Mm.269363 |
| 7.03 | 7.54 | 5.96 | 6.89 | DN3ab_Itr | DN3cd | 0.67 | 6.83 | 8.20E-05 | 0.0039 | 0.016 | ENSMUSG00000027496 | protein_coding | Aurka | aurora kinase A | 20878 | 2 | 172356190 | 172370535 | -1 | Mm.249363 |
| 6.73 | 7.37 | 5.66 | 7.14 | DN3ab_Itr | DN3cd | 0.98 | 5.14 | 0.00064 | 0.014 | 0.016 | ENSMUSG00000029366 | protein_coding | Dck | deoxycytidine kinase | 13178 | 5 | 88764996 | 88783281 | 1 | Mm.298892 |
| 7.96 | 6.63 | 9.34 | 7.38 | DN3ab_Itr | DN3cd | -1.68 | -11.68 | 1.10E-06 | 0.00037 | 0.017 | ENSMUSG00000015619 | protein_coding | Gata3 | GATA binding protein 3 | 14462 | 2 | 9857078 | 9890034 | -1 | Mm.313866 |
| 6.13 | 6.34 | 5.13 | 5.88 | DN3ab_Itr | DN3cd | 0.42 | 3.4 | 0.008 | 0.069 | 0.017 | ENSMUSG00000089726 | lincRNA | Mir17hg | Mir17 host gene (non-protein coding) | NA | 14 | 115042879 | 115046727 | 1 | |
| 5.89 | 5.69 | 6.58 | 5.98 | DN3ab_Itr | DN3cd | -0.4 | -4.15 | 0.0026 | 0.033 | 0.017 | ENSMUSG00000028826 | protein_coding | Maco1 | macoilin 1 | 66146 | 4 | 134802759 | 134853345 | -1 | Mm.99793 |
| -1.2 | 0.43 | -3.72 | 0.86 | DN3ab_Itr | DN3cd | 3.04 | 4.4 | 0.0018 | 0.027 | 0.017 | ENSMUSG00000020902 | protein_coding | Ntn1 | netrin 1 | 18208 | 11 | 68209364 | 68400823 | -1 | Mm.39095 |
| 7.21 | 7.41 | 6.51 | 7.19 | DN3ab_Itr | DN3cd | 0.4 | 3.61 | 0.0058 | 0.056 | 0.017 | ENSMUSG00000027968 | protein_coding | Larp7 | La ribonucleoprotein domain family, member 7 | 28036 | 3 | 127536714 | 127553349 | -1 | Mm.291032 |
| 5.22 | 4.08 | 6.74 | 4.82 | DN3ab_Itr | DN3cd | -1.59 | -8.87 | 1.10E-05 | 0.0012 | 0.017 | ENSMUSG00000054871 | protein_coding | Tmem158 | transmembrane protein 158 | 72309 | 9 | 123259053 | 123260764 | -1 | Mm.8569 |
| 7.34 | 7.68 | 6.94 | 7.6 | DN3ab_Itr | DN3cd | 0.48 | 6.35 | 0.00014 | 0.0057 | 0.017 | ENSMUSG00000020248 | protein_coding | Nfyb | nuclear transcription factor-Y beta | 18045 | 10 | 82748701 | 82764144 | -1 | Mm.245998 |
| 2.41 | 3.33 | 1.99 | 3.94 | DN3ab_Itr | DN3cd | 1.36 | 5.57 | 0.00037 | 0.0098 | 0.017 | ENSMUSG00000034164 | protein_coding | Emid1 | EMI domain containing 1 | 140703 | 11 | 5106265 | 5152257 | -1 | Mm.840 |
| 4.02 | 3.32 | 6.46 | 5.05 | DN3ab_Itr | DN3cd | -1.19 | -7.77 | 3.00E-05 | 0.0023 | 0.018 | ENSMUSG00000024235 | protein_coding | Map3k8 | mitogen-activated protein kinase kinase kinase 8 | 26410 | 18 | 4331327 | 4353015 | -1 | Mm.3275 |
| 7.07 | 6.64 | 7.82 | 6.89 | DN3ab_Itr | DN3cd | -0.68 | -5.69 | 0.00031 | 0.009 | 0.018 | ENSMUSG00000030987 | protein_coding | Stim1 | stromal interaction molecule 1 | 20866 | 7 | 102267806 | 102437319 | 1 | Mm.645 |
| 7.77 | 8.25 | 6.45 | 7.3 | DN3ab_Itr | DN3cd | 0.62 | 7.24 | 5.20E-05 | 0.0031 | 0.018 | ENSMUSG00000006398 | protein_coding | Cdc20 | cell division cycle 20 | 107995 | 4 | 118432901 | 118437352 | -1 | Mm.261764 Mm.289747 |
| 5.04 | 5.54 | 3.88 | 5.03 | DN3ab_Itr | DN3cd | 0.73 | 4.88 | 0.00091 | 0.017 | 0.018 | ENSMUSG00000027115 | protein_coding | Kif18a | kinesin family member 18A | 228421 | 2 | 109280738 | 109341747 | 1 | Mm.274086 |
| -4.01 | 0.15 | -1.33 | 0.12 | DN3ab_Itr | DN3cd | 2.86 | 4.42 | 0.0017 | 0.026 | 0.018 | ENSMUSG00000105263 | TEC | Gm42427 | predicted gene 42427 | NA | 5 | 44137980 | 44143271 | 1 | |
| 6.69 | 6.27 | 7.8 | 6.92 | DN3ab_Itr | DN3cd | -0.67 | -6.01 | 0.00021 | 0.0072 | 0.018 | ENSMUSG00000032479 | protein_coding | Map4 | microtubule-associated protein 4 | 17758 | 9 | 109931460 | 110083955 | 1 | Mm.217318 Mm.443428 Mm.484451 |
| -2.06 | -0.39 | -3.72 | 0.53 | DN3ab_Itr | DN3cd | 3.01 | 4.85 | 0.00094 | 0.018 | 0.018 | ENSMUSG00000030259 | protein_coding | Rassf8 | Ras association (RalGDS/AF-6) domain family (N-terminal) member 8 | 71323 | 6 | 145746748 | 145821079 | 1 | Mm.158669 Mm.444985 |
| 6.57 | 7.98 | 4.88 | 7.28 | DN3ab_Itr | DN3cd | 1.74 | 8 | 2.40E-05 | 0.002 | 0.018 | ENSMUSG00000028634 | protein_coding | Hivep3 | human immunodeficiency virus type I enhancer binding protein 3 | 16656 | 4 | 119733784 | 120138045 | 1 | Mm.302758 Mm.394479 Mm.422538 Mm.487430 Mm.488325 |
| 6.39 | 6.64 | 6.23 | 6.87 | DN3ab_Itr | DN3cd | 0.43 | 4.46 | 0.0016 | 0.025 | 0.019 | ENSMUSG00000040167 | protein_coding | Ikzf5 | IKAROS family zinc finger 5 | 67143 | 7 | 131388652 | 131410521 | -1 | Mm.334273 |
| 7.96 | 7.7 | 8.35 | 7.82 | DN3ab_Itr | DN3cd | -0.39 | -5.92 | 0.00024 | 0.0076 | 0.019 | ENSMUSG00000026219 | protein_coding | Trip12 | thyroid hormone receptor interactor 12 | 14897 | 1 | 84721189 | 84840516 | -1 | Mm.209265 |
| 8.62 | 9.02 | 7.5 | 8.72 | DN3ab_Itr | DN3cd | 0.73 | 3.92 | 0.0036 | 0.041 | 0.019 | ENSMUSG00000047139 | protein_coding | Cd24a | CD24a antigen | 12484 | 10 | 43578284 | 43584265 | 1 | Mm.29742 |
| 5.21 | 5.55 | 4.3 | 5.26 | DN3ab_Itr | DN3cd | 0.58 | 3.97 | 0.0033 | 0.039 | 0.019 | ENSMUSG00000030872 | protein_coding | Gga2 | golgi associated, gamma adaptin ear containing, ARF binding protein 2 | 74105 | 7 | 121986722 | 122021222 | -1 | Mm.29619 |
| 3.78 | 3.34 | 4.91 | 3.68 | DN3ab_Itr | DN3cd | -0.87 | -4.66 | 0.0012 | 0.021 | 0.019 | ENSMUSG00000037949 | protein_coding | Ano10 | anoctamin 10 | 102566 | 9 | 122175874 | 122294423 | -1 | Mm.87676 |
| 8.09 | 8.4 | 7.65 | 8.35 | DN3ab_Itr | DN3cd | 0.48 | 5.1 | 0.00068 | 0.014 | 0.019 | ENSMUSG00000002748 | protein_coding | Baz1b | bromodomain adjacent to zinc finger domain, 1B | 22385 | 5 | 135187264 | 135246129 | 1 | Mm.40331 |
| 4.13 | 4.59 | 3.13 | 4.34 | DN3ab_Itr | DN3cd | 0.73 | 4.24 | 0.0022 | 0.03 | 0.02 | ENSMUSG00000022833 | protein_coding | Ccdc14 | coiled-coil domain containing 14 | 239839 | 16 | 34690616 | 34725202 | 1 | Mm.272045 |
| 6.69 | 7.23 | 6.38 | 7.34 | DN3ab_Itr | DN3cd | 0.73 | 7.16 | 5.70E-05 | 0.0033 | 0.02 | ENSMUSG00000020455 | protein_coding | Trim11 | tripartite motif-containing 11 | 94091 | 11 | 58978093 | 58991458 | 1 | Mm.248049 |
| 5.23 | 4.61 | 6.77 | 5.47 | DN3ab_Itr | DN3cd | -1.02 | -6.28 | 0.00015 | 0.0059 | 0.02 | ENSMUSG00000031453 | protein_coding | Rasa3 | RAS p21 protein activator 3 | 19414 | 8 | 13566948 | 13677603 | -1 | Mm.18517 |
| -3.22 | -1.41 | -3.72 | 1.35 | DN3ab_Itr | DN3cd | 3.72 | 4.85 | 0.00095 | 0.018 | 0.02 | ENSMUSG00000036040 | protein_coding | Adamtsl2 | ADAMTS-like 2 | 77794 | 2 | 27079379 | 27108981 | 1 | Mm.330088 |
| 5.64 | 6.03 | 4.82 | 5.83 | DN3ab_Itr | DN3cd | 0.63 | 4.36 | 0.0019 | 0.027 | 0.02 | ENSMUSG00000074994 | protein_coding | Qser1 | glutamine and serine rich 1 | 99003 | 2 | 104754795 | 104816760 | -1 | Mm.274314 Mm.491693 |
| 5.37 | 6.05 | 4.18 | 5.37 | DN3ab_Itr | DN3cd | 0.86 | 7.3 | 4.90E-05 | 0.003 | 0.02 | ENSMUSG00000072082 | protein_coding | Ccnf | cyclin F | 12449 | 17 | 24222198 | 24251484 | -1 | Mm.77695 |
| 5.48 | 5.09 | 6.19 | 5.39 | DN3ab_Itr | DN3cd | -0.6 | -6.07 | 2.00E-04 | 0.0068 | 0.02 | ENSMUSG00000045176 | protein_coding | Borcs6 | BLOC-1 related complex subunit 6 | 71923 | 11 | 69059717 | 69061578 | 1 | Mm.340257 |
| 4.35 | 4.86 | 2.67 | 4.07 | DN3ab_Itr | DN3cd | 0.76 | 3.94 | 0.0035 | 0.04 | 0.02 | ENSMUSG00000023008 | protein_coding | Fmnl3 | formin-like 3 | 22379 | 15 | 99317225 | 99370482 | -1 | Mm.29043 |
| 6.43 | 7.03 | 6.07 | 6.24 | DN3ab_Itr | DN3cd | 0.43 | 4.15 | 0.0026 | 0.033 | 0.02 | ENSMUSG00000037278 | protein_coding | Tmem97 | transmembrane protein 97 | 69071 | 11 | 78541817 | 78550777 | -1 | Mm.29431 |
| 3.31 | 4.21 | 1.17 | 3.61 | DN3ab_Itr | DN3cd | 1.25 | 4.06 | 0.0029 | 0.036 | 0.021 | ENSMUSG00000033752 | protein_coding | Mnd1 | meiotic nuclear divisions 1 | 76915 | 3 | 84087933 | 84155786 | -1 | Mm.274829 |
| 5.03 | 4.41 | 6.09 | 4.83 | DN3ab_Itr | DN3cd | -0.96 | -6.39 | 0.00014 | 0.0055 | 0.021 | ENSMUSG00000098112 | protein_coding | Bin2 | bridging integrator 2 | 668218 | 15 | 100641077 | 100669553 | -1 | Mm.451225 |
| 6.48 | 4.69 | 7.94 | 5.38 | DN3ab_Itr | DN3cd | -2.22 | -12.23 | 7.50E-07 | 0.00033 | 0.021 | ENSMUSG00000062232 | protein_coding | Rapgef2 | Rap guanine nucleotide exchange factor (GEF) 2 | 76089 | 3 | 79062516 | 79286517 | -1 | Mm.31220 Mm.462396 |
| 6.9 | 7.46 | 5.93 | 6.86 | DN3ab_Itr | DN3cd | 0.7 | 7.97 | 2.50E-05 | 0.002 | 0.021 | ENSMUSG00000025747 | protein_coding | Tyms | thymidylate synthase | 22171 | 5 | 30058202 | 30073617 | -1 | Mm.268395 |
| 6.82 | 7.36 | 5.72 | 6.73 | DN3ab_Itr | DN3cd | 0.72 | 6.5 | 0.00012 | 0.005 | 0.021 | ENSMUSG00000005233 | protein_coding | Spc25 | SPC25, NDC80 kinetochore complex component, homolog (S. cerevisiae) | 66442 | 2 | 69193895 | 69206194 | -1 | Mm.272969 Mm.486261 |
| -3.22 | -1.11 | -3.72 | 0.96 | DN3ab_Itr | DN3cd | 3.56 | 5.72 | 3.00E-04 | 0.0088 | 0.021 | ENSMUSG00000110187 | TEC | Gm45496 | predicted gene 45496 | NA | 8 | 88560033 | 88562949 | 1 | |
| 4.72 | 2.95 | 8.66 | 5.74 | DN3ab_Itr | DN3cd | -2.67 | -11.93 | 9.20E-07 | 0.00036 | 0.021 | ENSMUSG00000055435 | protein_coding | Maf | avian musculoaponeurotic fibrosarcoma oncogene homolog | 17132 | 8 | 115682942 | 115707794 | -1 | Mm.275549 Mm.439772 Mm.488377 |
| 6.93 | 6.62 | 7.26 | 6.47 | DN3ab_Itr | DN3cd | -0.54 | -4.65 | 0.0012 | 0.021 | 0.022 | ENSMUSG00000040659 | protein_coding | Efhd2 | EF hand domain containing 2 | 27984 | 4 | 141858142 | 141874920 | -1 | Mm.395598 |
| 3.44 | 2.12 | 7.35 | 4.4 | DN3ab_Itr | DN3cd | -2.6 | -8.11 | 2.20E-05 | 0.0019 | 0.022 | ENSMUSG00000005672 | protein_coding | Kit | KIT proto-oncogene receptor tyrosine kinase | 16590 | 5 | 75574916 | 75656722 | 1 | Mm.247073 |
| 6.25 | 7.17 | 5.32 | 7.17 | DN3ab_Itr | DN3cd | 1.3 | 5.98 | 0.00022 | 0.0073 | 0.022 | ENSMUSG00000052949 | protein_coding | Rnf157 | ring finger protein 157 | 217340 | 11 | 116336353 | 116413032 | -1 | Mm.426344 |
| 5.56 | 5.83 | 4.48 | 5.28 | DN3ab_Itr | DN3cd | 0.46 | 3.68 | 0.0052 | 0.052 | 0.022 | ENSMUSG00000021569 | protein_coding | Trip13 | thyroid hormone receptor interactor 13 | 69716 | 13 | 73911347 | 73937784 | -1 | Mm.275095 |
| 2.99 | 4.53 | 2.69 | 3.58 | DN3ab_Itr | DN3cd | 1.29 | 8.02 | 2.40E-05 | 0.002 | 0.022 | ENSMUSG00000028602 | protein_coding | Tnfrsf8 | tumor necrosis factor receptor superfamily, member 8 | 21941 | 4 | 145267137 | 145315164 | -1 | Mm.12810 |
| 7.73 | 7 | 8.06 | 7.65 | DN3ab_Itr | DN3cd | -0.57 | -7.15 | 5.80E-05 | 0.0033 | 0.022 | ENSMUSG00000030629 | protein_coding | Zfand6 | zinc finger, AN1-type domain 6 | 65098 | 7 | 84613766 | 84689959 | -1 | Mm.209349 Mm.397115 |
| 7.05 | 7.41 | 6.44 | 7.2 | DN3ab_Itr | DN3cd | 0.53 | 5.57 | 0.00037 | 0.0098 | 0.022 | ENSMUSG00000044783 | protein_coding | Hjurp | Holliday junction recognition protein | 212427 | 1 | 88262471 | 88277633 | -1 | Mm.384762 |
| 4.02 | 1.83 | 4.88 | 3.51 | DN3ab_Itr | DN3cd | -1.64 | -8.68 | 1.30E-05 | 0.0014 | 0.022 | ENSMUSG00000019947 | protein_coding | Arid5b | AT rich interactive domain 5B (MRF1-like) | 71371 | 10 | 68092520 | 68278740 | -1 | Mm.440357 |
| 0.71 | 2.16 | 0.36 | 3.42 | DN3ab_Itr | DN3cd | 2.18 | 5.57 | 0.00037 | 0.0098 | 0.023 | ENSMUSG00000086527 | antisense | Gm15856 | predicted gene 15856 | NA | 6 | 121223692 | 121226153 | -1 | |
| 5.92 | 6.81 | 5.5 | 5.85 | DN3ab_Itr | DN3cd | 0.67 | 5.16 | 0.00062 | 0.014 | 0.023 | ENSMUSG00000027490 | protein_coding | E2f1 | E2F transcription factor 1 | 13555 | 2 | 154559407 | 154569892 | -1 | Mm.18036 Mm.441323 |
| 1.17 | 2.64 | 1.35 | 4.22 | DN3ab_Itr | DN3cd | 2.19 | 6.4 | 0.00013 | 0.0054 | 0.023 | ENSMUSG00000097604 | lincRNA | Gm17322 | predicted gene, 17322 | NA | 9 | 57998041 | 58007451 | 1 | |
| 9.61 | 8.25 | 9.43 | 7.75 | DN3ab_Itr | DN3cd | -1.5 | -19.23 | 1.50E-08 | 5.90E-05 | 0.023 | ENSMUSG00000066362 | processed_pseudogene | Rps13-ps1 | ribosomal protein S13, pseudogene 1 | NA | 8 | 87047689 | 87048226 | 1 | |
| 7.58 | 7.08 | 8.13 | 7.95 | DN3ab_Itr | DN3cd | -0.33 | -4.32 | 0.002 | 0.028 | 0.023 | ENSMUSG00000004040 | protein_coding | Stat3 | signal transducer and activator of transcription 3 | 20848 | 11 | 100885098 | 100939540 | -1 | Mm.249934 Mm.473190 |
| 8.5 | 8.99 | 7.71 | 8.85 | DN3ab_Itr | DN3cd | 0.76 | 5.04 | 0.00073 | 0.015 | 0.024 | ENSMUSG00000078619 | protein_coding | Smarcd2 | SWI/SNF related, matrix associated, actin dependent regulator of chromatin, subfamily d, member 2 | 83796 | 11 | 106263179 | 106272972 | -1 | Mm.21772 |
| 8 | 8.22 | 7.53 | 8.11 | DN3ab_Itr | DN3cd | 0.38 | 4.36 | 0.0019 | 0.027 | 0.024 | ENSMUSG00000024974 | protein_coding | Smc3 | structural maintenance of chromosomes 3 | 13006 | 19 | 53600398 | 53645833 | 1 | Mm.14910 |
| 6.58 | 4.75 | 6.62 | 5.26 | DN3ab_Itr | DN3cd | -1.59 | -13.72 | 2.80E-07 | 0.00024 | 0.024 | ENSMUSG00000027009 | protein_coding | Itga4 | integrin alpha 4 | 16401 | 2 | 79255426 | 79333123 | 1 | Mm.31903 |
| 6.1 | 5.76 | 6.9 | 6.1 | DN3ab_Itr | DN3cd | -0.58 | -5.15 | 0.00063 | 0.014 | 0.024 | ENSMUSG00000084817 | processed_pseudogene | Gm5526 | predicted pseudogene 5526 | NA | 1 | 45857332 | 45857754 | -1 | |
| -1.2 | 0.67 | -1.36 | 2.91 | DN3ab_Itr | DN3cd | 3.12 | 5.29 | 0.00053 | 0.012 | 0.024 | ENSMUSG00000109002 | antisense | Gm38405 | predicted gene, 38405 | NA | 7 | 99870666 | 99889358 | -1 | |
| 5.97 | 4.81 | 7.08 | 5.23 | DN3ab_Itr | DN3cd | -1.53 | -9.22 | 7.70E-06 | 0.00098 | 0.024 | ENSMUSG00000025026 | protein_coding | Add3 | adducin 3 (gamma) | 27360 | 19 | 53140445 | 53247399 | 1 | Mm.426080 Mm.491114 |
| 6.75 | 4.66 | 7.06 | 5.79 | DN3ab_Itr | DN3cd | -1.64 | -8.29 | 1.80E-05 | 0.0017 | 0.024 | ENSMUSG00000023828 | protein_coding | Slc22a3 | solute carrier family 22 (organic cation transporter), member 3 | 20519 | 17 | 12418960 | 12507704 | -1 | Mm.99252 |
| 4.82 | 4.13 | 6.17 | 4.85 | DN3ab_Itr | DN3cd | -1.05 | -6.88 | 7.80E-05 | 0.0038 | 0.025 | ENSMUSG00000029344 | protein_coding | Tpst2 | protein-tyrosine sulfotransferase 2 | 22022 | 5 | 112276691 | 112315361 | 1 | Mm.250631 |
| 2.49 | 1.82 | 3.44 | 1.78 | DN3ab_Itr | DN3cd | -1.18 | -4.74 | 0.0011 | 0.02 | 0.025 | ENSMUSG00000046808 | polymorphic_pseudogene | Atp10d | ATPase, class V, type 10D | 231287 | 5 | 72203329 | 72298775 | 1 | Mm.32775 Mm.441066 |
| 3.72 | 2.93 | 5.32 | 3.85 | DN3ab_Itr | DN3cd | -1.21 | -7.44 | 4.20E-05 | 0.0027 | 0.025 | ENSMUSG00000032440 | protein_coding | Tgfbr2 | transforming growth factor, beta receptor II | 21813 | 9 | 116084293 | 116175360 | -1 | Mm.172346 |
| 1.54 | 2.7 | 0.4 | 2.75 | DN3ab_Itr | DN3cd | 1.56 | 5.48 | 0.00041 | 0.01 | 0.025 | ENSMUSG00000041930 | protein_coding | Fam222a | family with sequence similarity 222, member A | 433940 | 5 | 114568016 | 114613220 | 1 | Mm.194212 |
| 8.27 | 8.43 | 8.03 | 8.46 | DN3ab_Itr | DN3cd | 0.28 | 4.16 | 0.0025 | 0.032 | 0.025 | ENSMUSG00000055436 | protein_coding | Srsf11 | serine/arginine-rich splicing factor 11 | 69207 | 3 | 158010473 | 158036639 | -1 | Mm.223946 |
| 5.98 | 4.46 | 8.9 | 6.53 | DN3ab_Itr | DN3cd | -2.1 | -10.93 | 1.90E-06 | 0.00048 | 0.025 | ENSMUSG00000040747 | protein_coding | Cd53 | CD53 antigen | 12508 | 3 | 106759921 | 106790149 | -1 | Mm.316861 |
| 5.57 | 7.14 | 5.96 | 7.08 | DN3ab_Itr | DN3cd | 1.35 | 12.14 | 7.90E-07 | 0.00034 | 0.025 | ENSMUSG00000040345 | protein_coding | Arhgap9 | Rho GTPase activating protein 9 | 216445 | 10 | 127321964 | 127329943 | 1 | Mm.227198 |
| 5.7 | 5.27 | 6.1 | 5.17 | DN3ab_Itr | DN3cd | -0.67 | -5.54 | 0.00038 | 0.01 | 0.025 | ENSMUSG00000073982 | protein_coding | Rhog | ras homolog family member G | 56212 | 7 | 102239123 | 102256054 | -1 | Mm.259795 |
| 4.1 | 2.86 | 4.91 | 4.21 | DN3ab_Itr | DN3cd | -0.9 | -6.79 | 8.60E-05 | 0.0041 | 0.025 | ENSMUSG00000036503 | protein_coding | Rnf13 | ring finger protein 13 | 24017 | 3 | 57736062 | 57835233 | 1 | Mm.274360 Mm.445604 |

| | | | | | | | | | | | | | | | | | | | | |
|---|---|---|---|---|---|---|---|---|---|---|---|---|---|---|---|---|---|---|---|---|
| 6.92 | 7.3 | 6.74 | 7.49 | DN3ab_ltr | DN3cd | 0.55 | 6.04 | 2.00E-04 | 0.007 | 0.026 | ENSMUSG00000038582 | protein_coding | Pptc7 | PTC7 protein phosphatase homolog | 320717 | 5 | 122284365 | 122324281 | 1 | Mm.489670 |
| 4.14 | 3.71 | 5.25 | 4.16 | DN3ab_ltr | DN3cd | -0.79 | -4.92 | 0.00085 | 0.017 | 0.026 | ENSMUSG00000074305 | protein_coding | Peak1 | pseudopodium-enriched atypical kinase 1 | 244895 | 9 | 56201126 | 56418067 | -1 | Mm.490331 |
| 7.68 | 8.15 | 6.98 | 7.8 | DN3ab_ltr | DN3cd | 0.61 | 7.06 | 6.30E-05 | 0.0034 | 0.026 | ENSMUSG00000028560 | protein_coding | Usp1 | ubiquitin specific peptidase 1 | 230484 | 4 | 98923810 | 98935543 | 1 | Mm.371692 |
| 6.06 | 6.54 | 4.92 | 5.89 | DN3ab_ltr | DN3cd | 0.66 | 5.69 | 0.00031 | 0.009 | 0.026 | ENSMUSG00000025574 | protein_coding | Tk1 | thymidine kinase 1 | 21877 | 11 | 117815526 | 117826092 | -1 | Mm.2661 Mm.443968 |
| 7.34 | 8 | 6.8 | 8.33 | DN3ab_ltr | DN3cd | 1.05 | 4.97 | 8.00E-04 | 0.016 | 0.026 | ENSMUSG00000045382 | protein_coding | Cxcr4 | chemokine (C-X-C motif) receptor 4 | 12767 | 1 | 128588199 | 128592293 | -1 | Mm.1401 |
| 6.49 | 7.37 | 5.53 | 6.8 | DN3ab_ltr | DN3cd | 1.02 | 10.69 | 2.30E-06 | 0.00053 | 0.026 | ENSMUSG00000038943 | protein_coding | Prc1 | protein regulator of cytokinesis 1 | 233406 | 7 | 80294450 | 80316259 | 1 | Mm.227274 |
| 6.1 | 6.51 | 5.51 | 6.33 | DN3ab_ltr | DN3cd | 0.58 | 5.71 | 0.00031 | 0.0088 | 0.026 | ENSMUSG00000031922 | protein_coding | Cep57 | centrosomal protein 57 | 74360 | 9 | 13807792 | 13827107 | -1 | Mm.157212 |
| 3.43 | 1.73 | 4.2 | 1.28 | DN3ab_ltr | DN3cd | -2.24 | -7.45 | 4.20E-05 | 0.0027 | 0.026 | ENSMUSG00000003849 | protein_coding | Nqo1 | NAD(P)H dehydrogenase, quinone 1 | 18104 | 8 | 107388225 | 107403206 | -1 | Mm.252 |
| 8.14 | 8.66 | 7.73 | 8.56 | DN3ab_ltr | DN3cd | 0.66 | 8.41 | 1.60E-05 | 0.0016 | 0.026 | ENSMUSG00000022710 | protein_coding | Usp7 | ubiquitin specific peptidase 7 | 252870 | 16 | 8689595 | 8792308 | -1 | Mm.295330 |
| 7.2 | 7.37 | 6.52 | 7 | DN3ab_ltr | DN3cd | 0.3 | 3.82 | 0.0042 | 0.045 | 0.027 | ENSMUSG00000029427 | protein_coding | Zcchc8 | zinc finger, CCHC domain containing 8 | 70650 | 5 | 123698294 | 123721100 | -1 | Mm.279427 |
| 6.27 | 5.93 | 7.03 | 6.39 | DN3ab_ltr | DN3cd | -0.5 | -6.29 | 0.00015 | 0.0058 | 0.027 | ENSMUSG00000022867 | protein_coding | Usp25 | ubiquitin specific peptidase 25 | 30940 | 16 | 77013706 | 77116780 | 1 | Mm.40986 |
| 5.09 | 4.28 | 4.49 | 4.2 | DN3ab_ltr | DN3cd | -0.59 | -4.5 | 0.0015 | 0.025 | 0.027 | ENSMUSG00000017210 | protein_coding | Med24 | mediator complex subunit 24 | 23989 | 11 | 98704591 | 98729435 | -1 | Mm.246493 |
| -0.24 | -2.34 | 3.55 | -1.12 | DN3ab_ltr | DN3cd | -3.67 | -5.86 | 0.00026 | 0.008 | 0.027 | ENSMUSG00000037613 | protein_coding | Tnfrsf23 | tumor necrosis factor receptor superfamily, member 23 | 79201 | 7 | 143665809 | 143685872 | -1 | Mm.290780 |
| 6.08 | 7.85 | 4.45 | 8.16 | DN3ab_ltr | DN3cd | 2.43 | 5.39 | 0.00046 | 0.011 | 0.027 | ENSMUSG00000000031 | lincRNA | H19 | H19, imprinted maternally expressed transcript | NA | 7 | 142575529 | 142578143 | -1 | |
| 4.94 | 5.38 | 4.12 | 5.09 | DN3ab_ltr | DN3cd | 0.64 | 4.97 | 8.00E-04 | 0.016 | 0.028 | ENSMUSG00000037108 | protein_coding | Zcwpw1 | zinc finger, CW type with PWWP domain 1 | 381678 | 5 | 137787798 | 137822621 | 1 | Mm.332765 |
| 4.68 | 5.16 | 3.79 | 4.87 | DN3ab_ltr | DN3cd | 0.7 | 4.77 | 0.0011 | 0.019 | 0.028 | ENSMUSG00000051278 | protein_coding | Zgrf1 | zinc finger, GRF-type containing 1 | 71643 | 3 | 127553489 | 127618023 | 1 | Mm.274884 Mm.454006 |
| 9.06 | 8.65 | 10.17 | 9.43 | DN3ab_ltr | DN3cd | -0.59 | -7.19 | 5.50E-05 | 0.0032 | 0.028 | ENSMUSG00000060802 | protein_coding | B2m | beta-2 microglobulin | 12010 | 2 | 122147686 | 122153083 | 1 | Mm.163 |
| 4.57 | 4.9 | 4.42 | 5.26 | DN3ab_ltr | DN3cd | 0.56 | 4.28 | 0.0021 | 0.029 | 0.03 | ENSMUSG00000083757 | transcribed_processed_pseudogene | Rps15a-ps4 | ribosomal protein S15A, pseudogene 4 | NA | 4 | 132219988 | 132220489 | -1 | |
| 3.95 | 5.2 | 3.88 | 4.31 | DN3ab_ltr | DN3cd | 0.91 | 4.5 | 0.0016 | 0.025 | 0.03 | ENSMUSG00000068855 | protein_coding | Hist2h2ac | histone cluster 2, H2ac | 319176 | 3 | 96220361 | 96220880 | -1 | Mm.358954 |
| -1.82 | -3.92 | 0.33 | -4.02 | DN3ab_ltr | DN3cd | -3.32 | -5.78 | 0.00028 | 0.0084 | 0.03 | ENSMUSG00000108068 | processed_pseudogene | Gm44075 | predicted gene, 44075 | NA | 6 | 63783852 | 63784067 | -1 | |

| GFP 4h | E47 4h | GFP 16h | E47 16h | sample_1 | sample_2 | logFC | test_stat | p_value | q_value | pvlltr | Geneid | biotype | SYMBOL | GENENAM | ENTREZID | Chr | Start | End | Strand | UniGene |
|---|---|---|---|---|---|---|---|---|---|---|---|---|---|---|---|---|---|---|---|---|
| 7.42 | 8.51 | 7.25 | 8.55 | GFP | E47 | 1.13 | 19.74 | 4.00E-09 | 6.40E-05 | 0.16 | ENSMUSG | protein_cc | Sla | src-like ada | 20491 | 15 | 66780819 | 66831829 | -1 | Mm.7601 |
| 6.84 | 7.98 | 7.01 | 8.06 | GFP | E47 | 1.12 | 19.6 | 4.30E-09 | 6.40E-05 | 0.51 | ENSMUSG | protein_cc | Sdc4 | syndecan 4 | 20971 | 2 | 1.64E+08 | 1.64E+08 | -1 | Mm.3815 |
| 9.38 | 10.15 | 9.38 | 10.29 | GFP | E47 | 0.8 | 19.13 | 5.40E-09 | 6.40E-05 | 0.19 | ENSMUSG | protein_cc | Larp1 | La ribonuc | 73158 | 11 | 58009064 | 58062034 | 1 | Mm.248843 |
| 4.55 | 6.65 | 3.98 | 6.7 | GFP | E47 | 2.18 | 18.24 | 8.50E-09 | 7.50E-05 | 0.064 | ENSMUSG | protein_cc | Spsb1 | splA/ryanc | 74646 | 4 | 1.5E+08 | 1.5E+08 | -1 | Mm.30 |
| -1.77 | 5.64 | -0.51 | 5.63 | GFP | E47 | 7.11 | 17.03 | 1.60E-08 | 0.00011 | 0.23 | ENSMUSG | protein_cc | Esr1 | estrogen r | 13982 | 10 | 4611593 | 5005614 | 1 | Mm.9213 Mm.463262 Mm.489063 |
| 0.68 | 3.87 | 0.58 | 4.59 | GFP | E47 | 3.31 | 16.7 | 1.90E-08 | 0.00011 | 0.13 | ENSMUSG | protein_cc | Cdk20 | cyclin-depe | 105278 | 13 | 64432314 | 64441773 | 1 | Mm.74982 |
| 9.69 | 10.55 | 9.92 | 10.85 | GFP | E47 | 0.88 | 16.57 | 2.10E-08 | 0.00011 | 0.68 | ENSMUSG | protein_cc | Id2 | inhibitor o | 15902 | 12 | 25093799 | 25097140 | -1 | Mm.34871 |
| 7.08 | 8.64 | 6.76 | 8.51 | GFP | E47 | 1.59 | 15.29 | 4.40E-08 | 0.00017 | 0.56 | ENSMUSG | protein_cc | F2r | coagulatio | 14062 | 13 | 95601803 | 95618487 | -1 | Mm.24816 |
| 4.4 | 5.93 | 4.85 | 6.73 | GFP | E47 | 1.6 | 15.1 | 4.90E-08 | 0.00018 | 0.19 | ENSMUSG | TR_C_gene | Trbc2 | T cell recep | NA | 6 | 41546730 | 41548352 | 1 | |
| -0.41 | 3.01 | -0.11 | 3.47 | GFP | E47 | 3.45 | 14.46 | 7.40E-08 | 0.00022 | 0.81 | ENSMUSG | protein_cc | Olfr523 | olfactory r | 258511 | 7 | 1.4E+08 | 1.4E+08 | 1 | Mm.334306 |
| 2.22 | 4.16 | 2.69 | 4.84 | GFP | E47 | 1.98 | 14.15 | 9.00E-08 | 0.00025 | 0.56 | ENSMUSG | TR_C_gene | Trbc1 | T cell recep | NA | 6 | 41538218 | 41539881 | 1 | |
| 1.53 | 3.64 | 1.78 | 3.88 | GFP | E47 | 2.11 | 13.93 | 1.00E-07 | 0.00027 | 0.97 | ENSMUSG | protein_cc | St8sia1 | ST8 alpha- | 20449 | 6 | 1.43E+08 | 1.43E+08 | -1 | Mm.260838 |
| -2.2 | 2.39 | -1.75 | 3.56 | GFP | E47 | 4.72 | 13.57 | 1.30E-07 | 0.00029 | 0.44 | ENSMUSG | protein_cc | Wnt10b | wingless-ty | 22410 | 15 | 98770712 | 98778150 | -1 | Mm.4709 |
| 4.72 | 6.02 | 4.22 | 5.61 | GFP | E47 | 1.32 | 13.54 | 1.40E-07 | 0.00029 | 0.76 | ENSMUSG | protein_cc | Cemip | cell migrat | 80982 | 7 | 83932857 | 84086502 | -1 | Mm.160389 |
| 11.79 | 11.42 | 11.87 | 11.4 | GFP | E47 | -0.38 | -13.44 | 1.50E-07 | 0.00029 | 0.14 | ENSMUSG | protein_cc | Ldha | lactate deh | 16828 | 7 | 46841475 | 46855627 | 1 | Mm.29324 |
| -5.08 | -0.89 | -4.42 | -0.54 | GFP | E47 | 4.12 | 13.43 | 1.50E-07 | 0.00029 | 0.68 | ENSMUSG | protein_cc | Sox8 | SRY (sex de | 20681 | 17 | 25565892 | 25570686 | -1 | Mm.258220 |
| -5.08 | 0.15 | -3.18 | 0.03 | GFP | E47 | 4.97 | 12.98 | 2.00E-07 | 0.00037 | 0.065 | ENSMUSG | protein_cc | Palmd | palmdelph | 114301 | 3 | 1.17E+08 | 1.17E+08 | -1 | Mm.253736 |
| 7.51 | 8.38 | 7.04 | 8.18 | GFP | E47 | 0.91 | 12.79 | 2.30E-07 | 0.00039 | 0.18 | ENSMUSG | protein_cc | Prss35 | protease, s | 244954 | 9 | 86743649 | 86758443 | 1 | Mm.257629 |
| 5.65 | 4.57 | 5.53 | 4.29 | GFP | E47 | -1.11 | -12.68 | 2.50E-07 | 4.00E-04 | 0.45 | ENSMUSG | protein_cc | Zbtb32 | zinc finger | 58206 | 7 | 30589681 | 30598909 | -1 | Mm.116789 |
| 7.53 | 8.32 | 7.32 | 8.39 | GFP | E47 | 0.84 | 12.55 | 2.70E-07 | 0.00042 | 0.11 | ENSMUSG | protein_cc | Egln3 | egl-9 famil | 112407 | 12 | 54178981 | 54203860 | -1 | Mm.133037 |
| 6.44 | 5.48 | 6.57 | 5.46 | GFP | E47 | -0.99 | -12.28 | 3.30E-07 | 0.00049 | 0.44 | ENSMUSG | protein_cc | Csn2 | casein beta | 12991 | 5 | 87692624 | 87699630 | -1 | Mm.268737 |
| 2.46 | 3.84 | 2.32 | 3.57 | GFP | E47 | 1.36 | 12.22 | 3.50E-07 | 0.00049 | 0.66 | ENSMUSG | protein_cc | Stom | stomatin | 13830 | 2 | 35313986 | 35336976 | -1 | Mm.295284 |
| 9.07 | 8.59 | 9.14 | 8.55 | GFP | E47 | -0.5 | -12.17 | 3.60E-07 | 0.00049 | 0.26 | ENSMUSG | protein_cc | Osm | oncostatin | 18413 | 11 | 4236420 | 4241026 | 1 | Mm.131422 |
| -5.08 | 0.86 | -1.84 | 5.4 | GFP | E47 | 6.12 | 11.92 | 4.30E-07 | 0.00054 | 0.41 | ENSMUSG | protein_cc | Gal | galanin | 14419 | 19 | 3409919 | 3414472 | -1 | Mm.4655 |
| 3.65 | 4.83 | 3.41 | 4.74 | GFP | E47 | 1.21 | 11.69 | 5.20E-07 | 0.00061 | 0.59 | ENSMUSG | protein_cc | Vamp1 | vesicle-ass | 22317 | 6 | 1.25E+08 | 1.25E+08 | 1 | Mm.32321 |
| 6.36 | 7.32 | 6.17 | 7.33 | GFP | E47 | 1 | 11.59 | 5.60E-07 | 0.00062 | 0.43 | ENSMUSG | protein_cc | Tcf3 | transcripti | 21423 | 10 | 80409514 | 80433647 | -1 | Mm.3406 |
| 5.89 | 6.76 | 5.5 | 6.55 | GFP | E47 | 0.89 | 11.47 | 6.20E-07 | 0.00064 | 0.39 | ENSMUSG | protein_cc | Bcl2l11 | BCL2-like 1 | 12125 | 2 | 1.28E+08 | 1.28E+08 | 1 | Mm.141083 Mm.453214 |
| -0.24 | 2.42 | -0.35 | 2.8 | GFP | E47 | 2.74 | 11.42 | 6.40E-07 | 0.00064 | 0.47 | ENSMUSG | processed_ | Gm38158 | predicted g | NA | 1 | 1.77E+08 | 1.77E+08 | 1 | |
| 4.18 | 5.77 | 3.71 | 5.99 | GFP | E47 | 1.69 | 11.36 | 6.70E-07 | 0.00064 | 0.098 | ENSMUSG | protein_cc | Irf8 | interferon | 15900 | 8 | 1.21E+08 | 1.21E+08 | 1 | Mm.334861 |
| 4.97 | 6.83 | 5.1 | 6.79 | GFP | E47 | 1.83 | 11.35 | 6.80E-07 | 0.00064 | 0.71 | ENSMUSG | protein_cc | Esm1 | endothelia | 71690 | 13 | 1.13E+08 | 1.13E+08 | 1 | Mm.38929 |
| 1.27 | 2.92 | 1.18 | 2.93 | GFP | E47 | 1.67 | 11.22 | 7.50E-07 | 0.00066 | 0.8 | ENSMUSG | protein_cc | Zfp941 | zinc finger | 407812 | 7 | 1.41E+08 | 1.41E+08 | -1 | Mm.359154 |
| 3.75 | 5.74 | 3.6 | 5.57 | GFP | E47 | 1.99 | 11.2 | 7.60E-07 | 0.00066 | 0.96 | ENSMUSG | protein_cc | Ccno | cyclin O | 218630 | 13 | 1.13E+08 | 1.13E+08 | 1 | Mm.25457 |
| 0.46 | 2.66 | 0.44 | 2.53 | GFP | E47 | 2.18 | 11.18 | 7.80E-07 | 0.00066 | 0.85 | ENSMUSG | protein_cc | Lipg | lipase, end | 16891 | 18 | 74939322 | 74961263 | -1 | Mm.299647 |
| 6.1 | 6.82 | 6.06 | 7.05 | GFP | E47 | 0.77 | 11.13 | 8.10E-07 | 0.00067 | 0.1 | ENSMUSG | protein_cc | Tox2 | TOX high m | 269389 | 2 | 1.63E+08 | 1.63E+08 | 1 | Mm.207709 |
| 6.74 | 7.55 | 6.46 | 7.57 | GFP | E47 | 0.86 | 11.09 | 8.40E-07 | 0.00068 | 0.15 | ENSMUSG | protein_cc | Tmem154 | transmem | 320782 | 3 | 84666192 | 84704575 | 1 | Mm.107370 |
| 4.74 | 6 | 4.56 | 5.65 | GFP | E47 | 1.23 | 10.98 | 9.10E-07 | 7.00E-04 | 0.59 | ENSMUSG | protein_cc | Sema7a | sema dom | 20361 | 9 | 57940112 | 57962865 | 1 | Mm.335187 |
| 1.4 | 3.15 | 1.13 | 3.34 | GFP | E47 | 1.82 | 10.97 | 9.30E-07 | 7.00E-04 | 0.33 | ENSMUSG | protein_cc | Fam43a | family with | 224093 | 16 | 30599723 | 30602797 | 1 | Mm.28244 |
| 2.84 | 3.99 | 2.56 | 3.79 | GFP | E47 | 1.16 | 10.79 | 1.10E-06 | 0.00079 | 0.78 | ENSMUSG | protein_cc | Nrp2 | neuropilin | 18187 | 1 | 62703285 | 62818695 | 1 | Mm.266341 |
| 3.27 | 4.33 | 3.66 | 4.56 | GFP | E47 | 1.03 | 10.55 | 1.30E-06 | 0.00093 | 0.49 | ENSMUSG | protein_cc | Rgcc | regulator c | 66214 | 14 | 79288756 | 79301645 | -1 | Mm.29811 |
| 5.61 | 4.64 | 5.61 | 4.52 | GFP | E47 | -1 | -10.5 | 1.40E-06 | 0.00095 | 0.63 | ENSMUSG | protein_cc | Dkk1 | dickkopf W | 13380 | 19 | 30545863 | 30549665 | -1 | Mm.214717 |
| 6.7 | 7.79 | 6.43 | 7.37 | GFP | E47 | 1.07 | 10.41 | 1.50E-06 | 0.001 | 0.61 | ENSMUSG | protein_cc | Man1a | mannosida | 17155 | 10 | 53904785 | 54076609 | -1 | Mm.117294 |
| 1.47 | 3.61 | 1.61 | 4.19 | GFP | E47 | 2.21 | 10.29 | 1.60E-06 | 0.0011 | 0.49 | ENSMUSG | protein_cc | Sema6d | sema dom | 214968 | 2 | 1.24E+08 | 1.25E+08 | 1 | Mm.330536 Mm.412083 |
| 2.73 | 3.96 | 2.82 | 3.92 | GFP | E47 | 1.21 | 10.27 | 1.70E-06 | 0.0011 | 0.66 | ENSMUSG | protein_cc | Gpx8 | glutathion | 69590 | 13 | 1.13E+08 | 1.13E+08 | -1 | Mm.12715 |
| 3.92 | 5.25 | 3.91 | 5.89 | GFP | E47 | 1.45 | 10.26 | 1.70E-06 | 0.0011 | 0.074 | ENSMUSG | protein_cc | Gimap7 | GTPase, IM | 231932 | 6 | 48718621 | 48724636 | 1 | Mm.30479 |
| -3.14 | 1.36 | -2.11 | 3.89 | GFP | E47 | 4.81 | 10.23 | 1.70E-06 | 0.0011 | 0.19 | ENSMUSG | protein_cc | Cbfa2t3 | CBFA2/RU | 12398 | 8 | 1.23E+08 | 1.23E+08 | -1 | Mm.194339 |
| 2.67 | 4.12 | 2.28 | 3.95 | GFP | E47 | 1.48 | 10.21 | 1.80E-06 | 0.0011 | 0.62 | ENSMUSG | protein_cc | Sprn | shadow of | 212518 | 7 | 1.4E+08 | 1.4E+08 | -1 | Mm.246858 |
| -0.85 | 2.54 | 0.34 | 3.05 | GFP | E47 | 3.24 | 10.02 | 2.10E-06 | 0.0013 | 0.41 | ENSMUSG | protein_cc | Il6ra | interleukin | 16194 | 3 | 89864059 | 89913196 | -1 | Mm.2856 |
| 6.26 | 7.1 | 6.17 | 7.31 | GFP | E47 | 0.9 | 9.89 | 2.30E-06 | 0.0014 | 0.24 | ENSMUSG | protein_cc | Rora | RAR-relate | 19883 | 9 | 68653786 | 69388246 | 1 | Mm.378450 Mm.391890 Mm.427266 Mm.4 |
| 7.53 | 8.09 | 7.33 | 8.1 | GFP | E47 | 0.6 | 9.84 | 2.40E-06 | 0.0014 | 0.2 | ENSMUSG | protein_cc | Gpr55 | G protein-c | 227326 | 1 | 85938318 | 85961007 | -1 | Mm.244158 Mm.396343 Mm.417775 |
| 3.24 | 4.32 | 3.36 | 4.32 | GFP | E47 | 1.06 | 9.81 | 2.50E-06 | 0.0014 | 0.67 | ENSMUSG | protein_cc | Lig4 | ligase IV, D | 319583 | 8 | 9969049 | 9977686 | -1 | Mm.80584 |
| 5.66 | 4.9 | 5.58 | 4.83 | GFP | E47 | -0.75 | -9.79 | 2.60E-06 | 0.0014 | 0.98 | ENSMUSG | protein_cc | Ak4 | adenylate | 11639 | 4 | 1.01E+08 | 1.01E+08 | 1 | Mm.42040 |
| 8.65 | 9.22 | 8.13 | 8.86 | GFP | E47 | 0.6 | 9.73 | 2.70E-06 | 0.0015 | 0.38 | ENSMUSG | protein_cc | Paxbp1 | PAX3 and P | 67367 | 16 | 91014037 | 91044543 | -1 | Mm.347 |
| 1 | 3.39 | 0.62 | 4.45 | GFP | E47 | 2.57 | 9.72 | 2.70E-06 | 0.0015 | 0.067 | ENSMUSG | protein_cc | Cd27 | CD27 antig | 21940 | 6 | 1.25E+08 | 1.25E+08 | -1 | Mm.367714 |
| 6.28 | 6.96 | 6.15 | 6.98 | GFP | E47 | 0.71 | 9.72 | 2.70E-06 | 0.0015 | 0.47 | ENSMUSG | protein_cc | Abhd17c | abhydrolas | 70178 | 7 | 84109356 | 84151893 | -1 | Mm.23914 |
| 3.53 | 4.87 | 3.04 | 4.81 | GFP | E47 | 1.4 | 9.62 | 3.00E-06 | 0.0016 | 0.34 | ENSMUSG | protein_cc | Mctp2 | multiple C2 | 244049 | 7 | 72077830 | 72306608 | -1 | Mm.217412 |
| 2.2 | 3.49 | 3.12 | 4.6 | GFP | E47 | 1.34 | 9.48 | 3.40E-06 | 0.0018 | 0.58 | ENSMUSG | protein_cc | Arl5c | ADP-ribosy | 217151 | 11 | 97989578 | 97996181 | -1 | Mm.277896 |
| 1.87 | 3.26 | 1.63 | 3.46 | GFP | E47 | 1.45 | 9.47 | 3.40E-06 | 0.0018 | 0.31 | ENSMUSG | protein_cc | Olfr60 | olfactory r | 18361 | 7 | 1.4E+08 | 1.4E+08 | -1 | Mm.222844 |
| 6.69 | 7.39 | 6.37 | 7.42 | GFP | E47 | 0.76 | 9.19 | 4.40E-06 | 0.0022 | 0.12 | ENSMUSG | protein_cc | Cerk | ceramide k | 223753 | 15 | 86139128 | 86186141 | -1 | Mm.222685 |
| 7.38 | 6.84 | 7.56 | 7.06 | GFP | E47 | -0.53 | -9.16 | 4.60E-06 | 0.0022 | 0.74 | ENSMUSG | protein_cc | Rilpl2 | Rab intera | 80291 | 5 | 1.24E+08 | 1.24E+08 | -1 | Mm.425280 |
| 3.48 | 1.82 | 3.44 | 2.01 | GFP | E47 | -1.6 | -9.13 | 4.70E-06 | 0.0023 | 0.61 | ENSMUSG | protein_cc | Slc7a11 | solute carr | 26570 | 3 | 49892526 | 50443614 | -1 | Mm.260988 |
| 11.75 | 11.47 | 11.76 | 11.49 | GFP | E47 | -0.28 | -8.88 | 6.00E-06 | 0.0028 | 0.89 | ENSMUSG | protein_cc | Aldoa | aldolase A, | 11674 | 7 | 1.27E+08 | 1.27E+08 | -1 | Mm.275831 |
| 3.11 | 5.34 | 3.21 | 6.27 | GFP | E47 | 2.37 | 8.85 | 6.20E-06 | 0.0029 | 0.3 | ENSMUSG | protein_cc | Id3 | inhibitor o | 15903 | 4 | 1.36E+08 | 1.36E+08 | 1 | Mm.110 |
| -1.77 | -4.85 | -1.04 | -3.37 | GFP | E47 | -2.93 | -8.8 | 6.50E-06 | 0.0029 | 0.34 | ENSMUSG | protein_cc | Vps37d | vacuolar p | 194309 | 5 | 1.35E+08 | 1.35E+08 | -1 | Mm.44356 |
| 3.24 | 4.29 | 3.34 | 4.79 | GFP | E47 | 1.12 | 8.79 | 6.60E-06 | 0.0029 | 0.22 | ENSMUSG | protein_cc | Tnf | tumor nec | 21926 | 17 | 35199381 | 35202007 | -1 | Mm.1293 |
| 5.64 | 6.21 | 5.9 | 6.68 | GFP | E47 | 0.62 | 8.72 | 7.00E-06 | 0.003 | 0.21 | ENSMUSG | protein_cc | Dhrs1 | dehydroge | 52585 | 14 | 55739020 | 55745690 | -1 | Mm.21623 |
| -1.77 | 1.73 | -1.94 | 2.86 | GFP | E47 | 3.69 | 8.47 | 9.00E-06 | 0.0036 | 0.33 | ENSMUSG | protein_cc | Wisp1 | WNT1 indu | 22402 | 15 | 66891320 | 66923201 | 1 | Mm.10222 |
| 7.49 | 8.02 | 7.44 | 7.84 | GFP | E47 | 0.51 | 8.46 | 9.10E-06 | 0.0036 | 0.41 | ENSMUSG | protein_cc | Lpar6 | lysophospl | 67168 | 14 | 73237895 | 73243294 | 1 | Mm.390681 |
| 4.73 | 5.76 | 4.36 | 5.69 | GFP | E47 | 1.08 | 8.46 | 9.10E-06 | 0.0036 | 0.43 | ENSMUSG | protein_cc | Errfi1 | ERBB recep | 74155 | 4 | 1.51E+08 | 1.51E+08 | 1 | Mm.318841 |
| 6.94 | 7.49 | 6.54 | 7.31 | GFP | E47 | 0.59 | 8.46 | 9.10E-06 | 0.0036 | 0.25 | ENSMUSG | protein_cc | Slc7a8 | solute carr | 50934 | 14 | 54722209 | 54781946 | -1 | Mm.276831 |
| 7.4 | 6.68 | 7.02 | 6.51 | GFP | E47 | -0.68 | -8.46 | 9.10E-06 | 0.0036 | 0.32 | ENSMUSG | protein_cc | Slc2a3 | solute carr | 20527 | 6 | 1.23E+08 | 1.23E+08 | -1 | Mm.395108 |
| 4.27 | 5.23 | 4.13 | 5.44 | GFP | E47 | 1.02 | 8.44 | 9.30E-06 | 0.0036 | 0.32 | ENSMUSG | protein_cc | Sla2 | Src-like ad | 77799 | 2 | 1.57E+08 | 1.57E+08 | [illegible] | Mm.29264 Mm.387161 |

| Con 24h | Con 72h | dKO 24h | dKO 72h | GFP_4h | GFP_16h | E47_4h | E47_16h | sample_1 | sample_2 | logFC | test_stat | p_value | q_value | Geneid | biotype | SYMBOL | GENENAM | ENTREZID | Chr | Start | End | Strand | UniGene |
|---|---|---|---|---|---|---|---|---|---|---|---|---|---|---|---|---|---|---|---|---|---|---|---|
| -3.04 | -0.6 | 2.15 | 3.65 | -1.78 | -1.75 | -3.27 | -3.79 | DN1_abVs | E47vsGFP | -12.97 | -7.91 | 2.20E-05 | 0.0021 | ENSMUSG( | protein_cc | Robo1 | roundabou | 19876 | 16 | 72308306 | 73046095 | 1 | Mm.310772 |
| -3.82 | -2 | -0.77 | -0.61 | -1.4 | -1.51 | -4.05 | -3.38 | DN1_abVs | E47vsGFP | -8.96 | -4.48 | 0.0031 | 0.048 | ENSMUSG( | protein_cc | Zfp532 | zinc finger | 328977 | 18 | 65580230 | 65689443 | 1 | Mm.286232 Mm.419081 |
| -3.82 | -3.96 | -2.17 | -1.03 | -1.76 | -2.45 | -4.08 | -4.15 | DN1_abVs | E47vsGFP | -8.58 | -5.11 | 0.00093 | 0.022 | ENSMUSG( | protein_cc | Shroom1 | shroom fa | 71774 | 11 | 53457205 | 53467766 | 1 | Mm.274370 |
| -0.14 | 2.22 | 3.28 | 5.01 | -5.09 | -2.54 | -4.85 | -4.97 | DN1_abVs | E47vsGFP | -8.41 | -9.01 | 4.20E-06 | 0.00071 | ENSMUSG( | TEC | Gm42992 | predicted g | NA | 5 | 1.32E+08 | 1.32E+08 | -1 | |
| 1.46 | 3.5 | 3.84 | 7.87 | 3.43 | 3.35 | 2.55 | 2.77 | DN3_abVs | E47vsGFP | -8.22 | -9.24 | 2.80E-05 | 0.0017 | ENSMUSG( | protein_cc | S1pr1 | sphingosin | 13609 | 3 | 1.16E+08 | 1.16E+08 | -1 | Mm.982 |
| -3.88 | -3.95 | -3.97 | -1.31 | -2.35 | -1.97 | -4.85 | -4.97 | DN3_abVs | E47vsGFP | -8.05 | -7.11 | 4.50E-05 | 0.0023 | ENSMUSG( | antisense | Hopxos | HOP home | NA | 5 | 77094532 | 77102303 | 1 | |
| -3.08 | -3.96 | -0.95 | -0.56 | -0.65 | -0.98 | -2.04 | -2.08 | DN3_abVs | E47vsGFP | -8.02 | -5.46 | 0.00097 | 0.018 | ENSMUSG( | protein_cc | Rnf208 | ring finger | 68846 | 2 | 25242255 | 25244262 | 1 | Mm.24777 Mm.490850 |
| 1.03 | 1.19 | 3.85 | 3.43 | -3.5 | -2.05 | -4.05 | -4.16 | DN3_abVs | E47vsGFP | -7.72 | -4.76 | 0.0021 | 0.029 | ENSMUSG( | processed | Gm15454 | predicted g | NA | 1 | 1.35E+08 | 1.35E+08 | 1 | |
| -3.82 | -3.17 | -3.75 | 0.48 | -3.14 | -3.13 | -4.85 | -4.97 | DN1_abVs | E47vsGFP | -7.26 | -4.49 | 0.0028 | 0.045 | ENSMUSG( | protein_cc | Pcdh9 | protocadh | 211712 | 14 | 93013410 | 93890679 | -1 | Mm.37126 Mm.458162 |
| 1.53 | 1.21 | 3.63 | 3.71 | -3.5 | -2.06 | -4.05 | -4.16 | DN1_abVs | E47vsGFP | -7.24 | -4.55 | 0.0031 | 0.048 | ENSMUSG( | processed | Gm15454 | predicted g | NA | 1 | 1.35E+08 | 1.35E+08 | 1 | |
| -3.88 | -1.14 | -2.01 | 1.01 | -3.49 | -3.21 | -4.85 | -4.97 | DN3_abVs | E47vsGFP | -7.13 | -4.79 | 0.0014 | 0.022 | ENSMUSG( | antisense | Gm19261 | predicted g | NA | 19 | 11516211 | 11516953 | -1 | |
| -2.29 | -1.1 | -0.39 | -0.77 | -2.19 | -1.36 | -4.06 | -4.18 | DN3_abVs | E47vsGFP | -6.92 | -5.28 | 0.001 | 0.018 | ENSMUSG( | processed | Gm4859 | predicted g | NA | 3 | 1.12E+08 | 1.12E+08 | 1 | |
| -1.06 | -1.07 | 0.97 | 0.73 | -3.13 | -2.89 | -4.85 | -4.18 | DN1_abVs | E47vsGFP | -6.84 | -5 | 0.001 | 0.024 | ENSMUSG( | transcribe | Rpl29-ps2 | ribosomal | NA | 13 | 4609334 | 4615025 | 1 | |
| 7.36 | 7.86 | 9.62 | 11.15 | 5.16 | 5.44 | 4.65 | 4.74 | DN1_abVs | E47vsGFP | -6.75 | -12.15 | 3.60E-06 | 0.00064 | ENSMUSG( | protein_cc | Lmo4 | LIM domai | 16911 | 3 | 1.44E+08 | 1.44E+08 | -1 | Mm.29187 |
| -1.47 | -1.76 | 0.36 | 0.35 | -2.88 | -0.77 | -2.29 | -4.17 | DN1_abVs | E47vsGFP | -6.74 | -4.18 | 0.0053 | 0.066 | ENSMUSG( | TEC | Gm42908 | predicted g | NA | 5 | 1.23E+08 | 1.23E+08 | 1 | |
| -3 | -1.39 | -0.44 | 0.38 | 5.43 | 5.61 | 4.46 | 4.28 | DN1_abVs | E47vsGFP | -6.64 | -7.66 | 4.10E-06 | 0.00069 | ENSMUSG( | protein_cc | Pkib | protein kin | 18768 | 10 | 57631981 | 57741112 | 1 | Mm.262135 |
| -1.14 | -0.84 | 1.12 | 0.36 | -3.13 | -2.89 | -4.85 | -4.18 | DN3_abVs | E47vsGFP | -6.47 | -5.66 | 0.00021 | 0.0066 | ENSMUSG( | transcribe | Rpl29-ps2 | ribosomal | NA | 13 | 4609334 | 4615025 | 1 | |
| -3.88 | -0.53 | -1.4 | 1.96 | 3.51 | 3.82 | 3 | 2.88 | DN3_abVs | E47vsGFP | -6.41 | -9.25 | 3.70E-06 | 0.00044 | ENSMUSG( | protein_cc | Tmem40 | transmeml | 94346 | 6 | 1.16E+08 | 1.16E+08 | -1 | Mm.29739 |
| -3.82 | -2.37 | -1.43 | -1.49 | -3.73 | -2.17 | -4.04 | -4.97 | DN1_abVs | E47vsGFP | -6.37 | -4.18 | 0.005 | 0.064 | ENSMUSG( | processed | Gm12058 | predicted g | NA | 11 | 22786811 | 22787247 | -1 | |
| 1.95 | 2.76 | 3.5 | 6.16 | 3.77 | 3.7 | 2.95 | 3.21 | DN3_abVs | E47vsGFP | -6.25 | -9.44 | 5.30E-06 | 0.00056 | ENSMUSG( | protein_cc | Deptor | DEP domai | 97998 | 15 | 55112317 | 55259271 | 1 | Mm.295397 Mm.393497 |
| -2.73 | -2.37 | -1.07 | -0.46 | -3.49 | -2.87 | -4.04 | -4.97 | DN1_abVs | E47vsGFP | -6.2 | -4.07 | 0.0066 | 0.06 | ENSMUSG( | protein_cc | Muc2 | mucin 2 | NA | 7 | 1.42E+08 | 1.42E+08 | 1 | |
| -3.88 | -0.89 | -2.01 | 0.58 | 1.55 | 2.16 | 0.67 | 0.26 | DN3_abVs | E47vsGFP | -6.12 | -4.63 | 0.0016 | 0.024 | ENSMUSG( | protein_cc | Arhgdig | Rho GDP d | 14570 | 17 | 26199183 | 26207786 | -1 | Mm.1383 |
| -1.14 | 0.14 | 0.93 | 2.98 | 7.74 | 7.88 | 7.33 | 7.13 | DN3_abVs | E47vsGFP | -6.06 | -9.42 | 1.30E-05 | 0.001 | ENSMUSG( | protein_cc | Lgmn | legumain | 19141 | 12 | 1.02E+08 | 1.02E+08 | -1 | Mm.17185 |
| -3.88 | -2.37 | -3.17 | -0.48 | -4.25 | -2.18 | -4.85 | -4.97 | DN3_abVs | E47vsGFP | -5.98 | -3.78 | 0.0085 | 0.07 | ENSMUSG( | protein_cc | Lrrc6 | leucine ricl | 54562 | 15 | 66379858 | 66500910 | -1 | Mm.244890 |
| 0.76 | 1.92 | 2.26 | 5.38 | 5.16 | 5.42 | 5.04 | 4.61 | DN3_abVs | E47vsGFP | -5.9 | -12.23 | 2.20E-06 | 0.00031 | ENSMUSG( | lincRNA | Gm14005 | predicted g | NA | 2 | 1.28E+08 | 1.29E+08 | -1 | |
| -1.31 | -0.38 | 1.32 | -0.36 | -2.09 | -1.93 | -4.85 | -2.41 | DN3_abVs | E47vsGFP | -5.89 | -5.65 | 0.00023 | 0.007 | ENSMUSG( | protein_cc | Klhdc9 | kelch dom | 68874 | 1 | 1.71E+08 | 1.71E+08 | -1 | Mm.10756 Mm.330288 |
| -3.83 | -3.96 | -1.19 | -3.05 | 4.89 | 5.12 | 4.09 | 3.57 | DN1_abVs | E47vsGFP | -5.88 | -4.79 | 0.0017 | 0.032 | ENSMUSG( | protein_cc | Ly6g | lymphocyt | 546644 | 15 | 75155240 | 75159126 | 1 | |
| -3.88 | -1.98 | -1.4 | -1.41 | -1.11 | -1.07 | -2.66 | -2.33 | DN3_abVs | E47vsGFP | -5.85 | -4.03 | 0.0061 | 0.057 | ENSMUSG( | protein_cc | Nos3 | nitric oxide | 18127 | 5 | 24364810 | 24384474 | 1 | Mm.258415 |
| -0.99 | -2.03 | -0.61 | -0.31 | -2.1 | -3.41 | -4.85 | -4.16 | DN3_abVs | E47vsGFP | -5.61 | -4.07 | 0.005 | 0.05 | ENSMUSG( | processed | Gm3724 | predicted g | NA | 5 | 24118316 | 24118664 | -1 | |
| 0.55 | 0.72 | 1.82 | 3.07 | 0.91 | 0.88 | 0 | -0.12 | DN1_abVs | E47vsGFP | -5.53 | -6.87 | 2.20E-05 | 0.0021 | ENSMUSG( | protein_cc | Syngr1 | synaptogy | 20972 | 15 | 80091334 | 80119501 | 1 | Mm.230301 Mm.396243 |
| -1.06 | -0.52 | 1.33 | 1.2 | 4.6 | 4.53 | 4.3 | 3.43 | DN1_abVs | E47vsGFP | -5.5 | -6.5 | 0.00024 | 0.0091 | ENSMUSG( | protein_cc | Slc17a8 | solute carr | 216227 | 10 | 89574020 | 89621253 | -1 | Mm.233921 |
| -1.88 | -1.39 | -1.01 | 1.64 | 1.58 | 2.58 | 1.31 | 1.36 | DN1_abVs | E47vsGFP | -5.4 | -5.34 | 0.0012 | 0.027 | ENSMUSG( | antisense | Gm14718 | predicted g | NA | X | 57200643 | 57204378 | 1 | |
| -0.1 | 1.51 | 2.29 | 2.72 | 3.9 | 3.65 | 3.3 | 2.69 | DN1_abVs | E47vsGFP | -5.15 | -5.92 | 0.00039 | 0.013 | ENSMUSG( | protein_cc | Gcnt4 | glucosamir | 218476 | 13 | 96924689 | 96950906 | 1 | Mm.483490 |
| -3.88 | -1.65 | -0.8 | -1.26 | 1.62 | 1.8 | 1.19 | 0.78 | DN3_abVs | E47vsGFP | -4.93 | -5.83 | 4.00E-04 | 0.01 | ENSMUSG( | protein_cc | Zfp503 | zinc finger | 218820 | 14 | 21983959 | 21989601 | -1 | Mm.292401 |
| -1.56 | -1.37 | -0.07 | -0.05 | -4.31 | -3.42 | -4.85 | -4.97 | DN3_abVs | E47vsGFP | -4.89 | -4.26 | 0.0046 | 0.047 | ENSMUSG( | processed | Gm13777 | predicted g | NA | 2 | 91030778 | 91031122 | -1 | |
| 1.1 | 3.99 | 2.84 | 5.91 | 0.79 | -0.14 | -0.15 | -0.41 | DN3_abVs | E47vsGFP | -4.87 | -6.77 | 0.00012 | 0.0046 | ENSMUSG( | protein_cc | Podxl | podocalyxi | 27205 | 6 | 31519488 | 31563981 | -1 | Mm.89918 |
| -2.24 | -1.76 | -1.43 | -0.22 | 0.04 | -0.38 | -1.31 | -1.43 | DN1_abVs | E47vsGFP | -4.74 | -4.17 | 0.0048 | 0.062 | ENSMUSG( | protein_cc | Aldh1l2 | aldehyde c | 216188 | 10 | 83487450 | 83534140 | -1 | Mm.263138 |
| 2.22 | 2.9 | 3.2 | 4.27 | -1.09 | 0.28 | -1.42 | -1.74 | DN1_abVs | E47vsGFP | -4.7 | -4.36 | 0.0034 | 0.051 | ENSMUSG( | TEC | Gm48207 | predicted g | NA | 10 | 1.06E+08 | 1.06E+08 | -1 | |
| 4.36 | 5.82 | 5.33 | 8.44 | 9.07 | 9.14 | 8.6 | 8.56 | DN3_abVs | E47vsGFP | -4.65 | -9.35 | 5.50E-06 | 0.00057 | ENSMUSG( | protein_cc | Osm | oncostatin | 18413 | 11 | 4236420 | 4241026 | 1 | Mm.131422 |
| 6.05 | 7.59 | 7.87 | 9.29 | 8.35 | 8.45 | 7.7 | 8.01 | DN3_abVs | E47vsGFP | -4.63 | -12.57 | 3.40E-07 | 0.00012 | ENSMUSG( | protein_cc | Klf6 | Kruppel-lik | 23849 | 13 | 5861482 | 5870394 | 1 | Mm.275036 |
| -3.82 | -3.96 | -3.75 | -1.47 | -0.14 | -0.39 | -0.6 | -1.98 | DN1_abVs | E47vsGFP | -4.61 | -4.49 | 0.0029 | 0.046 | ENSMUSG( | protein_cc | Moxd1 | monooxyg | 59012 | 10 | 24223517 | 24302790 | 1 | Mm.285934 |
| -0.17 | 0.53 | 0.4 | 2.62 | -1.85 | 0.29 | -1.97 | -1.51 | DN3_abVs | E47vsGFP | -4.58 | -4.05 | 0.0062 | 0.058 | ENSMUSG( | TEC | Gm44835 | predicted g | NA | 7 | 75560614 | 75563794 | 1 | |
| 6.39 | 8.43 | 8.06 | 10.83 | 13.07 | 13.08 | 12.91 | 12.74 | DN3_abVs | E47vsGFP | -4.57 | -15.35 | 2.80E-05 | 0.0017 | ENSMUSG( | protein_cc | Vim | vimentin | 22352 | 2 | 13573927 | 13582826 | 1 | Mm.268000 |
| 3.92 | 3.96 | 4.32 | 6.89 | 4.14 | 4.58 | 3.75 | 3.78 | DN1_abVs | E47vsGFP | -4.52 | -6.17 | 0.00038 | 0.013 | ENSMUSG( | protein_cc | Avpi1 | arginine va | 69534 | 19 | 42123273 | 42129059 | -1 | Mm.30060 |
| 1.45 | 1.17 | 2.66 | 2.67 | -3.5 | -4.5 | -4.85 | -4.97 | DN1_abVs | E47vsGFP | -4.51 | -5.9 | 0.00021 | 0.0083 | ENSMUSG( | processed | Gm6344 | predicted g | NA | 13 | 55085777 | 55086259 | 1 | |
| 7.34 | 7.07 | 8.42 | 9.3 | 5.16 | 5.44 | 4.65 | 4.74 | DN3_abVs | E47vsGFP | -4.51 | -8.42 | 6.20E-06 | 0.00062 | ENSMUSG( | protein_cc | Lmo4 | LIM domai | 16911 | 3 | 1.44E+08 | 1.44E+08 | -1 | Mm.29187 |
| 8.3 | 7.83 | 9.65 | 9.51 | 2.94 | 3.75 | 2.64 | 2.67 | DN3_abVs | E47vsGFP | -4.43 | -9.18 | 1.10E-06 | 2.00E-04 | ENSMUSG( | processed | Rps13-ps1 | ribosomal | NA | 8 | 87047689 | 87048226 | 1 | |
| 2.19 | 2.43 | 3.19 | 3.56 | -0.56 | 0 | -1.41 | -1.42 | DN1_abVs | E47vsGFP | -4.41 | -4.08 | 0.005 | 0.064 | ENSMUSG( | TEC | Gm45407 | predicted g | NA | 8 | 77368841 | 77372516 | -1 | |
| 1.38 | 1.06 | 2.7 | 2.27 | -3.5 | -4.5 | -4.85 | -4.97 | DN3_abVs | E47vsGFP | -4.35 | -5.16 | 0.00068 | 0.014 | ENSMUSG( | processed | Gm6344 | predicted g | NA | 13 | 55085777 | 55086259 | 1 | |
| 8.11 | 7.99 | 9.39 | 9.54 | 2.94 | 3.74 | 2.64 | 2.67 | DN1_abVs | E47vsGFP | -4.22 | -8.58 | 2.30E-06 | 0.00049 | ENSMUSG( | processed | Rps13-ps1 | ribosomal | NA | 8 | 87047689 | 87048226 | 1 | |
| -3.88 | -3.95 | -3.97 | -1.7 | -1.5 | -0.65 | -1.95 | -2.23 | DN3_abVs | E47vsGFP | -4.2 | -4.02 | 0.0063 | 0.058 | ENSMUSG( | protein_cc | Rorb | RAR-relate | 225998 | 19 | 18930605 | 19111196 | -1 | Mm.234641 |
| 6.16 | 7.37 | 7.94 | 8.68 | 8.35 | 8.45 | 7.7 | 8.01 | DN1_abVs | E47vsGFP | -4.19 | -8.81 | 6.00E-06 | 0.00089 | ENSMUSG( | protein_cc | Klf6 | Kruppel-lik | 23849 | 13 | 5861482 | 5870394 | 1 | Mm.275036 |
| 1.07 | 3.01 | 1.7 | 4.46 | 1.74 | 1.77 | 0.99 | 0.48 | DN3_abVs | E47vsGFP | -4.13 | -5.42 | 0.00039 | 0.01 | ENSMUSG( | protein_cc | Mgarp | mitochond | 67749 | 3 | 51388412 | 51396738 | -1 | Mm.273339 |
| -3.82 | -3.96 | -1.19 | -3.85 | -2.91 | -2.54 | -4.85 | -1.97 | DN1_abVs | E47vsGFP | -4.1 | -5.56 | 7.00E-04 | 0.019 | ENSMUSG( | lincRNA | A630023P | RIKEN cDN | NA | 5 | 1.11E+08 | 1.11E+08 | 1 | |
| 2.94 | 3.12 | 3.35 | 5.75 | 7.11 | 6.74 | 6.78 | 6.06 | DN3_abVs | E47vsGFP | -4.05 | -6.05 | 0.00066 | 0.014 | ENSMUSG( | protein_cc | Irf4 | interferon | 16364 | 13 | 30749226 | 30766976 | 1 | Mm.4677 |
| 0.12 | 1.25 | 1.37 | 2.19 | 2.69 | 2.8 | 2.19 | 1.53 | DN1_abVs | E47vsGFP | -3.96 | -6.25 | 7.10E-05 | 0.0045 | ENSMUSG( | protein_cc | Pgap1 | post-GPI a | 241062 | 1 | 54472994 | 54557684 | -1 | Mm.103539 |
| -3.88 | -3.97 | -3.97 | -1.32 | 1.83 | 1.58 | 0.87 | 1.17 | DN3_abVs | E47vsGFP | -3.94 | -5.39 | 0.00064 | 0.014 | ENSMUSG( | lincRNA | Gm35037 | predicted g | NA | 7 | 3451357 | 3475020 | 1 | |
| 2.04 | 2.03 | 2.37 | 3.86 | 7.03 | 7.22 | 6.05 | 6.55 | DN3_abVs | E47vsGFP | -3.8 | -5.44 | 0.00033 | 0.009 | ENSMUSG( | protein_cc | Pou2f2 | POU doma | 18987 | 7 | 25087344 | 25179726 | -1 | Mm.76683 |
| 1.76 | 3.75 | 2.13 | 5.66 | 5.66 | 5.61 | 4.91 | 4.83 | DN3_abVs | E47vsGFP | -3.8 | -4.63 | 0.0019 | 0.026 | ENSMUSG( | protein_cc | Ak4 | adenylate | 11639 | 4 | 1.01E+08 | 1.01E+08 | 1 | Mm.42040 |
| 0.87 | 2.15 | 2.1 | 3.4 | 3.24 | 3.26 | 2.64 | 2.62 | DN3_abVs | E47vsGFP | -3.71 | -4.71 | 0.0023 | 0.031 | ENSMUSG( | protein_cc | Spns3 | spinster h | 77577 | 11 | 72494919 | 72550506 | -1 | Mm.46185 |
| 6.76 | 7.4 | 7.99 | 9 | 3.09 | 2.83 | 2.72 | 2.38 | DN1_abVs | E47vsGFP | -3.66 | -7.98 | 7.60E-05 | 0.0046 | ENSMUSG( | protein_cc | Tox | thymocyte | 252838 | 4 | 6686353 | 6991557 | -1 | Mm.435828 Mm.488241 |
| -0.71 | -2.03 | -0.43 | -0.52 | -3.49 | -4.46 | -4.85 | -4.97 | DN3_abVs | E47vsGFP | -3.66 | -4.79 | 0.0011 | 0.019 | ENSMUSG( | processed | Gm10705 | predicted g | 1E+08 | 3 | 87207545 | 87208005 | -1 | |
| -3.88 | -3.96 | -2.38 | -3.64 | -4.25 | -3.76 | -4.85 | -4.96 | DN3_abVs | E47vsGFP | -3.62 | -3.8 | 0.0085 | 0.07 | ENSMUSG( | processed | Gm44180 | predicted g | NA | 6 | 1.16E+08 | 1.16E+08 | -1 | |
| 1.12 | 1.87 | 1.33 | 2.7 | 6.78 | 7.08 | 5.75 | 5.58 | DN1_abVs | E47vsGFP | -3.57 | -6.99 | 0.00014 | 0.0067 | ENSMUSG( | protein_cc | Phlda1 | pleckstrin | 21664 | 10 | 1.12E+08 | 1.12E+08 | 1 | Mm.3117 |
| 6.24 | 6.93 | 6.7 | 9.11 | 1.35 | 1.47 | 0.76 | 1.15 | DN1_abVs | E47vsGFP | -3.56 | -6.14 | 0.00041 | 0.013 | ENSMUSG( | protein_cc | Tubb2b | tubulin, be | 73710 | 13 | 34126748 | 34130466 | -1 | Mm.379227 Mm.472121 |
| 3.78 | 5.01 | 5.45 | 5.66 | 4.98 | 5.09 | 4.6 | 4.27 | DN3_abVs | E47vsGFP | -3.53 | -6.12 | 9.60E-05 | 0.0039 | ENSMUSG( | protein_cc | Tsc22d3 | TSC22 dor | 14605 | X | 1.41E+08 | 1.41E+08 | -1 | Mm.22216 |
| 2.96 | 4.56 | 4.31 | 5.53 | 4.62 | 4.69 | 4.35 | 3.82 | DN3_abVs | E47vsGFP | -3.47 | -5.89 | 0.00018 | 0.0061 | ENSMUSG( | protein_cc | Ccr7 | chemokine | 12775 | 11 | 99144196 | 99155077 | -1 | Mm.2932 |
| 4.02 | 5.26 | 4.27 | 7.22 | 7.41 | 7.03 | 6.68 | 6.53 | DN3_abVs | E47vsGFP | -3.45 | -5.24 | 0.00077 | 0.015 | ENSMUSG( | protein_cc | Slc2a3 | solute carr | 20527 | 6 | 1.23E+08 | 1.23E+08 | -1 | Mm.395108 |
| 5.18 | 5.16 | 6.44 | 6.79 | 6.35 | 6.28 | 6.12 | 5.97 | DN1_abVs | E47vsGFP | -3.43 | -10.05 | 4.50E-05 | 0.0034 | ENSMUSG( | protein_cc | Chn2 | chimerin 2 | 69993 | 6 | 54039554 | 54301810 | 1 | Mm.321147 Mm.448157 |
| 0.81 | 1.57 | 1.24 | 3.03 | 6.64 | 6.92 | 6.11 | 5.97 | DN1_abVs | E47vsGFP | -3.37 | -5.86 | 0.00016 | 0.007 | ENSMUSG( | protein_cc | Ckb | creatine ki | 12709 | 12 | 1.12E+08 | 1.12E+08 | -1 | Mm.16831 |
| 2.58 | 3.99 | 2.85 | 5.81 | 6.13 | 6.67 | 5.84 | 5.73 | DN3_abVs | E47vsGFP | -3.34 | -6.17 | 5.40E-05 | 0.0027 | ENSMUSG( | protein_cc | Klk8 | kallikrein r | 259277 | 7 | 43797577 | 43803826 | 1 | Mm.458023 |
| 0.23 | 0.9 | 0.89 | 2.17 | 0.37 | -0.2 | -0.77 | -0.45 | DN1_abVs | E47vsGFP | -3.31 | -4.41 | 0.0037 | 0.053 | ENSMUSG( | protein_cc | Hist1h2be | histone clu | 319179 | 13 | 23583670 | 23621124 | -1 | Mm.440391 |
| 0.29 | 1.8 | 1.85 | 2.63 | 2.38 | 2.31 | 1.99 | 1.82 | DN3_abVs | E47vsGFP | -3.28 | -6.18 | 0.00022 | 0.0067 | ENSMUSG( | protein_cc | Ube2e2 | ubiquitin-c | 218793 | 14 | 18573575 | 18894267 | -1 | Mm.23551 |
| 3.97 | 3.42 | 4.71 | 5.09 | 6.17 | 6.56 | 5.99 | 5.9 | DN1_abVs | E47vsGFP | -3.26 | -6.87 | 0.00013 | 0.0062 | ENSMUSG( | protein_cc | Prdx4 | peroxiredc | 53381 | X | 1.55E+08 | 1.55E+08 | -1 | Mm.247542 |
| 2.56 | 3.43 | 3.76 | 3.7 | 2.69 | 2.8 | 2.19 | 1.53 | DN3_abVs | E47vsGFP | -3.24 | -5.85 | 0.00018 | 0.0059 | ENSMUSG( | protein_cc | Pgap1 | post-GPI a | 241062 | 1 | 54472994 | 54557684 | -1 | Mm.103539 |
| 1.64 | 2.5 | 1.82 | 4.41 | 3.87 | 3.71 | 3.5 | 2.95 | DN3_abVs | E47vsGFP | -3.21 | -5 | 0.0011 | 0.019 | ENSMUSG( | protein_cc | Cpe | carboxype | 12876 | 8 | 64592542 | 64693054 | -1 | Mm.31395 Mm.470333 |
| 0.68 | 3 | 1.19 | 4.41 | 2.05 | 1.71 | 1.57 | 0.92 | DN3_abVs | E47vsGFP | -3.19 | -4.2 | 0.0046 | 0.048 | ENSMUSG( | protein_cc | Muc13 | mucin 13, | 17063 | 16 | 33794037 | 33819934 | 1 | Mm.3177 |
| 5.27 | 4.88 | 5.34 | 6.92 | 5.85 | 6.21 | 5.45 | 5.56 | DN1_abVs | E47vsGFP | -3.17 | -5.67 | 0.00055 | 0.016 | ENSMUSG( | protein_cc | Smco4 | single-pass | 170748 | 9 | 15493432 | 15545260 | 1 | Mm.86572 |

| PeakID (cn | Chr | Start | End | Strand | Peak Score | Focus Rati | Annotation | Detailed A | Distance t | Nearest Pr | Entrez ID | Nearest U | Nearest Re | Nearest E | Gene Nam | Gene Alias | Gene Desc | Gene Type | Conc | Conc_grou | Conc_grou | Fold | p.value | FDR |
|---|---|---|---|---|---|---|---|---|---|---|---|---|---|---|---|---|---|---|---|---|---|---|---|---|
| 1 | chr15 | 73118034 | 73118184 | + | 0 | NA | intron (NM | intron (NM | 27697 | NM_05306 | 93696 | Mm.23095 | NM_05306 | ENSMUSG | Chrac1 | 2410152E0 | chromatin | protein-co | 7.61 | 8.44 | 5.43 | 3 | 2.00E-37 | 1.19E-32 |
| 8 | chr2 | 1.04E+08 | 1.04E+08 | + | 0 | NA | intron (NM | intron (NM | 78714 | NM_15312 | 98956 | Mm.28800 | NM_15312 | ENSMUSG | Nat10 | AI429152 | N-acetyltra | protein-co | 8.52 | 9.16 | 7.36 | 1.8 | 8.31E-27 | 6.17E-23 |
| 11 | chr6 | 1.43E+08 | 1.43E+08 | + | 0 | NA | intron (NM | L1MA5\|LIN | 46775 | NM_01137 | 20449 | Mm.26083 | NM_01137 | ENSMUSG | St8sia1 | 9330109E0 | ST8 alpha- | protein-co | 8.5 | 9.1 | 7.44 | 1.66 | 1.21E-24 | 6.55E-21 |
| 13 | chr6 | 1.16E+08 | 1.16E+08 | + | 0 | NA | intron (NM | intron (NM | 5313 | NM_02637 | 67784 | Mm.3085 | NM_02637 | ENSMUSG | Plxnd1 | 6230425C2 | plexin D1 | protein-co | 7.28 | 8.04 | 5.58 | 2.46 | 8.26E-24 | 3.77E-20 |
| 18 | chr2 | 1.66E+08 | 1.66E+08 | + | 0 | NA | intron (NM | intron (NM | 64762 | NM_00125 | 228880 | Mm.22759 | NM_02723 | ENSMUSG | Zmynd8 | 1110013E2 | zinc finger | protein-co | 7.23 | 7.97 | 5.66 | 2.31 | 3.50E-21 | 1.15E-17 |
| 21 | chr6 | 1.09E+08 | 1.09E+08 | + | 0 | NA | intron (NM | intron (NM | 7554 | NM_13867 | 192193 | Mm.21596 | NM_13867 | ENSMUSG | Edem1 | A130059K | ER degrad | protein-co | 7.92 | 8.57 | 6.68 | 1.89 | 5.08E-20 | 1.43E-16 |
| 25 | chr3 | 1.33E+08 | 1.33E+08 | + | 0 | NA | intron (NM | intron (NM | 29451 | NM_00129 | 74776 | Mm.21030 | NM_14614 | ENSMUSG | Ppa2 | 1110013G | pyrophosp | protein-co | 6.21 | 7.1 | 3.48 | 3.62 | 3.77E-19 | 8.96E-16 |
| 27 | chr13 | 91541065 | 91541215 | + | 0 | NA | intron (NM | intron (NM | 80043 | NM_02427 | 66970 | Mm.34309 | NM_02418 | ENSMUSG | Ssbp2 | 1500004K0 | single-stra | protein-co | 8 | 8.61 | 6.91 | 1.7 | 1.15E-18 | 2.53E-15 |
| 34 | chr19 | 41039265 | 41039415 | + | 0 | NA | intron (NM | intron (NM | 10065 | NM_00934 | 21673 | Mm.25620 | NM_00934 | ENSMUSG | Dntt | Tdt | deoxynucle | protein-co | 6.21 | 7.05 | 3.92 | 3.14 | 3.47E-17 | 6.06E-14 |
| 38 | chr2 | 1.59E+08 | 1.59E+08 | + | 0 | NA | intron (NM | intron (NM | -48695 | NM_02797 | 71878 | Mm.85162 | NM_02797 | ENSMUSG | Fam83d | 2310007D0 | family with | protein-co | 8.64 | 9.18 | 7.76 | 1.42 | 1.42E-16 | 2.22E-13 |
| 39 | chr17 | 57194239 | 57194389 | + | 0 | NA | promoter- | promoter- | -133 | NM_01941 | 50930 | Mm.48336 | NM_01941 | ENSMUSG | Tnfsf14 | HVEM-L\|H | tumor nec | protein-co | 7.78 | 8.42 | 6.61 | 1.81 | 1.66E-16 | 2.53E-13 |
| 40 | chr17 | 47542178 | 47542328 | + | 0 | NA | intron (NM | intron (NM | 37202 | NM_00108 | 12445 | Mm.27291 | NM_00763 | ENSMUSG | Ccnd3 | 9230106B0 | cyclin D3 | protein-co | 6.26 | 7.09 | 4.11 | 2.98 | 1.75E-16 | 2.60E-13 |
| 43 | chr16 | 10668544 | 10668694 | + | 0 | NA | intron (NM | intron (NM | 116917 | NM_00127 | 12703 | Mm.130 | NM_00989 | ENSMUSG | Socs1 | Cish1\|Cish | suppressor | protein-co | 6.23 | 7.06 | 4.06 | 2.99 | 2.82E-16 | 3.89E-13 |
| 45 | chr12 | 73710961 | 73711111 | + | 0 | NA | intron (NM | intron (NM | 126240 | NM_00113 | 18755 | Mm.34167 | NM_00885 | ENSMUSG | Prkch | Pkch | protein kir | protein-co | 7.39 | 8.03 | 6.2 | 1.83 | 5.18E-16 | 6.83E-13 |
| 46 | chr19 | 32168208 | 32168358 | + | 0 | NA | intron (NM | RMER15\|L | 42730 | NM_00116 | 208449 | Mm.32981 | NM_14479 | ENSMUSG | Sgms1 | 9530058O | sphingomy | protein-co | 6.52 | 7.28 | 4.77 | 2.52 | 5.63E-16 | 7.20E-13 |
| 48 | chr2 | 1.59E+08 | 1.59E+08 | + | 0 | NA | intron (NM | intron (NM | -44638 | NM_02797 | 71878 | Mm.85162 | NM_02797 | ENSMUSG | Fam83d | 2310007D0 | family with | protein-co | 7.56 | 8.19 | 6.42 | 1.77 | 5.92E-16 | 7.32E-13 |
| 49 | chr10 | 60389655 | 60389805 | + | 0 | NA | intron (NR | intron (NR | -9996 | NM_00116 | 1E+08 | Mm.44413 | NM_00116 | ENSMUSG | Gm17455 | - | predicted | protein-co | 7.45 | 8.08 | 6.31 | 1.77 | 6.18E-16 | 7.49E-13 |
| 50 | chr2 | 1.02E+08 | 1.02E+08 | + | 0 | NA | intron (NM | MTEb\|LTR | 54512 | NM_00942 | 22034 | Mm.29272 | NM_00942 | ENSMUSG | Traf6 | 2310003F1 | TNF recept | protein-co | 8.31 | 8.84 | 7.46 | 1.38 | 7.29E-16 | 8.66E-13 |
| 52 | chr16 | 10788471 | 10788621 | + | 0 | NA | exon (NM_ | exon (NM_ | 109 | NM_01369 | 21959 | Mm.20679 | NM_01369 | ENSMUSG | Tnp2 | TP2\|Tp-2 | transition | protein-co | 6.85 | 7.56 | 5.4 | 2.16 | 1.55E-15 | 1.77E-12 |
| 53 | chr9 | 63737570 | 63737720 | + | 0 | NA | intron (NM | intron (NM | 20349 | NM_01676 | 17127 | Mm.7320 | NM_01676 | ENSMUSG | Smad3 | AU022421 | SMAD fam | protein-co | 7.41 | 8.04 | 6.28 | 1.76 | 1.65E-15 | 1.85E-12 |
| 56 | chr11 | 98000439 | 98000589 | + | 0 | NA | TTS (NM_1 | TTS (NM_1 | -4341 | NM_20723 | 217151 | Mm.27789 | NM_20723 | ENSMUSG | Arl5c | Arl12\|BC0 | ADP-ribosy | protein-co | 6.03 | 6.88 | 3.73 | 3.15 | 1.76E-15 | 1.87E-12 |
| 55 | chr7 | 1.26E+08 | 1.26E+08 | + | 0 | NA | intron (NM | intron (NM | 27081 | NM_02188 | 60504 | Mm.15564 | NM_02188 | ENSMUSG | Il21r | NILR | interleukir | protein-co | 7.74 | 8.36 | 6.64 | 1.72 | 1.74E-15 | 1.87E-12 |
| 59 | chr15 | 66789167 | 66789317 | + | 0 | NA | intron (NM | intron (NM | 42587 | NM_00919 | 20491 | Mm.7601 | NM_00919 | ENSMUSG | Sla | Slap\|Slap- | src-like ad | protein-co | 7.21 | 7.86 | 6 | 1.86 | 5.03E-15 | 5.06E-12 |
| 60 | chr10 | 1.21E+08 | 1.21E+08 | + | 0 | NA | intron (NM | intron (NM | 28753 | NM_02905 | 74694 | Mm.23383 | NM_02905 | ENSMUSG | Tbc1d30 | 4930505D0 | TBC1 dom | protein-co | 6.2 | 7 | 4.23 | 2.77 | 6.73E-15 | 6.66E-12 |
| 61 | chr13 | 51754170 | 51754320 | + | 0 | NA | intron (NM | intron (NM | 39502 | NM_00128 | 20354 | Mm.33903 | NM_01366 | ENSMUSG | Sema4d | CD100\|Se | sema dom | protein-co | 6.96 | 7.64 | 5.64 | 2 | 9.05E-15 | 8.81E-12 |
| 63 | chr8 | 11503991 | 11504141 | + | 0 | NA | intron (NM | intron (NM | 6498 | NM_02699 | 69225 | Mm.64911 | NM_02699 | ENSMUSG | Naxd | 0710008K0 | NAD(P)HX | protein-co | 6.37 | 7.14 | 4.6 | 2.54 | 9.83E-15 | 9.26E-12 |
| 65 | chr14 | 19962021 | 19962171 | + | 0 | NA | intron (NM | ID4\|SINE\| | 14597 | NM_00128 | 14702 | Mm.41737 | NM_01031 | ENSMUSG | Gng2 | - | guanine nu | protein-co | 6.39 | 7.16 | 4.65 | 2.52 | 1.02E-14 | 9.33E-12 |
| 64 | chr2 | 1.04E+08 | 1.04E+08 | + | 0 | NA | intron (NM | intron (NM | 76027 | NM_15312 | 98956 | Mm.28800 | NM_15312 | ENSMUSG | Nat10 | AI429152 | N-acetyltra | protein-co | 7.29 | 7.92 | 6.13 | 1.79 | 1.02E-14 | 9.33E-12 |
| 66 | chr2 | 1.32E+08 | 1.32E+08 | + | 0 | NA | intron (NM | intron (NM | 8853 | NM_17544 | 215653 | Mm.20473 | NM_17544 | ENSMUSG | Rassf2 | 3830431H0 | Ras associ | protein-co | 6.94 | 7.62 | 5.61 | 2 | 1.25E-14 | 1.12E-11 |
| 68 | chr8 | 25070421 | 25070571 | + | 0 | NA | intron (NM | intron (NM | 31315 | NM_03125 | 83436 | Mm.26112 | NM_03125 | ENSMUSG | Plekha2 | 6430512N | pleckstrin | protein-co | 7.26 | 7.92 | 6.02 | 1.9 | 1.46E-14 | 1.27E-11 |
| 69 | chr12 | 85624744 | 85624894 | + | 0 | NA | intron (NM | intron (NM | 24801 | NM_00120 | 81703 | Mm.10356 | NM_03088 | ENSMUSG | Jdp2 | Jundm2\|Ju | Jun dimeri | protein-co | 7.65 | 8.23 | 6.65 | 1.58 | 1.60E-14 | 1.38E-11 |
| 70 | chr11 | 1.17E+08 | 1.17E+08 | + | 0 | NA | intron (NM | intron (NM | -3886 | NM_01738 | 53860 | Mm.38450 | NM_01738 | ENSMUSG | 9-Sep | MSF1\|Msf | septin 9 | protein-co | 6.66 | 7.39 | 5.14 | 2.24 | 2.27E-14 | 1.91E-11 |
| 71 | chr12 | 25010520 | 25010670 | + | 0 | NA | intron (NM | intron (NM | 35663 | NM_00108 | 77480 | Mm.25064 | NM_00108 | ENSMUSG | Kidins220 | 3110039L1 | kinase D-ir | protein-co | 5.89 | 6.74 | 3.48 | 3.27 | 2.29E-14 | 1.91E-11 |
| 72 | chr17 | 50097964 | 50098114 | + | 0 | NA | intron (NM | intron (NM | 92458 | NM_18139 | 76438 | Mm.41854 | NM_18139 | ENSMUSG | Rftn1 | 2310015N | raftlin lipic | protein-co | 7.41 | 8.06 | 6.22 | 1.84 | 2.66E-14 | 2.20E-11 |
| 74 | chr18 | 84804431 | 84804581 | + | 0 | NA | intron (NM | intron (NM | 34135 | NM_00137 | 1.01E+08 | | NM_00137 | ENSMUSG | Gm17266 | - | predicted | protein-co | 7.74 | 8.31 | 6.81 | 1.5 | 3.61E-14 | 2.89E-11 |
| 75 | chr18 | 10568653 | 10568803 | + | 0 | NA | intron (NM | L1M4\|LINI | 41624 | NM_00108 | 77805 | Mm.21099 | NM_14454 | ENSMUSG | Esco1 | A930014I1 | establishm | protein-co | 6.03 | 6.85 | 3.92 | 2.92 | 3.78E-14 | 2.99E-11 |
| 76 | chr3 | 1.47E+08 | 1.47E+08 | + | 0 | NA | intron (NM | B4A\|SINE\| | 5158 | NM_00116 | 18749 | Mm.16766 | NM_01110 | ENSMUSG | Prkacb | CbPKA\|Pk | protein kir | protein-co | 8.79 | 9.26 | 8.09 | 1.16 | 4.29E-14 | 3.35E-11 |
| 77 | chr1 | 1.93E+08 | 1.93E+08 | + | 0 | NA | intron (NM | intron (NM | 31769 | NM_14481 | 215303 | Mm.40329 | NM_14481 | ENSMUSG | Camk1g | CLICK-III\|C | calcium/ca | protein-co | 7.59 | 8.21 | 6.48 | 1.74 | 4.44E-14 | 3.39E-11 |
| 82 | chr5 | 21354603 | 21354753 | + | 0 | NA | intron (NM | intron (NM | -17995 | NM_00801 | 14190 | Mm.29210 | NM_00801 | ENSMUSG | Fgl2 | AI385601\| | fibrinogen | protein-co | 7.76 | 8.31 | 6.85 | 1.47 | 7.26E-14 | 5.26E-11 |
| 85 | chr14 | 19797027 | 19797177 | + | 0 | NA | intron (NM | intron (NM | 26721 | NM_02652 | 68045 | Mm.21932 | NM_02652 | ENSMUSG | Rtraf | 2700060E0 | RNA transc | protein-co | 5.73 | 6.59 | 3.24 | 3.35 | 9.95E-14 | 6.95E-11 |
| 90 | chr7 | 27609814 | 27609964 | + | 0 | NA | intron (NM | intron (NM | 2553 | NM_00133 | 11652 | Mm.17719 | NM_00743 | ENSMUSG | Akt2 | 2410016A1 | thymoma | protein-co | 6.99 | 7.65 | 5.78 | 1.87 | 1.35E-13 | 8.89E-11 |
| 93 | chr4 | 1.01E+08 | 1.01E+08 | + | 0 | NA | intron (NM | intron (NM | 77693 | NM_01356 | 16451 | Mm.28965 | NM_01356 | ENSMUSG | Jak1 | AA960307 | Janus kina | protein-co | 6.53 | 7.25 | 5.03 | 2.22 | 1.54E-13 | 9.80E-11 |
| 94 | chr17 | 47531561 | 47531711 | + | 0 | NA | intron (NM | intron (NM | 26585 | NM_00108 | 12445 | Mm.27291 | NM_00763 | ENSMUSG | Ccnd3 | 9230106B0 | cyclin D3 | protein-co | 6.74 | 7.43 | 5.35 | 2.09 | 1.58E-13 | 9.96E-11 |
| 95 | chr18 | 84874299 | 84874449 | + | 0 | NA | intron (NM | intron (NM | 23036 | NM_00134 | 109672 | Mm.31018 | NM_02579 | ENSMUSG | Cyb5a | 0610009N | cytochrom | protein-co | 8.52 | 9.01 | 7.75 | 1.26 | 1.68E-13 | 1.05E-10 |
| 96 | chr8 | 14994499 | 14994649 | + | 0 | NA | intron (NM | intron (NM | -16451 | NM_02911 | 74901 | Mm.46675 | NM_02911 | ENSMUSG | Kbtbd11 | 2900016B0 | kelch repe | protein-co | 7 | 7.65 | 5.79 | 1.85 | 2.05E-13 | 1.27E-10 |
| 98 | chr1 | 87829806 | 87829956 | + | 0 | NA | intron (NM | (GA)n\|Sim | -23406 | NM_17764 | 227333 | Mm.27721 | NM_17764 | ENSMUSG | Dgkd | AI841987\| | diacylglyce | protein-co | 6.58 | 7.3 | 5.08 | 2.22 | 2.34E-13 | 1.42E-10 |
| 104 | chr15 | 97721413 | 97721563 | + | 0 | NA | promoter- | promoter- | -338 | NM_00890 | 19011 | Mm.1001 | NM_00890 | ENSMUSG | Endou | Pp11r\|Tcl- | endonucle | protein-co | 6.92 | 7.58 | 5.7 | 1.88 | 4.48E-13 | 2.56E-10 |
| 105 | chr9 | 73089616 | 73089766 | + | 0 | NA | intron (NM | intron (NM | -12707 | NM_00113 | 66991 | Mm.47174 | NM_02589 | ENSMUSG | Khdc3 | 2410004A1 | KH domair | protein-co | 7.15 | 7.77 | 6.04 | 1.73 | 4.54E-13 | 2.57E-10 |
| 107 | chr7 | 38189547 | 38189697 | + | 0 | NA | intron (NM | intron (NM | 5835 | NM_02816 | 72244 | Mm.18755 | NM_028166 | | 1600014C1 | AI428873 | RIKEN cDN | protein-co | 7.36 | 7.97 | 6.29 | 1.69 | 4.93E-13 | 2.73E-10 |
| 111 | chr15 | 57878511 | 57878661 | + | 0 | NA | intron (NM | intron (NM | 13832 | NM_02420 | 67819 | Mm.28938 | NM_02420 | ENSMUSG | Derl1 | 1110021N0 | Der1-like c | protein-co | 6.89 | 7.54 | 5.65 | 1.89 | 6.18E-13 | 3.31E-10 |
| 112 | chr11 | 58883880 | 58884030 | + | 0 | NA | 3' UTR (NM | 3' UTR (NM | 16715 | NM_17240 | 69944 | Mm.32118 | NM_17240 | ENSMUSG | 2810021J2 | AI449137 | RIKEN cDN | protein-co | 5.73 | 6.58 | 3.34 | 3.24 | 6.39E-13 | 3.39E-10 |
| 114 | chr3 | 1.53E+08 | 1.53E+08 | + | 0 | NA | intron (NM | ORR1E\|LT | 55794 | NM_01202 | 26938 | Mm.40915 | NM_01202 | ENSMUSG | St6galnac5 | AI851940\| | ST6 (alpha | protein-co | 6.85 | 7.52 | 5.55 | 1.97 | 8.42E-13 | 4.38E-10 |
| 115 | chr6 | 1.2E+08 | 1.2E+08 | + | 0 | NA | intron (NM | intron (NM | -38714 | NM_00134 | 29862 | Mm.10366 | NM_01671 | ENSMUSG | Ninj2 | - | ninjurin 2 | protein-co | 8.06 | 8.59 | 7.21 | 1.38 | 8.88E-13 | 4.58E-10 |
| 118 | chr18 | 20957178 | 20957328 | + | 0 | NA | intron (NM | intron (NM | -4111 | NM_00136 | 67664 | Mm.45980 | NM_02630 | ENSMUSG | Rnf125 | 4930553F0 | ring finger | protein-co | 6.21 | 6.97 | 4.48 | 2.49 | 9.95E-13 | 5.01E-10 |
| 120 | chr14 | 63377447 | 63377597 | + | 0 | NA | intron (NM | intron (NM | 39665 | NM_00754 | 12143 | Mm.3962 | NM_00754 | ENSMUSG | Blk | - | B lymphoi | protein-co | 6.99 | 7.63 | 5.81 | 1.82 | 1.25E-12 | 6.20E-10 |
| 123 | chr1 | 39396498 | 39396648 | + | 0 | NA | intron (NM | intron (NM | 28722 | NM_00125 | 114641 | Mm.37930 | NM_05325 | ENSMUSG | Rpl31 | Dts | ribosomal | protein-co | 5.72 | 6.57 | 3.46 | 3.11 | 1.66E-12 | 8.04E-10 |
| 127 | chr2 | 17702893 | 17703043 | + | 0 | NA | intron (NM | intron (NM | 28166 | NR_15612 | 74103 | Mm.12029 | NM_02875 | ENSMUSG | Nebl | 1200007O | nebulette | protein-co | 7.23 | 7.83 | 6.19 | 1.65 | 2.47E-12 | 1.16E-09 |
| 130 | chr5 | 36979807 | 36979957 | + | 0 | NA | intron (NM | intron (NM | 9100 | NM_01171 | 22393 | Mm.20916 | NM_01171 | ENSMUSG | Wfs1 | AI481085\| | wolframin | protein-co | 6.02 | 6.82 | 4.08 | 2.75 | 3.41E-12 | 1.56E-09 |
| 136 | chr18 | 50016860 | 50017010 | + | 0 | NA | intron (NM | intron (NM | -14049 | NM_00136 | 106869 | Mm.27740 | NM_13413 | ENSMUSG | Tnfaip8 | AA987150 | tumor nec | protein-co | 7.72 | 8.24 | 6.89 | 1.36 | 5.38E-12 | 2.35E-09 |
| 138 | chr2 | 1.64E+08 | 1.64E+08 | + | 0 | NA | intron (NM | intron (NM | 1877 | NM_01152 | 20971 | Mm.3815 | NM_01152 | ENSMUSG | Sdc4 | AA959608 | syndecan | protein-co | 7.78 | 8.32 | 6.89 | 1.43 | 7.07E-12 | 3.04E-09 |
| 139 | chr5 | 62858402 | 62858552 | + | 0 | NA | intron (NM | intron (NM | 46821 | NM_00117 | 1E+08 | | NM_00117 | ENSMUSG | Dthd1 | Gm17384 | death dom | protein-co | 7.01 | 7.63 | 5.87 | 1.76 | 7.75E-12 | 3.31E-09 |
| 140 | chr13 | 51754791 | 51754941 | + | 0 | NA | intron (NM | intron (NM | 38881 | NM_00128 | 20354 | Mm.33903 | NM_01366 | ENSMUSG | Sema4d | CD100\|Se | sema dom | protein-co | 6.95 | 7.57 | 5.83 | 1.74 | 7.93E-12 | 3.36E-09 |
| 141 | chr2 | 24748028 | 24748178 | + | 0 | NA | intron (NM | intron (NM | 15049 | NM_00104 | 12287 | Mm.4424 | NM_00757 | ENSMUSG | Cacna1b | AW05027€ | calcium ch | protein-co | 5.93 | 6.71 | 4.11 | 2.61 | 8.14E-12 | 3.43E-09 |
| 145 | chr1 | 1.67E+08 | 1.67E+08 | + | 0 | NA | intron (NM | intron (NM | 39609 | NM_00307 | 80914 | Mm.28089 | NM_00307 | ENSMUSG | Uck2 | AA407809 | uridine-cyt | protein-co | 6.29 | 7.01 | 4.78 | 2.23 | 8.59E-12 | 3.52E-09 |
| 143 | chr8 | 1.17E+08 | 1.17E+08 | + | 0 | NA | intron (NM | B3A\|SINE\| | 85708 | NM_02929 | 75465 | Mm.23114 | NM_02929 | ENSMUSG | Dynlrb2 | 1700009A0 | dynein ligl | protein-co | 7.12 | 7.73 | 6.02 | 1.71 | 8.53E-12 | 3.52E-09 |
| 146 | chr6 | 1.25E+08 | 1.25E+08 | + | 0 | NA | TTS (NM_0 | TTS (NM_0 | 6790 | NM_00949 | 22317 | Mm.32321 | NM_00949 | ENSMUSG | Vamp1 | Syb-1\|Syb | vesicle-ass | protein-co | 6.15 | 6.89 | 4.51 | 2.38 | 8.83E-12 | 3.59E-09 |
| 147 | chr3 | 98336004 | 98336154 | + | 0 | NA | intron (NM | intron (NM | 3890 | NM_01696 | 236539 | Mm.16898 | NM_01696 | ENSMUSG | Phgdh | 3-PGDH\|3 | 3-phosphc | protein-co | 6.61 | 7.3 | 5.27 | 2.03 | 9.32E-12 | 3.76E-09 |
| 148 | chr11 | 95356789 | 95356939 | + | 0 | NA | intron (NM | intron (NM | 19846 | NM_17254 | 215512 | Mm.26211 | NM_17254 | ENSMUSG | Fam117a | 5730593F1 | family witl | protein-co | 7.24 | 7.85 | 6.15 | 1.7 | 9.89E-12 | 3.97E-09 |
| 153 | chr4 | 1.07E+08 | 1.07E+08 | + | 0 | NA | TTS (NM_C | TTS (NM_C | -7149 | NM_00108 | 546840 | Mm.33175 | NM_00108 | ENSMUSG | Ldlrad1 | - | low densit | protein-co | 6.88 | 7.53 | 5.66 | 1.88 | 1.61E-11 | 6.23E-09 |
| 154 | chr12 | 73720492 | 73720642 | + | 0 | NA | intron (NM | intron (NM | 135771 | NM_00113 | 18755 | Mm.34167 | NM_00885 | ENSMUSG | Prkch | Pkch | protein kir | protein-co | 7.58 | 8.15 | 6.62 | 1.53 | 1.73E-11 | 6.68E-09 |
| 156 | chr19 | 10064384 | 10064534 | + | 0 | NA | intron (NM | intron (NM | 22911 | NM_02189 | 60527 | Mm.25387 | NM_02189 | ENSMUSG | Fads3 | AI464531 | fatty acid | protein-co | 6.24 | 6.97 | 4.65 | 2.33 | 2.37E-11 | 9.03E-09 |
| 161 | chr2 | 1.53E+08 | 1.53E+08 | + | 0 | NA | intron (NM | intron (NM | 12015 | NR_14925 | 12048 | Mm.23821 | NM_00974 | ENSMUSG | Bcl2l1 | Bcl(X)L\|Bc | BCL2-like 1 | protein-co | 6.6 | 7.3 | 5.18 | 2.12 | 3.15E-11 | 1.16E-08 |
| 166 | chr5 | 1.12E+08 | 1.12E+08 | + | 0 | NA | intron (NM | intron (NM | 17230 | NM_00135 | 72898 | Mm.15174 | NM_02838 | ENSMUSG | Asphd2 | 2900006N | aspartate | protein-co | 6.44 | 7.16 | 4.92 | 2.25 | 4.29E-11 | 1.53E-08 |
| 167 | chr19 | 32138405 | 32138555 | + | 0 | NA | intron (NM | intron (NM | -35340 | NM_01883 | 54447 | Mm.49122 | NM_01883 | ENSMUSG | Asah2 | AI585898 | N-acylsphi | protein-co | 7.39 | 7.96 | 6.42 | 1.54 | 4.37E-11 | 1.55E-08 |
| 168 | chr16 | 23279948 | 23280098 | + | 0 | NA | intron (NM | intron (NM | -10447 | NM_00125 | 20440 | Mm.14902 | NM_14593 | ENSMUSG | St6gal1 | AW742324 | beta galac | protein-co | 5.98 | 6.73 | 4.28 | 2.46 | 4.46E-11 | 1.58E-08 |
| 170 | chr13 | 63108770 | 63108920 | + | 0 | NA | intron (NM | intron (NM | -131184 | NR_11052 | 72061 | Mm.18810 | NM_02807 | ENSMUSG | Aopep | 2010111I0 | aminopept | protein-co | 6.82 | 7.44 | 5.7 | 1.74 | 4.88E-11 | 1.70E-08 |
| 171 | chr4 | 1.03E+08 | 1.03E+08 | + | 0 | NA | promoter- | promoter- | -676 | NM_00116 | 67344 | Mm.11134 | NM_02610 | ENSMUSG | Tctex1d1 | 1700055O | Tctex1 dor | protein-co | 5.82 | 6.6 | 4.02 | 2.58 | 5.07E-11 | 1.76E-08 |
| 172 | chr12 | 1.02E+08 | 1.02E+08 | + | 0 | NA | intron (NM | intron (NM | 11858 | NM_00116 | 217835 | Mm.24145 | NM_17762 | ENSMUSG | Rin3 | 6430500K0 | Ras and Ra | protein-co | 8.6 | 9.03 | 7.97 | 1.07 | 5.29E-11 | 1.83E-08 |
| 174 | chr4 | 8799645 | 8799795 | + | 0 | NA | intron (NM | intron (NM | 48396 | NM_00135 | 320790 | Mm.13879 | NM_00103 | ENSMUSG | Chd7 | A730019I0 | chromodo | protein-co | 6.45 | 7.13 | 5.09 | 2.05 | 5.69E-11 | 1.94E-08 |
| 175 | chr10 | 1.29E+08 | 1.29E+08 | + | 0 | NA | exon (NM_ | exon (NM_ | 610 | NM_14686 | 18373 | Mm.37711 | NM_14686 | ENSMUSG | Olfr9 | MOR269-3 | olfactory r | protein-co | 6.95 | 7.56 | 5.85 | 1.71 | 5.99E-11 | 2.03E-08 |
| 177 | chr10 | 99127803 | 99127953 | + | 0 | NA | intron (NR | intron (NR | 19743 | NM_01573 | 14426 | Mm.314 | NM_01573 | ENSMUSG | Galnt4 | AV011803 | polypeptic | protein-co | 7.62 | 8.15 | 6.76 | 1.39 | 7.06E-11 | 2.37E-08 |
| 181 | chr9 | 1.16E+08 | 1.16E+08 | + | 0 | NA | intron (NM | intron (NM | 36462 | NM_02957 | 21813 | Mm.17234 | NM_00937 | ENSMUSG | Tgfbr2 | 1110020H | transformi | protein-co | 5.99 | 6.73 | 4.4 | 2.33 | 8.70E-11 | 2.85E-08 |
| 184 | chr6 | 1.28E+08 | 1.28E+08 | + | 0 | NA | intron (NM | intron (NM | 4838 | NM_00103 | 381812 | Mm.29609 | NM_00103 | ENSMUSG | Cracr2a | Efcab4b\|G | calcium re | protein-co | 6.65 | 7.3 | 5.42 | 1.88 | 1.13E-10 | 3.61E-08 |
| 186 | chr8 | 1.21E+08 | 1.21E+08 | + | 0 | NA | intron (NM | (ATG)n\|Sir | 10148 | NM_00130 | 15900 | Mm.33486 | NM_00832 | ENSMUSG | Irf8 | AI893568\| | interferon | protein-co | 5.74 | 6.52 | 3.91 | 2.62 | 1.32E-10 | 4.19E-08 |
| 187 | chr9 | 75645555 | 75645705 | + | 0 | NA | intron (NM | intron (NM | 19898 | NM_02730 | 70082 | Mm.19119 | NM_02730 | ENSMUSG | Lysmd2 | 2210402C1 | LysM, puta | protein-co | 6.04 | 6.78 | 4.46 | 2.32 | 1.49E-10 | 4.70E-08 |
| 189 | chr18 | 82479918 | 82480068 | + | 0 | NA | intron (NM | intron (NM | 4870 | NM_00102 | 17196 | Mm.25206 | NM_01077 | ENSMUSG | Mbp | C76307\|H | myelin bas | protein-co | 6.81 | 7.43 | 5.7 | 1.73 | 1.58E-10 | 4.93E-08 |
| 190 | chr15 | 97750824 | 97750974 | + | 0 | NA | intron (NM | intron (NM | 16767 | NM_00135 | 223864 | Mm.24028 | NM_14485 | ENSMUSG | Rapgef3 | 2310016P2 | Rap guanir | protein-co | 6.91 | 7.52 | 5.85 | 1.67 | 1.64E-10 | 5.08E-08 |
| 191 | chr2 | 93192241 | 93192391 | + | 0 | NA | intron (NM | intron (NM | 4768 | NM_00135 | 277414 | Mm.41033 | NM_00102 | ENSMUSG | Trp53i11 | Tp53i11 | transforma | protein-co | 7.56 | 8.08 | 6.72 | 1.36 | 1.69E-10 | 5.20E-08 |
| 194 | chr9 | 1.03E+08 | 1.03E+08 | + | 0 | NA | exon (NM_ | exon (NM_ | 36479 | NM_02429 | 16716 | Mm.10169 | NM_02429 | ENSMUSG | Ky | CD1\|D9Mg | kyphoscoli | protein-co | 7.06 | 7.66 | 6.02 | 1.64 | 1.77E-10 | 5.39E-08 |
| 195 | chr9 | 21328091 | 21328241 | + | 0 | NA | intron (NM | intron (NM | 7447 | NM_00135 | 68682 | Mm.14842 | NM_15280 | ENSMUSG | Slc44a2 | 1110028E1 | solute carr | protein-co | 6.21 | 6.91 | 4.77 | 2.14 | 2.08E-10 | 6.30E-08 |
| 198 | chr10 | 76293330 | 76293480 | + | 0 | NA | intron (NM | (TGGG)n\|S | 39569 | NM_00911 | 20203 | Mm.23599 | NM_00911 | ENSMUSG | S100b | AI850290\| | S100 prote | protein-co | 6.49 | 7.17 | 5.13 | 2.04 | 2.31E-10 | 6.88E-08 |
| 199 | chr8 | 1.17E+08 | 1.17E+08 | + | 0 | NA | intron (NM | intron (NM | 94929 | NM_02929 | 75465 | Mm.23114 | NM_02929 | ENSMUSG | Dynlrb2 | 1700009A0 | dynein ligl | protein-co | 6.32 | 7.02 | 4.91 | 2.11 | 2.32E-10 | 6.88E-08 |
| 200 | chr2 | 1.28E+08 | 1.28E+08 | + | 0 | NA | intron (NM | intron (NM | 73062 | NM_02876 | 74121 | Mm.24483 | NM_02876 | ENSMUSG | Acoxl | 1200014P0 | acyl-Coenz | protein-co | 5.33 | 6.19 | 2.92 | 3.26 | 2.50E-10 | 7.39E-08 |
| 201 | chr8 | 14985651 | 14985801 | + | 0 | NA | intron (NM | intron (NM | -25299 | NM_02911 | 74901 | Mm.46675 | NM_02911 | ENSMUSG | Kbtbd11 | 2900016B0 | kelch repe | protein-co | 7.36 | 7.91 | 6.46 | 1.46 | 2.71E-10 | 7.98E-08 |
| 203 | chr10 | 43980175 | 43980325 | + | 0 | NA | intron (NM | intron (NM | 24596 | NM_17239 | 11630 | Mm.29208 | NM_17239 | ENSMUSG | Crybg1 | Aim1 | crystallin | protein-co | 7.01 | 7.6 | 5.99 | 1.61 | 2.90E-10 | 8.45E-08 |
| 205 | chr18 | 80697119 | 80697269 | + | 0 | NA | intron (NM | intron (NM | 10979 | NM_00116 | 18018 | Mm.32956 | NM_01679 | ENSMUSG | Nfatc1 | 2210017P0 | nuclear fac | protein-co | 7.31 | 7.85 | 6.43 | 1.42 | 3.39E-10 | 9.72E-08 |
| 206 | chr4 | 1.38E+08 | 1.38E+08 | + | 0 | NA | intron (NM | intron (NM | -4744 | NM_00119 | 16559 | Mm.27193 | NM_01062 | ENSMUSG | Kif17 | 5930435E0 | kinesin far | protein-co | 6.37 | 7.04 | 5.1 | 1.94 | 3.42E-10 | 9.77E-08 |
| 207 | chr7 | 49601687 | 49601837 | + | 0 | NA | intron (NM | intron (NM | 35073 | NM_00100 | 13172 | Mm.30717 | NM_00100 | ENSMUSG | Dbx1 | AI426026\| | developing | protein-co | 7.4 | 7.95 | 6.53 | 1.42 | 3.74E-10 | 1.06E-07 |
| 209 | chr2 | 1.59E+08 | 1.59E+08 | + | 0 | NA | promoter- | promoter- | -168 | NM_15308 | 228852 | Mm.15054 | NM_15308 | ENSMUSG | Ppp1r16b | ANKRD4\|C | protein ph | protein-co | 8.36 | 8.81 | 7.71 | 1.1 | 3.86E-10 | 1.09E-07 |
| 211 | chr3 | 89871126 | 89871276 | + | 0 | NA | exon (NM_ | exon (NM_ | 39831 | NM_17253 | 214547 | Mm.225 | NM_17253 | ENSMUSG | She | 9430022A1 | src homolo | protein-co | 6.67 | 7.31 | 5.51 | 1.8 | 3.99E-10 | 1.11E-07 |
| 214 | chr3 | 1.27E+08 | 1.27E+08 | + | 0 | NA | intron (NM | intron (NM | 63786 | NM_00134 | 108058 | Mm.25582 | NM_02381 | ENSMUSG | Camk2d | 2810011D1 | calcium/ca | protein-co | 6.59 | 7.25 | 5.36 | 1.89 | 4.35E-10 | 1.20E-07 |
| 217 | chr2 | 32062298 | 32062448 | + | 0 | NA | intron (NR | RLTR8\|LTF | 1791 | NR_15149 | 241303 | Mm.25044 | NM_17551 | ENSMUSG | Fam78a | A130092J0 | family witl | protein-co | 6.13 | 6.84 | 4.68 | 2.16 | 5.51E-10 | 1.49E-07 |
| 219 | chr18 | 80689959 | 80690109 | + | 0 | NA | intron (NM | intron (NM | 18139 | NM_00116 | 18018 | Mm.32956 | NM_01679 | ENSMUSG | Nfatc1 | 2210017P0 | nuclear fac | protein-co | 7.15 | 7.71 | 6.22 | 1.5 | 5.82E-10 | 1.56E-07 |
| 222 | chr8 | 11506957 | 11507107 | + | 0 | NA | intron (NM | intron (NM | 9464 | NM_02699 | 69225 | Mm.64911 | NM_02699 | ENSMUSG | Naxd | 0710008K0 | NAD(P)HX | protein-co | 6.2 | 6.88 | 4.83 | 2.06 | 6.60E-10 | 1.75E-07 |
| 224 | chr11 | 95834371 | 95834521 | + | 0 | NA | intron (NM | intron (NM | -7849 | NM_00128 | 14710 | Mm.46299 | NM_02312 | ENSMUSG | Gngt2 | AV096848 | guanine nu | protein-co | 6.8 | 7.4 | 5.78 | 1.62 | 7.40E-10 | 1.94E-07 |
| 227 | chr15 | 67103470 | 67103620 | + | 0 | NA | 3' UTR (NM | 3' UTR (NM | 73337 | NM_00917 | 20442 | Mm.24833 | NM_00917 | ENSMUSG | St3gal1 | 5330418N | ST3 beta-g | protein-co | 6.23 | 6.91 | 4.92 | 1.99 | 8.47E-10 | 2.20E-07 |
| 229 | chr10 | 75978249 | 75978399 | + | 0 | NA | exon (NM_ | exon (NM_ | 23810 | NM_19863 | 333669 | Mm.31077 | NM_19863 | ENSMUSG | Gm5134 | EG333669 | predicted | protein-co | 5.47 | 6.29 | 3.4 | 2.89 | 9.49E-10 | 2.44E-07 |
| 231 | chr4 | 1.36E+08 | 1.36E+08 | + | 0 | NA | intron (NM | intron (NM | -2827 | NM_00130 | 242705 | Mm.30793 | NM_17773 | ENSMUSG | E2f2 | E2F-2 | E2F transc | protein-co | 7.6 | 8.1 | 6.82 | 1.27 | 1.25E-09 | 3.16E-07 |
| 235 | chr4 | 1.35E+08 | 1.35E+08 | + | 0 | NA | intron (NM | intron (NM | -15716 | NM_00136 | 12399 | Mm.37889 | NM_01973 | ENSMUSG | Runx3 | AML2\|Cbf | runt relate | protein-co | 5.41 | 6.21 | 3.42 | 2.79 | 1.31E-09 | 3.28E-07 |
| 236 | chr15 | 86094845 | 86094995 | + | 0 | NA | intron (NM | intron (NM | 36193 | NM_00120 | 223752 | Mm.24442 | NM_17261 | ENSMUSG | Gramd4 | 9930106O | GRAM dor | protein-co | 5.95 | 6.68 | 4.4 | 2.29 | 1.41E-09 | 3.51E-07 |
| 237 | chr18 | 82484966 | 82485116 | + | 0 | NA | intron (NM | intron (NM | 9918 | NM_00102 | 17196 | Mm.25206 | NM_01077 | ENSMUSG | Mbp | C76307\|H | myelin bas | protein-co | 6.57 | 7.21 | 5.42 | 1.79 | 1.60E-09 | 3.97E-07 |
| 239 | chr3 | 1.06E+08 | 1.06E+08 | + | 0 | NA | intron (NM | intron (NM | -9580 | NM_00963 | 11542 | Mm.23502 | NM_00963 | ENSMUSG | Adora3 | A3AR\|A3R | adenosine | protein-co | 8.4 | 8.82 | 7.8 | 1.02 | 2.11E-09 | 5.17E-07 |
| 240 | chr2 | 28540123 | 28540273 | + | 0 | NA | intron (NM | intron (NM | 6734 | NM_00114 | 19730 | Mm.5236 | NM_00905 | ENSMUSG | Ralgds | Gnds\|Rgd: | ral guanine | protein-co | 6.86 | 7.45 | 5.87 | 1.57 | 2.13E-09 | 5.21E-07 |
| 242 | chr5 | 34249262 | 34249412 | + | 0 | NA | intron (NM | intron (NM | 38987 | NM_00101 | 231125 | Mm.11338 | NM_00101 | ENSMUSG | Zfyve28 | 9630058O | zinc finger | protein-co | 5.64 | 6.4 | 3.9 | 2.51 | 2.25E-09 | 5.46E-07 |
| 243 | chr15 | 66824788 | 66824938 | + | 0 | NA | intron (NM | intron (NM | 6966 | NM_00919 | 20491 | Mm.7601 | NM_00919 | ENSMUSG | Sla | Slap\|Slap- | src-like ad | protein-co | 5.78 | 6.52 | 4.18 | 2.34 | 2.46E-09 | 5.93E-07 |

| | | | | | | | | | | | | | | | | | | | | | | | | |
|---|---|---|---|---|---|---|---|---|---|---|---|---|---|---|---|---|---|---|---|---|---|---|---|---|
| 249 | chr6 | 50437856 | 50438006 | + | 0 | NA | intron (NM | RMER15-ir | 18239 | NM_0011 | 71720 | Mm.44153 | NM_0278 | ENSMUSG | Osbpl3 | 1200014M | oxysterol b | protein-co | 6.23 | 6.91 | 4.88 | 2.03 | 2.72E-09 | 6.42E-07 |
| 250 | chrX | 1.6E+08 | 1.6E+08 | + | 0 | NA | intron (NM | intron (NM | -11388 | NR_11097 | 58194 | Mm.28649 | NM_0213 | ENSMUSG | Sh3kbp1 | 1200007H | SH3-doma | protein-co | 6.47 | 7.13 | 5.22 | 1.91 | 2.90E-09 | 6.81E-07 |
| 251 | chr7 | 1.27E+08 | 1.27E+08 | + | 0 | NA | promoter- | promoter- | 42 | NM_0174 | 54204 | Mm.18 | NM_0174 | ENSMUSG | 1-Sep | Diff6\|Pnut | septin 1 | protein-co | 6.44 | 7.07 | 5.28 | 1.8 | 3.10E-09 | 7.26E-07 |
| 254 | chr9 | 63504034 | 63504184 | + | 0 | NA | intron (NM | MTD\|LTR\| | 98384 | NM_0013 | 78250 | Mm.44461 | NM_0300 | ENSMUSG | Iqch | 4921504K | IQ motif c | protein-co | 5.54 | 6.31 | 3.78 | 2.53 | 3.25E-09 | 7.48E-07 |
| 256 | chr10 | 26325405 | 26325555 | + | 0 | NA | intron (NM | RLTR14_RI | 48592 | NM_0013 | 237339 | Mm.33843 | NM_1727 | ENSMUSG | L3mbtl3 | AI481284\| | L3MBTL3 h | protein-co | 6.66 | 7.29 | 5.52 | 1.77 | 3.29E-09 | 7.54E-07 |
| 258 | chr2 | 1.02E+08 | 1.02E+08 | + | 0 | NA | intron (NM | intron (NM | 41355 | NM_0010 | 72446 | Mm.24138 | NM_1751 | ENSMUSG | Prr5l | 2600010E | proline ric | protein-co | 6.84 | 7.44 | 5.79 | 1.65 | 3.41E-09 | 7.75E-07 |
| 260 | chr2 | 57116448 | 57116598 | + | 0 | NA | intron (NM | CpG | -1485 | NM_0136 | 18227 | Mm.3507 | NM_0136 | ENSMUSG | Nr4a2 | HZF-3\|NO | nuclear re | protein-co | 7.52 | 8.01 | 6.78 | 1.23 | 3.78E-09 | 8.53E-07 |
| 262 | chr2 | 1.57E+08 | 1.57E+08 | + | 0 | NA | promoter- | promoter- | 41 | NM_0013 | 77799 | Mm.29264 | NM_0299 | ENSMUSG | Sla2 | A930009E | Src-like-ad | protein-co | 7.54 | 8.08 | 6.67 | 1.41 | 3.93E-09 | 8.81E-07 |
| 261 | chr5 | 1.18E+08 | 1.18E+08 | + | 0 | NA | intron (NM | intron (NM | 54438 | NR_15158 | 57816 | Mm.27328 | NM_0213 | ENSMUSG | Tesc | 1010001A | tescalcin | protein-co | 8.37 | 8.8 | 7.76 | 1.04 | 3.93E-09 | 8.81E-07 |
| 264 | chr18 | 24596886 | 24597036 | + | 0 | NA | intron (NM | intron (NM | 6856 | NM_1391 | 106957 | Mm.21688 | NM_1391 | ENSMUSG | Slc39a6 | Ermelin\|Zi | solute carr | protein-co | 5.66 | 6.4 | 4.04 | 2.36 | 4.19E-09 | 9.32E-07 |
| 265 | chr4 | 1.55E+08 | 1.55E+08 | + | 0 | NA | intron (NM | intron (NM | 59966 | NM_0113 | 20481 | Mm.28520 | NM_0113 | ENSMUSG | Ski | 2310012I0 | ski sarcom | protein-co | 6.8 | 7.38 | 5.81 | 1.57 | 4.23E-09 | 9.34E-07 |
| 266 | chr7 | 1.11E+08 | 1.11E+08 | + | 0 | NA | intron (NM | ORR1E\|LT | -10470 | NM_0108 | 17540 | Mm.25002 | NM_0108 | ENSMUSG | Mrvi1 | BB115629 | MRV integ | protein-co | 6.24 | 6.9 | 4.98 | 1.91 | 4.24E-09 | 9.34E-07 |
| 271 | chr8 | 1.2E+08 | 1.2E+08 | + | 0 | NA | intron (NM | intron (NM | 40287 | NM_0302 | 78892 | Mm.26468 | NM_0302 | ENSMUSG | Crispld2 | 1810049K | cysteine-ri | protein-co | 5.37 | 6.17 | 3.41 | 2.76 | 4.78E-09 | 1.03E-06 |
| 272 | chr11 | 88164444 | 88164594 | + | 0 | NA | intron (NM | intron (NM | -5046 | NM_0011 | 103841 | Mm.37484 | NM_1980 | ENSMUSG | Cuedc1 | AI841487\| | CUE doma | protein-co | 6.02 | 6.71 | 4.63 | 2.08 | 4.87E-09 | 1.05E-06 |
| 277 | chr5 | 1.14E+08 | 1.14E+08 | + | 0 | NA | exon (NM_ | exon (NM_ | -33491 | NM_0013 | 231646 | Mm.23828 | NM_1461 | ENSMUSG | Myo1h | 4631401O | myosin 1H | protein-co | 6.46 | 7.09 | 5.34 | 1.74 | 5.32E-09 | 1.12E-06 |
| 278 | chr2 | 66102114 | 66102264 | + | 0 | NA | intron (NM | MIR\|SINE\| | 22604 | NM_0157 | 14425 | Mm.43976 | NM_0157 | ENSMUSG | Galnt3 | - | polypeptic | protein-co | 7.5 | 7.99 | 6.76 | 1.23 | 5.78E-09 | 1.21E-06 |
| 279 | chr19 | 47526623 | 47526773 | + | 0 | NA | intron (NM | intron (NM | 10421 | NM_1753 | 108689 | Mm.21804 | NM_1753 | ENSMUSG | Stn1 | 0610009H | STN1, CST | protein-co | 7.21 | 7.73 | 6.39 | 1.34 | 5.88E-09 | 1.23E-06 |
| 282 | chr9 | 57148309 | 57148459 | + | 0 | NA | promoter- | promoter- | -79 | NR_15168 | 72774 | Mm.35749 | NM_0283 | ENSMUSG | Neil1 | 2810450N | nei endon | protein-co | 7.32 | 7.86 | 6.43 | 1.43 | 6.30E-09 | 1.30E-06 |
| 283 | chr13 | 63528466 | 63528616 | + | 0 | NA | intron (NM | intron (NM | 37194 | NM_0089 | 19206 | Mm.22879 | NM_0089 | ENSMUSG | Ptch1 | A230106A | patched 1 | protein-co | 7.97 | 8.43 | 7.28 | 1.14 | 6.42E-09 | 1.32E-06 |
| 284 | chr1 | 43500213 | 43500363 | + | 0 | NA | intron (NM | intron (NM | 54537 | NM_0108 | 17974 | Mm.38990 | NM_0108 | ENSMUSG | Nck2 | 4833426I1 | non-cataly | protein-co | 7.22 | 7.74 | 6.39 | 1.36 | 6.71E-09 | 1.38E-06 |
| 286 | chr4 | 46592223 | 46592373 | + | 0 | NA | intron (NM | intron (NM | 9631 | NM_0011 | 107684 | Mm.41631 | NM_1788 | ENSMUSG | Coro2a | 9030208C | coronin, a | protein-co | 6.12 | 6.79 | 4.84 | 1.95 | 7.92E-09 | 1.62E-06 |
| 287 | chr19 | 57131543 | 57131693 | + | 0 | NA | intron (NM | intron (NM | -12594 | NM_0011 | 226251 | Mm.21716 | NM_1786 | ENSMUSG | Ablim1 | 2210411C | actin-bindi | protein-co | 7.14 | 7.67 | 6.29 | 1.37 | 8.25E-09 | 1.68E-06 |
| 290 | chr2 | 1.21E+08 | 1.21E+08 | + | 0 | NA | intron (NM | intron (NM | 23311 | NM_0011 | 12335 | Mm.45802 | NM_0076 | ENSMUSG | Capn3 | AI323605\| | calpain 3 | protein-co | 6.61 | 7.25 | 5.43 | 1.82 | 9.00E-09 | 1.81E-06 |
| 294 | chr2 | 1.59E+08 | 1.59E+08 | + | 0 | NA | intron (NM | intron (NM | -33922 | NM_0279 | 71878 | Mm.85162 | NM_0279 | ENSMUSG | Fam83d | 2310007D | family wit | protein-co | 7.62 | 8.13 | 6.82 | 1.31 | 9.70E-09 | 1.93E-06 |
| 295 | chr6 | 83312642 | 83312792 | + | 0 | NA | intron (NM | intron (NM | 4887 | NM_0086 | 17768 | Mm.443 | NM_0086 | ENSMUSG | Mthfd2 | AW558851 | methylene | protein-co | 7.72 | 8.18 | 7.05 | 1.13 | 9.74E-09 | 1.93E-06 |
| 296 | chr2 | 1.63E+08 | 1.63E+08 | + | 0 | NA | intron (NM | intron (NM | 27967 | NM_0013 | 269389 | Mm.20770 | NM_0010 | ENSMUSG | Tox2 | AI851523\| | TOX high r | protein-co | 7.98 | 8.41 | 7.38 | 1.03 | 1.01E-08 | 1.99E-06 |
| 297 | chr11 | 11716078 | 11716228 | + | 0 | NA | intron (NM | intron (NM | 25976 | NM_0013 | 22778 | Mm.10354 | NM_0095 | ENSMUSG | Ikzf1 | 5832432G | IKAROS fa | protein-co | 5.94 | 6.63 | 4.57 | 2.07 | 1.03E-08 | 2.02E-06 |
| 299 | chr4 | 1.55E+08 | 1.55E+08 | + | 0 | NA | intron (NM | intron (NM | 59140 | NM_0113 | 20481 | Mm.28520 | NM_0113 | ENSMUSG | Ski | 2310012I0 | ski sarcom | protein-co | 5.92 | 6.62 | 4.54 | 2.08 | 1.04E-08 | 2.03E-06 |
| 306 | chr2 | 25035073 | 25035223 | + | 0 | NA | intron (NM | intron (NM | -19231 | NM_0011 | 56876 | Mm.42956 | NM_0202 | ENSMUSG | Nsmf | Jacob\|Nel | NMDA rec | protein-co | 4.85 | 5.74 | 2.07 | 3.67 | 1.32E-08 | 2.51E-06 |
| 308 | chr10 | 1.29E+08 | 1.29E+08 | + | 0 | NA | exon (NM_ | exon (NM_ | 473 | NM_1468 | 258861 | Mm.37772 | NM_1468 | ENSMUSG | Olfr763 | MOR269-2 | olfactory r | protein-co | 5.7 | 6.42 | 4.17 | 2.25 | 1.35E-08 | 2.56E-06 |
| 309 | chr5 | 1.39E+08 | 1.39E+08 | + | 0 | NA | intron (NM | intron (NM | 11269 | NM_0284 | 73212 | Mm.84294 | NM_0284 | ENSMUSG | 3110082I1 | AW41190 | RIKEN cDN | protein-co | 5.54 | 6.29 | 3.9 | 2.39 | 1.36E-08 | 2.57E-06 |
| 310 | chr2 | 1.45E+08 | 1.45E+08 | + | 0 | NA | intron (NM | intron (NM | -33628 | NM_0531 | 94249 | Mm.21717 | NM_0531 | ENSMUSG | Slc24a3 | NCKX3 | solute carr | protein-co | 6.75 | 7.33 | 5.76 | 1.56 | 1.39E-08 | 2.61E-06 |
| 311 | chr11 | 1.2E+08 | 1.2E+08 | + | 0 | NA | intron (NM | intron (NM | 35737 | NM_0013 | 268515 | Mm.30949 | NM_1984 | ENSMUSG | Bahcc1 | B930044J | BAH doma | protein-co | 5.81 | 6.52 | 4.35 | 2.17 | 1.48E-08 | 2.77E-06 |
| 315 | chr1 | 1.31E+08 | 1.31E+08 | + | 0 | NA | intron (NM | intron (NM | 11298 | NM_0013 | 54354 | Mm.24829 | NM_0187 | ENSMUSG | Rassf5 | 1300019G | Ras associ | protein-co | 7.49 | 7.96 | 6.78 | 1.18 | 1.56E-08 | 2.89E-06 |
| 314 | chr7 | 99120923 | 99121073 | + | 0 | NA | intron (NM | intron (NM | 20146 | NM_1786 | 78610 | Mm.32307 | NM_1786 | ENSMUSG | Uvrag | 9530039D | UV radiati | protein-co | 5.79 | 6.5 | 4.31 | 2.19 | 1.56E-08 | 2.89E-06 |
| 316 | chr8 | 14986768 | 14986918 | + | 0 | NA | intron (NM | intron (NM | -24182 | NM_0291 | 74901 | Mm.46675 | NM_0291 | ENSMUSG | Kbtbd11 | 2900016B | kelch repe | protein-co | 5.67 | 6.39 | 4.15 | 2.24 | 1.68E-08 | 3.09E-06 |
| 320 | chr15 | 55520884 | 55521034 | + | 0 | NA | TTS (NM_1 | TTS (NM_1 | 36353 | NM_0267 | 68537 | Mm.23679 | NM_0267 | ENSMUSG | Mrpl13 | 1110002D | mitochon | protein-co | 6.44 | 7.05 | 5.38 | 1.67 | 1.81E-08 | 3.28E-06 |
| 321 | chr2 | 17365248 | 17365398 | + | 0 | NA | intron (NR | ORR1E\|LT | 97533 | NM_0013 | 74103 | Mm.12029 | NM_0287 | ENSMUSG | Nebl | 1200007O | nebulette | protein-co | 7.87 | 8.33 | 7.17 | 1.16 | 1.81E-08 | 3.28E-06 |
| 323 | chr1 | 95605597 | 95605747 | + | 0 | NA | intron (NM | intron (NM | 61922 | NM_0091 | 20452 | Mm.30622 | NM_0091 | ENSMUSG | St8sia4 | PST\|PST-1 | ST8 alpha- | protein-co | 7.22 | 7.73 | 6.43 | 1.3 | 1.88E-08 | 3.36E-06 |
| 324 | chr16 | 10836440 | 10836590 | + | 0 | NA | intron (NM | intron (NM | 1456 | NM_0011 | 223970 | Mm.32577 | NM_0010 | ENSMUSG | Rmi2 | A630055G | RecQ med | protein-co | 6.55 | 7.14 | 5.54 | 1.6 | 1.88E-08 | 3.36E-06 |
| 325 | chr2 | 1.81E+08 | 1.81E+08 | + | 0 | NA | 5' UTR (NM | 5' UTR (NM | 840 | NM_0236 | 72699 | Mm.44013 | NM_0236 | ENSMUSG | Lime1 | 2810038K | Lck interac | protein-co | 7.81 | 8.26 | 7.15 | 1.12 | 1.88E-08 | 3.36E-06 |
| 327 | chr8 | 9975409 | 9975559 | + | 0 | NA | intron (NM | intron (NM | 839 | NM_1769 | 319583 | Mm.80584 | NM_1769 | ENSMUSG | Lig4 | 5830471N | ligase IV, D | protein-co | 6 | 6.67 | 4.72 | 1.95 | 1.89E-08 | 3.37E-06 |
| 329 | chr12 | 91668625 | 91668775 | + | 0 | NA | intron (NM | intron (NM | -78213 | NM_1753 | 83602 | Mm.27572 | NM_0313 | ENSMUSG | Gtf2a1 | 6330549H | general tra | protein-co | 7.52 | 8 | 6.79 | 1.22 | 2.01E-08 | 3.55E-06 |
| 331 | chr9 | 66560398 | 66560548 | + | 0 | NA | intron (NM | intron (NM | 32642 | NM_1449 | 235441 | Mm.32465 | NM_1449 | ENSMUSG | Usp3 | AA409661 | ubiquitin s | protein-co | 5.96 | 6.64 | 4.64 | 2 | 2.05E-08 | 3.60E-06 |
| 332 | chr9 | 32507542 | 32507692 | + | 0 | NA | intron (NM | intron (NM | 33835 | NM_0080 | 14247 | Mm.25890 | NM_0080 | ENSMUSG | Fli1 | EWSR2\|Fli | Friend leu | protein-co | 7.21 | 7.71 | 6.43 | 1.28 | 2.09E-08 | 3.66E-06 |
| 335 | chr11 | 77887524 | 77887674 | + | 0 | NA | intron (NM | AmnSINE1 | 6273 | NM_0089 | 19193 | Mm.8543 | NM_0089 | ENSMUSG | Pipox | LPIPOX\|Ps | pipecolic a | protein-co | 7.44 | 7.92 | 6.7 | 1.22 | 2.28E-08 | 3.95E-06 |
| 338 | chr12 | 85971791 | 85971941 | + | 0 | NA | intron (NM | intron (NM | 107175 | NM_0093 | 21809 | Mm.3992 | NM_0093 | ENSMUSG | Tgfb3 | TGF-beta- | transformi | protein-co | 5.88 | 6.58 | 4.47 | 2.11 | 2.59E-08 | 4.45E-06 |
| 342 | chr15 | 66815519 | 66815669 | + | 0 | NA | intron (NM | intron (NM | 16235 | NM_0091 | 20491 | Mm.7601 | NM_0091 | ENSMUSG | Sla | Slap\|Slap- | src-like ad | protein-co | 6.89 | 7.42 | 6.01 | 1.41 | 2.97E-08 | 5.02E-06 |
| 345 | chr2 | 1.02E+08 | 1.02E+08 | + | 0 | NA | intron (NM | intron (NM | 55526 | NM_0010 | 72446 | Mm.24138 | NM_1751 | ENSMUSG | Prr5l | 2600010E | proline ric | protein-co | 5.86 | 6.55 | 4.48 | 2.07 | 3.22E-08 | 5.40E-06 |
| 348 | chr9 | 21327699 | 21327849 | + | 0 | NA | intron (NM | B3\|SINE\|B | 7055 | NM_0013 | 68682 | Mm.14842 | NM_1528 | ENSMUSG | Slc44a2 | 1110028E | solute carr | protein-co | 5.97 | 6.66 | 4.61 | 2.05 | 3.39E-08 | 5.63E-06 |
| 351 | chr9 | 75303799 | 75303949 | + | 0 | NA | intron (NM | intron (NM | -2414 | NM_0103 | 14697 | Mm.17604 | NM_0103 | ENSMUSG | Gnb5 | GBS\|Gbet | guanine n | protein-co | 5.9 | 6.57 | 4.61 | 1.96 | 3.62E-08 | 5.95E-06 |
| 352 | chr15 | 80192409 | 80192559 | + | 0 | NA | intron (NM | intron (NM | 18763 | NM_0107 | 17309 | Mm.29969 | NM_0107 | ENSMUSG | Mgat3 | 1110038J1 | mannosid | protein-co | 6.18 | 6.82 | 5 | 1.81 | 3.64E-08 | 5.97E-06 |
| 356 | chr15 | 89131506 | 89131656 | + | 0 | NA | intron (NM | intron (NM | -2881 | NR_02844 | 170787 | Mm.20395 | NM_1991 | ENSMUSG | Hdac10 | AW548891 | histone de | protein-co | 7.33 | 7.82 | 6.6 | 1.22 | 4.19E-08 | 6.80E-06 |
| 358 | chr5 | 1.34E+08 | 1.34E+08 | + | 0 | NA | intron (NM | intron (NM | 9921 | NM_0335 | 94254 | Mm.29508 | NM_0335 | ENSMUSG | Rcc1l | 5730496C | reculator | protein-co | 7.65 | 8.1 | 7.01 | 1.09 | 4.35E-08 | 7.02E-06 |
| 360 | chr8 | 1.05E+08 | 1.05E+08 | + | 0 | NA | intron (NM | intron (NM | 1311 | NM_1489 | 104394 | Mm.34554 | NM_1489 | ENSMUSG | E2f4 | 2010111M | E2F transc | protein-co | 6.64 | 7.22 | 5.64 | 1.58 | 4.87E-08 | 7.79E-06 |
| 362 | chr1 | 36797614 | 36797764 | + | 0 | NA | intron (NM | ORR1F\|LT | 18705 | NM_0012 | 22637 | Mm.8038 | NM_0095 | ENSMUSG | Zap70 | Srk\|ZAP-7 | zeta-chain | protein-co | 5.45 | 6.2 | 3.79 | 2.41 | 5.07E-08 | 8.08E-06 |
| 361 | chr10 | 75137229 | 75137379 | + | 0 | NA | intron (NM | ORR1B2\|L | -74769 | NM_1534 | 74392 | Mm.27716 | NM_1534 | ENSMUSG | Specc1l | 4930470P | sperm anti | protein-co | 6.15 | 6.79 | 4.97 | 1.82 | 5.07E-08 | 8.08E-06 |
| 363 | chr7 | 1.28E+08 | 1.28E+08 | + | 0 | NA | intron (NM | intron (NM | 25399 | NM_0264 | 67865 | Mm.18635 | NM_0264 | ENSMUSG | Rgs10 | 2310010N | regulator | protein-co | 6.77 | 7.33 | 5.85 | 1.48 | 5.16E-08 | 8.20E-06 |
| 364 | chr11 | 1.2E+08 | 1.2E+08 | + | 0 | NA | intron (NM | intron (NM | 34904 | NM_0013 | 268515 | Mm.30949 | NM_1984 | ENSMUSG | Bahcc1 | B930044J | BAH doma | protein-co | 6.74 | 7.3 | 5.82 | 1.49 | 5.19E-08 | 8.22E-06 |
| 366 | chr6 | 8712025 | 8712175 | + | 0 | NA | intron (NM | intron (NM | 46358 | NM_0012 | 15893 | Mm.27568 | NM_0104 | ENSMUSG | Ica1 | 69kDa\|ICA | islet cell a | protein-co | 5.2 | 5.99 | 3.27 | 2.72 | 5.47E-08 | 8.62E-06 |
| 369 | chr11 | 1.02E+08 | 1.02E+08 | + | 0 | NA | intron (NM | intron (NM | 2112 | NM_0081 | 14824 | Mm.1568 | NM_0081 | ENSMUSG | Grn | GP88\|PCD | granulin | protein-co | 6.43 | 7.03 | 5.4 | 1.64 | 5.85E-08 | 9.14E-06 |
| 373 | chr8 | 1.22E+08 | 1.22E+08 | + | 0 | NA | promoter- | promoter- | -65 | NM_0078 | 13057 | Mm.27167 | NM_0078 | ENSMUSG | Cyba | b558\|nmf | cytochrom | protein-co | 8.19 | 8.6 | 7.59 | 1.01 | 6.67E-08 | 1.03E-05 |
| 374 | chr8 | 1.24E+08 | 1.24E+08 | + | 0 | NA | intron (NM | intron (NM | 42439 | NM_1392 | 108148 | Mm.33808 | NM_1392 | ENSMUSG | Galnt2 | AI480629 | polypeptic | protein-co | 6.48 | 7.08 | 5.45 | 1.63 | 6.76E-08 | 1.04E-05 |
| 380 | chr2 | 1.65E+08 | 1.65E+08 | + | 0 | NA | intron (NM | intron (NM | 39071 | NM_1749 | 104010 | Mm.15181 | NM_1749 | ENSMUSG | Cdh22 | - | cadherin 2 | protein-co | 6.38 | 7.02 | 5.2 | 1.82 | 8.05E-08 | 1.21E-05 |
| 379 | chr9 | 1.24E+08 | 1.24E+08 | + | 0 | NA | promoter- | promoter- | 76 | NM_0099 | 12774 | Mm.14302 | NM_0099 | ENSMUSG | Ccr5 | AM4-7\|CD | chemokin | protein-co | 6.96 | 7.48 | 6.15 | 1.32 | 7.97E-08 | 1.21E-05 |
| 386 | chr17 | 56444952 | 56445102 | + | 0 | NA | intron (NM | intron (NM | 17194 | NM_0012 | 19280 | Mm.25877 | NM_0112 | ENSMUSG | Ptprs | AL022616\| | protein ty | protein-co | 4.71 | 5.58 | 2.11 | 3.47 | 8.30E-08 | 1.24E-05 |
| 387 | chr18 | 39083942 | 39084092 | + | 0 | NA | intron (NM | intron (NM | 90872 | NM_0013 | 71302 | Mm.32939 | NM_1751 | ENSMUSG | Arhgap26 | 1810044B | Rho GTPas | protein-co | 6.67 | 7.22 | 5.75 | 1.48 | 8.38E-08 | 1.24E-05 |
| 388 | chr11 | 67588994 | 67589144 | + | 0 | NA | intron (NM | intron (NM | 2569 | NM_0011 | 14457 | Mm.40338 | NM_0080 | ENSMUSG | Gas7 | AW12476 | growth ar | protein-co | 6.45 | 7.03 | 5.46 | 1.57 | 8.85E-08 | 1.31E-05 |
| 394 | chr5 | 35817394 | 35817544 | + | 0 | NA | intron (NM | intron (NM | 59589 | NM_0011 | 231148 | Mm.25444 | NM_1776 | ENSMUSG | Ablim2 | AI606905\| | actin-bindi | protein-co | 5.74 | 6.42 | 4.4 | 2.02 | 9.92E-08 | 1.44E-05 |
| 396 | chr12 | 73715993 | 73716143 | + | 0 | NA | intron (NM | intron (NM | 131272 | NM_0013 | 18755 | Mm.34167 | NM_0088 | ENSMUSG | Prkch | Pkch | protein ki | protein-co | 7.68 | 8.11 | 7.07 | 1.05 | 1.03E-07 | 1.49E-05 |
| 397 | chr18 | 5610124 | 5610274 | + | 0 | NA | intron (NM | intron (NM | 17552 | NM_0013 | 21417 | Mm.3929 | NM_0115 | ENSMUSG | Zeb1 | 3110032K | zinc finger | protein-co | 6.43 | 7.03 | 5.39 | 1.63 | 1.05E-07 | 1.52E-05 |
| 400 | chr4 | 1.24E+08 | 1.24E+08 | + | 0 | NA | intron (NM | intron (NM | 10704 | NM_0174 | 54170 | Mm.22092 | NM_0174 | ENSMUSG | Rragc | AU041672 | Ras-relate | protein-co | 6.56 | 7.13 | 5.6 | 1.54 | 1.09E-07 | 1.55E-05 |
| 404 | chr14 | 31531912 | 31532062 | + | 0 | NA | intron (NM | intron (NM | 36908 | NM_0289 | 74427 | Mm.49120 | NM_0289 | ENSMUSG | Eaf1 | 4933403C | ELL associ | protein-co | 6.34 | 6.94 | 5.29 | 1.65 | 1.17E-07 | 1.64E-05 |
| 405 | chr19 | 6398572 | 6398722 | + | 0 | NA | TTS (NM_ | TTS (NM_ | -1228 | NM_0013 | 19395 | Mm.77017 | NM_0112 | ENSMUSG | Rasgrp2 | CDC25L\|C | RAS, guan | protein-co | 5.82 | 6.49 | 4.54 | 1.95 | 1.19E-07 | 1.68E-05 |
| 407 | chr7 | 83661029 | 83661179 | + | 0 | NA | exon (NM_ | exon (NM_ | -5609 | NM_0013 | 16170 | Mm.10137 | NM_0105 | ENSMUSG | Il16 | mKIAA404 | interleuki | protein-co | 6.26 | 6.87 | 5.19 | 1.68 | 1.27E-07 | 1.78E-05 |
| 408 | chr2 | 1.68E+08 | 1.68E+08 | + | 0 | NA | intron (NM | intron (NM | 101605 | NM_0012 | 18019 | Mm.11680 | NM_0108 | ENSMUSG | Nfatc2 | AI607462\| | nuclear fa | protein-co | 7.43 | 7.9 | 6.72 | 1.17 | 1.29E-07 | 1.80E-05 |
| 410 | chr6 | 1.2E+08 | 1.2E+08 | + | 0 | NA | intron (NM | intron (NM | 10417 | NM_0083 | 16172 | Mm.4481 | NM_0083 | ENSMUSG | Il17ra | AW53815 | interleuki | protein-co | 6.72 | 7.27 | 5.81 | 1.46 | 1.32E-07 | 1.83E-05 |
| 411 | chr10 | 88584559 | 88584709 | + | 0 | NA | intron (NM | intron (NM | 20518 | NM_1754 | 109272 | Mm.14720 | NM_1754 | ENSMUSG | Mybpc1 | 8030451F | myosin bi | protein-co | 6.92 | 7.46 | 6.03 | 1.43 | 1.34E-07 | 1.85E-05 |
| 413 | chr1 | 1.53E+08 | 1.53E+08 | + | 0 | NA | intron (NM | intron (NM | 103297 | NM_1754 | 226518 | Mm.40548 | NM_1754 | ENSMUSG | Nmnat2 | AI843915\| | nicotinami | protein-co | 4.84 | 5.68 | 2.62 | 3.06 | 1.41E-07 | 1.95E-05 |
| 414 | chr7 | 80718611 | 80718761 | + | 0 | NA | intron (NM | L1MB2\|LI | -29809 | NM_1738 | 70461 | Mm.39217 | NM_1738 | ENSMUSG | Crtc3 | 2610312F | CREB regu | protein-co | 6.5 | 7.08 | 5.48 | 1.6 | 1.49E-07 | 2.05E-05 |
| 415 | chr1 | 1.93E+08 | 1.93E+08 | + | 0 | NA | intron (NM | intron (NM | 32343 | NM_1448 | 215303 | Mm.40329 | NM_1448 | ENSMUSG | Camk1g | CLICK-III\|C | calcium/ca | protein-co | 7.6 | 8.04 | 6.96 | 1.08 | 1.51E-07 | 2.07E-05 |
| 419 | chr18 | 20946391 | 20946541 | + | 0 | NA | intron (NM | intron (NM | 1851 | NM_0263 | 67664 | Mm.45980 | NM_0263 | ENSMUSG | Rnf125 | 4930553F | ring finger | protein-co | 6.72 | 7.29 | 5.79 | 1.5 | 1.66E-07 | 2.26E-05 |
| 420 | chr7 | 1.45E+08 | 1.45E+08 | + | 0 | NA | intron (NR | intron (NR | -14064 | NM_0013 | 1E+08 | Mm.10223 | NM_0013 | ENSMUSG | Smim38 | 1810010D | small inte | protein-co | 6.91 | 7.42 | 6.09 | 1.34 | 1.68E-07 | 2.27E-05 |
| 421 | chr8 | 83740069 | 83740219 | + | 0 | NA | intron (NM | intron (NM | 1167 | NM_0011 | 26364 | Mm.33464 | NM_0119 | ENSMUSG | Adgre5 | AA409984 | adhesion | protein-co | 6.73 | 7.29 | 5.79 | 1.5 | 1.70E-07 | 2.29E-05 |
| 422 | chr8 | 1.17E+08 | 1.17E+08 | + | 0 | NA | intron (NM | intron (NM | 81208 | NM_0292 | 75465 | Mm.23114 | NM_0292 | ENSMUSG | Dynlrb2 | 1700009A | dynein lig | protein-co | 5.85 | 6.51 | 4.62 | 1.89 | 1.79E-07 | 2.41E-05 |
| 424 | chr11 | 46380337 | 46380487 | + | 0 | NA | intron (NM | Lx9\|LINE\| | 9103 | NM_0012 | 16428 | Mm.33992 | NM_0105 | ENSMUSG | Itk | Emt\|Tcsk\| | IL2 induci | protein-co | 5.15 | 5.92 | 3.38 | 2.54 | 1.80E-07 | 2.42E-05 |
| 426 | chr1 | 1.38E+08 | 1.38E+08 | + | 0 | NA | intron (NM | intron (NM | 17881 | NM_0011 | 19264 | Mm.39157 | NM_0112 | ENSMUSG | Ptprc | B220\|CD4 | protein ty | protein-co | 5.99 | 6.63 | 4.8 | 1.83 | 1.83E-07 | 2.44E-05 |
| 427 | chr15 | 96583381 | 96583531 | + | 0 | NA | intron (NM | intron (NM | 58506 | NM_1340 | 105727 | Mm.10356 | NM_1340 | ENSMUSG | Slc38a1 | AA408026 | solute carr | protein-co | 4.72 | 5.57 | 2.38 | 3.19 | 1.84E-07 | 2.45E-05 |
| 428 | chr14 | 45155632 | 45155782 | + | 0 | NA | intron (NM | MT2A\|LTR | 63718 | NM_0013 | 70561 | Mm.28129 | NM_1725 | ENSMUSG | Txndc16 | 5730420B | thioredoxi | protein-co | 6.03 | 6.66 | 4.87 | 1.8 | 1.85E-07 | 2.46E-05 |
| 429 | chr2 | 1.61E+08 | 1.61E+08 | + | 0 | NA | intron (NM | intron (NM | 65828 | NM_0212 | 18803 | Mm.44463 | NM_0212 | ENSMUSG | Plcg1 | AI894140\| | phospholi | protein-co | 5.82 | 6.48 | 4.55 | 1.93 | 1.87E-07 | 2.47E-05 |
| 430 | chr4 | 1.02E+08 | 1.02E+08 | + | 0 | NA | intron (NM | intron (NM | 1150 | NM_0011 | 72685 | Mm.76494 | NM_1984 | ENSMUSG | Dnajc6 | 2810027M | DnaJ heat | protein-co | 6.17 | 6.78 | 5.1 | 1.68 | 1.89E-07 | 2.50E-05 |
| 433 | chr19 | 40257521 | 40257671 | + | 0 | NA | intron (NM | intron (NM | 14020 | NM_0168 | 54132 | Mm.5567 | NM_0168 | ENSMUSG | Pdlim1 | CLP36\|Cli | PDZ and L | protein-co | 7.25 | 7.74 | 6.52 | 1.22 | 1.96E-07 | 2.57E-05 |
| 434 | chr1 | 1.34E+08 | 1.34E+08 | + | 0 | NA | intron (NM | intron (NM | 2306 | NM_0167 | 53311 | Mm.37906 | NM_0167 | ENSMUSG | Mybph | AI385643\| | myosin bi | protein-co | 7.42 | 7.88 | 6.72 | 1.16 | 1.98E-07 | 2.59E-05 |
| 435 | chr1 | 1.27E+08 | 1.27E+08 | + | 0 | NA | intron (NM | intron (NM | 103555 | NM_1451 | 107895 | Mm.21468 | NM_1451 | ENSMUSG | Mgat5 | 4930471A | mannosid | protein-co | 6.42 | 7.02 | 5.39 | 1.63 | 2.00E-07 | 2.61E-05 |
| 436 | chr17 | 56819757 | 56819907 | + | 0 | NA | intron (NM | intron (NM | 11176 | NM_0090 | 19725 | Mm.102 | NM_0090 | ENSMUSG | Rfx2 | 5430432H | regulatory | protein-co | 6.35 | 6.93 | 5.35 | 1.58 | 2.01E-07 | 2.63E-05 |
| 438 | chr17 | 88581726 | 88581876 | + | 0 | NA | intron (NM | intron (NM | -44754 | NM_0298 | 77057 | Mm.48220 | NM_0298 | ENSMUSG | Ston1 | 4921524J0 | stonin 1 | protein-co | 5.37 | 6.09 | 3.82 | 2.27 | 2.10E-07 | 2.73E-05 |
| 439 | chr5 | 1.18E+08 | 1.18E+08 | + | 0 | NA | intron (NM | intron (NM | 21365 | NR_15158 | 57816 | Mm.27328 | NM_0213 | ENSMUSG | Tesc | 1010001A | tescalcin | protein-co | 6.69 | 7.26 | 5.75 | 1.5 | 2.12E-07 | 2.74E-05 |
| 444 | chr2 | 93461254 | 93461404 | + | 0 | NA | intron (NR | intron (NR | 522 | NM_0012 | 12521 | Mm.4261 | NM_0076 | ENSMUSG | Cd82 | AA682076 | CD82 anti | protein-co | 7.52 | 8.01 | 6.77 | 1.24 | 2.28E-07 | 2.93E-05 |
| 446 | chr15 | 73732060 | 73732210 | + | 0 | NA | intron (NM | intron (NM | 7205 | NM_0011 | 19245 | Mm.39080 | NM_0089 | ENSMUSG | Ptp4a3 | AV088979 | protein ty | protein-co | 7.18 | 7.66 | 6.46 | 1.2 | 2.31E-07 | 2.95E-05 |
| 448 | chr10 | 95313621 | 95313771 | + | 0 | NA | intron (NM | MER21B\|L | 10013 | NM_0013 | 12905 | Mm.47373 | NM_0099 | ENSMUSG | Cradd | RAIDD | CASP2 and | protein-co | 6.39 | 6.98 | 5.4 | 1.58 | 2.37E-07 | 3.01E-05 |
| 449 | chr8 | 1.17E+08 | 1.17E+08 | + | 0 | NA | intron (NM | intron (NM | -28321 | NM_0289 | 74440 | Mm.23318 | NM_0289 | ENSMUSG | Cmip | 4933407C | c-Maf ind | protein-co | 7.39 | 7.85 | 6.7 | 1.14 | 2.37E-07 | 3.01E-05 |
| 451 | chr10 | 81329360 | 81329510 | + | 0 | NA | intron (NM | intron (NM | 732 | NM_0093 | 21390 | Mm.4545 | NM_0093 | ENSMUSG | Tbxa2r | TP\|TXA2-F | thrombox | protein-co | 5.34 | 6.08 | 3.75 | 2.33 | 2.43E-07 | 3.06E-05 |
| 452 | chr17 | 86969745 | 86969895 | + | 0 | NA | intron (NM | intron (NM | 6709 | NM_1454 | 104215 | Mm.826 | NM_1454 | ENSMUSG | Rhoq | Arhq\|TC1 | ras homol | protein-co | 6.17 | 6.78 | 5.11 | 1.66 | 2.50E-07 | 3.13E-05 |
| 453 | chr1 | 55524115 | 55524265 | + | 0 | NA | intron (NM | intron (NM | 118244 | NM_0011 | 227120 | Mm.19418 | NM_0011 | ENSMUSG | Plcl1 | C230017K | phospholi | protein-co | 4.86 | 5.7 | 2.67 | 3.03 | 2.52E-07 | 3.16E-05 |
| 457 | chr7 | 16362188 | 16362338 | + | 0 | NA | intron (NM | intron (NM | 25633 | NM_0197 | 56459 | Mm.25853 | NM_0197 | ENSMUSG | Sae1 | 2400010M | SUMO1 ac | protein-co | 5.39 | 6.11 | 3.85 | 2.27 | 2.77E-07 | 3.44E-05 |
| 460 | chrX | 13516532 | 13516682 | + | 0 | NA | TTS (NM_ | TTS (NM_ | 55517 | NM_1734 | 236690 | Mm.90233 | NM_1734 | ENSMUSG | Nyx | CLNP\|CSN | nyctalopin | protein-co | 5.13 | 5.9 | 3.36 | 2.54 | 2.87E-07 | 3.53E-05 |
| 461 | chr7 | 24378417 | 24378567 | + | 0 | NA | intron (NM | intron (NM | 8229 | NM_0011 | 16534 | Mm.9911 | NM_0084 | ENSMUSG | Kcnn4 | IK1\|IKCA1 | potassium | protein-co | 7.1 | 7.6 | 6.33 | 1.27 | 2.94E-07 | 3.61E-05 |
| 465 | chr16 | 48594591 | 48594741 | + | 0 | NA | intron (NM | intron (NM | 163429 | NM_0108 | 17450 | Mm.25006 | NM_0108 | ENSMUSG | Morc1 | Morc | microrchid | protein-co | 7.22 | 7.69 | 6.5 | 1.19 | 3.01E-07 | 3.66E-05 |
| 468 | chr19 | 60946850 | 60947000 | + | 0 | NA | intron (NM | intron (NM | 57176 | NM_0188 | 14773 | Mm.27940 | NM_0188 | ENSMUSG | Grk5 | Gprk5 | G protein- | protein-co | 7.72 | 8.16 | 7.08 | 1.07 | 3.16E-07 | 3.82E-05 |
| 469 | chr15 | 81033252 | 81033402 | + | 0 | NA | intron (NM | intron (NM | 55562 | NM_0013 | 105835 | Mm.27494 | NM_1340 | ENSMUSG | Sgsm3 | 1810012I0 | small G pr | protein-co | 6.42 | 6.99 | 5.48 | 1.51 | 3.26E-07 | 3.93E-05 |
| 470 | chr8 | 1.06E+08 | 1.06E+08 | + | 0 | NA | promoter- | promoter- | -148 | NM_0013 | 234695 | Mm.47164 | NM_0010 | ENSMUSG | Carmil2 | CG1399-P | capping pr | protein-co | 7.23 | 7.69 | 6.53 | 1.17 | 3.28E-07 | 3.95E-05 |
| 471 | chr13 | 1.02E+08 | 1.02E+08 | + | 0 | NA | intron (NM | intron (NM | -18844 | NM_0010 | 18708 | Mm.25933 | NM_0010 | ENSMUSG | Pik3r1 | PI3K\|p50a | phosphoin | protein-co | 5.96 | 6.64 | 4.63 | 2.01 | 3.37E-07 | 4.05E-05 |
| 475 | chr13 | 1.2E+08 | 1.2E+08 | + | 0 | NA | promoter- | promoter- | -234 | NM_0010 | 625360 | Mm.32750 | NM_0010 | ENSMUSG | BC147527 | - | cDNA seq | protein-co | 6.07 | 6.69 | 4.96 | 1.72 | 3.44E-07 | 4.10E-05 |
| 476 | chr16 | 30066941 | 30067091 | + | 0 | NA | exon (NM_ | exon (NM_ | 1659 | NM_0082 | 15205 | Mm.39085 | NM_0082 | ENSMUSG | Hes1 | Hry\|bHLH | hes family | protein-co | 7.45 | 7.89 | 6.82 | 1.07 | 3.56E-07 | 4.23E-05 |
| 480 | chr19 | 53893706 | 53893856 | + | 0 | NA | intron (NM | intron (NM | 1550 | NM_0110 | 18569 | Mm.1605 | NM_0110 | ENSMUSG | Pdcd4 | D19Ucla1\| | programm | protein-co | 6.72 | 7.25 | 5.88 | 1.38 | 3.91E-07 | 4.59E-05 |
| 481 | chr10 | 60284770 | 60284920 | + | 0 | NA | intron (NM | intron (NM | 384 | NM_0011 | 19156 | Mm.29978 | NM_0111 | ENSMUSG | Psap | AI037048\| | prosaposi | protein-co | 5.49 | 6.23 | 3.87 | 2.36 | 4.01E-07 | 4.69E-05 |

| | | | | | | | | | | | | | | | | | | | | | | | | |
|---|---|---|---|---|---|---|---|---|---|---|---|---|---|---|---|---|---|---|---|---|---|---|---|---|
| 483 | chr5 | 92499583 | 92499733 | + | 0 | NA | intron (NM | intron (NM | 5999 | NM_00764 | 12492 | Mm.29796 | NM_00764 | ENSMUSG | Scarb2 | 9330185J1 | scavenger | protein-co | 6.93 | 7.43 | 6.15 | 1.28 | 4.02E-07 | 4.69E-05 |
| 484 | chr16 | 94519190 | 94519340 | + | 0 | NA | intron (NM | intron (NM | 7364 | NM_00783 | 13185 | Mm.7472 | NM_00783 | ENSMUSG | Vps26c | AW53812S | VPS26 end | protein-co | 5.86 | 6.5 | 4.69 | 1.81 | 4.12E-07 | 4.79E-05 |
| 485 | chr5 | 34196504 | 34196654 | + | 0 | NA | exon (NM_ | exon (NM_ | -8869 | NM_01079 | 17122 | Mm.25926 | NM_01079 | ENSMUSG | Mxd4 | 2810410A | Max dimer | protein-co | 7.59 | 8.01 | 6.98 | 1.03 | 4.14E-07 | 4.80E-05 |
| 486 | chr7 | 1.26E+08 | 1.26E+08 | + | 0 | NA | promoter- | promoter- | 18 | NM_02188 | 60504 | Mm.15564 | NM_02188 | ENSMUSG | Il21r | NILR | interleukir | protein-co | 7.45 | 7.89 | 6.8 | 1.09 | 4.21E-07 | 4.87E-05 |
| 487 | chr5 | 1.44E+08 | 1.44E+08 | + | 0 | NA | intron (NM | intron (NM | -12173 | NM_17768 | 231874 | Mm.32043 | NM_17768 | ENSMUSG | Ccz1 | AU022870 | CCZ1 vacu | protein-co | 5.65 | 6.32 | 4.32 | 2 | 4.24E-07 | 4.90E-05 |
| 488 | chr10 | 93876876 | 93877026 | + | 0 | NA | intron (NM | RSINE1\|SI | 14251 | NM_01964 | 56307 | Mm.28932 | NM_01964 | ENSMUSG | Metap2 | 4930584B | methionin | protein-co | 6.37 | 6.96 | 5.38 | 1.58 | 4.26E-07 | 4.91E-05 |
| 489 | chr4 | 89369239 | 89369389 | + | 0 | NA | intron (NR | intron (NR | -58282 | NM_00767 | 12579 | Mm.42309 | NM_00767 | ENSMUSG | Cdkn2b | AV083695 | cyclin depe | protein-co | 5.43 | 6.13 | 4 | 2.14 | 4.27E-07 | 4.91E-05 |
| 490 | chr11 | 1.17E+08 | 1.17E+08 | + | 0 | NA | intron (NM | intron (NM | -4505 | NM_01738 | 53860 | Mm.38450 | NM_01738 | ENSMUSG | 9-Sep | MSF1\|Msf | septin 9 | protein-co | 6.54 | 7.09 | 5.63 | 1.46 | 4.40E-07 | 5.05E-05 |
| 491 | chr7 | 43791010 | 43791160 | + | 0 | NA | promoter- | promoter- | -806 | NM_02866 | 101533 | Mm.19942 | NM_02866 | ENSMUSG | Klk9 | 1200016C | kallikrein r | protein-co | 7 | 6.26 | 7.48 | -1.22 | 4.47E-07 | 5.13E-05 |
| 494 | chr2 | 1.03E+08 | 1.03E+08 | + | 0 | NA | intron (NM | intron (NM | 34027 | NM_00117 | 12505 | Mm.42362 | NM_00989 | ENSMUSG | Cd44 | AU023126 | CD44 antig | protein-co | 6.18 | 6.78 | 5.13 | 1.65 | 4.60E-07 | 5.24E-05 |
| 496 | chr2 | 6376193 | 6376343 | + | 0 | NA | intron (NM | intron (NM | 23532 | NM_18139 | 98910 | Mm.40673 | NM_18139 | ENSMUSG | Usp6nl | AI316785\| | USP6 N-te | protein-co | 7.07 | 7.56 | 6.31 | 1.25 | 4.78E-07 | 5.43E-05 |
| 498 | chr15 | 1.02E+08 | 1.02E+08 | + | 0 | NA | intron (NM | intron (NM | 6545 | NM_01356 | 16421 | Mm.58 | NM_01356 | ENSMUSG | Itgb7 | Ly69 | integrin be | protein-co | 8.13 | 8.55 | 7.53 | 1.02 | 4.84E-07 | 5.47E-05 |
| 500 | chr1 | 78538489 | 78538639 | + | 0 | NA | TTS (NM_( | TTS (NM_( | 27504 | NM_02671 | 68393 | Mm.41325 | NM_02671 | ENSMUSG | Mogat1 | 0610030A | monoacylg | protein-co | 5.84 | 6.49 | 4.62 | 1.87 | 4.98E-07 | 5.60E-05 |
| 501 | chrX | 13576759 | 13576909 | + | 0 | NA | intron (NM | intron (NM | -55935 | NM_01182 | 23890 | Mm.48732 | NM_01182 | ENSMUSG | Gpr34 | Lypsr1 | G protein- | protein-co | 5.52 | 6.25 | 4 | 2.25 | 4.98E-07 | 5.60E-05 |
| 502 | chr17 | 6063651 | 6063801 | + | 0 | NA | intron (NM | intron (NM | 16013 | NM_00111 | 321007 | Mm.5548 | NM_17731 | ENSMUSG | Serac1 | 4930511N | serine acti | protein-co | 5.28 | 6.02 | 3.71 | 2.31 | 5.02E-07 | 5.61E-05 |
| 505 | chr7 | 1.16E+08 | 1.16E+08 | + | 0 | NA | intron (NM | intron (NM | 6402 | NM_00130 | 233765 | Mm.3741 | NM_17274 | ENSMUSG | Plekha7 | A430081P | pleckstrin | protein-co | 7.56 | 8.01 | 6.92 | 1.09 | 5.43E-07 | 6.03E-05 |
| 507 | chr11 | 60130619 | 60130769 | + | 0 | NA | intron (NM | intron (NM | -9388 | NM_00103 | 19377 | Mm.29636 | NM_00902 | ENSMUSG | Rai1 | Gt1 | retinoic ac | protein-co | 6.36 | 6.93 | 5.41 | 1.52 | 5.57E-07 | 6.14E-05 |
| 509 | chr5 | 1.4E+08 | 1.4E+08 | + | 0 | NA | intron (NM | intron (NM | 56215 | NM_00135 | 17120 | Mm.27250 | NM_01079 | ENSMUSG | Mad1l1 | AI173502\| | MAD1 mit | protein-co | 6 | 6.61 | 4.9 | 1.71 | 5.73E-07 | 6.28E-05 |
| 510 | chr9 | 1.21E+08 | 1.21E+08 | + | 0 | NA | intron (NM | intron (NM | -5849 | NM_17511 | 67095 | Mm.49111 | NM_17511 | ENSMUSG | Trak1 | 2310001H | trafficking | protein-co | 7.35 | 7.81 | 6.69 | 1.11 | 5.73E-07 | 6.28E-05 |
| 513 | chr8 | 27159344 | 27159494 | + | 0 | NA | intron (NM | intron (NM | 15227 | NM_02942 | 75767 | Mm.20533 | NM_02942 | ENSMUSG | Rab11fip1 | 2010200K | RAB11 fam | protein-co | 7.31 | 7.78 | 6.61 | 1.16 | 5.93E-07 | 6.44E-05 |
| 515 | chr2 | 1.59E+08 | 1.59E+08 | + | 0 | NA | intron (NM | ID_B1\|SIN | -48376 | NM_02797 | 71878 | Mm.85162 | NM_02797 | ENSMUSG | Fam83d | 2310007D | family with | protein-co | 6.74 | 7.26 | 5.93 | 1.33 | 5.99E-07 | 6.47E-05 |
| 518 | chr5 | 1.15E+08 | 1.15E+08 | + | 0 | NA | intron (NM | intron (NM | 26740 | NM_00100 | 433940 | Mm.19421 | NM_00100 | ENSMUSG | Fam222a | - | family with | protein-co | 5.71 | 6.38 | 4.41 | 1.98 | 6.22E-07 | 6.69E-05 |
| 520 | chr18 | 39519426 | 39519576 | + | 0 | NA | promoter- | promoter- | -80 | NM_00136 | 14815 | Mm.12948 | NM_00817 | ENSMUSG | Nr3c1 | GR\|Grl-1\| | nuclear rec | protein-co | 7.83 | 8.24 | 7.23 | 1.01 | 6.77E-07 | 7.22E-05 |
| 521 | chr11 | 54778528 | 54778678 | + | 0 | NA | intron (NM | intron (NM | 9100 | NM_17862 | 72729 | Mm.29476 | NM_17862 | ENSMUSG | Cdc42se2 | 2810404F | CDC42 sma | protein-co | 5.86 | 6.49 | 4.7 | 1.79 | 6.82E-07 | 7.27E-05 |
| 522 | chr13 | 1.02E+08 | 1.02E+08 | + | 0 | NA | intron (NM | intron (NM | -26937 | NM_00102 | 18708 | Mm.25933 | NM_00102 | ENSMUSG | Pik3r1 | PI3K\|p50a | phosphoin | protein-co | 6.94 | 7.43 | 6.19 | 1.24 | 6.95E-07 | 7.39E-05 |
| 524 | chr15 | 86094447 | 86094597 | + | 0 | NA | intron (NM | intron (NM | 35795 | NM_00120 | 223752 | Mm.24442 | NM_17261 | ENSMUSG | Gramd4 | 9930016O | GRAM dor | protein-co | 6.61 | 7.14 | 5.76 | 1.38 | 7.04E-07 | 7.46E-05 |
| 528 | chr14 | 8128126 | 8128276 | + | 0 | NA | intron (NM | ORR1D2\|L | 29990 | NM_17827 | 218699 | Mm.29515 | NM_14549 | ENSMUSG | Pxk | C230080L | PX domain | protein-co | 7.62 | 6.98 | 8.07 | -1.09 | 7.17E-07 | 7.55E-05 |
| 526 | chr10 | 1.28E+08 | 1.28E+08 | + | 0 | NA | promoter- | promoter- | -570 | NM_05407 | 116848 | Mm.25221 | NM_05407 | ENSMUSG | Baz2a | AA415431 | bromodon | protein-co | 7.23 | 7.69 | 6.53 | 1.16 | 7.15E-07 | 7.55E-05 |
| 525 | chr6 | 34156207 | 34156357 | + | 0 | NA | 3' UTR (NM | 3' UTR (NM | 20772 | NM_02143 | 58246 | Mm.24552 | NM_02143 | ENSMUSG | Slc35b4 | 4930474D | solute carr | protein-co | 7.13 | 7.6 | 6.44 | 1.16 | 7.14E-07 | 7.55E-05 |
| 530 | chr8 | 12957146 | 12957296 | + | 0 | NA | intron (NM | intron (NM | 7798 | NM_00119 | 17207 | Mm.33441 | NM_17807 | ENSMUSG | Mcf2l | C130040G | mcf.2 tran | protein-co | 6.34 | 6.91 | 5.4 | 1.5 | 7.43E-07 | 7.78E-05 |
| 538 | chr2 | 69002414 | 69002564 | + | 0 | NA | intron (NM | intron (NM | -133311 | NM_18154 | 329416 | Mm.90047 | NM_18154 | ENSMUSG | Nostrin | Daip2\|mD | nitric oxide | protein-co | 5.54 | 6.23 | 4.2 | 2.02 | 7.97E-07 | 8.21E-05 |
| 539 | chr11 | 1.2E+08 | 1.2E+08 | + | 0 | NA | intron (NM | intron (NM | 11845 | NM_00136 | 72055 | Mm.25340 | NM_02424 | ENSMUSG | Slc38a10 | 1810073N | solute carr | protein-co | 7.34 | 7.79 | 6.7 | 1.09 | 8.52E-07 | 8.75E-05 |
| 548 | chr12 | 85969672 | 85969822 | + | 0 | NA | intron (NM | intron (NM | 109294 | NM_00936 | 21809 | Mm.3992 | NM_00936 | ENSMUSG | Tgfb3 | TGF-beta-3 | transformi | protein-co | 6.7 | 7.21 | 5.91 | 1.3 | 9.55E-07 | 9.66E-05 |
| 549 | chr5 | 1.12E+08 | 1.12E+08 | + | 0 | NA | intron (NM | intron (NM | 18422 | NM_00135 | 72898 | Mm.15174 | NM_02838 | ENSMUSG | Asphd2 | 2900006N | aspartate l | protein-co | 5.01 | 5.77 | 3.31 | 2.46 | 9.76E-07 | 9.85E-05 |
| 551 | chr14 | 60709328 | 60709478 | + | 0 | NA | intron (NM | intron (NM | -23501 | NM_00113 | 219140 | Mm.14977 | NM_00103 | ENSMUSG | Spata13 | C79050\|E | spermatog | protein-co | 7.25 | 7.71 | 6.58 | 1.13 | 9.86E-07 | 9.92E-05 |
| 554 | chr4 | 1.55E+08 | 1.55E+08 | + | 0 | NA | intron (NM | intron (NM | -6949 | NM_00103 | 18762 | Mm.28561 | NM_00880 | ENSMUSG | Prkcz | AI098070\| | protein kir | protein-co | 5.08 | 5.86 | 3.25 | 2.61 | 1.02E-06 | 1.02E-04 |
| 553 | chr8 | 40577608 | 40577758 | + | 0 | NA | intron (NM | MLT1E\|LT | 57109 | NM_00104 | 54384 | Mm.29487 | NM_01943 | ENSMUSG | Mtmr7 | - | myotubula | protein-co | 6.24 | 6.82 | 5.26 | 1.57 | 1.01E-06 | 1.02E-04 |
| 556 | chr2 | 27122121 | 27122271 | + | 0 | NA | intron (NM | intron (NM | 20281 | NM_17542 | 109349 | Mm.31408 | NM_17542 | ENSMUSG | Fam163b | C630035N | family with | protein-co | 4.95 | 5.72 | 3.17 | 2.54 | 1.04E-06 | 1.04E-04 |
| 558 | chr2 | 3731375 | 3731525 | + | 0 | NA | intron (NM | intron (NM | 17992 | NM_02562 | 66540 | Mm.27786 | NM_02562 | ENSMUSG | Fam107b | 3110001A | family with | protein-co | 6.73 | 7.25 | 5.93 | 1.31 | 1.06E-06 | 1.05E-04 |
| 557 | chr2 | 1.61E+08 | 1.61E+08 | + | 0 | NA | intron (NM | intron (NM | 2112 | NM_02128 | 18803 | Mm.44463 | NM_02128 | ENSMUSG | Plcg1 | AI894140\| | phospholi | protein-co | 6.65 | 7.18 | 5.82 | 1.36 | 1.05E-06 | 1.05E-04 |
| 559 | chr7 | 1.42E+08 | 1.42E+08 | + | 0 | NA | intron (NM | intron (NM | 6452 | NM_00998 | 13033 | Mm.23139 | NM_00998 | ENSMUSG | Ctsd | CD\|CatD | cathepsin | protein-co | 7.51 | 7.94 | 6.88 | 1.07 | 1.06E-06 | 1.05E-04 |
| 563 | chr2 | 3717364 | 3717514 | + | 0 | NA | intron (NM | intron (NM | 3981 | NM_02562 | 66540 | Mm.27786 | NM_02562 | ENSMUSG | Fam107b | 3110001A | family with | protein-co | 7.07 | 7.55 | 6.36 | 1.19 | 1.11E-06 | 1.09E-04 |
| 566 | chr12 | 87196492 | 87196642 | + | 0 | NA | intron (NM | intron (NM | 3662 | NM_00103 | 382620 | Mm.37491 | NM_00103 | ENSMUSG | Tmed8 | 6430595O | transmem | protein-co | 5.18 | 5.91 | 3.61 | 2.3 | 1.14E-06 | 1.12E-04 |
| 564 | chr4 | 1.27E+08 | 1.27E+08 | + | 0 | NA | intron (NM | intron (NM | 61302 | NM_13388 | 100317 | Mm.20620 | NM_13388 | ENSMUSG | AU040320 | A730047D | expressed | protein-co | 5.87 | 6.49 | 4.74 | 1.76 | 1.14E-06 | 1.12E-04 |
| 568 | chr14 | 25614440 | 25614590 | + | 0 | NA | intron (NM | MIRb\|SINE | 11002 | NM_00130 | 1E+08 | Mm.47381 | NM_001309430 | | Gm45915 | Israa | predicted | protein-co | 5.98 | 6.59 | 4.9 | 1.69 | 1.16E-06 | 1.13E-04 |
| 567 | chr4 | 1.24E+08 | 1.24E+08 | + | 0 | NA | intron (NM | ORR1E\|LT | 7220 | NM_02342 | 68050 | Mm.27903 | NM_02342 | ENSMUSG | Akirin1 | 6330407G | akirin 1 | protein-co | 6.28 | 6.86 | 5.32 | 1.53 | 1.16E-06 | 1.13E-04 |
| 573 | chr14 | 31531667 | 31531817 | + | 0 | NA | intron (NM | intron (NM | 36663 | NM_02893 | 74427 | Mm.49120 | NM_02893 | ENSMUSG | Eaf1 | 4933403C | ELL associ | protein-co | 5.03 | 5.78 | 3.36 | 2.42 | 1.21E-06 | 1.17E-04 |
| 572 | chr15 | 6588238 | 6588388 | + | 0 | NA | intron (NM | intron (NM | 8466 | NM_01181 | 23880 | Mm.17090 | NM_01181 | ENSMUSG | Fyb | ADAP\|B63 | FYN bindir | protein-co | 6.89 | 7.4 | 6.08 | 1.32 | 1.21E-06 | 1.17E-04 |
| 578 | chr19 | 45236656 | 45236806 | + | 0 | NA | 5' UTR (NM | 5' UTR (NM | 110 | NM_01069 | 16814 | Mm.11681 | NM_01069 | ENSMUSG | Lbx1 | Lbx1h | ladybird h | protein-co | 6.56 | 7.08 | 5.73 | 1.36 | 1.23E-06 | 1.18E-04 |
| 579 | chr3 | 1.27E+08 | 1.27E+08 | + | 0 | NA | intron (NM | L2\|LINE\|L | 66738 | NM_00134 | 108058 | Mm.25582 | NM_02381 | ENSMUSG | Camk2d | 2810011D | calcium/ca | protein-co | 6.42 | 6.96 | 5.55 | 1.41 | 1.23E-06 | 1.18E-04 |
| 575 | chr3 | 1.54E+08 | 1.54E+08 | + | 0 | NA | intron (NM | intron (NM | 9231 | NM_00128 | 55993 | Mm.27222 | NM_03187 | ENSMUSG | Msh4 | 4930485C | mutS hom | protein-co | 6.56 | 7.11 | 5.67 | 1.44 | 1.22E-06 | 1.18E-04 |
| 576 | chr5 | 1.48E+08 | 1.48E+08 | + | 0 | NA | promoter- | promoter- | -898 | NM_00135 | 71706 | Mm.15321 | NM_02787 | ENSMUSG | Slc46a3 | 1200006F | solute carr | protein-co | 6.5 | 7.03 | 5.65 | 1.39 | 1.23E-06 | 1.18E-04 |
| 581 | chr8 | 1.2E+08 | 1.2E+08 | + | 0 | NA | promoter- | promoter- | -14 | NM_00131 | 78892 | Mm.26468 | NM_03020 | ENSMUSG | Crispld2 | 1810049K | cysteine-ri | protein-co | 6.88 | 7.38 | 6.12 | 1.27 | 1.25E-06 | 1.19E-04 |
| 583 | chrX | 48136988 | 48137138 | + | 0 | NA | TTS (NM_1 | TTS (NM_1 | -9373 | NM_02877 | 74131 | Mm.27613 | NM_02877 | ENSMUSG | Sash3 | 1200013B | SAM and S | protein-co | 6.8 | 7.3 | 6.04 | 1.25 | 1.28E-06 | 1.22E-04 |
| 586 | chr11 | 98493126 | 98493276 | + | 0 | NA | intron (NM | intron (NM | 46367 | NM_01034 | 14786 | Mm.27670 | NM_01034 | ENSMUSG | Grb7 | mKIAA402 | growth fac | protein-co | 5.39 | 6.09 | 3.94 | 2.15 | 1.33E-06 | 1.26E-04 |
| 588 | chr7 | 1.11E+08 | 1.11E+08 | + | 0 | NA | intron (NM | MTD\|LTR\| | 4269 | NM_19446 | 17540 | Mm.25002 | NM_01082 | ENSMUSG | Mrvi1 | BB115629 | MRV integ | protein-co | 5.7 | 6.36 | 4.47 | 1.89 | 1.36E-06 | 1.28E-04 |
| 589 | chr10 | 1.27E+08 | 1.27E+08 | + | 0 | NA | intron (NM | L1M2\|LINI | 13454 | NM_00117 | 216440 | Mm.29524 | NM_17761 | ENSMUSG | Os9 | 4632413K | amplified i | protein-co | 6.87 | 7.36 | 6.11 | 1.25 | 1.39E-06 | 1.30E-04 |
| 590 | chr17 | 84419888 | 84420038 | + | 0 | NA | intron (NM | intron (NM | 46245 | NM_18302 | 240174 | Mm.24676 | NM_17782 | ENSMUSG | Thada | 6530405K | thyroid ad | protein-co | 6.89 | 7.37 | 6.15 | 1.22 | 1.39E-06 | 1.31E-04 |
| 597 | chr1 | 36831456 | 36831606 | + | 0 | NA | intron (NM | intron (NM | 52547 | NM_00128 | 22637 | Mm.8038 | NM_00951 | ENSMUSG | Zap70 | Srk\|ZAP-7 | zeta-chain | protein-co | 5.11 | 5.85 | 3.51 | 2.34 | 1.48E-06 | 1.37E-04 |
| 598 | chr19 | 18619790 | 18619940 | + | 0 | NA | intron (NM | intron (NM | 11948 | NM_01737 | 20409 | Mm.17222 | NM_01737 | ENSMUSG | Ostf1 | C78236\|S | osteoclast | protein-co | 6.76 | 7.28 | 5.93 | 1.34 | 1.49E-06 | 1.38E-04 |
| 599 | chr7 | 99674936 | 99675086 | + | 0 | NA | intron (NM | intron (NM | 20798 | NM_00129 | 101488 | Mm.11249 | NM_17531 | ENSMUSG | Slco2b1 | AI060904\| | solute carr | protein-co | 6.54 | 7.07 | 5.69 | 1.38 | 1.50E-06 | 1.38E-04 |
| 600 | chr2 | 1.04E+08 | 1.04E+08 | + | 0 | NA | intron (NM | intron (NM | 78184 | NM_15312 | 98956 | Mm.28800 | NM_15312 | ENSMUSG | Nat10 | AI429152 | N-acetyltr | protein-co | 7.24 | 7.72 | 6.53 | 1.19 | 1.50E-06 | 1.39E-04 |
| 605 | chr1 | 1.6E+08 | 1.6E+08 | + | 0 | NA | intron (NM | intron (NM | -202051 | NM_00116 | 67647 | Mm.17151 | NM_00102 | ENSMUSG | 4930523C( | - | RIKEN cDN | protein-co | 6.08 | 5.1 | 6.65 | -1.55 | 1.59E-06 | 1.46E-04 |
| 607 | chr1 | 1.83E+08 | 1.83E+08 | + | 0 | NA | intron (NM | intron (NM | 2148 | NM_00979 | 12334 | Mm.19306 | NM_00979 | ENSMUSG | Capn2 | AI326419\| | calpain 2 | protein-co | 6.8 | 7.29 | 6.04 | 1.25 | 1.62E-06 | 1.48E-04 |
| 611 | chr10 | 5384256 | 5384406 | + | 0 | NA | intron (NM | intron (NM | 103983 | NM_00134 | 64009 | Mm.33162 | NM_02202 | ENSMUSG | Syne1 | 8B\|A3300 | spectrin re | protein-co | 5.66 | 6.31 | 4.47 | 1.84 | 1.68E-06 | 1.53E-04 |
| 610 | chr5 | 1.49E+08 | 1.49E+08 | + | 0 | NA | intron (NM | intron (NM | -11127 | NM_02946 | 75860 | Mm.15960 | NM_02946 | ENSMUSG | Tex26 | 4930588N | testis expr | protein-co | 6.59 | 7.12 | 5.76 | 1.36 | 1.68E-06 | 1.53E-04 |
| 612 | chr16 | 18073751 | 18073901 | + | 0 | NA | intron (NM | intron (NM | 7423 | NR_11038 | 13353 | Mm.27155 | NM_01004 | ENSMUSG | Dgcr6 | - | DiGeorge s | protein-co | 5.63 | 6.3 | 4.35 | 1.95 | 1.72E-06 | 1.56E-04 |
| 614 | chr1 | 92069595 | 92069745 | + | 0 | NA | intron (NM | intron (NM | 110671 | NM_20722 | 208727 | Mm.31856 | NM_20722 | ENSMUSG | Hdac4 | 4932408F | histone de | protein-co | 5.04 | 5.78 | 3.44 | 2.34 | 1.78E-06 | 1.61E-04 |
| 620 | chr5 | 43184938 | 43185088 | + | 0 | NA | intron (NR | ORR1E\|LT | -48240 | NM_17593 | 231207 | Mm.7233 | NM_17593 | ENSMUSG | Cpeb2 | A630055H | cytoplasm | protein-co | 4.92 | 5.71 | 3.04 | 2.67 | 1.91E-06 | 1.70E-04 |
| 622 | chr15 | 66812132 | 66812282 | + | 0 | NA | intron (NM | MIRb\|SINE | 19622 | NM_00919 | 20491 | Mm.7601 | NM_00919 | ENSMUSG | Sla | Slap\|Slap- | src-like ad | protein-co | 6.44 | 6.97 | 5.59 | 1.38 | 1.97E-06 | 1.75E-04 |
| 624 | chr2 | 93192673 | 93192823 | + | 0 | NA | intron (NM | intron (NM | 5200 | NM_00135 | 277414 | Mm.41033 | NM_00102 | ENSMUSG | Trp53i11 | Tp53i11 | transform | protein-co | 7.99 | 8.61 | 6.87 | 1.74 | 1.98E-06 | 1.76E-04 |
| 625 | chr14 | 27229621 | 27229771 | + | 0 | NA | intron (NM | intron (NM | -8314 | NM_02787 | 71704 | Mm.24860 | NM_02787 | ENSMUSG | Arhgef3 | 1200004I2 | Rho guani | protein-co | 6.08 | 6.67 | 5.07 | 1.6 | 2.01E-06 | 1.77E-04 |
| 627 | chr1 | 66694370 | 66694520 | + | 0 | NA | exon (NM_ | exon (NM_ | -6386 | NM_00131 | 66646 | Mm.24091 | NM_02568 | ENSMUSG | Rpe | 2610429B | ribulose-5- | protein-co | 5.56 | 6.21 | 4.33 | 1.88 | 2.05E-06 | 1.81E-04 |
| 628 | chr5 | 1.39E+08 | 1.39E+08 | + | 0 | NA | exon (NM_ | exon (NM_ | -11140 | NM_00135 | 19085 | Mm.30616 | NM_00892 | ENSMUSG | Prkar1b | AI385716\| | protein kir | protein-co | 4.86 | 5.65 | 2.97 | 2.67 | 2.12E-06 | 1.87E-04 |
| 629 | chr8 | 71587057 | 71587207 | + | 0 | NA | 3' UTR (NM | 3' UTR (NM | -5052 | NM_00129 | 66171 | Mm.28228 | NM_02539 | ENSMUSG | Pgls | 1110030K | 6-phospho | protein-co | 5.74 | 6.38 | 4.55 | 1.83 | 2.13E-06 | 1.87E-04 |
| 631 | chr6 | 1.16E+08 | 1.16E+08 | + | 0 | NA | intron (NM | intron (NM | 20578 | NM_00129 | 67492 | Mm.27277 | NM_00108 | ENSMUSG | Zfand4 | Anubl1 | zinc finger | protein-co | 6.44 | 6.98 | 5.59 | 1.39 | 2.18E-06 | 1.91E-04 |
| 637 | chr1 | 1.27E+08 | 1.27E+08 | + | 0 | NA | intron (NM | intron (NM | 97928 | NM_14512 | 107895 | Mm.21468 | NM_14512 | ENSMUSG | Mgat5 | 4930471A | mannoside | protein-co | 5.87 | 6.48 | 4.8 | 1.68 | 2.29E-06 | 1.98E-04 |
| 638 | chr16 | 33687822 | 33687972 | + | 0 | NA | intron (NM | intron (NM | 3431 | NM_17525 | 77446 | Mm.24574 | NM_17299 | ENSMUSG | Heg1 | 4632417D | heart deve | protein-co | 6.66 | 7.18 | 5.86 | 1.31 | 2.31E-06 | 2.00E-04 |
| 639 | chr15 | 1.01E+08 | 1.01E+08 | + | 0 | NA | intron (NM | intron (NM | 5427 | NM_01687 | 53622 | Mm.34793 | NM_016879 | | Krt85 | AA589597 | keratin 85 | protein-co | 5.6 | 6.25 | 4.38 | 1.87 | 2.34E-06 | 2.02E-04 |
| 642 | chr3 | 1.31E+08 | 1.31E+08 | + | 0 | NA | intron (NM | intron (NM | 41961 | NM_00127 | 16842 | Mm.25521 | NM_01070 | ENSMUSG | Lef1 | 3000002B | lymphoid | protein-co | 5.38 | 6.06 | 4.05 | 2.01 | 2.45E-06 | 2.09E-04 |
| 645 | chr3 | 1.54E+08 | 1.54E+08 | + | 0 | NA | exon (NM_ | exon (NM_ | 7474 | NM_00128 | 55993 | Mm.27222 | NM_03187 | ENSMUSG | Msh4 | 4930485C | mutS hom | protein-co | 5.93 | 6.54 | 4.85 | 1.69 | 2.66E-06 | 2.26E-04 |
| 648 | chr3 | 89437395 | 89437545 | + | 0 | NA | intron (NM | intron (NM | 766 | NM_14613 | 229534 | Mm.65906 | NM_14613 | ENSMUSG | Pbxip1 | 4732463H | pre B cell l | protein-co | 6.56 | 7.07 | 5.77 | 1.3 | 2.74E-06 | 2.32E-04 |
| 650 | chr9 | 98574088 | 98574238 | + | 0 | NA | exon (NM_ | exon (NM_ | 10432 | NM_01582 | 50797 | Mm.40046 | NM_01582 | ENSMUSG | Copb2 | AI256832 | coatomer | protein-co | 6.25 | 5.17 | 6.86 | -1.69 | 2.82E-06 | 2.38E-04 |
| 649 | chr16 | 45786804 | 45786954 | + | 0 | NA | intron (NM | intron (NM | -43924 | NM_17251 | 213012 | Mm.24745 | NM_17251 | ENSMUSG | Abhd10 | - | abhydrola | protein-co | 7.33 | 7.77 | 6.67 | 1.1 | 2.82E-06 | 2.38E-04 |
| 653 | chr11 | 1.15E+08 | 1.15E+08 | + | 0 | NA | intron (NM | intron (NM | 3542 | NM_00134 | 68572 | Mm.27106 | NM_02672 | ENSMUSG | Mrpl58 | 1110001A | mitochonc | protein-co | 5.87 | 6.5 | 4.73 | 1.77 | 2.95E-06 | 2.48E-04 |
| 654 | chr5 | 97983487 | 97983637 | + | 0 | NA | intron (NM | B3\|SINE\|E | 47455 | NM_13373 | 71914 | Mm.24842 | NM_13373 | ENSMUSG | Antxr2 | 2310046B | anthrax to | protein-co | 5.77 | 6.4 | 4.64 | 1.76 | 3.01E-06 | 2.53E-04 |
| 655 | chr15 | 98776421 | 98776571 | + | 0 | NA | intron (NM | CpG | 1654 | NM_01171 | 22410 | Mm.4709 | NM_01171 | ENSMUSG | Wnt10b | Wnt12 | wingless-t | protein-co | 6.5 | 7.03 | 5.66 | 1.37 | 3.06E-06 | 2.56E-04 |
| 656 | chr10 | 1.18E+08 | 1.18E+08 | + | 0 | NA | intron (NM | intron (NM | 24986 | NM_01078 | 17245 | Mm.10119 | NM_01078 | ENSMUSG | Mdm1 | Arrd2\|Md | transform | protein-co | 4.55 | 5.37 | 2.5 | 2.87 | 3.10E-06 | 2.59E-04 |
| 659 | chr8 | 15018219 | 15018369 | + | 0 | NA | intron (NM | intron (NM | 7269 | NM_02911 | 74901 | Mm.46675 | NM_02911 | ENSMUSG | Kbtbd11 | 2900016B | kelch repe | protein-co | 5.85 | 6.45 | 4.78 | 1.68 | 3.30E-06 | 2.74E-04 |
| 661 | chr9 | 32706215 | 32706365 | + | 0 | NA | intron (NM | intron (NM | 10248 | NM_00103 | 23871 | Mm.29241 | NM_01180 | ENSMUSG | Ets1 | AI196000\| | E26 avian l | protein-co | 6.82 | 7.3 | 6.1 | 1.2 | 3.42E-06 | 2.83E-04 |
| 662 | chr3 | 1.53E+08 | 1.53E+08 | + | 0 | NA | intron (NM | intron (NM | 53393 | NM_01202 | 26938 | Mm.40915 | NM_01202 | ENSMUSG | St6galnac5 | AI851940\| | ST6 (alpha | protein-co | 6.8 | 7.31 | 6 | 1.31 | 3.45E-06 | 2.85E-04 |
| 669 | chr11 | 1.17E+08 | 1.17E+08 | + | 0 | NA | intron (NM | intron (NM | 26368 | NM_00111 | 53860 | Mm.38450 | NM_01738 | ENSMUSG | 9-Sep | MSF1\|Msf | septin 9 | protein-co | 7.24 | 7.68 | 6.61 | 1.07 | 3.87E-06 | 3.16E-04 |
| 668 | chr9 | 44047158 | 44047308 | + | 0 | NA | intron (NM | intron (NM | 3849 | NM_00938 | 21838 | Mm.3951 | NM_00938 | ENSMUSG | Thy1 | CD90\|T25 | thymus ce | protein-co | 5.16 | 5.86 | 3.71 | 2.15 | 3.87E-06 | 3.16E-04 |
| 670 | chr6 | 22019818 | 22019968 | + | 0 | NA | intron (NM | intron (NM | 33983 | NM_00108 | 214642 | Mm.22140 | NM_00108 | ENSMUSG | Cped1 | 6720481P | cadherin-li | protein-co | 5.46 | 6.12 | 4.23 | 1.89 | 3.92E-06 | 3.20E-04 |
| 671 | chr4 | 1.49E+08 | 1.49E+08 | + | 0 | NA | exon (NM_ | exon (NM_ | -7687 | NM_00136 | 1.03E+08 | | NM_001366402 | | Tmem274 | Gm13097 | transmem | protein-co | 4.47 | 5.3 | 2.27 | 3.03 | 3.97E-06 | 3.23E-04 |
| 672 | chr3 | 1.42E+08 | 1.42E+08 | + | 0 | NA | intron (NM | intron (NM | 152789 | NM_00135 | 12167 | Mm.39089 | NM_00750 | ENSMUSG | Bmpr1b | AI385617\| | bone morp | protein-co | 6.27 | 6.84 | 5.32 | 1.52 | 4.00E-06 | 3.25E-04 |
| 675 | chr1 | 1.36E+08 | 1.36E+08 | + | 0 | NA | intron (NM | intron (NM | 30577 | NM_17343 | 215690 | Mm.34977 | NM_17343 | ENSMUSG | Nav1 | 9530089B | neuron na | protein-co | 6.84 | 7.31 | 6.14 | 1.17 | 4.05E-06 | 3.28E-04 |
| 674 | chr2 | 1.28E+08 | 1.28E+08 | + | 0 | NA | intron (NR | intron (NR | 2933 | NM_00129 | 56461 | Mm.31529 | NM_01978 | ENSMUSG | Kcnip3 | 4933407H | Kv channe | protein-co | 5.8 | 6.41 | 4.73 | 1.68 | 4.05E-06 | 3.28E-04 |
| 676 | chr3 | 59149768 | 59149918 | + | 0 | NA | non-codin | non-codin | 3649 | NM_00128 | 140795 | Mm.43902 | NM_13320 | ENSMUSG | P2ry14 | A330108O | purinergic | protein-co | 6.36 | 6.9 | 5.49 | 1.41 | 4.06E-06 | 3.28E-04 |
| 677 | chr8 | 1.2E+08 | 1.2E+08 | + | 0 | NA | 3' UTR (NM | 3' UTR (NM | 51090 | NM_13396 | 102193 | Mm.24007 | NM_13396 | ENSMUSG | Zdhhc7 | AL024087\| | zinc finger | protein-co | 6.06 | 6.63 | 5.11 | 1.52 | 4.07E-06 | 3.28E-04 |
| 679 | chr3 | 52000200 | 52000350 | + | 0 | NA | intron (NM | intron (NM | 104731 | NM_00100 | 433586 | Mm.23497 | NM_00100 | ENSMUSG | Maml3 | AV234550 | mastermir | protein-co | 7.32 | 7.74 | 6.71 | 1.03 | 4.13E-06 | 3.32E-04 |
| 683 | chr10 | 81538735 | 81538885 | + | 0 | NA | intron (NM | intron (NM | 6352 | NM_01030 | 14672 | Mm.26092 | NM_01030 | ENSMUSG | Gna11 | Dsk7\|E43C | guanine nu | protein-co | 5.51 | 6.16 | 4.29 | 1.87 | 4.21E-06 | 3.36E-04 |
| 682 | chr7 | 16364446 | 16364596 | + | 0 | NA | intron (NM | intron (NM | 23375 | NM_01974 | 56459 | Mm.25853 | NM_01974 | ENSMUSG | Sae1 | 2400010M | SUMO1 ac | protein-co | 4.96 | 5.7 | 3.37 | 2.33 | 4.21E-06 | 3.36E-04 |
| 684 | chr11 | 1.18E+08 | 1.18E+08 | + | 0 | NA | exon (NM_ | exon (NM_ | 763 | NM_02996 | 77727 | Mm.15998 | NM_02996 | ENSMUSG | 6030468B | Il-40 | RIKEN cDN | protein-co | 4.61 | 5.42 | 2.61 | 2.82 | 4.29E-06 | 3.41E-04 |
| 686 | chr2 | 1.31E+08 | 1.31E+08 | + | 0 | NA | intron (NM | intron (NM | -5682 | NM_00129 | 140629 | Mm.48600 | NM_08056 | ENSMUSG | Ubox5 | 1500010O | U box dom | protein-co | 4.99 | 5.73 | 3.4 | 2.34 | 4.29E-06 | 3.41E-04 |
| 688 | chr7 | 72275602 | 72275752 | + | 0 | NA | intron (NM | intron (NM | 30918 | NM_00102 | 244049 | Mm.21741 | NM_00102 | ENSMUSG | Mctp2 | Gm489 | multiple C | protein-co | 7.31 | 7.73 | 6.71 | 1.02 | 4.31E-06 | 3.41E-04 |
| 685 | chr9 | 1.12E+08 | 1.12E+08 | + | 0 | NA | intron (NM | intron (NM | 9264 | NM_00117 | 74100 | Mm.29744 | NM_02875 | ENSMUSG | Arpp21 | 0710001E | cyclic AMP | protein-co | 5.62 | 6.29 | 4.34 | 1.95 | 4.29E-06 | 3.41E-04 |
| 690 | chr8 | 1.25E+08 | 1.25E+08 | + | 0 | NA | intron (NM | intron (NM | 20845 | NM_00135 | 234875 | Mm.38436 | NM_14560 | ENSMUSG | Ttc13 | BC017545 | tetratricop | protein-co | 6.46 | 6.98 | 5.63 | 1.34 | 4.34E-06 | 3.43E-04 |
| 697 | chr2 | 76253647 | 76253797 | + | 0 | NA | intron (NM | MLT1C\|LT | 85071 | NM_00108 | 241489 | Mm.24661 | NM_00108 | ENSMUSG | Pde11a | 6330414F | phosphodi | protein-co | 6.08 | 6.65 | 5.11 | 1.54 | 4.55E-06 | 3.56E-04 |
| 696 | chr2 | 1.03E+08 | 1.03E+08 | + | 0 | NA | intron (NM | intron (NM | 15026 | NM_00117 | 12505 | Mm.42362 | NM_00989 | ENSMUSG | Cd44 | AU023126 | CD44 antig | protein-co | 7.37 | 7.81 | 6.72 | 1.09 | 4.54E-06 | 3.56E-04 |
| 698 | chr12 | 1.16E+08 | 1.16E+08 | + | 0 | NA | intron (NM | intron (NM | 63075 | NM_00951 | 22355 | Mm.39524 | NM_00951 | ENSMUSG | Vipr2 | VPAC2\|VP | vasoactive | protein-co | 6.55 | 7.06 | 5.75 | 1.3 | 4.59E-06 | 3.59E-04 |
| 699 | chr10 | 57666641 | 57666791 | + | 0 | NA | intron (NM | ERVL-B4-ir | 15735 | NM_00119 | 18768 | Mm.26213 | NM_00880 | ENSMUSG | Pkib | Pkibeta\|P | protein kir | protein-co | 6.37 | 5.48 | 6.91 | -1.43 | 4.64E-06 | 3.63E-04 |
| 700 | chr2 | 1.26E+08 | 1.26E+08 | + | 0 | NA | intron (NM | intron (NM | 24654 | NM_00108 | 241633 | Mm.22223 | NM_00108 | ENSMUSG | Atp8b4 | Im | ATPase, cl | protein-co | 5.55 | 6.2 | 4.35 | 1.85 | 4.69E-06 | 3.66E-04 |
| 701 | chr3 | 1.31E+08 | 1.31E+08 | + | 0 | NA | intron (NM | intron (NM | 449 | NM_00127 | 16842 | Mm.25521 | NM_01070 | ENSMUSG | Lef1 | 3000002B | lymphoid | protein-co | 7.19 | 7.62 | 6.58 | 1.05 | 4.70E-06 | 3.66E-04 |
| 702 | chr6 | 1.21E+08 | 1.21E+08 | + | 0 | NA | intron (NM | intron (NM | 2054 | NM_14481 | 214932 | Mm.25532 | NM_14481 | ENSMUSG | Hdhd5 | A930002G | haloacid d | protein-co | 6.04 | 6.61 | 5.06 | 1.55 | 4.71E-06 | 3.66E-04 |
| 708 | chr1 | 86186858 | 86187008 | + | 0 | NA | intron (NM | intron (NM | 32153 | NM_30188 | 78795 | Mm.37926 | NM_02745 | ENSMUSG | Armc9 | 3830422A | armadillo | protein-co | 6.66 | 7.16 | 5.89 | 1.27 | 4.90E-06 | 3.78E-04 |

| | | | | | | | | | | | | | | | | | | | | | | | | |
|---|---|---|---|---|---|---|---|---|---|---|---|---|---|---|---|---|---|---|---|---|---|---|---|---|
| 712 | chr12 | 1.09E+08 | 1.09E+08 | + | 0 | NA | promoter- | promoter- | -35 | NM_0011€ | 14026 | Mm.23884 | NM_0079€ | ENSMUSG | Evl | AI528774\| | Ena-vasod | protein-co | 6.47 | 6.99 | 5.67 | 1.32 | 5.02E-06 | 3.85E-04 |
| 716 | chr14 | 52140847 | 52140997 | + | 0 | NA | intron (NM | intron (NM | 30019 | NM_0011€ | 77945 | Mm.21662 | NM_02387 | ENSMUSG | Rpgrip1 | 0610005A | retinitis pi | protein-co | 6.15 | 6.71 | 5.23 | 1.48 | 5.20E-06 | 3.97E-04 |
| 717 | chr1 | 43542449 | 43542599 | + | 0 | NA | intron (NM | intron (NM | 96773 | NM_01087 | 17974 | Mm.38990 | NM_01087 | ENSMUSG | Nck2 | 4833426I1 | non-cataly | protein-co | 6.63 | 7.14 | 5.84 | 1.29 | 5.30E-06 | 4.03E-04 |
| 718 | chr10 | 1.11E+08 | 1.11E+08 | + | 0 | NA | intron (NM | intron (NM | 6589 | NM_17548 | 237542 | Mm.22020 | NM_17548 | ENSMUSG | Osbpl8 | AA536976 | oxysterol b | protein-co | 6.57 | 7.07 | 5.77 | 1.3 | 5.30E-06 | 4.03E-04 |
| 721 | chr5 | 1.19E+08 | 1.19E+08 | + | 0 | NA | intron (NM | intron (NM | 103523 | NM_00134 | 76199 | Mm.20623 | NM_17242 | ENSMUSG | Med13l | 2210413I1 | mediator c | protein-co | 6.35 | 6.88 | 5.5 | 1.38 | 5.35E-06 | 4.04E-04 |
| 723 | chr2 | 1.53E+08 | 1.53E+08 | + | 0 | NA | intron (NM | intron (NM | 12477 | NR_14925 | 12048 | Mm.23821 | NM_00974 | ENSMUSG | Bcl2l1 | Bcl(X)L\|Bc | BCL2-like 1 | protein-co | 5.13 | 5.83 | 3.71 | 2.11 | 5.54E-06 | 4.17E-04 |
| 725 | chr16 | 95999403 | 95999553 | + | 0 | NA | intron (NM | intron (NM | -8518 | NM_01953 | 56088 | Mm.39624 | NM_01953 | ENSMUSG | Psmg1 | AW55210 | proteasom | protein-co | 6.92 | 7.39 | 6.21 | 1.18 | 5.58E-06 | 4.19E-04 |
| 724 | chr9 | 44043329 | 44043479 | + | 0 | NA | promoter- | promoter- | 20 | NM_00938 | 21838 | Mm.3951 | NM_00938 | ENSMUSG | Thy1 | CD90\|T25 | thymus ce | protein-co | 5.41 | 6.08 | 4.13 | 1.95 | 5.58E-06 | 4.19E-04 |
| 728 | chr9 | 44515602 | 44515752 | + | 0 | NA | intron (NM | intron (NM | 10767 | NM_00755 | 12145 | Mm.6246 | NM_00755 | ENSMUSG | Cxcr5 | Blr1\|CXC- | chemokine | protein-co | 5.67 | 6.29 | 4.58 | 1.71 | 5.65E-06 | 4.23E-04 |
| 729 | chr7 | 1.08E+08 | 1.08E+08 | + | 0 | NA | intron (NM | intron (NM | 16085 | NM_00135 | 19024 | Mm.2817 | NM_00890 | ENSMUSG | Ppfibp2 | Cclp1 | PTPRF inte | protein-co | 7.22 | 7.67 | 6.55 | 1.12 | 5.68E-06 | 4.25E-04 |
| 732 | chr2 | 1.59E+08 | 1.59E+08 | + | 0 | NA | intron (NM | intron (NM | -34746 | NM_02797 | 71878 | Mm.85162 | NM_02797 | ENSMUSG | Fam83d | 2310007D | family with | protein-co | 5.71 | 6.34 | 4.6 | 1.74 | 5.79E-06 | 4.30E-04 |
| 735 | chr2 | 73495951 | 73496101 | + | 0 | NA | intron (NM | intron (NM | -10273 | NM_00128 | 215280 | Mm.22350 | NM_15313 | ENSMUSG | Wipf1 | AI115543\| | WAS/WAS | protein-co | 7.01 | 7.48 | 6.31 | 1.17 | 5.97E-06 | 4.42E-04 |
| 736 | chr12 | 70102532 | 70102682 | + | 0 | NA | intron (NM | intron (NM | 247 | NM_00108 | 18080 | Mm.24404 | NM_00869 | ENSMUSG | Nin | 3110068G | ninein | protein-co | 5.33 | 6.01 | 3.98 | 2.03 | 6.01E-06 | 4.45E-04 |
| 737 | chr1 | 91352547 | 91352697 | + | 0 | NA | intron (NM | intron (NM | 1549 | NM_02755 | 70788 | Mm.68617 | NM_02755 | ENSMUSG | Klhl30 | 4631423F | kelch-like 3 | protein-co | 5.12 | 5.82 | 3.71 | 2.1 | 6.27E-06 | 4.62E-04 |
| 738 | chr6 | 8299188 | 8299338 | + | 0 | NA | intron (NM | intron (NM | 39975 | NM_00130 | 1E+08 | Mm.44610 | NM_00130 | ENSMUSG | Umad1 | Gm16039 | UMAP1-M | protein-co | 6.5 | 7.01 | 5.72 | 1.29 | 6.50E-06 | 4.78E-04 |
| 739 | chr9 | 63507209 | 63507359 | + | 0 | NA | intron (NM | intron (NM | 95209 | NM_00131 | 78250 | Mm.44461 | NM_03006 | ENSMUSG | Iqch | 4921504K | IQ motif co | protein-co | 5.39 | 6.05 | 4.14 | 1.91 | 6.65E-06 | 4.88E-04 |
| 748 | chr12 | 3671207 | 3671357 | + | 0 | NA | intron (NM | intron (NM | 98298 | NM_00788 | 13528 | Mm.28620 | NM_00788 | ENSMUSG | Dtnb | - | dystrobrev | protein-co | 6.31 | 6.85 | 5.45 | 1.4 | 7.09E-06 | 5.15E-04 |
| 749 | chr2 | 1.73E+08 | 1.73E+08 | + | 0 | NA | intron (NM | intron (NM | 5007 | NM_01954 | 56190 | Mm.48578 | NM_01954 | ENSMUSG | Rbm38 | Rnpc1\|Sel | RNA bindi | protein-co | 6.83 | 7.33 | 6.04 | 1.29 | 7.22E-06 | 5.23E-04 |
| 750 | chr8 | 39020563 | 39020713 | + | 0 | NA | intron (NM | intron (NM | 14793 | NM_03025 | 80286 | Mm.27272 | NM_03025 | ENSMUSG | Tusc3 | AU022242 | tumor sup | protein-co | 6.82 | 7.29 | 6.12 | 1.17 | 7.22E-06 | 5.23E-04 |
| 751 | chr13 | 42095945 | 42096095 | + | 0 | NA | intron (NM | intron (NM | 43999 | NM_00777 | 110521 | Mm.4808 | NM_00777 | ENSMUSG | Hivep1 | Cryabp1 | human imi | protein-co | 7.27 | 7.69 | 6.67 | 1.02 | 7.46E-06 | 5.38E-04 |
| 753 | chr5 | 30799058 | 30799208 | + | 0 | NA | 3' UTR (NM | 3' UTR (NM | -15508 | NM_13335 | 100732 | Mm.33229 | NM_13335 | ENSMUSG | Mapre3 | AI790651\| | microtubu | protein-co | 6.01 | 6.58 | 5.03 | 1.56 | 7.53E-06 | 5.42E-04 |
| 755 | chr16 | 97016622 | 97016772 | + | 0 | NA | intron (NM | ORR1E\|LT | 154038 | NM_03117 | 13508 | Mm.10243 | NM_03117 | ENSMUSG | Dscam | 4932410A | DS cell adh | protein-co | 5.72 | 6.36 | 4.57 | 1.79 | 7.63E-06 | 5.48E-04 |
| 756 | chr2 | 3717959 | 3718109 | + | 0 | NA | intron (NM | B4A\|SINE | 4576 | NM_02562 | 66540 | Mm.27786 | NM_02562 | ENSMUSG | Fam107b | 3110001A | family with | protein-co | 6.73 | 7.21 | 6.01 | 1.2 | 7.66E-06 | 5.50E-04 |
| 757 | chr5 | 1.34E+08 | 1.34E+08 | + | 0 | NA | intron (NM | intron (NM | 15006 | NR_10438 | 17969 | Mm.42529 | NM_01087 | ENSMUSG | Ncf1 | NCF-47K\| | neutrophil | protein-co | 5.97 | 6.53 | 5.02 | 1.52 | 7.78E-06 | 5.57E-04 |
| 758 | chr7 | 80782218 | 80782368 | + | 0 | NA | intron (NM | intron (NM | 21038 | NM_01672 | 29875 | Mm.20761 | NM_01672 | ENSMUSG | Iqgap1 | AA682088 | IQ motif co | protein-co | 6.88 | 7.37 | 6.14 | 1.23 | 7.91E-06 | 5.64E-04 |
| 760 | chr5 | 1.24E+08 | 1.24E+08 | + | 0 | NA | intron (NM | intron (NM | 7805 | NM_02660 | 68184 | Mm.28549 | NM_02660 | ENSMUSG | Denr | 1500003K | density-re | protein-co | 6.42 | 6.93 | 5.62 | 1.31 | 7.94E-06 | 5.65E-04 |
| 761 | chr13 | 92523469 | 92523619 | + | 0 | NA | intron (NM | intron (NM | 7266 | NM_17339 | 218441 | Mm.25958 | NM_17339 | ENSMUSG | Zfyve16 | AI035632\| | zinc finger | protein-co | 7.07 | 7.51 | 6.43 | 1.07 | 8.00E-06 | 5.69E-04 |
| 764 | chr11 | 1.2E+08 | 1.2E+08 | + | 0 | NA | intron (NM | intron (NM | 36621 | NM_00134 | 268515 | Mm.30949 | NM_19842 | ENSMUSG | Bahcc1 | B930044J | BAH doma | protein-co | 6.52 | 7.02 | 5.74 | 1.28 | 8.33E-06 | 5.88E-04 |
| 767 | chr9 | 32745977 | 32746127 | + | 0 | NA | intron (NM | intron (NM | 50010 | NM_00103 | 23871 | Mm.29241 | NM_01180 | ENSMUSG | Ets1 | AI196000\| | E26 avian l | protein-co | 5.71 | 6.32 | 4.62 | 1.7 | 8.45E-06 | 5.94E-04 |
| 768 | chr8 | 68627310 | 68627460 | + | 0 | NA | intron (NM | intron (NM | 107700 | NM_00136 | 234356 | Mm.33456 | NM_17275 | ENSMUSG | Csgalnact1 | 4732435N | chondroiti | protein-co | 6.68 | 7.17 | 5.94 | 1.22 | 8.48E-06 | 5.96E-04 |
| 769 | chr2 | 1.61E+08 | 1.61E+08 | + | 0 | NA | intron (NM | intron (NM | 9690 | NM_02128 | 18803 | Mm.44463 | NM_02128 | ENSMUSG | Plcg1 | AI894140\| | phospholip | protein-co | 6.67 | 7.16 | 5.93 | 1.23 | 8.65E-06 | 6.07E-04 |
| 770 | chr17 | 56021291 | 56021441 | + | 0 | NA | intron (NM | intron (NM | 12165 | NM_02653 | 68047 | Mm.35871 | NM_02653 | ENSMUSG | Mpnd | E130307M | MPN doma | protein-co | 6.38 | 6.9 | 5.57 | 1.34 | 8.69E-06 | 6.09E-04 |
| 771 | chr4 | 1.3E+08 | 1.3E+08 | + | 0 | NA | intron (NM | intron (NM | 27531 | NM_02564 | 66585 | Mm.42301 | NM_02564 | ENSMUSG | Snrnp40 | 0610009C | small nucle | protein-co | 5.81 | 6.43 | 4.67 | 1.76 | 8.78E-06 | 6.14E-04 |
| 772 | chr13 | 63119147 | 63119297 | + | 0 | NA | intron (NM | intron (NM | -120807 | NR_11052 | 72061 | Mm.18810 | NM_02807 | ENSMUSG | Aopep | 2010111I0 | aminopept | protein-co | 5.28 | 5.95 | 3.98 | 1.97 | 8.85E-06 | 6.18E-04 |
| 773 | chr7 | 35729749 | 35729899 | + | 0 | NA | intron (NM | intron (NM | 24630 | NM_17870 | 233115 | Mm.25780 | NM_17870 | ENSMUSG | Dpy19l3 | 6030410G | dpy-19-lik | protein-co | 7.42 | 7.83 | 6.83 | 1 | 8.86E-06 | 6.18E-04 |
| 774 | chr2 | 1.26E+08 | 1.26E+08 | + | 0 | NA | intron (NM | MER58A\|D | 82143 | NM_00129 | 69976 | Mm.20216 | NM_17515 | ENSMUSG | Galk2 | 2810017M | galactokin | protein-co | 6.18 | 6.74 | 5.25 | 1.49 | 9.00E-06 | 6.27E-04 |
| 777 | chr3 | 87742856 | 87743006 | + | 0 | NA | intron (NM | intron (NM | 5692 | NM_02897 | 74485 | Mm.47449 | NM_02897 | ENSMUSG | Lrrc71 | 4933430H | leucine ric | protein-co | 6.53 | 7.03 | 5.75 | 1.28 | 9.28E-06 | 6.43E-04 |
| 779 | chr7 | 81074681 | 81074831 | + | 0 | NA | intron (NM | intron (NM | 17156 | NM_05408 | 116904 | Mm.47576 | NM_05408 | ENSMUSG | Alpk3 | AW31948 | alpha-kina | protein-co | 6.48 | 7 | 5.64 | 1.36 | 9.37E-06 | 6.48E-04 |
| 781 | chr9 | 1.03E+08 | 1.03E+08 | + | 0 | NA | intron (NM | ORR1E\|LT | 15453 | NM_02429 | 16716 | Mm.10169 | NM_02429 | ENSMUSG | Ky | CD1\|D9Mi | kyphoscoli | protein-co | 5.38 | 6.04 | 4.11 | 1.93 | 9.47E-06 | 6.53E-04 |
| 782 | chr6 | 1.21E+08 | 1.21E+08 | + | 0 | NA | TTS (NM_( | TTS (NM_( | 16198 | NM_01737 | 53857 | Mm.32884 | NM_01737 | ENSMUSG | Tuba8 | - | tubulin, al | protein-co | 5.46 | 6.1 | 4.27 | 1.83 | 9.52E-06 | 6.56E-04 |
| 786 | chr11 | 4268619 | 4268769 | + | 0 | NA | intron (NM | intron (NM | 1898 | NM_00103 | 16878 | Mm.4964 | NM_00850 | ENSMUSG | Lif | - | leukemia i | protein-co | 6.43 | 6.94 | 5.62 | 1.32 | 9.63E-06 | 6.60E-04 |
| 787 | chr5 | 1.48E+08 | 1.48E+08 | + | 0 | NA | intron (NM | intron (NM | 20581 | NM_00130 | 11987 | Mm.27548 | NM_00751 | ENSMUSG | Slc7a1 | 4831426K | solute carr | protein-co | 6.32 | 6.84 | 5.49 | 1.35 | 9.65E-06 | 6.61E-04 |
| 790 | chr16 | 52091027 | 52091177 | + | 0 | NA | intron (NM | intron (NM | 59553 | NM_00103 | 208650 | Mm.32820 | NM_00103 | ENSMUSG | Cblb | AI429560\| | Casitas B-l | protein-co | 5.2 | 5.88 | 3.88 | 2.01 | 9.90E-06 | 6.75E-04 |
| 791 | chr7 | 25000277 | 25000427 | + | 0 | NA | intron (NM | ID_B1\|SIN | 5585 | NM_00129 | 232975 | Mm.44101 | NM_14492 | ENSMUSG | Atp1a3 | Atpa-2 | ATPase, Na | protein-co | 5.34 | 6.01 | 4.05 | 1.95 | 9.96E-06 | 6.78E-04 |
| 792 | chr10 | 1.27E+08 | 1.27E+08 | + | 0 | NA | intron (NM | intron (NM | -19274 | NM_00135 | 117150 | Mm.22682 | NM_05409 | ENSMUSG | Pip4k2c | Pip5k2c | phosphatid | protein-co | 5.53 | 6.15 | 4.41 | 1.75 | 1.00E-05 | 6.81E-04 |
| 794 | chr4 | 1.55E+08 | 1.55E+08 | + | 0 | NA | intron (NM | intron (NM | 4968 | NM_01138 | 20481 | Mm.28520 | NM_01138 | ENSMUSG | Ski | 2310012I0 | ski sarcom | protein-co | 6.82 | 7.29 | 6.11 | 1.18 | 1.03E-05 | 6.96E-04 |
| 795 | chr2 | 1.25E+08 | 1.25E+08 | + | 0 | NA | intron (NM | MTE2a\|LT | 12046 | NM_00129 | 214968 | Mm.33053 | NM_17255 | ENSMUSG | Sema6d | 1110067B | sema dom | protein-co | 5.97 | 6.53 | 5.04 | 1.49 | 1.03E-05 | 6.98E-04 |
| 799 | chr12 | 1.13E+08 | 1.13E+08 | + | 0 | NA | intron (NM | intron (NM | 3489 | NR_15724 | 116870 | Mm.21257 | NM_05408 | ENSMUSG | Mta1 | - | metastasis | protein-co | 4.96 | 5.68 | 3.48 | 2.2 | 1.09E-05 | 7.30E-04 |
| 802 | chr1 | 1.64E+08 | 1.64E+08 | + | 0 | NA | intron (NM | intron (NM | 12635 | NM_05408 | 116914 | Mm.35444 | NM_05408 | ENSMUSG | Slc19a2 | AV276020 | solute carr | protein-co | 6.37 | 5.56 | 6.89 | -1.32 | 1.12E-05 | 7.51E-04 |
| 804 | chr11 | 82947542 | 82947692 | + | 0 | NA | intron (NR | intron (NR | -4485 | NM_18320 | 327978 | Mm.42147 | NM_18320 | ENSMUSG | Slfn5 | AI956840 | schlafen 5 | protein-co | 6.21 | 6.75 | 5.33 | 1.42 | 1.17E-05 | 7.79E-04 |
| 806 | chr12 | 32015492 | 32015642 | + | 0 | NA | intron (NM | intron (NM | 45022 | NM_00136 | 19088 | Mm.25594 | NM_01115 | ENSMUSG | Prkar2b | AI451071\| | protein kir | protein-co | 6.72 | 7.19 | 6.02 | 1.16 | 1.19E-05 | 7.92E-04 |
| 808 | chr15 | 1E+08 | 1E+08 | + | 0 | NA | intron (NM | intron (NM | 8821 | NM_00873 | 18174 | Mm.23460 | NM_00873 | ENSMUSG | Slc11a2 | DCT1\|DM | solute carr | protein-co | 5.53 | 6 | 4.12 | 1.88 | 1.22E-05 | 8.11E-04 |
| 812 | chr15 | 1.01E+08 | 1.01E+08 | + | 0 | NA | intron (NM | ORR1E\|LT | 4901 | NM_01687 | 53622 | Mm.34793 | NM_016879 | | Krt85 | AA589597 | keratin 85 | protein-co | 5.76 | 6.35 | 4.76 | 1.58 | 1.24E-05 | 8.16E-04 |
| 810 | chr9 | 57831491 | 57831641 | + | 0 | NA | intron (NM | intron (NM | 2668 | NM_01968 | 56380 | Mm.42558 | NM_01968 | ENSMUSG | Arid3b | Bdp\|Dri2 | AT rich int | protein-co | 7.13 | 7.56 | 6.52 | 1.04 | 1.23E-05 | 8.16E-04 |
| 817 | chr11 | 1.19E+08 | 1.19E+08 | + | 0 | NA | intron (NM | intron (NM | 2299 | NM_00762 | 12418 | Mm.26807 | NM_00762 | ENSMUSG | Cbx4 | MPc2\|PC2 | chromobo | protein-co | 6.09 | 6.64 | 5.2 | 1.44 | 1.26E-05 | 8.28E-04 |
| 821 | chr8 | 1.12E+08 | 1.12E+08 | + | 0 | NA | intron (NM | intron (NM | 37247 | NM_02757 | 52815 | Mm.27589 | NM_02757 | ENSMUSG | Ldhd | 4733401P | lactate del | protein-co | 6.35 | 6.86 | 5.56 | 1.3 | 1.28E-05 | 8.37E-04 |
| 825 | chr13 | 12274073 | 12274223 | + | 0 | NA | intron (NM | B1F2\|SINE | -16035 | NM_00108 | 238505 | Mm.40335 | NM_00108 | ENSMUSG | Mtr | AI894170\| | 5-methylte | protein-co | 7.01 | 7.45 | 6.37 | 1.08 | 1.35E-05 | 8.74E-04 |
| 827 | chr4 | 59895265 | 59895415 | + | 0 | NA | intron (NM | intron (NM | 19716 | NM_02105 | 30936 | Mm.28443 | NM_02105 | ENSMUSG | Slc46a2 | 5430429N | solute carr | protein-co | 7.24 | 7.7 | 6.57 | 1.13 | 1.35E-05 | 8.77E-04 |
| 828 | chr14 | 25473276 | 25473426 | + | 0 | NA | intron (NM | intron (NM | 14166 | NM_18320 | 328365 | Mm.22748 | NM_18320 | ENSMUSG | Zmiz1 | BC065120 | zinc finger | protein-co | 6.01 | 6.56 | 5.1 | 1.46 | 1.36E-05 | 8.79E-04 |
| 833 | chr1 | 1.76E+08 | 1.76E+08 | + | 0 | NA | intron (NM | intron (NM | 14556 | NM_02135 | 12663 | Mm.44049 | NM_02135 | ENSMUSG | Chml | AI450814\| | choroidere | protein-co | 6.42 | 6.93 | 5.64 | 1.28 | 1.39E-05 | 8.92E-04 |
| 830 | chr4 | 1.3E+08 | 1.3E+08 | + | 0 | NA | intron (NM | intron (NM | 25332 | NM_00129 | 230775 | Mm.26224 | NM_17307 | ENSMUSG | Adgrb2 | Bai2 | adhesion G | protein-co | 5.1 | 5.8 | 3.69 | 2.11 | 1.38E-05 | 8.92E-04 |
| 832 | chr8 | 14965684 | 14965834 | + | 0 | NA | intron (NM | intron (NM | -45266 | NM_02911 | 74901 | Mm.46675 | NM_02911 | ENSMUSG | Kbtbd11 | 2900016B | kelch repe | protein-co | 5.63 | 6.26 | 4.5 | 1.76 | 1.38E-05 | 8.92E-04 |
| 834 | chr4 | 1.18E+08 | 1.18E+08 | + | 0 | NA | intron (NM | intron (NM | 29304 | NM_02317 | 66233 | Mm.29142 | NM_02317 | ENSMUSG | Dmap1 | 1500016M | DNA meth | protein-co | 5.05 | 3.71 | 5.73 | -2.02 | 1.39E-05 | 8.94E-04 |
| 836 | chr1 | 93389052 | 93389202 | + | 0 | NA | intron (NM | intron (NM | 15231 | NM_00127 | 404545 | Mm.29741 | NM_20703 | ENSMUSG | Ano7 | Ngep\|Nge | anoctamin | protein-co | 6.19 | 6.72 | 5.36 | 1.35 | 1.42E-05 | 9.13E-04 |
| 838 | chr12 | 70078720 | 70078870 | + | 0 | NA | intron (NM | intron (NM | 24059 | NM_00108 | 18080 | Mm.24404 | NM_00869 | ENSMUSG | Nin | 3110068G | ninein | protein-co | 7.03 | 7.46 | 6.39 | 1.07 | 1.43E-05 | 9.16E-04 |
| 841 | chr10 | 81298818 | 81298968 | + | 0 | NA | intron (NM | L1MB7\|LIN | 5930 | NM_00884 | 18717 | Mm.24222 | NM_00884 | ENSMUSG | Pip5k1c | AI115456\| | phosphati | protein-co | 7.14 | 7.59 | 6.49 | 1.11 | 1.45E-05 | 9.26E-04 |
| 845 | chr4 | 1.29E+08 | 1.29E+08 | + | 0 | NA | intron (NM | intron (NM | 7809 | NM_00119 | 54383 | Mm.47373 | NM_01877 | ENSMUSG | Phc2 | A3galt2\|A | polyhome | protein-co | 5.33 | 5.98 | 4.09 | 1.89 | 1.53E-05 | 9.69E-04 |
| 846 | chr3 | 1.31E+08 | 1.31E+08 | + | 0 | NA | TTS (NM_( | TTS (NM_( | 39438 | NM_00821 | 15107 | Mm.26016 | NM_00821 | ENSMUSG | Hadh | AA409008 | hydroxyac | protein-co | 6.82 | 7.27 | 6.17 | 1.1 | 1.54E-05 | 9.73E-04 |
| 847 | chr14 | 67784413 | 67784563 | + | 0 | NA | intron (NM | intron (NM | 39307 | NM_00814 | 14714 | Mm.35830 | NM_00814 | ENSMUSG | Gnrh1 | Gnrh\|Gnrl | gonadotro | protein-co | 5.42 | 6.05 | 4.26 | 1.78 | 1.55E-05 | 9.75E-04 |
| 848 | chr17 | 50084049 | 50084199 | + | 0 | NA | intron (NM | intron (NM | 106373 | NM_18139 | 76438 | Mm.41854 | NM_18139 | ENSMUSG | Rftn1 | 2310015N | raftlin lipi | protein-co | 6.25 | 6.77 | 5.42 | 1.35 | 1.55E-05 | 9.77E-04 |
| 849 | chr9 | 69757361 | 69757511 | + | 0 | NA | TTS (NM_( | TTS (NM_( | 3504 | NM_02237 | 64290 | Mm.44089 | NM_02237 | ENSMUSG | Foxb1 | C43\|Fkh5 | forkhead b | protein-co | 4.9 | 5.64 | 3.3 | 2.34 | 1.55E-05 | 9.79E-04 |
| 853 | chr9 | 35122964 | 35123114 | + | 0 | NA | intron (NR | intron (NR | -6175 | NM_00917 | 20443 | Mm.27597 | NM_00917 | ENSMUSG | St3gal4 | SIAT4-C\|S | ST3 beta-g | protein-co | 6.59 | 7.07 | 5.88 | 1.19 | 1.57E-05 | 9.84E-04 |
| 855 | chr19 | 47531346 | 47531496 | + | 0 | NA | intron (NM | intron (NM | 5698 | NM_17536 | 108689 | Mm.21804 | NM_17536 | ENSMUSG | Stn1 | 0610009H | STN1, CST | protein-co | 5.74 | 6.32 | 4.75 | 1.58 | 1.60E-05 | 0.001 |
| 858 | chr1 | 1.32E+08 | 1.32E+08 | + | 0 | NA | promoter- | promoter- | 66 | NM_00779 | 13034 | Mm.23024 | NM_00779 | ENSMUSG | Ctse | A430072O | cathepsin | protein-co | 5.85 | 6.42 | 4.89 | 1.52 | 1.62E-05 | 0.00101 |
| 859 | chr11 | 71230813 | 71230963 | + | 0 | NA | promoter- | promoter- | -155 | NM_00104 | 637515 | Mm.39040 | NM_00103 | ENSMUSG | Nlrp1b | Nalp1b | NLR family | protein-co | 6.76 | 7.24 | 6.04 | 1.21 | 1.63E-05 | 0.00101 |
| 861 | chr1 | 1.28E+08 | 1.28E+08 | + | 0 | NA | exon (NM_ | exon (NM_ | 9922 | NM_01173 | 22625 | Mm.24656 | NM_01173 | ENSMUSG | Map3k19 | Ysk4 | mitogen-a | protein-co | 6.92 | 7.37 | 6.25 | 1.12 | 1.65E-05 | 0.00102 |
| 860 | chr10 | 39023743 | 39023893 | + | 0 | NA | intron (NM | intron (NM | 58303 | NM_01068 | 16775 | Mm.25806 | NM_01068 | ENSMUSG | Lama4 | - | laminin, al | protein-co | 6.65 | 7.12 | 5.94 | 1.18 | 1.64E-05 | 0.00102 |
| 862 | chr5 | 24826497 | 24826647 | + | 0 | NA | intron (NM | intron (NM | 15789 | NM_05307 | 19744 | Mm.31917 | NM_05307 | ENSMUSG | Rheb | Rheb1 | Ras homol | protein-co | 5.49 | 6.15 | 4.24 | 1.92 | 1.65E-05 | 0.00102 |
| 863 | chr3 | 51882069 | 51882219 | + | 0 | NA | intron (NM | intron (NM | 220965 | NM_00131 | 211666 | Mm.24679 | NM_17499 | ENSMUSG | Mgst2 | GST2\|MGS | microsoma | protein-co | 5.32 | 5.97 | 4.12 | 1.85 | 1.66E-05 | 0.00103 |
| 866 | chr2 | 26936532 | 26936682 | + | 0 | NA | intron (NM | intron (NM | 2538 | NM_19862 | 279029 | Mm.33007 | NM_19862 | ENSMUSG | Stkld1 | Gm711\|Sg | serine/thre | protein-co | 6.18 | 6.7 | 5.35 | 1.35 | 1.72E-05 | 0.00106 |
| 867 | chr7 | 1.18E+08 | 1.18E+08 | + | 0 | NA | 3' UTR (NM | 3' UTR (NM | -3926 | NM_17066 | 267019 | Mm.28821 | NM_17066 | ENSMUSG | Rps15a | A630031B | ribosomal | protein-co | 5.85 | 6.42 | 4.9 | 1.53 | 1.73E-05 | 0.00106 |
| 872 | chr11 | 53528482 | 53528632 | + | 0 | NA | intron (NM | intron (NM | 8369 | NM_00136 | 20362 | Mm.27439 | NM_03314 | ENSMUSG | 8-Sep | AW04616 | septin 8 | protein-co | 7.05 | 7.47 | 6.46 | 1.01 | 1.79E-05 | 0.00109 |
| 873 | chr16 | 24826482 | 24826632 | + | 0 | NA | intron (NM | intron (NM | 104702 | NM_00114 | 210126 | Mm.20938 | NM_17866 | ENSMUSG | Lpp | 9430020K | LIM domai | protein-co | 5.56 | 6.18 | 4.46 | 1.72 | 1.81E-05 | 0.00111 |
| 875 | chr5 | 1.34E+08 | 1.34E+08 | + | 0 | NA | intron (NM | intron (NM | -40321 | NM_00128 | 17969 | Mm.42529 | NM_01087 | ENSMUSG | Ncf1 | NCF-47K\| | neutrophil | protein-co | 7.29 | 7.71 | 6.69 | 1.02 | 1.83E-05 | 0.00111 |
| 877 | chr2 | 1.63E+08 | 1.63E+08 | + | 0 | NA | intron (NM | intron (NM | 23652 | NM_00136 | 269389 | Mm.20770 | NM_00109 | ENSMUSG | Tox2 | AI851523\| | TOX high n | protein-co | 5.88 | 6.48 | 4.83 | 1.65 | 1.84E-05 | 0.00112 |
| 876 | chr4 | 33072935 | 33073085 | + | 0 | NA | intron (NM | intron (NM | 9898 | NM_00807 | 14409 | Mm.6227 | NM_00807 | ENSMUSG | Gabrr2 | - | gamma-an | protein-co | 6.46 | 6.95 | 5.72 | 1.23 | 1.84E-05 | 0.00112 |
| 878 | chr9 | 75253844 | 75253994 | + | 0 | NA | intron (NM | intron (NM | 21905 | NM_00108 | 208943 | Mm.41339 | NM_00108 | ENSMUSG | Myo5c | 9130003O | myosin VC | protein-co | 5.69 | 6.29 | 4.63 | 1.66 | 1.86E-05 | 0.00113 |
| 881 | chr6 | 91108381 | 91108531 | + | 0 | NA | intron (NM | MLT1A0\|L | 8370 | NM_01881 | 54563 | Mm.28162 | NM_01881 | ENSMUSG | Nup210 | 9830001L | nucleopori | protein-co | 6.7 | 7.16 | 6.01 | 1.15 | 1.90E-05 | 0.00115 |
| 882 | chr2 | 1.45E+08 | 1.45E+08 | + | 0 | NA | intron (NM | intron (NM | 30691 | NM_00135 | 94249 | Mm.21717 | NM_05319 | ENSMUSG | Slc24a3 | NCKX3 | solute carr | protein-co | 7 | 7.43 | 6.39 | 1.04 | 1.94E-05 | 0.00117 |
| 886 | chr13 | 19357426 | 19357576 | + | 0 | NA | TTS (NM_( | TTS (NM_( | 38312 | NM_02427 | 76205 | Mm.41940 | NM_02427 | ENSMUSG | Stard3nl | 0610035N | STARD3 N- | protein-co | 5.75 | 6.34 | 4.75 | 1.59 | 2.03E-05 | 0.00121 |
| 887 | chr6 | 1.35E+08 | 1.35E+08 | + | 0 | NA | intron (NM | intron (NM | 20408 | NM_00136 | 74525 | Mm.34182 | NM_02898 | ENSMUSG | Fam234b | 8430419L | family with | protein-co | 6.62 | 7.1 | 5.91 | 1.2 | 2.04E-05 | 0.00122 |
| 889 | chr6 | 1.34E+08 | 1.34E+08 | + | 0 | NA | intron (NM | intron (NM | 23996 | NM_00796 | 14011 | Mm.26999 | NM_00796 | ENSMUSG | Etv6 | AW12310 | ets variant | protein-co | 7.09 | 7.51 | 6.49 | 1.02 | 2.07E-05 | 0.00123 |
| 890 | chr7 | 1.4E+08 | 1.4E+08 | + | 0 | NA | exon (NM_ | exon (NM_ | -5139 | NM_19930 | 212508 | Mm.29608 | NM_19930 | ENSMUSG | Mtg1 | Gm169\|Gt | mitochonc | protein-co | 4.73 | 5.47 | 3.17 | 2.3 | 2.08E-05 | 0.00124 |
| 891 | chr19 | 4155079 | 4155229 | + | 0 | NA | intron (NM | intron (NM | 508 | NM_01693 | 19265 | Mm.27193 | NM_01693 | ENSMUSG | Ptprcap | AI605264\| | protein ty | protein-co | 6 | 6.56 | 5.09 | 1.47 | 2.10E-05 | 0.00125 |
| 892 | chr7 | 1.13E+08 | 1.13E+08 | + | 0 | NA | intron (NM | intron (NM | 34463 | NM_00136 | 11865 | Mm.33970 | NM_00748 | ENSMUSG | Arntl | Arnt3\|BM | aryl hydro | protein-co | 5.09 | 5.77 | 3.76 | 2.01 | 2.10E-05 | 0.00125 |
| 893 | chr9 | 32509929 | 32510079 | + | 0 | NA | intron (NM | intron (NM | 31448 | NM_00802 | 14247 | Mm.25890 | NM_00802 | ENSMUSG | Fli1 | EWSR2\|Fli | Friend leu | protein-co | 6.85 | 7.29 | 6.22 | 1.08 | 2.12E-05 | 0.00126 |
| 896 | chr2 | 71356683 | 71356833 | + | 0 | NA | intron (NM | intron (NM | 10997 | NM_17243 | 78830 | Mm.30928 | NM_17243 | ENSMUSG | Slc25a12 | 2610001D | solute carr | protein-co | 4.82 | 5.55 | 3.27 | 2.27 | 2.17E-05 | 0.00128 |
| 901 | chr8 | 1.12E+08 | 1.12E+08 | + | 0 | NA | intron (NM | MLT1H1\|L | 25541 | NM_00136 | 170737 | Mm.27157 | NM_13320 | ENSMUSG | Znrf1 | B830022L | zinc and ri | protein-co | 5.32 | 5.96 | 4.15 | 1.81 | 2.21E-05 | 0.0013 |
| 904 | chr1 | 95657390 | 95657540 | + | 0 | NA | intron (NM | intron (NM | 10129 | NM_00918 | 20452 | Mm.30622 | NM_00918 | ENSMUSG | St8sia4 | PST\|PST-1 | ST8 alpha- | protein-co | 6.72 | 7.18 | 6.03 | 1.15 | 2.24E-05 | 0.00131 |
| 903 | chr7 | 75745086 | 75745236 | + | 0 | NA | intron (NM | intron (NM | -103177 | NM_00112 | 207952 | Mm.20622 | NM_02965 | ENSMUSG | Klhl25 | 2810402K | kelch-like 2 | protein-co | 6.98 | 7.42 | 6.35 | 1.08 | 2.23E-05 | 0.00131 |
| 911 | chr1 | 87830067 | 87830217 | + | 0 | NA | intron (NM | intron (NM | -23145 | NM_17764 | 227333 | Mm.27721 | NM_17764 | ENSMUSG | Dgkd | AI841987\| | diacylglyce | protein-co | 5.19 | 5.85 | 3.94 | 1.91 | 2.36E-05 | 0.00137 |
| 910 | chr7 | 16246077 | 16246227 | + | 0 | NA | TTS (NM_( | TTS (NM_( | -1998 | NM_17691 | 319430 | Mm.23957 | NM_17691 | ENSMUSG | C5ar2 | C5L2\|E030 | compleme | protein-co | 5.35 | 5.99 | 4.14 | 1.85 | 2.36E-05 | 0.00137 |
| 914 | chr19 | 8592277 | 8592427 | + | 0 | NA | promoter- | promoter- | -830 | NM_0011€ | 19879 | Mm.28529 | NM_03119 | ENSMUSG | Slc22a8 | Oat3\|Roct | solute carr | protein-co | 6.39 | 6.89 | 5.63 | 1.26 | 2.39E-05 | 0.00138 |
| 919 | chr10 | 8287462 | 8287612 | + | 0 | NA | intron (NM | intron (NM | 231288 | NM_17738 | 338362 | Mm.41163 | NM_17738 | ENSMUSG | Ust | D930010O | uronyl-2-s | protein-co | 4.89 | 5.63 | 3.29 | 2.34 | 2.49E-05 | 0.00143 |
| 921 | chr7 | 63925503 | 63925653 | + | 0 | NA | intron (NM | intron (NM | 8721 | NM_00129 | 381994 | Mm.42546 | NM_00129 | ENSMUSG | E030018B | Gm1697 | RIKEN cDN | protein-co | 6.34 | 6.88 | 5.46 | 1.41 | 2.49E-05 | 0.00143 |
| 923 | chr8 | 1.17E+08 | 1.17E+08 | + | 0 | NA | intron (NM | intron (NM | 97535 | NM_02944 | 75796 | Mm.15957 | NM_02944 | ENSMUSG | Cdyl2 | 1700029M | chromodo | protein-co | 5.44 | 6.06 | 4.35 | 1.72 | 2.50E-05 | 0.00143 |
| 925 | chr10 | 76355079 | 76355229 | + | 0 | NA | intron (NM | intron (NM | -9863 | NM_00108 | 64451 | Mm.33183 | NM_00108 | ENSMUSG | Dip2a | 4931420H | disco inter | protein-co | 6.47 | 6.95 | 5.75 | 1.2 | 2.53E-05 | 0.00144 |
| 927 | chr2 | 1.64E+08 | 1.64E+08 | + | 0 | NA | TTS (NM_( | TTS (NM_( | 23959 | NM_00127 | 11486 | Mm.388 | NM_00739 | ENSMUSG | Ada | - | adenosine | protein-co | 5.34 | 6 | 4.08 | 1.92 | 2.55E-05 | 0.00145 |
| 926 | chr4 | 1.33E+08 | 1.33E+08 | + | 0 | NA | promoter- | promoter- | -62 | NM_03139 | 269589 | Mm.25660 | NM_03139 | ENSMUSG | Sytl1 | 9030417L | synaptota | protein-co | 6.93 | 7.38 | 6.27 | 1.1 | 2.54E-05 | 0.00145 |
| 929 | chr13 | 1.13E+08 | 1.13E+08 | + | 0 | NA | promoter- | promoter- | -57 | NM_01037 | 14938 | Mm.15510 | NM_01037 | ENSMUSG | Gzma | AW49411 | granzyme | protein-co | 5.48 | 6.11 | 4.33 | 1.78 | 2.63E-05 | 0.00149 |
| 934 | chr3 | 1.22E+08 | 1.22E+08 | + | 0 | NA | intron (NM | intron (NM | 18891 | NM_01386 | 29815 | Mm.45815 | NM_01386 | ENSMUSG | Bcar3 | AI131758\| | breast can | protein-co | 6.4 | 6.92 | 5.59 | 1.33 | 2.70E-05 | 0.00152 |
| 938 | chr16 | 21632056 | 21632206 | + | 0 | NA | intron (NM | intron (NM | 62534 | NM_00100 | 72190 | Mm.13160 | NM_00100 | ENSMUSG | 2510009E | AI194952 | RIKEN cDN | protein-co | 6.36 | 6.86 | 5.6 | 1.26 | 2.73E-05 | 0.00153 |
| 939 | chr4 | 46614783 | 46614933 | + | 0 | NA | intron (NM | intron (NM | -12929 | NM_0011€ | 107684 | Mm.41631 | NM_17889 | ENSMUSG | Coro2a | 9030208C | coronin, a | protein-co | 6.1 | 6.63 | 5.26 | 1.37 | 2.79E-05 | 0.00156 |
| 942 | chr11 | 11763512 | 11763662 | + | 0 | NA | intron (NM | intron (NM | 45375 | NM_0011€ | 60530 | Mm.23611 | NM_02189 | ENSMUSG | Fignl1 | - | fidgetin-lik | protein-co | 6.12 | 6.64 | 5.29 | 1.35 | 2.82E-05 | 0.00158 |
| 941 | chr11 | 80489941 | 80490091 | + | 0 | NA | intron (NM | intron (NM | 12993 | NM_00987 | 12569 | Mm.14227 | NM_00987 | ENSMUSG | Cdk5r1 | Cdk5r\|D11 | cyclin-dep | protein-co | 6.52 | 7 | 5.82 | 1.18 | 2.82E-05 | 0.00158 |
| 943 | chr4 | 44022408 | 44022558 | + | 0 | NA | intron (NM | intron (NM | 9813 | NM_00136 | 12757 | Mm.29887 | NM_01676 | ENSMUSG | Clta | AV026556 | clathrin, li | protein-co | 6.4 | 6.89 | 5.65 | 1.24 | 2.86E-05 | 0.0016 |

| | | | | | | | | | | | | | | | | | | | | | | | |
|---|---|---|---|---|---|---|---|---|---|---|---|---|---|---|---|---|---|---|---|---|---|---|---|
| 944 | chr19 | 10698430 | 10698580 | + | 0 | NA | intron (NM | intron (NM | -2263 | NM_00136 | 107305 | Mm.19091 | NM_1814( | ENSMUSG | Vps37c | 5730409F2 | vacuolar p | protein-co | 6.42 | 6.91 | 5.68 | 1.23 | 2.92E-05 | 0.00162 |
| 945 | chr11 | 75537497 | 75537647 | + | 0 | NA | intron (NM | intron (NM | 5460 | NM_00119 | 215113 | Mm.11186 | NM_17338 | ENSMUSG | Slc43a2 | 7630402D | solute carr | protein-co | 5.52 | 6.13 | 4.45 | 1.68 | 2.94E-05 | 0.00163 |
| 947 | chr13 | 63112703 | 63112853 | + | 0 | NA | intron (NM | intron (NM | -127251 | NR_11052 | 72061 | Mm.1881( | NM_02807 | ENSMUSG | Aopep | 2010111I0 | aminopept | protein-co | 5.54 | 6.18 | 4.39 | 1.79 | 2.98E-05 | 0.00165 |
| 948 | chr2 | 1.32E+08 | 1.32E+08 | + | 0 | NA | intron (NM | intron (NM | 18050 | NM_00117 | 228608 | Mm.13658 | NM_14553 | ENSMUSG | Smox | B130066H | spermine ( | protein-co | 7.24 | 7.68 | 6.62 | 1.06 | 3.01E-05 | 0.00166 |
| 951 | chr1 | 1.27E+08 | 1.27E+08 | + | 0 | NA | intron (NM | intron (NM | 104244 | NM_14512 | 107895 | Mm.21468 | NM_14512 | ENSMUSG | Mgat5 | 4930471A | mannoside | protein-co | 6.3 | 6.81 | 5.5 | 1.31 | 3.03E-05 | 0.00167 |
| 955 | chr10 | 67888138 | 67888288 | + | 0 | NA | 3' UTR (NM | 3' UTR (NM | 24449 | NM_17867 | 216049 | Mm.26834 | NM_17867 | ENSMUSG | Zfp365 | AI839779\| | zinc finger | protein-co | 5.01 | 5.71 | 3.57 | 2.14 | 3.11E-05 | 0.00171 |
| 956 | chr16 | 62807747 | 62807897 | + | 0 | NA | intron (NM | intron (NM | -6854 | NM_02658 | 68159 | Mm.38679 | NM_02658 | ENSMUSG | Stx19 | A030009B | syntaxin 1 | protein-co | 5.33 | 5.96 | 4.2 | 1.76 | 3.22E-05 | 0.00176 |
| 958 | chr4 | 1.03E+08 | 1.03E+08 | + | 0 | NA | intron (NM | PB1D7\|SIN | 15469 | NM_00135 | 242585 | Mm.2818( | NM_17773 | ENSMUSG | Slc35d1 | AI834976\| | solute carr | protein-co | 4.35 | 2.57 | 5.12 | -2.56 | 3.32E-05 | 0.00181 |
| 957 | chr15 | 93486990 | 93487140 | + | 0 | NA | intron (NM | intron (NM | 32455 | NM_00136 | 106042 | Mm.15031 | NM_00103 | ENSMUSG | Prickle1 | 1110058P | prickle pla | protein-co | 7.11 | 7.54 | 6.5 | 1.04 | 3.31E-05 | 0.00181 |
| 963 | chr8 | 1.06E+08 | 1.06E+08 | + | 0 | NA | intron (NM | intron (NM | -5833 | NM_02803 | 71986 | Mm.30751 | NM_02803 | ENSMUSG | Ddx28 | 2410004K | DEAD (Asp | protein-co | 5.67 | 6.26 | 4.65 | 1.61 | 3.44E-05 | 0.00187 |
| 964 | chr2 | 1.52E+08 | 1.52E+08 | + | 0 | NA | intron (NR | intron (NR | 4320 | NM_00137 | 228777 | Mm.33066 | NM_0010( | ENSMUSG | Nrsn2 | Gm123\|N | neurensin | protein-co | 6.36 | 6.85 | 5.59 | 1.26 | 3.46E-05 | 0.00188 |
| 967 | chr3 | 94388203 | 94388353 | + | 0 | NA | intron (NM | intron (NM | 10846 | NR_12165 | 19885 | Mm.4372 | NM_01128 | ENSMUSG | Rorc | Nr1f3\|ROI | RAR-relate | protein-co | 6.75 | 7.5 | 5.11 | 2.39 | 3.51E-05 | 0.00189 |
| 968 | chr6 | 97538835 | 97538985 | + | 0 | NA | intron (NM | MER21C\|L | -51090 | NM_00134 | 232288 | Mm.27789 | NM_14514 | ENSMUSG | Frmd4b | 6030440G | FERM dom | protein-co | 5.24 | 5.9 | 4.01 | 1.89 | 3.52E-05 | 0.00189 |
| 970 | chr5 | 1.08E+08 | 1.08E+08 | + | 0 | NA | intron (NM | ID_B1\|SIN | 15821 | NM_02606 | 67266 | Mm.27789 | NM_02606 | ENSMUSG | Dipk1a | 2900024C | divergent | protein-co | 5.69 | 6.27 | 4.71 | 1.56 | 3.55E-05 | 0.00191 |
| 973 | chr14 | 1.01E+08 | 1.01E+08 | + | 0 | NA | intron (NM | intron (NM | 131822 | NM_00108 | 210789 | Mm.32063 | NM_17338 | ENSMUSG | Tbc1d4 | 5930406J0 | TBC1 dom | protein-co | 6.67 | 7.12 | 6.01 | 1.11 | 3.66E-05 | 0.00196 |
| 974 | chr2 | 1.25E+08 | 1.25E+08 | + | 0 | NA | intron (NM | intron (NM | 8494 | NM_00129 | 214968 | Mm.33053 | NM_17253 | ENSMUSG | Sema6d | 1110067B( | sema dom | protein-co | 4.81 | 5.53 | 3.3 | 2.22 | 3.67E-05 | 0.00196 |
| 975 | chr11 | 68462822 | 68462972 | + | 0 | NA | intron (NM | ID_B1\|SIN | 30772 | NM_17732 | 320207 | Mm.24496 | NM_17732 | ENSMUSG | Pik3r5 | AV230647 | phosphoin | protein-co | 4.23 | 5.04 | 2.22 | 2.82 | 3.69E-05 | 0.00197 |
| 976 | chr15 | 58022991 | 58023141 | + | 0 | NA | intron (NM | intron (NM | 11228 | NM_02941 | 75758 | Mm.45579 | NM_02941 | ENSMUSG | 9130401M | AI849328 | RIKEN cDN | protein-co | 4.91 | 5.61 | 3.54 | 2.07 | 3.70E-05 | 0.00197 |
| 978 | chr3 | 1.01E+08 | 1.01E+08 | + | 0 | NA | TTS (NM_( | TTS (NM_( | 12257 | NM_01348 | 12481 | Mm.22842 | NM_01348 | ENSMUSG | Cd2 | LFA-2\|Ly-3 | CD2 antige | protein-co | 5.96 | 6.5 | 5.08 | 1.42 | 3.72E-05 | 0.00198 |
| 979 | chr14 | 26924645 | 26924795 | + | 0 | NA | intron (NM | MLTR11B\| | 30106 | NM_00117 | 142687 | Mm.13825 | NM_08085 | ENSMUSG | Asb14 | Asb-14 | ankyrin rep | protein-co | 4.71 | 5.45 | 3.16 | 2.29 | 3.74E-05 | 0.00199 |
| 980 | chr7 | 75586236 | 75586386 | + | 0 | NA | exon (NM_ | exon (NM_ | 130777 | NM_02933 | 75547 | Mm.2161( | NM_02933 | ENSMUSG | Akap13 | 1700026G | A kinase (F | protein-co | 6.57 | 7.03 | 5.88 | 1.15 | 3.75E-05 | 0.00199 |
| 982 | chr7 | 80205111 | 80205261 | + | 0 | NA | intron (NM | RMER4B\|L | 6745 | NM_00136 | 20352 | Mm.2759( | NM_01365 | ENSMUSG | Sema4b | SemC\|Sem | sema dom | protein-co | 6.18 | 6.68 | 5.39 | 1.29 | 3.80E-05 | 0.00201 |
| 985 | chr5 | 54101446 | 54101596 | + | 0 | NA | intron (NM | intron (NM | 102998 | NM_05408 | 116873 | Mm.26598 | NM_05408 | ENSMUSG | Stim2 | - | stromal int | protein-co | 6.21 | 6.71 | 5.42 | 1.29 | 3.87E-05 | 0.00205 |
| 987 | chrX | 1.07E+08 | 1.07E+08 | + | 0 | NA | promoter- | promoter- | -78 | NM_00135 | 78826 | Mm.74639 | NM_17243 | ENSMUSG | P2ry10 | 5830408N | purinergic | protein-co | 6.21 | 6.72 | 5.42 | 1.31 | 3.89E-05 | 0.00205 |
| 988 | chr2 | 90021503 | 90021653 | + | 0 | NA | exon (NM_ | exon (NM_ | 486 | NM_02136 | 258206 | Mm.37737 | NM_02136 | ENSMUSG | Olfr1264 | MOR236-1 | olfactory r | protein-co | 6.03 | 6.56 | 5.17 | 1.4 | 3.99E-05 | 0.0021 |
| 990 | chr10 | 26324851 | 26325001 | + | 0 | NA | intron (NM | intron (NM | 49146 | NM_00135 | 237339 | Mm.33843 | NM_17278 | ENSMUSG | L3mbtl3 | AI481284\| | L3MBTL3 h | protein-co | 4.78 | 5.51 | 3.27 | 2.24 | 4.05E-05 | 0.00212 |
| 991 | chr5 | 1.25E+08 | 1.25E+08 | + | 0 | NA | intron (NM | intron (NM | 52649 | NM_02844 | 73121 | Mm.29613 | NM_02844 | ENSMUSG | Rflna | 3110032G | refilin A | protein-co | 5.75 | 6.33 | 4.77 | 1.56 | 4.09E-05 | 0.00214 |
| 993 | chr7 | 25384970 | 25385120 | + | 0 | NA | intron (NM | intron (NM | 5067 | NM_00103 | 16890 | Mm.15854 | NM_01071 | ENSMUSG | Lipe | 4933403G | lipase, hor | protein-co | 6.6 | 7.07 | 5.91 | 1.16 | 4.16E-05 | 0.00217 |
| 996 | chr8 | 33306682 | 33306832 | + | 0 | NA | intron (NM | RSINE1\|SII | 78770 | NM_00112 | 22427 | Mm.2288( | NM_01172 | ENSMUSG | Wrn | AI846146 | Werner sy | protein-co | 5.65 | 6.23 | 4.67 | 1.56 | 4.17E-05 | 0.00217 |
| 997 | chr6 | 1.19E+08 | 1.19E+08 | + | 0 | NA | intron (NM | intron (NM | -7693 | NM_00135 | 68465 | Mm.29182 | NM_19798 | ENSMUSG | Adipor2 | 1110001I1 | adiponecti | protein-co | 7.01 | 7.42 | 6.41 | 1.01 | 4.17E-05 | 0.00218 |
| 999 | chr5 | 34908340 | 34908490 | + | 0 | NA | intron (NM | intron (NM | -7500 | NM_20727 | 403174 | Mm.29122 | NM_20727 | ENSMUSG | Msantd1 | A930005I0 | Myb/SANT | protein-co | 5.92 | 6.46 | 5.03 | 1.43 | 4.22E-05 | 0.00219 |
| 998 | chr5 | 72752762 | 72752912 | + | 0 | NA | promoter- | promoter- | -60 | NM_00128 | 22165 | Mm.3264 | NM_01369 | ENSMUSG | Txk | A130089B | TXK tyrosi | protein-co | 6.38 | 6.87 | 5.64 | 1.23 | 4.20E-05 | 0.00219 |
| 1002 | chr2 | 1.03E+08 | 1.03E+08 | + | 0 | NA | intron (NM | intron (NM | 31515 | NM_00791 | 13661 | Mm.10724 | NM_00791 | ENSMUSG | Ehf | 9030625L1 | ets homol | protein-co | 6.55 | 5.85 | 7.02 | -1.16 | 4.30E-05 | 0.00223 |
| 1007 | chr3 | 1.58E+08 | 1.58E+08 | + | 0 | NA | intron (NM | intron (NM | 7242 | NM_01119 | 19218 | Mm.30424 | NM_01119 | ENSMUSG | Ptger3 | EP3\|Pgere | prostaglan | protein-co | 6.4 | 5.6 | 6.92 | -1.32 | 4.43E-05 | 0.00228 |
| 1008 | chr9 | 1.2E+08 | 1.2E+08 | + | 0 | NA | intron (NM | intron (NM | 13350 | NM_00998 | 13051 | Mm.44065 | NM_00998 | ENSMUSG | Cx3cr1 | - | chemokine | protein-co | 5.58 | 6.16 | 4.58 | 1.58 | 4.52E-05 | 0.00231 |
| 1010 | chr6 | 1.21E+08 | 1.21E+08 | + | 0 | NA | TTS (NM_( | TTS (NM_( | 15489 | NM_01737 | 53857 | Mm.32884 | NM_01737 | ENSMUSG | Tuba8 | - | tubulin, al | protein-co | 5.87 | 6.41 | 5 | 1.41 | 4.60E-05 | 0.00235 |
| 1011 | chr7 | 46897610 | 46897760 | + | 0 | NA | intron (NM | intron (NM | 22284 | NM_00134 | 22088 | Mm.24133 | NM_02188 | ENSMUSG | Tsg101 | AI255943\| | tumor sus | protein-co | 5.12 | 5.81 | 3.76 | 2.04 | 4.65E-05 | 0.00237 |
| 1015 | chr12 | 85629808 | 85629958 | + | 0 | NA | intron (NM | intron (NM | 29865 | NM_0012( | 81703 | Mm.10356 | NM_03088 | ENSMUSG | Jdp2 | Jundm2\|J | Jun dimeri | protein-co | 6.36 | 6.86 | 5.58 | 1.28 | 4.77E-05 | 0.00242 |
| 1018 | chr13 | 1.13E+08 | 1.13E+08 | + | 0 | NA | intron (NM | intron (NM | 1898 | NM_02361 | 71690 | Mm.38929 | NM_02361 | ENSMUSG | Esm1 | 0610042H | endothelia | protein-co | 4.87 | 5.56 | 3.51 | 2.05 | 4.87E-05 | 0.00246 |
| 1017 | chr7 | 1.27E+08 | 1.27E+08 | + | 0 | NA | promoter- | promoter- | 35 | NM_0084( | 16408 | Mm.1618 | NM_0084( | ENSMUSG | Itgal | (p180)\|Cd | integrin al | protein-co | 6.82 | 7.25 | 6.19 | 1.06 | 4.87E-05 | 0.00246 |
| 1019 | chr12 | 54198438 | 54198588 | + | 0 | NA | intron (NM | intron (NM | 5361 | NM_02813 | 112407 | Mm.13303 | NM_02813 | ENSMUSG | Egln3 | 2610021G | egl-9 famil | protein-co | 5.65 | 6.24 | 4.66 | 1.58 | 4.92E-05 | 0.00248 |
| 1020 | chr17 | 65929463 | 65929613 | + | 0 | NA | intron (NM | intron (NM | 21649 | NM_02305 | 65960 | Mm.10153 | NM_02305 | ENSMUSG | Twsg1 | 1810013J1 | twisted ga | protein-co | 5.72 | 6.28 | 4.8 | 1.49 | 4.94E-05 | 0.00248 |
| 1022 | chr17 | 53658851 | 53659001 | + | 0 | NA | intron (NR | intron (NR | 30389 | NM_02823 | 72415 | Mm.1532( | NM_02823 | ENSMUSG | Sgo1 | 3300001M | shugoshin | protein-co | 5.11 | 3.9 | 5.76 | -1.86 | 5.00E-05 | 0.00251 |
| 1025 | chr12 | 1.11E+08 | 1.11E+08 | + | 0 | NA | intron (NM | MER21B\|L | 21134 | NM_00136 | 69641 | Mm.47576 | NM_02714 | ENSMUSG | Wdr20 | 2310040A | WD repeat | protein-co | 6.11 | 6.61 | 5.32 | 1.29 | 5.03E-05 | 0.00252 |
| 1024 | chr5 | 36553987 | 36554137 | + | 0 | NA | intron (NM | B3A\|SINE\| | -23439 | NM_00111 | 100855 | Mm.4862( | NM_13391 | ENSMUSG | Tbc1d14 | 2810413P | TBC1 dom | protein-co | 6.27 | 6.76 | 5.51 | 1.25 | 5.03E-05 | 0.00252 |
| 1029 | chr5 | 88381799 | 88381949 | + | 0 | NA | intron (NM | intron (NM | 5766 | NM_02779 | 71421 | Mm.7970( | NM_02779 | ENSMUSG | Amtn | 5430427O | amelotin | protein-co | 6.44 | 5.73 | 6.91 | -1.17 | 5.15E-05 | 0.00257 |
| 1031 | chr7 | 24617932 | 24618082 | + | 0 | NA | intron (NM | ID_B1\|SIN | 6679 | NM_0011( | 232970 | Mm.33366 | NM_0011( | ENSMUSG | Phldb3 | EG232970 | pleckstrin | protein-co | 6.81 | 7.25 | 6.17 | 1.08 | 5.18E-05 | 0.00258 |
| 1032 | chr13 | 12502741 | 12502891 | + | 0 | NA | intron (NM | B1F\|SINE\| | 17634 | NM_13364 | 171211 | Mm.15967 | NM_13364 | ENSMUSG | Edaradd | 1810032E( | EDAR (ect | protein-co | 5.27 | 5.9 | 4.15 | 1.75 | 5.20E-05 | 0.00259 |
| 1033 | chr15 | 93318907 | 93319057 | + | 0 | NA | intron (NR | intron (NR | 17953 | NR_02831 | 67057 | Mm.4714 | NM_02418 | ENSMUSG | Yaf2 | 2810021M | YY1 associ | protein-co | 6.72 | 7.18 | 6.03 | 1.14 | 5.25E-05 | 0.00261 |
| 1034 | chr9 | 1.15E+08 | 1.15E+08 | + | 0 | NA | intron (NM | intron (NM | 6507 | NM_13397 | 102545 | Mm.3560( | NM_13397 | ENSMUSG | Cmtm7 | AI481279\| | CKLF-like I | protein-co | 6.65 | 7.09 | 6.01 | 1.08 | 5.26E-05 | 0.00261 |
| 1035 | chr12 | 1.08E+08 | 1.08E+08 | + | 0 | NA | intron (NM | intron (NM | 72608 | NM_02139 | 58208 | Mm.39269 | NM_02139 | ENSMUSG | Bcl11b | 9130430L1 | B cell leuk | protein-co | 7.04 | 7.46 | 6.44 | 1.02 | 5.28E-05 | 0.00262 |
| 1036 | chr17 | 57188667 | 57188817 | + | 0 | NA | TTS (NM_( | TTS (NM_( | 5439 | NM_01941 | 50930 | Mm.48336 | NM_01941 | ENSMUSG | Tnfsf14 | HVEM-L\|H | tumor nec | protein-co | 6.51 | 6.98 | 5.81 | 1.17 | 5.29E-05 | 0.00262 |
| 1038 | chr1 | 1.34E+08 | 1.34E+08 | + | 0 | NA | promoter- | promoter- | -46 | NM_00115 | 240754 | Mm.36534 | NM_17284 | ENSMUSG | Lax1 | A530029E( | lymphocyt | protein-co | 6.8 | 7.24 | 6.16 | 1.08 | 5.36E-05 | 0.00265 |
| 1040 | chr1 | 1.54E+08 | 1.54E+08 | + | 0 | NA | intron (NM | intron (NM | 25674 | NM_00134 | 67792 | Mm.37914 | NM_02638 | ENSMUSG | Rgs8 | 6530413N | regulator ( | protein-co | 5.61 | 6.18 | 4.64 | 1.54 | 5.39E-05 | 0.00266 |
| 1041 | chr12 | 1.08E+08 | 1.08E+08 | + | 0 | NA | intron (NM | intron (NM | 38587 | NM_02139 | 58208 | Mm.39269 | NM_02139 | ENSMUSG | Bcl11b | 9130430L1 | B cell leuk | protein-co | 7.2 | 7.61 | 6.61 | 1.01 | 5.40E-05 | 0.00266 |
| 1039 | chr7 | 66116077 | 66116227 | + | 0 | NA | intron (NM | intron (NM | 6660 | NM_00108 | 269941 | Mm.33391 | NM_00108 | ENSMUSG | Chsy1 | - | chondroiti | protein-co | 6.35 | 6.83 | 5.63 | 1.2 | 5.38E-05 | 0.00266 |
| 1042 | chr6 | 15780267 | 15780417 | + | 0 | NA | intron (NM | L2\|LINE\|L | 59681 | NM_17508 | 16543 | Mm.1314 | NM_17508 | ENSMUSG | Mdfic | Kdt1\|Mdfi | MyoD fam | protein-co | 5.8 | 6.35 | 4.9 | 1.45 | 5.45E-05 | 0.00268 |
| 1043 | chr11 | 97977913 | 97978063 | + | 0 | NA | intron (NM | intron (NM | 8458 | NM_00116 | 72324 | Mm.39617 | NM_02819 | ENSMUSG | Plxdc1 | 2410003I0 | plexin don | protein-co | 4.51 | 5.28 | 2.78 | 2.5 | 5.52E-05 | 0.00271 |
| 1044 | chr14 | 1.21E+08 | 1.21E+08 | + | 0 | NA | intron (NM | intron (NM | 54156 | NM_14546 | 223255 | Mm.39075 | NM_14546 | ENSMUSG | Stk24 | 1810013H | serine/thr | protein-co | 6.95 | 7.39 | 6.32 | 1.07 | 5.54E-05 | 0.00272 |
| 1050 | chr16 | 76354428 | 76354578 | + | 0 | NA | intron (NM | intron (NM | 18546 | NM_17344 | 268903 | Mm.74711 | NM_17344 | ENSMUSG | Nrip1 | 6030458L | nuclear re | protein-co | 4.61 | 5.34 | 3.07 | 2.27 | 5.72E-05 | 0.00279 |
| 1049 | chr18 | 74295004 | 74295154 | + | 0 | NA | intron (NR | intron (NR | 11979 | NM_02894 | 74453 | Mm.18229 | NM_02894 | ENSMUSG | Cfap53 | 4933415I0 | cilia and fl | protein-co | 5.91 | 6.45 | 5.05 | 1.4 | 5.70E-05 | 0.00279 |
| 1051 | chr4 | 1.03E+08 | 1.03E+08 | + | 0 | NA | intron (NM | intron (NM | -58405 | NM_00117 | 18578 | Mm.20181 | NM_01984 | ENSMUSG | Pde4b | Dpde4\|R7 | phosphodi | protein-co | 7.11 | 7.53 | 6.52 | 1.01 | 5.73E-05 | 0.0028 |
| 1052 | chr2 | 1.53E+08 | 1.53E+08 | + | 0 | NA | intron (NM | intron (NM | 33782 | NR_14925 | 12048 | Mm.23821 | NM_00974 | ENSMUSG | Bcl2l1 | Bcl(X)L\|Bc | BCL2-like 1 | protein-co | 6.17 | 6.72 | 5.25 | 1.46 | 5.76E-05 | 0.00281 |
| 1054 | chr1 | 1.61E+08 | 1.61E+08 | + | 0 | NA | intron (NM | intron (NM | 69613 | NM_00114 | 620246 | | NM_001146330 | | Gpr52 | EG620246 | G protein- | protein-co | 5.78 | 4.86 | 6.34 | -1.48 | 5.82E-05 | 0.00283 |
| 1056 | chr1 | 91181538 | 91181688 | + | 0 | NA | intron (NM | intron (NM | 1791 | NM_01689 | 51801 | Mm.3272 | NM_01689 | ENSMUSG | Ramp1 | 9130218E | receptor (c | protein-co | 5.38 | 5.99 | 4.31 | 1.68 | 5.85E-05 | 0.00284 |
| 1055 | chr17 | 34129463 | 34129613 | + | 0 | NA | intron (NM | B1F\|SINE\| | 4020 | NM_00136 | 14998 | Mm.16373 | NM_01038 | ENSMUSG | H2-DMa | H-2Ma\|H2 | histocomp | protein-co | 5.6 | 6.18 | 4.6 | 1.59 | 5.85E-05 | 0.00284 |
| 1057 | chr11 | 1.22E+08 | 1.22E+08 | + | 0 | NA | intron (NM | intron (NM | 17175 | NM_17876 | 319530 | Mm.11395 | NM_17876 | ENSMUSG | Zfp750 | A030007D | zinc finger | protein-co | 5.12 | 5.79 | 3.84 | 1.95 | 5.89E-05 | 0.00286 |
| 1058 | chr4 | 1.42E+08 | 1.42E+08 | + | 0 | NA | intron (NM | intron (NM | 17907 | NM_00101 | 68817 | Mm.39372 | NM_00101 | ENSMUSG | Ddi2 | 1110056G | DNA-dama | protein-co | 5.45 | 4.43 | 6.04 | -1.6 | 5.92E-05 | 0.00287 |
| 1060 | chr15 | 6602099 | 6602249 | + | 0 | NA | intron (NM | intron (NM | 22327 | NM_01181 | 23880 | Mm.1709( | NM_01181 | ENSMUSG | Fyb | ADAP\|B63 | FYN bindir | protein-co | 5.92 | 6.44 | 5.08 | 1.37 | 6.04E-05 | 0.00292 |
| 1063 | chr17 | 35115539 | 35115689 | + | 0 | NA | promoter- | promoter- | -105 | NM_14893 | 266614 | Mm.10468 | NM_14893 | ENSMUSG | Ly6g5b | - | lymphocyt | protein-co | 5.99 | 6.51 | 5.17 | 1.33 | 6.06E-05 | 0.00292 |
| 1064 | chr1 | 52856383 | 52856533 | + | 0 | NA | intron (NM | RLTR21\|LT | 11420 | NM_1461( | 227095 | Mm.22206 | NM_1461( | ENSMUSG | Hibch | 2610509I1 | 3-hydroxyi | protein-co | 6.28 | 5.55 | 6.77 | -1.21 | 6.12E-05 | 0.00294 |
| 1065 | chr14 | 75192883 | 75193033 | + | 0 | NA | intron (NM | intron (NM | -49329 | NM_01977 | 56373 | Mm.24242 | NM_01977 | ENSMUSG | Cpb2 | 1110032P( | carboxype | protein-co | 6.27 | 6.78 | 5.46 | 1.32 | 6.23E-05 | 0.00299 |
| 1066 | chr6 | 22347621 | 22347771 | + | 0 | NA | intron (NM | intron (NM | 8385 | NM_13858 | 27999 | Mm.4903( | NM_13858 | ENSMUSG | Fam3c | D6Wsu176 | family with | protein-co | 6.67 | 7.12 | 6.03 | 1.09 | 6.24E-05 | 0.003 |
| 1069 | chr7 | 76408281 | 76408431 | + | 0 | NA | intron (NM | RMER15\|L | 178469 | NM_00119 | 244071 | Mm.32282 | NM_1729( | ENSMUSG | Agbl1 | Ccp4\|D43( | ATP/GTP b | protein-co | 4.93 | 5.62 | 3.59 | 2.02 | 6.31E-05 | 0.00302 |
| 1071 | chr19 | 36375484 | 36375634 | + | 0 | NA | intron (NM | intron (NM | -3508 | NM_0295( | 76073 | Mm.25918 | NM_0295( | ENSMUSG | Pcgf5 | 0610009F( | polycomb | protein-co | 6.43 | 6.89 | 5.74 | 1.15 | 6.35E-05 | 0.00303 |
| 1070 | chr19 | 56296839 | 56296989 | + | 0 | NA | intron (NM | intron (NM | 9002 | NM_1461( | 226243 | Mm.25791 | NM_1461( | ENSMUSG | Habp2 | AI035669\| | hyaluronic | protein-co | 6.22 | 6.74 | 5.4 | 1.34 | 6.33E-05 | 0.00303 |
| 1073 | chr11 | 1.03E+08 | 1.03E+08 | + | 0 | NA | intron (NM | intron (NM | -11695 | NM_0012( | 544817 | Mm.25911 | NM_13371 | ENSMUSG | Arhgap27 | 2310069I0 | Rho GTPas | protein-co | 5.92 | 6.46 | 5.07 | 1.39 | 6.47E-05 | 0.00308 |
| 1077 | chr11 | 74577161 | 74577311 | + | 0 | NA | intron (NR | intron (NR | 12950 | NM_00101 | 380711 | Mm.42569 | NM_00101 | ENSMUSG | Rap1gap2 | AU067654 | RAP1 GTPa | protein-co | 5.28 | 5.9 | 4.17 | 1.73 | 6.57E-05 | 0.00311 |
| 1078 | chr10 | 84702108 | 84702258 | + | 0 | NA | intron (NM | B4A\|SINE\| | -53865 | NM_00102 | 71137 | Mm.7907( | NM_02768 | ENSMUSG | Rfx4 | 4933412G | regulatory | protein-co | 6.42 | 5.72 | 6.89 | -1.17 | 6.62E-05 | 0.00314 |
| 1080 | chr15 | 73156531 | 73156681 | + | 0 | NA | intron (NM | ID_B1\|SIN | 28341 | NM_15317 | 239528 | Mm.23095 | NM_15317 | ENSMUSG | Ago2 | 1110029L1 | argonaute | protein-co | 6.13 | 6.64 | 5.32 | 1.31 | 6.73E-05 | 0.00318 |
| 1081 | chr7 | 63922048 | 63922198 | + | 0 | NA | intron (NM | intron (NM | 5266 | NM_00125 | 381994 | Mm.42546 | NM_00125 | ENSMUSG | E030018B | Gm1697 | RIKEN cDN | protein-co | 5.13 | 5.78 | 3.9 | 1.88 | 6.76E-05 | 0.00319 |
| 1083 | chr9 | 73073312 | 73073462 | + | 0 | NA | intron (NM | intron (NM | 9239 | NM_0013( | 11891 | Mm.34867 | NM_02363 | ENSMUSG | Rab27a | 2210402C( | RAB27A, m | protein-co | 6.04 | 6.55 | 5.24 | 1.31 | 6.78E-05 | 0.0032 |
| 1085 | chr11 | 34050802 | 34050952 | + | 0 | NA | intron (NM | intron (NM | 3949 | NM_01069 | 16822 | Mm.26535 | NM_01069 | ENSMUSG | Lcp2 | AI323664\| | lymphocyt | protein-co | 6.83 | 7.25 | 6.24 | 1.01 | 6.87E-05 | 0.00323 |
| 1084 | chr4 | 9526322 | 9526472 | + | 0 | NA | intron (NM | intron (NM | 117320 | NM_00117 | 65973 | Mm.2222( | NM_02306 | ENSMUSG | Asph | 2310005F1 | aspartate- | protein-co | 5.18 | 4.07 | 5.81 | -1.74 | 6.86E-05 | 0.00323 |
| 1089 | chr15 | 66722039 | 66722189 | + | 0 | NA | intron (NM | intron (NM | 51344 | NM_00937 | 21819 | Mm.44133 | NM_00937 | ENSMUSG | Tg | Tgn\|cog | thyroglobu | protein-co | 5.06 | 5.72 | 3.83 | 1.89 | 6.93E-05 | 0.00324 |
| 1087 | chr7 | 1.26E+08 | 1.26E+08 | + | 0 | NA | intron (NM | intron (NM | 3272 | NM_02188 | 60504 | Mm.15564 | NM_02188 | ENSMUSG | Il21r | NILR | interleukir | protein-co | 5.88 | 6.42 | 4.99 | 1.43 | 6.90E-05 | 0.00324 |
| 1091 | chr1 | 1.66E+08 | 1.66E+08 | + | 0 | NA | intron (NM | intron (NM | 16406 | NM_17859 | 226594 | Mm.26555 | NM_17859 | ENSMUSG | Rcsd1 | A430105K | RCSD dom | protein-co | 6.55 | 7.01 | 5.86 | 1.15 | 7.08E-05 | 0.00331 |
| 1094 | chr4 | 62686811 | 62686961 | + | 0 | NA | intron (NM | intron (NM | 12912 | NM_00131 | 50780 | Mm.28675 | NM_01949 | ENSMUSG | Rgs3 | 4930506N | regulator ( | protein-co | 5.65 | 6.21 | 4.71 | 1.51 | 7.13E-05 | 0.00332 |
| 1095 | chr16 | 55887986 | 55888136 | + | 0 | NA | intron (NM | intron (NM | 7218 | NM_00113 | 385658 | Mm.31063 | NM_00113 | ENSMUSG | Nxpe3 | Fam55c\|G | neurexoph | protein-co | 6.81 | 7.23 | 6.22 | 1.01 | 7.16E-05 | 0.00333 |
| 1096 | chr11 | 46310640 | 46310790 | + | 0 | NA | intron (NM | intron (NM | 1643 | NM_13376 | 76884 | Mm.15435 | NM_13376 | ENSMUSG | Cyfip2 | 1500004I0 | cytoplasm | protein-co | 6.38 | 6.88 | 5.62 | 1.25 | 7.25E-05 | 0.00337 |
| 1098 | chr16 | 13845134 | 13845284 | + | 0 | NA | intron (NM | intron (NM | 25958 | NM_00134 | 18203 | Mm.38041 | NM_01094 | ENSMUSG | Ntan1 | PNAA\|PN | N-termina | protein-co | 6.76 | 7.2 | 6.13 | 1.07 | 7.31E-05 | 0.00339 |
| 1099 | chr18 | 67212755 | 67212905 | + | 0 | NA | intron (NM | intron (NM | 7471 | NM_02419 | 67064 | Mm.73777 | NM_02419 | ENSMUSG | Chmp1b | 2810405I1 | charged m | protein-co | 6.64 | 7.09 | 5.99 | 1.11 | 7.33E-05 | 0.0034 |
| 1100 | chr2 | 1.02E+08 | 1.02E+08 | + | 0 | NA | intron (NM | intron (NM | 146905 | NM_17888 | 241576 | Mm.48974 | NM_17888 | ENSMUSG | Ldlrad3 | 6430500P( | low densit | protein-co | 5.1 | 5.75 | 3.91 | 1.84 | 7.47E-05 | 0.00346 |
| 1102 | chr3 | 1.54E+08 | 1.54E+08 | + | 0 | NA | intron (NM | intron (NM | 1283 | NM_01137 | 20447 | Mm.29645 | NM_01137 | ENSMUSG | St6galnac3 | Siat7c | ST6 (alpha | protein-co | 6.79 | 7.22 | 6.17 | 1.05 | 7.53E-05 | 0.00348 |
| 1104 | chr4 | 1.23E+08 | 1.23E+08 | + | 0 | NA | intron (NM | intron (NM | 11983 | NM_00891 | 19063 | Mm.27771 | NM_00891 | ENSMUSG | Ppt1 | 9530043G | palmitoyl- | protein-co | 6.02 | 6.54 | 5.19 | 1.34 | 7.69E-05 | 0.00354 |
| 1105 | chr11 | 1.05E+08 | 1.05E+08 | + | 0 | NA | 3' UTR (NM | 3' UTR (NM | 59600 | NM_01678 | 16416 | Mm.8715( | NM_01678 | ENSMUSG | Itgb3 | CD61\|GP3 | integrin be | protein-co | 6.27 | 6.74 | 5.55 | 1.19 | 7.74E-05 | 0.00356 |
| 1106 | chr13 | 59456312 | 59456462 | + | 0 | NA | intron (NM | intron (NM | 88064 | NM_00128 | 67269 | Mm.1530( | NM_02332 | ENSMUSG | Agtpbp1 | 1700020N | ATP/GTP b | protein-co | 6.36 | 6.83 | 5.66 | 1.18 | 7.74E-05 | 0.00356 |
| 1107 | chr16 | 94589128 | 94589278 | + | 0 | NA | intron (NM | intron (NM | 18296 | NM_00111 | 13548 | Mm.31097 | NM_00789 | ENSMUSG | Dyrk1a | 2310043O | dual-speci | protein-co | 5.45 | 6.04 | 4.44 | 1.59 | 7.84E-05 | 0.0036 |
| 1109 | chr7 | 1.16E+08 | 1.16E+08 | + | 0 | NA | intron (NM | intron (NM | 35614 | NM_0013( | 233765 | Mm.3741 | NM_17274 | ENSMUSG | Plekha7 | A430081P | pleckstrin | protein-co | 5.05 | 5.7 | 3.83 | 1.87 | 7.96E-05 | 0.00365 |
| 1110 | chr7 | 75724174 | 75724324 | + | 0 | NA | intron (NM | intron (NM | -124089 | NM_00112 | 207952 | Mm.20622 | NM_02965 | ENSMUSG | Klhl25 | 2810402K | kelch-like | protein-co | 5.64 | 4.73 | 6.2 | -1.47 | 8.00E-05 | 0.00366 |
| 1114 | chr9 | 21909557 | 21909707 | + | 0 | NA | 3' UTR (NM | 3' UTR (NM | 8551 | NM_03187 | 19340 | Mm.26015 | NM_03187 | ENSMUSG | Rab3d | C130057E | RAB3D, me | protein-co | 5.65 | 6.21 | 4.72 | 1.49 | 8.18E-05 | 0.00373 |
| 1119 | chr2 | 1.32E+08 | 1.32E+08 | + | 0 | NA | intron (NM | B1_Mus2\| | -9810 | NM_00129 | 110911 | Mm.2845( | NM_13865 | ENSMUSG | Cds2 | 5730450N | CDP-diacyl | protein-co | 4.39 | 5.14 | 2.72 | 2.42 | 8.34E-05 | 0.00378 |
| 1123 | chr13 | 59672415 | 59672565 | + | 0 | NA | intron (NM | intron (NM | 3294 | NM_00103 | 105348 | Mm.17133 | NM_0273( | ENSMUSG | Golm1 | 2310001L( | golgi mem | protein-co | 6.13 | 6.62 | 5.37 | 1.25 | 8.45E-05 | 0.00382 |
| 1125 | chr15 | 25865632 | 25865782 | + | 0 | NA | intron (NM | L2b\|LINE\| | 22409 | NM_00103 | 66270 | Mm.25311 | NM_02545 | ENSMUSG | Retreg1 | 1810015C( | reticulophagy | protein-co | 6.66 | 7.12 | 5.98 | 1.13 | 8.51E-05 | 0.00384 |
| 1129 | chr1 | 1.65E+08 | 1.65E+08 | + | 0 | NA | intron (NM | intron (NM | 6449 | NM_0320( | 83993 | Mm.19064 | NM_0320( | ENSMUSG | Tbx19 | AU019250 | T-box 19 | protein-co | 5.2 | 5.83 | 4.06 | 1.77 | 8.66E-05 | 0.00389 |
| 1132 | chr1 | 87828321 | 87828471 | + | 0 | NA | intron (NM | intron (NM | 24716 | NM_00911 | 20215 | Mm.1276 | NM_00911 | ENSMUSG | Sag | A930001K | S-antigen, | protein-co | 6.24 | 6.72 | 5.52 | 1.21 | 8.73E-05 | 0.00391 |
| 1130 | chr2 | 1.27E+08 | 1.27E+08 | + | 0 | NA | intron (NM | intron (NM | 12406 | NM_00823 | 15186 | Mm.18603 | NM_00823 | ENSMUSG | Hdc | AW10818 | histidine d | protein-co | 6.82 | 7.28 | 6.17 | 1.11 | 8.71E-05 | 0.00391 |
| 1133 | chr1 | 91195733 | 91195883 | + | 0 | NA | intron (NM | intron (NM | 15986 | NM_01689 | 51801 | Mm.3272 | NM_01689 | ENSMUSG | Ramp1 | 9130218E | receptor (c | protein-co | 5.34 | 5.94 | 4.3 | 1.65 | 8.75E-05 | 0.00392 |
| 1135 | chr10 | 75697125 | 75697275 | + | 0 | NA | intron (NM | intron (NM | -53192 | NM_02789 | 71733 | Mm.24795 | NM_02789 | ENSMUSG | Susd2 | 1200011D | sushi dom | protein-co | 5.86 | 6.39 | 5.01 | 1.39 | 8.81E-05 | 0.00394 |
| 1136 | chr7 | 17024345 | 17024495 | + | 0 | NA | intron (NM | intron (NM | 3494 | NM_01115 | 19060 | Mm.3294 | NM_01115 | ENSMUSG | Ppp5c | AU020526 | protein ph | protein-co | 6.59 | 7.04 | 5.94 | 1.1 | 8.90E-05 | 0.00397 |
| 1138 | chr13 | 24613561 | 24613711 | + | 0 | NA | promoter- | promoter- | -970 | NM_00128 | 193385 | Mm.21731 | NM_02967 | ENSMUSG | Ripor2 | Fam65b | RHO famil | protein-co | 6.32 | 6.8 | 5.59 | 1.2 | 8.99E-05 | 0.00401 |
| 1137 | chr2 | 65551877 | 65552027 | + | 0 | NA | intron (NM | MER5A\|DI | 15540 | NM_00135 | 20269 | Mm.33025 | NM_01873 | ENSMUSG | Scn3a | Gm1000\| | sodium ch | protein-co | 5.31 | 5.93 | 4.21 | 1.72 | 8.97E-05 | 0.00401 |
| 1140 | chr4 | 1.39E+08 | 1.39E+08 | + | 0 | NA | intron (NM | RSINE1\|SII | 20574 | NM_00127 | 12345 | Mm.2945 | NM_00979 | ENSMUSG | Capzb | 1700120C( | capping pr | protein-co | 5.51 | 6.08 | 4.55 | 1.53 | 9.17E-05 | 0.00408 |
| 1141 | chr5 | 1.49E+08 | 1.49E+08 | + | 0 | NA | intron (NM | intron (NM | -10783 | NM_02946 | 75860 | Mm.1596( | NM_02946 | ENSMUSG | Tex26 | 4930588N | testis expr | protein-co | 5.56 | 6.13 | 4.6 | 1.53 | 9.20E-05 | 0.00409 |
| 1143 | chr16 | 91490980 | 91491130 | + | 0 | NA | intron (NM | intron (NM | 5840 | NM_0105( | 15975 | Mm.502 | NM_0105( | ENSMUSG | Ifnar1 | Ifar\|Ifnar\| | interferon | protein-co | 6.7 | 7.13 | 6.1 | 1.03 | 9.30E-05 | 0.00413 |
| 1144 | chr1 | 52150750 | 52150900 | + | 0 | NA | intron (NM | intron (NM | 31203 | NM_0012( | 20846 | Mm.2774( | NM_00928 | ENSMUSG | Stat1 | 2010005J0 | signal tran | protein-co | 5.04 | 5.7 | 3.79 | 1.91 | 9.42E-05 | 0.00418 |

| | | | | | | | | | | | | | | | | | | | | | | | | |
|---|---|---|---|---|---|---|---|---|---|---|---|---|---|---|---|---|---|---|---|---|---|---|---|---|
| 1145 | chr9 | 42028946 | 42029096 | + | 0 | NA | intron (NM | intron (NM | 48192 | NM_00135 | 20660 | Mm.12192 | NM_0114: | ENSMUSG | Sorl1 | 2900010L1 | sortilin-rel | protein-co | 6.21 | 6.7 | 5.45 | 1.26 | 9.46E-05 | 0.00419 |
| 1147 | chr18 | 20982556 | 20982706 | + | 0 | NA | intron (NM | intron (NM | -18669 | NM_0197( | 56515 | Mm.25354 | NM_0197( | ENSMUSG | Rnf138 | 2410015A: | ring finger | protein-co | 5.17 | 5.8 | 4.04 | 1.75 | 9.50E-05 | 0.0042 |
| 1150 | chr1 | 1.28E+08 | 1.28E+08 | + | 0 | NA | intron (NM | MTD\|LTR\| | 66191 | NM_02813 | 72160 | Mm.28128 | NM_02813 | ENSMUSG | Tmem163 | 2610024A( | transmeml | protein-co | 6.14 | 6.63 | 5.38 | 1.25 | 9.59E-05 | 0.00423 |
| 1153 | chr14 | 66189059 | 66189209 | + | 0 | NA | intron (NM | MER63A\|D | 24690 | NM_00136 | 19229 | Mm.21613 | NM_17249 | ENSMUSG | Ptk2b | CADTK\|CA | PTK2 prote | protein-co | 5.29 | 5.89 | 4.23 | 1.66 | 9.64E-05 | 0.00424 |
| 1155 | chr13 | 92244862 | 92245012 | + | 0 | NA | intron (NM | B3A\|SINE\| | -109846 | NM_01004 | 13361 | Mm.23695 | NM_01004 | ENSMUSG | Dhfr | 8430436I0 | dihydrofol | protein-co | 5.59 | 6.15 | 4.66 | 1.49 | 9.68E-05 | 0.00425 |
| 1154 | chr17 | 12756069 | 12756219 | + | 0 | NA | intron (NM | intron (NM | 13562 | NM_01051 | 16004 | Mm.26553 | NM_01051 | ENSMUSG | Igf2r | AI661837\| | insulin-like | protein-co | 6.46 | 6.91 | 5.8 | 1.11 | 9.68E-05 | 0.00425 |
| 1156 | chr17 | 13715632 | 13715782 | + | 0 | NA | TTS (NM_( | TTS (NM_( | -44834 | NR_15173 | 17356 | Mm.59167 | NM_0108( | ENSMUSG | Afdn | 5033403D: | afadin, adh | protein-co | 6.86 | 7.32 | 6.18 | 1.13 | 9.79E-05 | 0.00429 |
| 1158 | chr2 | 30257875 | 30258025 | + | 0 | NA | intron (NM | intron (NM | -8253 | NM_17226 | 227696 | Mm.29794 | NM_17226 | ENSMUSG | Phyhd1 | - | phytanoyl- | protein-co | 5.44 | 6.02 | 4.46 | 1.56 | 9.84E-05 | 0.00431 |
| 1157 | chr7 | 25006730 | 25006880 | + | 0 | NA | promoter- | promoter- | -868 | NM_00129 | 232975 | Mm.44101 | NM_14492 | ENSMUSG | Atp1a3 | Atpa-2 | ATPase, Na | protein-co | 5.84 | 6.4 | 4.94 | 1.46 | 9.84E-05 | 0.00431 |
| 1159 | chr1 | 66841979 | 66842129 | + | 0 | NA | intron (NM | intron (NM | 21255 | NM_00738 | 11363 | Mm.2445 | NM_00738 | ENSMUSG | Acadl | AA960361 | acyl-Coenz | protein-co | 6.61 | 7.06 | 5.96 | 1.1 | 9.92E-05 | 0.00433 |
| 1160 | chr16 | 10700280 | 10700430 | + | 0 | NA | intron (NM | intron (NM | 85181 | NM_00127 | 12703 | Mm.130 | NM_00989 | ENSMUSG | Socs1 | Cish1\|Cish | suppresso | protein-co | 5.95 | 6.47 | 5.12 | 1.34 | 9.93E-05 | 0.00433 |
| 1163 | chr10 | 39564772 | 39564922 | + | 0 | NA | 3' UTR (NM | 3' UTR (NM | -48087 | NM_1340( | 103213 | Mm.43668 | NM_1340( | ENSMUSG | Traf3ip2 | AI429613\| | TRAF3 inte | protein-co | 4.29 | 5.06 | 2.53 | 2.54 | 1.01E-04 | 0.00438 |
| 1164 | chr7 | 80204484 | 80204634 | + | 0 | NA | intron (NM | intron (NM | 6118 | NM_00136 | 20352 | Mm.2759( | NM_01365 | ENSMUSG | Sema4b | SemC\|Sem | sema dom | protein-co | 6.43 | 6.89 | 5.76 | 1.13 | 1.02E-04 | 0.00441 |
| 1169 | chr14 | 30875548 | 30875698 | + | 0 | NA | 3' UTR (NM | 3' UTR (NM | -3634 | NM_18139 | 66175 | Mm.22089 | NM_18139 | ENSMUSG | Mustn1 | 1110028G( | musculosk | protein-co | 5.78 | 6.31 | 4.92 | 1.4 | 1.03E-04 | 0.00447 |
| 1172 | chr5 | 1.04E+08 | 1.04E+08 | + | 0 | NA | intron (NM | intron (NM | 1887 | NM_00108 | 17355 | Mm.6949 | NM_13391 | ENSMUSG | Aff1 | 9630032B( | AF4/FMR2 | protein-co | 6.77 | 7.21 | 6.13 | 1.08 | 1.05E-04 | 0.00455 |
| 1173 | chr11 | 59421588 | 59421738 | + | 0 | NA | intron (NM | intron (NM | 28371 | NM_00135 | 67826 | Mm.2668( | NM_14452 | ENSMUSG | Snap47 | 1110031B( | synaptoso | protein-co | 6.5 | 6.96 | 5.83 | 1.13 | 1.07E-04 | 0.0046 |
| 1175 | chr17 | 45558142 | 45558292 | + | 0 | NA | intron (NM | intron (NM | 2517 | NM_0013( | 18037 | Mm.57043 | NM_00869 | ENSMUSG | Nfkbie | I-kappa-B- | nuclear fac | protein-co | 6.48 | 6.93 | 5.82 | 1.11 | 1.09E-04 | 0.0047 |
| 1176 | chr6 | 71871515 | 71871665 | + | 0 | NA | intron (NM | intron (NM | 37172 | NM_02727 | 69956 | Mm.30256 | NM_02727 | ENSMUSG | Ptcd3 | 2610034F1 | pentatrico | protein-co | 5.57 | 6.13 | 4.63 | 1.51 | 1.10E-04 | 0.00471 |
| 1179 | chr4 | 6389535 | 6389685 | + | 0 | NA | intron (NM | intron (NM | 23930 | NM_00109 | 53378 | Mm.24747 | NM_0168( | ENSMUSG | Sdcbp | MDA-9\|Sy | syndecan b | protein-co | 5.17 | 5.79 | 4.07 | 1.72 | 1.11E-04 | 0.00474 |
| 1181 | chr7 | 19790035 | 19790185 | + | 0 | NA | intron (NM | intron (NM | 6699 | NM_00116 | 80794 | Mm.81698 | NM_02322 | ENSMUSG | Cblc | 2310076I2 | Casitas B-l | protein-co | 6.69 | 7.11 | 6.08 | 1.04 | 1.12E-04 | 0.00479 |
| 1182 | chr8 | 25611976 | 25612126 | + | 0 | NA | intron (NM | B4A\|SINE\| | 9767 | NM_0010( | 234135 | Mm.21733 | NM_0010( | ENSMUSG | Nsd3 | 6720429E( | nuclear rec | protein-co | 5.08 | 5.71 | 3.92 | 1.79 | 1.12E-04 | 0.00479 |
| 1183 | chr11 | 90548682 | 90548832 | + | 0 | NA | intron (NM | intron (NM | 89351 | NM_0115( | 20913 | Mm.2072( | NM_0115( | ENSMUSG | Stxbp4 | 6030470M | syntaxin bi | protein-co | 5.51 | 6.08 | 4.56 | 1.52 | 1.12E-04 | 0.0048 |
| 1185 | chr4 | 1.45E+08 | 1.45E+08 | + | 0 | NA | intron (NM | intron (NM | 133664 | NM_00135 | 20148 | Mm.14063 | NM_0113( | ENSMUSG | Dhrs3 | Rsdr1\|ret5 | dehydroge | protein-co | 5.83 | 6.38 | 4.96 | 1.42 | 1.14E-04 | 0.00486 |
| 1189 | chr16 | 92818642 | 92818792 | + | 0 | NA | intron (NM | intron (NM | 7357 | NM_00111 | 12394 | Mm.4081 | NM_00982 | ENSMUSG | Runx1 | AML1\|CBF | runt relate | protein-co | 6.17 | 6.65 | 5.45 | 1.21 | 1.15E-04 | 0.0049 |
| 1194 | chr19 | 45486318 | 45486468 | + | 0 | NA | intron (NM | intron (NM | 74171 | NM_00133 | 56626 | Mm.46509 | NM_02003 | ENSMUSG | Poll | 1110003P( | polymeras | protein-co | 5.67 | 6.26 | 4.67 | 1.6 | 1.17E-04 | 0.00494 |
| 1193 | chr2 | 1.22E+08 | 1.22E+08 | + | 0 | NA | exon (NM_ | exon (NM_ | -7451 | NM_14553 | 214585 | Mm.2223( | NM_14553 | ENSMUSG | Spg11 | 6030465E2 | SPG11, spa | protein-co | 5.48 | 6.06 | 4.48 | 1.58 | 1.17E-04 | 0.00494 |
| 1192 | chr7 | 1.25E+08 | 1.25E+08 | + | 0 | NA | intron (NM | L1ME3E\|L | 2578 | NM_00136 | 67711 | Mm.4467 | NM_02633 | ENSMUSG | Nsmce1 | 2510027N | NSE1 hom | protein-co | 6.42 | 6.88 | 5.74 | 1.14 | 1.17E-04 | 0.00494 |
| 1197 | chr12 | 1.08E+08 | 1.08E+08 | + | 0 | NA | intron (NM | intron (NM | 75201 | NM_02139 | 58208 | Mm.39269 | NM_02139 | ENSMUSG | Bcl11b | 9130430L1 | B cell leuk | protein-co | 6.04 | 6.55 | 5.26 | 1.29 | 1.18E-04 | 0.00496 |
| 1200 | chr11 | 1.17E+08 | 1.17E+08 | + | 0 | NA | intron (NM | intron (NM | -13730 | NM_01738 | 53860 | Mm.3845( | NM_01738 | ENSMUSG | 9-Sep | MSF1\|Msf | septin 9 | protein-co | 6.64 | 7.07 | 6.03 | 1.04 | 1.19E-04 | 0.00498 |
| 1199 | chr6 | 87740624 | 87740774 | + | 0 | NA | intron (NM | intron (NM | 9830 | NM_00115 | 58229 | Mm.33322 | NM_02141 | ENSMUSG | Efcc1 | Ccdc48 | EF hand ar | protein-co | 6.4 | 6.85 | 5.73 | 1.12 | 1.18E-04 | 0.00498 |
| 1201 | chr16 | 94846129 | 94846279 | + | 0 | NA | intron (NM | intron (NM | 10486 | NM_00102 | 16522 | Mm.32872 | NM_0106( | ENSMUSG | Kcnj6 | BIR1\|GIRK | potassium | protein-co | 4.98 | 5.62 | 3.78 | 1.84 | 1.19E-04 | 0.00499 |
| 1202 | chr11 | 11737064 | 11737214 | + | 0 | NA | intron (NM | intron (NM | 46962 | NM_0013( | 22778 | Mm.10354 | NM_00957 | ENSMUSG | Ikzf1 | 5832432G | IKAROS far | protein-co | 6.6 | 7.04 | 5.96 | 1.08 | 1.20E-04 | 0.00503 |
| 1203 | chr18 | 80869009 | 80869159 | + | 0 | NA | intron (NM | intron (NM | 64974 | NM_0012( | 50771 | Mm.24713 | NM_0158( | ENSMUSG | Atp9b | AA934181 | ATPase, cla | protein-co | 6.18 | 5.43 | 6.67 | -1.24 | 1.21E-04 | 0.00505 |
| 1204 | chr2 | 1.58E+08 | 1.58E+08 | + | 0 | NA | intron (NM | intron (NM | 81417 | NM_02568 | 66642 | Mm.45193 | NM_02568 | ENSMUSG | Ctnnbl1 | 5730471K( | catenin, be | protein-co | 5.48 | 6.05 | 4.53 | 1.52 | 1.21E-04 | 0.00507 |
| 1205 | chr6 | 1.28E+08 | 1.28E+08 | + | 0 | NA | intron (NM | intron (NM | 117768 | NM_17541 | 109246 | Mm.21814 | NM_17541 | ENSMUSG | Tspan9 | 6720474K: | tetraspani | protein-co | 5.67 | 6.21 | 4.79 | 1.42 | 1.21E-04 | 0.00508 |
| 1208 | chr9 | 57956982 | 57957132 | + | 0 | NA | intron (NM | intron (NM | 16945 | NM_01135 | 20361 | Mm.33518 | NM_01135 | ENSMUSG | Sema7a | 2900057C( | sema dom | protein-co | 5.19 | 5.81 | 4.1 | 1.71 | 1.24E-04 | 0.00514 |
| 1212 | chr9 | 42465685 | 42465835 | + | 0 | NA | intron (NM | intron (NM | -4299 | NM_00135 | 272589 | Mm.34483 | NM_17303 | ENSMUSG | Tbcel | D330014K: | tubulin fol | protein-co | 6.29 | 6.78 | 5.54 | 1.23 | 1.27E-04 | 0.00526 |
| 1214 | chr13 | 81690650 | 81690800 | + | 0 | NA | intron (NM | intron (NM | 20288 | NM_00108 | 67486 | Mm.27978 | NM_00108 | ENSMUSG | Polr3g | 2310047G: | polymeras | protein-co | 6.92 | 7.34 | 6.33 | 1 | 1.28E-04 | 0.00529 |
| 1219 | chr11 | 75652248 | 75652398 | + | 0 | NA | 5' UTR (NM | 5' UTR (NM | 173 | NM_00108 | 17913 | Mm.2345( | NM_00865 | ENSMUSG | Myo1c | C80397\|M | myosin IC | protein-co | 6.22 | 6.71 | 5.47 | 1.24 | 1.29E-04 | 0.00531 |
| 1221 | chr11 | 1.09E+08 | 1.09E+08 | + | 0 | NA | intron (NM | MTEa\|LTR | 943 | NM_00136 | 74008 | Mm.48222 | NM_02871 | ENSMUSG | Arsg | 6330406P( | arylsulfata | protein-co | 6.85 | 7.26 | 6.26 | 1 | 1.29E-04 | 0.00531 |
| 1222 | chr15 | 10662271 | 10662421 | + | 0 | NA | intron (NM | intron (NM | 51194 | NM_03069 | 75646 | Mm.21239 | NM_03069 | ENSMUSG | Rai14 | 1700008J1 | retinoic ac | protein-co | 6.02 | 6.54 | 5.2 | 1.34 | 1.30E-04 | 0.00531 |
| 1223 | chr13 | 1.09E+08 | 1.09E+08 | + | 0 | NA | intron (NM | intron (NM | 39458 | NM_01105 | 238871 | Mm.43442 | NM_01105 | ENSMUSG | Pde4d | 9630011N: | phosphodi | protein-co | 5.59 | 4.63 | 6.17 | -1.53 | 1.30E-04 | 0.00533 |
| 1224 | chr3 | 22151254 | 22151404 | + | 0 | NA | intron (NM | intron (NM | 74677 | NM_03073 | 81004 | Mm.20296 | NM_03073 | ENSMUSG | Tbl1xr1 | 8030499H( | transducin | protein-co | 5.43 | 6.01 | 4.47 | 1.54 | 1.32E-04 | 0.00541 |
| 1226 | chr11 | 4823998 | 4824148 | + | 0 | NA | intron (NM | intron (NM | 9062 | NM_00125 | 18016 | Mm.2971( | NM_01089 | ENSMUSG | Nf2 | merlin | neurofibro | protein-co | 6.3 | 6.76 | 5.61 | 1.14 | 1.33E-04 | 0.00543 |
| 1225 | chr17 | 34630319 | 34630469 | + | 0 | NA | intron (NM | intron (NM | 830 | NM_03089 | 260297 | Mm.39096 | NM_03089 | ENSMUSG | Prrt1 | Bat-4\|Bat4 | proline-ric | protein-co | 5.62 | 6.2 | 4.62 | 1.58 | 1.33E-04 | 0.00543 |
| 1228 | chr6 | 58843694 | 58843844 | + | 0 | NA | exon (NM_ | exon (NM_ | 10069 | NM_0287( | 73998 | Mm.19189 | NM_0287( | ENSMUSG | Herc3 | 5730409F1 | hect doma | protein-co | 5.67 | 6.22 | 4.79 | 1.42 | 1.34E-04 | 0.00546 |
| 1229 | chr9 | 62020375 | 62020525 | + | 0 | NA | intron (NM | L1MB3\|LIN | 6340 | NM_02874 | 74090 | Mm.27326 | NM_02874 | ENSMUSG | Paqr5 | 0610010I1 | progestin a | protein-co | 5.67 | 6.22 | 4.77 | 1.45 | 1.34E-04 | 0.00547 |
| 1231 | chr11 | 76496485 | 76496635 | + | 0 | NA | intron (NM | intron (NM | -2065 | NM_00134 | 109934 | Mm.25893 | NM_19801 | ENSMUSG | Abr | - | active BCR | protein-co | 5.92 | 6.44 | 5.1 | 1.34 | 1.37E-04 | 0.00554 |
| 1230 | chr2 | 18937815 | 18937965 | + | 0 | NA | intron (NM | intron (NM | 60231 | NM_00135 | 18718 | Mm.31397 | NM_00884 | ENSMUSG | Pip4k2a | AW742916 | phosphatic | protein-co | 5.56 | 6.12 | 4.62 | 1.5 | 1.36E-04 | 0.00554 |
| 1232 | chr18 | 84857045 | 84857195 | + | 0 | NA | intron (NM | intron (NM | 5782 | NM_00134 | 109672 | Mm.31018 | NM_02579 | ENSMUSG | Cyb5a | 0610009N | cytochrom | protein-co | 5.21 | 5.82 | 4.13 | 1.69 | 1.37E-04 | 0.00556 |
| 1233 | chr3 | 41604913 | 41605063 | + | 0 | NA | exon (NM_ | exon (NM_ | 40108 | NM_00113 | 269424 | Mm.28628 | NM_1723( | ENSMUSG | Jade1 | AU041499 | jade family | protein-co | 6.71 | 7.16 | 6.07 | 1.09 | 1.38E-04 | 0.00556 |
| 1236 | chr15 | 96583689 | 96583839 | + | 0 | NA | intron (NM | intron (NM | 58198 | NM_13408 | 105727 | Mm.10356 | NM_13408 | ENSMUSG | Slc38a1 | AA408026 | solute carr | protein-co | 4.24 | 5.01 | 2.48 | 2.52 | 1.40E-04 | 0.00565 |
| 1237 | chr12 | 70853176 | 70853326 | + | 0 | NA | intron (NM | intron (NM | 27737 | NM_02812 | 319710 | Mm.2962 | NM_02812 | ENSMUSG | Frmd6 | 2610019M | FERM dom | protein-co | 5.99 | 6.49 | 5.23 | 1.26 | 1.41E-04 | 0.00566 |
| 1239 | chr2 | 71866709 | 71866859 | + | 0 | NA | intron (NR | intron (NR | -6440 | NM_17266 | 228026 | Mm.34411 | NM_17266 | ENSMUSG | Pdk1 | B830012B( | pyruvate d | protein-co | 5.45 | 6.02 | 4.48 | 1.54 | 1.41E-04 | 0.00566 |
| 1240 | chr2 | 79270192 | 79270342 | + | 0 | NA | intron (NM | intron (NM | 14841 | NM_01057 | 16401 | Mm.31903 | NM_01057 | ENSMUSG | Itga4 | CD49D\|Itg | integrin al | protein-co | 5.32 | 5.91 | 4.31 | 1.6 | 1.41E-04 | 0.00567 |
| 1242 | chr14 | 60738639 | 60738789 | + | 0 | NA | intron (NM | L1MB2\|LIN | 5810 | NM_00131 | 219140 | Mm.14977 | NM_00103 | ENSMUSG | Spata13 | C79050\|ES | spermatog | protein-co | 6.74 | 7.16 | 6.15 | 1.01 | 1.42E-04 | 0.00568 |
| 1243 | chr14 | 66232287 | 66232437 | + | 0 | NA | intron (NM | intron (NM | -18538 | NM_00136 | 19229 | Mm.21613 | NM_17249 | ENSMUSG | Ptk2b | CADTK\|CA | PTK2 prote | protein-co | 6.46 | 6.9 | 5.82 | 1.08 | 1.42E-04 | 0.00569 |
| 1244 | chr13 | 40971372 | 40971522 | + | 0 | NA | intron (NR | intron (NR | -29563 | NM_02655 | 68083 | Mm.24789 | NM_02655 | ENSMUSG | Pak1ip1 | 5830431I1 | PAK1 inter | protein-co | 5.05 | 5.7 | 3.86 | 1.84 | 1.42E-04 | 0.0057 |
| 1245 | chr9 | 15300119 | 15300269 | + | 0 | NA | intron (NM | B3\|SINE\|B | 1413 | NM_13373 | 70984 | Mm.26184 | NM_13373 | ENSMUSG | 4931406C( | - | RIKEN cDN | protein-co | 5.54 | 6.1 | 4.59 | 1.51 | 1.43E-04 | 0.00571 |
| 1248 | chr5 | 73241519 | 73241669 | + | 0 | NA | intron (NM | intron (NM | 15024 | NM_02819 | 72313 | Mm.26059 | NM_02819 | ENSMUSG | Fryl | 2010313D: | FRY like tra | protein-co | 6.41 | 6.86 | 5.74 | 1.12 | 1.44E-04 | 0.00576 |
| 1249 | chr8 | 1.17E+08 | 1.17E+08 | + | 0 | NA | intron (NM | RLTR12C\|L | 99053 | NM_02944 | 75796 | Mm.15957 | NM_02944 | ENSMUSG | Cdyl2 | 1700029M | chromodo | protein-co | 5.73 | 6.26 | 4.89 | 1.37 | 1.46E-04 | 0.0058 |
| 1250 | chr9 | 90249197 | 90249347 | + | 0 | NA | intron (NM | intron (NM | 21497 | NM_19433 | 67016 | Mm.33534 | NM_19433 | ENSMUSG | Tbc1d2b | 1810061M | TBC1 doma | protein-co | 5.92 | 5.14 | 6.42 | -1.29 | 1.46E-04 | 0.00582 |
| 1253 | chr18 | 57167645 | 57167795 | + | 0 | NA | intron (NM | intron (NM | 34630 | NM_0010( | 70417 | Mm.29786 | NM_0010( | ENSMUSG | Megf10 | 3000002B( | multiple E( | protein-co | 6.46 | 5.78 | 6.92 | -1.15 | 1.48E-04 | 0.00587 |
| 1254 | chr11 | 1.09E+08 | 1.09E+08 | + | 0 | NA | intron (NM | intron (NM | 6386 | NM_00125 | 13929 | Mm.21185 | NM_02527 | ENSMUSG | Amz2 | AA408420 | archaelysir | protein-co | 5.11 | 5.74 | 3.97 | 1.77 | 1.48E-04 | 0.00588 |
| 1255 | chr11 | 1.16E+08 | 1.16E+08 | + | 0 | NA | TTS (NM_( | TTS (NM_( | -15170 | NM_00125 | 67283 | Mm.38342 | NM_02607 | ENSMUSG | Slc25a19 | 2900089E1 | solute carr | protein-co | 6.73 | 7.16 | 6.13 | 1.02 | 1.49E-04 | 0.0059 |
| 1257 | chr12 | 1.1E+08 | 1.1E+08 | + | 0 | NA | intron (NM | intron (NM | 7949 | NM_00136 | 26931 | Mm.24039 | NM_01202 | ENSMUSG | Ppp2r5c | 2610043M | protein ph | protein-co | 6.12 | 6.61 | 5.39 | 1.21 | 1.50E-04 | 0.00592 |
| 1259 | chr6 | 54187205 | 54187355 | + | 0 | NA | intron (NM | Lx7\|LINE\|L | -77578 | NM_00131 | 69993 | Mm.32114 | NM_02354 | ENSMUSG | Chn2 | 1700026N: | chimerin 2 | protein-co | 6.23 | 5.45 | 6.74 | -1.28 | 1.50E-04 | 0.00593 |
| 1260 | chr6 | 1.19E+08 | 1.19E+08 | + | 0 | NA | exon (NM_ | exon (NM_ | -12794 | NM_17249 | 211187 | Mm.12149 | NM_17249 | ENSMUSG | Lrtm2 | A230084J2 | leucine-ric | protein-co | 5.53 | 6.09 | 4.6 | 1.5 | 1.51E-04 | 0.00594 |
| 1264 | chr7 | 89253447 | 89253597 | + | 0 | NA | intron (NM | ORR1F\|LTI | 85265 | NM_02834 | 72759 | Mm.20847 | NM_02834 | ENSMUSG | Tmem135 | 2810439K( | transmeml | protein-co | 4.33 | 5.07 | 2.72 | 2.35 | 1.54E-04 | 0.00603 |
| 1265 | chr3 | 89961509 | 89961659 | + | 0 | NA | promoter- | promoter- | -987 | NM_00136 | 54667 | Mm.30455 | NM_00108 | ENSMUSG | Atp8b2 | Id | ATPase, cla | protein-co | 6.45 | 6.89 | 5.79 | 1.1 | 1.56E-04 | 0.00609 |
| 1267 | chr16 | 24886993 | 24887143 | + | 0 | NA | intron (NM | intron (NM | 165213 | NM_00114 | 210126 | Mm.20938 | NM_17866 | ENSMUSG | Lpp | 9430020K: | LIM domai | protein-co | 5.15 | 5.77 | 4.06 | 1.7 | 1.56E-04 | 0.0061 |
| 1266 | chr5 | 34976990 | 34977140 | + | 0 | NA | intron (NM | intron (NM | -22013 | NM_00116 | 71729 | Mm.1962( | NM_1734( | ENSMUSG | Rgs12 | 1200016K: | regulator c | protein-co | 4.8 | 5.5 | 3.42 | 2.08 | 1.56E-04 | 0.0061 |
| 1269 | chr18 | 67308153 | 67308303 | + | 0 | NA | intron (NM | intron (NM | 18981 | NM_05326 | 114663 | Mm.34079 | NM_05326 | ENSMUSG | Impa2 | 2210415D: | inositol (m | protein-co | 6.48 | 6.92 | 5.85 | 1.07 | 1.58E-04 | 0.00616 |
| 1271 | chr7 | 82582641 | 82582791 | + | 0 | NA | intron (NM | intron (NM | 65812 | NM_17789 | 330577 | Mm.22281 | NM_17789 | ENSMUSG | Saxo2 | 1700129I0 | stabilizer c | protein-co | 6.07 | 5.34 | 6.55 | -1.21 | 1.60E-04 | 0.00621 |
| 1274 | chr1 | 91942741 | 91942891 | + | 0 | NA | intron (NM | intron (NM | 141355 | NM_00785 | 13345 | Mm.9474 | NM_00785 | ENSMUSG | Twist2 | Dermo1\|b | twist basic | protein-co | 5.52 | 4.56 | 6.1 | -1.53 | 1.60E-04 | 0.00623 |
| 1276 | chr17 | 86615730 | 86615880 | + | 0 | NA | intron (NM | intron (NM | -138059 | NM_01013 | 13819 | Mm.1415 | NM_01013 | ENSMUSG | Epas1 | HIF-2alpha | endothelia | protein-co | 4.62 | 3.29 | 5.31 | -2.01 | 1.61E-04 | 0.00625 |
| 1278 | chr12 | 71933091 | 71933241 | + | 0 | NA | intron (NM | MTD\|LTR\| | 43623 | NM_00128 | 208846 | Mm.87417 | NM_0261( | ENSMUSG | Daam1 | - | dishevelle | protein-co | 6.25 | 6.72 | 5.55 | 1.17 | 1.62E-04 | 0.00628 |
| 1281 | chr18 | 58560954 | 58561104 | + | 0 | NA | intron (NM | intron (NM | 4789 | NM_00108 | 225579 | Mm.25851 | NM_00108 | ENSMUSG | Slc27a6 | 4732438L2 | solute carr | protein-co | 5.6 | 6.15 | 4.7 | 1.45 | 1.63E-04 | 0.00628 |
| 1282 | chr1 | 1.3E+08 | 1.3E+08 | + | 0 | NA | intron (NM | intron (NM | 377303 | NM_17248 | 210417 | Mm.38051 | NM_17248 | ENSMUSG | Thsd7b | 1700074E1 | thrombosp | protein-co | 5.74 | 6.28 | 4.86 | 1.42 | 1.63E-04 | 0.00629 |
| 1285 | chr10 | 99182275 | 99182425 | + | 0 | NA | intron (NR | intron (NR | 74215 | NM_01573 | 14426 | Mm.314 | NM_01573 | ENSMUSG | Galnt4 | AV011803 | polypeptid | protein-co | 5.92 | 6.43 | 5.13 | 1.3 | 1.63E-04 | 0.00629 |
| 1286 | chr14 | 99126856 | 99127006 | + | 0 | NA | intron (NM | Lx7\|LINE\|L | -27161 | NM_00136 | 72662 | Mm.16333 | NM_02831 | ENSMUSG | Dis3 | 2810028N( | DIS3 homc | protein-co | 5.3 | 4.3 | 5.88 | -1.58 | 1.65E-04 | 0.00633 |
| 1289 | chr9 | 65217234 | 65217384 | + | 0 | NA | intron (NM | intron (NM | 2619 | NM_17746 | 214424 | Mm.31129 | NM_17746 | ENSMUSG | Parp16 | ARTD15\|B | poly (ADP- | protein-co | 5.96 | 6.45 | 5.2 | 1.26 | 1.65E-04 | 0.00634 |
| 1294 | chr10 | 34172767 | 34172917 | + | 0 | NA | intron (NM | intron (NM | 34393 | NM_1725( | 212898 | Mm.34557 | NM_1725( | ENSMUSG | Dse | 6030499O( | dermatan | protein-co | 6.42 | 6.87 | 5.77 | 1.09 | 1.70E-04 | 0.00649 |
| 1295 | chr3 | 1.49E+08 | 1.49E+08 | + | 0 | NA | exon (NM_ | exon (NM_ | 101815 | NM_00108 | 99633 | Mm.9776 | NM_00108 | ENSMUSG | Adgrl2 | AI450192\| | adhesion G | protein-co | 6.01 | 6.53 | 5.19 | 1.34 | 1.70E-04 | 0.00649 |
| 1297 | chr2 | 1.64E+08 | 1.64E+08 | + | 0 | NA | intron (NM | MIRc\|SINE | 1383 | NM_01152 | 20971 | Mm.3815 | NM_01152 | ENSMUSG | Sdc4 | AA959608 | syndecan 4 | protein-co | 5.49 | 6.06 | 4.54 | 1.51 | 1.73E-04 | 0.00657 |
| 1298 | chr13 | 12274405 | 12274555 | + | 0 | NA | intron (NM | B3A\|SINE\| | -16367 | NM_00108 | 238505 | Mm.40335 | NM_00108 | ENSMUSG | Mtr | AI894170\| | 5-methylte | protein-co | 5.72 | 6.26 | 4.85 | 1.4 | 1.75E-04 | 0.00666 |
| 1301 | chr13 | 95685842 | 95685992 | + | 0 | NA | intron (NM | intron (NM | -11003 | NM_01017 | 14064 | Mm.44095 | NM_01017 | ENSMUSG | F2rl2 | F730031A( | coagulatio | protein-co | 5.88 | 6.39 | 5.07 | 1.32 | 1.77E-04 | 0.00669 |
| 1303 | chr7 | 1.23E+08 | 1.23E+08 | + | 0 | NA | intron (NM | intron (NM | 58357 | NM_00112 | 70497 | Mm.40331 | NM_14452 | ENSMUSG | Arhgap17 | 5730403H: | Rho GTPas | protein-co | 6.55 | 5.92 | 6.99 | -1.07 | 1.78E-04 | 0.00672 |
| 1302 | chr9 | 46215621 | 46215771 | + | 0 | NA | intron (NM | intron (NM | -12934 | NM_00969 | 11806 | Mm.26743 | NM_00969 | ENSMUSG | Apoa1 | Alp-1\|Apo | apolipopro | protein-co | 5.62 | 4.75 | 6.15 | -1.4 | 1.78E-04 | 0.00672 |
| 1306 | chr16 | 45108348 | 45108498 | + | 0 | NA | intron (NM | intron (NM | 14412 | NM_02643 | 67896 | Mm.18107 | NM_02643 | ENSMUSG | Ccdc80 | 2610001E1 | coiled-coil | protein-co | 6.58 | 7.01 | 5.96 | 1.05 | 1.79E-04 | 0.00675 |
| 1309 | chr1 | 1.71E+08 | 1.71E+08 | + | 0 | NA | intron (NM | RMER4B\|L | 21875 | NM_00108 | 226641 | Mm.37704 | NM_00108 | ENSMUSG | Atf6 | 9130025P: | activating | protein-co | 6.21 | 6.69 | 5.49 | 1.2 | 1.81E-04 | 0.00679 |
| 1310 | chr5 | 77384075 | 77384225 | + | 0 | NA | intron (NM | intron (NM | 23895 | NM_00115 | 29817 | Mm.23347 | NM_00804 | ENSMUSG | Igfbp7 | AGM\|Fstl2 | insulin-like | protein-co | 6.45 | 6.9 | 5.79 | 1.1 | 1.81E-04 | 0.0068 |
| 1314 | chr11 | 80323392 | 80323542 | + | 0 | NA | exon (NM_ | exon (NM_ | 22555 | NM_13922 | 246104 | Mm.19414 | NM_13922 | ENSMUSG | Rhbdl3 | AI847581\| | rhomboid | protein-co | 6.55 | 6.98 | 5.95 | 1.03 | 1.86E-04 | 0.00697 |
| 1319 | chr15 | 39937817 | 39937967 | + | 0 | NA | intron (NM | intron (NM | 6102 | NM_17281 | 239393 | Mm.39874 | NM_17281 | ENSMUSG | Lrp12 | AI848829\| | low densit | protein-co | 6.36 | 6.81 | 5.7 | 1.11 | 1.90E-04 | 0.00709 |
| 1320 | chr5 | 35662754 | 35662904 | + | 0 | NA | intron (NM | intron (NM | 16951 | NM_00104 | 78558 | Mm.27425 | NM_03012 | ENSMUSG | Htra3 | 2210021K2 | HtrA serin | protein-co | 5.39 | 4.44 | 5.97 | -1.53 | 1.91E-04 | 0.0071 |
| 1321 | chr19 | 46887981 | 46888131 | + | 0 | NA | non-codin | non-codin | 74295 | NM_02981 | 76952 | Mm.40965 | NM_02981 | ENSMUSG | Nt5c2 | 2010002I2 | 5'-nucleoti | protein-co | 5.92 | 6.42 | 5.14 | 1.28 | 1.91E-04 | 0.00712 |
| 1324 | chr7 | 1.43E+08 | 1.43E+08 | + | 0 | NA | intron (NM | CpG | 1718 | NM_00855 | 17173 | Mm.19641 | NM_00855 | ENSMUSG | Ascl2 | 2410083I1 | achaete-sc | protein-co | 5.5 | 4.61 | 6.05 | -1.44 | 1.93E-04 | 0.00717 |
| 1326 | chr1 | 1.93E+08 | 1.93E+08 | + | 0 | NA | intron (NM | RLTR40\|LT | 171304 | NM_14488 | 226861 | Mm.14585 | NM_14488 | ENSMUSG | Hhat | 2810432O: | hedgehog | protein-co | 4.27 | 5.02 | 2.62 | 2.4 | 1.96E-04 | 0.00727 |
| 1327 | chr19 | 6399593 | 6399743 | + | 0 | NA | promoter- | promoter- | -207 | NM_00136 | 19395 | Mm.77017 | NM_01124 | ENSMUSG | Rasgrp2 | CDC25L\|C | RAS, guany | protein-co | 6.66 | 7.08 | 6.06 | 1.02 | 1.97E-04 | 0.00728 |
| 1328 | chr6 | 1.42E+08 | 1.42E+08 | + | 0 | NA | TTS (NM_1 | TTS (NM_1 | 9282 | NM_00124 | 319552 | Mm.39649 | NM_00124 | ENSMUSG | Spx | B230216G | spexin hor | protein-co | 6.24 | 5.58 | 6.7 | -1.12 | 1.98E-04 | 0.00731 |
| 1330 | chr12 | 73715679 | 73715829 | + | 0 | NA | intron (NM | intron (NM | 130958 | NM_00131 | 18755 | Mm.34167 | NM_00885 | ENSMUSG | Prkch | Pkch | protein kir | protein-co | 5.96 | 6.45 | 5.2 | 1.25 | 1.99E-04 | 0.00733 |
| 1333 | chr16 | 32434189 | 32434339 | + | 0 | NA | intron (NM | intron (NM | 3034 | NM_00116 | 13026 | Mm.98775 | NM_00998 | ENSMUSG | Pcyt1a | CTalpha\|C | phosphate | protein-co | 6.23 | 6.69 | 5.54 | 1.15 | 2.07E-04 | 0.0076 |
| 1335 | chr5 | 66550547 | 66550697 | + | 0 | NA | intron (NM | intron (NM | 68195 | NM_0012( | 11787 | Mm.5159 | NM_00968 | ENSMUSG | Apbb2 | 2310007D( | amyloid be | protein-co | 4.97 | 3.84 | 5.59 | -1.75 | 2.08E-04 | 0.00761 |
| 1334 | chr6 | 1.34E+08 | 1.34E+08 | + | 0 | NA | intron (NM | intron (NM | -178350 | NM_02577 | 66813 | Mm.47447 | NM_02577 | ENSMUSG | Bcl2l14 | 4930452K2 | BCL2-like 1 | protein-co | 6.71 | 6.1 | 7.14 | -1.04 | 2.08E-04 | 0.00761 |
| 1336 | chr5 | 35819724 | 35819874 | + | 0 | NA | intron (NM | intron (NM | 61919 | NM_00117 | 231148 | Mm.25444 | NM_17767 | ENSMUSG | Ablim2 | AI606905\| | actin-bindi | protein-co | 4.89 | 5.54 | 3.71 | 1.83 | 2.11E-04 | 0.0077 |
| 1337 | chr18 | 82485414 | 82485564 | + | 0 | NA | intron (NM | intron (NM | 10366 | NM_00102 | 17196 | Mm.25206 | NM_01077 | ENSMUSG | Mbp | C76307\|H | myelin bas | protein-co | 5.37 | 5.94 | 4.4 | 1.55 | 2.12E-04 | 0.00773 |
| 1339 | chr12 | 99381043 | 99381193 | + | 0 | NA | intron (NM | Lx4A\|LINE | 68956 | NM_18318 | 71375 | Mm.34197 | NM_18318 | ENSMUSG | Foxn3 | 5430426H: | forkhead b | protein-co | 5.66 | 6.2 | 4.77 | 1.43 | 2.13E-04 | 0.00776 |
| 1341 | chr17 | 8342810 | 8342960 | + | 0 | NA | non-codin | non-codin | 2146 | NM_17877 | 320111 | Mm.44251 | NM_17877 | ENSMUSG | Prr18 | 9630019K: | proline ricl | protein-co | 5.31 | 5.9 | 4.32 | 1.58 | 2.14E-04 | 0.0078 |
| 1344 | chr6 | 88769797 | 88769947 | + | 0 | NA | intron (NM | intron (NM | 45168 | NM_00116 | 23945 | Mm.27219 | NM_01184 | ENSMUSG | Mgll | AA589436 | monoglyce | protein-co | 6.58 | 7.01 | 5.97 | 1.03 | 2.23E-04 | 0.00806 |
| 1345 | chr16 | 89972804 | 89972954 | + | 0 | NA | intron (NM | MIR\|SINE\| | 1820 | NM_00114 | 21844 | Mm.3109( | NM_00938 | ENSMUSG | Tiam1 | AI847750\| | T cell lymp | protein-co | 6.45 | 6.9 | 5.79 | 1.1 | 2.23E-04 | 0.00808 |
| 1347 | chr9 | 1.2E+08 | 1.2E+08 | + | 0 | NA | intron (NM | intron (NM | 1055 | NM_02873 | 74052 | Mm.18957 | NM_02873 | ENSMUSG | Ttc21a | 4921538N | tetratricop | protein-co | 5.58 | 6.13 | 4.66 | 1.47 | 2.25E-04 | 0.00812 |
| 1350 | chr11 | 1.17E+08 | 1.17E+08 | + | 0 | NA | intron (NM | intron (NM | 17312 | NM_00111 | 53860 | Mm.3845( | NM_01738 | ENSMUSG | 9-Sep | MSF1\|Msf | septin 9 | protein-co | 6.54 | 6.96 | 5.95 | 1.02 | 2.28E-04 | 0.0082 |
| 1351 | chr18 | 65051874 | 65052024 | + | 0 | NA | intron (NM | intron (NM | 28422 | NM_03188 | 83814 | Mm.98668 | NM_03188 | ENSMUSG | Nedd4l | 1300012C( | neural pre | protein-co | 5.12 | 4.06 | 5.73 | -1.67 | 2.28E-04 | 0.0082 |
| 1355 | chr1 | 51513141 | 51513291 | + | 0 | NA | intron (NR | intron (NR | -34817 | NM_02869 | 109019 | Mm.19629 | NM_02869 | ENSMUSG | Nabp1 | 4930434H( | nucleic aci | protein-co | 5.15 | 5.76 | 4.09 | 1.67 | 2.29E-04 | 0.00822 |
| 1354 | chr16 | 36821626 | 36821776 | + | 0 | NA | intron (NM | intron (NM | 6558 | NM_00111 | 106389 | Mm.43207 | NM_13411 | ENSMUSG | Eaf2 | AW048865 | ELL associa | protein-co | 5.88 | 6.38 | 5.11 | 1.27 | 2.29E-04 | 0.00822 |
| 1358 | chr3 | 1.09E+08 | 1.09E+08 | + | 0 | NA | intron (NM | intron (NM | 10059 | NM_17268 | 229731 | Mm.33574 | NM_17268 | ENSMUSG | Slc25a24 | 2610016M | solute carr | protein-co | 6.36 | 6.81 | 5.69 | 1.12 | 2.30E-04 | 0.00825 |

| | | | | | | | | | | | | | | | | | | | | | | | | |
|---|---|---|---|---|---|---|---|---|---|---|---|---|---|---|---|---|---|---|---|---|---|---|---|---|
| 1359 | chr7 | 49633429 | 49633579 | + | 0 | NA | exon (NM_ | exon (NM_ | 3331 | NM_0010( | 13172 | Mm.30717 | NM_0010( | ENSMUSG( | Dbx1 | AI426026\| | developing | protein-co | 5.6 | 6.13 | 4.74 | 1.39 | 2.31E-04 | 0.00825 |
| 1356 | chr2 | 1.29E+08 | 1.29E+08 | + | 0 | NA | intron (NM | MER58A\|D | 14896 | NM_0013! | 72477 | Mm.25968 | NM_0282 | ENSMUSG( | Tmem87b | 2610301K1 | transmem | protein-co | 6.02 | 5.3 | 6.5 | -1.2 | 2.30E-04 | 0.00825 |
| 1365 | chr7 | 46899476 | 46899626 | + | 0 | NA | intron (NM | L2\|LINE\|L1 | 20418 | NM_0013 | 22088 | Mm.24133 | NM_0218 | ENSMUSG( | Tsg101 | AI255943\| | tumor sus | protein-co | 6.65 | 7.07 | 6.07 | 1 | 2.36E-04 | 0.00841 |
| 1366 | chr7 | 1.13E+08 | 1.13E+08 | + | 0 | NA | intron (NM | intron (NM | 115366 | NM_0093 | 21676 | Mm.24685 | NM_0093 | ENSMUSG( | Tead1 | 2610024B( | TEA domai | protein-co | 4.92 | 5.56 | 3.74 | 1.82 | 2.36E-04 | 0.00841 |
| 1368 | chr8 | 71535794 | 71535944 | + | 0 | NA | intron (NM | intron (NM | 1603 | NM_1980 | 69550 | Mm.26032 | NM_1980 | ENSMUSG( | Bst2 | 2310015I1 | bone marr | protein-co | 6.06 | 6.54 | 5.33 | 1.21 | 2.37E-04 | 0.00843 |
| 1370 | chr13 | 41381644 | 41381794 | + | 0 | NA | intron (NM | intron (NM | -22718 | NM_0174 | 18003 | Mm.28898 | NM_0174 | ENSMUSG( | Nedd9 | Cas-L\|CasL | neural pre | protein-co | 6.35 | 6.8 | 5.69 | 1.11 | 2.40E-04 | 0.00849 |
| 1372 | chr17 | 34196622 | 34196772 | + | 0 | NA | exon (NM_ | exon (NM_ | -1498 | NM_0107 | 16913 | Mm.18019 | NM_0107 | ENSMUSG( | Psmb8 | Lmp-7\|Lm | proteasom | protein-co | 6.52 | 6.94 | 5.91 | 1.03 | 2.41E-04 | 0.00853 |
| 1373 | chr14 | 1.21E+08 | 1.21E+08 | + | 0 | NA | intron (NM | intron (NM | 6015 | NM_1454 | 223255 | Mm.39075 | NM_1454 | ENSMUSG( | Stk24 | 1810013H( | serine/thr | protein-co | 6.32 | 6.77 | 5.67 | 1.1 | 2.42E-04 | 0.00857 |
| 1376 | chr15 | 99233877 | 99234027 | + | 0 | NA | exon (NM_ | exon (NM_ | 9360 | NM_0106( | 16512 | Mm.37479 | NM_0106( | ENSMUSG( | Kcnh3 | AU019351 | potassium | protein-co | 4.46 | 5.17 | 2.98 | 2.19 | 2.46E-04 | 0.00866 |
| 1377 | chr4 | 1.41E+08 | 1.41E+08 | + | 0 | NA | intron (NM | intron (NM | 18831 | NM_1738 | 108911 | Mm.253 | NM_1738 | ENSMUSG( | Rcc2 | 2610510H( | regulator ( | protein-co | 6.01 | 5.28 | 6.5 | -1.22 | 2.46E-04 | 0.00866 |
| 1380 | chr14 | 21181201 | 21181351 | + | 0 | NA | intron (NM | intron (NM | 105124 | NM_0012 | 11534 | Mm.18873 | NM_1340 | ENSMUSG( | Adk | 2310026J0 | adenosine | protein-co | 4.11 | 4.89 | 2.29 | 2.6 | 2.52E-04 | 0.00885 |
| 1381 | chr13 | 24442451 | 24442601 | + | 0 | NA | intron (NM | intron (NM | 27404 | NM_0077 | 12763 | Mm.8396 | NM_0077 | ENSMUSG( | Cmah | - | cytidine m | protein-co | 5.26 | 5.85 | 4.25 | 1.59 | 2.53E-04 | 0.00887 |
| 1384 | chr11 | 44545354 | 44545504 | + | 0 | NA | intron (NM | intron (NM | 25140 | NM_0013 | 74315 | Mm.15677 | NM_0288 | ENSMUSG( | Rnf145 | 3732413I1 | ring finger | protein-co | 7.05 | 7.47 | 6.45 | 1.01 | 2.56E-04 | 0.00897 |
| 1386 | chr17 | 62672171 | 62672321 | + | 0 | NA | intron (NM | intron (NM | 209071 | NM_0101( | 13640 | Mm.7978 | NM_0101( | ENSMUSG( | Efna5 | AL-1\|AV15 | ephrin A5 | protein-co | 5.19 | 4.18 | 5.78 | -1.6 | 2.59E-04 | 0.00903 |
| 1389 | chr9 | 7859052 | 7859202 | + | 0 | NA | intron (NM | intron (NM | 14043 | NM_0074 | 11796 | Mm.2026 | NM_0074 | ENSMUSG( | Birc3 | AW10767( | baculovira | protein-co | 4.84 | 5.48 | 3.66 | 1.82 | 2.65E-04 | 0.00922 |
| 1391 | chr10 | 17920065 | 17920215 | + | 0 | NA | intron (NM | B4A\|SINE\| | 27927 | NM_0010 | 380629 | Mm.27643 | NM_0010 | ENSMUSG( | Heca | Gm869\|HI | hdc homol | protein-co | 5.34 | 5.93 | 4.34 | 1.59 | 2.70E-04 | 0.00934 |
| 1394 | chr16 | 75865659 | 75865809 | + | 0 | NA | intron (NM | intron (NM | 43532 | NM_0233 | 67742 | Mm.1314( | NM_0233 | ENSMUSG( | Samsn1 | 4930571B1 | SAM doma | protein-co | 5.62 | 6.15 | 4.79 | 1.36 | 2.72E-04 | 0.00937 |
| 1392 | chr7 | 16201267 | 16201417 | + | 0 | NA | intron (NM | intron (NM | 20651 | NM_0012 | 71723 | Mm.75259 | NM_0278 | ENSMUSG( | Dhx34 | 1200013B( | DEAH (Asp | protein-co | 5.84 | 6.35 | 5.04 | 1.32 | 2.71E-04 | 0.00937 |
| 1393 | chr9 | 63707951 | 63708101 | + | 0 | NA | intron (NM | intron (NM | 49968 | NM_0167 | 17127 | Mm.7320 | NM_0167 | ENSMUSG( | Smad3 | AU022421 | SMAD fam | protein-co | 6.26 | 6.71 | 5.61 | 1.1 | 2.71E-04 | 0.00937 |
| 1395 | chr6 | 82744575 | 82744725 | + | 0 | NA | intron (NM | intron (NM | 29804 | NM_0138 | 15277 | Mm.25584 | NM_0138 | ENSMUSG( | Hk2 | AI642394\| | hexokinase | protein-co | 6.21 | 6.66 | 5.54 | 1.12 | 2.73E-04 | 0.00941 |
| 1396 | chr14 | 1.19E+08 | 1.19E+08 | + | 0 | NA | intron (NM | intron (NM | 8497 | NM_0011 | 58187 | Mm.39075 | NM_0213 | ENSMUSG( | Cldn10 | 6720456I1 | claudin 10 | protein-co | 5.57 | 6.11 | 4.7 | 1.41 | 2.73E-04 | 0.00942 |
| 1400 | chr13 | 55529977 | 55530127 | + | 0 | NA | TTS (NM_1 | TTS (NM_1 | -1514 | NM_0137 | 27261 | Mm.3391( | NM_0137 | ENSMUSG( | Dok3 | AI450713\| | docking pr | protein-co | 5.21 | 5.81 | 4.15 | 1.67 | 2.78E-04 | 0.00955 |
| 1403 | chr1 | 64088046 | 64088196 | + | 0 | NA | intron (NM | intron (NM | 33268 | NM_0335 | 93691 | Mm.29466 | NM_0335 | ENSMUSG( | Klf7 | 9830124P( | Kruppel-lik | protein-co | 5.29 | 5.87 | 4.32 | 1.55 | 2.80E-04 | 0.00959 |
| 1404 | chr2 | 60809710 | 60809860 | + | 0 | NA | intron (NM | ORR1E\|LT | 71653 | NM_0011 | 56878 | Mm.25966 | NM_0202 | ENSMUSG( | Rbms1 | 2600014B1 | RNA bindi | protein-co | 6.51 | 6.94 | 5.91 | 1.03 | 2.81E-04 | 0.00961 |
| 1405 | chr2 | 60278767 | 60278917 | + | 0 | NA | intron (NM | ORR1E\|LT | 5642 | NM_0254 | 66205 | Mm.22728 | NM_0254 | ENSMUSG( | Cd302 | 1110055L2 | CD302 ant | protein-co | 6.48 | 6.91 | 5.87 | 1.04 | 2.85E-04 | 0.00975 |
| 1411 | chr9 | 1.15E+08 | 1.15E+08 | + | 0 | NA | intron (NM | intron (NM | 941 | NM_0260 | 67213 | Mm.28858 | NM_0260 | ENSMUSG( | Cmtm6 | 2810051A1 | CKLF-like M | protein-co | 6.22 | 6.68 | 5.56 | 1.12 | 2.90E-04 | 0.00988 |
| 1413 | chr2 | 1.03E+08 | 1.03E+08 | + | 0 | NA | intron (NM | intron (NM | 27574 | NM_0011 | 12505 | Mm.42362 | NM_0098 | ENSMUSG( | Cd44 | AU023126 | CD44 antig | protein-co | 5.9 | 6.41 | 5.12 | 1.29 | 2.91E-04 | 0.00989 |
| 1412 | chr5 | 1.1E+08 | 1.1E+08 | + | 0 | NA | intron (NM | intron (NM | 2783 | NM_0013 | 269682 | Mm.9392 | NM_0081 | ENSMUSG( | Golga3 | 5330413L0 | golgi autoa | protein-co | 5.53 | 4.62 | 6.08 | -1.46 | 2.91E-04 | 0.00989 |

| PeakID (cn | Chr | Start | End | Strand | Peak Score | Focus Rati | Annotation | Detailed A | Distance t | Nearest Pr | Entrez ID | Nearest U | Nearest Re | Nearest En | Gene Nam | Gene Alias | Gene Desc | Gene Type | Conc | Conc_grou | Conc_grou | Fold | p.value | FDR |
|---|---|---|---|---|---|---|---|---|---|---|---|---|---|---|---|---|---|---|---|---|---|---|---|---|
| 1 | chr15 | 73118034 | 73118184 | + | 0 | NA | intron (NM | intron (NM | 27697 | NM_05306 | 93696 | Mm.23095 | NM_05306 | ENSMUSG | Chrac1 | 2410152E | chromatin | protein-co | 7.94 | 8.78 | 5.69 | 3.09 | 2.16E-42 | 1.31E-37 |
| 2 | chr14 | 16173388 | 16173538 | + | 0 | NA | Intergenic | Intergenic | 69868 | NM_00135 | 71147 | Mm.19796 | NM_02769 | ENSMUSG | Oxsm | 4933425A | 3-oxoacyl- | protein-co | 6.89 | 7.83 | 3.23 | 4.6 | 1.73E-29 | 5.25E-25 |
| 3 | chr7 | 67500516 | 67500666 | + | 0 | NA | Intergenic | Intergenic | -127733 | NM_00103 | 17258 | Mm.13278 | NM_00103 | ENSMUSG | Mef2a | A430079H | myocyte e | protein-co | 7 | 7.9 | 4.06 | 3.84 | 1.50E-28 | 3.03E-24 |
| 4 | chr4 | 40875976 | 40876126 | + | 0 | NA | Intergenic | Intergenic | -21514 | NM_0223( | 14595 | Mm.15622 | NM_0223( | ENSMUSG | B4galt1 | B-1,4-GalT | UDP-Gal:b | protein-co | 7.49 | 8.29 | 5.56 | 2.73 | 8.78E-26 | 1.33E-21 |
| 6 | chr13 | 64410702 | 64410852 | + | 0 | NA | Intergenic | Intergenic | -21776 | NM_05318 | 105278 | Mm.74982 | NM_05318 | ENSMUSG | Cdk20 | 4932702G | cyclin-dep | protein-co | 7.43 | 8.2 | 5.67 | 2.53 | 1.89E-23 | 1.91E-19 |
| 7 | chr4 | 1.36E+08 | 1.36E+08 | + | 0 | NA | Intergenic | Intergenic | -28503 | NM_02591 | 67025 | Mm.27685 | NM_02591 | ENSMUSG | Rpl11 | 2010203J1 | ribosomal | protein-co | 6.91 | 7.75 | 4.65 | 3.1 | 5.57E-23 | 4.81E-19 |
| 9 | chr1 | 95268203 | 95268353 | + | 0 | NA | Intergenic | Intergenic | -45350 | NM_02632 | 67698 | Mm.27270 | NM_02632 | ENSMUSG | Fam174a | 2310044D | family witl | protein-co | 8.12 | 8.83 | 6.67 | 2.16 | 1.25E-22 | 8.44E-19 |
| 11 | chr12 | 34990447 | 34990597 | + | 0 | NA | Intergenic | RLTR19\|LT | 5761 | NM_02929 | 75456 | Mm.79179 | NM_02929 | ENSMUSG | Prps1l1 | 1700011K | phosphoril | protein-co | 7.12 | 7.93 | 5.09 | 2.84 | 1.70E-22 | 9.35E-19 |
| 12 | chr7 | 88206549 | 88206699 | + | 0 | NA | Intergenic | Intergenic | -71461 | NM_00131 | 13032 | Mm.89295 | NM_00998 | ENSMUSG | Ctsc | AI047818\| | cathepsin | protein-co | 8.21 | 8.86 | 6.99 | 1.87 | 1.94E-22 | 9.80E-19 |
| 13 | chr11 | 1.18E+08 | 1.18E+08 | + | 0 | NA | intron (NM | intron (NM | 1403 | NM_02791 | 71776 | Mm.3200 | NM_02791 | ENSMUSG | Tha1 | 1300017K | threonine | protein-co | 7.3 | 8.06 | 5.56 | 2.51 | 3.60E-22 | 1.68E-18 |
| 14 | chr5 | 65929293 | 65929443 | + | 0 | NA | Intergenic | Intergenic | -37756 | NM_00108 | 231252 | Mm.14001 | NM_00108 | ENSMUSG | Chrna9 | 2410015I0 | cholinergic | protein-co | 6.92 | 7.76 | 4.68 | 3.08 | 2.71E-21 | 1.17E-17 |
| 16 | chr17 | 57194239 | 57194389 | + | 0 | NA | promoter- | promoter- | -133 | NM_01941 | 50930 | Mm.48336 | NM_01941 | ENSMUSG | Tnfsf14 | HVEM-L\|H | tumor nec | protein-co | 7.76 | 8.45 | 6.38 | 2.06 | 2.33E-20 | 8.83E-17 |
| 17 | chr4 | 1.36E+08 | 1.36E+08 | + | 0 | NA | Intergenic | Intergenic | -35744 | NM_02591 | 67025 | Mm.27685 | NM_02591 | ENSMUSG | Rpl11 | 2010203J1 | ribosomal | protein-co | 6.86 | 7.66 | 4.88 | 2.78 | 7.42E-20 | 2.64E-16 |
| 18 | chr9 | 1.24E+08 | 1.24E+08 | + | 0 | NA | Intergenic | Intergenic | -12658 | NM_00116 | 12769 | Mm.44060 | NM_00991 | ENSMUSG | Ccr9 | A130091K | chemokine | protein-co | 6.93 | 7.73 | 4.99 | 2.74 | 3.39E-19 | 1.14E-15 |
| 19 | chr3 | 9887055 | 9887205 | + | 0 | NA | Intergenic | Intergenic | -53451 | NM_00119 | 94212 | Mm.23897 | NM_05318 | ENSMUSG | Pag1 | Cbp\|F730 | phosphopı | protein-co | 7.9 | 8.56 | 6.64 | 1.91 | 3.99E-19 | 1.27E-15 |
| 21 | chr16 | 30184964 | 30185114 | + | 0 | NA | Intergenic | Intergenic | 82493 | NM_0279( | 71756 | Mm.29061 | NM_0279( | ENSMUSG | Cpn2 | 1300018K | carboxype | protein-co | 7.23 | 7.96 | 5.65 | 2.32 | 2.51E-18 | 7.24E-15 |
| 22 | chr2 | 1.04E+08 | 1.04E+08 | + | 0 | NA | intron (NM | intron (NM | 78714 | NM_15312 | 98956 | Mm.28800 | NM_15312 | ENSMUSG | Nat10 | AI429152 | N-acetyltra | protein-co | 8.54 | 9.09 | 7.65 | 1.44 | 3.31E-18 | 9.11E-15 |
| 23 | chr8 | 1.23E+08 | 1.23E+08 | + | 0 | NA | Intergenic | Intergenic | -3346 | NM_00982 | 12398 | Mm.19433 | NM_00982 | ENSMUSG | Cbfa2t3 | A630044F | CBFA2/RU | protein-co | 6.83 | 7.62 | 4.94 | 2.68 | 5.33E-18 | 1.40E-14 |
| 24 | chr13 | 91541065 | 91541215 | + | 0 | NA | intron (NM | intron (NM | 80043 | NM_00136 | 66970 | Mm.34309 | NM_02418 | ENSMUSG | Ssbp2 | 1500004K | single-stra | protein-co | 7.93 | 8.54 | 6.86 | 1.68 | 3.47E-17 | 8.76E-14 |
| 25 | chr19 | 3451852 | 3452002 | + | 0 | NA | Intergenic | Intergenic | -37383 | NM_00132 | 14419 | Mm.4655 | NM_01025 | ENSMUSG | Gal | Galn | galanin an | protein-co | 6.64 | 7.46 | 4.49 | 2.98 | 4.78E-17 | 1.16E-13 |
| 26 | chr17 | 79954888 | 79955038 | + | 0 | NA | Intergenic | Intergenic | -58840 | NM_00135 | 56298 | Mm.17540 | NM_01971 | ENSMUSG | Atl2 | 2010110I2 | atlastin GT | protein-co | 6.37 | 7.2 | 4.14 | 3.07 | 5.69E-17 | 1.33E-13 |
| 28 | chr4 | 1.5E+08 | 1.5E+08 | + | 0 | NA | Intergenic | Intergenic | -82348 | NM_02746 | 70556 | Mm.41877 | NM_02746 | ENSMUSG | Slc25a33 | 5730438N | solute carr | protein-co | 7.97 | 8.56 | 6.95 | 1.61 | 2.24E-16 | 4.85E-13 |
| 29 | chr8 | 1.2E+08 | 1.2E+08 | + | 0 | NA | Intergenic | Intergenic | 32986 | NM_13396 | 102193 | Mm.24007 | NM_13396 | ENSMUSG | Zdhhc7 | AL024087\| | zinc finger | protein-co | 7.05 | 7.78 | 5.54 | 2.24 | 2.78E-16 | 5.80E-13 |
| 31 | chr5 | 1.49E+08 | 1.49E+08 | + | 0 | NA | Intergenic | Intergenic | -43523 | NR_15159 | 70717 | Mm.25213 | NM_02751 | ENSMUSG | Medag | 6330406I1 | mesenteri | protein-co | 6.26 | 7.12 | 3.91 | 3.2 | 3.95E-16 | 7.71E-13 |
| 32 | chr3 | 1.33E+08 | 1.33E+08 | + | 0 | NA | intron (NM | intron (NM | 29451 | NM_14614 | 74776 | Mm.21030 | NM_14614 | ENSMUSG | Ppa2 | 1110013G | pyrophosp | protein-co | 6.47 | 7.3 | 4.28 | 3.02 | 4.16E-16 | 7.87E-13 |
| 33 | chr6 | 1.16E+08 | 1.16E+08 | + | 0 | NA | intron (NM | intron (NM | 5313 | NM_02637 | 67784 | Mm.3085 | NM_02637 | ENSMUSG | Plxnd1 | 6230425C | plexin D1 | protein-co | 7.55 | 8.2 | 6.34 | 1.85 | 4.40E-16 | 8.08E-13 |
| 34 | chr5 | 1.14E+08 | 1.14E+08 | + | 0 | NA | Intergenic | Intergenic | -2206 | NM_00915 | 20345 | Mm.33259 | NM_00915 | ENSMUSG | Selplg | CD162\|Ps | selectin, pl | protein-co | 6.54 | 7.34 | 4.62 | 2.72 | 5.99E-16 | 1.07E-12 |
| 35 | chr7 | 98563097 | 98563247 | + | 0 | NA | Intergenic | Intergenic | 68950 | NM_00111 | 434215 | Mm.32305 | NM_00111 | ENSMUSG | Lrrc32 | AI426318\| | leucine ric | protein-co | 7.23 | 7.94 | 5.77 | 2.16 | 7.78E-16 | 1.35E-12 |
| 36 | chr17 | 47531561 | 47531711 | + | 0 | NA | intron (NM | intron (NM | 26585 | NM_00108 | 12445 | Mm.27291 | NM_00763 | ENSMUSG | Ccnd3 | 9230106B | cyclin D3 | protein-co | 7.05 | 7.77 | 5.59 | 2.18 | 8.07E-16 | 1.36E-12 |
| 38 | chr16 | 10788471 | 10788621 | + | 0 | NA | exon (NM_ | exon (NM_ | 109 | NM_01369 | 21959 | Mm.20679 | NM_01369 | ENSMUSG | Tnp2 | TP2\|Tp-2 | transition | protein-co | 7.27 | 7.97 | 5.86 | 2.11 | 1.88E-15 | 3.00E-12 |
| 39 | chr18 | 39866824 | 39866974 | + | 0 | NA | Intergenic | Intergenic | 93402 | NM_01103 | 18459 | Mm.9493 | NM_01103 | ENSMUSG | Pabpc2 | Pabp\|Pab | poly(A) bir | protein-co | 6.8 | 7.55 | 5.15 | 2.4 | 3.42E-15 | 5.30E-12 |
| 42 | chr2 | 1.66E+08 | 1.66E+08 | + | 0 | NA | intron (NM | intron (NM | 64762 | NM_00125 | 228880 | Mm.22759 | NM_02723 | ENSMUSG | Zmynd8 | 1110013E | zinc finger | protein-co | 7.25 | 7.93 | 5.94 | 1.99 | 3.87E-15 | 5.58E-12 |
| 41 | chr11 | 57958379 | 57958529 | + | 0 | NA | Intergenic | Intergenic | -50610 | NM_02845 | 73158 | Mm.24884 | NM_02845 | ENSMUSG | Larp1 | 1810024J1 | La ribonuc | protein-co | 6.62 | 7.41 | 4.75 | 2.65 | 3.86E-15 | 5.58E-12 |
| 43 | chr16 | 30227430 | 30227580 | + | 0 | NA | Intergenic | Intergenic | 40027 | NM_0279( | 71756 | Mm.29061 | NM_0279( | ENSMUSG | Cpn2 | 1300018K | carboxype | protein-co | 7.03 | 7.74 | 5.55 | 2.19 | 4.79E-15 | 6.74E-12 |
| 44 | chr7 | 1.4E+08 | 1.4E+08 | + | 0 | NA | Intergenic | Intergenic | 9708 | NM_00101 | 258055 | Mm.37733 | NM_00101 | ENSMUSG | Olfr524 | MOR103-1 | olfactory r | protein-co | 6.68 | 7.44 | 4.94 | 2.5 | 5.05E-15 | 6.95E-12 |
| 48 | chr1 | 1.34E+08 | 1.34E+08 | + | 0 | NA | Intergenic | Intergenic | 60264 | NM_00922 | 20643 | Mm.24911 | NM_00922 | ENSMUSG | Snrpe | AL022645\| | small nucl | protein-co | 6.66 | 7.42 | 4.96 | 2.45 | 1.04E-14 | 1.31E-11 |
| 49 | chr11 | 21180761 | 21180911 | + | 0 | NA | Intergenic | Intergenic | -58196 | NM_00129 | 245944 | Mm.17010 | NM_13906 | ENSMUSG | Vps54 | Hcc8\|Vps5 | VPS54 GAI | protein-co | 6.42 | 7.22 | 4.45 | 2.77 | 1.13E-14 | 1.39E-11 |
| 50 | chr5 | 1.12E+08 | 1.12E+08 | + | 0 | NA | Intergenic | Intergenic | -5471 | NM_00941 | 22022 | Mm.25063 | NM_00941 | ENSMUSG | Tpst2 | AI448750\| | protein-tyı | protein-co | 7.66 | 8.27 | 6.58 | 1.69 | 2.07E-14 | 2.51E-11 |
| 55 | chr4 | 1.36E+08 | 1.36E+08 | + | 0 | NA | intron (NM | intron (NM | -2827 | NM_00130 | 242705 | Mm.30793 | NM_17773 | ENSMUSG | E2f2 | E2F-2 | E2F transc | protein-co | 7.89 | 8.49 | 6.85 | 1.63 | 3.38E-14 | 3.72E-11 |
| 56 | chr1 | 1.64E+08 | 1.64E+08 | + | 0 | NA | intron (NM | intron (NM | 31908 | NM_00972 | 11931 | Mm.4550 | NM_00972 | ENSMUSG | Atp1b1 | Atp4b\|Atp | ATPase, N | protein-co | 5.66 | 6.6 | 1.95 | 4.65 | 5.28E-14 | 5.70E-11 |
| 57 | chr2 | 1.21E+08 | 1.21E+08 | + | 0 | NA | intron (NM | intron (NM | 23311 | NM_00110 | 12335 | Mm.45802 | NM_00760 | ENSMUSG | Capn3 | AI323605\| | calpain 3 | protein-co | 6.98 | 7.67 | 5.6 | 2.07 | 6.60E-14 | 6.91E-11 |
| 59 | chr2 | 27122121 | 27122271 | + | 0 | NA | intron (NM | intron (NM | 20281 | NM_17542 | 109349 | Mm.31408 | NM_17542 | ENSMUSG | Fam163b | C630035N | family witl | protein-co | 5.65 | 6.59 | 2.05 | 4.54 | 8.42E-14 | 8.64E-11 |
| 60 | chr8 | 83748792 | 83748942 | + | 0 | NA | Intergenic | Intergenic | -7556 | NM_00116 | 26364 | Mm.33464 | NM_01192 | ENSMUSG | Adgre5 | AA409984 | adhesion ( | protein-co | 6.78 | 7.51 | 5.2 | 2.31 | 1.01E-13 | 1.02E-10 |
| 63 | chr8 | 1.23E+08 | 1.23E+08 | + | 0 | NA | Intergenic | Intergenic | -2617 | NM_14493 | 257633 | Mm.33487 | NM_14493 | ENSMUSG | Acsf3 | BB101783 | acyl-CoA s | protein-co | 5.6 | 6.55 | 1.77 | 4.78 | 1.20E-13 | 1.15E-10 |
| 65 | chr8 | 1.22E+08 | 1.22E+08 | + | 0 | NA | Intergenic | RMER4A\|L | -3830 | NM_01391 | 30927 | Mm.10367 | NM_01391 | ENSMUSG | Snai3 | AI643946\| | snail famil | protein-co | 6.36 | 7.13 | 4.56 | 2.57 | 1.93E-13 | 1.80E-10 |
| 68 | chr8 | 1.17E+08 | 1.17E+08 | + | 0 | NA | intron (NM | intron (NM | 94929 | NM_02929 | 75465 | Mm.23114 | NM_02929 | ENSMUSG | Dynlrb2 | 1700009A | dynein ligł | protein-co | 6.7 | 7.43 | 5.15 | 2.28 | 3.24E-13 | 2.88E-10 |
| 71 | chr19 | 10064384 | 10064534 | + | 0 | NA | intron (NM | intron (NM | 22911 | NM_02189 | 60527 | Mm.25387 | NM_02189 | ENSMUSG | Fads3 | AI464531 | fatty acid ( | protein-co | 6.3 | 7.08 | 4.51 | 2.57 | 5.50E-13 | 4.69E-10 |
| 72 | chr6 | 1.25E+08 | 1.25E+08 | + | 0 | NA | TTS (NM_( | TTS (NM_( | 6790 | NM_00108 | 22317 | Mm.32321 | NM_00949 | ENSMUSG | Vamp1 | Syb-1\|Syb | vesicle-ass | protein-co | 6.14 | 6.93 | 4.22 | 2.71 | 7.69E-13 | 6.39E-10 |
| 74 | chr8 | 1.23E+08 | 1.23E+08 | + | 0 | NA | Intergenic | Intergenic | -1723 | NM_00982 | 12398 | Mm.19433 | NM_00982 | ENSMUSG | Cbfa2t3 | A630044F | CBFA2/RU | protein-co | 6.37 | 7.16 | 4.52 | 2.64 | 8.11E-13 | 6.63E-10 |
| 75 | chr4 | 55343335 | 55343485 | + | 0 | NA | Intergenic | RLTR19\|LT | -6632 | NM_00901 | 19359 | Mm.19684 | NM_00901 | ENSMUSG | Rad23b | 0610007D | RAD23 hor | protein-co | 6.42 | 7.19 | 4.66 | 2.53 | 8.78E-13 | 7.09E-10 |
| 77 | chr16 | 23279948 | 23280098 | + | 0 | NA | intron (NM | intron (NM | -10447 | NM_00125 | 20440 | Mm.14902 | NM_14593 | ENSMUSG | St6gal1 | AW742324 | beta galac | protein-co | 6.28 | 7.06 | 4.48 | 2.57 | 1.00E-12 | 7.89E-10 |
| 78 | chr8 | 1.23E+08 | 1.23E+08 | + | 0 | NA | Intergenic | Intergenic | -13577 | NM_00982 | 12398 | Mm.19433 | NM_00982 | ENSMUSG | Cbfa2t3 | A630044F | CBFA2/RU | protein-co | 5.48 | 6.44 | 1.49 | 4.94 | 1.23E-12 | 9.57E-10 |
| 80 | chr8 | 95156092 | 95156242 | + | 0 | NA | Intergenic | Intergenic | -13627 | NM_01063 | 16582 | Mm.37895 | NM_01063 | ENSMUSG | Kifc3 | AI325457\| | kinesin far | protein-co | 5.97 | 6.82 | 3.59 | 3.22 | 1.48E-12 | 1.12E-09 |
| 81 | chr3 | 94388203 | 94388353 | + | 0 | NA | intron (NM | intron (NM | 10846 | NR_12165 | 19885 | Mm.4372 | NM_01128 | ENSMUSG | Rorc | Nr1f3\|ROI | RAR-relate | protein-co | 6.65 | 7.36 | 5.2 | 2.16 | 1.55E-12 | 1.16E-09 |
| 84 | chr5 | 36979807 | 36979957 | + | 0 | NA | intron (NM | intron (NM | 9100 | NM_01171 | 22393 | Mm.20916 | NM_01171 | ENSMUSG | Wfs1 | AI481085\| | wolframin | protein-co | 6.3 | 7.07 | 4.58 | 2.48 | 1.88E-12 | 1.35E-09 |
| 85 | chr19 | 41039265 | 41039415 | + | 0 | NA | intron (NM | intron (NM | 10065 | NM_00104 | 21673 | Mm.25620 | NM_00934 | ENSMUSG | Dntt | Tdt | deoxynucl | protein-co | 6.27 | 7.06 | 4.39 | 2.67 | 2.27E-12 | 1.61E-09 |
| 87 | chr15 | 66883918 | 66884068 | + | 0 | NA | Intergenic | Intergenic | -7376 | NM_01886 | 22402 | Mm.10222 | NM_01886 | ENSMUSG | Ccn4 | AW146261 | cellular co | protein-co | 7.53 | 8.11 | 6.53 | 1.58 | 2.61E-12 | 1.81E-09 |
| 89 | chr8 | 95154810 | 95154960 | + | 0 | NA | Intergenic | Intergenic | -12345 | NM_01063 | 16582 | Mm.37895 | NM_01063 | ENSMUSG | Kifc3 | AI325457\| | kinesin far | protein-co | 6.07 | 6.87 | 4.06 | 2.82 | 3.29E-12 | 2.24E-09 |
| 91 | chr16 | 10668544 | 10668694 | + | 0 | NA | intron (NM | intron (NM | 116917 | NM_00127 | 12703 | Mm.130 | NM_00989 | ENSMUSG | Socs1 | Cish1\|Cish | suppresso | protein-co | 6.45 | 7.22 | 4.73 | 2.48 | 3.82E-12 | 2.51E-09 |
| 92 | chr13 | 64408128 | 64408278 | + | 0 | NA | Intergenic | ORR1C2\|L | -24350 | NM_05318 | 105278 | Mm.74982 | NM_05318 | ENSMUSG | Cdk20 | 4932702G | cyclin-dep | protein-co | 5.5 | 6.39 | 2.72 | 3.67 | 3.82E-12 | 2.51E-09 |
| 93 | chr18 | 6280702 | 6280852 | + | 0 | NA | Intergenic | Intergenic | -39253 | NM_00844 | 16573 | Mm.22374 | NM_00844 | ENSMUSG | Kif5b | AL022807\| | kinesin far | protein-co | 5.45 | 6.37 | 2.35 | 4.02 | 4.46E-12 | 2.91E-09 |
| 95 | chr5 | 1.4E+08 | 1.4E+08 | + | 0 | NA | Intergenic | Intergenic | -1452 | NM_02674 | 68510 | Mm.29294 | NM_02674 | ENSMUSG | Ints1 | 1110015K | integrator | protein-co | 6.08 | 6.86 | 4.25 | 2.61 | 4.78E-12 | 3.05E-09 |
| 98 | chr2 | 25035073 | 25035223 | + | 0 | NA | intron (NM | intron (NM | -19231 | NM_02027 | 56876 | Mm.42956 | NM_02027 | ENSMUSG | Nsmf | Jacob\|Nel | NMDA rec | protein-co | 5.43 | 6.38 | 1.27 | 5.12 | 5.13E-12 | 3.17E-09 |
| 102 | chr2 | 24748028 | 24748178 | + | 0 | NA | intron (NM | intron (NM | 15049 | NM_00757 | 12287 | Mm.4424 | NM_00757 | ENSMUSG | Cacna1b | AW050276 | calcium ch | protein-co | 5.91 | 6.72 | 3.94 | 2.78 | 7.25E-12 | 4.30E-09 |
| 104 | chr15 | 52753442 | 52753592 | + | 0 | NA | Intergenic | ORR1B1\|L | 41072 | NM_02721 | 69790 | Mm.19556 | NM_02721 | ENSMUSG | Med30 | 1810038N | mediator ( | protein-co | 6.1 | 6.88 | 4.33 | 2.55 | 8.62E-12 | 5.02E-09 |
| 105 | chr5 | 34976990 | 34977140 | + | 0 | NA | intron (NM | intron (NM | -22013 | NM_00116 | 71729 | Mm.19620 | NM_17340 | ENSMUSG | Rgs12 | 1200016K | regulator ( | protein-co | 5.47 | 6.37 | 2.48 | 3.89 | 8.87E-12 | 5.12E-09 |
| 109 | chr3 | 1.53E+08 | 1.53E+08 | + | 0 | NA | intron (NM | ORR1E\|LT | 55794 | NM_01202 | 26938 | Mm.40915 | NM_01202 | ENSMUSG | St6galnac5 | AI851940\| | ST6 (alpha | protein-co | 7.09 | 7.72 | 5.93 | 1.79 | 1.02E-11 | 5.67E-09 |
| 111 | chr13 | 51754170 | 51754320 | + | 0 | NA | intron (NM | intron (NM | 39502 | NM_00128 | 20354 | Mm.33903 | NM_01366 | ENSMUSG | Sema4d | CD100\|Se | sema dom | protein-co | 6.99 | 7.63 | 5.8 | 1.83 | 1.23E-11 | 6.73E-09 |
| 119 | chr13 | 13557104 | 13557254 | + | 0 | NA | Intergenic | Intergenic | -33230 | NM_01074 | 17101 | Mm.34233 | NM_01074 | ENSMUSG | Lyst | D13Sfk13\| | lysosomal | protein-co | 6.24 | 7 | 4.53 | 2.47 | 2.08E-11 | 1.06E-08 |
| 121 | chr11 | 1.02E+08 | 1.02E+08 | + | 0 | NA | intron (NM | intron (NM | 2112 | NM_00817 | 14824 | Mm.1568 | NM_00817 | ENSMUSG | Grn | GP88\|PCD | granulin | protein-co | 6.67 | 7.37 | 5.24 | 2.13 | 2.15E-11 | 1.07E-08 |
| 122 | chr2 | 1.28E+08 | 1.28E+08 | + | 0 | NA | intron (NM | intron (NM | 73062 | NM_02876 | 74121 | Mm.24483 | NM_02876 | ENSMUSG | Acoxl | 1200014P | acyl-Coenz | protein-co | 5.57 | 6.42 | 3.25 | 3.17 | 2.26E-11 | 1.12E-08 |
| 124 | chr17 | 47542178 | 47542328 | + | 0 | NA | intron (NM | intron (NM | 37202 | NM_00108 | 12445 | Mm.27291 | NM_00763 | ENSMUSG | Ccnd3 | 9230106B | cyclin D3 | protein-co | 6.61 | 7.31 | 5.21 | 2.1 | 2.41E-11 | 1.17E-08 |
| 127 | chr15 | 80192409 | 80192559 | + | 0 | NA | intron (NM | intron (NM | 18763 | NM_01079 | 17309 | Mm.29969 | NM_01079 | ENSMUSG | Mgat3 | 1110038J1 | mannoside | protein-co | 6.48 | 7.2 | 4.98 | 2.22 | 2.68E-11 | 1.27E-08 |
| 134 | chr17 | 79982715 | 79982865 | + | 0 | NA | Intergenic | Intergenic | 79478 | NR_15211 | 72692 | Mm.64579 | NM_14480 | ENSMUSG | Hnrnpll | 2510028H | heterogen | protein-co | 5.32 | 6.23 | 2.28 | 3.94 | 3.27E-11 | 1.48E-08 |
| 137 | chr7 | 1.26E+08 | 1.26E+08 | + | 0 | NA | intron (NM | intron (NM | 27081 | NM_02188 | 60504 | Mm.15564 | NM_02188 | ENSMUSG | Il21r | NILR | interleukir | protein-co | 7.59 | 8.15 | 6.63 | 1.52 | 3.53E-11 | 1.56E-08 |
| 138 | chr5 | 65910465 | 65910615 | + | 0 | NA | Intergenic | Intergenic | 46956 | NM_00108 | 74734 | Mm.20323 | NM_00108 | ENSMUSG | Rhoh | 5830400A | ras homolo | protein-co | 5.69 | 6.55 | 3.27 | 3.28 | 3.56E-11 | 1.56E-08 |
| 147 | chr19 | 32138405 | 32138555 | + | 0 | NA | intron (NM | intron (NM | -35340 | NM_01883 | 54447 | Mm.49122 | NM_01883 | ENSMUSG | Asah2 | AI585898 | N-acylsphi | protein-co | 7.64 | 8.18 | 6.76 | 1.43 | 5.81E-11 | 2.37E-08 |
| 146 | chr18 | 20957178 | 20957328 | + | 0 | NA | intron (NM | intron (NM | -4111 | NM_00136 | 67664 | Mm.45980 | NM_02630 | ENSMUSG | Rnf125 | 4930553F | ring finger | protein-co | 6.24 | 6.97 | 4.65 | 2.32 | 5.80E-11 | 2.37E-08 |
| 151 | chr1 | 1.93E+08 | 1.93E+08 | + | 0 | NA | intron (NM | intron (NM | 31769 | NM_14481 | 215303 | Mm.40329 | NM_14481 | ENSMUSG | Camk1g | CLICK-III\|C | calcium/ca | protein-co | 7.37 | 7.95 | 6.41 | 1.54 | 7.22E-11 | 2.88E-08 |
| 153 | chr7 | 63999149 | 63999299 | + | 0 | NA | Intergenic | Intergenic | -60309 | NM_02136 | 50794 | Mm.24047 | NM_02136 | ENSMUSG | Klf13 | 0610043C | Kruppel-lik | protein-co | 6.56 | 7.25 | 5.16 | 2.09 | 7.52E-11 | 2.96E-08 |
| 157 | chr15 | 97750824 | 97750974 | + | 0 | NA | intron (NM | intron (NM | 16767 | NM_00135 | 223864 | Mm.24028 | NM_14485 | ENSMUSG | Rapgef3 | 2310016P | Rap guanir | protein-co | 7.16 | 7.76 | 6.1 | 1.66 | 7.91E-11 | 3.03E-08 |
| 160 | chr9 | 57769876 | 57770026 | + | 0 | NA | Intergenic | Intergenic | -4091 | NM_00771 | 102414 | Mm.25720 | NM_00771 | ENSMUSG | Clk3 | AI256811 | CDC-like ki | protein-co | 6.03 | 6.82 | 4.19 | 2.63 | 8.89E-11 | 3.33E-08 |
| 175 | chr1 | 1.38E+08 | 1.38E+08 | + | 0 | NA | Intergenic | Intergenic | 29729 | NM_17739 | 338375 | Mm.11442 | NM_17739 | ENSMUSG | Atp6v1g3 | - | ATPase, H- | protein-co | 7.21 | 7.8 | 6.19 | 1.6 | 1.38E-10 | 4.76E-08 |
| 176 | chr13 | 51754791 | 51754941 | + | 0 | NA | intron (NM | intron (NM | 38881 | NM_00128 | 20354 | Mm.33903 | NM_01366 | ENSMUSG | Sema4d | CD100\|Se | sema dom | protein-co | 7.18 | 7.77 | 6.16 | 1.61 | 1.43E-10 | 4.90E-08 |
| 177 | chr15 | 67103470 | 67103620 | + | 0 | NA | 3' UTR (NM | 3' UTR (NM | 73337 | NM_00917 | 20442 | Mm.24833 | NM_00917 | ENSMUSG | St3gal1 | 5330418N | ST3 beta-g | protein-co | 6.24 | 6.98 | 4.67 | 2.31 | 1.48E-10 | 5.03E-08 |
| 184 | chr16 | 10806256 | 10806406 | + | 0 | NA | Intergenic | Intergenic | -9508 | NM_01363 | 19118 | Mm.42733 | NM_01363 | ENSMUSG | Prm1 | Prm-1 | protamine | protein-co | 6.29 | 7.01 | 4.77 | 2.24 | 1.76E-10 | 5.77E-08 |
| 187 | chr9 | 44649000 | 44649150 | + | 0 | NA | Intergenic | Intergenic | -24158 | NM_02148 | 58866 | Mm.45380 | NM_02148 | ENSMUSG | Treh | 2210412M | trehalase ( | protein-co | 5.74 | 6.54 | 3.77 | 2.77 | 1.97E-10 | 6.35E-08 |
| 194 | chr7 | 49601687 | 49601837 | + | 0 | NA | intron (NM | intron (NM | 35073 | NM_00100 | 13172 | Mm.30717 | NM_00100 | ENSMUSG | Dbx1 | AI426026\| | developing | protein-co | 7.26 | 7.85 | 6.24 | 1.61 | 2.20E-10 | 6.85E-08 |
| 195 | chr15 | 66789167 | 66789317 | + | 0 | NA | intron (NM | intron (NM | 42587 | NM_00102 | 20491 | Mm.7601 | NM_00919 | ENSMUSG | Sla | Slap\|Slap- | src-like ad | protein-co | 7.05 | 7.67 | 5.94 | 1.73 | 2.39E-10 | 7.37E-08 |
| 196 | chr7 | 1.13E+08 | 1.13E+08 | + | 0 | NA | intron (NM | intron (NM | 34463 | NM_00136 | 11865 | Mm.33970 | NM_00748 | ENSMUSG | Arntl | Arnt3\|BM | aryl hydro | protein-co | 5.56 | 6.39 | 3.4 | 2.99 | 2.45E-10 | 7.53E-08 |
| 197 | chr5 | 1.49E+08 | 1.49E+08 | + | 0 | NA | Intergenic | MER74A\|L | -37481 | NR_15159 | 70717 | Mm.25213 | NM_02751 | ENSMUSG | Medag | 6330406I1 | mesenteri | protein-co | 6.87 | 7.5 | 5.72 | 1.77 | 2.50E-10 | 7.64E-08 |
| 200 | chr2 | 84964149 | 84964299 | + | 0 | NA | Intergenic | Intergenic | -16237 | NM_00892 | 19074 | Mm.14272 | NM_00892 | ENSMUSG | Prg2 | EMBP\|MB | proteoglyc | protein-co | 6.47 | 7.16 | 5.14 | 2.02 | 2.59E-10 | 7.79E-08 |
| 201 | chr19 | 32168208 | 32168358 | + | 0 | NA | intron (NM | RMER15\|L | 42730 | NM_00116 | 208449 | Mm.32981 | NM_14479 | ENSMUSG | Sgms1 | 9530058O | sphingomy | protein-co | 6.49 | 7.17 | 5.15 | 2.02 | 2.71E-10 | 8.11E-08 |
| 204 | chr1 | 36025079 | 36025229 | + | 0 | NA | Intergenic | Intergenic | -43246 | NM_01581 | 50785 | Mm.21356 | NM_01581 | ENSMUSG | Hs6st1 | 6Ost1 | heparan su | protein-co | 6.03 | 6.81 | 4.24 | 2.56 | 2.93E-10 | 8.65E-08 |
| 206 | chr17 | 29304467 | 29304617 | + | 0 | NA | Intergenic | Intergenic | -14340 | NM_02373 | 74116 | Mm.30950 | NM_02373 | ENSMUSG | Pi16 | 1200009H | peptidase | protein-co | 6.47 | 7.17 | 5.07 | 2.1 | 3.00E-10 | 8.79E-08 |
| 208 | chr1 | 87829806 | 87829956 | + | 0 | NA | intron (NM | (GA)n\|Sim | -23406 | NM_17764 | 227333 | Mm.27721 | NM_17764 | ENSMUSG | Dgkd | AI841987\| | diacylglyce | protein-co | 6.67 | 7.32 | 5.45 | 1.87 | 3.11E-10 | 9.01E-08 |
| 214 | chr10 | 1.21E+08 | 1.21E+08 | + | 0 | NA | intron (NM | intron (NM | 28753 | NM_02905 | 74694 | Mm.23383 | NM_02905 | ENSMUSG | Tbc1d30 | 4930505D | TBC1 dom | protein-co | 6.2 | 6.96 | 4.49 | 2.48 | 3.58E-10 | 1.01E-07 |
| 216 | chr13 | 85507017 | 85507167 | + | 0 | NA | Intergenic | Intergenic | -217606 | NM_14545 | 218397 | Mm.25965 | NM_14545 | ENSMUSG | Rasa1 | Gap\|RasG | RAS p21 pı | protein-co | 7.17 | 7.79 | 6.04 | 1.75 | 3.78E-10 | 1.05E-07 |
| 218 | chr7 | 83661029 | 83661179 | + | 0 | NA | exon (NM_ | exon (NM_ | -5609 | NM_00136 | 16170 | Mm.10137 | NM_01055 | ENSMUSG | Il16 | mKIAA404 | interleukir | protein-co | 6.46 | 7.17 | 5.02 | 2.14 | 3.90E-10 | 1.08E-07 |
| 220 | chr10 | 75978249 | 75978399 | + | 0 | NA | exon (NM_ | exon (NM_ | 23810 | NM_19863 | 333669 | Mm.31077 | NM_19863 | ENSMUSG | Gm5134 | EG333669 | predicted | protein-co | 5.64 | 6.46 | 3.61 | 2.85 | 3.93E-10 | 1.08E-07 |
| 223 | chr4 | 1.5E+08 | 1.5E+08 | + | 0 | NA | Intergenic | Intergenic | -80178 | NM_02746 | 70556 | Mm.41877 | NM_02746 | ENSMUSG | Slc25a33 | 5730438N | solute carr | protein-co | 7.41 | 7.97 | 6.5 | 1.47 | 4.06E-10 | 1.10E-07 |
| 222 | chr6 | 1.37E+08 | 1.37E+08 | + | 0 | NA | Intergenic | B1F\|SINE\| | -21888 | NM_02580 | 66857 | Mm.3311 | NM_02580 | ENSMUSG | Plbd1 | 1100001H | phospholiı | protein-co | 5.85 | 6.64 | 4 | 2.64 | 4.05E-10 | 1.10E-07 |
| 228 | chr15 | 52753784 | 52753934 | + | 0 | NA | Intergenic | Intergenic | 41414 | NM_02721 | 69790 | Mm.19556 | NM_02721 | ENSMUSG | Med30 | 1810038N | mediator ( | protein-co | 6.94 | 7.55 | 5.86 | 1.69 | 4.50E-10 | 1.19E-07 |
| 238 | chr17 | 6661214 | 6661364 | + | 0 | NA | Intergenic | Intergenic | -5350 | NM_00119 | 1E+08 | Mm.45798 | NM_00116 | ENSMUSG | Dynlt1f | 10004063 | dynein ligł | protein-co | 6.74 | 7.42 | 5.42 | 2 | 5.75E-10 | 1.46E-07 |
| 243 | chr16 | 30145641 | 30145791 | + | 0 | NA | Intergenic | Intergenic | 80359 | NM_00823 | 15205 | Mm.39085 | NM_00823 | ENSMUSG | Hes1 | Hry\|bHLH | hes family | protein-co | 6.17 | 6.93 | 4.5 | 2.43 | 6.36E-10 | 1.58E-07 |
| 245 | chr5 | 62858402 | 62858552 | + | 0 | NA | intron (NM | intron (NM | 46821 | NM_00117 | 1E+08 |  | NM_00117 | ENSMUSG | Dthd1 | Gm17384 | death dom | protein-co | 6.87 | 7.49 | 5.75 | 1.75 | 6.53E-10 | 1.61E-07 |
| 246 | chr6 | 41541895 | 41542045 | + | 0 | NA | Intergenic | Intergenic | 20194 | NM_00943 | 22072 | Mm.27692 | NM_00943 | ENSMUSG | Prss2 | TRY8\|TRY | protease, s | protein-co | 6.07 | 6.81 | 4.47 | 2.34 | 6.76E-10 | 1.66E-07 |
| 250 | chr18 | 38495858 | 38496008 | + | 0 | NA | Intergenic | Intergenic | 77026 | NM_02299 | 65113 | Mm.10249 | NM_02299 | ENSMUSG | Ndfip1 | 0610010M | Nedd4 fan | protein-co | 6.38 | 7.08 | 5 | 2.07 | 7.21E-10 | 1.74E-07 |
| 253 | chr3 | 1.27E+08 | 1.27E+08 | + | 0 | NA | intron (NM | intron (NM | 63786 | NM_00102 | 108058 | Mm.25582 | NM_02381 | ENSMUSG | Camk2d | 2810011D | calcium/ca | protein-co | 6.78 | 7.41 | 5.65 | 1.75 | 8.36E-10 | 1.99E-07 |
| 262 | chr1 | 1.82E+08 | 1.82E+08 | + | 0 | NA | Intergenic | Intergenic | -10833 | NM_13381 | 98386 | Mm.4538 | NM_13381 | ENSMUSG | Lbr | AI505894\| | lamin B re | protein-co | 6.96 | 7.56 | 5.9 | 1.67 | 1.10E-09 | 2.54E-07 |
| 264 | chr11 | 1.2E+08 | 1.2E+08 | + | 0 | NA | intron (NM | intron (NM | 35737 | NM_19842 | 268515 | Mm.30949 | NM_19842 | ENSMUSG | Bahcc1 | B930044J | BAH doma | protein-co | 5.76 | 6.53 | 3.99 | 2.54 | 1.11E-09 | 2.54E-07 |
| 266 | chr5 | 1.18E+08 | 1.18E+08 | + | 0 | NA | Intergenic | Intergenic | 26705 | NM_00108 | 76792 | Mm.10342 | NM_00108 | ENSMUSG | 2410131K | - | RIKEN cDN | protein-co | 5.64 | 6.43 | 3.74 | 2.69 | 1.22E-09 | 2.77E-07 |
| 267 | chr18 | 81400285 | 81400435 | + | 0 | NA | Intergenic | Intergenic | -413339 | NM_17828 | 20689 | Mm.21591 | NM_17828 | ENSMUSG | Sall3 | B130022O | spalt like t | protein-co | 7.67 | 8.2 | 6.81 | 1.38 | 1.29E-09 | 2.91E-07 |
| 268 | chr12 | 25010520 | 25010670 | + | 0 | NA | intron (NM | intron (NM | 35663 | NM_00108 | 77480 | Mm.25064 | NM_00108 | ENSMUSG | Kidins220 | 3110039L1 | kinase D-ir | protein-co | 5.9 | 6.66 | 4.18 | 2.48 | 1.29E-09 | 2.91E-07 |
| 269 | chr4 | 1.03E+08 | 1.03E+08 | + | 0 | NA | promoter- | promoter- | -676 | NM_00116 | 67344 | Mm.11134 | NM_02610 | ENSMUSG | Tctex1d1 | 1700055O | Tctex1 dor | protein-co | 5.99 | 6.74 | 4.38 | 2.36 | 1.37E-09 | 3.05E-07 |

| | | | | | | | | | | | | | | | | | | | | | | | | |
|---|---|---|---|---|---|---|---|---|---|---|---|---|---|---|---|---|---|---|---|---|---|---|---|---|
| 273 | chr2 | 1.02E+08 | 1.02E+08 | + | 0 | NA | intron (NM | intron (NM | 146905 | NM_00129 | 241576 | Mm.48974 | NM_17888 | ENSMUSG | Ldlrad3 | 6430500P | low densit | protein-co | 5.45 | 6.29 | 3.23 | 3.06 | 1.47E-09 | 3.25E-07 |
| 275 | chr4 | 1.41E+08 | 1.41E+08 | + | 0 | NA | Intergenic | Intergenic | -3281 | NM_02909 | 74772 | Mm.20562 | NM_02909 | ENSMUSG | Atp13a2 | 1110012E | ATPase typ | protein-co | 7.96 | 8.44 | 7.23 | 1.21 | 1.57E-09 | 3.44E-07 |
| 282 | chr1 | 39396498 | 39396648 | + | 0 | NA | intron (NM | intron (NM | 28722 | NM_00129 | 114641 | Mm.37930 | NM_05329 | ENSMUSG | Rpl31 | Dts | ribosomal | protein-co | 5.8 | 6.57 | 4.08 | 2.48 | 1.78E-09 | 3.79E-07 |
| 297 | chr2 | 1.02E+08 | 1.02E+08 | + | 0 | NA | intron (NM | intron (NM | 41355 | NM_00108 | 72446 | Mm.24138 | NM_17518 | ENSMUSG | Prr5l | 2600010E | proline ric | protein-co | 6.95 | 7.55 | 5.89 | 1.66 | 2.36E-09 | 4.79E-07 |
| 302 | chr2 | 1.28E+08 | 1.28E+08 | + | 0 | NA | intron (NR | intron (NR | 2933 | NM_00129 | 56461 | Mm.31529 | NM_01978 | ENSMUSG | Kcnip3 | 4933407H | Kv channe | protein-co | 5.85 | 6.6 | 4.22 | 2.38 | 2.46E-09 | 4.90E-07 |
| 304 | chr17 | 43232913 | 43233063 | + | 0 | NA | Intergenic | Intergenic | -37359 | NM_13337 | 77596 | Mm.15995 | NM_13337 | ENSMUSG | Adgrf1 | 5031409J1 | adhesion G | protein-co | 6.95 | 7.56 | 5.88 | 1.68 | 2.52E-09 | 4.97E-07 |
| 308 | chr18 | 24596886 | 24597036 | + | 0 | NA | intron (NM | intron (NM | 6856 | NM_13914 | 106957 | Mm.21688 | NM_13914 | ENSMUSG | Slc39a6 | Ermelin\|Zi | solute carr | protein-co | 5.9 | 6.64 | 4.29 | 2.35 | 2.58E-09 | 5.05E-07 |
| 309 | chr1 | 1.38E+08 | 1.38E+08 | + | 0 | NA | intron (NM | intron (NM | 17881 | NM_00111 | 19264 | Mm.39157 | NM_01112 | ENSMUSG | Ptprc | B220\|CD4 | protein tyr | protein-co | 6.11 | 6.82 | 4.62 | 2.2 | 2.79E-09 | 5.43E-07 |
| 315 | chr13 | 1.17E+08 | 1.17E+08 | + | 0 | NA | Intergenic | Intergenic | -4227 | NM_01033 | 13723 | Mm.27492 | NM_01033 | ENSMUSG | Emb | AL022799 | embigin | protein-co | 8.19 | 8.63 | 7.56 | 1.07 | 3.07E-09 | 5.86E-07 |
| 318 | chr1 | 1.67E+08 | 1.67E+08 | + | 0 | NA | intron (NM | intron (NM | 39609 | NM_03072 | 80914 | Mm.28089 | NM_03072 | ENSMUSG | Uck2 | AA407809 | uridine-cyt | protein-co | 6.47 | 7.14 | 5.19 | 1.95 | 3.20E-09 | 6.04E-07 |
| 322 | chr10 | 76293330 | 76293480 | + | 0 | NA | intron (NM | (TGGG)n\|S | 39569 | NM_00911 | 20203 | Mm.23599 | NM_00911 | ENSMUSG | S100b | AI850290 | S100 prote | protein-co | 6.75 | 7.43 | 5.43 | 2 | 3.26E-09 | 6.10E-07 |
| 323 | chr17 | 56444952 | 56445102 | + | 0 | NA | intron (NM | intron (NM | 17194 | NM_00129 | 19280 | Mm.25877 | NM_01112 | ENSMUSG | Ptprs | AL022616 | protein tyr | protein-co | 5.42 | 6.24 | 3.29 | 2.95 | 3.47E-09 | 6.46E-07 |
| 328 | chr12 | 25092722 | 25092872 | + | 0 | NA | Intergenic | Intergenic | 3295 | NM_01049 | 15902 | Mm.34871 | NM_01049 | ENSMUSG | Id2 | AI255428 | inhibitor o | protein-co | 8.14 | 8.65 | 7.36 | 1.28 | 3.60E-09 | 6.61E-07 |
| 331 | chr2 | 32062298 | 32062448 | + | 0 | NA | intron (NR | RLTR8\|LTF | 1791 | NR_15149 | 241303 | Mm.25044 | NM_17551 | ENSMUSG | Fam78a | A130092J | family with | protein-co | 6.2 | 6.89 | 4.86 | 2.03 | 3.82E-09 | 6.94E-07 |
| 332 | chr1 | 55524115 | 55524265 | + | 0 | NA | intron (NM | intron (NM | 118244 | NM_00111 | 227120 | Mm.19418 | NM_00111 | ENSMUSG | Plcl1 | C230017K | phospholip | protein-co | 5 | 5.92 | 1.88 | 4.04 | 3.87E-09 | 7.02E-07 |
| 335 | chr9 | 75303799 | 75303949 | + | 0 | NA | Intergenic | Intergenic | -2414 | NM_01031 | 14697 | Mm.17604 | NM_01031 | ENSMUSG | Gnb5 | GBS\|Gbet | guanine nu | protein-co | 6.25 | 6.93 | 4.9 | 2.03 | 4.29E-09 | 7.69E-07 |
| 336 | chr15 | 66824788 | 66824938 | + | 0 | NA | intron (NM | intron (NM | 6966 | NM_00102 | 20491 | Mm.7601 | NM_00919 | ENSMUSG | Sla | Slap\|Slap- | src-like ad | protein-co | 6.17 | 6.88 | 4.73 | 2.15 | 4.30E-09 | 7.69E-07 |
| 337 | chr11 | 69460166 | 69460316 | + | 0 | NA | intron (NM | intron (NM | -44047 | NM_00101 | 216850 | Mm.26120 | NM_00101 | ENSMUSG | Kdm6b | 1700064E | KDM1 lysir | protein-co | 5.03 | 5.92 | 2.25 | 3.66 | 4.46E-09 | 7.97E-07 |
| 338 | chr12 | 3548589 | 3548739 | + | 0 | NA | Intergenic | Intergenic | -23727 | NM_00135 | 13528 | Mm.28620 | NM_00788 | ENSMUSG | Dtnb | - | dystrobrev | protein-co | 6.01 | 6.74 | 4.46 | 2.27 | 4.51E-09 | 8.03E-07 |
| 341 | chr11 | 1.17E+08 | 1.17E+08 | + | 0 | NA | intron (NM | intron (NM | -3886 | NM_01738 | 53860 | Mm.38450 | NM_01738 | ENSMUSG | 9-Sep | MSF1\|Msf | septin 9 | protein-co | 6.76 | 7.37 | 5.68 | 1.69 | 4.87E-09 | 8.60E-07 |
| 343 | chr11 | 98459378 | 98459528 | + | 0 | NA | Intergenic | Intergenic | 12619 | NM_01034 | 14786 | Mm.27670 | NM_01034 | ENSMUSG | Grb7 | mKIAA402 | growth fac | protein-co | 5.56 | 6.37 | 3.54 | 2.83 | 4.92E-09 | 8.63E-07 |
| 346 | chr3 | 1.07E+08 | 1.07E+08 | + | 0 | NA | Intergenic | Intergenic | -11109 | NM_00841 | 16490 | Mm.39285 | NM_00841 | ENSMUSG | Kcna2 | Akr6a4\|Gr | potassium | protein-co | 5.51 | 6.32 | 3.49 | 2.83 | 5.13E-09 | 8.93E-07 |
| 350 | chr1 | 1.34E+08 | 1.34E+08 | + | 0 | NA | Intergenic | Intergenic | 11805 | NM_00757 | 12227 | Mm.39264 | NM_00757 | ENSMUSG | Btg2 | AA959598 | B cell trans | protein-co | 6.52 | 7.22 | 5.11 | 2.12 | 5.68E-09 | 9.77E-07 |
| 352 | chr1 | 1.84E+08 | 1.84E+08 | + | 0 | NA | Intergenic | Intergenic | 28177 | NM_02851 | 73363 | Mm.50115 | NM_02851 | ENSMUSG | 1700056E2 | - | RIKEN cDN | protein-co | 7.3 | 7.83 | 6.44 | 1.39 | 5.94E-09 | 1.02E-06 |
| 353 | chr16 | 10815025 | 10815175 | + | 0 | NA | Intergenic | Intergenic | -18277 | NM_01363 | 19118 | Mm.42733 | NM_01363 | ENSMUSG | Prm1 | Prm-1 | protamine | protein-co | 6.98 | 7.59 | 5.92 | 1.67 | 6.07E-09 | 1.03E-06 |
| 363 | chr18 | 84804431 | 84804581 | + | 0 | NA | intron (NM | intron (NM | 34135 | NM_00137 | 1.01E+08 | | NM_00137 | ENSMUSG | Gm17266 | - | predicted | protein-co | 7.93 | 8.43 | 7.15 | 1.29 | 7.34E-09 | 1.22E-06 |
| 367 | chr10 | 80914356 | 80914506 | + | 0 | NA | intron (NM | intron (NM | 3814 | NM_01072 | 16907 | Mm.7362 | NM_01072 | ENSMUSG | Lmnb2 | - | lamin B2 | protein-co | 5.23 | 6.06 | 3.09 | 2.97 | 7.93E-09 | 1.30E-06 |
| 373 | chr5 | 1.44E+08 | 1.44E+08 | + | 0 | NA | intron (NM | intron (NM | -12173 | NM_17768 | 231874 | Mm.32043 | NM_17768 | ENSMUSG | Ccz1 | AU022870 | CCZ1 vacu | protein-co | 6.19 | 6.87 | 4.84 | 2.03 | 8.46E-09 | 1.36E-06 |
| 374 | chr14 | 19962021 | 19962171 | + | 0 | NA | intron (NM | ID4\|SINE\| | 14597 | NM_00128 | 14702 | Mm.41737 | NM_01031 | ENSMUSG | Gng2 | - | guanine nu | protein-co | 6.37 | 7.02 | 5.16 | 1.86 | 8.66E-09 | 1.39E-06 |
| 375 | chr13 | 63108770 | 63108920 | + | 0 | NA | intron (NR | intron (NM | -131184 | NR_11052 | 72061 | Mm.18810 | NM_02807 | ENSMUSG | Aopep | 2010111I0 | aminopept | protein-co | 6.94 | 7.54 | 5.9 | 1.64 | 8.94E-09 | 1.43E-06 |
| 376 | chr6 | 72993449 | 72993599 | + | 0 | NA | Intergenic | Intergenic | -34776 | NM_00119 | 19240 | Mm.3532 | NM_02528 | ENSMUSG | Tmsb10 | Ptmb10\|T | thymosin, | protein-co | 6.53 | 7.17 | 5.34 | 1.83 | 9.01E-09 | 1.44E-06 |
| 379 | chr9 | 63737570 | 63737720 | + | 0 | NA | intron (NM | intron (NM | 20349 | NM_01676 | 17127 | Mm.7320 | NM_01676 | ENSMUSG | Smad3 | AU022421 | SMAD fam | protein-co | 7.56 | 8.07 | 6.77 | 1.3 | 9.61E-09 | 1.52E-06 |
| 382 | chr4 | 1.41E+08 | 1.41E+08 | + | 0 | NA | Intergenic | Intergenic | -1726 | NM_01013 | 13836 | Mm.2581 | NM_01013 | ENSMUSG | Epha2 | AW545284 | Eph recept | protein-co | 6.97 | 7.54 | 6.01 | 1.53 | 9.90E-09 | 1.56E-06 |
| 390 | chr8 | 25070421 | 25070571 | + | 0 | NA | intron (NM | intron (NM | 31315 | NM_03125 | 83436 | Mm.26112 | NM_03125 | ENSMUSG | Plekha2 | 6430512N | pleckstrin | protein-co | 7.33 | 7.86 | 6.47 | 1.39 | 1.10E-08 | 1.69E-06 |
| 398 | chr9 | 73089616 | 73089766 | + | 0 | NA | Intergenic | Intergenic | -12707 | NM_02589 | 66991 | Mm.47174 | NM_02589 | ENSMUSG | Khdc3 | 2410004A | KH domair | protein-co | 7.27 | 7.82 | 6.36 | 1.46 | 1.23E-08 | 1.86E-06 |
| 400 | chr2 | 1.02E+08 | 1.02E+08 | + | 0 | NA | intron (NM | intron (NM | 55526 | NM_00108 | 72446 | Mm.24138 | NM_17518 | ENSMUSG | Prr5l | 2600010E | proline ric | protein-co | 5.74 | 6.48 | 4.15 | 2.33 | 1.27E-08 | 1.91E-06 |
| 401 | chr14 | 1.22E+08 | 1.22E+08 | + | 0 | NA | Intergenic | ORR1D2\|L | -32512 | NM_13407 | 105445 | Mm.37859 | NM_13407 | ENSMUSG | Dock9 | AA959601 | dedicator | protein-co | 5.68 | 6.42 | 4.06 | 2.36 | 1.28E-08 | 1.92E-06 |
| 411 | chr7 | 48937682 | 48937832 | + | 0 | NA | Intergenic | Intergenic | -21316 | NM_00111 | 78286 | Mm.33388 | NM_17527 | ENSMUSG | Nav2 | 5330421F | neuron na | protein-co | 7.6 | 8.14 | 6.73 | 1.42 | 1.41E-08 | 2.06E-06 |
| 421 | chr2 | 1.59E+08 | 1.59E+08 | + | 0 | NA | intron (NM | intron (NM | -44638 | NM_02797 | 71878 | Mm.85162 | NM_02797 | ENSMUSG | Fam83d | 2310007D | family with | protein-co | 7.44 | 7.97 | 6.59 | 1.37 | 1.61E-08 | 2.29E-06 |
| 420 | chr11 | 95356789 | 95356939 | + | 0 | NA | intron (NM | intron (NM | 19846 | NM_17254 | 215512 | Mm.26211 | NM_17254 | ENSMUSG | Fam117a | 5730593F | family with | protein-co | 7.22 | 7.76 | 6.35 | 1.42 | 1.60E-08 | 2.29E-06 |
| 429 | chr5 | 1.12E+08 | 1.12E+08 | + | 0 | NA | intron (NM | intron (NM | 17230 | NM_00135 | 72898 | Mm.15174 | NM_02838 | ENSMUSG | Asphd2 | 2900006N | aspartate | protein-co | 6.46 | 7.09 | 5.32 | 1.77 | 1.71E-08 | 2.39E-06 |
| 437 | chr4 | 1.36E+08 | 1.36E+08 | + | 0 | NA | Intergenic | CpG | 4173 | NM_00832 | 15903 | Mm.110 | NM_00832 | ENSMUSG | Id3 | Hlh462\|Id | inhibitor o | protein-co | 7.48 | 7.99 | 6.68 | 1.32 | 1.82E-08 | 2.49E-06 |
| 440 | chr15 | 86094447 | 86094597 | + | 0 | NA | intron (NM | intron (NM | 35795 | NM_00120 | 223752 | Mm.24442 | NM_17261 | ENSMUSG | Gramd4 | 9930016O | GRAM dor | protein-co | 6.91 | 7.48 | 5.96 | 1.52 | 1.91E-08 | 2.60E-06 |
| 446 | chr11 | 58883880 | 58884030 | + | 0 | NA | 3' UTR (NM | 3' UTR (NM | 16715 | NM_17240 | 69944 | Mm.32118 | NM_17240 | ENSMUSG | 2810021J2 | AI449137 | RIKEN cDN | protein-co | 5.59 | 6.36 | 3.82 | 2.55 | 2.01E-08 | 2.70E-06 |
| 455 | chr5 | 34249262 | 34249412 | + | 0 | NA | intron (NM | intron (NM | 38987 | NM_00101 | 231125 | Mm.11338 | NM_00101 | ENSMUSG | Zfyve28 | 9630058O | zinc finger | protein-co | 5.58 | 6.33 | 3.92 | 2.41 | 2.18E-08 | 2.86E-06 |
| 460 | chr10 | 8287462 | 8287612 | + | 0 | NA | intron (NM | intron (NM | 231288 | NM_17738 | 338362 | Mm.41163 | NM_17738 | ENSMUSG | Ust | D930010O | uronyl-2-s | protein-co | 5.58 | 6.39 | 3.62 | 2.77 | 2.24E-08 | 2.91E-06 |
| 467 | chr11 | 1.15E+08 | 1.15E+08 | + | 0 | NA | intron (NM | intron (NM | 3542 | NM_02672 | 68572 | Mm.27106 | NM_02672 | ENSMUSG | Mrpl58 | 1110001A | mitochonc | protein-co | 5.81 | 6.55 | 4.19 | 2.37 | 2.40E-08 | 3.07E-06 |
| 471 | chr18 | 82484966 | 82485116 | + | 0 | NA | intron (NM | intron (NM | 9918 | NM_00102 | 17196 | Mm.25206 | NM_01077 | ENSMUSG | Mbp | C76307\|H | myelin bas | protein-co | 6.8 | 7.4 | 5.74 | 1.65 | 2.43E-08 | 3.09E-06 |
| 470 | chr18 | 74295004 | 74295154 | + | 0 | NA | intron (NR | intron (NR | 11979 | NM_02894 | 74453 | Mm.18229 | NM_02894 | ENSMUSG | Cfap53 | 4933415I0 | cilia and fl | protein-co | 6.09 | 6.77 | 4.77 | 2 | 2.43E-08 | 3.09E-06 |
| 475 | chr12 | 1.02E+08 | 1.02E+08 | + | 0 | NA | intron (NM | intron (NM | 60183 | NM_17215 | 238384 | Mm.31244 | NM_17215 | ENSMUSG | Slc24a4 | A930002M | solute carr | protein-co | 4.7 | 5.64 | 1.24 | 4.4 | 2.50E-08 | 3.15E-06 |
| 480 | chr2 | 93192673 | 93192823 | + | 0 | NA | intron (NM | intron (NM | 5200 | NM_00135 | 277414 | Mm.41033 | NM_00102 | ENSMUSG | Trp53i11 | Tp53i11 | transform | protein-co | 7.98 | 8.42 | 7.35 | 1.08 | 2.59E-08 | 3.24E-06 |
| 481 | chr4 | 1.38E+08 | 1.38E+08 | + | 0 | NA | intron (NM | intron (NM | -4744 | NM_01062 | 16559 | Mm.27193 | NM_01062 | ENSMUSG | Kif17 | 5930435E | kinesin far | protein-co | 6.35 | 7.01 | 5.14 | 1.87 | 2.63E-08 | 3.28E-06 |
| 485 | chr11 | 1.18E+08 | 1.18E+08 | + | 0 | NA | Intergenic | Intergenic | -6473 | NM_17257 | 217364 | Mm.47396 | NM_17257 | ENSMUSG | Engase | C130099A | endo-beta | protein-co | 5.72 | 6.44 | 4.17 | 2.28 | 2.70E-08 | 3.34E-06 |
| 487 | chr4 | 47186645 | 47186795 | + | 0 | NA | Intergenic | Intergenic | -21292 | NM_00992 | 12819 | Mm.23354 | NM_00992 | ENSMUSG | Col15a1 | - | collagen, t | protein-co | 5.81 | 6.55 | 4.23 | 2.31 | 2.73E-08 | 3.36E-06 |
| 493 | chr7 | 67419705 | 67419855 | + | 0 | NA | Intergenic | Intergenic | -46922 | NM_00103 | 17258 | Mm.13278 | NM_00103 | ENSMUSG | Mef2a | A430079H | myocyte e | protein-co | 4.9 | 5.77 | 2.41 | 3.36 | 2.88E-08 | 3.49E-06 |
| 509 | chrX | 1.6E+08 | 1.6E+08 | + | 0 | NA | intron (NM | intron (NM | -11388 | NR_11096 | 58194 | Mm.28649 | NM_02138 | ENSMUSG | Sh3kbp1 | 1200007H | SH3-doma | protein-co | 6.69 | 7.28 | 5.68 | 1.6 | 3.38E-08 | 3.96E-06 |
| 516 | chr18 | 10568653 | 10568803 | + | 0 | NA | intron (NM | L1M4\|LINI | 41624 | NM_00108 | 77805 | Mm.21099 | NM_14454 | ENSMUSG | Esco1 | A930014I | establishm | protein-co | 5.71 | 6.45 | 4.12 | 2.32 | 3.57E-08 | 4.14E-06 |
| 515 | chr6 | 8712025 | 8712175 | + | 0 | NA | intron (NM | intron (NM | 46358 | NM_00129 | 15893 | Mm.27568 | NM_01049 | ENSMUSG | Ica1 | 69kDa\|ICA | islet cell a | protein-co | 5.52 | 6.29 | 3.78 | 2.51 | 3.57E-08 | 4.14E-06 |
| 521 | chr16 | 97016622 | 97016772 | + | 0 | NA | intron (NM | ORR1E\|LT | 154038 | NM_03117 | 13508 | Mm.10243 | NM_03117 | ENSMUSG | Dscam | 4932410A | DS cell adh | protein-co | 5.73 | 6.47 | 4.11 | 2.36 | 3.75E-08 | 4.30E-06 |
| 522 | chr5 | 1.4E+08 | 1.4E+08 | + | 0 | NA | intron (NM | intron (NM | 3380 | NM_17489 | 231830 | Mm.24051 | NM_17489 | ENSMUSG | Micall2 | A930021H | MICAL-like | protein-co | 5.92 | 6.61 | 4.54 | 2.08 | 3.77E-08 | 4.31E-06 |
| 529 | chr1 | 36831456 | 36831606 | + | 0 | NA | intron (NM | intron (NM | 52547 | NM_00128 | 22637 | Mm.8038 | NM_00951 | ENSMUSG | Zap70 | Srk\|ZAP-7 | zeta-chain | protein-co | 5.72 | 6.46 | 4.14 | 2.32 | 3.90E-08 | 4.41E-06 |
| 532 | chr17 | 84881050 | 84881200 | + | 0 | NA | Intergenic | LTR33\|LTF | -75616 | NM_00115 | 19043 | Mm.24969 | NM_01115 | ENSMUSG | Ppm1b | PP2CB | protein ph | protein-co | 5.76 | 6.47 | 4.29 | 2.18 | 4.03E-08 | 4.53E-06 |
| 539 | chr15 | 78486704 | 78486854 | + | 0 | NA | intron (NM | CT-rich\|Lo | 8287 | NM_00836 | 16185 | Mm.35287 | NM_00836 | ENSMUSG | Il2rb | CD122\|IL- | interleukir | protein-co | 4.82 | 5.69 | 2.25 | 3.44 | 4.22E-08 | 4.67E-06 |
| 550 | chr12 | 87196492 | 87196642 | + | 0 | NA | intron (NM | intron (NM | 3662 | NM_00103 | 382620 | Mm.37491 | NM_00103 | ENSMUSG | Tmed8 | 6430595O | transmem | protein-co | 5.21 | 6.03 | 3.09 | 2.94 | 4.51E-08 | 4.90E-06 |
| 555 | chr12 | 16008323 | 16008473 | + | 0 | NA | Intergenic | Intergenic | -191613 | NM_14455 | 217410 | Mm.49113 | NM_14455 | ENSMUSG | Trib2 | AW319517 | tribbles ps | protein-co | 6.79 | 7.39 | 5.76 | 1.62 | 4.59E-08 | 4.94E-06 |
| 557 | chr6 | 1.09E+08 | 1.09E+08 | + | 0 | NA | intron (NM | intron (NM | 7554 | NM_13867 | 192193 | Mm.21596 | NM_13867 | ENSMUSG | Edem1 | A130059K | ER degrad | protein-co | 7.87 | 8.35 | 7.16 | 1.19 | 4.65E-08 | 4.99E-06 |
| 560 | chr18 | 82479918 | 82480068 | + | 0 | NA | intron (NM | intron (NM | 4870 | NM_00102 | 17196 | Mm.25206 | NM_01077 | ENSMUSG | Mbp | C76307\|H | myelin bas | protein-co | 6.86 | 7.43 | 5.91 | 1.52 | 4.79E-08 | 5.11E-06 |
| 564 | chr13 | 13529219 | 13529369 | + | 0 | NA | Intergenic | Intergenic | -61115 | NM_01074 | 17101 | Mm.34233 | NM_01074 | ENSMUSG | Lyst | D13Sfk13\| | lysosomal | protein-co | 7.5 | 7.99 | 6.76 | 1.23 | 4.96E-08 | 5.25E-06 |
| 565 | chr7 | 84173537 | 84173687 | + | 0 | NA | Intergenic | MLT1G3\|L | -21719 | NM_13337 | 70178 | Mm.23914 | NM_13337 | ENSMUSG | Abhd17c | 2210412D | abhydrola | protein-co | 5.85 | 6.59 | 4.27 | 2.32 | 5.00E-08 | 5.29E-06 |
| 573 | chr11 | 97977913 | 97978063 | + | 0 | NA | intron (NM | intron (NM | 8458 | NM_00116 | 72324 | Mm.39617 | NM_02819 | ENSMUSG | Plxdc1 | 2410003I0 | plexin don | protein-co | 5.08 | 5.91 | 2.91 | 2.99 | 5.21E-08 | 5.44E-06 |
| 574 | chr8 | 1.24E+08 | 1.24E+08 | + | 0 | NA | Intergenic | Intergenic | 18901 | NM_13399 | 69581 | Mm.16825 | NM_13399 | ENSMUSG | Rhou | 2310026M | ras homolc | protein-co | 5.69 | 6.42 | 4.17 | 2.25 | 5.31E-08 | 5.53E-06 |
| 580 | chr2 | 1.64E+08 | 1.64E+08 | + | 0 | NA | intron (NM | intron (NM | 1877 | NM_01152 | 20971 | Mm.3815 | NM_01152 | ENSMUSG | Sdc4 | AA959608 | syndecan | protein-co | 7.94 | 8.38 | 7.33 | 1.05 | 5.49E-08 | 5.67E-06 |
| 594 | chr2 | 35504133 | 35504283 | + | 0 | NA | Intergenic | Intergenic | -42759 | NM_00130 | 14594 | Mm.28112 | NM_01028 | ENSMUSG | Ggta1 | AW108479 | glycoprote | protein-co | 5.89 | 6.6 | 4.43 | 2.18 | 6.16E-08 | 6.21E-06 |
| 595 | chr2 | 1.28E+08 | 1.28E+08 | + | 0 | NA | intron (NM | intron (NM | 25426 | NM_00128 | 12125 | Mm.14108 | NM_00975 | ENSMUSG | Bcl2l11 | 1500006F | BCL2-like 1 | protein-co | 5.05 | 5.88 | 2.92 | 2.95 | 6.19E-08 | 6.21E-06 |
| 602 | chrX | 11845654 | 11845804 | + | 0 | NA | Intergenic | Intergenic | 234824 | NM_02951 | 71458 | Mm.19632 | NM_02951 | ENSMUSG | Bcor | 2900008C | BCL6 inter | protein-co | 5.11 | 5.93 | 3.03 | 2.9 | 6.49E-08 | 6.46E-06 |
| 607 | chr1 | 1.38E+08 | 1.38E+08 | + | 0 | NA | Intergenic | Intergenic | 25942 | NM_17739 | 338375 | Mm.11442 | NM_17739 | ENSMUSG | Atp6v1g3 | - | ATPase, H | protein-co | 6.86 | 7.47 | 5.77 | 1.7 | 6.62E-08 | 6.53E-06 |
| 612 | chr15 | 57878511 | 57878661 | + | 0 | NA | intron (NM | intron (NM | 13832 | NM_02420 | 67819 | Mm.28938 | NM_02420 | ENSMUSG | Derl1 | 1110021N | Der1-like c | protein-co | 6.73 | 7.32 | 5.74 | 1.58 | 6.80E-08 | 6.66E-06 |
| 611 | chr16 | 62807747 | 62807897 | + | 0 | NA | Intergenic | Intergenic | -6854 | NM_02658 | 68159 | Mm.38679 | NM_02658 | ENSMUSG | Stx19 | A030009B | syntaxin 1 | protein-co | 5.84 | 6.57 | 4.3 | 2.27 | 6.80E-08 | 6.66E-06 |
| 616 | chr4 | 1.34E+08 | 1.34E+08 | + | 0 | NA | Intergenic | Intergenic | -6234 | NM_00116 | 230806 | Mm.24008 | NM_00116 | ENSMUSG | Crybg2 | Aim1l | crystallin | protein-co | 6.13 | 6.84 | 4.65 | 2.19 | 6.93E-08 | 6.73E-06 |
| 628 | chr19 | 18619790 | 18619940 | + | 0 | NA | intron (NM | intron (NM | 11948 | NM_01737 | 20409 | Mm.17222 | NM_01737 | ENSMUSG | Ostf1 | C78236\|SI | osteoclast | protein-co | 6.87 | 7.44 | 5.92 | 1.52 | 7.37E-08 | 7.03E-06 |
| 631 | chr17 | 45488972 | 45489122 | + | 0 | NA | Intergenic | B3A\|SINE\| | -14109 | NM_02764 | 71020 | Mm.44257 | NM_02764 | ENSMUSG | Spats1 | 1700011H | spermatog | protein-co | 5.13 | 5.94 | 3.05 | 2.9 | 7.57E-08 | 7.18E-06 |
| 635 | chr7 | 72280845 | 72280995 | + | 0 | NA | intron (NM | intron (NM | 25675 | NM_00102 | 244049 | Mm.21741 | NM_00102 | ENSMUSG | Mctp2 | Gm489 | multiple C | protein-co | 5.64 | 6.38 | 4.03 | 2.35 | 7.81E-08 | 7.35E-06 |
| 643 | chr9 | 44515602 | 44515752 | + | 0 | NA | intron (NM | intron (NM | 10767 | NM_00755 | 12145 | Mm.6246 | NM_00755 | ENSMUSG | Cxcr5 | Blr1\|CXC- | chemokine | protein-co | 6.03 | 6.7 | 4.74 | 1.96 | 8.17E-08 | 7.59E-06 |
| 649 | chr8 | 14994499 | 14994649 | + | 0 | NA | intron (NM | intron (NM | -16451 | NM_02911 | 74901 | Mm.46675 | NM_02911 | ENSMUSG | Kbtbd11 | 2900016B | kelch repe | protein-co | 7.07 | 7.6 | 6.22 | 1.38 | 8.36E-08 | 7.70E-06 |
| 670 | chr15 | 4816884 | 4817034 | + | 0 | NA | Intergenic | Intergenic | -81778 | NM_00116 | 223825 | Mm.15928 | NM_00116 | ENSMUSG | Mroh2b | 4930455B | maestro he | protein-co | 6.74 | 7.33 | 5.74 | 1.59 | 9.29E-08 | 8.30E-06 |
| 674 | chr1 | 30793054 | 30793204 | + | 0 | NA | Intergenic | Intergenic | 70127 | NM_00108 | 213109 | Mm.19448 | NM_02587 | ENSMUSG | Phf3 | 2310061N | PHD finger | protein-co | 6.24 | 6.88 | 5.08 | 1.8 | 9.39E-08 | 8.33E-06 |
| 676 | chr10 | 95314197 | 95314347 | + | 0 | NA | intron (NM | intron (NM | 9437 | NM_00133 | 12905 | Mm.47373 | NM_00999 | ENSMUSG | Cradd | RAIDD | CASP2 and | protein-co | 5.09 | 5.9 | 3.09 | 2.81 | 9.49E-08 | 8.39E-06 |
| 698 | chr5 | 1.39E+08 | 1.39E+08 | + | 0 | NA | exon (NM | exon (NM_ | -11140 | NM_00135 | 19085 | Mm.30616 | NM_00892 | ENSMUSG | Prkar1b | AI385716 | protein kir | protein-co | 5.69 | 6.41 | 4.17 | 2.24 | 1.07E-07 | 9.14E-06 |
| 699 | chr5 | 1.12E+08 | 1.12E+08 | + | 0 | NA | intron (NM | intron (NM | 18422 | NM_00135 | 72898 | Mm.15174 | NM_02838 | ENSMUSG | Asphd2 | 2900006N | aspartate | protein-co | 5.33 | 6.1 | 3.56 | 2.54 | 1.07E-07 | 9.18E-06 |
| 702 | chr1 | 53791463 | 53791613 | + | 0 | NA | Intergenic | Intergenic | -6323 | NM_13381 | 98267 | Mm.25559 | NM_13381 | ENSMUSG | Stk17b | 3110009A | serine/thr | protein-co | 7.61 | 8.08 | 6.91 | 1.17 | 1.08E-07 | 9.22E-06 |
| 714 | chr17 | 23617150 | 23617300 | + | 0 | NA | Intergenic | MLTR11B\| | 16399 | NM_00128 | 332221 | Mm.27031 | NM_00103 | ENSMUSG | Zscan10 | BC068284 | zinc finger | protein-co | 6.05 | 6.7 | 4.81 | 1.89 | 1.15E-07 | 9.66E-06 |
| 725 | chr8 | 1.24E+08 | 1.24E+08 | + | 0 | NA | intron (NM | intron (NM | 86605 | NM_13927 | 108148 | Mm.33808 | NM_13927 | ENSMUSG | Galnt2 | AI480629 | polypeptic | protein-co | 7.15 | 7.68 | 6.29 | 1.38 | 1.22E-07 | 1.01E-05 |
| 729 | chr2 | 1.53E+08 | 1.53E+08 | + | 0 | NA | intron (NR | intron (NM | 12015 | NR_14925 | 12048 | Mm.23821 | NM_00974 | ENSMUSG | Bcl2l1 | Bcl(X)L\|Bc | BCL2-like 1 | protein-co | 6.49 | 7.12 | 5.34 | 1.78 | 1.24E-07 | 1.01E-05 |
| 737 | chr9 | 75645555 | 75645705 | + | 0 | NA | intron (NM | intron (NM | 19898 | NM_02730 | 70082 | Mm.19119 | NM_02730 | ENSMUSG | Lysmd2 | 2210402C | LysM, puta | protein-co | 6.19 | 6.83 | 4.98 | 1.85 | 1.29E-07 | 1.05E-05 |
| 739 | chr11 | 95834371 | 95834521 | + | 0 | NA | intron (NM | intron (NM | -7849 | NM_00128 | 14710 | Mm.46299 | NM_02312 | ENSMUSG | Gngt2 | AV096488 | guanine nu | protein-co | 7.01 | 7.54 | 6.16 | 1.39 | 1.32E-07 | 1.07E-05 |
| 760 | chr3 | 1.31E+08 | 1.31E+08 | + | 0 | NA | intron (NM | intron (NM | 41295 | NM_00127 | 16842 | Mm.25521 | NM_01070 | ENSMUSG | Lef1 | 3000002B | lymphoid | protein-co | 5.11 | 5.9 | 3.18 | 2.72 | 1.46E-07 | 1.15E-05 |
| 758 | chr2 | 1.28E+08 | 1.28E+08 | + | 0 | NA | intron (NM | intron (NM | 84703 | NM_02876 | 74121 | Mm.24483 | NM_02876 | ENSMUSG | Acoxl | 1200014P | acyl-Coenz | protein-co | 4.82 | 5.68 | 2.42 | 3.25 | 1.46E-07 | 1.15E-05 |
| 769 | chr11 | 52318522 | 52318672 | + | 0 | NA | Intergenic | Intergenic | -35583 | NM_00131 | 21414 | Mm.31630 | NM_00933 | ENSMUSG | Tcf7 | AI465550 | transcripti | protein-co | 6.37 | 7.01 | 5.19 | 1.82 | 1.56E-07 | 1.21E-05 |
| 775 | chr6 | 48693999 | 48694149 | + | 0 | NA | Intergenic | RMER1B\| | 9496 | NM_17499 | 107526 | Mm.33304 | NM_17499 | ENSMUSG | Gimap4 | AU019574 | GTPase, IN | protein-co | 5.01 | 5.83 | 2.91 | 2.92 | 1.59E-07 | 1.23E-05 |
| 792 | chr15 | 86094845 | 86094995 | + | 0 | NA | intron (NM | intron (NM | 36193 | NM_00120 | 223752 | Mm.24442 | NM_17261 | ENSMUSG | Gramd4 | 9930016O | GRAM dor | protein-co | 5.9 | 6.58 | 4.57 | 2.01 | 1.78E-07 | 1.34E-05 |
| 802 | chr8 | 85618897 | 85619047 | + | 0 | NA | Intergenic | RMER13B\| | -63611 | NM_01979 | 56445 | Mm.47557 | NM_01979 | ENSMUSG | Dnaja2 | 1500017M | DnaJ heat | protein-co | 6.7 | 7.32 | 5.59 | 1.72 | 1.86E-07 | 1.39E-05 |
| 821 | chr17 | 15678379 | 15678529 | + | 0 | NA | Intergenic | Intergenic | -26513 | NM_00769 | 12648 | Mm.8137 | NM_00769 | ENSMUSG | Chd1 | 4930525N | chromodo | protein-co | 4.89 | 5.71 | 2.74 | 2.97 | 2.10E-07 | 1.53E-05 |
| 833 | chr16 | 30066941 | 30067091 | + | 0 | NA | exon (NM | exon (NM_ | 1659 | NM_00823 | 15205 | Mm.39085 | NM_00823 | ENSMUSG | Hes1 | Hry\|bHLHI | hes family | protein-co | 7.78 | 8.22 | 7.14 | 1.08 | 2.28E-07 | 1.64E-05 |
| 840 | chr12 | 1.13E+08 | 1.13E+08 | + | 0 | NA | Intergenic | Intergenic | 11898 | NM_13404 | 104732 | Mm.87616 | NM_13404 | ENSMUSG | Tedc1 | 4930427A | tubulin ep | protein-co | 7.78 | 8.22 | 7.14 | 1.08 | 2.33E-07 | 1.66E-05 |
| 836 | chr7 | 1.25E+08 | 1.25E+08 | + | 0 | NA | intron (NM | intron (NM | 4115 | NM_02631 | 67711 | Mm.4467 | NM_02631 | ENSMUSG | Nsmce1 | 2510027N | NSE1 hom | protein-co | 5.5 | 6.22 | 3.98 | 2.25 | 2.32E-07 | 1.66E-05 |
| 863 | chr4 | 1.01E+08 | 1.01E+08 | + | 0 | NA | intron (NM | intron (NM | 77693 | NM_01356 | 16451 | Mm.28965 | NM_01356 | ENSMUSG | Jak1 | AA960307 | Janus kina | protein-co | 6.65 | 7.22 | 5.71 | 1.51 | 2.62E-07 | 1.81E-05 |
| 875 | chr4 | 47186899 | 47187049 | + | 0 | NA | Intergenic | Intergenic | -21038 | NM_00992 | 12819 | Mm.23354 | NM_00992 | ENSMUSG | Col15a1 | - | collagen, t | protein-co | 5.07 | 5.88 | 3.01 | 2.87 | 2.79E-07 | 1.91E-05 |
| 892 | chr14 | 45155632 | 45155782 | + | 0 | NA | intron (NM | MT2A\|LTR | 63718 | NM_17259 | 70561 | Mm.28129 | NM_17259 | ENSMUSG | Txndc16 | 5730420B | thioredoxi | protein-co | 5.81 | 6.51 | 4.41 | 2.1 | 3.09E-07 | 2.07E-05 |
| 906 | chr5 | 24826497 | 24826647 | + | 0 | NA | intron (NM | intron (NM | 15789 | NM_05307 | 19744 | Mm.31917 | NM_05307 | ENSMUSG | Rheb | Rheb1 | Ras homol | protein-co | 5.85 | 6.53 | 4.51 | 2.02 | 3.28E-07 | 2.16E-05 |
| 910 | chr10 | 60389655 | 60389805 | + | 0 | NA | intron (NR | intron (NR | -9996 | NM_00116 | 1E+08 | Mm.44413 | NM_00116 | ENSMUSG | Gm17455 | - | predicted | protein-co | 7.36 | 7.85 | 6.59 | 1.26 | 3.37E-07 | 2.21E-05 |
| 909 | chr9 | 1.24E+08 | 1.24E+08 | + | 0 | NA | Intergenic | Intergenic | -10264 | NM_00116 | 12769 | Mm.44060 | NM_00991 | ENSMUSG | Ccr9 | A130091K | chemokine | protein-co | 6.14 | 6.77 | 5.01 | 1.76 | 3.37E-07 | 2.21E-05 |
| 918 | chr12 | 3547569 | 3547719 | + | 0 | NA | Intergenic | RLTR19A\|I | -24747 | NM_00135 | 13528 | Mm.28620 | NM_00788 | ENSMUSG | Dtnb | - | dystrobrev | protein-co | 5.62 | 6.35 | 4.07 | 2.28 | 3.48E-07 | 2.26E-05 |
| 928 | chr11 | 68462822 | 68462972 | + | 0 | NA | intron (NM | ID_B1\|SIN | 30772 | NM_17732 | 320207 | Mm.24496 | NM_17732 | ENSMUSG | Pik3r5 | AV230647 | phosphoin | protein-co | 4.63 | 5.5 | 2.11 | 3.39 | 3.55E-07 | 2.28E-05 |
| 931 | chr8 | 70695469 | 70695619 | + | 0 | NA | Intergenic | Intergenic | -2195 | NM_01059 | 16478 | Mm.41985 | NM_01059 | ENSMUSG | Jund | Jund1 | jun D prot | protein-co | 7.68 | 8.15 | 6.97 | 1.17 | 3.59E-07 | 2.30E-05 |
| 955 | chr17 | 44893702 | 44893852 | + | 0 | NA | intron (NR | intron (NR | -12892 | NM_02929 | 75341 | Mm.38682 | NM_029257 | | 4930564C | | RIKEN cDN | protein-co | 5.37 | 6.11 | 3.77 | 2.34 | 4.06E-07 | 2.53E-05 |
| 978 | chr11 | 52314234 | 52314384 | + | 0 | NA | Intergenic | CpG | -31295 | NM_00131 | 21414 | Mm.31630 | NM_00933 | ENSMUSG | Tcf7 | AI465550 | transcripti | protein-co | 5.13 | 5.9 | 3.33 | 2.58 | 4.50E-07 | 2.74E-05 |

| | | | | | | | | | | | | | | | | | | | | | | | | |
|---|---|---|---|---|---|---|---|---|---|---|---|---|---|---|---|---|---|---|---|---|---|---|---|---|
| 981 | chr10 | 26325405 | 26325555 | + | 0 | NA | intron (NM | RLTR14_RI | 48592 | NM_00135 | 237339 | Mm.33843 | NM_17278 | ENSMUSG | L3mbtl3 | AI481284 | | L3MBTL3 h | protein-co | 6.72 | 7.29 | 5.77 | 1.52 | 4.58E-07 | 2.78E-05 |
| 985 | chr8 | 1.22E+08 | 1.22E+08 | + | 0 | NA | promoter- | promoter- | -69 | NM_01391 | 30927 | Mm.10367 | NM_01391 | ENSMUSG | Snai3 | AI643946 | | snail famil | protein-co | 7.62 | 8.08 | 6.96 | 1.12 | 4.67E-07 | 2.82E-05 |
| 992 | chr11 | 1.18E+08 | 1.18E+08 | + | 0 | NA | Intergenic | Intergenic | 16989 | NM_00770 | 12702 | Mm.3468 | NM_00770 | ENSMUSG | Socs3 | Cis3 | Cish3 | suppressor | protein-co | 5.75 | 6.43 | 4.45 | 1.98 | 4.81E-07 | 2.89E-05 |
| 1002 | chr7 | 25000277 | 25000427 | + | 0 | NA | intron (NM | ID_B1 | SIN | 5585 | NM_00129 | 232975 | Mm.44101 | NM_14492 | ENSMUSG | Atp1a3 | Atpa-2 | ATPase, Na | protein-co | 5.78 | 6.46 | 4.43 | 2.02 | 4.98E-07 | 2.96E-05 |
| 1012 | chr14 | 15746944 | 15747094 | + | 0 | NA | Intergenic | Intergenic | -308032 | NM_14605 | 218763 | Mm.13803 | NM_14605 | ENSMUSG | Lrrc3b | BC019794 | leucine ric | protein-co | 5.78 | 6.46 | 4.43 | 2.04 | 5.20E-07 | 3.06E-05 |
| 1030 | chr12 | 1.13E+08 | 1.13E+08 | + | 0 | NA | Intergenic | Intergenic | -7566 | NM_01992 | 56696 | Mm.20455 | NM_01992 | ENSMUSG | Gpr132 | G2a | G protein- | protein-co | 6.45 | 7.05 | 5.42 | 1.62 | 5.49E-07 | 3.18E-05 |
| 1034 | chr1 | 1.84E+08 | 1.84E+08 | + | 0 | NA | Intergenic | Intergenic | 19716 | NM_02851 | 73363 | Mm.50115 | NM_02851 | ENSMUSG | 1700056E2 | - | RIKEN cDN | protein-co | 6.15 | 6.78 | 5 | 1.78 | 5.62E-07 | 3.24E-05 |
| 1047 | chr6 | 48725594 | 48725744 | + | 0 | NA | Intergenic | Intergenic | 7048 | NM_14616 | 231932 | Mm.30479 | NM_14616 | ENSMUSG | Gimap7 | IAN7 | Ian3 | GTPase, IM | protein-co | 6.37 | 6.96 | 5.37 | 1.59 | 5.86E-07 | 3.33E-05 |
| 1055 | chr16 | 18073751 | 18073901 | + | 0 | NA | intron (NM | intron (NM | 7423 | NM_00128 | 13353 | Mm.27155 | NM_01004 | ENSMUSG | Dgcr6 | - | DiGeorge s | protein-co | 5.7 | 6.39 | 4.32 | 2.07 | 6.19E-07 | 3.50E-05 |
| 1058 | chr1 | 1.66E+08 | 1.66E+08 | + | 0 | NA | intron (NM | intron (NM | 16406 | NM_17859 | 226594 | Mm.26555 | NM_17859 | ENSMUSG | Rcsd1 | A430105K | RCSD dom | protein-co | 6.75 | 7.29 | 5.88 | 1.41 | 6.24E-07 | 3.51E-05 |
| 1061 | chr10 | 81329360 | 81329510 | + | 0 | NA | intron (NM | intron (NM | 732 | NM_00127 | 21390 | Mm.4545 | NM_00932 | ENSMUSG | Tbxa2r | TP | TXA2-R | thromboxa | protein-co | 5.76 | 6.44 | 4.44 | 2 | 6.27E-07 | 3.52E-05 |
| 1071 | chr4 | 32258234 | 32258384 | + | 0 | NA | Intergenic | Intergenic | -159126 | NM_00110 | 12014 | Mm.49122 | NM_00752 | ENSMUSG | Bach2 | E030004N | BTB and C | protein-co | 7.06 | 7.56 | 6.28 | 1.28 | 6.60E-07 | 3.67E-05 |
| 1078 | chr10 | 60370741 | 60370891 | + | 0 | NA | 3' UTR (NM | 3' UTR (NM | 23965 | NM_00115 | 74048 | Mm.27358 | NM_02873 | ENSMUSG | Vsir | 4632428N | V-set imm | protein-co | 7.71 | 8.17 | 7.05 | 1.12 | 6.75E-07 | 3.72E-05 |
| 1084 | chr12 | 52824868 | 52825018 | + | 0 | NA | intron (NM | intron (NM | 125560 | NM_19811 | 238161 | Mm.31082 | NM_19811 | ENSMUSG | Akap6 | AI482140 | | A kinase (P | protein-co | 4.93 | 5.73 | 3 | 2.73 | 6.91E-07 | 3.79E-05 |
| 1085 | chr13 | 1.14E+08 | 1.14E+08 | + | 0 | NA | Intergenic | Intergenic | 44131 | NM_13079 | 170625 | Mm.33721 | NM_13079 | ENSMUSG | Snx18 | Snag1 | sorting ne | protein-co | 5.47 | 6.18 | 4.01 | 2.17 | 7.02E-07 | 3.85E-05 |
| 1111 | chr3 | 95668697 | 95668847 | + | 0 | NA | Intergenic | Intergenic | 10051 | NM_00856 | 17210 | Mm.1639 | NM_00856 | ENSMUSG | Mcl1 | AW55680 | myeloid ce | protein-co | 7.06 | 7.57 | 6.27 | 1.3 | 8.09E-07 | 4.32E-05 |
| 1113 | chr17 | 56819757 | 56819907 | + | 0 | NA | intron (NM | intron (NM | 11176 | NM_02778 | 19725 | Mm.102 | NM_00905 | ENSMUSG | Rfx2 | 5430432H | regulatory | protein-co | 6.6 | 7.17 | 5.65 | 1.52 | 8.13E-07 | 4.34E-05 |
| 1116 | chr7 | 3264419 | 3264569 | + | 0 | NA | Intergenic | Intergenic | -14754 | NM_00103 | 378425 | Mm.27715 | NM_00103 | ENSMUSG | Nlrp12 | Nalp12 | PY | NLR family | protein-co | 5.57 | 6.29 | 4.06 | 2.24 | 8.24E-07 | 4.38E-05 |
| 1121 | chr9 | 50416793 | 50416943 | + | 0 | NA | Intergenic | RMER20C_ | -77657 | NM_02963 | 76509 | Mm.29959 | NM_02963 | ENSMUSG | Plet1 | 0610037B | placenta e | protein-co | 7.14 | 7.65 | 6.36 | 1.29 | 8.34E-07 | 4.42E-05 |
| 1127 | chr2 | 1.73E+08 | 1.73E+08 | + | 0 | NA | Intergenic | Intergenic | -2295 | NM_01204 | 26972 | Mm.23495 | NM_01204 | ENSMUSG | Spo11 | AI449549 | | SPO11 me | protein-co | 7.66 | 8.15 | 6.9 | 1.25 | 8.46E-07 | 4.46E-05 |
| 1132 | chr17 | 86969745 | 86969895 | + | 0 | NA | intron (NM | intron (NM | 6709 | NM_14549 | 104215 | Mm.826 | NM_14549 | ENSMUSG | Rhoq | Arhq | TC1 | ras homol | protein-co | 6.33 | 6.95 | 5.21 | 1.74 | 8.81E-07 | 4.62E-05 |
| 1138 | chr13 | 24442451 | 24442601 | + | 0 | NA | intron (NM | intron (NM | 27404 | NM_00128 | 12763 | Mm.8396 | NM_00777 | ENSMUSG | Cmah | - | cytidine m | protein-co | 5.28 | 6.02 | 3.68 | 2.35 | 9.11E-07 | 4.75E-05 |
| 1159 | chr9 | 21328091 | 21328241 | + | 0 | NA | intron (NM | intron (NM | 7447 | NM_00119 | 68682 | Mm.14842 | NM_15280 | ENSMUSG | Slc44a2 | 1110028E | solute carr | protein-co | 6.1 | 6.71 | 5.01 | 1.7 | 9.64E-07 | 4.93E-05 |
| 1180 | chr7 | 63984408 | 63984558 | + | 0 | NA | Intergenic | PB1D9 | SIN | -45568 | NM_02136 | 50794 | Mm.24047 | NM_02136 | ENSMUSG | Klf13 | 0610043C | Kruppel-lik | protein-co | 5.25 | 5.99 | 3.68 | 2.3 | 1.06E-06 | 5.33E-05 |
| 1183 | chr6 | 50437856 | 50438006 | + | 0 | NA | intron (NM | RMER15-ir | 18239 | NM_00116 | 71720 | Mm.44153 | NM_02788 | ENSMUSG | Osbpl3 | 1200014M | oxysterol b | protein-co | 6.54 | 7.14 | 5.5 | 1.64 | 1.07E-06 | 5.36E-05 |
| 1187 | chrX | 1.14E+08 | 1.14E+08 | + | 0 | NA | Intergenic | Intergenic | 569787 | NM_03360 | 93837 | Mm.79760 | NM_03360 | ENSMUSG | Dach2 | 9430028N | dachshunc | protein-co | 7.54 | 7.99 | 6.89 | 1.1 | 1.08E-06 | 5.41E-05 |
| 1216 | chr17 | 35205237 | 35205387 | + | 0 | NA | promoter- | promoter- | 39 | NM_01073 | 16992 | Mm.87787 | NM_01073 | ENSMUSG | Lta | LT | LT-[a] | | lymphotox | protein-co | 6.14 | 6.75 | 5.08 | 1.67 | 1.20E-06 | 5.86E-05 |
| 1243 | chr11 | 51494333 | 51494483 | + | 0 | NA | intron (NM | intron (NM | -90349 | NM_02839 | 72947 | Mm.25687 | NM_02839 | ENSMUSG | Phykpl | 2900006B | 5-phospho | protein-co | 5.18 | 5.93 | 3.49 | 2.44 | 1.36E-06 | 6.47E-05 |
| 1244 | chr1 | 1.93E+08 | 1.93E+08 | + | 0 | NA | Intergenic | Intergenic | 25137 | NM_14481 | 215303 | Mm.40329 | NM_14481 | ENSMUSG | Camk1g | CLICK-III | C | calcium/ca | protein-co | 5.25 | 6.01 | 3.52 | 2.49 | 1.36E-06 | 6.47E-05 |
| 1245 | chr3 | 1.31E+08 | 1.31E+08 | + | 0 | NA | Intergenic | Intergenic | -162815 | NM_01070 | 16842 | Mm.25521 | NM_01070 | ENSMUSG | Lef1 | 3000002B | lymphoid e | protein-co | 5.16 | 5.92 | 3.46 | 2.46 | 1.36E-06 | 6.48E-05 |
| 1248 | chr10 | 43980175 | 43980325 | + | 0 | NA | intron (NM | intron (NM | 24596 | NM_17239 | 11630 | Mm.29208 | NM_17239 | ENSMUSG | Crybg1 | Aim1 | crystallin b | protein-co | 6.63 | 7.19 | 5.72 | 1.46 | 1.38E-06 | 6.58E-05 |
| 1250 | chr3 | 1.53E+08 | 1.53E+08 | + | 0 | NA | intron (NM | intron (NM | 40376 | NM_01202 | 26938 | Mm.40915 | NM_01202 | ENSMUSG | St6galnac5 | AI851940 | | ST6 (alpha | protein-co | 5.84 | 6.48 | 4.65 | 1.83 | 1.39E-06 | 6.58E-05 |
| 1280 | chr16 | 30962627 | 30962777 | + | 0 | NA | intron (NM | intron (NM | 118730 | NM_19862 | 268880 | Mm.28669 | NM_19862 | ENSMUSG | Xxylt1 | AI480653 | xyloside xy | protein-co | 4.79 | 5.6 | 2.83 | 2.76 | 1.54E-06 | 7.14E-05 |
| 1291 | chr3 | 98336004 | 98336154 | + | 0 | NA | intron (NM | intron (NM | 3890 | NM_01696 | 236539 | Mm.16898 | NM_01696 | ENSMUSG | Phgdh | 3-PGDH | 3 | 3-phospho | protein-co | 6.74 | 7.28 | 5.87 | 1.41 | 1.60E-06 | 7.34E-05 |
| 1303 | chr2 | 93132292 | 93132442 | + | 0 | NA | Intergenic | Intergenic | -55181 | NR_15150 | 277414 | Mm.41033 | NM_00102 | ENSMUSG | Trp53i11 | Tp53i11 | transforma | protein-co | 6.84 | 7.38 | 5.97 | 1.41 | 1.67E-06 | 7.59E-05 |
| 1308 | chr7 | 19844919 | 19845069 | + | 0 | NA | Intergenic | Intergenic | -7009 | NM_00127 | 1.01E+08 | Mm.44535 | NM_00127 | ENSMUSG | Gm19345 | - | predicted | protein-co | 6.5 | 7.05 | 5.59 | 1.46 | 1.70E-06 | 7.68E-05 |
| 1324 | chr11 | 52268553 | 52268703 | + | 0 | NA | intron (NM | intron (NM | 13967 | NM_00933 | 21414 | Mm.31630 | NM_00933 | ENSMUSG | Tcf7 | AI465550 | | transcripti | protein-co | 5 | 5.77 | 3.27 | 2.5 | 1.76E-06 | 7.90E-05 |
| 1326 | chr10 | 7976620 | 7976770 | + | 0 | NA | Intergenic | Intergenic | -20572 | NM_13860 | 68652 | Mm.19304 | NM_13860 | ENSMUSG | Tab2 | 1110030N | TGF-beta a | protein-co | 7 | 7.53 | 6.16 | 1.37 | 1.78E-06 | 7.93E-05 |
| 1340 | chr10 | 95313621 | 95313771 | + | 0 | NA | intron (NM | MER21B | L | 10013 | NM_00133 | 12905 | Mm.47373 | NM_00999 | ENSMUSG | Cradd | RAIDD | CASP2 and | protein-co | 6.56 | 7.11 | 5.65 | 1.46 | 1.82E-06 | 8.03E-05 |
| 1348 | chr15 | 36614809 | 36614959 | + | 0 | NA | Intergenic | Intergenic | -5911 | NM_00877 | 18458 | Mm.37157 | NM_00877 | ENSMUSG | Pabpc1 | PABP | Pab | poly(A) bir | protein-co | 4.73 | 5.53 | 2.74 | 2.79 | 1.85E-06 | 8.14E-05 |
| 1350 | chr12 | 85545764 | 85545914 | + | 0 | NA | Intergenic | Intergenic | -53266 | NM_00120 | 81703 | Mm.10356 | NM_03088 | ENSMUSG | Jdp2 | Jundm2 | Ju | Jun dimeri | protein-co | 5.8 | 6.46 | 4.55 | 1.91 | 1.87E-06 | 8.19E-05 |
| 1354 | chr1 | 1.37E+08 | 1.37E+08 | + | 0 | NA | Intergenic | Intergenic | -258373 | NM_03067 | 26424 | Mm.16794 | NM_03067 | ENSMUSG | Nr5a2 | AU020803 | nuclear rec | protein-co | 5.53 | 6.22 | 4.12 | 2.1 | 1.88E-06 | 8.24E-05 |
| 1362 | chr17 | 15107371 | 15107521 | + | 0 | NA | Intergenic | Intergenic | 65806 | NM_00136 | 381062 | Mm.39376 | NM_18313 | ENSMUSG | Ermard | 2210404J1 | ER membr | protein-co | 7.29 | 7.8 | 6.49 | 1.3 | 1.91E-06 | 8.30E-05 |
| 1363 | chr4 | 1.55E+08 | 1.55E+08 | + | 0 | NA | intron (NM | intron (NM | 59140 | NM_00135 | 20481 | Mm.28520 | NM_01138 | ENSMUSG | Ski | 2310012I0 | ski sarcom | protein-co | 6.12 | 6.71 | 5.08 | 1.63 | 1.91E-06 | 8.31E-05 |
| 1366 | chr11 | 57958680 | 57958830 | + | 0 | NA | Intergenic | Intergenic | -50309 | NM_02845 | 73158 | Mm.24884 | NM_02845 | ENSMUSG | Larp1 | 1810024J1 | La ribonuc | protein-co | 5.68 | 6.35 | 4.41 | 1.94 | 1.93E-06 | 8.35E-05 |
| 1374 | chr1 | 1.34E+08 | 1.34E+08 | + | 0 | NA | Intergenic | Intergenic | -4959 | NM_00115 | 240754 | Mm.36534 | NM_17284 | ENSMUSG | Lax1 | A530029E | lymphocyt | protein-co | 5.62 | 6.32 | 4.2 | 2.12 | 1.97E-06 | 8.47E-05 |
| 1382 | chr4 | 46592223 | 46592373 | + | 0 | NA | intron (NM | intron (NM | 9631 | NM_00116 | 107684 | Mm.41631 | NM_17889 | ENSMUSG | Coro2a | 9030208C | coronin, ac | protein-co | 6.43 | 6.99 | 5.49 | 1.5 | 2.04E-06 | 8.73E-05 |
| 1387 | chr3 | 19954030 | 19954180 | + | 0 | NA | Intergenic | Intergenic | -2949 | NM_00127 | 12870 | Mm.13787 | NM_00775 | ENSMUSG | Cp | D3Ertd555 | ceruloplas | protein-co | 4.77 | 5.59 | 2.72 | 2.87 | 2.07E-06 | 8.84E-05 |
| 1396 | chr6 | 1.13E+08 | 1.13E+08 | + | 0 | NA | Intergenic | Intergenic | -57621 | NM_00108 | 18430 | Mm.33331 | NM_00108 | ENSMUSG | Oxtr | OTR | oxytocin re | protein-co | 6.48 | 7.05 | 5.54 | 1.52 | 2.14E-06 | 9.09E-05 |
| 1402 | chr11 | 1.17E+08 | 1.17E+08 | + | 0 | NA | intron (NM | intron (NM | -32139 | NM_00111 | 53860 | Mm.38450 | NM_01738 | ENSMUSG | 9-Sep | MSF1 | Msf | septin 9 | protein-co | 5.16 | 5.89 | 3.58 | 2.31 | 2.19E-06 | 9.23E-05 |
| 1406 | chr17 | 84161286 | 84161436 | + | 0 | NA | Intergenic | Intergenic | 26586 | NM_00100 | 12193 | Mm.25932 | NM_00100 | ENSMUSG | Zfp36l2 | Brf2 | ERF2 | zinc finger | protein-co | 6.12 | 6.75 | 4.97 | 1.78 | 2.20E-06 | 9.27E-05 |
| 1418 | chr13 | 92738874 | 92739024 | + | 0 | NA | Intergenic | Intergenic | 55869 | NM_01158 | 21828 | Mm.20865 | NM_01158 | ENSMUSG | Thbs4 | TSP4 | thrombosp | protein-co | 5.09 | 5.86 | 3.31 | 2.55 | 2.28E-06 | 9.52E-05 |
| 1424 | chr14 | 73404311 | 73404461 | + | 0 | NA | Intergenic | Intergenic | -19097 | NM_00841 | 16432 | Mm.4266 | NM_00841 | ENSMUSG | Itm2b | AI256040 | | integral m | protein-co | 4.93 | 5.7 | 3.13 | 2.58 | 2.32E-06 | 9.63E-05 |
| 1433 | chr10 | 81538735 | 81538885 | + | 0 | NA | intron (NM | intron (NM | 6352 | NM_01030 | 14672 | Mm.26092 | NM_01030 | ENSMUSG | Gna11 | Dsk7 | E430 | guanine nu | protein-co | 5.75 | 6.43 | 4.43 | 2 | 2.41E-06 | 9.96E-05 |
| 1439 | chr7 | 3580028 | 3580178 | + | 0 | NA | Intergenic | Intergenic | 36054 | NM_17563 | 232790 | Mm.35888 | NM_17563 | ENSMUSG | Oscar | mOSCAR | | osteoclast | protein-co | 4.39 | 5.27 | 1.79 | 3.48 | 2.44E-06 | 1.00E-04 |
| 1443 | chr1 | 1.38E+08 | 1.38E+08 | + | 0 | NA | Intergenic | (TC)n | Simp | 29390 | NM_17739 | 338375 | Mm.11442 | NM_17739 | ENSMUSG | Atp6v1g3 | - | ATPase, H- | protein-co | 6.2 | 6.82 | 5.12 | 1.7 | 2.47E-06 | 1.01E-04 |
| 1455 | chr12 | 3338993 | 3339143 | + | 0 | NA | Intergenic | Intergenic | -26064 | NM_00844 | 16570 | Mm.7688 | NM_00844 | ENSMUSG | Kif3c | mKIAA405 | kinesin far | protein-co | 5.2 | 5.95 | 3.51 | 2.44 | 2.55E-06 | 1.04E-04 |
| 1465 | chr15 | 74930651 | 74930801 | + | 0 | NA | Intergenic | MurERV4- | -24325 | NM_00116 | 17069 | Mm.788 | NM_00852 | ENSMUSG | Ly6e | Ly67 | RIG- | lymphocyt | protein-co | 6.45 | 7.01 | 5.52 | 1.49 | 2.64E-06 | 1.07E-04 |
| 1472 | chr10 | 26421270 | 26421420 | + | 0 | NA | Intergenic | RMER6A | L | -46126 | NM_17278 | 237339 | Mm.33843 | NM_17278 | ENSMUSG | L3mbtl3 | AI481284 | | L3MBTL3 h | protein-co | 4.53 | 5.36 | 2.28 | 3.08 | 2.68E-06 | 1.08E-04 |
| 1477 | chr3 | 1.54E+08 | 1.54E+08 | + | 0 | NA | intron (NM | intron (NM | 9231 | NM_00128 | 55993 | Mm.27222 | NM_03187 | ENSMUSG | Msh4 | 4930485C | mutS hom | protein-co | 6.65 | 7.21 | 5.72 | 1.48 | 2.75E-06 | 1.10E-04 |
| 1490 | chr17 | 84166574 | 84166724 | + | 0 | NA | Intergenic | Intergenic | 21298 | NM_00100 | 12193 | Mm.25932 | NM_00100 | ENSMUSG | Zfp36l2 | Brf2 | ERF2 | zinc finger | protein-co | 5.57 | 6.24 | 4.28 | 1.96 | 2.81E-06 | 1.12E-04 |
| 1494 | chr9 | 89885897 | 89886047 | + | 0 | NA | Intergenic | Intergenic | -23803 | NM_01124 | 19417 | Mm.44561 | NM_01124 | ENSMUSG | Rasgrf1 | AI844718 | | RAS protei | protein-co | 6.97 | 7.49 | 6.16 | 1.33 | 2.87E-06 | 1.13E-04 |
| 1495 | chr5 | 1.39E+08 | 1.39E+08 | + | 0 | NA | intron (NM | intron (NM | 11269 | NM_02840 | 73212 | Mm.84294 | NM_02840 | ENSMUSG | 3110082I1 | AW41190 | RIKEN cDN | protein-co | 5.87 | 6.5 | 4.73 | 1.77 | 2.91E-06 | 1.15E-04 |
| 1515 | chr8 | 14986768 | 14986918 | + | 0 | NA | intron (NM | intron (NM | -24182 | NM_02911 | 74901 | Mm.46675 | NM_02911 | ENSMUSG | Kbtbd11 | 2900016B | kelch repe | protein-co | 5.96 | 6.59 | 4.8 | 1.8 | 3.04E-06 | 1.19E-04 |
| 1518 | chr8 | 1.12E+08 | 1.12E+08 | + | 0 | NA | intron (NM | MLT1H1 | L | 25541 | NM_00116 | 170737 | Mm.27157 | NM_13320 | ENSMUSG | Znrf1 | B830022L | zinc and ri | protein-co | 5.34 | 6.05 | 3.88 | 2.17 | 3.05E-06 | 1.19E-04 |
| 1527 | chr6 | 99910037 | 99910187 | + | 0 | NA | Intergenic | Intergenic | -183720 | NM_01576 | 50501 | Mm.87365 | NM_01576 | ENSMUSG | Prok2 | Bv8 | PK2 | I | prokinetici | protein-co | 7.34 | 7.8 | 6.67 | 1.12 | 3.11E-06 | 1.20E-04 |
| 1545 | chr6 | 86605336 | 86605486 | + | 0 | NA | Intergenic | Intergenic | -22763 | NM_02641 | 67855 | Mm.18304 | NM_02641 | ENSMUSG | Asprv1 | 2300003P | aspartic pe | protein-co | 5.88 | 6.53 | 4.69 | 1.84 | 3.25E-06 | 1.25E-04 |
| 1558 | chr18 | 38205115 | 38205265 | + | 0 | NA | intron (NM | intron (NM | 4572 | NM_02935 | 75599 | Mm.36560 | NM_02935 | ENSMUSG | Pcdh1 | 2010005A | protocadh | protein-co | 4.67 | 5.5 | 2.53 | 2.97 | 3.39E-06 | 1.29E-04 |
| 1574 | chrX | 13576759 | 13576909 | + | 0 | NA | intron (NM | intron (NM | -55935 | NM_01182 | 23890 | Mm.48732 | NM_01182 | ENSMUSG | Gpr34 | Lypsr1 | G protein- | protein-co | 5.69 | 6.35 | 4.46 | 1.89 | 3.50E-06 | 1.31E-04 |
| 1586 | chr13 | 92244862 | 92245012 | + | 0 | NA | intron (NM | B3A | SINE | | -109846 | NM_01004 | 13361 | Mm.23695 | NM_01004 | ENSMUSG | Dhfr | 8430436I0 | dihydrofol | protein-co | 5.74 | 6.4 | 4.46 | 1.94 | 3.62E-06 | 1.35E-04 |
| 1602 | chr2 | 1.59E+08 | 1.59E+08 | + | 0 | NA | intron (NM | intron (NM | -34746 | NM_02797 | 71878 | Mm.85162 | NM_02797 | ENSMUSG | Fam83d | 2310007D | family with | protein-co | 5.78 | 6.42 | 4.59 | 1.84 | 3.78E-06 | 1.39E-04 |
| 1607 | chrX | 1.3E+08 | 1.3E+08 | + | 0 | NA | Intergenic | RLTR26B_I | -58891 | NM_17249 | 54004 | Mm.10763 | NM_01739 | ENSMUSG | Diaph2 | Dia3 | Diap | diaphanou | protein-co | 4.95 | 5.71 | 3.27 | 2.44 | 3.83E-06 | 1.41E-04 |
| 1616 | chr17 | 12756069 | 12756219 | + | 0 | NA | intron (NM | intron (NM | 13562 | NM_01051 | 16004 | Mm.26553 | NM_01051 | ENSMUSG | Igf2r | AI661837 | | insulin-like | protein-co | 6.81 | 7.32 | 6.01 | 1.31 | 3.94E-06 | 1.44E-04 |
| 1625 | chr8 | 11526194 | 11526344 | + | 0 | NA | intron (NM | intron (NM | 24502 | NM_02424 | 71941 | Mm.19657 | NM_02424 | ENSMUSG | Cars2 | 2310051N | cysteinyl-t | protein-co | 4.91 | 5.69 | 3.1 | 2.59 | 4.00E-06 | 1.45E-04 |
| 1632 | chr19 | 47654928 | 47655078 | + | 0 | NA | intron (NM | intron (NM | 37039 | NM_00129 | 12821 | Mm.1225 | NM_00773 | ENSMUSG | Col17a1 | BP180 | Bp | collagen, t | protein-co | 5.75 | 6.4 | 4.55 | 1.85 | 4.09E-06 | 1.48E-04 |
| 1634 | chr4 | 94533293 | 94533443 | + | 0 | NA | intron (NM | intron (NM | 23428 | NM_02636 | 67770 | Mm.35170 | NM_02636 | ENSMUSG | Caap1 | 5830433M | caspase ac | protein-co | 5.05 | 5.79 | 3.46 | 2.32 | 4.11E-06 | 1.48E-04 |
| 1673 | chr5 | 43824739 | 43824889 | + | 0 | NA | intron (NM | MLT1I | LTF | 5921 | NM_00976 | 12182 | Mm.24633 | NM_00976 | ENSMUSG | Bst1 | 114/A10 | / | bone marr | protein-co | 5.08 | 5.82 | 3.5 | 2.32 | 4.51E-06 | 1.59E-04 |
| 1679 | chr14 | 73408345 | 73408495 | + | 0 | NA | Intergenic | Lx8 | LINE | I | -23131 | NM_00841 | 16432 | Mm.4266 | NM_00841 | ENSMUSG | Itm2b | AI256040 | | integral m | protein-co | 5.3 | 6 | 3.87 | 2.13 | 4.56E-06 | 1.60E-04 |
| 1685 | chr16 | 95432733 | 95432883 | + | 0 | NA | intron (NM | intron (NM | 26540 | NM_00130 | 13876 | Mm.16453 | NM_13369 | ENSMUSG | Erg | D030036I2 | ETS transc | protein-co | 5.79 | 6.41 | 4.66 | 1.75 | 4.61E-06 | 1.61E-04 |
| 1717 | chr10 | 88584559 | 88584709 | + | 0 | NA | intron (NM | intron (NM | 20518 | NM_00129 | 109272 | Mm.14720 | NM_17541 | ENSMUSG | Mybpc1 | 8030451F | myosin bir | protein-co | 6.91 | 7.42 | 6.1 | 1.31 | 5.01E-06 | 1.72E-04 |
| 1722 | chr19 | 3419414 | 3419564 | + | 0 | NA | Intergenic | Intergenic | -4945 | NM_00132 | 14419 | Mm.4655 | NM_01025 | ENSMUSG | Gal | Galn | galanin an | protein-co | 5.88 | 6.52 | 4.71 | 1.81 | 5.07E-06 | 1.73E-04 |
| 1729 | chr7 | 1.28E+08 | 1.28E+08 | + | 0 | NA | Intergenic | RLTR26_M | -3880 | NM_00840 | 16409 | Mm.26210 | NM_00840 | ENSMUSG | Itgam | CD11b/CD | integrin al | protein-co | 5.68 | 6.34 | 4.39 | 1.95 | 5.17E-06 | 1.76E-04 |
| 1732 | chr2 | 1.32E+08 | 1.32E+08 | + | 0 | NA | intron (NM | intron (NM | 8853 | NM_17544 | 215653 | Mm.20473 | NM_17544 | ENSMUSG | Rassf2 | 3830431H | Ras associ | protein-co | 7.08 | 7.56 | 6.37 | 1.18 | 5.20E-06 | 1.77E-04 |
| 1747 | chr12 | 1.08E+08 | 1.08E+08 | + | 0 | NA | intron (NM | intron (NM | 38587 | NM_00107 | 58208 | Mm.39269 | NM_02139 | ENSMUSG | Bcl11b | 9130430L | B cell leuk | protein-co | 7.28 | 7.72 | 6.63 | 1.09 | 5.38E-06 | 1.81E-04 |
| 1751 | chr15 | 66877872 | 66878022 | + | 0 | NA | Intergenic | Intergenic | -13422 | NM_01886 | 22402 | Mm.10222 | NM_01886 | ENSMUSG | Ccn4 | AW14626 | cellular co | protein-co | 7.58 | 8 | 6.99 | 1.01 | 5.44E-06 | 1.83E-04 |
| 1764 | chr6 | 1.47E+08 | 1.47E+08 | + | 0 | NA | Intergenic | MTEa | LTR | -12688 | NM_01058 | 16439 | Mm.7800 | NM_01058 | ENSMUSG | Itpr2 | AI649341 | | inositol 1,4 | protein-co | 6.56 | 7.13 | 5.61 | 1.51 | 5.61E-06 | 1.87E-04 |
| 1768 | chr14 | 63377447 | 63377597 | + | 0 | NA | intron (NM | intron (NM | 39665 | NM_00754 | 12143 | Mm.3962 | NM_00754 | ENSMUSG | Blk | - | B lymphoi | protein-co | 7.27 | 7.71 | 6.62 | 1.09 | 5.67E-06 | 1.89E-04 |
| 1769 | chr8 | 1.06E+08 | 1.06E+08 | + | 0 | NA | intron (NM | intron (NM | -5833 | NM_02803 | 71986 | Mm.30751 | NM_02803 | ENSMUSG | Ddx28 | 2410004K | DEAD (Asp | protein-co | 5.8 | 6.43 | 4.67 | 1.76 | 5.67E-06 | 1.89E-04 |
| 1793 | chr8 | 11506957 | 11507107 | + | 0 | NA | intron (NM | intron (NM | 9464 | NM_02699 | 69225 | Mm.64911 | NM_02699 | ENSMUSG | Naxd | 0710008K | NAD(P)HX | protein-co | 6.25 | 6.85 | 5.19 | 1.67 | 6.03E-06 | 1.98E-04 |
| 1809 | chr6 | 1.13E+08 | 1.13E+08 | + | 0 | NA | Intergenic | Intergenic | -87500 | NM_00108 | 18430 | Mm.33331 | NM_00108 | ENSMUSG | Oxtr | OTR | oxytocin re | protein-co | 5.94 | 6.54 | 4.87 | 1.67 | 6.28E-06 | 2.04E-04 |
| 1811 | chr2 | 1.57E+08 | 1.57E+08 | + | 0 | NA | promoter- | promoter- | 41 | NM_02998 | 77799 | Mm.29264 | NM_02998 | ENSMUSG | Sla2 | A930009E | Src-like-ad | protein-co | 7.5 | 7.93 | 6.88 | 1.05 | 6.31E-06 | 2.05E-04 |
| 1818 | chr4 | 95303496 | 95303646 | + | 0 | NA | Intergenic | Intergenic | -251349 | NM_01059 | 16476 | Mm.27507 | NM_01059 | ENSMUSG | Jun | AP-1 | Junc | jun proto-c | protein-co | 6.53 | 7.06 | 5.67 | 1.39 | 6.45E-06 | 2.08E-04 |
| 1830 | chr11 | 46380337 | 46380487 | + | 0 | NA | intron (NM | Lx9 | LINE | I | 9103 | NM_01058 | 16428 | Mm.33992 | NM_01058 | ENSMUSG | Itk | Emt | Tcsk | I | IL2 inducib | protein-co | 5.5 | 6.16 | 4.26 | 1.9 | 6.71E-06 | 2.15E-04 |
| 1862 | chr10 | 60381133 | 60381283 | + | 0 | NA | intron (NR | intron (NR | -18518 | NM_00116 | 1E+08 | Mm.44413 | NM_00116 | ENSMUSG | Gm17455 | - | predicted | protein-co | 5.67 | 6.31 | 4.52 | 1.79 | 7.23E-06 | 2.28E-04 |
| 1876 | chr3 | 89871126 | 89871276 | + | 0 | NA | exon (NM_ | exon (NM_ | 39831 | NM_17253 | 214547 | Mm.225 | NM_17253 | ENSMUSG | She | 9430022A | src homolo | protein-co | 6.67 | 7.24 | 5.71 | 1.52 | 7.42E-06 | 2.32E-04 |
| 1880 | chr14 | 1.22E+08 | 1.22E+08 | + | 0 | NA | Intergenic | Intergenic | 31865 | NM_00103 | 223262 | Mm.29763 | NM_00103 | ENSMUSG | Timm8a2 | - | transloca | protein-co | 5.79 | 6.4 | 4.69 | 1.71 | 7.50E-06 | 2.34E-04 |
| 1883 | chr5 | 1.34E+08 | 1.34E+08 | + | 0 | NA | intron (NM | intron (NM | 15006 | NM_00128 | 17969 | Mm.42529 | NM_01087 | ENSMUSG | Ncf1 | NCF-47K | | neutrophil | protein-co | 6.15 | 6.72 | 5.19 | 1.53 | 7.55E-06 | 2.35E-04 |
| 1886 | chr8 | 1.2E+08 | 1.2E+08 | + | 0 | NA | 3' UTR (NM | 3' UTR (NM | 51090 | NM_13390 | 102193 | Mm.24007 | NM_13390 | ENSMUSG | Zdhhc7 | AL024087 | zinc finger | protein-co | 6.24 | 6.79 | 5.32 | 1.48 | 7.61E-06 | 2.37E-04 |
| 1889 | chr4 | 47187510 | 47187660 | + | 0 | NA | Intergenic | Intergenic | -20427 | NM_00992 | 12819 | Mm.23354 | NM_00992 | ENSMUSG | Col15a1 | - | collagen, t | protein-co | 5.2 | 5.9 | 3.78 | 2.12 | 7.62E-06 | 2.37E-04 |
| 1900 | chr8 | 1.2E+08 | 1.2E+08 | + | 0 | NA | intron (NM | intron (NM | 40287 | NM_03020 | 78892 | Mm.26468 | NM_03020 | ENSMUSG | Crispld2 | 1810049K | cysteine-ri | protein-co | 5.61 | 6.25 | 4.4 | 1.85 | 7.74E-06 | 2.39E-04 |
| 1909 | chr6 | 1.25E+08 | 1.25E+08 | + | 0 | NA | Intergenic | Intergenic | -4705 | NM_00104 | 21940 | Mm.36771 | NM_00103 | ENSMUSG | Cd27 | S152 | Tnfr | CD27 antig | protein-co | 5.74 | 6.37 | 4.59 | 1.78 | 7.96E-06 | 2.44E-04 |
| 1916 | chr15 | 25303469 | 25303619 | + | 0 | NA | Intergenic | Intergenic | 110220 | NM_02739 | 70350 | Mm.29586 | NM_02739 | ENSMUSG | Basp1 | 2610024P | brain abur | protein-co | 6.19 | 6.76 | 5.24 | 1.52 | 8.08E-06 | 2.47E-04 |
| 1919 | chr2 | 1.02E+08 | 1.02E+08 | + | 0 | NA | Intergenic | Intergenic | -72988 | NM_17888 | 241576 | Mm.48974 | NM_17888 | ENSMUSG | Ldlrad3 | 6430500P | low densit | protein-co | 5.59 | 6.24 | 4.37 | 1.87 | 8.15E-06 | 2.49E-04 |
| 1927 | chr9 | 21327699 | 21327849 | + | 0 | NA | intron (NM | B3 | SINE | E | 7055 | NM_00119 | 68682 | Mm.14842 | NM_15280 | ENSMUSG | Slc44a2 | 1110028E | solute carr | protein-co | 6.22 | 6.81 | 5.22 | 1.58 | 8.38E-06 | 2.55E-04 |
| 1952 | chr10 | 45020436 | 45020586 | + | 0 | NA | Intergenic | Intergenic | -46705 | NM_01115 | 19072 | Mm.37294 | NM_01115 | ENSMUSG | Prep | AI047692 | | prolyl endo | protein-co | 7.61 | 8.04 | 6.98 | 1.06 | 8.84E-06 | 2.65E-04 |
| 1953 | chr2 | 1.63E+08 | 1.63E+08 | + | 0 | NA | Intergenic | MTE2b | LT | -39574 | NM_00136 | 269389 | Mm.20770 | NM_00109 | ENSMUSG | Tox2 | AI851523 | | TOX high r | protein-co | 6.55 | 7.06 | 5.73 | 1.33 | 8.86E-06 | 2.66E-04 |
| 1956 | chr4 | 1.34E+08 | 1.34E+08 | + | 0 | NA | intron (NM | intron (NM | 518 | NM_14583 | 83557 | Mm.30256 | NM_14583 | ENSMUSG | Lin28a | AL024421 | | lin-28 hom | protein-co | 4.53 | 5.34 | 2.51 | 2.83 | 8.92E-06 | 2.67E-04 |
| 1981 | chr11 | 60099935 | 60100085 | + | 0 | NA | Intergenic | Intergenic | -5003 | NM_00902 | 19377 | Mm.29636 | NM_00902 | ENSMUSG | Rai1 | Gt1 | retinoic ac | protein-co | 4.79 | 5.55 | 3.09 | 2.46 | 9.58E-06 | 2.83E-04 |
| 1989 | chr19 | 53891042 | 53891192 | + | 0 | NA | Intergenic | Intergenic | -1114 | NM_01105 | 18569 | Mm.1605 | NM_01105 | ENSMUSG | Pdcd4 | D19Ucla1 | | programm | protein-co | 6.43 | 6.96 | 5.58 | 1.38 | 9.72E-06 | 2.86E-04 |
| 1995 | chr9 | 63742423 | 63742573 | + | 0 | NA | intron (NM | intron (NM | 15496 | NM_01676 | 17127 | Mm.7320 | NM_01676 | ENSMUSG | Smad3 | AU022421 | SMAD fam | protein-co | 5.78 | 6.42 | 4.58 | 1.84 | 9.85E-06 | 2.89E-04 |
| 2000 | chr7 | 4948515 | 4948665 | + | 0 | NA | Intergenic | Intergenic | 15816 | NM_00114 | 381836 | | NM_00114 | ENSMUSG | Sbk2 | Gm1079 | S | SH3-bindir | protein-co | 5.4 | 6.08 | 4.07 | 2.01 | 1.00E-05 | 2.93E-04 |
| 2010 | chr1 | 91352547 | 91352697 | + | 0 | NA | intron (NM | intron (NM | 1549 | NM_02755 | 70788 | Mm.68617 | NM_02755 | ENSMUSG | Klhl30 | 4631423F | kelch-like | protein-co | 5.4 | 6.07 | 4.11 | 1.96 | 1.02E-05 | 2.96E-04 |
| 2011 | chr1 | 86301442 | 86301592 | + | 0 | NA | Intergenic | Intergenic | -1704 | NM_14522 | 227327 | Mm.86467 | NM_14522 | ENSMUSG | B3gnt7 | C330001H | UDP-GlcNA | protein-co | 5.83 | 6.44 | 4.74 | 1.69 | 1.02E-05 | 2.97E-04 |
| 2013 | chr7 | 80718611 | 80718761 | + | 0 | NA | intron (NM | L1MB2 | LII | -29809 | NM_17380 | 70461 | Mm.39217 | NM_17380 | ENSMUSG | Crtc3 | 2610312F | CREB regu | protein-co | 6.72 | 7.21 | 5.96 | 1.25 | 1.03E-05 | 2.98E-04 |
| 2019 | chr5 | 9014918 | 9015068 | + | 0 | NA | Intergenic | Intergenic | -17847 | NM_02373 | 74114 | Mm.28197 | NM_02373 | ENSMUSG | Crot | 1200003H | carnitine C | protein-co | 5.06 | 5.77 | 3.58 | 2.18 | 1.05E-05 | 3.03E-04 |
| 2038 | chr6 | 1.28E+08 | 1.28E+08 | + | 0 | NA | intron (NM | intron (NM | 4838 | NM_00136 | 381812 | Mm.29609 | NM_00103 | ENSMUSG | Cracr2a | Efcab4b | G | calcium re | protein-co | 6.78 | 7.33 | 5.87 | 1.46 | 1.09E-05 | 3.13E-04 |

| | | | | | | | | | | | | | | | | | | | | | | | | |
|---|---|---|---|---|---|---|---|---|---|---|---|---|---|---|---|---|---|---|---|---|---|---|---|---|
| 2043 | chr4 | 57958223 | 57958373 | + | 0 | NA | Intergenic | Intergenic | -1887 | NM_0116 | 22166 | Mm.26061 | NM_0116 | ENSMUSG | Txn1 | ADF\|AW5 | thioredoxi | protein-co | 6.47 | 7.04 | 5.54 | 1.5 | 1.12E-05 | 3.19E-04 |
| 2049 | chr5 | 1.22E+08 | 1.22E+08 | + | 0 | NA | Intergenic | Intergenic | -25150 | NM_0010 | 74096 | Mm.28804 | NM_0287 | ENSMUSG | Hvcn1 | 0610039P | hydrogen | protein-co | 4.97 | 5.7 | 3.46 | 2.24 | 1.12E-05 | 3.19E-04 |
| 2051 | chr2 | 1.63E+08 | 1.63E+08 | + | 0 | NA | intron (NM | intron (NM | 23652 | NM_0013 | 269389 | Mm.20770 | NM_0010 | ENSMUSG | Tox2 | AI851523 | TOX high r | protein-co | 5.91 | 6.51 | 4.86 | 1.65 | 1.12E-05 | 3.20E-04 |
| 2052 | chr15 | 1.01E+08 | 1.01E+08 | + | 0 | NA | intron (NM | ORR1E\|LT | 4901 | NM_0168 | 53622 | Mm.34793 | NM_016879 | | Krt85 | AA589597 | keratin 85 | protein-co | 5.85 | 6.45 | 4.81 | 1.63 | 1.13E-05 | 3.21E-04 |
| 2055 | chr7 | 16362188 | 16362338 | + | 0 | NA | intron (NM | intron (NM | 25633 | NM_0012 | 56459 | Mm.25853 | NM_0197 | ENSMUSG | Sae1 | 2400010M | SUMO1 ac | protein-co | 5.59 | 6.26 | 4.28 | 1.97 | 1.13E-05 | 3.22E-04 |
| 2064 | chr9 | 1.18E+08 | 1.18E+08 | + | 0 | NA | Intergenic | Intergenic | -177007 | NM_0137 | 27215 | Mm.92705 | NM_0137 | ENSMUSG | Azi2 | AA410145 | 5-azacytidi | protein-co | 5.51 | 6.17 | 4.27 | 1.9 | 1.16E-05 | 3.29E-04 |
| 2086 | chr1 | 92069595 | 92069745 | + | 0 | NA | intron (NM | intron (NM | 110671 | NM_2072 | 208727 | Mm.31856 | NM_2072 | ENSMUSG | Hdac4 | 4932408F | histone de | protein-co | 5.16 | 5.87 | 3.72 | 2.15 | 1.20E-05 | 3.37E-04 |
| 2101 | chr2 | 17702893 | 17703043 | + | 0 | NA | intron (NM | intron (NM | 28166 | NM_0287 | 74103 | Mm.12029 | NM_0287 | ENSMUSG | Nebl | 1200007O | nebulette | protein-co | 7.17 | 7.61 | 6.53 | 1.07 | 1.24E-05 | 3.44E-04 |
| 2124 | chr2 | 1.64E+08 | 1.64E+08 | + | 0 | NA | TTS (NM_( | TTS (NM_( | 23959 | NM_0012 | 11486 | Mm.388 | NM_0073 | ENSMUSG | Ada | - | adenosine | protein-co | 5.66 | 6.31 | 4.46 | 1.85 | 1.30E-05 | 3.58E-04 |
| 2126 | chr4 | 1.45E+08 | 1.45E+08 | + | 0 | NA | intron (NM | intron (NM | 60563 | NM_0012 | 230895 | Mm.24031 | NM_0010 | ENSMUSG | Vps13d | - | vacuolar p | protein-co | 5.11 | 5.83 | 3.61 | 2.22 | 1.30E-05 | 3.58E-04 |
| 2135 | chr10 | 1.27E+08 | 1.27E+08 | + | 0 | NA | intron (NM | intron (NM | -19274 | NM_0013 | 117150 | Mm.22682 | NM_0540 | ENSMUSG | Pip4k2c | Pip5k2c | phosphati | protein-co | 5.92 | 6.53 | 4.85 | 1.68 | 1.33E-05 | 3.63E-04 |
| 2136 | chr6 | 42385771 | 42385921 | + | 0 | NA | Intergenic | Intergenic | -12578 | NM_0235 | 13835 | Mm.13333 | NM_0235 | ENSMUSG | Epha1 | 5730453L | Eph recept | protein-co | 4.57 | 5.41 | 2.25 | 3.16 | 1.33E-05 | 3.63E-04 |
| 2137 | chr2 | 90030413 | 90030563 | + | 0 | NA | Intergenic | Lx2B2\|LIN | -6433 | NM_1463 | 258340 | Mm.37744 | NM_1463 | ENSMUSG | Olfr1265 | MOR228-2 | olfactory r | protein-co | 6.75 | 7.24 | 6.01 | 1.23 | 1.33E-05 | 3.64E-04 |
| 2156 | chr19 | 3441283 | 3441433 | + | 0 | NA | Intergenic | Intergenic | -26814 | NM_0013 | 14419 | Mm.4655 | NM_0102 | ENSMUSG | Gal | Galn | galanin an | protein-co | 5.66 | 6.31 | 4.43 | 1.88 | 1.40E-05 | 3.78E-04 |
| 2159 | chr11 | 95367188 | 95367338 | + | 0 | NA | intron (NM | intron (NM | -17593 | NM_0013 | 110172 | Mm.4593 | NM_0167 | ENSMUSG | Slc35b1 | UGTrel1\| | solute carr | protein-co | 5.13 | 5.83 | 3.72 | 2.11 | 1.41E-05 | 3.80E-04 |
| 2182 | chr13 | 19272896 | 19273046 | + | 0 | NA | Intergenic | Intergenic | 122842 | NM_0013 | 76205 | Mm.41940 | NM_0242 | ENSMUSG | Stard3nl | 0610035N | STARD3 N- | protein-co | 5.87 | 6.47 | 4.82 | 1.65 | 1.49E-05 | 3.97E-04 |
| 2187 | chr17 | 31835773 | 31835923 | + | 0 | NA | Intergenic | Intergenic | 19956 | NM_0108 | 17691 | Mm.29094 | NM_0108 | ENSMUSG | Sik1 | Hrt-20\|Ms | salt induci | protein-co | 6.48 | 7 | 5.66 | 1.34 | 1.51E-05 | 4.01E-04 |
| 2202 | chr4 | 1.03E+08 | 1.03E+08 | + | 0 | NA | Intergenic | MER102b\| | 45674 | NM_0011 | 18578 | Mm.20181 | NM_0198 | ENSMUSG | Pde4b | Dpde4\|R7 | phosphodi | protein-co | 6.72 | 7.23 | 5.94 | 1.29 | 1.56E-05 | 4.12E-04 |
| 2208 | chr8 | 1.17E+08 | 1.17E+08 | + | 0 | NA | intron (NM | intron (NM | 97535 | NM_0294 | 75796 | Mm.15957 | NM_0294 | ENSMUSG | Cdyl2 | 1700029M | chromodo | protein-co | 5.57 | 6.24 | 4.27 | 1.96 | 1.58E-05 | 4.16E-04 |
| 2216 | chr9 | 20636398 | 20636548 | + | 0 | NA | Intergenic | Intergenic | -6244 | NM_0013 | 66177 | Mm.39016 | NM_0254 | ENSMUSG | Ubl5 | 1110030M | ubiquitin-l | protein-co | 4.73 | 5.49 | 3.02 | 2.47 | 1.61E-05 | 4.24E-04 |
| 2230 | chr19 | 46518572 | 46518722 | + | 0 | NA | Intergenic | Intergenic | 16999 | NM_0531 | 93679 | Mm.39217 | NM_0531 | ENSMUSG | Trim8 | AA408830 | tripartite r | protein-co | 5.76 | 6.37 | 4.67 | 1.69 | 1.64E-05 | 4.28E-04 |
| 2232 | chr18 | 38464186 | 38464336 | + | 0 | NA | 3' UTR (NM | 3' UTR (NM | 45354 | NM_0229 | 65113 | Mm.10249 | NM_0229 | ENSMUSG | Ndfip1 | 0610010M | Nedd4 fam | protein-co | 4.48 | 5.3 | 2.35 | 2.96 | 1.64E-05 | 4.28E-04 |
| 2258 | chr18 | 70643632 | 70643782 | + | 0 | NA | Intergenic | Intergenic | 75415 | NM_0013 | 17191 | Mm.322 | NM_0107 | ENSMUSG | Mbd2 | MBD2a | methyl-Cp | protein-co | 6.81 | 7.33 | 6 | 1.33 | 1.72E-05 | 4.42E-04 |
| 2272 | chr9 | 66560398 | 66560548 | + | 0 | NA | intron (NM | intron (NM | 32642 | NM_0013 | 235441 | Mm.32465 | NM_1449 | ENSMUSG | Usp3 | AA409661 | ubiquitin s | protein-co | 5.72 | 6.34 | 4.62 | 1.72 | 1.74E-05 | 4.45E-04 |
| 2276 | chr1 | 1.66E+08 | 1.66E+08 | + | 0 | NA | Intergenic | Intergenic | -1117 | NM_0010 | 226594 | Mm.26555 | NM_1785 | ENSMUSG | Rcsd1 | A430105K | RCSD dom | protein-co | 6.87 | 7.34 | 6.17 | 1.16 | 1.74E-05 | 4.46E-04 |
| 2279 | chr6 | 1.2E+08 | 1.2E+08 | + | 0 | NA | intron (NM | intron (NM | 10417 | NM_0083 | 16172 | Mm.4481 | NM_0083 | ENSMUSG | Il17ra | AW53815 | interleukir | protein-co | 6.61 | 7.1 | 5.85 | 1.25 | 1.75E-05 | 4.48E-04 |
| 2298 | chr1 | 1.34E+08 | 1.34E+08 | + | 0 | NA | promoter- | promoter- | -46 | NM_0011 | 240754 | Mm.36534 | NM_1728 | ENSMUSG | Lax1 | A530029E | lymphocyt | protein-co | 6.77 | 7.26 | 6.03 | 1.23 | 1.82E-05 | 4.59E-04 |
| 2296 | chr9 | 1.16E+08 | 1.16E+08 | + | 0 | NA | intron (NM | intron (NM | 36462 | NM_0295 | 21813 | Mm.17234 | NM_0093 | ENSMUSG | Tgfbr2 | 1110020H | transformi | protein-co | 5.94 | 6.58 | 4.79 | 1.79 | 1.82E-05 | 4.59E-04 |
| 2305 | chr2 | 27297160 | 27297310 | + | 0 | NA | intron (NR | intron (NR | -49932 | NM_1386 | 192166 | Mm.27846 | NM_1386 | ENSMUSG | Sardh | - | sarcosine | protein-co | 4.46 | 5.26 | 2.52 | 2.74 | 1.86E-05 | 4.70E-04 |
| 2328 | chr5 | 1.18E+08 | 1.18E+08 | + | 0 | NA | intron (NM | intron (NM | 21365 | NM_0213 | 57816 | Mm.27328 | NM_0213 | ENSMUSG | Tesc | 1010001A | tescalcin | protein-co | 7 | 7.45 | 6.34 | 1.11 | 1.97E-05 | 4.91E-04 |
| 2340 | chr3 | 9865852 | 9866002 | + | 0 | NA | Intergenic | Intergenic | -32248 | NM_0011 | 94212 | Mm.23897 | NM_0531 | ENSMUSG | Pag1 | Cbp\|F730 | phosphopr | protein-co | 6.62 | 7.12 | 5.86 | 1.26 | 2.00E-05 | 4.96E-04 |
| 2341 | chr16 | 10836440 | 10836590 | + | 0 | NA | intron (NM | intron (NM | 1456 | NM_0011 | 223970 | Mm.32577 | NM_0010 | ENSMUSG | Rmi2 | A630055G | RecQ med | protein-co | 6.46 | 6.99 | 5.6 | 1.4 | 2.00E-05 | 4.96E-04 |
| 2354 | chr13 | 1.2E+08 | 1.2E+08 | + | 0 | NA | promoter- | promoter- | -234 | NM_0010 | 625360 | Mm.32750 | NM_0010 | ENSMUSG | BC147527 | - | cDNA sequ | protein-co | 5.83 | 6.44 | 4.78 | 1.66 | 2.07E-05 | 5.10E-04 |
| 2360 | chr3 | 31082934 | 31083084 | + | 0 | NA | Intergenic | PB1\|SINE\| | -12049 | NM_0113 | 20482 | Mm.15406 | NM_0113 | ENSMUSG | Skil | Skir\|SnoN | SKI-like | protein-co | 4.4 | 5.2 | 2.41 | 2.79 | 2.10E-05 | 5.17E-04 |
| 2365 | chr1 | 1.62E+08 | 1.62E+08 | + | 0 | NA | Intergenic | Intergenic | -16294 | NM_0101 | 14103 | Mm.3355 | NM_0101 | ENSMUSG | Fasl | APT1LG1\| | Fas ligand | protein-co | 6.75 | 7.24 | 6.02 | 1.22 | 2.12E-05 | 5.21E-04 |
| 2382 | chr3 | 97831680 | 97831830 | + | 0 | NA | intron (NM | intron (NM | 56952 | NM_0012 | 83679 | Mm.12984 | NM_0314 | ENSMUSG | Pde4dip | 4732458A | phosphodi | protein-co | 5.29 | 5.95 | 4.01 | 1.94 | 2.20E-05 | 5.35E-04 |
| 2389 | chr2 | 1.67E+08 | 1.67E+08 | + | 0 | NA | intron (NM | intron (NM | 40541 | NM_0084 | 16500 | Mm.20534 | NM_0084 | ENSMUSG | Kcnb1 | Kcr1-1\|Kv | potassium | protein-co | 4.72 | 5.47 | 3.09 | 2.38 | 2.24E-05 | 5.43E-04 |
| 2406 | chr7 | 80204484 | 80204634 | + | 0 | NA | intron (NM | intron (NM | 6118 | NM_0013 | 20352 | Mm.27590 | NM_0136 | ENSMUSG | Sema4b | SemC\|Sem | sema dom | protein-co | 6.64 | 7.13 | 5.89 | 1.24 | 2.29E-05 | 5.52E-04 |
| 2416 | chr10 | 81434616 | 81434766 | + | 0 | NA | Intergenic | PB1D11\|S | -7518 | NM_0086 | 18029 | Mm.42693 | NM_0086 | ENSMUSG | Nfic | 1110019L | nuclear fac | protein-co | 6.68 | 7.17 | 5.93 | 1.24 | 2.33E-05 | 5.58E-04 |
| 2427 | chr2 | 1.73E+08 | 1.73E+08 | + | 0 | NA | Intergenic | MLTR12\|L | -90633 | NM_0281 | 72221 | Mm.46132 | NM_0281 | ENSMUSG | 1700021F | - | RIKEN cDN | protein-co | 5.14 | 5.85 | 3.71 | 2.14 | 2.36E-05 | 5.64E-04 |
| 2440 | chr5 | 1.22E+08 | 1.22E+08 | + | 0 | NA | intron (NM | B3A\|SINE\| | 24694 | NM_0287 | 74096 | Mm.28804 | NM_0287 | ENSMUSG | Hvcn1 | 0610039P | hydrogen | protein-co | 5.82 | 6.43 | 4.75 | 1.68 | 2.44E-05 | 5.78E-04 |
| 2443 | chr19 | 6399593 | 6399743 | + | 0 | NA | promoter- | promoter- | -207 | NM_0013 | 19395 | Mm.77017 | NM_0112 | ENSMUSG | Rasgrp2 | CDC25L\|C | RAS, guany | protein-co | 6.68 | 7.21 | 5.86 | 1.35 | 2.45E-05 | 5.81E-04 |
| 2444 | chr9 | 1.19E+08 | 1.19E+08 | + | 0 | NA | intron (NM | intron (NM | -41775 | NM_0011 | 104099 | Mm.33552 | NM_1337 | ENSMUSG | Itga9 | (alpha)9\|2 | integrin al | protein-co | 4.92 | 5.63 | 3.46 | 2.17 | 2.45E-05 | 5.81E-04 |
| 2461 | chr17 | 88142909 | 88143059 | + | 0 | NA | Intergenic | Intergenic | 17472 | NM_0011 | 225058 | Mm.31205 | NM_001190356 | | Gm4832 | EG225058 | predicted | protein-co | 4.21 | 5.05 | 1.93 | 3.12 | 2.52E-05 | 5.94E-04 |
| 2480 | chr2 | 1.32E+08 | 1.32E+08 | + | 0 | NA | intron (NM | B1_Mus2\| | -9810 | NM_0012 | 110911 | Mm.28450 | NM_1386 | ENSMUSG | Cds2 | 5730450N | CDP-diacyl | protein-co | 4.58 | 5.35 | 2.82 | 2.53 | 2.59E-05 | 6.05E-04 |
| 2487 | chr9 | 57956982 | 57957132 | + | 0 | NA | intron (NM | intron (NM | 16945 | NM_0113 | 20361 | Mm.33518 | NM_0113 | ENSMUSG | Sema7a | 2900057C | sema dom | protein-co | 5.5 | 6.13 | 4.35 | 1.78 | 2.62E-05 | 6.09E-04 |
| 2509 | chr3 | 38440624 | 38440774 | + | 0 | NA | Intergenic | MIR\|SINE\| | 44117 | NM_0011 | 99696 | Mm.29093 | NM_0010 | ENSMUSG | Ankrd50 | AI662170\| | ankyrin rep | protein-co | 4.77 | 5.51 | 3.18 | 2.33 | 2.70E-05 | 6.23E-04 |
| 2515 | chr4 | 1.35E+08 | 1.35E+08 | + | 0 | NA | intron (NM | intron (NM | -15716 | NM_0013 | 12399 | Mm.37889 | NM_0019 | ENSMUSG | Runx3 | AML2\|Cbf | runt relate | protein-co | 5.84 | 6.45 | 4.79 | 1.66 | 2.71E-05 | 6.24E-04 |
| 2517 | chr2 | 17390998 | 17391148 | + | 0 | NA | intron (NM | intron (NM | 71783 | NM_0013 | 74103 | Mm.12029 | NM_0287 | ENSMUSG | Nebl | 1200007O | nebulette | protein-co | 4.8 | 5.53 | 3.25 | 2.28 | 2.74E-05 | 6.30E-04 |
| 2519 | chr17 | 80112993 | 80113143 | + | 0 | NA | Intergenic | Intergenic | -14403 | NM_1769 | 319625 | Mm.29098 | NM_1769 | ENSMUSG | Galm | A530057M | galactose | protein-co | 7.08 | 7.53 | 6.43 | 1.09 | 2.75E-05 | 6.31E-04 |
| 2528 | chr8 | 1.06E+08 | 1.06E+08 | + | 0 | NA | promoter- | promoter- | -148 | NM_0013 | 234695 | Mm.47164 | NM_0010 | ENSMUSG | Carmil2 | CG1399-Pl | capping pr | protein-co | 7.33 | 7.75 | 6.74 | 1.01 | 2.79E-05 | 6.37E-04 |
| 2538 | chr12 | 70102532 | 70102682 | + | 0 | NA | intron (NM | intron (NM | 247 | NM_0010 | 18080 | Mm.24404 | NM_0086 | ENSMUSG | Nin | 3110068G | ninein | protein-co | 5.56 | 6.2 | 4.38 | 1.82 | 2.83E-05 | 6.44E-04 |
| 2590 | chr2 | 71356683 | 71356833 | + | 0 | NA | intron (NM | intron (NM | 10997 | NM_1724 | 78830 | Mm.30928 | NM_1724 | ENSMUSG | Slc25a12 | 2610002D | solute carr | protein-co | 4.9 | 5.64 | 3.3 | 2.34 | 3.09E-05 | 6.89E-04 |
| 2598 | chr1 | 52019757 | 52019907 | + | 0 | NA | intron (NM | intron (NM | 11592 | NM_0114 | 20849 | Mm.1550 | NM_0114 | ENSMUSG | Stat4 | - | signal tran | protein-co | 5.66 | 6.26 | 4.6 | 1.66 | 3.12E-05 | 6.92E-04 |
| 2619 | chr6 | 1.16E+08 | 1.16E+08 | + | 0 | NA | Intergenic | Intergenic | -25263 | NM_1990 | 381802 | Mm.29120 | NM_1990 | ENSMUSG | Tsen2 | AU067695 | tRNA splic | protein-co | 6.97 | 7.42 | 6.29 | 1.13 | 3.27E-05 | 7.19E-04 |
| 2630 | chr2 | 1.61E+08 | 1.61E+08 | + | 0 | NA | intron (NR | intron (NR | 55127 | NM_1733 | 71389 | Mm.12273 | NM_1733 | ENSMUSG | Chd6 | 5430439G | chromodo | protein-co | 4.21 | 5.04 | 2.09 | 2.94 | 3.32E-05 | 7.27E-04 |
| 2634 | chr7 | 16201267 | 16201417 | + | 0 | NA | intron (NM | intron (NM | 20651 | NM_0012 | 71723 | Mm.75259 | NM_0278 | ENSMUSG | Dhx34 | 1200013B | DEAH (Asp | protein-co | 6.21 | 6.76 | 5.33 | 1.43 | 3.34E-05 | 7.30E-04 |
| 2638 | chr1 | 1.27E+08 | 1.27E+08 | + | 0 | NA | intron (NM | intron (NM | 103555 | NM_1451 | 107895 | Mm.21468 | NM_1451 | ENSMUSG | Mgat5 | 4930471A | mannoside | protein-co | 6.37 | 6.88 | 5.58 | 1.3 | 3.39E-05 | 7.39E-04 |
| 2645 | chr10 | 5384256 | 5384406 | + | 0 | NA | intron (NM | intron (NM | 103983 | NM_0013 | 64009 | Mm.33162 | NM_0220 | ENSMUSG | Syne1 | 8B\|A3300 | spectrin re | protein-co | 5.74 | 6.34 | 4.69 | 1.64 | 3.44E-05 | 7.49E-04 |
| 2650 | chr3 | 19487877 | 19488027 | + | 0 | NA | Intergenic | ORR1E\|LT | -20643 | NM_0011 | 66326 | Mm.45615 | NM_0254 | ENSMUSG | Dnajc5b | 1700008A | DnaJ heat | protein-co | 4.91 | 5.62 | 3.45 | 2.17 | 3.46E-05 | 7.53E-04 |
| 2663 | chr14 | 73105494 | 73105644 | + | 0 | NA | Intergenic | Intergenic | -36941 | NM_1340 | 105670 | Mm.28006 | NM_1340 | ENSMUSG | Rcbtb2 | 2610028E | regulator | protein-co | 6.18 | 6.73 | 5.28 | 1.45 | 3.53E-05 | 7.64E-04 |
| 2667 | chr2 | 26936532 | 26936682 | + | 0 | NA | intron (NM | intron (NM | 2538 | NM_1986 | 279029 | Mm.33007 | NM_1986 | ENSMUSG | Stkld1 | Gm711\|Sg | serine/thre | protein-co | 6.21 | 6.74 | 5.37 | 1.37 | 3.56E-05 | 7.68E-04 |
| 2672 | chr11 | 1.15E+08 | 1.15E+08 | + | 0 | NA | intron (NM | intron (NM | -39057 | NM_0013 | 66176 | Mm.86823 | NM_0254 | ENSMUSG | Nat9 | 1110028N | N-acetyltra | protein-co | 5.49 | 6.12 | 4.35 | 1.77 | 3.57E-05 | 7.69E-04 |
| 2670 | chr19 | 44578144 | 44578294 | + | 0 | NA | Intergenic | MIR\|SINE\| | 15365 | NM_1769 | 319594 | Mm.14527 | NM_1769 | ENSMUSG | Hif1an | 2310046M | hypoxia-in | protein-co | 4.78 | 5.52 | 3.18 | 2.33 | 3.56E-05 | 7.69E-04 |
| 2673 | chr8 | 9975409 | 9975559 | + | 0 | NA | intron (NM | intron (NM | 839 | NM_1769 | 319583 | Mm.80584 | NM_1769 | ENSMUSG | Lig4 | 5830471N | ligase IV, D | protein-co | 5.76 | 6.34 | 4.77 | 1.57 | 3.57E-05 | 7.70E-04 |
| 2680 | chr15 | 66815519 | 66815669 | + | 0 | NA | intron (NM | intron (NM | 16235 | NM_0010 | 20491 | Mm.7601 | NM_0091 | ENSMUSG | Sla | Slap\|Slap- | src-like ad | protein-co | 6.88 | 7.35 | 6.19 | 1.16 | 3.61E-05 | 7.76E-04 |
| 2689 | chr19 | 6398572 | 6398722 | + | 0 | NA | TTS (NM_( | TTS (NM_( | -1228 | NM_0013 | 19395 | Mm.77017 | NM_0112 | ENSMUSG | Rasgrp2 | CDC25L\|C | RAS, guany | protein-co | 5.91 | 6.48 | 4.97 | 1.51 | 3.64E-05 | 7.78E-04 |
| 2724 | chr12 | 99381043 | 99381193 | + | 0 | NA | intron (NM | Lx4A\|LINE | 68956 | NM_1831 | 71375 | Mm.34197 | NM_1831 | ENSMUSG | Foxn3 | 5430426H | forkhead | protein-co | 5.56 | 6.17 | 4.49 | 1.68 | 3.86E-05 | 8.13E-04 |
| 2737 | chr12 | 1.11E+08 | 1.11E+08 | + | 0 | NA | intron (NM | intron (NM | 15550 | NM_0119 | 26448 | Mm.14094 | NM_0119 | ENSMUSG | Mok | RAGE1\|Ra | MOK prote | protein-co | 5.63 | 6.25 | 4.51 | 1.74 | 3.99E-05 | 8.37E-04 |
| 2739 | chr5 | 35819724 | 35819874 | + | 0 | NA | intron (NM | intron (NM | 61919 | NM_0011 | 231148 | Mm.25444 | NM_1776 | ENSMUSG | Ablim2 | AI606905\| | actin-bindi | protein-co | 5.08 | 5.76 | 3.75 | 2.01 | 4.00E-05 | 8.38E-04 |
| 2741 | chr9 | 57972194 | 57972344 | + | 0 | NA | intron (NM | intron (NM | -25755 | NM_0013 | 13070 | Mm.30286 | NM_0197 | ENSMUSG | Cyp11a1 | Cyp11a\|C | cytochrom | protein-co | 6.33 | 6.87 | 5.43 | 1.44 | 4.00E-05 | 8.39E-04 |
| 2744 | chr16 | 18220146 | 18220296 | + | 0 | NA | TTS (NM_ | TTS (NM_ | 2342 | NM_0013 | 1.03E+08 | | NM_0013 | ENSMUSG | Ccdc188 | EG665975 | coiled-coil | protein-co | 6.18 | 6.74 | 5.26 | 1.48 | 4.01E-05 | 8.39E-04 |
| 2746 | chr8 | 1.17E+08 | 1.17E+08 | + | 0 | NA | intron (NM | intron (NM | 113700 | NM_0292 | 75465 | Mm.23114 | NM_0292 | ENSMUSG | Dynlrb2 | 1700009A | dynein ligh | protein-co | 4.93 | 5.64 | 3.45 | 2.19 | 4.02E-05 | 8.40E-04 |
| 2751 | chr4 | 1.55E+08 | 1.55E+08 | + | 0 | NA | non-codin | non-codin | 5620 | NM_0013 | 544678 | Mm.47279 | NM_1776 | ENSMUSG | Cfap74 | 2010015L | cilia and fl | protein-co | 4.85 | 5.56 | 3.4 | 2.16 | 4.04E-05 | 8.44E-04 |
| 2754 | chr1 | 1.62E+08 | 1.62E+08 | + | 0 | NA | Intergenic | Intergenic | 9231 | NM_0012 | 14103 | Mm.3355 | NM_0101 | ENSMUSG | Fasl | APT1LG1\| | Fas ligand | protein-co | 5.99 | 6.55 | 5.05 | 1.5 | 4.08E-05 | 8.51E-04 |
| 2768 | chr1 | 38883936 | 38884086 | + | 0 | NA | intron (NM | intron (NM | 14149 | NM_1451 | 98388 | Mm.26005 | NM_1451 | ENSMUSG | Chst10 | AI507003\| | carbohydr | protein-co | 6.89 | 7.34 | 6.24 | 1.1 | 4.14E-05 | 8.58E-04 |
| 2769 | chr12 | 76095372 | 76095522 | + | 0 | NA | intron (NM | intron (NM | 81812 | NR_10438 | 13983 | Mm.2561 | NM_0101 | ENSMUSG | Esr2 | ER[b]\|ERb | estrogen r | protein-co | 5.15 | 5.84 | 3.79 | 2.05 | 4.14E-05 | 8.58E-04 |
| 2770 | chr3 | 37805718 | 37805868 | + | 0 | NA | Intergenic | Intergenic | -81433 | NM_1986 | 381438 | Mm.49054 | NM_1986 | ENSMUSG | Gm5148 | EG381438 | predicted | protein-co | 6.72 | 7.21 | 5.99 | 1.22 | 4.15E-05 | 8.60E-04 |
| 2795 | chr15 | 55520884 | 55521034 | + | 0 | NA | TTS (NM_ | TTS (NM_ | 36353 | NM_0267 | 68537 | Mm.23679 | NM_0267 | ENSMUSG | Mrpl13 | 1110002D | mitochonc | protein-co | 6.57 | 7.07 | 5.79 | 1.29 | 4.31E-05 | 8.84E-04 |
| 2802 | chr3 | 22151254 | 22151404 | + | 0 | NA | intron (NM | intron (NM | 74677 | NM_0307 | 81004 | Mm.20296 | NM_0307 | ENSMUSG | Tbl1xr1 | 8030499H | transducin | protein-co | 5.57 | 6.17 | 4.51 | 1.66 | 4.36E-05 | 8.92E-04 |
| 2807 | chr7 | 38189547 | 38189697 | + | 0 | NA | intron (NM | intron (NM | 5835 | NM_0281 | 72244 | Mm.18755 | NM_028166 | | 1600014C | AI428873 | RIKEN cDN | protein-co | 7.14 | 7.56 | 6.55 | 1.01 | 4.38E-05 | 8.94E-04 |
| 2817 | chr17 | 88581726 | 88581876 | + | 0 | NA | intron (NM | intron (NM | -44754 | NM_0298 | 77057 | Mm.48220 | NM_0298 | ENSMUSG | Ston1 | 4921524J | stonin 1 | protein-co | 5.38 | 6.08 | 3.98 | 2.09 | 4.44E-05 | 9.04E-04 |
| 2828 | chr5 | 48331543 | 48331693 | + | 0 | NA | intron (NM | intron (NM | -40774 | NM_0257 | 66768 | Mm.29409 | NM_0257 | ENSMUSG | Pacrgl | 4933428G | PARK2 co- | protein-co | 5.43 | 6.12 | 4.04 | 2.08 | 4.58E-05 | 9.27E-04 |
| 2845 | chr16 | 91366723 | 91366873 | + | 0 | NA | Intergenic | Intergenic | -5985 | NM_0013 | 15976 | Mm.6834 | NM_0105 | ENSMUSG | Ifnar2 | AI747302\| | interferon | protein-co | 7.04 | 7.48 | 6.42 | 1.05 | 4.72E-05 | 9.50E-04 |
| 2884 | chr6 | 99901304 | 99901454 | + | 0 | NA | Intergenic | Intergenic | -174987 | NM_0157 | 50501 | Mm.87365 | NM_0157 | ENSMUSG | Prok2 | Bv8\|PK2\| | prokinetic | protein-co | 4.21 | 5.02 | 2.14 | 2.88 | 5.03E-05 | 9.97E-04 |
| 2886 | chr16 | 43832054 | 43832204 | + | 0 | NA | Intergenic | ID4\|SINE\| | 57547 | NM_0291 | 106248 | Mm.27535 | NM_0291 | ENSMUSG | Qtrt2 | 3110012M | queuine tR | protein-co | 4.97 | 5.66 | 3.62 | 2.04 | 5.04E-05 | 9.98E-04 |
| 2888 | chr1 | 1.28E+08 | 1.28E+08 | + | 0 | NA | exon (NM_ | exon (NM_ | 9922 | NM_0117 | 22625 | Mm.24656 | NM_0117 | ENSMUSG | Map3k19 | Ysk4 | mitogen-a | protein-co | 7.1 | 7.54 | 6.46 | 1.08 | 5.06E-05 | 0.001 |
| 2895 | chr10 | 75697125 | 75697275 | + | 0 | NA | intron (NM | intron (NM | -53192 | NM_0278 | 71733 | Mm.24795 | NM_0278 | ENSMUSG | Susd2 | 1200011D | sushi dom | protein-co | 5.65 | 6.24 | 4.64 | 1.6 | 5.09E-05 | 0.001 |
| 2898 | chr8 | 86052064 | 86052214 | + | 0 | NA | intron (NM | intron (NM | 211151 | NM_0013 | 102093 | Mm.23729 | NM_1994 | ENSMUSG | Phkb | AI463271 | phosphory | protein-co | 5.13 | 5.79 | 3.89 | 1.9 | 5.10E-05 | 0.00101 |
| 2904 | chr12 | 51886507 | 51886657 | + | 0 | NA | intron (NM | B4\|SINE\|E | -57046 | NM_1447 | 207304 | Mm.24939 | NM_1447 | ENSMUSG | Hectd1 | A630086P | HECT dom | protein-co | 5.53 | 6.15 | 4.44 | 1.71 | 5.21E-05 | 0.00103 |
| 2935 | chr3 | 51882069 | 51882219 | + | 0 | NA | intron (NM | intron (NM | 220965 | NM_1749 | 211666 | Mm.24679 | NM_1749 | ENSMUSG | Mgst2 | GST2\|MGS | microsoma | protein-co | 5.64 | 6.27 | 4.51 | 1.76 | 5.50E-05 | 0.00107 |
| 2938 | chr3 | 1.42E+08 | 1.42E+08 | + | 0 | NA | intron (NM | intron (NM | 152789 | NM_0013 | 12167 | Mm.39089 | NM_0075 | ENSMUSG | Bmpr1b | AI385617\| | bone morp | protein-co | 5.93 | 6.48 | 5.04 | 1.44 | 5.53E-05 | 0.00108 |
| 2948 | chr13 | 1.02E+08 | 1.02E+08 | + | 0 | NA | intron (NM | intron (NM | -18844 | NM_0010 | 18708 | Mm.25933 | NM_0010 | ENSMUSG | Pik3r1 | PI3K\|p50a | phosphoin | protein-co | 5.93 | 6.49 | 5 | 1.49 | 5.62E-05 | 0.00109 |
| 2957 | chrX | 13516532 | 13516682 | + | 0 | NA | TTS (NM_( | TTS (NM_( | 55517 | NM_1734 | 236690 | Mm.90233 | NM_1734 | ENSMUSG | Nyx | CLNP\|CSN | nyctalopin | protein-co | 5.28 | 5.92 | 4.11 | 1.81 | 5.67E-05 | 0.0011 |
| 2960 | chr7 | 67421162 | 67421312 | + | 0 | NA | Intergenic | Intergenic | -48379 | NM_0010 | 17258 | Mm.13278 | NM_0010 | ENSMUSG | Mef2a | A430079H | myocyte e | protein-co | 4.9 | 5.6 | 3.52 | 2.07 | 5.68E-05 | 0.0011 |
| 2973 | chr18 | 4294669 | 4294819 | + | 0 | NA | Intergenic | Intergenic | 58209 | NM_0077 | 26410 | Mm.3275 | NM_0077 | ENSMUSG | Map3k8 | Cot\|Cot/T | mitogen-a | protein-co | 5.98 | 6.54 | 5.07 | 1.48 | 5.78E-05 | 0.00111 |
| 2982 | chr8 | 95047079 | 95047229 | + | 0 | NA | Intergenic | Intergenic | -7949 | NM_0010 | 330830 | Mm.27711 | NM_0010 | ENSMUSG | Drc7 | Ccdc135\|C | dynein reg | protein-co | 5.77 | 6.35 | 4.78 | 1.58 | 5.83E-05 | 0.00112 |
| 2991 | chr2 | 1.59E+08 | 1.59E+08 | + | 0 | NA | intron (NM | intron (NM | -49040 | NM_0279 | 71878 | Mm.85162 | NM_0279 | ENSMUSG | Fam83d | 2310007D | family with | protein-co | 7.24 | 7.67 | 6.62 | 1.05 | 5.95E-05 | 0.00113 |
| 2989 | chr1 | 66694370 | 66694520 | + | 0 | NA | exon (NM_ | exon (NM_ | -6386 | NM_0013 | 66646 | Mm.24091 | NM_0256 | ENSMUSG | Rpe | 2810429B | ribulose-5- | protein-co | 5.75 | 6.33 | 4.78 | 1.55 | 5.91E-05 | 0.00113 |
| 3007 | chr6 | 1.46E+08 | 1.46E+08 | + | 0 | NA | Intergenic | ID_B1\|SIN | 19616 | NM_0012 | 79362 | Mm.15452 | NM_0244 | ENSMUSG | Bhlhe41 | 6430520M | basic helix | protein-co | 4.46 | 5.23 | 2.67 | 2.56 | 6.05E-05 | 0.00115 |
| 3013 | chr1 | 1.27E+08 | 1.27E+08 | + | 0 | NA | intron (NM | intron (NM | 234270 | NM_0281 | 72160 | Mm.28128 | NM_0281 | ENSMUSG | Tmem163 | 2610024A | transmem | protein-co | 5.27 | 5.93 | 4 | 1.93 | 6.14E-05 | 0.00116 |
| 3027 | chr4 | 1.07E+08 | 1.07E+08 | + | 0 | NA | TTS (NM_( | TTS (NM_( | -7149 | NM_0010 | 546840 | Mm.33175 | NM_0010 | ENSMUSG | Ldlrad1 | - | low densit | protein-co | 6.87 | 7.36 | 6.13 | 1.24 | 6.30E-05 | 0.00119 |
| 3038 | chr2 | 18937815 | 18937965 | + | 0 | NA | intron (NM | intron (NM | 60231 | NM_0088 | 18718 | Mm.31397 | NM_0088 | ENSMUSG | Pip4k2a | AW74291 | phosphati | protein-co | 5.31 | 5.95 | 4.12 | 1.84 | 6.44E-05 | 0.00121 |
| 3046 | chr9 | 1.03E+08 | 1.03E+08 | + | 0 | NA | exon (NM_ | exon (NM_ | 36479 | NM_0242 | 16716 | Mm.10169 | NM_0242 | ENSMUSG | Ky | CD1\|D9M | kyphoscoli | protein-co | 7.06 | 7.48 | 6.45 | 1.03 | 6.55E-05 | 0.00122 |
| 3054 | chr2 | 1.03E+08 | 1.03E+08 | + | 0 | NA | intron (NM | intron (NM | 34027 | NM_0011 | 12505 | Mm.42362 | NM_0098 | ENSMUSG | Cd44 | AU023126 | CD44 antig | protein-co | 6.18 | 6.7 | 5.36 | 1.34 | 6.66E-05 | 0.00124 |
| 3068 | chr19 | 57131543 | 57131693 | + | 0 | NA | Intergenic | Intergenic | -12594 | NM_0011 | 226251 | Mm.21716 | NM_1786 | ENSMUSG | Ablim1 | 2210411C | actin-bindi | protein-co | 7.17 | 7.6 | 6.57 | 1.03 | 6.85E-05 | 0.00127 |
| 3090 | chr2 | 1.26E+08 | 1.26E+08 | + | 0 | NA | intron (NM | MER58A\|E | 82143 | NM_0012 | 69976 | Mm.20216 | NM_1751 | ENSMUSG | Galk2 | 2810017M | galactokin | protein-co | 6.51 | 7.02 | 5.72 | 1.3 | 7.05E-05 | 0.0013 |
| 3107 | chr8 | 1.23E+08 | 1.23E+08 | + | 0 | NA | promoter- | promoter- | -378 | NM_1772 | 12398 | Mm.19433 | NM_0098 | ENSMUSG | Cbfa2t3 | A630044F | CBFA2/RU | protein-co | 6.77 | 7.24 | 6.06 | 1.19 | 7.18E-05 | 0.00131 |
| 3118 | chr16 | 46022343 | 46022493 | + | 0 | NA | Intergenic | Intergenic | -12005 | NM_0011 | 433022 | Mm.38099 | NM_0011 | ENSMUSG | Plcxd2 | AW49463 | phosphati | protein-co | 7.09 | 7.51 | 6.5 | 1.01 | 7.30E-05 | 0.00133 |
| 3119 | chr12 | 1.13E+08 | 1.13E+08 | + | 0 | NA | TTS (NM_( | TTS (NM_( | -16881 | NM_0011 | 399566 | Mm.44014 | NM_2016 | ENSMUSG | Btbd6 | - | BTB (POZ) | protein-co | 4.62 | 5.35 | 3.09 | 2.25 | 7.31E-05 | 0.00133 |
| 3130 | chr14 | 30273123 | 30273273 | + | 0 | NA | intron (NM | intron (NM | 80288 | NM_0013 | 12289 | Mm.9772 | NM_0289 | ENSMUSG | Cacna1d | 8430418G | calcium ch | protein-co | 4.72 | 5.43 | 3.24 | 2.19 | 7.37E-05 | 0.00134 |
| 3145 | chr17 | 28388748 | 28388898 | + | 0 | NA | Intergenic | Intergenic | -23607 | NM_0214 | 22157 | Mm.42102 | NM_0214 | ENSMUSG | Tulp1 | Tulp1l | tubby like | protein-co | 4.48 | 5.23 | 2.84 | 2.38 | 7.56E-05 | 0.00136 |
| 3165 | chr1 | 1.81E+08 | 1.81E+08 | + | 0 | NA | Intergenic | Intergenic | -12543 | NM_0100 | 13590 | Mm.37891 | NM_0100 | ENSMUSG | Lefty1 | AI450052\| | left right d | protein-co | 4.84 | 5.53 | 3.46 | 2.07 | 7.75E-05 | 0.00139 |
| 3175 | chr12 | 73794252 | 73794402 | + | 0 | NA | Intergenic | Intergenic | -106997 | NM_0013 | 15251 | Mm.3879 | NM_0104 | ENSMUSG | Hif1a | AA959795 | hypoxia in | protein-co | 6.23 | 6.74 | 5.41 | 1.33 | 7.84E-05 | 0.0014 |
| 3171 | chr15 | 1.01E+08 | 1.01E+08 | + | 0 | NA | intron (NM | intron (NM | 5427 | NM_0168 | 53622 | Mm.34793 | NM_016879 | | Krt85 | AA589597 | keratin 85 | protein-co | 5.85 | 6.43 | 4.87 | 1.56 | 7.82E-05 | 0.0014 |
| 3177 | chr9 | 63504034 | 63504184 | + | 0 | NA | intron (NM | MTD\|LTR\| | 98384 | NM_0013 | 78250 | Mm.44461 | NM_0300 | ENSMUSG | Iqch | 4921504K | IQ motif c | protein-co | 5.22 | 5.85 | 4.06 | 1.8 | 7.85E-05 | 0.0014 |

| | | | | | | | | | | | | | | | | | | | | | | | | |
|---|---|---|---|---|---|---|---|---|---|---|---|---|---|---|---|---|---|---|---|---|---|---|---|---|
| 3213 | chr12 | 80736849 | 80736999 | + | 0 | NA | Intergenic | Intergenic | 23791 | NM_0010( | 380768 | Mm.29367 | NM_0010( | ENSMUSG( | Ccdc177 | Gm1568 | coiled-coil | protein-co | 5.38 | 6 | 4.28 | 1.72 | 8.46E-05 | 0.00149 |
| 3218 | chr2 | 71866709 | 71866859 | + | 0 | NA | intron (NR | intron (NR | -6440 | NM_17266 | 228026 | Mm.34411 | NM_17266 | ENSMUSG( | Pdk1 | B830012B( | pyruvate d | protein-co | 5.39 | 6 | 4.32 | 1.68 | 8.51E-05 | 0.0015 |
| 3223 | chr3 | 22138158 | 22138308 | + | 0 | NA | intron (NM | intron (NM | 61581 | NM_03073 | 81004 | Mm.20296 | NM_03073 | ENSMUSG( | Tbl1xr1 | 8030499H( | transducin | protein-co | 4.54 | 5.29 | 2.92 | 2.36 | 8.56E-05 | 0.0015 |
| 3237 | chr2 | 1.32E+08 | 1.32E+08 | + | 0 | NA | intron (NM | intron (NM | 7961 | NM_00129 | 110911 | Mm.2845( | NM_13865 | ENSMUSG( | Cds2 | 5730450N( | CDP-diacyl | protein-co | 4.58 | 5.31 | 3 | 2.31 | 8.84E-05 | 0.00154 |
| 3243 | chr12 | 32088721 | 32088871 | + | 0 | NA | Intergenic | Intergenic | -27519 | NM_01115 | 19088 | Mm.25594 | NM_01115 | ENSMUSG( | Prkar2b | AI451071\| | protein kir | protein-co | 4.01 | 4.85 | 1.79 | 3.06 | 8.93E-05 | 0.00156 |
| 3256 | chr5 | 35817394 | 35817544 | + | 0 | NA | intron (NM | intron (NM | 59589 | NM_00117 | 231148 | Mm.25444 | NM_17767 | ENSMUSG( | Ablim2 | AI606905\| | actin-bindi | protein-co | 5.87 | 6.43 | 4.94 | 1.49 | 9.04E-05 | 0.00157 |
| 3258 | chr13 | 19219755 | 19219905 | + | 0 | NA | Intergenic | Intergenic | 175983 | NM_00134 | 76205 | Mm.4194( | NM_02427 | ENSMUSG( | Stard3nl | 0610035N( | STARD3 N- | protein-co | 5.68 | 6.25 | 4.7 | 1.55 | 9.08E-05 | 0.00157 |
| 3265 | chr2 | 1.04E+08 | 1.04E+08 | + | 0 | NA | Intergenic | Intergenic | -8887 | NM_00136 | 241589 | Mm.3589( | NM_00103 | ENSMUSG( | D430041D | 96300566( | RIKEN cDN | protein-co | 3.72 | 4.6 | 0.97 | 3.64 | 9.10E-05 | 0.00158 |
| 3271 | chr1 | 1.53E+08 | 1.53E+08 | + | 0 | NA | Intergenic | MER58A\|[ | -11098 | NM_00848 | 16782 | Mm.32927 | NM_00848 | ENSMUSG( | Lamc2 | AA589349 | laminin, ga | protein-co | 6.15 | 6.68 | 5.33 | 1.35 | 9.17E-05 | 0.00159 |
| 3279 | chr18 | 70633730 | 70633880 | + | 0 | NA | Intergenic | MLT1L\|LTI | 65513 | NM_00131 | 17191 | Mm.322 | NM_01077 | ENSMUSG( | Mbd2 | MBD2a | methyl-Cp | protein-co | 5.35 | 5.98 | 4.21 | 1.77 | 9.25E-05 | 0.0016 |
| 3290 | chr4 | 1.55E+08 | 1.55E+08 | + | 0 | NA | intron (NM | intron (NM | -6949 | NM_00103 | 18762 | Mm.28561 | NM_00886 | ENSMUSG( | Prkcz | AI098070\| | protein kir | protein-co | 5.15 | 5.82 | 3.85 | 1.97 | 9.47E-05 | 0.00163 |
| 3296 | chr15 | 98777239 | 98777389 | + | 0 | NA | intron (NM | intron (NM | 836 | NM_01171 | 22410 | Mm.4709 | NM_01171 | ENSMUSG( | Wnt10b | Wnt12 | wingless-t | protein-co | 5.39 | 6.03 | 4.22 | 1.81 | 9.57E-05 | 0.00164 |
| 3341 | chr17 | 28488360 | 28488510 | + | 0 | NA | Intergenic | Intergenic | -2286 | NM_01022 | 14229 | Mm.2764( | NM_01022 | ENSMUSG( | Fkbp5 | D17Ertd59 | FK506 binc | protein-co | 6.56 | 7.04 | 5.81 | 1.23 | 1.02E-04 | 0.00172 |
| 3340 | chr7 | 1.11E+08 | 1.11E+08 | + | 0 | NA | intron (NM | MTD\|LTR\| | 4269 | NM_19446 | 17540 | Mm.25002 | NM_01082 | ENSMUSG( | Mrvi1 | BB115629 | MRV integ | protein-co | 5.87 | 6.42 | 4.96 | 1.46 | 1.02E-04 | 0.00172 |
| 3333 | chr16 | 24886993 | 24887143 | + | 0 | NA | intron (NM | intron (NM | 165213 | NM_00114 | 210126 | Mm.20938 | NM_17866 | ENSMUSG( | Lpp | 9430020K( | LIM domai | protein-co | 5.14 | 5.78 | 3.97 | 1.81 | 1.01E-04 | 0.00172 |
| 3346 | chr12 | 1.09E+08 | 1.09E+08 | + | 0 | NA | Intergenic | Intergenic | -11267 | NR_03381 | 13386 | Mm.15706 | NM_01005 | ENSMUSG( | Dlk1 | AW742678 | delta like r | protein-co | 5.83 | 6.4 | 4.87 | 1.53 | 1.03E-04 | 0.00173 |
| 3350 | chr18 | 70445091 | 70445241 | + | 0 | NA | Intergenic | Intergenic | -27288 | NM_02901 | 170461 | Mm.83623 | NM_02901 | ENSMUSG( | Stard6 | 1700011K( | StAR-relat | protein-co | 5.3 | 5.94 | 4.11 | 1.82 | 1.03E-04 | 0.00174 |
| 3353 | chr10 | 26324851 | 26325001 | + | 0 | NA | intron (NM | intron (NM | 49146 | NM_00135 | 237339 | Mm.33843 | NM_17278 | ENSMUSG( | L3mbtl3 | AI481284\| | L3MBTL3 h | protein-co | 4.73 | 5.43 | 3.32 | 2.1 | 1.03E-04 | 0.00174 |
| 3414 | chr2 | 1.31E+08 | 1.31E+08 | + | 0 | NA | intron (NM | intron (NM | -5682 | NM_08056 | 140629 | Mm.4860( | NM_08056 | ENSMUSG( | Ubox5 | 1500010O( | U box dom | protein-co | 5.05 | 5.71 | 3.81 | 1.9 | 1.12E-04 | 0.00184 |
| 3425 | chr5 | 37044926 | 37045076 | + | 0 | NA | Intergenic | Intergenic | -5856 | NM_17839 | 76071 | Mm.8528( | NM_17839 | ENSMUSG( | Jakmip1 | 5830437M | janus kinas | protein-co | 7 | 7.42 | 6.4 | 1.02 | 1.14E-04 | 0.00188 |
| 3433 | chr9 | 50407458 | 50407608 | + | 0 | NA | Intergenic | Intergenic | -86992 | NM_02963 | 76509 | Mm.29959 | NM_02963 | ENSMUSG( | Plet1 | 0610037P( | placenta e | protein-co | 5.31 | 5.93 | 4.21 | 1.72 | 1.15E-04 | 0.00188 |
| 3449 | chr13 | 19300596 | 19300746 | + | 0 | NA | Intergenic | Intergenic | 95142 | NM_00134 | 76205 | Mm.4194( | NM_02427 | ENSMUSG( | Stard3nl | 0610035N( | STARD3 N- | protein-co | 6.26 | 6.76 | 5.49 | 1.27 | 1.17E-04 | 0.0019 |
| 3458 | chr16 | 52091027 | 52091177 | + | 0 | NA | intron (NM | intron (NM | 59553 | NM_00103 | 208650 | Mm.3282( | NM_00103 | ENSMUSG( | Cblb | AI429560\| | Casitas B-l | protein-co | 5.49 | 6.13 | 4.31 | 1.82 | 1.18E-04 | 0.00192 |
| 3467 | chr5 | 37079899 | 37080049 | + | 0 | NA | intron (NM | intron (NM | -5262 | NM_00134 | 76071 | Mm.8528( | NM_17839 | ENSMUSG( | Jakmip1 | 5830437M | janus kinas | protein-co | 5.37 | 5.99 | 4.27 | 1.71 | 1.19E-04 | 0.00193 |
| 3499 | chr10 | 93876876 | 93877026 | + | 0 | NA | intron (NM | RSINE1\|SII | 14251 | NM_01964 | 56307 | Mm.28932 | NM_01964 | ENSMUSG( | Metap2 | 4930584B( | methionin | protein-co | 6.1 | 6.62 | 5.27 | 1.34 | 1.24E-04 | 0.00199 |
| 3506 | chr16 | 5429561 | 5429711 | + | 0 | NA | Intergenic | RMER15\|L | -173680 | NM_02744 | 70511 | Mm.4578( | NM_02744 | ENSMUSG( | Eef2kmt | 5730409G( | eukaryotic | protein-co | 6.64 | 7.12 | 5.93 | 1.18 | 1.26E-04 | 0.00201 |
| 3516 | chr18 | 58560954 | 58561104 | + | 0 | NA | intron (NM | intron (NM | 4789 | NM_00108 | 225579 | Mm.25851 | NM_00108 | ENSMUSG( | Slc27a6 | 4732438L( | solute carr | protein-co | 5.83 | 6.37 | 4.96 | 1.42 | 1.27E-04 | 0.00203 |
| 3534 | chr18 | 46381045 | 46381195 | + | 0 | NA | Intergenic | Intergenic | -69192 | NM_00116 | 240261 | Mm.32941 | NM_00116 | ENSMUSG( | Ccdc112 | 8430438M | coiled-coil | protein-co | 6.31 | 6.8 | 5.55 | 1.25 | 1.29E-04 | 0.00205 |
| 3546 | chr7 | 1.16E+08 | 1.16E+08 | + | 0 | NA | intron (NM | intron (NM | 35614 | NM_00130 | 233765 | Mm.3741 | NM_17274 | ENSMUSG( | Plekha7 | A430081P( | pleckstrin | protein-co | 4.89 | 5.56 | 3.56 | 2 | 1.32E-04 | 0.00209 |
| 3552 | chr15 | 97721413 | 97721563 | + | 0 | NA | promoter- | promoter- | -338 | NM_00890 | 19011 | Mm.1001 | NM_00890 | ENSMUSG( | Endou | Pp11r\|Tcl- | endonucle | protein-co | 6.84 | 7.28 | 6.2 | 1.09 | 1.34E-04 | 0.00212 |
| 3572 | chr18 | 58571124 | 58571274 | + | 0 | NA | intron (NM | intron (NM | 14959 | NM_00108 | 225579 | Mm.25851 | NM_00108 | ENSMUSG( | Slc27a6 | 4732438L( | solute carr | protein-co | 5.11 | 5.79 | 3.8 | 1.98 | 1.38E-04 | 0.00216 |
| 3584 | chr1 | 39025251 | 39025401 | + | 0 | NA | Intergenic | Intergenic | 37512 | NM_02685 | 68833 | Mm.28121 | NM_02685 | ENSMUSG( | Pdcl3 | 1110061A( | phosducin | protein-co | 6.78 | 7.22 | 6.14 | 1.08 | 1.41E-04 | 0.00221 |
| 3603 | chr7 | 1.26E+08 | 1.26E+08 | + | 0 | NA | Intergenic | Intergenic | 32213 | NM_02188 | 60504 | Mm.15564 | NM_02188 | ENSMUSG( | Il21r | NILR | interleukir | protein-co | 7 | 7.42 | 6.41 | 1 | 1.44E-04 | 0.00224 |
| 3609 | chr9 | 66094803 | 66094953 | + | 0 | NA | intron (NM | intron (NM | -25147 | NM_00102 | 382083 | Mm.4771( | NM_00102 | ENSMUSG( | Snx22 | Gm1120 | sorting ne: | protein-co | 4.82 | 5.52 | 3.42 | 2.09 | 1.45E-04 | 0.00225 |
| 3613 | chr8 | 40577608 | 40577758 | + | 0 | NA | intron (NM | MLT1E\|LT | 57109 | NM_00104 | 54384 | Mm.29487 | NM_01943 | ENSMUSG( | Mtmr7 | - | myotubula | protein-co | 6.15 | 6.65 | 5.39 | 1.27 | 1.46E-04 | 0.00226 |
| 3616 | chr2 | 89930340 | 89930490 | + | 0 | NA | exon (NM_ | exon (NM_ | 604 | NM_14697 | 258980 | Mm.22337 | NM_14697 | ENSMUSG( | Olfr1258 | MOR232-3 | olfactory r | protein-co | 6.06 | 6.6 | 5.18 | 1.42 | 1.46E-04 | 0.00226 |
| 3631 | chr14 | 31531912 | 31532062 | + | 0 | NA | intron (NM | intron (NM | 36908 | NM_02893 | 74427 | Mm.4912( | NM_02893 | ENSMUSG( | Eaf1 | 4933403C( | ELL associ | protein-co | 6.48 | 6.96 | 5.76 | 1.19 | 1.49E-04 | 0.0023 |
| 3636 | chr17 | 23610560 | 23610710 | + | 0 | NA | exon (NM_ | exon (NM_ | 9809 | NM_00128 | 332221 | Mm.27031 | NM_00103 | ENSMUSG( | Zscan10 | BC068284 | zinc finger | protein-co | 7.29 | 7.72 | 6.68 | 1.04 | 1.50E-04 | 0.00231 |
| 3648 | chr2 | 1.45E+08 | 1.45E+08 | + | 0 | NA | intron (NM | intron (NM | -33628 | NM_05319 | 94249 | Mm.21717 | NM_05319 | ENSMUSG( | Slc24a3 | NCKX3 | solute carr | protein-co | 6.77 | 7.22 | 6.11 | 1.11 | 1.51E-04 | 0.00232 |
| 3672 | chr5 | 37047217 | 37047367 | + | 0 | NA | Intergenic | (TG)n\|Sim | -3565 | NM_17839 | 76071 | Mm.8528( | NM_17839 | ENSMUSG( | Jakmip1 | 5830437M | janus kinas | protein-co | 6.83 | 7.26 | 6.19 | 1.07 | 1.55E-04 | 0.00236 |
| 3671 | chr13 | 19178392 | 19178542 | + | 0 | NA | Intergenic | Intergenic | 217346 | NM_00134 | 76205 | Mm.4194( | NM_02427 | ENSMUSG( | Stard3nl | 0610035N( | STARD3 N- | protein-co | 5.99 | 6.51 | 5.17 | 1.34 | 1.55E-04 | 0.00236 |
| 3687 | chr14 | 30291883 | 30292033 | + | 0 | NA | intron (NM | intron (NM | 61528 | NM_00130 | 12289 | Mm.9772 | NM_02898 | ENSMUSG( | Cacna1d | 8430418G( | calcium ch | protein-co | 4.5 | 5.25 | 2.84 | 2.4 | 1.58E-04 | 0.00239 |
| 3698 | chr11 | 80489941 | 80490091 | + | 0 | NA | intron (NM | intron (NM | 12993 | NM_00987 | 12569 | Mm.14227 | NM_00987 | ENSMUSG( | Cdk5r1 | Cdk5r\|D11 | cyclin-dep | protein-co | 6.97 | 7.39 | 6.37 | 1.01 | 1.60E-04 | 0.00241 |
| 3715 | chr1 | 64949480 | 64949630 | + | 0 | NA | intron (NM | Lx7\|LINE\|I | 7269 | NM_00103 | 241075 | Mm.23953 | NM_1755( | ENSMUSG( | Plekhm3 | 9430067K( | pleckstrin | protein-co | 5.15 | 5.79 | 4.01 | 1.78 | 1.65E-04 | 0.00247 |
| 3727 | chr4 | 1.3E+08 | 1.3E+08 | + | 0 | NA | intron (NM | intron (NM | 27531 | NM_02564 | 66585 | Mm.42301 | NM_02564 | ENSMUSG( | Snrnp40 | 0610009C( | small nucle | protein-co | 5.85 | 6.4 | 4.98 | 1.42 | 1.68E-04 | 0.00251 |
| 3730 | chr16 | 48627767 | 48627917 | + | 0 | NA | intron (NM | MTE2b\|LT | 144220 | NM_19829 | 77647 | Mm.16729 | NM_19829 | ENSMUSG( | Trat1 | C030046M | T cell rece | protein-co | 4.95 | 5.6 | 3.74 | 1.86 | 1.68E-04 | 0.00251 |
| 3740 | chr18 | 38974032 | 38974182 | + | 0 | NA | Intergenic | MLT1K\|LT | -19038 | NM_00136 | 71302 | Mm.32939 | NM_17516 | ENSMUSG( | Arhgap26 | 1810044A( | Rho GTPas | protein-co | 4.97 | 5.64 | 3.65 | 1.99 | 1.70E-04 | 0.00253 |
| 3747 | chr16 | 23144443 | 23144593 | + | 0 | NA | Intergenic | Intergenic | -2018 | NM_00960 | 11450 | Mm.3969 | NM_00960 | ENSMUSG( | Adipoq | 30kDa\|API | adiponecti | protein-co | 5.9 | 6.44 | 5.04 | 1.39 | 1.71E-04 | 0.00254 |
| 3763 | chr14 | 1.19E+08 | 1.19E+08 | + | 0 | NA | intron (NM | intron (NM | 8497 | NM_00116 | 58187 | Mm.39075 | NM_02138 | ENSMUSG( | Cldn10 | 6720456I( | claudin 10 | protein-co | 5.44 | 6.04 | 4.37 | 1.67 | 1.74E-04 | 0.00257 |
| 3769 | chr13 | 91091542 | 91091692 | + | 0 | NA | intron (NM | intron (NM | 132370 | NM_02577 | 66795 | Mm.23538 | NM_02577 | ENSMUSG( | Atg10 | 5330424L( | autophagy | protein-co | 6.14 | 6.66 | 5.33 | 1.33 | 1.75E-04 | 0.00258 |
| 3772 | chr1 | 1.84E+08 | 1.84E+08 | + | 0 | NA | Intergenic | Intergenic | 27761 | NM_02851 | 73363 | Mm.50115 | NM_02851 | ENSMUSG( | 1700056E( | - | RIKEN cDN | protein-co | 5.92 | 6.45 | 5.07 | 1.38 | 1.75E-04 | 0.00258 |
| 3777 | chr11 | 11763512 | 11763662 | + | 0 | NA | intron (NM | intron (NM | 45375 | NM_00116 | 60530 | Mm.23611 | NM_02189 | ENSMUSG( | Fignl1 | - | fidgetin-lik | protein-co | 5.89 | 6.42 | 5.04 | 1.39 | 1.75E-04 | 0.00258 |
| 3778 | chr11 | 32558782 | 32558932 | + | 0 | NA | intron (NM | intron (NM | 25591 | NM_00928 | 20868 | Mm.8235 | NM_00928 | ENSMUSG( | Stk10 | Gek1\|Lok\| | serine/thre | protein-co | 5.72 | 6.28 | 4.8 | 1.48 | 1.75E-04 | 0.00258 |
| 3822 | chr7 | 83661428 | 83661578 | + | 0 | NA | intron (NM | intron (NM | -6008 | NM_00136 | 16170 | Mm.10137 | NM_01055 | ENSMUSG( | Il16 | mKIAA404 | interleukir | protein-co | 5.55 | 6.12 | 4.6 | 1.52 | 1.85E-04 | 0.00269 |
| 3828 | chr13 | 46845048 | 46845198 | + | 0 | NA | intron (NM | B3A\|SINE\| | 84595 | NM_01061 | 16553 | Mm.3427( | NM_01061 | ENSMUSG( | Kif13a | 4930505I( | kinesin far | protein-co | 4.6 | 5.32 | 3.07 | 2.25 | 1.86E-04 | 0.0027 |
| 3837 | chr4 | 1.24E+08 | 1.24E+08 | + | 0 | NA | intron (NM | intron (NM | 10704 | NM_01747 | 54170 | Mm.22092 | NM_01747 | ENSMUSG( | Rragc | AU041672 | Ras-relate | protein-co | 6.57 | 7.02 | 5.92 | 1.1 | 1.89E-04 | 0.00273 |
| 3843 | chr8 | 83748027 | 83748177 | + | 0 | NA | Intergenic | Intergenic | -6791 | NM_00116 | 26364 | Mm.33464 | NM_01192 | ENSMUSG( | Adgre5 | AA409984 | adhesion ( | protein-co | 6.33 | 6.83 | 5.54 | 1.29 | 1.91E-04 | 0.00276 |
| 3860 | chr15 | 37885238 | 37885388 | + | 0 | NA | Intergenic | Intergenic | 75543 | NM_00135 | 382985 | Mm.24738 | NM_19947 | ENSMUSG( | Rrm2b | p53R2 | ribonuclec | protein-co | 4.81 | 5.52 | 3.36 | 2.16 | 1.94E-04 | 0.00279 |
| 3876 | chr1 | 1.39E+08 | 1.39E+08 | + | 0 | NA | Intergenic | L2\|LINE\|L | 56360 | NM_00104 | 16876 | Mm.25073 | NM_01077 | ENSMUSG( | Lhx9 | 3110009O( | LIM homeo | protein-co | 6.31 | 6.81 | 5.53 | 1.28 | 1.98E-04 | 0.00284 |
| 3877 | chr8 | 11503991 | 11504141 | + | 0 | NA | intron (NM | intron (NM | 6498 | NM_02699 | 69225 | Mm.64911 | NM_02699 | ENSMUSG( | Naxd | 0710008K( | NAD(P)HX | protein-co | 7.18 | 7.93 | 5.55 | 2.38 | 1.98E-04 | 0.00284 |
| 3890 | chr11 | 82947542 | 82947692 | + | 0 | NA | intron (NR | intron (NR | -4485 | NM_18320 | 327978 | Mm.42147 | NM_18320 | ENSMUSG( | Slfn5 | AI956840 | schlafen 5 | protein-co | 6.18 | 6.68 | 5.43 | 1.25 | 2.01E-04 | 0.00286 |
| 3883 | chr10 | 95781052 | 95781202 | + | 0 | NA | Intergenic | Intergenic | -159536 | NM_00100 | 216238 | Mm.21003 | NM_00100 | ENSMUSG( | Eea1 | A430109M | early endo | protein-co | 5.07 | 5.71 | 3.91 | 1.79 | 2.00E-04 | 0.00286 |
| 3893 | chr12 | 70088438 | 70088588 | + | 0 | NA | intron (NM | MER91B\|[ | 14341 | NM_00108 | 18080 | Mm.24404 | NM_00869 | ENSMUSG( | Nin | 3110068G( | ninein | protein-co | 4.75 | 5.43 | 3.45 | 1.98 | 2.02E-04 | 0.00288 |
| 3903 | chr16 | 36985697 | 36985847 | + | 0 | NA | intron (NM | (CAGG)n\|S | 4695 | NM_00135 | 207215 | Mm.21189 | NM_00103 | ENSMUSG( | Fbxo40 | 9830003A( | F-box prot | protein-co | 6.74 | 7.17 | 6.13 | 1.03 | 2.04E-04 | 0.0029 |
| 3913 | chr7 | 1.13E+08 | 1.13E+08 | + | 0 | NA | intron (NM | intron (NM | 115366 | NM_00116 | 21676 | Mm.24685 | NM_00934 | ENSMUSG( | Tead1 | 2610024B( | TEA domai | protein-co | 4.96 | 5.65 | 3.61 | 2.04 | 2.07E-04 | 0.00293 |
| 3937 | chr7 | 1.02E+08 | 1.02E+08 | + | 0 | NA | intron (NM | intron (NM | 17349 | NM_00910 | 20133 | Mm.19748 | NM_00910 | ENSMUSG( | Rrm1 | RnrM1 | ribonuclec | protein-co | 4.35 | 5.14 | 2.48 | 2.65 | 2.10E-04 | 0.00296 |
| 3944 | chr10 | 1.18E+08 | 1.18E+08 | + | 0 | NA | Intergenic | Intergenic | 11196 | NM_01697 | 50929 | Mm.10358 | NM_01697 | ENSMUSG( | Il22 | IL-22\|IL-2 | interleukir | protein-co | 3.51 | 4.42 | 0.49 | 3.92 | 2.12E-04 | 0.00298 |
| 3945 | chr8 | 1.21E+08 | 1.21E+08 | + | 0 | NA | Intergenic | Intergenic | 33706 | NM_00129 | 12857 | Mm.38679 | NM_00994 | ENSMUSG( | Cox4i1 | AL024441\| | cytochrom | protein-co | 6.14 | 6.64 | 5.36 | 1.28 | 2.13E-04 | 0.00299 |
| 3958 | chr11 | 1.18E+08 | 1.18E+08 | + | 0 | NA | Intergenic | Intergenic | -6916 | NM_00770 | 12702 | Mm.3468 | NM_00770 | ENSMUSG( | Socs3 | Cis3\|Cish3 | suppresso | protein-co | 5.18 | 5.79 | 4.09 | 1.7 | 2.15E-04 | 0.00301 |
| 3967 | chr9 | 57638156 | 57638306 | + | 0 | NA | intron (NM | intron (NM | 7422 | NM_00778 | 12988 | Mm.21974 | NM_00778 | ENSMUSG( | Csk | AW21263( | c-src tyros | protein-co | 5.8 | 6.33 | 4.96 | 1.38 | 2.18E-04 | 0.00304 |
| 3988 | chr7 | 1.11E+08 | 1.11E+08 | + | 0 | NA | intron (NM | ORR1E\|LT | -10470 | NM_01082 | 17540 | Mm.25002 | NM_01082 | ENSMUSG( | Mrvi1 | BB115629 | MRV integ | protein-co | 6.42 | 6.88 | 5.75 | 1.13 | 2.22E-04 | 0.00309 |
| 3987 | chr10 | 75137229 | 75137379 | + | 0 | NA | intron (NM | ORR1B2\|L | -74769 | NM_1534( | 74392 | Mm.27716 | NM_1534( | ENSMUSG( | Specc1l | 4930470P( | sperm anti | protein-co | 6.15 | 6.65 | 5.39 | 1.26 | 2.22E-04 | 0.00309 |
| 4055 | chr4 | 1.34E+08 | 1.34E+08 | + | 0 | NA | Intergenic | Intergenic | -15700 | NM_00116 | 230806 | Mm.24008 | NM_00116 | ENSMUSG( | Crybg2 | Aim1l | crystallin b | protein-co | 4.67 | 5.35 | 3.33 | 2.02 | 2.42E-04 | 0.00331 |
| 4083 | chr15 | 84684707 | 84684857 | + | 0 | NA | intron (NM | MLT1D\|LT | 3784 | NM_14606 | 109270 | Mm.29137 | NM_14606 | ENSMUSG( | Prr5 | AU043908 | proline ricl | protein-co | 6.13 | 6.63 | 5.35 | 1.29 | 2.51E-04 | 0.0034 |
| 4111 | chr3 | 21943060 | 21943210 | + | 0 | NA | Intergenic | Intergenic | -133517 | NM_03073 | 81004 | Mm.20296 | NM_03073 | ENSMUSG( | Tbl1xr1 | 8030499H( | transducin | protein-co | 5.69 | 6.26 | 4.74 | 1.52 | 2.60E-04 | 0.00348 |
| 4115 | chr8 | 68121478 | 68121628 | + | 0 | NA | Intergenic | Intergenic | -146979 | NM_17769 | 234353 | Mm.32525 | NM_02762 | ENSMUSG( | Psd3 | 4931420C( | pleckstrin | protein-co | 6.61 | 7.06 | 5.97 | 1.09 | 2.61E-04 | 0.00349 |
| 4124 | chr19 | 53893706 | 53893856 | + | 0 | NA | intron (NM | intron (NM | 1550 | NM_01105 | 18569 | Mm.1605 | NM_01105 | ENSMUSG( | Pdcd4 | D19Ucla1\| | programm | protein-co | 6.8 | 7.22 | 6.21 | 1.01 | 2.63E-04 | 0.00352 |
| 4123 | chr18 | 35759576 | 35759726 | + | 0 | NA | Intergenic | Intergenic | -11908 | NM_01991 | 56550 | Mm.18005 | NM_01991 | ENSMUSG( | Ube2d2a | 1500034D( | ubiquitin-( | protein-co | 4.44 | 5.21 | 2.72 | 2.49 | 2.63E-04 | 0.00352 |
| 4128 | chr8 | 1.21E+08 | 1.21E+08 | + | 0 | NA | intron (NM | (ATG)n\|Sir | 10148 | NM_00130 | 15900 | Mm.33486 | NM_00832 | ENSMUSG( | Irf8 | AI893568\| | interferon | protein-co | 5.53 | 6.13 | 4.49 | 1.64 | 2.65E-04 | 0.00353 |
| 4131 | chr9 | 1.04E+08 | 1.04E+08 | + | 0 | NA | intron (NM | intron (NM | 6245 | NM_01980 | 56318 | Mm.19941 | NM_01980 | ENSMUSG( | Acpp | 5'-NT\|A03 | acid phosp | protein-co | 5.24 | 5.85 | 4.18 | 1.66 | 2.66E-04 | 0.00355 |
| 4136 | chr4 | 1.35E+08 | 1.35E+08 | + | 0 | NA | intron (NM | intron (NM | 10675 | NM_01973 | 12399 | Mm.37889 | NM_01973 | ENSMUSG( | Runx3 | AML2\|Cbf | runt relate | protein-co | 6.47 | 6.95 | 5.76 | 1.19 | 2.69E-04 | 0.00358 |
| 4158 | chr15 | 66705407 | 66705557 | + | 0 | NA | intron (NM | intron (NM | 34712 | NM_00937 | 21819 | Mm.44133 | NM_00937 | ENSMUSG( | Tg | Tgn\|cog | thyroglobu | protein-co | 5.07 | 5.7 | 3.94 | 1.76 | 2.77E-04 | 0.00367 |
| 4174 | chr7 | 1.17E+08 | 1.17E+08 | + | 0 | NA | intron (NM | intron (NM | 105636 | NM_17564 | 233781 | Mm.46142 | NM_17564 | ENSMUSG( | Xylt1 | 8030490L( | xylosyltrar | protein-co | 5.88 | 6.4 | 5.06 | 1.35 | 2.81E-04 | 0.0037 |
| 4176 | chr3 | 1.27E+08 | 1.27E+08 | + | 0 | NA | intron (NM | L2\|LINE\|L | 66738 | NM_00102 | 108058 | Mm.25582 | NM_02381 | ENSMUSG( | Camk2d | 2810011D( | calcium/ca | protein-co | 6.57 | 7.05 | 5.85 | 1.2 | 2.81E-04 | 0.00371 |
| 4231 | chr1 | 95451068 | 95451218 | + | 0 | NA | Intergenic | Intergenic | 137515 | NM_02632 | 67698 | Mm.2727( | NM_02632 | ENSMUSG( | Fam174a | 2310044D( | family with | protein-co | 6.7 | 7.12 | 6.09 | 1.03 | 2.98E-04 | 0.00388 |
| 4240 | chr4 | 89369239 | 89369389 | + | 0 | NA | intron (NR | intron (NR | -58282 | NM_00767 | 12579 | Mm.42309 | NM_00767 | ENSMUSG( | Cdkn2b | AV083695 | cyclin depe | protein-co | 5.66 | 6.2 | 4.79 | 1.41 | 3.04E-04 | 0.00394 |
| 4241 | chr11 | 71670138 | 71670288 | + | 0 | NA | Intergenic | Intergenic | -80472 | NM_00135 | 216881 | Mm.48648 | NM_17767 | ENSMUSG( | Wscd1 | mKIAA052 | WSC doma | protein-co | 5.26 | 5.86 | 4.21 | 1.65 | 3.04E-04 | 0.00394 |
| 4243 | chr7 | 1.4E+08 | 1.4E+08 | + | 0 | NA | exon (NM_ | exon (NM_ | -5139 | NM_1993( | 212508 | Mm.29608 | NM_1993( | ENSMUSG( | Mtg1 | Gm169\|Gt | mitochonc | protein-co | 4.88 | 5.57 | 3.53 | 2.03 | 3.04E-04 | 0.00394 |
| 4274 | chr7 | 80205111 | 80205261 | + | 0 | NA | intron (NM | RMER4B\|L | 6745 | NM_00136 | 20352 | Mm.27590 | NM_01365 | ENSMUSG( | Sema4b | SemC\|Sem | sema dom | protein-co | 6.52 | 6.96 | 5.86 | 1.11 | 3.14E-04 | 0.00404 |
| 4288 | chr7 | 25006730 | 25006880 | + | 0 | NA | promoter- | promoter- | -868 | NM_00129 | 232975 | Mm.44101 | NM_14492 | ENSMUSG( | Atp1a3 | Atpa-2 | ATPase, Na | protein-co | 6.24 | 6.72 | 5.52 | 1.19 | 3.18E-04 | 0.00408 |
| 4340 | chr7 | 47140031 | 47140181 | + | 0 | NA | Intergenic | ORR1D2\|L | -6422 | NM_01364 | 19259 | Mm.4654 | NM_01364 | ENSMUSG( | Ptpn5 | Step | protein tyr | protein-co | 4.96 | 5.6 | 3.77 | 1.83 | 3.37E-04 | 0.00425 |
| 4357 | chr4 | 63702490 | 63702640 | + | 0 | NA | Intergenic | Intergenic | 42494 | NM_17737 | 326623 | Mm.20815 | NM_17737 | ENSMUSG( | Tnfsf15 | Tl1\|Tl1a\|T | tumor nec | protein-co | 5.12 | 5.72 | 4.06 | 1.66 | 3.44E-04 | 0.00433 |
| 4369 | chr8 | 1.24E+08 | 1.24E+08 | + | 0 | NA | intron (NM | intron (NM | 42439 | NM_13927 | 108148 | Mm.33808 | NM_13927 | ENSMUSG( | Galnt2 | AI480629 | polypeptic | protein-co | 6.7 | 7.12 | 6.09 | 1.04 | 3.47E-04 | 0.00436 |
| 4377 | chr1 | 87847307 | 87847457 | + | 0 | NA | Intergenic | Intergenic | -5905 | NM_17764 | 227333 | Mm.27721 | NM_17764 | ENSMUSG( | Dgkd | AI841987\| | diacylglyce | protein-co | 6.13 | 6.62 | 5.39 | 1.23 | 3.51E-04 | 0.0044 |
| 4436 | chr5 | 1.43E+08 | 1.43E+08 | + | 0 | NA | intron (NM | intron (NM | 28085 | NM_0090( | 19353 | Mm.29251 | NM_0090( | ENSMUSG( | Rac1 | AL023026\| | Rac family | protein-co | 5.74 | 6.27 | 4.9 | 1.37 | 3.75E-04 | 0.00463 |
| 4441 | chr5 | 1.08E+08 | 1.08E+08 | + | 0 | NA | intron (NM | intron (NM | 6142 | NM_01027 | 14581 | Mm.2078 | NM_01027 | ENSMUSG( | Gfi1 | AW495828 | growth fac | protein-co | 5.25 | 5.86 | 4.16 | 1.7 | 3.77E-04 | 0.00465 |
| 4446 | chr4 | 1.41E+08 | 1.41E+08 | + | 0 | NA | intron (NM | intron (NM | 911 | NM_01013 | 13836 | Mm.2581 | NM_01013 | ENSMUSG( | Epha2 | AW545284 | Eph recept | protein-co | 6.06 | 6.55 | 5.31 | 1.24 | 3.79E-04 | 0.00466 |
| 4453 | chr1 | 36797614 | 36797764 | + | 0 | NA | intron (NM | ORR1F\|LTI | 18705 | NM_00128 | 22637 | Mm.8038 | NM_00953 | ENSMUSG( | Zap70 | Srk\|ZAP-7( | zeta-chain | protein-co | 5.65 | 6.18 | 4.79 | 1.4 | 3.82E-04 | 0.00469 |
| 4460 | chr19 | 4304992 | 4305142 | + | 0 | NA | intron (NM | intron (NM | 1155 | NM_00129 | 110355 | Mm.25414 | NM_13086 | ENSMUSG( | Grk2 | Adrbk-1\|A | G protein- | protein-co | 6.1 | 6.58 | 5.39 | 1.2 | 3.84E-04 | 0.00471 |
| 4475 | chr5 | 9015485 | 9015635 | + | 0 | NA | Intergenic | ORR1E\|LTI | -18414 | NM_02373 | 74114 | Mm.28197 | NM_02373 | ENSMUSG( | Crot | 1200003H( | carnitine C | protein-co | 5.37 | 5.96 | 4.38 | 1.58 | 3.89E-04 | 0.00476 |
| 4477 | chr7 | 1.26E+08 | 1.26E+08 | + | 0 | NA | intron (NM | intron (NM | 3272 | NM_02188 | 60504 | Mm.15564 | NM_02188 | ENSMUSG( | Il21r | NILR | interleukir | protein-co | 5.97 | 6.46 | 5.22 | 1.24 | 3.91E-04 | 0.00477 |
| 4483 | chr4 | 1.49E+08 | 1.49E+08 | + | 0 | NA | exon (NM_ | exon (NM_ | -7687 | NM_00136 | 1.03E+08 | | NM_001366402 | | Tmem274 | Gm13097\| | transmeml | protein-co | 4.95 | 5.57 | 3.83 | 1.74 | 3.94E-04 | 0.0048 |
| 4489 | chr2 | 1.65E+08 | 1.65E+08 | + | 0 | NA | intron (NM | intron (NM | 39071 | NM_17498 | 104010 | Mm.15181 | NM_17498 | ENSMUSG( | Cdh22 | - | cadherin 2 | protein-co | 6.35 | 6.8 | 5.69 | 1.11 | 3.96E-04 | 0.00482 |
| 4500 | chr1 | 1.57E+08 | 1.57E+08 | + | 0 | NA | Intergenic | Intergenic | -51692 | NM_02388 | 78255 | Mm.28376 | NM_02388 | ENSMUSG( | Ralgps2 | 1810020P( | Ral GEF wi | protein-co | 5.76 | 6.28 | 4.95 | 1.33 | 4.00E-04 | 0.00485 |
| 4510 | chr19 | 10070396 | 10070546 | + | 0 | NA | intron (NM | intron (NM | 28923 | NM_02189 | 60527 | Mm.25387 | NM_02189 | ENSMUSG( | Fads3 | AI464531 | fatty acid ( | protein-co | 4.55 | 5.25 | 3.15 | 2.1 | 4.03E-04 | 0.00487 |
| 4519 | chr16 | 5383644 | 5383794 | + | 0 | NA | Intergenic | Intergenic | -127763 | NM_02744 | 70511 | Mm.4578( | NM_02744 | ENSMUSG( | Eef2kmt | 5730409G( | eukaryotic | protein-co | 5 | 5.65 | 3.82 | 1.83 | 4.04E-04 | 0.00488 |
| 4512 | chr14 | 19797027 | 19797177 | + | 0 | NA | intron (NM | intron (NM | 26721 | NM_02652 | 68045 | Mm.21932 | NM_02652 | ENSMUSG( | Rtraf | 2700060E( | RNA transc | protein-co | 5.73 | 6.53 | 3.8 | 2.73 | 4.03E-04 | 0.00488 |
| 4513 | chr11 | 98000439 | 98000589 | + | 0 | NA | TTS (NM_1 | TTS (NM_1 | -4341 | NM_20723 | 217151 | Mm.27789 | NM_20723 | ENSMUSG( | Arl5c | Arl12\|BC0 | ADP-ribosy | protein-co | 6.55 | 7.35 | 4.62 | 2.73 | 4.03E-04 | 0.00488 |
| 4539 | chr12 | 31392398 | 31392548 | + | 0 | NA | Intergenic | Intergenic | -41002 | NM_00786 | 13382 | Mm.3131 | NM_00786 | ENSMUSG( | Dld | - | dihydrolip( | protein-co | 4.98 | 5.62 | 3.81 | 1.81 | 4.11E-04 | 0.00495 |
| 4545 | chr5 | 1.48E+08 | 1.48E+08 | + | 0 | NA | promoter- | promoter- | -898 | NM_00135 | 71706 | Mm.15321 | NM_02787 | ENSMUSG( | Slc46a3 | 1200006F( | solute carr | protein-co | 6.4 | 6.86 | 5.73 | 1.13 | 4.15E-04 | 0.00498 |
| 4546 | chr1 | 1.27E+08 | 1.27E+08 | + | 0 | NA | intron (NM | intron (NM | 97928 | NM_14512 | 107895 | Mm.21468 | NM_14512 | ENSMUSG( | Mgat5 | 4930471A( | mannoside | protein-co | 5.86 | 6.36 | 5.08 | 1.28 | 4.15E-04 | 0.00498 |
| 4548 | chr8 | 1.05E+08 | 1.05E+08 | + | 0 | NA | intron (NM | intron (NM | 1311 | NM_14895 | 104394 | Mm.34554 | NM_14895 | ENSMUSG( | E2f4 | 2010111M | E2F transc | protein-co | 6.68 | 7.12 | 6.04 | 1.07 | 4.16E-04 | 0.005 |
| 4577 | chr1 | 20740622 | 20740772 | + | 0 | NA | Intergenic | Intergenic | 9792 | NM_01055 | 16171 | Mm.5419 | NM_01055 | ENSMUSG( | Il17a | Ctla-8\|Ctl | interleukir | protein-co | 5.49 | 6.09 | 4.45 | 1.64 | 4.30E-04 | 0.00512 |
| 4591 | chr4 | 1.25E+08 | 1.25E+08 | + | 0 | NA | Intergenic | RMER15\|L | -84164 | NM_15315 | 230738 | Mm.402 | NM_15315 | ENSMUSG( | Zc3h12a | BC036563 | zinc finger | protein-co | 5.46 | 6.03 | 4.49 | 1.54 | 4.36E-04 | 0.00518 |
| 4594 | chr4 | 10919203 | 10919353 | + | 0 | NA | Intergenic | Intergenic | 44780 | NM_0260( | 67157 | Mm.44055 | NM_0260( | ENSMUSG( | 2610301B( | AI428449 | RIKEN cDN | protein-co | 5.84 | 6.36 | 5.02 | 1.34 | 4.38E-04 | 0.0052 |
| 4602 | chr1 | 52150750 | 52150900 | + | 0 | NA | intron (NM | intron (NM | 31203 | NM_0012( | 20846 | Mm.2774( | NM_00928 | ENSMUSG( | Stat1 | 2010005J( | signal tran | protein-co | 5.16 | 5.77 | 4.1 | 1.67 | 4.41E-04 | 0.00522 |

| | | | | | | | | | | | | | | | | | | | | | | | | |
|---|---|---|---|---|---|---|---|---|---|---|---|---|---|---|---|---|---|---|---|---|---|---|---|---|
| 4607 | chr11 | 57954917 | 57955067 | + | 0 | NA | Intergenic | Intergenic | -54072 | NM_02845 | 73158 | Mm.24884 | NM_02845 | ENSMUSG | Larp1 | 1810024J1 | La ribonuc | protein-co | 6.16 | 6.64 | 5.44 | 1.19 | 4.43E-04 | 0.00525 |
| 4609 | chr1 | 97640590 | 97640740 | + | 0 | NA | Intergenic | Intergenic | 21011 | NM_00135 | 52392 | Mm.12309 | NM_13382 | ENSMUSG | D1Ertd622 | AI987691\| | DNA segm | protein-co | 4.91 | 5.53 | 3.78 | 1.75 | 4.45E-04 | 0.00526 |
| 4617 | chr10 | 1.29E+08 | 1.29E+08 | + | 0 | NA | TTS (NM_1 | TTS (NM_1 | 1139 | NM_14686 | 258861 | Mm.37772 | NM_14686 | ENSMUSG | Olfr763 | MOR269-2 | olfactory r | protein-co | 5.1 | 5.69 | 4.06 | 1.63 | 4.49E-04 | 0.0053 |
| 4621 | chr2 | 70098632 | 70098782 | + | 0 | NA | intron (NM | ORR1F\|LTI | 59581 | NM_17737 | 329421 | Mm.99648 | NM_17737 | ENSMUSG | Myo3b | A430065P1 | myosin IIIE | protein-co | 4.74 | 5.39 | 3.52 | 1.87 | 4.51E-04 | 0.00531 |
| 4623 | chr6 | 29838600 | 29838750 | + | 0 | NA | intron (NM | intron (NM | -15066 | NM_00134 | 74340 | Mm.21089 | NM_02141 | ENSMUSG | Ahcyl2 | 4631427C1 | S-adenosyl | protein-co | 3.74 | 4.55 | 1.73 | 2.82 | 4.52E-04 | 0.00533 |
| 4637 | chr10 | 45004225 | 45004375 | + | 0 | NA | Intergenic | Intergenic | -62916 | NM_01115 | 19072 | Mm.37294 | NM_01115 | ENSMUSG | Prep | AI047692\| | prolyl endo | protein-co | 5.55 | 6.13 | 4.6 | 1.53 | 4.56E-04 | 0.00536 |
| 4648 | chr7 | 19790035 | 19790185 | + | 0 | NA | intron (NM | intron (NM | 6699 | NM_02322 | 80794 | Mm.81698 | NM_02322 | ENSMUSG | Cblc | 2310076I2 | Casitas B-l | protein-co | 6.88 | 7.31 | 6.27 | 1.04 | 4.61E-04 | 0.00539 |
| 4650 | chr2 | 1.63E+08 | 1.63E+08 | + | 0 | NA | Intergenic | Intergenic | -12547 | NM_02663 | 68236 | Mm.33648 | NM_02663 | ENSMUSG | Gtsf1l | 2410116G | gametocyt | protein-co | 6.42 | 6.91 | 5.68 | 1.24 | 4.61E-04 | 0.0054 |
| 4667 | chr3 | 51446869 | 51447019 | + | 0 | NA | intron (NM | intron (NM | 30928 | NM_05308 | 74838 | Mm.27528 | NM_05308 | ENSMUSG | Naa15 | 5730450D1 | N(alpha)-a | protein-co | 4.52 | 5.21 | 3.15 | 2.06 | 4.69E-04 | 0.00547 |
| 4673 | chr5 | 1.4E+08 | 1.4E+08 | + | 0 | NA | intron (NM | intron (NM | 56215 | NR_15261 | 17120 | Mm.27250 | NM_01075 | ENSMUSG | Mad1l1 | AI173502\| | MAD1 mito | protein-co | 5.85 | 6.37 | 5.02 | 1.34 | 4.71E-04 | 0.00549 |
| 4676 | chr16 | 45044512 | 45044662 | + | 0 | NA | Intergenic | Intergenic | -49424 | NM_02643 | 67896 | Mm.18107 | NM_02643 | ENSMUSG | Ccdc80 | 2610001E1 | coiled-coil | protein-co | 4.75 | 5.43 | 3.43 | 2 | 4.72E-04 | 0.00549 |
| 4698 | chr7 | 1.26E+08 | 1.26E+08 | + | 0 | NA | intron (NM | intron (NM | 157047 | NM_00108 | 233865 | Mm.27438 | NM_00108 | ENSMUSG | D430042O | Kiaa0556 | RIKEN cDN | protein-co | 5.37 | 5.98 | 4.29 | 1.69 | 4.81E-04 | 0.00556 |
| 4724 | chr3 | 1.36E+08 | 1.36E+08 | + | 0 | NA | Intergenic | Intergenic | 87735 | NM_02728 | 110173 | Mm.28053 | NM_02728 | ENSMUSG | Manba | 24100300 | mannosida | protein-co | 4.72 | 5.37 | 3.52 | 1.85 | 4.96E-04 | 0.0057 |
| 4729 | chr2 | 70931018 | 70931168 | + | 0 | NA | Intergenic | Intergenic | -105613 | NR_15642 | 228012 | Mm.13651 | NM_17266 | ENSMUSG | Tlk1 | 4930545J1 | tousled-lik | protein-co | 4.46 | 5.15 | 3.09 | 2.06 | 4.99E-04 | 0.00572 |
| 4750 | chr9 | 1.09E+08 | 1.09E+08 | + | 0 | NA | Intergenic | Intergenic | -3090 | NM_02963 | 76500 | Mm.27633 | NM_02963 | ENSMUSG | Ip6k2 | 1500005N | inositol he | protein-co | 6.15 | 6.62 | 5.46 | 1.16 | 5.09E-04 | 0.00581 |
| 4757 | chr5 | 1.15E+08 | 1.15E+08 | + | 0 | NA | Intergenic | ORR1D2\|L | 61198 | NM_00130 | 17855 | Mm.28088 | NM_02355 | ENSMUSG | Mvk | 2310010A | mevalonat | protein-co | 5.98 | 6.47 | 5.23 | 1.24 | 5.12E-04 | 0.00583 |
| 4765 | chr11 | 46127654 | 46127804 | + | 0 | NA | promoter- | promoter- | -175 | NM_00131 | 11492 | Mm.89940 | NM_00961 | ENSMUSG | Adam19 | AL024287\| | a disintegr | protein-co | 5.63 | 6.21 | 4.64 | 1.57 | 5.15E-04 | 0.00586 |
| 4766 | chr17 | 50084049 | 50084199 | + | 0 | NA | intron (NM | intron (NM | 106373 | NM_18139 | 76438 | Mm.41854 | NM_18139 | ENSMUSG | Rftn1 | 2310015N | raftlin lipic | protein-co | 6.44 | 6.89 | 5.79 | 1.1 | 5.16E-04 | 0.00587 |
| 4772 | chr6 | 72588314 | 72588464 | + | 0 | NA | intron (NM | intron (NM | 9908 | NM_00136 | 232089 | Mm.16723 | NM_14491 | ENSMUSG | Elmod3 | AI844780\| | ELMO/CED | protein-co | 5.03 | 5.64 | 3.93 | 1.71 | 5.19E-04 | 0.00589 |
| 4779 | chr10 | 76355079 | 76355229 | + | 0 | NA | intron (NM | intron (NM | -9863 | NM_00108 | 64451 | Mm.33183 | NM_00108 | ENSMUSG | Dip2a | 4931420H1 | disco inter | protein-co | 6.59 | 7.02 | 5.97 | 1.05 | 5.21E-04 | 0.00591 |
| 4785 | chr8 | 95149594 | 95149744 | + | 0 | NA | Intergenic | Intergenic | -7129 | NM_01063 | 16582 | Mm.37895 | NM_01063 | ENSMUSG | Kifc3 | AI325457\| | kinesin far | protein-co | 6.86 | 7.28 | 6.27 | 1.01 | 5.24E-04 | 0.00594 |
| 4799 | chr3 | 1.08E+08 | 1.08E+08 | + | 0 | NA | Intergenic | Intergenic | -30862 | NM_13386 | 99662 | Mm.10849 | NM_13386 | ENSMUSG | Eps8l3 | AI504489\| | EPS8-like 3 | protein-co | 6.33 | 6.77 | 5.69 | 1.09 | 5.35E-04 | 0.00604 |
| 4816 | chr1 | 1.33E+08 | 1.33E+08 | + | 0 | NA | Intergenic | Intergenic | -8239 | NM_18293 | 240753 | Mm.25355 | NM_18293 | ENSMUSG | Plekha6 | 9530028L | pleckstrin | protein-co | 5.21 | 5.8 | 4.19 | 1.62 | 5.43E-04 | 0.0061 |
| 4826 | chr11 | 1.17E+08 | 1.17E+08 | + | 0 | NA | intron (NM | intron (NM | -4505 | NM_01738 | 53860 | Mm.38450 | NM_01738 | ENSMUSG | 9-Sep | MSF1\|Msf | septin 9 | protein-co | 6.54 | 6.96 | 5.94 | 1.03 | 5.46E-04 | 0.00612 |
| 4831 | chr5 | 1.43E+08 | 1.43E+08 | + | 0 | NA | exon (NM_ | exon (NM_ | 5558 | NM_00137 | 330230 | | NM_00137 | ENSMUSG | Zfp853 | Gm792 | zinc finger | protein-co | 6.07 | 6.54 | 5.36 | 1.18 | 5.51E-04 | 0.00616 |
| 4832 | chr7 | 1.13E+08 | 1.13E+08 | + | 0 | NA | intron (NM | intron (NM | 35105 | NM_00136 | 11865 | Mm.33970 | NM_00748 | ENSMUSG | Arntl | Arnt3\|BM | aryl hydro | protein-co | 4.76 | 5.41 | 3.56 | 1.85 | 5.51E-04 | 0.00616 |
| 4841 | chr15 | 1.01E+08 | 1.01E+08 | + | 0 | NA | Intergenic | Intergenic | -8278 | NM_01044 | 15370 | Mm.119 | NM_01044 | ENSMUSG | Nr4a1 | GFRP1\|Gf | nuclear re | protein-co | 5.66 | 6.22 | 4.73 | 1.48 | 5.55E-04 | 0.00619 |
| 4858 | chr15 | 66722039 | 66722189 | + | 0 | NA | intron (NM | intron (NM | 51344 | NM_00937 | 21819 | Mm.44133 | NM_00937 | ENSMUSG | Tg | Tgn\|cog | thyroglobu | protein-co | 4.45 | 5.14 | 3.11 | 2.03 | 5.63E-04 | 0.00626 |
| 4873 | chr14 | 77861338 | 77861488 | + | 0 | NA | intron (NM | RLTR8\|LTF | 13504 | NM_02538 | 66148 | Mm.24804 | NM_02538 | ENSMUSG | Dnajc15 | 1110003P1 | DnaJ heat | protein-co | 5.41 | 5.97 | 4.47 | 1.5 | 5.71E-04 | 0.00633 |
| 4882 | chr10 | 94848504 | 94848654 | + | 0 | NA | intron (NM | intron (NM | 96266 | NM_01879 | 54712 | Mm.25671 | NM_01879 | ENSMUSG | Plxnc1 | 2510048K1 | plexin C1 | protein-co | 5.6 | 6.13 | 4.74 | 1.39 | 5.77E-04 | 0.00638 |
| 4889 | chr17 | 29152700 | 29152850 | + | 0 | NA | Intergenic | Intergenic | 38594 | NM_00136 | 442827 | Mm.20023 | NM_00100 | ENSMUSG | Rab44 | 9830134C1 | RAB44, me | protein-co | 4.51 | 5.19 | 3.19 | 2 | 5.81E-04 | 0.00641 |
| 4900 | chr16 | 38503642 | 38503792 | + | 0 | NA | intron (NM | RMER1A\|( | 18946 | NM_02427 | 76916 | Mm.13223 | NM_02427 | ENSMUSG | Timmdc1 | 2810021C2 | transloca | protein-co | 5.89 | 6.38 | 5.14 | 1.24 | 5.88E-04 | 0.00648 |
| 4919 | chr9 | 35122964 | 35123114 | + | 0 | NA | intron (NR | intron (NR | -6175 | NM_00917 | 20443 | Mm.27597 | NM_00917 | ENSMUSG | St3gal4 | SIAT4-C\|S | ST3 beta-g | protein-co | 6.67 | 7.09 | 6.07 | 1.01 | 5.96E-04 | 0.00653 |
| 4927 | chr5 | 75840531 | 75840681 | + | 0 | NA | Intergenic | Intergenic | 137849 | NM_00136 | 16542 | Mm.285 | NM_01061 | ENSMUSG | Kdr | 6130401C | kinase inse | protein-co | 6.42 | 6.88 | 5.76 | 1.12 | 6.00E-04 | 0.00656 |
| 4925 | chr1 | 1.27E+08 | 1.27E+08 | + | 0 | NA | intron (NM | intron (NM | 158900 | NM_14512 | 107895 | Mm.21468 | NM_14512 | ENSMUSG | Mgat5 | 4930471A | mannoside | protein-co | 5.21 | 5.82 | 4.11 | 1.71 | 5.99E-04 | 0.00656 |
| 4964 | chr8 | 33306682 | 33306832 | + | 0 | NA | intron (NM | RSINE1\|SII | 78770 | NM_00112 | 22427 | Mm.22880 | NM_01172 | ENSMUSG | Wrn | AI846146 | Werner sy | protein-co | 5.97 | 6.46 | 5.22 | 1.23 | 6.20E-04 | 0.00673 |
| 4973 | chr4 | 9681274 | 9681424 | + | 0 | NA | Intergenic | ORR1E\|LTI | -12005 | NM_00129 | 65973 | Mm.22220 | NM_02306 | ENSMUSG | Asph | 2310005F1 | aspartate- | protein-co | 5.65 | 6.18 | 4.79 | 1.39 | 6.26E-04 | 0.00678 |
| 4992 | chr4 | 99169090 | 99169240 | + | 0 | NA | Intergenic | RLTR19A\|I | -24769 | NM_17502 | 242557 | Mm.24166 | NM_17502 | ENSMUSG | Atg4c | Apg4-C\|Ap | autophagy | protein-co | 6.18 | 6.64 | 5.52 | 1.12 | 6.35E-04 | 0.00684 |
| 4989 | chr15 | 6602099 | 6602249 | + | 0 | NA | intron (NM | intron (NM | 22327 | NM_00127 | 23880 | Mm.17090 | NM_01181 | ENSMUSG | Fyb | ADAP\|B63 | FYN bindir | protein-co | 6.03 | 6.51 | 5.32 | 1.19 | 6.34E-04 | 0.00684 |
| 4997 | chr2 | 93131972 | 93132122 | + | 0 | NA | Intergenic | Intergenic | -55501 | NR_15150 | 277414 | Mm.41033 | NM_00102 | ENSMUSG | Trp53i11 | Tp53i11 | transforma | protein-co | 6.55 | 6.96 | 5.96 | 1 | 6.36E-04 | 0.00685 |
| 4996 | chr6 | 97538835 | 97538985 | + | 0 | NA | intron (NM | MER21C\|L | -51090 | NM_00134 | 232288 | Mm.27789 | NM_14514 | ENSMUSG | Frmd4b | 6030440G | FERM dom | protein-co | 4.94 | 5.56 | 3.8 | 1.76 | 6.36E-04 | 0.00685 |
| 5002 | chr7 | 90286266 | 90286416 | + | 0 | NA | Intergenic | Intergenic | -16014 | NM_00128 | 83671 | Mm.26751 | NM_03139 | ENSMUSG | Sytl2 | AI266830\| | synaptotag | protein-co | 5.5 | 6.05 | 4.61 | 1.43 | 6.39E-04 | 0.00688 |
| 5014 | chr8 | 1.16E+08 | 1.16E+08 | + | 0 | NA | Intergenic | Intergenic | 192244 | NM_00102 | 17132 | Mm.27554 | NM_00102 | ENSMUSG | Maf | 2810401A | avian musc | protein-co | 6.04 | 6.56 | 5.21 | 1.35 | 6.45E-04 | 0.00692 |
| 5025 | chr5 | 34908340 | 34908490 | + | 0 | NA | intron (NM | intron (NM | -7500 | NM_00131 | 403174 | Mm.29122 | NM_20727 | ENSMUSG | Msantd1 | A930005I0 | Myb/SANT | protein-co | 5.96 | 6.44 | 5.23 | 1.21 | 6.51E-04 | 0.00697 |
| 5026 | chr8 | 71535794 | 71535944 | + | 0 | NA | intron (NM | intron (NM | 1603 | NM_19809 | 69550 | Mm.26032 | NM_19809 | ENSMUSG | Bst2 | 2310015I1 | bone marr | protein-co | 6.13 | 6.63 | 5.36 | 1.27 | 6.52E-04 | 0.00698 |
| 5031 | chr3 | 1.49E+08 | 1.49E+08 | + | 0 | NA | exon (NM_ | exon (NM_ | 101815 | NM_00108 | 99633 | Mm.9776 | NM_00108 | ENSMUSG | Adgrl2 | AI450192\| | adhesion ( | protein-co | 5.95 | 6.44 | 5.2 | 1.24 | 6.54E-04 | 0.00699 |
| 5036 | chr12 | 3671207 | 3671357 | + | 0 | NA | intron (NM | intron (NM | 98298 | NM_00788 | 13528 | Mm.28620 | NM_00788 | ENSMUSG | Dtnb | - | dystrobrev | protein-co | 6.3 | 6.75 | 5.65 | 1.1 | 6.56E-04 | 0.00701 |
| 5038 | chr7 | 67444541 | 67444691 | + | 0 | NA | Intergenic | Intergenic | -71758 | NM_00103 | 17258 | Mm.13278 | NM_00103 | ENSMUSG | Mef2a | A430079H | myocyte e | protein-co | 5.21 | 5.79 | 4.24 | 1.54 | 6.57E-04 | 0.00702 |
| 5042 | chr19 | 40818921 | 40819071 | + | 0 | NA | Intergenic | (TTTTG)n\| | -12283 | NM_17283 | 240665 | Mm.309 | NM_17283 | ENSMUSG | Ccnj | D430039C1 | cyclin J | protein-co | 5.11 | 5.69 | 4.11 | 1.58 | 6.60E-04 | 0.00704 |
| 5056 | chr9 | 96299712 | 96299862 | + | 0 | NA | intron (NM | intron (NM | 40539 | NM_00118 | 211586 | Mm.17977 | NM_17866 | ENSMUSG | Tfdp2 | 1110029I0 | transcripti | protein-co | 4.33 | 5.02 | 2.94 | 2.08 | 6.70E-04 | 0.00712 |
| 5062 | chr8 | 1.12E+08 | 1.12E+08 | + | 0 | NA | intron (NM | intron (NM | 37247 | NM_02757 | 52815 | Mm.27589 | NM_02757 | ENSMUSG | Ldhd | 4733401P2 | lactate del | protein-co | 6.53 | 6.94 | 5.94 | 1.01 | 6.72E-04 | 0.00714 |
| 5064 | chr2 | 1.53E+08 | 1.53E+08 | + | 0 | NA | intron (NM | intron (NM | 33782 | NR_14925 | 12048 | Mm.23821 | NM_00974 | ENSMUSG | Bcl2l1 | Bcl(X)L\|Bc | BCL2-like 1 | protein-co | 5.99 | 6.48 | 5.24 | 1.24 | 6.73E-04 | 0.00714 |
| 5069 | chr9 | 80156871 | 80157021 | + | 0 | NA | Intergenic | ORR1E\|LTI | -8088 | NM_00103 | 17920 | Mm.4040 | NM_00866 | ENSMUSG | Myo6 | BC029719 | myosin VI | protein-co | 5.71 | 6.23 | 4.89 | 1.33 | 6.76E-04 | 0.00717 |
| 5075 | chr12 | 21480349 | 21480499 | + | 0 | NA | Intergenic | Intergenic | -62988 | NM_01173 | 22630 | Mm.28963 | NM_01173 | ENSMUSG | Ywhaq | 2700028P | tyrosine 3- | protein-co | 6.04 | 6.52 | 5.33 | 1.19 | 6.78E-04 | 0.00718 |
| 5086 | chr13 | 19336465 | 19336615 | + | 0 | NA | Intergenic | Intergenic | 59273 | NM_00134 | 76205 | Mm.41940 | NM_02427 | ENSMUSG | Stard3nl | 0610035N | STARD3 N- | protein-co | 5.14 | 5.72 | 4.15 | 1.57 | 6.85E-04 | 0.00723 |
| 5100 | chr6 | 86620073 | 86620223 | + | 0 | NA | Intergenic | Intergenic | -8026 | NM_02641 | 67855 | Mm.18304 | NM_02641 | ENSMUSG | Asprv1 | 2300003P2 | aspartic pe | protein-co | 5.4 | 5.96 | 4.49 | 1.48 | 6.94E-04 | 0.0073 |
| 5108 | chr1 | 1.8E+08 | 1.8E+08 | + | 0 | NA | TTS (NM_( | TTS (NM_( | 19401 | NM_01118 | 19165 | Mm.33085 | NM_01118 | ENSMUSG | Psen2 | ALG-3\|Ad | presenilin | protein-co | 6.6 | 7.04 | 5.97 | 1.07 | 7.00E-04 | 0.00735 |
| 5110 | chr1 | 1.54E+08 | 1.54E+08 | + | 0 | NA | intron (NM | intron (NM | 1859 | NM_00124 | 240816 | Mm.32934 | NM_00124 | ENSMUSG | Rgsl1 | 4930415K1 | regulator ( | protein-co | 4.42 | 5.11 | 3.09 | 2.02 | 7.00E-04 | 0.00735 |
| 5115 | chr8 | 78622474 | 78622624 | + | 0 | NA | intron (NR | ORR1D2\|L | 113221 | NM_00128 | 76775 | Mm.33463 | NM_02973 | ENSMUSG | Slc10a7 | 2410193C | solute carr | protein-co | 5.58 | 6.11 | 4.76 | 1.34 | 7.03E-04 | 0.00737 |
| 5128 | chr7 | 1.4E+08 | 1.4E+08 | + | 0 | NA | intron (NM | RLTR35B_I | 6725 | NM_00112 | 244234 | Mm.85125 | NM_17553 | ENSMUSG | 5830411N | Scart2 | RIKEN cDN | protein-co | 5.78 | 6.3 | 4.95 | 1.35 | 7.09E-04 | 0.00741 |
| 5126 | chr8 | 46508716 | 46508866 | + | 0 | NA | intron (NM | intron (NM | 17825 | NM_00130 | 14081 | Mm.21032 | NM_00798 | ENSMUSG | Acsl1 | Acas\|Acas | acyl-CoA s | protein-co | 5 | 5.61 | 3.91 | 1.7 | 7.08E-04 | 0.00741 |
| 5142 | chr3 | 1.31E+08 | 1.31E+08 | + | 0 | NA | Intergenic | L2\|LINE\|L | -90069 | NM_00128 | 23971 | Mm.24491 | NM_01186 | ENSMUSG | Papss1 | AI325286\| | 3'-phospho | protein-co | 5.33 | 5.92 | 4.33 | 1.59 | 7.18E-04 | 0.00748 |
| 5161 | chr7 | 63999642 | 63999792 | + | 0 | NA | Intergenic | Intergenic | -60802 | NM_02136 | 50794 | Mm.24047 | NM_02136 | ENSMUSG | Klf13 | 0610043C1 | Kruppel-lik | protein-co | 5.18 | 5.75 | 4.21 | 1.54 | 7.29E-04 | 0.00757 |
| 5181 | chr17 | 72996656 | 72996806 | + | 0 | NA | Intergenic | Intergenic | 78426 | NM_02999 | 77889 | Mm.15410 | NM_02999 | ENSMUSG | Lbh | 1810009F1 | limb-bud a | protein-co | 6.32 | 6.76 | 5.69 | 1.07 | 7.43E-04 | 0.00768 |
| 5204 | chr9 | 75253844 | 75253994 | + | 0 | NA | intron (NM | intron (NM | 21905 | NM_00108 | 208943 | Mm.41339 | NM_00108 | ENSMUSG | Myo5c | 91300030 | myosin VC | protein-co | 5.73 | 6.25 | 4.92 | 1.32 | 7.57E-04 | 0.00779 |
| 5207 | chr1 | 87830067 | 87830217 | + | 0 | NA | intron (NM | intron (NM | -23145 | NM_17764 | 227333 | Mm.27721 | NM_17764 | ENSMUSG | Dgkd | AI841987\| | diacylglyce | protein-co | 5.14 | 5.73 | 4.15 | 1.58 | 7.59E-04 | 0.0078 |
| 5227 | chr17 | 35115539 | 35115689 | + | 0 | NA | promoter- | promoter- | -105 | NM_14893 | 266614 | Mm.10468 | NM_14893 | ENSMUSG | Ly6g5b | - | lymphocyt | protein-co | 5.94 | 6.42 | 5.23 | 1.2 | 7.69E-04 | 0.00787 |
| 5229 | chr2 | 1.52E+08 | 1.52E+08 | + | 0 | NA | intron (NR | intron (NR | 4320 | NM_00100 | 228777 | Mm.33066 | NM_00100 | ENSMUSG | Nrsn2 | Gm123\|Ne | neurensin | protein-co | 6.33 | 6.78 | 5.68 | 1.11 | 7.71E-04 | 0.00788 |
| 5238 | chr13 | 19139235 | 19139385 | + | 0 | NA | exon (NM_ | exon (NM_ | 190959 | NM_17500 | 218038 | Mm.10165 | NM_17500 | ENSMUSG | Amph | - | amphiphys | protein-co | 5.55 | 6.1 | 4.65 | 1.45 | 7.77E-04 | 0.00794 |
| 5244 | chr2 | 30257875 | 30258025 | + | 0 | NA | intron (NM | intron (NM | -8253 | NM_17226 | 227696 | Mm.29794 | NM_17226 | ENSMUSG | Phyhd1 | - | phytanoyl- | protein-co | 5.64 | 6.15 | 4.84 | 1.32 | 7.80E-04 | 0.00795 |
| 5247 | chr14 | 78356875 | 78357025 | + | 0 | NA | Intergenic | L4_A_Man | -48907 | NM_01161 | 21943 | Mm.24922 | NM_01161 | ENSMUSG | Tnfsf11 | Ly109l\|OD | tumor nec | protein-co | 5.52 | 6.05 | 4.68 | 1.36 | 7.80E-04 | 0.00795 |
| 5252 | chr15 | 25865632 | 25865782 | + | 0 | NA | intron (NM | L2b\|LINE\| | 22409 | NM_00103 | 66270 | Mm.25311 | NM_02545 | ENSMUSG | Retreg1 | 1810015C | reticulopha | protein-co | 6.49 | 6.94 | 5.84 | 1.1 | 7.87E-04 | 0.00801 |
| 5294 | chr8 | 70003257 | 70003407 | + | 0 | NA | Intergenic | Intergenic | -6948 | NM_00128 | 234366 | Mm.27004 | NM_14559 | ENSMUSG | Gatad2a | 1110066C1 | GATA zinc | protein-co | 6.11 | 6.57 | 5.43 | 1.15 | 8.17E-04 | 0.00825 |
| 5301 | chr11 | 1.18E+08 | 1.18E+08 | + | 0 | NA | exon (NM_ | exon (NM_ | 763 | NM_02996 | 77727 | Mm.15998 | NM_02996 | ENSMUSG | 6030468B | Il-40 | RIKEN cDN | protein-co | 4.83 | 5.46 | 3.68 | 1.78 | 8.20E-04 | 0.00827 |
| 5323 | chr14 | 47359692 | 47359842 | + | 0 | NA | Intergenic | RMER6B\|L | -14093 | NM_00114 | 16854 | Mm.24861 | NM_01070 | ENSMUSG | Lgals3 | GBP\|L-34\| | lectin, gala | protein-co | 5.63 | 6.15 | 4.79 | 1.36 | 8.34E-04 | 0.00837 |
| 5355 | chr11 | 1.16E+08 | 1.16E+08 | + | 0 | NA | intron (NM | intron (NM | -1508 | NM_18325 | 66528 | Mm.27574 | NM_18325 | ENSMUSG | Smim5 | 2210020M | small integ | protein-co | 5.69 | 6.23 | 4.82 | 1.41 | 8.54E-04 | 0.00852 |
| 5357 | chr11 | 74577161 | 74577311 | + | 0 | NA | intron (NR | intron (NR | 12950 | NM_00101 | 380711 | Mm.42569 | NM_00101 | ENSMUSG | Rap1gap2 | AU067654 | RAP1 GTPa | protein-co | 5.83 | 6.32 | 5.09 | 1.23 | 8.56E-04 | 0.00853 |
| 5360 | chr10 | 1.29E+08 | 1.29E+08 | + | 0 | NA | exon (NM_ | exon (NM_ | 473 | NM_14686 | 258861 | Mm.37772 | NM_14686 | ENSMUSG | Olfr763 | MOR269-2 | olfactory r | protein-co | 5.29 | 5.85 | 4.34 | 1.51 | 8.57E-04 | 0.00854 |
| 5371 | chr6 | 88758013 | 88758163 | + | 0 | NA | intron (NM | MIR\|SINE\| | 33384 | NM_00116 | 23945 | Mm.27219 | NM_01184 | ENSMUSG | Mgll | AA589436 | monoglyce | protein-co | 5.88 | 6.36 | 5.16 | 1.2 | 8.71E-04 | 0.00865 |
| 5379 | chr1 | 84915158 | 84915308 | + | 0 | NA | intron (NM | B3\|SINE\|E | 19850 | NM_02792 | 71781 | Mm.15875 | NM_02792 | ENSMUSG | Slc16a14 | 1110004H1 | solute carr | protein-co | 5.92 | 6.41 | 5.19 | 1.22 | 8.78E-04 | 0.00871 |
| 5413 | chr7 | 34933209 | 34933359 | + | 0 | NA | intron (NM | intron (NM | 20877 | NM_00882 | 18624 | Mm.69751 | NM_00882 | ENSMUSG | Pepd | Pep-4\|Pep | peptidase | protein-co | 4.53 | 5.2 | 3.23 | 1.98 | 9.05E-04 | 0.00892 |
| 5420 | chr2 | 1.25E+08 | 1.25E+08 | + | 0 | NA | intron (NM | intron (NM | 8494 | NM_00129 | 214968 | Mm.33053 | NM_17253 | ENSMUSG | Sema6d | 1110067B | sema dom | protein-co | 5 | 5.6 | 3.94 | 1.67 | 9.10E-04 | 0.00894 |
| 5436 | chr5 | 1.25E+08 | 1.25E+08 | + | 0 | NA | intron (NM | intron (NM | 52649 | NM_02844 | 73121 | Mm.29613 | NM_02844 | ENSMUSG | Rflna | 3110032G | refilin A | protein-co | 6.02 | 6.48 | 5.32 | 1.16 | 9.26E-04 | 0.00907 |
| 5442 | chr4 | 1.3E+08 | 1.3E+08 | + | 0 | NA | promoter- | promoter- | -13 | NM_00116 | 16818 | Mm.29375 | NM_01069 | ENSMUSG | Lck | Hck-3\|Lsk | lymphocyt | protein-co | 5.12 | 5.71 | 4.12 | 1.58 | 9.32E-04 | 0.00911 |
| 5451 | chr3 | 1.31E+08 | 1.31E+08 | + | 0 | NA | intron (NM | intron (NM | 41961 | NM_00127 | 16842 | Mm.25521 | NM_01070 | ENSMUSG | Lef1 | 3000002B | lymphoid e | protein-co | 5.35 | 5.89 | 4.48 | 1.42 | 9.39E-04 | 0.00917 |
| 5461 | chr5 | 1.07E+08 | 1.07E+08 | + | 0 | NA | intron (NM | intron (NM | 19012 | NM_00136 | 52397 | Mm.24947 | NM_02685 | ENSMUSG | Zfp644 | 1110068L | zinc finger | protein-co | 5.01 | 5.6 | 4.02 | 1.58 | 9.45E-04 | 0.0092 |
| 5471 | chr7 | 80143509 | 80143659 | + | 0 | NA | Intergenic | Intergenic | -28234 | NM_17301 | 269951 | Mm.24643 | NM_17301 | ENSMUSG | Idh2 | E430004F2 | isocitrate ( | protein-co | 5.39 | 5.93 | 4.5 | 1.43 | 9.50E-04 | 0.00924 |
| 5472 | chr1 | 1.53E+08 | 1.53E+08 | + | 0 | NA | intron (NM | intron (NM | 103297 | NM_17546 | 226518 | Mm.40548 | NM_17546 | ENSMUSG | Nmnat2 | AI843915\| | nicotinami | protein-co | 4.39 | 5.06 | 3.09 | 1.97 | 9.51E-04 | 0.00925 |
| 5481 | chr18 | 39831977 | 39832127 | + | 0 | NA | Intergenic | Intergenic | 58555 | NM_01103 | 18459 | Mm.9493 | NM_01103 | ENSMUSG | Pabpc2 | Pabp\|Pab | poly(A) bir | protein-co | 5.8 | 6.29 | 5.06 | 1.23 | 9.61E-04 | 0.00931 |
| 5495 | chr13 | 63119147 | 63119297 | + | 0 | NA | intron (NM | intron (NM | -120807 | NR_11052 | 72061 | Mm.18810 | NM_02807 | ENSMUSG | Aopep | 2010111I0 | aminopept | protein-co | 5.48 | 6.01 | 4.64 | 1.37 | 9.73E-04 | 0.0094 |
| 5494 | chr3 | 1.02E+08 | 1.02E+08 | + | 0 | NA | intron (NM | intron (NM | -24786 | NM_17229 | 242125 | Mm.32900 | NM_17229 | ENSMUSG | Mab21l3 | - | mab-21-lik | protein-co | 5 | 5.6 | 3.95 | 1.64 | 9.72E-04 | 0.0094 |
| 5498 | chr2 | 72318595 | 72318745 | + | 0 | NA | intron (NM | ERVL-int\|L | 32969 | NM_17808 | 65964 | Mm.30414 | NM_02305 | ENSMUSG | Map3k20 | AV006891 | mitogen-a | protein-co | 5.62 | 6.19 | 4.69 | 1.49 | 9.74E-04 | 0.00941 |
| 5504 | chr5 | 1.07E+08 | 1.07E+08 | + | 0 | NA | intron (NM | intron (NM | 33977 | NM_17787 | 330149 | Mm.12166 | NM_17787 | ENSMUSG | Hfm1 | A330009G | HFM1, ATF | protein-co | 4.58 | 5.23 | 3.38 | 1.85 | 9.79E-04 | 0.00944 |
| 5506 | chr11 | 76496485 | 76496635 | + | 0 | NA | intron (NM | intron (NM | -2065 | NM_00134 | 109934 | Mm.25893 | NM_19801 | ENSMUSG | Abr | - | active BCR | protein-co | 5.99 | 6.45 | 5.31 | 1.14 | 9.81E-04 | 0.00947 |
| 5507 | chr3 | 98335785 | 98335935 | + | 0 | NA | intron (NM | intron (NM | 4109 | NM_01696 | 236539 | Mm.16898 | NM_01696 | ENSMUSG | Phgdh | 3-PGDH\|3 | 3-phospho | protein-co | 6.01 | 6.47 | 5.32 | 1.15 | 9.82E-04 | 0.00947 |
| 5512 | chr13 | 19337077 | 19337227 | + | 0 | NA | Intergenic | Intergenic | 58661 | NM_00134 | 76205 | Mm.41940 | NM_02427 | ENSMUSG | Stard3nl | 0610035N | STARD3 N- | protein-co | 6.1 | 6.56 | 5.43 | 1.12 | 9.86E-04 | 0.0095 |
| 5532 | chr8 | 1.21E+08 | 1.21E+08 | + | 0 | NA | Intergenic | Intergenic | -10123 | NM_00832 | 15900 | Mm.33486 | NM_00832 | ENSMUSG | Irf8 | AI893568\| | interferon | protein-co | 5.91 | 6.38 | 5.21 | 1.17 | 0.00101 | 0.00972 |
| 5545 | chr2 | 58135122 | 58135272 | + | 0 | NA | intron (NM | intron (NM | 24925 | NM_13920 | 227929 | Mm.27390 | NM_13920 | ENSMUSG | Cytip | A130053M | cytohesin | protein-co | 6.44 | 6.87 | 5.81 | 1.06 | 0.00103 | 0.00982 |
| 5565 | chr19 | 45236656 | 45236806 | + | 0 | NA | 5' UTR (NM | 5' UTR (NM | 110 | NM_01069 | 16814 | Mm.11681 | NM_01069 | ENSMUSG | Lbx1 | Lbx1h | ladybird h | protein-co | 6.38 | 6.8 | 5.79 | 1.01 | 0.00104 | 0.00994 |
| 5566 | chr11 | 1.22E+08 | 1.22E+08 | + | 0 | NA | intron (NM | intron (NM | 17175 | NM_17876 | 319530 | Mm.11395 | NM_17876 | ENSMUSG | Zfp750 | A030007D | zinc finger | protein-co | 5.54 | 6.09 | 4.62 | 1.47 | 0.00105 | 0.00996 |
